# In mice and humans, the brain’s blood vessels mature postnatally to acquire barrier and contractile properties

**DOI:** 10.1101/2021.05.17.444486

**Authors:** Leila Slaoui, Alice Gilbert, Laetitia Federici, Armelle Rancillac, Antoinette Gelot, Maryline Favier, Noémie Robil, Gaëlle Letort, Karine Dias, Laurent Jourdren, Philippe Mailly, Sylvain Auvity, Salvatore Cisternino, Martine Cohen-Salmon, Anne-Cécile Boulay

## Abstract

The brain dense vascular network is essential for distributing oxygen and nutrients to neural cells. The network develops during embryogenesis and leads to the formation of the endothelial blood-brain barrier (BBB). This barrier is surrounded by mural cells (pericytes and vascular smooth muscle cells (VSMCs)) and fibroblasts. Here, we compared the molecular and functional properties of brain vascular cells on postnatal day (P)5 vs. P15, via a transcriptomic analysis of purified mouse cortical microvessels (MVs) and the identification of vascular-cell-type-specific or -preferentially expressed transcripts. We found that endothelial cells (ECs), VSMCs and fibroblasts follow specific molecular maturation programs over this time period. In particular, ECs acquire P-glycoprotein (P-gP)-mediated efflux capacities. The arterial VSMC network expands, acquires contractile proteins (such as smooth muscle actin (SMA) and myosin heavy chain 11 (Myh11)) and becomes contractile. We also analyzed samples of human brain cortex from the early prenatal stage through to adulthood: the expression of endothelial P-gP increased at birth and Myh11 in VSMCs acts as a developmental switch (as in the mouse) at birth and up to the age of 2 of 5 years. Thus, in both mice and humans, the early postnatal phase is a critical period during which the essential properties of cerebral blood vessels (i.e. the endothelial efflux of xenobiotics and other molecules, and the VSMC contractility required for vessel tone and brain perfusion) are acquired and mature.

## Introduction

The brain’s extremely dense vascular system sustains the neurons’ high metabolic demand by providing the cells with a regulated flow of blood and physically and functionally sheltering them from harmful components in the blood. These vessels form a complex, heterogeneous network. Firstly, the arteries at the pial surface branch into penetrating arterioles, which in turn ramify into a dense capillary tree that converges into venules and then veins. Large cortical veins within the subarachnoid space ultimately connect to venous sinuses and egress blood from the brain. The vessels are composed of endothelial cells (ECs), which form the blood-brain barrier (BBB) that separates the blood from the parenchyma in most areas of the brain. ECs are contacted by various types of cell (Vanlandewijck et al., 2018). In the arterial network, ECs are surrounded by concentric, contractile VSMCs, which regulate vascular tone and blood perfusion (Rungta et al., 2018). The VSMCs become progressively sparser as the vessels branch and are present as discrete, non-contractile cells in veins (Frosen and Joutel, 2018; Iadecola, 2017). Pericytes (PCs, another type of periendothelial mural cell) line the vessel walls of pre-capillary arterioles, capillaries, and post-capillary venules. Like VSMCs, PCs form a continuum that ranges from contractile cells around pre-capillary arterioles and capillaries to a mesh of star-shaped, non-contractile cells on post-capillary venules (Hartmann et al., 2015). Perivascular fibroblasts (FBs) are present around vessels other than capillaries (Saunders et al., 2018; Vanlandewijck et al., 2018). The FBs are highly heterogeneous and are thought to have various functions in extracellular matrix (ECM) composition, blood vessel contractility, and immune control (Manberg et al., 2021; Saunders et al., 2018). Lastly, perivascular astrocytic processes (also referred to as “endfeet”) form a continuous sheath around all the vessels (Mathiisen et al., 2010; McCaslin et al., 2011) and control several vascular properties (i.e. BBB integrity, cerebral blood flow, perivascular homeostasis and immunity) (Alvarez et al., 2013; Cohen-Salmon et al., 2021).

In rodents, the brain’s blood vessels start to develop on or around embryonic day (E) 9. Firstly, ECs from a vascular plexus formed by mesodermal angioblasts invade the neuroectoderm and form intraneural vessels (Coelho-Santos and Shih, 2020). This angiogenic phase is followed by a differentiation phase around E15, during which PCs and VSMCs are recruited to the endothelial surface, induce BBB properties (such as the formation of tight junctions (TJs)) between ECs, and thus reduce paracellular molecular transfer and increase transcellular transport (Daneman et al., 2010b; Hellstrom et al., 1999). Interestingly, a second wave of angiogenesis, vascular remodeling and BBB maturation occurs postnatally (Coelho-Santos and Shih, 2020) - suggesting that angiogenesis and BBB genesis are interlinked (Daneman et al., 2009; Daneman et al., 2010a; Liebner et al., 2008).

Although great efforts to characterize the embryonic phase of brain vascular development have been made, the exact time course, molecular bases and functional consequences of the system’s postnatal maturation have barely been described. By combining transcriptomic, biochemical, histochemical and functional assessments, we characterized the brain’s vascular compartment during postnatal development in mice and humans. Our results revealed that all vascular cells (except PCs) follow a specific maturation program. Importantly, we showed that ECs acquire efflux properties that help to protect the brain, while the VSMC network expands and becomes contractile.

## Results

### MVs follow a transcriptional maturation program between P5 and P15

Several studies in rodents have suggested that the brain’s vascular system architecture reorganizes after birth (Coelho-Santos and Shih, 2020). To characterize the molecular bases of these changes, we compared the transcriptome of brain-purified MVs on P5 and P15. We limited our study to parenchymal MVs purified from the dorsal part of the cortex and that were less than 100 µm in diameter, since vascular properties and developmental programs differ from one region of the brain to another. mRNAs extracted from cortical MVs were sequenced, and 15126 transcripts with more than 50 reads in at least one stage were identified (**Fig. 1A**; **Table S1**). 13986 of the transcripts (92 %) were equally expressed at both time points (−1 < log2 fold-change (FC) < 1 or p adjusted value (padj) > 0.05), and only 1140 (8 %) differed significantly between P5 and P15 (log2FC ≤ −1 or log2FC ≥ 1, and padj ≤ 0.05). 627 of these 1140 (55 %) transcripts were upregulated on P15 (log2FC ≥ 1 and padj ≤ 0.05), and 513 (45 %) were downregulated (log2FC ≤ −1 and padj ≤ 0.05) (**Fig. 1A**; **Table S1**). These results indicated that transcription in vascular cells is mostly stable but differs significantly in some respects between P5 and P15. To further characterize these differences, we performed a gene ontology (GO) analysis of our RNA sequencing (RNAseq) data (**Fig. 1B, C; Table S2**). Regarding biological processes, pathways related to mitosis (DNA replication, cell division, and cell cycle) were downregulated on P15, while pathways related to ECM composition, cell junctions and ion transport were upregulated **(Fig. 1B)**. Analysis of the cellular components highlighted lower expression on P15 of pathways related to the chromosomes’ centromeric regions and higher expression of pathways related to the membrane, ECM, cell junction, cell surface, and cytoskeleton **(Fig. 1C)**.

**Fig. 1.**
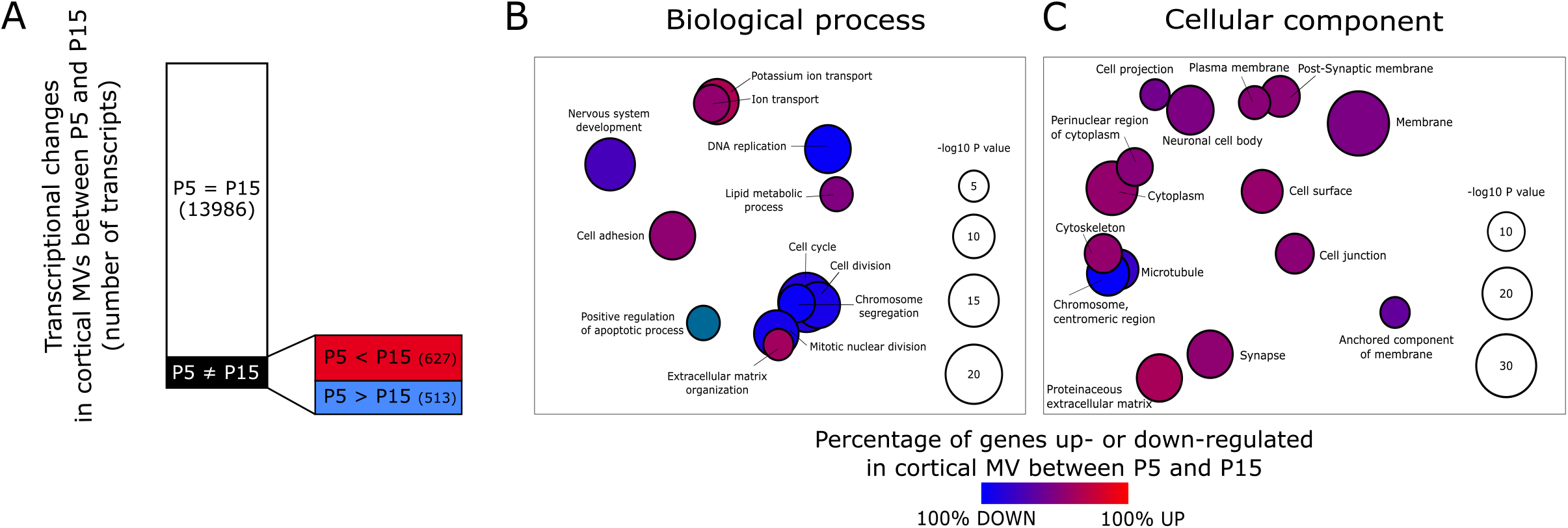
Brain MVs follow a transcriptional maturation program between P5 and P15. **A**. Graphical comparison of the RNAseq data for MV transcripts on P5 vs. P15. Transcripts with no changes (in white): −1 < log2FC < 1 or padj > 0.05; transcripts with changes (in black): log2FC ≤ −1 or log2FC ≥ 1, padj ≤ 0.05; transcripts upregulated on P15 (in red): log2FC ≥ 1, padj ≤ 0.05; transcripts downregulated on P15 (in blue): log2FC ≤ −1, padj ≤ 0.05. **B. C**. A gene ontology (GO) analysis of transcriptional changes in cortical MVs between P5 and P15 for “biological processes” (**B**) and “cellular components” (**C**). A REViGO representation of the results of GO analysis (the data are given in Table S2). n=3 libraries for each stage.

Hence, cortical MVs are mostly stable but follow a transcriptional maturation program between P5 and P15.

### Characterization of postnatal transcriptional maturation in brain vascular cells

We next sought to characterize the maturation of each vascular cell type. By taking advantage of recently published single cell RNASeq datasets for adult vascular cells, we started by identifying the genes preferentially or specifically expressed in ECs, PCs, VSMCs and FBs (He et al., 2016; Vanlandewijck et al., 2018). A transcript was considered to be specific for or preferentially expressed in a cell type when it was (i) detected in more than 60% of the single cells of that type and in a small percentage of other cells (the specific threshold was determined for each cell type, see the Material and Methods)) and (ii) expressed at a higher level than in other cells (LogFC ≥ 1.5). This approach led to the identification of 73 EC-, 14 PC-, 38 FB- and 21 VSMC-transcripts (**Table S3**). To determine the relative contribution of each cell type within the MVs, we compared the expression level of these transcripts in MVs on P5 and P15. To standardize high and low gene expression levels, the reads per kilo base per million mapped reads (RPKM) in MVs were normalized against the RPKM values in the single-cell RNASeq datasets (**Fig. 2A, B**). Our analysis showed that on both P5 and P15, the transcript expression level was higher in ECs than in other cell types - reflecting the fact that ECs are more numerous or more transcriptionally active (**Fig. 2A, B**). We then compared the P5 and P15 values (**Fig. 2C**). Expression levels were higher on P15 for all cell types except PCs, which showed no significant changes between P5 and P15 (**Fig. 2C**). Lastly, we analyzed the FC for each transcript in MVs between P5 and P15: 13, 10 and 10 transcripts were (log2FC ≥ 1, padj ≤ 0.05) upregulated on P15 in ECs, FBs and VSMCs, respectively, and 2 and 1 transcripts were downregulated (log2FC ≤ −1, padj ≤ 0.05) on P15 in FBs and VSMCs, respectively (**Fig. 2D, E, Table S3**).

**Fig. 2.**
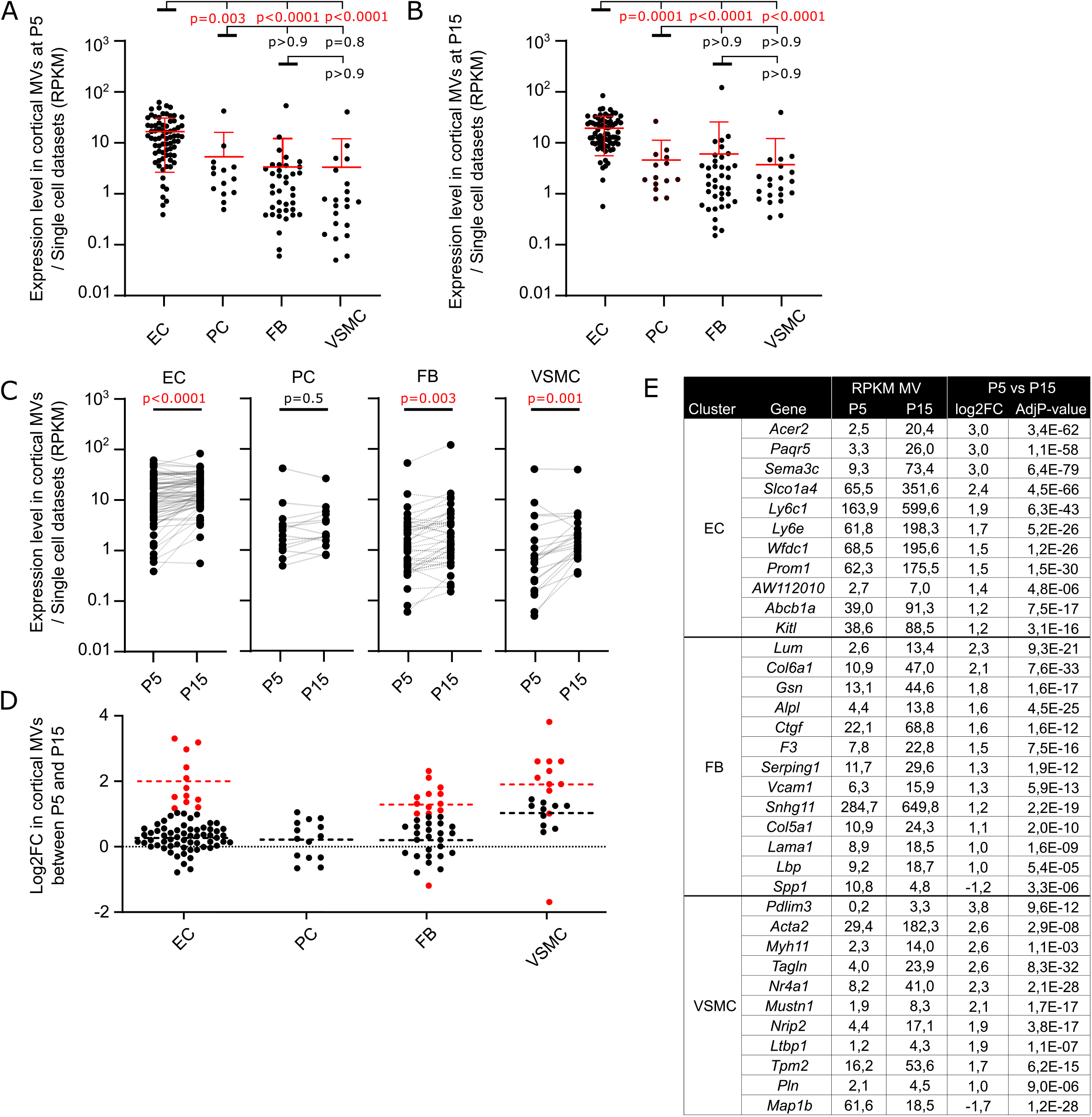
The postnatal transcriptional maturation of vascular cells. **A. B**. Ratio of the expression level (in RPKM) of cell-type-specific or -preferentially expressed transcripts in cortical MVs and in single-cell datasets on P5 (**A**) and P15 (**B**). Data are quoted as the mean ± SD. Dunn’s multiple comparison test (n=3 libraries for each stage). **C**. Comparison of the ratio calculated in **A** and **B** on P5 vs. P15. Data are quoted as the mean ± SD. Wilcoxon matched-pairs signed rank test (n=3 libraries for each stage). **D**. log2FC for cell-type-specific or -preferentially expressed transcripts in cortical MVs on P5 and P15. Each dot represents a transcript. Red dots correspond to significantly changes between P5 and P15 log2FC ≤ −1 or log2FC ≥ 1, padj ≤ 0.05). Black dots correspond to transcripts that did not change between P5 and P15 (−1 < log2FC <1 or padj > 0.05). Data are quoted as the mean ± SD. **E**. RNAseq data for significantly changed cell-type-specific or - preferentially expressed transcripts in cortical MVs between P5 and P15. EC, endothelial cell; PC, pericyte; FB, fibroblast: VSMC, vascular smooth muscle cell.

The brain’s perivascular FBs have not previously been extensively described. In particular, it is not known how and when these cells are recruited to the MV surface. Strikingly, 14 of the 38 FB-specific or -preferentially expressed genes encoded ECM proteins or ECM-associated proteins, such as the secreted integrin-binding glycoprotein osteopontin (the *Spp1* gene), the matricellular connective tissue growth factor (*Ctgf*), Serpin Family G Member 1 (*Serping1*), collagens (*Col1a1, 1a2, 3a1, 5a1, 6a1* and *6a2*), laminin (*Lama1*) or leucine-rich proteoglycans (decorin (*Dcn*), lumican (*Lum*), and osteoglycin (*Ogn*)) (**Table S3**). These data strongly suggested that FBs contribute to the perivascular basal lamina (BL). On P15, six of these genes (*Ctgf, Serping1, Lum, Col6a1, Col5a1* and *Lama1*) were upregulated and one (*Spp1*) was downregulated. The RNAseq results for a selection of genes were validated (using qPCRs) on mRNA extracted from whole-brain purified MVs on P5, P10 and P15 (**Fig. 3A**; **Table S4)**.

**Fig. 3.**
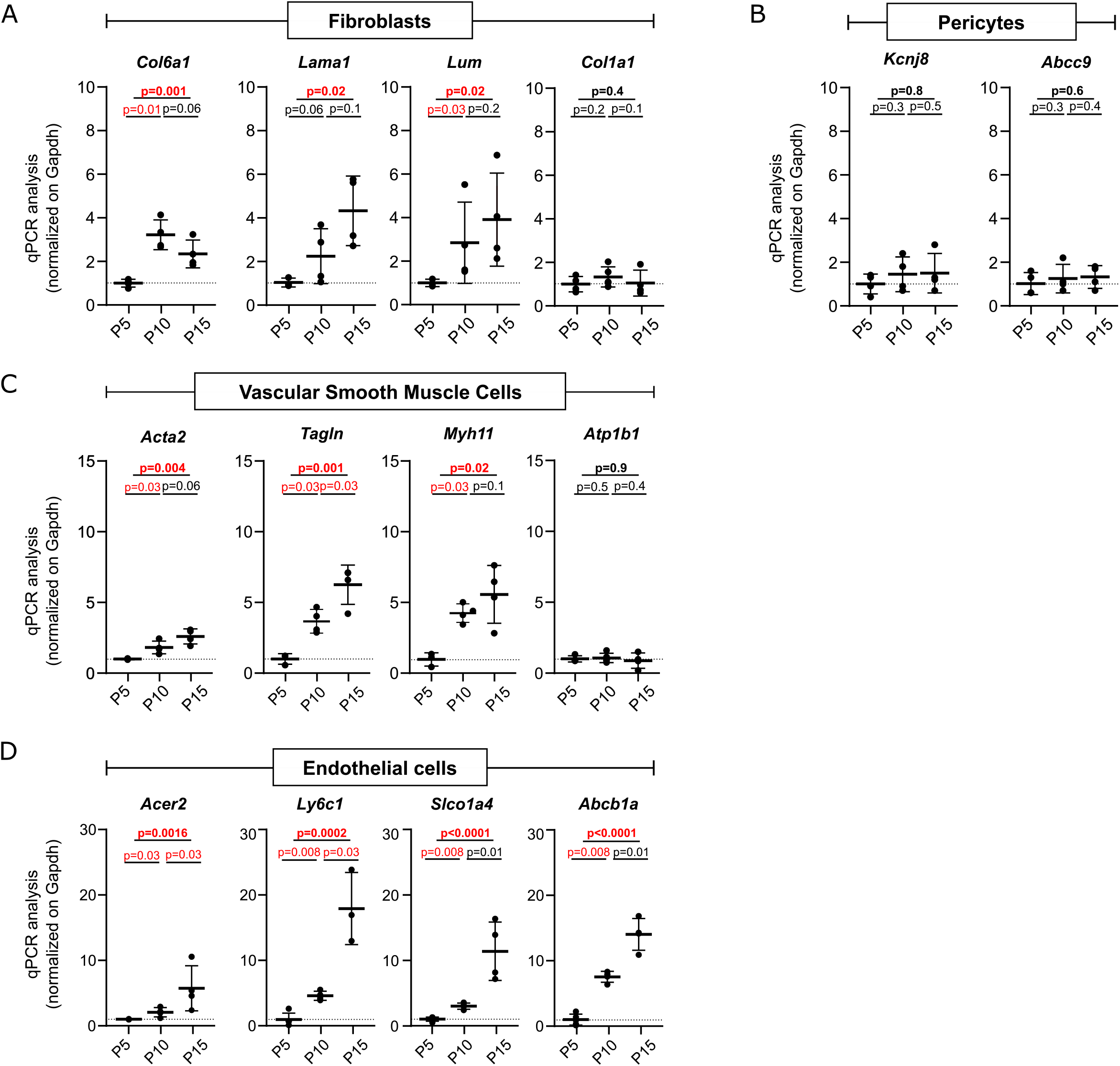
qPCR validations of transcriptional maturation in vascular cells from P5 to P15. The qPCR results for cell-type-specific or -preferentially expressed transcripts on P5, P10 and P15 purified MVs from whole brain are shown. Fibroblasts (**A**); pericytes (**B**); vascular smooth muscle cells (**C**); endothelial cells (**D**). Signals were normalized against *Gapdh*. The value on P5 was set to 1 (dotted line). The plotted data are quoted as the mean ± SD. Kruskal-Wallis test (overall, in bold) and one-tailed Mann-Whitney test (comparison of stages) for 3 to 5 samples per stage (mice per sample: 5 for P5; 3 for P15; 2 for P30; 2 for P60).

PCs have been the focus of intense research over recent years (Alarcon-Martinez et al., 2021). These cells are recruited shortly after vasculogenesis and induce and maintain the integrity of the BBB (Daneman et al., 2010b). The PCs’ molecular properties after birth have not previously been described. Of the 14 genes identified here as being specifically or preferentially expressed in PCs, several coded for channels/transporters (*Abcc9, Kcnj8, Slc6a20a*, and *Trpc3*) or enzymes (*Art3, Ggt1, Enpep*, and *Pla1a*). The expression levels did not change between P5 and P15. The results for a selection of genes were validated using qPCRs on MVs purified from whole brain (**Fig. 3B**; **Table S4)**.

VSMCs form a continuum that ranges from contractile cells in arteries and arterioles to non-contractile cells around veins (Iadecola, 2017). Our analysis identified 21 VSMC-specific or -preferentially expressed transcripts. Ten of these were upregulated on P15; all the corresponding gene products are important contributors to contractility, such as *Acta2* (coding for SMA), *Tpm2* (tropomyosin2), *Myh11* (myosin heavy chain 11) and *Tagln* (SM22/transgelin) (Alexander and Owens, 2012). Again, we validated these RNAseq results using qPCRs on MVs purified from whole brain (**Fig. 3C**; **Table S4)**.

ECs form the vascular wall and sustain certain particular properties of the brain (such as immune quiescence and the BBB) via the expression of specific TJ proteins and transporters regulating molecular transport between the blood and the brain (Daneman et al., 2010a). Eleven of the 73 identified specific or preferentially expressed endothelial transcripts were expressed at a higher level on P15; these included *Abcb1a* (coding for the efflux P-glycoprotein (P-gP) (Loscher and Potschka, 2005)) and *Slco1a4* (coding for the organic anion transporting polypeptide 1a4 (Oatp1a4)). Both genes regulate the transport of xenobiotics across the BBB (Strazielle and Ghersi-Egea, 2015). This upregulation was again confirmed using qPCRs on whole-brain MVs (**Fig. 3D; Table S4)**.

Considering our data as a whole, we identified a set of specific or preferentially-expressed transcripts for each vascular cell type and characterized their expression within MVs between P5 and P15. Our data highlighted specific functions, such as the BL for FBs, BBB transport for ECs, and contractility for VSMCs. These findings suggest that all brain vascular cells (except PCs) undergo a transcriptional maturation between P5 and P15.

### The properties of BBB ECs mature during postnatal development in mice and humans

We next addressed the postnatal maturation of EC functions. Our RNAseq data from cortical MVs indicated that the levels of mRNA coding for EC TJ proteins were similar on P5 and P15 (**Table S1**). Western blots of whole-brain purified MVs showed that the amount of claudin 5 (a TJ protein that is critical for BBB integrity (Nitta et al., 2003)) increased progressively from P5 to P60 (**Fig. 4A; Table S4**). We therefore compared the BBB’s integrity *in vivo* on P5, P15 and P30 by measuring the distribution volume (expressed as an apparent volume, Vvasc, in µL/g) of [^14^C]-sucrose as a marker of the vascular space and the BBB’s physical integrity (Dagenais et al., 2000). The [^14^C]-sucrose Vvasc was similar on P5, P15 and P30, which suggests that EC TJs are already fully functional on P5 (**Fig. 4B, C**).

**Fig. 4.**
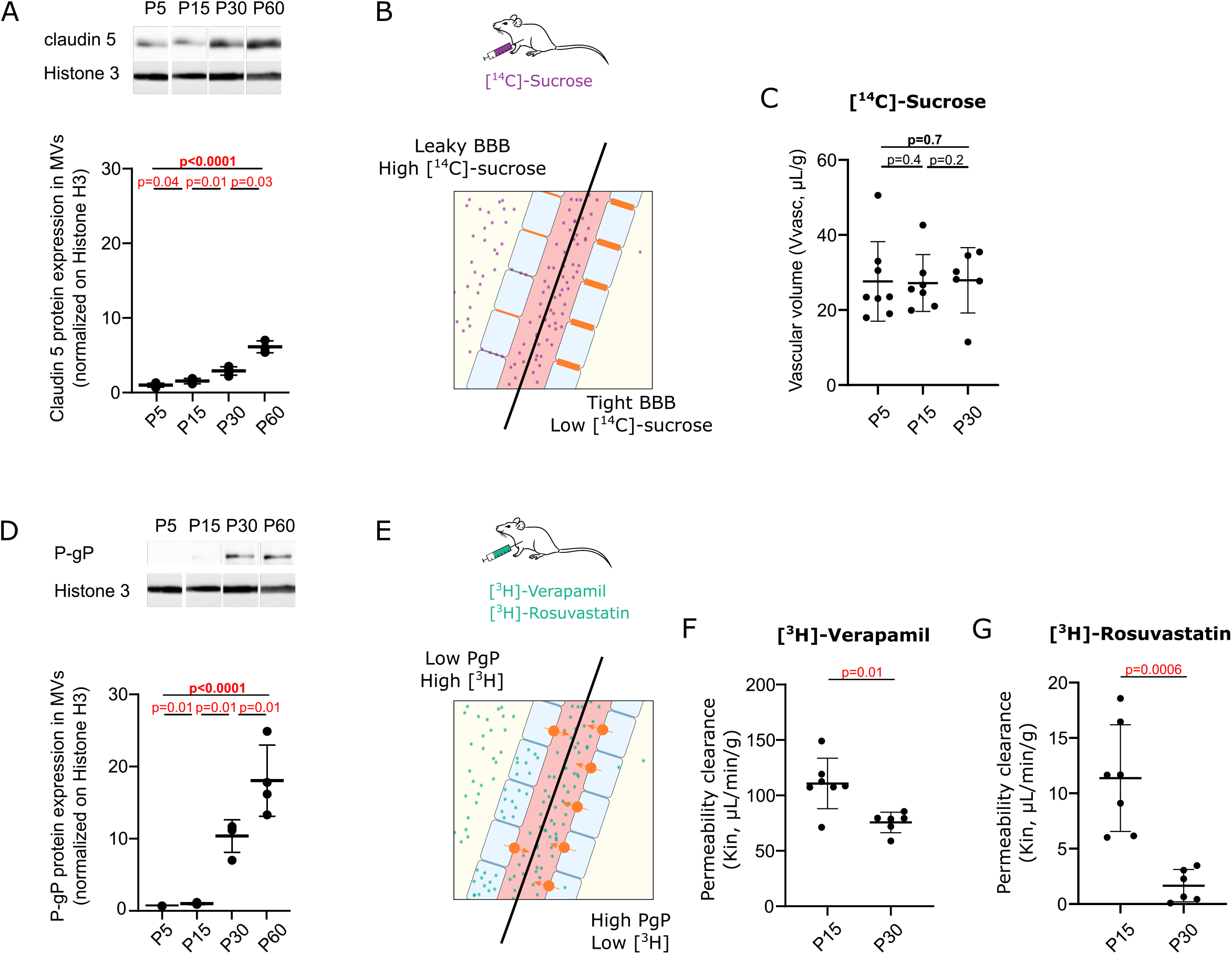
The BBB ECs’ properties mature during postnatal development in the mouse. **A**. Western blot detection and analysis of claudin 5 and histone 3 (H3) in protein extracts from MVs purified from whole brain on P5, P15, P30 and P60. Signals were normalized against H3. The results on P5 were set to 1. Kruskal-Wallis test (overall, in bold) and a one-tailed Mann-Whitney test (comparison of stages). The data are quoted as the mean ± SD (n = 4 samples per developmental stage (mice per sample: 5 for P5; 4 for P10; 3 for P15; 2 for P60)). **B, C**. *In vivo* evaluation of the integrity of BBB TJs on P15 and P30. **B**. Schematic representation of the protocol: mice were intraperitoneally injected with [^14^C]-sucrose as a marker of the vascular space. The amount of [^14^C]-sucrose in the brain was measured (Vvasc, µL/g). Higher levels indicated leakage of [^14^C]-sucrose into the parenchyma as a result of BBB leakage. **C**. Results. The data are quoted as the mean ± SD. Kruskal-Wallis test (overall, in bold) and a one-tailed Mann-Whitney test (comparison of stages) for n=8 mice on P5, 7 on P15 and 6 at P30. **D**. Western blot detection and analysis of P-gP and histone 3 in protein extracts from MVs purified from whole brain on P5, P15, P30 and P60. Signals were normalized against H3. The results on P5 were set to 1. Kruskal-Wallis test (overall, in bold) and a one-tailed Mann-Whitney test (comparison of stages). The data are quoted as the mean ± SD (n = 4 samples per developmental stage; mice per sample: 5 for P5; 4 for P10; 3 for P15; 2 for P60)). **E-G**. *In vivo* evaluation of P-gP activity on P15 and P30. **E**. Schematic representation of the protocol: mice were intraperitoneally injected with [^3^H]-verapamil or [^3^H]-rosuvastatin. The time course of brain transport (Kin, µL/min/g) of [^3^H]-verapamil or [^3^H]-rosuvastatin was measured (2.5-5 min for ^3^H]-verapamil; 5-7 min for [^3^H]-rosuvastatin. A lower level of brain transport corresponds to a higher net efflux across the BBB. **F. G**. The results for [^3^H]-verapamil (**F**) or [^3^H]-rosuvastatin (**G**). The data are quoted as the mean ± SD. One-tailed Mann-Whitney test (n= 7 mice on P15 and 6 at P30).

The BBB plays an important role by exerting two-way control (i.e. influx and efflux) on the transcellular passage of endobiotics and xenobiotics. ECs are equipped with a specific set of ATP-binding cassette (ABC) family transporters (involved in the unidirectional efflux from the brain to the blood) and solute carrier (SLC) transporters (mediating influx) (Loscher and Potschka, 2005; Strazielle and Ghersi-Egea, 2015). Our RNAseq data from cortical MVs indicated that the transcription of *Abcb1a* (encoding P-gP) was upregulated between P5 and P15 (log2Fc=1.4 Padj=1.1 × 10^−15^); these findings were confirmed with qPCRs (**Table S3, S4; Fig. 2E, 3D)**. This transcriptional change also influenced the protein level, albeit with a time lag: the P-gP level on Western blot of P5, P15, P30 and P60 whole-brain MV protein extracts was higher from P30 onwards with a significant shift between P15 and P30 (**Fig 4D; Table S5**). This result prompted us to investigate the *in vivo* function of P-gP by measuring the brain transport of [^3^H]-verapamil, a known P-gP substrate (Chapy et al., 2016). Mice were injected with [^3^H]-verapamil on P15 or P30 (**Fig. 4E-G**). Brain transport of [^3^H]-verapamil was lower on P30 than on P15, which is suggestive of higher P-gP-mediated efflux (**Fig. 4F**). [^3^H]-rosuvastatin (a substrate for both P-gP and Oatp1a4) (Ose et al., 2010; Veszelka et al., 2018) was also significantly less transported on P30 mice than on P15 mice, indicating that P-gP efflux predominates over Oatp1a4 influx during this period of development (**Fig. 4G**).

Our demonstration of greater postnatal P-gP expression and functional activity in the mouse brain prompted us to look at whether this maturation also occurred in humans. We performed an immunohistochemical analysis of P-gP in non-diseased human cortex slices from 15 weeks of gestation (wg) to 17 years of age (**Fig. 5**). P-gP was already present in the brain vessels at 15 wg (**Fig. 5A**). The level of P-gP in brain vessels increased at birth and then stabilized (**Fig. 5A, B**).

**Fig. 5.**
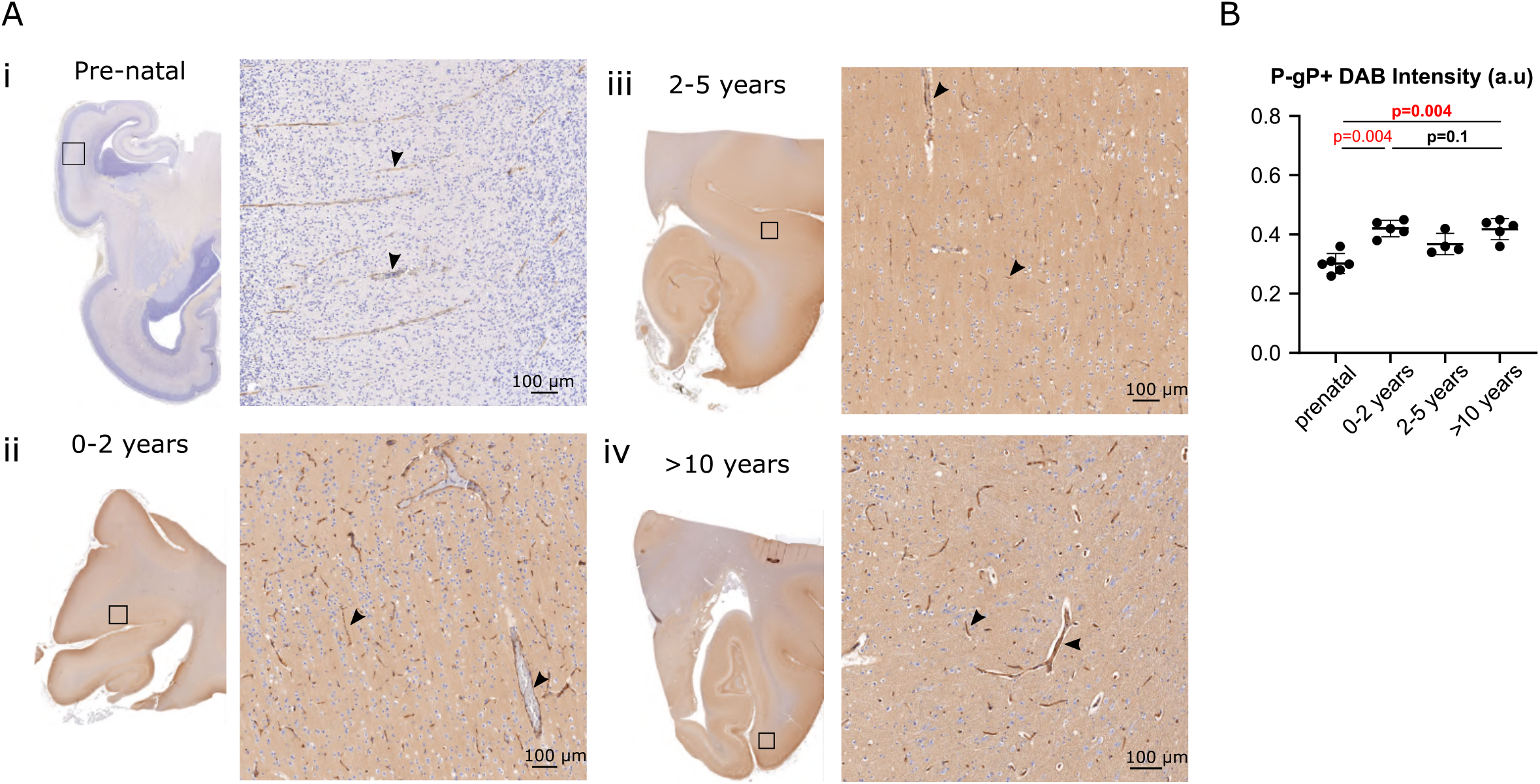
P-gP expression in the developing human cortex. **A, B**. Immunohistochemistry analysis of P-gP in the developing human cortex. **A**. Representative images of P-gP-immunostained cortical slices (left) and (at higher magnification) in the parenchyma of the boxed areas (right) for samples taken (i) in the prenatal period (ii), at 0 to 2 years of age, (iii) between 2 and 5 years of age, and (iv) after 10 years of age. P-gP immunostaining (arrowheads) was revealed by DAB staining. **B**. The DAB intensity was quantified and quoted as the mean ± SD. Kruskal-Wallis test (overall, in bold) and a one-tailed Mann-Whitney test (comparison of stages). Number of samples per developmental age: 6 for prenatal, 5 for 0-2 years, 4 for 2-5 years, and 5 for >10 years.

Taken as a whole, these data indicate that TJs in the murine BBB are fully functional on P15 but that ECs continue to acquire efflux capacity after P15. In humans, the BBB’s efflux properties might develop during early embryogenesis but continue to mature during the late prenatal and early postnatal periods.

### VSMCs acquire contractile properties during postnatal development in mice and humans

We next investigated the postnatal maturation of VSMCs. Our data indicated that the transcription of VSMC-specific genes encoding contractile proteins was upregulated between P5 and P15. To further study this progression, we performed a fluorescent *in situ* hybridization (FISH) analysis of *Myh11* (encoding myosin heavy chain 11) on cortical slices **(Fig. 6A**). On P5, few Myh11 FISH dots were found in the vessels or the parenchyma. In contrast, large vessels gave an intense signal on P15 **(Fig. 6A**). To quantify these differences, we next performed FISH on whole-brain purified MVs **(Fig. 6B)**. On P5, Myh11 FISH dots were detected in most MVs at a moderate density (10-100 × 10^−3^ dots.µm^-3^) (**Fig. 6C**). On P15, the proportion of unlabeled (0-10 × 10^−3^dots.µm^-3^) small-diameter vessels (mean: 5.9 µm ± 3.0) was higher (**Fig. 6C, D**), as was the proportion of densely labeled (>100×10^− 3^ dots.µm^-3^) larger-diameter (mean: 13.5 µm ± 5.6) vessels (**Fig. 6C, D; Table S4**). These data highlighted a change in Myh11 expression from low and diffuse on P5 to high and vessel-type-specific on P15. We next used Western blotting to analyze the protein levels of SMA and Myh11 in protein extracts from MVs purified from whole brain on P5, P15, P30 and P60. Levels of both proteins rose progressively (**Fig. 6E; Table S4**). Overall, these results showed that VSMCs progressively acquired their molecular contractile properties.

**Fig. 6.**
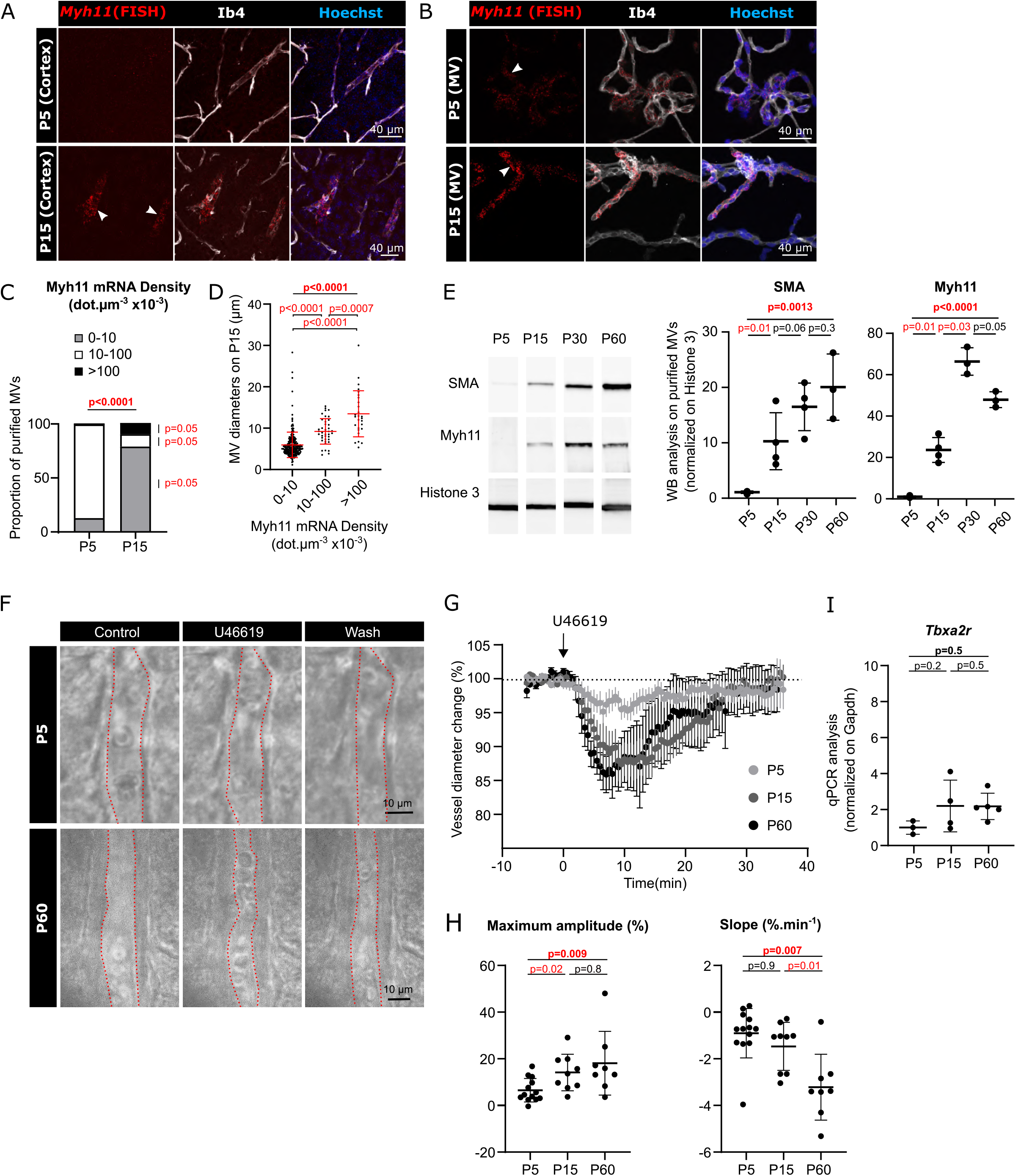
VSMCs acquire contractile properties during postnatal development. **A-D**. FISH analysis of Myh11 expression on P5 and P15. **A**. Confocal microscopy images (FISH detection) of Myh11 mRNAs (red dots) in the cortex (**A**) or in purified MVs (**B**). Vessels were stained with isolectin B4 (IB4). Nuclei were stained with Hoechst. White arrowheads indicate FISH dots in VSMCs. **C, D**. Quantification of (**B). C**. MVs were divided into three categories, according to their FISH dot density. The proportion of MVs in each category on P5 and P15 are shown. A chi^2^ test for changes in the distribution across vessel categories (bold) and a one-tailed Mann-Whitney test for changes over time of each category. **D**. MV diameter for each FISH dot density category on P15. Kruskal-Wallis test (overall, in bold) and a one-tailed Mann-Whitney test (comparison of categories). **E**. Western blot detection and analysis of SMA, Myh11 and histone 3 (H3) in protein extracts from MVs purified from whole brain on P5, P15, P30 and P60. Signals were normalized against H3. The results on P5 were set to 1. Kruskal-Wallis test (overall, in bold) and a one-tailed Mann-Whitney test (comparison of stages). The data are quoted as the mean ± SD (n = 4 samples per developmental stage; mice per sample: 5 for P5; 4 for P10; 3 for P15; 2 for P60). **F-H**. *Ex vivo* analysis of arteriolar constriction on slices of somatosensory cortex from mice on P5, P15 and P60. **F**. Representative infrared images of arteriole constriction in response to bath application of U46619 and dilation upon washing, for samples obtained on P5 and P60. The vessel lumen is indicated by a red dotted line. **G**. Time course of the change in vessel lumen size for P5, P15 and P60 samples. 0 min corresponds to the addition of U46619 in the recording chamber medium. The data are quoted as the mean ± SEM. **H**. Analysis of **(G)**. Maximal amplitude and slope of the contraction. The mean value before U466619 application was set as 0. Kruskal-Wallis test (overall, in bold) and a one-tailed Mann-Whitney test (comparison of stages). The data are quoted as the mean ± SD (n = 13 vessels on P5; 9 on P15; 8 at P60; 3 mice per group). **I**. qPCR results for Tbxa2r in mRNAs from MVs purified from whole brain on P5, P15 and P60. Signals were normalized against *Gapdh*. The results on P5 were set to 1. Kruskal-Wallis test (overall, in bold) and a one-tailed Mann-Whitney test (comparison of stages). The data are quoted as the mean ± SD. n=3 or 4 samples per stage (mice per sample: 5 for P5; 3 for P15; 2 for P30; 2 for P60).

To address the functional consequences of these changes, we compared the *ex vivo* contractility of VSMCs on brain slices sampled on P5, P15 and P60. To this end, we recorded the contractile change in lumen diameter of cortical arterioles upon exposure for 2 min to a thromboxane A_2_ receptor agonist U46619 (9,11-dideoxy-11a,9a-epoxymethanoprostaglandin F2α), which is known to induce reversible vasoconstriction (Perrenoud et al., 2012) (**Fig. 6F-J**). It should be noted that according to our RNAseq data, the level of transcription of *Tbxa2r* (encoding the thromboxane A_2_ receptor) did not change between P5 and P15 (log2Fc = −0.3; padj = 0.5) (**Table S1**). This finding was confirmed (using a qPCR assay) on whole brain MVs (**Fig. 6I; Table S4**). The amplitude of vasoconstriction was assessed as the percentage reduction in vessel diameter upon application of U46619. The slope indicated the velocity of this vasoconstriction. On P5, application of U46619 had a small effect on vessel diameter. In contrast, the amplitude of vasoconstriction was significantly higher on P15 (**Fig. 6F-J, Table S4**) and even higher again on P60 - indicating that VSMC contractility continues to progresses after P15. Thus, in the mouse, vascular contractility rose progressively after birth and was correlated with the expression of SMA and Myh11. In conclusion, our data constitute the first observation of marked postnatal maturation of VSMC contractility in the brain.

### The VSMC contractile network matures postnatally in the mouse cortex and the human cortex

To further characterize the postnatal maturation of VSMCs, we used an immunofluorescence assay to analyze the SMA-positive cortical vascular network on cleared brain samples from P5, P15 and P60 (**Fig. 7A, Table S4**). The endothelial network was counterstained with an anti-Pecam-1 antibody (**Fig. 7B**). In the parenchyma, we observed that the SMA-positive vascular network became progressively denser. Interestingly, the number of SMA-positive primary branches (penetrating arterioles) did not change significantly over time (**Fig. 7A, C**). However, the number of SMA-positive secondary branches and ramifications rose progressively, testifying to expansion and complexification of the arterial VSMC cortical network between P5 and P60 (**Fig. 7A-C)**. We next looked at whether the vascular network in humans also matured after birth by measuring Myh11 expression (using an immunohistochemical assay) on human cortex slices from 15 wg to 17 years of age (**Fig. 7D, E**). In prenatal samples (from 15 to 39 wg), a Myh11 signal was detected in the meninges but not in the parenchyma (**Fig. 7Di**). In contrast, penetrating arteries were intensely labeled after birth (**Fig. 7Dii**). After 2 years of age, a large number of Myh11-immunolabeled vessels were observed throughout the parenchyma (**Fig. 7Diii and iv**). Hence, we quantified a progressive increase in Myh11-positive surface area and the number of Myh11-positive vessels from birth to the age of 2 to 5 years (**Fig. 7E, Table S4**).

**Fig. 7.**
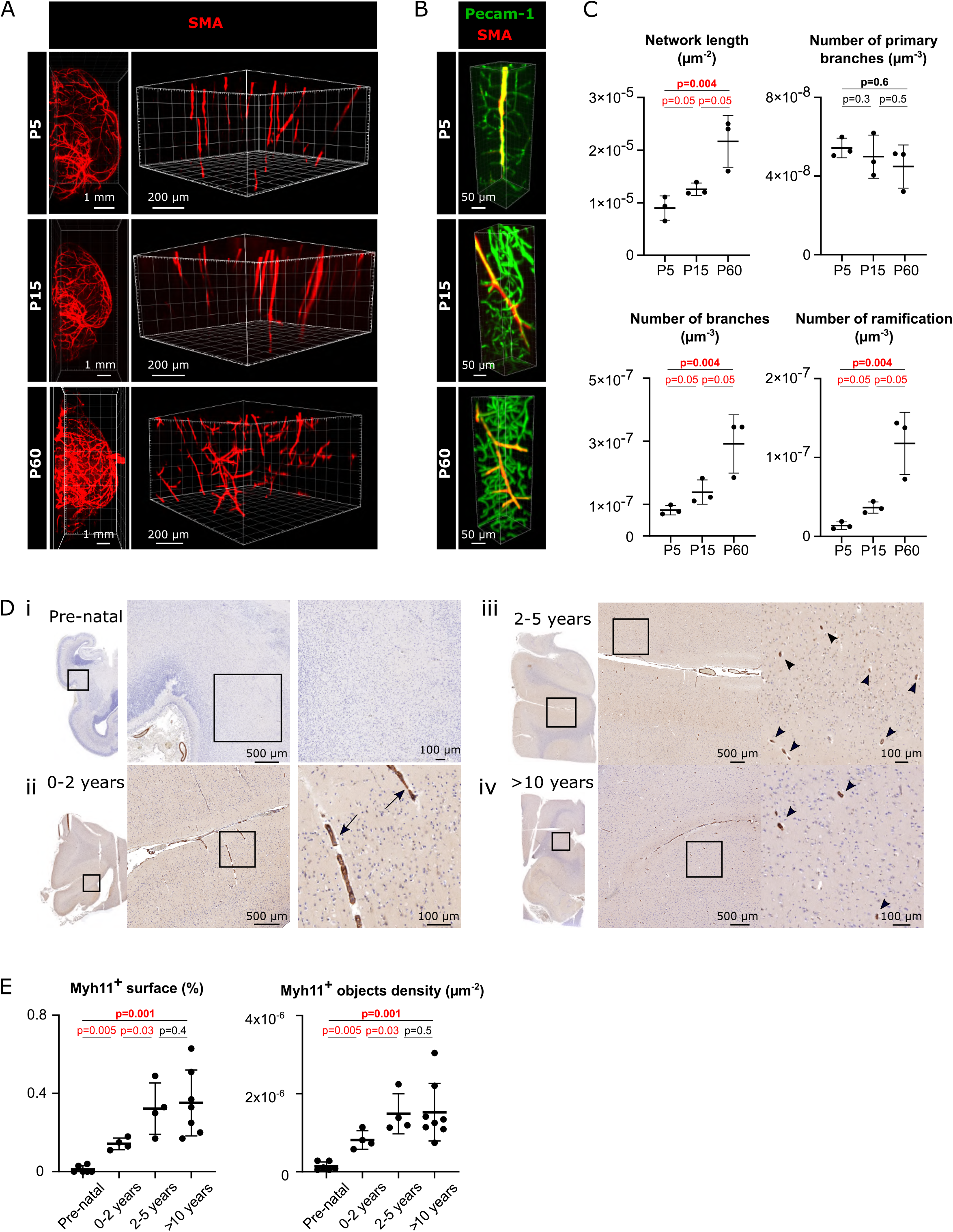
The network of contractile VSMCs matures postnatally in the mouse and human cortex. **A.B**. Representative 3D images of the arterial network in cleared brains and (at higher magnification) in the cortex on P5, P15 and P60, after immunolabeling for SMA (**A, B**) and Pecam-1 (**B**). **C**. Comparative analysis of the SMA-positive vessel length, number of primary or total branches, and ramification in the parenchymal cortex. The data are quoted as the mean ± SD (n = 3 mice per stage). Kruskal-Wallis test (overall, in bold) and a one-tailed Mann-Whitney test (comparison of stages). **D. E**. Immunohistochemistry analysis of Myh11 expression in the developing human cortex. **D**. Representative images of Myh11-stained brain cortical slices (left) and (at higher magnification) in the parenchyma of the boxed area (right) for samples taken (i) in the prenatal period (ii), at 0 to 2 years of age, (iii) between 2 and 5 years of age, and (iv) after 10 years of age. Myh11 expression in penetrating arterioles (arrow) and parenchymal arterioles (arrowheads) was revealed by DAB staining. **E**. Quantification of the stained surface area and the number of detected objects, quoted as the mean ± SD. Kruskal-Wallis test (overall, in bold) and a one-tailed Mann-Whitney test (comparison of stages). Number of samples per developmental age: 6 for prenatal, 5 for 0-2 years, 4 for 2-5 years and 5 for >10 years.

In conclusion, the VSMC network develops progressively after birth in both the mouse and the human. In humans, the Myh11-positive vascular network is not mature before the age of 2 at least.

## Discussion

The development of the brain’s vascular system during embryogenesis has been well described. In contrast, data on the vascular system’s molecular and functional postnatal maturation are scarce. Here, we demonstrated that vascular cells in the mouse and the human follow postnatal transcriptional programs. We also highlighted the endothelial maturation of efflux properties and the acquisition of contractility by VSMCs.

Our analysis of the postnatal maturation of brain vascular cells was firstly based on the transcriptional analysis of MVs mechanically purified from the mouse cortex on P5 and P15. This technique has several advantages. Firstly, the vascular cells were not exposed to enzyme treatments that might induce transcriptomic changes. Secondly, this MV preparation enabled us to probe the contribution of all the component cell types while maintaining the *in vivo* ratio between the latter. We showed that ECs are the main contributors to transcription in cortical MVs on both P5 and P15. Our comparison of these two time points revealed that the transcriptome was stable overall: changes were observed for only 8% of the genes. Nevertheless, our GO analysis highlighted the upregulation of pathways specifically related to ECM composition, cell junctions and ion transport and thus evidence the functional maturation of MVs between P5 and P15. In contrast, we observed the downregulation of pathways related to DNA replication, cell division and the cell cycle - suggesting that the putative angiogenic program involved in the postnatal densification of the capillary network was less active on P15.

We next focused on transcripts that were preferentially or specifically expressed in each vascular cell type. We identified these transcripts by checking brain vessel single-cell RNAseq databases (He et al., 2016; Vanlandewijck et al., 2018). Our analysis therefore focused on the transcriptional signature of each vascular cell type, rather than overlaps in molecular repertoires between cell types. Several transcripts encoding ECM components were specifically expressed by FBs, confirming that these cells actively contribute to the BL (Manberg et al., 2021) and (possibly) its postnatal development because some genes were more strongly expressed on P15 than on P5 (Coelho-Santos and Shih, 2020). The recruitment of perivascular FBs and their differentiation has not previously been fully described. The literature data suggest that perivascular FBs are recruited from the meninges after birth (Kelly et al., 2016). Here, we found that half of the FB transcripts were upregulated between P5 and P15; hence, postnatal molecular maturation might also cover the FBs. Our list of PC-preferentially expressed transcripts was consistent with previous transcriptomic analyses (Bondjers et al., 2006; Chasseigneaux et al., 2018); in particular, we identified *Kcnj8* (encoding the ATP-sensitive inward rectifier potassium channel Kir6.1) and *Abcc9* (encoding the ABC transporter 9) involved in ion transport and intercellular signaling (Nelson et al., 2015). Interestingly, expression of all the PC-preferentially expressed transcripts did not differ significantly when comparing P5 and P15; this suggests that PC differentiation and maturation are already complete on P5. Our P5-P15 comparison of MVs showed that some EC-specific or -preferentially expressed transcripts were upregulated and thus indicating that ECs continued to mature after birth. The expression levels of transcripts of TJ proteins or TJ-associated proteins were similar on P5 and P15 cortical MVs. However, we observed a progressive increase between P5 and P60 in claudin 5 expression in MVs purified from whole brain. This apparent discrepancy might be due to differences in transcriptional regulation between regions of the brain. Nevertheless, the BBB integrity did not change between P5 and P30. These results are in line with previous reports in which the barrier function was fully established during embryogenesis (by around E15) in the mouse (Daneman et al., 2010b) and also suggest that even intermediate levels of claudin 5 are enough to make the BBB’s paracellular impermeability fully operative.

The control of molecular access to the brain parenchyma also requires transcellular regulation via the expression of ABC and SLC transport proteins at the BBB. Importantly, P-gP levels increased 10-fold from P15 to P30 in mouse MVs. These results were in line with the greater P-gP substrate efflux observed on P30 (vs. P15), which showed that this functional property of brain ECs matured postnatally. The literature data on the presence of P-gP in the human cortex are contradictory, i.e. no expression at 22-26 wg in one study (Daood et al., 2008) but expression at 22 wg in another (Virgintino et al., 2013). Although an increase has already been documented between 33-43 wg and adulthood, the intermediate stages have not previously been described (Daood et al., 2008). Here, we demonstrated that P-gP expression in the cortex can be observed as early as 15 wg, increases at birth, and then stabilizes. At the placental barrier, P-gP has a role in protecting the fetus from exposure to harmful compounds present in the maternal circulation (Saunders et al., 2019). Hence, our results might suggest that a pre-term infant’s brain is more vulnerable to these threats.

The vast majority of VSMC-preferentially expressed or -specific transcripts identified in our study were related to muscle differentiation and contractility. Apart from well-known VSMCs transcripts (such as *Acta2, Myh11, desmin* and *Tgln*), our analysis highlighted the expression of markers like *NR4a1* (coding for nuclear receptor subfamily 4, group A, member 1, involved in SMC proliferation) (Hinze et al., 2013; Yu et al., 2015), *Pdlim3* (coding for actinin-associated LIM protein, involved in myocyte stability (Zheng et al., 2010), *Mustn1* (regulating myoblast differentiation) (Hadjiargyrou, 2018; Liu et al., 2010), and *Pln* (coding for phospholamban, which regulates the sarcoplasmic reticulum Ca^2+^ATPase (SERCA) activity and muscle contractility) (Kranias and Hajjar, 2012). Surprisingly, about half of the genes were strongly upregulated on P15; this indicates that VSMC contractility continued to differentiate between P5 and P15. These results were consistent with our *ex vivo* observations of the progressive acquisition of contractility by cortical arterioles. It is noteworthy that some of these transcripts were expressed (albeit at a much lower level) by PCs. Nevertheless, our FISH analysis showed that the strongest expression of *Myh11* was limited to the largest vessels, and therefore excluded the PCs. A postnatal phenotypic switch therefore occurred in murine VSMCs, with increased expression of contractile proteins and the acquisition of contractility. Importantly, this phenotypic switch probably also occurs in the human cortex at some point in the first two years after birth, as illustrated by the appearance of Myh11 immunoreactivity. Furthermore, we observed a progressive increase in the number of SMA-positive vessels (in the mouse, from P5 to P60) and Myh11-positive vessels (in humans, up until the age of two), which shows that the arteriolar network continues its densification postnatally. It is thought that during embryogenesis, VSMC are recruited to newly formed vessels in two ways: (i) the *de novo* formation of VSMCs via the induction of undifferentiated perivascular mesenchymal cells, and (ii) the migration of VSMCs from a preexisting pool (Hellstrom et al., 1999). The same mechanisms might operate during the VSMC network’s postnatal expansion. During embryogenesis, VSMC differentiation is regulated by several signaling pathways (including VEGFA, Notch, EphrinB2/B4, PDGFβ and TGBβ pathways) between ECs and mural cells (Owens et al., 2004). It would be interesting to address these pathways’ role during the postnatal period. Taken as a whole, our results in the mouse and the human suggest that the myogenic tone of parenchymal arterioles (which determines vascular tone and brain perfusion) is acquired progressively after birth. Non-invasive MRI studies of intracranial aneurysms have shown that blood flow in the cortex was more impaired in preterm newborns (before 36 wg) than in term newborns (Chalouhi et al., 2012). From a pharmacological point of view, the maturation of the brain vasculature’s responsiveness is associated with differential responses to vasoactive drugs in preterm newborns, term newborns, and adults. Thus, the best vasoactive drugs for the management of neonatal systemic hypotension might not be those used for the same purpose in adults (Levene et al., 1990; Toth-Heyn and Cataldi, 2012). Lastly, impairments of VSMC differentiation and contractility have been linked to several small-vessel diseases, such as cerebral aneurysm (Chalouhi et al., 2012), arteriovenous malformations, and cerebral cavernous malformations (Frosen and Joutel, 2018; Uranishi et al., 2001). In view of our results, impairments in the VSMCs’ postnatal differentiation might also be involved in these diseases.

In conclusion, our present results revealed that the parenchymal cerebral vasculature in the mouse and in the human undergoes a profound molecular maturation after birth. This maturation results notably in the acquisition of endothelial efflux properties and VSMC contractility. These properties govern blood-brain homeostasis, drug resistance, and cerebral perfusion; hence, their maturation is a crucial step in the acquisition of brain functions and might guide treatment approaches in pediatric medicine.

## Acknowledgments

This work was funded by grants from the *Association Européenne contre les Leucodystrophies* (ELA) (ELA2012-014C2B), the *Fondation pour la Recherche Médicale* (FRM) (AJE20171039094) and the *Fondation Maladies Rares* (20170603). A. Gilbert’s PhD fellowship was funded by the FRM (PLP20170939025p60) and ELA (ELA2012-014C2B). L. Slaoui’s fellowship PhD was funded by the Ecole Normale Supérieure. A.-C. Boulay’s work was funded by the FRM (AJE20171039094) and the *Foundation pour l’aide à la recherche sur la sclérose en plaques* (ARSEP). Despite our efforts, our work has not received any support from the French National Agency for Research (ANR).

## Methods

### Animal experiments and ethical approval

Mice were purchased from Janvier Labs (Le Genest-Saint-Isle, France) and kept in pathogen-free conditions. All animal experiments were carried out in compliance with (i) the European Directive 2010/63/EU on the protection of animals used for scientific purposes and (ii) the guidelines issued by the French National Animal Care and Use Committee (reference: 2013/118). The study was also approved by the French Ministry for Research and Higher Education’s institutional review board.

### MV purification

MVs were purified from dorsal cortex (for RNASeq analyses) or whole brain (for other experiments), as described previously (Boulay et al., 2015). Meningeal vessels were removed by applying a 100 µm-mesh negative filter, and cell debris was removed by then applying a 20 µm-mesh positive filter.

### RNA sequencing and analysis

Total mRNA was extracted from purified cortical MVs on P5 or P15, using the RNeasy Lipid Tissue Kit (Qiagen, Hilden, Germany). Ten ng of total RNA were amplified and converted into cDNA using a SMART-Seq v4 Ultra Low Input RNA Kit (Clontech). Next, an average of 150 pg of amplified cDNA per library were processed with a Nextera XT DNA Kit (Illumina). Libraries were multiplexed on two high-output flow cells and sequenced (using 75 bp reads) on a NextSeq 500 device (Illumina). The mean ± standard deviation (SD) number of reads meeting the Illumina quality criterion was 48 ± 5 million per sample. The RNA-seq gene expression data and raw fastq files are available on the GEO repository (www.ncbi.nlm.nih.gov/geo/) under the accession number GSE173844. The RNA-Seq data were analyzed using GenoSplice technology (www.genosplice.com). Sequencing, data quality checks, read distribution checks (e.g. for potential ribosomal contamination), and insert size estimation were performed using FastQC, Picard-Tools, Samtools and rseqc. Reads were mapped onto the mm10 mouse genome assembly using STARv2.4.0 (Dobin et al., 2013). The procedures for the gene expression regulation study have been described elsewhere (Noli et al., 2015). Briefly, for each gene present in the Mouse FAST DB v2018_1 annotations, we counted the reads aligning on constitutive regions (which are not prone to alternative splicing). Based on these counts, the level of differential gene expression was normalized using DESeq2 in R (v.3.2.5). Genes were considered to be expressed when their RPKM value was greater than 99% of the value for the background (intergenic region). Only genes expressed in at least one of the two experimental conditions were analyzed further. Expression was considered to have changed significantly when the log2 fold change was ≥ 1 or ≤ −1 and padj was ≤ 0.05.

### Pathway/Gene Ontology (GO) analysis

The GO analysis was performed using the DAVID functional annotation tool (version 6.8) (Huang da et al., 2009; Huang et al., 2007). GO terms and pathways were considered to be enriched if the following conditions were met: fold enrichment ≥ 2.0, uncorrected p-value ≤ 0.05, and minimum number of regulated genes in pathway/term ≥ 2.0. The analysis was performed three times: on all regulated genes, on upregulated genes only, and on downregulated genes only. The three sets of results were emerged to provide a single list. The transcription factor analysis was performed using mouse and human orthologs and the DAVID functional annotation Tool (version 6.8) (Huang da et al., 2009; Huang et al., 2007). The results were visualized using REViGO web tool (Supek et al., 2011).

### Single-cell RNA-seq analysis

Raw reads from GEO datasets GSE99058 and GSE98816 were downloaded (He et al., 2016; Vanlandewijck et al., 2018). Seurat 3.1.1 was used to normalize unique molecular identifiers (Butler et al., 2018), using a global-scaling method with a scale factor of 10000 and log-transformation of the data. This was followed by a linear transformation scaling step, so as to avoid highly expressed genes with an excessively high weight in the downstream analysis. With the exception of PCs, cell types were grouped by their level of identity: fibroblasts (fibroblast-like type 1 and 2); ECs (types 1, 2 and 3 ECs, arterial ECs, venous ECs, and capillary ECs), and VSMCs (arteriolar SMCs, arterial SMCs, and venous SMCs). A transcript was considered to be specific or preferentially expressed in a given cell type when it was detected in more than 60% of the corresponding single cells and in a small percentage of cells of other types (PCs, 15%; VSMCs, 40%; ECs, 25%; FBs, 20%) and had a higher expression level than in other cells (according to a Wilcoxon rank sum test, and with logFC > 1.5).

### Quantitative RT-PCR

RNA was extracted using the RNeasy Mini Kit (Qiagen). cDNA was then generated from 100 ng of RNA using the Superscript III Reverse Transcriptase Kit. Differential levels of cDNA expression were measured using droplet digital PCR (ddPCR)). Briefly, cDNA and primers were distributed into approximately 10000 to 20000 droplets. The nucleic acids were then PCR-amplified in a thermal cycler and read (as the number of positive and negative droplets) on a QX200 ddPCR system (Biorad, Hercules, CA, USA)). The ratio for each tested gene was normalized against the total number of positive droplets for *Gapdh* mRNA. The primer sequences are given in Table S5. Three to five independent samples were analyzed in each experiment.

### High-resolution fluorescent *in situ* hybridization

Fluorescent *in situ* hybridization (FISH) was performed on floating PBS/paraformaldehyde (PFA) 4% fixed brain sections or purified MVs immobilized on a glass slide coated with Cell-Tak (Corning) and fixed for 10 min in PBS/PFA 4%, according to the v2 Multiplex RNAscope technique (Advanced Cell Diagnostics, Inc., Newark, CA, USA) described previously (Oudart et al., 2020). Brain sections (but not MVs) were treated with protease at room temperature. After the FISH, MVs were stained with isolectin B4 (1/100) in PBS/ normal goat serum (NGS) 5%/Triton 0.5% overnight at 4°C. Nuclei were stained with Hoechst reagent (1/2000). The brain sections and purified MV were imaged using a Spinning Disk CSU-W1 microscope and Metamorph Premier 7.8 software.

### FISH quantification

The mRNA density in vessels was analyzed using a newly developed “Vessel_Scope” ImageJ plugin (Rueden et al., 2017). In a calibration step, the intensity of a single mRNA dot was estimated in each experimental condition. Isolated dots were detected using the cell counter ImageJ plugin and segmented using the mcib3D library (Ollion et al., 2013). The background intensity was calculated for regions of interest drawn near each dot:

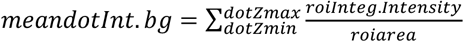

where dotZmin and dotZmax correspond to the dot’s lower and upper z positions, respectively. The background-corrected “single mRNA” intensity was then determined as:

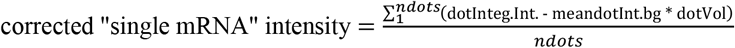

Each 3D Z-stack image was then analyzed. MVs were segmented using 3D median filter and Li threshold. RNAScope dots were segmented using difference of Gaussian filter and Triangle threshold. Segmented objects were detected using the mcib3D library (Ollion et al., 2013). For each MV, the number of single mRNAs was defined as the total intensity of dots in the MV divided by the corrected “single mRNA” intensity. The mRNA density of each MV was calculated as the number of single mRNAs divided by the MV’s volume. The MV diameters were measured manually using the straight line tool in ImageJ.

### Western blots

MV pellets were sonicated three times for 10 s at 20 Hz (Vibra cell VCX130) in 2% SDS and heated at 56°C in Laemmli loading buffer (Biorad). The protein content was measured using the Pierce 660 nm protein assay kit (Thermo Scientific, Waltham, MA, USA). 10 µg of proteins were separated by denaturing electrophoresis on a 4– 15% Criterion™ TGX™ Precast Midi Protein Gel (Biorad) and then electrotransferred to nitrocellulose membranes using the Trans-blot Turbo Transfer System (Biorad). Membranes were hybridized, as described previously (Ezan et al., 2012). The antibodies used in the present study are listed in Table **S5**. Horseradish peroxidase activity was visualized by enhanced chemiluminescence in a Western Lightning Plus system (Perkin Elmer, Waltham, MA, USA). Chemiluminescent imaging was performed on a FUSION FX system (Vilber, South Korea). The level of chemiluminescence for each antibody was normalized against the histone 3 staining on the membrane.

### Clearing and immunohistochemical analysis of murine tissue samples

Mice were killed with pentobarbital (600 mg/kg, i.p.). Brains were removed and post-fixed in 4% PFA for 24 h at 4°C and then assessed using the “immunolabeling-enabled three-dimensional imaging of solvent-cleared organs” technique (Renier et al., 2014). The samples were first dehydrated with increasingly concentrated aqueous Methanol (MetOH) solutions (MetOH: 20%, 40%, 60%, 80%, and twice 100%, for 1 h each) at RT and then incubated in 66% dichloromethane (DCM, Sigma Aldrich)/33% MetOH overnight. After 2 washes in 100% MetOH, brains were incubated in 5% H_2_O_2_/MetOH overnight at RT, rehydrated with increasingly dilute aqueous MetOH solutions (80%, 60%, 40%, and 20%; 1h each). Before immunostaining, brains were permeabilized first for 2 × 1h at RT in 0.2% Triton X-100/PBS, for 24 h at 37°C in 0.16% Triton X-100/2.3% glycine/20% DMSO/PBS, and then for 2 days at 37°C in 0.16% Triton X-100/6% donkey serum/10% DMSO/PBS. Brains were incubated for 3 days at 37°C with primary antibody diluted in a 0.2 Tween/1% heparin/3% donkey serum/5% DMSO/PBS solution, washed 5 times during 24h at 37°C in 0.2% Tween20/1% heparin/PBS solution, incubated for 3 days at 37°C with secondary antibody diluted in a 0.2 Tween/1% heparin/3% donkey serum/PBS solution, and another washed five times. The brain samples were then dehydrated again with a MetOH/H_2_O series (20%, 40%, 60%, 80% and 100% for 1h each, and then 100% overnight) at RT. On the following day, brains were incubated for 3h in 66% DCM/33% MetOH and then twice for 15 min at RT in 100% DCM and lastly cleared overnight in dibenzyl ether.

The cleared tissue were imaged using a light sheet microscope and Inspector pro software (Lavision Biotec GmbH, Bielefeld, Germany). 3D reconstructions were visualized with Imaris software (Bitplane). The length and number of branch points of SMA-immunolabeled brain vessels were quantified using the “Surface” and “Filament” tools in Imaris software (Oxford instruments, Oxford). Three brains were analyzed per developmental stage.

### Immunohistochemical analysis of human tissue samples

The specimens described here are part of the “Hôpitaux Universitaires de l’Est Parisien – Neuropathologie du développement” brain collection (biobank identification number: BB-0033-00082). Informed consent was obtained for brain autopsy and histological examination. Fetal brains were obtained from spontaneous or medical abortions. The fetuses did not display any significant brain disorders or diseases. The same technical procedures were applied to all brain samples: after removal, brains were fixed with formalin for 5–12 weeks. A macroscopic analysis enabled the samples to be selected and processed (paraffin embedding, preparation of 7-micron slices, and staining with hematein reagent) for histological analysis. Coronal slices (including the temporal telencephalic parenchyma and the hippocampus) were deparaffinized and unmasked in citrate buffer (pH 6.0). Expression of Myh11 or P-gP was detected using the Bond Polymer Refine Detection kit (Leica) and processed on automated immunostaining systems (the Bond RX Leica for P-gP and Myh11, and the LEICA BOND III for SMA). Pictures were acquired using a slide scanner (Lamina, Perkin Elmer).

Stained samples were analyzed using QuPath (Bankhead et al., 2017). For each sample, a QuPath “pixel classifier” was trained to discriminate between DAB-positive spots and the background. This “classifier” consisted in an artificial neural network based on four features: a Gaussian filter to select for the intensity, and three structure tensor eigenvalues to favor thin elongated objects. To train the classifier, we defined manually annotated spots and background area on one image per developmental stage. When the results were visually satisfactory, the trained pixel classifier was used to detect positive spots in manually defined regions of interest. For P-gP, the presence of false positives was handled by keeping only positive spots above 80 µm^2^ in area.

### VSMC contractility *ex vivo*

Mice were rapidly decapitated, and the brains were quickly removed and placed in cold (∼4°C) artificial cerebrospinal fluid (aCSF) solution containing 119 mM NaCl, 2.5 mM KCl, 2.5 mM CaCl_2_, 26.2 mM NaHCO_3_, 1 mM NaH_2_PO_4_, 1.3 mM MgSO_4_, 11 mM D-glucose (pH = 7.35). Brains were constantly oxygenated with 95% O_2_ –5% CO_2_. Brain cortex slices (400 µm thick) were cut with a vibratome (VT2000S, Leica) and transferred to a constantly oxygenated (95% O_2_–5% CO_2_) holding chamber containing aCSF. Subsequently, individual slices were placed in a submerged recording chamber maintained at RT under an upright microscope (Zeiss) equipped with a CCD camera (Qimaging) and perfused at 2 mL/min with oxygenated aCSF. Only one vessel per slice was selected for measurements of vascular responsiveness, at the junction between layers I and II of the somatosensory cortex and with a well-defined luminal diameter (10–15 µm). An image was acquired every 30 s. Each recording started with the establishment of a control baseline for 5 min. Vessels with an unstable baseline (i.e. a change in diameter of more than 5%) were discarded from analysis. Vasoconstriction was induced by the application of the thromboxane A_2_ receptor agonist U46619 (9,11-dideoxy-11a,9a-epoxymethanoprostaglandin F2α, 50 nM, Sigma) for 2 min. The signal was recorded until it had returned to the baseline.

Any drift in the images during the recording time was corrected either online (for z-drift) or off-line (for the x and y drift), using Image Pro Plus 7.0. To minimize the differences between two consecutive frames, images were manually repositioned using the subtraction tool in Image Pro Plus. Vasoconstriction was measured using a custom routine running in IgorPro (Wavemetrics).

### Analysis of radiotracer passage into the brain

#### Radioactive compounds

[^14^C]-Sucrose (20.1 GBq mmol^-1^), and [^3^H]-verapamil (3049 GBq mmol^-1^) were purchased from Perkin Elmer (Paris, France). [^3^H]-rosuvastatin (203.5 GBq mmol^-1^) was purchased by Moravek Inc (Brea, CA, USA).

#### Brain transport of [^3^H]-verapamil and [^3^H]-rosuvastatin in P15 and P30 mice

The permeation (Kin; µL s^-1^ g^-1^) of [^3^H]-verapamil and [^3^H]-rosuvastatin across the BBB in P15 and P30 mouse brains was determined with an integration plot analysis, as described previously (Patlak et al., 1983). Briefly, the mouse was injected intraperitoneally with saline (10 mL kg^-1^) containing [^3^H]-verapamil (∼0.074 MBq mL^−1^) or [^3^H]-rosuvastatin (∼0.130 MBq mL^−1^) and the vascular marker [^14^C]-sucrose (∼0.031 MBq mL^−1^). On P15 (n = 6-7 animals) and P30 (n = 6-7 animals), mice were decapitated at each selected time point (e.g. 2.5, 5 and 7 min) and blood samples were obtained. The brain was removed from the skull. For each mouse, the brain hemispheres and two blood samples were weighed in Econo® glass vials for liquid scintillation counting (Perkin Elmer). All samples were treated with a tissue solubilizer (Solvable^®^, Perkin Elmer). As recommended by the manufacturer (Perkin Elmer), blood samples were decolored with 0.2 mL 30% hydrogen peroxide solution so as to avoid color quench problems. After complete tissue digestion of the samples (24 h in a water bath at 45°C), the vials were cooled and mixed with Ultima-gold XR^®^ (Perkin Elmer). The radioactivity in each vial (^3^H and/or ^14^C disintegrations per minute (dpm)) was counted in Perkin Elmer Tri-Carb counter.

In the integration plot analysis, brain permeability was defined as an apparent brain distribution volume (Vd) corrected for the vascular space over the duration of the experiments. Vd(t) can be calculated as the dpm for the [^3^H]-compound per gram of brain tissue after correction for the amount of [^3^H]-compound in the vascular space (estimated from the [^14^C]-sucrose brain volume (Vv)). The brain clearance of the [^3^H]-compound (Kin, µL s^-1^ g^-1^) was determined by applying the following equation:

K_in_ = Vd(t). Cb(t) / AUC(t), where AUC(t) (dpm.min/µL) is the area under the blood concentration time curve of the [^3^H]-compound from time 0 to time t, and Cb(t) (dpm/µL) is the corresponding blood concentration.

#### BBB integrity in mice on P5, P15 and P30

Sucrose is a very hydrophilic, low-molecular-weight (342 Da) disaccharide that does not cross lipid membrane by passive diffusion and does not have a dedicated transporter. In this context, variations in the level of sucrose brain distribution volume (i.e. Vv) reflect changes in the BBB’s integrity/leakiness. The mouse Vv (in µL per gram) was calculated using the brain distribution of [^14^C]-sucrose (i.e. the level of [^14^C]-sucrose (dpm per gram) in the midbrain divided by the [^14^C]-sucrose blood concentration at 5 min), using the technique described above.

## Supplementary tables

**Table S1: Comparison of cortical MV transcriptomes on P5 and P15**. Selected mRNAs have a mean number of reads ≥ 50 in at least one condition. The fold-change (FC) between expression on P5 and P15 and associated p adjusted (padj) values are indicated. Base mean: mean reads for each transcript. N=3 libraries for each stage (related to Figure 1).

**Table S2: Gene ontology**

In cortical MVs, the “biological process” and “cellular component” GO pathways significantly changed between P5 and P15 (related to Figure 1).

**Table S3: Identification of vascular-cell-type-specific or -preferentially expressed transcripts**

Cell-type-specific or -preferentially expressed transcripts are listed for each cluster (related to Figure 2). *Left-hand columns*: identification of transcripts -preferentially expressed in or specific to each cell. % of single-cells within and outside each cluster (Vanlandewijck et al., 2018); logFC between these % and the associated adjusted P-value; RPKM in cells within the cluster. *Right-hand columns:* cortical MV RNA-Seq analysis. log2FC on P15 versus P5 and the associated adjusted P-values; RPKM on P5 or P15, and RPKM normalized against the RPKM in the single cell cluster. EC, endothelial cell; PC, pericyte; FB, fibroblast: VSMC, vascular smooth muscle cell.

**Table S4: Western blot, FISH, immunohistology and qPCR data**

Data on the observations and the associated statistical tests (for Figures 2-7)

**Table S5: List of resources**

## Supplementary videos

**Video S1, S2 and S3: Visualization of the SMA-positive vascular network** (related to Figure 7), generated by Imaris software. **S1** on P5; **S2** on P15; **S3** at P60. The videos show the meningeal arterial network, followed by the cortical penetrating arterioles and the intraparenchymal vascular network.

