## Supplementary Figures and Tables for "In mice and humans, the brain’s blood vessels mature postnatally to acquire barrier and contractile properties"

Supplementary Table S1

| Ensembl reference name | id | Gene symbol | Description | baseMean P15 | baseMean P5 | log2FoldChange | padj |
|---|---|---|---|---|---|---|---|
| ENSMUSG00000025903 | GSMG0000005 | Lypla1 | lysophospholipase 1 | 778.59 | 884.62 | -0.18 | 4.59E-01 |
| ENSMUSG00000033793 | GSMG0000009 | Atp6v1h | ATPase, H+ transporting, lysosomal V1 subunit H | 701.66 | 812.36 | -0.21 | 3.92E-01 |
| ENSMUSG00000025907 | GSMG0000012 | Rb1cc1 | RB1-inducible coiled-coil 1 | 475.71 | 385.32 | 0.30 | 1.91E-01 |
| ENSMUSG00000051285 | GSMG0000016 | Pcmtd1 | protein-L-isoaspartate (D-aspartate) O-methyltransferase domain contai | 879.28 | 608.16 | 0.53 | 9.44E-03 |
| NULL | GSMG0000017 | NULL | NULL | 456.42 | 268.78 | 0.76 | 2.40E-04 |
| ENSMUSG00000061024 | GSMG0000023 | Rrs1 | ribosome biogenesis regulator 1 | 618.37 | 907.43 | -0.55 | 3.12E-03 |
| ENSMUSG00000025911 | GSMG0000024 | Adhfe1 | alcohol dehydrogenase, iron containing, 1 | 178.26 | 261.39 | -0.55 | 4.59E-02 |
| ENSMUSG00000025915 | GSMG0000027 | Sgk3 | serum/glucocorticoid regulated kinase 3 | 571.46 | 783.11 | -0.46 | 3.36E-02 |
| ENSMUSG00000056763 | GSMG0000030 | Cspp1 | centrosome and spindle pole associated protein 1 | 338.49 | 333.35 | 0.02 | 9.63E-01 |
| ENSMUSG00000048960 | GSMG0000033 | Prex2 | phosphatidylinositol-3,4,5-trisphosphate-dependent Rac exchange facto | 176.25 | 154.44 | 0.19 | 6.22E-01 |
| ENSMUSG00000057715 | GSMG0000035 | A830018L16Rik | RIKEN cDNA A830018L16 gene | 172.27 | 236.37 | -0.46 | 7.70E-02 |
| ENSMUSG00000092083 | GSMG0000049 | Kcnb2 | potassium voltage gated channel, Shab-related subfamily, member 2 | 59.90 | 108.90 | -0.87 | 1.15E-02 |
| NULL | GSMG0000050 | NULL | NULL | 128.80 | 159.44 | -0.30 | 5.23E-01 |
| NULL | GSMG0000051 | NULL | NULL | 179.07 | 190.59 | -0.09 | 8.34E-01 |
| NULL | GSMG0000052 | NULL | NULL | 109.86 | 105.14 | 0.07 | 9.03E-01 |
| ENSMUSG00000025925 | GSMG0000057 | Terf1 | telomeric repeat binding factor 1 | 404.27 | 407.15 | -0.01 | 9.83E-01 |
| ENSMUSG00000025921 | GSMG0000059 | Rdh10 | retinol dehydrogenase 10 (all-trans) | 271.83 | 244.86 | 0.16 | 6.67E-01 |
| ENSMUSG00000025940 | GSMG0000062 | Tmem70 | transmembrane protein 70 | 707.15 | 726.28 | -0.04 | 9.16E-01 |
| ENSMUSG00000025779 | GSMG0000063 | Ly96 | lymphocyte antigen 96 | 96.59 | 91.89 | 0.07 | 8.96E-01 |
| ENSMUSG00000025777 | GSMG0000066 | Gdap1 | ganglioside-induced differentiation-associated-protein 1 | 237.81 | 298.94 | -0.33 | 1.99E-01 |
| ENSMUSG00000025776 | GSMG0000069 | Crispld1 | cysteine-rich secretory protein LCCL domain containing 1 | 407.59 | 90.05 | 2.18 | 2.43E-18 |
| ENSMUSG00000025931 | GSMG0000077 | Paqr8 | progestin and adipoQ receptor family member VIII | 1810.58 | 2222.36 | -0.30 | 8.33E-02 |
| ENSMUSG00000025933 | GSMG0000081 | Tmem14a | transmembrane protein 14A | 246.19 | 119.49 | 1.04 | 6.55E-05 |
| ENSMUSG00000026156 | GSMG0000091 | B3gat2 | beta-1,3-glucuronyltransferase 2 (glucuronosyltransferase S) | 707.41 | 683.19 | 0.05 | 8.70E-01 |
| ENSMUSG00000073725 | GSMG0000097 | Lmbrd1 | LMBR1 domain containing 1 | 2526.82 | 2259.77 | 0.16 | 4.28E-01 |
| NULL | GSMG0000102 | NULL | NULL | 172.96 | 193.93 | -0.17 | 6.42E-01 |
| ENSMUSG00000026058 | GSMG0000107 | Khdrbs2 | KH domain containing, RNA binding, signal transduction associated 2 | 383.75 | 937.73 | -1.29 | 5.40E-16 |
| NULL | GSMG0000109 | NULL | NULL | 121.42 | 208.23 | -0.78 | 1.98E-03 |
| ENSMUSG00000004768 | GSMG0000116 | Rab23 | RAB23, member RAS oncogene family | 170.59 | 202.52 | -0.25 | 4.32E-01 |
| ENSMUSG00000026131 | GSMG0000117 | Dst | dystonin | 6417.42 | 5942.23 | 0.11 | 6.26E-01 |
| NULL | GSMG0000119 | NULL | NULL | 437.57 | 349.17 | 0.33 | 1.72E-01 |
| ENSMUSG00000026127 | GSMG0000122 | Imp4 | IMP4, U3 small nucleolar ribonucleoprotein | 687.35 | 812.57 | -0.24 | 3.48E-01 |

| | | | | | | | |
|---|---|---|---|---|---|---|---|
| ENSMUSG00000026126 | GSMG0000124 | Ptpn18 | protein tyrosine phosphatase, non-receptor type 18 | 95.15 | 137.35 | -0.54 | 1.40E-01 |
| ENSMUSG00000045174 | GSMG0000127 | Amer3 | APC membrane recruitment 3 | 278.71 | 540.13 | -0.96 | 1.56E-03 |
| ENSMUSG00000026123 | GSMG0000132 | Plekhb2 | pleckstrin homology domain containing, family B (evectins) member 2 | 322.66 | 350.99 | -0.12 | 7.28E-01 |
| ENSMUSG00000045216 | GSMG0000135 | Hs6st1 | heparan sulfate 6-O-sulfotransferase 1 | 776.38 | 937.29 | -0.27 | 4.70E-01 |
| ENSMUSG00000037447 | GSMG0000137 | Arid5a | AT rich interactive domain 5A (MRF1-like) | 332.96 | 131.09 | 1.35 | 5.08E-09 |
| ENSMUSG00000001138 | GSMG0000143 | Cnnm3 | cyclin M3 | 289.03 | 174.51 | 0.73 | 5.43E-02 |
| ENSMUSG00000026117 | GSMG0000148 | Zap70 | zeta-chain (TCR) associated protein kinase | 124.32 | 209.32 | -0.76 | 1.69E-02 |
| ENSMUSG00000026113 | GSMG0000152 | Inpp4a | inositol polyphosphate-4-phosphatase, type I | 508.68 | 690.17 | -0.44 | 1.39E-01 |
| ENSMUSG00000026111 | GSMG0000155 | Unc50 | unc-50 homolog (C. elegans) | 1845.14 | 1758.50 | 0.07 | 7.85E-01 |
| ENSMUSG00000026087 | GSMG0000159 | Mrpl30 | mitochondrial ribosomal protein L30 | 447.23 | 608.72 | -0.44 | 1.72E-02 |
| ENSMUSG00000026083 | GSMG0000161 | Eif5b | eukaryotic translation initiation factor 5B | 1803.24 | 2162.21 | -0.26 | 3.82E-01 |
| ENSMUSG00000026078 | GSMG0000168 | Pdcl3 | phosducin-like 3 | 1043.26 | 1172.24 | -0.17 | 5.12E-01 |
| NULL | GSMG0000169 | NULL | NULL | 226.73 | 226.05 | 0.00 | 9.95E-01 |
| ENSMUSG00000026077 | GSMG0000170 | Npas2 | neuronal PAS domain protein 2 | 377.10 | 368.26 | 0.03 | 9.29E-01 |
| ENSMUSG00000003135 | GSMG0000172 | Cnot11 | CCR4-NOT transcription complex, subunit 11 | 359.16 | 440.57 | -0.29 | 1.94E-01 |
| ENSMUSG00000026074 | GSMG0000174 | Map4k4 | mitogen-activated protein kinase kinase kinase kinase 4 | 2400.12 | 2645.02 | -0.14 | 6.64E-01 |
| ENSMUSG00000026072 | GSMG0000179 | Il1r1 | interleukin 1 receptor, type I | 413.28 | 359.78 | 0.20 | 4.80E-01 |
| ENSMUSG00000026062 | GSMG0000186 | Slc9a2 | solute carrier family 9 (sodium/hydrogen exchanger), member 2 | 198.48 | 82.52 | 1.27 | 4.48E-05 |
| NULL | GSMG0000193 | NULL | NULL | 276.34 | 578.36 | -1.07 | 3.90E-07 |
| NULL | GSMG0000194 | 2900092D14Rik | RIKEN cDNA 2900092D14 gene | 149.22 | 360.44 | -1.28 | 1.54E-07 |
| ENSMUSG00000060679 | GSMG0000196 | Mrps9 | mitochondrial ribosomal protein S9 | 508.74 | 531.61 | -0.06 | 8.53E-01 |
| ENSMUSG00000010290 | GSMG0000200 | AI597479 | expressed sequence AI597479 | 155.55 | 134.55 | 0.21 | 5.59E-01 |
| ENSMUSG00000066877 | GSMG0000203 | Nck2 | non-catalytic region of tyrosine kinase adaptor protein 2 | 1035.49 | 1239.87 | -0.26 | 1.73E-01 |
| NULL | GSMG0000204 | NULL | NULL | 307.33 | 324.22 | -0.08 | 8.43E-01 |
| NULL | GSMG0000208 | NULL | NULL | 129.52 | 180.74 | -0.48 | 1.03E-01 |
| ENSMUSG00000026051 | GSMG0000209 | 1500015O10Rik | RIKEN cDNA 1500015O10 gene | 51.03 | 215.52 | -2.08 | 6.30E-02 |
| ENSMUSG00000041763 | GSMG0000210 | Tpp2 | tripeptidyl peptidase II | 576.40 | 650.65 | -0.17 | 4.42E-01 |
| ENSMUSG00000041684 | GSMG0000211 | Bivm | basic, immunoglobulin-like variable motif containing | 228.86 | 242.15 | -0.08 | 8.50E-01 |
| ENSMUSG00000026048 | GSMG0000212 | Ercc5 | excision repair cross-complementing rodent repair deficiency, complem | 1271.98 | 1266.60 | 0.01 | 9.81E-01 |
| ENSMUSG00000056870 | GSMG0000214 | Gulp1 | GULP, engulfment adaptor PTB domain containing 1 | 230.01 | 187.90 | 0.29 | 4.71E-01 |
| ENSMUSG00000026043 | GSMG0000219 | Col3a1 | collagen, type III, alpha 1 | 9085.97 | 5712.87 | 0.67 | 1.93E-04 |
| ENSMUSG00000025995 | GSMG0000221 | Wdr75 | WD repeat domain 75 | 509.72 | 757.32 | -0.57 | 3.02E-03 |
| ENSMUSG00000026109 | GSMG0000229 | Tmeff2 | transmembrane protein with EGF-like and two follistatin-like domains 2 | 1008.95 | 705.69 | 0.52 | 3.50E-02 |

| | | | | | | | |
|---|---|---|---|---|---|---|---|
| NULL | GSMG0000234 | NULL | NULL | 159.81 | 122.95 | 0.38 | 2.54E-01 |
| ENSMUSG00000045954 | GSMG0000236 | Sdpr | serum deprivation response | 4353.86 | 2278.21 | 0.93 | 1.72E-05 |
| ENSMUSG00000026104 | GSMG0000238 | Stat1 | signal transducer and activator of transcription 1 | 1368.66 | 1830.10 | -0.42 | 1.09E-01 |
| ENSMUSG00000043015 | GSMG0000243 | Tmem194b | transmembrane protein 194B | 53.87 | 56.91 | -0.07 | 9.21E-01 |
| ENSMUSG00000041426 | GSMG0000244 | Hibch | 3-hydroxyisobutyryl-Coenzyme A hydrolase | 224.53 | 276.33 | -0.29 | 3.46E-01 |
| ENSMUSG00000026097 | GSMG0000248 | Ormdl1 | ORM1-like 1 (S. cerevisiae) | 372.45 | 379.42 | -0.02 | 9.56E-01 |
| ENSMUSG00000026096 | GSMG0000249 | Osgepl1 | O-sialoglycoprotein endopeptidase-like 1 | 470.69 | 409.70 | 0.20 | 4.73E-01 |
| ENSMUSG00000025981 | GSMG0000253 | Coq10b | coenzyme Q10B | 706.86 | 439.73 | 0.69 | 7.45E-04 |
| ENSMUSG00000073676 | GSMG0000255 | Hspe1 | heat shock protein 1 (chaperonin 10) | 393.97 | 737.31 | -0.91 | 3.44E-05 |
| ENSMUSG00000025979 | GSMG0000256 | Mob4 | MOB family member 4, phocein | 1349.06 | 1475.15 | -0.13 | 6.48E-01 |
| ENSMUSG00000038349 | GSMG0000262 | Plcl1 | phospholipase C-like 1 | 766.66 | 979.78 | -0.36 | 1.21E-01 |
| NULL | GSMG0000264 | NULL | NULL | 430.97 | 273.19 | 0.65 | 7.98E-03 |
| ENSMUSG00000025971 | GSMG0000277 | 9430016H08Rik | RIKEN cDNA 9430016H08 gene | 501.80 | 735.87 | -0.55 | 2.77E-03 |
| ENSMUSG00000038305 | GSMG0000278 | Spats2l | spermatogenesis associated, serine-rich 2-like | 358.07 | 412.58 | -0.20 | 4.12E-01 |
| NULL | GSMG0000280 | NULL | NULL | 68.18 | 59.42 | 0.19 | 7.33E-01 |
| ENSMUSG00000051223 | GSMG0000287 | Bzw1 | basic leucine zipper and W2 domains 1 | 4005.30 | 5724.68 | -0.52 | 2.08E-03 |
| ENSMUSG00000026036 | GSMG0000288 | Nif3l1 | Ngg1 interacting factor 3-like 1 (S. pombe) | 348.40 | 469.61 | -0.43 | 7.39E-02 |
| ENSMUSG00000026032 | GSMG0000290 | Ndufb3 | NADH dehydrogenase (ubiquinone) 1 beta subcomplex 3 | 1300.72 | 1373.82 | -0.08 | 8.08E-01 |
| ENSMUSG00000026029 | GSMG0000294 | Casp8 | caspase 8 | 2186.77 | 2495.34 | -0.19 | 4.10E-01 |
| ENSMUSG00000026027 | GSMG0000296 | Stradb | STE20-related kinase adaptor beta | 288.34 | 383.03 | -0.41 | 1.06E-01 |
| ENSMUSG00000026020 | GSMG0000304 | Nop58 | NOP58 ribonucleoprotein | 757.19 | 1296.03 | -0.78 | 5.18E-05 |
| ENSMUSG00000067336 | GSMG0000306 | Bmpr2 | bone morphogenetic protein receptor, type II (serine/threonine kinase) | 698.53 | 443.76 | 0.66 | 7.75E-04 |
| ENSMUSG00000041040 | GSMG0000310 | Fam117b | family with sequence similarity 117, member B | 360.31 | 281.05 | 0.36 | 2.02E-01 |
| ENSMUSG00000073664 | GSMG0000315 | Nbeal1 | neurobeachin like 1 | 538.54 | 478.06 | 0.17 | 5.87E-01 |
| NULL | GSMG0000318 | NULL | NULL | 215.00 | 146.73 | 0.55 | 1.55E-01 |
| ENSMUSG00000049439 | GSMG0000319 | Cyp20a1 | cytochrome P450, family 20, subfamily a, polypeptide 1 | 406.05 | 571.55 | -0.49 | 2.84E-02 |
| ENSMUSG00000026782 | GSMG0000320 | Abi2 | abl-interactor 2 | 616.14 | 841.95 | -0.45 | 1.32E-02 |
| ENSMUSG00000052062 | GSMG0000336 | Pard3b | par-3 family cell polarity regulator beta | 180.89 | 158.31 | 0.20 | 5.99E-01 |
| ENSMUSG00000025969 | GSMG0000350 | Nrp2 | neuropilin 2 | 1287.81 | 1400.95 | -0.12 | 7.10E-01 |
| ENSMUSG00000084799 | GSMG0000356 | Ino80dos | INO80 complex subunit D, opposite strand | 177.13 | 174.16 | 0.02 | 9.65E-01 |
| ENSMUSG00000025967 | GSMG0000359 | Eef1b2 | eukaryotic translation elongation factor 1 beta 2 | 1598.22 | 2241.39 | -0.49 | 1.85E-03 |
| ENSMUSG00000025964 | GSMG0000362 | Adam23 | a disintegrin and metallopeptidase domain 23 | 1022.91 | 1352.36 | -0.40 | 7.97E-02 |
| ENSMUSG00000025962 | GSMG0000363 | Fastkd2 | FAST kinase domains 2 | 513.80 | 558.72 | -0.12 | 6.95E-01 |

| | | | | | | | |
|---|---|---|---|---|---|---|---|
| ENSMUSG00000025958 | GSMG0000365 | Creb1 | cAMP responsive element binding protein 1 | 269.77 | 267.90 | 0.01 | 9.73E-01 |
| ENSMUSG00000025949 | GSMG0000374 | Pikfyve | phosphoinositide kinase, FYVE finger containing | 225.18 | 228.64 | -0.02 | 9.62E-01 |
| ENSMUSG00000015222 | GSMG0000378 | Map2 | microtubule-associated protein 2 | 452.29 | 869.20 | -0.94 | 8.78E-08 |
| NULL | GSMG0000380 | NULL | NULL | 80.03 | 145.13 | -0.85 | 9.98E-03 |
| ENSMUSG00000055567 | GSMG0000381 | Unc80 | unc-80, NALCN activator | 135.16 | 58.63 | 1.21 | 7.06E-04 |
| ENSMUSG00000026005 | GSMG0000384 | Rpe | ribulose-5-phosphate-3-epimerase | 840.96 | 622.92 | 0.43 | 1.08E-01 |
| ENSMUSG00000026192 | GSMG0000398 | Atic | 5-aminoimidazole-4-carboxamide ribonucleotide formyltransferase/IMI | 110.83 | 269.55 | -1.29 | 5.19E-08 |
| ENSMUSG00000026188 | GSMG0000401 | Tmem169 | transmembrane protein 169 | 247.97 | 335.78 | -0.44 | 1.18E-01 |
| ENSMUSG00000026187 | GSMG0000402 | Xrcc5 | X-ray repair complementing defective repair in Chinese hamster cells 5 | 289.53 | 465.79 | -0.69 | 8.58E-04 |
| ENSMUSG00000039354 | GSMG0000405 | Smarcal1 | SWI/SNF related matrix associated, actin dependent regulator of chrom | 695.87 | 830.03 | -0.25 | 2.03E-01 |
| ENSMUSG00000039323 | GSMG0000409 | Igfbp2 | insulin-like growth factor binding protein 2 | 3214.29 | 3962.19 | -0.30 | 2.65E-01 |
| ENSMUSG00000006304 | GSMG0000417 | Arpc2 | actin related protein 2/3 complex, subunit 2 | 4916.00 | 6317.22 | -0.36 | 2.40E-02 |
| NULL | GSMG0000418 | NULL | NULL | 590.54 | 471.81 | 0.32 | 3.65E-01 |
| ENSMUSG00000026179 | GSMG0000420 | Pnkd | paroxysmal nonkinesiogenic dyskinesia | 7556.47 | 6328.70 | 0.26 | 1.04E-01 |
| ENSMUSG00000026176 | GSMG0000426 | Ctdsp1 | CTD (carboxy-terminal domain, RNA polymerase II, polypeptide A) sn | 103.42 | 72.76 | 0.51 | 2.00E-01 |
| ENSMUSG00000026175 | GSMG0000427 | Vil1 | villin 1 | 248.59 | 230.36 | 0.11 | 8.36E-01 |
| ENSMUSG00000026174 | GSMG0000428 | Cnot9 | CCR4-NOT transcription complex, subunit 9 | 319.76 | 426.03 | -0.41 | 5.62E-02 |
| ENSMUSG00000026173 | GSMG0000429 | Plcd4 | phospholipase C, delta 4 | 276.60 | 123.95 | 1.16 | 3.94E-05 |
| ENSMUSG00000026172 | GSMG0000430 | Bcs1l | BCS1-like (yeast) | 227.33 | 269.84 | -0.24 | 5.42E-01 |
| ENSMUSG00000033276 | GSMG0000431 | Stk36 | serine/threonine kinase 36 | 98.84 | 191.73 | -0.96 | 3.30E-03 |
| ENSMUSG00000033257 | GSMG0000432 | Ttll4 | tubulin tyrosine ligase-like family, member 4 | 131.87 | 199.74 | -0.60 | 3.37E-02 |
| ENSMUSG00000026170 | GSMG0000433 | Cyp27a1 | cytochrome P450, family 27, subfamily a, polypeptide 1 | 169.23 | 51.96 | 1.70 | 1.97E-08 |
| NULL | GSMG0000437 | NULL | NULL | 180.22 | 131.48 | 0.45 | 1.42E-01 |
| ENSMUSG00000049339 | GSMG0000440 | Fam134a | family with sequence similarity 134, member A | 1553.60 | 1623.93 | -0.06 | 8.24E-01 |
| ENSMUSG00000026197 | GSMG0000441 | Zfand2b | zinc finger, AN1 type domain 2B | 822.96 | 687.15 | 0.26 | 2.06E-01 |
| ENSMUSG00000026199 | GSMG0000442 | Ankzf1 | ankyrin repeat and zinc finger domain containing 1 | 296.81 | 215.93 | 0.46 | 5.33E-02 |
| ENSMUSG00000026201 | GSMG0000443 | Stk16 | serine/threonine kinase 16 | 901.33 | 854.29 | 0.08 | 7.92E-01 |
| ENSMUSG00000026203 | GSMG0000444 | Dnajb2 | DnaJ heat shock protein family (Hsp40) member B2 | 738.79 | 606.71 | 0.29 | 1.57E-01 |
| ENSMUSG00000026208 | GSMG0000445 | Des | desmin | 895.86 | 547.95 | 0.71 | 1.11E-04 |
| ENSMUSG00000026207 | GSMG0000446 | Speg | SPEG complex locus | 1226.77 | 966.89 | 0.34 | 1.09E-01 |
| ENSMUSG00000033021 | GSMG0000447 | Gmppa | GDP-mannose pyrophosphorylase A | 1825.71 | 1609.31 | 0.18 | 4.05E-01 |
| ENSMUSG00000051703 | GSMG0000449 | Tmem198 | transmembrane protein 198 | 213.16 | 247.58 | -0.22 | 5.59E-01 |
| ENSMUSG00000032968 | GSMG0000450 | Inha | inhibin alpha | 1671.37 | 1688.33 | -0.01 | 9.68E-01 |

| | | | | | | | |
|---|---|---|---|---|---|---|---|
| ENSMUSG00000026213 | GSMG0000451 | Stk11ip | serine/threonine kinase 11 interacting protein | 1125.65 | 863.50 | 0.38 | 8.45E-02 |
| ENSMUSG00000006576 | GSMG0000452 | Slc4a3 | solute carrier family 4 (anion exchanger), member 3 | 5378.22 | 4772.46 | 0.17 | 6.20E-01 |
| ENSMUSG00000032908 | GSMG0000457 | Sgpp2 | sphingosine-1-phosphate phosphotase 2 | 1345.81 | 161.43 | 3.06 | 1.16E-48 |
| G00000032883,ENSMUSG000 | GSMG0000461 | Acsl3 // Utp14b | acyl-CoA synthetase long-chain family member 3 // UTP14B small subu | 2755.70 | 2722.17 | 0.02 | 9.54E-01 |
| ENSMUSG00000047330 | GSMG0000462 | Kcne4 | potassium voltage-gated channel, Isk-related subfamily, gene 4 | 569.60 | 370.59 | 0.62 | 4.87E-03 |
| ENSMUSG00000026248 | GSMG0000464 | Mrpl44 | mitochondrial ribosomal protein L44 | 347.62 | 411.86 | -0.25 | 3.33E-01 |
| ENSMUSG00000054976 | GSMG0000467 | Nyap2 | neuronal tyrosine-phophorylated phosphoinositide 3-kinase adaptor 2 | 240.45 | 266.72 | -0.14 | 7.66E-01 |
| ENSMUSG00000026142 | GSMG0000471 | Rhbdd1 | rhomboid domain containing 1 | 330.39 | 240.21 | 0.46 | 5.78E-02 |
| ENSMUSG00000026150 | GSMG0000475 | Mff | mitochondrial fission factor | 5908.70 | 7443.45 | -0.33 | 2.60E-02 |
| ENSMUSG00000026159 | GSMG0000478 | Agfg1 | ArfGAP with FG repeats 1 | 1576.92 | 1121.07 | 0.49 | 4.24E-03 |
| ENSMUSG00000026222 | GSMG0000490 | Sp100 | nuclear antigen Sp100 | 380.74 | 129.57 | 1.56 | 2.21E-11 |
| ENSMUSG00000036707 | GSMG0000493 | Cab39 | calcium binding protein 39 | 1301.25 | 1366.11 | -0.07 | 8.08E-01 |
| ENSMUSG00000026223 | GSMG0000496 | Itm2c | integral membrane protein 2C | 4554.45 | 3201.32 | 0.51 | 2.54E-03 |
| ENSMUSG00000026227 | GSMG0000499 | 2810459M11Rik | RIKEN cDNA 2810459M11 gene | 70.77 | 107.17 | -0.60 | 8.44E-02 |
| ENSMUSG00000026229 | GSMG0000500 | Psmd1 | proteasome (prosome, macropain) 26S subunit, non-ATPase, 1 | 1094.82 | 1249.09 | -0.19 | 4.32E-01 |
| ENSMUSG00000062590 | GSMG0000502 | Armc9 | armadillo repeat containing 9 | 1185.20 | 1021.32 | 0.21 | 3.28E-01 |
| ENSMUSG00000026240 | GSMG0000510 | Cops7b | COP9 signalosome subunit 7B | 938.78 | 1080.40 | -0.20 | 3.18E-01 |
| ENSMUSG00000053333 | GSMG0000512 | Dis3l2 | DIS3 like 3'-5' exoribonuclease 2 | 1098.79 | 1249.63 | -0.18 | 3.96E-01 |
| ENSMUSG00000026254 | GSMG0000526 | Eif4e2 | eukaryotic translation initiation factor 4E member 2 | 929.93 | 1205.00 | -0.37 | 4.69E-02 |
| ENSMUSG00000048000 | GSMG0000528 | Gigyf2 | GRB10 interacting GYF protein 2 | 1006.93 | 1191.55 | -0.24 | 2.09E-01 |
| ENSMUSG00000026288 | GSMG0000533 | Inpp5d | inositol polyphosphate-5-phosphatase D | 764.72 | 770.63 | -0.01 | 9.73E-01 |
| ENSMUSG00000026289 | GSMG0000535 | Atg16l1 | autophagy related 16-like 1 (S. cerevisiae) | 1503.95 | 1481.81 | 0.02 | 9.44E-01 |
| NULL | GSMG0000539 | NULL | NULL | 263.35 | 226.28 | 0.22 | 4.39E-01 |
| ENSMUSG00000070738 | GSMG0000540 | Dgkd | diacylglycerol kinase, delta | 296.73 | 419.47 | -0.50 | 4.19E-02 |
| ENSMUSG00000036206 | GSMG0000550 | Sh3bp4 | SH3-domain binding protein 4 | 3678.34 | 4135.70 | -0.17 | 3.54E-01 |
| NULL | GSMG0000551 | NULL | NULL | 215.97 | 167.12 | 0.37 | 2.18E-01 |
| ENSMUSG00000055013 | GSMG0000552 | Agap1 | ArfGAP with GTPase domain, ankyrin repeat and PH domain 1 | 725.79 | 789.03 | -0.12 | 7.42E-01 |
| ENSMUSG00000044337 | GSMG0000561 | Ackr3 | atypical chemokine receptor 3 | 7112.41 | 1421.76 | 2.32 | 1.02E-34 |
| ENSMUSG00000034432 | GSMG0000565 | Cops8 | COP9 signalosome subunit 8 | 2402.30 | 2652.96 | -0.14 | 4.77E-01 |
| ENSMUSG00000026303 | GSMG0000568 | Mlph | melanophilin | 272.05 | 219.52 | 0.31 | 2.98E-01 |
| ENSMUSG00000026305 | GSMG0000569 | Lrrfip1 | leucine rich repeat (in FLII) interacting protein 1 | 1249.82 | 1098.12 | 0.19 | 4.05E-01 |
| ENSMUSG00000034353 | GSMG0000571 | Ramp1 | receptor (calcitonin) activity modifying protein 1 | 1323.61 | 654.89 | 1.02 | 1.71E-09 |
| NULL | GSMG0000574 | NULL | NULL | 214.43 | 55.27 | 1.96 | 6.49E-09 |

| | | | | | | | |
|---|---|---|---|---|---|---|---|
| ENSMUSG00000034343 | GSMG0000575 | Ube2f | ubiquitin-conjugating enzyme E2F (putative) | 218.67 | 238.63 | -0.13 | 7.37E-01 |
| ENSMUSG00000026307 | GSMG0000576 | Scly | selenocysteine lyase | 495.40 | 467.21 | 0.08 | 7.92E-01 |
| ENSMUSG00000034292 | GSMG0000581 | Traf3ip1 | TRAF3 interacting protein 1 | 100.25 | 106.63 | -0.08 | 8.69E-01 |
| ENSMUSG00000026311 | GSMG0000582 | Asb1 | ankyrin repeat and SOCS box-containing 1 | 79.26 | 47.63 | 0.73 | 1.09E-01 |
| ENSMUSG00000034220 | GSMG0000592 | Gpc1 | glypican 1 | 397.99 | 739.41 | -0.90 | 4.68E-05 |
| ENSMUSG00000047067 | GSMG0000594 | Dusp28 | dual specificity phosphatase 28 | 192.37 | 163.98 | 0.23 | 6.01E-01 |
| ENSMUSG00000026270 | GSMG0000596 | Capn10 | calpain 10 | 353.30 | 376.22 | -0.09 | 7.76E-01 |
| ENSMUSG00000047793 | GSMG0000602 | Sned1 | sushi, nidogen and EGF-like domains 1 | 620.06 | 381.16 | 0.70 | 2.89E-01 |
| ENSMUSG00000026275 | GSMG0000604 | Ppp1r7 | protein phosphatase 1, regulatory (inhibitor) subunit 7 | 596.37 | 734.33 | -0.30 | 2.55E-01 |
| ENSMUSG00000026276 | GSMG0000607 | Sept2 | septin 2 | 3251.71 | 3522.55 | -0.12 | 6.30E-01 |
| ENSMUSG00000026278 | GSMG0000611 | Bok | BCL2-related ovarian killer | 1467.62 | 2468.08 | -0.75 | 2.35E-07 |
| NULL | GSMG0000612 | NULL | NULL | 200.60 | 295.29 | -0.56 | 2.84E-02 |
| ENSMUSG00000026280 | GSMG0000613 | Atg4b | autophagy related 4B, cysteine peptidase | 1116.43 | 1213.11 | -0.12 | 6.51E-01 |
| NULL | GSMG0000614 | NULL | NULL | 210.32 | 286.75 | -0.44 | 8.72E-02 |
| ENSMUSG00000026283 | GSMG0000615 | Ing5 | inhibitor of growth family, member 5 | 727.97 | 622.14 | 0.23 | 2.76E-01 |
| ENSMUSG00000073609 | GSMG0000616 | D2hgdh | D-2-hydroxyglutarate dehydrogenase | 211.21 | 186.95 | 0.18 | 5.91E-01 |
| ENSMUSG00000051185 | GSMG0000624 | Fam174a | family with sequence similarity 174, member A | 1968.67 | 1405.77 | 0.49 | 7.67E-03 |
| ENSMUSG00000026333 | GSMG0000629 | Gin1 | gypsy retrotransposon integrase 1 | 200.80 | 196.17 | 0.03 | 9.39E-01 |
| ENSMUSG00000026319 | GSMG0000647 | 2310035C23Rik | RIKEN cDNA 2310035C23 gene | 659.40 | 746.63 | -0.18 | 4.72E-01 |
| ENSMUSG00000038866 | GSMG0000651 | Zcchc2 | zinc finger, CCHC domain containing 2 | 60.43 | 52.03 | 0.21 | 7.84E-01 |
| ENSMUSG00000044340 | GSMG0000652 | Phlpp1 | PH domain and leucine rich repeat protein phosphatase 1 | 210.37 | 143.68 | 0.55 | 1.53E-01 |
| ENSMUSG00000026377 | GSMG0000676 | Nifk | nucleolar protein interacting with the FHA domain of MKI67 | 430.37 | 620.90 | -0.53 | 1.07E-02 |
| ENSMUSG00000064302 | GSMG0000678 | Clasp1 | CLIP associating protein 1 | 1029.67 | 1452.24 | -0.50 | 6.10E-03 |
| ENSMUSG00000026339 | GSMG0000698 | Ccdc93 | coiled-coil domain containing 93 | 309.69 | 356.01 | -0.20 | 4.55E-01 |
| ENSMUSG00000026342 | GSMG0000705 | Slc35f5 | solute carrier family 35, member F5 | 1281.44 | 1236.51 | 0.05 | 8.63E-01 |
| ENSMUSG00000026343 | GSMG0000706 | Gpr39 | G protein-coupled receptor 39 | 467.95 | 430.23 | 0.12 | 6.91E-01 |
| ENSMUSG00000036155 | GSMG0000707 | Mgat5 | mannoside acetylglucosaminyltransferase 5 | 288.34 | 276.83 | 0.06 | 8.78E-01 |
| NULL | GSMG0000713 | NULL | NULL | 520.70 | 547.62 | -0.07 | 8.19E-01 |
| ENSMUSG00000026349 | GSMG0000715 | Ccnt2 | cyclin T2 | 941.92 | 790.13 | 0.25 | 1.93E-01 |
| ENSMUSG00000036104 | GSMG0000716 | Rab3gap1 | RAB3 GTPase activating protein subunit 1 | 442.04 | 370.16 | 0.26 | 3.52E-01 |
| ENSMUSG00000056211 | GSMG0000723 | R3hdm1 | R3H domain containing 1 | 1879.00 | 1221.77 | 0.62 | 4.52E-05 |
| ENSMUSG00000026353 | GSMG0000725 | Ubxn4 | UBX domain protein 4 | 900.76 | 994.18 | -0.14 | 6.04E-01 |
| ENSMUSG00000046404 | GSMG0000740 | Yod1 | YOD1 deubiquitinase | 661.21 | 556.62 | 0.25 | 3.21E-01 |

| | | | | | | | |
|---|---|---|---|---|---|---|---|
| NULL | GSMG0000741 | NULL | NULL | 293.20 | 332.14 | -0.18 | 6.91E-01 |
| ENSMUSG00000026427 | GSMG0000749 | Eif2d | eukaryotic translation initiation factor 2D | 1309.44 | 1288.04 | 0.02 | 9.32E-01 |
| NULL | GSMG0000750 | NULL | NULL | 652.41 | 851.37 | -0.38 | 3.16E-02 |
| ENSMUSG00000013275 | GSMG0000757 | Slc41a1 | solute carrier family 41, member 1 | 2478.59 | 2635.24 | -0.09 | 7.58E-01 |
| ENSMUSG00000026433 | GSMG0000758 | Rab29 | RAB29, member RAS oncogene family | 498.48 | 381.62 | 0.39 | 8.68E-02 |
| ENSMUSG00000026434 | GSMG0000759 | Nucks1 | nuclear casein kinase and cyclin-dependent kinase substrate 1 | 362.67 | 598.82 | -0.73 | 3.50E-03 |
| ENSMUSG00000026436 | GSMG0000762 | Elk4 | ELK4, member of ETS oncogene family | 749.42 | 794.43 | -0.08 | 7.58E-01 |
| ENSMUSG00000009772 | GSMG0000768 | Nuak2 | NUAK family, SNF1-like kinase, 2 | 59.62 | 43.69 | 0.46 | 4.42E-01 |
| ENSMUSG00000042046 | GSMG0000770 | Dstyk | dual serine/threonine and tyrosine protein kinase | 661.87 | 653.79 | 0.02 | 9.62E-01 |
| ENSMUSG00000026439 | GSMG0000772 | Rbbp5 | retinoblastoma binding protein 5 | 445.37 | 523.51 | -0.23 | 3.40E-01 |
| ENSMUSG00000026443 | GSMG0000776 | Lrrn2 | leucine rich repeat protein 2, neuronal | 1024.30 | 1760.56 | -0.78 | 1.65E-05 |
| ENSMUSG00000026447 | GSMG0000777 | Pik3c2b | phosphoinositide-3-kinase, class 2, beta polypeptide | 250.19 | 605.37 | -1.27 | 2.82E-09 |
| ENSMUSG00000046062 | GSMG0000778 | Ppp1r15b | protein phosphatase 1, regulatory (inhibitor) subunit 15b | 2397.01 | 2476.27 | -0.05 | 8.66E-01 |
| ENSMUSG00000041757 | GSMG0000780 | Plekha6 | pleckstrin homology domain containing, family A member 6 | 272.39 | 231.71 | 0.23 | 5.86E-01 |
| ENSMUSG00000070644 | GSMG0000783 | Etnk2 | ethanolamine kinase 2 | 83.93 | 86.09 | -0.03 | 9.64E-01 |
| ENSMUSG00000041559 | GSMG0000789 | Fmod | fibromodulin | 505.64 | 355.86 | 0.51 | 1.06E-01 |
| ENSMUSG00000026456 | GSMG0000796 | Cyb5r1 | cytochrome b5 reductase 1 | 1069.03 | 958.98 | 0.16 | 5.35E-01 |
| ENSMUSG00000026457 | GSMG0000797 | Adipor1 | adiponectin receptor 1 | 3857.93 | 3806.87 | 0.02 | 9.43E-01 |
| ENSMUSG00000026455 | GSMG0000799 | Klhl12 | kelch-like 12 | 376.35 | 393.63 | -0.06 | 8.80E-01 |
| ENSMUSG00000042229 | GSMG0000800 | Rabif | RAB interacting factor | 206.68 | 195.28 | 0.08 | 8.54E-01 |
| ENSMUSG00000042207 | GSMG0000801 | Kdm5b | lysine (K)-specific demethylase 5B | 3501.36 | 3832.80 | -0.13 | 5.41E-01 |
| ENSMUSG00000026429 | GSMG0000806 | Ube2t | ubiquitin-conjugating enzyme E2T | 32.44 | 57.04 | -0.81 | 1.12E-01 |
| ENSMUSG00000026426 | GSMG0000809 | Arl8a | ADP-ribosylation factor-like 8A | 1273.95 | 1978.82 | -0.64 | 2.82E-04 |
| ENSMUSG00000026421 | GSMG0000815 | Csrp1 | cysteine and glycine-rich protein 1 | 1752.00 | 1514.01 | 0.21 | 2.93E-01 |
| ENSMUSG00000041801 | GSMG0000816 | Phlda3 | pleckstrin homology like domain, family A, member 3 | 165.95 | 154.43 | 0.10 | 8.24E-01 |
| ENSMUSG00000026411 | GSMG0000821 | Tmem9 | transmembrane protein 9 | 1781.23 | 2105.81 | -0.24 | 2.48E-01 |
| ENSMUSG00000041642 | GSMG0000824 | Kif21b | kinesin family member 21B | 605.21 | 1048.12 | -0.79 | 2.61E-03 |
| ENSMUSG00000026404 | GSMG0000829 | Ddx59 | DEAD (Asp-Glu-Ala-Asp) box polypeptide 59 | 200.34 | 284.74 | -0.50 | 3.43E-02 |
| ENSMUSG00000041483 | GSMG0000833 | Zfp281 | zinc finger protein 281 | 978.61 | 983.58 | -0.01 | 9.81E-01 |
| NULL | GSMG0000836 | NULL | NULL | 265.54 | 413.94 | -0.64 | 2.13E-02 |
| NULL | GSMG0000838 | NULL | NULL | 301.00 | 289.17 | 0.06 | 8.83E-01 |
| NULL | GSMG0000840 | NULL | NULL | 123.60 | 162.54 | -0.40 | 2.93E-01 |
| NULL | GSMG0000845 | NULL | NULL | 126.78 | 195.56 | -0.62 | 2.75E-02 |

| | | | | | | | |
|---|---|---|---|---|---|---|---|
| ENSMUSG00000056268 | GSMG0000848 | Dennd1b | DENN/MADD domain containing 1B | 69.88 | 53.20 | 0.40 | 4.46E-01 |
| ENSMUSG00000033964 | GSMG0000857 | Zbtb41 | zinc finger and BTB domain containing 41 | 95.19 | 112.70 | -0.24 | 5.62E-01 |
| ENSMUSG00000052726 | GSMG0000862 | Kcnt2 | potassium channel, subfamily T, member 2 | 140.29 | 151.28 | -0.11 | 8.19E-01 |
| ENSMUSG00000033849 | GSMG0000864 | B3galt2 | UDP-Gal:betaGlcNAc beta 1,3-galactosyltransferase, polypeptide 2 | 318.12 | 493.70 | -0.63 | 1.42E-03 |
| ENSMUSG00000018196 | GSMG0000866 | Glrx2 | glutaredoxin 2 (thioltransferase) | 1045.62 | 1231.03 | -0.24 | 3.16E-01 |
| ENSMUSG00000018189 | GSMG0000867 | Uchl5 | ubiquitin carboxyl-terminal esterase L5 | 543.88 | 744.97 | -0.46 | 3.90E-02 |
| ENSMUSG00000035131 | GSMG0000869 | Brinp3 | bone morphogenetic protein/retinoic acid inducible neural specific 3 | 468.95 | 324.94 | 0.53 | 4.21E-02 |
| ENSMUSG00000006005 | GSMG0000876 | Tpr | translocated promoter region, nuclear basket protein | 1972.48 | 2480.70 | -0.33 | 1.14E-01 |
| ENSMUSG00000023150 | GSMG0000880 | Ivns1abp | influenza virus NS1A binding protein | 2406.28 | 3109.69 | -0.37 | 5.02E-02 |
| ENSMUSG00000053286 | GSMG0000882 | Trmt1l | tRNA methyltransferase 1 like | 560.35 | 465.89 | 0.27 | 2.48E-01 |
| NULL | GSMG0000883 | NULL | NULL | 252.78 | 90.20 | 1.48 | 6.73E-09 |
| ENSMUSG00000026483 | GSMG0000884 | Fam129a | family with sequence similarity 129, member A | 796.18 | 817.90 | -0.04 | 9.10E-01 |
| ENSMUSG00000043019 | GSMG0000885 | Edem3 | ER degradation enhancer, mannosidase alpha-like 3 | 914.55 | 950.75 | -0.06 | 8.45E-01 |
| ENSMUSG00000032649 | GSMG0000889 | Colgalt2 | collagen beta(1-O)galactosyltransferase 2 | 678.96 | 633.79 | 0.10 | 7.17E-01 |
| ENSMUSG00000008475 | GSMG0000892 | Arpc5 | actin related protein 2/3 complex, subunit 5 | 6069.22 | 8268.41 | -0.45 | 4.25E-02 |
| ENSMUSG00000026480 | GSMG0000893 | Ncf2 | neutrophil cytosolic factor 2 | 148.62 | 119.61 | 0.31 | 3.67E-01 |
| ENSMUSG00000042751 | GSMG0000896 | Nmnat2 | nicotinamide nucleotide adenylyltransferase 2 | 178.11 | 169.58 | 0.08 | 8.91E-01 |
| ENSMUSG00000026475 | GSMG0000901 | Rgs16 | regulator of G-protein signaling 16 | 283.92 | 176.48 | 0.69 | 1.62E-02 |
| ENSMUSG00000066800 | GSMG0000902 | Rnasel | ribonuclease L (2', 5'-oligoisoadenylate synthetase-dependent) | 201.01 | 375.46 | -0.89 | 1.16E-03 |
| ENSMUSG00000026473 | GSMG0000905 | Glul | glutamate-ammonia ligase (glutamine synthetase) | 3255.97 | 1597.83 | 1.03 | 1.95E-07 |
| NULL | GSMG0000908 | NULL | NULL | 204.70 | 129.54 | 0.65 | 3.97E-02 |
| ENSMUSG00000026470 | GSMG0000909 | Stx6 | syntaxin 6 | 1412.67 | 1868.45 | -0.40 | 1.83E-02 |
| ENSMUSG00000033701 | GSMG0000910 | Acbd6 | acyl-Coenzyme A binding domain containing 6 | 852.96 | 1162.42 | -0.45 | 2.56E-02 |
| ENSMUSG00000050565 | GSMG0000915 | Tor1aip2 | torsin A interacting protein 2 | 1847.61 | 2023.61 | -0.13 | 5.29E-01 |
| ENSMUSG00000026596 | GSMG0000922 | Abl2 | v-abl Abelson murine leukemia viral oncogene 2 (arg, Abelson-related | 787.82 | 1013.67 | -0.36 | 5.54E-02 |
| NULL | GSMG0000926 | NULL | NULL | 131.45 | 116.67 | 0.18 | 7.21E-01 |
| NULL | GSMG0000927 | NULL | NULL | 76.45 | 96.26 | -0.33 | 4.56E-01 |
| ENSMUSG00000049881 | GSMG0000932 | 2810025M15Rik | RIKEN cDNA 2810025M15 gene | 551.57 | 641.13 | -0.22 | 3.93E-01 |
| ENSMUSG00000033488 | GSMG0000933 | BC026585 | cDNA sequence BC026585 | 49.60 | 65.16 | -0.40 | 4.26E-01 |
| ENSMUSG00000026587 | GSMG0000935 | Astn1 | astrotactin 1 | 587.86 | 826.12 | -0.49 | 3.78E-02 |
| ENSMUSG00000015829 | GSMG0000941 | Tnr | tenascin R | 458.11 | 385.04 | 0.25 | 3.99E-01 |
| ENSMUSG00000090394 | GSMG0000945 | 4930523C07Rik | RIKEN cDNA 4930523C07 gene | 1530.58 | 1178.11 | 0.38 | 2.86E-02 |
| ENSMUSG00000058267 | GSMG0000946 | Mrps14 | mitochondrial ribosomal protein S14 | 855.05 | 865.51 | -0.02 | 9.63E-01 |

| | | | | | | | |
|---|---|---|---|---|---|---|---|
| ENSMUSG00000040423 | GSMG0000950 | Rc3h1 | RING CCCH (C3H) domains 1 | 715.89 | 652.33 | 0.14 | 6.42E-01 |
| ENSMUSG00000053332 | GSMG0000957 | Gas5 | growth arrest specific 5 | 1451.92 | 1473.00 | -0.02 | 9.48E-01 |
| ENSMUSG00000044835 | GSMG0000960 | Ankrd45 | ankyrin repeat domain 45 | 40.07 | 100.35 | -1.31 | 1.88E-04 |
| ENSMUSG00000026698 | GSMG0000965 | Pigc | phosphatidylinositol glycan anchor biosynthesis, class C | 1231.97 | 1150.45 | 0.10 | 7.23E-01 |
| ENSMUSG00000026696 | GSMG0000969 | Vamp4 | vesicle-associated membrane protein 4 | 1492.27 | 1306.51 | 0.19 | 5.06E-01 |
| ENSMUSG00000026585 | GSMG0000973 | Kifap3 | kinesin-associated protein 3 | 1626.94 | 1973.42 | -0.28 | 1.32E-01 |
| ENSMUSG00000026584 | GSMG0000977 | Scyl3 | SCY1-like 3 (S. cerevisiae) | 757.44 | 884.53 | -0.22 | 3.34E-01 |
| ENSMUSG00000026579 | GSMG0000982 | F5 | coagulation factor V | 201.88 | 50.15 | 2.01 | 4.10E-02 |
| ENSMUSG00000040918 | GSMG0000983 | Slc19a2 | solute carrier family 19 (thiamine transporter), member 2 | 323.22 | 279.46 | 0.21 | 5.00E-01 |
| ENSMUSG00000026578 | GSMG0000984 | Ccdc181 | coiled-coil domain containing 181 | 313.40 | 304.06 | 0.04 | 9.10E-01 |
| ENSMUSG00000026575 | GSMG0000986 | Nme7 | NME/NM23 family member 7 | 287.79 | 304.12 | -0.08 | 8.40E-01 |
| ENSMUSG00000026568 | GSMG0000997 | Mpc2 | mitochondrial pyruvate carrier 2 | 1370.61 | 1502.63 | -0.13 | 6.79E-01 |
| ENSMUSG00000040713 | GSMG0001002 | Creg1 | cellular repressor of E1A-stimulated genes 1 | 1907.17 | 1713.31 | 0.15 | 4.91E-01 |
| ENSMUSG00000040612 | GSMG0001009 | Ildr2 | immunoglobulin-like domain containing receptor 2 | 239.57 | 441.55 | -0.88 | 2.23E-01 |
| NULL | GSMG0001010 | NULL | NULL | 406.43 | 294.90 | 0.47 | 5.66E-02 |
| ENSMUSG00000026563 | GSMG0001012 | Tada1 | transcriptional adaptor 1 | 413.34 | 618.22 | -0.58 | 8.51E-03 |
| ENSMUSG00000060568 | GSMG0001013 | Fam78b | family with sequence similarity 78, member B | 51.95 | 105.29 | -1.02 | 4.06E-03 |
| NULL | GSMG0001017 | NULL | NULL | 49.65 | 100.15 | -1.01 | 1.42E-02 |
| ENSMUSG00000052428 | GSMG0001018 | Tmco1 | transmembrane and coiled-coil domains 1 | 2131.09 | 2028.19 | 0.07 | 8.12E-01 |
| ENSMUSG00000026687 | GSMG0001019 | Aldh9a1 | aldehyde dehydrogenase 9, subfamily A1 | 1438.35 | 1958.88 | -0.45 | 7.37E-03 |
| ENSMUSG00000015843 | GSMG0001021 | Rxrg | retinoid X receptor gamma | 45.18 | 96.05 | -1.08 | 7.93E-03 |
| ENSMUSG00000026678 | GSMG0001023 | Rgs5 | regulator of G-protein signaling 5 | 153669.94 | 87644.32 | 0.81 | 1.46E-04 |
| ENSMUSG00000038463 | GSMG0001032 | Olfml2b | olfactomedin-like 2B | 210.10 | 155.82 | 0.44 | 2.09E-01 |
| ENSMUSG00000005677 | GSMG0001038 | Nr1i3 | nuclear receptor subfamily 1, group I, member 3 | 338.25 | 324.73 | 0.06 | 8.63E-01 |
| ENSMUSG00000006403 | GSMG0001040 | Adamts4 | a disintegrin-like and metallopeptidase (reprolysin type) with thrombos[illegible] | 429.43 | 229.29 | 0.90 | 7.38E-04 |
| ENSMUSG00000052423 | GSMG0001041 | B4galt3 | UDP-Gal:betaGlcNAc beta 1,4-galactosyltransferase, polypeptide 3 | 667.28 | 930.18 | -0.48 | 8.43E-03 |
| ENSMUSG00000013973 | GSMG0001042 | Dedd | death effector domain-containing | 340.45 | 363.73 | -0.10 | 7.73E-01 |
| ENSMUSG00000006412 | GSMG0001043 | Pfdn2 | prefoldin 2 | 691.12 | 925.01 | -0.42 | 6.69E-02 |
| ENSMUSG00000026641 | GSMG0001048 | Usf1 | upstream transcription factor 1 | 532.98 | 535.41 | -0.01 | 9.88E-01 |
| ENSMUSG00000038235 | GSMG0001049 | F11r | F11 receptor | 2678.71 | 2271.75 | 0.24 | 1.99E-01 |
| ENSMUSG00000060244 | GSMG0001050 | Alyref2 | Aly/REF export factor 2 | 260.87 | 354.71 | -0.45 | 8.96E-02 |
| ENSMUSG00000026553 | GSMG0001059 | Copa | coatomer protein complex subunit alpha | 8963.97 | 9659.44 | -0.11 | 6.19E-01 |
| ENSMUSG00000003464 | GSMG0001060 | Pex19 | peroxisomal biogenesis factor 19 | 592.83 | 551.37 | 0.11 | 6.99E-01 |

| | | | | | | | |
|---|---|---|---|---|---|---|---|
| ENSMUSG00000026554 | GSMG0001061 | Dcaf8 | DDB1 and CUL4 associated factor 8 | 2689.22 | 2620.24 | 0.04 | 8.95E-01 |
| NULL | GSMG0001062 | NULL | NULL | 136.72 | 146.45 | -0.10 | 8.36E-01 |
| ENSMUSG00000038034 | GSMG0001063 | Igsf8 | immunoglobulin superfamily, member 8 | 2299.69 | 2660.16 | -0.21 | 3.32E-01 |
| ENSMUSG00000044708 | GSMG0001064 | Kcnj10 | potassium inwardly-rectifying channel, subfamily J, member 10 | 993.37 | 569.47 | 0.80 | 5.64E-05 |
| ENSMUSG00000037995 | GSMG0001069 | Igsf9 | immunoglobulin superfamily, member 9 | 294.22 | 969.19 | -1.72 | 3.34E-12 |
| ENSMUSG00000026547 | GSMG0001070 | Tagln2 | transgelin 2 | 5575.03 | 5971.98 | -0.10 | 6.57E-01 |
| ENSMUSG00000039748 | GSMG0001105 | Exo1 | exonuclease 1 | 102.00 | 265.22 | -1.37 | 2.14E-06 |
| ENSMUSG00000026504 | GSMG0001110 | Sdccag8 | serologically defined colon cancer antigen 8 | 541.84 | 411.85 | 0.39 | 6.64E-02 |
| NULL | GSMG0001111 | NULL | NULL | 71.25 | 43.93 | 0.70 | 1.04E-01 |
| ENSMUSG00000063659 | GSMG0001117 | Zbtb18 | zinc finger and BTB domain containing 18 | 1449.92 | 5871.30 | -2.02 | 4.02E-37 |
| ENSMUSG00000026502 | GSMG0001123 | Desi2 | desumoylating isopeptidase 2 | 1398.40 | 1674.80 | -0.26 | 1.93E-01 |
| ENSMUSG00000026500 | GSMG0001128 | Cox20 | COX20 Cox2 chaperone | 149.37 | 155.87 | -0.07 | 8.98E-01 |
| ENSMUSG00000026495 | GSMG0001130 | Efcab2 | EF-hand calcium binding domain 2 | 333.93 | 306.16 | 0.13 | 7.07E-01 |
| ENSMUSG00000038949 | GSMG0001137 | Cnst | consortin, connexin sorting protein | 538.42 | 501.73 | 0.10 | 7.80E-01 |
| ENSMUSG00000038936 | GSMG0001138 | Sccpdh | saccharopine dehydrogenase (putative) | 1171.57 | 1494.92 | -0.35 | 3.46E-02 |
| ENSMUSG00000026490 | GSMG0001140 | Cdc42bpa | CDC42 binding protein kinase alpha | 502.09 | 363.04 | 0.46 | 8.99E-02 |
| ENSMUSG00000058729 | GSMG0001144 | Lin9 | lin-9 homolog (C. elegans) | 93.87 | 122.20 | -0.38 | 4.03E-01 |
| ENSMUSG00000038855 | GSMG0001145 | Itpkb | inositol 1,4,5-trisphosphate 3-kinase B | 1106.57 | 843.02 | 0.39 | 6.39E-01 |
| NULL | GSMG0001149 | NULL | NULL | 87.18 | 75.82 | 0.19 | 7.35E-01 |
| ENSMUSG00000026496 | GSMG0001150 | Parp1 | poly (ADP-ribose) polymerase family, member 1 | 1722.17 | 2384.70 | -0.47 | 5.96E-03 |
| ENSMUSG00000026499 | GSMG0001151 | Acbd3 | acyl-Coenzyme A binding domain containing 3 | 649.71 | 697.12 | -0.10 | 7.10E-01 |
| ENSMUSG00000038806 | GSMG0001155 | Sde2 | SDE2 telomere maintenance homolog (S. pombe) | 738.05 | 816.85 | -0.15 | 5.36E-01 |
| ENSMUSG00000026520 | GSMG0001157 | Pycr2 | pyrroline-5-carboxylate reductase family, member 2 | 408.46 | 448.82 | -0.13 | 6.48E-01 |
| ENSMUSG00000026519 | GSMG0001159 | Tmem63a | transmembrane protein 63a | 242.27 | 161.90 | 0.59 | 3.34E-02 |
| ENSMUSG00000062169 | GSMG0001163 | Cnih4 | cornichon family AMPA receptor auxiliary protein 4 | 606.04 | 726.87 | -0.26 | 2.16E-01 |
| ENSMUSG00000026514 | GSMG0001165 | Cnih3 | cornichon family AMPA receptor auxiliary protein 3 | 423.14 | 625.96 | -0.56 | 9.35E-03 |
| ENSMUSG00000026510 | GSMG0001173 | Trp53bp2 | transformation related protein 53 binding protein 2 | 633.89 | 683.73 | -0.11 | 6.89E-01 |
| NULL | GSMG0001174 | NULL | NULL | 158.98 | 119.09 | 0.42 | 1.82E-01 |
| ENSMUSG00000038576 | GSMG0001178 | Susd4 | sushi domain containing 4 | 693.55 | 471.66 | 0.56 | 1.17E-02 |
| NULL | GSMG0001191 | NULL | NULL | 386.07 | 257.43 | 0.59 | 1.32E-02 |
| ENSMUSG00000039318 | GSMG0001192 | Rab3gap2 | RAB3 GTPase activating protein subunit 2 | 855.64 | 760.64 | 0.17 | 4.32E-01 |
| ENSMUSG00000026617 | GSMG0001195 | Bpnt1 | bisphosphate 3'-nucleotidase 1 | 305.42 | 331.49 | -0.12 | 7.70E-01 |
| ENSMUSG00000026615 | GSMG0001196 | Eprs | glutamyl-prolyl-tRNA synthetase | 3395.70 | 4417.97 | -0.38 | 1.28E-02 |

| | | | | | | | |
|---|---|---|---|---|---|---|---|
| ENSMUSG00000026614 | GSMG0001197 | Slc30a10 | solute carrier family 30, member 10 | 2116.09 | 2276.69 | -0.11 | 6.55E-01 |
| ENSMUSG00000039210 | GSMG0001205 | Gpatch2 | G patch domain containing 2 | 780.11 | 723.05 | 0.11 | 6.85E-01 |
| NULL | GSMG0001218 | NULL | NULL | 300.56 | 290.42 | 0.04 | 9.30E-01 |
| ENSMUSG00000026604 | GSMG0001219 | Ptpn14 | protein tyrosine phosphatase, non-receptor type 14 | 97.27 | 67.05 | 0.54 | 1.61E-01 |
| NULL | GSMG0001222 | NULL | NULL | 682.80 | 865.26 | -0.34 | 1.70E-01 |
| ENSMUSG00000026634 | GSMG0001225 | Angel2 | angel homolog 2 (Drosophila) | 1247.21 | 1305.32 | -0.07 | 8.17E-01 |
| ENSMUSG00000062510 | GSMG0001228 | Nsl1 | NSL1, MIS12 kinetochore complex component | 172.61 | 432.86 | -1.32 | 7.24E-08 |
| ENSMUSG00000073478 | GSMG0001230 | D730003I15Rik | RIKEN cDNA D730003I15 gene | 147.79 | 178.77 | -0.27 | 5.00E-01 |
| ENSMUSG00000026627 | GSMG0001233 | Tmem206 | transmembrane protein 206 | 400.41 | 600.79 | -0.58 | 1.91E-03 |
| ENSMUSG00000037461 | GSMG0001236 | Ints7 | integrator complex subunit 7 | 881.10 | 1234.37 | -0.49 | 1.79E-02 |
| ENSMUSG00000026623 | GSMG0001238 | Lpgat1 | lysophosphatidylglycerol acyltransferase 1 | 539.23 | 492.90 | 0.13 | 6.51E-01 |
| ENSMUSG00000026622 | GSMG0001241 | Nek2 | NIMA (never in mitosis gene a)-related expressed kinase 2 | 83.96 | 190.39 | -1.17 | 1.59E-04 |
| ENSMUSG00000037434 | GSMG0001242 | Slc30a1 | solute carrier family 30 (zinc transporter), member 1 | 3162.21 | 2549.41 | 0.31 | 2.84E-01 |
| ENSMUSG00000058248 | GSMG0001247 | Kcnh1 | potassium voltage-gated channel, subfamily H (eag-related), member 1 | 461.87 | 391.66 | 0.24 | 4.33E-01 |
| G00000026638,ENSMUSG000 | GSMG0001250 | Irf6 // A130010J15 | interferon regulatory factor 6 // RIKEN cDNA A130010J15 gene | 954.60 | 774.30 | 0.30 | 1.07E-01 |
| ENSMUSG00000026640 | GSMG0001256 | Plxna2 | plexin A2 | 450.11 | 460.20 | -0.03 | 9.46E-01 |
| NULL | GSMG0001257 | NULL | NULL | 491.74 | 402.59 | 0.29 | 4.49E-01 |
| ENSMUSG00000016494 | GSMG0001261 | Cd34 | CD34 antigen | 9203.09 | 9715.48 | -0.08 | 7.58E-01 |
| ENSMUSG00000096929 | GSMG0001262 | A330023F24Rik | RIKEN cDNA A330023F24 gene | 129.38 | 80.50 | 0.68 | 3.84E-02 |
| NULL | GSMG0001264 | NULL | NULL | 1648.53 | 920.85 | 0.84 | 7.47E-09 |
| NULL | GSMG0001266 | NULL | NULL | 321.88 | 325.12 | -0.01 | 9.71E-01 |
| NULL | GSMG0001267 | NULL | NULL | 587.85 | 644.45 | -0.13 | 6.35E-01 |
| NULL | GSMG0001268 | NULL | NULL | 442.00 | 559.08 | -0.34 | 3.08E-01 |
| NULL | GSMG0001270 | NULL | NULL | 192.44 | 248.78 | -0.37 | 1.86E-01 |
| NULL | GSMG0001272 | NULL | NULL | 482.59 | 522.13 | -0.11 | 7.04E-01 |
| NULL | GSMG0001274 | NULL | NULL | 531.33 | 494.97 | 0.10 | 7.27E-01 |
| ENSMUSG00000025902 | GSMG0001279 | Sox17 | SRY (sex determining region Y)-box 17 | 7152.67 | 3949.58 | 0.86 | 2.08E-09 |
| ENSMUSG00000033845 | GSMG0001281 | Mrpl15 | mitochondrial ribosomal protein L15 | 696.51 | 844.27 | -0.28 | 2.97E-01 |
| NULL | GSMG0001283 | NULL | NULL | 144.53 | 168.39 | -0.23 | 5.53E-01 |
| ENSMUSG00000025909 | GSMG0001292 | Sntg1 | syntrophin, gamma 1 | 75.74 | 46.54 | 0.70 | 1.32E-01 |
| ENSMUSG00000045210 | GSMG0001300 | Vcpip1 | valosin containing protein (p97)/p47 complex interacting protein 1 | 1518.89 | 1480.37 | 0.04 | 8.89E-01 |
| ENSMUSG00000098234 | GSMG0001301 | Snhg6 | small nucleolar RNA host gene 6 | 151.54 | 214.32 | -0.51 | 1.23E-01 |
| ENSMUSG00000025917 | GSMG0001304 | Cops5 | COP9 signalosome subunit 5 | 1332.89 | 1776.77 | -0.41 | 4.11E-02 |

| | | | | | | | |
|---|---|---|---|---|---|---|---|
| ENSMUSG00000025938 | GSMG0001318 | Slco5a1 | solute carrier organic anion transporter family, member 5A1 | 48.06 | 119.50 | -1.30 | 6.90E-05 |
| NULL | GSMG0001322 | NULL | NULL | 151.05 | 145.60 | 0.05 | 9.28E-01 |
| ENSMUSG00000025935 | GSMG0001329 | Tram1 | translocating chain-associating membrane protein 1 | 2744.53 | 3531.84 | -0.36 | 7.65E-02 |
| ENSMUSG00000025937 | GSMG0001332 | Lactb2 | lactamase, beta 2 | 288.02 | 240.81 | 0.26 | 3.95E-01 |
| ENSMUSG00000025920 | GSMG0001345 | Stau2 | staufen (RNA binding protein) homolog 2 (Drosophila) | 859.33 | 1282.70 | -0.58 | 1.98E-04 |
| NULL | GSMG0001350 | NULL | NULL | 115.77 | 188.76 | -0.72 | 5.76E-02 |
| ENSMUSG00000025939 | GSMG0001351 | Ube2w | ubiquitin-conjugating enzyme E2W (putative) | 834.23 | 788.84 | 0.08 | 8.36E-01 |
| ENSMUSG00000079658 | GSMG0001354 | Tceb1 | transcription elongation factor B (SIII), polypeptide 1 | 1251.63 | 1541.61 | -0.30 | 1.58E-01 |
| ENSMUSG00000042686 | GSMG0001357 | Jph1 | junctophilin 1 | 291.02 | 400.61 | -0.46 | 3.25E-02 |
| ENSMUSG00000041859 | GSMG0001370 | Mcm3 | minichromosome maintenance complex component 3 | 673.05 | 1686.55 | -1.32 | 2.37E-10 |
| NULL | GSMG0001372 | NULL | NULL | 1028.29 | 1282.31 | -0.32 | 1.31E-01 |
| ENSMUSG00000041779 | GSMG0001373 | Tram2 | translocating chain-associating membrane protein 2 | 285.87 | 363.99 | -0.35 | 1.93E-01 |
| ENSMUSG00000028033 | GSMG0001375 | Kcnq5 | potassium voltage-gated channel, subfamily Q, member 5 | 198.68 | 321.61 | -0.70 | 1.43E-02 |
| NULL | GSMG0001378 | NULL | NULL | 414.13 | 482.18 | -0.22 | 5.29E-01 |
| NULL | GSMG0001379 | NULL | NULL | 221.82 | 216.74 | 0.04 | 9.36E-01 |
| NULL | GSMG0001380 | NULL | NULL | 444.31 | 480.42 | -0.11 | 8.08E-01 |
| NULL | GSMG0001382 | NULL | NULL | 138.51 | 178.86 | -0.37 | 3.89E-01 |
| NULL | GSMG0001384 | NULL | NULL | 107.00 | 119.53 | -0.16 | 7.43E-01 |
| NULL | GSMG0001385 | NULL | NULL | 414.26 | 422.79 | -0.03 | 9.61E-01 |
| NULL | GSMG0001386 | NULL | NULL | 117.41 | 176.07 | -0.58 | 7.36E-02 |
| NULL | GSMG0001390 | NULL | NULL | 199.27 | 144.09 | 0.47 | 1.54E-01 |
| NULL | GSMG0001393 | NULL | NULL | 245.72 | 145.33 | 0.76 | 3.66E-03 |
| NULL | GSMG0001394 | NULL | NULL | 373.61 | 218.64 | 0.77 | 8.36E-04 |
| ENSMUSG00000026158 | GSMG0001398 | Ogfrl1 | opioid growth factor receptor-like 1 | 1056.98 | 920.01 | 0.20 | 4.23E-01 |
| ENSMUSG00000026155 | GSMG0001400 | Smap1 | small ArfGAP 1 | 665.00 | 680.16 | -0.04 | 9.29E-01 |
| ENSMUSG00000026154 | GSMG0001402 | Sdhaf4 | succinate dehydrogenase complex assembly factor 4 | 614.50 | 597.88 | 0.04 | 9.29E-01 |
| ENSMUSG00000026153 | GSMG0001403 | Fam135a | family with sequence similarity 135, member A | 229.73 | 228.46 | 0.01 | 9.85E-01 |
| ENSMUSG00000026141 | GSMG0001406 | Col19a1 | collagen, type XIX, alpha 1 | 1530.95 | 759.17 | 1.01 | 2.14E-08 |
| ENSMUSG00000104257 | GSMG0001409 | Gm20172 | predicted gene, 20172 | 192.12 | 159.42 | 0.27 | 5.06E-01 |
| ENSMUSG00000033569 | GSMG0001416 | Adgrb3 | adhesion G protein-coupled receptor B3 | 457.25 | 824.09 | -0.85 | 3.21E-05 |
| NULL | GSMG0001419 | NULL | NULL | 444.99 | 305.87 | 0.54 | 5.02E-02 |
| NULL | GSMG0001421 | NULL | NULL | 479.41 | 399.30 | 0.26 | 3.34E-01 |
| NULL | GSMG0001422 | NULL | NULL | 386.54 | 324.66 | 0.25 | 3.72E-01 |

| | | | | | | | |
|---|---|---|---|---|---|---|---|
| NULL | GSMG0001427 | NULL | NULL | 268.48 | 207.39 | 0.37 | 3.89E-01 |
| ENSMUSG00000048874 | GSMG0001435 | Phf3 | PHD finger protein 3 | 203.63 | 229.31 | -0.17 | 5.95E-01 |
| ENSMUSG00000026134 | GSMG0001441 | Prim2 | DNA primase, p58 subunit | 197.69 | 389.47 | -0.97 | 4.11E-05 |
| ENSMUSG00000042215 | GSMG0001444 | Bag2 | BCL2-associated athanogene 2 | 1006.89 | 983.65 | 0.03 | 9.08E-01 |
| ENSMUSG00000042197 | GSMG0001445 | Zfp451 | zinc finger protein 451 | 205.94 | 213.74 | -0.05 | 8.94E-01 |
| ENSMUSG00000042182 | GSMG0001447 | Bend6 | BEN domain containing 6 | 223.73 | 439.45 | -0.97 | 5.11E-07 |
| ENSMUSG00000042111 | GSMG0001454 | Ccdc115 | coiled-coil domain containing 115 | 534.64 | 760.89 | -0.51 | 4.53E-03 |
| NULL | GSMG0001460 | NULL | NULL | 265.83 | 231.29 | 0.20 | 5.93E-01 |
| ENSMUSG00000037503 | GSMG0001461 | Fam168b | family with sequence similarity 168, member B | 522.90 | 583.74 | -0.16 | 6.33E-01 |
| ENSMUSG00000037470 | GSMG0001463 | Uggt1 | UDP-glucose glycoprotein glucosyltransferase 1 | 2171.27 | 2535.06 | -0.22 | 2.49E-01 |
| ENSMUSG00000010453 | GSMG0001466 | Kansl3 | KAT8 regulatory NSL complex subunit 3 | 1663.79 | 1463.45 | 0.18 | 4.66E-01 |
| ENSMUSG00000001143 | GSMG0001469 | Lman2l | lectin, mannose-binding 2-like | 261.59 | 237.06 | 0.14 | 6.51E-01 |
| ENSMUSG00000026121 | GSMG0001472 | Sema4c | sema domain, immunoglobulin domain (Ig), transmembrane domain (T | 905.99 | 1150.49 | -0.34 | 6.54E-01 |
| ENSMUSG00000037351 | GSMG0001475 | Actr1b | ARP1 actin-related protein 1B, centractin beta | 5588.10 | 5040.09 | 0.15 | 4.50E-01 |
| ENSMUSG00000026116 | GSMG0001477 | Tmem131 | transmembrane protein 131 | 1572.63 | 2169.74 | -0.46 | 8.22E-03 |
| ENSMUSG00000026112 | GSMG0001484 | Coa5 | cytochrome C oxidase assembly factor 5 | 1590.86 | 1608.40 | -0.01 | 9.66E-01 |
| ENSMUSG00000026110 | GSMG0001486 | Mgat4a | mannoside acetylglucosaminyltransferase 4, isoenzyme A | 376.64 | 489.69 | -0.38 | 6.68E-02 |
| ENSMUSG00000026090 | GSMG0001489 | 2010300C02Rik | RIKEN cDNA 2010300C02 gene | 132.86 | 49.93 | 1.41 | 9.76E-06 |
| ENSMUSG00000060771 | GSMG0001491 | Tsga10 | testis specific 10 | 159.56 | 214.15 | -0.43 | 1.43E-01 |
| ENSMUSG00000026088 | GSMG0001492 | Mitd1 | MIT, microtubule interacting and transport, domain containing 1 | 335.26 | 316.34 | 0.08 | 8.32E-01 |
| ENSMUSG00000058407 | GSMG0001495 | Txndc9 | thioredoxin domain containing 9 | 745.85 | 841.23 | -0.17 | 5.46E-01 |
| ENSMUSG00000026082 | GSMG0001496 | Rev1 | REV1, DNA directed polymerase | 1642.62 | 1682.70 | -0.03 | 9.03E-01 |
| ENSMUSG00000037138 | GSMG0001498 | Aff3 | AF4/FMR2 family, member 3 | 296.18 | 482.63 | -0.71 | 2.21E-02 |
| NULL | GSMG0001500 | NULL | NULL | 91.86 | 157.52 | -0.77 | 1.39E-02 |
| NULL | GSMG0001509 | NULL | NULL | 231.38 | 312.08 | -0.43 | 7.76E-02 |
| ENSMUSG00000026080 | GSMG0001513 | Chst10 | carbohydrate sulfotransferase 10 | 187.36 | 361.45 | -0.95 | 8.71E-06 |
| ENSMUSG00000003134 | GSMG0001517 | Tbc1d8 | TBC1 domain family, member 8 | 543.96 | 575.60 | -0.08 | 8.18E-01 |
| ENSMUSG00000050967 | GSMG0001519 | Creg2 | cellular repressor of E1A-stimulated genes 2 | 643.95 | 595.52 | 0.11 | 6.65E-01 |
| ENSMUSG00000041945 | GSMG0001528 | Mfsd9 | major facilitator superfamily domain containing 9 | 380.96 | 339.94 | 0.16 | 5.53E-01 |
| ENSMUSG00000060424 | GSMG0001531 | Pantr1 | POU domain, class 3, transcription factor 3 adjacent noncoding transcri | 418.31 | 791.85 | -0.92 | 5.20E-08 |
| ENSMUSG00000070939 | GSMG0001533 | Tgfbrap1 | transforming growth factor, beta receptor associated protein 1 | 398.39 | 355.35 | 0.17 | 6.04E-01 |
| ENSMUSG00000008136 | GSMG0001534 | Fhl2 | four and a half LIM domains 2 | 218.79 | 160.55 | 0.45 | 1.39E-01 |
| ENSMUSG00000057363 | GSMG0001538 | Uxs1 | UDP-glucuronate decarboxylase 1 | 333.18 | 461.22 | -0.47 | 3.93E-02 |

| | | | | | | | |
|---|---|---|---|---|---|---|---|
| ENSMUSG00000026049 | GSMG0001540 | Tex30 | testis expressed 30 | 156.84 | 202.76 | -0.37 | 2.75E-01 |
| ENSMUSG00000026047 | GSMG0001541 | Kdelc1 | KDEL (Lys-Asp-Glu-Leu) containing 1 | 1113.78 | 1115.70 | 0.00 | 9.97E-01 |
| ENSMUSG00000026042 | GSMG0001545 | Col5a2 | collagen, type V, alpha 2 | 1783.98 | 1958.94 | -0.13 | 6.33E-01 |
| ENSMUSG00000025993 | GSMG0001548 | Slc40a1 | solute carrier family 40 (iron-regulated transporter), member 1 | 14595.54 | 14991.82 | -0.04 | 8.79E-01 |
| ENSMUSG00000025986 | GSMG0001550 | Slc39a10 | solute carrier family 39 (zinc transporter), member 10 | 3938.09 | 4118.04 | -0.06 | 8.47E-01 |
| NULL | GSMG0001551 | NULL | NULL | 103.40 | 60.94 | 0.76 | 4.27E-02 |
| ENSMUSG00000018417 | GSMG0001558 | Myo1b | myosin IB | 10530.82 | 13646.44 | -0.37 | 2.50E-02 |
| ENSMUSG00000026103 | GSMG0001561 | Gls | glutaminase | 532.96 | 480.42 | 0.15 | 5.96E-01 |
| NULL | GSMG0001562 | NULL | NULL | 348.68 | 211.60 | 0.72 | 5.05E-04 |
| ENSMUSG00000002881 | GSMG0001564 | Nab1 | Ngfi-A binding protein 1 | 699.06 | 570.88 | 0.29 | 2.29E-01 |
| ENSMUSG00000041439 | GSMG0001568 | Mfsd6 | major facilitator superfamily domain containing 6 | 180.80 | 196.56 | -0.12 | 7.59E-01 |
| ENSMUSG00000026102 | GSMG0001571 | Inpp1 | inositol polyphosphate-1-phosphatase | 486.75 | 347.25 | 0.49 | 1.43E-02 |
| ENSMUSG00000043629 | GSMG0001572 | 1700019D03Rik | RIKEN cDNA 1700019D03 gene | 171.88 | 161.79 | 0.09 | 8.44E-01 |
| ENSMUSG00000026095 | GSMG0001576 | Asnsd1 | asparagine synthetase domain containing 1 | 826.40 | 1162.80 | -0.49 | 1.02E-02 |
| ENSMUSG00000026094 | GSMG0001579 | Stk17b | serine/threonine kinase 17b (apoptosis-inducing) | 178.68 | 161.01 | 0.15 | 7.22E-01 |
| ENSMUSG00000042807 | GSMG0001581 | Hecw2 | HECT, C2 and WW domain containing E3 ubiquitin protein ligase 2 | 96.57 | 140.03 | -0.53 | 1.08E-01 |
| NULL | GSMG0001585 | NULL | NULL | 63.92 | 30.43 | 1.06 | 4.94E-02 |
| NULL | GSMG0001586 | NULL | NULL | 211.76 | 141.16 | 0.58 | 4.01E-02 |
| ENSMUSG00000041303 | GSMG0001587 | Gtf3c3 | general transcription factor IIIC, polypeptide 3 | 698.10 | 794.57 | -0.19 | 4.54E-01 |
| ENSMUSG00000073678 | GSMG0001590 | Pgap1 | post-GPI attachment to proteins 1 | 96.18 | 134.49 | -0.48 | 1.77E-01 |
| ENSMUSG00000052331 | GSMG0001592 | Ankrd44 | ankyrin repeat domain 44 | 123.37 | 102.60 | 0.27 | 5.48E-01 |
| ENSMUSG00000025982 | GSMG0001596 | Sf3b1 | splicing factor 3b, subunit 1 | 15082.22 | 15581.49 | -0.05 | 8.56E-01 |
| ENSMUSG00000025980 | GSMG0001599 | Hspd1 | heat shock protein 1 (chaperonin) | 335.05 | 496.73 | -0.57 | 9.55E-03 |
| ENSMUSG00000025978 | GSMG0001600 | Rftn2 | raftlin family member 2 | 425.12 | 505.76 | -0.25 | 3.22E-01 |
| ENSMUSG00000038331 | GSMG0001603 | Satb2 | special AT-rich sequence binding protein 2 | 154.03 | 591.46 | -1.93 | 1.69E-21 |
| ENSMUSG00000048495 | GSMG0001606 | Tyw5 | tRNA-yW synthesizing protein 5 | 205.22 | 208.62 | -0.03 | 9.55E-01 |
| ENSMUSG00000054770 | GSMG0001608 | Kctd18 | potassium channel tetramerisation domain containing 18 | 245.59 | 237.02 | 0.05 | 8.90E-01 |
| ENSMUSG00000026034 | GSMG0001611 | Clk1 | CDC-like kinase 1 | 5264.77 | 3639.53 | 0.53 | 2.78E-03 |
| ENSMUSG00000026035 | GSMG0001612 | Ppil3 | peptidylprolyl isomerase (cyclophilin)-like 3 | 168.67 | 196.00 | -0.21 | 5.66E-01 |
| ENSMUSG00000026037 | GSMG0001613 | Orc2 | origin recognition complex, subunit 2 | 488.71 | 536.51 | -0.13 | 6.20E-01 |
| ENSMUSG00000038174 | GSMG0001615 | Fam126b | family with sequence similarity 126, member B | 189.15 | 139.48 | 0.44 | 1.51E-01 |
| ENSMUSG00000026028 | GSMG0001619 | Trak2 | trafficking protein, kinesin binding 2 | 2331.26 | 1825.75 | 0.35 | 4.41E-02 |
| ENSMUSG00000038079 | GSMG0001622 | Tmem237 | transmembrane protein 237 | 744.41 | 1053.86 | -0.50 | 9.00E-03 |

| | | | | | | | |
|---|---|---|---|---|---|---|---|
| ENSMUSG00000026018 | GSMG0001631 | Ica1l | islet cell autoantigen 1-like | 101.67 | 144.86 | -0.51 | 1.28E-01 |
| ENSMUSG00000026019 | GSMG0001633 | Wdr12 | WD repeat domain 12 | 424.24 | 568.83 | -0.42 | 4.05E-02 |
| NULL | GSMG0001634 | NULL | NULL | 196.57 | 227.48 | -0.22 | 5.82E-01 |
| NULL | GSMG0001637 | NULL | NULL | 309.98 | 377.61 | -0.29 | 2.55E-01 |
| NULL | GSMG0001641 | NULL | NULL | 70.24 | 96.97 | -0.45 | 3.61E-01 |
| ENSMUSG00000025968 | GSMG0001653 | Ndufs1 | NADH dehydrogenase (ubiquinone) Fe-S protein 1 | 1903.78 | 1890.70 | 0.01 | 9.73E-01 |
| ENSMUSG00000025959 | GSMG0001659 | Klf7 | Kruppel-like factor 7 (ubiquitous) | 995.17 | 1411.98 | -0.50 | 1.06E-02 |
| NULL | GSMG0001660 | NULL | NULL | 627.43 | 947.26 | -0.60 | 1.97E-02 |
| NULL | GSMG0001661 | NULL | NULL | 661.51 | 731.26 | -0.14 | 5.68E-01 |
| ENSMUSG00000025956 | GSMG0001664 | Mettl21a | methyltransferase like 21A | 539.19 | 482.36 | 0.16 | 5.95E-01 |
| ENSMUSG00000051344 | GSMG0001668 | Plekhm3 | pleckstrin homology domain containing, family M, member 3 | 476.58 | 407.14 | 0.23 | 5.02E-01 |
| ENSMUSG00000044816 | GSMG0001677 | D630023F18Rik | RIKEN cDNA D630023F18 gene | 57.80 | 35.02 | 0.72 | 1.40E-01 |
| ENSMUSG00000025950 | GSMG0001678 | Idh1 | isocitrate dehydrogenase 1 (NADP+), soluble | 1500.60 | 2545.25 | -0.76 | 2.37E-06 |
| NULL | GSMG0001680 | NULL | NULL | 477.96 | 486.16 | -0.03 | 9.61E-01 |
| ENSMUSG00000026004 | GSMG0001683 | Kansl1l | KAT8 regulatory NSL complex subunit 1-like | 543.18 | 631.55 | -0.22 | 3.82E-01 |
| ENSMUSG00000026003 | GSMG0001689 | Acadl | acyl-Coenzyme A dehydrogenase, long-chain | 2729.83 | 3792.68 | -0.47 | 5.04E-03 |
| ENSMUSG00000026000 | GSMG0001691 | Lancl1 | LanC (bacterial lantibiotic synthetase component C)-like 1 | 2354.23 | 1386.99 | 0.76 | 1.64E-06 |
| ENSMUSG00000062209 | GSMG0001693 | Erbb4 | erb-b2 receptor tyrosine kinase 4 | 351.30 | 470.89 | -0.42 | 9.82E-02 |
| ENSMUSG00000026193 | GSMG0001727 | Fn1 | fibronectin 1 | 68359.91 | 51452.58 | 0.41 | 1.22E-01 |
| ENSMUSG00000026189 | GSMG0001735 | Pecr | peroxisomal trans-2-enoyl-CoA reductase | 70.85 | 119.04 | -0.74 | 2.13E-02 |
| ENSMUSG00000039372 | GSMG0001736 | March4 | membrane-associated ring finger (C3HC4) 4 | 114.38 | 271.66 | -1.25 | 4.31E-08 |
| ENSMUSG00000026185 | GSMG0001738 | Igfbp5 | insulin-like growth factor binding protein 5 | 1450.14 | 1585.25 | -0.13 | 6.84E-01 |
| ENSMUSG00000006299 | GSMG0001752 | Aamp | angio-associated migratory protein | 2068.13 | 2148.30 | -0.05 | 8.58E-01 |
| NULL | GSMG0001753 | NULL | NULL | 89.96 | 75.61 | 0.24 | 6.10E-01 |
| ENSMUSG00000006301 | GSMG0001754 | Tmbim1 | transmembrane BAX inhibitor motif containing 1 | 714.44 | 451.12 | 0.66 | 9.35E-05 |
| ENSMUSG00000026171 | GSMG0001755 | Rnf25 | ring finger protein 25 | 340.13 | 352.15 | -0.05 | 8.97E-01 |
| ENSMUSG00000033364 | GSMG0001758 | Usp37 | ubiquitin specific peptidase 37 | 361.87 | 367.19 | -0.02 | 9.61E-01 |
| ENSMUSG00000026135 | GSMG0001760 | Zfp142 | zinc finger protein 142 | 480.00 | 488.80 | -0.02 | 9.51E-01 |
| ENSMUSG00000033159 | GSMG0001767 | Cnppd1 | cyclin Pas1/PHO80 domain containing 1 | 741.22 | 673.64 | 0.14 | 5.95E-01 |
| ENSMUSG00000026198 | GSMG0001768 | Abcb6 | ATP-binding cassette, sub-family B (MDR/TAP), member 6 | 348.75 | 442.17 | -0.34 | 1.98E-01 |
| ENSMUSG00000033124 | GSMG0001769 | Atg9a | autophagy related 9A | 2737.26 | 2669.89 | 0.04 | 9.08E-01 |
| ENSMUSG00000026200 | GSMG0001770 | Glb1l | galactosidase, beta 1-like | 567.67 | 490.37 | 0.21 | 3.96E-01 |
| ENSMUSG00000026202 | GSMG0001771 | Tuba4a | tubulin, alpha 4A | 455.67 | 199.04 | 1.20 | 1.17E-08 |

| | | | | | | | |
|---|---|---|---|---|---|---|---|
| ENSMUSG00000026204 | GSMG0001773 | Ptprn | protein tyrosine phosphatase, receptor type, N | 2926.68 | 1558.64 | 0.91 | 1.81E-06 |
| ENSMUSG00000033061 | GSMG0001774 | Resp18 | regulated endocrine-specific protein 18 | 244.66 | 261.11 | -0.10 | 8.12E-01 |
| ENSMUSG00000026209 | GSMG0001775 | Dnpep | aspartyl aminopeptidase | 2718.27 | 2645.31 | 0.04 | 8.89E-01 |
| ENSMUSG00000032997 | GSMG0001779 | Chpf | chondroitin polymerizing factor | 1901.47 | 1901.39 | 0.00 | 1.00E+00 |
| ENSMUSG00000026211 | GSMG0001780 | Obsl1 | obscurin-like 1 | 667.01 | 682.18 | -0.03 | 9.38E-01 |
| ENSMUSG00000026235 | GSMG0001783 | Epha4 | Eph receptor A4 | 1187.04 | 1138.12 | 0.06 | 8.59E-01 |
| ENSMUSG00000026245 | GSMG0001789 | Farsb | phenylalanyl-tRNA synthetase, beta subunit | 1473.59 | 1906.53 | -0.37 | 8.40E-02 |
| NULL | GSMG0001790 | NULL | NULL | 119.24 | 136.60 | -0.19 | 6.89E-01 |
| ENSMUSG00000050711 | GSMG0001794 | Scg2 | secretogranin II | 421.74 | 422.48 | 0.00 | 9.93E-01 |
| ENSMUSG00000073643 | GSMG0001797 | Wdfy1 | WD repeat and FYVE domain containing 1 | 804.10 | 1045.66 | -0.38 | 3.49E-02 |
| ENSMUSG00000026249 | GSMG0001798 | Serpine2 | serine (or cysteine) peptidase inhibitor, clade E, member 2 | 14924.90 | 14620.72 | 0.03 | 9.30E-01 |
| ENSMUSG00000043230 | GSMG0001801 | Fam124b | family with sequence similarity 124, member B | 721.71 | 1050.15 | -0.54 | 3.23E-03 |
| ENSMUSG00000004364 | GSMG0001803 | Cul3 | cullin 3 | 1781.90 | 1162.26 | 0.62 | 1.64E-02 |
| ENSMUSG00000038608 | GSMG0001805 | Dock10 | dedicator of cytokinesis 10 | 1048.06 | 793.71 | 0.40 | 7.53E-02 |
| ENSMUSG00000038496 | GSMG0001823 | Slc19a3 | solute carrier family 19, member 3 | 1580.96 | 477.69 | 1.73 | 3.15E-26 |
| ENSMUSG00000026163 | GSMG0001826 | Sphkap | SPHK1 interactor, AKAP domain containing | 255.74 | 220.67 | 0.21 | 4.76E-01 |
| ENSMUSG00000045658 | GSMG0001830 | Pid1 | phosphotyrosine interaction domain containing 1 | 1291.00 | 1639.05 | -0.34 | 4.55E-02 |
| NULL | GSMG0001835 | NULL | NULL | 237.02 | 228.07 | 0.06 | 8.87E-01 |
| ENSMUSG00000036766 | GSMG0001836 | Dner | delta/notch-like EGF repeat containing | 537.36 | 1361.53 | -1.34 | 9.41E-10 |
| ENSMUSG00000026219 | GSMG0001840 | Trip12 | thyroid hormone receptor interactor 12 | 5143.37 | 6176.71 | -0.26 | 1.22E-01 |
| NULL | GSMG0001841 | NULL | NULL | 162.02 | 125.10 | 0.37 | 4.03E-01 |
| ENSMUSG00000026234 | GSMG0001857 | Ncl | nucleolin | 6536.97 | 10084.63 | -0.63 | 8.20E-04 |
| ENSMUSG00000026239 | GSMG0001860 | Pde6d | phosphodiesterase 6D, cGMP-specific, rod, delta | 1237.94 | 1513.42 | -0.29 | 1.95E-01 |
| ENSMUSG00000026241 | GSMG0001862 | Nppc | natriuretic peptide type C | 128.40 | 87.84 | 0.55 | 9.56E-02 |
| ENSMUSG00000026259 | GSMG0001868 | Ngef | neuronal guanine nucleotide exchange factor | 2895.79 | 1953.96 | 0.57 | 1.75E-03 |
| NULL | GSMG0001869 | NULL | NULL | 224.57 | 91.96 | 1.28 | 3.24E-05 |
| ENSMUSG00000005501 | GSMG0001871 | Usp40 | ubiquitin specific peptidase 40 | 485.97 | 561.01 | -0.21 | 3.98E-01 |
| ENSMUSG00000044783 | GSMG0001873 | Hjurp | Holliday junction recognition protein | 1289.31 | 2364.62 | -0.88 | 2.54E-09 |
| ENSMUSG00000026309 | GSMG0001892 | Ilkap | integrin-linked kinase-associated serine/threonine phosphatase 2C | 1287.62 | 1336.25 | -0.05 | 8.53E-01 |
| ENSMUSG00000067071 | GSMG0001894 | Hes6 | hairy and enhancer of split 6 | 99.20 | 183.51 | -0.90 | 8.34E-04 |
| ENSMUSG00000055866 | GSMG0001895 | Per2 | period circadian clock 2 | 856.50 | 380.55 | 1.17 | 2.85E-08 |
| NULL | GSMG0001896 | NULL | NULL | 122.34 | 112.22 | 0.12 | 8.23E-01 |
| ENSMUSG00000026313 | GSMG0001898 | Hdac4 | histone deacetylase 4 | 1612.31 | 1275.20 | 0.34 | 1.97E-01 |

| | | | | | | | |
|---|---|---|---|---|---|---|---|
| ENSMUSG00000026260 | GSMG0001901 | Ndufa10 | NADH dehydrogenase (ubiquinone) 1 alpha subcomplex 10 | 1837.69 | 2139.53 | -0.22 | 2.65E-01 |
| ENSMUSG00000073616 | GSMG0001905 | Myeov2 | myeloma overexpressed 2 | 1267.28 | 1405.67 | -0.15 | 5.41E-01 |
| ENSMUSG00000014602 | GSMG0001912 | Kif1a | kinesin family member 1A | 4670.43 | 7498.02 | -0.68 | 1.91E-03 |
| ENSMUSG00000026273 | GSMG0001914 | Mterf4 | mitochondrial transcription termination factor 4 | 206.82 | 209.11 | -0.01 | 9.77E-01 |
| ENSMUSG00000026274 | GSMG0001915 | Pask | PAS domain containing serine/threonine kinase | 49.55 | 130.25 | -1.38 | 4.78E-05 |
| ENSMUSG00000034088 | GSMG0001916 | Hdlbp | high density lipoprotein (HDL) binding protein | 5236.36 | 5943.70 | -0.18 | 3.19E-01 |
| ENSMUSG00000026277 | GSMG0001919 | Stk25 | serine/threonine kinase 25 (yeast) | 4212.16 | 4760.39 | -0.18 | 4.51E-01 |
| ENSMUSG00000026279 | GSMG0001920 | Thap4 | THAP domain containing 4 | 597.64 | 859.24 | -0.52 | 7.89E-03 |
| ENSMUSG00000026281 | GSMG0001921 | Dtymk | deoxythymidylate kinase | 858.97 | 1274.32 | -0.57 | 1.87E-03 |
| ENSMUSG00000040710 | GSMG0001924 | St8sia4 | ST8 alpha-N-acetyl-neuraminide alpha-2,8-sialyltransferase 4 | 930.59 | 777.48 | 0.26 | 3.12E-01 |
| ENSMUSG00000044768 | GSMG0001937 | D1Ertd622e | DNA segment, Chr 1, ERATO Doi 622, expressed | 746.63 | 939.34 | -0.33 | 8.99E-02 |
| ENSMUSG00000040648 | GSMG0001938 | Ppip5k2 | diphosphoinositol pentakisphosphate kinase 2 | 986.52 | 1464.61 | -0.57 | 8.63E-04 |
| ENSMUSG00000026335 | GSMG0001940 | Pam | peptidylglycine alpha-amidating monooxygenase | 2323.20 | 2656.10 | -0.19 | 8.69E-01 |
| NULL | GSMG0001941 | NULL | NULL | 200.47 | 227.86 | -0.18 | 6.05E-01 |
| ENSMUSG00000089706 | GSMG0001942 | B230216N24Rik | RIKEN cDNA B230216N24 gene | 207.02 | 265.10 | -0.36 | 3.23E-01 |
| NULL | GSMG0001945 | NULL | NULL | 401.74 | 554.76 | -0.47 | 2.45E-02 |
| ENSMUSG00000056536 | GSMG0001949 | Pign | phosphatidylinositol glycan anchor biosynthesis, class N | 218.48 | 213.39 | 0.03 | 9.41E-01 |
| ENSMUSG00000057329 | GSMG0001955 | Bcl2 | B cell leukemia/lymphoma 2 | 2010.55 | 1943.67 | 0.05 | 8.66E-01 |
| ENSMUSG00000009905 | GSMG0001962 | Kdsr | 3-ketodihydrosphingosine reductase | 700.99 | 690.25 | 0.02 | 9.42E-01 |
| ENSMUSG00000009907 | GSMG0001963 | Vps4b | vacuolar protein sorting 4B | 1502.90 | 1442.77 | 0.06 | 8.52E-01 |
| ENSMUSG00000038702 | GSMG0001974 | Dsel | dermatan sulfate epimerase-like | 67.48 | 85.46 | -0.34 | 4.47E-01 |
| ENSMUSG00000026374 | GSMG0001982 | Tsn | translin | 2898.44 | 3958.19 | -0.45 | 1.70E-02 |
| ENSMUSG00000004451 | GSMG0001987 | Ralb | v-ral simian leukemia viral oncogene B | 1341.83 | 2077.48 | -0.63 | 7.23E-05 |
| ENSMUSG00000026383 | GSMG0001991 | Epb41l5 | erythrocyte membrane protein band 4.1 like 5 | 89.86 | 168.72 | -0.90 | 1.99E-03 |
| ENSMUSG00000026384 | GSMG0001994 | Ptpn4 | protein tyrosine phosphatase, non-receptor type 4 | 385.05 | 351.44 | 0.13 | 6.54E-01 |
| ENSMUSG00000050777 | GSMG0002003 | Tmem37 | transmembrane protein 37 | 1356.64 | 1751.95 | -0.37 | 3.21E-02 |
| ENSMUSG00000026385 | GSMG0002004 | Dbi | diazepam binding inhibitor | 2906.86 | 4597.69 | -0.66 | 1.34E-05 |
| ENSMUSG00000026389 | GSMG0002005 | Steap3 | STEAP family member 3 | 799.56 | 650.28 | 0.30 | 2.46E-01 |
| ENSMUSG00000003721 | GSMG0002008 | Insig2 | insulin induced gene 2 | 533.39 | 418.65 | 0.35 | 1.30E-01 |
| ENSMUSG00000036815 | GSMG0002013 | Dpp10 | dipeptidylpeptidase 10 | 557.26 | 760.58 | -0.45 | 5.83E-02 |
| NULL | GSMG0002020 | NULL | NULL | 150.88 | 126.77 | 0.25 | 6.15E-01 |
| NULL | GSMG0002025 | NULL | NULL | 1376.22 | 532.58 | 1.37 | 3.65E-11 |
| NULL | GSMG0002026 | NULL | NULL | 143.85 | 61.46 | 1.24 | 4.36E-04 |

| | | | | | | | |
|---|---|---|---|---|---|---|---|
| NULL | GSMG0002027 | NULL | NULL | 534.22 | 218.57 | 1.29 | 4.45E-09 |
| ENSMUSG00000026341 | GSMG0002029 | Actr3 | ARP3 actin-related protein 3 | 2448.92 | 3308.21 | -0.43 | 1.17E-02 |
| ENSMUSG00000026344 | GSMG0002033 | Lypd1 | Ly6/Plaur domain containing 1 | 883.74 | 855.93 | 0.05 | 8.85E-01 |
| ENSMUSG00000049690 | GSMG0002034 | Nckap5 | NCK-associated protein 5 | 790.34 | 1694.46 | -1.10 | 5.37E-09 |
| ENSMUSG00000026347 | GSMG0002044 | Tmem163 | transmembrane protein 163 | 135.46 | 105.76 | 0.36 | 3.57E-01 |
| NULL | GSMG0002045 | NULL | NULL | 198.23 | 163.98 | 0.28 | 4.15E-01 |
| ENSMUSG00000026355 | GSMG0002057 | Mcm6 | minichromosome maintenance complex component 6 | 705.98 | 1212.88 | -0.78 | 5.34E-05 |
| ENSMUSG00000026356 | GSMG0002058 | Dars | aspartyl-tRNA synthetase | 1909.74 | 2665.61 | -0.48 | 1.17E-02 |
| ENSMUSG00000045382 | GSMG0002059 | Cxcr4 | chemokine (C-X-C motif) receptor 4 | 786.13 | 918.53 | -0.22 | 2.80E-01 |
| ENSMUSG00000026399 | GSMG0002064 | Cd55 | CD55 molecule, decay accelerating factor for complement | 499.26 | 271.36 | 0.88 | 4.84E-05 |
| G00000026409,ENSMUSG000 | GSMG0002067 | C4bp-ps1 // Pfkfb2 | complement component 4 binding protein, pseudogene 1 // 6-phosphofi | 302.63 | 285.49 | 0.09 | 8.05E-01 |
| ENSMUSG00000016528 | GSMG0002072 | Mapkapk2 | MAP kinase-activated protein kinase 2 | 2264.78 | 1863.61 | 0.28 | 3.76E-01 |
| ENSMUSG00000026425 | GSMG0002078 | Srgap2 | SLIT-ROBO Rho GTPase activating protein 2 | 1092.80 | 946.81 | 0.21 | 4.84E-01 |
| NULL | GSMG0002080 | NULL | NULL | 144.11 | 224.25 | -0.65 | 6.01E-02 |
| NULL | GSMG0002081 | NULL | NULL | 254.07 | 322.90 | -0.35 | 2.63E-01 |
| ENSMUSG00000059149 | GSMG0002084 | Mfsd4a | major facilitator superfamily domain containing 4A | 245.08 | 155.45 | 0.66 | 1.07E-02 |
| ENSMUSG00000026437 | GSMG0002085 | Cdk18 | cyclin-dependent kinase 18 | 227.52 | 151.00 | 0.59 | 5.14E-02 |
| ENSMUSG00000042066 | GSMG0002088 | Tmcc2 | transmembrane and coiled-coil domains 2 | 510.99 | 678.70 | -0.41 | 1.47E-01 |
| NULL | GSMG0002091 | NULL | NULL | 916.03 | 841.66 | 0.12 | 6.45E-01 |
| ENSMUSG00000053024 | GSMG0002093 | Cntn2 | contactin 2 | 260.35 | 460.02 | -0.82 | 9.91E-04 |
| ENSMUSG00000026442 | GSMG0002094 | Nfasc | neurofascin | 1460.22 | 1251.23 | 0.22 | 5.26E-01 |
| NULL | GSMG0002095 | NULL | NULL | 180.85 | 165.26 | 0.13 | 7.85E-01 |
| ENSMUSG00000054387 | GSMG0002097 | Mdm4 | transformed mouse 3T3 cell double minute 4 | 279.40 | 243.09 | 0.21 | 5.62E-01 |
| NULL | GSMG0002098 | NULL | NULL | 1256.85 | 1286.94 | -0.03 | 9.09E-01 |
| ENSMUSG00000070643 | GSMG0002101 | Sox13 | SRY (sex determining region Y)-box 13 | 1487.89 | 1016.76 | 0.55 | 1.81E-02 |
| ENSMUSG00000090553 | GSMG0002102 | Snrpe | small nuclear ribonucleoprotein E | 573.30 | 847.44 | -0.56 | 1.53E-03 |
| ENSMUSG00000102976 | GSMG0002103 | Zc3h11a | zinc finger CCCH type containing 11A | 1369.20 | 1430.53 | -0.06 | 8.29E-01 |
| ENSMUSG00000026463 | GSMG0002107 | Atp2b4 | ATPase, Ca++ transporting, plasma membrane 4 | 421.19 | 361.08 | 0.22 | 5.82E-01 |
| ENSMUSG00000041577 | GSMG0002110 | Prelp | proline arginine-rich end leucine-rich repeat | 377.01 | 103.56 | 1.87 | 1.53E-14 |
| ENSMUSG00000020423 | GSMG0002111 | Btg2 | B cell translocation gene 2, anti-proliferative | 3394.19 | 1131.95 | 1.58 | 6.82E-19 |
| ENSMUSG00000042429 | GSMG0002113 | Adora1 | adenosine A1 receptor | 1810.88 | 1764.03 | 0.04 | 9.10E-01 |
| ENSMUSG00000026458 | GSMG0002114 | Ppfia4 | protein tyrosine phosphatase, receptor type, f polypeptide (PTPRF), inte | 67.25 | 183.20 | -1.45 | 1.49E-07 |
| ENSMUSG00000042305 | GSMG0002115 | Tmem183a | transmembrane protein 183A | 844.79 | 1122.03 | -0.41 | 1.46E-02 |

| | | | | | | | |
|---|---|---|---|---|---|---|---|
| NULL | GSMG0002119 | NULL | NULL | 1145.22 | 952.21 | 0.27 | 1.89E-01 |
| ENSMUSG00000073557 | GSMG0002120 | Ppp1r12b | protein phosphatase 1, regulatory (inhibitor) subunit 12B | 314.39 | 271.33 | 0.21 | 5.75E-01 |
| NULL | GSMG0002121 | NULL | NULL | 209.84 | 181.03 | 0.21 | 5.43E-01 |
| NULL | GSMG0002122 | NULL | NULL | 634.07 | 632.74 | 0.00 | 9.99E-01 |
| ENSMUSG00000026424 | GSMG0002127 | Gpr37l1 | G protein-coupled receptor 37-like 1 | 2954.73 | 885.66 | 1.74 | 3.82E-26 |
| ENSMUSG00000041926 | GSMG0002129 | Rnpep | arginyl aminopeptidase (aminopeptidase B) | 2226.52 | 1778.00 | 0.33 | 1.08E-01 |
| ENSMUSG00000062580 | GSMG0002130 | Timm17a | translocase of inner mitochondrial membrane 17a | 1692.01 | 1906.01 | -0.17 | 4.83E-01 |
| ENSMUSG00000041889 | GSMG0002131 | Shisa4 | shisa family member 4 | 1349.70 | 950.64 | 0.51 | 4.48E-03 |
| ENSMUSG00000041879 | GSMG0002133 | Ipo9 | importin 9 | 6955.27 | 7077.35 | -0.02 | 9.41E-01 |
| NULL | GSMG0002136 | NULL | NULL | 142.85 | 142.79 | 0.00 | 1.00E+00 |
| NULL | GSMG0002137 | NULL | NULL | 99.12 | 111.15 | -0.16 | 7.30E-01 |
| NULL | GSMG0002138 | NULL | NULL | 352.12 | 353.65 | 0.00 | 9.95E-01 |
| NULL | GSMG0002139 | NULL | NULL | 284.64 | 247.63 | 0.20 | 6.22E-01 |
| NULL | GSMG0002141 | NULL | NULL | 639.32 | 530.34 | 0.27 | 2.79E-01 |
| ENSMUSG00000041570 | GSMG0002151 | Camsap2 | calmodulin regulated spectrin-associated protein family, member 2 | 1901.98 | 2070.17 | -0.12 | 6.57E-01 |
| ENSMUSG00000026393 | GSMG0002165 | Nek7 | NIMA (never in mitosis gene a)-related expressed kinase 7 | 606.55 | 407.47 | 0.58 | 4.79E-03 |
| ENSMUSG00000026365 | GSMG0002181 | Cfh | complement component factor h | 1103.38 | 335.76 | 1.72 | 7.87E-18 |
| ENSMUSG00000026361 | GSMG0002186 | Cdc73 | cell division cycle 73, Paf1/RNA polymerase II complex component | 535.29 | 405.87 | 0.40 | 1.52E-01 |
| ENSMUSG00000018199 | GSMG0002189 | Trove2 | TROVE domain family, member 2 | 979.94 | 1174.33 | -0.26 | 2.47E-01 |
| ENSMUSG00000026360 | GSMG0002190 | Rgs2 | regulator of G-protein signaling 2 | 606.09 | 810.80 | -0.42 | 3.87E-02 |
| ENSMUSG00000056220 | GSMG0002206 | Pla2g4a | phospholipase A2, group IVA (cytosolic, calcium-dependent) | 346.77 | 111.41 | 1.64 | 2.79E-11 |
| ENSMUSG00000006010 | GSMG0002210 | BC003331 | cDNA sequence BC003331 | 607.81 | 647.50 | -0.09 | 7.59E-01 |
| ENSMUSG00000066842 | GSMG0002213 | Hmcn1 | hemicentin 1 | 3077.86 | 2796.18 | 0.14 | 5.39E-01 |
| ENSMUSG00000052748 | GSMG0002217 | Swt1 | SWT1 RNA endoribonuclease homolog (S. cerevisiae) | 1003.34 | 736.61 | 0.45 | 6.23E-02 |
| ENSMUSG00000026484 | GSMG0002218 | Rnf2 | ring finger protein 2 | 466.92 | 722.57 | -0.63 | 1.60E-03 |
| ENSMUSG00000014980 | GSMG0002226 | Tsen15 | tRNA splicing endonuclease subunit 15 | 105.06 | 83.66 | 0.33 | 4.29E-01 |
| ENSMUSG00000026482 | GSMG0002227 | Rgl1 | ral guanine nucleotide dissociation stimulator,-like 1 | 3611.13 | 4417.71 | -0.29 | 1.10E-01 |
| NULL | GSMG0002228 | NULL | NULL | 554.35 | 710.63 | -0.36 | 1.46E-01 |
| ENSMUSG00000042772 | GSMG0002231 | Smg7 | Smg-7 homolog, nonsense mediated mRNA decay factor (C. elegans) | 1747.19 | 1115.14 | 0.65 | 4.19E-05 |
| ENSMUSG00000026478 | GSMG0002234 | Lamc1 | laminin, gamma 1 | 3790.52 | 3528.25 | 0.10 | 7.61E-01 |
| ENSMUSG00000042699 | GSMG0002238 | Dhx9 | DEAH (Asp-Glu-Ala-His) box polypeptide 9 | 325.68 | 453.62 | -0.48 | 5.20E-02 |
| ENSMUSG00000042684 | GSMG0002239 | Npl | N-acetylneuraminate pyruvate lyase | 128.71 | 273.10 | -1.08 | 1.16E-05 |
| ENSMUSG00000004110 | GSMG0002248 | Cacna1e | calcium channel, voltage-dependent, R type, alpha 1E subunit | 1212.14 | 1102.39 | 0.14 | 6.34E-01 |

| | | | | | | | |
|---|---|---|---|---|---|---|---|
| NULL | GSMG0002249 | NULL | NULL | 260.30 | 144.04 | 0.85 | 1.15E-04 |
| ENSMUSG00000056708 | GSMG0002251 | Ier5 | immediate early response 5 | 2088.89 | 1531.72 | 0.45 | 6.42E-03 |
| ENSMUSG00000026471 | GSMG0002252 | Mr1 | major histocompatibility complex, class I-related | 238.41 | 165.83 | 0.53 | 3.26E-02 |
| ENSMUSG00000026469 | GSMG0002255 | Xpr1 | xenotropic and polytropic retrovirus receptor 1 | 983.21 | 987.54 | -0.01 | 9.88E-01 |
| ENSMUSG00000033684 | GSMG0002260 | Qsox1 | quiescin Q6 sulfhydryl oxidase 1 | 1476.52 | 860.74 | 0.78 | 1.93E-05 |
| ENSMUSG00000026466 | GSMG0002265 | Tor1aip1 | torsin A interacting protein 1 | 1705.00 | 1365.67 | 0.32 | 9.62E-02 |
| ENSMUSG00000026600 | GSMG0002271 | Soat1 | sterol O-acyltransferase 1 | 285.95 | 253.36 | 0.18 | 6.05E-01 |
| ENSMUSG00000060519 | GSMG0002272 | Tor3a | torsin family 3, member A | 382.39 | 322.71 | 0.24 | 3.74E-01 |
| ENSMUSG00000033557 | GSMG0002273 | Fam20b | family with sequence similarity 20, member B | 463.61 | 397.31 | 0.22 | 3.87E-01 |
| ENSMUSG00000026594 | GSMG0002276 | Ralgps2 | Ral GEF with PH domain and SH3 binding motif 2 | 561.55 | 910.39 | -0.70 | 4.68E-05 |
| ENSMUSG00000070565 | GSMG0002283 | Rasal2 | RAS protein activator like 2 | 1613.45 | 1207.24 | 0.42 | 1.22E-02 |
| NULL | GSMG0002284 | NULL | NULL | 76.04 | 44.93 | 0.75 | 8.95E-02 |
| NULL | GSMG0002289 | NULL | NULL | 441.32 | 199.84 | 1.15 | 2.25E-07 |
| ENSMUSG00000004031 | GSMG0002290 | Brinp2 | bone morphogenic protein/retinoic acid inducible neural-specific 2 | 487.63 | 426.95 | 0.19 | 5.79E-01 |
| ENSMUSG00000026729 | GSMG0002296 | 4930562F07Rik | RIKEN cDNA 4930562F07 gene | 184.11 | 155.05 | 0.25 | 4.88E-01 |
| ENSMUSG00000014226 | GSMG0002298 | Cacybp | calcyclin binding protein | 1215.31 | 1913.52 | -0.66 | 7.22E-04 |
| ENSMUSG00000043467 | GSMG0002314 | Zbtb37 | zinc finger and BTB domain containing 37 | 164.91 | 197.97 | -0.27 | 4.36E-01 |
| ENSMUSG00000026709 | GSMG0002315 | Dars2 | aspartyl-tRNA synthetase 2 (mitochondrial) | 383.58 | 375.08 | 0.03 | 9.29E-01 |
| ENSMUSG00000026705 | GSMG0002316 | Klhl20 | kelch-like 20 | 677.94 | 617.27 | 0.14 | 6.15E-01 |
| ENSMUSG00000026701 | GSMG0002319 | Prdx6 | peroxiredoxin 6 | 3563.77 | 3049.35 | 0.23 | 2.36E-01 |
| ENSMUSG00000040297 | GSMG0002322 | Suco | SUN domain containing ossification factor | 919.18 | 1053.55 | -0.20 | 4.36E-01 |
| ENSMUSG00000040265 | GSMG0002324 | Dnm3 | dynamin 3 | 1698.85 | 1609.29 | 0.08 | 7.72E-01 |
| NULL | GSMG0002334 | NULL | NULL | 79.71 | 99.55 | -0.32 | 4.83E-01 |
| ENSMUSG00000026694 | GSMG0002335 | Mettl13 | methyltransferase like 13 | 360.91 | 518.51 | -0.52 | 3.05E-02 |
| ENSMUSG00000040181 | GSMG0002342 | Fmo1 | flavin containing monooxygenase 1 | 342.16 | 305.52 | 0.17 | 6.47E-01 |
| ENSMUSG00000040124 | GSMG0002351 | Gorab | golgin, RAB6-interacting | 74.47 | 96.95 | -0.39 | 4.44E-01 |
| ENSMUSG00000041406 | GSMG0002355 | BC055324 | cDNA sequence BC055324 | 130.80 | 234.71 | -0.84 | 3.20E-03 |
| ENSMUSG00000026577 | GSMG0002358 | Blzf1 | basic leucine zipper nuclear factor 1 | 616.08 | 649.71 | -0.08 | 7.96E-01 |
| ENSMUSG00000026576 | GSMG0002359 | Atp1b1 | ATPase, Na+/K+ transporting, beta 1 polypeptide | 3040.01 | 3100.86 | -0.03 | 9.26E-01 |
| NULL | GSMG0002360 | NULL | NULL | 887.94 | 687.93 | 0.37 | 9.24E-02 |
| ENSMUSG00000040848 | GSMG0002365 | Sft2d2 | SFT2 domain containing 2 | 2687.51 | 2575.28 | 0.06 | 8.03E-01 |
| ENSMUSG00000040843 | GSMG0002366 | Tiprl | TIP41, TOR signalling pathway regulator-like (S. cerevisiae) | 641.79 | 720.51 | -0.17 | 5.03E-01 |
| ENSMUSG00000026571 | GSMG0002368 | Dcaf6 | DDB1 and CUL4 associated factor 6 | 420.28 | 316.77 | 0.41 | 8.63E-02 |

| | | | | | | | |
|---|---|---|---|---|---|---|---|
| ENSMUSG00000026566 | GSMG0002373 | Mpzl1 | myelin protein zero-like 1 | 3438.61 | 3803.52 | -0.15 | 5.16E-01 |
| NULL | GSMG0002374 | NULL | NULL | 443.22 | 335.43 | 0.40 | 2.22E-01 |
| ENSMUSG00000040723 | GSMG0002375 | Rcsd1 | RCSD domain containing 1 | 780.00 | 629.42 | 0.31 | 1.23E-01 |
| ENSMUSG00000026565 | GSMG0002382 | Pou2f1 | POU domain, class 2, transcription factor 1 | 553.06 | 700.75 | -0.34 | 7.83E-02 |
| ENSMUSG00000026564 | GSMG0002388 | Dusp27 | dual specificity phosphatase 27 (putative) | 65.43 | 195.17 | -1.57 | 3.66E-09 |
| ENSMUSG00000040596 | GSMG0002393 | Pogk | pogo transposable element with KRAB domain | 233.51 | 439.44 | -0.91 | 5.05E-05 |
| NULL | GSMG0002399 | NULL | NULL | 236.53 | 349.15 | -0.56 | 1.35E-02 |
| ENSMUSG00000026558 | GSMG0002400 | Uck2 | uridine-cytidine kinase 2 | 183.84 | 382.33 | -1.06 | 2.31E-07 |
| NULL | GSMG0002401 | NULL | NULL | 365.21 | 291.86 | 0.32 | 3.66E-01 |
| NULL | GSMG0002402 | NULL | NULL | 817.64 | 690.22 | 0.24 | 3.43E-01 |
| ENSMUSG00000026688 | GSMG0002404 | Mgst3 | microsomal glutathione S-transferase 3 | 709.79 | 667.14 | 0.09 | 7.49E-01 |
| ENSMUSG00000052534 | GSMG0002407 | Pbx1 | pre B cell leukemia homeobox 1 | 84.65 | 124.22 | -0.56 | 1.10E-01 |
| NULL | GSMG0002409 | NULL | NULL | 249.94 | 365.95 | -0.55 | 1.97E-02 |
| NULL | GSMG0002412 | NULL | NULL | 280.46 | 414.80 | -0.56 | 1.15E-02 |
| NULL | GSMG0002413 | NULL | NULL | 284.15 | 389.41 | -0.46 | 7.27E-02 |
| NULL | GSMG0002417 | NULL | NULL | 167.50 | 245.48 | -0.55 | 7.08E-02 |
| NULL | GSMG0002418 | NULL | NULL | 190.96 | 318.76 | -0.73 | 2.72E-03 |
| NULL | GSMG0002419 | NULL | NULL | 567.41 | 763.57 | -0.43 | 2.38E-02 |
| ENSMUSG00000026683 | GSMG0002421 | Nuf2 | NUF2, NDC80 kinetochore complex component | 199.36 | 569.86 | -1.51 | 3.34E-11 |
| ENSMUSG00000038530 | GSMG0002422 | Rgs4 | regulator of G-protein signaling 4 | 9886.84 | 4923.44 | 1.01 | 2.69E-08 |
| ENSMUSG00000026675 | GSMG0002424 | Hsd17b7 | hydroxysteroid (17-beta) dehydrogenase 7 | 148.32 | 273.02 | -0.88 | 3.18E-04 |
| ENSMUSG00000026674 | GSMG0002426 | Ddr2 | discoidin domain receptor family, member 2 | 1232.32 | 960.79 | 0.36 | 1.75E-01 |
| ENSMUSG00000026670 | GSMG0002427 | Uap1 | UDP-N-acetylglucosamine pyrophosphorylase 1 | 1150.29 | 1065.78 | 0.11 | 6.57E-01 |
| ENSMUSG00000026667 | GSMG0002429 | Uhmk1 | U2AF homology motif (UHM) kinase 1 | 300.47 | 286.44 | 0.07 | 8.85E-01 |
| ENSMUSG00000038473 | GSMG0002431 | Nos1ap | nitric oxide synthase 1 (neuronal) adaptor protein | 383.47 | 224.08 | 0.77 | 2.26E-04 |
| ENSMUSG00000026663 | GSMG0002433 | Atf6 | activating transcription factor 6 | 548.37 | 637.60 | -0.22 | 4.10E-01 |
| ENSMUSG00000026659 | GSMG0002436 | Dusp12 | dual specificity phosphatase 12 | 260.88 | 363.82 | -0.48 | 3.19E-02 |
| ENSMUSG00000026656 | GSMG0002439 | Fcgr2b | Fc receptor, IgG, low affinity IIb | 126.39 | 33.90 | 1.91 | 4.54E-08 |
| ENSMUSG00000059498 | GSMG0002440 | Fcgr3 | Fc receptor, IgG, low affinity III | 170.07 | 57.28 | 1.57 | 1.24E-08 |
| ENSMUSG00000058076 | GSMG0002441 | Sdhc | succinate dehydrogenase complex, subunit C, integral membrane protei | 2084.46 | 2194.68 | -0.07 | 8.08E-01 |
| ENSMUSG00000038370 | GSMG0002444 | Pcp4l1 | Purkinje cell protein 4-like 1 | 139.41 | 39.66 | 1.81 | 1.12E-07 |
| ENSMUSG00000005674 | GSMG0002445 | Tomm40l | translocase of outer mitochondrial membrane 40 homolog-like (yeast) | 188.18 | 169.89 | 0.15 | 7.21E-01 |
| ENSMUSG00000058715 | GSMG0002446 | Fcer1g | Fc receptor, IgE, high affinity I, gamma polypeptide | 294.57 | 152.66 | 0.95 | 7.33E-05 |

| | | | | | | | |
|---|---|---|---|---|---|---|---|
| ENSMUSG00000013593 | GSMG0002447 | Ndufs2 | NADH dehydrogenase (ubiquinone) Fe-S protein 2 | 4539.83 | 5179.38 | -0.19 | 3.16E-01 |
| ENSMUSG00000062729 | GSMG0002449 | Ppox | protoporphyrinogen oxidase | 1126.70 | 1026.51 | 0.14 | 5.77E-01 |
| ENSMUSG00000053483 | GSMG0002450 | Usp21 | ubiquitin specific peptidase 21 | 1607.71 | 1987.22 | -0.31 | 8.25E-02 |
| ENSMUSG00000062963 | GSMG0002451 | Ufc1 | ubiquitin-fold modifier conjugating enzyme 1 | 925.20 | 1120.75 | -0.28 | 1.97E-01 |
| ENSMUSG00000013997 | GSMG0002453 | Nit1 | nitrilase 1 | 366.74 | 351.55 | 0.06 | 8.65E-01 |
| ENSMUSG00000045259 | GSMG0002454 | Klhdc9 | kelch domain containing 9 | 150.97 | 101.59 | 0.57 | 1.33E-01 |
| ENSMUSG00000026556 | GSMG0002464 | Vangl2 | vang-like 2 (van gogh, Drosophila) | 684.77 | 754.23 | -0.14 | 5.84E-01 |
| ENSMUSG00000003458 | GSMG0002466 | Ncstn | nicastrin | 4483.65 | 4626.35 | -0.04 | 8.67E-01 |
| ENSMUSG00000013698 | GSMG0002468 | Pea15a | phosphoprotein enriched in astrocytes 15A | 18050.83 | 19870.24 | -0.14 | 5.13E-01 |
| ENSMUSG00000007097 | GSMG0002470 | Atp1a2 | ATPase, Na+/K+ transporting, alpha 2 polypeptide | 16145.16 | 9232.65 | 0.81 | 1.82E-06 |
| ENSMUSG00000038026 | GSMG0002471 | Kcnj9 | potassium inwardly-rectifying channel, subfamily J, member 9 | 1092.62 | 388.68 | 1.49 | 9.00E-12 |
| ENSMUSG00000026544 | GSMG0002478 | Dusp23 | dual specificity phosphatase 23 | 310.15 | 223.01 | 0.47 | 6.01E-02 |
| ENSMUSG00000037872 | GSMG0002483 | Ackr1 | atypical chemokine receptor 1 (Duffy blood group) | 630.02 | 1193.44 | -0.92 | 7.00E-08 |
| ENSMUSG00000005338 | GSMG0002484 | Cadm3 | cell adhesion molecule 3 | 284.02 | 400.22 | -0.49 | 4.22E-02 |
| NULL | GSMG0002496 | NULL | NULL | 304.99 | 198.14 | 0.63 | 7.72E-03 |
| ENSMUSG00000026527 | GSMG0002498 | Rgs7 | regulator of G protein signaling 7 | 556.17 | 659.45 | -0.24 | 2.99E-01 |
| NULL | GSMG0002500 | NULL | NULL | 197.41 | 134.42 | 0.55 | 8.19E-02 |
| NULL | GSMG0002501 | NULL | NULL | 380.14 | 262.60 | 0.54 | 3.11E-02 |
| ENSMUSG00000026526 | GSMG0002503 | Fh1 | fumarate hydratase 1 | 1529.17 | 1902.30 | -0.31 | 1.12E-01 |
| ENSMUSG00000026525 | GSMG0002504 | Opn3 | opsin 3 | 65.83 | 63.33 | 0.06 | 9.36E-01 |
| NULL | GSMG0002509 | NULL | NULL | 205.73 | 259.96 | -0.34 | 2.60E-01 |
| ENSMUSG00000019699 | GSMG0002521 | Akt3 | thymoma viral proto-oncogene 3 | 960.94 | 1109.39 | -0.21 | 3.18E-01 |
| ENSMUSG00000015961 | GSMG0002532 | Adss | adenylosuccinate synthetase, non muscle | 663.16 | 891.71 | -0.43 | 2.04E-02 |
| ENSMUSG00000039630 | GSMG0002538 | Hnrnpu | heterogeneous nuclear ribonucleoprotein U | 8365.30 | 10112.60 | -0.27 | 1.26E-01 |
| ENSMUSG00000055067 | GSMG0002542 | Smyd3 | SET and MYND domain containing 3 | 301.56 | 425.16 | -0.50 | 2.18E-02 |
| ENSMUSG00000026491 | GSMG0002550 | Ahctf1 | AT hook containing transcription factor 1 | 1713.13 | 1782.89 | -0.06 | 8.31E-01 |
| ENSMUSG00000026489 | GSMG0002552 | Adck3 | aarF domain containing kinase 3 | 146.68 | 88.57 | 0.73 | 1.50E-02 |
| ENSMUSG00000010609 | GSMG0002553 | Psen2 | presenilin 2 | 244.48 | 210.66 | 0.22 | 5.36E-01 |
| ENSMUSG00000038776 | GSMG0002562 | Ephx1 | epoxide hydrolase 1, microsomal | 1520.72 | 523.11 | 1.54 | 2.81E-15 |
| ENSMUSG00000026516 | GSMG0002565 | Nvl | nuclear VCP-like | 246.09 | 566.20 | -1.21 | 1.33E-10 |
| ENSMUSG00000038733 | GSMG0002566 | Wdr26 | WD repeat domain 26 | 3072.21 | 2664.06 | 0.21 | 3.29E-01 |
| ENSMUSG00000004880 | GSMG0002572 | Lbr | lamin B receptor | 1378.17 | 1770.02 | -0.36 | 8.92E-02 |
| ENSMUSG00000022995 | GSMG0002575 | Enah | enabled homolog (Drosophila) | 279.33 | 263.64 | 0.08 | 8.32E-01 |

| | | | | | | | |
|---|---|---|---|---|---|---|---|
| NULL | GSMG0002576 | NULL | NULL | 388.35 | 434.30 | -0.16 | 6.33E-01 |
| ENSMUSG00000038633 | GSMG0002581 | Degs1 | delta(4)-desaturase, sphingolipid 1 | 2273.42 | 2760.82 | -0.28 | 1.32E-01 |
| ENSMUSG00000047539 | GSMG0002582 | Fbxo28 | F-box protein 28 | 1515.12 | 1681.72 | -0.15 | 5.12E-01 |
| ENSMUSG00000026509 | GSMG0002584 | Capn2 | calpain 2 | 5098.50 | 4575.62 | 0.16 | 4.05E-01 |
| ENSMUSG00000039377 | GSMG0002589 | Hlx | H2.0-like homeobox | 333.74 | 310.53 | 0.10 | 8.31E-01 |
| ENSMUSG00000073481 | GSMG0002591 | March2 | mitochondrial amidoxime reducing component 2 | 3289.70 | 3512.47 | -0.09 | 7.17E-01 |
| ENSMUSG00000039349 | GSMG0002592 | C130074G19Rik | RIKEN cDNA C130074G19 gene | 523.41 | 482.16 | 0.12 | 7.56E-01 |
| ENSMUSG00000026620 | GSMG0002593 | Mark1 | MAP/microtubule affinity regulating kinase 1 | 820.03 | 466.48 | 0.81 | 2.50E-01 |
| ENSMUSG00000026618 | GSMG0002594 | Iars2 | isoleucine-tRNA synthetase 2, mitochondrial | 1954.13 | 1985.89 | -0.02 | 9.40E-01 |
| ENSMUSG00000039246 | GSMG0002598 | Lyplal1 | lysophospholipase-like 1 | 236.83 | 258.08 | -0.12 | 7.56E-01 |
| ENSMUSG00000039239 | GSMG0002601 | Tgfb2 | transforming growth factor, beta 2 | 489.89 | 407.75 | 0.27 | 3.03E-01 |
| ENSMUSG00000001305 | GSMG0002605 | Rrp15 | ribosomal RNA processing 15 homolog (S. cerevisiae) | 370.36 | 544.84 | -0.56 | 1.38E-02 |
| ENSMUSG00000026608 | GSMG0002617 | Kctd3 | potassium channel tetramerisation domain containing 3 | 786.82 | 981.68 | -0.32 | 9.62E-02 |
| ENSMUSG00000037624 | GSMG0002619 | Kcnk2 | potassium channel, subfamily K, member 2 | 107.37 | 120.72 | -0.16 | 7.31E-01 |
| ENSMUSG00000026605 | GSMG0002622 | Cenpf | centromere protein F | 98.34 | 232.51 | -1.24 | 1.75E-06 |
| ENSMUSG00000026603 | GSMG0002624 | Smyd2 | SET and MYND domain containing 2 | 1459.77 | 1701.54 | -0.22 | 2.85E-01 |
| ENSMUSG00000089872 | GSMG0002628 | Rps6kc1 | ribosomal protein S6 kinase polypeptide 1 | 481.17 | 545.54 | -0.18 | 4.72E-01 |
| ENSMUSG00000037568 | GSMG0002630 | Vash2 | vasohibin 2 | 25.13 | 169.02 | -2.77 | 8.86E-17 |
| ENSMUSG00000066595 | GSMG0002631 | Mfsd7b | major facilitator superfamily domain containing 7B | 139.23 | 117.70 | 0.24 | 5.42E-01 |
| ENSMUSG00000026632 | GSMG0002632 | Tatdn3 | TatD DNase domain containing 3 | 276.61 | 253.41 | 0.13 | 7.21E-01 |
| ENSMUSG00000026628 | GSMG0002634 | Atf3 | activating transcription factor 3 | 618.91 | 367.71 | 0.75 | 1.51E-03 |
| ENSMUSG00000037499 | GSMG0002635 | Nenf | neuron derived neurotrophic factor | 1560.66 | 1645.73 | -0.08 | 7.78E-01 |
| ENSMUSG00000026626 | GSMG0002636 | Ppp2r5a | protein phosphatase 2, regulatory subunit B', alpha | 1312.00 | 980.68 | 0.42 | 1.13E-01 |
| ENSMUSG00000037474 | GSMG0002641 | Dtl | denticleless E3 ubiquitin protein ligase | 136.73 | 374.65 | -1.45 | 3.65E-11 |
| ENSMUSG00000026637 | GSMG0002645 | Traf5 | TNF receptor-associated factor 5 | 273.05 | 179.91 | 0.60 | 1.36E-02 |
| ENSMUSG00000037395 | GSMG0002646 | Rcor3 | REST corepressor 3 | 310.28 | 370.75 | -0.26 | 3.02E-01 |
| ENSMUSG00000037375 | GSMG0002647 | Hhat | hedgehog acyltransferase | 153.21 | 175.64 | -0.19 | 6.37E-01 |
| ENSMUSG00000016262 | GSMG0002649 | Sertad4 | SERTA domain containing 4 | 307.42 | 432.15 | -0.49 | 2.75E-02 |
| ENSMUSG00000016181 | GSMG0002652 | Diexf | digestive organ expansion factor homolog (zebrafish) | 143.95 | 177.86 | -0.30 | 3.90E-01 |
| ENSMUSG00000016194 | GSMG0002654 | Hsd11b1 | hydroxysteroid 11-beta dehydrogenase 1 | 169.01 | 80.57 | 1.07 | 4.64E-05 |
| ENSMUSG00000009633 | GSMG0002657 | G0s2 | G0/G1 switch gene 2 | 520.16 | 259.04 | 1.01 | 7.95E-07 |
| ENSMUSG00000016179 | GSMG0002659 | Camk1g | calcium/calmodulin-dependent protein kinase I gamma | 643.50 | 668.67 | -0.05 | 8.86E-01 |
| ENSMUSG00000097325 | GSMG0002664 | Gm16897 | predicted gene, 16897 | 208.76 | 148.94 | 0.49 | 1.47E-01 |

| | | | | | | | |
|---|---|---|---|---|---|---|---|
| NULL | GSMG0002666 | NULL | NULL | 255.58 | 188.35 | 0.44 | 1.31E-01 |
| ENSMUSG00000016493 | GSMG0002667 | Cd46 | CD46 antigen, complement regulatory protein | 98.83 | 104.97 | -0.08 | 8.70E-01 |
| ENSMUSG00000016481 | GSMG0002669 | Cr1l | complement component (3b/4b) receptor 1-like | 2132.95 | 1884.91 | 0.18 | 4.54E-01 |
| ENSMUSG00000019774 | GSMG0002680 | Mtrf1l | mitochondrial translational release factor 1-like | 179.75 | 145.70 | 0.31 | 3.48E-01 |
| ENSMUSG00000019773 | GSMG0002681 | Fbxo5 | F-box protein 5 | 150.54 | 293.90 | -0.96 | 8.19E-05 |
| NULL | GSMG0002686 | NULL | NULL | 564.11 | 289.21 | 0.96 | 8.66E-08 |
| NULL | GSMG0002688 | NULL | NULL | 185.96 | 189.55 | -0.02 | 9.54E-01 |
| ENSMUSG00000075327 | GSMG0002689 | Zbtb2 | zinc finger and BTB domain containing 2 | 762.78 | 704.02 | 0.12 | 6.49E-01 |
| ENSMUSG00000019796 | GSMG0002696 | Lrp11 | low density lipoprotein receptor-related protein 11 | 932.36 | 1264.41 | -0.44 | 3.59E-02 |
| NULL | GSMG0002697 | A630066F11Rik | RIKEN cDNA A630066F11 gene | 114.17 | 133.66 | -0.23 | 5.71E-01 |
| ENSMUSG00000040034 | GSMG0002698 | Nup43 | nucleoporin 43 | 92.50 | 151.89 | -0.70 | 4.42E-02 |
| ENSMUSG00000040021 | GSMG0002699 | Lats1 | large tumor suppressor | 560.71 | 521.88 | 0.10 | 8.29E-01 |
| ENSMUSG00000019794 | GSMG0002701 | Katna1 | katanin p60 (ATPase-containing) subunit A1 | 718.37 | 691.86 | 0.06 | 8.65E-01 |
| ENSMUSG00000015757 | GSMG0002704 | Ppil4 | peptidylprolyl isomerase (cyclophilin)-like 4 | 417.25 | 405.72 | 0.04 | 9.02E-01 |
| ENSMUSG00000090112 | GSMG0002716 | Shprh | SNF2 histone linker PHD RING helicase | 271.41 | 298.98 | -0.14 | 6.91E-01 |
| ENSMUSG00000047648 | GSMG0002717 | Fbxo30 | F-box protein 30 | 895.02 | 1083.41 | -0.28 | 1.78E-01 |
| ENSMUSG00000055493 | GSMG0002719 | Epm2a | epilepsy, progressive myoclonic epilepsy, type 2 gene alpha | 130.29 | 119.54 | 0.12 | 7.97E-01 |
| ENSMUSG00000078348 | GSMG0002721 | Sf3b5 | splicing factor 3b, subunit 5 | 938.23 | 1076.36 | -0.20 | 4.65E-01 |
| ENSMUSG00000019810 | GSMG0002723 | Fuca2 | fucosidase, alpha-L- 2, plasma | 958.98 | 774.37 | 0.31 | 1.05E-01 |
| ENSMUSG00000015501 | GSMG0002728 | Hivep2 | human immunodeficiency virus type I enhancer binding protein 2 | 365.57 | 373.53 | -0.03 | 9.32E-01 |
| NULL | GSMG0002729 | NULL | NULL | 830.22 | 388.45 | 1.10 | 3.01E-11 |
| NULL | GSMG0002730 | NULL | NULL | 306.09 | 132.47 | 1.21 | 3.15E-08 |
| NULL | GSMG0002733 | NULL | NULL | 404.31 | 379.46 | 0.09 | 8.05E-01 |
| ENSMUSG00000039910 | GSMG0002739 | Cited2 | Cbp/p300-interacting transactivator, with Glu/Asp-rich carboxy-termina | 510.34 | 451.89 | 0.17 | 5.57E-01 |
| ENSMUSG00000019854 | GSMG0002741 | Reps1 | RalBP1 associated Eps domain containing protein | 637.39 | 729.20 | -0.20 | 5.37E-01 |
| ENSMUSG00000019851 | GSMG0002749 | Perp | PERP, TP53 apoptosis effector | 1023.35 | 376.88 | 1.44 | 1.31E-16 |
| ENSMUSG00000020009 | GSMG0002755 | Ifngr1 | interferon gamma receptor 1 | 1282.69 | 922.82 | 0.48 | 8.06E-03 |
| ENSMUSG00000071369 | GSMG0002759 | Map3k5 | mitogen-activated protein kinase kinase kinase 5 | 324.86 | 198.87 | 0.71 | 1.61E-02 |
| ENSMUSG00000019996 | GSMG0002761 | Map7 | microtubule-associated protein 7 | 1076.82 | 833.53 | 0.37 | 6.98E-02 |
| NULL | GSMG0002765 | NULL | NULL | 137.38 | 55.28 | 1.31 | 6.92E-06 |
| ENSMUSG00000037608 | GSMG0002766 | Bclaf1 | BCL2-associated transcription factor 1 | 3684.89 | 4041.29 | -0.13 | 5.62E-01 |
| ENSMUSG00000019992 | GSMG0002767 | Mtfr2 | mitochondrial fission regulator 2 | 49.77 | 110.17 | -1.16 | 3.36E-03 |
| NULL | GSMG0002768 | NULL | NULL | 267.88 | 155.14 | 0.79 | 1.81E-03 |

| | | | | | | | |
|---|---|---|---|---|---|---|---|
| ENSMUSG00000019986 | GSMG0002769 | Ahi1 | Abelson helper integration site 1 | 570.01 | 608.52 | -0.09 | 7.31E-01 |
| ENSMUSG00000019977 | GSMG0002772 | Hbs1l | Hbs1-like (S. cerevisiae) | 1747.02 | 1978.07 | -0.18 | 3.78E-01 |
| ENSMUSG00000019970 | GSMG0002778 | Sgk1 | serum/glucocorticoid regulated kinase 1 | 3110.61 | 1851.37 | 0.75 | 7.62E-06 |
| ENSMUSG00000037490 | GSMG0002785 | Slc2a12 | solute carrier family 2 (facilitated glucose transporter), member 12 | 149.66 | 212.91 | -0.50 | 1.35E-01 |
| ENSMUSG00000037455 | GSMG0002790 | Slc18b1 | solute carrier family 18, subfamily B, member 1 | 367.73 | 478.29 | -0.38 | 6.43E-02 |
| ENSMUSG00000019998 | GSMG0002803 | Stx7 | syntaxin 7 | 993.76 | 1190.48 | -0.26 | 2.61E-01 |
| ENSMUSG00000019997 | GSMG0002808 | Ctgf | connective tissue growth factor | 1473.42 | 478.25 | 1.62 | 8.12E-11 |
| ENSMUSG00000019984 | GSMG0002809 | Med23 | mediator complex subunit 23 | 1111.72 | 1367.20 | -0.30 | 1.43E-01 |
| ENSMUSG00000019978 | GSMG0002811 | Epb41l2 | erythrocyte membrane protein band 4.1 like 2 | 1262.91 | 1272.74 | -0.01 | 9.76E-01 |
| ENSMUSG00000039031 | GSMG0002820 | Arhgap18 | Rho GTPase activating protein 18 | 759.89 | 1304.31 | -0.78 | 6.03E-05 |
| ENSMUSG00000019889 | GSMG0002823 | Ptprk | protein tyrosine phosphatase, receptor type, K | 5003.33 | 9817.37 | -0.97 | 2.35E-11 |
| NULL | GSMG0002826 | NULL | NULL | 124.91 | 218.37 | -0.80 | 3.57E-03 |
| NULL | GSMG0002828 | NULL | NULL | 139.39 | 274.34 | -0.98 | 1.20E-03 |
| ENSMUSG00000038916 | GSMG0002844 | Soga3 | SOGA family member 3 | 3298.33 | 3047.89 | 0.11 | 6.09E-01 |
| NULL | GSMG0002845 | NULL | NULL | 750.23 | 768.34 | -0.03 | 9.13E-01 |
| ENSMUSG00000004360 | GSMG0002846 | 9330159F19Rik | RIKEN cDNA 9330159F19 gene | 518.22 | 448.42 | 0.21 | 4.03E-01 |
| ENSMUSG00000019883 | GSMG0002847 | Echdc1 | enoyl Coenzyme A hydratase domain containing 1 | 984.30 | 990.63 | -0.01 | 9.81E-01 |
| ENSMUSG00000000295 | GSMG0002851 | Hddc2 | HD domain containing 2 | 419.47 | 507.58 | -0.28 | 2.24E-01 |
| ENSMUSG00000047514 | GSMG0002863 | Tspyl1 | testis-specific protein, Y-encoded-like 1 | 2345.18 | 2318.03 | 0.02 | 9.53E-01 |
| ENSMUSG00000039485 | GSMG0002865 | Tspyl4 | TSPY-like 4 | 1737.91 | 1897.50 | -0.13 | 6.20E-01 |
| ENSMUSG00000019779 | GSMG0002867 | Frk | fyn-related kinase | 211.12 | 136.63 | 0.63 | 2.07E-02 |
| ENSMUSG00000044499 | GSMG0002869 | Hs3st5 | heparan sulfate (glucosamine) 3-O-sulfotransferase 5 | 159.93 | 333.99 | -1.06 | 4.49E-07 |
| ENSMUSG00000019777 | GSMG0002875 | Hdac2 | histone deacetylase 2 | 2232.61 | 2934.50 | -0.39 | 2.03E-02 |
| ENSMUSG00000019846 | GSMG0002884 | Lama4 | laminin, alpha 4 | 3400.60 | 5545.38 | -0.71 | 4.76E-05 |
| NULL | GSMG0002885 | NULL | NULL | 466.72 | 574.92 | -0.30 | 2.06E-01 |
| ENSMUSG00000019845 | GSMG0002887 | Tube1 | epsilon-tubulin 1 | 71.96 | 117.36 | -0.71 | 4.00E-02 |
| ENSMUSG00000019843 | GSMG0002889 | Fyn | Fyn proto-oncogene | 2857.64 | 4018.45 | -0.49 | 2.51E-03 |
| ENSMUSG00000019841 | GSMG0002894 | Rev3l | REV3 like, DNA directed polymerase zeta catalytic subunit | 1287.51 | 2045.62 | -0.67 | 5.16E-05 |
| ENSMUSG00000038481 | GSMG0002906 | Cdk19 | cyclin-dependent kinase 19 | 400.14 | 238.29 | 0.75 | 3.38E-04 |
| ENSMUSG00000019831 | GSMG0002912 | Wasf1 | WAS protein family, member 1 | 628.60 | 951.11 | -0.60 | 2.45E-03 |
| ENSMUSG00000019826 | GSMG0002914 | Zbtb24 | zinc finger and BTB domain containing 24 | 500.48 | 534.34 | -0.10 | 7.32E-01 |
| ENSMUSG00000019823 | GSMG0002915 | Mical1 | microtubule associated monooxygenase, calponin and LIM domain con | 1368.86 | 1805.63 | -0.40 | 7.17E-02 |
| ENSMUSG00000078451 | GSMG0002916 | Ppil6 | peptidylprolyl isomerase (cyclophilin)-like 6 | 104.48 | 88.56 | 0.23 | 6.06E-01 |

| | | | | | | | |
|---|---|---|---|---|---|---|---|
| ENSMUSG00000019818 | GSMG0002917 | Cd164 | CD164 antigen | 3453.82 | 3726.79 | -0.11 | 6.79E-01 |
| ENSMUSG00000038332 | GSMG0002920 | Sesn1 | sestrin 1 | 1963.16 | 1830.78 | 0.10 | 7.15E-01 |
| NULL | GSMG0002921 | NULL | NULL | 399.00 | 329.36 | 0.28 | 3.07E-01 |
| ENSMUSG00000019804 | GSMG0002927 | Snx3 | sorting nexin 3 | 1950.79 | 2176.63 | -0.16 | 5.95E-01 |
| ENSMUSG00000038280 | GSMG0002929 | Ostm1 | osteopetrosis associated transmembrane protein 1 | 566.31 | 448.31 | 0.34 | 1.07E-01 |
| ENSMUSG00000019802 | GSMG0002930 | Sec63 | SEC63-like (S. cerevisiae) | 2012.36 | 2310.03 | -0.20 | 3.35E-01 |
| ENSMUSG00000038240 | GSMG0002935 | Pdss2 | prenyl (solanesyl) diphosphate synthase, subunit 2 | 113.56 | 162.13 | -0.51 | 1.01E-01 |
| ENSMUSG00000038214 | GSMG0002940 | Bend3 | BEN domain containing 3 | 358.91 | 613.65 | -0.77 | 5.27E-03 |
| ENSMUSG00000047139 | GSMG0002942 | Cd24a | CD24a antigen | 797.55 | 3666.08 | -2.20 | 3.98E-57 |
| ENSMUSG00000019864 | GSMG0002948 | Rtn4ip1 | reticulon 4 interacting protein 1 | 185.49 | 161.76 | 0.20 | 5.67E-01 |
| ENSMUSG00000038160 | GSMG0002951 | Atg5 | autophagy related 5 | 1259.09 | 1336.01 | -0.09 | 7.62E-01 |
| ENSMUSG00000019849 | GSMG0002955 | Prep | prolyl endopeptidase | 1171.41 | 1697.05 | -0.53 | 3.82E-03 |
| ENSMUSG00000038822 | GSMG0002960 | Hace1 | HECT domain and ankyrin repeat containing, E3 ubiquitin protein ligas | 331.42 | 285.18 | 0.22 | 4.56E-01 |
| ENSMUSG00000019891 | GSMG0002982 | Dcbld1 | discoidin, CUB and LCCL domain containing 1 | 492.59 | 1228.88 | -1.32 | 4.29E-20 |
| NULL | GSMG0002984 | NULL | NULL | 87.74 | 91.66 | -0.07 | 9.00E-01 |
| ENSMUSG00000023068 | GSMG0002986 | Nus1 | NUS1 dehydrodolichyl diphosphate synthase subunit | 1351.88 | 1698.01 | -0.33 | 6.69E-02 |
| ENSMUSG00000038602 | GSMG0002988 | Slc35f1 | solute carrier family 35, member F1 | 277.59 | 733.60 | -1.40 | 1.32E-13 |
| ENSMUSG00000038583 | GSMG0002996 | Pln | phospholamban | 254.00 | 122.88 | 1.05 | 9.03E-06 |
| ENSMUSG00000019857 | GSMG0002999 | Asf1a | anti-silencing function 1A histone chaperone | 637.36 | 853.62 | -0.42 | 5.55E-02 |
| ENSMUSG00000047669 | GSMG0003005 | Msl3l2 | male-specific lethal 3-like 2 (Drosophila) | 231.28 | 309.44 | -0.42 | 8.98E-02 |
| ENSMUSG00000050953 | GSMG0003006 | Gja1 | gap junction protein, alpha 1 | 12859.38 | 11888.92 | 0.11 | 6.39E-01 |
| ENSMUSG00000019878 | GSMG0003011 | Hsf2 | heat shock factor 2 | 268.38 | 277.08 | -0.04 | 9.29E-01 |
| NULL | GSMG0003012 | NULL | NULL | 601.03 | 649.89 | -0.11 | 6.79E-01 |
| ENSMUSG00000019874 | GSMG0003014 | Fabp7 | fatty acid binding protein 7, brain | 740.25 | 8214.77 | -3.47 | 3.36E-130 |
| ENSMUSG00000019872 | GSMG0003015 | Smpdl3a | sphingomyelin phosphodiesterase, acid-like 3A | 2369.79 | 1164.65 | 1.03 | 1.77E-10 |
| ENSMUSG00000038039 | GSMG0003017 | Gcc2 | GRIP and coiled-coil domain containing 2 | 742.09 | 763.87 | -0.04 | 9.03E-01 |
| NULL | GSMG0003018 | NULL | NULL | 110.17 | 82.46 | 0.42 | 2.73E-01 |
| ENSMUSG00000019920 | GSMG0003019 | Lims1 | LIM and senescent cell antigen-like domains 1 | 3020.41 | 3726.01 | -0.30 | 1.33E-01 |
| ENSMUSG00000003226 | GSMG0003022 | Ranbp2 | RAN binding protein 2 | 1754.49 | 1811.01 | -0.05 | 8.57E-01 |
| ENSMUSG00000037990 | GSMG0003030 | Sh3rf3 | SH3 domain containing ring finger 3 | 230.79 | 357.16 | -0.63 | 2.17E-02 |
| NULL | GSMG0003031 | NULL | NULL | 515.37 | 769.94 | -0.58 | 6.83E-03 |
| ENSMUSG00000019916 | GSMG0003034 | P4ha1 | procollagen-proline, 2-oxoglutarate 4-dioxygenase (proline 4-hydroxyla | 824.87 | 564.96 | 0.55 | 9.76E-03 |
| ENSMUSG00000020111 | GSMG0003038 | Micu1 | mitochondrial calcium uptake 1 | 1054.88 | 1110.48 | -0.07 | 7.78E-01 |

| | | | | | | | |
|---|---|---|---|---|---|---|---|
| NULL | GSMG0003039 | NULL | NULL | 220.22 | 144.17 | 0.61 | 2.79E-02 |
| ENSMUSG00000020109 | GSMG0003040 | Dnajb12 | DnaJ heat shock protein family (Hsp40) member B12 | 335.75 | 314.80 | 0.09 | 8.12E-01 |
| ENSMUSG00000044475 | GSMG0003041 | Ascc1 | activating signal cointegrator 1 complex subunit 1 | 522.75 | 512.09 | 0.03 | 9.28E-01 |
| ENSMUSG00000058297 | GSMG0003043 | Spock2 | sparc/osteonectin, cwcv and kazal-like domains proteoglycan 2 | 7903.79 | 8368.79 | -0.08 | 8.20E-01 |
| ENSMUSG00000004207 | GSMG0003044 | Psap | prosaposin | 9901.69 | 8691.77 | 0.19 | 3.84E-01 |
| ENSMUSG00000020101 | GSMG0003045 | Vsir | V-set immunoregulatory receptor | 1217.68 | 896.11 | 0.44 | 2.87E-02 |
| NULL | GSMG0003054 | NULL | NULL | 125.04 | 99.33 | 0.33 | 3.72E-01 |
| NULL | GSMG0003055 | NULL | NULL | 158.56 | 105.82 | 0.58 | 6.06E-02 |
| ENSMUSG00000037151 | GSMG0003056 | Lrrc20 | leucine rich repeat containing 20 | 316.43 | 368.75 | -0.22 | 4.32E-01 |
| ENSMUSG00000020089 | GSMG0003061 | Ppa1 | pyrophosphatase (inorganic) 1 | 553.19 | 827.51 | -0.58 | 3.10E-03 |
| ENSMUSG00000020088 | GSMG0003062 | Sar1a | secretion associated Ras related GTPase 1A | 5718.02 | 5925.34 | -0.05 | 8.54E-01 |
| ENSMUSG00000020087 | GSMG0003063 | Tysnd1 | trypsin domain containing 1 | 714.85 | 898.12 | -0.33 | 8.43E-02 |
| ENSMUSG00000020085 | GSMG0003064 | Aifm2 | apoptosis-inducing factor, mitochondrion-associated 2 | 116.18 | 111.90 | 0.05 | 9.20E-01 |
| NULL | GSMG0003065 | NULL | NULL | 279.31 | 250.69 | 0.15 | 6.80E-01 |
| NULL | GSMG0003071 | NULL | NULL | 138.19 | 107.69 | 0.35 | 4.67E-01 |
| NULL | GSMG0003072 | NULL | NULL | 124.54 | 107.67 | 0.20 | 6.42E-01 |
| ENSMUSG00000071253 | GSMG0003074 | Slc25a16 | solute carrier family 25 (mitochondrial carrier, Graves disease autoantig | 283.84 | 274.91 | 0.05 | 9.13E-01 |
| ENSMUSG00000036875 | GSMG0003075 | Dna2 | DNA replication helicase/nuclease 2 | 149.95 | 296.99 | -0.98 | 3.01E-06 |
| ENSMUSG00000020070 | GSMG0003076 | Rufy2 | RUN and FYVE domain-containing 2 | 1814.48 | 1895.61 | -0.06 | 8.35E-01 |
| ENSMUSG00000020064 | GSMG0003080 | Herc4 | hect domain and RLD 4 | 488.66 | 469.47 | 0.06 | 8.79E-01 |
| ENSMUSG00000036764 | GSMG0003085 | Dnajc12 | DnaJ heat shock protein family (Hsp40) member C12 | 149.25 | 126.42 | 0.24 | 5.35E-01 |
| NULL | GSMG0003093 | NULL | NULL | 455.32 | 244.73 | 0.89 | 7.36E-06 |
| NULL | GSMG0003103 | NULL | NULL | 205.97 | 178.75 | 0.21 | 5.69E-01 |
| ENSMUSG00000037876 | GSMG0003107 | Jmjd1c | jumonji domain containing 1C | 4123.22 | 4861.27 | -0.24 | 2.38E-01 |
| ENSMUSG00000019944 | GSMG0003116 | Rhobtb1 | Rho-related BTB domain containing 1 | 1417.99 | 1500.07 | -0.08 | 7.37E-01 |
| NULL | GSMG0003119 | NULL | NULL | 285.32 | 331.74 | -0.22 | 4.54E-01 |
| NULL | GSMG0003120 | NULL | NULL | 460.47 | 484.43 | -0.07 | 8.25E-01 |
| NULL | GSMG0003121 | NULL | NULL | 215.29 | 318.28 | -0.56 | 1.43E-02 |
| NULL | GSMG0003123 | NULL | NULL | 308.38 | 401.73 | -0.38 | 1.04E-01 |
| NULL | GSMG0003124 | NULL | NULL | 591.75 | 763.82 | -0.37 | 1.09E-01 |
| NULL | GSMG0003125 | NULL | NULL | 450.50 | 341.82 | 0.40 | 2.15E-01 |
| NULL | GSMG0003127 | NULL | NULL | 534.00 | 367.83 | 0.54 | 1.76E-02 |
| NULL | GSMG0003128 | NULL | NULL | 139.47 | 124.18 | 0.17 | 7.25E-01 |

| | | | | | | | |
|---|---|---|---|---|---|---|---|
| NULL | GSMG0003129 | NULL | NULL | 129.83 | 123.99 | 0.07 | 8.94E-01 |
| ENSMUSG00000069601 | GSMG0003130 | Ank3 | ankyrin 3, epithelial | 846.53 | 1214.77 | -0.52 | 1.58E-03 |
| NULL | GSMG0003131 | NULL | NULL | 546.11 | 513.54 | 0.09 | 7.97E-01 |
| ENSMUSG00000037762 | GSMG0003136 | Slc16a9 | solute carrier family 16 (monocarboxylic acid transporters), member 9 | 1288.32 | 1155.36 | 0.16 | 5.10E-01 |
| ENSMUSG00000043259 | GSMG0003137 | Fam13c | family with sequence similarity 13, member C | 1802.55 | 1348.13 | 0.42 | 2.02E-02 |
| ENSMUSG00000019923 | GSMG0003144 | Zwint | ZW10 interactor | 1889.08 | 2323.15 | -0.30 | 1.28E-01 |
| NULL | GSMG0003146 | NULL | NULL | 514.02 | 586.89 | -0.19 | 5.06E-01 |
| ENSMUSG00000052613 | GSMG0003147 | Pcdh15 | protocadherin 15 | 456.02 | 477.32 | -0.06 | 8.85E-01 |
| ENSMUSG00000040009 | GSMG0003154 | Gnaz | guanine nucleotide binding protein, alpha z subunit | 293.38 | 481.26 | -0.71 | 7.23E-04 |
| ENSMUSG00000020175 | GSMG0003155 | Rab36 | RAB36, member RAS oncogene family | 371.48 | 301.64 | 0.30 | 2.91E-01 |
| ENSMUSG00000009681 | GSMG0003156 | Bcr | breakpoint cluster region | 715.65 | 794.54 | -0.15 | 5.71E-01 |
| NULL | GSMG0003157 | NULL | NULL | 100.27 | 102.33 | -0.03 | 9.50E-01 |
| ENSMUSG00000033444 | GSMG0003158 | Specc1l | sperm antigen with calponin homology and coiled-coil domains 1-like | 2826.56 | 2707.58 | 0.06 | 8.21E-01 |
| ENSMUSG00000020178 | GSMG0003159 | Adora2a | adenosine A2a receptor | 1540.90 | 2703.60 | -0.81 | 3.40E-08 |
| ENSMUSG00000020180 | GSMG0003163 | Snrpd3 | small nuclear ribonucleoprotein D3 | 1291.84 | 2037.37 | -0.66 | 2.06E-04 |
| ENSMUSG00000006345 | GSMG0003164 | Ggt1 | gamma-glutamyltransferase 1 | 277.39 | 168.07 | 0.72 | 3.76E-03 |
| ENSMUSG00000006344 | GSMG0003165 | Ggt5 | gamma-glutamyltransferase 5 | 669.35 | 445.60 | 0.59 | 2.89E-03 |
| NULL | GSMG0003169 | NULL | NULL | 135.82 | 128.35 | 0.08 | 8.69E-01 |
| ENSMUSG00000049422 | GSMG0003170 | Chchd10 | coiled-coil-helix-coiled-coil-helix domain containing 10 | 583.12 | 323.01 | 0.85 | 1.66E-04 |
| ENSMUSG00000049764 | GSMG0003173 | Zfp280b | zinc finger protein 280B | 1176.49 | 1235.39 | -0.07 | 7.99E-01 |
| ENSMUSG00000033208 | GSMG0003177 | S100b | S100 protein, beta polypeptide, neural | 568.62 | 161.64 | 1.81 | 9.53E-22 |
| ENSMUSG00000001150 | GSMG0003182 | Mcm3ap | minichromosome maintenance complex component 3 associated protein | 2281.98 | 2300.01 | -0.01 | 9.77E-01 |
| ENSMUSG00000033105 | GSMG0003183 | Lss | lanosterol synthase | 352.47 | 703.98 | -1.00 | 6.54E-09 |
| ENSMUSG00000001436 | GSMG0003189 | Slc19a1 | solute carrier family 19 (folate transporter), member 1 | 3508.51 | 2958.05 | 0.25 | 1.75E-01 |
| ENSMUSG00000020260 | GSMG0003191 | Pofut2 | protein O-fucosyltransferase 2 | 1373.75 | 1516.92 | -0.14 | 4.93E-01 |
| ENSMUSG00000009291 | GSMG0003195 | Pttg1ip | pituitary tumor-transforming 1 interacting protein | 13434.45 | 10782.59 | 0.32 | 6.17E-02 |
| ENSMUSG00000020265 | GSMG0003196 | Sumo3 | small ubiquitin-like modifier 3 | 3697.72 | 4605.46 | -0.32 | 6.57E-02 |
| ENSMUSG00000009293 | GSMG0003197 | Ube2g2 | ubiquitin-conjugating enzyme E2G 2 | 1585.73 | 1466.18 | 0.11 | 6.51E-01 |
| ENSMUSG00000020284 | GSMG0003202 | 1810043G02Rik | RIKEN cDNA 1810043G02 gene | 428.33 | 347.58 | 0.30 | 2.16E-01 |
| ENSMUSG00000000732 | GSMG0003205 | Icosl | icos ligand | 115.28 | 57.96 | 0.99 | 2.68E-03 |
| ENSMUSG00000005054 | GSMG0003209 | Cstb | cystatin B | 1643.48 | 1708.14 | -0.06 | 8.65E-01 |
| NULL | GSMG0003211 | NULL | NULL | 158.25 | 96.94 | 0.70 | 2.32E-02 |
| ENSMUSG00000032763 | GSMG0003212 | Ilvbl | ilvB (bacterial acetolactate synthase)-like | 1003.10 | 1132.68 | -0.17 | 4.41E-01 |

| | | | | | | | |
|---|---|---|---|---|---|---|---|
| ENSMUSG00000020310 | GSMG0003224 | Madcam1 | mucosal vascular addressin cell adhesion molecule 1 | 165.69 | 237.12 | -0.52 | 4.55E-02 |
| ENSMUSG00000020308 | GSMG0003225 | Tpgs1 | tubulin polyglutamylase complex subunit 1 | 1471.69 | 1459.34 | 0.01 | 9.70E-01 |
| ENSMUSG00000020307 | GSMG0003226 | Cdc34 | cell division cycle 34 | 832.88 | 1002.89 | -0.27 | 1.91E-01 |
| ENSMUSG00000023175 | GSMG0003228 | Bsg | basigin | 224441.93 | 148317.42 | 0.60 | 1.16E-04 |
| ENSMUSG00000020331 | GSMG0003229 | Hcn2 | hyperpolarization-activated, cyclic nucleotide-gated K+ 2 | 848.45 | 395.23 | 1.10 | 1.74E-04 |
| ENSMUSG00000035863 | GSMG0003233 | Palm | paralemmin | 5446.60 | 4928.42 | 0.14 | 5.37E-01 |
| ENSMUSG00000006498 | GSMG0003235 | Ptbp1 | polypyrimidine tract binding protein 1 | 2508.17 | 2594.55 | -0.05 | 8.68E-01 |
| ENSMUSG00000019564 | GSMG0003240 | Arid3a | AT rich interactive domain 3A (BRIGHT-like) | 208.16 | 249.85 | -0.27 | 4.11E-01 |
| ENSMUSG00000035754 | GSMG0003241 | Wdr18 | WD repeat domain 18 | 1043.43 | 1279.44 | -0.29 | 1.01E-01 |
| ENSMUSG00000004665 | GSMG0003243 | Cnn2 | calponin 2 | 5573.74 | 4621.44 | 0.27 | 1.40E-01 |
| ENSMUSG00000035722 | GSMG0003244 | Abca7 | ATP-binding cassette, sub-family A (ABC1), member 7 | 516.91 | 337.84 | 0.61 | 8.11E-03 |
| ENSMUSG00000035697 | GSMG0003245 | Hmha1 | histocompatibility (minor) HA-1 | 233.28 | 137.73 | 0.76 | 4.78E-03 |
| ENSMUSG00000003068 | GSMG0003247 | Stk11 | serine/threonine kinase 11 | 455.63 | 526.82 | -0.21 | 4.75E-01 |
| ENSMUSG00000003072 | GSMG0003250 | Atp5d | ATP synthase, H+ transporting, mitochondrial F1 complex, delta subun | 4810.51 | 5017.26 | -0.06 | 8.20E-01 |
| ENSMUSG00000045193 | GSMG0003252 | Cirbp | cold inducible RNA binding protein | 763.87 | 1213.72 | -0.67 | 1.27E-03 |
| ENSMUSG00000035595 | GSMG0003253 | 1600002K03Rik | RIKEN cDNA 1600002K03 gene | 96.96 | 94.37 | 0.03 | 9.55E-01 |
| ENSMUSG00000003070 | GSMG0003254 | Efna2 | ephrin A2 | 158.59 | 296.72 | -0.90 | 3.56E-04 |
| ENSMUSG00000020156 | GSMG0003255 | Mum1 | melanoma associated antigen (mutated) 1 | 1192.77 | 1365.33 | -0.19 | 3.57E-01 |
| ENSMUSG00000020153 | GSMG0003256 | Ndufs7 | NADH dehydrogenase (ubiquinone) Fe-S protein 7 | 2578.82 | 2607.41 | -0.02 | 9.56E-01 |
| ENSMUSG00000069565 | GSMG0003257 | Dazap1 | DAZ associated protein 1 | 572.95 | 846.24 | -0.57 | 1.62E-02 |
| ENSMUSG00000063457 | GSMG0003258 | Rps15 | ribosomal protein S15 | 1766.40 | 2477.49 | -0.49 | 1.42E-02 |
| ENSMUSG00000020135 | GSMG0003259 | Apc2 | adenomatosis polyposis coli 2 | 2024.43 | 4393.61 | -1.12 | 5.41E-07 |
| G00000113640,ENSMUSG000 | GSMG0003271 | Adat3 // Scamp4 | adenosine deaminase, tRNA-specific 3 // secretory carrier membrane pr | 640.34 | 737.80 | -0.21 | 3.87E-01 |
| ENSMUSG00000003345 | GSMG0003272 | Csnk1g2 | casein kinase 1, gamma 2 | 4315.32 | 4011.66 | 0.11 | 7.33E-01 |
| ENSMUSG00000055862 | GSMG0003274 | Izumo4 | IZUMO family member 4 | 886.80 | 735.70 | 0.27 | 2.13E-01 |
| ENSMUSG00000061589 | GSMG0003275 | Dot1l | DOT1-like, histone H3 methyltransferase (S. cerevisiae) | 335.95 | 453.08 | -0.43 | 2.58E-01 |
| ENSMUSG00000020211 | GSMG0003276 | Sf3a2 | splicing factor 3a, subunit 2 | 513.63 | 689.24 | -0.43 | 1.06E-01 |
| ENSMUSG00000035206 | GSMG0003280 | Sppl2b | signal peptide peptidase like 2B | 1066.82 | 901.52 | 0.24 | 3.60E-01 |
| ENSMUSG00000015312 | GSMG0003283 | Gadd45b | growth arrest and DNA-damage-inducible 45 beta | 378.20 | 184.35 | 1.04 | 1.60E-04 |
| ENSMUSG00000004929 | GSMG0003286 | Thop1 | thimet oligopeptidase 1 | 965.44 | 1197.01 | -0.31 | 1.17E-01 |
| ENSMUSG00000035027 | GSMG0003287 | Map2k2 | mitogen-activated protein kinase kinase 2 | 3628.85 | 3695.60 | -0.03 | 9.23E-01 |
| ENSMUSG00000035011 | GSMG0003289 | Zbtb7a | zinc finger and BTB domain containing 7a | 635.42 | 506.60 | 0.33 | 2.87E-01 |
| ENSMUSG00000034994 | GSMG0003291 | Eef2 | eukaryotic translation elongation factor 2 | 14152.99 | 15538.47 | -0.13 | 4.75E-01 |

| | | | | | | | |
|---|---|---|---|---|---|---|---|
| ENSMUSG00000034974 | GSMG0003292 | Dapk3 | death-associated protein kinase 3 | 1162.03 | 1087.90 | 0.09 | 7.20E-01 |
| ENSMUSG00000034949 | GSMG0003294 | Zfr2 | zinc finger RNA binding protein 2 | 907.66 | 855.17 | 0.09 | 8.10E-01 |
| ENSMUSG00000004933 | GSMG0003295 | Matk | megakaryocyte-associated tyrosine kinase | 9153.46 | 6429.52 | 0.51 | 2.64E-03 |
| ENSMUSG00000004931 | GSMG0003296 | Apba3 | amyloid beta (A4) precursor protein-binding, family A, member 3 | 1861.38 | 1385.77 | 0.43 | 1.10E-02 |
| ENSMUSG00000034902 | GSMG0003297 | Pip5k1c | phosphatidylinositol-4-phosphate 5-kinase, type 1 gamma | 3110.51 | 3222.86 | -0.05 | 8.83E-01 |
| ENSMUSG00000034889 | GSMG0003298 | Cactin | cactin, spliceosome C complex subunit | 612.87 | 676.26 | -0.14 | 5.70E-01 |
| ENSMUSG00000034881 | GSMG0003299 | Tbxa2r | thromboxane A2 receptor | 846.61 | 894.81 | -0.08 | 7.87E-01 |
| ENSMUSG00000034854 | GSMG0003300 | Mfsd12 | major facilitator superfamily domain containing 12 | 364.95 | 315.16 | 0.21 | 6.18E-01 |
| ENSMUSG00000078440 | GSMG0003302 | Dohh | deoxyhypusine hydroxylase/monooxygenase | 734.67 | 652.65 | 0.17 | 5.06E-01 |
| NULL | GSMG0003304 | NULL | NULL | 176.68 | 115.91 | 0.61 | 2.46E-02 |
| ENSMUSG00000054452 | GSMG0003305 | Aes | amino-terminal enhancer of split | 4344.40 | 4316.67 | 0.01 | 9.79E-01 |
| ENSMUSG00000034771 | GSMG0003306 | Tle2 | transducin-like enhancer of split 2 | 1279.75 | 1850.03 | -0.53 | 1.06E-02 |
| ENSMUSG00000054708 | GSMG0003308 | Ankrd24 | ankyrin repeat domain 24 | 701.41 | 836.54 | -0.25 | 2.85E-01 |
| NULL | GSMG0003310 | NULL | NULL | 172.80 | 206.75 | -0.26 | 4.25E-01 |
| NULL | GSMG0003313 | NULL | NULL | 215.15 | 273.25 | -0.35 | 2.29E-01 |
| ENSMUSG00000078435 | GSMG0003316 | AU041133 | expressed sequence AU041133 | 232.70 | 264.26 | -0.18 | 5.74E-01 |
| ENSMUSG00000020246 | GSMG0003324 | Hcfc2 | host cell factor C2 | 513.17 | 523.25 | -0.03 | 9.33E-01 |
| ENSMUSG00000020250 | GSMG0003326 | Txnrd1 | thioredoxin reductase 1 | 2279.19 | 3099.79 | -0.44 | 6.10E-03 |
| ENSMUSG00000020255 | GSMG0003332 | D10Wsu102e | DNA segment, Chr 10, Wayne State University 102, expressed | 257.00 | 388.56 | -0.59 | 7.07E-03 |
| ENSMUSG00000034560 | GSMG0003334 | A230046K03Rik | RIKEN cDNA A230046K03 gene | 990.11 | 1056.00 | -0.09 | 7.23E-01 |
| ENSMUSG00000087651 | GSMG0003335 | 1500009L16Rik | RIKEN cDNA 1500009L16 gene | 232.03 | 281.52 | -0.28 | 3.29E-01 |
| ENSMUSG00000020034 | GSMG0003340 | Tcp11l2 | t-complex 11 (mouse) like 2 | 1545.08 | 1062.30 | 0.54 | 5.11E-04 |
| ENSMUSG00000034453 | GSMG0003341 | Polr3b | polymerase (RNA) III (DNA directed) polypeptide B | 174.78 | 192.63 | -0.14 | 7.13E-01 |
| ENSMUSG00000020037 | GSMG0003342 | Rfx4 | regulatory factor X, 4 (influences HLA class II expression) | 152.40 | 115.93 | 0.40 | 3.67E-01 |
| ENSMUSG00000035620 | GSMG0003343 | Ric8b | RIC8 guanine nucleotide exchange factor B | 351.66 | 346.75 | 0.02 | 9.62E-01 |
| ENSMUSG00000060935 | GSMG0003345 | Tmem263 | transmembrane protein 263 | 253.52 | 287.87 | -0.18 | 5.33E-01 |
| ENSMUSG00000020042 | GSMG0003346 | Btbd11 | BTB (POZ) domain containing 11 | 104.35 | 139.22 | -0.41 | 3.30E-01 |
| ENSMUSG00000001785 | GSMG0003347 | Pwp1 | PWP1 homolog, endonuclein | 781.99 | 1100.82 | -0.49 | 9.98E-03 |
| ENSMUSG00000001786 | GSMG0003349 | Fbxo7 | F-box protein 7 | 328.84 | 237.88 | 0.47 | 3.65E-02 |
| ENSMUSG00000020044 | GSMG0003352 | Timp3 | tissue inhibitor of metalloproteinase 3 | 10284.94 | 3714.47 | 1.47 | 1.21E-28 |
| ENSMUSG00000035351 | GSMG0003369 | Nup37 | nucleoporin 37 | 267.57 | 356.32 | -0.41 | 1.49E-01 |
| ENSMUSG00000020056 | GSMG0003370 | Ccdc53 | coiled-coil domain containing 53 | 757.30 | 819.68 | -0.11 | 6.66E-01 |
| ENSMUSG00000035311 | GSMG0003371 | Gnptab | N-acetylglucosamine-1-phosphate transferase, alpha and beta subunits | 1142.36 | 1487.32 | -0.38 | 3.43E-02 |

| | | | | | | | |
|---|---|---|---|---|---|---|---|
| ENSMUSG00000019951 | GSMG0003379 | Uhrf1bp1l | UHRF1 (ICBP90) binding protein 1-like | 1468.33 | 1473.46 | 0.00 | 9.86E-01 |
| ENSMUSG00000058589 | GSMG0003389 | Anks1b | ankyrin repeat and sterile alpha motif domain containing 1B | 2053.37 | 2038.89 | 0.01 | 9.77E-01 |
| NULL | GSMG0003393 | NULL | NULL | 381.35 | 256.33 | 0.57 | 2.33E-02 |
| NULL | GSMG0003397 | NULL | NULL | 691.01 | 366.14 | 0.92 | 2.28E-07 |
| ENSMUSG00000019975 | GSMG0003399 | Ikbip | IKBKB interacting protein | 852.95 | 961.68 | -0.17 | 5.16E-01 |
| NULL | GSMG0003400 | NULL | NULL | 4220.46 | 4047.68 | 0.06 | 7.93E-01 |
| NULL | GSMG0003404 | NULL | NULL | 50.58 | 34.16 | 0.57 | 2.73E-01 |
| ENSMUSG00000020015 | GSMG0003405 | Cdk17 | cyclin-dependent kinase 17 | 4787.82 | 4219.70 | 0.18 | 3.69E-01 |
| ENSMUSG00000111531 | GSMG0003408 | Gm32688 | predicted gene, 32688 | 518.84 | 487.27 | 0.09 | 7.93E-01 |
| ENSMUSG00000015889 | GSMG0003410 | Lta4h | leukotriene A4 hydrolase | 3216.47 | 3842.26 | -0.26 | 1.61E-01 |
| ENSMUSG00000020019 | GSMG0003415 | Ntn4 | netrin 4 | 860.44 | 167.10 | 2.37 | 6.97E-34 |
| ENSMUSG00000020021 | GSMG0003422 | Fgd6 | FYVE, RhoGEF and PH domain containing 6 | 551.95 | 578.29 | -0.07 | 8.26E-01 |
| ENSMUSG00000005897 | GSMG0003424 | Nr2c1 | nuclear receptor subfamily 2, group C, member 1 | 331.06 | 325.46 | 0.03 | 9.43E-01 |
| ENSMUSG00000020024 | GSMG0003430 | Cep83 | centrosomal protein 83 | 498.21 | 525.75 | -0.08 | 8.44E-01 |
| ENSMUSG00000036499 | GSMG0003442 | Eea1 | early endosome antigen 1 | 290.38 | 311.66 | -0.11 | 7.58E-01 |
| ENSMUSG00000036478 | GSMG0003446 | Btg1 | B cell translocation gene 1, anti-proliferative | 1376.93 | 1264.67 | 0.12 | 6.57E-01 |
| ENSMUSG00000019929 | GSMG0003452 | Dcn | decorin | 2055.65 | 1361.90 | 0.60 | 5.35E-03 |
| ENSMUSG00000036446 | GSMG0003453 | Lum | lumican | 707.60 | 140.10 | 2.34 | 1.34E-18 |
| ENSMUSG00000019943 | GSMG0003459 | Atp2b1 | ATPase, Ca++ transporting, plasma membrane 1 | 9150.06 | 10000.26 | -0.13 | 5.20E-01 |
| ENSMUSG00000019952 | GSMG0003465 | Poc1b | POC1 centriolar protein B | 265.59 | 349.62 | -0.40 | 7.57E-02 |
| ENSMUSG00000090035 | GSMG0003466 | Galnt4 | UDP-N-acetyl-alpha-D-galactosamine:polypeptide N-acetylgalactosami | 946.86 | 1131.64 | -0.26 | 2.45E-01 |
| ENSMUSG00000019960 | GSMG0003469 | Dusp6 | dual specificity phosphatase 6 | 3489.50 | 3261.91 | 0.10 | 6.69E-01 |
| ENSMUSG00000019966 | GSMG0003474 | Kitl | kit ligand | 2317.20 | 1018.11 | 1.19 | 2.73E-14 |
| NULL | GSMG0003476 | NULL | NULL | 512.75 | 277.05 | 0.89 | 2.39E-05 |
| ENSMUSG00000019971 | GSMG0003478 | Cep290 | centrosomal protein 290 | 987.85 | 757.66 | 0.38 | 2.03E-01 |
| ENSMUSG00000044921 | GSMG0003491 | Rassf9 | Ras association (RalGDS/AF-6) domain family (N-terminal) member 9 | 2159.10 | 2621.30 | -0.28 | 1.28E-01 |
| ENSMUSG00000019894 | GSMG0003493 | Slc6a15 | solute carrier family 6 (neurotransmitter transporter), member 15 | 162.00 | 613.96 | -1.92 | 2.04E-17 |
| ENSMUSG00000019897 | GSMG0003496 | Ccdc59 | coiled-coil domain containing 59 | 985.39 | 1269.19 | -0.37 | 2.10E-01 |
| ENSMUSG00000053825 | GSMG0003497 | Ppfia2 | protein tyrosine phosphatase, receptor type, f polypeptide (PTPRF), inte | 1021.88 | 1192.18 | -0.22 | 2.79E-01 |
| ENSMUSG00000019906 | GSMG0003498 | Lin7a | lin-7 homolog A (C. elegans) | 130.82 | 81.71 | 0.68 | 6.83E-02 |
| ENSMUSG00000019907 | GSMG0003503 | Ppp1r12a | protein phosphatase 1, regulatory (inhibitor) subunit 12A | 1076.72 | 1070.04 | 0.01 | 9.82E-01 |
| ENSMUSG00000035873 | GSMG0003507 | Pawr | PRKC, apoptosis, WT1, regulator | 446.09 | 598.60 | -0.43 | 3.67E-02 |
| NULL | GSMG0003510 | NULL | NULL | 673.92 | 517.71 | 0.38 | 1.49E-01 |

| | | | | | | | |
|---|---|---|---|---|---|---|---|
| ENSMUSG00000020185 | GSMG0003516 | E2f7 | E2F transcription factor 7 | 201.17 | 369.39 | -0.87 | 1.92E-05 |
| ENSMUSG00000020186 | GSMG0003517 | Csrp2 | cysteine and glycine-rich protein 2 | 2396.50 | 2543.94 | -0.09 | 7.46E-01 |
| ENSMUSG00000020189 | GSMG0003518 | Osbpl8 | oxysterol binding protein-like 8 | 651.75 | 653.63 | -0.01 | 9.88E-01 |
| ENSMUSG00000020205 | GSMG0003529 | Phlda1 | pleckstrin homology like domain, family A, member 1 | 1153.69 | 895.00 | 0.36 | 1.39E-01 |
| ENSMUSG00000063334 | GSMG0003531 | Krr1 | KRR1, small subunit (SSU) processome component, homolog (yeast) | 428.30 | 509.76 | -0.25 | 3.56E-01 |
| ENSMUSG00000035681 | GSMG0003536 | Kcnc2 | potassium voltage gated channel, Shaw-related subfamily, member 2 | 324.33 | 293.73 | 0.15 | 7.15E-01 |
| NULL | GSMG0003537 | NULL | NULL | 273.03 | 231.55 | 0.24 | 4.42E-01 |
| NULL | GSMG0003542 | NULL | NULL | 851.29 | 1081.45 | -0.35 | 2.20E-01 |
| ENSMUSG00000034163 | GSMG0003548 | Zfc3h1 | zinc finger, C3H1-type containing | 211.18 | 193.83 | 0.12 | 7.32E-01 |
| ENSMUSG00000020151 | GSMG0003553 | Ptprr | protein tyrosine phosphatase, receptor type, R | 231.48 | 226.83 | 0.03 | 9.38E-01 |
| ENSMUSG00000020154 | GSMG0003557 | Ptprb | protein tyrosine phosphatase, receptor type, B | 5839.40 | 3522.31 | 0.73 | 9.45E-06 |
| ENSMUSG00000020164 | GSMG0003560 | Kcnmb4os1 | potassium large conductance calcium-activated channel, subfamily M, b | 84.36 | 75.57 | 0.16 | 7.71E-01 |
| NULL | GSMG0003562 | NULL | NULL | 307.20 | 304.62 | 0.01 | 9.88E-01 |
| ENSMUSG00000020183 | GSMG0003577 | Cpm | carboxypeptidase M | 315.71 | 121.82 | 1.37 | 6.42E-11 |
| ENSMUSG00000020212 | GSMG0003581 | Mdm1 | transformed mouse 3T3 cell double minute 1 | 250.54 | 400.14 | -0.68 | 2.38E-03 |
| ENSMUSG00000034813 | GSMG0003590 | Grip1 | glutamate receptor interacting protein 1 | 211.30 | 365.11 | -0.79 | 1.01E-03 |
| NULL | GSMG0003591 | NULL | NULL | 194.14 | 222.33 | -0.19 | 5.73E-01 |
| NULL | GSMG0003592 | NULL | NULL | 351.79 | 491.10 | -0.48 | 1.93E-02 |
| NULL | GSMG0003594 | NULL | NULL | 304.45 | 479.31 | -0.66 | 9.32E-04 |
| NULL | GSMG0003595 | NULL | NULL | 196.12 | 283.66 | -0.53 | 4.09E-02 |
| NULL | GSMG0003596 | NULL | NULL | 270.21 | 357.99 | -0.41 | 1.33E-01 |
| NULL | GSMG0003599 | NULL | NULL | 63.94 | 33.79 | 0.92 | 3.35E-02 |
| ENSMUSG00000020225 | GSMG0003600 | Tmbim4 | transmembrane BAX inhibitor motif containing 4 | 1869.12 | 1639.27 | 0.19 | 3.28E-01 |
| ENSMUSG00000034707 | GSMG0003609 | Gns | glucosamine (N-acetyl)-6-sulfatase | 529.11 | 547.42 | -0.05 | 8.87E-01 |
| ENSMUSG00000034613 | GSMG0003617 | Ppm1h | protein phosphatase 1H (PP2C domain containing) | 278.74 | 463.83 | -0.74 | 2.88E-03 |
| NULL | GSMG0003619 | NULL | NULL | 263.89 | 175.77 | 0.59 | 1.53E-02 |
| NULL | GSMG0003621 | NULL | NULL | 305.11 | 252.35 | 0.27 | 3.58E-01 |
| NULL | GSMG0003622 | NULL | NULL | 193.66 | 181.58 | 0.10 | 8.10E-01 |
| NULL | GSMG0003623 | NULL | NULL | 113.54 | 120.61 | -0.09 | 8.54E-01 |
| ENSMUSG00000044071 | GSMG0003626 | Fam19a2 | family with sequence similarity 19, member A2 | 303.35 | 662.19 | -1.13 | 3.94E-11 |
| ENSMUSG00000006732 | GSMG0003640 | Mettl1 | methyltransferase like 1 | 163.15 | 250.71 | -0.62 | 9.67E-03 |
| ENSMUSG00000006728 | GSMG0003642 | Cdk4 | cyclin-dependent kinase 4 | 2048.18 | 3329.72 | -0.70 | 1.22E-05 |
| ENSMUSG00000025422 | GSMG0003645 | Agap2 | ArfGAP with GTPase domain, ankyrin repeat and PH domain 2 | 403.22 | 239.61 | 0.75 | 3.00E-03 |

| | | | | | | | |
|---|---|---|---|---|---|---|---|
| NULL | GSMG0003646 | NULL | NULL | 513.63 | 574.58 | -0.16 | 5.38E-01 |
| ENSMUSG00000006731 | GSMG0003647 | B4galnt1 | beta-1,4-N-acetyl-galactosaminyl transferase 1 | 995.84 | 1359.36 | -0.45 | 3.38E-02 |
| ENSMUSG00000025410 | GSMG0003650 | Dctn2 | dynactin 2 | 3402.20 | 3436.37 | -0.01 | 9.58E-01 |
| ENSMUSG00000025408 | GSMG0003652 | Ddit3 | DNA-damage inducible transcript 3 | 74.41 | 60.51 | 0.30 | 5.66E-01 |
| ENSMUSG00000025404 | GSMG0003655 | R3hdm2 | R3H domain containing 2 | 1737.30 | 1277.95 | 0.44 | 4.09E-02 |
| NULL | GSMG0003657 | NULL | NULL | 447.46 | 343.23 | 0.39 | 1.34E-01 |
| ENSMUSG00000040280 | GSMG0003660 | Ndufa4l2 | NADH dehydrogenase (ubiquinone) 1 alpha subcomplex, 4-like 2 | 307.33 | 291.15 | 0.08 | 8.59E-01 |
| ENSMUSG00000002147 | GSMG0003662 | Stat6 | signal transducer and activator of transcription 6 | 2735.67 | 1887.66 | 0.54 | 2.57E-03 |
| ENSMUSG00000040195 | GSMG0003664 | 1700012D01Rik // | RIKEN cDNA 1700012D01 gene // transmembrane protein 194 | 76.62 | 81.35 | -0.09 | 8.73E-01 |
| ENSMUSG00000025400 | GSMG0003666 | Tac2 | tachykinin 2 | 81.11 | 35.99 | 1.17 | 2.99E-03 |
| ENSMUSG00000025395 | GSMG0003673 | Prim1 | DNA primase, p49 subunit | 169.77 | 417.43 | -1.30 | 9.51E-07 |
| ENSMUSG00000025393 | GSMG0003678 | Atp5b | ATP synthase, H+ transporting mitochondrial F1 complex, beta subunit | 21569.08 | 24339.07 | -0.17 | 4.01E-01 |
| NULL | GSMG0003679 | NULL | NULL | 215.07 | 169.67 | 0.34 | 2.85E-01 |
| ENSMUSG00000040054 | GSMG0003680 | Baz2a | bromodomain adjacent to zinc finger domain, 2A | 1098.85 | 1149.18 | -0.07 | 8.67E-01 |
| ENSMUSG00000044005 | GSMG0003682 | Gls2 | glutaminase 2 (liver, mitochondrial) | 233.69 | 88.55 | 1.40 | 4.05E-07 |
| ENSMUSG00000039994 | GSMG0003685 | Timeless | timeless circadian clock 1 | 830.76 | 994.05 | -0.26 | 2.18E-01 |
| ENSMUSG00000040033 | GSMG0003688 | Stat2 | signal transducer and activator of transcription 2 | 1139.74 | 1226.84 | -0.11 | 6.71E-01 |
| ENSMUSG00000005682 | GSMG0003689 | Pan2 | PAN2 poly(A) specific ribonuclease subunit | 990.63 | 1070.91 | -0.11 | 6.81E-01 |
| ENSMUSG00000025381 | GSMG0003690 | Cnpy2 | canopy FGF signaling regulator 2 | 1575.82 | 1733.36 | -0.14 | 6.09E-01 |
| ENSMUSG00000005683 | GSMG0003691 | Cs | citrate synthase | 2330.34 | 2665.77 | -0.19 | 3.72E-01 |
| ENSMUSG00000014498 | GSMG0003693 | Ankrd52 | ankyrin repeat domain 52 | 195.43 | 240.05 | -0.29 | 3.79E-01 |
| ENSMUSG00000025373 | GSMG0003695 | Rnf41 | ring finger protein 41 | 559.79 | 686.43 | -0.29 | 2.41E-01 |
| NULL | GSMG0003696 | NULL | NULL | 121.73 | 120.28 | 0.01 | 9.79E-01 |
| ENSMUSG00000025369 | GSMG0003697 | Smarcc2 | SWI/SNF related, matrix associated, actin dependent regulator of chron | 5617.63 | 7119.92 | -0.34 | 6.69E-02 |
| NULL | GSMG0003699 | NULL | NULL | 246.89 | 186.76 | 0.40 | 1.42E-01 |
| ENSMUSG00000097353 | GSMG0003700 | A430046D13Rik | Riken cDNA A430046D13 gene | 268.76 | 192.30 | 0.48 | 6.90E-02 |
| ENSMUSG00000064030 | GSMG0003706 | Pym1 | PYM homolog 1, exon junction complex associated factor | 214.78 | 327.55 | -0.61 | 2.22E-02 |
| ENSMUSG00000025354 | GSMG0003708 | Dnajc14 | DnaJ heat shock protein family (Hsp40) member C14 | 2043.96 | 1952.31 | 0.07 | 8.00E-01 |
| NULL | GSMG0003709 | NULL | NULL | 146.26 | 173.50 | -0.24 | 5.33E-01 |
| ENSMUSG00000025351 | GSMG0003712 | Cd63 | CD63 antigen | 1573.95 | 1550.21 | 0.02 | 9.46E-01 |
| ENSMUSG00000025348 | GSMG0003713 | Itga7 | integrin alpha 7 | 2375.33 | 798.05 | 1.57 | 8.40E-21 |
| ENSMUSG00000019772 | GSMG0003738 | Vip | vasoactive intestinal polypeptide | 454.53 | 200.33 | 1.18 | 2.21E-09 |
| ENSMUSG00000046916 | GSMG0003739 | Myct1 | myc target 1 | 782.42 | 1016.72 | -0.38 | 9.74E-02 |

| | | | | | | | |
|---|---|---|---|---|---|---|---|
| ENSMUSG00000061759 | GSMG0003742 | Armt1 | acidic residue methyltransferase 1 | 109.38 | 113.59 | -0.05 | 9.20E-01 |
| ENSMUSG00000038587 | GSMG0003746 | Akap12 | A kinase (PRKA) anchor protein (gravin) 12 | 7676.52 | 6601.92 | 0.22 | 2.73E-01 |
| NULL | GSMG0003747 | NULL | NULL | 365.92 | 203.04 | 0.85 | 6.68E-05 |
| ENSMUSG00000040675 | GSMG0003748 | Mthfd1l | methylenetetrahydrofolate dehydrogenase (NADP+ dependent) 1-like | 340.80 | 492.35 | -0.53 | 1.26E-02 |
| ENSMUSG00000040624 | GSMG0003750 | Plekhg1 | pleckstrin homology domain containing, family G (with RhoGef domai | 2685.15 | 3501.49 | -0.38 | 4.28E-02 |
| NULL | GSMG0003753 | NULL | NULL | 219.01 | 191.58 | 0.20 | 6.59E-01 |
| ENSMUSG00000040653 | GSMG0003757 | Ppp1r14c | protein phosphatase 1, regulatory (inhibitor) subunit 14c | 371.48 | 907.75 | -1.29 | 5.94E-12 |
| ENSMUSG00000019795 | GSMG0003760 | Pcmt1 | protein-L-isoaspartate (D-aspartate) O-methyltransferase 1 | 1046.04 | 1035.67 | 0.02 | 9.62E-01 |
| ENSMUSG00000040006 | GSMG0003764 | Ginm1 | glycoprotein integral membrane 1 | 2447.30 | 1949.78 | 0.33 | 1.03E-01 |
| NULL | GSMG0003765 | NULL | NULL | 231.28 | 235.27 | -0.03 | 9.51E-01 |
| ENSMUSG00000015755 | GSMG0003767 | Tab2 | TGF-beta activated kinase 1/MAP3K7 binding protein 2 | 691.96 | 705.15 | -0.03 | 9.37E-01 |
| ENSMUSG00000047712 | GSMG0003769 | Ust | uronyl-2-sulfotransferase | 92.88 | 110.44 | -0.24 | 6.13E-01 |
| ENSMUSG00000015305 | GSMG0003778 | Sash1 | SAM and SH3 domain containing 1 | 313.35 | 254.43 | 0.30 | 2.44E-01 |
| NULL | GSMG0003785 | NULL | NULL | 217.91 | 297.94 | -0.45 | 8.87E-02 |
| ENSMUSG00000060487 | GSMG0003795 | Samd5 | sterile alpha motif domain containing 5 | 697.84 | 782.41 | -0.16 | 5.64E-01 |
| ENSMUSG00000019790 | GSMG0003796 | Stxbp5 | syntaxin binding protein 5 (tomosyn) | 247.74 | 245.23 | 0.02 | 9.71E-01 |
| ENSMUSG00000019832 | GSMG0003800 | Rab32 | RAB32, member RAS oncogene family | 168.83 | 185.92 | -0.14 | 7.26E-01 |
| ENSMUSG00000019820 | GSMG0003807 | Utrn | utrophin | 3560.55 | 2092.60 | 0.77 | 5.00E-06 |
| ENSMUSG00000019814 | GSMG0003816 | Ltv1 | LTV1 ribosome biogenesis factor | 689.48 | 954.70 | -0.47 | 1.25E-02 |
| ENSMUSG00000062866 | GSMG0003818 | Phactr2 | phosphatase and actin regulator 2 | 1063.99 | 903.72 | 0.24 | 2.43E-01 |
| ENSMUSG00000019809 | GSMG0003822 | Pex3 | peroxisomal biogenesis factor 3 | 539.68 | 515.32 | 0.07 | 8.32E-01 |
| ENSMUSG00000019806 | GSMG0003823 | Aig1 | androgen-induced 1 | 473.80 | 508.98 | -0.10 | 7.54E-01 |
| NULL | GSMG0003825 | NULL | NULL | 290.29 | 283.64 | 0.03 | 9.46E-01 |
| NULL | GSMG0003830 | NULL | NULL | 275.40 | 233.16 | 0.24 | 4.08E-01 |
| ENSMUSG00000019868 | GSMG0003835 | Vta1 | vesicle (multivesicular body) trafficking 1 | 605.65 | 740.32 | -0.29 | 1.55E-01 |
| ENSMUSG00000039879 | GSMG0003843 | Heca | hdc homolog, cell cycle regulator | 271.94 | 109.24 | 1.31 | 8.31E-04 |
| ENSMUSG00000078453 | GSMG0003844 | Abracl | ABRA C-terminal like | 310.54 | 401.89 | -0.38 | 2.65E-01 |
| ENSMUSG00000059554 | GSMG0003847 | Ccdc28a | coiled-coil domain containing 28A | 195.12 | 143.63 | 0.44 | 1.50E-01 |
| NULL | GSMG0003850 | NULL | NULL | 334.81 | 313.38 | 0.10 | 8.09E-01 |
| ENSMUSG00000019852 | GSMG0003851 | Arfgef3 | ARFGEF family member 3 | 310.12 | 399.79 | -0.36 | 2.42E-01 |
| ENSMUSG00000019850 | GSMG0003855 | Tnfaip3 | tumor necrosis factor, alpha-induced protein 3 | 313.26 | 288.35 | 0.12 | 7.22E-01 |
| ENSMUSG00000020003 | GSMG0003863 | Pex7 | peroxisomal biogenesis factor 7 | 315.77 | 278.78 | 0.18 | 5.47E-01 |
| ENSMUSG00000019990 | GSMG0003870 | Pde7b | phosphodiesterase 7B | 141.08 | 116.96 | 0.28 | 5.04E-01 |

| | | | | | | | |
|---|---|---|---|---|---|---|---|
| NULL | GSMG0003874 | NULL | NULL | 261.50 | 128.94 | 1.02 | 2.15E-05 |
| ENSMUSG00000071359 | GSMG0003888 | Tbpl1 | TATA box binding protein-like 1 | 441.40 | 435.60 | 0.02 | 9.64E-01 |
| ENSMUSG00000037370 | GSMG0003905 | Enpp1 | ectonucleotide pyrophosphatase/phosphodiesterase 1 | 494.38 | 146.83 | 1.75 | 3.58E-19 |
| ENSMUSG00000019989 | GSMG0003906 | Enpp3 | ectonucleotide pyrophosphatase/phosphodiesterase 3 | 100.87 | 241.56 | -1.26 | 2.17E-05 |
| ENSMUSG00000039166 | GSMG0003909 | Akap7 | A kinase (PRKA) anchor protein 7 | 1178.52 | 1766.76 | -0.58 | 3.34E-04 |
| ENSMUSG00000039089 | GSMG0003919 | L3mbtl3 | l(3)mbt-like 3 (Drosophila) | 552.95 | 734.44 | -0.41 | 3.77E-02 |
| NULL | GSMG0003921 | NULL | NULL | 272.25 | 324.46 | -0.26 | 4.76E-01 |
| ENSMUSG00000019899 | GSMG0003923 | Lama2 | laminin, alpha 2 | 859.33 | 667.39 | 0.37 | 6.36E-02 |
| ENSMUSG00000038876 | GSMG0003933 | Rnf146 | ring finger protein 146 | 1431.81 | 1364.15 | 0.07 | 7.98E-01 |
| ENSMUSG00000019792 | GSMG0003941 | Trmt11 | tRNA methyltransferase 11 | 311.86 | 362.50 | -0.22 | 4.26E-01 |
| ENSMUSG00000111080 | GSMG0003942 | Gm20300 | predicted gene, 20300 | 436.80 | 367.28 | 0.25 | 3.62E-01 |
| ENSMUSG00000019791 | GSMG0003943 | Hint3 | histidine triad nucleotide binding protein 3 | 334.41 | 371.37 | -0.15 | 6.63E-01 |
| ENSMUSG00000039697 | GSMG0003945 | Ncoa7 | nuclear receptor coactivator 7 | 765.63 | 759.13 | 0.01 | 9.70E-01 |
| ENSMUSG00000019789 | GSMG0003947 | Hey2 | hairy/enhancer-of-split related with YRPW motif 2 | 381.00 | 234.80 | 0.70 | 4.74E-03 |
| ENSMUSG00000000296 | GSMG0003949 | Tpd52l1 | tumor protein D52-like 1 | 1741.40 | 1562.26 | 0.16 | 6.13E-01 |
| NULL | GSMG0003962 | NULL | NULL | 56.59 | 42.63 | 0.41 | 5.00E-01 |
| NULL | GSMG0003963 | NULL | NULL | 150.46 | 146.04 | 0.04 | 9.48E-01 |
| NULL | GSMG0003965 | NULL | NULL | 128.54 | 166.57 | -0.38 | 2.78E-01 |
| NULL | GSMG0003966 | NULL | NULL | 75.15 | 73.06 | 0.05 | 9.46E-01 |
| NULL | GSMG0003967 | NULL | NULL | 487.62 | 471.01 | 0.05 | 8.98E-01 |
| ENSMUSG00000019785 | GSMG0003972 | Clvs2 | clavesin 2 | 68.48 | 98.53 | -0.53 | 1.71E-01 |
| ENSMUSG00000039531 | GSMG0003975 | Zufsp | zinc finger with UFM1-specific peptidase domain | 492.11 | 422.97 | 0.22 | 3.46E-01 |
| ENSMUSG00000019782 | GSMG0003978 | Rwdd1 | RWD domain containing 1 | 755.90 | 1057.37 | -0.49 | 4.15E-02 |
| ENSMUSG00000039497 | GSMG0003982 | Dse | dermatan sulfate epimerase | 329.06 | 322.80 | 0.03 | 9.37E-01 |
| NULL | GSMG0003985 | NULL | NULL | 266.56 | 295.35 | -0.15 | 7.02E-01 |
| ENSMUSG00000039480 | GSMG0003987 | Nt5dc1 | 5'-nucleotidase domain containing 1 | 243.24 | 235.00 | 0.05 | 9.06E-01 |
| ENSMUSG00000069662 | GSMG0003990 | Marcks | myristoylated alanine rich protein kinase C substrate | 5087.35 | 6924.18 | -0.45 | 1.01E-01 |
| NULL | GSMG0003993 | NULL | NULL | 594.67 | 720.97 | -0.28 | 2.39E-01 |
| ENSMUSG00000051736 | GSMG0003994 | Fam229b | family with sequence similarity 229, member B | 182.94 | 164.63 | 0.15 | 6.85E-01 |
| ENSMUSG00000087177 | GSMG0003998 | E130307A14Rik | RIKEN cDNA E130307A14 gene | 573.03 | 777.99 | -0.44 | 1.24E-02 |
| ENSMUSG00000038510 | GSMG0004012 | Rpf2 | ribosome production factor 2 homolog | 214.68 | 368.87 | -0.78 | 4.59E-04 |
| ENSMUSG00000019837 | GSMG0004014 | Gtf3c6 | general transcription factor IIIC, polypeptide 6, alpha | 907.76 | 1183.01 | -0.38 | 3.15E-02 |
| ENSMUSG00000038446 | GSMG0004018 | Cdc40 | cell division cycle 40 | 410.40 | 386.64 | 0.09 | 7.95E-01 |

| | | | | | | | |
|---|---|---|---|---|---|---|---|
| ENSMUSG00000046922 | GSMG0004019 | Gpr6 | G protein-coupled receptor 6 | 187.48 | 103.96 | 0.85 | 1.19E-02 |
| ENSMUSG00000038417 | GSMG0004020 | Fig4 | FIG4 phosphoinositide 5-phosphatase | 786.70 | 867.15 | -0.14 | 6.13E-01 |
| ENSMUSG00000019822 | GSMG0004022 | Smpd2 | sphingomyelin phosphodiesterase 2, neutral | 1410.90 | 1185.05 | 0.25 | 2.68E-01 |
| ENSMUSG00000019813 | GSMG0004024 | Cep57l1 | centrosomal protein 57-like 1 | 105.99 | 135.10 | -0.35 | 3.38E-01 |
| ENSMUSG00000038302 | GSMG0004032 | Lace1 | lactation elevated 1 | 249.74 | 205.83 | 0.28 | 3.24E-01 |
| ENSMUSG00000038248 | GSMG0004040 | Sobp | sine oculis-binding protein homolog (Drosophila) | 411.78 | 494.44 | -0.27 | 5.03E-01 |
| NULL | GSMG0004041 | NULL | NULL | 116.75 | 153.52 | -0.40 | 3.54E-01 |
| NULL | GSMG0004042 | NULL | NULL | 282.60 | 447.96 | -0.66 | 1.21E-02 |
| NULL | GSMG0004043 | NULL | NULL | 638.15 | 848.34 | -0.41 | 7.42E-02 |
| NULL | GSMG0004044 | NULL | NULL | 217.35 | 306.77 | -0.50 | 1.12E-01 |
| ENSMUSG00000019797 | GSMG0004047 | 1700021F05Rik | RIKEN cDNA 1700021F05 gene | 331.96 | 404.08 | -0.29 | 2.56E-01 |
| ENSMUSG00000019863 | GSMG0004048 | Qrsl1 | glutaminyl-tRNA synthase (glutamine-hydrolyzing)-like 1 | 294.63 | 353.82 | -0.26 | 3.52E-01 |
| ENSMUSG00000038151 | GSMG0004052 | Prdm1 | PR domain containing 1, with ZNF domain | 507.20 | 716.90 | -0.50 | 1.26E-02 |
| NULL | GSMG0004056 | NULL | NULL | 134.91 | 146.80 | -0.12 | 7.69E-01 |
| ENSMUSG00000056073 | GSMG0004061 | Grik2 | glutamate receptor, ionotropic, kainate 2 (beta 2) | 505.39 | 766.39 | -0.60 | 7.06E-03 |
| NULL | GSMG0004065 | NULL | NULL | 186.91 | 276.93 | -0.56 | 6.11E-02 |
| NULL | GSMG0004066 | NULL | NULL | 297.88 | 383.09 | -0.37 | 2.94E-01 |
| ENSMUSG00000019861 | GSMG0004083 | Gopc | golgi associated PDZ and coiled-coil motif containing | 430.05 | 444.38 | -0.05 | 8.91E-01 |
| NULL | GSMG0004094 | NULL | NULL | 234.40 | 156.75 | 0.58 | 3.11E-02 |
| ENSMUSG00000003746 | GSMG0004102 | Man1a | mannosidase 1, alpha | 655.82 | 575.16 | 0.19 | 4.70E-01 |
| ENSMUSG00000019877 | GSMG0004113 | Serinc1 | serine incorporator 1 | 4781.98 | 5203.25 | -0.12 | 6.49E-01 |
| NULL | GSMG0004118 | NULL | NULL | 188.52 | 275.63 | -0.55 | 8.30E-02 |
| ENSMUSG00000019917 | GSMG0004120 | Sept10 | septin 10 | 836.92 | 1008.46 | -0.27 | 2.60E-01 |
| ENSMUSG00000009647 | GSMG0004123 | Mcu | mitochondrial calcium uniporter | 761.53 | 1112.85 | -0.55 | 6.27E-04 |
| NULL | GSMG0004127 | NULL | NULL | 217.94 | 228.56 | -0.07 | 8.83E-01 |
| ENSMUSG00000020108 | GSMG0004130 | Ddit4 | DNA-damage-inducible transcript 4 | 785.05 | 220.94 | 1.83 | 2.26E-11 |
| ENSMUSG00000020107 | GSMG0004131 | Anapc16 | anaphase promoting complex subunit 16 | 1452.77 | 1609.83 | -0.15 | 5.53E-01 |
| ENSMUSG00000020100 | GSMG0004136 | Slc29a3 | solute carrier family 29 (nucleoside transporters), member 3 | 98.18 | 72.30 | 0.44 | 3.29E-01 |
| ENSMUSG00000020099 | GSMG0004138 | Unc5b | unc-5 netrin receptor B | 1890.44 | 1746.59 | 0.11 | 6.11E-01 |
| ENSMUSG00000020097 | GSMG0004141 | Sgpl1 | sphingosine phosphate lyase 1 | 2238.92 | 2625.96 | -0.23 | 2.16E-01 |
| ENSMUSG00000020092 | GSMG0004143 | Pald1 | phosphatase domain containing, paladin 1 | 5010.05 | 7043.82 | -0.49 | 2.14E-03 |
| ENSMUSG00000020091 | GSMG0004145 | Eif4ebp2 | eukaryotic translation initiation factor 4E binding protein 2 | 829.54 | 719.46 | 0.20 | 3.72E-01 |
| ENSMUSG00000020086 | GSMG0004151 | H2afy2 | H2A histone family, member Y2 | 314.93 | 740.86 | -1.24 | 2.38E-13 |

| | | | | | | | |
|---|---|---|---|---|---|---|---|
| ENSMUSG00000058806 | GSMG0004152 | Col13a1 | collagen, type XIII, alpha 1 | 196.01 | 106.02 | 0.89 | 8.49E-04 |
| ENSMUSG00000020083 | GSMG0004155 | 2010107G23Rik | RIKEN cDNA 2010107G23 gene | 339.21 | 552.22 | -0.71 | 2.68E-04 |
| ENSMUSG00000037031 | GSMG0004156 | Tspan15 | tetraspanin 15 | 764.77 | 1283.85 | -0.75 | 1.30E-05 |
| ENSMUSG00000037012 | GSMG0004158 | Hk1 | hexokinase 1 | 4911.80 | 5171.76 | -0.07 | 7.57E-01 |
| ENSMUSG00000020079 | GSMG0004160 | Supv3l1 | suppressor of var1, 3-like 1 (S. cerevisiae) | 450.12 | 537.72 | -0.25 | 3.17E-01 |
| ENSMUSG00000020078 | GSMG0004161 | Vps26a | VPS26 retromer complex component A | 687.39 | 645.55 | 0.09 | 7.92E-01 |
| ENSMUSG00000020077 | GSMG0004162 | Srgn | serglycin | 3740.81 | 3047.81 | 0.30 | 1.98E-01 |
| ENSMUSG00000036955 | GSMG0004164 | Kif1bp | KIF1 binding protein | 561.80 | 502.53 | 0.16 | 5.18E-01 |
| ENSMUSG00000020075 | GSMG0004165 | Ddx21 | DEAD (Asp-Glu-Ala-Asp) box polypeptide 21 | 532.27 | 787.96 | -0.56 | 2.63E-03 |
| ENSMUSG00000020076 | GSMG0004166 | Ddx50 | DEAD (Asp-Glu-Ala-Asp) box polypeptide 50 | 2403.38 | 2470.54 | -0.04 | 8.92E-01 |
| ENSMUSG00000020074 | GSMG0004168 | Ccar1 | cell division cycle and apoptosis regulator 1 | 3037.28 | 3509.56 | -0.21 | 3.12E-01 |
| ENSMUSG00000020069 | GSMG0004174 | Hnrnph3 | heterogeneous nuclear ribonucleoprotein H3 | 279.83 | 306.16 | -0.13 | 6.77E-01 |
| ENSMUSG00000020063 | GSMG0004178 | Sirt1 | sirtuin 1 | 272.21 | 174.97 | 0.63 | 2.64E-02 |
| ENSMUSG00000042846 | GSMG0004184 | Lrrtm3 | leucine rich repeat transmembrane neuronal 3 | 1728.94 | 1011.00 | 0.77 | 3.88E-07 |
| NULL | GSMG0004187 | NULL | NULL | 180.98 | 87.13 | 1.05 | 3.99E-05 |
| NULL | GSMG0004188 | NULL | NULL | 129.10 | 72.30 | 0.84 | 8.13E-03 |
| NULL | GSMG0004189 | NULL | NULL | 273.53 | 160.97 | 0.76 | 2.15E-03 |
| ENSMUSG00000019873 | GSMG0004195 | Reep3 | receptor accessory protein 3 | 2016.12 | 2096.82 | -0.06 | 8.38E-01 |
| ENSMUSG00000057134 | GSMG0004201 | Ado | 2-aminoethanethiol (cysteamine) dioxygenase | 1191.10 | 1540.86 | -0.37 | 8.00E-02 |
| ENSMUSG00000037855 | GSMG0004204 | Zfp365 | zinc finger protein 365 | 419.28 | 175.37 | 1.26 | 3.40E-09 |
| ENSMUSG00000019947 | GSMG0004206 | Arid5b | AT rich interactive domain 5B (MRF1-like) | 354.09 | 436.90 | -0.30 | 2.07E-01 |
| ENSMUSG00000090622 | GSMG0004219 | A930033H14Rik | RIKEN cDNA A930033H14 gene | 563.67 | 586.96 | -0.06 | 8.57E-01 |
| ENSMUSG00000019942 | GSMG0004222 | Cdk1 | cyclin-dependent kinase 1 | 768.50 | 1805.21 | -1.23 | 5.32E-12 |
| ENSMUSG00000037747 | GSMG0004226 | Phyhipl | phytanoyl-CoA hydroxylase interacting protein-like | 840.73 | 1143.82 | -0.44 | 1.57E-02 |
| ENSMUSG00000014329 | GSMG0004231 | Bicc1 | BicC family RNA binding protein 1 | 608.92 | 439.61 | 0.47 | 2.16E-02 |
| ENSMUSG00000003923 | GSMG0004235 | Tfam | transcription factor A, mitochondrial | 742.13 | 954.32 | -0.36 | 5.51E-02 |
| ENSMUSG00000019927 | GSMG0004236 | Ube2d1 | ubiquitin-conjugating enzyme E2D 1 | 816.76 | 1363.41 | -0.74 | 6.55E-06 |
| ENSMUSG00000037710 | GSMG0004237 | Cisd1 | CDGSH iron sulfur domain 1 | 1691.43 | 2001.87 | -0.24 | 2.85E-01 |
| ENSMUSG00000033416 | GSMG0004248 | Gucd1 | guanylyl cyclase domain containing 1 | 200.91 | 180.78 | 0.16 | 6.91E-01 |
| ENSMUSG00000046807 | GSMG0004251 | Lrrc75b | leucine rich repeat containing 75B | 17.18 | 53.28 | -1.61 | 1.12E-03 |
| ENSMUSG00000020196 | GSMG0004254 | Cabin1 | calcineurin binding protein 1 | 1682.14 | 1489.22 | 0.18 | 4.78E-01 |
| ENSMUSG00000001666 | GSMG0004255 | Ddt | D-dopachrome tautomerase | 350.84 | 369.89 | -0.08 | 8.50E-01 |
| ENSMUSG00000001663 | GSMG0004257 | Gstt1 | glutathione S-transferase, theta 1 | 102.11 | 74.81 | 0.45 | 2.33E-01 |

| | | | | | | | |
|---|---|---|---|---|---|---|---|
| ENSMUSG00000033318 | GSMG0004259 | Gstt2 | glutathione S-transferase, theta 2 | 1084.44 | 1067.33 | 0.02 | 9.42E-01 |
| ENSMUSG00000000902 | GSMG0004262 | Smarcb1 | SWI/SNF related, matrix associated, actin dependent regulator of chron | 1215.61 | 1673.40 | -0.46 | 8.70E-03 |
| ENSMUSG00000000901 | GSMG0004263 | Mmp11 | matrix metallopeptidase 11 | 201.37 | 274.55 | -0.45 | 8.73E-02 |
| ENSMUSG00000020230 | GSMG0004266 | Prmt2 | protein arginine N-methyltransferase 2 | 1442.31 | 2555.93 | -0.83 | 6.05E-09 |
| ENSMUSG00000020231 | GSMG0004267 | Dip2a | disco interacting protein 2 homolog A | 865.94 | 863.69 | 0.00 | 9.95E-01 |
| ENSMUSG00000001151 | GSMG0004268 | Pcnt | pericentrin (kendrin) | 651.01 | 627.17 | 0.05 | 8.90E-01 |
| ENSMUSG00000020241 | GSMG0004272 | Col6a2 | collagen, type VI, alpha 2 | 1091.10 | 713.33 | 0.61 | 1.47E-04 |
| ENSMUSG00000001119 | GSMG0004273 | Col6a1 | collagen, type VI, alpha 1 | 5794.40 | 1353.10 | 2.10 | 2.88E-30 |
| ENSMUSG00000001120 | GSMG0004275 | Pcbp3 | poly(rC) binding protein 3 | 426.83 | 513.66 | -0.27 | 2.85E-01 |
| ENSMUSG00000001435 | GSMG0004282 | Col18a1 | collagen, type XVIII, alpha 1 | 6563.59 | 8546.73 | -0.38 | 1.08E-01 |
| ENSMUSG00000078444 | GSMG0004283 | Gm10941 | predicted gene 10941 | 27.48 | 72.60 | -1.42 | 6.26E-04 |
| ENSMUSG00000020262 | GSMG0004284 | Adarb1 | adenosine deaminase, RNA-specific, B1 | 237.59 | 292.68 | -0.30 | 3.11E-01 |
| ENSMUSG00000032977 | GSMG0004285 | Fam207a | family with sequence similarity 207, member A | 488.78 | 545.36 | -0.16 | 5.36E-01 |
| ENSMUSG00000009292 | GSMG0004290 | Trpm2 | transient receptor potential cation channel, subfamily M, member 2 | 246.66 | 129.78 | 0.93 | 2.44E-03 |
| ENSMUSG00000020277 | GSMG0004291 | Pfkl | phosphofructokinase, liver, B-type | 4726.49 | 4404.91 | 0.10 | 6.63E-01 |
| ENSMUSG00000053329 | GSMG0004293 | D10Jhu81e | DNA segment, Chr 10, Johns Hopkins University 81 expressed | 317.84 | 356.26 | -0.17 | 5.74E-01 |
| ENSMUSG00000032834 | GSMG0004294 | Pwp2 | PWP2 periodic tryptophan protein homolog (yeast) | 173.86 | 209.04 | -0.26 | 4.42E-01 |
| ENSMUSG00000000374 | GSMG0004296 | Trappc10 | trafficking protein particle complex 10 | 484.12 | 496.88 | -0.04 | 9.33E-01 |
| ENSMUSG00000001211 | GSMG0004298 | Agpat3 | 1-acylglycerol-3-phosphate O-acyltransferase 3 | 2385.63 | 1961.53 | 0.28 | 2.68E-01 |
| ENSMUSG00000061032 | GSMG0004299 | Rrp1 | ribosomal RNA processing 1 homolog (S. cerevisiae) | 6295.77 | 7001.39 | -0.15 | 4.05E-01 |
| ENSMUSG00000032788 | GSMG0004300 | Pdxk | pyridoxal (pyridoxine, vitamin B6) kinase | 981.54 | 544.21 | 0.85 | 1.94E-05 |
| ENSMUSG00000032714 | GSMG0004303 | Syde1 | synapse defective 1, Rho GTPase, homolog 1 (C. elegans) | 1825.63 | 1698.99 | 0.10 | 6.47E-01 |
| ENSMUSG00000060301 | GSMG0004308 | 2610008E11Rik | RIKEN cDNA 2610008E11 gene | 1778.45 | 1449.33 | 0.30 | 1.01E-01 |
| ENSMUSG00000042570 | GSMG0004310 | Mier2 | MIER family member 2 | 1006.48 | 990.70 | 0.02 | 9.50E-01 |
| ENSMUSG00000020312 | GSMG0004313 | Shc2 | SHC (Src homology 2 domain containing) transforming protein 2 | 1274.81 | 1143.86 | 0.16 | 5.88E-01 |
| ENSMUSG00000020329 | GSMG0004316 | Polrmt | polymerase (RNA) mitochondrial (DNA directed) | 962.24 | 1000.92 | -0.06 | 8.82E-01 |
| ENSMUSG00000035890 | GSMG0004317 | Rnf126 | ring finger protein 126 | 629.63 | 749.90 | -0.25 | 4.09E-01 |
| NULL | GSMG0004319 | NULL | NULL | 187.92 | 135.27 | 0.47 | 1.43E-01 |
| ENSMUSG00000035835 | GSMG0004321 | Plppr3 | phospholipid phosphatase related 3 | 1234.72 | 4920.44 | -2.00 | 1.34E-33 |
| ENSMUSG00000013833 | GSMG0004322 | Med16 | mediator complex subunit 16 | 1670.20 | 1856.86 | -0.15 | 5.00E-01 |
| ENSMUSG00000035781 | GSMG0004323 | R3hdm4 | R3H domain containing 4 | 1645.59 | 1546.56 | 0.09 | 7.12E-01 |
| ENSMUSG00000013858 | GSMG0004324 | Tmem259 | transmembrane protein 259 | 975.78 | 750.22 | 0.38 | 6.60E-02 |
| ENSMUSG00000004667 | GSMG0004325 | Polr2e | polymerase (RNA) II (DNA directed) polypeptide E | 1858.93 | 1901.62 | -0.03 | 9.09E-01 |

| | | | | | | | |
|---|---|---|---|---|---|---|---|
| ENSMUSG00000035673 | GSMG0004328 | Sbno2 | strawberry notch homolog 2 (Drosophila) | 1230.23 | 1204.62 | 0.03 | 9.41E-01 |
| ENSMUSG00000020150 | GSMG0004330 | Gamt | guanidinoacetate methyltransferase | 475.21 | 334.98 | 0.50 | 1.39E-02 |
| ENSMUSG00000020133 | GSMG0004331 | 2310011J03Rik | RIKEN cDNA 2310011J03 gene | 990.33 | 836.19 | 0.24 | 2.20E-01 |
| ENSMUSG00000043822 | GSMG0004333 | Adamtsl5 | ADAMTS-like 5 | 84.44 | 34.46 | 1.30 | 1.35E-03 |
| ENSMUSG00000048696 | GSMG0004334 | Mex3d | mex3 RNA binding family member D | 293.10 | 284.75 | 0.04 | 9.54E-01 |
| ENSMUSG00000035478 | GSMG0004335 | Mbd3 | methyl-CpG binding domain protein 3 | 3351.04 | 4067.85 | -0.28 | 9.19E-02 |
| ENSMUSG00000020163 | GSMG0004336 | Uqcr11 | ubiquinol-cytochrome c reductase, complex III subunit XI | 2445.84 | 2002.64 | 0.29 | 9.09E-02 |
| ENSMUSG00000020167 | GSMG0004337 | Tcf3 | transcription factor 3 | 2867.17 | 3600.81 | -0.33 | 5.84E-02 |
| ENSMUSG00000047417 | GSMG0004339 | Rexo1 | REX1, RNA exonuclease 1 | 645.07 | 683.42 | -0.08 | 8.22E-01 |
| ENSMUSG00000003346 | GSMG0004341 | Abhd17a | abhydrolase domain containing 17A | 4265.33 | 5572.03 | -0.39 | 8.90E-03 |
| ENSMUSG00000003344 | GSMG0004342 | Btbd2 | BTB (POZ) domain containing 2 | 499.87 | 491.47 | 0.02 | 9.62E-01 |
| ENSMUSG00000020190 | GSMG0004343 | Mknk2 | MAP kinase-interacting serine/threonine kinase 2 | 2019.68 | 2211.23 | -0.13 | 7.52E-01 |
| ENSMUSG00000020198 | GSMG0004345 | Ap3d1 | adaptor-related protein complex 3, delta 1 subunit | 5352.68 | 5785.09 | -0.11 | 6.64E-01 |
| ENSMUSG00000035278 | GSMG0004346 | Plekhj1 | pleckstrin homology domain containing, family J member 1 | 996.90 | 1058.61 | -0.09 | 7.79E-01 |
| ENSMUSG00000051067 | GSMG0004348 | Lingo3 | leucine rich repeat and Ig domain containing 3 | 369.16 | 400.62 | -0.12 | 7.74E-01 |
| ENSMUSG00000020219 | GSMG0004350 | Timm13 | translocase of inner mitochondrial membrane 13 | 2066.54 | 2447.83 | -0.24 | 2.60E-01 |
| ENSMUSG00000062075 | GSMG0004351 | Lmnb2 | lamin B2 | 559.46 | 792.42 | -0.50 | 1.25E-02 |
| ENSMUSG00000048240 | GSMG0004352 | Gng7 | guanine nucleotide binding protein (G protein), gamma 7 | 2193.40 | 1593.43 | 0.46 | 1.82E-02 |
| ENSMUSG00000043670 | GSMG0004353 | Diras1 | DIRAS family, GTP-binding RAS-like 1 | 346.71 | 498.06 | -0.52 | 6.10E-02 |
| ENSMUSG00000046822 | GSMG0004354 | Slc39a3 | solute carrier family 39 (zinc transporter), member 3 | 2170.19 | 1780.57 | 0.29 | 2.15E-01 |
| ENSMUSG00000004937 | GSMG0004355 | Sgta | small glutamine-rich tetratricopeptide repeat (TPR)-containing, alpha | 2024.86 | 2251.42 | -0.15 | 5.13E-01 |
| ENSMUSG00000004934 | GSMG0004357 | Pias4 | protein inhibitor of activated STAT 4 | 531.22 | 515.39 | 0.04 | 8.95E-01 |
| ENSMUSG00000034958 | GSMG0004360 | Atcay | ataxia, cerebellar, Cayman type | 429.05 | 601.77 | -0.49 | 2.82E-02 |
| NULL | GSMG0004361 | NULL | NULL | 3211.15 | 2349.65 | 0.45 | 1.35E-02 |
| ENSMUSG00000034932 | GSMG0004362 | Mrpl54 | mitochondrial ribosomal protein L54 | 767.37 | 978.15 | -0.35 | 8.37E-02 |
| NULL | GSMG0004364 | NULL | NULL | 328.06 | 299.13 | 0.13 | 7.25E-01 |
| ENSMUSG00000020232 | GSMG0004366 | Hmg20b | high mobility group 20B | 855.95 | 928.66 | -0.12 | 7.18E-01 |
| ENSMUSG00000020235 | GSMG0004368 | Fzr1 | fizzy/cell division cycle 20 related 1 (Drosophila) | 1681.56 | 2190.23 | -0.38 | 4.33E-02 |
| ENSMUSG00000055053 | GSMG0004369 | Nfic | nuclear factor I/C | 744.15 | 601.41 | 0.31 | 1.23E-01 |
| ENSMUSG00000034818 | GSMG0004370 | Celf5 | CUGBP, Elav-like family member 5 | 244.25 | 305.11 | -0.32 | 3.70E-01 |
| ENSMUSG00000020238 | GSMG0004371 | Ncln | nicalin | 1909.52 | 2295.07 | -0.26 | 2.03E-01 |
| ENSMUSG00000034781 | GSMG0004374 | Gna11 | guanine nucleotide binding protein, alpha 11 | 3759.32 | 3683.96 | 0.03 | 9.15E-01 |
| ENSMUSG00000034748 | GSMG0004376 | Sirt6 | sirtuin 6 | 214.70 | 216.50 | -0.01 | 9.77E-01 |

| | | | | | | | |
|---|---|---|---|---|---|---|---|
| ENSMUSG00000062931 | GSMG0004382 | Zfp938 | zinc finger protein 938 | 440.29 | 398.27 | 0.14 | 5.95E-01 |
| ENSMUSG00000063320 | GSMG0004387 | 1190007I07Rik | RIKEN cDNA 1190007I07 gene | 100.35 | 100.83 | -0.01 | 9.85E-01 |
| ENSMUSG00000020248 | GSMG0004389 | Nfyb | nuclear transcription factor-Y beta | 449.96 | 492.08 | -0.13 | 6.34E-01 |
| ENSMUSG00000034591 | GSMG0004394 | Slc41a2 | solute carrier family 41, member 2 | 82.87 | 104.13 | -0.34 | 4.97E-01 |
| ENSMUSG00000020256 | GSMG0004395 | Aldh1l2 | aldehyde dehydrogenase 1 family, member L2 | 114.68 | 564.82 | -2.30 | 9.18E-23 |
| ENSMUSG00000020263 | GSMG0004397 | Appl2 | adaptor protein, phosphotyrosine interaction, PH domain and leucine zi | 1522.31 | 1252.85 | 0.28 | 2.14E-01 |
| ENSMUSG00000046841 | GSMG0004402 | Ckap4 | cytoskeleton-associated protein 4 | 1875.90 | 2518.79 | -0.43 | 7.70E-03 |
| ENSMUSG00000049038 | GSMG0004407 | Mterf2 | mitochondrial transcription termination factor 2 | 299.28 | 188.01 | 0.67 | 7.73E-03 |
| ENSMUSG00000020038 | GSMG0004408 | Cry1 | cryptochrome 1 (photolyase-like) | 907.77 | 949.92 | -0.06 | 8.17E-01 |
| ENSMUSG00000035529 | GSMG0004412 | Prdm4 | PR domain containing 4 | 592.14 | 814.19 | -0.46 | 5.57E-02 |
| ENSMUSG00000001783 | GSMG0004413 | Rtcb | RNA 2',3'-cyclic phosphate and 5'-OH ligase | 2024.29 | 2398.25 | -0.24 | 2.58E-01 |
| ENSMUSG00000059602 | GSMG0004418 | Syn3 | synapsin III | 119.83 | 399.31 | -1.73 | 7.87E-18 |
| ENSMUSG00000020048 | GSMG0004421 | Hsp90b1 | heat shock protein 90, beta (Grp94), member 1 | 14784.52 | 16869.18 | -0.19 | 3.94E-01 |
| ENSMUSG00000060002 | GSMG0004433 | Chpt1 | choline phosphotransferase 1 | 480.76 | 302.45 | 0.67 | 4.98E-04 |
| ENSMUSG00000035189 | GSMG0004441 | Ano4 | anoctamin 4 | 108.37 | 96.74 | 0.16 | 7.46E-01 |
| ENSMUSG00000074802 | GSMG0004444 | Gas2l3 | growth arrest-specific 2 like 3 | 223.61 | 460.50 | -1.04 | 2.65E-07 |
| ENSMUSG00000069539 | GSMG0004448 | Scyl2 | SCY1-like 2 (S. cerevisiae) | 531.36 | 555.07 | -0.06 | 8.53E-01 |
| ENSMUSG00000019948 | GSMG0004449 | Actr6 | ARP6 actin-related protein 6 | 286.91 | 381.59 | -0.41 | 9.82E-02 |
| ENSMUSG00000019979 | GSMG0004452 | Apaf1 | apoptotic peptidase activating factor 1 | 442.64 | 583.92 | -0.40 | 4.10E-02 |
| ENSMUSG00000061904 | GSMG0004454 | Slc25a3 | solute carrier family 25 (mitochondrial carrier, phosphate carrier), mem | 5306.66 | 4902.02 | 0.11 | 6.07E-01 |
| NULL | GSMG0004457 | NULL | NULL | 104.53 | 81.55 | 0.35 | 4.60E-01 |
| ENSMUSG00000019961 | GSMG0004458 | Tmpo | thymopoietin | 846.14 | 1230.96 | -0.54 | 1.22E-03 |
| ENSMUSG00000019988 | GSMG0004466 | Nedd1 | neural precursor cell expressed, developmentally down-regulated gene 1 | 579.79 | 595.01 | -0.04 | 9.21E-01 |
| NULL | GSMG0004471 | NULL | NULL | 123.04 | 81.56 | 0.60 | 6.88E-02 |
| ENSMUSG00000008398 | GSMG0004472 | Elk3 | ELK3, member of ETS oncogene family | 606.85 | 880.59 | -0.54 | 9.57E-03 |
| ENSMUSG00000020018 | GSMG0004475 | Snrpf | small nuclear ribonucleoprotein polypeptide F | 160.93 | 212.10 | -0.40 | 2.44E-01 |
| ENSMUSG00000036112 | GSMG0004477 | Metap2 | methionine aminopeptidase 2 | 1981.06 | 2247.60 | -0.18 | 4.41E-01 |
| NULL | GSMG0004480 | NULL | NULL | 94.09 | 105.66 | -0.16 | 7.37E-01 |
| ENSMUSG00000097164 | GSMG0004485 | Cep83os | centrosomal protein 83, opposite strand | 566.63 | 523.84 | 0.11 | 6.83E-01 |
| ENSMUSG00000074785 | GSMG0004486 | Plxnc1 | plexin C1 | 195.83 | 278.67 | -0.51 | 7.20E-02 |
| ENSMUSG00000062981 | GSMG0004497 | Mrpl42 | mitochondrial ribosomal protein L42 | 1027.03 | 1255.86 | -0.29 | 1.51E-01 |
| ENSMUSG00000020029 | GSMG0004499 | Nudt4 | nudix (nucleoside diphosphate linked moiety X)-type motif 4 | 3444.57 | 3361.69 | 0.03 | 8.98E-01 |
| NULL | GSMG0004500 | NULL | NULL | 185.66 | 229.04 | -0.31 | 3.63E-01 |

| | | | | | | | |
|---|---|---|---|---|---|---|---|
| ENSMUSG00000036676 | GSMG0004518 | Tmtc3 | transmembrane and tetratricopeptide repeat containing 3 | 607.43 | 573.64 | 0.08 | 7.73E-01 |
| ENSMUSG00000046567 | GSMG0004519 | 4930430F08Rik | RIKEN cDNA 4930430F08 gene | 151.80 | 135.56 | 0.16 | 6.94E-01 |
| ENSMUSG00000019890 | GSMG0004523 | Nts | neurotensin | 113.26 | 389.29 | -1.78 | 1.68E-13 |
| ENSMUSG00000036019 | GSMG0004529 | Tmtc2 | transmembrane and tetratricopeptide repeat containing 2 | 486.67 | 299.18 | 0.70 | 4.79E-04 |
| NULL | GSMG0004537 | NULL | NULL | 328.37 | 223.21 | 0.56 | 1.66E-02 |
| NULL | GSMG0004539 | NULL | NULL | 261.33 | 168.35 | 0.63 | 1.44E-02 |
| ENSMUSG00000036009 | GSMG0004542 | Mettl25 | methyltransferase like 25 | 240.92 | 265.78 | -0.14 | 6.66E-01 |
| ENSMUSG00000035864 | GSMG0004558 | Syt1 | synaptotagmin I | 2559.40 | 3256.69 | -0.35 | 3.12E-02 |
| NULL | GSMG0004562 | NULL | NULL | 866.62 | 477.21 | 0.86 | 8.66E-08 |
| NULL | GSMG0004563 | NULL | NULL | 210.24 | 154.61 | 0.44 | 9.16E-02 |
| NULL | GSMG0004567 | NULL | NULL | 155.75 | 78.58 | 0.98 | 2.19E-03 |
| NULL | GSMG0004568 | NULL | NULL | 508.55 | 340.11 | 0.58 | 1.91E-02 |
| NULL | GSMG0004569 | NULL | NULL | 669.29 | 410.20 | 0.71 | 2.37E-03 |
| NULL | GSMG0004570 | NULL | NULL | 242.71 | 167.81 | 0.53 | 4.06E-02 |
| NULL | GSMG0004571 | NULL | NULL | 606.68 | 333.09 | 0.87 | 1.23E-05 |
| NULL | GSMG0004574 | NULL | NULL | 324.34 | 222.98 | 0.54 | 3.72E-02 |
| NULL | GSMG0004575 | NULL | NULL | 395.00 | 258.87 | 0.61 | 2.00E-02 |
| NULL | GSMG0004576 | NULL | NULL | 311.34 | 247.04 | 0.34 | 2.80E-01 |
| NULL | GSMG0004589 | NULL | NULL | 855.52 | 677.57 | 0.34 | 1.18E-01 |
| NULL | GSMG0004592 | NULL | NULL | 907.83 | 644.61 | 0.49 | 2.84E-02 |
| NULL | GSMG0004593 | NULL | NULL | 929.51 | 639.40 | 0.54 | 1.12E-02 |
| NULL | GSMG0004596 | NULL | NULL | 133.91 | 197.05 | -0.57 | 8.53E-02 |
| ENSMUSG00000035798 | GSMG0004597 | Zdhhc17 | zinc finger, DHHC domain containing 17 | 395.13 | 453.65 | -0.20 | 4.30E-01 |
| NULL | GSMG0004598 | NULL | NULL | 1215.21 | 919.21 | 0.40 | 8.88E-02 |
| ENSMUSG00000074748 | GSMG0004603 | Atxn7l3b | ataxin 7-like 3B | 6003.45 | 7491.73 | -0.32 | 1.33E-01 |
| ENSMUSG00000020130 | GSMG0004610 | Tbc1d15 | TBC1 domain family, member 15 | 756.21 | 795.93 | -0.07 | 8.22E-01 |
| NULL | GSMG0004611 | NULL | NULL | 153.08 | 99.77 | 0.61 | 1.43E-01 |
| ENSMUSG00000020132 | GSMG0004612 | Rab21 | RAB21, member RAS oncogene family | 324.17 | 201.37 | 0.68 | 3.64E-02 |
| ENSMUSG00000069520 | GSMG0004613 | Tmem19 | transmembrane protein 19 | 715.20 | 633.71 | 0.17 | 5.02E-01 |
| ENSMUSG00000054934 | GSMG0004620 | Kcnmb4 | potassium large conductance calcium-activated channel, subfamily M, b | 215.77 | 212.77 | 0.02 | 9.74E-01 |
| ENSMUSG00000020166 | GSMG0004621 | Cnot2 | CCR4-NOT transcription complex, subunit 2 | 1269.18 | 1557.48 | -0.29 | 1.18E-01 |
| ENSMUSG00000064181 | GSMG0004628 | Rab3ip | RAB3A interacting protein | 191.32 | 216.91 | -0.18 | 6.09E-01 |
| ENSMUSG00000034024 | GSMG0004629 | Cct2 | chaperonin containing Tcp1, subunit 2 (beta) | 1556.04 | 2302.26 | -0.56 | 1.83E-03 |

| | | | | | | | |
|---|---|---|---|---|---|---|---|
| ENSMUSG00000020170 | GSMG0004630 | Frs2 | fibroblast growth factor receptor substrate 2 | 1140.40 | 1101.81 | 0.05 | 8.79E-01 |
| ENSMUSG00000020171 | GSMG0004634 | Yeats4 | YEATS domain containing 4 | 667.33 | 968.24 | -0.54 | 1.19E-02 |
| ENSMUSG00000055531 | GSMG0004640 | Cpsf6 | cleavage and polyadenylation specific factor 6 | 2957.88 | 3566.49 | -0.27 | 2.25E-01 |
| ENSMUSG00000020184 | GSMG0004642 | Mdm2 | transformed mouse 3T3 cell double minute 2 | 1763.38 | 1450.72 | 0.28 | 1.14E-01 |
| ENSMUSG00000060181 | GSMG0004643 | Slc35e3 | solute carrier family 35, member E3 | 634.93 | 805.68 | -0.34 | 6.77E-02 |
| ENSMUSG00000052798 | GSMG0004644 | Nup107 | nucleoporin 107 | 703.90 | 1085.83 | -0.63 | 3.54E-04 |
| ENSMUSG00000052681 | GSMG0004645 | Rap1b | RAS related protein 1b | 3198.55 | 3127.18 | 0.03 | 9.13E-01 |
| ENSMUSG00000028630 | GSMG0004653 | Dyrk2 | dual-specificity tyrosine-(Y)-phosphorylation regulated kinase 2 | 1055.42 | 1517.04 | -0.52 | 1.35E-02 |
| ENSMUSG00000020114 | GSMG0004659 | Cand1 | cullin associated and neddylation disassociated 1 | 442.21 | 518.99 | -0.23 | 3.19E-01 |
| ENSMUSG00000020228 | GSMG0004667 | Helb | helicase (DNA) B | 51.96 | 56.78 | -0.11 | 8.94E-01 |
| ENSMUSG00000051236 | GSMG0004677 | Msrb3 | methionine sulfoxide reductase B3 | 5733.46 | 4416.77 | 0.38 | 1.55E-02 |
| ENSMUSG00000048661 | GSMG0004680 | Lemd3 | LEM domain containing 3 | 255.05 | 248.26 | 0.04 | 9.36E-01 |
| ENSMUSG00000052302 | GSMG0004686 | Tbc1d30 | TBC1 domain family, member 30 | 232.30 | 156.67 | 0.57 | 2.82E-02 |
| ENSMUSG00000025795 | GSMG0004687 | Rassf3 | Ras association (RalGDS/AF-6) domain family member 3 | 5927.44 | 4500.67 | 0.40 | 1.92E-02 |
| ENSMUSG00000020115 | GSMG0004689 | Tbk1 | TANK-binding kinase 1 | 1375.43 | 1432.71 | -0.06 | 8.34E-01 |
| ENSMUSG00000020121 | GSMG0004696 | Srgap1 | SLIT-ROBO Rho GTPase activating protein 1 | 139.17 | 268.57 | -0.94 | 2.38E-03 |
| NULL | GSMG0004699 | NULL | NULL | 134.44 | 210.40 | -0.65 | 9.45E-03 |
| ENSMUSG00000034620 | GSMG0004700 | Tmem5 | transmembrane protein 5 | 689.76 | 784.99 | -0.19 | 4.06E-01 |
| ENSMUSG00000065406 | GSMG0004705 | Mirlet7i | microRNA let7i | 64.34 | 56.40 | 0.18 | 8.12E-01 |
| ENSMUSG00000034602 | GSMG0004707 | Mon2 | MON2 homolog, regulator of endosome to Golgi trafficking | 1215.76 | 916.84 | 0.41 | 1.93E-02 |
| ENSMUSG00000020124 | GSMG0004710 | Usp15 | ubiquitin specific peptidase 15 | 784.44 | 900.47 | -0.20 | 4.19E-01 |
| ENSMUSG00000020102 | GSMG0004717 | Slc16a7 | solute carrier family 16 (monocarboxylic acid transporters), member 7 | 126.64 | 169.74 | -0.42 | 1.67E-01 |
| ENSMUSG00000025436 | GSMG0004725 | Xrcc6bp1 | XRCC6 binding protein 1 | 114.58 | 159.73 | -0.48 | 1.44E-01 |
| ENSMUSG00000040521 | GSMG0004728 | Tsfm | Ts translation elongation factor, mitochondrial | 268.23 | 370.86 | -0.47 | 3.76E-02 |
| ENSMUSG00000040502 | GSMG0004729 | March9 | membrane-associated ring finger (C3HC4) 9 | 68.98 | 129.00 | -0.91 | 8.90E-03 |
| ENSMUSG00000006736 | GSMG0004730 | Tspan31 | tetraspanin 31 | 4221.49 | 4435.35 | -0.07 | 7.76E-01 |
| ENSMUSG00000040462 | GSMG0004732 | Os9 | amplified in osteosarcoma | 1147.04 | 1232.64 | -0.10 | 6.70E-01 |
| ENSMUSG00000019467 | GSMG0004735 | Arhgef25 | Rho guanine nucleotide exchange factor (GEF) 25 | 1950.33 | 1254.17 | 0.64 | 5.89E-04 |
| ENSMUSG00000040415 | GSMG0004736 | Dtx3 | deltex 3, E3 ubiquitin ligase | 5563.48 | 4265.73 | 0.38 | 1.32E-02 |
| ENSMUSG00000025417 | GSMG0004737 | Pip4k2c | phosphatidylinositol-5-phosphate 4-kinase, type II, gamma | 412.59 | 309.67 | 0.41 | 1.36E-01 |
| ENSMUSG00000074657 | GSMG0004738 | Kif5a | kinesin family member 5A | 1075.32 | 868.63 | 0.31 | 2.35E-01 |
| ENSMUSG00000040354 | GSMG0004742 | Mars | methionine-tRNA synthetase | 2930.23 | 3866.85 | -0.40 | 1.01E-02 |
| ENSMUSG00000025403 | GSMG0004746 | Shmt2 | serine hydroxymethyltransferase 2 (mitochondrial) | 954.39 | 1334.31 | -0.48 | 3.34E-03 |

| | | | | | | | |
|---|---|---|---|---|---|---|---|
| ENSMUSG00000040258 | GSMG0004747 | Nxph4 | neurexophilin 4 | 179.79 | 252.31 | -0.50 | 8.22E-02 |
| ENSMUSG00000040249 | GSMG0004748 | Lrp1 | low density lipoprotein receptor-related protein 1 | 2641.74 | 2375.01 | 0.15 | 6.74E-01 |
| ENSMUSG00000025402 | GSMG0004750 | Nab2 | Ngfi-A binding protein 2 | 607.22 | 294.49 | 1.04 | 3.66E-05 |
| ENSMUSG00000058396 | GSMG0004751 | Gpr182 | G protein-coupled receptor 182 | 389.25 | 295.53 | 0.40 | 2.43E-01 |
| ENSMUSG00000040043 | GSMG0004756 | Rbms2 | RNA binding motif, single stranded interacting protein 2 | 655.96 | 590.77 | 0.15 | 5.63E-01 |
| ENSMUSG00000051346 | GSMG0004758 | Spryd4 | SPRY domain containing 4 | 427.80 | 506.64 | -0.24 | 3.23E-01 |
| ENSMUSG00000039914 | GSMG0004765 | Coq10a | coenzyme Q10A | 1117.03 | 947.71 | 0.24 | 2.35E-01 |
| ENSMUSG00000025374 | GSMG0004767 | Nabp2 | nucleic acid binding protein 2 | 510.94 | 575.59 | -0.17 | 5.06E-01 |
| ENSMUSG00000025366 | GSMG0004770 | Esyt1 | extended synaptotagmin-like protein 1 | 5356.30 | 5182.06 | 0.05 | 8.53E-01 |
| ENSMUSG00000025364 | GSMG0004773 | Pa2g4 | proliferation-associated 2G4 | 514.50 | 901.10 | -0.81 | 3.71E-05 |
| ENSMUSG00000049858 | GSMG0004778 | Suox | sulfite oxidase | 599.77 | 485.82 | 0.30 | 1.71E-01 |
| ENSMUSG00000025358 | GSMG0004780 | Cdk2 | cyclin-dependent kinase 2 | 501.61 | 667.49 | -0.41 | 4.18E-02 |
| ENSMUSG00000025357 | GSMG0004781 | Dgka | diacylglycerol kinase, alpha | 862.83 | 790.66 | 0.13 | 6.24E-01 |
| ENSMUSG00000047090 | GSMG0004782 | Tmem198b | transmembrane protein 198b | 834.20 | 750.21 | 0.15 | 5.38E-01 |
| ENSMUSG00000025353 | GSMG0004783 | Ormdl2 | ORM1-like 2 (S. cerevisiae) | 138.40 | 125.05 | 0.15 | 7.27E-01 |
| ENSMUSG00000025350 | GSMG0004787 | Rdh5 | retinol dehydrogenase 5 | 265.61 | 140.67 | 0.92 | 1.13E-03 |
| ENSMUSG00000090247 | GSMG0004788 | Bloc1s1 | biogenesis of lysosomal organelles complex-1, subunit 1 | 1695.77 | 1751.20 | -0.05 | 8.81E-01 |
| ENSMUSG00000020454 | GSMG0004809 | Eif4enif1 | eukaryotic translation initiation factor 4E nuclear import factor 1 | 1124.81 | 1273.59 | -0.18 | 4.27E-01 |
| ENSMUSG00000020453 | GSMG0004812 | Patz1 | POZ (BTB) and AT hook containing zinc finger 1 | 1092.06 | 1220.84 | -0.16 | 5.48E-01 |
| ENSMUSG00000034614 | GSMG0004815 | Pik3ip1 | phosphoinositide-3-kinase interacting protein 1 | 987.91 | 534.23 | 0.89 | 1.48E-07 |
| ENSMUSG00000034587 | GSMG0004816 | 8430429K09Rik | RIKEN cDNA 8430429K09 gene | 242.78 | 296.25 | -0.29 | 3.03E-01 |
| ENSMUSG00000034579 | GSMG0004819 | Pla2g3 | phospholipase A2, group III | 125.14 | 274.18 | -1.13 | 3.53E-06 |
| ENSMUSG00000075702 | GSMG0004820 | Selm | selenoprotein M | 2231.92 | 2739.50 | -0.30 | 2.48E-01 |
| NULL | GSMG0004821 | NULL | NULL | 403.37 | 423.53 | -0.07 | 8.34E-01 |
| ENSMUSG00000034543 | GSMG0004823 | Morc2a | microrchidia 2A | 928.91 | 1240.80 | -0.42 | 6.48E-02 |
| ENSMUSG00000047205 | GSMG0004825 | Dusp18 | dual specificity phosphatase 18 | 404.60 | 517.45 | -0.36 | 1.72E-01 |
| ENSMUSG00000020430 | GSMG0004826 | Pes1 | pescadillo ribosomal biogenesis factor 1 | 361.49 | 475.95 | -0.40 | 6.22E-02 |
| ENSMUSG00000049721 | GSMG0004827 | Gal3st1 | galactose-3-O-sulfotransferase 1 | 289.65 | 90.41 | 1.68 | 5.51E-10 |
| ENSMUSG00000003581 | GSMG0004832 | Rnf215 | ring finger protein 215 | 1951.65 | 1732.75 | 0.17 | 4.06E-01 |
| ENSMUSG00000002129 | GSMG0004833 | Sf3a1 | splicing factor 3a, subunit 1 | 1951.02 | 2420.84 | -0.31 | 5.08E-02 |
| ENSMUSG00000034412 | GSMG0004834 | Tbc1d10a | TBC1 domain family, member 10a | 969.07 | 805.32 | 0.27 | 1.94E-01 |
| ENSMUSG00000020424 | GSMG0004835 | Gatsl3 | GATS protein-like 3 | 1199.21 | 523.37 | 1.20 | 3.25E-15 |
| ENSMUSG00000020412 | GSMG0004840 | Ascc2 | activating signal cointegrator 1 complex subunit 2 | 1250.18 | 1603.59 | -0.36 | 4.93E-02 |

| | | | | | | | |
|---|---|---|---|---|---|---|---|
| ENSMUSG00000009076 | GSMG0004841 | Zmat5 | zinc finger, matrin type 5 | 442.06 | 502.03 | -0.19 | 5.46E-01 |
| NULL | GSMG0004843 | NULL | NULL | 149.92 | 180.85 | -0.27 | 4.49E-01 |
| NULL | GSMG0004844 | NULL | NULL | 124.50 | 163.52 | -0.39 | 2.86E-01 |
| ENSMUSG00000034285 | GSMG0004845 | Nipsnap1 | nipsnap homolog 1 (C. elegans) | 363.87 | 448.30 | -0.30 | 2.25E-01 |
| ENSMUSG00000034274 | GSMG0004847 | Thoc5 | THO complex 5 | 970.69 | 1230.66 | -0.34 | 6.55E-02 |
| ENSMUSG00000009090 | GSMG0004848 | Ap1b1 | adaptor protein complex AP-1, beta 1 subunit | 2778.88 | 2956.25 | -0.09 | 7.41E-01 |
| ENSMUSG00000034209 | GSMG0004850 | Rasl10a | RAS-like, family 10, member A | 415.24 | 259.86 | 0.68 | 1.06E-03 |
| ENSMUSG00000034175 | GSMG0004851 | Rhbdd3 | rhomboid domain containing 3 | 626.33 | 668.19 | -0.09 | 7.78E-01 |
| ENSMUSG00000020484 | GSMG0004852 | Xbp1 | X-box binding protein 1 | 6257.71 | 6467.23 | -0.05 | 8.72E-01 |
| ENSMUSG00000083844 | GSMG0004855 | Ube2d-ps | ubiquitin-conjugating enzyme E2D, pseudogene | 483.75 | 467.12 | 0.05 | 8.80E-01 |
| ENSMUSG00000020476 | GSMG0004856 | Dbnl | drebrin-like | 1910.63 | 2055.85 | -0.11 | 6.36E-01 |
| ENSMUSG00000020473 | GSMG0004858 | Aebp1 | AE binding protein 1 | 653.04 | 907.26 | -0.47 | 9.65E-02 |
| ENSMUSG00000002741 | GSMG0004860 | Ykt6 | YKT6 v-SNARE homolog (S. cerevisiae) | 2021.57 | 2555.54 | -0.34 | 1.06E-01 |
| ENSMUSG00000020456 | GSMG0004863 | Ogdh | oxoglutarate (alpha-ketoglutarate) dehydrogenase (lipoamide) | 1526.17 | 1417.72 | 0.11 | 6.58E-01 |
| ENSMUSG00000041164 | GSMG0004865 | Zmiz2 | zinc finger, MIZ-type containing 2 | 583.31 | 552.28 | 0.08 | 8.73E-01 |
| NULL | GSMG0004867 | NULL | NULL | 720.62 | 219.53 | 1.71 | 4.71E-02 |
| ENSMUSG00000000378 | GSMG0004868 | Ccm2 | cerebral cavernous malformation 2 | 1003.24 | 1018.33 | -0.02 | 9.43E-01 |
| NULL | GSMG0004871 | NULL | NULL | 478.83 | 583.04 | -0.28 | 3.80E-01 |
| ENSMUSG00000041046 | GSMG0004873 | Ramp3 | receptor (calcitonin) activity modifying protein 3 | 391.48 | 311.70 | 0.33 | 1.61E-01 |
| ENSMUSG00000020431 | GSMG0004874 | Adcy1 | adenylate cyclase 1 | 109.05 | 139.45 | -0.35 | 5.35E-01 |
| ENSMUSG00000048834 | GSMG0004898 | Vstm2a | V-set and transmembrane domain containing 2A | 291.94 | 432.59 | -0.57 | 1.78E-02 |
| ENSMUSG00000020122 | GSMG0004899 | Egfr | epidermal growth factor receptor | 146.42 | 189.96 | -0.37 | 2.33E-01 |
| ENSMUSG00000044629 | GSMG0004905 | Cnrip1 | cannabinoid receptor interacting protein 1 | 947.90 | 1414.02 | -0.58 | 1.64E-04 |
| G00000033953,ENSMUSG000 | GSMG0004906 | Ppp3r1 // Wdr92 | protein phosphatase 3, regulatory subunit B, alpha isoform (calcineurin | 4853.41 | 4127.73 | 0.23 | 2.84E-01 |
| ENSMUSG00000000581 | GSMG0004907 | C1d | C1D nuclear receptor co-repressor | 1273.65 | 1701.68 | -0.42 | 8.35E-02 |
| ENSMUSG00000045671 | GSMG0004916 | Spred2 | sprouty-related, EVH1 domain containing 2 | 151.82 | 180.67 | -0.25 | 4.57E-01 |
| NULL | GSMG0004918 | NULL | NULL | 311.07 | 182.14 | 0.78 | 3.90E-04 |
| ENSMUSG00000020149 | GSMG0004920 | Rab1a | RAB1A, member RAS oncogene family | 5630.60 | 6573.42 | -0.22 | 2.24E-01 |
| ENSMUSG00000049800 | GSMG0004922 | Sertad2 | SERTA domain containing 2 | 1264.89 | 1187.31 | 0.09 | 7.23E-01 |
| ENSMUSG00000020134 | GSMG0004925 | Peli1 | pellino 1 | 1196.31 | 1242.21 | -0.05 | 8.57E-01 |
| ENSMUSG00000020128 | GSMG0004930 | Vps54 | VPS54 GARP complex subunit | 740.01 | 571.54 | 0.37 | 1.43E-01 |
| ENSMUSG00000020319 | GSMG0004934 | Wdpcp | WD repeat containing planar cell polarity effector | 116.64 | 114.61 | 0.03 | 9.55E-01 |
| ENSMUSG00000049904 | GSMG0004938 | Tmem17 | transmembrane protein 17 | 173.33 | 280.54 | -0.69 | 1.46E-02 |

| | | | | | | | |
|---|---|---|---|---|---|---|---|
| ENSMUSG00000044068 | GSMG0004942 | Zrsr1 | zinc finger (CCCH type), RNA binding motif and serine/arginine rich 1 | 1072.62 | 734.59 | 0.55 | 7.28E-04 |
| ENSMUSG00000007739 | GSMG0004945 | Cct4 | chaperonin containing Tcp1, subunit 4 (delta) | 2746.32 | 3558.09 | -0.37 | 4.56E-02 |
| ENSMUSG00000049811 | GSMG0004947 | Fam161a | family with sequence similarity 161, member A | 230.75 | 134.18 | 0.78 | 7.99E-03 |
| ENSMUSG00000020290 | GSMG0004950 | Xpo1 | exportin 1 | 1577.44 | 2444.53 | -0.63 | 3.04E-05 |
| ENSMUSG00000020280 | GSMG0004959 | Pus10 | pseudouridylate synthase 10 | 424.64 | 553.42 | -0.38 | 1.03E-01 |
| ENSMUSG00000000861 | GSMG0004963 | Bcl11a | B cell CLL/lymphoma 11A (zinc finger protein) | 209.68 | 502.46 | -1.27 | 1.52E-09 |
| NULL | GSMG0004964 | NULL | NULL | 454.71 | 513.11 | -0.18 | 5.38E-01 |
| ENSMUSG00000004018 | GSMG0004970 | Fancl | Fanconi anemia, complementation group L | 233.18 | 331.43 | -0.50 | 4.12E-02 |
| ENSMUSG00000020467 | GSMG0004975 | Efemp1 | epidermal growth factor-containing fibulin-like extracellular matrix pro | 131.45 | 118.46 | 0.16 | 7.56E-01 |
| ENSMUSG00000020464 | GSMG0004976 | Pnpt1 | polyribonucleotide nucleotidyltransferase 1 | 1051.18 | 1479.72 | -0.49 | 5.94E-03 |
| ENSMUSG00000020463 | GSMG0004977 | Smek2 | SMEK homolog 2, suppressor of mek1 (Dictyostelium) | 1649.22 | 1771.25 | -0.10 | 7.26E-01 |
| ENSMUSG00000032740 | GSMG0004982 | Ccdc88a | coiled coil domain containing 88A | 581.29 | 779.41 | -0.42 | 2.47E-01 |
| ENSMUSG00000020459 | GSMG0004983 | Mtif2 | mitochondrial translational initiation factor 2 | 567.87 | 599.21 | -0.08 | 8.32E-01 |
| ENSMUSG00000020458 | GSMG0004986 | Rtn4 | reticulon 4 | 661.16 | 682.75 | -0.05 | 9.02E-01 |
| NULL | GSMG0004991 | NULL | NULL | 108.87 | 104.58 | 0.05 | 9.22E-01 |
| ENSMUSG00000040850 | GSMG0004994 | Psme4 | proteasome (prosome, macropain) activator subunit 4 | 285.34 | 216.92 | 0.40 | 2.04E-01 |
| ENSMUSG00000020305 | GSMG0004996 | Asb3 | ankyrin repeat and SOCS box-containing 3 | 372.33 | 384.27 | -0.04 | 9.05E-01 |
| NULL | GSMG0005000 | NULL | NULL | 287.78 | 263.10 | 0.13 | 7.18E-01 |
| ENSMUSG00000020300 | GSMG0005001 | Cpeb4 | cytoplasmic polyadenylation element binding protein 4 | 143.27 | 164.52 | -0.20 | 5.87E-01 |
| ENSMUSG00000020297 | GSMG0005004 | Nsg2 | neuron specific gene family member 2 | 2411.92 | 5701.06 | -1.24 | 5.40E-16 |
| ENSMUSG00000040767 | GSMG0005005 | Snrnp25 | small nuclear ribonucleoprotein 25 (U11/U12) | 368.04 | 373.36 | -0.02 | 9.65E-01 |
| ENSMUSG00000020287 | GSMG0005006 | Mpg | N-methylpurine-DNA glycosylase | 1073.57 | 1188.68 | -0.15 | 5.06E-01 |
| ENSMUSG00000040711 | GSMG0005012 | Sh3pxd2b | SH3 and PX domains 2B | 81.00 | 151.44 | -0.90 | 1.55E-03 |
| ENSMUSG00000044949 | GSMG0005014 | Ubtd2 | ubiquitin domain containing 2 | 573.30 | 805.92 | -0.49 | 2.69E-02 |
| ENSMUSG00000020271 | GSMG0005019 | Fbxw11 | F-box and WD-40 domain protein 11 | 2146.40 | 2494.43 | -0.22 | 2.74E-01 |
| ENSMUSG00000020155 | GSMG0005027 | Kcnmb1 | potassium large conductance calcium-activated channel, subfamily M, b | 113.22 | 72.83 | 0.64 | 6.41E-02 |
| ENSMUSG00000002699 | GSMG0005028 | Lcp2 | lymphocyte cytosolic protein 2 | 186.05 | 288.00 | -0.62 | 3.85E-02 |
| ENSMUSG00000056427 | GSMG0005034 | Slit3 | slit homolog 3 (Drosophila) | 1047.44 | 552.16 | 0.92 | 1.81E-05 |
| ENSMUSG00000018846 | GSMG0005042 | Pank3 | pantothenate kinase 3 | 322.29 | 416.05 | -0.37 | 1.87E-01 |
| ENSMUSG00000020328 | GSMG0005050 | Nudcd2 | NudC domain containing 2 | 602.91 | 822.94 | -0.45 | 5.66E-02 |
| ENSMUSG00000007653 | GSMG0005055 | Gabrb2 | gamma-aminobutyric acid (GABA) A receptor, subunit beta 2 | 230.34 | 308.11 | -0.42 | 1.28E-01 |
| ENSMUSG00000020409 | GSMG0005064 | Slu7 | SLU7 splicing factor homolog (S. cerevisiae) | 1635.75 | 1544.96 | 0.08 | 7.69E-01 |
| ENSMUSG00000046491 | GSMG0005065 | C1qtnf2 | C1q and tumor necrosis factor related protein 2 | 448.07 | 336.88 | 0.41 | 6.55E-02 |

| | | | | | | | |
|---|---|---|---|---|---|---|---|
| ENSMUSG00000044707 | GSMG0005066 | Ccnjl | cyclin J-like | 257.00 | 420.92 | -0.71 | 2.68E-04 |
| ENSMUSG00000044950 | GSMG0005069 | Pwwp2a | PWWP domain containing 2A | 610.30 | 606.37 | 0.01 | 9.83E-01 |
| ENSMUSG00000019189 | GSMG0005074 | Rnf145 | ring finger protein 145 | 817.24 | 850.17 | -0.06 | 8.69E-01 |
| ENSMUSG00000057098 | GSMG0005077 | Ebf1 | early B cell factor 1 | 1894.27 | 1330.96 | 0.51 | 8.32E-03 |
| NULL | GSMG0005078 | NULL | NULL | 746.11 | 490.31 | 0.61 | 2.93E-02 |
| NULL | GSMG0005080 | NULL | NULL | 595.81 | 437.54 | 0.45 | 1.53E-01 |
| NULL | GSMG0005086 | NULL | NULL | 290.85 | 185.73 | 0.65 | 1.57E-02 |
| NULL | GSMG0005091 | NULL | NULL | 261.72 | 203.00 | 0.37 | 1.81E-01 |
| ENSMUSG00000006169 | GSMG0005092 | Clint1 | clathrin interactor 1 | 1665.07 | 1613.83 | 0.05 | 8.75E-01 |
| ENSMUSG00000011256 | GSMG0005094 | Adam19 | a disintegrin and metallopeptidase domain 19 (meltrin beta) | 709.58 | 1286.00 | -0.86 | 2.50E-06 |
| NULL | GSMG0005096 | NULL | NULL | 61.25 | 45.14 | 0.45 | 3.88E-01 |
| ENSMUSG00000020397 | GSMG0005097 | Med7 | mediator complex subunit 7 | 623.77 | 613.83 | 0.02 | 9.50E-01 |
| ENSMUSG00000020372 | GSMG0005106 | Rack1 | receptor for activated C kinase 1 | 2682.02 | 3716.95 | -0.47 | 1.06E-03 |
| ENSMUSG00000078920 | GSMG0005108 | Ifi47 | interferon gamma inducible protein 47 | 437.93 | 435.38 | 0.01 | 9.85E-01 |
| ENSMUSG00000046311 | GSMG0005110 | Zfp62 | zinc finger protein 62 | 320.29 | 287.28 | 0.16 | 5.99E-01 |
| ENSMUSG00000020346 | GSMG0005111 | Mgat1 | mannoside acetylglucosaminyltransferase 1 | 2871.52 | 3248.49 | -0.18 | 3.60E-01 |
| ENSMUSG00000020357 | GSMG0005124 | Flt4 | FMS-like tyrosine kinase 4 | 3001.26 | 4238.26 | -0.50 | 2.58E-02 |
| ENSMUSG00000064057 | GSMG0005125 | Scgb3a1 | secretoglobin, family 3A, member 1 | 251.89 | 271.67 | -0.11 | 7.58E-01 |
| ENSMUSG00000020363 | GSMG0005127 | Gfpt2 | glutamine fructose-6-phosphate transaminase 2 | 144.29 | 203.22 | -0.50 | 1.40E-01 |
| ENSMUSG00000020366 | GSMG0005128 | Mapk9 | mitogen-activated protein kinase 9 | 929.92 | 1027.16 | -0.14 | 5.47E-01 |
| ENSMUSG00000020374 | GSMG0005130 | Rasgef1c | RasGEF domain family, member 1C | 240.88 | 186.74 | 0.36 | 2.14E-01 |
| ENSMUSG00000020376 | GSMG0005131 | Rnf130 | ring finger protein 130 | 2122.06 | 1745.53 | 0.28 | 1.17E-01 |
| ENSMUSG00000036644 | GSMG0005134 | Tbc1d9b | TBC1 domain family, member 9B | 5363.22 | 4422.70 | 0.28 | 2.01E-01 |
| ENSMUSG00000020381 | GSMG0005135 | 3010026O09Rik | RIKEN cDNA 3010026O09 gene | 873.50 | 808.86 | 0.11 | 6.75E-01 |
| ENSMUSG00000036620 | GSMG0005137 | Mgat4b | mannoside acetylglucosaminyltransferase 4, isoenzyme B | 4025.81 | 4677.39 | -0.22 | 3.30E-01 |
| ENSMUSG00000007850 | GSMG0005142 | Hnrnph1 | heterogeneous nuclear ribonucleoprotein H1 | 2267.23 | 2915.62 | -0.36 | 8.01E-02 |
| ENSMUSG00000036545 | GSMG0005143 | Adamts2 | a disintegrin-like and metallopeptidase (reprolysin type) with thrombos | 240.77 | 155.93 | 0.63 | 5.32E-02 |
| ENSMUSG00000020364 | GSMG0005153 | Zfp354a | zinc finger protein 354A | 583.01 | 591.41 | -0.02 | 9.57E-01 |
| ENSMUSG00000020385 | GSMG0005155 | Clk4 | CDC like kinase 4 | 1951.96 | 1448.69 | 0.43 | 1.49E-02 |
| ENSMUSG00000063564 | GSMG0005156 | Col23a1 | collagen, type XXIII, alpha 1 | 468.63 | 271.62 | 0.79 | 1.02E-03 |
| ENSMUSG00000020359 | GSMG0005161 | Phykpl | 5-phosphohydroxy-L-lysine phospholyase | 262.80 | 208.25 | 0.34 | 2.29E-01 |
| ENSMUSG00000001056 | GSMG0005163 | Nhp2 | NHP2 ribonucleoprotein | 709.10 | 1271.71 | -0.84 | 2.48E-07 |
| ENSMUSG00000052563 | GSMG0005164 | D930048N14Rik | RIKEN cDNA D930048N14 gene | 746.31 | 650.00 | 0.20 | 3.84E-01 |

| | | | | | | | |
|---|---|---|---|---|---|---|---|
| ENSMUSG00000020386 | GSMG0005166 | Sar1b | secretion associated Ras related GTPase 1B | 1383.69 | 1787.28 | -0.37 | 5.57E-02 |
| ENSMUSG00000020392 | GSMG0005168 | Cdkn2aipnl | CDKN2A interacting protein N-terminal like | 1106.18 | 1650.54 | -0.58 | 2.57E-04 |
| NULL | GSMG0005169 | NULL | NULL | 141.47 | 117.09 | 0.28 | 4.97E-01 |
| ENSMUSG00000020389 | GSMG0005170 | Cdkl3 | cyclin-dependent kinase-like 3 | 134.98 | 248.64 | -0.88 | 2.68E-04 |
| ENSMUSG00000020349 | GSMG0005171 | Ppp2ca | protein phosphatase 2 (formerly 2A), catalytic subunit, alpha isoform | 3207.75 | 3591.94 | -0.16 | 4.89E-01 |
| ENSMUSG00000036275 | GSMG0005172 | 9530068E07Rik | RIKEN cDNA 9530068E07 gene | 1305.65 | 1514.31 | -0.21 | 3.01E-01 |
| ENSMUSG00000036309 | GSMG0005174 | Skp1a | S-phase kinase-associated protein 1A | 5688.51 | 8440.69 | -0.57 | 2.57E-03 |
| ENSMUSG00000020402 | GSMG0005176 | Vdac1 | voltage-dependent anion channel 1 | 1658.43 | 2197.77 | -0.41 | 1.88E-02 |
| ENSMUSG00000036264 | GSMG0005178 | Fstl4 | follistatin-like 4 | 338.46 | 254.38 | 0.41 | 1.77E-01 |
| ENSMUSG00000018239 | GSMG0005181 | Zcchc10 | zinc finger, CCHC domain containing 10 | 272.31 | 284.46 | -0.06 | 8.74E-01 |
| ENSMUSG00000049470 | GSMG0005182 | Aff4 | AF4/FMR2 family, member 4 | 2228.54 | 2279.44 | -0.03 | 9.13E-01 |
| NULL | GSMG0005184 | NULL | NULL | 1883.11 | 2216.04 | -0.24 | 2.89E-01 |
| ENSMUSG00000018387 | GSMG0005186 | Shroom1 | shroom family member 1 | 2135.59 | 1944.44 | 0.14 | 5.72E-01 |
| NULL | GSMG0005187 | NULL | NULL | 327.61 | 91.36 | 1.84 | 1.30E-14 |
| ENSMUSG00000018398 | GSMG0005188 | Sept8 | septin 8 | 2163.24 | 1679.57 | 0.37 | 3.21E-02 |
| ENSMUSG00000018395 | GSMG0005189 | Kif3a | kinesin family member 3A | 1672.11 | 2091.68 | -0.32 | 1.08E-01 |
| ENSMUSG00000018899 | GSMG0005191 | Irf1 | interferon regulatory factor 1 | 473.80 | 337.68 | 0.49 | 8.69E-02 |
| ENSMUSG00000018906 | GSMG0005193 | P4ha2 | procollagen-proline, 2-oxoglutarate 4-dioxygenase (proline 4-hydroxyla | 393.19 | 216.71 | 0.86 | 9.56E-05 |
| ENSMUSG00000020333 | GSMG0005194 | Acsl6 | acyl-CoA synthetase long-chain family member 6 | 1545.75 | 943.53 | 0.71 | 1.41E-05 |
| ENSMUSG00000035992 | GSMG0005196 | Fnip1 | folliculin interacting protein 1 | 407.71 | 374.44 | 0.12 | 6.97E-01 |
| ENSMUSG00000037533 | GSMG0005198 | Rapgef6 | Rap guanine nucleotide exchange factor (GEF) 6 | 1432.17 | 1866.49 | -0.38 | 1.46E-02 |
| NULL | GSMG0005202 | NULL | NULL | 59.34 | 54.54 | 0.11 | 8.67E-01 |
| NULL | GSMG0005203 | NULL | NULL | 336.98 | 298.95 | 0.18 | 6.01E-01 |
| ENSMUSG00000020267 | GSMG0005208 | Hint1 | histidine triad nucleotide binding protein 1 | 2604.14 | 3963.20 | -0.61 | 1.31E-03 |
| ENSMUSG00000018339 | GSMG0005209 | Gpx3 | glutathione peroxidase 3 | 288.51 | 312.85 | -0.12 | 7.28E-01 |
| ENSMUSG00000000594 | GSMG0005211 | Gm2a | GM2 ganglioside activator protein | 4180.02 | 3420.42 | 0.29 | 9.84E-02 |
| ENSMUSG00000018583 | GSMG0005216 | G3bp1 | GTPase activating protein (SH3 domain) binding protein 1 | 2688.84 | 3426.30 | -0.35 | 4.70E-02 |
| NULL | GSMG0005217 | NULL | NULL | 411.96 | 429.01 | -0.06 | 8.69E-01 |
| ENSMUSG00000020524 | GSMG0005219 | Gria1 | glutamate receptor, ionotropic, AMPA1 (alpha 1) | 1396.20 | 1975.66 | -0.50 | 8.80E-03 |
| NULL | GSMG0005220 | NULL | NULL | 747.99 | 664.74 | 0.17 | 4.82E-01 |
| ENSMUSG00000020522 | GSMG0005222 | Mfap3 | microfibrillar-associated protein 3 | 2811.82 | 2997.25 | -0.09 | 7.18E-01 |
| ENSMUSG00000020520 | GSMG0005223 | Galnt10 | UDP-N-acetyl-alpha-D-galactosamine:polypeptide N-acetylgalactosami | 426.08 | 522.90 | -0.30 | 1.94E-01 |
| ENSMUSG00000020519 | GSMG0005227 | Sap30l | SAP30-like | 451.12 | 324.95 | 0.47 | 1.28E-01 |

| | | | | | | | |
|---|---|---|---|---|---|---|---|
| ENSMUSG00000037331 | GSMG0005230 | Larp1 | La ribonucleoprotein domain family, member 1 | 331.70 | 335.57 | -0.02 | 9.70E-01 |
| ENSMUSG00000020515 | GSMG0005231 | Cnot8 | CCR4-NOT transcription complex, subunit 8 | 889.53 | 1124.18 | -0.34 | 7.71E-02 |
| ENSMUSG00000020514 | GSMG0005232 | Mrpl22 | mitochondrial ribosomal protein L22 | 366.23 | 421.55 | -0.20 | 4.21E-01 |
| ENSMUSG00000078853 | GSMG0005234 | Igtp | interferon gamma induced GTPase | 507.96 | 716.07 | -0.50 | 5.64E-01 |
| ENSMUSG00000069874 | GSMG0005235 | Irgm2 | immunity-related GTPase family M member 2 | 642.89 | 650.68 | -0.02 | 9.74E-01 |
| ENSMUSG00000037243 | GSMG0005238 | Zfp692 | zinc finger protein 692 | 1902.43 | 1980.83 | -0.06 | 8.47E-01 |
| ENSMUSG00000013646 | GSMG0005240 | Sh3bp5l | SH3 binding domain protein 5 like | 729.75 | 637.11 | 0.20 | 4.26E-01 |
| NULL | GSMG0005252 | NULL | NULL | 871.67 | 784.86 | 0.15 | 5.57E-01 |
| ENSMUSG00000056895 | GSMG0005255 | Hist3h2ba | histone cluster 3, H2ba | 422.07 | 553.07 | -0.39 | 2.82E-01 |
| ENSMUSG00000036964 | GSMG0005256 | Trim17 | tripartite motif-containing 17 | 769.76 | 664.53 | 0.21 | 3.48E-01 |
| NULL | GSMG0005257 | NULL | NULL | 103.37 | 121.97 | -0.23 | 6.23E-01 |
| ENSMUSG00000020455 | GSMG0005258 | Trim11 | tripartite motif-containing 11 | 1042.64 | 896.81 | 0.22 | 3.40E-01 |
| ENSMUSG00000036860 | GSMG0005260 | Mrpl55 | mitochondrial ribosomal protein L55 | 649.22 | 749.61 | -0.21 | 3.80E-01 |
| ENSMUSG00000020441 | GSMG0005261 | 2310033P09Rik | RIKEN cDNA 2310033P09 gene | 858.15 | 816.23 | 0.07 | 8.20E-01 |
| ENSMUSG00000036819 | GSMG0005263 | Jmjd4 | jumonji domain containing 4 | 107.86 | 136.83 | -0.34 | 4.00E-01 |
| ENSMUSG00000072893 | GSMG0005265 | 4933439C10Rik | RIKEN cDNA 4933439C10 gene | 4150.53 | 2439.22 | 0.77 | 6.92E-06 |
| ENSMUSG00000005417 | GSMG0005268 | Mprip | myosin phosphatase Rho interacting protein | 15671.97 | 10907.87 | 0.52 | 6.82E-02 |
| ENSMUSG00000087249 | GSMG0005274 | Gm16062 | predicted gene 16062 | 95.68 | 115.33 | -0.27 | 5.39E-01 |
| ENSMUSG00000032615 | GSMG0005275 | Nt5m | 5',3'-nucleotidase, mitochondrial | 812.85 | 822.01 | -0.01 | 9.62E-01 |
| ENSMUSG00000061650 | GSMG0005278 | Med9 | mediator complex subunit 9 | 1212.25 | 1148.82 | 0.08 | 7.71E-01 |
| ENSMUSG00000062115 | GSMG0005281 | Rai1 | retinoic acid induced 1 | 1193.31 | 1617.43 | -0.44 | 1.38E-01 |
| NULL | GSMG0005283 | NULL | NULL | 330.33 | 213.05 | 0.63 | 4.58E-03 |
| NULL | GSMG0005284 | NULL | NULL | 166.31 | 110.52 | 0.59 | 1.37E-01 |
| ENSMUSG00000056598 | GSMG0005286 | Lrrc48 | leucine rich repeat containing 48 | 96.60 | 62.39 | 0.63 | 2.30E-01 |
| ENSMUSG00000018415 | GSMG0005288 | Gid4 | GID complex subunit 4, VID24 homolog | 354.36 | 399.91 | -0.18 | 5.14E-01 |
| ENSMUSG00000020537 | GSMG0005289 | Drg2 | developmentally regulated GTP binding protein 2 | 1000.98 | 1091.05 | -0.12 | 6.28E-01 |
| ENSMUSG00000042650 | GSMG0005291 | Alkbh5 | alkB homolog 5, RNA demethylase | 1745.61 | 1663.01 | 0.07 | 7.99E-01 |
| ENSMUSG00000020536 | GSMG0005293 | Llgl1 | lethal giant larvae homolog 1 (Drosophila) | 2090.24 | 1799.05 | 0.22 | 3.49E-01 |
| ENSMUSG00000018599 | GSMG0005294 | Mief2 | mitochondrial elongation factor 2 | 559.94 | 559.42 | 0.00 | 9.97E-01 |
| ENSMUSG00000042569 | GSMG0005297 | Dhrs7b | dehydrogenase/reductase (SDR family) member 7B | 653.96 | 685.22 | -0.07 | 8.37E-01 |
| ENSMUSG00000018932 | GSMG0005300 | Map2k3 | mitogen-activated protein kinase kinase 3 | 1154.36 | 1140.35 | 0.02 | 9.55E-01 |
| ENSMUSG00000042436 | GSMG0005306 | Mfap4 | microfibrillar-associated protein 4 | 273.01 | 153.36 | 0.83 | 6.25E-04 |
| ENSMUSG00000001039 | GSMG0005307 | B9d1 | B9 protein domain 1 | 346.24 | 355.06 | -0.03 | 9.24E-01 |

| | | | | | | | |
|---|---|---|---|---|---|---|---|
| NULL | GSMG0005308 | NULL | NULL | 423.05 | 303.31 | 0.48 | 2.35E-02 |
| ENSMUSG00000004837 | GSMG0005310 | Grap | GRB2-related adaptor protein | 7222.24 | 6207.26 | 0.22 | 2.52E-01 |
| ENSMUSG00000042331 | GSMG0005313 | Specc1 | sperm antigen with calponin homology and coiled-coil domains 1 | 584.26 | 422.21 | 0.47 | 1.20E-02 |
| ENSMUSG00000018500 | GSMG0005316 | Adora2b | adenosine A2b receptor | 334.46 | 271.87 | 0.30 | 3.03E-01 |
| ENSMUSG00000042298 | GSMG0005318 | Ttc19 | tetratricopeptide repeat domain 19 | 413.54 | 316.13 | 0.38 | 1.08E-01 |
| NULL | GSMG0005323 | NULL | NULL | 35.47 | 71.58 | -1.04 | 3.37E-02 |
| ENSMUSG00000018507 | GSMG0005325 | Trpv2 | transient receptor potential cation channel, subfamily V, member 2 | 343.89 | 369.79 | -0.10 | 7.88E-01 |
| ENSMUSG00000086841 | GSMG0005326 | 2410006H16Rik | RIKEN cDNA 2410006H16 gene | 198.31 | 250.64 | -0.34 | 2.71E-01 |
| ENSMUSG00000048497 | GSMG0005329 | Mmgt2 | membrane magnesium transporter 2 | 112.34 | 108.53 | 0.05 | 9.18E-01 |
| G00000047821,ENSMUSG000 | GSMG0005331 | Trim16 // Fbxw10 | tripartite motif-containing 16 // F-box and WD-40 domain protein 10 | 472.61 | 482.69 | -0.03 | 9.31E-01 |
| ENSMUSG00000014177 | GSMG0005332 | Tvp23b | trans-golgi network vesicle protein 23B | 589.52 | 431.32 | 0.45 | 3.58E-02 |
| ENSMUSG00000018217 | GSMG0005335 | Pmp22 | peripheral myelin protein 22 | 3775.56 | 2223.49 | 0.76 | 1.10E-06 |
| ENSMUSG00000087497 | GSMG0005340 | 2810001G20Rik | RIKEN cDNA 2810001G20 gene | 322.88 | 322.96 | 0.00 | 9.96E-01 |
| ENSMUSG00000020549 | GSMG0005345 | Elac2 | elaC ribonuclease Z 2 | 1119.83 | 1255.69 | -0.16 | 4.98E-01 |
| ENSMUSG00000018347 | GSMG0005347 | Zkscan6 | zinc finger with KRAB and SCAN domains 6 | 590.56 | 666.97 | -0.18 | 4.43E-01 |
| ENSMUSG00000069844 | GSMG0005355 | Sco1 | SCO1 cytochrome c oxidase assembly protein | 423.00 | 514.83 | -0.29 | 2.25E-01 |
| ENSMUSG00000033066 | GSMG0005364 | Gas7 | growth arrest specific 7 | 182.52 | 181.63 | 0.01 | 9.84E-01 |
| ENSMUSG00000020903 | GSMG0005371 | Stx8 | syntaxin 8 | 901.21 | 973.44 | -0.11 | 6.59E-01 |
| ENSMUSG00000046207 | GSMG0005376 | Pik3r6 | phosphoinositide-3-kinase, regulatory subunit 6 | 2116.23 | 2134.10 | -0.01 | 9.70E-01 |
| ENSMUSG00000020900 | GSMG0005379 | Myh10 | myosin, heavy polypeptide 10, non-muscle | 8073.62 | 17715.89 | -1.13 | 1.31E-11 |
| NULL | GSMG0005385 | NULL | NULL | 253.48 | 218.61 | 0.21 | 5.98E-01 |
| ENSMUSG00000018740 | GSMG0005386 | Slc25a35 | solute carrier family 25, member 35 | 175.38 | 90.04 | 0.96 | 1.25E-03 |
| ENSMUSG00000020898 | GSMG0005389 | Ctc1 | CTS telomere maintenance complex component 1 | 1200.58 | 1731.46 | -0.53 | 1.08E-02 |
| ENSMUSG00000020897 | GSMG0005390 | Aurkb | aurora kinase B | 271.42 | 723.35 | -1.41 | 1.64E-13 |
| ENSMUSG00000045176 | GSMG0005391 | Borcs6 | BLOC-1 related complex subunit 6 | 210.16 | 180.99 | 0.21 | 5.71E-01 |
| ENSMUSG00000020895 | GSMG0005392 | Tmem107 | transmembrane protein 107 | 251.74 | 260.72 | -0.05 | 9.10E-01 |
| ENSMUSG00000020894 | GSMG0005394 | Vamp2 | vesicle-associated membrane protein 2 | 6175.24 | 4734.14 | 0.38 | 1.86E-02 |
| ENSMUSG00000020893 | GSMG0005395 | Per1 | period circadian clock 1 | 447.07 | 175.65 | 1.35 | 9.10E-07 |
| ENSMUSG00000049299 | GSMG0005401 | Trappc1 | trafficking protein particle complex 1 | 1377.48 | 1531.05 | -0.15 | 5.04E-01 |
| ENSMUSG00000018470 | GSMG0005402 | Kcnab3 | potassium voltage-gated channel, shaker-related subfamily, beta membe | 276.75 | 173.94 | 0.67 | 1.09E-02 |
| ENSMUSG00000043419 | GSMG0005403 | Chd3os | chromodomain helicase DNA binding protein 3, opposite strand | 264.97 | 254.54 | 0.06 | 8.93E-01 |
| ENSMUSG00000059278 | GSMG0005404 | Naa38 | N(alpha)-acetyltransferase 38, NatC auxiliary subunit | 768.40 | 890.58 | -0.21 | 3.99E-01 |
| ENSMUSG00000059552 | GSMG0005408 | Trp53 | transformation related protein 53 | 996.78 | 1397.16 | -0.49 | 4.80E-02 |

| | | | | | | | |
|---|---|---|---|---|---|---|---|
| ENSMUSG00000069835 | GSMG0005410 | Sat2 | spermidine/spermine N1-acetyl transferase 2 | 317.17 | 293.98 | 0.11 | 7.27E-01 |
| ENSMUSG00000018765 | GSMG0005411 | Fxr2 | fragile X mental retardation, autosomal homolog 2 | 2027.59 | 1412.04 | 0.52 | 7.45E-02 |
| NULL | GSMG0005413 | NULL | NULL | 378.22 | 390.56 | -0.05 | 8.97E-01 |
| NULL | GSMG0005419 | NULL | NULL | 140.17 | 142.98 | -0.03 | 9.50E-01 |
| ENSMUSG00000070394 | GSMG0005420 | Tmem256 | transmembrane protein 256 | 1346.90 | 1406.15 | -0.06 | 8.70E-01 |
| ENSMUSG00000019461 | GSMG0005421 | Plscr3 | phospholipid scramblase 3 | 1850.35 | 2379.50 | -0.36 | 4.82E-02 |
| ENSMUSG00000047284 | GSMG0005423 | Neurl4 | neuralized E3 ubiquitin protein ligase 4 | 421.67 | 504.48 | -0.26 | 3.98E-01 |
| ENSMUSG00000023170 | GSMG0005424 | Gps2 | G protein pathway suppressor 2 | 1086.66 | 1069.17 | 0.02 | 9.37E-01 |
| ENSMUSG00000018559 | GSMG0005427 | Ctdnep1 | CTD nuclear envelope phosphatase 1 | 418.11 | 327.74 | 0.35 | 3.22E-01 |
| ENSMUSG00000018567 | GSMG0005428 | Gabarap | gamma-aminobutyric acid receptor associated protein | 3553.87 | 3697.53 | -0.06 | 8.19E-01 |
| ENSMUSG00000018572 | GSMG0005429 | Phf23 | PHD finger protein 23 | 708.33 | 767.77 | -0.12 | 6.51E-01 |
| ENSMUSG00000020888 | GSMG0005430 | Dvl2 | dishevelled segment polarity protein 2 | 401.78 | 386.49 | 0.05 | 8.86E-01 |
| ENSMUSG00000020886 | GSMG0005431 | Dlg4 | discs, large homolog 4 (Drosophila) | 758.15 | 601.41 | 0.33 | 1.37E-01 |
| ENSMUSG00000040938 | GSMG0005436 | Slc16a11 | solute carrier family 16 (monocarboxylic acid transporters), member 11 | 721.78 | 552.16 | 0.39 | 1.35E-01 |
| ENSMUSG00000060216 | GSMG0005437 | Arrb2 | arrestin, beta 2 | 406.62 | 441.52 | -0.12 | 6.69E-01 |
| ENSMUSG00000018923 | GSMG0005438 | Med11 | mediator complex subunit 11 | 157.52 | 186.12 | -0.24 | 4.92E-01 |
| ENSMUSG00000018286 | GSMG0005442 | Psmb6 | proteasome (prosome, macropain) subunit, beta type 6 | 2580.18 | 3346.64 | -0.38 | 1.74E-02 |
| ENSMUSG00000020828 | GSMG0005443 | Pld2 | phospholipase D2 | 1167.73 | 1317.29 | -0.17 | 4.81E-01 |
| ENSMUSG00000020827 | GSMG0005444 | Mink1 | misshapen-like kinase 1 (zebrafish) | 1751.11 | 1424.77 | 0.30 | 3.77E-01 |
| ENSMUSG00000040746 | GSMG0005448 | Rnf167 | ring finger protein 167 | 1442.28 | 1283.32 | 0.17 | 4.41E-01 |
| NULL | GSMG0005449 | NULL | NULL | 477.37 | 719.18 | -0.59 | 9.25E-03 |
| ENSMUSG00000060600 | GSMG0005451 | Eno3 | enolase 3, beta muscle | 246.45 | 197.85 | 0.32 | 2.92E-01 |
| ENSMUSG00000043602 | GSMG0005455 | Zfp3 | zinc finger protein 3 | 344.23 | 435.51 | -0.34 | 1.66E-01 |
| ENSMUSG00000020817 | GSMG0005456 | Rabep1 | rabaptin, RAB GTPase binding effector protein 1 | 1533.15 | 1647.21 | -0.10 | 6.70E-01 |
| ENSMUSG00000018449 | GSMG0005460 | Rpain | RPA interacting protein | 1123.32 | 1039.41 | 0.11 | 6.22E-01 |
| NULL | GSMG0005461 | NULL | NULL | 251.97 | 116.28 | 1.11 | 2.65E-06 |
| ENSMUSG00000040599 | GSMG0005462 | Mis12 | MIS12 kinetochore complex component | 932.42 | 987.92 | -0.08 | 8.03E-01 |
| ENSMUSG00000020811 | GSMG0005464 | Wscd1 | WSC domain containing 1 | 1458.08 | 1986.27 | -0.45 | 3.39E-02 |
| ENSMUSG00000020808 | GSMG0005465 | Fam64a | family with sequence similarity 64, member A | 207.17 | 509.79 | -1.29 | 2.81E-07 |
| ENSMUSG00000020803 | GSMG0005466 | Txndc17 | thioredoxin domain containing 17 | 1583.00 | 1627.94 | -0.04 | 8.91E-01 |
| ENSMUSG00000040483 | GSMG0005469 | Xaf1 | XIAP associated factor 1 | 284.47 | 228.95 | 0.31 | 3.10E-01 |
| ENSMUSG00000040463 | GSMG0005471 | Mybbp1a | MYB binding protein (P160) 1a | 2980.90 | 4195.04 | -0.49 | 2.44E-03 |
| ENSMUSG00000020794 | GSMG0005472 | Ube2g1 | ubiquitin-conjugating enzyme E2G 1 | 3676.10 | 4341.68 | -0.24 | 2.96E-01 |

| | | | | | | | |
|---|---|---|---|---|---|---|---|
| ENSMUSG00000020790 | GSMG0005478 | Ankfy1 | ankyrin repeat and FYVE domain containing 1 | 296.58 | 398.32 | -0.42 | 8.25E-02 |
| ENSMUSG00000020788 | GSMG0005483 | Atp2a3 | ATPase, Ca++ transporting, ubiquitous | 6123.12 | 2913.77 | 1.07 | 5.54E-11 |
| ENSMUSG00000020787 | GSMG0005484 | P2rx1 | purinergic receptor P2X, ligand-gated ion channel, 1 | 384.48 | 186.33 | 1.04 | 2.57E-07 |
| ENSMUSG00000020785 | GSMG0005485 | Camkk1 | calcium/calmodulin-dependent protein kinase kinase 1, alpha | 1740.01 | 1289.24 | 0.43 | 2.13E-02 |
| ENSMUSG00000020783 | GSMG0005486 | Ncbp3 | nuclear cap binding subunit 3 | 646.69 | 757.62 | -0.23 | 3.16E-01 |
| ENSMUSG00000005951 | GSMG0005491 | Shpk | sedoheptulokinase | 98.39 | 58.67 | 0.74 | 4.46E-02 |
| ENSMUSG00000020741 | GSMG0005506 | Cluh | clustered mitochondria (cluA/CLU1) homolog | 998.94 | 1014.49 | -0.02 | 9.61E-01 |
| ENSMUSG00000010554 | GSMG0005507 | Mettl16 | methyltransferase like 16 | 313.20 | 322.35 | -0.04 | 9.20E-01 |
| ENSMUSG00000038335 | GSMG0005511 | Tsr1 | TSR1 20S rRNA accumulation | 305.52 | 533.50 | -0.80 | 3.03E-05 |
| ENSMUSG00000038290 | GSMG0005512 | Smg6 | Smg-6 homolog, nonsense mediated mRNA decay factor (C. elegans) | 731.87 | 741.46 | -0.02 | 9.58E-01 |
| NULL | GSMG0005517 | NULL | NULL | 137.70 | 52.47 | 1.39 | 3.40E-06 |
| ENSMUSG00000045287 | GSMG0005519 | Rtn4rl1 | reticulon 4 receptor-like 1 | 1848.71 | 1817.94 | 0.02 | 9.35E-01 |
| NULL | GSMG0005520 | NULL | NULL | 322.67 | 180.48 | 0.84 | 2.42E-04 |
| NULL | GSMG0005521 | NULL | NULL | 91.37 | 127.11 | -0.47 | 1.94E-01 |
| NULL | GSMG0005522 | NULL | NULL | 43.27 | 61.46 | -0.49 | 3.37E-01 |
| ENSMUSG00000018809 | GSMG0005523 | Smyd4 | SET and MYND domain containing 4 | 358.96 | 430.23 | -0.26 | 2.95E-01 |
| ENSMUSG00000085148 | GSMG0005525 | Mir22hg | Mir22 host gene (non-protein coding) | 638.75 | 409.54 | 0.64 | 7.06E-04 |
| ENSMUSG00000020850 | GSMG0005527 | Prpf8 | pre-mRNA processing factor 8 | 1427.71 | 1626.57 | -0.19 | 3.67E-01 |
| ENSMUSG00000038195 | GSMG0005528 | Rilp | Rab interacting lysosomal protein | 127.50 | 99.30 | 0.36 | 3.36E-01 |
| ENSMUSG00000038188 | GSMG0005529 | Scarf1 | scavenger receptor class F, member 1 | 2306.93 | 2361.98 | -0.03 | 9.07E-01 |
| ENSMUSG00000038178 | GSMG0005530 | Slc43a2 | solute carrier family 43, member 2 | 1121.59 | 1759.88 | -0.65 | 2.33E-03 |
| ENSMUSG00000017781 | GSMG0005531 | Pitpna | phosphatidylinositol transfer protein, alpha | 4129.74 | 4493.51 | -0.12 | 5.87E-01 |
| ENSMUSG00000006127 | GSMG0005533 | Inpp5k | inositol polyphosphate 5-phosphatase K | 2454.76 | 2123.99 | 0.21 | 2.89E-01 |
| ENSMUSG00000017774 | GSMG0005534 | Myo1c | myosin IC | 4020.58 | 3982.66 | 0.01 | 9.62E-01 |
| ENSMUSG00000017776 | GSMG0005535 | Crk | v-crk avian sarcoma virus CT10 oncogene homolog | 6993.28 | 6774.03 | 0.05 | 8.59E-01 |
| ENSMUSG00000020849 | GSMG0005539 | Ywhae | tyrosine 3-monooxygenase/tryptophan 5-monooxygenase activation pro | 3928.16 | 6019.79 | -0.62 | 6.21E-03 |
| ENSMUSG00000069808 | GSMG0005545 | Fam57a | family with sequence similarity 57, member A | 812.73 | 859.91 | -0.08 | 7.87E-01 |
| ENSMUSG00000038046 | GSMG0005547 | Rnmtl1 | RNA methyltransferase like 1 | 557.98 | 575.63 | -0.04 | 8.96E-01 |
| ENSMUSG00000020843 | GSMG0005548 | Timm22 | translocase of inner mitochondrial membrane 22 | 1692.70 | 1840.85 | -0.12 | 6.01E-01 |
| ENSMUSG00000020840 | GSMG0005552 | Blmh | bleomycin hydrolase | 1334.99 | 1608.93 | -0.27 | 1.52E-01 |
| ENSMUSG00000037926 | GSMG0005560 | Ssh2 | slingshot homolog 2 (Drosophila) | 724.00 | 810.14 | -0.16 | 5.68E-01 |
| ENSMUSG00000020836 | GSMG0005563 | Coro6 | coronin 6 | 349.52 | 26.00 | 3.75 | 4.81E-36 |
| NULL | GSMG0005564 | NULL | NULL | 58.82 | 93.60 | -0.68 | 1.59E-01 |

| | | | | | | | |
|---|---|---|---|---|---|---|---|
| ENSMUSG00000011877 | GSMG0005565 | Git1 | G protein-coupled receptor kinase-interactor 1 | 5258.99 | 3656.38 | 0.52 | 8.34E-02 |
| ENSMUSG00000037857 | GSMG0005568 | Nufip2 | nuclear fragile X mental retardation protein interacting protein 2 | 1555.99 | 1346.01 | 0.21 | 4.96E-01 |
| ENSMUSG00000000632 | GSMG0005572 | Sez6 | seizure related gene 6 | 2630.23 | 5358.70 | -1.03 | 1.27E-07 |
| ENSMUSG00000037791 | GSMG0005573 | Phf12 | PHD finger protein 12 | 1119.70 | 1175.65 | -0.07 | 8.48E-01 |
| ENSMUSG00000020834 | GSMG0005576 | Dhrs13 | dehydrogenase/reductase (SDR family) member 13 | 84.30 | 214.76 | -1.35 | 2.55E-07 |
| ENSMUSG00000061981 | GSMG0005577 | Flot2 | flotillin 2 | 859.66 | 1214.70 | -0.50 | 3.12E-03 |
| ENSMUSG00000037750 | GSMG0005581 | Fam222b | family with sequence similarity 222, member B | 1901.13 | 2050.29 | -0.11 | 6.20E-01 |
| NULL | GSMG0005582 | NULL | NULL | 478.30 | 351.89 | 0.44 | 6.13E-01 |
| ENSMUSG00000002059 | GSMG0005586 | Rab34 | RAB34, member RAS oncogene family | 1222.50 | 1269.41 | -0.05 | 8.70E-01 |
| ENSMUSG00000002064 | GSMG0005588 | Sdf2 | stromal cell derived factor 2 | 1539.81 | 1331.93 | 0.21 | 3.23E-01 |
| ENSMUSG00000010277 | GSMG0005589 | 2610507B11Rik | RIKEN cDNA 2610507B11 gene | 3070.15 | 3343.29 | -0.12 | 6.68E-01 |
| ENSMUSG00000002055 | GSMG0005591 | Spag5 | sperm associated antigen 5 | 264.16 | 373.61 | -0.49 | 1.10E-01 |
| ENSMUSG00000017390 | GSMG0005592 | Aldoc | aldolase C, fructose-bisphosphate | 3793.60 | 1961.45 | 0.95 | 2.10E-06 |
| ENSMUSG00000041958 | GSMG0005593 | Pigs | phosphatidylinositol glycan anchor biosynthesis, class S | 1043.37 | 1020.62 | 0.03 | 9.26E-01 |
| ENSMUSG00000002058 | GSMG0005594 | Unc119 | unc-119 lipid binding chaperone | 239.37 | 462.83 | -0.96 | 7.82E-07 |
| ENSMUSG00000001105 | GSMG0005595 | Ift20 | intraflagellar transport 20 | 1478.20 | 1262.44 | 0.23 | 3.22E-01 |
| ENSMUSG00000020829 | GSMG0005597 | Slc46a1 | solute carrier family 46, member 1 | 141.97 | 187.10 | -0.39 | 2.12E-01 |
| ENSMUSG00000017344 | GSMG0005598 | Vtn | vitronectin | 31497.28 | 17134.47 | 0.88 | 6.88E-06 |
| ENSMUSG00000001100 | GSMG0005600 | Poldip2 | polymerase (DNA-directed), delta interacting protein 2 | 3263.96 | 3485.92 | -0.09 | 6.84E-01 |
| NULL | GSMG0005602 | NULL | NULL | 88.14 | 65.14 | 0.44 | 2.92E-01 |
| ENSMUSG00000072640 | GSMG0005603 | Lyrm9 | LYR motif containing 9 | 997.88 | 928.60 | 0.10 | 6.98E-01 |
| ENSMUSG00000020826 | GSMG0005604 | Nos2 | nitric oxide synthase 2, inducible | 1421.73 | 1517.03 | -0.09 | 7.12E-01 |
| ENSMUSG00000017639 | GSMG0005613 | Rab11fip4 | RAB11 family interacting protein 4 (class II) | 248.48 | 319.64 | -0.36 | 2.00E-01 |
| ENSMUSG00000017548 | GSMG0005616 | Suz12 | suppressor of zeste 12 homolog (Drosophila) | 264.18 | 257.37 | 0.03 | 9.48E-01 |
| ENSMUSG00000017550 | GSMG0005617 | Atad5 | ATPase family, AAA domain containing 5 | 276.44 | 560.46 | -1.02 | 5.92E-08 |
| ENSMUSG00000020709 | GSMG0005618 | Adap2 | ArfGAP with dual PH domains 2 | 523.30 | 348.51 | 0.59 | 1.34E-02 |
| ENSMUSG00000020707 | GSMG0005619 | Rnf135 | ring finger protein 135 | 134.85 | 122.72 | 0.14 | 7.66E-01 |
| ENSMUSG00000017686 | GSMG0005620 | Rhot1 | ras homolog family member T1 | 943.53 | 883.50 | 0.10 | 7.26E-01 |
| ENSMUSG00000017692 | GSMG0005624 | Rhbdl3 | rhomboid, veinlet-like 3 (Drosophila) | 138.57 | 41.16 | 1.75 | 3.58E-08 |
| ENSMUSG00000017421 | GSMG0005627 | Zfp207 | zinc finger protein 207 | 3803.90 | 3183.02 | 0.26 | 2.11E-01 |
| ENSMUSG00000017428 | GSMG0005629 | Psmd11 | proteasome (prosome, macropain) 26S subunit, non-ATPase, 11 | 832.23 | 1020.12 | -0.29 | 2.44E-01 |
| NULL | GSMG0005630 | NULL | NULL | 138.27 | 240.75 | -0.81 | 3.24E-03 |
| NULL | GSMG0005631 | NULL | NULL | 955.33 | 1481.13 | -0.64 | 1.44E-02 |

| | | | | | | | |
|---|---|---|---|---|---|---|---|
| ENSMUSG00000035413 | GSMG0005632 | Tmem98 | transmembrane protein 98 | 3123.07 | 2495.39 | 0.32 | 9.92E-02 |
| NULL | GSMG0005635 | NULL | NULL | 354.67 | 224.53 | 0.66 | 6.42E-03 |
| ENSMUSG00000035352 | GSMG0005642 | Ccl12 | chemokine (C-C motif) ligand 12 | 134.41 | 23.66 | 2.52 | 6.98E-09 |
| ENSMUSG00000046010 | GSMG0005647 | Zfp830 | zinc finger protein 830 | 727.08 | 727.52 | 0.00 | 9.98E-01 |
| ENSMUSG00000020697 | GSMG0005648 | Lig3 | ligase III, DNA, ATP-dependent | 676.57 | 833.88 | -0.30 | 2.10E-01 |
| ENSMUSG00000018845 | GSMG0005650 | Unc45b | unc-45 myosin chaperone B | 977.20 | 1125.71 | -0.20 | 4.06E-01 |
| ENSMUSG00000054404 | GSMG0005651 | Slfn5 | schlafen 5 | 638.66 | 425.02 | 0.59 | 9.71E-03 |
| ENSMUSG00000035152 | GSMG0005660 | Ap2b1 | adaptor-related protein complex 2, beta 1 subunit | 2327.06 | 2744.59 | -0.24 | 2.32E-01 |
| NULL | GSMG0005661 | NULL | NULL | 137.98 | 152.04 | -0.14 | 7.21E-01 |
| ENSMUSG00000020680 | GSMG0005665 | Taf15 | TATA-box binding protein associated factor 15 | 91.59 | 83.75 | 0.14 | 8.13E-01 |
| ENSMUSG00000000976 | GSMG0005672 | Heatr6 | HEAT repeat containing 6 | 678.79 | 871.12 | -0.36 | 8.30E-02 |
| ENSMUSG00000020677 | GSMG0005674 | Ddx52 | DEAD (Asp-Glu-Ala-Asp) box polypeptide 52 | 757.51 | 909.99 | -0.26 | 2.36E-01 |
| ENSMUSG00000034940 | GSMG0005675 | Synrg | synergin, gamma | 159.25 | 182.94 | -0.20 | 5.95E-01 |
| NULL | GSMG0005677 | NULL | NULL | 115.49 | 248.27 | -1.11 | 9.28E-07 |
| ENSMUSG00000020532 | GSMG0005679 | Acaca | acetyl-Coenzyme A carboxylase alpha | 924.17 | 1636.55 | -0.82 | 1.37E-06 |
| ENSMUSG00000020527 | GSMG0005687 | Myo19 | myosin XIX | 246.51 | 266.55 | -0.11 | 7.68E-01 |
| ENSMUSG00000000805 | GSMG0005688 | Car4 | carbonic anhydrase 4 | 16696.25 | 10611.69 | 0.65 | 8.35E-05 |
| ENSMUSG00000020525 | GSMG0005692 | Ppm1d | protein phosphatase 1D magnesium-dependent, delta isoform | 242.61 | 269.85 | -0.15 | 7.15E-01 |
| ENSMUSG00000059439 | GSMG0005694 | Bcas3 | breast carcinoma amplified sequence 3 | 591.64 | 567.62 | 0.06 | 8.54E-01 |
| ENSMUSG00000000093 | GSMG0005699 | Tbx2 | T-box 2 | 1442.40 | 850.76 | 0.76 | 4.72E-03 |
| ENSMUSG00000000094 | GSMG0005701 | Tbx4 | T-box 4 | 118.27 | 320.69 | -1.43 | 2.43E-08 |
| ENSMUSG00000085208 | GSMG0005702 | Brip1os | BRCA1 interacting protein C-terminal helicase 1, opposite strand | 220.92 | 278.96 | -0.33 | 2.23E-01 |
| ENSMUSG00000020521 | GSMG0005706 | Rnft1 | ring finger protein, transmembrane 1 | 868.34 | 676.37 | 0.36 | 9.86E-02 |
| ENSMUSG00000020513 | GSMG0005708 | Tubd1 | tubulin, delta 1 | 224.11 | 211.86 | 0.09 | 8.43E-01 |
| ENSMUSG00000072582 | GSMG0005711 | Ptrh2 | peptidyl-tRNA hydrolase 2 | 527.18 | 723.48 | -0.46 | 2.59E-02 |
| ENSMUSG00000020492 | GSMG0005714 | Ska2 | spindle and kinetochore associated complex subunit 2 | 155.47 | 205.05 | -0.40 | 1.77E-01 |
| ENSMUSG00000018548 | GSMG0005716 | Trim37 | tripartite motif-containing 37 | 387.23 | 493.81 | -0.35 | 1.07E-01 |
| ENSMUSG00000020486 | GSMG0005721 | Sept4 | septin 4 | 9479.19 | 8596.94 | 0.14 | 4.81E-01 |
| ENSMUSG00000018401 | GSMG0005722 | Mtmr4 | myotubularin related protein 4 | 1211.72 | 1484.83 | -0.29 | 2.03E-01 |
| ENSMUSG00000034121 | GSMG0005731 | Mks1 | Meckel syndrome, type 1 | 114.58 | 137.62 | -0.26 | 5.28E-01 |
| ENSMUSG00000018379 | GSMG0005732 | Srsf1 | serine/arginine-rich splicing factor 1 | 6113.64 | 6633.54 | -0.12 | 6.28E-01 |
| ENSMUSG00000018377 | GSMG0005735 | Vezf1 | vascular endothelial zinc finger 1 | 569.49 | 540.54 | 0.08 | 8.17E-01 |
| ENSMUSG00000018378 | GSMG0005737 | Cuedc1 | CUE domain containing 1 | 681.69 | 798.34 | -0.23 | 2.81E-01 |

| | | | | | | | |
|---|---|---|---|---|---|---|---|
| ENSMUSG00000023723 | GSMG0005743 | Mrps23 | mitochondrial ribosomal protein S23 | 609.67 | 723.97 | -0.25 | 2.63E-01 |
| ENSMUSG00000033983 | GSMG0005753 | Coil | coilin | 387.72 | 505.41 | -0.38 | 7.23E-02 |
| ENSMUSG00000000275 | GSMG0005754 | Trim25 | tripartite motif-containing 25 | 407.99 | 420.46 | -0.04 | 9.21E-01 |
| ENSMUSG00000069763 | GSMG0005757 | Tmem100 | transmembrane protein 100 | 875.26 | 307.49 | 1.51 | 1.73E-11 |
| ENSMUSG00000003948 | GSMG0005758 | Mmd | monocyte to macrophage differentiation-associated | 2639.51 | 3347.64 | -0.34 | 8.02E-02 |
| NULL | GSMG0005759 | NULL | NULL | 305.10 | 260.59 | 0.23 | 4.66E-01 |
| ENSMUSG00000020544 | GSMG0005761 | Cox11 | cytochrome c oxidase assembly protein 11 | 409.04 | 451.40 | -0.14 | 6.24E-01 |
| ENSMUSG00000056158 | GSMG0005764 | Car10 | carbonic anhydrase 10 | 87.15 | 24.83 | 1.81 | 2.07E-06 |
| NULL | GSMG0005771 | NULL | NULL | 160.74 | 20.67 | 2.96 | 6.31E-04 |
| ENSMUSG00000059474 | GSMG0005773 | Mbtd1 | mbt domain containing 1 | 144.28 | 168.56 | -0.22 | 5.76E-01 |
| ENSMUSG00000020859 | GSMG0005778 | Spag9 | sperm associated antigen 9 | 3657.16 | 3879.66 | -0.09 | 8.10E-01 |
| ENSMUSG00000037573 | GSMG0005781 | Tob1 | transducer of ErbB-2.1 | 2508.19 | 1135.80 | 1.14 | 5.60E-13 |
| ENSMUSG00000020864 | GSMG0005783 | Ankrd40 | ankyrin repeat domain 40 | 843.05 | 544.03 | 0.63 | 7.76E-03 |
| NULL | GSMG0005784 | NULL | NULL | 224.02 | 159.55 | 0.50 | 1.40E-01 |
| ENSMUSG00000020869 | GSMG0005791 | Lrrc59 | leucine rich repeat containing 59 | 3872.08 | 5305.36 | -0.45 | 3.38E-03 |
| ENSMUSG00000024414 | GSMG0005792 | Mrpl27 | mitochondrial ribosomal protein L27 | 1057.42 | 1294.06 | -0.29 | 1.33E-01 |
| ENSMUSG00000001506 | GSMG0005795 | Col1a1 | collagen, type I, alpha 1 | 6534.33 | 7734.39 | -0.24 | 4.10E-01 |
| ENSMUSG00000038976 | GSMG0005797 | Ppp1r9b | protein phosphatase 1, regulatory subunit 9B | 308.52 | 332.71 | -0.11 | 7.97E-01 |
| ENSMUSG00000047181 | GSMG0005798 | Samd14 | sterile alpha motif domain containing 14 | 562.06 | 1635.43 | -1.54 | 6.67E-15 |
| ENSMUSG00000020873 | GSMG0005803 | Slc35b1 | solute carrier family 35, member B1 | 2307.32 | 2659.46 | -0.20 | 2.82E-01 |
| ENSMUSG00000057522 | GSMG0005804 | Spop | speckle-type POZ protein | 3318.13 | 2784.38 | 0.25 | 1.60E-01 |
| NULL | GSMG0005806 | NULL | NULL | 131.93 | 169.79 | -0.37 | 3.32E-01 |
| G00000050860,ENSMUSG000 | GSMG0005810 | Zfp652 // Phospho | zinc finger protein 652 // phosphatase, orphan 1 | 59.34 | 65.05 | -0.13 | 8.57E-01 |
| NULL | GSMG0005813 | NULL | NULL | 1069.06 | 856.27 | 0.32 | 2.94E-01 |
| ENSMUSG00000038811 | GSMG0005815 | Gngt2 | guanine nucleotide binding protein (G protein), gamma transducing acti | 112.41 | 81.83 | 0.46 | 2.56E-01 |
| ENSMUSG00000006058 | GSMG0005818 | Snf8 | SNF8, ESCRT-II complex subunit, homolog (S. cerevisiae) | 932.27 | 1073.58 | -0.20 | 3.09E-01 |
| ENSMUSG00000018666 | GSMG0005844 | Cbx1 | chromobox 1 | 998.80 | 1558.23 | -0.64 | 3.96E-04 |
| ENSMUSG00000018672 | GSMG0005848 | Copz2 | coatomer protein complex, subunit zeta 2 | 580.80 | 433.20 | 0.42 | 6.01E-02 |
| ENSMUSG00000078700 | GSMG0005849 | D030028A08Rik | RIKEN cDNA D030028A08 gene | 83.73 | 127.90 | -0.60 | 7.79E-02 |
| ENSMUSG00000020877 | GSMG0005853 | Scrn2 | secernin 2 | 159.52 | 189.09 | -0.25 | 4.63E-01 |
| ENSMUSG00000001445 | GSMG0005854 | Mrpl10 | mitochondrial ribosomal protein L10 | 323.74 | 449.03 | -0.47 | 4.16E-02 |
| ENSMUSG00000038534 | GSMG0005855 | Osbpl7 | oxysterol binding protein-like 7 | 228.21 | 229.10 | -0.01 | 9.86E-01 |
| ENSMUSG00000018882 | GSMG0005857 | Mrpl45 | mitochondrial ribosomal protein L45 | 344.87 | 432.52 | -0.32 | 1.76E-01 |

| | | | | | | | |
|---|---|---|---|---|---|---|---|
| ENSMUSG00000038485 | GSMG0005858 | Socs7 | suppressor of cytokine signaling 7 | 765.89 | 597.22 | 0.36 | 9.62E-02 |
| ENSMUSG00000049807 | GSMG0005860 | Arhgap23 | Rho GTPase activating protein 23 | 1822.71 | 1882.89 | -0.05 | 9.14E-01 |
| ENSMUSG00000078695 | GSMG0005864 | Cisd3 | CDGSH iron sulfur domain 3 | 171.08 | 178.82 | -0.06 | 8.88E-01 |
| ENSMUSG00000038366 | GSMG0005867 | Lasp1 | LIM and SH3 protein 1 | 640.74 | 590.73 | 0.12 | 7.19E-01 |
| ENSMUSG00000003119 | GSMG0005874 | Cdk12 | cyclin-dependent kinase 12 | 496.15 | 520.74 | -0.07 | 8.21E-01 |
| ENSMUSG00000061718 | GSMG0005877 | Ppp1r1b | protein phosphatase 1, regulatory (inhibitor) subunit 1B | 815.68 | 419.07 | 0.96 | 2.18E-01 |
| ENSMUSG00000018167 | GSMG0005878 | Stard3 | START domain containing 3 | 1195.51 | 1310.06 | -0.13 | 5.71E-01 |
| ENSMUSG00000062312 | GSMG0005881 | Erbb2 | erb-b2 receptor tyrosine kinase 2 | 361.55 | 372.55 | -0.04 | 9.14E-01 |
| ENSMUSG00000019312 | GSMG0005882 | Grb7 | growth factor receptor bound protein 7 | 701.58 | 507.04 | 0.47 | 7.99E-02 |
| ENSMUSG00000017221 | GSMG0005887 | Psmd3 | proteasome (prosome, macropain) 26S subunit, non-ATPase, 3 | 3601.42 | 3949.49 | -0.13 | 5.28E-01 |
| ENSMUSG00000058756 | GSMG0005889 | Thra | thyroid hormone receptor alpha | 3057.14 | 4949.15 | -0.70 | 5.32E-04 |
| G00000052915,ENSMUSG000 | GSMG0005890 | Msl1 // Casc3 | male-specific lethal 1 homolog (Drosophila) // cancer susceptibility can | 387.47 | 369.47 | 0.06 | 8.91E-01 |
| ENSMUSG00000038020 | GSMG0005891 | Rapgefl1 | Rap guanine nucleotide exchange factor (GEF)-like 1 | 531.93 | 493.07 | 0.11 | 7.94E-01 |
| ENSMUSG00000017499 | GSMG0005895 | Cdc6 | cell division cycle 6 | 98.86 | 303.86 | -1.61 | 2.58E-10 |
| ENSMUSG00000037992 | GSMG0005896 | Rara | retinoic acid receptor, alpha | 277.55 | 346.75 | -0.33 | 2.70E-01 |
| ENSMUSG00000017493 | GSMG0005899 | Igfbp4 | insulin-like growth factor binding protein 4 | 6667.05 | 12014.02 | -0.85 | 1.12E-09 |
| NULL | GSMG0005902 | NULL | NULL | 480.79 | 103.16 | 2.22 | 9.64E-02 |
| ENSMUSG00000001555 | GSMG0005914 | Fkbp10 | FK506 binding protein 10 | 2098.27 | 2865.91 | -0.45 | 2.45E-02 |
| ENSMUSG00000006782 | GSMG0005919 | Cnp | 2',3'-cyclic nucleotide 3' phosphodiesterase | 1279.45 | 656.26 | 0.96 | 2.45E-08 |
| ENSMUSG00000017837 | GSMG0005920 | Nkiras2 | NFKB inhibitor interacting Ras-like protein 2 | 189.86 | 211.16 | -0.15 | 6.62E-01 |
| NULL | GSMG0005924 | NULL | NULL | 2622.96 | 2216.49 | 0.24 | 3.30E-01 |
| ENSMUSG00000019302 | GSMG0005925 | Atp6v0a1 | ATPase, H+ transporting, lysosomal V0 subunit A1 | 1214.44 | 1403.85 | -0.21 | 2.86E-01 |
| NULL | GSMG0005926 | NULL | NULL | 385.43 | 259.37 | 0.57 | 1.29E-02 |
| NULL | GSMG0005927 | NULL | NULL | 929.61 | 611.12 | 0.60 | 9.86E-04 |
| ENSMUSG00000001751 | GSMG0005928 | Naglu | alpha-N-acetylglucosaminidase (Sanfilippo disease IIIB) | 274.36 | 227.96 | 0.27 | 4.21E-01 |
| ENSMUSG00000017801 | GSMG0005931 | Mlx | MAX-like protein X | 407.08 | 343.43 | 0.25 | 3.17E-01 |
| ENSMUSG00000035198 | GSMG0005932 | Tubg1 | tubulin, gamma 1 | 841.74 | 1114.79 | -0.40 | 4.18E-02 |
| ENSMUSG00000045007 | GSMG0005933 | Tubg2 | tubulin, gamma 2 | 348.32 | 390.30 | -0.16 | 5.83E-01 |
| ENSMUSG00000017167 | GSMG0005934 | Cntnap1 | contactin associated protein-like 1 | 414.23 | 90.96 | 2.19 | 3.40E-15 |
| ENSMUSG00000035112 | GSMG0005936 | Wnk4 | WNK lysine deficient protein kinase 4 | 145.54 | 195.65 | -0.42 | 2.11E-01 |
| ENSMUSG00000078653 | GSMG0005937 | Cntd1 | cyclin N-terminal domain containing 1 | 610.87 | 609.69 | 0.00 | 9.93E-01 |
| ENSMUSG00000078652 | GSMG0005938 | Psme3 | proteaseome (prosome, macropain) activator subunit 3 (PA28 gamma, K | 822.19 | 1016.31 | -0.31 | 2.09E-01 |
| ENSMUSG00000019326 | GSMG0005940 | Aoc3 | amine oxidase, copper containing 3 | 124.58 | 49.34 | 1.34 | 9.02E-04 |

| | | | | | | | |
|---|---|---|---|---|---|---|---|
| ENSMUSG00000035007 | GSMG0005943 | Rundc1 | RUN domain containing 1 | 199.94 | 171.35 | 0.22 | 5.06E-01 |
| ENSMUSG00000010358 | GSMG0005945 | Ifi35 | interferon-induced protein 35 | 95.12 | 76.51 | 0.31 | 4.67E-01 |
| ENSMUSG00000001313 | GSMG0005946 | Rnd2 | Rho family GTPase 2 | 1003.92 | 710.35 | 0.50 | 6.11E-03 |
| ENSMUSG00000017119 | GSMG0005947 | Nbr1 | neighbor of Brca1 gene 1 | 3842.56 | 3745.00 | 0.04 | 8.87E-01 |
| ENSMUSG00000010362 | GSMG0005949 | Rdm1 | RAD52 motif 1 | 442.26 | 364.18 | 0.28 | 3.84E-01 |
| ENSMUSG00000034936 | GSMG0005950 | Arl4d | ADP-ribosylation factor-like 4D | 560.85 | 333.55 | 0.75 | 1.40E-04 |
| ENSMUSG00000034931 | GSMG0005951 | Dhx8 | DEAH (Asp-Glu-Ala-His) box polypeptide 8 | 691.32 | 783.47 | -0.18 | 4.57E-01 |
| ENSMUSG00000034793 | GSMG0005957 | G6pc3 | glucose 6 phosphatase, catalytic, 3 | 687.09 | 704.49 | -0.04 | 9.19E-01 |
| ENSMUSG00000034757 | GSMG0005960 | Tmub2 | transmembrane and ubiquitin-like domain containing 2 | 802.88 | 779.00 | 0.05 | 8.87E-01 |
| ENSMUSG00000006575 | GSMG0005961 | Rundc3a | RUN domain containing 3A | 3012.69 | 2359.30 | 0.35 | 1.37E-01 |
| ENSMUSG00000034708 | GSMG0005962 | Grn | granulin | 4756.38 | 3781.35 | 0.33 | 9.87E-02 |
| NULL | GSMG0005964 | NULL | NULL | 607.48 | 656.11 | -0.11 | 6.75E-01 |
| ENSMUSG00000020926 | GSMG0005968 | Adam11 | a disintegrin and metallopeptidase domain 11 | 1813.31 | 2165.65 | -0.26 | 3.42E-01 |
| ENSMUSG00000020928 | GSMG0005969 | Higd1b | HIG1 domain family, member 1B | 2565.47 | 2260.01 | 0.18 | 3.83E-01 |
| NULL | GSMG0005970 | NULL | NULL | 147.23 | 141.12 | 0.06 | 8.95E-01 |
| ENSMUSG00000020936 | GSMG0005974 | Nmt1 | N-myristoyltransferase 1 | 1395.34 | 1534.02 | -0.14 | 5.58E-01 |
| ENSMUSG00000056938 | GSMG0005975 | Acbd4 | acyl-Coenzyme A binding domain containing 4 | 729.47 | 363.36 | 1.01 | 1.19E-08 |
| ENSMUSG00000048878 | GSMG0005976 | Hexim1 | hexamethylene bis-acetamide inducible 1 | 429.40 | 311.66 | 0.46 | 5.67E-02 |
| ENSMUSG00000055805 | GSMG0005978 | Fmnl1 | formin-like 1 | 1061.86 | 455.72 | 1.22 | 2.39E-10 |
| NULL | GSMG0005980 | NULL | NULL | 115.81 | 115.12 | 0.01 | 9.83E-01 |
| NULL | GSMG0005982 | NULL | NULL | 136.33 | 134.31 | 0.01 | 9.79E-01 |
| ENSMUSG00000046215 | GSMG0005988 | Rprml | reprimo-like | 832.58 | 586.06 | 0.51 | 2.94E-02 |
| ENSMUSG00000062421 | GSMG0005992 | Arf2 | ADP-ribosylation factor 2 | 4270.83 | 4473.31 | -0.07 | 7.85E-01 |
| ENSMUSG00000018634 | GSMG0005993 | Crhr1 | corticotropin releasing hormone receptor 1 | 322.06 | 315.51 | 0.03 | 9.48E-01 |
| ENSMUSG00000018411 | GSMG0005995 | Mapt | microtubule-associated protein tau | 1071.88 | 4482.07 | -2.07 | 8.31E-45 |
| ENSMUSG00000061086 | GSMG0005998 | Myl4 | myosin, light polypeptide 4 | 386.46 | 277.43 | 0.48 | 3.06E-02 |
| ENSMUSG00000020691 | GSMG0006003 | Mettl2 | methyltransferase like 2 | 290.73 | 380.97 | -0.39 | 1.43E-01 |
| ENSMUSG00000020694 | GSMG0006005 | Tlk2 | tousled-like kinase 2 (Arabidopsis) | 436.66 | 467.22 | -0.10 | 7.67E-01 |
| ENSMUSG00000020695 | GSMG0006010 | Mrc2 | mannose receptor, C type 2 | 949.00 | 844.57 | 0.17 | 6.34E-01 |
| ENSMUSG00000053580 | GSMG0006015 | Tanc2 | tetratricopeptide repeat, ankyrin repeat and coiled-coil containing 2 | 448.08 | 596.37 | -0.41 | 1.63E-01 |
| NULL | GSMG0006018 | NULL | NULL | 80.94 | 73.93 | 0.12 | 8.43E-01 |
| ENSMUSG00000020681 | GSMG0006023 | Ace | angiotensin I converting enzyme (peptidyl-dipeptidase A) 1 | 625.53 | 240.66 | 1.38 | 2.39E-02 |
| ENSMUSG00000049354 | GSMG0006026 | Dcaf7 | DDB1 and CUL4 associated factor 7 | 724.90 | 721.64 | 0.01 | 9.85E-01 |

| | | | | | | | |
|---|---|---|---|---|---|---|---|
| ENSMUSG00000001983 | GSMG0006027 | Taco1 | translational activator of mitochondrially encoded cytochrome c oxidase | 134.79 | 155.20 | -0.21 | 6.32E-01 |
| ENSMUSG00000020700 | GSMG0006030 | Map3k3 | mitogen-activated protein kinase kinase kinase 3 | 2755.32 | 2058.26 | 0.42 | 1.12E-01 |
| ENSMUSG00000020705 | GSMG0006033 | Ddx42 | DEAD (Asp-Glu-Ala-Asp) box polypeptide 42 | 1614.89 | 1894.51 | -0.23 | 2.23E-01 |
| ENSMUSG00000020708 | GSMG0006034 | Psmc5 | protease (prosome, macropain) 26S subunit, ATPase 5 | 3272.06 | 3577.88 | -0.13 | 5.79E-01 |
| NULL | GSMG0006038 | NULL | NULL | 110.86 | 141.40 | -0.36 | 4.30E-01 |
| ENSMUSG00000018372 | GSMG0006039 | Cep95 | centrosomal protein 95 | 504.09 | 438.74 | 0.20 | 4.64E-01 |
| ENSMUSG00000020720 | GSMG0006043 | Psmd12 | proteasome (prosome, macropain) 26S subunit, non-ATPase, 12 | 2184.26 | 2665.83 | -0.29 | 1.74E-01 |
| ENSMUSG00000020721 | GSMG0006045 | Helz | helicase with zinc finger domain | 1026.11 | 1070.26 | -0.06 | 8.50E-01 |
| ENSMUSG00000000142 | GSMG0006054 | Axin2 | axin 2 | 515.83 | 498.30 | 0.05 | 9.65E-01 |
| ENSMUSG00000020611 | GSMG0006057 | Gna13 | guanine nucleotide binding protein, alpha 13 | 1082.48 | 1088.34 | -0.01 | 9.80E-01 |
| ENSMUSG00000020610 | GSMG0006060 | Amz2 // LOC1005 | archaelysin family metallopeptidase 2 // uncharacterized LOC10050349 | 518.31 | 628.08 | -0.27 | 2.95E-01 |
| ENSMUSG00000020604 | GSMG0006062 | Arsg | arylsulfatase G | 66.93 | 75.75 | -0.17 | 8.26E-01 |
| ENSMUSG00000020612 | GSMG0006064 | Prkar1a | protein kinase, cAMP dependent regulatory, type I, alpha | 9887.26 | 9823.60 | 0.01 | 9.76E-01 |
| ENSMUSG00000020623 | GSMG0006069 | Map2k6 | mitogen-activated protein kinase kinase 6 | 138.46 | 146.42 | -0.07 | 8.79E-01 |
| ENSMUSG00000051497 | GSMG0006073 | Kcnj16 | potassium inwardly-rectifying channel, subfamily J, member 16 | 272.18 | 364.08 | -0.42 | 1.23E-01 |
| ENSMUSG00000041695 | GSMG0006074 | Kcnj2 | potassium inwardly-rectifying channel, subfamily J, member 2 | 2616.58 | 5642.66 | -1.11 | 2.33E-14 |
| ENSMUSG00000000567 | GSMG0006075 | Sox9 | SRY (sex determining region Y)-box 9 | 954.88 | 1196.95 | -0.33 | 2.07E-01 |
| ENSMUSG00000047904 | GSMG0006080 | Sstr2 | somatostatin receptor 2 | 257.10 | 285.97 | -0.15 | 6.61E-01 |
| ENSMUSG00000018661 | GSMG0006081 | Cog1 | component of oligomeric golgi complex 1 | 2004.14 | 2144.61 | -0.10 | 7.07E-01 |
| ENSMUSG00000041623 | GSMG0006083 | D11Wsu47e | DNA segment, Chr 11, Wayne State University 47, expressed | 430.04 | 525.68 | -0.29 | 1.81E-01 |
| ENSMUSG00000034714 | GSMG0006088 | Ttyh2 | tweety family member 2 | 7908.74 | 10145.30 | -0.36 | 2.20E-01 |
| ENSMUSG00000051043 | GSMG0006093 | Gprc5c | G protein-coupled receptor, family C, group 5, member C | 821.70 | 630.42 | 0.38 | 6.42E-02 |
| ENSMUSG00000020733 | GSMG0006095 | Slc9a3r1 | solute carrier family 9 (sodium/hydrogen exchanger), member 3 regulat | 465.61 | 526.30 | -0.18 | 4.88E-01 |
| ENSMUSG00000045980 | GSMG0006096 | Tmem104 | transmembrane protein 104 | 224.18 | 176.15 | 0.34 | 3.60E-01 |
| ENSMUSG00000050910 | GSMG0006099 | Cdr2l | cerebellar degeneration-related protein 2-like | 1372.68 | 1610.05 | -0.23 | 3.32E-01 |
| ENSMUSG00000018858 | GSMG0006101 | Ict1 | immature colon carcinoma transcript 1 | 669.24 | 882.66 | -0.40 | 2.85E-02 |
| ENSMUSG00000016940 | GSMG0006102 | Kctd2 | potassium channel tetramerisation domain containing 2 | 1494.57 | 1351.19 | 0.15 | 5.89E-01 |
| ENSMUSG00000020739 | GSMG0006107 | Nup85 | nucleoporin 85 | 1352.26 | 1850.38 | -0.45 | 5.54E-03 |
| ENSMUSG00000020747 | GSMG0006111 | Tmem94 | transmembrane protein 94 | 466.80 | 508.08 | -0.12 | 7.20E-01 |
| ENSMUSG00000020781 | GSMG0006112 | Tsen54 | tRNA splicing endonuclease subunit 54 | 672.02 | 726.35 | -0.11 | 6.53E-01 |
| ENSMUSG00000075420 | GSMG0006117 | Smim6 | small integral membrane protein 6 | 74.21 | 61.57 | 0.27 | 5.82E-01 |
| ENSMUSG00000020755 | GSMG0006118 | Sap30bp | SAP30 binding protein | 456.84 | 533.41 | -0.22 | 4.09E-01 |
| ENSMUSG00000020758 | GSMG0006120 | Itgb4 | integrin beta 4 | 215.77 | 157.88 | 0.45 | 2.10E-01 |

| | | | | | | | |
|---|---|---|---|---|---|---|---|
| ENSMUSG00000020770 | GSMG0006121 | Unk | unkempt family zinc finger | 1571.64 | 1677.22 | -0.09 | 7.06E-01 |
| ENSMUSG00000020778 | GSMG0006126 | Ten1 | TEN1 telomerase capping complex subunit | 114.45 | 135.83 | -0.24 | 5.49E-01 |
| ENSMUSG00000020812 | GSMG0006138 | 1810032O08Rik | RIKEN cDNA 1810032O08 gene | 88.53 | 136.09 | -0.62 | 4.59E-02 |
| ENSMUSG00000090266 | GSMG0006142 | Mettl23 | methyltransferase like 23 | 979.22 | 864.46 | 0.18 | 3.91E-01 |
| ENSMUSG00000020818 | GSMG0006143 | Mfsd11 | major facilitator superfamily domain containing 11 | 698.01 | 756.31 | -0.12 | 6.40E-01 |
| ENSMUSG00000043857 | GSMG0006144 | Mgat5b | mannoside acetylglucosaminyltransferase 5, isoenzyme B | 367.87 | 723.39 | -0.98 | 1.94E-05 |
| NULL | GSMG0006146 | NULL | NULL | 262.63 | 379.93 | -0.53 | 1.75E-02 |
| ENSMUSG00000086859 | GSMG0006147 | Snhg20 | small nucleolar RNA host gene 20 | 1838.66 | 1431.90 | 0.36 | 1.44E-01 |
| ENSMUSG00000020823 | GSMG0006148 | Sec14l1 | SEC14-like lipid binding 1 | 1284.64 | 1244.60 | 0.05 | 8.73E-01 |
| ENSMUSG00000059248 | GSMG0006150 | Sept9 | septin 9 | 724.09 | 1221.88 | -0.76 | 6.66E-05 |
| ENSMUSG00000025571 | GSMG0006158 | Tnrc6c | trinucleotide repeat containing 6C | 228.49 | 179.06 | 0.35 | 2.39E-01 |
| NULL | GSMG0006159 | NULL | NULL | 216.33 | 184.78 | 0.22 | 5.32E-01 |
| NULL | GSMG0006160 | NULL | NULL | 97.87 | 100.76 | -0.03 | 9.63E-01 |
| ENSMUSG00000050106 | GSMG0006161 | Tmc8 | transmembrane channel-like gene family 8 | 135.62 | 182.89 | -0.42 | 1.86E-01 |
| G00000017718,ENSMUSG000 | GSMG0006163 | Syngr2 // Afmid | synaptogyrin 2 // arylformamidase | 2423.82 | 2040.15 | 0.25 | 1.97E-01 |
| ENSMUSG00000017716 | GSMG0006164 | Birc5 | baculoviral IAP repeat-containing 5 | 423.32 | 1148.30 | -1.44 | 7.67E-13 |
| ENSMUSG00000017715 | GSMG0006168 | Pgs1 | phosphatidylglycerophosphate synthase 1 | 1518.53 | 1478.12 | 0.04 | 8.98E-01 |
| ENSMUSG00000017446 | GSMG0006174 | C1qtnf1 | C1q and tumor necrosis factor related protein 1 | 1631.56 | 1452.85 | 0.17 | 4.70E-01 |
| ENSMUSG00000033857 | GSMG0006175 | Engase | endo-beta-N-acetylglucosaminidase | 514.44 | 593.00 | -0.20 | 4.86E-01 |
| ENSMUSG00000025577 | GSMG0006179 | Cbx2 | chromobox 2 | 341.32 | 575.36 | -0.75 | 3.23E-05 |
| ENSMUSG00000025579 | GSMG0006182 | Gaa | glucosidase, alpha, acid | 3003.88 | 2871.85 | 0.07 | 8.51E-01 |
| ENSMUSG00000039908 | GSMG0006184 | Slc26a11 | solute carrier family 26, member 11 | 291.15 | 268.67 | 0.12 | 7.49E-01 |
| ENSMUSG00000039850 | GSMG0006188 | Endov | endonuclease V | 230.27 | 263.64 | -0.19 | 5.34E-01 |
| ENSMUSG00000025583 | GSMG0006190 | Rptor | regulatory associated protein of MTOR, complex 1 | 248.11 | 252.94 | -0.03 | 9.57E-01 |
| ENSMUSG00000025371 | GSMG0006197 | Chmp6 | charged multivesicular body protein 6 | 2200.64 | 3149.62 | -0.52 | 2.91E-04 |
| ENSMUSG00000025372 | GSMG0006199 | Baiap2 | brain-specific angiogenesis inhibitor 1-associated protein 2 | 2371.79 | 2117.60 | 0.16 | 4.10E-01 |
| ENSMUSG00000078572 | GSMG0006201 | 1810043H04Rik | RIKEN cDNA 1810043H04 gene | 391.46 | 404.66 | -0.05 | 8.87E-01 |
| ENSMUSG00000043644 | GSMG0006206 | 0610009L18Rik | RIKEN cDNA 0610009L18 gene | 89.24 | 91.18 | -0.03 | 9.57E-01 |
| NULL | GSMG0006209 | NULL | NULL | 53.78 | 60.46 | -0.18 | 7.72E-01 |
| ENSMUSG00000049957 | GSMG0006212 | Ccdc137 | coiled-coil domain containing 137 | 155.98 | 175.67 | -0.17 | 6.43E-01 |
| ENSMUSG00000025793 | GSMG0006213 | Hgs | HGF-regulated tyrosine kinase substrate | 2377.45 | 2706.72 | -0.19 | 3.83E-01 |
| ENSMUSG00000025792 | GSMG0006215 | Slc25a10 | solute carrier family 25 (mitochondrial carrier, dicarboxylate transporte | 361.82 | 489.17 | -0.43 | 4.89E-02 |
| NULL | GSMG0006218 | NULL | NULL | 3651.12 | 4115.47 | -0.17 | 3.66E-01 |

| | | | | | | | |
|---|---|---|---|---|---|---|---|
| ENSMUSG00000025135 | GSMG0006219 | Anapc11 | anaphase promoting complex subunit 11 | 1190.11 | 1396.43 | -0.23 | 3.48E-01 |
| NULL | GSMG0006221 | NULL | NULL | 781.66 | 465.52 | 0.75 | 9.55E-05 |
| NULL | GSMG0006222 | NULL | NULL | 113.40 | 98.60 | 0.20 | 6.57E-01 |
| NULL | GSMG0006223 | NULL | NULL | 108.45 | 132.66 | -0.30 | 4.84E-01 |
| NULL | GSMG0006225 | NULL | NULL | 921.54 | 610.90 | 0.59 | 1.95E-02 |
| ENSMUSG00000025142 | GSMG0006226 | Aspscr1 | alveolar soft part sarcoma chromosome region, candidate 1 (human) | 3547.00 | 3666.74 | -0.05 | 8.44E-01 |
| ENSMUSG00000025145 | GSMG0006227 | Lrrc45 | leucine rich repeat containing 45 | 5401.87 | 5023.09 | 0.11 | 6.57E-01 |
| ENSMUSG00000018012 | GSMG0006228 | Rac3 | RAS-related C3 botulinum substrate 3 | 503.28 | 2484.91 | -2.30 | 2.79E-53 |
| ENSMUSG00000039307 | GSMG0006236 | Hexdc | hexosaminidase (glycosyl hydrolase family 20, catalytic domain) contai | 620.65 | 742.90 | -0.26 | 3.38E-01 |
| ENSMUSG00000039230 | GSMG0006241 | Tbcd | tubulin-specific chaperone d | 2373.23 | 2759.98 | -0.22 | 2.89E-01 |
| NULL | GSMG0006249 | NULL | NULL | 102.26 | 120.71 | -0.25 | 5.56E-01 |
| ENSMUSG00000020457 | GSMG0006251 | Drg1 | developmentally regulated GTP binding protein 1 | 1058.32 | 1153.25 | -0.12 | 6.37E-01 |
| ENSMUSG00000020451 | GSMG0006256 | Limk2 | LIM motif-containing protein kinase 2 | 1920.47 | 2192.26 | -0.19 | 3.33E-01 |
| ENSMUSG00000020448 | GSMG0006257 | Rnf185 | ring finger protein 185 | 496.09 | 591.82 | -0.26 | 2.95E-01 |
| ENSMUSG00000034570 | GSMG0006258 | Inpp5j | inositol polyphosphate 5-phosphatase J | 1513.80 | 527.61 | 1.52 | 6.20E-18 |
| ENSMUSG00000020439 | GSMG0006259 | Smtn | smoothelin | 7273.42 | 7458.87 | -0.04 | 8.90E-01 |
| ENSMUSG00000056579 | GSMG0006260 | Tug1 | taurine upregulated gene 1 | 1804.94 | 2761.51 | -0.61 | 4.78E-04 |
| ENSMUSG00000020435 | GSMG0006261 | Osbp2 | oxysterol binding protein 2 | 559.57 | 412.84 | 0.44 | 5.89E-02 |
| ENSMUSG00000048807 | GSMG0006264 | Slc35e4 | solute carrier family 35, member E4 | 1165.34 | 804.94 | 0.53 | 1.18E-02 |
| ENSMUSG00000020432 | GSMG0006265 | Tcn2 | transcobalamin 2 | 4939.01 | 3341.02 | 0.56 | 5.13E-04 |
| ENSMUSG00000004748 | GSMG0006267 | Mtfp1 | mitochondrial fission process 1 | 133.12 | 50.45 | 1.40 | 3.96E-06 |
| ENSMUSG00000003585 | GSMG0006268 | Sec14l2 | SEC14-like lipid binding 2 | 109.16 | 72.30 | 0.59 | 1.32E-01 |
| ENSMUSG00000051427 | GSMG0006269 | Ccdc157 | coiled-coil domain containing 157 | 283.55 | 196.10 | 0.53 | 1.26E-01 |
| ENSMUSG00000034354 | GSMG0006272 | Mtmr3 | myotubularin related protein 3 | 1303.96 | 1379.75 | -0.08 | 8.21E-01 |
| NULL | GSMG0006278 | NULL | NULL | 56.01 | 66.68 | -0.25 | 6.36E-01 |
| ENSMUSG00000059534 | GSMG0006279 | Uqcr10 | ubiquinol-cytochrome c reductase, complex III subunit X | 2259.53 | 2520.49 | -0.16 | 5.78E-01 |
| ENSMUSG00000009073 | GSMG0006281 | Nf2 | neurofibromatosis 2 | 1307.49 | 1433.45 | -0.13 | 5.76E-01 |
| ENSMUSG00000020396 | GSMG0006282 | Nefh | neurofilament, heavy polypeptide | 281.34 | 137.25 | 1.03 | 2.79E-06 |
| ENSMUSG00000034201 | GSMG0006284 | Gas2l1 | growth arrest-specific 2 like 1 | 1042.64 | 761.60 | 0.45 | 2.11E-02 |
| ENSMUSG00000009079 | GSMG0006286 | Ewsr1 | Ewing sarcoma breakpoint region 1 | 8912.72 | 9150.78 | -0.04 | 8.94E-01 |
| ENSMUSG00000034164 | GSMG0006287 | Emid1 | EMI domain containing 1 | 1145.03 | 1309.19 | -0.19 | 3.91E-01 |
| ENSMUSG00000020393 | GSMG0006288 | Kremen1 | kringle containing transmembrane protein 1 | 326.99 | 341.00 | -0.06 | 8.72E-01 |
| ENSMUSG00000041961 | GSMG0006290 | Znrf3 | zinc and ring finger 3 | 127.86 | 77.92 | 0.72 | 1.46E-01 |

| | | | | | | | |
|---|---|---|---|---|---|---|---|
| ENSMUSG00000020482 | GSMG0006298 | Ccdc117 | coiled-coil domain containing 117 | 1308.96 | 1160.30 | 0.17 | 4.30E-01 |
| ENSMUSG00000020477 | GSMG0006300 | Mrps24 | mitochondrial ribosomal protein S24 | 1160.85 | 1375.97 | -0.25 | 2.44E-01 |
| ENSMUSG00000049680 | GSMG0006301 | Urgcp | upregulator of cell proliferation | 1838.48 | 1418.83 | 0.37 | 6.45E-02 |
| NULL | GSMG0006305 | NULL | NULL | 60.20 | 44.35 | 0.43 | 4.26E-01 |
| ENSMUSG00000020475 | GSMG0006306 | Pgam2 | phosphoglycerate mutase 2 | 83.02 | 53.33 | 0.64 | 2.12E-01 |
| ENSMUSG00000020471 | GSMG0006308 | Pold2 | polymerase (DNA directed), delta 2, regulatory subunit | 386.36 | 586.73 | -0.60 | 2.02E-03 |
| ENSMUSG00000057897 | GSMG0006311 | Camk2b | calcium/calmodulin-dependent protein kinase II, beta | 3888.84 | 3558.52 | 0.13 | 6.94E-01 |
| ENSMUSG00000053838 | GSMG0006313 | Nudcd3 | NudC domain containing 3 | 637.23 | 722.70 | -0.18 | 4.13E-01 |
| ENSMUSG00000004393 | GSMG0006317 | Ddx56 | DEAD (Asp-Glu-Ala-Asp) box polypeptide 56 | 1315.64 | 1655.17 | -0.33 | 4.34E-02 |
| ENSMUSG00000004394 | GSMG0006318 | Tmed4 | transmembrane emp24 protein transport domain containing 4 | 3619.67 | 3385.01 | 0.10 | 7.06E-01 |
| ENSMUSG00000041126 | GSMG0006321 | H2afv | H2A histone family, member V | 2565.14 | 2800.22 | -0.13 | 5.95E-01 |
| ENSMUSG00000085156 | GSMG0006327 | Snhg15 | small nucleolar RNA host gene 15 | 155.02 | 191.98 | -0.31 | 3.36E-01 |
| ENSMUSG00000041073 | GSMG0006328 | Nacad | NAC alpha domain containing | 1240.72 | 1734.59 | -0.48 | 6.97E-02 |
| ENSMUSG00000000384 | GSMG0006329 | Tbrg4 | transforming growth factor beta regulated gene 4 | 477.04 | 534.43 | -0.17 | 5.54E-01 |
| ENSMUSG00000020427 | GSMG0006333 | Igfbp3 | insulin-like growth factor binding protein 3 | 6384.63 | 1283.70 | 2.31 | 1.43E-41 |
| ENSMUSG00000020422 | GSMG0006335 | Tns3 | tensin 3 | 566.08 | 526.25 | 0.10 | 8.32E-01 |
| ENSMUSG00000020413 | GSMG0006338 | Hus1 | HUS1 checkpoint clamp component | 146.14 | 183.86 | -0.33 | 3.04E-01 |
| ENSMUSG00000020182 | GSMG0006346 | Ddc | dopa decarboxylase | 766.85 | 582.21 | 0.40 | 8.56E-02 |
| ENSMUSG00000020176 | GSMG0006347 | Grb10 | growth factor receptor bound protein 10 | 1821.54 | 2630.77 | -0.53 | 3.13E-03 |
| ENSMUSG00000020116 | GSMG0006362 | Pno1 | partner of NOB1 homolog | 218.24 | 340.98 | -0.64 | 8.71E-03 |
| ENSMUSG00000016984 | GSMG0006363 | Etaa1 | Ewing tumor-associated antigen 1 | 277.71 | 395.98 | -0.51 | 2.71E-02 |
| ENSMUSG00000020152 | GSMG0006379 | Actr2 | ARP2 actin-related protein 2 | 966.80 | 1061.41 | -0.14 | 7.06E-01 |
| ENSMUSG00000044066 | GSMG0006384 | Cep68 | centrosomal protein 68 | 2256.61 | 3087.73 | -0.45 | 3.29E-03 |
| ENSMUSG00000020142 | GSMG0006386 | Slc1a4 | solute carrier family 1 (glutamate/neutral amino acid transporter), memb | 1081.83 | 5498.99 | -2.34 | 2.30E-44 |
| NULL | GSMG0006387 | NULL | NULL | 286.42 | 1226.53 | -2.10 | 2.21E-25 |
| NULL | GSMG0006389 | NULL | NULL | 151.83 | 136.45 | 0.15 | 7.93E-01 |
| ENSMUSG00000049659 | GSMG0006390 | Aftph | aftiphilin | 1502.87 | 1294.88 | 0.22 | 2.71E-01 |
| ENSMUSG00000042363 | GSMG0006392 | Lgalsl | lectin, galactoside binding-like | 857.12 | 825.38 | 0.05 | 8.89E-01 |
| ENSMUSG00000001891 | GSMG0006393 | Ugp2 | UDP-glucose pyrophosphorylase 2 | 2321.68 | 2918.29 | -0.33 | 1.54E-01 |
| ENSMUSG00000020321 | GSMG0006395 | Mdh1 | malate dehydrogenase 1, NAD (soluble) | 4864.80 | 4139.24 | 0.23 | 2.59E-01 |
| NULL | GSMG0006396 | NULL | NULL | 262.18 | 170.82 | 0.62 | 2.25E-02 |
| ENSMUSG00000042302 | GSMG0006398 | Ehbp1 | EH domain binding protein 1 | 310.56 | 320.81 | -0.05 | 9.06E-01 |
| NULL | GSMG0006401 | NULL | NULL | 331.29 | 263.25 | 0.33 | 2.19E-01 |

| | | | | | | | |
|---|---|---|---|---|---|---|---|
| G00000051355,ENSMUSG000 | GSMG0006414 | B3gnt2 // Commd1 | UDP-GlcNAc:betaGal beta-1,3-N-acetylglucosaminyltransferase 2 // C( | 6486.31 | 5147.62 | 0.33 | 1.37E-01 |
| NULL | GSMG0006416 | NULL | NULL | 344.87 | 173.27 | 0.99 | 1.22E-04 |
| ENSMUSG00000020288 | GSMG0006422 | Ahsa2 | AHA1, activator of heat shock protein ATPase 2 | 858.46 | 1043.16 | -0.28 | 1.49E-01 |
| ENSMUSG00000042208 | GSMG0006425 | 0610010F05Rik | RIKEN cDNA 0610010F05 gene | 808.88 | 1080.29 | -0.42 | 3.51E-02 |
| ENSMUSG00000020283 | GSMG0006427 | Pex13 | peroxisomal biogenesis factor 13 | 388.87 | 402.20 | -0.05 | 8.99E-01 |
| ENSMUSG00000020273 | GSMG0006431 | Papolg | poly(A) polymerase gamma | 424.26 | 431.33 | -0.02 | 9.50E-01 |
| NULL | GSMG0006439 | NULL | NULL | 137.89 | 162.69 | -0.24 | 4.96E-01 |
| ENSMUSG00000064090 | GSMG0006443 | Vrk2 | vaccinia related kinase 2 | 81.93 | 72.12 | 0.19 | 7.25E-01 |
| ENSMUSG00000032878 | GSMG0006448 | Ccdc85a | coiled-coil domain containing 85A | 1174.38 | 967.16 | 0.28 | 1.94E-01 |
| NULL | GSMG0006449 | NULL | NULL | 272.20 | 222.92 | 0.29 | 2.93E-01 |
| ENSMUSG00000020462 | GSMG0006451 | Cfap36 | cilia and flagella associated protein 36 | 827.67 | 931.76 | -0.17 | 5.23E-01 |
| NULL | GSMG0006452 | NULL | NULL | 174.87 | 245.54 | -0.49 | 6.58E-02 |
| ENSMUSG00000032673 | GSMG0006453 | Prorsd1 | prolyl-tRNA synthetase domain containing 1 | 259.81 | 204.73 | 0.34 | 2.81E-01 |
| ENSMUSG00000020315 | GSMG0006460 | Sptbn1 | spectrin beta, non-erythrocytic 1 | 7429.10 | 6175.39 | 0.27 | 2.98E-01 |
| ENSMUSG00000020311 | GSMG0006471 | Erlec1 | endoplasmic reticulum lectin 1 | 1215.31 | 1212.68 | 0.00 | 9.91E-01 |
| ENSMUSG00000020309 | GSMG0006473 | Chac2 | ChaC, cation transport regulator 2 | 217.43 | 248.58 | -0.20 | 6.21E-01 |
| ENSMUSG00000044502 | GSMG0006476 | Bod1 | biorientation of chromosomes in cell division 1 | 462.17 | 528.48 | -0.19 | 4.19E-01 |
| ENSMUSG00000020282 | GSMG0006482 | Rhbdf1 | rhomboid 5 homolog 1 | 3630.15 | 3957.00 | -0.12 | 5.64E-01 |
| ENSMUSG00000020289 | GSMG0006483 | Nprl3 | nitrogen permease regulator-like 3 | 1089.31 | 1125.78 | -0.05 | 8.74E-01 |
| NULL | GSMG0006491 | NULL | NULL | 52.15 | 104.49 | -1.01 | 3.62E-03 |
| ENSMUSG00000057113 | GSMG0006492 | Npm1 | nucleophosmin 1 | 697.57 | 1055.69 | -0.60 | 3.76E-03 |
| ENSMUSG00000040594 | GSMG0006494 | Ranbp17 | RAN binding protein 17 | 255.29 | 308.34 | -0.27 | 2.87E-01 |
| ENSMUSG00000053519 | GSMG0006498 | Kcnip1 | Kv channel-interacting protein 1 | 66.70 | 83.89 | -0.33 | 4.62E-01 |
| ENSMUSG00000069910 | GSMG0006511 | Spdl1 | spindle apparatus coiled-coil protein 1 | 46.53 | 120.32 | -1.35 | 2.84E-04 |
| ENSMUSG00000051062 | GSMG0006514 | Fbll1 | fibrillarin-like 1 | 137.87 | 159.19 | -0.22 | 6.81E-01 |
| ENSMUSG00000018848 | GSMG0006515 | Rars | arginyl-tRNA synthetase | 1091.88 | 1489.03 | -0.45 | 4.16E-02 |
| ENSMUSG00000018849 | GSMG0006516 | Wwc1 | WW, C2 and coiled-coil domain containing 1 | 900.65 | 1078.58 | -0.26 | 3.25E-01 |
| ENSMUSG00000049336 | GSMG0006519 | Tenm2 | teneurin transmembrane protein 2 | 1033.53 | 1947.43 | -0.91 | 9.25E-06 |
| NULL | GSMG0006524 | NULL | NULL | 114.92 | 130.51 | -0.18 | 7.13E-01 |
| NULL | GSMG0006525 | NULL | NULL | 377.63 | 557.09 | -0.56 | 8.02E-03 |
| NULL | GSMG0006528 | NULL | NULL | 483.82 | 690.99 | -0.51 | 8.42E-03 |
| NULL | GSMG0006529 | NULL | NULL | 748.15 | 971.93 | -0.38 | 1.59E-01 |
| NULL | GSMG0006530 | NULL | NULL | 349.34 | 484.10 | -0.47 | 6.33E-02 |

| | | | | | | | |
|---|---|---|---|---|---|---|---|
| NULL | GSMG0006531 | NULL | NULL | 1905.27 | 2615.96 | -0.46 | 2.37E-02 |
| NULL | GSMG0006532 | NULL | NULL | 278.25 | 504.67 | -0.86 | 4.05E-05 |
| NULL | GSMG0006533 | NULL | NULL | 297.03 | 388.92 | -0.39 | 1.83E-01 |
| NULL | GSMG0006534 | NULL | NULL | 490.44 | 754.56 | -0.62 | 4.17E-02 |
| NULL | GSMG0006535 | NULL | NULL | 315.09 | 547.41 | -0.80 | 1.47E-03 |
| NULL | GSMG0006536 | NULL | NULL | 282.22 | 687.63 | -1.29 | 2.24E-10 |
| NULL | GSMG0006538 | NULL | NULL | 526.01 | 852.39 | -0.69 | 1.18E-02 |
| NULL | GSMG0006539 | NULL | NULL | 148.67 | 237.74 | -0.68 | 1.03E-02 |
| ENSMUSG00000042032 | GSMG0006546 | Mat2b | methionine adenosyltransferase II, beta | 2498.77 | 1995.67 | 0.32 | 9.54E-02 |
| ENSMUSG00000020330 | GSMG0006547 | Hmmr | hyaluronan mediated motility receptor (RHAMM) | 68.22 | 159.48 | -1.23 | 5.62E-05 |
| ENSMUSG00000020326 | GSMG0006548 | Ccng1 | cyclin G1 | 744.94 | 627.15 | 0.25 | 3.46E-01 |
| ENSMUSG00000020436 | GSMG0006549 | Gabrg2 | gamma-aminobutyric acid (GABA) A receptor, subunit gamma 2 | 211.37 | 353.66 | -0.74 | 5.94E-04 |
| ENSMUSG00000010803 | GSMG0006553 | Gabra1 | gamma-aminobutyric acid (GABA) A receptor, subunit alpha 1 | 405.33 | 266.38 | 0.61 | 1.44E-02 |
| NULL | GSMG0006554 | NULL | NULL | 179.25 | 192.53 | -0.11 | 7.94E-01 |
| ENSMUSG00000020415 | GSMG0006559 | Pttg1 | pituitary tumor-transforming gene 1 | 286.12 | 742.08 | -1.37 | 1.30E-13 |
| ENSMUSG00000041278 | GSMG0006563 | Ttc1 | tetratricopeptide repeat domain 1 | 711.32 | 745.89 | -0.07 | 8.26E-01 |
| ENSMUSG00000050541 | GSMG0006564 | Adra1b | adrenergic receptor, alpha 1b | 94.89 | 112.40 | -0.25 | 5.79E-01 |
| NULL | GSMG0006566 | NULL | NULL | 538.92 | 439.22 | 0.29 | 4.48E-01 |
| ENSMUSG00000044847 | GSMG0006572 | Lsm11 | U7 snRNP-specific Sm-like protein LSM11 | 156.24 | 227.38 | -0.54 | 4.34E-02 |
| ENSMUSG00000011254 | GSMG0006573 | Thg1l | tRNA-histidine guanylyltransferase 1-like (S. cerevisiae) | 34.65 | 57.64 | -0.73 | 1.49E-01 |
| ENSMUSG00000020340 | GSMG0006575 | Cyfip2 | cytoplasmic FMR1 interacting protein 2 | 478.19 | 454.97 | 0.07 | 8.45E-01 |
| NULL | GSMG0006576 | NULL | NULL | 571.30 | 366.79 | 0.64 | 1.21E-03 |
| NULL | GSMG0006581 | NULL | NULL | 233.82 | 248.82 | -0.09 | 8.33E-01 |
| ENSMUSG00000046879 | GSMG0006599 | Irgm1 | immunity-related GTPase family M member 1 | 876.08 | 1948.35 | -1.15 | 7.97E-02 |
| ENSMUSG00000020362 | GSMG0006610 | Cnot6 | CCR4-NOT transcription complex, subunit 6 | 2170.01 | 2305.84 | -0.09 | 7.47E-01 |
| ENSMUSG00000015837 | GSMG0006611 | Sqstm1 | sequestosome 1 | 3775.41 | 3276.42 | 0.20 | 3.56E-01 |
| ENSMUSG00000020377 | GSMG0006613 | Ltc4s | leukotriene C4 synthase | 206.14 | 98.87 | 1.05 | 5.09E-04 |
| ENSMUSG00000020368 | GSMG0006617 | Canx | calnexin | 10561.00 | 12690.09 | -0.26 | 1.34E-01 |
| ENSMUSG00000020375 | GSMG0006619 | Rufy1 | RUN and FYVE domain containing 1 | 927.84 | 850.68 | 0.13 | 5.89E-01 |
| ENSMUSG00000044807 | GSMG0006622 | Zfp354c | zinc finger protein 354C | 797.84 | 1197.36 | -0.59 | 3.62E-04 |
| ENSMUSG00000048728 | GSMG0006624 | Zfp454 | zinc finger protein 454 | 179.98 | 203.90 | -0.18 | 6.11E-01 |
| ENSMUSG00000049321 | GSMG0006625 | Zfp2 | zinc finger protein 2 | 443.87 | 571.90 | -0.37 | 6.48E-02 |
| ENSMUSG00000020358 | GSMG0006634 | Hnrnpab | heterogeneous nuclear ribonucleoprotein A/B | 2581.98 | 3874.35 | -0.59 | 2.54E-03 |

| | | | | | | | |
|---|---|---|---|---|---|---|---|
| ENSMUSG00000001054 | GSMG0006635 | Rmnd5b | required for meiotic nuclear division 5 homolog B | 1605.68 | 1563.81 | 0.04 | 8.97E-01 |
| ENSMUSG00000001053 | GSMG0006637 | N4bp3 | NEDD4 binding protein 3 | 1737.51 | 1846.21 | -0.09 | 7.39E-01 |
| ENSMUSG00000007777 | GSMG0006638 | 0610009B22Rik | RIKEN cDNA 0610009B22 gene | 546.93 | 542.92 | 0.01 | 9.85E-01 |
| ENSMUSG00000036391 | GSMG0006639 | Sec24a | Sec24 related gene family, member A (S. cerevisiae) | 205.43 | 209.26 | -0.03 | 9.53E-01 |
| ENSMUSG00000020390 | GSMG0006642 | Ube2b | ubiquitin-conjugating enzyme E2B | 2741.73 | 2332.73 | 0.23 | 2.70E-01 |
| ENSMUSG00000108218 | GSMG0006645 | Olfr1372-ps1 | olfactory receptor 1372, pseudogene 1 | 629.50 | 227.17 | 1.47 | 6.71E-12 |
| ENSMUSG00000000782 | GSMG0006647 | Tcf7 | transcription factor 7, T cell specific | 931.57 | 909.49 | 0.03 | 9.19E-01 |
| ENSMUSG00000020361 | GSMG0006653 | Hspa4 | heat shock protein 4 | 2076.85 | 2814.16 | -0.44 | 1.01E-02 |
| ENSMUSG00000044894 | GSMG0006655 | Uqcrq | ubiquinol-cytochrome c reductase, complex III subunit VII | 2350.19 | 2875.87 | -0.29 | 2.06E-01 |
| NULL | GSMG0006658 | NULL | NULL | 211.57 | 71.39 | 1.56 | 2.08E-07 |
| ENSMUSG00000020380 | GSMG0006664 | Rad50 | RAD50 double strand break repair protein | 379.62 | 480.25 | -0.34 | 2.00E-01 |
| ENSMUSG00000018900 | GSMG0006668 | Slc22a5 | solute carrier family 22 (organic cation transporter), member 5 | 489.13 | 414.65 | 0.24 | 3.99E-01 |
| ENSMUSG00000020388 | GSMG0006672 | Pdlim4 | PDZ and LIM domain 4 | 136.21 | 135.86 | 0.01 | 9.89E-01 |
| ENSMUSG00000052298 | GSMG0006678 | Cdc42se2 | CDC42 small effector 2 | 1123.07 | 1102.58 | 0.03 | 9.31E-01 |
| ENSMUSG00000020400 | GSMG0006681 | Tnip1 | TNFAIP3 interacting protein 1 | 974.91 | 932.68 | 0.06 | 8.34E-01 |
| ENSMUSG00000018340 | GSMG0006683 | Anxa6 | annexin A6 | 4643.96 | 8339.64 | -0.84 | 1.11E-08 |
| ENSMUSG00000018593 | GSMG0006689 | Sparc | secreted acidic cysteine rich glycoprotein | 107258.43 | 88095.75 | 0.28 | 1.24E-01 |
| ENSMUSG00000018585 | GSMG0006690 | Atox1 | antioxidant 1 copper chaperone | 4701.37 | 4410.05 | 0.09 | 7.96E-01 |
| ENSMUSG00000020523 | GSMG0006694 | Fam114a2 | family with sequence similarity 114, member A2 | 1342.08 | 1244.80 | 0.11 | 6.32E-01 |
| ENSMUSG00000037275 | GSMG0006702 | Gemin5 | gem (nuclear organelle) associated protein 5 | 445.36 | 620.75 | -0.48 | 1.93E-02 |
| ENSMUSG00000049755 | GSMG0006703 | Zfp672 | zinc finger protein 672 | 229.39 | 182.89 | 0.33 | 2.50E-01 |
| ENSMUSG00000037001 | GSMG0006714 | Zfp39 | zinc finger protein 39 | 379.56 | 433.57 | -0.19 | 4.94E-01 |
| ENSMUSG00000020496 | GSMG0006715 | Rnf187 | ring finger protein 187 | 1295.86 | 1495.90 | -0.21 | 3.37E-01 |
| ENSMUSG00000020444 | GSMG0006725 | Guk1 | guanylate kinase 1 | 1126.36 | 1113.57 | 0.02 | 9.58E-01 |
| ENSMUSG00000048076 | GSMG0006728 | Arf1 | ADP-ribosylation factor 1 | 11464.48 | 12671.90 | -0.14 | 5.27E-01 |
| ENSMUSG00000009894 | GSMG0006731 | Snap47 | synaptosomal-associated protein, 47 | 2551.75 | 3164.64 | -0.31 | 1.04E-01 |
| ENSMUSG00000054519 | GSMG0006733 | Zfp867 | zinc finger protein 867 | 420.89 | 473.06 | -0.17 | 5.73E-01 |
| ENSMUSG00000020472 | GSMG0006734 | Zkscan17 | zinc finger with KRAB and SCAN domains 17 | 210.75 | 230.72 | -0.13 | 7.07E-01 |
| NULL | GSMG0006739 | NULL | NULL | 109.39 | 73.43 | 0.58 | 2.26E-01 |
| NULL | GSMG0006740 | NULL | NULL | 627.32 | 470.64 | 0.41 | 8.88E-02 |
| ENSMUSG00000032633 | GSMG0006742 | Flcn | folliculin | 931.61 | 676.29 | 0.46 | 1.52E-02 |
| ENSMUSG00000019373 | GSMG0006743 | Cops3 | COP9 signalosome subunit 3 | 1587.42 | 1825.27 | -0.20 | 3.13E-01 |
| ENSMUSG00000049892 | GSMG0006745 | Rasd1 | RAS, dexamethasone-induced 1 | 409.26 | 72.33 | 2.50 | 1.81E-16 |

| | | | | | | | |
|---|---|---|---|---|---|---|---|
| ENSMUSG00000000301 | GSMG0006746 | Pemt | phosphatidylethanolamine N-methyltransferase | 49.25 | 82.98 | -0.75 | 5.36E-02 |
| ENSMUSG00000020538 | GSMG0006748 | Srebf1 | sterol regulatory element binding transcription factor 1 | 1671.27 | 1663.18 | 0.01 | 9.85E-01 |
| ENSMUSG00000000538 | GSMG0006749 | Tom1l2 | target of myb1-like 2 (chicken) | 1913.30 | 1283.07 | 0.58 | 1.32E-03 |
| ENSMUSG00000042709 | GSMG0006751 | Atpaf2 | ATP synthase mitochondrial F1 complex assembly factor 2 | 316.25 | 486.37 | -0.62 | 1.71E-03 |
| ENSMUSG00000002812 | GSMG0006756 | Flii | flightless I actin binding protein | 5216.76 | 4950.47 | 0.08 | 7.58E-01 |
| ENSMUSG00000002814 | GSMG0006757 | Top3a | topoisomerase (DNA) III alpha | 280.15 | 381.56 | -0.44 | 1.12E-01 |
| ENSMUSG00000020534 | GSMG0006759 | Shmt1 | serine hydroxymethyltransferase 1 (soluble) | 83.63 | 170.94 | -1.02 | 4.05E-04 |
| ENSMUSG00000043284 | GSMG0006760 | Tmem11 | transmembrane protein 11 | 639.81 | 761.58 | -0.25 | 2.42E-01 |
| ENSMUSG00000018931 | GSMG0006762 | Natd1 | N-acetyltransferase domain containing 1 | 984.59 | 978.90 | 0.01 | 9.79E-01 |
| ENSMUSG00000042506 | GSMG0006764 | Usp22 | ubiquitin specific peptidase 22 | 4316.11 | 7824.47 | -0.86 | 1.06E-09 |
| ENSMUSG00000010025 | GSMG0006766 | Aldh3a2 | aldehyde dehydrogenase family 3, subfamily A2 | 947.41 | 887.70 | 0.09 | 7.39E-01 |
| ENSMUSG00000010086 | GSMG0006769 | Rnf112 | ring finger protein 112 | 4528.34 | 1460.20 | 1.63 | 3.13E-25 |
| ENSMUSG00000001034 | GSMG0006770 | Mapk7 | mitogen-activated protein kinase 7 | 1677.06 | 1615.35 | 0.05 | 8.65E-01 |
| NULL | GSMG0006773 | NULL | NULL | 91.93 | 197.80 | -1.10 | 3.10E-04 |
| ENSMUSG00000001036 | GSMG0006774 | Epn2 | epsin 2 | 2291.38 | 1994.88 | 0.20 | 2.96E-01 |
| ENSMUSG00000020528 | GSMG0006776 | Prpsap2 | phosphoribosyl pyrophosphate synthetase-associated protein 2 | 168.33 | 218.08 | -0.37 | 1.82E-01 |
| ENSMUSG00000004798 | GSMG0006777 | Ulk2 | unc-51 like kinase 2 | 2264.44 | 2426.53 | -0.10 | 7.10E-01 |
| ENSMUSG00000047804 | GSMG0006781 | Akap10 | A kinase (PRKA) anchor protein 10 | 219.44 | 233.98 | -0.09 | 7.97E-01 |
| ENSMUSG00000014243 | GSMG0006785 | Zswim7 | zinc finger SWIM-type containing 7 | 209.95 | 160.09 | 0.39 | 1.51E-01 |
| ENSMUSG00000018501 | GSMG0006786 | Ncor1 | nuclear receptor co-repressor 1 | 3517.68 | 3825.29 | -0.12 | 6.31E-01 |
| ENSMUSG00000018509 | GSMG0006790 | Cenpv | centromere protein V | 296.47 | 377.02 | -0.35 | 1.76E-01 |
| ENSMUSG00000046417 | GSMG0006794 | Lrrc75a | leucine rich repeat containing 75A | 343.75 | 206.55 | 0.74 | 3.47E-03 |
| ENSMUSG00000005267 | GSMG0006795 | Zfp287 | zinc finger protein 287 | 411.24 | 585.82 | -0.51 | 4.33E-02 |
| ENSMUSG00000047342 | GSMG0006797 | Zfp286 | zinc finger protein 286 | 272.70 | 384.29 | -0.49 | 3.40E-02 |
| ENSMUSG00000042148 | GSMG0006806 | Cox10 | cytochrome c oxidase assembly protein 10 | 660.56 | 786.68 | -0.25 | 2.59E-01 |
| NULL | GSMG0006813 | NULL | NULL | 279.64 | 251.81 | 0.15 | 6.70E-01 |
| ENSMUSG00000033352 | GSMG0006819 | Map2k4 | mitogen-activated protein kinase kinase 4 | 613.25 | 687.90 | -0.17 | 4.66E-01 |
| NULL | GSMG0006823 | NULL | NULL | 313.23 | 216.79 | 0.53 | 2.09E-02 |
| ENSMUSG00000020910 | GSMG0006830 | Adprm | ADP-ribose/CDP-alcohol diphosphatase, manganese dependent | 286.49 | 268.28 | 0.10 | 8.07E-01 |
| ENSMUSG00000020905 | GSMG0006835 | Usp43 | ubiquitin specific peptidase 43 | 269.44 | 478.24 | -0.83 | 2.30E-04 |
| ENSMUSG00000020902 | GSMG0006840 | Ntn1 | netrin 1 | 1371.31 | 1019.78 | 0.43 | 1.89E-01 |
| ENSMUSG00000018736 | GSMG0006846 | Ndel1 | nudE neurodevelopment protein 1 like 1 | 1587.98 | 1529.16 | 0.06 | 8.53E-01 |
| ENSMUSG00000052921 | GSMG0006848 | Arhgef15 | Rho guanine nucleotide exchange factor (GEF) 15 | 1012.75 | 933.01 | 0.12 | 6.58E-01 |

| | | | | | | | |
|---|---|---|---|---|---|---|---|
| ENSMUSG00000032782 | GSMG0006857 | Cntrob | centrobin, centrosomal BRCA2 interacting protein | 307.20 | 433.16 | -0.50 | 2.89E-02 |
| ENSMUSG00000045377 | GSMG0006862 | Tmem88 | transmembrane protein 88 | 2197.08 | 1825.61 | 0.27 | 1.53E-01 |
| ENSMUSG00000018476 | GSMG0006864 | Kdm6b | KDM1 lysine (K)-specific demethylase 6B | 309.53 | 289.04 | 0.10 | 7.74E-01 |
| ENSMUSG00000003934 | GSMG0006868 | Efnb3 | ephrin B3 | 687.52 | 1555.88 | -1.18 | 1.87E-08 |
| ENSMUSG00000041346 | GSMG0006869 | Wrap53 | WD repeat containing, antisense to Trp53 | 385.05 | 419.18 | -0.12 | 6.76E-01 |
| ENSMUSG00000041329 | GSMG0006872 | Atp1b2 | ATPase, Na+/K+ transporting, beta 2 polypeptide | 4789.27 | 3702.67 | 0.37 | 7.68E-02 |
| ENSMUSG00000018761 | GSMG0006874 | Mpdu1 | mannose-P-dolichol utilization defect 1 | 507.00 | 684.97 | -0.43 | 1.80E-02 |
| ENSMUSG00000018774 | GSMG0006875 | Cd68 | CD68 antigen | 270.54 | 156.09 | 0.80 | 8.54E-04 |
| ENSMUSG00000005204 | GSMG0006877 | Senp3 | SUMO/sentrin specific peptidase 3 | 512.82 | 656.95 | -0.36 | 1.42E-01 |
| 52,ENSMUSG00000089669,EI | GSMG0006878 | Tnfsfm13 // Tnfsf1 | tumor necrosis factor (ligand) superfamily, membrane-bound member 1 | 236.25 | 144.83 | 0.71 | 5.45E-03 |
| ENSMUSG00000005198 | GSMG0006879 | Polr2a | polymerase (RNA) II (DNA directed) polypeptide A | 1055.83 | 1105.24 | -0.07 | 8.88E-01 |
| ENSMUSG00000041189 | GSMG0006882 | Chrnb1 | cholinergic receptor, nicotinic, beta polypeptide 1 (muscle) | 441.71 | 286.95 | 0.62 | 5.78E-03 |
| NULL | GSMG0006884 | NULL | NULL | 130.60 | 202.80 | -0.63 | 1.82E-02 |
| ENSMUSG00000051790 | GSMG0006888 | Nlgn2 | neuroligin 2 | 118.30 | 121.82 | -0.04 | 9.41E-01 |
| ENSMUSG00000018570 | GSMG0006892 | 2810408A11Rik | RIKEN cDNA 2810408A11 gene | 58.20 | 109.21 | -0.89 | 1.51E-02 |
| ENSMUSG00000078812 | GSMG0006893 | Eif5a | eukaryotic translation initiation factor 5A | 8757.83 | 12899.40 | -0.56 | 8.15E-04 |
| ENSMUSG00000018566 | GSMG0006894 | Slc2a4 | solute carrier family 2 (facilitated glucose transporter), member 4 | 152.93 | 84.01 | 0.86 | 4.62E-03 |
| ENSMUSG00000018565 | GSMG0006895 | Elp5 | elongator acetyltransferase complex subunit 5 | 1447.84 | 1690.40 | -0.22 | 2.67E-01 |
| ENSMUSG00000018574 | GSMG0006897 | Acadvl | acyl-Coenzyme A dehydrogenase, very long chain | 1938.42 | 1802.98 | 0.10 | 6.78E-01 |
| ENSMUSG00000044367 | GSMG0006898 | Slc16a13 | solute carrier family 16 (monocarboxylic acid transporters), member 13 | 348.75 | 395.03 | -0.18 | 5.18E-01 |
| ENSMUSG00000000317 | GSMG0006899 | Bcl6b | B cell CLL/lymphoma 6, member B | 472.63 | 428.45 | 0.14 | 6.02E-01 |
| G00000020831,ENSMUSG000 | GSMG0006900 | 0610010K14Rik // | RIKEN cDNA 0610010K14 gene // ribonuclease, RNase K | 3272.14 | 3086.17 | 0.08 | 7.38E-01 |
| ENSMUSG00000000320 | GSMG0006901 | Alox12 | arachidonate 12-lipoxygenase | 162.51 | 49.84 | 1.71 | 1.17E-06 |
| ENSMUSG00000018921 | GSMG0006905 | Pelp1 | proline, glutamic acid and leucine rich protein 1 | 352.77 | 442.58 | -0.33 | 3.30E-01 |
| ENSMUSG00000040712 | GSMG0006906 | Camta2 | calmodulin binding transcription activator 2 | 3230.59 | 2673.93 | 0.27 | 2.32E-01 |
| ENSMUSG00000018920 | GSMG0006907 | Cxcl16 | chemokine (C-X-C motif) ligand 16 | 93.05 | 74.02 | 0.34 | 4.94E-01 |
| ENSMUSG00000014606 | GSMG0006911 | Slc25a11 | solute carrier family 25 (mitochondrial carrier oxoglutarate carrier), mer | 1051.03 | 1018.33 | 0.05 | 8.70E-01 |
| ENSMUSG00000018293 | GSMG0006912 | Pfn1 | profilin 1 | 10385.87 | 16127.73 | -0.63 | 4.81E-04 |
| ENSMUSG00000018287 | GSMG0006913 | Spag7 | sperm associated antigen 7 | 1523.54 | 1696.06 | -0.15 | 4.53E-01 |
| ENSMUSG00000040667 | GSMG0006919 | Nup88 | nucleoporin 88 | 2578.97 | 2616.54 | -0.02 | 9.43E-01 |
| ENSMUSG00000018446 | GSMG0006920 | C1qbp | complement component 1, q subcomponent binding protein | 1048.29 | 1778.92 | -0.76 | 2.46E-06 |
| ENSMUSG00000040620 | GSMG0006921 | Dhx33 | DEAH (Asp-Glu-Ala-His) box polypeptide 33 | 311.37 | 308.07 | 0.01 | 9.73E-01 |
| ENSMUSG00000018442 | GSMG0006922 | Derl2 | Der1-like domain family, member 2 | 575.00 | 560.24 | 0.04 | 9.34E-01 |

| | | | | | | | |
|---|---|---|---|---|---|---|---|
| ENSMUSG00000018451 | GSMG0006923 | 6330403K07Rik | RIKEN cDNA 6330403K07 gene | 1076.96 | 1900.27 | -0.82 | 3.52E-07 |
| ENSMUSG00000040543 | GSMG0006931 | Pitpnm3 | PITPNM family member 3 | 305.20 | 107.31 | 1.50 | 1.93E-06 |
| ENSMUSG00000020807 | GSMG0006932 | 4933427D14Rik | RIKEN cDNA 4933427D14 gene | 777.10 | 792.56 | -0.03 | 9.31E-01 |
| ENSMUSG00000020801 | GSMG0006933 | Med31 | mediator complex subunit 31 | 400.66 | 385.35 | 0.06 | 8.78E-01 |
| ENSMUSG00000040447 | GSMG0006938 | Spns2 | spinster homolog 2 | 1437.42 | 1351.23 | 0.09 | 8.37E-01 |
| ENSMUSG00000057778 | GSMG0006941 | Cyb5d2 | cytochrome b5 domain containing 2 | 331.96 | 293.28 | 0.18 | 5.68E-01 |
| ENSMUSG00000050107 | GSMG0006946 | Gsg2 | germ cell-specific gene 2 | 219.00 | 403.30 | -0.88 | 7.75E-06 |
| ENSMUSG00000047260 | GSMG0006947 | Emc6 | ER membrane protein complex subunit 6 | 1402.14 | 1849.09 | -0.40 | 3.85E-02 |
| ENSMUSG00000005949 | GSMG0006948 | Ctns | cystinosis, nephropathic | 463.67 | 419.83 | 0.14 | 5.91E-01 |
| ENSMUSG00000038807 | GSMG0006959 | Rap1gap2 | RAP1 GTPase activating protein 2 | 197.80 | 200.46 | -0.02 | 9.70E-01 |
| NULL | GSMG0006962 | NULL | NULL | 114.24 | 149.59 | -0.38 | 2.92E-01 |
| NULL | GSMG0006963 | NULL | NULL | 393.61 | 316.60 | 0.31 | 3.38E-01 |
| ENSMUSG00000020745 | GSMG0006967 | Pafah1b1 | platelet-activating factor acetylhydrolase, isoform 1b, subunit 1 | 951.00 | 925.82 | 0.04 | 9.03E-01 |
| NULL | GSMG0006968 | NULL | NULL | 507.65 | 419.34 | 0.27 | 4.55E-01 |
| ENSMUSG00000038351 | GSMG0006971 | Sgsm2 | small G protein signaling modulator 2 | 1505.50 | 1553.04 | -0.04 | 9.10E-01 |
| ENSMUSG00000001323 | GSMG0006974 | Srr | serine racemase | 461.00 | 457.99 | 0.01 | 9.74E-01 |
| ENSMUSG00000043099 | GSMG0006978 | Hic1 | hypermethylated in cancer 1 | 1895.82 | 1373.89 | 0.46 | 1.56E-02 |
| G00000038268,ENSMUSG000 | GSMG0006979 | Ovca2 // Dph1 | candidate tumor suppressor in ovarian cancer 2 // diphthamide biosynth | 285.25 | 338.45 | -0.24 | 4.16E-01 |
| ENSMUSG00000000751 | GSMG0006980 | Rpa1 | replication protein A1 | 1702.82 | 2378.59 | -0.48 | 1.66E-02 |
| ENSMUSG00000000753 | GSMG0006982 | Serpinf1 | serine (or cysteine) peptidase inhibitor, clade F, member 1 | 1835.34 | 1430.56 | 0.36 | 1.30E-01 |
| ENSMUSG00000045374 | GSMG0006984 | Wdr81 | WD repeat domain 81 | 221.64 | 209.45 | 0.08 | 8.87E-01 |
| NULL | GSMG0006987 | NULL | NULL | 509.53 | 494.28 | 0.04 | 8.91E-01 |
| NULL | GSMG0006988 | NULL | NULL | 735.43 | 757.38 | -0.04 | 8.86E-01 |
| NULL | GSMG0006990 | NULL | NULL | 755.21 | 744.02 | 0.02 | 9.45E-01 |
| NULL | GSMG0006991 | NULL | NULL | 373.30 | 277.70 | 0.43 | 5.35E-02 |
| ENSMUSG00000020847 | GSMG0006994 | Rph3al | rabphilin 3A-like (without C2 domains) | 146.83 | 111.60 | 0.39 | 2.40E-01 |
| ENSMUSG00000020846 | GSMG0006996 | Fam101b | family with sequence similarity 101, member B | 15334.14 | 14275.56 | 0.10 | 7.00E-01 |
| NULL | GSMG0006999 | NULL | NULL | 360.49 | 337.65 | 0.09 | 7.62E-01 |
| 86,ENSMUSG00000017288,EI | GSMG0007000 | Vps53 // Gemin4 // | VPS53 GARP complex subunit // gem (nuclear organelle) associated pr | 3858.52 | 4083.63 | -0.08 | 7.17E-01 |
| ENSMUSG00000020844 | GSMG0007001 | Nxn | nucleoredoxin | 370.28 | 244.06 | 0.60 | 7.56E-03 |
| NULL | GSMG0007007 | NULL | NULL | 204.19 | 182.37 | 0.16 | 7.09E-01 |
| ENSMUSG00000017631 | GSMG0007008 | Abr | active BCR-related gene | 2137.40 | 2182.44 | -0.03 | 9.32E-01 |
| ENSMUSG00000010392 | GSMG0007013 | Gosr1 | golgi SNAP receptor complex member 1 | 408.95 | 454.61 | -0.15 | 5.74E-01 |

| | | | | | | | |
|---|---|---|---|---|---|---|---|
| ENSMUSG00000020841 | GSMG0007014 | Cpd | carboxypeptidase D | 2352.80 | 2084.92 | 0.17 | 5.20E-01 |
| ENSMUSG00000037958 | GSMG0007018 | Nsrp1 | nuclear speckle regulatory protein 1 | 376.41 | 415.98 | -0.15 | 6.98E-01 |
| ENSMUSG00000037907 | GSMG0007022 | Ankrd13b | ankyrin repeat domain 13b | 469.16 | 857.43 | -0.87 | 1.66E-03 |
| ENSMUSG00000044328 | GSMG0007024 | Trp53i13 | transformation related protein 53 inducible protein 13 | 647.11 | 637.73 | 0.02 | 9.56E-01 |
| ENSMUSG00000017291 | GSMG0007026 | Taok1 | TAO kinase 1 | 701.90 | 706.93 | -0.01 | 9.79E-01 |
| ENSMUSG00000020832 | GSMG0007034 | Eral1 | Era (G-protein)-like 1 (E. coli) | 455.04 | 512.57 | -0.17 | 4.88E-01 |
| ENSMUSG00000017386 | GSMG0007036 | Traf4 | TNF receptor associated factor 4 | 279.33 | 561.92 | -1.01 | 1.89E-07 |
| ENSMUSG00000017405 | GSMG0007037 | Nek8 | NIMA (never in mitosis gene a)-related expressed kinase 8 | 368.11 | 394.74 | -0.10 | 7.58E-01 |
| ENSMUSG00000002052 | GSMG0007040 | Supt6 | suppressor of Ty 6 | 349.16 | 363.88 | -0.06 | 8.84E-01 |
| ENSMUSG00000050132 | GSMG0007045 | Sarm1 | sterile alpha and HEAT/Armadillo motif containing 1 | 294.38 | 610.47 | -1.05 | 1.31E-05 |
| ENSMUSG00000051232 | GSMG0007046 | Tmem199 | transmembrane protein 199 | 1194.32 | 1319.86 | -0.14 | 5.35E-01 |
| ENSMUSG00000017615 | GSMG0007047 | Tnfaip1 | tumor necrosis factor, alpha-induced protein 1 (endothelial) | 4322.51 | 5009.93 | -0.21 | 3.04E-01 |
| ENSMUSG00000037278 | GSMG0007048 | Tmem97 | transmembrane protein 97 | 515.37 | 879.76 | -0.77 | 1.78E-05 |
| ENSMUSG00000017376 | GSMG0007050 | Nlk | nemo like kinase | 344.89 | 300.60 | 0.20 | 4.68E-01 |
| ENSMUSG00000001123 | GSMG0007060 | Lgals9 | lectin, galactose binding, soluble 9 | 2733.36 | 2964.82 | -0.12 | 6.60E-01 |
| ENSMUSG00000017677 | GSMG0007064 | Wsb1 | WD repeat and SOCS box-containing 1 | 3382.77 | 3863.69 | -0.19 | 3.35E-01 |
| ENSMUSG00000049612 | GSMG0007067 | Omg | oligodendrocyte myelin glycoprotein | 990.84 | 417.34 | 1.25 | 1.64E-12 |
| NULL | GSMG0007071 | NULL | NULL | 59.15 | 32.15 | 0.88 | 7.22E-02 |
| ENSMUSG00000035575 | GSMG0007073 | Utp6 | UTP6 small subunit processome component | 744.31 | 797.21 | -0.10 | 7.33E-01 |
| ENSMUSG00000017561 | GSMG0007076 | Crlf3 | cytokine receptor-like factor 3 | 429.36 | 413.49 | 0.05 | 8.77E-01 |
| ENSMUSG00000046909 | GSMG0007077 | Tefm | transcription elongation factor, mitochondrial | 148.50 | 165.86 | -0.16 | 6.88E-01 |
| NULL | GSMG0007078 | NULL | NULL | 162.94 | 106.84 | 0.60 | 4.66E-02 |
| ENSMUSG00000057181 | GSMG0007079 | 5730455P16Rik | RIKEN cDNA 5730455P16 gene | 420.22 | 454.20 | -0.11 | 7.01E-01 |
| NULL | GSMG0007081 | NULL | NULL | 427.51 | 483.03 | -0.18 | 4.87E-01 |
| ENSMUSG00000035441 | GSMG0007084 | Myo1d | myosin ID | 748.18 | 557.32 | 0.43 | 2.32E-02 |
| ENSMUSG00000020704 | GSMG0007087 | Asic2 | acid-sensing (proton-gated) ion channel 2 | 227.07 | 222.18 | 0.03 | 9.51E-01 |
| NULL | GSMG0007089 | NULL | NULL | 526.52 | 306.44 | 0.78 | 1.06E-04 |
| NULL | GSMG0007091 | NULL | NULL | 273.93 | 180.00 | 0.61 | 9.69E-03 |
| NULL | GSMG0007092 | NULL | NULL | 337.37 | 223.41 | 0.59 | 2.04E-02 |
| ENSMUSG00000020696 | GSMG0007098 | Rffl | ring finger and FYVE like domain containing protein | 190.03 | 193.96 | -0.03 | 9.45E-01 |
| ENSMUSG00000018841 | GSMG0007101 | Rad51d | RAD51 paralog D | 117.96 | 147.48 | -0.32 | 3.68E-01 |
| ENSMUSG00000020692 | GSMG0007104 | Nle1 | notchless homolog 1 (Drosophila) | 460.29 | 578.14 | -0.33 | 3.37E-01 |
| NULL | GSMG0007105 | NULL | NULL | 96.01 | 128.78 | -0.42 | 3.12E-01 |

| | | | | | | | |
|---|---|---|---|---|---|---|---|
| ENSMUSG00000086327 | GSMG0007106 | Slfn5os | schlafen 5, opposite strand | 79.50 | 90.12 | -0.19 | 7.02E-01 |
| ENSMUSG00000069793 | GSMG0007108 | Slfn9 | schlafen 9 | 108.64 | 338.20 | -1.63 | 4.52E-09 |
| ENSMUSG00000018733 | GSMG0007115 | Pex12 | peroxisomal biogenesis factor 12 | 134.08 | 149.35 | -0.15 | 7.44E-01 |
| ENSMUSG00000020682 | GSMG0007118 | Mmp28 | matrix metallopeptidase 28 (epilysin) | 196.36 | 201.81 | -0.04 | 9.43E-01 |
| ENSMUSG00000018648 | GSMG0007126 | Dusp14 | dual specificity phosphatase 14 | 394.90 | 102.82 | 1.94 | 6.55E-15 |
| ENSMUSG00000018651 | GSMG0007127 | Tada2a | transcriptional adaptor 2A | 363.25 | 388.73 | -0.10 | 7.55E-01 |
| ENSMUSG00000018697 | GSMG0007130 | Aatf | apoptosis antagonizing transcription factor | 783.52 | 1006.40 | -0.36 | 4.32E-02 |
| ENSMUSG00000018405 | GSMG0007135 | Mrm1 | mitochondrial rRNA methyltransferase 1 | 307.24 | 292.70 | 0.07 | 8.54E-01 |
| ENSMUSG00000034449 | GSMG0007136 | Dhrs11 | dehydrogenase/reductase (SDR family) member 11 | 223.10 | 196.41 | 0.18 | 5.65E-01 |
| ENSMUSG00000020530 | GSMG0007137 | Ggnbp2 | gametogenetin binding protein 2 | 2703.10 | 2634.53 | 0.04 | 8.97E-01 |
| ENSMUSG00000020526 | GSMG0007140 | Znhit3 | zinc finger, HIT type 3 | 446.83 | 524.26 | -0.23 | 3.98E-01 |
| ENSMUSG00000000804 | GSMG0007141 | Usp32 | ubiquitin specific peptidase 32 | 326.13 | 354.16 | -0.12 | 6.94E-01 |
| ENSMUSG00000018481 | GSMG0007142 | Appbp2 | amyloid beta precursor protein (cytoplasmic tail) binding protein 2 | 1020.32 | 1257.69 | -0.30 | 1.37E-01 |
| NULL | GSMG0007150 | NULL | NULL | 81.22 | 51.74 | 0.65 | 1.24E-01 |
| ENSMUSG00000018068 | GSMG0007155 | Ints2 | integrator complex subunit 2 | 1079.30 | 1590.75 | -0.56 | 7.20E-04 |
| ENSMUSG00000034297 | GSMG0007157 | Med13 | mediator complex subunit 13 | 575.05 | 442.61 | 0.38 | 2.09E-01 |
| ENSMUSG00000020516 | GSMG0007162 | Rps6kb1 | ribosomal protein S6 kinase, polypeptide 1 | 403.49 | 484.61 | -0.26 | 3.38E-01 |
| ENSMUSG00000018171 | GSMG0007166 | Vmp1 | vacuole membrane protein 1 | 1515.03 | 1700.30 | -0.17 | 4.54E-01 |
| ENSMUSG00000047126 | GSMG0007170 | Cltc | clathrin, heavy polypeptide (Hc) | 11441.54 | 13165.41 | -0.20 | 2.79E-01 |
| ENSMUSG00000018425 | GSMG0007172 | Dhx40 | DEAH (Asp-Glu-Ala-His) box polypeptide 40 | 1216.35 | 968.75 | 0.33 | 6.19E-02 |
| ENSMUSG00000018427 | GSMG0007175 | Ypel2 | yippee-like 2 (Drosophila) | 696.93 | 307.30 | 1.18 | 1.24E-06 |
| NULL | GSMG0007177 | NULL | NULL | 228.31 | 61.35 | 1.90 | 3.98E-14 |
| ENSMUSG00000061666 | GSMG0007179 | Gdpd1 | glycerophosphodiester phosphodiesterase domain containing 1 | 396.26 | 740.17 | -0.90 | 1.13E-06 |
| ENSMUSG00000020495 | GSMG0007181 | Smg8 | smg-8 homolog, nonsense mediated mRNA decay factor (C. elegans) | 203.97 | 222.26 | -0.13 | 7.28E-01 |
| ENSMUSG00000046442 | GSMG0007185 | Ppm1e | protein phosphatase 1E (PP2C domain containing) | 344.42 | 320.92 | 0.10 | 7.78E-01 |
| NULL | GSMG0007186 | NULL | NULL | 369.50 | 288.23 | 0.36 | 2.75E-01 |
| NULL | GSMG0007189 | NULL | NULL | 443.53 | 233.71 | 0.93 | 3.02E-06 |
| NULL | GSMG0007193 | NULL | NULL | 569.09 | 452.06 | 0.33 | 1.44E-01 |
| ENSMUSG00000020483 | GSMG0007200 | Dynll2 | dynein light chain LC8-type 2 | 698.37 | 740.66 | -0.09 | 8.06E-01 |
| ENSMUSG00000069769 | GSMG0007204 | Msi2 | musashi RNA-binding protein 2 | 133.31 | 127.19 | 0.07 | 8.87E-01 |
| NULL | GSMG0007211 | NULL | NULL | 784.54 | 666.58 | 0.23 | 4.34E-01 |
| ENSMUSG00000018428 | GSMG0007212 | Akap1 | A kinase (PRKA) anchor protein 1 | 372.06 | 432.77 | -0.22 | 3.92E-01 |
| ENSMUSG00000000278 | GSMG0007213 | Scpep1 | serine carboxypeptidase 1 | 1236.12 | 1319.54 | -0.09 | 7.19E-01 |

| | | | | | | | |
|---|---|---|---|---|---|---|---|
| ENSMUSG00000000276 | GSMG0007217 | Dgke | diacylglycerol kinase, epsilon | 175.27 | 198.04 | -0.17 | 6.21E-01 |
| ENSMUSG00000003949 | GSMG0007227 | Hlf | hepatic leukemia factor | 127.54 | 120.54 | 0.08 | 8.72E-01 |
| NULL | GSMG0007228 | NULL | NULL | 246.31 | 223.97 | 0.13 | 7.41E-01 |
| ENSMUSG00000020546 | GSMG0007230 | Stxbp4 | syntaxin binding protein 4 | 315.09 | 314.89 | 0.00 | 9.96E-01 |
| ENSMUSG00000020541 | GSMG0007234 | Tom1l1 | target of myb1-like 1 (chicken) | 178.42 | 170.03 | 0.07 | 8.72E-01 |
| ENSMUSG00000054079 | GSMG0007238 | Utp18 | UTP18 small subunit processome component | 726.64 | 989.37 | -0.45 | 3.30E-02 |
| ENSMUSG00000020857 | GSMG0007239 | Nme2 | NME/NM23 nucleoside diphosphate kinase 2 | 2096.09 | 3016.02 | -0.53 | 7.89E-03 |
| ENSMUSG00000037601 | GSMG0007240 | Nme1 | NME/NM23 nucleoside diphosphate kinase 1 | 2370.93 | 4076.55 | -0.78 | 7.80E-08 |
| NULL | GSMG0007243 | NULL | NULL | 217.06 | 442.14 | -1.03 | 3.40E-06 |
| ENSMUSG00000020863 | GSMG0007244 | Luc7l3 | LUC7-like 3 (S. cerevisiae) | 4716.01 | 4796.26 | -0.02 | 9.47E-01 |
| ENSMUSG00000020866 | GSMG0007246 | Cacna1g | calcium channel, voltage-dependent, T type, alpha 1G subunit | 648.41 | 866.70 | -0.42 | 5.66E-02 |
| ENSMUSG00000039096 | GSMG0007250 | Rsad1 | radical S-adenosyl methionine domain containing 1 | 154.16 | 204.49 | -0.41 | 1.36E-01 |
| ENSMUSG00000076435 | GSMG0007251 | Acsf2 | acyl-CoA synthetase family member 2 | 152.46 | 165.02 | -0.11 | 7.97E-01 |
| ENSMUSG00000020868 | GSMG0007254 | Xylt2 | xylosyltransferase II | 1532.39 | 1199.12 | 0.35 | 9.18E-02 |
| ENSMUSG00000038967 | GSMG0007262 | Pdk2 | pyruvate dehydrogenase kinase, isoenzyme 2 | 718.59 | 555.38 | 0.37 | 6.69E-02 |
| ENSMUSG00000001507 | GSMG0007263 | Itga3 | integrin alpha 3 | 143.99 | 105.32 | 0.45 | 1.89E-01 |
| ENSMUSG00000038909 | GSMG0007266 | Kat7 | K(lysine) acetyltransferase 7 | 2382.27 | 2906.89 | -0.29 | 1.14E-01 |
| ENSMUSG00000014349 | GSMG0007281 | Ube2z | ubiquitin-conjugating enzyme E2Z | 468.45 | 426.96 | 0.13 | 7.28E-01 |
| ENSMUSG00000020876 | GSMG0007289 | Snx11 | sorting nexin 11 | 372.66 | 330.85 | 0.17 | 5.35E-01 |
| ENSMUSG00000038615 | GSMG0007290 | Nfe2l1 | nuclear factor, erythroid derived 2,-like 1 | 4744.31 | 3909.37 | 0.28 | 2.00E-01 |
| ENSMUSG00000018669 | GSMG0007291 | Cdk5rap3 | CDK5 regulatory subunit associated protein 3 | 1813.57 | 2016.53 | -0.15 | 4.66E-01 |
| ENSMUSG00000018659 | GSMG0007293 | Pnpo | pyridoxine 5'-phosphate oxidase | 607.60 | 439.68 | 0.47 | 4.68E-02 |
| ENSMUSG00000018678 | GSMG0007294 | Sp2 | Sp2 transcription factor | 441.23 | 473.28 | -0.10 | 7.87E-01 |
| ENSMUSG00000038517 | GSMG0007301 | Tbkbp1 | TBK1 binding protein 1 | 338.05 | 341.79 | -0.02 | 9.69E-01 |
| ENSMUSG00000001440 | GSMG0007302 | Kpnb1 | karyopherin (importin) beta 1 | 1218.17 | 1549.59 | -0.35 | 1.69E-01 |
| ENSMUSG00000001441 | GSMG0007303 | Npepps | aminopeptidase puromycin sensitive | 902.53 | 995.38 | -0.14 | 5.24E-01 |
| ENSMUSG00000038453 | GSMG0007307 | Srcin1 | SRC kinase signaling inhibitor 1 | 560.89 | 452.62 | 0.31 | 4.44E-01 |
| ENSMUSG00000043439 | GSMG0007308 | E130012A19Rik | RIKEN cDNA E130012A19 gene | 278.54 | 330.81 | -0.25 | 4.14E-01 |
| ENSMUSG00000018537 | GSMG0007310 | Pcgf2 | polycomb group ring finger 2 | 235.05 | 274.36 | -0.22 | 5.21E-01 |
| ENSMUSG00000018541 | GSMG0007312 | Cwc25 | CWC25 spliceosome-associated protein | 362.94 | 371.57 | -0.04 | 9.22E-01 |
| ENSMUSG00000017417 | GSMG0007316 | Plxdc1 | plexin domain containing 1 | 1093.07 | 883.53 | 0.31 | 1.11E-01 |
| ENSMUSG00000038352 | GSMG0007318 | Arl5c | ADP-ribosylation factor-like 5C | 117.52 | 103.27 | 0.18 | 7.30E-01 |
| ENSMUSG00000020882 | GSMG0007319 | Cacnb1 | calcium channel, voltage-dependent, beta 1 subunit | 388.75 | 338.63 | 0.20 | 5.25E-01 |

| | | | | | | | |
|---|---|---|---|---|---|---|---|
| ENSMUSG00000017400 | GSMG0007320 | Stac2 | SH3 and cysteine rich domain 2 | 264.68 | 129.90 | 1.03 | 1.25E-05 |
| ENSMUSG00000020883 | GSMG0007321 | Fbxl20 | F-box and leucine-rich repeat protein 20 | 608.27 | 560.33 | 0.12 | 6.85E-01 |
| NULL | GSMG0007322 | NULL | NULL | 201.54 | 142.42 | 0.50 | 7.63E-02 |
| ENSMUSG00000018160 | GSMG0007323 | Med1 | mediator complex subunit 1 | 481.52 | 617.39 | -0.36 | 2.47E-01 |
| ENSMUSG00000038255 | GSMG0007324 | Neurod2 | neurogenic differentiation 2 | 554.92 | 779.69 | -0.49 | 3.03E-02 |
| ENSMUSG00000038208 | GSMG0007326 | Pgap3 | post-GPI attachment to proteins 3 | 116.99 | 115.18 | 0.03 | 9.63E-01 |
| ENSMUSG00000002580 | GSMG0007327 | Mien1 | migration and invasion enhancer 1 | 1116.10 | 1381.14 | -0.31 | 1.30E-01 |
| ENSMUSG00000038150 | GSMG0007330 | Ormdl3 | ORM1-like 3 (S. cerevisiae) | 3215.28 | 3363.65 | -0.07 | 7.89E-01 |
| ENSMUSG00000017210 | GSMG0007333 | Med24 | mediator complex subunit 24 | 3594.24 | 3603.83 | 0.00 | 9.91E-01 |
| ENSMUSG00000020889 | GSMG0007334 | Nr1d1 | nuclear receptor subfamily 1, group D, member 1 | 975.64 | 581.79 | 0.75 | 2.93E-04 |
| ENSMUSG00000020914 | GSMG0007337 | Top2a | topoisomerase (DNA) II alpha | 312.54 | 804.05 | -1.36 | 8.15E-10 |
| ENSMUSG00000035849 | GSMG0007343 | Krt222 | keratin 222 | 152.00 | 102.04 | 0.57 | 1.03E-01 |
| ENSMUSG00000019761 | GSMG0007350 | Krt10 | keratin 10 | 266.39 | 318.05 | -0.26 | 4.34E-01 |
| ENSMUSG00000006930 | GSMG0007387 | Hap1 | huntingtin-associated protein 1 | 1248.99 | 531.41 | 1.23 | 1.04E-01 |
| ENSMUSG00000001552 | GSMG0007388 | Jup | junction plakoglobin | 4756.19 | 5034.57 | -0.08 | 7.96E-01 |
| ENSMUSG00000006931 | GSMG0007389 | P3h4 | prolyl 3-hydroxylase family member 4 (non-enzymatic) | 550.38 | 625.03 | -0.18 | 4.76E-01 |
| ENSMUSG00000017176 | GSMG0007390 | Nt5c3b | 5'-nucleotidase, cytosolic IIIB | 776.58 | 999.36 | -0.36 | 8.71E-02 |
| ENSMUSG00000048732 | GSMG0007392 | Klhl11 | kelch-like 11 | 827.16 | 785.33 | 0.07 | 8.15E-01 |
| ENSMUSG00000020917 | GSMG0007393 | Acly | ATP citrate lyase | 3407.49 | 5142.36 | -0.59 | 4.87E-04 |
| ENSMUSG00000014195 | GSMG0007394 | Dnajc7 | DnaJ heat shock protein family (Hsp40) member C7 | 1835.39 | 1739.68 | 0.08 | 7.60E-01 |
| ENSMUSG00000020918 | GSMG0007398 | Kat2a | K(lysine) acetyltransferase 2A | 2453.54 | 2669.22 | -0.12 | 6.02E-01 |
| ENSMUSG00000019173 | GSMG0007399 | Rab5c | RAB5C, member RAS oncogene family | 281.04 | 345.14 | -0.29 | 2.92E-01 |
| ENSMUSG00000035355 | GSMG0007401 | Kcnh4 | potassium voltage-gated channel, subfamily H (eag-related), member 4 | 543.42 | 425.14 | 0.35 | 2.02E-01 |
| ENSMUSG00000017747 | GSMG0007403 | Ghdc | GH3 domain containing | 244.21 | 227.97 | 0.10 | 7.92E-01 |
| ENSMUSG00000020919 | GSMG0007404 | Stat5b | signal transducer and activator of transcription 5B | 998.28 | 942.62 | 0.08 | 7.76E-01 |
| ENSMUSG00000004040 | GSMG0007408 | Stat3 | signal transducer and activator of transcription 3 | 3486.47 | 3848.14 | -0.14 | 4.72E-01 |
| ENSMUSG00000004044 | GSMG0007409 | Ptrf | polymerase I and transcript release factor | 7123.02 | 6220.34 | 0.20 | 4.65E-01 |
| ENSMUSG00000019303 | GSMG0007410 | Psmc3ip | proteasome (prosome, macropain) 26S subunit, ATPase 3, interacting p | 67.67 | 196.72 | -1.52 | 1.29E-07 |
| ENSMUSG00000017802 | GSMG0007411 | Fam134c | family with sequence similarity 134, member C | 467.77 | 482.23 | -0.05 | 9.05E-01 |
| ENSMUSG00000035172 | GSMG0007412 | Plekhh3 | pleckstrin homology domain containing, family H (with MyTH4 domai | 1048.20 | 794.69 | 0.40 | 5.20E-02 |
| ENSMUSG00000006920 | GSMG0007414 | Ezh1 | enhancer of zeste 1 polycomb repressive complex 2 subunit | 1498.56 | 1151.37 | 0.38 | 2.59E-02 |
| NULL | GSMG0007415 | NULL | NULL | 497.95 | 482.94 | 0.04 | 9.29E-01 |
| ENSMUSG00000017188 | GSMG0007417 | Coa3 | cytochrome C oxidase assembly factor 3 | 1672.10 | 1912.47 | -0.19 | 3.58E-01 |

| | | | | | | | |
|---|---|---|---|---|---|---|---|
| ENSMUSG00000035086 | GSMG0007418 | Becn1 | beclin 1, autophagy related | 2451.88 | 2426.09 | 0.02 | 9.56E-01 |
| ENSMUSG00000075528 | GSMG0007420 | Aarsd1 | alanyl-tRNA synthetase domain containing 1 | 962.12 | 1104.03 | -0.20 | 3.67E-01 |
| NULL | GSMG0007422 | NULL | NULL | 391.10 | 344.39 | 0.18 | 5.19E-01 |
| ENSMUSG00000034993 | GSMG0007423 | Vat1 | vesicle amine transport 1 | 229.60 | 216.09 | 0.08 | 8.88E-01 |
| ENSMUSG00000017724 | GSMG0007427 | Etv4 | ets variant 4 | 190.03 | 294.54 | -0.63 | 1.28E-02 |
| ENSMUSG00000001493 | GSMG0007428 | Meox1 | mesenchyme homeobox 1 | 1732.57 | 2141.66 | -0.31 | 1.14E-01 |
| ENSMUSG00000003518 | GSMG0007431 | Dusp3 | dual specificity phosphatase 3 (vaccinia virus phosphatase VH1-related | 4592.49 | 3851.59 | 0.25 | 3.42E-01 |
| ENSMUSG00000052373 | GSMG0007433 | Mpp3 | membrane protein, palmitoylated 3 (MAGUK p55 subfamily member 3 | 399.53 | 497.07 | -0.31 | 2.42E-01 |
| ENSMUSG00000017314 | GSMG0007435 | Mpp2 | membrane protein, palmitoylated 2 (MAGUK p55 subfamily member 2 | 496.33 | 305.51 | 0.70 | 2.38E-03 |
| ENSMUSG00000020921 | GSMG0007438 | Tmem101 | transmembrane protein 101 | 606.67 | 723.41 | -0.25 | 2.21E-01 |
| ENSMUSG00000020922 | GSMG0007439 | Lsm12 | LSM12 homolog | 740.08 | 1202.85 | -0.70 | 3.51E-04 |
| ENSMUSG00000008855 | GSMG0007440 | Hdac5 | histone deacetylase 5 | 2378.44 | 2019.27 | 0.24 | 3.60E-01 |
| ENSMUSG00000059995 | GSMG0007441 | Atxn7l3 | ataxin 7-like 3 | 235.91 | 281.94 | -0.26 | 5.49E-01 |
| ENSMUSG00000020923 | GSMG0007443 | Ubtf | upstream binding transcription factor, RNA polymerase I | 2679.29 | 2379.13 | 0.17 | 3.96E-01 |
| NULL | GSMG0007444 | NULL | NULL | 57.65 | 37.51 | 0.61 | 3.26E-01 |
| ENSMUSG00000006574 | GSMG0007445 | Slc4a1 | solute carrier family 4 (anion exchanger), member 1 | 155.06 | 343.91 | -1.16 | 3.30E-06 |
| NULL | GSMG0007446 | NULL | NULL | 65.54 | 63.40 | 0.04 | 9.48E-01 |
| ENSMUSG00000018677 | GSMG0007447 | Slc25a39 | solute carrier family 25, member 39 | 2903.36 | 3194.19 | -0.14 | 5.04E-01 |
| ENSMUSG00000034685 | GSMG0007448 | Fam171a2 | family with sequence similarity 171, member A2 | 722.94 | 1333.52 | -0.88 | 5.34E-04 |
| ENSMUSG00000034621 | GSMG0007450 | Gpatch8 | G patch domain containing 8 | 2903.68 | 2754.25 | 0.08 | 7.42E-01 |
| NULL | GSMG0007451 | NULL | NULL | 534.84 | 439.47 | 0.29 | 2.43E-01 |
| NULL | GSMG0007453 | NULL | NULL | 107.60 | 91.45 | 0.24 | 6.27E-01 |
| NULL | GSMG0007454 | NULL | NULL | 163.21 | 156.61 | 0.06 | 9.13E-01 |
| ENSMUSG00000020925 | GSMG0007460 | Ccdc43 | coiled-coil domain containing 43 | 786.06 | 993.85 | -0.34 | 2.22E-01 |
| ENSMUSG00000034520 | GSMG0007461 | Gjc1 | gap junction protein, gamma 1 | 3764.03 | 4134.73 | -0.14 | 5.53E-01 |
| ENSMUSG00000020929 | GSMG0007462 | Eftud2 | elongation factor Tu GTP binding domain containing 2 | 4383.93 | 4948.05 | -0.17 | 3.89E-01 |
| ENSMUSG00000020932 | GSMG0007463 | Gfap | glial fibrillary acidic protein | 1410.94 | 967.21 | 0.55 | 1.87E-02 |
| ENSMUSG00000051378 | GSMG0007464 | Kif18b | kinesin family member 18B | 77.91 | 207.13 | -1.40 | 6.26E-07 |
| ENSMUSG00000045532 | GSMG0007465 | C1ql1 | complement component 1, q subcomponent-like 1 | 122.80 | 119.98 | 0.04 | 9.42E-01 |
| ENSMUSG00000020935 | GSMG0007467 | Dcakd | dephospho-CoA kinase domain containing | 1181.73 | 1846.41 | -0.64 | 2.65E-05 |
| ENSMUSG00000020937 | GSMG0007468 | Plcd3 | phospholipase C, delta 3 | 306.33 | 271.43 | 0.18 | 5.83E-01 |
| NULL | GSMG0007469 | NULL | NULL | 911.17 | 684.67 | 0.41 | 2.52E-02 |
| ENSMUSG00000034255 | GSMG0007474 | Arhgap27 | Rho GTPase activating protein 27 | 1271.79 | 1470.11 | -0.21 | 3.00E-01 |

| | | | | | | | |
|---|---|---|---|---|---|---|---|
| ENSMUSG00000034247 | GSMG0007475 | Plekhm1 | pleckstrin homology domain containing, family M (with RUN domain) | 891.13 | 679.20 | 0.39 | 1.01E-01 |
| ENSMUSG00000020946 | GSMG0007479 | Gosr2 | golgi SNAP receptor complex member 2 | 6639.86 | 6217.04 | 0.10 | 7.07E-01 |
| ENSMUSG00000034187 | GSMG0007482 | Nsf | N-ethylmaleimide sensitive fusion protein | 2796.09 | 2603.58 | 0.10 | 6.63E-01 |
| NULL | GSMG0007485 | NULL | NULL | 296.14 | 274.88 | 0.11 | 7.56E-01 |
| NULL | GSMG0007488 | NULL | NULL | 181.31 | 217.12 | -0.26 | 4.05E-01 |
| ENSMUSG00000018412 | GSMG0007491 | Kansl1 | KAT8 regulatory NSL complex subunit 1 | 1220.35 | 1266.82 | -0.05 | 8.64E-01 |
| ENSMUSG00000020687 | GSMG0007494 | Cdc27 | cell division cycle 27 | 973.28 | 1074.01 | -0.14 | 5.74E-01 |
| ENSMUSG00000099681 | GSMG0007498 | 1700052K11Rik | RIKEN cDNA 1700052K11 gene | 418.48 | 501.01 | -0.26 | 3.73E-01 |
| ENSMUSG00000019590 | GSMG0007501 | Cyb561 | cytochrome b-561 | 386.39 | 258.87 | 0.58 | 1.34E-02 |
| G00000069631,ENSMUSG000 | GSMG0007507 | Strada // Ccdc47 | STE20-related kinase adaptor alpha // coiled-coil domain containing 47 | 6620.66 | 6511.89 | 0.02 | 9.30E-01 |
| ENSMUSG00000020706 | GSMG0007508 | Ftsj3 | FtsJ RNA methyltransferase homolog 3 (E. coli) | 1261.98 | 1586.96 | -0.33 | 7.96E-02 |
| ENSMUSG00000078619 | GSMG0007509 | Smarcd2 | SWI/SNF related, matrix associated, actin dependent regulator of chron | 1338.45 | 1482.27 | -0.15 | 5.30E-01 |
| ENSMUSG00000001029 | GSMG0007513 | Icam2 | intercellular adhesion molecule 2 | 5292.82 | 3708.40 | 0.51 | 5.12E-03 |
| ENSMUSG00000020715 | GSMG0007516 | Ern1 | endoplasmic reticulum (ER) to nucleus signalling 1 | 126.34 | 118.10 | 0.09 | 8.61E-01 |
| ENSMUSG00000040548 | GSMG0007518 | Tex2 | testis expressed gene 2 | 625.20 | 541.04 | 0.21 | 3.60E-01 |
| ENSMUSG00000020717 | GSMG0007522 | Pecam1 | platelet/endothelial cell adhesion molecule 1 | 10823.02 | 9948.59 | 0.12 | 5.62E-01 |
| ENSMUSG00000020718 | GSMG0007523 | Polg2 | polymerase (DNA directed), gamma 2, accessory subunit | 66.14 | 86.10 | -0.38 | 3.64E-01 |
| ENSMUSG00000020719 | GSMG0007524 | Ddx5 | DEAD (Asp-Glu-Ala-Asp) box polypeptide 5 | 7170.05 | 6661.95 | 0.11 | 7.08E-01 |
| ENSMUSG00000018363 | GSMG0007525 | Smurf2 | SMAD specific E3 ubiquitin protein ligase 2 | 219.73 | 161.96 | 0.44 | 1.00E-01 |
| NULL | GSMG0007527 | NULL | NULL | 190.06 | 217.54 | -0.20 | 5.94E-01 |
| ENSMUSG00000018433 | GSMG0007533 | Nol11 | nucleolar protein 11 | 537.48 | 636.17 | -0.24 | 3.37E-01 |
| ENSMUSG00000040430 | GSMG0007535 | Pitpnc1 | phosphatidylinositol transfer protein, cytoplasmic 1 | 587.23 | 497.91 | 0.24 | 2.90E-01 |
| NULL | GSMG0007541 | NULL | NULL | 754.07 | 617.81 | 0.29 | 2.18E-01 |
| ENSMUSG00000020723 | GSMG0007548 | Cacng4 | calcium channel, voltage-dependent, gamma subunit 4 | 191.45 | 492.28 | -1.36 | 3.53E-02 |
| ENSMUSG00000050965 | GSMG0007552 | Prkca | protein kinase C, alpha | 204.76 | 287.48 | -0.49 | 4.75E-02 |
| NULL | GSMG0007556 | NULL | NULL | 282.30 | 237.24 | 0.25 | 4.22E-01 |
| NULL | GSMG0007560 | NULL | NULL | 322.98 | 357.97 | -0.15 | 6.34E-01 |
| ENSMUSG00000020599 | GSMG0007566 | Rgs9 | regulator of G-protein signaling 9 | 169.58 | 157.31 | 0.10 | 8.43E-01 |
| ENSMUSG00000041920 | GSMG0007568 | Slc16a6 | solute carrier family 16 (monocarboxylic acid transporters), member 6 | 371.89 | 442.84 | -0.25 | 3.25E-01 |
| ENSMUSG00000041895 | GSMG0007569 | Wipi1 | WD repeat domain, phosphoinositide interacting 1 | 722.97 | 589.04 | 0.30 | 1.97E-01 |
| ENSMUSG00000020614 | GSMG0007570 | Fam20a | family with sequence similarity 20, member A | 477.06 | 352.62 | 0.44 | 2.87E-02 |
| ENSMUSG00000041797 | GSMG0007573 | Abca9 | ATP-binding cassette, sub-family A (ABC1), member 9 | 743.90 | 462.94 | 0.68 | 1.02E-04 |
| ENSMUSG00000018800 | GSMG0007579 | Abca5 | ATP-binding cassette, sub-family A (ABC1), member 5 | 434.91 | 558.47 | -0.36 | 1.40E-01 |

| | | | | | | | |
|---|---|---|---|---|---|---|---|
| ENSMUSG00000041654 | GSMG0007594 | Slc39a11 | solute carrier family 39 (metal ion transporter), member 11 | 617.95 | 1131.22 | -0.87 | 1.04E-08 |
| ENSMUSG00000041629 | GSMG0007597 | Fam104a | family with sequence similarity 104, member A | 875.00 | 766.01 | 0.19 | 3.99E-01 |
| ENSMUSG00000041598 | GSMG0007600 | Cdc42ep4 | CDC42 effector protein (Rho GTPase binding) 4 | 5736.23 | 4905.51 | 0.23 | 3.54E-01 |
| ENSMUSG00000041592 | GSMG0007602 | Sdk2 | sidekick cell adhesion molecule 2 | 145.62 | 136.62 | 0.09 | 8.77E-01 |
| NULL | GSMG0007607 | NULL | NULL | 539.81 | 677.51 | -0.33 | 4.00E-01 |
| NULL | GSMG0007608 | NULL | NULL | 321.53 | 410.54 | -0.35 | 3.62E-01 |
| ENSMUSG00000000202 | GSMG0007609 | Btbd17 | BTB (POZ) domain containing 17 | 425.04 | 296.68 | 0.52 | 3.67E-02 |
| ENSMUSG00000015542 | GSMG0007617 | Nat9 | N-acetyltransferase 9 (GCN5-related, putative) | 436.50 | 378.44 | 0.21 | 4.18E-01 |
| ENSMUSG00000020734 | GSMG0007618 | Grin2c | glutamate receptor, ionotropic, NMDA2C (epsilon 3) | 327.42 | 40.78 | 3.01 | 3.10E-20 |
| ENSMUSG00000018861 | GSMG0007619 | Fdxr | ferredoxin reductase | 308.37 | 311.24 | -0.01 | 9.72E-01 |
| ENSMUSG00000034586 | GSMG0007622 | Hid1 | HID1 domain containing | 1932.85 | 2278.98 | -0.24 | 2.73E-01 |
| ENSMUSG00000034566 | GSMG0007625 | Atp5h | ATP synthase, H+ transporting, mitochondrial F0 complex, subunit D | 1501.92 | 1695.06 | -0.18 | 5.35E-01 |
| ENSMUSG00000020736 | GSMG0007626 | Nt5c | 5',3'-nucleotidase, cytosolic | 1330.12 | 1465.95 | -0.14 | 5.47E-01 |
| ENSMUSG00000020737 | GSMG0007627 | Hn1 | hematological and neurological expressed sequence 1 | 2402.12 | 6773.31 | -1.50 | 2.83E-25 |
| NULL | GSMG0007631 | NULL | NULL | 136.33 | 140.73 | -0.04 | 9.44E-01 |
| ENSMUSG00000020740 | GSMG0007632 | Gga3 | golgi associated, gamma adaptin ear containing, ARF binding protein 3 | 756.10 | 785.79 | -0.06 | 8.60E-01 |
| ENSMUSG00000059923 | GSMG0007634 | Grb2 | growth factor receptor bound protein 2 | 850.31 | 967.86 | -0.19 | 4.07E-01 |
| NULL | GSMG0007636 | NULL | NULL | 126.18 | 77.24 | 0.71 | 4.62E-02 |
| NULL | GSMG0007637 | NULL | NULL | 777.09 | 566.73 | 0.46 | 3.11E-02 |
| ENSMUSG00000034471 | GSMG0007639 | Caskin2 | CASK-interacting protein 2 | 1886.53 | 1727.10 | 0.13 | 7.00E-01 |
| ENSMUSG00000020752 | GSMG0007641 | Recql5 | RecQ protein-like 5 | 299.25 | 236.79 | 0.34 | 2.80E-01 |
| NULL | GSMG0007644 | NULL | NULL | 92.27 | 88.35 | 0.06 | 9.08E-01 |
| ENSMUSG00000020766 | GSMG0007647 | Galk1 | galactokinase 1 | 794.95 | 965.00 | -0.28 | 3.02E-01 |
| ENSMUSG00000016559 | GSMG0007648 | H3f3b | H3 histone, family 3B | 18080.81 | 23315.01 | -0.37 | 1.03E-01 |
| ENSMUSG00000034341 | GSMG0007650 | Wbp2 | WW domain binding protein 2 | 2621.33 | 2374.66 | 0.14 | 4.74E-01 |
| ENSMUSG00000020773 | GSMG0007651 | Trim47 | tripartite motif-containing 47 | 968.07 | 471.03 | 1.04 | 2.15E-11 |
| ENSMUSG00000020775 | GSMG0007653 | Mrpl38 | mitochondrial ribosomal protein L38 | 1020.65 | 1016.30 | 0.00 | 9.88E-01 |
| ENSMUSG00000020776 | GSMG0007654 | Fbf1 | Fas (TNFRSF6) binding factor 1 | 580.04 | 665.03 | -0.20 | 5.28E-01 |
| ENSMUSG00000020777 | GSMG0007655 | Acox1 | acyl-Coenzyme A oxidase 1, palmitoyl | 3262.33 | 3698.32 | -0.18 | 3.68E-01 |
| ENSMUSG00000057286 | GSMG0007656 | St6galnac2 | ST6 (alpha-N-acetyl-neuraminyl-2,3-beta-galactosyl-1,3)-N-acetylgalac | 1125.01 | 338.19 | 1.73 | 4.15E-20 |
| ENSMUSG00000020780 | GSMG0007658 | Srp68 | signal recognition particle 68 | 506.40 | 560.96 | -0.15 | 5.61E-01 |
| ENSMUSG00000020792 | GSMG0007660 | Exoc7 | exocyst complex component 7 | 2129.52 | 2425.89 | -0.19 | 3.25E-01 |
| ENSMUSG00000015869 | GSMG0007665 | Prpsap1 | phosphoribosyl pyrophosphate synthetase-associated protein 1 | 3316.69 | 3556.57 | -0.10 | 6.62E-01 |

| | | | | | | | |
|---|---|---|---|---|---|---|---|
| NULL | GSMG0007666 | NULL | NULL | 372.19 | 401.68 | -0.11 | 7.52E-01 |
| ENSMUSG00000020802 | GSMG0007668 | Ube2o | ubiquitin-conjugating enzyme E2O | 1089.99 | 1014.47 | 0.10 | 7.82E-01 |
| ENSMUSG00000020806 | GSMG0007670 | Rhbdf2 | rhomboid 5 homolog 2 | 1561.10 | 1078.00 | 0.53 | 9.90E-03 |
| ENSMUSG00000020810 | GSMG0007671 | Cygb | cytoglobin | 188.22 | 126.42 | 0.57 | 5.59E-02 |
| ENSMUSG00000020814 | GSMG0007675 | Mxra7 | matrix-remodelling associated 7 | 1163.24 | 803.82 | 0.53 | 2.16E-03 |
| ENSMUSG00000056962 | GSMG0007676 | Jmjd6 | jumonji domain containing 6 | 1054.88 | 1049.33 | 0.01 | 9.80E-01 |
| ENSMUSG00000034120 | GSMG0007677 | Srsf2 | serine/arginine-rich splicing factor 2 | 11141.01 | 13856.09 | -0.31 | 1.19E-01 |
| NULL | GSMG0007683 | NULL | NULL | 370.91 | 307.42 | 0.27 | 3.36E-01 |
| ENSMUSG00000025572 | GSMG0007687 | Tmc6 | transmembrane channel-like gene family 6 | 337.02 | 291.03 | 0.21 | 4.47E-01 |
| ENSMUSG00000025574 | GSMG0007688 | Tk1 | thymidine kinase 1 | 308.96 | 825.36 | -1.41 | 6.83E-14 |
| ENSMUSG00000017713 | GSMG0007689 | Tha1 | threonine aldolase 1 | 97.08 | 80.47 | 0.26 | 5.89E-01 |
| ENSMUSG00000053113 | GSMG0007690 | Socs3 | suppressor of cytokine signaling 3 | 239.90 | 179.24 | 0.42 | 1.65E-01 |
| ENSMUSG00000017132 | GSMG0007693 | Cyth1 | cytohesin 1 | 1419.13 | 1280.10 | 0.15 | 5.40E-01 |
| ENSMUSG00000033909 | GSMG0007695 | Usp36 | ubiquitin specific peptidase 36 | 520.78 | 495.14 | 0.07 | 8.04E-01 |
| ENSMUSG00000017466 | GSMG0007696 | Timp2 | tissue inhibitor of metalloproteinase 2 | 5794.08 | 7182.89 | -0.31 | 1.29E-01 |
| ENSMUSG00000033880 | GSMG0007698 | Lgals3bp | lectin, galactoside-binding, soluble, 3 binding protein | 638.03 | 405.26 | 0.66 | 1.16E-03 |
| ENSMUSG00000025575 | GSMG0007699 | Cant1 | calcium activated nucleotidase 1 | 765.00 | 820.79 | -0.10 | 7.21E-01 |
| ENSMUSG00000025576 | GSMG0007703 | Rbfox3 | RNA binding protein, fox-1 homolog (C. elegans) 3 | 459.37 | 671.23 | -0.55 | 1.38E-02 |
| NULL | GSMG0007704 | NULL | NULL | 358.55 | 202.71 | 0.82 | 3.11E-03 |
| NULL | GSMG0007705 | NULL | NULL | 491.07 | 314.70 | 0.64 | 3.60E-03 |
| ENSMUSG00000025578 | GSMG0007706 | Cbx8 | chromobox 8 | 281.43 | 152.24 | 0.88 | 2.44E-04 |
| ENSMUSG00000039976 | GSMG0007710 | Tbc1d16 | TBC1 domain family, member 16 | 262.73 | 583.77 | -1.15 | 5.08E-06 |
| ENSMUSG00000025580 | GSMG0007712 | Eif4a3 | eukaryotic translation initiation factor 4A3 | 1974.47 | 2481.62 | -0.33 | 4.87E-02 |
| ENSMUSG00000005043 | GSMG0007713 | Sgsh | N-sulfoglucosamine sulfohydrolase (sulfamidase) | 134.25 | 137.96 | -0.04 | 9.48E-01 |
| ENSMUSG00000025582 | GSMG0007717 | Nptx1 | neuronal pentraxin 1 | 127.12 | 38.62 | 1.72 | 7.18E-05 |
| NULL | GSMG0007719 | NULL | NULL | 131.18 | 184.30 | -0.49 | 8.16E-02 |
| NULL | GSMG0007720 | NULL | NULL | 199.79 | 185.32 | 0.10 | 8.24E-01 |
| ENSMUSG00000025375 | GSMG0007722 | Aatk | apoptosis-associated tyrosine kinase | 2671.87 | 2335.03 | 0.19 | 5.66E-01 |
| ENSMUSG00000039781 | GSMG0007724 | Cep131 | centrosomal protein 131 | 1506.36 | 1641.20 | -0.12 | 7.12E-01 |
| ENSMUSG00000025377 | GSMG0007725 | Enthd2 | ENTH domain containing 2 | 351.01 | 330.41 | 0.09 | 7.90E-01 |
| ENSMUSG00000061306 | GSMG0007726 | Slc38a10 | solute carrier family 38, member 10 | 1574.25 | 2003.01 | -0.35 | 7.56E-02 |
| ENSMUSG00000043993 | GSMG0007729 | 2900052L18Rik | RIKEN cDNA 2900052L18 gene | 544.77 | 680.76 | -0.32 | 2.30E-01 |
| ENSMUSG00000025384 | GSMG0007731 | Faap100 | Fanconi anemia core complex associated protein 100 | 563.82 | 946.07 | -0.75 | 2.14E-05 |

| | | | | | | | |
|---|---|---|---|---|---|---|---|
| ENSMUSG00000039703 | GSMG0007732 | Nploc4 | NPL4 homolog, ubiquitin recognition factor | 1227.72 | 1280.32 | -0.06 | 8.63E-01 |
| ENSMUSG00000039670 | GSMG0007735 | Oxld1 | oxidoreductase like domain containing 1 | 184.91 | 120.82 | 0.61 | 2.74E-02 |
| ENSMUSG00000057594 | GSMG0007736 | Arl16 | ADP-ribosylation factor-like 16 | 821.99 | 733.10 | 0.17 | 5.41E-01 |
| ENSMUSG00000061111 | GSMG0007738 | Fam195b | family with sequence similarity 195, member B | 3653.24 | 4013.95 | -0.14 | 5.04E-01 |
| ENSMUSG00000025130 | GSMG0007740 | P4hb | prolyl 4-hydroxylase, beta polypeptide | 8244.72 | 10104.85 | -0.29 | 1.00E-01 |
| ENSMUSG00000025132 | GSMG0007742 | Arhgdia | Rho GDP dissociation inhibitor (GDI) alpha | 6574.82 | 7470.17 | -0.18 | 3.30E-01 |
| ENSMUSG00000025134 | GSMG0007744 | Alyref | Aly/REF export factor | 1550.23 | 2361.32 | -0.61 | 1.89E-04 |
| ENSMUSG00000025137 | GSMG0007745 | Pcyt2 | phosphate cytidylyltransferase 2, ethanolamine | 3017.81 | 2184.35 | 0.47 | 1.44E-02 |
| ENSMUSG00000025138 | GSMG0007746 | Sirt7 | sirtuin 7 | 1810.24 | 1370.95 | 0.40 | 2.03E-02 |
| ENSMUSG00000051510 | GSMG0007747 | Mafg | v-maf musculoaponeurotic fibrosarcoma oncogene family, protein G (av | 2696.20 | 1910.55 | 0.50 | 3.90E-03 |
| ENSMUSG00000025140 | GSMG0007748 | Pycr1 | pyrroline-5-carboxylate reductase 1 | 67.53 | 92.57 | -0.45 | 2.61E-01 |
| ENSMUSG00000042988 | GSMG0007750 | Notum | notum pectinacetylesterase homolog (Drosophila) | 2034.22 | 1391.83 | 0.55 | 2.43E-03 |
| ENSMUSG00000025144 | GSMG0007752 | Stra13 | stimulated by retinoic acid 13 | 745.30 | 854.50 | -0.20 | 4.60E-01 |
| ENSMUSG00000039450 | GSMG0007753 | Dcxr | dicarbonyl L-xylulose reductase | 150.11 | 93.85 | 0.68 | 3.47E-02 |
| ENSMUSG00000025150 | GSMG0007754 | Cbr2 | carbonyl reductase 2 | 356.74 | 24.14 | 3.89 | 6.57E-45 |
| ENSMUSG00000025158 | GSMG0007755 | Rfng | RFNG O-fucosylpeptide 3-beta-N-acetylglucosaminyltransferase | 1437.93 | 1470.73 | -0.03 | 9.18E-01 |
| ENSMUSG00000025155 | GSMG0007756 | Dus1l | dihydrouridine synthase 1-like (S. cerevisiae) | 1304.17 | 1204.00 | 0.12 | 6.23E-01 |
| ENSMUSG00000025153 | GSMG0007757 | Fasn | fatty acid synthase | 329.99 | 687.83 | -1.06 | 1.20E-06 |
| ENSMUSG00000048445 | GSMG0007758 | Ccdc57 | coiled-coil domain containing 57 | 153.16 | 166.30 | -0.12 | 7.78E-01 |
| ENSMUSG00000025162 | GSMG0007759 | Csnk1d | casein kinase 1, delta | 2164.43 | 2485.74 | -0.20 | 3.12E-01 |
| ENSMUSG00000025169 | GSMG0007765 | Ogfod3 | 2-oxoglutarate and iron-dependent oxygenase domain containing 3 | 316.85 | 356.96 | -0.17 | 5.89E-01 |
| ENSMUSG00000025173 | GSMG0007767 | Wdr45b | WD repeat domain 45B | 1054.01 | 1204.70 | -0.19 | 3.63E-01 |
| ENSMUSG00000025170 | GSMG0007768 | Rab40b | Rab40B, member RAS oncogene family | 291.56 | 134.51 | 1.11 | 4.40E-06 |
| ENSMUSG00000079179 | GSMG0007771 | Rab10os | RAB10, member RAS oncogene family, opposite strand | 1085.50 | 852.03 | 0.35 | 1.62E-01 |
| ENSMUSG00000046605 | GSMG0007772 | B3gntl1 | UDP-GlcNAc:betaGal beta-1,3-N-acetylglucosaminyltransferase-like 1 | 180.42 | 178.31 | 0.02 | 9.68E-01 |
| ENSMUSG00000020668 | GSMG0007773 | Kif3c | kinesin family member 3C | 256.61 | 356.50 | -0.48 | 4.84E-02 |
| ENSMUSG00000037486 | GSMG0007774 | Asxl2 | additional sex combs like 2 (Drosophila) | 179.18 | 198.00 | -0.15 | 6.82E-01 |
| ENSMUSG00000071454 | GSMG0007778 | Dtnb | dystrobrevin, beta | 535.75 | 560.64 | -0.07 | 8.42E-01 |
| NULL | GSMG0007781 | NULL | NULL | 130.41 | 139.38 | -0.10 | 8.35E-01 |
| ENSMUSG00000020661 | GSMG0007782 | Dnmt3a | DNA methyltransferase 3A | 1084.39 | 949.35 | 0.19 | 6.03E-01 |
| NULL | GSMG0007783 | NULL | NULL | 168.03 | 77.52 | 1.12 | 1.83E-04 |
| ENSMUSG00000020660 | GSMG0007784 | Pomc | pro-opiomelanocortin-alpha | 551.61 | 206.68 | 1.42 | 2.79E-11 |
| ENSMUSG00000020657 | GSMG0007785 | Dnajc27 | DnaJ heat shock protein family (Hsp40) member C27 | 225.85 | 162.25 | 0.48 | 7.94E-02 |

| | | | | | | | |
|---|---|---|---|---|---|---|---|
| ENSMUSG00000020654 | GSMG0007786 | Adcy3 | adenylate cyclase 3 | 1046.95 | 1095.31 | -0.07 | 8.54E-01 |
| ENSMUSG00000096199 | GSMG0007787 | Ptrhd1 | peptidyl-tRNA hydrolase domain containing 1 | 426.51 | 519.02 | -0.28 | 2.32E-01 |
| ENSMUSG00000020640 | GSMG0007793 | Itsn2 | intersectin 2 | 1217.67 | 1238.66 | -0.03 | 9.31E-01 |
| ENSMUSG00000037361 | GSMG0007795 | Sf3b6 | splicing factor 3B, subunit 6 | 1282.31 | 1599.52 | -0.32 | 1.22E-01 |
| ENSMUSG00000037735 | GSMG0007805 | 2810032G03Rik | RIKEN cDNA 2810032G03 gene | 207.40 | 315.14 | -0.61 | 6.68E-02 |
| ENSMUSG00000037669 | GSMG0007816 | Ldah | lipid droplet associated hydrolase | 210.07 | 240.60 | -0.19 | 6.12E-01 |
| ENSMUSG00000020605 | GSMG0007819 | Hs1bp3 | HCLS1 binding protein 3 | 78.95 | 83.47 | -0.08 | 8.82E-01 |
| NULL | GSMG0007821 | NULL | NULL | 226.10 | 178.60 | 0.34 | 2.84E-01 |
| NULL | GSMG0007822 | NULL | NULL | 239.12 | 153.36 | 0.64 | 4.12E-02 |
| NULL | GSMG0007823 | NULL | NULL | 336.31 | 269.72 | 0.32 | 3.52E-01 |
| ENSMUSG00000020594 | GSMG0007825 | Pum2 | pumilio RNA-binding family member 2 | 461.91 | 620.67 | -0.43 | 4.64E-02 |
| ENSMUSG00000020592 | GSMG0007826 | Sdc1 | syndecan 1 | 219.54 | 211.84 | 0.05 | 9.17E-01 |
| ENSMUSG00000020585 | GSMG0007828 | Laptm4a | lysosomal-associated protein transmembrane 4A | 7154.06 | 6543.30 | 0.13 | 5.89E-01 |
| ENSMUSG00000066643 | GSMG0007831 | Wdr35 | WD repeat domain 35 | 157.47 | 177.70 | -0.18 | 6.84E-01 |
| ENSMUSG00000066637 | GSMG0007832 | Ttc32 | tetratricopeptide repeat domain 32 | 202.12 | 196.70 | 0.04 | 9.45E-01 |
| ENSMUSG00000048387 | GSMG0007834 | Osr1 | odd-skipped related 1 (Drosophila) | 58.08 | 86.99 | -0.58 | 1.71E-01 |
| ENSMUSG00000020621 | GSMG0007836 | Rdh14 | retinol dehydrogenase 14 (all-trans and 9-cis) | 892.34 | 956.62 | -0.10 | 6.96E-01 |
| ENSMUSG00000020608 | GSMG0007839 | Smc6 | structural maintenance of chromosomes 6 | 545.46 | 688.63 | -0.34 | 1.77E-01 |
| ENSMUSG00000020589 | GSMG0007842 | Fam49a | family with sequence similarity 49, member A | 386.21 | 742.21 | -0.94 | 5.83E-08 |
| ENSMUSG00000020591 | GSMG0007856 | Ntsr2 | neurotensin receptor 2 | 4070.85 | 381.43 | 3.42 | 3.45E-95 |
| ENSMUSG00000057469 | GSMG0007857 | E2f6 | E2F transcription factor 6 | 356.41 | 385.28 | -0.11 | 7.28E-01 |
| ENSMUSG00000020580 | GSMG0007858 | Rock2 | Rho-associated coiled-coil containing protein kinase 2 | 3069.96 | 2586.00 | 0.25 | 3.20E-01 |
| ENSMUSG00000020571 | GSMG0007861 | Pdia6 | protein disulfide isomerase associated 6 | 5276.91 | 8441.36 | -0.68 | 2.47E-05 |
| ENSMUSG00000061458 | GSMG0007862 | Nol10 | nucleolar protein 10 | 452.88 | 632.43 | -0.48 | 1.11E-02 |
| ENSMUSG00000011179 | GSMG0007863 | Odc1 | ornithine decarboxylase, structural 1 | 1219.12 | 2216.70 | -0.86 | 1.67E-06 |
| ENSMUSG00000071379 | GSMG0007864 | Hpcal1 | hippocalcin-like 1 | 342.21 | 258.58 | 0.40 | 1.56E-01 |
| NULL | GSMG0007867 | NULL | NULL | 1017.72 | 910.88 | 0.16 | 4.84E-01 |
| NULL | GSMG0007868 | NULL | NULL | 387.01 | 439.95 | -0.19 | 5.74E-01 |
| NULL | GSMG0007870 | NULL | NULL | 334.06 | 488.22 | -0.55 | 7.03E-03 |
| ENSMUSG00000052632 | GSMG0007879 | Asap2 | ArfGAP with SH3 domain, ankyrin repeat and PH domain 2 | 427.31 | 656.37 | -0.62 | 1.44E-03 |
| ENSMUSG00000054309 | GSMG0007880 | Cpsf3 | cleavage and polyadenylation specificity factor 3 | 1558.98 | 1900.94 | -0.29 | 1.36E-01 |
| ENSMUSG00000062054 | GSMG0007881 | Iah1 | isoamyl acetate-hydrolyzing esterase 1 homolog | 439.43 | 423.13 | 0.06 | 8.76E-01 |
| ENSMUSG00000059669 | GSMG0007889 | Taf1b | TATA-box binding protein associated factor, RNA polymerase I, B | 737.17 | 849.15 | -0.20 | 3.61E-01 |

| | | | | | | | |
|---|---|---|---|---|---|---|---|
| NULL | GSMG0007890 | NULL | NULL | 287.84 | 315.24 | -0.13 | 6.93E-01 |
| ENSMUSG00000020653 | GSMG0007893 | Klf11 | Kruppel-like factor 11 | 797.76 | 792.51 | 0.01 | 9.79E-01 |
| ENSMUSG00000020649 | GSMG0007894 | Rrm2 | ribonucleotide reductase M2 | 565.30 | 1392.01 | -1.30 | 1.87E-10 |
| ENSMUSG00000020646 | GSMG0007896 | Mboat2 | membrane bound O-acyltransferase domain containing 2 | 385.94 | 548.74 | -0.50 | 1.82E-02 |
| NULL | GSMG0007897 | NULL | NULL | 116.99 | 50.10 | 1.23 | 1.94E-04 |
| NULL | GSMG0007900 | NULL | NULL | 496.50 | 510.14 | -0.04 | 9.07E-01 |
| ENSMUSG00000036333 | GSMG0007901 | Kidins220 | kinase D-interacting substrate 220 | 520.84 | 627.68 | -0.27 | 2.74E-01 |
| ENSMUSG00000020638 | GSMG0007909 | Cmpk2 | cytidine monophosphate (UMP-CMP) kinase 2, mitochondrial | 363.60 | 265.93 | 0.45 | 1.28E-01 |
| NULL | GSMG0007911 | NULL | NULL | 101.75 | 371.66 | -1.88 | 4.74E-12 |
| ENSMUSG00000020630 | GSMG0007913 | Rnaseh1 | ribonuclease H1 | 202.38 | 229.27 | -0.18 | 5.88E-01 |
| ENSMUSG00000020629 | GSMG0007914 | Adi1 | acireductone dioxygenase 1 | 915.58 | 771.35 | 0.25 | 2.31E-01 |
| ENSMUSG00000036613 | GSMG0007916 | Tssc1 | tumor suppressing subtransferable candidate 1 | 373.33 | 425.27 | -0.19 | 5.37E-01 |
| NULL | GSMG0007917 | NULL | NULL | 117.29 | 86.18 | 0.45 | 2.05E-01 |
| ENSMUSG00000061911 | GSMG0007921 | Myt1l | myelin transcription factor 1-like | 2279.80 | 3078.12 | -0.43 | 1.37E-02 |
| NULL | GSMG0007922 | NULL | NULL | 233.86 | 184.46 | 0.34 | 2.33E-01 |
| NULL | GSMG0007925 | NULL | NULL | 243.96 | 251.21 | -0.04 | 9.30E-01 |
| NULL | GSMG0007926 | NULL | NULL | 294.35 | 335.95 | -0.19 | 5.58E-01 |
| ENSMUSG00000020674 | GSMG0007928 | Pxdn | peroxidasin | 2503.02 | 4979.89 | -0.99 | 1.73E-09 |
| ENSMUSG00000043061 | GSMG0007929 | Tmem18 | transmembrane protein 18 | 1736.56 | 1666.79 | 0.06 | 8.19E-01 |
| ENSMUSG00000020669 | GSMG0007932 | Sh3yl1 | Sh3 domain YSC-like 1 | 142.07 | 265.87 | -0.91 | 1.25E-03 |
| ENSMUSG00000002900 | GSMG0007935 | Lamb1 | laminin B1 | 4903.90 | 9291.33 | -0.92 | 3.64E-11 |
| ENSMUSG00000035933 | GSMG0007939 | Cog5 | component of oligomeric golgi complex 5 | 824.22 | 901.11 | -0.13 | 6.44E-01 |
| NULL | GSMG0007945 | NULL | NULL | 843.58 | 786.10 | 0.10 | 7.48E-01 |
| ENSMUSG00000020572 | GSMG0007946 | Nampt | nicotinamide phosphoribosyltransferase | 5369.31 | 7078.55 | -0.40 | 1.50E-02 |
| ENSMUSG00000020570 | GSMG0007949 | Sypl | synaptophysin-like protein | 1975.42 | 1895.83 | 0.06 | 8.31E-01 |
| ENSMUSG00000020564 | GSMG0007950 | Atxn7l1 | ataxin 7-like 1 | 190.71 | 231.06 | -0.28 | 3.59E-01 |
| ENSMUSG00000020561 | GSMG0007958 | Twistnb | twist basic helix-loop-helix transcription factor 1 neighbor | 724.75 | 854.49 | -0.24 | 2.89E-01 |
| ENSMUSG00000020590 | GSMG0007964 | Snx13 | sorting nexin 13 | 408.77 | 413.62 | -0.02 | 9.65E-01 |
| ENSMUSG00000036188 | GSMG0007975 | Ankmy2 | ankyrin repeat and MYND domain containing 2 | 429.31 | 462.48 | -0.11 | 7.20E-01 |
| ENSMUSG00000043153 | GSMG0007978 | Ispd | isoprenoid synthase domain containing | 117.39 | 124.24 | -0.07 | 8.93E-01 |
| ENSMUSG00000050103 | GSMG0007986 | Agmo | alkylglycerol monooxygenase | 65.68 | 68.40 | -0.05 | 9.43E-01 |
| ENSMUSG00000036095 | GSMG0007990 | Dgkb | diacylglycerol kinase, beta | 587.40 | 235.87 | 1.32 | 1.71E-12 |
| ENSMUSG00000035954 | GSMG0008008 | Dock4 | dedicator of cytokinesis 4 | 2781.29 | 2883.41 | -0.05 | 8.71E-01 |

| | | | | | | | |
|---|---|---|---|---|---|---|---|
| NULL | GSMG0008009 | NULL | NULL | 805.40 | 740.46 | 0.12 | 7.37E-01 |
| NULL | GSMG0008011 | NULL | NULL | 524.20 | 391.16 | 0.42 | 7.56E-02 |
| NULL | GSMG0008015 | NULL | NULL | 236.10 | 171.96 | 0.46 | 2.19E-01 |
| ENSMUSG00000036257 | GSMG0008032 | Pnpla8 | patatin-like phospholipase domain containing 8 | 977.25 | 842.96 | 0.21 | 3.18E-01 |
| ENSMUSG00000020598 | GSMG0008034 | Nrcam | neuronal cell adhesion molecule | 3404.90 | 3930.24 | -0.21 | 2.49E-01 |
| NULL | GSMG0008035 | NULL | NULL | 196.51 | 163.09 | 0.27 | 4.38E-01 |
| NULL | GSMG0008041 | NULL | NULL | 260.71 | 588.45 | -1.17 | 8.79E-09 |
| ENSMUSG00000035293 | GSMG0008046 | G2e3 | G2/M-phase specific E3 ubiquitin ligase | 255.69 | 363.24 | -0.50 | 4.12E-02 |
| ENSMUSG00000020952 | GSMG0008047 | Scfd1 | Sec1 family domain containing 1 | 953.55 | 1049.23 | -0.14 | 6.34E-01 |
| ENSMUSG00000020953 | GSMG0008048 | Coch | cochlin | 680.65 | 410.42 | 0.73 | 2.76E-04 |
| ENSMUSG00000020955 | GSMG0008049 | Ap4s1 | adaptor-related protein complex AP-4, sigma 1 | 364.68 | 369.73 | -0.02 | 9.59E-01 |
| NULL | GSMG0008051 | NULL | NULL | 523.41 | 440.23 | 0.25 | 3.23E-01 |
| ENSMUSG00000035142 | GSMG0008054 | Nubpl | nucleotide binding protein-like | 225.66 | 219.72 | 0.04 | 9.34E-01 |
| ENSMUSG00000035133 | GSMG0008056 | Arhgap5 | Rho GTPase activating protein 5 | 2885.05 | 2503.33 | 0.20 | 3.00E-01 |
| ENSMUSG00000061603 | GSMG0008058 | Akap6 | A kinase (PRKA) anchor protein 6 | 640.06 | 697.49 | -0.12 | 7.41E-01 |
| ENSMUSG00000021010 | GSMG0008064 | Npas3 | neuronal PAS domain protein 3 | 157.33 | 254.96 | -0.70 | 4.64E-03 |
| ENSMUSG00000021023 | GSMG0008081 | 1110008L16Rik | RIKEN cDNA 1110008L16 gene | 126.57 | 161.21 | -0.35 | 2.97E-01 |
| ENSMUSG00000021024 | GSMG0008083 | Psma6 | proteasome (prosome, macropain) subunit, alpha type 6 | 2811.45 | 3625.69 | -0.37 | 3.20E-02 |
| NULL | GSMG0008086 | NULL | NULL | 92.40 | 85.97 | 0.09 | 8.74E-01 |
| ENSMUSG00000012076 | GSMG0008087 | Brms1l | breast cancer metastasis-suppressor 1-like | 470.41 | 443.93 | 0.09 | 7.85E-01 |
| G00000047022,ENSMUSG000 | GSMG0008092 | Prps1l3 // Mipol1 | phosphoribosyl pyrophosphate synthetase 1-like 3 // mirror-image polyd | 122.40 | 125.48 | -0.03 | 9.46E-01 |
| ENSMUSG00000060121 | GSMG0008102 | Gemin2 | gem (nuclear organelle) associated protein 2 | 164.61 | 230.71 | -0.49 | 6.94E-02 |
| ENSMUSG00000020994 | GSMG0008103 | Pnn | pinin | 2920.09 | 3077.03 | -0.08 | 7.75E-01 |
| ENSMUSG00000035653 | GSMG0008106 | Lrfn5 | leucine rich repeat and fibronectin type III domain containing 5 | 476.69 | 902.84 | -0.92 | 3.47E-08 |
| ENSMUSG00000035614 | GSMG0008111 | Fam179b | family with sequence similarity 179, member B | 793.86 | 804.33 | -0.02 | 9.52E-01 |
| ENSMUSG00000035597 | GSMG0008113 | Prpf39 | pre-mRNA processing factor 39 | 1194.48 | 1049.58 | 0.19 | 4.74E-01 |
| ENSMUSG00000055884 | GSMG0008114 | Fancm | Fanconi anemia, complementation group M | 174.15 | 217.62 | -0.32 | 2.76E-01 |
| NULL | GSMG0008122 | NULL | NULL | 268.08 | 290.26 | -0.12 | 7.52E-01 |
| ENSMUSG00000097061 | GSMG0008123 | 9330151L19Rik | RIKEN cDNA 9330151L19 gene | 1121.22 | 827.14 | 0.44 | 4.01E-02 |
| ENSMUSG00000020978 | GSMG0008126 | Klhdc2 | kelch domain containing 2 | 868.80 | 1138.06 | -0.39 | 3.40E-02 |
| ENSMUSG00000044147 | GSMG0008128 | Arf6 | ADP-ribosylation factor 6 | 3337.24 | 3619.08 | -0.12 | 6.42E-01 |
| ENSMUSG00000054894 | GSMG0008131 | Atp5s | ATP synthase, H+ transporting, mitochondrial F0 complex, subunit S | 202.07 | 190.92 | 0.08 | 8.38E-01 |
| ENSMUSG00000021072 | GSMG0008138 | Tmx1 | thioredoxin-related transmembrane protein 1 | 818.14 | 1015.04 | -0.31 | 1.50E-01 |

| | | | | | | | |
|---|---|---|---|---|---|---|---|
| ENSMUSG00000048285 | GSMG0008139 | Frmd6 | FERM domain containing 6 | 836.96 | 1129.39 | -0.43 | 2.23E-02 |
| ENSMUSG00000021076 | GSMG0008142 | Actr10 | ARP10 actin-related protein 10 | 1557.71 | 1739.90 | -0.16 | 4.62E-01 |
| ENSMUSG00000048118 | GSMG0008147 | Arid4a | AT rich interactive domain 4A (RBP1-like) | 534.85 | 436.77 | 0.29 | 1.84E-01 |
| ENSMUSG00000034601 | GSMG0008149 | 2700049A03Rik | RIKEN cDNA 2700049A03 gene | 91.83 | 101.11 | -0.13 | 8.03E-01 |
| ENSMUSG00000044548 | GSMG0008151 | Dact1 | dishevelled-binding antagonist of beta-catenin 1 | 691.05 | 816.79 | -0.24 | 3.19E-01 |
| ENSMUSG00000034574 | GSMG0008155 | Daam1 | dishevelled associated activator of morphogenesis 1 | 322.05 | 337.79 | -0.07 | 8.47E-01 |
| NULL | GSMG0008156 | NULL | NULL | 50.50 | 49.83 | 0.01 | 9.90E-01 |
| NULL | GSMG0008158 | NULL | NULL | 333.63 | 363.14 | -0.12 | 7.25E-01 |
| ENSMUSG00000005078 | GSMG0008160 | Jkamp | JNK1/MAPK8-associated membrane protein | 1203.62 | 1167.87 | 0.04 | 8.86E-01 |
| ENSMUSG00000034501 | GSMG0008165 | Pcnx4 | pecanex homolog 4 | 679.23 | 660.41 | 0.04 | 9.01E-01 |
| ENSMUSG00000021096 | GSMG0008167 | Ppm1a | protein phosphatase 1A, magnesium dependent, alpha isoform | 1314.41 | 1282.60 | 0.03 | 9.22E-01 |
| ENSMUSG00000021103 | GSMG0008173 | Mnat1 | menage a trois 1 | 372.68 | 415.96 | -0.16 | 6.20E-01 |
| ENSMUSG00000044712 | GSMG0008178 | Slc38a6 | solute carrier family 38, member 6 | 173.58 | 188.64 | -0.12 | 7.83E-01 |
| ENSMUSG00000021108 | GSMG0008182 | Prkch | protein kinase C, eta | 1740.84 | 1436.64 | 0.28 | 1.52E-01 |
| ENSMUSG00000021109 | GSMG0008186 | Hif1a | hypoxia inducible factor 1, alpha subunit | 2361.10 | 2694.62 | -0.19 | 3.25E-01 |
| ENSMUSG00000021113 | GSMG0008188 | Snapc1 | small nuclear RNA activating complex, polypeptide 1 | 306.40 | 275.68 | 0.15 | 6.81E-01 |
| NULL | GSMG0008189 | NULL | NULL | 233.51 | 381.46 | -0.71 | 1.22E-03 |
| ENSMUSG00000044912 | GSMG0008190 | Syt16 | synaptotagmin XVI | 368.03 | 667.93 | -0.86 | 3.90E-06 |
| ENSMUSG00000071265 | GSMG0008192 | 1700086L19Rik | RIKEN cDNA 1700086L19 gene | 190.62 | 234.75 | -0.30 | 3.33E-01 |
| ENSMUSG00000046768 | GSMG0008196 | Rhoj | ras homolog family member J | 6211.16 | 8248.88 | -0.41 | 1.48E-02 |
| NULL | GSMG0008201 | NULL | NULL | 57.47 | 81.21 | -0.50 | 2.42E-01 |
| ENSMUSG00000021048 | GSMG0008211 | Mthfd1 | methylenetetrahydrofolate dehydrogenase (NADP+ dependent), methen | 1013.89 | 1625.25 | -0.68 | 1.69E-05 |
| ENSMUSG00000033454 | GSMG0008214 | Zbtb1 | zinc finger and BTB domain containing 1 | 1375.98 | 1271.10 | 0.11 | 6.43E-01 |
| NULL | GSMG0008215 | NULL | NULL | 167.28 | 144.87 | 0.20 | 6.28E-01 |
| ENSMUSG00000059970 | GSMG0008216 | Hspa2 | heat shock protein 2 | 1037.08 | 541.05 | 0.94 | 2.06E-07 |
| ENSMUSG00000052609 | GSMG0008219 | Plekhg3 | pleckstrin homology domain containing, family G (with RhoGef domai | 136.64 | 70.60 | 0.95 | 6.73E-03 |
| ENSMUSG00000090258 | GSMG0008221 | Churc1 | churchill domain containing 1 | 739.67 | 590.63 | 0.32 | 9.25E-02 |
| ENSMUSG00000033373 | GSMG0008222 | Fntb | farnesyltransferase, CAAX box, beta | 264.55 | 376.63 | -0.51 | 2.75E-02 |
| ENSMUSG00000021065 | GSMG0008224 | Fut8 | fucosyltransferase 8 | 2307.88 | 1932.01 | 0.26 | 2.10E-01 |
| ENSMUSG00000047454 | GSMG0008229 | Gphn | gephyrin | 1149.32 | 1346.54 | -0.23 | 2.30E-01 |
| NULL | GSMG0008230 | NULL | NULL | 195.61 | 221.35 | -0.18 | 7.12E-01 |
| ENSMUSG00000021112 | GSMG0008242 | Mpp5 | membrane protein, palmitoylated 5 (MAGUK p55 subfamily member 5 | 585.46 | 537.85 | 0.12 | 7.00E-01 |
| ENSMUSG00000021116 | GSMG0008243 | Eif2s1 | eukaryotic translation initiation factor 2, subunit 1 alpha | 924.73 | 1192.19 | -0.37 | 1.07E-01 |

| | | | | | | | |
|---|---|---|---|---|---|---|---|
| ENSMUSG00000060716 | GSMG0008245 | Plekhh1 | pleckstrin homology domain containing, family H (with MyTH4 domai | 215.73 | 488.51 | -1.18 | 2.04E-07 |
| ENSMUSG00000032705 | GSMG0008262 | Exd2 | exonuclease 3'-5' domain containing 2 | 2665.08 | 1820.56 | 0.55 | 1.73E-04 |
| ENSMUSG00000021130 | GSMG0008264 | Galnt16 | UDP-N-acetyl-alpha-D-galactosamine:polypeptide N-acetylgalactosami | 173.03 | 154.15 | 0.17 | 7.07E-01 |
| ENSMUSG00000048833 | GSMG0008265 | Slc39a9 | solute carrier family 39 (zinc transporter), member 9 | 292.90 | 334.62 | -0.19 | 5.35E-01 |
| ENSMUSG00000021133 | GSMG0008268 | Susd6 | sushi domain containing 6 | 743.85 | 604.16 | 0.30 | 1.94E-01 |
| ENSMUSG00000021134 | GSMG0008273 | Srsf5 | serine/arginine-rich splicing factor 5 | 7190.48 | 6861.44 | 0.07 | 8.34E-01 |
| ENSMUSG00000021140 | GSMG0008280 | Pcnx | pecanex homolog (Drosophila) | 1654.02 | 1617.33 | 0.03 | 9.39E-01 |
| ENSMUSG00000042700 | GSMG0008282 | Sipa1l1 | signal-induced proliferation-associated 1 like 1 | 1516.80 | 1302.49 | 0.22 | 3.59E-01 |
| ENSMUSG00000021219 | GSMG0008285 | Rgs6 | regulator of G-protein signaling 6 | 420.71 | 503.22 | -0.26 | 3.68E-01 |
| NULL | GSMG0008291 | NULL | NULL | 411.65 | 353.78 | 0.22 | 4.41E-01 |
| NULL | GSMG0008293 | NULL | NULL | 314.89 | 366.33 | -0.21 | 4.96E-01 |
| ENSMUSG00000021222 | GSMG0008295 | Dcaf4 | DDB1 and CUL4 associated factor 4 | 373.22 | 572.48 | -0.62 | 1.31E-03 |
| ENSMUSG00000010608 | GSMG0008296 | Rbm25 | RNA binding motif protein 25 | 5990.24 | 6323.62 | -0.08 | 8.09E-01 |
| ENSMUSG00000019969 | GSMG0008301 | Psen1 | presenilin 1 | 1241.93 | 1340.73 | -0.11 | 7.07E-01 |
| ENSMUSG00000021223 | GSMG0008303 | Papln | papilin, proteoglycan-like sulfated glycoprotein | 363.67 | 233.76 | 0.64 | 4.20E-02 |
| ENSMUSG00000046791 | GSMG0008304 | 2410016O06Rik | RIKEN cDNA 2410016O06 gene | 213.73 | 244.12 | -0.19 | 5.49E-01 |
| ENSMUSG00000072946 | GSMG0008313 | Ptgr2 | prostaglandin reductase 2 | 734.54 | 604.12 | 0.28 | 2.95E-01 |
| ENSMUSG00000042472 | GSMG0008315 | Zfp410 | zinc finger protein 410 | 965.85 | 1033.98 | -0.10 | 7.65E-01 |
| ENSMUSG00000021235 | GSMG0008316 | Coq6 | coenzyme Q6 monooxygenase | 672.97 | 679.05 | -0.01 | 9.70E-01 |
| ENSMUSG00000057265 | GSMG0008317 | Bbof1 | basal body orientation factor 1 | 176.26 | 132.24 | 0.41 | 2.16E-01 |
| NULL | GSMG0008318 | NULL | NULL | 482.95 | 431.23 | 0.16 | 5.92E-01 |
| ENSMUSG00000085793 | GSMG0008319 | Lin52 | lin-52 homolog (C. elegans) | 424.20 | 388.39 | 0.13 | 6.51E-01 |
| ENSMUSG00000021241 | GSMG0008324 | Isca2 | iron-sulfur cluster assembly 2 | 1639.84 | 1475.81 | 0.15 | 5.25E-01 |
| ENSMUSG00000021243 | GSMG0008326 | Fcf1 | FCF1 rRNA processing protein | 507.96 | 558.90 | -0.14 | 5.93E-01 |
| ENSMUSG00000021244 | GSMG0008327 | Ylpm1 | YLP motif containing 1 | 2904.58 | 3301.33 | -0.18 | 4.16E-01 |
| ENSMUSG00000004789 | GSMG0008330 | Dlst | dihydrolipoamide S-succinyltransferase (E2 component of 2-oxo-glutar | 1503.97 | 1589.92 | -0.08 | 7.66E-01 |
| ENSMUSG00000004788 | GSMG0008331 | Eif2b2 | eukaryotic translation initiation factor 2B, subunit 2 beta | 843.81 | 1037.37 | -0.30 | 1.17E-01 |
| NULL | GSMG0008333 | NULL | NULL | 59.39 | 85.95 | -0.54 | 1.89E-01 |
| ENSMUSG00000021250 | GSMG0008335 | Fos | FBJ osteosarcoma oncogene | 727.82 | 77.92 | 3.22 | 1.07E-39 |
| ENSMUSG00000034271 | GSMG0008336 | Jdp2 | Jun dimerization protein 2 | 172.95 | 101.06 | 0.77 | 4.61E-03 |
| ENSMUSG00000034258 | GSMG0008338 | Mfsd7c | major facilitator superfamily domain containing 7C | 3203.31 | 3879.62 | -0.28 | 1.18E-01 |
| ENSMUSG00000012609 | GSMG0008340 | Ttll5 | tubulin tyrosine ligase-like family, member 5 | 111.49 | 107.97 | 0.05 | 9.28E-01 |
| ENSMUSG00000007867 | GSMG0008343 | Ift43 | intraflagellar transport 43 | 673.09 | 579.19 | 0.22 | 4.15E-01 |

| | | | | | | | |
|---|---|---|---|---|---|---|---|
| ENSMUSG00000021254 | GSMG0008344 | Gpatch2l | G patch domain containing 2 like | 419.06 | 493.26 | -0.24 | 3.66E-01 |
| NULL | GSMG0008345 | NULL | NULL | 272.14 | 323.12 | -0.25 | 3.30E-01 |
| ENSMUSG00000021256 | GSMG0008348 | Vash1 | vasohibin 1 | 124.10 | 112.85 | 0.13 | 7.93E-01 |
| ENSMUSG00000034157 | GSMG0008351 | Cipc | CLOCK interacting protein, circadian | 1423.59 | 1244.16 | 0.19 | 3.36E-01 |
| ENSMUSG00000034145 | GSMG0008352 | Tmem63c | transmembrane protein 63c | 339.54 | 357.09 | -0.07 | 8.71E-01 |
| ENSMUSG00000021033 | GSMG0008353 | Gstz1 | glutathione transferase zeta 1 (maleylacetoacetate isomerase) | 349.22 | 398.82 | -0.19 | 4.55E-01 |
| ENSMUSG00000021037 | GSMG0008356 | Ahsa1 | AHA1, activator of heat shock protein ATPase 1 | 1323.29 | 1431.33 | -0.11 | 6.47E-01 |
| ENSMUSG00000021040 | GSMG0008357 | Slirp | SRA stem-loop interacting RNA binding protein | 688.20 | 804.47 | -0.23 | 3.71E-01 |
| ENSMUSG00000021044 | GSMG0008365 | Adck1 | aarF domain containing kinase 1 | 270.62 | 247.41 | 0.13 | 7.71E-01 |
| ENSMUSG00000066392 | GSMG0008369 | Nrxn3 | neurexin III | 2367.82 | 1900.23 | 0.32 | 1.35E-01 |
| NULL | GSMG0008371 | NULL | NULL | 540.58 | 649.00 | -0.26 | 5.10E-01 |
| NULL | GSMG0008380 | NULL | NULL | 1155.66 | 704.78 | 0.71 | 2.14E-04 |
| NULL | GSMG0008381 | NULL | NULL | 793.61 | 498.38 | 0.67 | 2.27E-04 |
| NULL | GSMG0008382 | NULL | NULL | 939.35 | 627.49 | 0.58 | 5.97E-04 |
| NULL | GSMG0008384 | NULL | NULL | 363.40 | 243.82 | 0.58 | 1.26E-02 |
| NULL | GSMG0008387 | NULL | NULL | 766.39 | 498.61 | 0.62 | 4.62E-04 |
| NULL | GSMG0008388 | NULL | NULL | 213.16 | 143.44 | 0.58 | 1.64E-01 |
| NULL | GSMG0008389 | NULL | NULL | 986.29 | 595.56 | 0.73 | 1.09E-05 |
| NULL | GSMG0008394 | NULL | NULL | 228.79 | 144.38 | 0.67 | 1.82E-02 |
| NULL | GSMG0008399 | NULL | NULL | 252.36 | 180.23 | 0.48 | 4.80E-02 |
| NULL | GSMG0008402 | NULL | NULL | 465.86 | 255.13 | 0.87 | 6.06E-05 |
| ENSMUSG00000047414 | GSMG0008411 | Flrt2 | fibronectin leucine rich transmembrane protein 2 | 3753.04 | 2799.36 | 0.42 | 9.90E-03 |
| NULL | GSMG0008413 | NULL | NULL | 353.10 | 303.49 | 0.22 | 4.57E-01 |
| ENSMUSG00000021007 | GSMG0008418 | Spata7 | spermatogenesis associated 7 | 153.20 | 140.74 | 0.12 | 7.83E-01 |
| ENSMUSG00000021012 | GSMG0008420 | Zc3h14 | zinc finger CCCH type containing 14 | 4020.48 | 4734.02 | -0.24 | 1.81E-01 |
| ENSMUSG00000021013 | GSMG0008427 | Ttc8 | tetratricopeptide repeat domain 8 | 446.61 | 365.04 | 0.29 | 2.28E-01 |
| ENSMUSG00000021177 | GSMG0008435 | Tdp1 | tyrosyl-DNA phosphodiesterase 1 | 513.39 | 748.51 | -0.54 | 1.08E-02 |
| ENSMUSG00000021178 | GSMG0008437 | Psmc1 | protease (prosome, macropain) 26S subunit, ATPase 1 | 4039.16 | 4998.74 | -0.31 | 1.03E-01 |
| ENSMUSG00000001175 | GSMG0008440 | Calm1 | calmodulin 1 | 42121.26 | 45069.57 | -0.10 | 7.37E-01 |
| ENSMUSG00000041781 | GSMG0008450 | Cpsf2 | cleavage and polyadenylation specific factor 2 | 407.98 | 600.11 | -0.55 | 4.98E-03 |
| ENSMUSG00000041771 | GSMG0008451 | Slc24a4 | solute carrier family 24 (sodium/potassium/calcium exchanger), membe | 66.46 | 20.93 | 1.66 | 2.69E-03 |
| NULL | GSMG0008452 | NULL | NULL | 808.08 | 270.14 | 1.58 | 9.89E-10 |
| ENSMUSG00000044456 | GSMG0008453 | Rin3 | Ras and Rab interactor 3 | 749.94 | 839.07 | -0.16 | 5.90E-01 |

| | | | | | | | |
|---|---|---|---|---|---|---|---|
| ENSMUSG00000021192 | GSMG0008458 | Golga5 | golgi autoantigen, golgin subfamily a, 5 | 1764.22 | 1708.76 | 0.05 | 8.72E-01 |
| ENSMUSG00000021194 | GSMG0008459 | Chga | chromogranin A | 1130.30 | 1137.33 | -0.01 | 9.80E-01 |
| ENSMUSG00000046675 | GSMG0008460 | Tmem251 | transmembrane protein 251 | 339.17 | 286.00 | 0.25 | 3.34E-01 |
| ENSMUSG00000041712 | GSMG0008461 | Ubr7 | ubiquitin protein ligase E3 component n-recognin 7 (putative) | 496.65 | 602.51 | -0.28 | 2.16E-01 |
| ENSMUSG00000021198 | GSMG0008463 | Unc79 | unc-79 homolog (C. elegans) | 699.99 | 583.19 | 0.26 | 3.08E-01 |
| NULL | GSMG0008464 | NULL | NULL | 133.18 | 136.70 | -0.04 | 9.45E-01 |
| ENSMUSG00000064215 | GSMG0008468 | Ifi27 | interferon, alpha-inducible protein 27 | 729.67 | 517.79 | 0.49 | 2.15E-02 |
| NULL | GSMG0008484 | NULL | NULL | 61.24 | 43.41 | 0.49 | 3.61E-01 |
| ENSMUSG00000021102 | GSMG0008486 | Glrx5 | glutaredoxin 5 | 2020.38 | 2534.81 | -0.33 | 5.52E-02 |
| ENSMUSG00000094910 | GSMG0008494 | D430019H16Rik | RIKEN cDNA D430019H16 gene | 288.14 | 697.35 | -1.27 | 1.54E-12 |
| ENSMUSG00000044715 | GSMG0008498 | Gskip | GSK3B interacting protein | 987.72 | 842.26 | 0.23 | 3.09E-01 |
| ENSMUSG00000021111 | GSMG0008500 | Papola | poly (A) polymerase alpha | 872.81 | 1098.34 | -0.33 | 1.99E-01 |
| ENSMUSG00000021115 | GSMG0008504 | Vrk1 | vaccinia related kinase 1 | 554.46 | 782.41 | -0.50 | 1.40E-02 |
| ENSMUSG00000021258 | GSMG0008514 | Ccnk | cyclin K | 125.75 | 98.95 | 0.33 | 4.75E-01 |
| NULL | GSMG0008515 | NULL | NULL | 161.70 | 67.11 | 1.26 | 1.68E-05 |
| ENSMUSG00000021259 | GSMG0008517 | Cyp46a1 | cytochrome P450, family 46, subfamily a, polypeptide 1 | 4406.77 | 2358.14 | 0.90 | 1.97E-08 |
| ENSMUSG00000058070 | GSMG0008518 | Eml1 | echinoderm microtubule associated protein like 1 | 2297.30 | 1949.33 | 0.24 | 1.78E-01 |
| ENSMUSG00000021262 | GSMG0008519 | Evl | Ena-vasodilator stimulated phosphoprotein | 1410.90 | 2070.69 | -0.55 | 1.31E-02 |
| NULL | GSMG0008521 | NULL | NULL | 716.09 | 929.01 | -0.37 | 1.08E-01 |
| ENSMUSG00000021264 | GSMG0008522 | Yy1 | YY1 transcription factor | 272.10 | 290.06 | -0.10 | 8.20E-01 |
| NULL | GSMG0008523 | NULL | NULL | 270.81 | 200.64 | 0.44 | 1.16E-01 |
| ENSMUSG00000097451 | GSMG0008537 | Rian | RNA imprinted and accumulated in nucleus | 1371.84 | 1374.05 | 0.00 | 9.96E-01 |
| NULL | GSMG0008541 | NULL | NULL | 470.52 | 419.64 | 0.16 | 5.53E-01 |
| ENSMUSG00000097391 | GSMG0008542 | Mirg | miRNA containing gene | 437.18 | 278.23 | 0.65 | 1.66E-03 |
| ENSMUSG00000017843 | GSMG0008544 | Ppp2r5c | protein phosphatase 2, regulatory subunit B', gamma | 1683.32 | 2183.33 | -0.38 | 6.83E-02 |
| ENSMUSG00000018707 | GSMG0008553 | Dync1h1 | dynein cytoplasmic 1 heavy chain 1 | 2560.81 | 2914.57 | -0.19 | 3.32E-01 |
| ENSMUSG00000037957 | GSMG0008555 | Wdr20 | WD repeat domain 20 | 792.71 | 762.12 | 0.06 | 8.54E-01 |
| ENSMUSG00000021271 | GSMG0008558 | Zfp839 | zinc finger protein 839 | 326.21 | 218.62 | 0.58 | 4.32E-02 |
| ENSMUSG00000021275 | GSMG0008560 | Tecpr2 | tectonin beta-propeller repeat containing 2 | 480.25 | 352.50 | 0.45 | 1.65E-01 |
| ENSMUSG00000037896 | GSMG0008563 | Rcor1 | REST corepressor 1 | 529.92 | 437.65 | 0.27 | 4.26E-01 |
| ENSMUSG00000021277 | GSMG0008569 | Traf3 | TNF receptor-associated factor 3 | 182.26 | 193.19 | -0.09 | 8.63E-01 |
| ENSMUSG00000021278 | GSMG0008571 | Amn | amnionless | 129.38 | 195.11 | -0.59 | 1.12E-01 |
| ENSMUSG00000021282 | GSMG0008576 | Eif5 | eukaryotic translation initiation factor 5 | 2365.33 | 3010.65 | -0.35 | 5.61E-02 |

| | | | | | | | |
|---|---|---|---|---|---|---|---|
| ENSMUSG00000007411 | GSMG0008577 | Mark3 | MAP/microtubule affinity regulating kinase 3 | 5785.20 | 5391.14 | 0.10 | 6.84E-01 |
| ENSMUSG00000060950 | GSMG0008578 | Trmt61a | tRNA methyltransferase 61A | 130.14 | 166.47 | -0.36 | 2.53E-01 |
| ENSMUSG00000037787 | GSMG0008579 | Apopt1 | apoptogenic, mitochondrial 1 | 950.21 | 901.66 | 0.08 | 7.98E-01 |
| ENSMUSG00000021288 | GSMG0008580 | Klc1 | kinesin light chain 1 | 4758.96 | 5670.36 | -0.25 | 1.58E-01 |
| NULL | GSMG0008581 | NULL | NULL | 257.38 | 218.81 | 0.23 | 5.87E-01 |
| ENSMUSG00000021286 | GSMG0008582 | Zfyve21 | zinc finger, FYVE domain containing 21 | 568.82 | 496.24 | 0.20 | 4.46E-01 |
| ENSMUSG00000021294 | GSMG0008587 | Kif26a | kinesin family member 26A | 1219.41 | 1160.35 | 0.07 | 8.79E-01 |
| ENSMUSG00000043122 | GSMG0008588 | A530016L24Rik | RIKEN cDNA A530016L24 gene | 211.85 | 278.60 | -0.40 | 2.46E-01 |
| ENSMUSG00000037679 | GSMG0008589 | Inf2 | inverted formin, FH2 and WH2 domain containing | 337.33 | 212.73 | 0.66 | 1.46E-03 |
| ENSMUSG00000064326 | GSMG0008591 | Siva1 | SIVA1, apoptosis-inducing factor | 403.60 | 508.78 | -0.33 | 1.65E-01 |
| ENSMUSG00000072825 | GSMG0008593 | Cep170b | centrosomal protein 170B | 1499.69 | 1388.61 | 0.11 | 8.99E-01 |
| ENSMUSG00000052160 | GSMG0008594 | Pld4 | phospholipase D family, member 4 | 276.60 | 133.35 | 1.06 | 3.79E-05 |
| ENSMUSG00000037594 | GSMG0008595 | BC022687 | cDNA sequence BC022687 | 289.49 | 392.74 | -0.44 | 4.41E-02 |
| ENSMUSG00000002803 | GSMG0008597 | Btbd6 | BTB (POZ) domain containing 6 | 120.64 | 115.49 | 0.06 | 9.09E-01 |
| ENSMUSG00000021143 | GSMG0008599 | Pacs2 | phosphofurin acidic cluster sorting protein 2 | 302.88 | 279.07 | 0.11 | 8.25E-01 |
| ENSMUSG00000021144 | GSMG0008602 | Mta1 | metastasis associated 1 | 1086.50 | 671.74 | 0.69 | 3.40E-01 |
| ENSMUSG00000006356 | GSMG0008603 | Crip2 | cysteine rich protein 2 | 6992.67 | 5453.72 | 0.36 | 3.64E-02 |
| ENSMUSG00000006360 | GSMG0008604 | Crip1 | cysteine-rich protein 1 (intestinal) | 1887.64 | 543.88 | 1.79 | 1.98E-19 |
| ENSMUSG00000037466 | GSMG0008605 | 4930427A07Rik | RIKEN cDNA 4930427A07 gene | 71.34 | 148.50 | -1.05 | 3.47E-04 |
| ENSMUSG00000049036 | GSMG0008606 | Tmem121 | transmembrane protein 121 | 243.78 | 484.14 | -0.99 | 3.71E-05 |
| ENSMUSG00000042063 | GSMG0008615 | Zfp386 | zinc finger protein 386 (Kruppel-like) | 1292.76 | 1253.45 | 0.04 | 8.73E-01 |
| NULL | GSMG0008616 | NULL | NULL | 37.66 | 227.74 | -2.57 | 7.00E-20 |
| ENSMUSG00000021171 | GSMG0008618 | Esyt2 | extended synaptotagmin-like protein 2 | 2573.95 | 1785.36 | 0.53 | 1.84E-03 |
| NULL | GSMG0008620 | NULL | NULL | 103.35 | 206.58 | -1.00 | 4.79E-04 |
| ENSMUSG00000042029 | GSMG0008621 | Ncapg2 | non-SMC condensin II complex, subunit G2 | 257.04 | 470.56 | -0.87 | 1.99E-05 |
| ENSMUSG00000056553 | GSMG0008622 | Ptprn2 | protein tyrosine phosphatase, receptor type, N polypeptide 2 | 235.11 | 373.76 | -0.67 | 6.31E-03 |
| ENSMUSG00000041992 | GSMG0008628 | Rapgef5 | Rap guanine nucleotide exchange factor (GEF) 5 | 1294.79 | 1683.76 | -0.38 | 8.53E-02 |
| ENSMUSG00000021175 | GSMG0008634 | Cdca7l | cell division cycle associated 7 like | 187.35 | 259.87 | -0.47 | 1.12E-01 |
| NULL | GSMG0008640 | NULL | NULL | 443.03 | 378.41 | 0.23 | 3.82E-01 |
| ENSMUSG00000020671 | GSMG0008643 | Rab10 | RAB10, member RAS oncogene family | 4666.20 | 5975.02 | -0.36 | 5.23E-02 |
| ENSMUSG00000020658 | GSMG0008654 | Efr3b | EFR3 homolog B | 2687.99 | 2188.83 | 0.30 | 2.05E-01 |
| NULL | GSMG0008655 | NULL | NULL | 230.23 | 146.02 | 0.65 | 1.24E-02 |
| NULL | GSMG0008656 | NULL | NULL | 66.31 | 79.42 | -0.26 | 6.29E-01 |

| | | | | | | | |
|---|---|---|---|---|---|---|---|
| ENSMUSG00000020652 | GSMG0008657 | Cenpo | centromere protein O | 83.07 | 92.78 | -0.16 | 7.37E-01 |
| ENSMUSG00000020647 | GSMG0008659 | Ncoa1 | nuclear receptor coactivator 1 | 2889.05 | 2053.98 | 0.49 | 5.91E-02 |
| ENSMUSG00000020635 | GSMG0008667 | Fkbp1b | FK506 binding protein 1b | 199.36 | 510.58 | -1.35 | 1.26E-12 |
| ENSMUSG00000020634 | GSMG0008669 | Ubxn2a | UBX domain protein 2A | 444.13 | 417.78 | 0.09 | 8.15E-01 |
| NULL | GSMG0008671 | NULL | NULL | 96.07 | 96.25 | 0.00 | 9.99E-01 |
| ENSMUSG00000020627 | GSMG0008672 | Klhl29 | kelch-like 29 | 245.38 | 263.94 | -0.11 | 8.42E-01 |
| NULL | GSMG0008673 | NULL | NULL | 183.05 | 199.01 | -0.12 | 7.71E-01 |
| NULL | GSMG0008676 | NULL | NULL | 265.26 | 281.66 | -0.09 | 8.04E-01 |
| NULL | GSMG0008678 | NULL | NULL | 724.08 | 588.27 | 0.30 | 1.81E-01 |
| ENSMUSG00000054364 | GSMG0008686 | Rhob | ras homolog family member B | 3810.66 | 2981.14 | 0.35 | 6.21E-02 |
| NULL | GSMG0008690 | NULL | NULL | 277.35 | 259.54 | 0.09 | 8.20E-01 |
| NULL | GSMG0008694 | NULL | NULL | 109.23 | 109.94 | -0.01 | 9.82E-01 |
| ENSMUSG00000054459 | GSMG0008700 | Vsnl1 | visinin-like 1 | 648.93 | 522.20 | 0.31 | 3.87E-01 |
| ENSMUSG00000037169 | GSMG0008705 | Mycn | v-myc avian myelocytomatosis viral related oncogene, neuroblastoma d | 455.81 | 850.42 | -0.90 | 4.53E-07 |
| ENSMUSG00000037149 | GSMG0008707 | Ddx1 | DEAD (Asp-Glu-Ala-Asp) box polypeptide 1 | 2834.54 | 4112.69 | -0.54 | 7.47E-04 |
| ENSMUSG00000020607 | GSMG0008712 | Fam84a | family with sequence similarity 84, member A | 348.59 | 544.56 | -0.65 | 1.55E-02 |
| ENSMUSG00000020601 | GSMG0008715 | Trib2 | tribbles pseudokinase 2 | 6611.69 | 6348.29 | 0.06 | 8.31E-01 |
| ENSMUSG00000020593 | GSMG0008719 | Lpin1 | lipin 1 | 654.72 | 671.35 | -0.03 | 9.16E-01 |
| ENSMUSG00000051726 | GSMG0008727 | Kcnf1 | potassium voltage-gated channel, subfamily F, member 1 | 932.24 | 605.85 | 0.62 | 2.72E-03 |
| ENSMUSG00000062352 | GSMG0008736 | Itgb1bp1 | integrin beta 1 binding protein 1 | 519.48 | 518.64 | 0.00 | 9.91E-01 |
| ENSMUSG00000052593 | GSMG0008737 | Adam17 | a disintegrin and metallopeptidase domain 17 | 2472.23 | 1673.42 | 0.56 | 2.90E-04 |
| NULL | GSMG0008744 | NULL | NULL | 285.98 | 303.40 | -0.09 | 8.08E-01 |
| NULL | GSMG0008747 | NULL | NULL | 279.65 | 649.64 | -1.21 | 2.14E-08 |
| ENSMUSG00000020644 | GSMG0008749 | Id2 | inhibitor of DNA binding 2 | 872.14 | 1100.69 | -0.34 | 1.69E-01 |
| ENSMUSG00000020642 | GSMG0008752 | Rnf144a | ring finger protein 144A | 196.38 | 341.64 | -0.79 | 9.03E-04 |
| NULL | GSMG0008753 | NULL | NULL | 606.65 | 531.90 | 0.19 | 4.61E-01 |
| ENSMUSG00000020641 | GSMG0008754 | Rsad2 | radical S-adenosyl methionine domain containing 2 | 110.02 | 224.80 | -1.04 | 6.27E-05 |
| ENSMUSG00000020628 | GSMG0008766 | Trappc12 | trafficking protein particle complex 12 | 764.72 | 798.79 | -0.06 | 8.41E-01 |
| ENSMUSG00000020664 | GSMG0008777 | Dld | dihydrolipoamide dehydrogenase | 1979.89 | 2335.97 | -0.24 | 3.29E-01 |
| ENSMUSG00000020659 | GSMG0008778 | Cbll1 | Casitas B-lineage lymphoma-like 1 | 504.65 | 593.10 | -0.23 | 3.50E-01 |
| ENSMUSG00000020650 | GSMG0008781 | Bcap29 | B cell receptor associated protein 29 | 383.08 | 367.63 | 0.06 | 9.03E-01 |
| ENSMUSG00000044067 | GSMG0008783 | Gpr22 | G protein-coupled receptor 22 | 183.56 | 113.29 | 0.70 | 1.54E-02 |
| ENSMUSG00000002996 | GSMG0008785 | Hbp1 | high mobility group box transcription factor 1 | 3339.87 | 2728.09 | 0.29 | 7.12E-02 |

| | | | | | | | |
|---|---|---|---|---|---|---|---|
| ENSMUSG00000002997 | GSMG0008786 | Prkar2b | protein kinase, cAMP dependent regulatory, type II beta | 335.70 | 595.25 | -0.83 | 3.51E-06 |
| ENSMUSG00000020573 | GSMG0008789 | Pik3cg | phosphoinositide-3-kinase, catalytic, gamma polypeptide | 294.41 | 662.17 | -1.17 | 2.49E-09 |
| ENSMUSG00000052125 | GSMG0008794 | F730043M19Rik | RIKEN cDNA F730043M19 gene | 519.09 | 623.73 | -0.27 | 2.40E-01 |
| ENSMUSG00000004698 | GSMG0008798 | Hdac9 | histone deacetylase 9 | 463.66 | 499.44 | -0.11 | 7.18E-01 |
| ENSMUSG00000019256 | GSMG0008804 | Ahr | aryl-hydrocarbon receptor | 257.44 | 350.11 | -0.45 | 6.54E-02 |
| ENSMUSG00000020577 | GSMG0008806 | Tspan13 | tetraspanin 13 | 13845.71 | 13975.49 | -0.01 | 9.70E-01 |
| NULL | GSMG0008807 | NULL | NULL | 97.91 | 71.96 | 0.45 | 2.92E-01 |
| ENSMUSG00000020547 | GSMG0008808 | Bzw2 | basic leucine zipper and W2 domains 2 | 1043.35 | 2531.29 | -1.28 | 4.73E-18 |
| ENSMUSG00000047446 | GSMG0008816 | Arl4a | ADP-ribosylation factor-like 4A | 2666.67 | 2287.18 | 0.22 | 2.83E-01 |
| ENSMUSG00000001627 | GSMG0008820 | Ifrd1 | interferon-related developmental regulator 1 | 880.10 | 967.67 | -0.14 | 5.87E-01 |
| ENSMUSG00000055917 | GSMG0008822 | Zfp277 | zinc finger protein 277 | 660.73 | 767.64 | -0.22 | 3.42E-01 |
| ENSMUSG00000036295 | GSMG0008829 | Lrrn3 | leucine rich repeat protein 3, neuronal | 4149.28 | 3620.37 | 0.20 | 4.24E-01 |
| ENSMUSG00000014905 | GSMG0008832 | Dnajb9 | DnaJ heat shock protein family (Hsp40) member B9 | 670.66 | 533.59 | 0.33 | 2.78E-01 |
| ENSMUSG00000046314 | GSMG0008833 | Stxbp6 | syntaxin binding protein 6 (amisyn) | 229.55 | 114.75 | 1.00 | 9.56E-04 |
| ENSMUSG00000002688 | GSMG0008844 | Prkd1 | protein kinase D1 | 283.17 | 286.57 | -0.01 | 9.72E-01 |
| ENSMUSG00000020954 | GSMG0008853 | Strn3 | striatin, calmodulin binding protein 3 | 1059.95 | 701.23 | 0.60 | 3.52E-04 |
| ENSMUSG00000035181 | GSMG0008862 | Heatr5a | HEAT repeat containing 5A | 237.75 | 220.59 | 0.11 | 7.72E-01 |
| ENSMUSG00000020956 | GSMG0008865 | Dtd2 | D-tyrosyl-tRNA deacylase 2 | 286.50 | 322.84 | -0.17 | 5.74E-01 |
| ENSMUSG00000044408 | GSMG0008877 | Sptssa | serine palmitoyltransferase, small subunit A | 3236.09 | 3304.96 | -0.03 | 9.40E-01 |
| ENSMUSG00000054302 | GSMG0008878 | Eapp | E2F-associated phosphoprotein | 1160.09 | 894.32 | 0.38 | 5.67E-02 |
| ENSMUSG00000005656 | GSMG0008879 | Snx6 | sorting nexin 6 | 2026.12 | 2398.33 | -0.24 | 3.99E-01 |
| ENSMUSG00000062929 | GSMG0008881 | Cfl2 | cofilin 2, muscle | 1833.63 | 2411.75 | -0.40 | 2.40E-02 |
| ENSMUSG00000062198 | GSMG0008884 | 2700097O09Rik | RIKEN cDNA 2700097O09 gene | 250.46 | 264.53 | -0.08 | 8.28E-01 |
| ENSMUSG00000021022 | GSMG0008885 | Ppp2r3c | protein phosphatase 2, regulatory subunit B", gamma | 465.22 | 646.20 | -0.47 | 1.13E-02 |
| ENSMUSG00000021025 | GSMG0008888 | Nfkbia | nuclear factor of kappa light polypeptide gene enhancer in B cells inhib | 5365.95 | 3614.00 | 0.57 | 4.15E-04 |
| NULL | GSMG0008890 | NULL | NULL | 408.01 | 530.84 | -0.38 | 1.75E-01 |
| ENSMUSG00000021027 | GSMG0008891 | Ralgapa1 | Ral GTPase activating protein, alpha subunit 1 | 4789.82 | 4222.13 | 0.18 | 3.25E-01 |
| NULL | GSMG0008894 | NULL | NULL | 244.85 | 198.47 | 0.30 | 2.82E-01 |
| ENSMUSG00000021028 | GSMG0008896 | Mbip | MAP3K12 binding inhibitory protein 1 | 584.19 | 589.06 | -0.01 | 9.69E-01 |
| ENSMUSG00000045930 | GSMG0008916 | Clec14a | C-type lectin domain family 14, member a | 3418.34 | 2214.93 | 0.63 | 3.51E-04 |
| ENSMUSG00000020986 | GSMG0008918 | Sec23a | SEC23 homolog A, COPII coat complex component | 1498.40 | 1720.27 | -0.20 | 3.18E-01 |
| ENSMUSG00000020993 | GSMG0008919 | Trappc6b | trafficking protein particle complex 6B | 1666.82 | 1561.55 | 0.09 | 7.13E-01 |
| ENSMUSG00000035329 | GSMG0008920 | Fbxo33 | F-box protein 33 | 191.75 | 126.39 | 0.60 | 2.28E-02 |

| | | | | | | | |
|---|---|---|---|---|---|---|---|
| ENSMUSG00000113429 | GSMG0008924 | Gm20063 | predicted gene, 20063 | 111.21 | 80.23 | 0.47 | 2.15E-01 |
| ENSMUSG00000020948 | GSMG0008927 | Klhl28 | kelch-like 28 | 335.05 | 407.68 | -0.28 | 2.34E-01 |
| NULL | GSMG0008928 | NULL | NULL | 58.05 | 59.25 | -0.03 | 9.63E-01 |
| ENSMUSG00000020949 | GSMG0008929 | Fkbp3 | FK506 binding protein 3 | 950.37 | 1313.11 | -0.47 | 3.38E-02 |
| ENSMUSG00000047534 | GSMG0008931 | Mis18bp1 | MIS18 binding protein 1 | 53.99 | 110.73 | -1.03 | 9.97E-03 |
| NULL | GSMG0008938 | NULL | NULL | 205.71 | 152.18 | 0.43 | 1.88E-01 |
| NULL | GSMG0008939 | NULL | NULL | 502.33 | 414.94 | 0.28 | 2.77E-01 |
| NULL | GSMG0008942 | NULL | NULL | 501.67 | 436.88 | 0.20 | 4.63E-01 |
| NULL | GSMG0008943 | NULL | NULL | 221.72 | 204.62 | 0.12 | 7.87E-01 |
| NULL | GSMG0008944 | NULL | NULL | 198.50 | 127.67 | 0.63 | 4.44E-02 |
| ENSMUSG00000034912 | GSMG0008945 | Mdga2 | MAM domain containing glycosylphosphatidylinositol anchor 2 | 407.05 | 585.24 | -0.52 | 1.80E-02 |
| G00000020973,ENSMUSG000 | GSMG0008947 | Rpl36al // Dnaaf2 | ribosomal protein L36A-like // dynein, axonemal assembly factor 2 | 1173.66 | 1458.14 | -0.31 | 1.02E-01 |
| ENSMUSG00000020974 | GSMG0008948 | Pole2 | polymerase (DNA directed), epsilon 2 (p59 subunit) | 96.60 | 168.62 | -0.81 | 3.51E-03 |
| ENSMUSG00000020982 | GSMG0008949 | Nemf | nuclear export mediator factor | 2599.25 | 2677.57 | -0.04 | 8.84E-01 |
| NULL | GSMG0008950 | NULL | NULL | 2350.93 | 2533.98 | -0.11 | 6.79E-01 |
| ENSMUSG00000049882 | GSMG0008952 | Vcpkmt | valosin containing protein lysine (K) methyltransferase | 57.58 | 76.11 | -0.41 | 4.14E-01 |
| ENSMUSG00000034801 | GSMG0008953 | Sos2 | son of sevenless homolog 2 (Drosophila) | 1397.76 | 643.49 | 1.12 | 3.43E-06 |
| ENSMUSG00000020988 | GSMG0008956 | L2hgdh | L-2-hydroxyglutarate dehydrogenase | 108.93 | 128.91 | -0.25 | 5.39E-01 |
| ENSMUSG00000020990 | GSMG0008958 | Cdkl1 | cyclin-dependent kinase-like 1 (CDC2-related kinase) | 80.04 | 129.90 | -0.70 | 3.05E-02 |
| ENSMUSG00000034761 | GSMG0008959 | Map4k5 | mitogen-activated protein kinase kinase kinase kinase 5 | 864.60 | 1201.38 | -0.47 | 2.76E-02 |
| ENSMUSG00000021067 | GSMG0008962 | Sav1 | salvador family WW domain containing 1 | 268.08 | 176.30 | 0.60 | 4.00E-02 |
| ENSMUSG00000021068 | GSMG0008964 | Nin | ninein | 1147.78 | 1634.98 | -0.51 | 1.34E-02 |
| NULL | GSMG0008965 | NULL | NULL | 87.08 | 164.34 | -0.92 | 2.19E-03 |
| ENSMUSG00000021071 | GSMG0008967 | Trim9 | tripartite motif-containing 9 | 893.48 | 720.01 | 0.31 | 2.57E-01 |
| ENSMUSG00000085622 | GSMG0008968 | 3110056K07Rik | RIKEN cDNA 3110056K07 gene | 113.51 | 78.02 | 0.54 | 1.46E-01 |
| ENSMUSG00000021079 | GSMG0008970 | Timm9 | translocase of inner mitochondrial membrane 9 | 88.28 | 104.60 | -0.25 | 5.83E-01 |
| NULL | GSMG0008972 | NULL | NULL | 180.39 | 205.94 | -0.20 | 5.98E-01 |
| ENSMUSG00000043398 | GSMG0008976 | Gpr135 | G protein-coupled receptor 135 | 279.47 | 311.40 | -0.16 | 6.13E-01 |
| ENSMUSG00000019718 | GSMG0008977 | L3hypdh | L-3-hydroxyproline dehydratase (trans-) | 336.20 | 313.56 | 0.10 | 7.77E-01 |
| ENSMUSG00000021087 | GSMG0008980 | Rtn1 | reticulon 1 | 4295.91 | 9731.08 | -1.18 | 4.73E-14 |
| NULL | GSMG0008981 | NULL | NULL | 213.28 | 486.35 | -1.19 | 1.09E-07 |
| ENSMUSG00000021094 | GSMG0008984 | Dhrs7 | dehydrogenase/reductase (SDR family) member 7 | 1191.64 | 912.86 | 0.38 | 5.84E-02 |
| NULL | GSMG0008986 | NULL | NULL | 121.09 | 114.41 | 0.08 | 8.94E-01 |

| | | | | | | | |
|---|---|---|---|---|---|---|---|
| ENSMUSG00000034442 | GSMG0008992 | Trmt5 | TRM5 tRNA methyltransferase 5 | 275.64 | 315.03 | -0.19 | 5.13E-01 |
| NULL | GSMG0008996 | NULL | NULL | 155.33 | 175.11 | -0.17 | 6.41E-01 |
| ENSMUSG00000034402 | GSMG0009000 | Kcnh5 | potassium voltage-gated channel, subfamily H (eag-related), member 5 | 496.41 | 368.98 | 0.43 | 1.30E-01 |
| ENSMUSG00000021051 | GSMG0009002 | Ppp2r5e | protein phosphatase 2, regulatory subunit B', epsilon | 395.13 | 517.43 | -0.39 | 1.69E-01 |
| ENSMUSG00000045690 | GSMG0009008 | Wdr89 | WD repeat domain 89 | 877.59 | 1373.22 | -0.65 | 5.56E-03 |
| ENSMUSG00000021054 | GSMG0009009 | Sgpp1 | sphingosine-1-phosphate phosphatase 1 | 1759.65 | 1409.45 | 0.32 | 8.60E-02 |
| ENSMUSG00000056459 | GSMG0009015 | Zbtb25 | zinc finger and BTB domain containing 25 | 317.38 | 374.24 | -0.24 | 3.44E-01 |
| NULL | GSMG0009016 | NULL | NULL | 145.82 | 159.33 | -0.13 | 7.68E-01 |
| ENSMUSG00000021062 | GSMG0009022 | Rab15 | RAB15, member RAS oncogene family | 314.36 | 227.56 | 0.47 | 8.16E-02 |
| NULL | GSMG0009023 | NULL | NULL | 286.95 | 129.22 | 1.16 | 3.54E-06 |
| ENSMUSG00000059436 | GSMG0009025 | Max | Max protein | 921.89 | 842.94 | 0.13 | 7.44E-01 |
| ENSMUSG00000021114 | GSMG0009034 | Atp6v1d | ATPase, H+ transporting, lysosomal V1 subunit D | 1018.86 | 1025.29 | -0.01 | 9.79E-01 |
| ENSMUSG00000046157 | GSMG0009037 | Tmem229b | transmembrane protein 229B | 1289.57 | 1762.25 | -0.45 | 1.92E-02 |
| ENSMUSG00000021124 | GSMG0009041 | Vti1b | vesicle transport through interaction with t-SNAREs 1B | 2134.70 | 2251.65 | -0.08 | 7.93E-01 |
| NULL | GSMG0009042 | NULL | NULL | 310.88 | 225.76 | 0.46 | 6.21E-02 |
| ENSMUSG00000066441 | GSMG0009043 | Rdh11 | retinol dehydrogenase 11 | 303.27 | 441.66 | -0.54 | 2.06E-02 |
| ENSMUSG00000066440 | GSMG0009044 | Zfyve26 | zinc finger, FYVE domain containing 26 | 973.83 | 955.14 | 0.03 | 9.50E-01 |
| ENSMUSG00000021127 | GSMG0009050 | Zfp36l1 | zinc finger protein 36, C3H type-like 1 | 4806.53 | 5819.65 | -0.28 | 3.10E-01 |
| ENSMUSG00000015143 | GSMG0009052 | Actn1 | actinin, alpha 1 | 756.77 | 654.85 | 0.21 | 5.21E-01 |
| ENSMUSG00000049106 | GSMG0009055 | Dcaf5 | DDB1 and CUL4 associated factor 5 | 466.83 | 380.02 | 0.30 | 4.20E-01 |
| NULL | GSMG0009060 | NULL | NULL | 127.67 | 212.53 | -0.74 | 2.73E-02 |
| ENSMUSG00000062961 | GSMG0009063 | Ccdc177 | coiled-coil domain containing 177 | 480.95 | 634.58 | -0.40 | 2.27E-01 |
| 39,ENSMUSG00000090935,El | GSMG0009076 | Synj2bp-cox16 // C | Synj2bp-cox16 readthrough // cytochrome c oxidase assembly protein 1 | 3076.03 | 2957.53 | 0.06 | 8.31E-01 |
| ENSMUSG00000002679 | GSMG0009080 | Med6 | mediator complex subunit 6 | 365.35 | 427.83 | -0.23 | 3.62E-01 |
| ENSMUSG00000042724 | GSMG0009084 | Map3k9 | mitogen-activated protein kinase kinase kinase 9 | 124.58 | 177.09 | -0.50 | 1.48E-01 |
| ENSMUSG00000021221 | GSMG0009091 | Dpf3 | D4, zinc and double PHD fingers, family 3 | 134.24 | 139.85 | -0.06 | 9.11E-01 |
| ENSMUSG00000042628 | GSMG0009096 | Zfyve1 | zinc finger, FYVE domain containing 1 | 934.92 | 1049.41 | -0.17 | 5.28E-01 |
| ENSMUSG00000021224 | GSMG0009100 | Numb | numb homolog (Drosophila) | 1063.58 | 824.13 | 0.37 | 4.85E-02 |
| NULL | GSMG0009101 | NULL | NULL | 188.61 | 85.00 | 1.14 | 3.50E-04 |
| NULL | GSMG0009102 | NULL | NULL | 758.28 | 360.24 | 1.07 | 7.24E-09 |
| NULL | GSMG0009105 | NULL | NULL | 84.76 | 24.29 | 1.80 | 4.21E-05 |
| ENSMUSG00000042507 | GSMG0009111 | Elmsan1 | ELM2 and Myb/SANT-like domain containing 1 | 462.39 | 349.52 | 0.40 | 2.48E-01 |
| ENSMUSG00000021236 | GSMG0009116 | Entpd5 | ectonucleoside triphosphate diphosphohydrolase 5 | 561.32 | 534.69 | 0.07 | 8.12E-01 |

| | | | | | | | |
|---|---|---|---|---|---|---|---|
| ENSMUSG00000021238 | GSMG0009117 | Aldh6a1 | aldehyde dehydrogenase family 6, subfamily A1 | 982.56 | 814.84 | 0.27 | 1.79E-01 |
| ENSMUSG00000021240 | GSMG0009118 | Abcd4 | ATP-binding cassette, sub-family D (ALD), member 4 | 527.20 | 423.12 | 0.32 | 2.02E-01 |
| ENSMUSG00000071234 | GSMG0009119 | Syndig1l | synapse differentiation inducing 1 like | 33.65 | 82.78 | -1.29 | 6.98E-04 |
| ENSMUSG00000021242 | GSMG0009120 | Npc2 | Niemann-Pick type C2 | 2463.86 | 2373.57 | 0.05 | 8.56E-01 |
| ENSMUSG00000042350 | GSMG0009122 | Arel1 | apoptosis resistant E3 ubiquitin protein ligase 1 | 817.54 | 935.30 | -0.19 | 4.05E-01 |
| ENSMUSG00000019235 | GSMG0009127 | Rps6kl1 | ribosomal protein S6 kinase-like 1 | 590.58 | 534.65 | 0.15 | 6.22E-01 |
| ENSMUSG00000004791 | GSMG0009128 | Pgf | placental growth factor | 375.93 | 284.65 | 0.40 | 1.68E-01 |
| ENSMUSG00000021245 | GSMG0009129 | Mlh3 | mutL homolog 3 | 113.76 | 113.18 | 0.01 | 9.81E-01 |
| ENSMUSG00000008822 | GSMG0009130 | Acyp1 | acylphosphatase 1, erythrocyte (common) type | 260.40 | 222.35 | 0.22 | 5.94E-01 |
| ENSMUSG00000034290 | GSMG0009132 | Nek9 | NIMA (never in mitosis gene a)-related expressed kinase 9 | 890.04 | 906.88 | -0.03 | 9.48E-01 |
| ENSMUSG00000021252 | GSMG0009138 | 0610007P14Rik | RIKEN cDNA 0610007P14 gene | 839.12 | 1270.77 | -0.60 | 4.08E-04 |
| ENSMUSG00000021253 | GSMG0009140 | Tgfb3 | transforming growth factor, beta 3 | 249.97 | 148.44 | 0.76 | 1.27E-02 |
| ENSMUSG00000021257 | GSMG0009142 | Angel1 | angel homolog 1 (Drosophila) | 891.45 | 1146.41 | -0.36 | 4.43E-02 |
| NULL | GSMG0009143 | NULL | NULL | 494.77 | 512.26 | -0.05 | 8.78E-01 |
| ENSMUSG00000034126 | GSMG0009148 | Pomt2 | protein-O-mannosyltransferase 2 | 809.96 | 951.65 | -0.23 | 3.42E-01 |
| ENSMUSG00000113918 | GSMG0009149 | Gm6566 | predicted gene 6566 | 140.25 | 133.18 | 0.08 | 8.73E-01 |
| ENSMUSG00000034111 | GSMG0009150 | Tmed8 | transmembrane emp24 domain containing 8 | 82.03 | 66.23 | 0.31 | 5.74E-01 |
| ENSMUSG00000021036 | GSMG0009153 | Sptlc2 | serine palmitoyltransferase, long chain base subunit 2 | 1237.12 | 1395.89 | -0.17 | 5.11E-01 |
| ENSMUSG00000079036 | GSMG0009156 | Alkbh1 | alkB homolog 1, histone H2A dioxygenase | 413.08 | 517.41 | -0.32 | 2.00E-01 |
| ENSMUSG00000021039 | GSMG0009157 | Snw1 | SNW domain containing 1 | 1525.61 | 1655.01 | -0.12 | 6.34E-01 |
| ENSMUSG00000007682 | GSMG0009166 | Dio2 | deiodinase, iodothyronine, type II | 643.92 | 410.38 | 0.65 | 6.18E-04 |
| ENSMUSG00000061533 | GSMG0009168 | Cep128 | centrosomal protein 128 | 118.79 | 169.29 | -0.52 | 8.17E-02 |
| ENSMUSG00000020962 | GSMG0009170 | Gtf2a1 | general transcription factor II A, 1 | 294.63 | 369.23 | -0.33 | 1.74E-01 |
| ENSMUSG00000020964 | GSMG0009174 | Sel1l | sel-1 suppressor of lin-12-like (C. elegans) | 2024.78 | 2224.43 | -0.14 | 5.75E-01 |
| ENSMUSG00000021003 | GSMG0009179 | Galc | galactosylceramidase | 427.45 | 450.11 | -0.07 | 8.18E-01 |
| ENSMUSG00000033854 | GSMG0009182 | Kcnk10 | potassium channel, subfamily K, member 10 | 51.33 | 188.80 | -1.90 | 8.57E-11 |
| ENSMUSG00000021009 | GSMG0009185 | Ptpn21 | protein tyrosine phosphatase, non-receptor type 21 | 445.39 | 332.60 | 0.42 | 9.29E-02 |
| ENSMUSG00000051166 | GSMG0009186 | Eml5 | echinoderm microtubule associated protein like 5 | 1100.95 | 968.99 | 0.18 | 4.61E-01 |
| NULL | GSMG0009191 | NULL | NULL | 247.12 | 285.00 | -0.21 | 5.29E-01 |
| ENSMUSG00000021179 | GSMG0009207 | Nrde2 | nrde-2 necessary for RNA interference, domain containing | 430.04 | 512.91 | -0.26 | 3.89E-01 |
| ENSMUSG00000033530 | GSMG0009209 | Ttc7b | tetratricopeptide repeat domain 7B | 1475.79 | 810.37 | 0.86 | 6.57E-04 |
| ENSMUSG00000021180 | GSMG0009213 | Rps6ka5 | ribosomal protein S6 kinase, polypeptide 5 | 182.17 | 204.70 | -0.16 | 6.80E-01 |
| ENSMUSG00000021182 | GSMG0009219 | Ccdc88c | coiled-coil domain containing 88C | 86.39 | 122.59 | -0.51 | 1.72E-01 |

| | | | | | | | |
|---|---|---|---|---|---|---|---|
| ENSMUSG00000041846 | GSMG0009220 | Smek1 | SMEK homolog 1, suppressor of mek1 (Dictyostelium) | 880.89 | 1090.53 | -0.31 | 1.18E-01 |
| ENSMUSG00000021186 | GSMG0009222 | Fbln5 | fibulin 5 | 4081.80 | 1328.31 | 1.62 | 2.28E-11 |
| NULL | GSMG0009223 | NULL | NULL | 215.09 | 186.79 | 0.20 | 6.00E-01 |
| ENSMUSG00000021188 | GSMG0009224 | Trip11 | thyroid hormone receptor interactor 11 | 226.60 | 203.77 | 0.15 | 6.81E-01 |
| ENSMUSG00000021189 | GSMG0009226 | Atxn3 | ataxin 3 | 276.10 | 275.99 | 0.00 | 9.96E-01 |
| ENSMUSG00000021190 | GSMG0009230 | Lgmn | legumain | 3601.70 | 2599.99 | 0.47 | 3.61E-03 |
| G00000057963,ENSMUSG000 | GSMG0009231 | Itpk1 // Gm20604 / | inositol 1,3,4-triphosphate 5/6 kinase // predicted gene 20604 // modula | 2039.56 | 2136.85 | -0.07 | 8.09E-01 |
| NULL | GSMG0009233 | NULL | NULL | 840.84 | 716.41 | 0.23 | 3.51E-01 |
| ENSMUSG00000041702 | GSMG0009234 | Btbd7 | BTB (POZ) domain containing 7 | 397.80 | 333.41 | 0.26 | 4.23E-01 |
| ENSMUSG00000041645 | GSMG0009245 | Ddx24 | DEAD (Asp-Glu-Ala-Asp) box polypeptide 24 | 3735.82 | 3904.23 | -0.06 | 7.83E-01 |
| ENSMUSG00000041415 | GSMG0009262 | Dicer1 | dicer 1, ribonuclease type III | 374.63 | 424.78 | -0.18 | 5.28E-01 |
| ENSMUSG00000021097 | GSMG0009263 | Clmn | calmin | 336.09 | 90.82 | 1.89 | 1.31E-14 |
| ENSMUSG00000113722 | GSMG0009268 | Snhg10 | small nucleolar RNA host gene 10 | 164.69 | 166.52 | -0.02 | 9.65E-01 |
| ENSMUSG00000041341 | GSMG0009274 | Atg2b | autophagy related 2B | 350.64 | 367.93 | -0.07 | 8.57E-01 |
| ENSMUSG00000048251 | GSMG0009287 | Bcl11b | B cell leukemia/lymphoma 11B | 121.42 | 195.44 | -0.69 | 4.13E-02 |
| NULL | GSMG0009291 | NULL | NULL | 324.88 | 321.59 | 0.01 | 9.72E-01 |
| ENSMUSG00000056770 | GSMG0009294 | Setd3 | SET domain containing 3 | 528.01 | 589.18 | -0.16 | 5.28E-01 |
| ENSMUSG00000021263 | GSMG0009303 | Degs2 | delta(4)-desaturase, sphingolipid 2 | 1222.67 | 91.36 | 3.74 | 2.27E-73 |
| NULL | GSMG0009304 | NULL | NULL | 228.03 | 284.67 | -0.32 | 3.20E-01 |
| ENSMUSG00000021265 | GSMG0009305 | Slc25a29 | solute carrier family 25 (mitochondrial carrier, palmitoylcarnitine transp | 320.03 | 381.70 | -0.25 | 3.90E-01 |
| ENSMUSG00000021266 | GSMG0009306 | Wars | tryptophanyl-tRNA synthetase | 1582.54 | 1837.04 | -0.22 | 2.55E-01 |
| ENSMUSG00000040867 | GSMG0009308 | Begain | brain-enriched guanylate kinase-associated | 2816.88 | 2448.88 | 0.20 | 5.84E-01 |
| NULL | GSMG0009316 | NULL | NULL | 74.45 | 52.31 | 0.50 | 3.47E-01 |
| ENSMUSG00000056458 | GSMG0009323 | Mok | MOK protein kinase | 138.45 | 159.48 | -0.20 | 5.95E-01 |
| ENSMUSG00000021276 | GSMG0009325 | Cinp | cyclin-dependent kinase 2 interacting protein | 1086.97 | 1127.83 | -0.05 | 8.50E-01 |
| ENSMUSG00000021279 | GSMG0009330 | Cdc42bpb | CDC42 binding protein kinase beta | 1144.06 | 1250.99 | -0.13 | 7.65E-01 |
| NULL | GSMG0009331 | NULL | NULL | 311.86 | 365.79 | -0.23 | 5.25E-01 |
| ENSMUSG00000010529 | GSMG0009334 | Gm266 | predicted gene 266 | 185.34 | 56.98 | 1.70 | 3.90E-06 |
| ENSMUSG00000096995 | GSMG0009335 | 2810029C07Rik | RIKEN cDNA 2810029C07 gene | 255.94 | 123.19 | 1.06 | 3.77E-06 |
| ENSMUSG00000001270 | GSMG0009336 | Ckb | creatine kinase, brain | 8398.70 | 5850.00 | 0.52 | 1.23E-03 |
| ENSMUSG00000049792 | GSMG0009337 | Bag5 | BCL2-associated athanogene 5 | 1928.61 | 1828.90 | 0.08 | 7.68E-01 |
| ENSMUSG00000021287 | GSMG0009338 | Xrcc3 | X-ray repair complementing defective repair in Chinese hamster cells 3 | 298.61 | 283.66 | 0.07 | 8.70E-01 |
| ENSMUSG00000021285 | GSMG0009339 | Ppp1r13b | protein phosphatase 1, regulatory (inhibitor) subunit 13B | 833.81 | 807.46 | 0.05 | 8.89E-01 |

| | | | | | | | |
|---|---|---|---|---|---|---|---|
| NULL | GSMG0009340 | NULL | NULL | 251.06 | 128.75 | 0.96 | 4.39E-05 |
| ENSMUSG00000113769 | GSMG0009342 | 5033406O09Rik | RIKEN cDNA 5033406O09 gene | 218.33 | 159.34 | 0.46 | 1.63E-01 |
| ENSMUSG00000021290 | GSMG0009343 | 2010107E04Rik | RIKEN cDNA 2010107E04 gene | 1437.65 | 1654.68 | -0.20 | 3.46E-01 |
| ENSMUSG00000054013 | GSMG0009349 | Tmem179 | transmembrane protein 179 | 1768.43 | 1273.95 | 0.47 | 1.80E-02 |
| ENSMUSG00000001729 | GSMG0009350 | Akt1 | thymoma viral proto-oncogene 1 | 6711.70 | 6417.16 | 0.06 | 8.65E-01 |
| NULL | GSMG0009353 | NULL | NULL | 66.77 | 346.86 | -2.36 | 7.29E-18 |
| ENSMUSG00000047832 | GSMG0009354 | Cdca4 | cell division cycle associated 4 | 724.45 | 1030.78 | -0.51 | 6.17E-03 |
| ENSMUSG00000002799 | GSMG0009357 | Jag2 | jagged 2 | 3365.76 | 1734.27 | 0.96 | 1.38E-01 |
| ENSMUSG00000002804 | GSMG0009358 | Nudt14 | nudix (nucleoside diphosphate linked moiety X)-type motif 14 | 946.04 | 1043.83 | -0.14 | 5.57E-01 |
| ENSMUSG00000011158 | GSMG0009361 | Brf1 | BRF1, RNA polymerase III transcription initiation factor 90 kDa subun | 891.17 | 882.66 | 0.02 | 9.62E-01 |
| NULL | GSMG0009363 | NULL | NULL | 1710.09 | 1289.61 | 0.41 | 2.67E-02 |
| ENSMUSG00000042050 | GSMG0009367 | Wdr60 | WD repeat domain 60 | 462.82 | 351.01 | 0.40 | 5.21E-02 |
| NULL | GSMG0009368 | NULL | NULL | 147.05 | 134.86 | 0.12 | 8.07E-01 |
| ENSMUSG00000021218 | GSMG0009385 | Gdi2 | guanosine diphosphate (GDP) dissociation inhibitor 2 | 3754.87 | 4563.72 | -0.28 | 1.41E-01 |
| ENSMUSG00000033781 | GSMG0009387 | Asb13 | ankyrin repeat and SOCS box-containing 13 | 257.59 | 198.97 | 0.37 | 1.85E-01 |
| ENSMUSG00000033715 | GSMG0009392 | Akr1c14 | aldo-keto reductase family 1, member C14 | 589.10 | 366.74 | 0.68 | 1.79E-03 |
| ENSMUSG00000000078 | GSMG0009400 | Klf6 | Kruppel-like factor 6 | 2110.68 | 1558.94 | 0.44 | 3.46E-02 |
| ENSMUSG00000021193 | GSMG0009402 | Pitrm1 | pitrilysin metallepetidase 1 | 1566.22 | 1838.17 | -0.23 | 2.28E-01 |
| ENSMUSG00000052551 | GSMG0009403 | Adarb2 | adenosine deaminase, RNA-specific, B2 | 221.78 | 310.44 | -0.48 | 1.49E-01 |
| NULL | GSMG0009404 | NULL | NULL | 159.94 | 210.97 | -0.40 | 1.89E-01 |
| ENSMUSG00000033499 | GSMG0009414 | Larp4b | La ribonucleoprotein domain family, member 4B | 1126.45 | 1115.30 | 0.01 | 9.64E-01 |
| ENSMUSG00000048264 | GSMG0009418 | Dip2c | disco interacting protein 2 homolog C | 178.35 | 223.01 | -0.32 | 3.75E-01 |
| NULL | GSMG0009419 | NULL | NULL | 114.89 | 481.60 | -2.06 | 5.11E-23 |
| NULL | GSMG0009423 | NULL | NULL | 412.41 | 343.06 | 0.27 | 4.82E-01 |
| ENSMUSG00000050244 | GSMG0009432 | Heatr1 | HEAT repeat containing 1 | 89.37 | 107.44 | -0.27 | 5.12E-01 |
| ENSMUSG00000057069 | GSMG0009434 | Ero1lb | ERO1-like beta (S. cerevisiae) | 211.88 | 172.94 | 0.30 | 3.76E-01 |
| NULL | GSMG0009435 | NULL | NULL | 260.00 | 487.29 | -0.91 | 3.29E-06 |
| ENSMUSG00000005397 | GSMG0009442 | Nid1 | nidogen 1 | 15129.20 | 21466.79 | -0.50 | 5.08E-03 |
| ENSMUSG00000019726 | GSMG0009444 | Lyst | lysosomal trafficking regulator | 482.31 | 329.34 | 0.55 | 8.63E-03 |
| ENSMUSG00000021303 | GSMG0009445 | Gng4 | guanine nucleotide binding protein (G protein), gamma 4 | 616.25 | 922.84 | -0.58 | 4.14E-03 |
| ENSMUSG00000039242 | GSMG0009447 | B3galnt2 | UDP-GalNAc:betaGlcNAc beta 1,3-galactosaminyltransferase, polypep | 370.81 | 335.59 | 0.15 | 6.64E-01 |
| ENSMUSG00000039219 | GSMG0009448 | Arid4b | AT rich interactive domain 4B (RBP1-like) | 431.54 | 414.25 | 0.06 | 8.65E-01 |
| NULL | GSMG0009449 | NULL | NULL | 261.18 | 242.63 | 0.10 | 8.04E-01 |

| | | | | | | | |
|---|---|---|---|---|---|---|---|
| ENSMUSG00000015671 | GSMG0009451 | Psma2 | proteasome (prosome, macropain) subunit, alpha type 2 | 2205.52 | 2569.00 | -0.22 | 3.00E-01 |
| ENSMUSG00000039182 | GSMG0009452 | AW209491 | expressed sequence AW209491 | 902.19 | 773.17 | 0.22 | 3.94E-01 |
| ENSMUSG00000041324 | GSMG0009465 | Inhba | inhibin beta-A | 24.78 | 71.19 | -1.54 | 2.17E-04 |
| ENSMUSG00000012429 | GSMG0009469 | Mplkip | M-phase specific PLK1 intereacting protein | 627.96 | 749.00 | -0.25 | 3.06E-01 |
| ENSMUSG00000041236 | GSMG0009475 | Vps41 | VPS41 HOPS complex subunit | 3293.63 | 3169.00 | 0.06 | 8.16E-01 |
| ENSMUSG00000021314 | GSMG0009477 | Amph | amphiphysin | 211.71 | 304.78 | -0.52 | 2.67E-02 |
| NULL | GSMG0009478 | NULL | NULL | 241.87 | 168.83 | 0.52 | 4.00E-02 |
| ENSMUSG00000041112 | GSMG0009483 | Elmo1 | engulfment and cell motility 1 | 956.00 | 1239.44 | -0.38 | 3.01E-02 |
| ENSMUSG00000021326 | GSMG0009495 | Trim27 | tripartite motif-containing 27 | 423.66 | 572.58 | -0.43 | 3.18E-02 |
| ENSMUSG00000036721 | GSMG0009498 | Zscan12 | zinc finger and SCAN domain containing 12 | 66.84 | 95.31 | -0.51 | 2.05E-01 |
| ENSMUSG00000006720 | GSMG0009509 | Zfp184 | zinc finger protein 184 (Kruppel-like) | 32.50 | 72.63 | -1.14 | 1.14E-02 |
| ENSMUSG00000061482 | GSMG0009534 | Hist1h4d | histone cluster 1, H4d | 128.53 | 70.86 | 0.85 | 4.54E-02 |
| ENSMUSG00000018102 | GSMG0009538 | Hist1h2bc | histone cluster 1, H2bc | 395.21 | 213.07 | 0.89 | 3.11E-04 |
| ENSMUSG00000036181 | GSMG0009540 | Hist1h1c | histone cluster 1, H1c | 374.06 | 509.76 | -0.45 | 4.36E-02 |
| ENSMUSG00000036006 | GSMG0009554 | Fam65b | family with sequence similarity 65, member B | 373.47 | 443.17 | -0.25 | 3.80E-01 |
| ENSMUSG00000019132 | GSMG0009556 | BC005537 | cDNA sequence BC005537 | 1699.25 | 1595.90 | 0.09 | 7.97E-01 |
| ENSMUSG00000035958 | GSMG0009557 | Tdp2 | tyrosyl-DNA phosphodiesterase 2 | 595.02 | 611.79 | -0.04 | 9.23E-01 |
| ENSMUSG00000006711 | GSMG0009558 | D130043K22Rik | RIKEN cDNA D130043K22 gene | 962.94 | 700.32 | 0.46 | 3.05E-02 |
| ENSMUSG00000086448 | GSMG0009561 | 9330162012Rik | cDNA RIKEN 9330162012 gene | 402.01 | 373.81 | 0.10 | 7.47E-01 |
| ENSMUSG00000021340 | GSMG0009562 | Gpld1 | glycosylphosphatidylinositol specific phospholipase D1 | 295.09 | 204.18 | 0.54 | 6.47E-02 |
| ENSMUSG00000035910 | GSMG0009563 | Dcdc2a | doublecortin domain containing 2a | 253.88 | 157.84 | 0.69 | 7.47E-03 |
| ENSMUSG00000086363 | GSMG0009587 | A330102I10Rik | RIKEN cDNA A330102I10 gene | 107.02 | 201.11 | -0.92 | 6.33E-04 |
| NULL | GSMG0009589 | NULL | NULL | 84.85 | 134.45 | -0.66 | 3.11E-02 |
| ENSMUSG00000038732 | GSMG0009595 | Mboat1 | membrane bound O-acyltransferase domain containing 1 | 219.16 | 172.29 | 0.34 | 2.92E-01 |
| ENSMUSG00000069255 | GSMG0009600 | Dusp22 | dual specificity phosphatase 22 | 622.59 | 507.16 | 0.29 | 2.43E-01 |
| ENSMUSG00000038415 | GSMG0009604 | Foxq1 | forkhead box Q1 | 6280.79 | 7084.68 | -0.17 | 4.68E-01 |
| ENSMUSG00000038402 | GSMG0009606 | Foxf2 | forkhead box F2 | 4526.85 | 5682.72 | -0.33 | 9.91E-02 |
| ENSMUSG00000050295 | GSMG0009607 | Foxc1 | forkhead box C1 | 3063.59 | 2155.57 | 0.51 | 6.56E-02 |
| ENSMUSG00000021400 | GSMG0009610 | Wrnip1 | Werner helicase interacting protein 1 | 692.50 | 612.83 | 0.18 | 4.87E-01 |
| ENSMUSG00000042842 | GSMG0009612 | Serpinb6b | serine (or cysteine) peptidase inhibitor, clade B, member 6b | 416.99 | 402.07 | 0.05 | 8.90E-01 |
| ENSMUSG00000045827 | GSMG0009614 | Serpinb9 | serine (or cysteine) peptidase inhibitor, clade B, member 9 | 438.45 | 290.41 | 0.59 | 9.34E-03 |
| ENSMUSG00000046949 | GSMG0009624 | Nqo2 | NAD(P)H dehydrogenase, quinone 2 | 225.40 | 275.19 | -0.29 | 3.21E-01 |
| ENSMUSG00000021408 | GSMG0009625 | Ripk1 | receptor (TNFRSF)-interacting serine-threonine kinase 1 | 1276.78 | 1096.36 | 0.22 | 3.02E-01 |

| | | | | | | | |
|---|---|---|---|---|---|---|---|
| ENSMUSG00000038286 | GSMG0009626 | Bphl | biphenyl hydrolase-like (serine hydrolase, breast epithelial mucin-assoc | 381.11 | 314.14 | 0.28 | 2.45E-01 |
| ENSMUSG00000071451 | GSMG0009629 | Psmg4 | proteasome (prosome, macropain) assembly chaperone 4 | 356.90 | 746.05 | -1.07 | 3.09E-08 |
| ENSMUSG00000021413 | GSMG0009632 | Prpf4b | pre-mRNA processing factor 4B | 4184.14 | 3709.16 | 0.17 | 4.06E-01 |
| ENSMUSG00000059288 | GSMG0009638 | Cdyl | chromodomain protein, Y chromosome-like | 294.56 | 291.99 | 0.01 | 9.81E-01 |
| ENSMUSG00000021420 | GSMG0009641 | Fars2 | phenylalanine-tRNA synthetase 2 (mitochondrial) | 561.05 | 647.75 | -0.21 | 4.23E-01 |
| ENSMUSG00000021423 | GSMG0009648 | Ly86 | lymphocyte antigen 86 | 107.87 | 57.99 | 0.90 | 1.97E-02 |
| ENSMUSG00000039087 | GSMG0009653 | Rreb1 | ras responsive element binding protein 1 | 920.35 | 789.18 | 0.22 | 5.70E-01 |
| NULL | GSMG0009655 | NULL | NULL | 335.34 | 363.51 | -0.12 | 7.74E-01 |
| ENSMUSG00000021428 | GSMG0009658 | Riok1 | RIO kinase 1 (yeast) | 954.06 | 991.12 | -0.05 | 8.55E-01 |
| ENSMUSG00000021431 | GSMG0009661 | Snrnp48 | small nuclear ribonucleoprotein 48 (U11/U12) | 462.95 | 428.13 | 0.11 | 7.37E-01 |
| ENSMUSG00000039004 | GSMG0009662 | Bmp6 | bone morphogenetic protein 6 | 219.98 | 170.67 | 0.37 | 3.06E-01 |
| ENSMUSG00000021360 | GSMG0009672 | Gcnt2 | glucosaminyl (N-acetyl) transferase 2, I-branching enzyme | 1603.94 | 707.88 | 1.18 | 3.29E-12 |
| ENSMUSG00000038683 | GSMG0009674 | Pak1ip1 | PAK1 interacting protein 1 | 1157.38 | 1103.71 | 0.07 | 8.02E-01 |
| ENSMUSG00000021361 | GSMG0009675 | Tmem14c | transmembrane protein 14C | 458.07 | 820.57 | -0.84 | 3.12E-01 |
| ENSMUSG00000091264 | GSMG0009676 | Smim13 | small integral membrane protein 13 | 2147.17 | 2062.08 | 0.06 | 8.23E-01 |
| ENSMUSG00000021366 | GSMG0009683 | Hivep1 | human immunodeficiency virus type I enhancer binding protein 1 | 133.70 | 136.89 | -0.03 | 9.53E-01 |
| ENSMUSG00000021367 | GSMG0009688 | Edn1 | endothelin 1 | 1945.95 | 800.33 | 1.28 | 5.97E-16 |
| ENSMUSG00000054728 | GSMG0009689 | Phactr1 | phosphatase and actin regulator 1 | 1125.69 | 895.88 | 0.33 | 8.30E-02 |
| NULL | GSMG0009690 | NULL | NULL | 1103.48 | 574.78 | 0.94 | 1.46E-06 |
| ENSMUSG00000054021 | GSMG0009693 | Sirt5 | sirtuin 5 | 364.12 | 364.14 | 0.00 | 9.99E-01 |
| ENSMUSG00000063200 | GSMG0009694 | Nol7 | nucleolar protein 7 | 1514.30 | 1906.19 | -0.33 | 1.96E-01 |
| ENSMUSG00000015396 | GSMG0009699 | Cd83 | CD83 antigen | 1254.73 | 1369.51 | -0.13 | 6.23E-01 |
| ENSMUSG00000038518 | GSMG0009707 | Jarid2 | jumonji, AT rich interactive domain 2 | 2187.03 | 2100.99 | 0.06 | 8.48E-01 |
| ENSMUSG00000038175 | GSMG0009716 | Mylip | myosin regulatory light chain interacting protein | 527.30 | 342.32 | 0.62 | 9.05E-03 |
| ENSMUSG00000000253 | GSMG0009717 | Gmpr | guanosine monophosphate reductase | 1239.31 | 1787.93 | -0.53 | 4.62E-04 |
| NULL | GSMG0009719 | NULL | NULL | 82.96 | 139.12 | -0.75 | 1.65E-02 |
| NULL | GSMG0009720 | NULL | NULL | 39.36 | 65.30 | -0.74 | 9.46E-02 |
| ENSMUSG00000021373 | GSMG0009722 | Cap2 | CAP, adenylate cyclase-associated protein, 2 (yeast) | 270.42 | 95.55 | 1.50 | 1.97E-09 |
| ENSMUSG00000069237 | GSMG0009723 | Fam8a1 | family with sequence similarity 8, member A1 | 1336.61 | 1101.19 | 0.28 | 2.61E-01 |
| ENSMUSG00000038080 | GSMG0009726 | Kdm1b | lysine (K)-specific demethylase 1B | 154.20 | 188.19 | -0.28 | 3.98E-01 |
| ENSMUSG00000038068 | GSMG0009728 | Rnf144b | ring finger protein 144B | 483.41 | 371.67 | 0.38 | 2.68E-01 |
| ENSMUSG00000021379 | GSMG0009732 | Id4 | inhibitor of DNA binding 4 | 495.25 | 338.53 | 0.55 | 9.85E-03 |
| ENSMUSG00000097059 | GSMG0009737 | Fam120aos | family with sequence similarity 120A, opposite strand | 295.82 | 229.05 | 0.36 | 2.48E-01 |

| | | | | | | | |
|---|---|---|---|---|---|---|---|
| ENSMUSG00000037966 | GSMG0009739 | Ninj1 | ninjurin 1 | 1427.22 | 1122.20 | 0.35 | 5.37E-02 |
| ENSMUSG00000037933 | GSMG0009741 | Bicd2 | bicaudal D homolog 2 (Drosophila) | 441.56 | 476.59 | -0.11 | 7.01E-01 |
| ENSMUSG00000021385 | GSMG0009742 | Ippk | inositol 1,3,4,5,6-pentakisphosphate 2-kinase | 330.63 | 248.34 | 0.41 | 1.06E-01 |
| ENSMUSG00000043631 | GSMG0009744 | Ecm2 | extracellular matrix protein 2, female organ and adipocyte specific | 322.06 | 287.94 | 0.16 | 6.23E-01 |
| ENSMUSG00000021388 | GSMG0009745 | Aspn | asporin | 719.84 | 425.01 | 0.76 | 7.52E-03 |
| ENSMUSG00000021390 | GSMG0009748 | Ogn | osteoglycin | 190.64 | 223.28 | -0.23 | 4.64E-01 |
| ENSMUSG00000021392 | GSMG0009749 | Nol8 | nucleolar protein 8 | 592.21 | 568.01 | 0.06 | 8.85E-01 |
| ENSMUSG00000037851 | GSMG0009750 | Iars | isoleucine-tRNA synthetase | 2432.45 | 2928.71 | -0.27 | 1.33E-01 |
| ENSMUSG00000037816 | GSMG0009756 | Fbxw17 | F-box and WD-40 domain protein 17 | 462.04 | 504.42 | -0.12 | 6.93E-01 |
| ENSMUSG00000021395 | GSMG0009761 | Spin1 | spindlin 1 | 315.11 | 383.69 | -0.28 | 3.01E-01 |
| ENSMUSG00000067586 | GSMG0009764 | S1pr3 | sphingosine-1-phosphate receptor 3 | 8009.75 | 8767.41 | -0.13 | 5.53E-01 |
| ENSMUSG00000035139 | GSMG0009767 | Secisbp2 | SECIS binding protein 2 | 613.47 | 613.49 | 0.00 | 1.00E+00 |
| ENSMUSG00000021453 | GSMG0009770 | Gadd45g | growth arrest and DNA-damage-inducible 45 gamma | 743.17 | 331.00 | 1.17 | 3.27E-08 |
| ENSMUSG00000021457 | GSMG0009774 | Syk | spleen tyrosine kinase | 478.56 | 542.67 | -0.18 | 4.79E-01 |
| ENSMUSG00000021474 | GSMG0009782 | Sfxn1 | sideroflexin 1 | 1009.88 | 1700.24 | -0.75 | 1.04E-06 |
| ENSMUSG00000025867 | GSMG0009784 | Cplx2 | complexin 2 | 405.04 | 285.03 | 0.50 | 5.80E-02 |
| ENSMUSG00000043183 | GSMG0009786 | Simc1 | SUMO-interacting motifs containing 1 | 130.94 | 239.72 | -0.88 | 3.14E-04 |
| ENSMUSG00000025870 | GSMG0009787 | Arl10 | ADP-ribosylation factor-like 10 | 245.15 | 348.89 | -0.51 | 7.16E-02 |
| ENSMUSG00000025868 | GSMG0009789 | Higd2a | HIG1 domain family, member 2A | 1180.30 | 1305.70 | -0.15 | 5.84E-01 |
| ENSMUSG00000025873 | GSMG0009790 | Faf2 | Fas associated factor family member 2 | 1858.51 | 2032.10 | -0.13 | 5.83E-01 |
| ENSMUSG00000025875 | GSMG0009794 | Tspan17 | tetraspanin 17 | 860.94 | 849.26 | 0.02 | 9.49E-01 |
| ENSMUSG00000025876 | GSMG0009796 | Unc5a | unc-5 netrin receptor A | 754.71 | 913.30 | -0.28 | 3.49E-01 |
| ENSMUSG00000021481 | GSMG0009797 | Zfp346 | zinc finger protein 346 | 111.26 | 102.88 | 0.12 | 8.10E-01 |
| ENSMUSG00000021488 | GSMG0009801 | Nsd1 | nuclear receptor-binding SET-domain protein 1 | 301.59 | 360.99 | -0.26 | 2.93E-01 |
| ENSMUSG00000021486 | GSMG0009805 | Prelid1 | PRELI domain containing 1 | 4119.55 | 4839.42 | -0.23 | 2.53E-01 |
| ENSMUSG00000052087 | GSMG0009806 | Rgs14 | regulator of G-protein signaling 14 | 216.73 | 115.67 | 0.90 | 2.09E-03 |
| ENSMUSG00000074886 | GSMG0009809 | Grk6 | G protein-coupled receptor kinase 6 | 1934.43 | 2295.75 | -0.25 | 2.23E-01 |
| ENSMUSG00000058569 | GSMG0009813 | Tmed9 | transmembrane emp24 protein transport domain containing 9 | 3796.59 | 4418.86 | -0.22 | 4.83E-01 |
| ENSMUSG00000021504 | GSMG0009814 | B4galt7 | xylosylprotein beta1,4-galactosyltransferase, polypeptide 7 (galactosylt | 143.61 | 177.14 | -0.30 | 3.51E-01 |
| ENSMUSG00000021501 | GSMG0009816 | Caml | calcium modulating ligand | 704.13 | 743.81 | -0.08 | 8.10E-01 |
| ENSMUSG00000021500 | GSMG0009817 | Ddx46 | DEAD (Asp-Glu-Ala-Asp) box polypeptide 46 | 1095.46 | 1209.19 | -0.14 | 5.30E-01 |
| ENSMUSG00000045767 | GSMG0009819 | B230219D22Rik | RIKEN cDNA B230219D22 gene | 3978.98 | 3751.27 | 0.08 | 7.15E-01 |
| ENSMUSG00000021497 | GSMG0009820 | Txndc15 | thioredoxin domain containing 15 | 1473.08 | 1665.86 | -0.18 | 4.30E-01 |

| | | | | | | | |
|---|---|---|---|---|---|---|---|
| ENSMUSG00000021496 | GSMG0009821 | Pcbd2 | pterin 4 alpha carbinolamine dehydratase/dimerization cofactor of hepat | 217.81 | 203.12 | 0.10 | 8.43E-01 |
| NULL | GSMG0009825 | NULL | NULL | 236.37 | 215.81 | 0.13 | 7.08E-01 |
| NULL | GSMG0009827 | NULL | NULL | 96.52 | 137.82 | -0.52 | 3.43E-01 |
| ENSMUSG00000035493 | GSMG0009836 | Tgfbi | transforming growth factor, beta induced | 290.54 | 152.48 | 0.93 | 5.43E-04 |
| ENSMUSG00000021540 | GSMG0009837 | Smad5 | SMAD family member 5 | 504.12 | 693.08 | -0.46 | 2.97E-02 |
| ENSMUSG00000050002 | GSMG0009841 | Idnk | idnK gluconokinase homolog (E. coli) | 225.49 | 234.52 | -0.06 | 8.88E-01 |
| ENSMUSG00000035367 | GSMG0009845 | Rmi1 | RecQ mediated genome instability 1 | 548.13 | 639.91 | -0.22 | 4.03E-01 |
| ENSMUSG00000055254 | GSMG0009846 | Ntrk2 | neurotrophic tyrosine kinase, receptor, type 2 | 2693.10 | 2657.99 | 0.02 | 9.51E-01 |
| NULL | GSMG0009847 | NULL | NULL | 475.68 | 276.67 | 0.78 | 5.57E-04 |
| NULL | GSMG0009848 | NULL | NULL | 821.09 | 429.83 | 0.94 | 3.79E-05 |
| ENSMUSG00000021555 | GSMG0009851 | Naa35 | N(alpha)-acetyltransferase 35, NatC auxiliary subunit | 1097.35 | 1319.82 | -0.27 | 1.54E-01 |
| ENSMUSG00000021559 | GSMG0009859 | Dapk1 | death associated protein kinase 1 | 2083.99 | 2570.28 | -0.30 | 1.83E-01 |
| NULL | GSMG0009860 | NULL | NULL | 253.73 | 310.87 | -0.30 | 4.37E-01 |
| NULL | GSMG0009861 | NULL | NULL | 599.30 | 501.69 | 0.26 | 2.42E-01 |
| ENSMUSG00000021458 | GSMG0009869 | 2010111I01Rik | RIKEN cDNA 2010111I01 gene | 3696.28 | 2305.15 | 0.68 | 1.02E-06 |
| NULL | GSMG0009873 | NULL | NULL | 375.27 | 302.50 | 0.31 | 1.95E-01 |
| NULL | GSMG0009874 | NULL | NULL | 274.98 | 191.41 | 0.53 | 8.74E-02 |
| ENSMUSG00000021476 | GSMG0009881 | Habp4 | hyaluronic acid binding protein 4 | 1321.39 | 893.99 | 0.56 | 4.74E-04 |
| ENSMUSG00000021483 | GSMG0009884 | Cdk20 | cyclin-dependent kinase 20 | 208.15 | 266.46 | -0.35 | 2.18E-01 |
| ENSMUSG00000021514 | GSMG0009888 | Zfp369 | zinc finger protein 369 | 212.72 | 213.32 | 0.00 | 9.96E-01 |
| ENSMUSG00000021518 | GSMG0009891 | Ptdss1 | phosphatidylserine synthase 1 | 1787.43 | 1917.92 | -0.10 | 7.22E-01 |
| ENSMUSG00000021532 | GSMG0009906 | Fastkd3 | FAST kinase domains 3 | 374.33 | 313.20 | 0.26 | 3.17E-01 |
| ENSMUSG00000021595 | GSMG0009911 | Nsun2 | NOL1/NOP2/Sun domain family member 2 | 1641.90 | 2302.92 | -0.49 | 1.28E-03 |
| ENSMUSG00000021598 | GSMG0009913 | Med10 | mediator complex subunit 10 | 395.60 | 537.44 | -0.44 | 4.84E-02 |
| ENSMUSG00000021607 | GSMG0009930 | Mrpl36 | mitochondrial ribosomal protein L36 | 553.64 | 605.47 | -0.13 | 6.63E-01 |
| ENSMUSG00000021608 | GSMG0009931 | Lpcat1 | lysophosphatidylcholine acyltransferase 1 | 1048.36 | 2113.98 | -1.01 | 1.24E-12 |
| ENSMUSG00000021610 | GSMG0009933 | Clptm1l | CLPTM1-like | 2735.90 | 3250.25 | -0.25 | 1.87E-01 |
| ENSMUSG00000017756 | GSMG0009936 | Slc12a7 | solute carrier family 12, member 7 | 724.73 | 795.58 | -0.13 | 6.54E-01 |
| ENSMUSG00000057649 | GSMG0009937 | Brd9 | bromodomain containing 9 | 2481.75 | 2250.33 | 0.14 | 5.30E-01 |
| ENSMUSG00000021573 | GSMG0009939 | Tppp | tubulin polymerization promoting protein | 1406.02 | 439.56 | 1.68 | 4.88E-20 |
| ENSMUSG00000021578 | GSMG0009942 | Ccdc127 | coiled-coil domain containing 127 | 594.97 | 732.72 | -0.30 | 1.66E-01 |
| ENSMUSG00000021583 | GSMG0009945 | Erap1 | endoplasmic reticulum aminopeptidase 1 | 1134.44 | 1048.73 | 0.11 | 6.81E-01 |
| ENSMUSG00000001542 | GSMG0009948 | Ell2 | elongation factor RNA polymerase II 2 | 155.29 | 170.34 | -0.14 | 7.36E-01 |

| | | | | | | | |
|---|---|---|---|---|---|---|---|
| ENSMUSG00000021591 | GSMG0009954 | Glrx | glutaredoxin | 1221.70 | 1137.36 | 0.10 | 7.07E-01 |
| ENSMUSG00000033991 | GSMG0009957 | Ttc37 | tetratricopeptide repeat domain 37 | 182.36 | 212.72 | -0.22 | 5.00E-01 |
| ENSMUSG00000021596 | GSMG0009960 | Mctp1 | multiple C2 domains, transmembrane 1 | 354.80 | 336.23 | 0.08 | 8.15E-01 |
| ENSMUSG00000071252 | GSMG0009963 | 2210408I21Rik | RIKEN cDNA 2210408I21 gene | 136.33 | 162.17 | -0.25 | 4.83E-01 |
| ENSMUSG00000064138 | GSMG0009965 | Fam172a | family with sequence similarity 172, member A | 441.74 | 525.27 | -0.25 | 3.16E-01 |
| ENSMUSG00000074794 | GSMG0009980 | Arrdc3 | arrestin domain containing 3 | 302.44 | 309.71 | -0.03 | 9.39E-01 |
| ENSMUSG00000035840 | GSMG0009985 | Lysmd3 | LysM, putative peptidoglycan-binding, domain containing 3 | 842.65 | 824.72 | 0.03 | 9.19E-01 |
| ENSMUSG00000051098 | GSMG0009987 | Mblac2 | metallo-beta-lactamase domain containing 2 | 582.14 | 829.72 | -0.51 | 8.72E-03 |
| ENSMUSG00000021537 | GSMG0009989 | Cetn3 | centrin 3 | 2403.05 | 3088.45 | -0.36 | 9.49E-02 |
| ENSMUSG00000005583 | GSMG0009991 | Mef2c | myocyte enhancer factor 2C | 2912.82 | 5494.26 | -0.91 | 2.40E-09 |
| NULL | GSMG0009992 | NULL | NULL | 615.83 | 995.70 | -0.69 | 1.33E-02 |
| NULL | GSMG0009993 | NULL | NULL | 213.86 | 305.87 | -0.52 | 1.47E-01 |
| NULL | GSMG0009994 | NULL | NULL | 270.59 | 456.31 | -0.75 | 2.94E-01 |
| NULL | GSMG0009995 | NULL | NULL | 324.55 | 542.55 | -0.74 | 3.31E-03 |
| NULL | GSMG0009997 | NULL | NULL | 213.86 | 398.66 | -0.90 | 1.23E-05 |
| ENSMUSG00000050334 | GSMG0009999 | C130071C03Rik | RIKEN cDNA C130071C03 gene | 518.40 | 949.83 | -0.88 | 6.86E-07 |
| ENSMUSG00000035762 | GSMG0010003 | Tmem161b | transmembrane protein 161B | 359.80 | 501.90 | -0.48 | 4.03E-02 |
| NULL | GSMG0010004 | NULL | NULL | 198.16 | 210.59 | -0.09 | 8.47E-01 |
| ENSMUSG00000021548 | GSMG0010009 | Ccnh | cyclin H | 444.30 | 509.96 | -0.20 | 4.84E-01 |
| ENSMUSG00000034488 | GSMG0010011 | Edil3 | EGF-like repeats and discoidin I-like domains 3 | 305.44 | 455.31 | -0.57 | 1.05E-02 |
| ENSMUSG00000021613 | GSMG0010019 | Hapln1 | hyaluronan and proteoglycan link protein 1 | 128.97 | 82.85 | 0.65 | 8.95E-02 |
| ENSMUSG00000012422 | GSMG0010024 | Tmem167 | transmembrane protein 167 | 2082.83 | 2169.31 | -0.06 | 8.28E-01 |
| ENSMUSG00000003992 | GSMG0010030 | Ssbp2 | single-stranded DNA binding protein 2 | 706.95 | 1261.45 | -0.83 | 7.01E-07 |
| ENSMUSG00000021707 | GSMG0010040 | Dhfr | dihydrofolate reductase | 153.25 | 258.65 | -0.76 | 6.55E-03 |
| ENSMUSG00000021703 | GSMG0010045 | Serinc5 | serine incorporator 5 | 1652.95 | 1107.38 | 0.58 | 5.49E-03 |
| ENSMUSG00000007617 | GSMG0010051 | Homer1 | homer scaffolding protein 1 | 169.17 | 204.05 | -0.27 | 3.74E-01 |
| NULL | GSMG0010056 | NULL | NULL | 434.00 | 276.86 | 0.65 | 1.87E-02 |
| ENSMUSG00000042082 | GSMG0010060 | Arsb | arylsulfatase B | 503.47 | 412.06 | 0.29 | 2.72E-01 |
| ENSMUSG00000045312 | GSMG0010064 | Lhfpl2 | lipoma HMGIC fusion partner-like 2 | 6885.72 | 7240.97 | -0.07 | 7.57E-01 |
| ENSMUSG00000021686 | GSMG0010066 | Ap3b1 | adaptor-related protein complex 3, beta 1 subunit | 1306.70 | 1340.45 | -0.04 | 8.95E-01 |
| ENSMUSG00000042043 | GSMG0010072 | Tbca | tubulin cofactor A | 800.89 | 1051.82 | -0.39 | 1.25E-01 |
| ENSMUSG00000042015 | GSMG0010075 | Wdr41 | WD repeat domain 41 | 268.23 | 293.15 | -0.13 | 6.98E-01 |
| NULL | GSMG0010076 | NULL | NULL | 655.18 | 442.80 | 0.56 | 8.12E-03 |

| | | | | | | | |
|---|---|---|---|---|---|---|---|
| ENSMUSG00000041995 | GSMG0010079 | Zbed3 | zinc finger, BED type containing 3 | 934.09 | 1159.05 | -0.31 | 1.24E-01 |
| ENSMUSG00000021671 | GSMG0010087 | Poc5 | POC5 centriolar protein | 378.71 | 500.44 | -0.40 | 5.58E-02 |
| ENSMUSG00000021669 | GSMG0010089 | Col4a3bp | collagen, type IV, alpha 3 (Goodpasture antigen) binding protein | 576.32 | 293.74 | 0.97 | 4.25E-08 |
| ENSMUSG00000041817 | GSMG0010095 | Fam169a | family with sequence similarity 169, member A | 84.54 | 145.62 | -0.78 | 1.20E-02 |
| ENSMUSG00000021666 | GSMG0010096 | Gfm2 | G elongation factor, mitochondrial 2 | 920.10 | 877.27 | 0.07 | 8.06E-01 |
| ENSMUSG00000041773 | GSMG0010097 | Enc1 | ectodermal-neural cortex 1 | 354.65 | 377.66 | -0.09 | 8.17E-01 |
| ENSMUSG00000021661 | GSMG0010107 | Ankra2 | ankyrin repeat, family A (RFXANK-like), 2 | 436.61 | 561.46 | -0.36 | 7.66E-02 |
| ENSMUSG00000048603 | GSMG0010108 | LOC105246056 // | uncharacterized LOC105246056 // predicted gene 9828 | 1312.50 | 1728.04 | -0.40 | 1.37E-01 |
| NULL | GSMG0010109 | NULL | NULL | 304.71 | 173.26 | 0.81 | 1.85E-02 |
| ENSMUSG00000050919 | GSMG0010115 | Zfp366 | zinc finger protein 366 | 3267.07 | 2353.11 | 0.47 | 1.71E-02 |
| ENSMUSG00000041632 | GSMG0010117 | Mrps27 | mitochondrial ribosomal protein S27 | 263.47 | 497.94 | -0.92 | 3.32E-07 |
| ENSMUSG00000021643 | GSMG0010120 | Serf1 | small EDRK-rich factor 1 | 196.97 | 704.92 | -1.85 | 2.50E-12 |
| ENSMUSG00000021645 | GSMG0010121 | Smn1 | survival motor neuron 1 | 245.98 | 372.14 | -0.60 | 6.81E-03 |
| G00000052293,ENSMUSG000 | GSMG0010123 | Ak6 // Taf9 | adenylate kinase 6 // TATA-box binding protein associated factor 9 | 102.65 | 146.86 | -0.52 | 1.08E-01 |
| ENSMUSG00000048924 | GSMG0010124 | Ccdc125 | coiled-coil domain containing 125 | 185.21 | 314.94 | -0.76 | 9.12E-04 |
| ENSMUSG00000042743 | GSMG0010140 | Sgtb | small glutamine-rich tetratricopeptide repeat (TPR)-containing, beta | 211.13 | 180.83 | 0.22 | 4.67E-01 |
| ENSMUSG00000021712 | GSMG0010141 | Trim23 | tripartite motif-containing 23 | 339.28 | 384.20 | -0.18 | 5.16E-01 |
| ENSMUSG00000021714 | GSMG0010142 | Cenpk | centromere protein K | 42.33 | 110.46 | -1.38 | 3.76E-05 |
| ENSMUSG00000021716 | GSMG0010148 | Srek1ip1 | splicing regulatory glutamine/lysine-rich protein 1interacting protein 1 | 312.22 | 337.54 | -0.12 | 7.91E-01 |
| NULL | GSMG0010149 | NULL | NULL | 250.74 | 194.08 | 0.37 | 1.64E-01 |
| ENSMUSG00000021692 | GSMG0010156 | Dimt1 | DIM1 dimethyladenosine transferase 1-like (S. cerevisiae) | 353.44 | 312.90 | 0.18 | 5.48E-01 |
| ENSMUSG00000071181 | GSMG0010157 | 3830408C21Rik | RIKEN cDNA 3830408C21 gene | 224.28 | 385.06 | -0.77 | 4.98E-04 |
| ENSMUSG00000071180 | GSMG0010163 | Smim15 | small integral membrane protein 15 | 944.63 | 862.79 | 0.13 | 6.53E-01 |
| ENSMUSG00000021694 | GSMG0010164 | Ercc8 | excision repaiross-complementing rodent repair deficiency, complemen | 204.32 | 284.01 | -0.47 | 5.35E-02 |
| ENSMUSG00000021696 | GSMG0010166 | Elovl7 | ELOVL family member 7, elongation of long chain fatty acids (yeast) | 517.71 | 260.12 | 1.00 | 5.57E-06 |
| ENSMUSG00000021697 | GSMG0010168 | Depdc1b | DEP domain containing 1B | 166.10 | 283.03 | -0.77 | 2.82E-03 |
| NULL | GSMG0010193 | NULL | NULL | 476.81 | 623.69 | -0.39 | 1.54E-01 |
| ENSMUSG00000021701 | GSMG0010199 | Plk2 | polo-like kinase 2 | 6049.10 | 4273.82 | 0.50 | 2.56E-03 |
| ENSMUSG00000032727 | GSMG0010204 | Mier3 | MIER family member 3 | 217.11 | 183.62 | 0.24 | 4.68E-01 |
| ENSMUSG00000021756 | GSMG0010210 | Il6st | interleukin 6 signal transducer | 6004.91 | 4284.45 | 0.49 | 3.34E-03 |
| NULL | GSMG0010212 | NULL | NULL | 280.13 | 267.21 | 0.07 | 8.77E-01 |
| ENSMUSG00000021759 | GSMG0010215 | Plpp1 | phospholipid phosphatase 1 | 949.91 | 750.83 | 0.34 | 1.65E-01 |
| ENSMUSG00000042426 | GSMG0010217 | Dhx29 | DEAH (Asp-Glu-Ala-His) box polypeptide 29 | 223.69 | 257.98 | -0.21 | 5.23E-01 |

| | | | | | | | |
|---|---|---|---|---|---|---|---|
| NULL | GSMG0010225 | NULL | NULL | 196.18 | 95.09 | 1.04 | 3.61E-05 |
| ENSMUSG00000042348 | GSMG0010227 | Arl15 | ADP-ribosylation factor-like 15 | 1867.05 | 1717.29 | 0.12 | 5.83E-01 |
| NULL | GSMG0010228 | NULL | NULL | 553.33 | 409.58 | 0.44 | 5.55E-02 |
| NULL | GSMG0010229 | NULL | NULL | 294.77 | 212.75 | 0.47 | 4.16E-02 |
| NULL | GSMG0010232 | NULL | NULL | 328.56 | 246.99 | 0.41 | 1.57E-01 |
| NULL | GSMG0010234 | NULL | NULL | 253.31 | 240.21 | 0.07 | 8.45E-01 |
| NULL | GSMG0010237 | NULL | NULL | 662.89 | 473.16 | 0.49 | 3.42E-02 |
| NULL | GSMG0010238 | NULL | NULL | 317.79 | 293.86 | 0.11 | 7.34E-01 |
| ENSMUSG00000015536 | GSMG0010244 | Mocs2 | molybdenum cofactor synthesis 2 | 1122.58 | 785.43 | 0.51 | 4.89E-03 |
| ENSMUSG00000021728 | GSMG0010249 | Emb | embigin | 101.28 | 96.34 | 0.07 | 9.01E-01 |
| ENSMUSG00000021730 | GSMG0010251 | Hcn1 | hyperpolarization-activated, cyclic nucleotide-gated K+ 1 | 191.73 | 225.24 | -0.23 | 5.67E-01 |
| ENSMUSG00000025451 | GSMG0010267 | Paip1 | polyadenylate binding protein-interacting protein 1 | 81.68 | 104.58 | -0.36 | 3.51E-01 |
| ENSMUSG00000062822 | GSMG0010269 | 4833420G17Rik | RIKEN cDNA 4833420G17 gene | 2075.07 | 2099.87 | -0.02 | 9.49E-01 |
| ENSMUSG00000033799 | GSMG0010270 | Fam208b | family with sequence similarity 208, member B | 239.55 | 292.62 | -0.29 | 2.63E-01 |
| ENSMUSG00000021215 | GSMG0010273 | Net1 | neuroepithelial cell transforming gene 1 | 2150.23 | 1439.47 | 0.58 | 4.12E-03 |
| ENSMUSG00000045410 | GSMG0010281 | Akr1e1 | aldo-keto reductase family 1, member E1 | 244.80 | 217.28 | 0.17 | 7.12E-01 |
| ENSMUSG00000021196 | GSMG0010285 | Pfkp | phosphofructokinase, platelet | 4883.04 | 16772.86 | -1.78 | 2.77E-45 |
| ENSMUSG00000021147 | GSMG0010290 | Wdr37 | WD repeat domain 37 | 720.31 | 711.37 | 0.02 | 9.55E-01 |
| ENSMUSG00000021149 | GSMG0010294 | Gtpbp4 | GTP binding protein 4 | 826.98 | 976.88 | -0.24 | 3.02E-01 |
| ENSMUSG00000021156 | GSMG0010297 | Zmynd11 | zinc finger, MYND domain containing 11 | 6026.61 | 6566.49 | -0.12 | 5.65E-01 |
| ENSMUSG00000021313 | GSMG0010306 | Ryr2 | ryanodine receptor 2, cardiac | 650.68 | 416.03 | 0.65 | 4.37E-02 |
| ENSMUSG00000021311 | GSMG0010312 | Mtr | 5-methyltetrahydrofolate-homocysteine methyltransferase | 227.75 | 305.60 | -0.42 | 1.03E-01 |
| ENSMUSG00000052374 | GSMG0010313 | Actn2 | actinin alpha 2 | 114.46 | 143.58 | -0.33 | 3.69E-01 |
| ENSMUSG00000057554 | GSMG0010314 | Lgals8 | lectin, galactose binding, soluble 8 | 592.65 | 561.23 | 0.08 | 8.04E-01 |
| ENSMUSG00000021302 | GSMG0010326 | Ggps1 | geranylgeranyl diphosphate synthase 1 | 486.67 | 489.11 | -0.01 | 9.85E-01 |
| ENSMUSG00000039233 | GSMG0010327 | Tbce | tubulin-specific chaperone E | 1785.85 | 2118.21 | -0.25 | 1.61E-01 |
| ENSMUSG00000021301 | GSMG0010329 | Hecw1 | HECT, C2 and WW domain containing E3 ubiquitin protein ligase 1 | 597.17 | 1007.44 | -0.75 | 2.13E-05 |
| ENSMUSG00000015672 | GSMG0010335 | Mrpl32 | mitochondrial ribosomal protein L32 | 553.24 | 598.12 | -0.11 | 7.78E-01 |
| NULL | GSMG0010346 | NULL | NULL | 256.21 | 136.31 | 0.91 | 5.58E-03 |
| ENSMUSG00000041297 | GSMG0010351 | Cdk13 | cyclin-dependent kinase 13 | 1203.12 | 1022.34 | 0.23 | 5.07E-01 |
| ENSMUSG00000008859 | GSMG0010358 | Rala | v-ral simian leukemia viral oncogene A (ras related) | 1360.76 | 2011.64 | -0.56 | 3.32E-04 |
| ENSMUSG00000003062 | GSMG0010367 | Stard3nl | STARD3 N-terminal like | 1540.98 | 1308.68 | 0.24 | 2.93E-01 |
| ENSMUSG00000002808 | GSMG0010370 | Epdr1 | ependymin related protein 1 (zebrafish) | 1309.71 | 962.98 | 0.44 | 5.73E-03 |

| | | | | | | | |
|---|---|---|---|---|---|---|---|
| ENSMUSG00000021327 | GSMG0010379 | Zkscan3 | zinc finger with KRAB and SCAN domains 3 | 700.41 | 662.99 | 0.08 | 7.79E-01 |
| ENSMUSG00000022228 | GSMG0010381 | Zscan26 | zinc finger and SCAN domain containing 26 | 818.24 | 701.96 | 0.22 | 3.61E-01 |
| ENSMUSG00000058773 | GSMG0010397 | Hist1h1b | histone cluster 1, H1b | 23.46 | 83.36 | -1.85 | 7.84E-06 |
| ENSMUSG00000046351 | GSMG0010415 | Zfp322a | zinc finger protein 322A | 1601.89 | 1666.29 | -0.06 | 8.49E-01 |
| ENSMUSG00000036376 | GSMG0010416 | Abt1 | activator of basal transcription 1 | 641.80 | 693.50 | -0.11 | 7.11E-01 |
| ENSMUSG00000006611 | GSMG0010431 | Hfe | hemochromatosis | 416.14 | 495.03 | -0.25 | 3.83E-01 |
| ENSMUSG00000021338 | GSMG0010439 | Lrrc16a | leucine rich repeat containing 16A | 474.30 | 695.09 | -0.55 | 8.21E-03 |
| ENSMUSG00000006715 | GSMG0010448 | Gmnn | geminin | 219.27 | 399.59 | -0.86 | 1.01E-03 |
| ENSMUSG00000006717 | GSMG0010450 | Acot13 | acyl-CoA thioesterase 13 | 567.75 | 408.09 | 0.48 | 2.25E-02 |
| ENSMUSG00000035936 | GSMG0010451 | Aldh5a1 | aldhehyde dehydrogenase family 5, subfamily A1 | 144.21 | 166.01 | -0.20 | 6.06E-01 |
| NULL | GSMG0010452 | NULL | NULL | 1425.32 | 1049.73 | 0.44 | 1.78E-02 |
| ENSMUSG00000021339 | GSMG0010453 | Mrs2 | MRS2 magnesium transporter | 894.91 | 721.96 | 0.31 | 1.30E-01 |
| ENSMUSG00000048978 | GSMG0010454 | Nrsn1 | neurensin 1 | 2153.80 | 899.97 | 1.26 | 6.34E-15 |
| NULL | GSMG0010471 | NULL | NULL | 2507.17 | 5371.55 | -1.10 | 3.96E-10 |
| NULL | GSMG0010482 | NULL | NULL | 163.44 | 245.65 | -0.59 | 4.07E-02 |
| ENSMUSG00000076431 | GSMG0010488 | Sox4 | SRY (sex determining region Y)-box 4 | 4068.18 | 6625.70 | -0.70 | 2.88E-04 |
| ENSMUSG00000006191 | GSMG0010489 | Cdkal1 | CDK5 regulatory subunit associated protein 1-like 1 | 685.47 | 714.63 | -0.06 | 8.55E-01 |
| ENSMUSG00000016477 | GSMG0010494 | E2f3 | E2F transcription factor 3 | 90.06 | 119.79 | -0.42 | 2.97E-01 |
| ENSMUSG00000038462 | GSMG0010496 | Uqcrfs1 | ubiquinol-cytochrome c reductase, Rieske iron-sulfur polypeptide 1 | 2532.75 | 2764.47 | -0.13 | 5.80E-01 |
| NULL | GSMG0010497 | NULL | NULL | 58.77 | 52.38 | 0.16 | 8.23E-01 |
| ENSMUSG00000021357 | GSMG0010500 | Exoc2 | exocyst complex component 2 | 548.66 | 611.35 | -0.16 | 5.55E-01 |
| NULL | GSMG0010507 | NULL | NULL | 1184.27 | 1272.15 | -0.10 | 7.26E-01 |
| ENSMUSG00000038408 | GSMG0010508 | 1700018A04Rik | RIKEN cDNA 1700018A04 gene | 528.29 | 431.69 | 0.29 | 2.39E-01 |
| ENSMUSG00000038372 | GSMG0010509 | Gmds | GDP-mannose 4, 6-dehydratase | 249.88 | 353.56 | -0.50 | 3.03E-02 |
| ENSMUSG00000044734 | GSMG0010518 | Serpinb1a | serine (or cysteine) peptidase inhibitor, clade B, member 1a | 72.30 | 36.93 | 0.97 | 2.15E-02 |
| ENSMUSG00000038267 | GSMG0010530 | Slc22a23 | solute carrier family 22, member 23 | 150.61 | 306.93 | -1.02 | 2.18E-04 |
| NULL | GSMG0010532 | NULL | NULL | 215.51 | 184.44 | 0.23 | 5.39E-01 |
| ENSMUSG00000021417 | GSMG0010538 | Eci2 | enoyl-Coenzyme A delta isomerase 2 | 2728.73 | 3011.42 | -0.14 | 5.10E-01 |
| ENSMUSG00000046573 | GSMG0010545 | Lyrm4 | LYR motif containing 4 | 101.50 | 147.24 | -0.54 | 1.32E-01 |
| ENSMUSG00000039114 | GSMG0010550 | Nrn1 | neuritin 1 | 566.56 | 71.89 | 2.98 | 6.02E-45 |
| ENSMUSG00000039109 | GSMG0010552 | F13a1 | coagulation factor XIII, A1 subunit | 213.70 | 38.69 | 2.47 | 1.74E-17 |
| ENSMUSG00000021427 | GSMG0010558 | Ssr1 | signal sequence receptor, alpha | 2269.73 | 2984.53 | -0.40 | 6.02E-02 |
| ENSMUSG00000038991 | GSMG0010563 | Txndc5 | thioredoxin domain containing 5 | 3912.86 | 4261.83 | -0.12 | 6.48E-01 |

| | | | | | | | |
|---|---|---|---|---|---|---|---|
| ENSMUSG00000038982 | GSMG0010564 | Bloc1s5 | biogenesis of lysosomal organelles complex-1, subunit 5, muted | 684.94 | 860.25 | -0.33 | 9.02E-02 |
| ENSMUSG00000001707 | GSMG0010565 | Eef1e1 | eukaryotic translation elongation factor 1 epsilon 1 | 250.11 | 461.86 | -0.88 | 1.78E-05 |
| ENSMUSG00000021432 | GSMG0010567 | Slc35b3 | solute carrier family 35, member B3 | 814.47 | 917.20 | -0.17 | 4.63E-01 |
| ENSMUSG00000021364 | GSMG0010578 | Elovl2 | elongation of very long chain fatty acids (FEN1/Elo2, SUR4/Elo3, yeas | 110.20 | 148.99 | -0.43 | 1.95E-01 |
| ENSMUSG00000021365 | GSMG0010579 | Nedd9 | neural precursor cell expressed, developmentally down-regulated gene 9 | 1456.35 | 1265.53 | 0.20 | 3.70E-01 |
| ENSMUSG00000021368 | GSMG0010587 | Tbc1d7 | TBC1 domain family, member 7 | 358.73 | 395.81 | -0.14 | 6.73E-01 |
| NULL | GSMG0010589 | NULL | NULL | 321.68 | 173.38 | 0.89 | 3.69E-05 |
| NULL | GSMG0010590 | NULL | NULL | 277.09 | 363.93 | -0.40 | 1.23E-01 |
| ENSMUSG00000038546 | GSMG0010591 | Ranbp9 | RAN binding protein 9 | 519.49 | 219.54 | 1.24 | 3.53E-05 |
| ENSMUSG00000021371 | GSMG0010595 | Mcur1 | mitochondrial calcium uniporter regulator 1 | 1952.13 | 1567.53 | 0.32 | 7.51E-02 |
| ENSMUSG00000057531 | GSMG0010599 | Dtnbp1 | dystrobrevin binding protein 1 | 959.36 | 1040.44 | -0.12 | 6.44E-01 |
| ENSMUSG00000046876 | GSMG0010603 | Atxn1 | ataxin 1 | 886.93 | 1083.78 | -0.29 | 2.10E-01 |
| NULL | GSMG0010604 | NULL | NULL | 220.14 | 248.97 | -0.17 | 6.59E-01 |
| NULL | GSMG0010607 | NULL | NULL | 117.38 | 130.43 | -0.15 | 7.62E-01 |
| NULL | GSMG0010610 | NULL | NULL | 268.37 | 357.86 | -0.42 | 1.29E-01 |
| ENSMUSG00000021374 | GSMG0010612 | Nup153 | nucleoporin 153 | 258.32 | 314.01 | -0.28 | 3.36E-01 |
| ENSMUSG00000021375 | GSMG0010613 | Kif13a | kinesin family member 13A | 384.08 | 284.15 | 0.43 | 2.29E-01 |
| ENSMUSG00000044231 | GSMG0010617 | Nhlrc1 | NHL repeat containing 1 | 239.94 | 252.69 | -0.08 | 8.44E-01 |
| ENSMUSG00000021376 | GSMG0010618 | Tpmt | thiopurine methyltransferase | 161.31 | 226.90 | -0.49 | 6.88E-02 |
| ENSMUSG00000021377 | GSMG0010619 | Dek | DEK oncogene (DNA binding) | 1468.76 | 2203.04 | -0.59 | 1.20E-02 |
| ENSMUSG00000038042 | GSMG0010631 | Ptpdc1 | protein tyrosine phosphatase domain containing 1 | 146.78 | 118.92 | 0.30 | 4.25E-01 |
| ENSMUSG00000038014 | GSMG0010632 | Fam120a | family with sequence similarity 120, member A | 1604.27 | 852.18 | 0.91 | 1.55E-01 |
| ENSMUSG00000038025 | GSMG0010633 | Phf2 | PHD finger protein 2 | 178.76 | 110.91 | 0.69 | 4.83E-01 |
| NULL | GSMG0010636 | NULL | NULL | 667.86 | 542.26 | 0.30 | 1.90E-01 |
| ENSMUSG00000037989 | GSMG0010638 | Wnk2 | WNK lysine deficient protein kinase 2 | 546.00 | 584.63 | -0.10 | 7.90E-01 |
| ENSMUSG00000037960 | GSMG0010639 | Card19 | caspase recruitment domain family, member 19 | 1024.58 | 726.79 | 0.49 | 8.84E-03 |
| ENSMUSG00000021391 | GSMG0010645 | Cenpp | centromere protein P | 60.05 | 136.75 | -1.18 | 8.54E-04 |
| NULL | GSMG0010655 | 9430083A17Rik | RIKEN cDNA 9430083A17 gene | 520.99 | 663.40 | -0.35 | 1.17E-01 |
| NULL | GSMG0010659 | NULL | NULL | 369.68 | 368.90 | 0.00 | 9.91E-01 |
| NULL | GSMG0010660 | NULL | NULL | 373.89 | 369.86 | 0.02 | 9.70E-01 |
| ENSMUSG00000021448 | GSMG0010661 | Shc3 | src homology 2 domain-containing transforming protein C3 | 367.16 | 343.31 | 0.10 | 8.26E-01 |
| ENSMUSG00000021451 | GSMG0010663 | Sema4d | sema domain, immunoglobulin domain (Ig), transmembrane domain (T | 389.05 | 245.22 | 0.67 | 2.79E-02 |
| ENSMUSG00000047842 | GSMG0010666 | Diras2 | DIRAS family, GTP-binding RAS-like 2 | 1066.56 | 934.97 | 0.19 | 4.35E-01 |

| | | | | | | | |
|---|---|---|---|---|---|---|---|
| ENSMUSG00000021460 | GSMG0010669 | Auh | AU RNA binding protein/enoyl-coenzyme A hydratase | 516.60 | 419.89 | 0.30 | 1.70E-01 |
| ENSMUSG00000056749 | GSMG0010671 | Nfil3 | nuclear factor, interleukin 3, regulated | 190.66 | 360.97 | -0.92 | 3.64E-05 |
| ENSMUSG00000021468 | GSMG0010675 | Sptlc1 | serine palmitoyltransferase, long chain base subunit 1 | 651.27 | 757.33 | -0.22 | 3.62E-01 |
| ENSMUSG00000021478 | GSMG0010680 | Drd1 | dopamine receptor D1 | 342.76 | 357.22 | -0.06 | 8.85E-01 |
| ENSMUSG00000025872 | GSMG0010684 | Thoc3 | THO complex 3 | 529.38 | 747.56 | -0.50 | 4.03E-03 |
| ENSMUSG00000025871 | GSMG0010685 | 4833439L19Rik | RIKEN cDNA 4833439L19 gene | 1954.20 | 1983.93 | -0.02 | 9.41E-01 |
| ENSMUSG00000025869 | GSMG0010687 | Nop16 | NOP16 nucleolar protein | 665.78 | 1028.80 | -0.63 | 6.67E-04 |
| ENSMUSG00000047547 | GSMG0010688 | Cltb | clathrin, light polypeptide (Lcb) | 4166.33 | 3229.96 | 0.37 | 3.06E-02 |
| ENSMUSG00000034928 | GSMG0010689 | Rnf44 | ring finger protein 44 | 596.15 | 655.91 | -0.14 | 7.39E-01 |
| ENSMUSG00000069227 | GSMG0010691 | Gprin1 | G protein-regulated inducer of neurite outgrowth 1 | 754.21 | 1480.84 | -0.97 | 1.02E-04 |
| ENSMUSG00000034891 | GSMG0010692 | Sncb | synuclein, beta | 717.28 | 476.78 | 0.59 | 7.56E-02 |
| ENSMUSG00000025878 | GSMG0010696 | Uimc1 | ubiquitin interaction motif containing 1 | 876.28 | 896.23 | -0.03 | 9.21E-01 |
| ENSMUSG00000034789 | GSMG0010698 | Rab24 | RAB24, member RAS oncogene family | 2833.15 | 1958.17 | 0.53 | 2.66E-04 |
| ENSMUSG00000021485 | GSMG0010699 | Mxd3 | Max dimerization protein 3 | 173.21 | 286.62 | -0.72 | 7.62E-03 |
| ENSMUSG00000021484 | GSMG0010700 | Lman2 | lectin, mannose-binding 2 | 2066.72 | 2158.91 | -0.06 | 8.10E-01 |
| ENSMUSG00000034675 | GSMG0010704 | Dbn1 | drebrin 1 | 5113.43 | 7068.73 | -0.47 | 2.00E-03 |
| ENSMUSG00000021493 | GSMG0010705 | Pdlim7 | PDZ and LIM domain 7 | 1593.32 | 1444.91 | 0.14 | 5.79E-01 |
| ENSMUSG00000035711 | GSMG0010706 | Dok3 | docking protein 3 | 63.14 | 46.16 | 0.45 | 3.49E-01 |
| ENSMUSG00000021494 | GSMG0010707 | Ddx41 | DEAD (Asp-Glu-Ala-Asp) box polypeptide 41 | 1791.42 | 1939.71 | -0.11 | 6.25E-01 |
| ENSMUSG00000021495 | GSMG0010708 | Fam193b | family with sequence similarity 193, member B | 2437.30 | 2009.54 | 0.28 | 2.43E-01 |
| NULL | GSMG0010709 | NULL | NULL | 312.25 | 324.17 | -0.06 | 8.90E-01 |
| NULL | GSMG0010710 | NULL | NULL | 238.30 | 231.33 | 0.04 | 9.38E-01 |
| ENSMUSG00000015937 | GSMG0010714 | H2afy | H2A histone family, member Y | 3190.74 | 4269.13 | -0.42 | 1.21E-02 |
| ENSMUSG00000021508 | GSMG0010718 | Cxcl14 | chemokine (C-X-C motif) ligand 14 | 842.99 | 769.94 | 0.13 | 6.66E-01 |
| ENSMUSG00000056222 | GSMG0010723 | Spock1 | sparc/osteonectin, cwcv and kazal-like domains proteoglycan 1 | 400.83 | 626.68 | -0.65 | 1.05E-02 |
| NULL | GSMG0010725 | NULL | NULL | 835.28 | 679.56 | 0.30 | 1.97E-01 |
| NULL | GSMG0010726 | NULL | NULL | 661.86 | 854.21 | -0.37 | 2.71E-01 |
| ENSMUSG00000007836 | GSMG0010728 | Hnrnpa0 | heterogeneous nuclear ribonucleoprotein A0 | 414.88 | 675.23 | -0.71 | 2.60E-03 |
| ENSMUSG00000005312 | GSMG0010730 | Ubqln1 | ubiquilin 1 | 2101.01 | 2534.08 | -0.27 | 1.21E-01 |
| ENSMUSG00000021552 | GSMG0010731 | Gkap1 | G kinase anchoring protein 1 | 300.60 | 302.33 | -0.01 | 9.81E-01 |
| ENSMUSG00000021550 | GSMG0010733 | 2210016F16Rik | RIKEN cDNA 2210016F16 gene | 323.81 | 292.80 | 0.15 | 6.58E-01 |
| ENSMUSG00000021546 | GSMG0010734 | Hnrnpk | heterogeneous nuclear ribonucleoprotein K | 6363.72 | 7378.61 | -0.21 | 3.04E-01 |
| NULL | GSMG0010737 | NULL | NULL | 95.88 | 63.32 | 0.60 | 1.58E-01 |

| | | | | | | | |
|---|---|---|---|---|---|---|---|
| NULL | GSMG0010742 | NULL | NULL | 137.85 | 161.88 | -0.23 | 5.46E-01 |
| ENSMUSG00000021557 | GSMG0010743 | Agtpbp1 | ATP/GTP binding protein 1 | 681.55 | 820.29 | -0.27 | 1.76E-01 |
| ENSMUSG00000021556 | GSMG0010747 | Golm1 | golgi membrane protein 1 | 2002.80 | 1951.31 | 0.04 | 9.01E-01 |
| ENSMUSG00000035248 | GSMG0010753 | Zcchc6 | zinc finger, CCHC domain containing 6 | 894.62 | 906.14 | -0.02 | 9.53E-01 |
| NULL | GSMG0010755 | NULL | NULL | 216.61 | 274.44 | -0.34 | 2.03E-01 |
| NULL | GSMG0010757 | NULL | NULL | 548.06 | 463.28 | 0.24 | 2.96E-01 |
| ENSMUSG00000050345 | GSMG0010760 | 4930486L24Rik | RIKEN cDNA 4930486L24 gene | 113.91 | 75.32 | 0.60 | 1.43E-01 |
| ENSMUSG00000074874 | GSMG0010761 | Ctla2b | cytotoxic T lymphocyte-associated protein 2 beta | 807.61 | 609.46 | 0.41 | 4.01E-02 |
| ENSMUSG00000044258 | GSMG0010763 | Ctla2a | cytotoxic T lymphocyte-associated protein 2 alpha | 12057.80 | 9930.38 | 0.28 | 9.58E-02 |
| NULL | GSMG0010779 | NULL | NULL | 132.19 | 132.51 | -0.01 | 9.90E-01 |
| ENSMUSG00000021461 | GSMG0010789 | Fancc | Fanconi anemia, complementation group C | 245.12 | 259.57 | -0.08 | 8.23E-01 |
| ENSMUSG00000097242 | GSMG0010790 | Gm16907 | predicted gene, 16907 | 487.01 | 313.13 | 0.64 | 1.77E-03 |
| ENSMUSG00000021466 | GSMG0010792 | Ptch1 | patched 1 | 117.11 | 138.18 | -0.24 | 5.84E-01 |
| ENSMUSG00000033114 | GSMG0010798 | Slc35d2 | solute carrier family 35, member D2 | 159.85 | 120.56 | 0.40 | 1.96E-01 |
| ENSMUSG00000021482 | GSMG0010801 | Aaed1 | AhpC/TSA antioxidant enzyme domain containing 1 | 515.59 | 423.00 | 0.28 | 2.05E-01 |
| ENSMUSG00000021477 | GSMG0010802 | Ctsl | cathepsin L | 6141.77 | 5592.37 | 0.14 | 5.43E-01 |
| ENSMUSG00000038212 | GSMG0010806 | Mfsd14b | major facilitator superfamily domain containing 14B | 2117.03 | 1828.85 | 0.21 | 3.00E-01 |
| ENSMUSG00000021520 | GSMG0010813 | Uqcrb | ubiquinol-cytochrome c reductase binding protein | 2187.62 | 2368.91 | -0.12 | 6.66E-01 |
| ENSMUSG00000021519 | GSMG0010814 | Mterf3 | mitochondrial transcription termination factor 3 | 775.33 | 757.92 | 0.03 | 9.34E-01 |
| ENSMUSG00000059839 | GSMG0010826 | Zfp874b | zinc finger protein 874b | 131.58 | 117.37 | 0.17 | 6.86E-01 |
| G00000071291,ENSMUSG000 | GSMG0010827 | Zfp58 // Zfp87 | zinc finger protein 58 // zinc finger protein 87 | 953.80 | 854.44 | 0.16 | 4.79E-01 |
| ENSMUSG00000095432 | GSMG0010828 | Zfp748 | zinc finger protein 748 | 439.14 | 428.84 | 0.04 | 9.25E-01 |
| ENSMUSG00000034617 | GSMG0010840 | Mtrr | 5-methyltetrahydrofolate-homocysteine methyltransferase reductase | 357.03 | 435.26 | -0.28 | 2.78E-01 |
| ENSMUSG00000021536 | GSMG0010841 | Adcy2 | adenylate cyclase 2 | 2650.20 | 1161.34 | 1.19 | 2.42E-10 |
| ENSMUSG00000021594 | GSMG0010847 | Srd5a1 | steroid 5 alpha-reductase 1 | 216.64 | 359.68 | -0.73 | 1.72E-03 |
| ENSMUSG00000052981 | GSMG0010850 | Ube2ql1 | ubiquitin-conjugating enzyme E2Q family-like 1 | 736.57 | 721.06 | 0.03 | 9.48E-01 |
| ENSMUSG00000049538 | GSMG0010856 | Adamts16 | a disintegrin-like and metallopeptidase (reprolysin type) with thrombos | 675.24 | 487.28 | 0.47 | 2.69E-02 |
| ENSMUSG00000021606 | GSMG0010865 | Ndufs6 | NADH dehydrogenase (ubiquinone) Fe-S protein 6 | 1356.97 | 1548.46 | -0.19 | 3.75E-01 |
| ENSMUSG00000021567 | GSMG0010869 | Nkd2 | naked cuticle 2 homolog (Drosophila) | 423.52 | 385.64 | 0.13 | 6.40E-01 |
| ENSMUSG00000021569 | GSMG0010871 | Trip13 | thyroid hormone receptor interactor 13 | 38.13 | 117.06 | -1.61 | 6.51E-07 |
| ENSMUSG00000021572 | GSMG0010873 | Cep72 | centrosomal protein 72 | 137.74 | 156.71 | -0.19 | 6.44E-01 |
| ENSMUSG00000034152 | GSMG0010875 | Exoc3 | exocyst complex component 3 | 1433.02 | 1729.80 | -0.27 | 1.79E-01 |
| ENSMUSG00000021576 | GSMG0010878 | Pdcd6 | programmed cell death 6 | 1759.76 | 1542.58 | 0.19 | 3.82E-01 |

| | | | | | | | |
|---|---|---|---|---|---|---|---|
| ENSMUSG00000021577 | GSMG0010880 | Sdha | succinate dehydrogenase complex, subunit A, flavoprotein (Fp) | 5265.92 | 5197.13 | 0.02 | 9.51E-01 |
| ENSMUSG00000069208 | GSMG0010885 | Zfp825 | zinc finger protein 825 | 263.40 | 219.16 | 0.27 | 3.36E-01 |
| ENSMUSG00000021585 | GSMG0010888 | Cast | calpastatin | 1378.48 | 1148.99 | 0.26 | 2.83E-01 |
| ENSMUSG00000021589 | GSMG0010892 | Rhobtb3 | Rho-related BTB domain containing 3 | 351.23 | 394.72 | -0.17 | 5.53E-01 |
| ENSMUSG00000043190 | GSMG0010894 | Rfesd | Rieske (Fe-S) domain containing | 152.36 | 169.12 | -0.15 | 6.98E-01 |
| ENSMUSG00000021592 | GSMG0010896 | Arsk | arylsulfatase K | 210.17 | 150.10 | 0.49 | 8.42E-02 |
| ENSMUSG00000069171 | GSMG0010903 | Nr2f1 | nuclear receptor subfamily 2, group F, member 1 | 154.45 | 236.11 | -0.61 | 1.02E-02 |
| ENSMUSG00000035834 | GSMG0010909 | Polr3g | polymerase (RNA) III (DNA directed) polypeptide G | 44.32 | 54.42 | -0.30 | 6.17E-01 |
| NULL | GSMG0010912 | NULL | NULL | 579.38 | 945.04 | -0.71 | 8.51E-03 |
| NULL | GSMG0010913 | NULL | NULL | 128.26 | 202.36 | -0.65 | 2.04E-02 |
| ENSMUSG00000098087 | GSMG0010916 | Gm17750 | predicted gene, 17750 | 66.90 | 193.79 | -1.54 | 9.63E-09 |
| ENSMUSG00000021549 | GSMG0010919 | Rasa1 | RAS p21 protein activator 1 | 2061.22 | 2124.52 | -0.04 | 8.72E-01 |
| ENSMUSG00000021614 | GSMG0010923 | Vcan | versican | 31.97 | 130.00 | -2.01 | 1.78E-08 |
| ENSMUSG00000021615 | GSMG0010925 | Xrcc4 | X-ray repair complementing defective repair in Chinese hamster cells 4 | 245.08 | 259.39 | -0.08 | 8.58E-01 |
| NULL | GSMG0010935 | NULL | NULL | 114.39 | 126.64 | -0.15 | 7.98E-01 |
| ENSMUSG00000021621 | GSMG0010940 | Zcchc9 | zinc finger, CCHC domain containing 9 | 383.74 | 442.51 | -0.21 | 4.90E-01 |
| ENSMUSG00000014850 | GSMG0010943 | Msh3 | mutS homolog 3 | 183.10 | 211.05 | -0.20 | 5.75E-01 |
| ENSMUSG00000021706 | GSMG0010946 | Zfyve16 | zinc finger, FYVE domain containing 16 | 529.19 | 527.05 | 0.01 | 9.82E-01 |
| ENSMUSG00000042167 | GSMG0010953 | Papd4 | PAP associated domain containing 4 | 622.43 | 745.54 | -0.26 | 2.24E-01 |
| ENSMUSG00000021690 | GSMG0010957 | Jmy | junction-mediating and regulatory protein | 160.30 | 188.64 | -0.24 | 4.96E-01 |
| NULL | GSMG0010963 | NULL | NULL | 265.74 | 285.28 | -0.10 | 7.68E-01 |
| ENSMUSG00000021687 | GSMG0010964 | Scamp1 | secretory carrier membrane protein 1 | 1239.93 | 1322.89 | -0.09 | 7.13E-01 |
| ENSMUSG00000021684 | GSMG0010971 | Pde8b | phosphodiesterase 8B | 1904.87 | 1342.68 | 0.50 | 4.65E-03 |
| NULL | GSMG0010972 | NULL | NULL | 762.44 | 393.42 | 0.96 | 9.96E-08 |
| NULL | GSMG0010973 | NULL | NULL | 1248.19 | 722.62 | 0.79 | 1.16E-05 |
| ENSMUSG00000021681 | GSMG0010974 | Aggf1 | angiogenic factor with G patch and FHA domains 1 | 1362.53 | 1133.08 | 0.27 | 1.40E-01 |
| ENSMUSG00000021680 | GSMG0010975 | Crhbp | corticotropin releasing hormone binding protein | 72.24 | 22.16 | 1.71 | 4.93E-05 |
| ENSMUSG00000048376 | GSMG0010978 | F2r | coagulation factor II (thrombin) receptor | 2424.31 | 4592.73 | -0.92 | 2.54E-11 |
| ENSMUSG00000021676 | GSMG0010979 | Iqgap2 | IQ motif containing GTPase activating protein 2 | 474.83 | 355.62 | 0.41 | 9.09E-02 |
| ENSMUSG00000021668 | GSMG0010985 | Polk | polymerase (DNA directed), kappa | 519.36 | 227.46 | 1.19 | 2.36E-10 |
| ENSMUSG00000021670 | GSMG0010988 | Hmgcr | 3-hydroxy-3-methylglutaryl-Coenzyme A reductase | 1929.95 | 3594.81 | -0.90 | 1.49E-09 |
| ENSMUSG00000021665 | GSMG0010993 | Hexb | hexosaminidase B | 1618.56 | 1418.02 | 0.19 | 3.25E-01 |
| ENSMUSG00000021662 | GSMG0011001 | Arhgef28 | Rho guanine nucleotide exchange factor (GEF) 28 | 1394.70 | 1473.91 | -0.08 | 7.69E-01 |

| | | | | | | | |
|---|---|---|---|---|---|---|---|
| ENSMUSG00000041747 | GSMG0011002 | Utp15 | UTP15 small subunit processome component | 444.06 | 585.89 | -0.40 | 5.34E-02 |
| ENSMUSG00000041685 | GSMG0011007 | Fcho2 | FCH domain only 2 | 2308.79 | 1751.30 | 0.40 | 3.42E-02 |
| ENSMUSG00000009470 | GSMG0011010 | Tnpo1 | transportin 1 | 613.18 | 729.67 | -0.25 | 3.46E-01 |
| NULL | GSMG0011011 | NULL | NULL | 265.47 | 366.78 | -0.46 | 3.68E-02 |
| ENSMUSG00000021650 | GSMG0011015 | Ptcd2 | pentatricopeptide repeat domain 2 | 576.18 | 558.71 | 0.04 | 8.95E-01 |
| ENSMUSG00000052727 | GSMG0011019 | Map1b | microtubule-associated protein 1B | 4792.24 | 16072.68 | -1.75 | 1.17E-28 |
| NULL | GSMG0011021 | NULL | NULL | 278.78 | 252.77 | 0.14 | 6.88E-01 |
| NULL | GSMG0011022 | NULL | NULL | 756.13 | 650.50 | 0.22 | 3.89E-01 |
| ENSMUSG00000021647 | GSMG0011023 | Cartpt | CART prepropeptide | 249.24 | 42.86 | 2.54 | 1.48E-20 |
| ENSMUSG00000021646 | GSMG0011024 | Mccc2 | methylcrotonoyl-Coenzyme A carboxylase 2 (beta) | 282.65 | 323.55 | -0.19 | 5.15E-01 |
| ENSMUSG00000049658 | GSMG0011025 | Bdp1 | B double prime 1, subunit of RNA polymerase III transcription initiatio | 128.88 | 121.49 | 0.09 | 8.53E-01 |
| ENSMUSG00000021639 | GSMG0011032 | Gtf2h2 | general transcription factor II H, polypeptide 2 | 597.89 | 754.10 | -0.33 | 1.21E-01 |
| ENSMUSG00000021638 | GSMG0011033 | Ocln | occludin | 7244.67 | 4120.37 | 0.81 | 4.65E-08 |
| ENSMUSG00000021636 | GSMG0011034 | Marveld2 | MARVEL (membrane-associating) domain containing 2 | 583.14 | 417.74 | 0.48 | 1.77E-02 |
| ENSMUSG00000021635 | GSMG0011035 | Rad17 | RAD17 checkpoint clamp loader component | 513.50 | 516.30 | -0.01 | 9.82E-01 |
| ENSMUSG00000069089 | GSMG0011037 | Cdk7 | cyclin-dependent kinase 7 | 819.02 | 908.79 | -0.15 | 5.10E-01 |
| ENSMUSG00000045273 | GSMG0011040 | Cenph | centromere protein H | 33.34 | 90.71 | -1.44 | 2.09E-04 |
| ENSMUSG00000021629 | GSMG0011042 | Slc30a5 | solute carrier family 30 (zinc transporter), member 5 | 1380.27 | 1356.63 | 0.03 | 9.34E-01 |
| ENSMUSG00000041417 | GSMG0011047 | Pik3r1 | phosphatidylinositol 3-kinase, regulatory subunit, polypeptide 1 (p85 al | 409.87 | 582.54 | -0.50 | 1.46E-02 |
| NULL | GSMG0011049 | NULL | NULL | 883.65 | 690.67 | 0.35 | 1.99E-01 |
| ENSMUSG00000034751 | GSMG0011052 | Mast4 | microtubule associated serine/threonine kinase family member 4 | 3494.79 | 3176.85 | 0.14 | 5.32E-01 |
| NULL | GSMG0011053 | NULL | NULL | 294.17 | 237.33 | 0.31 | 3.85E-01 |
| NULL | GSMG0011062 | NULL | NULL | 620.26 | 680.05 | -0.13 | 6.97E-01 |
| NULL | GSMG0011064 | NULL | NULL | 403.52 | 522.54 | -0.37 | 1.28E-01 |
| ENSMUSG00000032621 | GSMG0011071 | Srek1 | splicing regulatory glutamine/lysine-rich protein 1 | 863.35 | 876.99 | -0.02 | 9.45E-01 |
| NULL | GSMG0011072 | NULL | NULL | 701.91 | 748.31 | -0.09 | 7.89E-01 |
| ENSMUSG00000021709 | GSMG0011075 | Erbb2ip | Erbb2 interacting protein | 2182.28 | 2105.96 | 0.05 | 8.85E-01 |
| ENSMUSG00000021710 | GSMG0011079 | Nln | neurolysin (metallopeptidase M3 family) | 119.77 | 285.63 | -1.25 | 3.00E-08 |
| ENSMUSG00000021711 | GSMG0011081 | Trappc13 | trafficking protein particle complex 13 | 715.17 | 696.02 | 0.04 | 9.02E-01 |
| ENSMUSG00000021713 | GSMG0011082 | Ppwd1 | peptidylprolyl isomerase domain and WD repeat containing 1 | 140.33 | 214.41 | -0.60 | 7.16E-02 |
| ENSMUSG00000021715 | GSMG0011085 | Cwc27 | CWC27 spliceosome-associated protein | 509.62 | 598.54 | -0.23 | 3.02E-01 |
| ENSMUSG00000021719 | GSMG0011089 | Rgs7bp | regulator of G-protein signalling 7 binding protein | 558.56 | 272.13 | 1.04 | 2.40E-09 |
| NULL | GSMG0011090 | NULL | NULL | 173.31 | 68.34 | 1.34 | 3.72E-07 |

| | | | | | | | |
|---|---|---|---|---|---|---|---|
| NULL | GSMG0011091 | NULL | NULL | 256.90 | 92.55 | 1.47 | 1.29E-10 |
| ENSMUSG00000021720 | GSMG0011092 | Rnf180 | ring finger protein 180 | 201.72 | 408.36 | -1.01 | 1.04E-06 |
| ENSMUSG00000042590 | GSMG0011096 | Ipo11 | importin 11 | 974.43 | 1369.88 | -0.49 | 1.50E-03 |
| ENSMUSG00000021693 | GSMG0011098 | Kif2a | kinesin family member 2A | 579.88 | 854.62 | -0.56 | 4.02E-03 |
| NULL | GSMG0011100 | NULL | NULL | 87.85 | 91.07 | -0.04 | 9.62E-01 |
| NULL | GSMG0011105 | NULL | NULL | 163.85 | 185.17 | -0.17 | 6.94E-01 |
| NULL | GSMG0011106 | NULL | NULL | 64.71 | 71.46 | -0.14 | 8.08E-01 |
| NULL | GSMG0011107 | NULL | NULL | 614.32 | 622.76 | -0.02 | 9.62E-01 |
| NULL | GSMG0011108 | NULL | NULL | 417.12 | 478.30 | -0.19 | 5.18E-01 |
| NULL | GSMG0011116 | NULL | NULL | 126.85 | 176.21 | -0.48 | 9.97E-02 |
| ENSMUSG00000021700 | GSMG0011118 | Rab3c | RAB3C, member RAS oncogene family | 544.50 | 856.41 | -0.65 | 2.17E-04 |
| ENSMUSG00000032745 | GSMG0011124 | Gpbp1 | GC-rich promoter binding protein 1 | 1916.51 | 2079.01 | -0.12 | 7.53E-01 |
| ENSMUSG00000021754 | GSMG0011125 | Map3k1 | mitogen-activated protein kinase kinase kinase 1 | 64.38 | 74.12 | -0.20 | 7.45E-01 |
| ENSMUSG00000016018 | GSMG0011136 | Skiv2l2 | superkiller viralicidic activity 2-like 2 (S. cerevisiae) | 2068.40 | 2451.50 | -0.25 | 1.66E-01 |
| ENSMUSG00000021760 | GSMG0011139 | Gpx8 | glutathione peroxidase 8 (putative) | 743.45 | 809.06 | -0.12 | 7.21E-01 |
| ENSMUSG00000042364 | GSMG0011145 | Snx18 | sorting nexin 18 | 3160.58 | 2356.37 | 0.42 | 4.06E-03 |
| ENSMUSG00000051456 | GSMG0011147 | Hspb3 | heat shock protein 3 | 86.85 | 132.66 | -0.62 | 5.34E-02 |
| NULL | GSMG0011150 | NULL | NULL | 268.41 | 277.19 | -0.05 | 9.30E-01 |
| ENSMUSG00000021764 | GSMG0011153 | Ndufs4 | NADH dehydrogenase (ubiquinone) Fe-S protein 4 | 1931.57 | 2267.88 | -0.23 | 2.22E-01 |
| NULL | GSMG0011162 | NULL | NULL | 2267.67 | 1764.93 | 0.36 | 8.15E-02 |
| G00000042275,ENSMUSG000 | GSMG0011164 | Itga1 // Pelo | integrin alpha 1 // pelota homolog (Drosophila) | 11925.96 | 11292.84 | 0.08 | 7.73E-01 |
| NULL | GSMG0011165 | NULL | NULL | 560.77 | 395.38 | 0.51 | 2.54E-02 |
| NULL | GSMG0011169 | NULL | NULL | 89.68 | 139.53 | -0.63 | 5.49E-02 |
| ENSMUSG00000021725 | GSMG0011170 | Parp8 | poly (ADP-ribose) polymerase family, member 8 | 1101.53 | 984.29 | 0.16 | 5.57E-01 |
| ENSMUSG00000021731 | GSMG0011178 | Mrps30 | mitochondrial ribosomal protein S30 | 821.82 | 1130.19 | -0.46 | 8.70E-03 |
| ENSMUSG00000025453 | GSMG0011181 | Nnt | nicotinamide nucleotide transhydrogenase | 1112.86 | 1479.35 | -0.41 | 1.73E-02 |
| NULL | GSMG0011183 | NULL | NULL | 82.59 | 62.10 | 0.41 | 3.76E-01 |
| NULL | GSMG0011184 | NULL | NULL | 494.50 | 604.58 | -0.29 | 2.06E-01 |
| ENSMUSG00000094870 | GSMG0011185 | Zfp131 | zinc finger protein 131 | 285.34 | 316.54 | -0.15 | 6.24E-01 |
| ENSMUSG00000025277 | GSMG0011210 | Abhd6 | abhydrolase domain containing 6 | 1070.32 | 1020.50 | 0.07 | 8.24E-01 |
| ENSMUSG00000023156 | GSMG0011213 | Rpp14 | ribonuclease P 14 subunit | 631.65 | 670.03 | -0.08 | 7.92E-01 |
| ENSMUSG00000033885 | GSMG0011214 | Pxk | PX domain containing serine/threonine kinase | 1624.64 | 1765.05 | -0.12 | 6.23E-01 |
| ENSMUSG00000021752 | GSMG0011216 | Kctd6 | potassium channel tetramerisation domain containing 6 | 810.01 | 616.48 | 0.39 | 9.87E-02 |

| | | | | | | | |
|---|---|---|---|---|---|---|---|
| ENSMUSG00000021745 | GSMG0011224 | Ptprg | protein tyrosine phosphatase, receptor type, G | 4899.13 | 4165.45 | 0.23 | 3.32E-01 |
| NULL | GSMG0011225 | NULL | NULL | 522.76 | 484.22 | 0.11 | 7.47E-01 |
| ENSMUSG00000033111 | GSMG0011230 | 3830406C13Rik | RIKEN cDNA 3830406C13 gene | 536.07 | 454.84 | 0.24 | 5.26E-01 |
| ENSMUSG00000056296 | GSMG0011233 | Synpr | synaptoporin | 154.92 | 156.69 | -0.01 | 9.83E-01 |
| ENSMUSG00000021738 | GSMG0011237 | Atxn7 | ataxin 7 | 940.18 | 671.19 | 0.49 | 5.27E-02 |
| ENSMUSG00000068758 | GSMG0011241 | Il3ra | interleukin 3 receptor, alpha chain | 146.53 | 105.63 | 0.47 | 1.59E-01 |
| ENSMUSG00000021733 | GSMG0011242 | Slc4a7 | solute carrier family 4, sodium bicarbonate cotransporter, member 7 | 661.48 | 991.41 | -0.58 | 4.86E-04 |
| ENSMUSG00000021785 | GSMG0011248 | Ngly1 | N-glycanase 1 | 816.45 | 641.29 | 0.35 | 1.29E-01 |
| ENSMUSG00000017485 | GSMG0011250 | Top2b | topoisomerase (DNA) II beta | 1499.62 | 1852.90 | -0.31 | 1.08E-01 |
| NULL | GSMG0011257 | NULL | NULL | 585.27 | 404.18 | 0.54 | 2.57E-02 |
| ENSMUSG00000021772 | GSMG0011259 | Nkiras1 | NFKB inhibitor interacting Ras-like protein 1 | 996.42 | 665.31 | 0.58 | 4.09E-04 |
| ENSMUSG00000021806 | GSMG0011263 | Nid2 | nidogen 2 | 7514.96 | 11013.23 | -0.55 | 1.73E-03 |
| NULL | GSMG0011265 | NULL | NULL | 121.38 | 332.50 | -1.46 | 3.61E-11 |
| ENSMUSG00000021809 | GSMG0011266 | Nudt13 | nudix (nucleoside diphosphate linked moiety X)-type motif 13 | 187.42 | 200.71 | -0.10 | 8.03E-01 |
| ENSMUSG00000039599 | GSMG0011268 | Fam149b | family with sequence similarity 149, member B | 376.04 | 326.02 | 0.21 | 4.46E-01 |
| NULL | GSMG0011273 | NULL | NULL | 222.80 | 274.15 | -0.30 | 4.48E-01 |
| ENSMUSG00000039367 | GSMG0011274 | Sec24c | Sec24 related gene family, member C (S. cerevisiae) | 2758.88 | 3017.66 | -0.13 | 6.33E-01 |
| ENSMUSG00000039357 | GSMG0011275 | Fut11 | fucosyltransferase 11 | 744.98 | 645.57 | 0.21 | 3.39E-01 |
| ENSMUSG00000063787 | GSMG0011276 | Chchd1 | coiled-coil-helix-coiled-coil-helix domain containing 1 | 738.67 | 1132.05 | -0.62 | 5.01E-03 |
| ENSMUSG00000021819 | GSMG0011277 | Zswim8 | zinc finger SWIM-type containing 8 | 479.73 | 391.41 | 0.29 | 3.19E-01 |
| ENSMUSG00000021823 | GSMG0011280 | Vcl | vinculin | 1845.35 | 1733.13 | 0.09 | 7.04E-01 |
| ENSMUSG00000039197 | GSMG0011282 | Adk | adenosine kinase | 1152.34 | 1000.70 | 0.20 | 4.30E-01 |
| ENSMUSG00000021767 | GSMG0011295 | Kat6b | K(lysine) acetyltransferase 6B | 1168.91 | 1347.64 | -0.20 | 3.63E-01 |
| ENSMUSG00000021770 | GSMG0011297 | Samd8 | sterile alpha motif domain containing 8 | 221.59 | 178.06 | 0.32 | 2.97E-01 |
| ENSMUSG00000021771 | GSMG0011298 | Vdac2 | voltage-dependent anion channel 2 | 2417.16 | 2721.30 | -0.17 | 4.31E-01 |
| ENSMUSG00000007817 | GSMG0011324 | Zmiz1 | zinc finger, MIZ-type containing 1 | 1465.74 | 1628.96 | -0.15 | 6.31E-01 |
| ENSMUSG00000021868 | GSMG0011329 | Ppif | peptidylprolyl isomerase F (cyclophilin F) | 459.00 | 525.53 | -0.20 | 4.48E-01 |
| ENSMUSG00000021866 | GSMG0011330 | Anxa11 | annexin A11 | 982.53 | 575.37 | 0.77 | 8.62E-04 |
| ENSMUSG00000040818 | GSMG0011335 | Dennd6a | DENN/MADD domain containing 6A | 1288.46 | 1241.23 | 0.05 | 8.50E-01 |
| NULL | GSMG0011336 | NULL | NULL | 431.81 | 407.92 | 0.08 | 8.20E-01 |
| ENSMUSG00000021877 | GSMG0011337 | Arf4 | ADP-ribosylation factor 4 | 6319.54 | 7723.53 | -0.29 | 1.48E-01 |
| ENSMUSG00000040717 | GSMG0011344 | Il17rd | interleukin 17 receptor D | 221.07 | 242.91 | -0.14 | 7.51E-01 |
| ENSMUSG00000021895 | GSMG0011350 | Arhgef3 | Rho guanine nucleotide exchange factor (GEF) 3 | 2120.70 | 1106.21 | 0.94 | 2.77E-10 |

| | | | | | | | |
|---|---|---|---|---|---|---|---|
| NULL | GSMG0011351 | NULL | NULL | 581.46 | 202.48 | 1.52 | 4.17E-11 |
| ENSMUSG00000040651 | GSMG0011353 | Fam208a | family with sequence similarity 208, member A | 228.79 | 270.14 | -0.24 | 3.87E-01 |
| ENSMUSG00000040640 | GSMG0011354 | Erc2 | ELKS/RAB6-interacting/CAST family member 2 | 2083.40 | 1949.25 | 0.10 | 7.04E-01 |
| NULL | GSMG0011359 | NULL | NULL | 416.59 | 324.90 | 0.36 | 1.27E-01 |
| NULL | GSMG0011360 | NULL | NULL | 663.97 | 504.66 | 0.40 | 9.40E-02 |
| NULL | GSMG0011362 | NULL | NULL | 340.60 | 244.50 | 0.48 | 3.38E-02 |
| ENSMUSG00000021994 | GSMG0011363 | Wnt5a | wingless-type MMTV integration site family, member 5A | 420.33 | 268.57 | 0.65 | 5.51E-03 |
| ENSMUSG00000042682 | GSMG0011368 | Selk | selenoprotein K | 3395.74 | 3391.13 | 0.00 | 9.96E-01 |
| ENSMUSG00000015971 | GSMG0011369 | Actr8 | ARP8 actin-related protein 8 | 1418.87 | 1416.40 | 0.00 | 9.91E-01 |
| ENSMUSG00000021962 | GSMG0011373 | Dcp1a | decapping mRNA 1A | 341.25 | 384.48 | -0.17 | 6.29E-01 |
| ENSMUSG00000021957 | GSMG0011374 | Tkt | transketolase | 2504.17 | 4342.07 | -0.79 | 6.93E-09 |
| ENSMUSG00000052395 | GSMG0011377 | Rft1 | RFT1 homolog | 440.78 | 399.70 | 0.14 | 6.27E-01 |
| ENSMUSG00000006527 | GSMG0011378 | Sfmbt1 | Scm-like with four mbt domains 1 | 353.58 | 391.62 | -0.15 | 6.27E-01 |
| ENSMUSG00000006526 | GSMG0011382 | Tmem110 | transmembrane protein 110 | 377.12 | 424.06 | -0.17 | 5.41E-01 |
| ENSMUSG00000042485 | GSMG0011384 | Mustn1 | musculoskeletal, embryonic nuclear protein 1 | 305.32 | 71.40 | 2.10 | 1.67E-17 |
| ENSMUSG00000021918 | GSMG0011386 | Nek4 | NIMA (never in mitosis gene a)-related expressed kinase 4 | 310.80 | 308.04 | 0.02 | 9.69E-01 |
| ENSMUSG00000021916 | GSMG0011387 | Glt8d1 | glycosyltransferase 8 domain containing 1 | 679.19 | 695.28 | -0.04 | 9.19E-01 |
| ENSMUSG00000042323 | GSMG0011388 | Pbrm1 | polybromo 1 | 2745.08 | 3079.71 | -0.17 | 3.93E-01 |
| ENSMUSG00000071547 | GSMG0011390 | Nt5dc2 | 5'-nucleotidase domain containing 2 | 763.01 | 1913.86 | -1.33 | 1.05E-13 |
| ENSMUSG00000021904 | GSMG0011392 | Sema3g | sema domain, immunoglobulin domain (Ig), short basic domain, secrete | 610.09 | 150.07 | 2.02 | 2.43E-14 |
| ENSMUSG00000021901 | GSMG0011393 | Bap1 | Brca1 associated protein 1 | 2614.48 | 2874.46 | -0.14 | 5.51E-01 |
| ENSMUSG00000021893 | GSMG0011394 | Capn7 | calpain 7 | 1304.10 | 1435.66 | -0.14 | 5.86E-01 |
| ENSMUSG00000021890 | GSMG0011396 | Eaf1 | ELL associated factor 1 | 220.03 | 226.62 | -0.04 | 9.26E-01 |
| ENSMUSG00000021900 | GSMG0011398 | Btd | biotinidase | 1446.38 | 1442.45 | 0.00 | 9.89E-01 |
| ENSMUSG00000021903 | GSMG0011400 | Galnt15 | UDP-N-acetyl-alpha-D-galactosamine:polypeptide N-acetylgalactosami | 157.72 | 82.65 | 0.93 | 1.19E-02 |
| ENSMUSG00000021906 | GSMG0011401 | Oxnad1 | oxidoreductase NAD-binding domain containing 1 | 123.82 | 121.78 | 0.03 | 9.55E-01 |
| ENSMUSG00000021911 | GSMG0011405 | Parg | poly (ADP-ribose) glycohydrolase | 889.41 | 996.46 | -0.16 | 5.04E-01 |
| ENSMUSG00000021913 | GSMG0011408 | Ogdhl | oxoglutarate dehydrogenase-like | 709.76 | 390.60 | 0.86 | 8.44E-05 |
| ENSMUSG00000054051 | GSMG0011410 | Ercc6 | excision repair cross-complementing rodent repair deficiency, complem | 864.67 | 912.36 | -0.08 | 7.67E-01 |
| ENSMUSG00000050666 | GSMG0011415 | Vstm4 | V-set and transmembrane domain containing 4 | 1008.02 | 899.19 | 0.16 | 4.74E-01 |
| NULL | GSMG0011416 | NULL | NULL | 117.75 | 67.63 | 0.81 | 2.13E-02 |
| ENSMUSG00000021943 | GSMG0011421 | Gdf10 | growth differentiation factor 10 | 168.30 | 111.76 | 0.59 | 1.14E-01 |
| ENSMUSG00000041479 | GSMG0011427 | Syt15 | synaptotagmin XV | 408.59 | 143.54 | 1.51 | 9.16E-11 |

| | | | | | | | |
|---|---|---|---|---|---|---|---|
| ENSMUSG00000021794 | GSMG0011428 | Glud1 | glutamate dehydrogenase 1 | 4465.37 | 5111.51 | -0.19 | 2.59E-01 |
| ENSMUSG00000041445 | GSMG0011431 | Mmrn2 | multimerin 2 | 4090.69 | 2850.02 | 0.52 | 1.14E-03 |
| ENSMUSG00000041408 | GSMG0011433 | Wapl | WAPL cohesin release factor | 2282.62 | 1987.33 | 0.20 | 4.01E-01 |
| ENSMUSG00000041078 | GSMG0011436 | Grid1 | glutamate receptor, ionotropic, delta 1 | 524.73 | 429.50 | 0.29 | 4.30E-01 |
| ENSMUSG00000021832 | GSMG0011472 | Psmc6 | proteasome (prosome, macropain) 26S subunit, ATPase, 6 | 1566.36 | 1646.59 | -0.07 | 8.28E-01 |
| ENSMUSG00000037628 | GSMG0011480 | Cdkn3 | cyclin-dependent kinase inhibitor 3 | 88.01 | 172.49 | -0.97 | 5.82E-04 |
| ENSMUSG00000055128 | GSMG0011481 | Cgrrf1 | cell growth regulator with ring finger domain 1 | 474.66 | 360.99 | 0.40 | 1.03E-01 |
| ENSMUSG00000021840 | GSMG0011488 | Mapk1ip1l | mitogen-activated protein kinase 1 interacting protein 1-like | 313.45 | 384.03 | -0.29 | 2.09E-01 |
| ENSMUSG00000037536 | GSMG0011491 | Fbxo34 | F-box protein 34 | 1533.44 | 1169.58 | 0.39 | 2.43E-02 |
| ENSMUSG00000021843 | GSMG0011493 | Ktn1 | kinectin 1 | 1485.32 | 1673.52 | -0.17 | 5.30E-01 |
| ENSMUSG00000021846 | GSMG0011496 | Peli2 | pellino 2 | 147.33 | 185.14 | -0.33 | 3.32E-01 |
| ENSMUSG00000036339 | GSMG0011502 | Tmem260 | transmembrane protein 260 | 425.55 | 374.38 | 0.19 | 4.88E-01 |
| ENSMUSG00000036291 | GSMG0011509 | Ap5m1 | adaptor-related protein complex 5, mu 1 subunit | 193.34 | 161.83 | 0.26 | 4.63E-01 |
| ENSMUSG00000036282 | GSMG0011510 | Naa30 | N(alpha)-acetyltransferase 30, NatC catalytic subunit | 277.91 | 267.89 | 0.05 | 8.88E-01 |
| ENSMUSG00000036023 | GSMG0011521 | Parp2 | poly (ADP-ribose) polymerase family, member 2 | 476.53 | 558.03 | -0.23 | 3.53E-01 |
| ENSMUSG00000035960 | GSMG0011523 | Apex1 | apurinic/apyrimidinic endonuclease 1 | 441.13 | 978.30 | -1.15 | 2.80E-10 |
| G00000021876,ENSMUSG000 | GSMG0011527 | Rnase4 // Ang | ribonuclease, RNase A family 4 // angiogenin, ribonuclease, RNase A fa | 567.76 | 327.40 | 0.80 | 2.79E-04 |
| ENSMUSG00000004561 | GSMG0011537 | Mettl17 | methyltransferase like 17 | 1798.26 | 1814.27 | -0.01 | 9.65E-01 |
| ENSMUSG00000004562 | GSMG0011539 | Arhgef40 | Rho guanine nucleotide exchange factor (GEF) 40 | 1570.87 | 1700.69 | -0.12 | 7.95E-01 |
| ENSMUSG00000016831 | GSMG0011544 | Tox4 | TOX high mobility group box family member 4 | 957.21 | 984.12 | -0.04 | 8.98E-01 |
| NULL | GSMG0011566 | NULL | NULL | 263.17 | 303.47 | -0.20 | 5.01E-01 |
| ENSMUSG00000040997 | GSMG0011567 | Abhd4 | abhydrolase domain containing 4 | 1123.80 | 1154.63 | -0.04 | 9.02E-01 |
| ENSMUSG00000000959 | GSMG0011568 | Oxa1l | oxidase assembly 1-like | 1055.26 | 1235.09 | -0.23 | 2.82E-01 |
| ENSMUSG00000010406 | GSMG0011569 | Mrpl52 | mitochondrial ribosomal protein L52 | 688.81 | 956.49 | -0.48 | 1.08E-01 |
| ENSMUSG00000000957 | GSMG0011570 | Mmp14 | matrix metallopeptidase 14 (membrane-inserted) | 1950.37 | 2721.87 | -0.48 | 1.41E-03 |
| ENSMUSG00000022175 | GSMG0011572 | Lrp10 | low-density lipoprotein receptor-related protein 10 | 4436.74 | 4202.45 | 0.08 | 7.78E-01 |
| ENSMUSG00000022176 | GSMG0011573 | Rem2 | rad and gem related GTP binding protein 2 | 283.59 | 358.49 | -0.34 | 2.99E-01 |
| ENSMUSG00000040822 | GSMG0011578 | 1700123O20Rik | RIKEN cDNA 1700123O20 gene | 1258.27 | 1413.18 | -0.17 | 5.13E-01 |
| ENSMUSG00000022194 | GSMG0011581 | Pabpn1 | poly(A) binding protein, nuclear 1 | 6632.77 | 4983.17 | 0.41 | 1.38E-01 |
| ENSMUSG00000040759 | GSMG0011583 | Cmtm5 | CKLF-like MARVEL transmembrane domain containing 5 | 717.10 | 220.61 | 1.70 | 3.29E-21 |
| ENSMUSG00000022204 | GSMG0011587 | Ngdn | neuroguidin, EIF4E binding protein | 577.27 | 649.69 | -0.17 | 4.85E-01 |
| G00000045691,ENSMUSG000 | GSMG0011589 | Zfhx2os // Thtpa | zinc finger homeobox 2, opposite strand // thiamine triphosphatase | 880.86 | 1051.98 | -0.26 | 2.30E-01 |
| ENSMUSG00000022210 | GSMG0011595 | Dhrs4 | dehydrogenase/reductase (SDR family) member 4 | 416.41 | 401.97 | 0.05 | 8.88E-01 |

| | | | | | | | |
|---|---|---|---|---|---|---|---|
| ENSMUSG00000022211 | GSMG0011597 | Lrrc16b | leucine rich repeat containing 16B | 1547.02 | 2612.53 | -0.76 | 3.50E-03 |
| ENSMUSG00000022212 | GSMG0011598 | Cpne6 | copine VI | 4396.16 | 2415.28 | 0.86 | 3.98E-06 |
| ENSMUSG00000040618 | GSMG0011599 | Pck2 | phosphoenolpyruvate carboxykinase 2 (mitochondrial) | 1016.95 | 924.35 | 0.14 | 5.82E-01 |
| ENSMUSG00000022214 | GSMG0011600 | Dcaf11 | DDB1 and CUL4 associated factor 11 | 2442.39 | 1880.28 | 0.38 | 1.88E-02 |
| ENSMUSG00000022216 | GSMG0011602 | Psme1 | proteasome (prosome, macropain) activator subunit 1 (PA28 alpha) | 1238.58 | 1435.68 | -0.21 | 3.80E-01 |
| ENSMUSG00000047098 | GSMG0011603 | Rnf31 | ring finger protein 31 | 698.87 | 585.94 | 0.25 | 3.06E-01 |
| ENSMUSG00000002325 | GSMG0011604 | Irf9 | interferon regulatory factor 9 | 614.98 | 615.43 | 0.00 | 9.98E-01 |
| ENSMUSG00000002326 | GSMG0011609 | Gmpr2 | guanosine monophosphate reductase 2 | 435.60 | 454.23 | -0.06 | 8.53E-01 |
| ENSMUSG00000019297 | GSMG0011610 | Nop9 | NOP9 nucleolar protein | 821.46 | 1061.86 | -0.37 | 6.01E-02 |
| ENSMUSG00000023411 | GSMG0011613 | Nfatc4 | nuclear factor of activated T cells, cytoplasmic, calcineurin dependent 4 | 144.42 | 130.55 | 0.14 | 7.67E-01 |
| ENSMUSG00000075592 | GSMG0011614 | Nynrin | NYN domain and retroviral integrase containing | 220.38 | 310.48 | -0.50 | 9.48E-02 |
| ENSMUSG00000047153 | GSMG0011615 | Khnyn | KH and NYN domain containing | 1624.96 | 1414.30 | 0.20 | 4.07E-01 |
| ENSMUSG00000079184 | GSMG0011625 | Mphosph8 | M-phase phosphoprotein 8 | 1107.69 | 1047.11 | 0.08 | 8.34E-01 |
| ENSMUSG00000100017 | GSMG0011628 | 2410022M11Rik | RIKEN cDNA 2410022M11 gene | 315.46 | 343.98 | -0.13 | 7.15E-01 |
| ENSMUSG00000021945 | GSMG0011629 | Zmym2 | zinc finger, MYM-type 2 | 336.80 | 362.31 | -0.10 | 7.61E-01 |
| ENSMUSG00000040040 | GSMG0011633 | Ift88 | intraflagellar transport 88 | 427.29 | 392.69 | 0.12 | 6.99E-01 |
| ENSMUSG00000050222 | GSMG0011634 | Il17d | interleukin 17D | 152.99 | 131.73 | 0.21 | 6.95E-01 |
| ENSMUSG00000021967 | GSMG0011640 | Mrpl57 | mitochondrial ribosomal protein L57 | 474.49 | 582.25 | -0.30 | 2.05E-01 |
| NULL | GSMG0011641 | NULL | NULL | 171.92 | 204.47 | -0.26 | 4.68E-01 |
| ENSMUSG00000035469 | GSMG0011646 | Rcbtb1 | regulator of chromosome condensation (RCC1) and BTB (POZ) domai | 368.87 | 370.78 | -0.01 | 9.84E-01 |
| ENSMUSG00000021981 | GSMG0011647 | Cab39l | calcium binding protein 39-like | 615.03 | 791.39 | -0.36 | 1.48E-01 |
| ENSMUSG00000021987 | GSMG0011652 | Mtmr6 | myotubularin related protein 6 | 2901.76 | 2783.68 | 0.06 | 8.27E-01 |
| ENSMUSG00000021986 | GSMG0011653 | Amer2 | APC membrane recruitment 2 | 638.02 | 962.47 | -0.60 | 5.58E-03 |
| ENSMUSG00000021990 | GSMG0011657 | Spata13 | spermatogenesis associated 13 | 401.78 | 437.17 | -0.12 | 7.31E-01 |
| ENSMUSG00000021993 | GSMG0011660 | Mipep | mitochondrial intermediate peptidase | 180.01 | 234.59 | -0.38 | 1.49E-01 |
| NULL | GSMG0011661 | NULL | NULL | 77.93 | 57.19 | 0.45 | 3.22E-01 |
| NULL | GSMG0011662 | NULL | NULL | 485.45 | 414.39 | 0.23 | 3.94E-01 |
| NULL | GSMG0011664 | NULL | NULL | 537.65 | 622.01 | -0.21 | 3.96E-01 |
| ENSMUSG00000035235 | GSMG0011669 | Trim13 | tripartite motif-containing 13 | 384.82 | 496.47 | -0.37 | 1.68E-01 |
| ENSMUSG00000021932 | GSMG0011672 | Rnaseh2b | ribonuclease H2, subunit B | 293.02 | 407.53 | -0.47 | 6.09E-02 |
| ENSMUSG00000035184 | GSMG0011674 | Fam124a | family with sequence similarity 124, member A | 518.15 | 1522.17 | -1.55 | 2.48E-19 |
| ENSMUSG00000014547 | GSMG0011679 | Wdfy2 | WD repeat and FYVE domain containing 2 | 179.05 | 154.51 | 0.21 | 5.88E-01 |
| ENSMUSG00000021939 | GSMG0011684 | Ctsb | cathepsin B | 12613.79 | 12256.17 | 0.04 | 8.75E-01 |

| | | | | | | | |
|---|---|---|---|---|---|---|---|
| ENSMUSG00000035067 | GSMG0011689 | Xkr6 | X-linked Kx blood group related 6 | 390.81 | 354.41 | 0.14 | 7.37E-01 |
| NULL | GSMG0011693 | NULL | NULL | 264.14 | 442.50 | -0.75 | 1.66E-03 |
| ENSMUSG00000021958 | GSMG0011697 | Pinx1 | PIN2/TERF1 interacting, telomerase inhibitor 1 | 275.26 | 382.22 | -0.48 | 1.16E-01 |
| ENSMUSG00000063060 | GSMG0011699 | Sox7 | SRY (sex determining region Y)-box 7 | 3423.23 | 3640.98 | -0.09 | 7.16E-01 |
| NULL | GSMG0011710 | NULL | NULL | 134.00 | 71.81 | 0.90 | 3.58E-03 |
| ENSMUSG00000097545 | GSMG0011712 | Mir124a-1hg | Mir124-1 host gene (non-protein coding) | 3366.34 | 1795.96 | 0.91 | 9.63E-07 |
| ENSMUSG00000021975 | GSMG0011716 | Ints9 | integrator complex subunit 9 | 674.20 | 628.64 | 0.10 | 7.17E-01 |
| ENSMUSG00000034532 | GSMG0011718 | Fbxo16 | F-box protein 16 | 71.76 | 99.85 | -0.48 | 2.26E-01 |
| ENSMUSG00000034522 | GSMG0011721 | Zfp395 | zinc finger protein 395 | 239.41 | 174.27 | 0.46 | 1.17E-01 |
| ENSMUSG00000022033 | GSMG0011725 | Pbk | PDZ binding kinase | 163.16 | 377.04 | -1.20 | 1.95E-06 |
| ENSMUSG00000022035 | GSMG0011726 | Ccdc25 | coiled-coil domain containing 25 | 877.24 | 891.21 | -0.02 | 9.43E-01 |
| ENSMUSG00000022037 | GSMG0011727 | Clu | clusterin | 8388.11 | 3225.11 | 1.38 | 1.85E-11 |
| ENSMUSG00000022043 | GSMG0011729 | Trim35 | tripartite motif-containing 35 | 6124.19 | 7433.18 | -0.28 | 8.66E-02 |
| ENSMUSG00000022044 | GSMG0011730 | Stmn4 | stathmin-like 4 | 629.30 | 1691.30 | -1.43 | 6.82E-14 |
| ENSMUSG00000046204 | GSMG0011735 | Pnma2 | paraneoplastic antigen MA2 | 1878.62 | 1930.32 | -0.04 | 9.00E-01 |
| ENSMUSG00000034327 | GSMG0011743 | Kctd9 | potassium channel tetramerisation domain containing 9 | 122.06 | 113.45 | 0.11 | 8.30E-01 |
| ENSMUSG00000022055 | GSMG0011746 | Nefl | neurofilament, light polypeptide | 1535.63 | 1108.23 | 0.47 | 2.00E-02 |
| ENSMUSG00000014813 | GSMG0011748 | Stc1 | stanniocalcin 1 | 360.38 | 305.61 | 0.24 | 4.94E-01 |
| ENSMUSG00000022089 | GSMG0011763 | Bin3 | bridging integrator 3 | 550.71 | 522.08 | 0.08 | 7.92E-01 |
| ENSMUSG00000003469 | GSMG0011769 | Phyhip | phytanoyl-CoA hydroxylase interacting protein | 361.94 | 97.97 | 1.88 | 8.53E-11 |
| ENSMUSG00000033595 | GSMG0011770 | Lgi3 | leucine-rich repeat LGI family, member 3 | 309.96 | 53.07 | 2.55 | 3.66E-20 |
| ENSMUSG00000033589 | GSMG0011771 | Reep4 | receptor accessory protein 4 | 452.26 | 444.97 | 0.02 | 9.45E-01 |
| ENSMUSG00000022096 | GSMG0011772 | Hr | hairless | 393.98 | 201.48 | 0.97 | 1.42E-03 |
| ENSMUSG00000045211 | GSMG0011773 | Nudt18 | nudix (nucleoside diphosphate linked moiety X)-type motif 18 | 854.40 | 597.36 | 0.52 | 8.81E-03 |
| NULL | GSMG0011776 | NULL | NULL | 138.96 | 122.96 | 0.18 | 6.94E-01 |
| ENSMUSG00000022103 | GSMG0011779 | Gfra2 | glial cell line derived neurotrophic factor family receptor alpha 2 | 301.98 | 217.60 | 0.47 | 4.88E-02 |
| ENSMUSG00000022106 | GSMG0011786 | Rcbtb2 | regulator of chromosome condensation (RCC1) and BTB (POZ) domai | 677.71 | 814.92 | -0.26 | 2.42E-01 |
| ENSMUSG00000033446 | GSMG0011787 | Lpar6 | lysophosphatidic acid receptor 6 | 2943.23 | 2221.94 | 0.41 | 5.86E-02 |
| NULL | GSMG0011789 | NULL | NULL | 947.94 | 581.96 | 0.70 | 7.53E-03 |
| ENSMUSG00000022109 | GSMG0011790 | Med4 | mediator complex subunit 4 | 797.42 | 1055.28 | -0.40 | 3.17E-02 |
| ENSMUSG00000022110 | GSMG0011791 | Sucla2 | succinate-Coenzyme A ligase, ADP-forming, beta subunit | 1401.71 | 1530.45 | -0.13 | 6.23E-01 |
| ENSMUSG00000021996 | GSMG0011799 | Esd | esterase D/formylglutathione hydrolase | 1551.89 | 1661.88 | -0.10 | 7.34E-01 |
| NULL | GSMG0011801 | NULL | NULL | 672.49 | 665.51 | 0.01 | 9.68E-01 |

| | | | | | | | |
|---|---|---|---|---|---|---|---|
| ENSMUSG00000021998 | GSMG0011805 | Lcp1 | lymphocyte cytosolic protein 1 | 1532.83 | 3084.14 | -1.01 | 1.96E-10 |
| ENSMUSG00000022000 | GSMG0011808 | Zc3h13 | zinc finger CCCH type containing 13 | 1036.91 | 1151.55 | -0.15 | 5.36E-01 |
| ENSMUSG00000022009 | GSMG0011817 | Nufip1 | nuclear fragile X mental retardation protein interacting protein 1 | 317.83 | 424.06 | -0.41 | 9.33E-02 |
| ENSMUSG00000022010 | GSMG0011819 | Tsc22d1 | TSC22 domain family, member 1 | 28661.08 | 31796.43 | -0.15 | 6.98E-01 |
| ENSMUSG00000022012 | GSMG0011828 | Enox1 | ecto-NOX disulfide-thiol exchanger 1 | 353.02 | 499.44 | -0.50 | 1.83E-02 |
| ENSMUSG00000058997 | GSMG0011842 | Vwa8 | von Willebrand factor A domain containing 8 | 871.44 | 853.62 | 0.03 | 9.37E-01 |
| ENSMUSG00000022022 | GSMG0011851 | Mtrf1 | mitochondrial translational release factor 1 | 173.81 | 178.42 | -0.04 | 9.34E-01 |
| NULL | GSMG0011852 | NULL | NULL | 268.11 | 290.02 | -0.11 | 7.53E-01 |
| ENSMUSG00000036461 | GSMG0011854 | Elf1 | E74-like factor 1 | 444.42 | 398.44 | 0.16 | 5.57E-01 |
| ENSMUSG00000022024 | GSMG0011856 | Sugt1 | SGT1, suppressor of G2 allele of SKP1 (S. cerevisiae) | 1129.25 | 1254.62 | -0.15 | 5.99E-01 |
| ENSMUSG00000035566 | GSMG0011862 | Pcdh17 | protocadherin 17 | 1109.95 | 2061.08 | -0.89 | 2.08E-08 |
| ENSMUSG00000022019 | GSMG0011865 | Tdrd3 | tudor domain containing 3 | 844.34 | 1069.95 | -0.34 | 5.62E-02 |
| ENSMUSG00000022070 | GSMG0011873 | Bora | bora, aurora kinase A activator | 210.44 | 392.04 | -0.89 | 1.04E-04 |
| ENSMUSG00000022064 | GSMG0011874 | Pibf1 | progesterone immunomodulatory binding factor 1 | 406.35 | 494.90 | -0.28 | 3.47E-01 |
| ENSMUSG00000005148 | GSMG0011878 | Klf5 | Kruppel-like factor 5 | 194.21 | 223.28 | -0.20 | 5.58E-01 |
| ENSMUSG00000022111 | GSMG0011884 | Uchl3 | ubiquitin carboxyl-terminal esterase L3 (ubiquitin thiolesterase) | 292.70 | 532.61 | -0.86 | 2.14E-05 |
| ENSMUSG00000033060 | GSMG0011887 | Lmo7 | LIM domain only 7 | 412.26 | 520.96 | -0.34 | 1.46E-01 |
| NULL | GSMG0011888 | NULL | NULL | 317.76 | 254.27 | 0.32 | 2.32E-01 |
| ENSMUSG00000022125 | GSMG0011891 | Cln5 | ceroid-lipofuscinosis, neuronal 5 | 828.16 | 938.36 | -0.18 | 5.11E-01 |
| ENSMUSG00000055717 | GSMG0011898 | Slain1 | SLAIN motif family, member 1 | 147.54 | 243.52 | -0.72 | 3.64E-03 |
| ENSMUSG00000053253 | GSMG0011903 | Ndfip2 | Nedd4 family interacting protein 2 | 1940.93 | 1819.88 | 0.09 | 6.98E-01 |
| ENSMUSG00000033214 | GSMG0011912 | Slitrk5 | SLIT and NTRK-like family, member 5 | 1325.04 | 2144.05 | -0.69 | 4.49E-05 |
| ENSMUSG00000022112 | GSMG0011918 | Gpc5 | glypican 5 | 200.67 | 359.99 | -0.84 | 6.14E-04 |
| ENSMUSG00000058571 | GSMG0011924 | Gpc6 | glypican 6 | 813.34 | 699.86 | 0.22 | 3.55E-01 |
| NULL | GSMG0011930 | NULL | NULL | 385.48 | 337.60 | 0.19 | 5.45E-01 |
| ENSMUSG00000022131 | GSMG0011945 | Gpr180 | G protein-coupled receptor 180 | 965.18 | 952.03 | 0.02 | 9.50E-01 |
| NULL | GSMG0011952 | NULL | NULL | 289.41 | 174.77 | 0.72 | 4.01E-03 |
| ENSMUSG00000022132 | GSMG0011953 | Cldn10 | claudin 10 | 652.23 | 333.77 | 0.97 | 9.14E-08 |
| ENSMUSG00000022136 | GSMG0011955 | Dnajc3 | DnaJ heat shock protein family (Hsp40) member C3 | 1759.02 | 1684.31 | 0.06 | 8.29E-01 |
| NULL | GSMG0011963 | NULL | NULL | 266.15 | 196.10 | 0.44 | 1.49E-01 |
| ENSMUSG00000022139 | GSMG0011966 | Mbnl2 | muscleblind-like 2 | 3026.14 | 2580.88 | 0.23 | 3.11E-01 |
| NULL | GSMG0011967 | NULL | NULL | 274.08 | 111.75 | 1.30 | 6.62E-08 |
| NULL | GSMG0011968 | NULL | NULL | 323.65 | 172.47 | 0.91 | 3.08E-05 |

| | | | | | | | |
|---|---|---|---|---|---|---|---|
| NULL | GSMG0011969 | NULL | NULL | 418.19 | 323.87 | 0.37 | 1.51E-01 |
| ENSMUSG00000051615 | GSMG0011970 | Rap2a | RAS related protein 2a | 3027.33 | 2511.41 | 0.27 | 1.91E-01 |
| ENSMUSG00000030662 | GSMG0011973 | Ipo5 | importin 5 | 578.38 | 981.23 | -0.76 | 9.95E-05 |
| ENSMUSG00000025555 | GSMG0011974 | Farp1 | FERM, RhoGEF (Arhgef) and pleckstrin domain protein 1 (chondrocyt | 449.46 | 721.39 | -0.68 | 1.72E-04 |
| NULL | GSMG0011977 | NULL | NULL | 1428.63 | 2424.92 | -0.76 | 1.11E-03 |
| ENSMUSG00000041765 | GSMG0011981 | Ubac2 | ubiquitin associated domain containing 2 | 716.34 | 930.98 | -0.38 | 4.25E-02 |
| ENSMUSG00000025544 | GSMG0011986 | Tm9sf2 | transmembrane 9 superfamily member 2 | 4883.56 | 4906.75 | -0.01 | 9.82E-01 |
| ENSMUSG00000025545 | GSMG0011988 | Clybl | citrate lyase beta like | 251.69 | 228.58 | 0.14 | 7.21E-01 |
| ENSMUSG00000041650 | GSMG0011995 | Pcca | propionyl-Coenzyme A carboxylase, alpha polypeptide | 1879.34 | 1927.10 | -0.04 | 8.94E-01 |
| ENSMUSG00000032925 | GSMG0012002 | Itgbl1 | integrin, beta-like 1 | 70.60 | 108.05 | -0.61 | 1.76E-01 |
| NULL | GSMG0012026 | NULL | NULL | 121.84 | 111.69 | 0.13 | 7.74E-01 |
| ENSMUSG00000021748 | GSMG0012027 | Pdhb | pyruvate dehydrogenase (lipoamide) beta | 1507.55 | 1446.79 | 0.06 | 8.19E-01 |
| ENSMUSG00000021750 | GSMG0012029 | Fam107a | family with sequence similarity 107, member A | 718.91 | 55.32 | 3.69 | 8.67E-50 |
| ENSMUSG00000021743 | GSMG0012040 | Fezf2 | Fez family zinc finger 2 | 188.66 | 333.44 | -0.82 | 6.57E-05 |
| ENSMUSG00000054423 | GSMG0012041 | Cadps | Ca2+-dependent secretion activator | 1145.08 | 810.50 | 0.50 | 1.56E-02 |
| NULL | GSMG0012042 | NULL | NULL | 378.98 | 247.07 | 0.62 | 2.93E-03 |
| NULL | GSMG0012043 | NULL | NULL | 199.84 | 138.45 | 0.53 | 1.62E-01 |
| NULL | GSMG0012044 | NULL | NULL | 194.30 | 151.14 | 0.37 | 2.59E-01 |
| NULL | GSMG0012046 | NULL | NULL | 120.20 | 82.94 | 0.54 | 1.25E-01 |
| ENSMUSG00000053453 | GSMG0012051 | Thoc7 | THO complex 7 | 621.89 | 749.42 | -0.27 | 2.89E-01 |
| ENSMUSG00000021737 | GSMG0012052 | Psmd6 | proteasome (prosome, macropain) 26S subunit, non-ATPase, 6 | 2382.59 | 2758.67 | -0.21 | 2.60E-01 |
| ENSMUSG00000045201 | GSMG0012057 | Lrrc3b | leucine rich repeat containing 3B | 140.93 | 180.92 | -0.36 | 2.51E-01 |
| ENSMUSG00000017491 | GSMG0012060 | Rarb | retinoic acid receptor, beta | 98.44 | 99.23 | -0.01 | 9.89E-01 |
| ENSMUSG00000021775 | GSMG0012066 | Nr1d2 | nuclear receptor subfamily 1, group D, member 2 | 1120.06 | 776.55 | 0.53 | 4.02E-03 |
| ENSMUSG00000021774 | GSMG0012068 | Ube2e1 | ubiquitin-conjugating enzyme E2E 1 | 296.75 | 411.78 | -0.47 | 2.61E-02 |
| ENSMUSG00000058317 | GSMG0012070 | Ube2e2 | ubiquitin-conjugating enzyme E2E 2 | 428.52 | 458.28 | -0.10 | 7.69E-01 |
| NULL | GSMG0012074 | NULL | NULL | 431.00 | 356.23 | 0.27 | 2.50E-01 |
| ENSMUSG00000021807 | GSMG0012077 | 2700060E02Rik | RIKEN cDNA 2700060E02 gene | 543.63 | 867.68 | -0.68 | 9.24E-04 |
| ENSMUSG00000043004 | GSMG0012078 | Gng2 | guanine nucleotide binding protein (G protein), gamma 2 | 1643.18 | 4129.07 | -1.33 | 4.41E-20 |
| ENSMUSG00000045107 | GSMG0012080 | Saysd1 | SAYSVFN motif domain containing 1 | 821.49 | 878.57 | -0.10 | 7.29E-01 |
| ENSMUSG00000021810 | GSMG0012083 | Ecd | ecdysoneless homolog (Drosophila) | 1067.50 | 1396.14 | -0.39 | 2.18E-02 |
| ENSMUSG00000021811 | GSMG0012085 | Dnajc9 | DnaJ heat shock protein family (Hsp40) member C9 | 474.80 | 811.92 | -0.77 | 6.62E-06 |
| ENSMUSG00000049960 | GSMG0012087 | Mrps16 | mitochondrial ribosomal protein S16 | 590.75 | 698.41 | -0.24 | 3.83E-01 |

| | | | | | | | |
|---|---|---|---|---|---|---|---|
| ENSMUSG00000021814 | GSMG0012089 | Anxa7 | annexin A7 | 1521.23 | 1200.60 | 0.34 | 5.87E-02 |
| ENSMUSG00000021816 | GSMG0012092 | Ppp3cb | protein phosphatase 3, catalytic subunit, beta isoform | 519.18 | 623.72 | -0.27 | 2.59E-01 |
| NULL | GSMG0012101 | NULL | NULL | 695.58 | 682.36 | 0.03 | 9.30E-01 |
| ENSMUSG00000039308 | GSMG0012103 | Ndst2 | N-deacetylase/N-sulfotransferase (heparan glucosaminyl) 2 | 1608.86 | 1778.62 | -0.14 | 5.40E-01 |
| ENSMUSG00000021820 | GSMG0012104 | Camk2g | calcium/calmodulin-dependent protein kinase II gamma | 525.39 | 433.29 | 0.28 | 3.16E-01 |
| ENSMUSG00000021773 | GSMG0012111 | Comtd1 | catechol-O-methyltransferase domain containing 1 | 33.90 | 54.86 | -0.70 | 1.56E-01 |
| NULL | GSMG0012117 | NULL | NULL | 1062.40 | 777.82 | 0.45 | 1.23E-02 |
| ENSMUSG00000063142 | GSMG0012118 | Kcnma1 | potassium large conductance calcium-activated channel, subfamily M, a | 1656.17 | 1211.18 | 0.45 | 1.86E-02 |
| NULL | GSMG0012119 | NULL | NULL | 1045.30 | 1044.76 | 0.00 | 1.00E+00 |
| NULL | GSMG0012121 | NULL | NULL | 354.72 | 230.99 | 0.62 | 1.42E-02 |
| ENSMUSG00000021782 | GSMG0012124 | Dlg5 | discs, large homolog 5 (Drosophila) | 590.81 | 799.00 | -0.44 | 4.54E-02 |
| ENSMUSG00000025280 | GSMG0012127 | Polr3a | polymerase (RNA) III (DNA directed) polypeptide A | 488.88 | 563.06 | -0.20 | 3.87E-01 |
| ENSMUSG00000114922 | GSMG0012137 | Ppifos | peptidylprolyl isomerase F, opposite strand | 177.03 | 258.80 | -0.55 | 3.22E-02 |
| ENSMUSG00000055538 | GSMG0012139 | Zcchc24 | zinc finger, CCHC domain containing 24 | 356.26 | 280.69 | 0.34 | 3.04E-01 |
| ENSMUSG00000021870 | GSMG0012143 | Slmap | sarcolemma associated protein | 416.47 | 421.54 | -0.02 | 9.56E-01 |
| ENSMUSG00000043702 | GSMG0012145 | Pde12 | phosphodiesterase 12 | 691.05 | 831.24 | -0.27 | 1.70E-01 |
| ENSMUSG00000040760 | GSMG0012146 | Appl1 | adaptor protein, phosphotyrosine interaction, PH domain and leucine zi | 941.91 | 1081.04 | -0.20 | 3.48E-01 |
| ENSMUSG00000046753 | GSMG0012151 | Ccdc66 | coiled-coil domain containing 66 | 547.81 | 527.17 | 0.06 | 8.66E-01 |
| ENSMUSG00000021991 | GSMG0012164 | Cacna2d3 | calcium channel, voltage-dependent, alpha2/delta subunit 3 | 1586.66 | 1438.86 | 0.14 | 6.42E-01 |
| ENSMUSG00000015968 | GSMG0012167 | Cacna1d | calcium channel, voltage-dependent, L type, alpha 1D subunit | 435.59 | 437.22 | -0.01 | 9.90E-01 |
| ENSMUSG00000021948 | GSMG0012170 | Prkcd | protein kinase C, delta | 926.24 | 795.98 | 0.22 | 3.05E-01 |
| ENSMUSG00000006522 | GSMG0012172 | Itih3 | inter-alpha trypsin inhibitor, heavy chain 3 | 511.17 | 228.85 | 1.16 | 4.11E-08 |
| ENSMUSG00000021917 | GSMG0012174 | Spcs1 | signal peptidase complex subunit 1 homolog (S. cerevisiae) | 654.94 | 746.83 | -0.19 | 5.02E-01 |
| ENSMUSG00000042354 | GSMG0012175 | Gnl3 | guanine nucleotide binding protein-like 3 (nucleolar) | 990.89 | 1503.32 | -0.60 | 8.02E-04 |
| ENSMUSG00000042286 | GSMG0012178 | Stab1 | stabilin 1 | 3535.35 | 3462.94 | 0.03 | 9.32E-01 |
| ENSMUSG00000021910 | GSMG0012179 | Nisch | nischarin | 4852.18 | 4576.99 | 0.08 | 7.17E-01 |
| ENSMUSG00000021902 | GSMG0012180 | Phf7 | PHD finger protein 7 | 167.40 | 182.21 | -0.13 | 7.53E-01 |
| ENSMUSG00000021892 | GSMG0012185 | Sh3bp5 | SH3-domain binding protein 5 (BTK-associated) | 1197.61 | 2385.46 | -0.99 | 3.03E-11 |
| ENSMUSG00000021891 | GSMG0012188 | Mettl6 | methyltransferase like 6 | 464.26 | 515.97 | -0.15 | 5.72E-01 |
| ENSMUSG00000021884 | GSMG0012193 | Hacl1 | 2-hydroxyacyl-CoA lyase 1 | 385.63 | 302.87 | 0.35 | 1.24E-01 |
| ENSMUSG00000014496 | GSMG0012195 | Ankrd28 | ankyrin repeat domain 28 | 622.01 | 756.87 | -0.28 | 3.12E-01 |
| ENSMUSG00000021905 | GSMG0012202 | Dph3 | diphthamine biosynthesis 3 | 1252.55 | 1278.04 | -0.03 | 9.25E-01 |
| NULL | GSMG0012204 | NULL | NULL | 106.34 | 149.85 | -0.49 | 1.04E-01 |

| | | | | | | | |
|---|---|---|---|---|---|---|---|
| ENSMUSG00000021936 | GSMG0012216 | Mapk8 | mitogen-activated protein kinase 8 | 329.61 | 380.60 | -0.21 | 5.17E-01 |
| ENSMUSG00000041471 | GSMG0012224 | Fam35a | family with sequence similarity 35, member A | 192.42 | 201.82 | -0.07 | 8.71E-01 |
| ENSMUSG00000021796 | GSMG0012227 | Bmpr1a | bone morphogenetic protein receptor, type 1A | 469.63 | 417.67 | 0.17 | 6.07E-01 |
| ENSMUSG00000058690 | GSMG0012237 | Ccser2 | coiled-coil serine rich 2 | 1571.59 | 1680.67 | -0.10 | 7.59E-01 |
| ENSMUSG00000041028 | GSMG0012243 | Ghitm | growth hormone inducible transmembrane protein | 2947.08 | 2562.33 | 0.20 | 3.30E-01 |
| ENSMUSG00000041014 | GSMG0012245 | Nrg3 | neuregulin 3 | 73.54 | 73.06 | 0.01 | 9.85E-01 |
| NULL | GSMG0012246 | NULL | NULL | 282.82 | 258.08 | 0.14 | 7.21E-01 |
| NULL | GSMG0012247 | NULL | NULL | 216.23 | 167.99 | 0.36 | 3.03E-01 |
| NULL | GSMG0012250 | NULL | NULL | 556.71 | 502.71 | 0.15 | 6.40E-01 |
| NULL | GSMG0012251 | NULL | NULL | 716.56 | 672.52 | 0.09 | 8.12E-01 |
| NULL | GSMG0012252 | NULL | NULL | 520.01 | 532.61 | -0.03 | 9.31E-01 |
| NULL | GSMG0012253 | NULL | NULL | 250.83 | 296.65 | -0.24 | 4.32E-01 |
| NULL | GSMG0012254 | NULL | NULL | 488.91 | 582.00 | -0.25 | 3.96E-01 |
| NULL | GSMG0012256 | NULL | NULL | 306.36 | 262.11 | 0.23 | 5.29E-01 |
| NULL | GSMG0012257 | NULL | NULL | 430.55 | 503.69 | -0.23 | 4.28E-01 |
| NULL | GSMG0012260 | NULL | NULL | 595.94 | 519.19 | 0.20 | 5.32E-01 |
| NULL | GSMG0012261 | NULL | NULL | 563.94 | 553.34 | 0.03 | 9.42E-01 |
| NULL | GSMG0012262 | NULL | NULL | 520.88 | 532.74 | -0.03 | 9.50E-01 |
| NULL | GSMG0012263 | NULL | NULL | 267.09 | 252.92 | 0.08 | 8.75E-01 |
| NULL | GSMG0012264 | NULL | NULL | 440.86 | 520.61 | -0.24 | 3.57E-01 |
| NULL | GSMG0012265 | NULL | NULL | 178.72 | 188.12 | -0.07 | 8.86E-01 |
| ENSMUSG00000037824 | GSMG0012273 | Tspan14 | tetraspanin 14 | 1746.16 | 2143.21 | -0.30 | 8.32E-02 |
| ENSMUSG00000021792 | GSMG0012276 | Fam213a | family with sequence similarity 213, member A | 1414.32 | 1631.60 | -0.21 | 4.08E-01 |
| ENSMUSG00000021830 | GSMG0012294 | Txndc16 | thioredoxin domain containing 16 | 223.51 | 211.69 | 0.08 | 8.48E-01 |
| ENSMUSG00000021831 | GSMG0012296 | Ero1l | ERO1-like (S. cerevisiae) | 697.64 | 680.74 | 0.03 | 9.15E-01 |
| ENSMUSG00000037722 | GSMG0012297 | Gnpnat1 | glucosamine-phosphate N-acetyltransferase 1 | 470.33 | 489.88 | -0.06 | 8.82E-01 |
| ENSMUSG00000037712 | GSMG0012298 | Fermt2 | fermitin family member 2 | 6559.42 | 6166.22 | 0.09 | 7.54E-01 |
| ENSMUSG00000037697 | GSMG0012300 | Ddhd1 | DDHD domain containing 1 | 544.79 | 433.51 | 0.33 | 2.02E-01 |
| ENSMUSG00000021835 | GSMG0012303 | Bmp4 | bone morphogenetic protein 4 | 135.29 | 94.86 | 0.51 | 1.18E-01 |
| ENSMUSG00000015759 | GSMG0012306 | Cnih1 | cornichon family AMPA receptor auxiliary protein 1 | 1804.60 | 1967.71 | -0.12 | 5.87E-01 |
| ENSMUSG00000062014 | GSMG0012308 | Gmfb | glia maturation factor, beta | 3718.94 | 4099.95 | -0.14 | 5.34E-01 |
| ENSMUSG00000037580 | GSMG0012315 | Gch1 | GTP cyclohydrolase 1 | 1311.50 | 649.81 | 1.01 | 2.64E-09 |
| ENSMUSG00000037572 | GSMG0012316 | Wdhd1 | WD repeat and HMG-box DNA binding protein 1 | 60.06 | 102.31 | -0.76 | 3.89E-02 |

| | | | | | | | |
|---|---|---|---|---|---|---|---|
| ENSMUSG00000037544 | GSMG0012318 | Dlgap5 | discs, large (Drosophila) homolog-associated protein 5 | 155.69 | 362.20 | -1.22 | 1.21E-07 |
| ENSMUSG00000037526 | GSMG0012319 | Atg14 | autophagy related 14 | 971.99 | 921.34 | 0.08 | 7.81E-01 |
| ENSMUSG00000061244 | GSMG0012332 | Exoc5 | exocyst complex component 5 | 2225.93 | 2919.57 | -0.39 | 3.75E-02 |
| ENSMUSG00000021852 | GSMG0012335 | Slc35f4 | solute carrier family 35, member F4 | 110.75 | 172.43 | -0.64 | 2.45E-02 |
| ENSMUSG00000036242 | GSMG0012337 | 3632451O06Rik | RIKEN cDNA 3632451O06 gene | 286.90 | 346.17 | -0.27 | 3.45E-01 |
| ENSMUSG00000006288 | GSMG0012349 | Ttc5 | tetratricopeptide repeat domain 5 | 1172.72 | 1221.03 | -0.06 | 8.35E-01 |
| NULL | GSMG0012353 | NULL | NULL | 193.69 | 102.99 | 0.91 | 1.54E-03 |
| ENSMUSG00000006289 | GSMG0012354 | Osgep | O-sialoglycoprotein endopeptidase | 809.17 | 836.91 | -0.05 | 8.79E-01 |
| ENSMUSG00000035953 | GSMG0012355 | Tmem55b | transmembrane protein 55b | 754.33 | 720.81 | 0.07 | 8.16E-01 |
| ENSMUSG00000004558 | GSMG0012369 | Ndrg2 | N-myc downstream regulated gene 2 | 5567.79 | 4368.41 | 0.35 | 6.88E-02 |
| ENSMUSG00000097633 | GSMG0012371 | Gm16617 | predicted gene, 16617 | 97.54 | 139.09 | -0.52 | 1.28E-01 |
| ENSMUSG00000049295 | GSMG0012372 | Zfp219 | zinc finger protein 219 | 476.80 | 389.80 | 0.29 | 2.27E-01 |
| ENSMUSG00000060373 | GSMG0012374 | Hnrnpc | heterogeneous nuclear ribonucleoprotein C | 2146.93 | 2746.94 | -0.36 | 7.21E-02 |
| NULL | GSMG0012375 | NULL | NULL | 808.72 | 713.35 | 0.18 | 5.66E-01 |
| ENSMUSG00000035726 | GSMG0012376 | Supt16 | suppressor of Ty 16 | 546.46 | 777.25 | -0.51 | 7.43E-03 |
| ENSMUSG00000053754 | GSMG0012378 | Chd8 | chromodomain helicase DNA binding protein 8 | 228.79 | 226.77 | 0.01 | 9.83E-01 |
| ENSMUSG00000022159 | GSMG0012380 | Rab2b | RAB2B, member RAS oncogene family | 773.39 | 740.21 | 0.06 | 8.30E-01 |
| ENSMUSG00000022160 | GSMG0012381 | Mettl3 | methyltransferase like 3 | 720.77 | 688.70 | 0.07 | 8.21E-01 |
| ENSMUSG00000022174 | GSMG0012389 | Dad1 | defender against cell death 1 | 3595.36 | 4202.28 | -0.23 | 2.68E-01 |
| ENSMUSG00000000958 | GSMG0012393 | Slc7a7 | solute carrier family 7 (cationic amino acid transporter, y+ system), mer | 209.45 | 557.64 | -1.41 | 1.73E-11 |
| NULL | GSMG0012394 | NULL | NULL | 2493.82 | 2486.66 | 0.00 | 9.90E-01 |
| ENSMUSG00000023110 | GSMG0012395 | Prmt5 | protein arginine N-methyltransferase 5 | 1497.56 | 2163.76 | -0.53 | 7.66E-04 |
| ENSMUSG00000022177 | GSMG0012396 | Haus4 | HAUS augmin-like complex, subunit 4 | 384.24 | 419.47 | -0.13 | 6.80E-01 |
| ENSMUSG00000022178 | GSMG0012397 | Ajuba | ajuba LIM protein | 1419.15 | 1373.89 | 0.05 | 8.74E-01 |
| ENSMUSG00000022185 | GSMG0012401 | Acin1 | apoptotic chromatin condensation inducer 1 | 286.83 | 279.86 | 0.03 | 9.55E-01 |
| ENSMUSG00000022180 | GSMG0012403 | Slc7a8 | solute carrier family 7 (cationic amino acid transporter, y+ system), mer | 1481.37 | 3456.42 | -1.22 | 2.10E-17 |
| ENSMUSG00000057156 | GSMG0012404 | Homez | homeodomain leucine zipper-encoding gene | 522.86 | 490.55 | 0.09 | 7.74E-01 |
| ENSMUSG00000022199 | GSMG0012406 | Slc22a17 | solute carrier family 22 (organic cation transporter), member 17 | 6551.72 | 6703.44 | -0.03 | 9.28E-01 |
| ENSMUSG00000022203 | GSMG0012407 | Efs | embryonal Fyn-associated substrate | 85.26 | 129.26 | -0.61 | 1.07E-01 |
| NULL | GSMG0012411 | NULL | NULL | 87.05 | 47.83 | 0.86 | 2.96E-02 |
| ENSMUSG00000040701 | GSMG0012413 | Ap1g2 | adaptor protein complex AP-1, gamma 2 subunit | 568.70 | 578.55 | -0.02 | 9.52E-01 |
| ENSMUSG00000022208 | GSMG0012414 | Jph4 | junctophilin 4 | 744.69 | 978.64 | -0.40 | 2.25E-01 |
| ENSMUSG00000022217 | GSMG0012420 | Emc9 | ER membrane protein complex subunit 9 | 432.97 | 493.71 | -0.19 | 4.90E-01 |

| | | | | | | | |
|---|---|---|---|---|---|---|---|
| ENSMUSG00000079197 | GSMG0012421 | Psme2 | proteasome (prosome, macropain) activator subunit 2 (PA28 beta) | 619.61 | 753.71 | -0.28 | 2.48E-01 |
| ENSMUSG00000002319 | GSMG0012422 | Ipo4 | importin 4 | 2213.02 | 3168.74 | -0.52 | 5.58E-03 |
| ENSMUSG00000002320 | GSMG0012423 | Tm9sf1 | transmembrane 9 superfamily member 1 | 2505.58 | 2414.87 | 0.05 | 8.38E-01 |
| ENSMUSG00000002329 | GSMG0012424 | Mdp1 | magnesium-dependent phosphatase 1 | 1550.12 | 1416.34 | 0.13 | 5.53E-01 |
| ENSMUSG00000010376 | GSMG0012425 | Nedd8 | neural precursor cell expressed, developmentally down-regulated gene 8 | 2288.56 | 2800.76 | -0.29 | 1.45E-01 |
| ENSMUSG00000007589 | GSMG0012427 | Tinf2 | Terf1 (TRF1)-interacting nuclear factor 2 | 558.17 | 656.28 | -0.23 | 3.29E-01 |
| ENSMUSG00000040472 | GSMG0012429 | Rabggta | Rab geranylgeranyl transferase, a subunit | 1302.80 | 1645.45 | -0.34 | 5.02E-02 |
| ENSMUSG00000002332 | GSMG0012430 | Dhrs1 | dehydrogenase/reductase (SDR family) member 1 | 1446.15 | 1304.22 | 0.15 | 5.36E-01 |
| ENSMUSG00000022220 | GSMG0012432 | Adcy4 | adenylate cyclase 4 | 2695.14 | 1983.52 | 0.44 | 8.80E-03 |
| ENSMUSG00000022221 | GSMG0012433 | Ripk3 | receptor-interacting serine-threonine kinase 3 | 126.40 | 169.47 | -0.42 | 1.67E-01 |
| ENSMUSG00000022223 | GSMG0012435 | Sdr39u1 | short chain dehydrogenase/reductase family 39U, member 1 | 209.95 | 184.59 | 0.19 | 5.99E-01 |
| ENSMUSG00000021938 | GSMG0012455 | Pspc1 | paraspeckle protein 1 | 448.74 | 666.96 | -0.57 | 1.93E-03 |
| ENSMUSG00000040123 | GSMG0012458 | Zmym5 | zinc finger, MYM-type 5 | 712.23 | 684.73 | 0.05 | 8.67E-01 |
| ENSMUSG00000046352 | GSMG0012462 | Gjb2 | gap junction protein, beta 2 | 622.28 | 554.62 | 0.17 | 6.67E-01 |
| ENSMUSG00000040055 | GSMG0012464 | Gjb6 | gap junction protein, beta 6 | 165.96 | 79.59 | 1.06 | 1.95E-04 |
| ENSMUSG00000021947 | GSMG0012465 | Cryl1 | crystallin, lambda 1 | 147.48 | 130.01 | 0.19 | 6.33E-01 |
| ENSMUSG00000021951 | GSMG0012468 | Eef1akmt1 | eukaryotic translation elongation factor 1 alpha lysine methyltransferase | 250.39 | 335.73 | -0.42 | 9.01E-02 |
| G00000021952,ENSMUSG000 | GSMG0012470 | Xpo4 // Lats2 | exportin 4 // large tumor suppressor 2 | 4505.06 | 3939.90 | 0.19 | 3.44E-01 |
| ENSMUSG00000021965 | GSMG0012472 | Ska3 | spindle and kinetochore associated complex subunit 3 | 63.91 | 216.60 | -1.76 | 1.55E-09 |
| ENSMUSG00000021969 | GSMG0012473 | Zdhhc20 | zinc finger, DHHC domain containing 20 | 1286.34 | 1492.14 | -0.21 | 3.80E-01 |
| ENSMUSG00000021973 | GSMG0012474 | Micu2 | mitochondrial calcium uptake 2 | 918.31 | 1056.07 | -0.20 | 4.38E-01 |
| ENSMUSG00000021982 | GSMG0012482 | Cdadc1 | cytidine and dCMP deaminase domain containing 1 | 1019.35 | 994.36 | 0.04 | 9.12E-01 |
| ENSMUSG00000021983 | GSMG0012484 | Atp8a2 | ATPase, aminophospholipid transporter-like, class I, type 8A, member 2 | 965.04 | 1448.24 | -0.59 | 2.74E-03 |
| ENSMUSG00000063895 | GSMG0012488 | Nupl1 | nucleoporin like 1 | 541.36 | 694.47 | -0.36 | 6.71E-02 |
| NULL | GSMG0012489 | NULL | NULL | 213.38 | 381.21 | -0.84 | 1.12E-03 |
| NULL | GSMG0012491 | NULL | NULL | 479.77 | 709.53 | -0.57 | 1.02E-02 |
| ENSMUSG00000060548 | GSMG0012493 | Tnfrsf19 | tumor necrosis factor receptor superfamily, member 19 | 3457.83 | 2472.21 | 0.48 | 2.95E-03 |
| ENSMUSG00000021928 | GSMG0012499 | Ebpl | emopamil binding protein-like | 595.76 | 900.71 | -0.60 | 1.26E-03 |
| ENSMUSG00000021929 | GSMG0012500 | Kpna3 | karyopherin (importin) alpha 3 | 729.68 | 950.23 | -0.38 | 3.70E-02 |
| NULL | GSMG0012503 | NULL | NULL | 90.06 | 59.03 | 0.61 | 1.10E-01 |
| ENSMUSG00000021930 | GSMG0012504 | Sprydd7 | SPRY domain containing 7 | 317.66 | 402.89 | -0.35 | 1.32E-01 |
| ENSMUSG00000035161 | GSMG0012512 | Ints6 | integrator complex subunit 6 | 1104.65 | 891.60 | 0.31 | 1.08E-01 |
| ENSMUSG00000021273 | GSMG0012518 | Fdft1 | farnesyl diphosphate farnesyl transferase 1 | 482.87 | 1193.00 | -1.30 | 2.09E-14 |

| | | | | | | | |
|---|---|---|---|---|---|---|---|
| ENSMUSG00000035078 | GSMG0012525 | Mtmr9 | myotubularin related protein 9 | 527.49 | 799.23 | -0.60 | 1.30E-03 |
| ENSMUSG00000054733 | GSMG0012532 | Msra | methionine sulfoxide reductase A | 331.43 | 428.15 | -0.37 | 8.88E-02 |
| NULL | GSMG0012534 | NULL | NULL | 256.20 | 381.61 | -0.57 | 1.13E-02 |
| NULL | GSMG0012535 | NULL | NULL | 113.57 | 158.16 | -0.48 | 1.86E-01 |
| ENSMUSG00000021972 | GSMG0012540 | Hmbox1 | homeobox containing 1 | 251.65 | 229.95 | 0.13 | 7.18E-01 |
| ENSMUSG00000021978 | GSMG0012543 | Extl3 | exostoses (multiple)-like 3 | 2983.80 | 2445.62 | 0.29 | 9.79E-02 |
| ENSMUSG00000007989 | GSMG0012548 | Fzd3 | frizzled class receptor 3 | 287.10 | 342.01 | -0.25 | 4.26E-01 |
| ENSMUSG00000045731 | GSMG0012552 | Pnoc | prepronociceptin | 88.86 | 82.15 | 0.12 | 8.42E-01 |
| ENSMUSG00000022031 | GSMG0012553 | Elp3 | elongator acetyltransferase complex subunit 3 | 590.63 | 735.24 | -0.31 | 1.54E-01 |
| ENSMUSG00000022034 | GSMG0012554 | Esco2 | establishment of sister chromatid cohesion N-acetyltransferase 2 | 94.63 | 233.50 | -1.31 | 2.80E-06 |
| ENSMUSG00000034463 | GSMG0012556 | Scara3 | scavenger receptor class A, member 3 | 669.79 | 511.11 | 0.39 | 2.11E-01 |
| ENSMUSG00000022040 | GSMG0012559 | Ephx2 | epoxide hydrolase 2, cytoplasmic | 156.05 | 93.28 | 0.75 | 8.09E-03 |
| ENSMUSG00000059456 | GSMG0012560 | Ptk2b | PTK2 protein tyrosine kinase 2 beta | 1118.99 | 851.44 | 0.39 | 9.03E-02 |
| ENSMUSG00000022048 | GSMG0012570 | Dpysl2 | dihydropyrimidinase-like 2 | 162.39 | 234.44 | -0.52 | 9.60E-02 |
| ENSMUSG00000022052 | GSMG0012574 | Ppp2r2a | protein phosphatase 2, regulatory subunit B, alpha | 969.63 | 1179.10 | -0.28 | 1.80E-01 |
| ENSMUSG00000048922 | GSMG0012581 | Cdca2 | cell division cycle associated 2 | 35.95 | 142.08 | -1.97 | 6.57E-09 |
| ENSMUSG00000022054 | GSMG0012585 | Nefm | neurofilament, medium polypeptide | 707.17 | 531.90 | 0.41 | 6.71E-02 |
| ENSMUSG00000034248 | GSMG0012592 | Slc25a37 | solute carrier family 25, member 37 | 1155.49 | 1054.71 | 0.13 | 6.00E-01 |
| ENSMUSG00000034194 | GSMG0012602 | R3hcc1 | R3H domain and coiled-coil containing 1 | 829.25 | 1183.11 | -0.51 | 2.09E-03 |
| ENSMUSG00000034190 | GSMG0012603 | Chmp7 | charged multivesicular body protein 7 | 1028.27 | 1223.13 | -0.25 | 1.80E-01 |
| ENSMUSG00000022075 | GSMG0012604 | Rhobtb2 | Rho-related BTB domain containing 2 | 484.07 | 358.29 | 0.43 | 6.45E-02 |
| ENSMUSG00000033712 | GSMG0012606 | Ccar2 | cell cycle activator and apoptosis regulator 2 | 2321.80 | 2777.56 | -0.26 | 1.74E-01 |
| ENSMUSG00000044551 | GSMG0012607 | 9930012K11Rik | RIKEN cDNA 9930012K11 gene | 404.14 | 414.51 | -0.04 | 9.14E-01 |
| ENSMUSG00000022090 | GSMG0012608 | Pdlim2 | PDZ and LIM domain 2 | 940.47 | 617.81 | 0.61 | 3.42E-04 |
| ENSMUSG00000022091 | GSMG0012609 | Sorbs3 | sorbin and SH3 domain containing 3 | 2334.57 | 2107.27 | 0.15 | 4.83E-01 |
| ENSMUSG00000022092 | GSMG0012610 | Ppp3cc | protein phosphatase 3, catalytic subunit, gamma isoform | 1091.78 | 750.27 | 0.54 | 1.15E-03 |
| ENSMUSG00000000776 | GSMG0012614 | Polr3d | polymerase (RNA) III (DNA directed) polypeptide D | 391.55 | 570.55 | -0.54 | 3.47E-03 |
| ENSMUSG00000022098 | GSMG0012615 | Bmp1 | bone morphogenetic protein 1 | 1554.47 | 1137.94 | 0.45 | 1.12E-02 |
| ENSMUSG00000022095 | GSMG0012617 | Fam160b2 | family with sequence similarity 160, member B2 | 1224.36 | 996.52 | 0.30 | 2.63E-01 |
| ENSMUSG00000022099 | GSMG0012618 | Dmtn | dematin actin binding protein | 3198.52 | 2936.63 | 0.12 | 5.81E-01 |
| ENSMUSG00000022100 | GSMG0012622 | Xpo7 | exportin 7 | 1691.70 | 2019.92 | -0.26 | 2.16E-01 |
| NULL | GSMG0012627 | NULL | NULL | 364.69 | 296.94 | 0.29 | 4.29E-01 |
| NULL | GSMG0012628 | NULL | NULL | 432.38 | 309.25 | 0.48 | 2.77E-02 |

| | | | | | | | |
|---|---|---|---|---|---|---|---|
| ENSMUSG00000033487 | GSMG0012632 | Fndc3a | fibronectin type III domain containing 3A | 886.45 | 954.42 | -0.11 | 6.59E-01 |
| ENSMUSG00000033470 | GSMG0012641 | Cysltr2 | cysteinyl leukotriene receptor 2 | 322.84 | 106.21 | 1.60 | 1.92E-11 |
| ENSMUSG00000022105 | GSMG0012642 | Rb1 | retinoblastoma 1 | 774.40 | 626.83 | 0.30 | 1.89E-01 |
| ENSMUSG00000022108 | GSMG0012650 | Itm2b | integral membrane protein 2B | 21127.46 | 15055.78 | 0.49 | 2.87E-03 |
| ENSMUSG00000068015 | GSMG0012658 | Lrch1 | leucine-rich repeats and calponin homology (CH) domain containing 1 | 647.56 | 653.22 | -0.01 | 9.69E-01 |
| ENSMUSG00000034893 | GSMG0012669 | Cog3 | component of oligomeric golgi complex 3 | 496.32 | 560.89 | -0.17 | 5.07E-01 |
| NULL | GSMG0012670 | NULL | NULL | 243.67 | 284.58 | -0.22 | 4.17E-01 |
| ENSMUSG00000022003 | GSMG0012671 | Slc25a30 | solute carrier family 25, member 30 | 106.91 | 119.27 | -0.15 | 7.63E-01 |
| ENSMUSG00000067995 | GSMG0012672 | Gtf2f2 | general transcription factor IIF, polypeptide 2 | 188.70 | 287.96 | -0.61 | 8.83E-03 |
| ENSMUSG00000022008 | GSMG0012676 | Gpalpp1 | GPALPP motifs containing 1 | 256.60 | 284.66 | -0.15 | 6.66E-01 |
| ENSMUSG00000052584 | GSMG0012682 | Serp2 | stress-associated endoplasmic reticulum protein family member 2 | 504.69 | 866.58 | -0.78 | 5.45E-06 |
| ENSMUSG00000044350 | GSMG0012687 | Lacc1 | laccase (multicopper oxidoreductase) domain containing 1 | 274.92 | 284.32 | -0.05 | 9.03E-01 |
| ENSMUSG00000022013 | GSMG0012694 | Dnajc15 | DnaJ heat shock protein family (Hsp40) member C15 | 539.31 | 649.29 | -0.27 | 2.97E-01 |
| ENSMUSG00000034731 | GSMG0012703 | Dgkh | diacylglycerol kinase, eta | 2202.47 | 1709.50 | 0.37 | 1.50E-01 |
| ENSMUSG00000022018 | GSMG0012711 | Rgcc | regulator of cell cycle | 2631.43 | 2339.57 | 0.17 | 5.14E-01 |
| ENSMUSG00000022020 | GSMG0012713 | Naa16 | N(alpha)-acetyltransferase 16, NatA auxiliary subunit | 619.66 | 657.24 | -0.09 | 8.05E-01 |
| ENSMUSG00000022023 | GSMG0012714 | Wbp4 | WW domain binding protein 4 | 580.89 | 576.01 | 0.01 | 9.81E-01 |
| ENSMUSG00000036422 | GSMG0012716 | Pcdh8 | protocadherin 8 | 589.91 | 1168.72 | -0.99 | 8.04E-06 |
| ENSMUSG00000055421 | GSMG0012733 | Pcdh9 | protocadherin 9 | 12560.15 | 9220.74 | 0.45 | 1.87E-02 |
| NULL | GSMG0012744 | NULL | NULL | 409.44 | 224.21 | 0.87 | 9.35E-05 |
| NULL | GSMG0012745 | NULL | NULL | 576.93 | 388.46 | 0.57 | 1.37E-02 |
| NULL | GSMG0012746 | NULL | NULL | 272.27 | 172.22 | 0.66 | 1.16E-02 |
| NULL | GSMG0012748 | NULL | NULL | 343.10 | 264.69 | 0.38 | 1.54E-01 |
| NULL | GSMG0012753 | NULL | NULL | 390.01 | 227.31 | 0.78 | 1.94E-02 |
| NULL | GSMG0012754 | NULL | NULL | 295.53 | 147.99 | 1.00 | 1.01E-03 |
| NULL | GSMG0012758 | NULL | NULL | 459.12 | 325.98 | 0.49 | 3.79E-02 |
| NULL | GSMG0012760 | NULL | NULL | 352.16 | 241.40 | 0.55 | 1.59E-02 |
| NULL | GSMG0012761 | NULL | NULL | 135.74 | 109.73 | 0.31 | 5.29E-01 |
| NULL | GSMG0012766 | NULL | NULL | 1061.74 | 889.50 | 0.26 | 2.40E-01 |
| ENSMUSG00000022076 | GSMG0012769 | Klhl1 | kelch-like 1 | 26.57 | 244.78 | -3.19 | 2.44E-28 |
| ENSMUSG00000055639 | GSMG0012771 | Dach1 | dachshund 1 (Drosophila) | 605.19 | 703.76 | -0.22 | 3.51E-01 |
| NULL | GSMG0012780 | NULL | NULL | 280.13 | 245.72 | 0.19 | 6.41E-01 |
| ENSMUSG00000033186 | GSMG0012787 | Mzt1 | mitotic spindle organizing protein 1 | 560.39 | 834.52 | -0.58 | 5.48E-02 |

| | | | | | | | |
|---|---|---|---|---|---|---|---|
| ENSMUSG00000033166 | GSMG0012789 | Dis3 | DIS3 homolog, exosome endoribonuclease and 3'-5' exoribonuclease | 458.99 | 643.96 | -0.49 | 1.12E-02 |
| ENSMUSG00000072294 | GSMG0012792 | Klf12 | Kruppel-like factor 12 | 649.25 | 450.80 | 0.53 | 2.06E-02 |
| NULL | GSMG0012794 | NULL | NULL | 104.71 | 87.91 | 0.26 | 5.63E-01 |
| NULL | GSMG0012798 | NULL | NULL | 245.47 | 208.74 | 0.24 | 4.37E-01 |
| ENSMUSG00000033083 | GSMG0012801 | Tbc1d4 | TBC1 domain family, member 4 | 350.61 | 130.79 | 1.42 | 3.47E-08 |
| ENSMUSG00000075486 | GSMG0012805 | Commd6 | COMM domain containing 6 | 649.42 | 648.80 | 0.00 | 1.00E+00 |
| ENSMUSG00000022124 | GSMG0012810 | Fbxl3 | F-box and leucine-rich repeat protein 3 | 518.88 | 468.48 | 0.15 | 5.95E-01 |
| ENSMUSG00000033004 | GSMG0012811 | Mycbp2 | MYC binding protein 2 | 5591.97 | 6849.13 | -0.29 | 6.04E-02 |
| ENSMUSG00000022122 | GSMG0012816 | Ednrb | endothelin receptor type B | 1090.10 | 1830.57 | -0.75 | 5.83E-06 |
| ENSMUSG00000022120 | GSMG0012824 | Rnf219 | ring finger protein 219 | 61.74 | 87.07 | -0.50 | 2.58E-01 |
| G00000022119,ENSMUSG000 | GSMG0012825 | Gm17066 // Rbm2 | predicted gene 17066 // RNA binding motif protein 26 | 1139.89 | 1105.14 | 0.05 | 9.00E-01 |
| ENSMUSG00000022114 | GSMG0012832 | Spry2 | sprouty homolog 2 (Drosophila) | 245.55 | 174.59 | 0.49 | 7.44E-02 |
| ENSMUSG00000075478 | GSMG0012841 | Slitrk1 | SLIT and NTRK-like family, member 1 | 521.10 | 576.54 | -0.15 | 6.58E-01 |
| NULL | GSMG0012848 | NULL | NULL | 442.48 | 365.27 | 0.27 | 3.42E-01 |
| ENSMUSG00000022130 | GSMG0012851 | Tgds | TDP-glucose 4,6-dehydratase | 314.63 | 375.22 | -0.25 | 3.42E-01 |
| NULL | GSMG0012854 | NULL | NULL | 166.60 | 256.26 | -0.62 | 1.58E-02 |
| ENSMUSG00000032849 | GSMG0012855 | Abcc4 | ATP-binding cassette, sub-family C (CFTR/MRP), member 4 | 5596.56 | 3096.86 | 0.85 | 7.66E-10 |
| ENSMUSG00000042156 | GSMG0012858 | Dzip1 | DAZ interacting protein 1 | 151.32 | 197.17 | -0.38 | 1.81E-01 |
| NULL | GSMG0012860 | NULL | NULL | 224.67 | 142.40 | 0.65 | 7.42E-02 |
| NULL | GSMG0012861 | NULL | NULL | 291.67 | 310.97 | -0.09 | 7.97E-01 |
| NULL | GSMG0012868 | NULL | NULL | 517.64 | 392.79 | 0.40 | 1.09E-01 |
| ENSMUSG00000063410 | GSMG0012871 | Stk24 | serine/threonine kinase 24 | 212.88 | 246.90 | -0.22 | 5.50E-01 |
| ENSMUSG00000025558 | GSMG0012875 | Dock9 | dedicator of cytokinesis 9 | 6654.73 | 8109.34 | -0.29 | 2.11E-01 |
| NULL | GSMG0012876 | NULL | NULL | 412.22 | 1003.47 | -1.28 | 2.04E-13 |
| NULL | GSMG0012877 | NULL | NULL | 94.03 | 73.94 | 0.34 | 4.45E-01 |
| NULL | GSMG0012886 | NULL | NULL | 70.59 | 66.33 | 0.08 | 9.03E-01 |
| ENSMUSG00000041703 | GSMG0012887 | Zic5 | zinc finger protein of the cerebellum 5 | 174.14 | 121.55 | 0.51 | 1.36E-01 |
| ENSMUSG00000041625 | GSMG0012889 | Ggact | gamma-glutamylamine cyclotransferase | 94.22 | 73.13 | 0.37 | 3.63E-01 |
| ENSMUSG00000041594 | GSMG0012890 | Tmtc4 | transmembrane and tetratricopeptide repeat containing 4 | 665.15 | 1271.61 | -0.93 | 5.12E-09 |
| ENSMUSG00000025551 | GSMG0012897 | Fgf14 | fibroblast growth factor 14 | 478.12 | 614.08 | -0.36 | 2.13E-01 |
| NULL | GSMG0012901 | NULL | NULL | 546.28 | 515.78 | 0.08 | 8.26E-01 |
| NULL | GSMG0012904 | NULL | NULL | 133.39 | 102.87 | 0.38 | 3.81E-01 |
| NULL | GSMG0012906 | NULL | NULL | 945.39 | 823.95 | 0.20 | 4.26E-01 |

| | | | | | | | |
|---|---|---|---|---|---|---|---|
| NULL | GSMG0012907 | NULL | NULL | 377.15 | 307.67 | 0.29 | 3.59E-01 |
| NULL | GSMG0012909 | NULL | NULL | 1043.51 | 883.66 | 0.24 | 2.53E-01 |
| NULL | GSMG0012910 | NULL | NULL | 255.96 | 198.77 | 0.36 | 2.70E-01 |
| ENSMUSG00000064373 | GSMG0012911 | Sepp1 | selenoprotein P, plasma, 1 | 20593.24 | 15610.75 | 0.40 | 2.70E-02 |
| NULL | GSMG0012913 | NULL | NULL | 629.98 | 344.61 | 0.87 | 1.06E-05 |
| ENSMUSG00000022186 | GSMG0012916 | Oxct1 | 3-oxoacid CoA transferase 1 | 1800.75 | 2206.09 | -0.29 | 8.71E-02 |
| ENSMUSG00000049148 | GSMG0012917 | Plcxd3 | phosphatidylinositol-specific phospholipase C, X domain containing 3 | 82.58 | 143.41 | -0.79 | 1.14E-02 |
| ENSMUSG00000050697 | GSMG0012925 | Prkaa1 | protein kinase, AMP-activated, alpha 1 catalytic subunit | 703.40 | 714.35 | -0.02 | 9.55E-01 |
| ENSMUSG00000022151 | GSMG0012926 | Ttc33 | tetratricopeptide repeat domain 33 | 986.62 | 935.82 | 0.08 | 8.23E-01 |
| ENSMUSG00000022150 | GSMG0012929 | Dab2 | disabled 2, mitogen-responsive phosphoprotein | 5525.10 | 5145.09 | 0.10 | 6.95E-01 |
| ENSMUSG00000050310 | GSMG0012937 | Rictor | RPTOR independent companion of MTOR, complex 2 | 2656.07 | 2372.95 | 0.16 | 3.90E-01 |
| ENSMUSG00000054263 | GSMG0012940 | Lifr | leukemia inhibitory factor receptor | 1952.31 | 1656.10 | 0.24 | 2.65E-01 |
| ENSMUSG00000022142 | GSMG0012946 | Nup155 | nucleoporin 155 | 173.95 | 224.05 | -0.36 | 2.19E-01 |
| ENSMUSG00000022253 | GSMG0012957 | Nadk2 | NAD kinase 2, mitochondrial | 351.81 | 422.94 | -0.26 | 3.59E-01 |
| ENSMUSG00000022248 | GSMG0012971 | Rad1 | RAD1 checkpoint DNA exonuclease | 146.73 | 188.90 | -0.36 | 2.23E-01 |
| ENSMUSG00000022244 | GSMG0012977 | Amacr | alpha-methylacyl-CoA racemase | 328.47 | 295.03 | 0.16 | 6.43E-01 |
| ENSMUSG00000047497 | GSMG0012980 | Adamts12 | a disintegrin-like and metallopeptidase (reprolysin type) with thrombos | 51.87 | 85.29 | -0.71 | 8.03E-02 |
| ENSMUSG00000022201 | GSMG0012986 | Zfr | zinc finger RNA binding protein | 2844.57 | 3030.09 | -0.09 | 6.82E-01 |
| ENSMUSG00000039458 | GSMG0012990 | Mtmr12 | myotubularin related protein 12 | 934.06 | 966.72 | -0.05 | 8.71E-01 |
| ENSMUSG00000022200 | GSMG0012991 | Golph3 | golgi phosphoprotein 3 | 2785.81 | 2467.92 | 0.17 | 4.44E-01 |
| ENSMUSG00000022191 | GSMG0012994 | Drosha | drosha, ribonuclease type III | 1944.13 | 2191.05 | -0.17 | 3.67E-01 |
| ENSMUSG00000022321 | GSMG0013000 | Cdh10 | cadherin 10 | 337.54 | 456.97 | -0.44 | 6.54E-02 |
| NULL | GSMG0013003 | NULL | NULL | 98.51 | 75.15 | 0.39 | 4.60E-01 |
| NULL | GSMG0013007 | NULL | NULL | 215.87 | 109.07 | 0.99 | 2.42E-04 |
| NULL | GSMG0013008 | NULL | NULL | 282.86 | 96.14 | 1.55 | 3.87E-11 |
| ENSMUSG00000040420 | GSMG0013012 | Cdh18 | cadherin 18 | 217.47 | 125.21 | 0.80 | 1.56E-02 |
| ENSMUSG00000022272 | GSMG0013024 | Myo10 | myosin X | 8946.24 | 9919.67 | -0.15 | 5.59E-01 |
| ENSMUSG00000022270 | GSMG0013028 | Fam134b | family with sequence similarity 134, member B | 193.97 | 162.68 | 0.25 | 5.04E-01 |
| ENSMUSG00000052253 | GSMG0013032 | Zfp622 | zinc finger protein 622 | 272.68 | 307.22 | -0.17 | 5.68E-01 |
| ENSMUSG00000022269 | GSMG0013033 | March11 | membrane-associated ring finger (C3HC4) 11 | 174.65 | 219.05 | -0.33 | 3.02E-01 |
| ENSMUSG00000022265 | GSMG0013036 | Ank | progressive ankylosis | 729.99 | 679.53 | 0.10 | 7.09E-01 |
| NULL | GSMG0013037 | NULL | NULL | 363.33 | 322.81 | 0.17 | 6.07E-01 |
| NULL | GSMG0013039 | NULL | NULL | 249.42 | 244.08 | 0.03 | 9.34E-01 |

| | | | | | | | |
|---|---|---|---|---|---|---|---|
| ENSMUSG00000022240 | GSMG0013044 | Ctnnd2 | catenin (cadherin associated protein), delta 2 | 1790.00 | 2200.66 | -0.30 | 2.15E-01 |
| NULL | GSMG0013045 | NULL | NULL | 662.82 | 652.04 | 0.03 | 9.42E-01 |
| NULL | GSMG0013052 | NULL | NULL | 97.16 | 157.61 | -0.71 | 4.03E-02 |
| NULL | GSMG0013054 | NULL | NULL | 206.30 | 191.24 | 0.11 | 8.04E-01 |
| ENSMUSG00000039168 | GSMG0013056 | Dap | death-associated protein | 1270.15 | 1208.52 | 0.07 | 8.33E-01 |
| ENSMUSG00000022235 | GSMG0013063 | Cmbl | carboxymethylenebutenolidase-like (Pseudomonas) | 202.25 | 173.47 | 0.23 | 5.28E-01 |
| ENSMUSG00000022231 | GSMG0013069 | Sema5a | sema domain, seven thrombospondin repeats (type 1 and type 1-like), tr | 173.75 | 115.58 | 0.59 | 1.26E-01 |
| ENSMUSG00000022261 | GSMG0013073 | Sdc2 | syndecan 2 | 4370.25 | 2973.14 | 0.56 | 2.75E-03 |
| ENSMUSG00000039007 | GSMG0013074 | Cpq | carboxypeptidase Q | 1470.63 | 995.49 | 0.56 | 6.57E-03 |
| ENSMUSG00000022255 | GSMG0013082 | Mtdh | metadherin | 1587.48 | 2183.05 | -0.46 | 1.19E-02 |
| ENSMUSG00000022257 | GSMG0013085 | Laptm4b | lysosomal-associated protein transmembrane 4B | 3844.48 | 4883.72 | -0.34 | 9.64E-02 |
| ENSMUSG00000022324 | GSMG0013087 | Matn2 | matrilin 2 | 140.82 | 130.68 | 0.11 | 8.27E-01 |
| ENSMUSG00000037617 | GSMG0013109 | Spag1 | sperm associated antigen 1 | 298.26 | 275.80 | 0.11 | 7.58E-01 |
| NULL | GSMG0013115 | NULL | NULL | 165.12 | 221.60 | -0.43 | 1.80E-01 |
| ENSMUSG00000022295 | GSMG0013122 | Atp6v1c1 | ATPase, H+ transporting, lysosomal V1 subunit C1 | 2107.98 | 1840.17 | 0.20 | 4.11E-01 |
| ENSMUSG00000022297 | GSMG0013124 | Fzd6 | frizzled class receptor 6 | 1854.32 | 1648.70 | 0.17 | 4.66E-01 |
| ENSMUSG00000054196 | GSMG0013126 | Cthrc1 | collagen triple helix repeat containing 1 | 1963.66 | 1886.14 | 0.06 | 8.63E-01 |
| ENSMUSG00000022300 | GSMG0013127 | Dcaf13 | DDB1 and CUL4 associated factor 13 | 1093.20 | 1248.86 | -0.19 | 3.87E-01 |
| NULL | GSMG0013133 | NULL | NULL | 398.06 | 318.30 | 0.33 | 2.13E-01 |
| ENSMUSG00000037386 | GSMG0013134 | Rims2 | regulating synaptic membrane exocytosis 2 | 851.05 | 757.13 | 0.17 | 5.55E-01 |
| ENSMUSG00000022306 | GSMG0013142 | Zfpm2 | zinc finger protein, multitype 2 | 46.23 | 80.01 | -0.78 | 4.57E-02 |
| ENSMUSG00000022307 | GSMG0013155 | Oxr1 | oxidation resistance 1 | 1695.46 | 1183.80 | 0.52 | 1.05E-03 |
| NULL | GSMG0013159 | NULL | NULL | 62.66 | 25.91 | 1.27 | 3.57E-03 |
| ENSMUSG00000022337 | GSMG0013165 | Emc2 | ER membrane protein complex subunit 2 | 1022.56 | 1027.83 | -0.01 | 9.83E-01 |
| ENSMUSG00000022338 | GSMG0013169 | Eny2 | enhancer of yellow 2 homolog (Drosophila) | 1146.69 | 1509.04 | -0.40 | 3.78E-02 |
| ENSMUSG00000022339 | GSMG0013172 | Ebag9 | estrogen receptor-binding fragment-associated gene 9 | 156.16 | 165.66 | -0.08 | 8.67E-01 |
| NULL | GSMG0013173 | 2310069G16Rik | RIKEN cDNA 2310069G16 gene | 53.50 | 80.72 | -0.60 | 1.31E-01 |
| ENSMUSG00000022313 | GSMG0013177 | Utp23 | UTP23 small subunit processome component | 331.26 | 340.37 | -0.04 | 9.22E-01 |
| ENSMUSG00000038622 | GSMG0013180 | Med30 | mediator complex subunit 30 | 374.17 | 518.76 | -0.47 | 5.19E-02 |
| ENSMUSG00000024479 | GSMG0013184 | Mal2 | mal, T cell differentiation protein 2 | 894.06 | 683.46 | 0.39 | 1.04E-01 |
| ENSMUSG00000037362 | GSMG0013185 | Nov | nephroblastoma overexpressed gene | 927.26 | 863.16 | 0.10 | 7.14E-01 |
| ENSMUSG00000022419 | GSMG0013187 | Deptor | DEP domain containing MTOR-interacting protein | 141.21 | 74.50 | 0.93 | 2.18E-03 |
| ENSMUSG00000022369 | GSMG0013193 | Mtbp | Mdm2, transformed 3T3 cell double minute p53 binding protein | 305.33 | 512.41 | -0.74 | 6.85E-04 |

| | | | | | | | |
|---|---|---|---|---|---|---|---|
| NULL | GSMG0013199 | NULL | NULL | 494.19 | 506.79 | -0.03 | 9.26E-01 |
| ENSMUSG00000022359 | GSMG0013203 | Wdyhv1 | WDYHV motif containing 1 | 276.83 | 233.01 | 0.25 | 4.01E-01 |
| ENSMUSG00000037119 | GSMG0013210 | Fam91a1 | family with sequence similarity 91, member A1 | 1304.86 | 1300.27 | 0.01 | 9.87E-01 |
| ENSMUSG00000037085 | GSMG0013211 | Trmt12 | tRNA methyltranferase 12 | 531.54 | 534.30 | -0.01 | 9.79E-01 |
| ENSMUSG00000022354 | GSMG0013213 | Ndufb9 | NADH dehydrogenase (ubiquinone) 1 beta subcomplex, 9 | 3499.63 | 3475.57 | 0.01 | 9.70E-01 |
| ENSMUSG00000022351 | GSMG0013218 | Sqle | squalene epoxidase | 1568.62 | 4455.92 | -1.51 | 2.14E-22 |
| ENSMUSG00000059586 | GSMG0013219 | Nsmce2 | NSE2/MMS21 homolog, SMC5-SMC6 complex SUMO ligase | 667.08 | 813.34 | -0.29 | 1.68E-01 |
| ENSMUSG00000032501 | GSMG0013223 | Trib1 | tribbles pseudokinase 1 | 672.38 | 716.81 | -0.09 | 7.69E-01 |
| NULL | GSMG0013225 | NULL | NULL | 58.30 | 72.74 | -0.32 | 5.24E-01 |
| ENSMUSG00000022346 | GSMG0013230 | Myc | myelocytomatosis oncogene | 585.07 | 1082.86 | -0.89 | 3.24E-07 |
| ENSMUSG00000097039 | GSMG0013231 | Pvt1 | plasmacytoma variant translocation 1 | 52.64 | 103.95 | -0.97 | 6.28E-03 |
| ENSMUSG00000015002 | GSMG0013244 | Efr3a | EFR3 homolog A | 2153.49 | 1861.13 | 0.21 | 3.18E-01 |
| ENSMUSG00000072501 | GSMG0013247 | Phf20l1 | PHD finger protein 20-like 1 | 2770.64 | 2888.07 | -0.06 | 8.19E-01 |
| NULL | GSMG0013252 | NULL | NULL | 267.04 | 357.14 | -0.42 | 9.01E-02 |
| ENSMUSG00000022332 | GSMG0013257 | Khdrbs3 | KH domain containing, RNA binding, signal transduction associated 3 | 1125.63 | 737.60 | 0.61 | 3.35E-01 |
| NULL | GSMG0013258 | NULL | NULL | 697.70 | 488.75 | 0.51 | 2.86E-02 |
| NULL | GSMG0013262 | NULL | NULL | 188.68 | 223.78 | -0.25 | 5.12E-01 |
| ENSMUSG00000068391 | GSMG0013264 | Chrac1 | chromatin accessibility complex 1 | 941.06 | 1080.81 | -0.20 | 3.11E-01 |
| NULL | GSMG0013266 | NULL | NULL | 123.01 | 107.93 | 0.19 | 6.47E-01 |
| ENSMUSG00000036661 | GSMG0013267 | Dennd3 | DENN/MADD domain containing 3 | 2873.57 | 2844.84 | 0.01 | 9.67E-01 |
| ENSMUSG00000059895 | GSMG0013270 | Ptp4a3 | protein tyrosine phosphatase 4a3 | 3580.25 | 4964.84 | -0.47 | 8.39E-04 |
| ENSMUSG00000034730 | GSMG0013271 | Adgrb1 | adhesion G protein-coupled receptor B1 | 3198.93 | 1977.24 | 0.69 | 4.23E-03 |
| ENSMUSG00000022587 | GSMG0013278 | Ly6e | lymphocyte antigen 6 complex, locus E | 12832.09 | 4035.14 | 1.67 | 1.17E-23 |
| ENSMUSG00000022579 | GSMG0013282 | Gpihbp1 | GPI-anchored HDL-binding protein 1 | 25.08 | 117.48 | -2.21 | 4.94E-10 |
| ENSMUSG00000047003 | GSMG0013283 | Zfp41 | zinc finger protein 41 | 194.70 | 387.11 | -0.98 | 1.00E-05 |
| ENSMUSG00000022580 | GSMG0013286 | Rhpn1 | rhophilin, Rho GTPase binding protein 1 | 479.61 | 259.61 | 0.89 | 1.02E-04 |
| ENSMUSG00000022575 | GSMG0013288 | Gsdmd | gasdermin D | 546.61 | 413.11 | 0.40 | 6.77E-02 |
| ENSMUSG00000050846 | GSMG0013290 | Zfp623 | zinc finger protein 623 | 607.01 | 615.13 | -0.02 | 9.56E-01 |
| ENSMUSG00000034429 | GSMG0013291 | Zfp707 | zinc finger protein 707 | 643.01 | 599.69 | 0.10 | 7.37E-01 |
| ENSMUSG00000063704 | GSMG0013292 | Mapk15 | mitogen-activated protein kinase 15 | 218.38 | 1032.07 | -2.24 | 9.18E-23 |
| ENSMUSG00000022564 | GSMG0013296 | Grina | glutamate receptor, ionotropic, N-methyl D-aspartate-associated protein | 6007.34 | 4546.89 | 0.40 | 2.71E-02 |
| ENSMUSG00000071724 | GSMG0013298 | Smpd5 | sphingomyelin phosphodiesterase 5 | 209.78 | 253.74 | -0.27 | 3.30E-01 |
| ENSMUSG00000034259 | GSMG0013299 | Exosc4 | exosome component 4 | 1370.62 | 1664.63 | -0.28 | 1.39E-01 |

| | | | | | | | |
|---|---|---|---|---|---|---|---|
| ENSMUSG00000022561 | GSMG0013300 | Gpaa1 | GPI anchor attachment protein 1 | 2510.97 | 2712.99 | -0.11 | 6.55E-01 |
| ENSMUSG00000022551 | GSMG0013301 | Cyc1 | cytochrome c-1 | 3531.70 | 3704.73 | -0.07 | 7.80E-01 |
| ENSMUSG00000022553 | GSMG0013302 | Maf1 | MAF1 homolog, negative regulator of RNA polymerase III | 1439.98 | 1508.46 | -0.07 | 7.90E-01 |
| ENSMUSG00000022554 | GSMG0013303 | Hgh1 | HGH1 homolog | 595.12 | 756.93 | -0.35 | 7.13E-02 |
| ENSMUSG00000022558 | GSMG0013304 | Mroh1 | maestro heat-like repeat family member 1 | 2414.62 | 1675.93 | 0.53 | 7.99E-03 |
| ENSMUSG00000022556 | GSMG0013307 | Hsf1 | heat shock factor 1 | 1974.68 | 1791.22 | 0.14 | 5.14E-01 |
| ENSMUSG00000022560 | GSMG0013311 | Slc52a2 | solute carrier protein 52, member 2 | 1866.67 | 2314.94 | -0.31 | 8.35E-02 |
| ENSMUSG00000022550 | GSMG0013313 | Adck5 | aarF domain containing kinase 5 | 993.98 | 800.65 | 0.31 | 1.24E-01 |
| ENSMUSG00000004187 | GSMG0013324 | Kifc2 | kinesin family member C2 | 8246.37 | 4907.22 | 0.75 | 2.91E-05 |
| ENSMUSG00000033819 | GSMG0013325 | Ppp1r16a | protein phosphatase 1, regulatory (inhibitor) subunit 16A | 1034.85 | 887.54 | 0.22 | 3.63E-01 |
| ENSMUSG00000022546 | GSMG0013326 | Gpt | glutamic pyruvic transaminase, soluble | 186.85 | 212.19 | -0.18 | 6.05E-01 |
| ENSMUSG00000019080 | GSMG0013327 | Mfsd3 | major facilitator superfamily domain containing 3 | 387.79 | 396.29 | -0.03 | 9.46E-01 |
| ENSMUSG00000033728 | GSMG0013328 | Lrrc14 | leucine rich repeat containing 14 | 446.53 | 443.51 | 0.01 | 9.80E-01 |
| NULL | GSMG0013329 | NULL | NULL | 125.24 | 144.58 | -0.21 | 6.11E-01 |
| ENSMUSG00000033669 | GSMG0013331 | Zfp7 | zinc finger protein 7 | 354.19 | 275.17 | 0.36 | 1.15E-01 |
| ENSMUSG00000055041 | GSMG0013332 | Commd5 | COMM domain containing 5 | 408.50 | 469.28 | -0.20 | 4.51E-01 |
| ENSMUSG00000003970 | GSMG0013333 | Rpl8 | ribosomal protein L8 | 2885.21 | 3628.70 | -0.33 | 5.64E-02 |
| ENSMUSG00000063236 | GSMG0013334 | 1110038F14Rik | RIKEN cDNA 1110038F14 gene | 1016.48 | 978.26 | 0.05 | 8.57E-01 |
| NULL | GSMG0013346 | NULL | NULL | 261.19 | 151.98 | 0.78 | 1.12E-03 |
| ENSMUSG00000033287 | GSMG0013352 | Kctd17 | potassium channel tetramerisation domain containing 17 | 1483.71 | 1597.10 | -0.11 | 7.69E-01 |
| ENSMUSG00000018008 | GSMG0013354 | Cyth4 | cytohesin 4 | 519.20 | 369.32 | 0.49 | 2.38E-02 |
| ENSMUSG00000049521 | GSMG0013356 | Cdc42ep1 | CDC42 effector protein (Rho GTPase binding) 1 | 3698.94 | 3409.89 | 0.12 | 6.57E-01 |
| ENSMUSG00000033128 | GSMG0013357 | Gga1 | golgi associated, gamma adaptin ear containing, ARF binding protein 1 | 1409.63 | 1521.92 | -0.11 | 7.52E-01 |
| ENSMUSG00000022436 | GSMG0013358 | Sh3bp1 | SH3-domain binding protein 1 | 270.98 | 210.05 | 0.37 | 2.22E-01 |
| NULL | GSMG0013359 | Pdxp | pyridoxal (pyridoxine, vitamin B6) phosphatase | 1253.52 | 1275.73 | -0.02 | 9.42E-01 |
| ENSMUSG00000068220 | GSMG0013360 | Lgals1 | lectin, galactose binding, soluble 1 | 899.36 | 632.22 | 0.51 | 3.11E-02 |
| ENSMUSG00000033099 | GSMG0013361 | Nol12 | nucleolar protein 12 | 698.69 | 815.95 | -0.22 | 3.24E-01 |
| ENSMUSG00000033088 | GSMG0013362 | Triobp | TRIO and F-actin binding protein | 2913.04 | 2879.63 | 0.02 | 9.58E-01 |
| ENSMUSG00000096210 | GSMG0013363 | H1f0 | H1 histone family, member 0 | 9293.90 | 14169.23 | -0.61 | 3.73E-04 |
| ENSMUSG00000033047 | GSMG0013366 | Eif3l | eukaryotic translation initiation factor 3, subunit L | 2315.44 | 2988.94 | -0.37 | 4.10E-02 |
| ENSMUSG00000033020 | GSMG0013368 | Polr2f | polymerase (RNA) II (DNA directed) polypeptide F | 929.77 | 1280.53 | -0.46 | 3.93E-02 |
| ENSMUSG00000068206 | GSMG0013372 | Pick1 | protein interacting with C kinase 1 | 865.22 | 933.16 | -0.11 | 7.13E-01 |
| ENSMUSG00000042622 | GSMG0013375 | Maff | v-maf musculoaponeurotic fibrosarcoma oncogene family, protein F (av | 281.87 | 231.41 | 0.29 | 2.74E-01 |

| | | | | | | | |
|---|---|---|---|---|---|---|---|
| ENSMUSG00000010830 | GSMG0013379 | Kdelr3 | KDEL (Lys-Asp-Glu-Leu) endoplasmic reticulum protein retention rece | 261.23 | 162.90 | 0.68 | 7.12E-03 |
| NULL | GSMG0013380 | NULL | NULL | 602.72 | 600.42 | 0.01 | 9.85E-01 |
| ENSMUSG00000022428 | GSMG0013381 | Cby1 | chibby homolog 1 (Drosophila) | 1064.86 | 1066.86 | 0.00 | 9.91E-01 |
| ENSMUSG00000022427 | GSMG0013382 | Tomm22 | translocase of outer mitochondrial membrane 22 homolog (yeast) | 1059.66 | 1247.43 | -0.24 | 3.46E-01 |
| ENSMUSG00000042535 | GSMG0013383 | Gtpbp1 | GTP binding protein 1 | 854.40 | 1046.74 | -0.29 | 1.12E-01 |
| ENSMUSG00000009585 | GSMG0013386 | Apobec3 | apolipoprotein B mRNA editing enzyme, catalytic polypeptide 3 | 411.82 | 426.86 | -0.05 | 8.80E-01 |
| ENSMUSG00000022415 | GSMG0013388 | Syngr1 | synaptogyrin 1 | 2219.51 | 1680.05 | 0.40 | 9.02E-02 |
| ENSMUSG00000022414 | GSMG0013389 | Tab1 | TGF-beta activated kinase 1/MAP3K7 binding protein 1 | 238.97 | 245.28 | -0.03 | 9.41E-01 |
| ENSMUSG00000042428 | GSMG0013390 | Mgat3 | mannoside acetylglucosaminyltransferase 3 | 755.22 | 545.04 | 0.47 | 8.12E-02 |
| ENSMUSG00000022412 | GSMG0013391 | Mief1 | mitochondrial elongation factor 1 | 212.89 | 234.13 | -0.14 | 7.33E-01 |
| ENSMUSG00000042406 | GSMG0013392 | Atf4 | activating transcription factor 4 | 2794.01 | 2814.12 | -0.01 | 9.72E-01 |
| ENSMUSG00000047888 | GSMG0013399 | Tnrc6b | trinucleotide repeat containing 6b | 283.01 | 314.41 | -0.15 | 6.48E-01 |
| NULL | GSMG0013403 | NULL | NULL | 175.52 | 127.20 | 0.47 | 1.14E-01 |
| ENSMUSG00000022407 | GSMG0013407 | Adsl | adenylosuccinate lyase | 1062.35 | 1453.00 | -0.45 | 5.67E-03 |
| ENSMUSG00000042303 | GSMG0013409 | Sgsm3 | small G protein signaling modulator 3 | 610.23 | 686.45 | -0.17 | 4.84E-01 |
| ENSMUSG00000050164 | GSMG0013412 | Mchr1 | melanin-concentrating hormone receptor 1 | 228.35 | 124.17 | 0.88 | 5.09E-04 |
| ENSMUSG00000022401 | GSMG0013413 | Xpnpep3 | X-prolyl aminopeptidase 3, mitochondrial | 71.42 | 61.18 | 0.22 | 6.78E-01 |
| ENSMUSG00000022400 | GSMG0013414 | Rbx1 | ring-box 1 | 2892.11 | 3510.38 | -0.28 | 1.51E-01 |
| ENSMUSG00000055024 | GSMG0013415 | Ep300 | E1A binding protein p300 | 307.37 | 277.45 | 0.15 | 7.58E-01 |
| ENSMUSG00000022477 | GSMG0013416 | Aco2 | aconitase 2, mitochondrial | 7301.15 | 6911.55 | 0.08 | 7.55E-01 |
| ENSMUSG00000022394 | GSMG0013417 | L3mbtl2 | l(3)mbt-like 2 (Drosophila) | 398.03 | 511.44 | -0.36 | 1.11E-01 |
| NULL | GSMG0013418 | NULL | NULL | 171.38 | 153.76 | 0.15 | 7.37E-01 |
| ENSMUSG00000022390 | GSMG0013419 | Zc3h7b | zinc finger CCCH type containing 7B | 144.27 | 170.01 | -0.24 | 5.27E-01 |
| ENSMUSG00000022389 | GSMG0013420 | Tef | thyrotroph embryonic factor | 516.13 | 331.53 | 0.64 | 1.19E-03 |
| ENSMUSG00000042109 | GSMG0013425 | Csdc2 | cold shock domain containing C2, RNA binding | 2022.38 | 3342.44 | -0.72 | 8.84E-05 |
| ENSMUSG00000022471 | GSMG0013427 | Xrcc6 | X-ray repair complementing defective repair in Chinese hamster cells 6 | 592.13 | 1023.03 | -0.79 | 5.45E-07 |
| ENSMUSG00000068114 | GSMG0013431 | Ccdc134 | coiled-coil domain containing 134 | 439.05 | 406.50 | 0.11 | 7.20E-01 |
| ENSMUSG00000022463 | GSMG0013432 | Srebf2 | sterol regulatory element binding factor 2 | 725.84 | 1070.13 | -0.56 | 9.64E-04 |
| NULL | GSMG0013434 | NULL | NULL | 303.00 | 295.52 | 0.04 | 9.34E-01 |
| ENSMUSG00000068099 | GSMG0013435 | 1500009C09Rik | RIKEN cDNA 1500009C09 gene | 273.51 | 366.32 | -0.42 | 1.80E-01 |
| ENSMUSG00000022456 | GSMG0013436 | Sept3 | septin 3 | 1630.99 | 3301.64 | -1.02 | 1.01E-12 |
| ENSMUSG00000022452 | GSMG0013440 | Smdt1 | single-pass membrane protein with aspartate rich tail 1 | 2214.08 | 2572.21 | -0.22 | 2.80E-01 |
| NULL | GSMG0013453 | NULL | NULL | 62.60 | 53.63 | 0.23 | 6.86E-01 |

| | | | | | | | |
|---|---|---|---|---|---|---|---|
| ENSMUSG00000041736 | GSMG0013456 | Tspo | translocator protein | 1096.86 | 530.15 | 1.05 | 2.46E-11 |
| ENSMUSG00000041708 | GSMG0013458 | Mpped1 | metallophosphoesterase domain containing 1 | 321.17 | 1139.00 | -1.82 | 7.67E-26 |
| ENSMUSG00000041653 | GSMG0013462 | Pnpla3 | patatin-like phospholipase domain containing 3 | 76.97 | 112.65 | -0.55 | 1.09E-01 |
| ENSMUSG00000022437 | GSMG0013463 | Samm50 | SAMM50 sorting and assembly machinery component | 2434.19 | 2860.96 | -0.23 | 1.74E-01 |
| ENSMUSG00000022438 | GSMG0013464 | Parvb | parvin, beta | 474.53 | 397.14 | 0.25 | 3.56E-01 |
| ENSMUSG00000022439 | GSMG0013466 | Parvg | parvin, gamma | 71.29 | 44.38 | 0.70 | 1.62E-01 |
| ENSMUSG00000036106 | GSMG0013467 | Prr5 | proline rich 5 (renal) | 405.77 | 628.76 | -0.63 | 6.10E-04 |
| ENSMUSG00000016619 | GSMG0013470 | Nup50 | nucleoporin 50 | 527.23 | 749.77 | -0.51 | 5.34E-03 |
| ENSMUSG00000022434 | GSMG0013474 | Fam118a | family with sequence similarity 118, member A | 1278.31 | 1156.38 | 0.15 | 5.68E-01 |
| ENSMUSG00000006369 | GSMG0013476 | Fbln1 | fibulin 1 | 931.54 | 1703.04 | -0.87 | 2.07E-06 |
| ENSMUSG00000016541 | GSMG0013477 | Atxn10 | ataxin 10 | 2584.06 | 3846.64 | -0.57 | 4.48E-04 |
| NULL | GSMG0013480 | NULL | NULL | 171.79 | 153.38 | 0.17 | 6.68E-01 |
| NULL | GSMG0013483 | NULL | NULL | 583.02 | 417.89 | 0.48 | 6.23E-02 |
| ENSMUSG00000022383 | GSMG0013486 | Ppara | peroxisome proliferator activated receptor alpha | 118.19 | 62.69 | 0.92 | 4.35E-03 |
| ENSMUSG00000035944 | GSMG0013491 | Ttc38 | tetratricopeptide repeat domain 38 | 128.36 | 103.57 | 0.31 | 3.97E-01 |
| ENSMUSG00000022386 | GSMG0013493 | Trmu | tRNA 5-methylaminomethyl-2-thiouridylate methyltransferase | 833.37 | 732.14 | 0.19 | 3.96E-01 |
| ENSMUSG00000035900 | GSMG0013495 | Gramd4 | GRAM domain containing 4 | 810.21 | 1043.50 | -0.37 | 1.28E-01 |
| NULL | GSMG0013496 | NULL | NULL | 277.37 | 304.46 | -0.14 | 7.56E-01 |
| NULL | GSMG0013498 | NULL | NULL | 56.56 | 46.92 | 0.26 | 6.71E-01 |
| NULL | GSMG0013499 | NULL | NULL | 95.38 | 81.65 | 0.23 | 6.88E-01 |
| ENSMUSG00000051864 | GSMG0013500 | Tbc1d22a | TBC1 domain family, member 22a | 344.95 | 387.89 | -0.17 | 5.52E-01 |
| ENSMUSG00000054863 | GSMG0013503 | Fam19a5 | family with sequence similarity 19, member A5 | 438.24 | 376.08 | 0.22 | 4.63E-01 |
| NULL | GSMG0013504 | NULL | NULL | 152.02 | 99.28 | 0.62 | 7.49E-02 |
| ENSMUSG00000023272 | GSMG0013508 | Creld2 | cysteine-rich with EGF-like domains 2 | 1333.59 | 1578.78 | -0.24 | 2.66E-01 |
| ENSMUSG00000035828 | GSMG0013509 | Pim3 | proviral integration site 3 | 1155.40 | 1387.81 | -0.26 | 2.10E-01 |
| ENSMUSG00000058441 | GSMG0013511 | Panx2 | pannexin 2 | 243.45 | 269.48 | -0.15 | 7.51E-01 |
| ENSMUSG00000015363 | GSMG0013512 | Trabd | TraB domain containing | 886.13 | 972.37 | -0.13 | 5.91E-01 |
| NULL | GSMG0013513 | NULL | NULL | 222.61 | 247.96 | -0.16 | 7.09E-01 |
| ENSMUSG00000035757 | GSMG0013514 | Selo | selenoprotein O | 584.60 | 537.25 | 0.12 | 6.80E-01 |
| ENSMUSG00000036561 | GSMG0013517 | Ppp6r2 | protein phosphatase 6, regulatory subunit 2 | 464.48 | 471.53 | -0.02 | 9.54E-01 |
| ENSMUSG00000008690 | GSMG0013522 | Ncaph2 | non-SMC condensin II complex, subunit H2 | 2434.39 | 2554.94 | -0.07 | 8.06E-01 |
| ENSMUSG00000022619 | GSMG0013525 | Mapk8ip2 | mitogen-activated protein kinase 8 interacting protein 2 | 1322.66 | 1381.13 | -0.06 | 8.52E-01 |
| ENSMUSG00000022623 | GSMG0013527 | Shank3 | SH3/ankyrin domain gene 3 | 1218.44 | 1039.80 | 0.23 | 5.17E-01 |

| | | | | | | | |
|---|---|---|---|---|---|---|---|
| ENSMUSG00000075470 | GSMG0013530 | Alg10b | asparagine-linked glycosylation 10B (alpha-1,2-glucosyltransferase) | 322.01 | 433.68 | -0.43 | 6.45E-02 |
| ENSMUSG00000036273 | GSMG0013534 | Lrrk2 | leucine-rich repeat kinase 2 | 862.35 | 679.64 | 0.34 | 7.45E-02 |
| NULL | GSMG0013537 | NULL | NULL | 342.17 | 295.71 | 0.21 | 4.91E-01 |
| ENSMUSG00000055022 | GSMG0013539 | Cntn1 | contactin 1 | 1152.60 | 2338.66 | -1.02 | 1.29E-07 |
| ENSMUSG00000036218 | GSMG0013544 | Pdzrn4 | PDZ domain containing RING finger 4 | 304.92 | 169.91 | 0.84 | 6.21E-03 |
| ENSMUSG00000036167 | GSMG0013551 | Pphln1 | periphilin 1 | 355.41 | 498.25 | -0.48 | 5.72E-02 |
| ENSMUSG00000064210 | GSMG0013560 | Ano6 | anoctamin 6 | 3127.66 | 2799.89 | 0.16 | 4.18E-01 |
| NULL | GSMG0013570 | NULL | NULL | 94.59 | 117.63 | -0.32 | 4.48E-01 |
| NULL | GSMG0013575 | NULL | NULL | 121.47 | 75.46 | 0.68 | 5.46E-02 |
| ENSMUSG00000081534 | GSMG0013576 | Slc48a1 | solute carrier family 48 (heme transporter), member 1 | 4045.92 | 2859.58 | 0.50 | 7.87E-04 |
| ENSMUSG00000052369 | GSMG0013578 | Tmem106c | transmembrane protein 106C | 489.78 | 622.43 | -0.34 | 1.69E-01 |
| ENSMUSG00000033065 | GSMG0013580 | Pfkm | phosphofructokinase, muscle | 3390.88 | 2531.34 | 0.42 | 2.26E-02 |
| ENSMUSG00000029875 | GSMG0013582 | Ccdc184 | coiled-coil domain containing 184 | 265.72 | 459.52 | -0.80 | 1.34E-03 |
| NULL | GSMG0013587 | NULL | NULL | 81.88 | 101.26 | -0.31 | 5.09E-01 |
| ENSMUSG00000003352 | GSMG0013591 | Cacnb3 | calcium channel, voltage-dependent, beta 3 subunit | 946.39 | 2011.78 | -1.09 | 3.53E-07 |
| NULL | GSMG0013593 | NULL | NULL | 432.02 | 341.39 | 0.34 | 3.54E-01 |
| NULL | GSMG0013594 | NULL | NULL | 192.19 | 148.82 | 0.37 | 2.80E-01 |
| NULL | GSMG0013597 | NULL | NULL | 232.51 | 192.59 | 0.27 | 3.56E-01 |
| NULL | GSMG0013604 | NULL | NULL | 96.91 | 400.64 | -2.06 | 9.62E-18 |
| ENSMUSG00000023484 | GSMG0013609 | Prph | peripherin | 99.04 | 126.46 | -0.35 | 3.30E-01 |
| ENSMUSG00000032783 | GSMG0013610 | Troap | trophinin associated protein | 82.78 | 236.05 | -1.50 | 6.18E-07 |
| ENSMUSG00000051934 | GSMG0013612 | Spats2 | spermatogenesis associated, serine-rich 2 | 499.40 | 572.74 | -0.20 | 4.23E-01 |
| ENSMUSG00000037579 | GSMG0013613 | Kcnh3 | potassium voltage-gated channel, subfamily H (eag-related), member 3 | 1530.92 | 815.31 | 0.91 | 5.19E-04 |
| ENSMUSG00000023007 | GSMG0013616 | Prpf40b | pre-mRNA processing factor 40B | 3537.24 | 3388.14 | 0.06 | 8.41E-01 |
| ENSMUSG00000023010 | GSMG0013619 | Tmbim6 | transmembrane BAX inhibitor motif containing 6 | 2458.23 | 2695.36 | -0.13 | 6.95E-01 |
| ENSMUSG00000023017 | GSMG0013624 | Asic1 | acid-sensing (proton-gated) ion channel 1 | 164.11 | 221.21 | -0.43 | 1.98E-01 |
| ENSMUSG00000023018 | GSMG0013625 | Smarcd1 | SWI/SNF related, matrix associated, actin dependent regulator of chron | 983.12 | 1256.92 | -0.36 | 9.88E-02 |
| ENSMUSG00000023020 | GSMG0013627 | Cox14 | cytochrome c oxidase assembly protein 14 | 1541.15 | 1496.54 | 0.04 | 8.92E-01 |
| NULL | GSMG0013628 | NULL | NULL | 170.10 | 72.18 | 1.24 | 5.44E-05 |
| ENSMUSG00000023025 | GSMG0013632 | Larp4 | La ribonucleoprotein domain family, member 4 | 733.87 | 878.50 | -0.26 | 2.07E-01 |
| ENSMUSG00000023026 | GSMG0013634 | Dip2b | disco interacting protein 2 homolog B | 826.92 | 744.90 | 0.15 | 6.64E-01 |
| ENSMUSG00000023027 | GSMG0013639 | Atf1 | activating transcription factor 1 | 181.53 | 122.85 | 0.56 | 3.48E-02 |
| ENSMUSG00000054619 | GSMG0013642 | Mettl7a1 | methyltransferase like 7A1 | 290.51 | 103.64 | 1.49 | 3.32E-11 |

| | | | | | | | |
|---|---|---|---|---|---|---|---|
| ENSMUSG00000037353 | GSMG0013646 | Letmd1 | LETM1 domain containing 1 | 469.05 | 503.70 | -0.10 | 7.09E-01 |
| ENSMUSG00000000346 | GSMG0013649 | Dazap2 | DAZ associated protein 2 | 6526.17 | 5983.21 | 0.13 | 5.80E-01 |
| ENSMUSG00000023032 | GSMG0013651 | Slc4a8 | solute carrier family 4 (anion exchanger), member 8 | 372.96 | 509.27 | -0.45 | 5.46E-02 |
| ENSMUSG00000023033 | GSMG0013654 | Scn8a | sodium channel, voltage-gated, type VIII, alpha | 1244.85 | 1089.52 | 0.19 | 3.79E-01 |
| NULL | GSMG0013657 | NULL | NULL | 380.68 | 257.39 | 0.57 | 3.73E-02 |
| ENSMUSG00000000530 | GSMG0013660 | Acvrl1 | activin A receptor, type II-like 1 | 5108.72 | 2837.81 | 0.85 | 3.08E-08 |
| ENSMUSG00000000532 | GSMG0013661 | Acvr1b | activin A receptor, type 1B | 97.02 | 181.26 | -0.89 | 9.71E-04 |
| ENSMUSG00000000531 | GSMG0013662 | Grasp | GRP1 (general receptor for phosphoinositides 1)-associated scaffold pr | 1532.37 | 892.62 | 0.78 | 6.26E-07 |
| ENSMUSG00000023034 | GSMG0013663 | Nr4a1 | nuclear receptor subfamily 4, group A, member 1 | 1983.46 | 399.54 | 2.31 | 2.12E-28 |
| ENSMUSG00000037204 | GSMG0013664 | Atg101 | autophagy related 101 | 1327.25 | 1159.49 | 0.19 | 3.12E-01 |
| ENSMUSG00000058655 | GSMG0013678 | Eif4b | eukaryotic translation initiation factor 4B | 2274.02 | 2671.93 | -0.23 | 1.94E-01 |
| ENSMUSG00000037003 | GSMG0013681 | Tns2 | tensin 2 | 5033.18 | 3982.30 | 0.34 | 6.00E-02 |
| ENSMUSG00000023046 | GSMG0013682 | Igfbp6 | insulin-like growth factor binding protein 6 | 204.40 | 135.78 | 0.59 | 2.77E-02 |
| ENSMUSG00000046897 | GSMG0013684 | Zfp740 | zinc finger protein 740 | 447.27 | 453.34 | -0.02 | 9.55E-01 |
| ENSMUSG00000058290 | GSMG0013688 | Espl1 | extra spindle pole bodies 1, separase | 278.59 | 731.98 | -1.39 | 2.67E-15 |
| ENSMUSG00000001289 | GSMG0013689 | Pfdn5 | prefoldin 5 | 1194.12 | 1434.45 | -0.27 | 2.42E-01 |
| ENSMUSG00000001285 | GSMG0013690 | Myg1 | melanocyte proliferating gene 1 | 445.17 | 495.00 | -0.15 | 5.71E-01 |
| ENSMUSG00000001280 | GSMG0013692 | Sp1 | trans-acting transcription factor 1 | 1129.21 | 1307.31 | -0.21 | 2.80E-01 |
| ENSMUSG00000023048 | GSMG0013695 | Prr13 | proline rich 13 | 271.68 | 252.33 | 0.11 | 7.94E-01 |
| ENSMUSG00000056851 | GSMG0013696 | Pcbp2 | poly(rC) binding protein 2 | 1172.74 | 1278.91 | -0.12 | 6.26E-01 |
| ENSMUSG00000023051 | GSMG0013697 | Tarbp2 | TAR (HIV) RNA binding protein 2 | 690.53 | 734.02 | -0.09 | 7.78E-01 |
| ENSMUSG00000060992 | GSMG0013713 | Copz1 | coatomer protein complex, subunit zeta 1 | 3875.00 | 4346.28 | -0.17 | 4.65E-01 |
| ENSMUSG00000022489 | GSMG0013718 | Pde1b | phosphodiesterase 1B, Ca2+-calmodulin dependent | 718.85 | 641.83 | 0.16 | 6.50E-01 |
| ENSMUSG00000055737 | GSMG0013719 | Ghr | growth hormone receptor | 438.91 | 317.28 | 0.47 | 1.98E-02 |
| ENSMUSG00000022184 | GSMG0013725 | Fbxo4 | F-box protein 4 | 428.23 | 332.20 | 0.37 | 9.95E-02 |
| ENSMUSG00000041935 | GSMG0013726 | AW549877 | expressed sequence AW549877 | 2412.07 | 2237.27 | 0.11 | 6.36E-01 |
| ENSMUSG00000041849 | GSMG0013735 | Card6 | caspase recruitment domain family, member 6 | 85.66 | 77.23 | 0.15 | 7.87E-01 |
| NULL | GSMG0013742 | NULL | NULL | 227.24 | 219.76 | 0.05 | 9.02E-01 |
| ENSMUSG00000022146 | GSMG0013744 | Osmr | oncostatin M receptor | 845.02 | 869.65 | -0.04 | 8.90E-01 |
| ENSMUSG00000042961 | GSMG0013747 | Egflam | EGF-like, fibronectin type III and laminin G domains | 2780.17 | 2556.59 | 0.12 | 5.95E-01 |
| ENSMUSG00000039828 | GSMG0013750 | Wdr70 | WD repeat domain 70 | 769.28 | 990.61 | -0.36 | 8.15E-02 |
| ENSMUSG00000022141 | GSMG0013751 | Nipbl | Nipped-B homolog (Drosophila) | 578.00 | 626.59 | -0.11 | 7.05E-01 |
| NULL | GSMG0013753 | NULL | NULL | 157.95 | 113.53 | 0.47 | 1.91E-01 |

| | | | | | | | |
|---|---|---|---|---|---|---|---|
| NULL | GSMG0013757 | NULL | NULL | 317.59 | 301.92 | 0.08 | 8.52E-01 |
| ENSMUSG00000005360 | GSMG0013758 | Slc1a3 | solute carrier family 1 (glial high affinity glutamate transporter), membe | 5146.15 | 5616.58 | -0.13 | 5.44E-01 |
| ENSMUSG00000054115 | GSMG0013761 | Skp2 | S-phase kinase-associated protein 2 (p45) | 139.89 | 297.73 | -1.08 | 8.91E-06 |
| ENSMUSG00000022247 | GSMG0013768 | Brix1 | BRX1, biogenesis of ribosomes | 579.55 | 796.97 | -0.46 | 6.20E-02 |
| ENSMUSG00000022246 | GSMG0013770 | Rai14 | retinoic acid induced 14 | 823.31 | 1068.25 | -0.38 | 3.98E-02 |
| NULL | GSMG0013771 | NULL | NULL | 243.03 | 225.40 | 0.11 | 8.13E-01 |
| ENSMUSG00000022241 | GSMG0013775 | Tars | threonyl-tRNA synthetase | 1932.15 | 2258.20 | -0.22 | 2.74E-01 |
| ENSMUSG00000022206 | GSMG0013776 | Npr3 | natriuretic peptide receptor 3 | 74.24 | 95.04 | -0.36 | 3.87E-01 |
| ENSMUSG00000022205 | GSMG0013778 | Sub1 | SUB1 homolog (S. cerevisiae) | 3592.48 | 5029.93 | -0.49 | 6.60E-02 |
| ENSMUSG00000022197 | GSMG0013782 | Pdzd2 | PDZ domain containing 2 | 4585.56 | 5668.61 | -0.31 | 1.24E-01 |
| NULL | GSMG0013784 | NULL | NULL | 122.69 | 211.34 | -0.79 | 1.89E-02 |
| ENSMUSG00000022195 | GSMG0013787 | 6030458C11Rik | RIKEN cDNA 6030458C11 gene | 407.81 | 425.13 | -0.06 | 8.67E-01 |
| ENSMUSG00000039385 | GSMG0013789 | Cdh6 | cadherin 6 | 198.97 | 278.58 | -0.48 | 5.41E-02 |
| ENSMUSG00000047565 | GSMG0013799 | Acot10 | acyl-CoA thioesterase 10 | 380.59 | 275.77 | 0.47 | 4.17E-02 |
| ENSMUSG00000045763 | GSMG0013812 | Basp1 | brain abundant, membrane attached signal protein 1 | 4137.99 | 7532.24 | -0.86 | 1.75E-04 |
| NULL | GSMG0013813 | NULL | NULL | 221.33 | 224.21 | -0.01 | 9.78E-01 |
| NULL | GSMG0013814 | NULL | NULL | 36.27 | 68.89 | -0.95 | 8.35E-02 |
| ENSMUSG00000043556 | GSMG0013816 | Fbxl7 | F-box and leucine-rich repeat protein 7 | 124.18 | 111.33 | 0.16 | 7.13E-01 |
| ENSMUSG00000046034 | GSMG0013828 | Otulin | OTU deubiquitinase with linear linkage specificity | 446.35 | 424.76 | 0.07 | 8.37E-01 |
| ENSMUSG00000056069 | GSMG0013829 | Fam105a | family with sequence similarity 105, member A | 266.51 | 209.34 | 0.35 | 1.84E-01 |
| ENSMUSG00000022263 | GSMG0013831 | Trio | triple functional domain (PTPRF interacting) | 437.19 | 464.86 | -0.09 | 7.94E-01 |
| NULL | GSMG0013832 | NULL | NULL | 262.44 | 208.08 | 0.34 | 3.10E-01 |
| NULL | GSMG0013835 | NULL | NULL | 664.77 | 665.29 | 0.00 | 9.98E-01 |
| NULL | GSMG0013839 | NULL | NULL | 267.11 | 237.87 | 0.17 | 6.23E-01 |
| NULL | GSMG0013840 | NULL | NULL | 198.07 | 184.02 | 0.11 | 8.04E-01 |
| ENSMUSG00000022237 | GSMG0013842 | Ankrd33b | ankyrin repeat domain 33B | 239.59 | 49.12 | 2.28 | 6.53E-15 |
| ENSMUSG00000039100 | GSMG0013844 | March6 | membrane-associated ring finger (C3HC4) 6 | 479.87 | 502.76 | -0.06 | 8.52E-01 |
| NULL | GSMG0013845 | NULL | NULL | 454.37 | 457.10 | -0.01 | 9.86E-01 |
| ENSMUSG00000022234 | GSMG0013846 | Cct5 | chaperonin containing Tcp1, subunit 5 (epsilon) | 4916.50 | 6909.96 | -0.49 | 5.33E-03 |
| ENSMUSG00000096956 | GSMG0013847 | Snhg18 | small nucleolar RNA host gene 18 | 121.26 | 63.88 | 0.92 | 3.94E-03 |
| ENSMUSG00000038984 | GSMG0013852 | Tspyl5 | testis-specific protein, Y-encoded-like 5 | 494.57 | 614.42 | -0.31 | 1.61E-01 |
| ENSMUSG00000022323 | GSMG0013857 | Rida | reactive intermediate imine deaminase A homolog | 354.10 | 259.13 | 0.45 | 1.09E-01 |
| ENSMUSG00000038879 | GSMG0013858 | Nipal2 | NIPA-like domain containing 2 | 93.99 | 55.78 | 0.75 | 4.64E-02 |

| | | | | | | | |
|---|---|---|---|---|---|---|---|
| ENSMUSG00000022329 | GSMG0013863 | Stk3 | serine/threonine kinase 3 | 795.56 | 755.26 | 0.08 | 7.91E-01 |
| NULL | GSMG0013865 | NULL | NULL | 157.07 | 141.90 | 0.15 | 7.59E-01 |
| ENSMUSG00000014313 | GSMG0013871 | Cox6c | cytochrome c oxidase subunit VIc | 4132.64 | 4250.17 | -0.04 | 8.97E-01 |
| ENSMUSG00000022280 | GSMG0013874 | Rnf19a | ring finger protein 19A | 984.44 | 962.35 | 0.03 | 9.18E-01 |
| NULL | GSMG0013875 | NULL | NULL | 357.32 | 258.04 | 0.47 | 8.19E-02 |
| ENSMUSG00000048307 | GSMG0013877 | Ankrd46 | ankyrin repeat domain 46 | 931.09 | 967.88 | -0.06 | 8.46E-01 |
| ENSMUSG00000022283 | GSMG0013880 | Pabpc1 | poly(A) binding protein, cytoplasmic 1 | 2228.15 | 3076.32 | -0.47 | 2.89E-03 |
| ENSMUSG00000062397 | GSMG0013884 | Zfp706 | zinc finger protein 706 | 4519.27 | 5526.55 | -0.29 | 1.50E-01 |
| NULL | GSMG0013889 | NULL | NULL | 518.54 | 286.27 | 0.86 | 9.85E-04 |
| ENSMUSG00000022292 | GSMG0013893 | Rrm2b | ribonucleotide reductase M2 B (TP53 inducible) | 191.99 | 148.54 | 0.37 | 2.96E-01 |
| ENSMUSG00000037487 | GSMG0013894 | Ubr5 | ubiquitin protein ligase E3 component n-recognin 5 | 3593.13 | 3329.25 | 0.11 | 7.62E-01 |
| ENSMUSG00000037465 | GSMG0013897 | Klf10 | Kruppel-like factor 10 | 1488.31 | 1457.11 | 0.03 | 9.19E-01 |
| ENSMUSG00000037458 | GSMG0013898 | Azin1 | antizyme inhibitor 1 | 1385.34 | 1468.51 | -0.08 | 7.56E-01 |
| ENSMUSG00000022299 | GSMG0013904 | Slc25a32 | solute carrier family 25, member 32 | 270.48 | 170.90 | 0.66 | 9.44E-03 |
| ENSMUSG00000022305 | GSMG0013908 | Lrp12 | low density lipoprotein-related protein 12 | 266.69 | 370.55 | -0.48 | 7.13E-02 |
| ENSMUSG00000022309 | GSMG0013916 | Angpt1 | angiopoietin 1 | 109.44 | 99.11 | 0.15 | 7.73E-01 |
| ENSMUSG00000022336 | GSMG0013924 | Eif3e | eukaryotic translation initiation factor 3, subunit E | 1286.92 | 1728.82 | -0.43 | 5.11E-02 |
| ENSMUSG00000038736 | GSMG0013929 | Nudcd1 | NudC domain containing 1 | 154.06 | 189.25 | -0.29 | 4.16E-01 |
| NULL | GSMG0013930 | NULL | NULL | 671.21 | 1830.03 | -1.45 | 8.49E-25 |
| NULL | GSMG0013931 | NULL | NULL | 848.43 | 1880.08 | -1.15 | 7.34E-13 |
| ENSMUSG00000022340 | GSMG0013933 | Sybu | syntabulin (syntaxin-interacting) | 326.38 | 918.64 | -1.49 | 3.96E-21 |
| ENSMUSG00000022342 | GSMG0013934 | Kcnv1 | potassium channel, subfamily V, member 1 | 152.07 | 117.49 | 0.37 | 3.09E-01 |
| ENSMUSG00000038679 | GSMG0013946 | Trps1 | trichorhinophalangeal syndrome I (human) | 733.14 | 604.26 | 0.28 | 2.48E-01 |
| ENSMUSG00000022312 | GSMG0013957 | Eif3h | eukaryotic translation initiation factor 3, subunit H | 1437.04 | 1793.00 | -0.32 | 9.27E-02 |
| ENSMUSG00000022314 | GSMG0013959 | Rad21 | RAD21 cohesin complex component | 902.91 | 1052.12 | -0.22 | 3.73E-01 |
| ENSMUSG00000061731 | GSMG0013962 | Ext1 | exostoses (multiple) 1 | 468.26 | 515.19 | -0.14 | 7.20E-01 |
| ENSMUSG00000063727 | GSMG0013975 | Tnfrsf11b | tumor necrosis factor receptor superfamily, member 11b (osteoprotegeri | 341.85 | 956.92 | -1.49 | 2.18E-18 |
| ENSMUSG00000022425 | GSMG0013976 | Enpp2 | ectonucleotide pyrophosphatase/phosphodiesterase 2 | 4644.82 | 7574.73 | -0.71 | 1.33E-04 |
| ENSMUSG00000037343 | GSMG0013978 | Taf2 | TATA-box binding protein associated factor 2 | 1603.73 | 1854.66 | -0.21 | 3.09E-01 |
| ENSMUSG00000022370 | GSMG0013984 | Mrpl13 | mitochondrial ribosomal protein L13 | 727.00 | 989.91 | -0.45 | 2.53E-02 |
| ENSMUSG00000060429 | GSMG0013986 | Sntb1 | syntrophin, basic 1 | 267.75 | 125.01 | 1.10 | 2.55E-05 |
| ENSMUSG00000022365 | GSMG0013996 | Derl1 | Der1-like domain family, member 1 | 2140.50 | 2147.75 | 0.00 | 9.88E-01 |
| G00000022361,ENSMUSG000 | GSMG0013997 | 9130401M01Rik // | RIKEN cDNA 9130401M01 gene // zinc fingers and homeoboxes 1 | 4207.53 | 4281.19 | -0.02 | 9.27E-01 |

| | | | | | | | |
|---|---|---|---|---|---|---|---|
| ENSMUSG00000022360 | GSMG0013999 | Atad2 | ATPase family, AAA domain containing 2 | 80.19 | 133.89 | -0.73 | 5.69E-02 |
| ENSMUSG00000022358 | GSMG0014001 | Fbxo32 | F-box protein 32 | 131.63 | 57.83 | 1.18 | 2.55E-04 |
| ENSMUSG00000062373 | GSMG0014008 | Tmem65 | transmembrane protein 65 | 1806.39 | 1974.96 | -0.13 | 6.06E-01 |
| ENSMUSG00000050891 | GSMG0014011 | Tatdn1 | TatD DNase domain containing 1 | 192.80 | 213.33 | -0.15 | 7.03E-01 |
| ENSMUSG00000022353 | GSMG0014013 | Mtss1 | metastasis suppressor 1 | 603.23 | 730.97 | -0.28 | 2.46E-01 |
| ENSMUSG00000022350 | GSMG0014017 | E430025E21Rik | RIKEN cDNA E430025E21 gene | 761.13 | 963.99 | -0.34 | 5.32E-02 |
| ENSMUSG00000072568 | GSMG0014025 | Fam84b | family with sequence similarity 84, member B | 1217.64 | 1523.68 | -0.32 | 1.05E-01 |
| ENSMUSG00000022378 | GSMG0014039 | Fam49b | family with sequence similarity 49, member B | 829.05 | 1213.88 | -0.55 | 7.60E-04 |
| ENSMUSG00000022377 | GSMG0014045 | Asap1 | ArfGAP with SH3 domain, ankyrin repeat and PH domain1 | 2378.19 | 2301.80 | 0.05 | 8.51E-01 |
| NULL | GSMG0014046 | NULL | NULL | 252.88 | 227.02 | 0.15 | 6.90E-01 |
| NULL | GSMG0014051 | NULL | NULL | 572.80 | 365.15 | 0.65 | 6.15E-04 |
| NULL | GSMG0014055 | NULL | NULL | 193.95 | 144.06 | 0.43 | 1.35E-01 |
| ENSMUSG00000022376 | GSMG0014056 | Adcy8 | adenylate cyclase 8 | 876.33 | 721.53 | 0.28 | 2.86E-01 |
| NULL | GSMG0014059 | NULL | NULL | 627.91 | 573.87 | 0.13 | 6.75E-01 |
| NULL | GSMG0014061 | NULL | NULL | 534.82 | 336.08 | 0.67 | 1.18E-03 |
| NULL | GSMG0014062 | NULL | NULL | 1016.86 | 707.68 | 0.52 | 1.79E-03 |
| NULL | GSMG0014063 | NULL | NULL | 491.73 | 359.05 | 0.45 | 4.12E-02 |
| NULL | GSMG0014064 | NULL | NULL | 330.49 | 214.71 | 0.62 | 3.57E-03 |
| ENSMUSG00000036944 | GSMG0014067 | Tmem71 | transmembrane protein 71 | 137.58 | 94.77 | 0.53 | 1.10E-01 |
| ENSMUSG00000022372 | GSMG0014069 | Sla | src-like adaptor | 37.22 | 232.28 | -2.66 | 8.39E-23 |
| ENSMUSG00000005125 | GSMG0014070 | Ndrg1 | N-myc downstream regulated gene 1 | 676.41 | 334.22 | 1.02 | 1.37E-06 |
| ENSMUSG00000013846 | GSMG0014071 | St3gal1 | ST3 beta-galactoside alpha-2,3-sialyltransferase 1 | 260.05 | 224.60 | 0.21 | 4.79E-01 |
| NULL | GSMG0014086 | NULL | NULL | 294.87 | 97.59 | 1.60 | 6.51E-12 |
| ENSMUSG00000036760 | GSMG0014088 | Kcnk9 | potassium channel, subfamily K, member 9 | 181.67 | 130.45 | 0.48 | 2.09E-01 |
| ENSMUSG00000047921 | GSMG0014089 | Trappc9 | trafficking protein particle complex 9 | 892.65 | 926.36 | -0.05 | 8.73E-01 |
| ENSMUSG00000106847 | GSMG0014092 | Peg13 | paternally expressed 13 | 1970.75 | 2078.22 | -0.08 | 8.46E-01 |
| NULL | GSMG0014095 | NULL | NULL | 116.42 | 93.77 | 0.31 | 4.46E-01 |
| NULL | GSMG0014096 | NULL | NULL | 258.30 | 252.13 | 0.03 | 9.34E-01 |
| ENSMUSG00000036698 | GSMG0014098 | Ago2 | argonaute RISC catalytic subunit 2 | 196.86 | 195.34 | 0.02 | 9.72E-01 |
| ENSMUSG00000022607 | GSMG0014099 | Ptk2 | PTK2 protein tyrosine kinase 2 | 3918.71 | 4597.45 | -0.23 | 2.05E-01 |
| NULL | GSMG0014101 | NULL | NULL | 403.14 | 261.51 | 0.63 | 9.96E-03 |
| NULL | GSMG0014102 | NULL | NULL | 268.46 | 246.73 | 0.13 | 7.58E-01 |
| ENSMUSG00000079020 | GSMG0014106 | Slc45a4 | solute carrier family 45, member 4 | 494.68 | 327.29 | 0.60 | 5.10E-02 |

| | | | | | | | |
|---|---|---|---|---|---|---|---|
| ENSMUSG00000022602 | GSMG0014115 | Arc | activity regulated cytoskeletal-associated protein | 834.77 | 34.65 | 4.58 | 2.38E-59 |
| ENSMUSG00000022595 | GSMG0014118 | Lypd2 | Ly6/Plaur domain containing 2 | 66.32 | 65.01 | 0.03 | 9.65E-01 |
| ENSMUSG00000022594 | GSMG0014120 | Lynx1 | Ly6/neurotoxin 1 | 1567.60 | 291.37 | 2.43 | 3.93E-40 |
| ENSMUSG00000079018 | GSMG0014131 | Ly6c1 | lymphocyte antigen 6 complex, locus C1 | 15098.31 | 4165.63 | 1.86 | 3.95E-40 |
| ENSMUSG00000022577 | GSMG0014134 | Ly6h | lymphocyte antigen 6 complex, locus H | 1432.58 | 2300.26 | -0.68 | 1.16E-03 |
| ENSMUSG00000000934 | GSMG0014139 | Top1mt | DNA topoisomerase 1, mitochondrial | 364.97 | 529.44 | -0.54 | 1.02E-02 |
| ENSMUSG00000022574 | GSMG0014143 | Naprt | nicotinate phosphoribosyltransferase | 296.85 | 183.51 | 0.70 | 1.69E-03 |
| ENSMUSG00000055762 | GSMG0014144 | Eef1d | eukaryotic translation elongation factor 1 delta (guanine nucleotide excl | 1314.33 | 1811.02 | -0.46 | 5.81E-03 |
| ENSMUSG00000022571 | GSMG0014146 | Pycrl | pyrroline-5-carboxylate reductase-like | 248.59 | 288.76 | -0.22 | 4.46E-01 |
| ENSMUSG00000022570 | GSMG0014147 | Tsta3 | tissue specific transplantation antigen P35B | 1242.82 | 1473.04 | -0.25 | 1.82E-01 |
| ENSMUSG00000022568 | GSMG0014152 | Scrib | scribbled planar cell polarity | 765.59 | 1036.49 | -0.44 | 1.07E-01 |
| ENSMUSG00000002524 | GSMG0014153 | Puf60 | poly-U binding splicing factor 60 | 2943.19 | 3120.99 | -0.08 | 7.37E-01 |
| ENSMUSG00000075590 | GSMG0014155 | Nrbp2 | nuclear receptor binding protein 2 | 4698.86 | 6543.57 | -0.48 | 5.18E-02 |
| G00000044361,ENSMUSG000 | GSMG0014156 | Eppk1 // BC02413 | epiplakin 1 // cDNA sequence BC024139 | 99.09 | 356.88 | -1.84 | 8.15E-14 |
| G00000022565,ENSMUSG000 | GSMG0014157 | Plec // Parp10 | plectin // poly (ADP-ribose) polymerase family, member 10 | 710.26 | 491.83 | 0.53 | 1.21E-01 |
| ENSMUSG00000022562 | GSMG0014160 | Oplah | 5-oxoprolinase (ATP-hydrolysing) | 552.33 | 424.80 | 0.38 | 1.54E-01 |
| ENSMUSG00000022552 | GSMG0014162 | Sharpin | SHANK-associated RH domain interacting protein | 1418.88 | 1459.58 | -0.04 | 8.81E-01 |
| ENSMUSG00000022555 | GSMG0014167 | Dgat1 | diacylglycerol O-acyltransferase 1 | 650.95 | 752.01 | -0.21 | 3.65E-01 |
| ENSMUSG00000022559 | GSMG0014169 | Fbxl6 | F-box and leucine-rich repeat protein 6 | 2246.31 | 2224.89 | 0.01 | 9.61E-01 |
| NULL | GSMG0014170 | NULL | NULL | 90.65 | 95.05 | -0.06 | 9.18E-01 |
| ENSMUSG00000034022 | GSMG0014172 | Cpsf1 | cleavage and polyadenylation specific factor 1 | 4144.40 | 4610.89 | -0.15 | 5.64E-01 |
| NULL | GSMG0014174 | Vps28 | vacuolar protein sorting 28 | 2348.92 | 2433.77 | -0.05 | 8.52E-01 |
| ENSMUSG00000059323 | GSMG0014175 | Tonsl | tonsoku-like, DNA repair protein | 523.65 | 780.52 | -0.58 | 7.63E-04 |
| ENSMUSG00000053929 | GSMG0014176 | Cyhr1 | cysteine and histidine rich 1 | 2886.67 | 1760.60 | 0.71 | 4.06E-03 |
| ENSMUSG00000033697 | GSMG0014180 | Arhgap39 | Rho GTPase activating protein 39 | 640.48 | 781.79 | -0.29 | 4.31E-01 |
| ENSMUSG00000022526 | GSMG0014181 | Zfp251 | zinc finger protein 251 | 892.64 | 824.07 | 0.12 | 6.94E-01 |
| ENSMUSG00000054967 | GSMG0014183 | Zfp647 | zinc finger protein 647 | 239.81 | 356.95 | -0.57 | 1.37E-02 |
| ENSMUSG00000033565 | GSMG0014186 | Rbfox2 | RNA binding protein, fox-1 homolog (C. elegans) 2 | 1198.32 | 1672.97 | -0.48 | 1.94E-02 |
| NULL | GSMG0014189 | NULL | NULL | 684.53 | 784.58 | -0.20 | 3.99E-01 |
| NULL | GSMG0014192 | NULL | NULL | 135.99 | 88.38 | 0.63 | 1.05E-01 |
| NULL | GSMG0014193 | NULL | NULL | 126.85 | 101.25 | 0.32 | 4.32E-01 |
| ENSMUSG00000022443 | GSMG0014202 | Myh9 | myosin, heavy polypeptide 9, non-muscle | 15596.59 | 15377.80 | 0.02 | 9.68E-01 |
| NULL | GSMG0014204 | NULL | NULL | 1121.71 | 769.47 | 0.54 | 5.52E-02 |

| | | | | | | | |
|---|---|---|---|---|---|---|---|
| ENSMUSG00000005354 | GSMG0014205 | Txn2 | thioredoxin 2 | 2500.82 | 2480.41 | 0.01 | 9.67E-01 |
| ENSMUSG00000016552 | GSMG0014207 | Foxred2 | FAD-dependent oxidoreductase domain containing 2 | 119.38 | 216.13 | -0.86 | 9.19E-04 |
| ENSMUSG00000016554 | GSMG0014208 | Eif3d | eukaryotic translation initiation factor 3, subunit D | 830.71 | 1091.51 | -0.39 | 3.34E-02 |
| ENSMUSG00000016637 | GSMG0014210 | Ift27 | intraflagellar transport 27 | 457.81 | 543.11 | -0.25 | 2.95E-01 |
| ENSMUSG00000005716 | GSMG0014211 | Pvalb | parvalbumin | 86.71 | 1.62 | 5.76 | 2.59E-13 |
| ENSMUSG00000044986 | GSMG0014214 | Tst | thiosulfate sulfurtransferase, mitochondrial | 197.29 | 301.58 | -0.61 | 1.07E-02 |
| ENSMUSG00000022440 | GSMG0014218 | C1qtnf6 | C1q and tumor necrosis factor related protein 6 | 1035.29 | 1324.17 | -0.36 | 5.48E-02 |
| ENSMUSG00000033220 | GSMG0014220 | Rac2 | RAS-related C3 botulinum substrate 2 | 80.97 | 45.96 | 0.82 | 3.57E-02 |
| ENSMUSG00000043460 | GSMG0014223 | Elfn2 | leucine rich repeat and fibronectin type III, extracellular 2 | 736.90 | 847.83 | -0.20 | 5.53E-01 |
| ENSMUSG00000018169 | GSMG0014224 | Mfng | MFNG O-fucosylpeptide 3-beta-N-acetylglucosaminyltransferase | 1208.40 | 1482.71 | -0.30 | 1.82E-01 |
| ENSMUSG00000033170 | GSMG0014225 | Card10 | caspase recruitment domain family, member 10 | 3289.45 | 2670.84 | 0.30 | 8.96E-02 |
| ENSMUSG00000033055 | GSMG0014235 | Ankrd54 | ankyrin repeat domain 54 | 1235.27 | 1198.21 | 0.04 | 8.78E-01 |
| ENSMUSG00000033006 | GSMG0014239 | Sox10 | SRY (sex determining region Y)-box 10 | 101.65 | 72.18 | 0.49 | 2.80E-01 |
| NULL | GSMG0014242 | NULL | NULL | 54.01 | 44.12 | 0.29 | 6.30E-01 |
| ENSMUSG00000042632 | GSMG0014245 | Pla2g6 | phospholipase A2, group VI | 912.80 | 1422.94 | -0.64 | 4.39E-05 |
| ENSMUSG00000009035 | GSMG0014246 | Tmem184b | transmembrane protein 184b | 7078.39 | 6332.51 | 0.16 | 4.69E-01 |
| ENSMUSG00000022433 | GSMG0014247 | Csnk1e | casein kinase 1, epsilon | 2478.24 | 3072.11 | -0.31 | 8.51E-02 |
| ENSMUSG00000044216 | GSMG0014249 | Kcnj4 | potassium inwardly-rectifying channel, subfamily J, member 4 | 286.99 | 112.68 | 1.34 | 2.39E-05 |
| ENSMUSG00000055065 | GSMG0014250 | Ddx17 | DEAD (Asp-Glu-Ala-Asp) box polypeptide 17 | 680.14 | 701.37 | -0.04 | 9.03E-01 |
| NULL | GSMG0014251 | NULL | NULL | 244.33 | 271.98 | -0.16 | 6.12E-01 |
| ENSMUSG00000022426 | GSMG0014256 | Josd1 | Josephin domain containing 1 | 2014.11 | 1588.32 | 0.34 | 6.36E-02 |
| ENSMUSG00000022420 | GSMG0014259 | Dnal4 | dynein, axonemal, light chain 4 | 832.26 | 785.11 | 0.08 | 7.72E-01 |
| 21,ENSMUSG00000089715,EI | GSMG0014260 | Npcd // Nptxr // Cb | neuronal pentraxin chromo domain // neuronal pentraxin receptor // chr | 601.97 | 555.19 | 0.12 | 8.19E-01 |
| ENSMUSG00000000489 | GSMG0014263 | Pdgfb | platelet derived growth factor, B polypeptide | 1781.82 | 1333.21 | 0.42 | 1.73E-02 |
| ENSMUSG00000051518 | GSMG0014268 | Rps19bp1 | ribosomal protein S19 binding protein 1 | 453.45 | 723.51 | -0.68 | 2.57E-03 |
| NULL | GSMG0014272 | NULL | NULL | 53.24 | 48.51 | 0.13 | 8.42E-01 |
| ENSMUSG00000042292 | GSMG0014273 | Mkl1 | MKL (megakaryoblastic leukemia)/myocardin-like 1 | 518.29 | 491.17 | 0.08 | 8.57E-01 |
| NULL | GSMG0014278 | NULL | NULL | 214.80 | 149.99 | 0.51 | 7.97E-02 |
| ENSMUSG00000022404 | GSMG0014280 | Slc25a17 | solute carrier family 25 (mitochondrial carrier, peroxisomal membrane p | 757.97 | 911.60 | -0.27 | 2.20E-01 |
| ENSMUSG00000022403 | GSMG0014281 | St13 | suppression of tumorigenicity 13 | 1053.36 | 1532.15 | -0.54 | 6.84E-03 |
| ENSMUSG00000063765 | GSMG0014284 | Chadl | chondroadherin-like | 367.92 | 392.28 | -0.10 | 8.12E-01 |
| ENSMUSG00000022391 | GSMG0014285 | Rangap1 | RAN GTPase activating protein 1 | 355.68 | 521.64 | -0.55 | 3.18E-03 |
| ENSMUSG00000061360 | GSMG0014288 | Phf5a | PHD finger protein 5A | 1017.27 | 1245.59 | -0.29 | 1.51E-01 |

| | | | | | | | |
|---|---|---|---|---|---|---|---|
| ENSMUSG00000022476 | GSMG0014289 | Polr3h | polymerase (RNA) III (DNA directed) polypeptide H | 702.52 | 795.72 | -0.18 | 5.09E-01 |
| ENSMUSG00000022474 | GSMG0014290 | Pmm1 | phosphomannomutase 1 | 567.64 | 628.77 | -0.15 | 5.50E-01 |
| ENSMUSG00000022472 | GSMG0014291 | Desi1 | desumoylating isopeptidase 1 | 1403.10 | 1211.89 | 0.21 | 2.87E-01 |
| ENSMUSG00000068101 | GSMG0014295 | Cenpm | centromere protein M | 233.76 | 534.31 | -1.19 | 1.13E-09 |
| ENSMUSG00000022453 | GSMG0014296 | Naga | N-acetyl galactosaminidase, alpha | 776.10 | 551.14 | 0.49 | 7.73E-03 |
| ENSMUSG00000022450 | GSMG0014297 | Ndufa6 | NADH dehydrogenase (ubiquinone) 1 alpha subcomplex, 6 (B14) | 1688.66 | 1752.07 | -0.05 | 8.87E-01 |
| ENSMUSG00000061740 | GSMG0014298 | Cyp2d22 | cytochrome P450, family 2, subfamily d, polypeptide 22 | 1744.66 | 1047.46 | 0.74 | 2.41E-06 |
| ENSMUSG00000058099 | GSMG0014312 | Nfam1 | Nfat activating molecule with ITAM motif 1 | 81.88 | 78.52 | 0.07 | 9.18E-01 |
| ENSMUSG00000018040 | GSMG0014313 | Rrp7a | ribosomal RNA processing 7 homolog A (S. cerevisiae) | 456.63 | 571.59 | -0.32 | 1.43E-01 |
| ENSMUSG00000041815 | GSMG0014314 | Poldip3 | polymerase (DNA-directed), delta interacting protein 3 | 1250.10 | 1408.22 | -0.17 | 3.92E-01 |
| ENSMUSG00000018042 | GSMG0014315 | Cyb5r3 | cytochrome b5 reductase 3 | 4331.14 | 3250.27 | 0.41 | 1.70E-02 |
| ENSMUSG00000047878 | GSMG0014317 | A4galt | alpha 1,4-galactosyltransferase | 723.21 | 444.94 | 0.70 | 1.98E-04 |
| ENSMUSG00000054277 | GSMG0014319 | Arfgap3 | ADP-ribosylation factor GTPase activating protein 3 | 943.75 | 852.91 | 0.15 | 5.54E-01 |
| ENSMUSG00000016664 | GSMG0014321 | Pacsin2 | protein kinase C and casein kinase substrate in neurons 2 | 919.78 | 819.32 | 0.17 | 5.20E-01 |
| NULL | GSMG0014322 | NULL | NULL | 110.53 | 89.50 | 0.30 | 4.44E-01 |
| ENSMUSG00000022442 | GSMG0014324 | Ttll1 | tubulin tyrosine ligase-like 1 | 335.40 | 524.45 | -0.64 | 2.37E-03 |
| ENSMUSG00000016757 | GSMG0014326 | Ttll12 | tubulin tyrosine ligase-like family, member 12 | 355.61 | 546.29 | -0.62 | 1.71E-03 |
| ENSMUSG00000016763 | GSMG0014327 | Scube1 | signal peptide, CUB domain, EGF-like 1 | 601.86 | 528.85 | 0.19 | 5.64E-01 |
| ENSMUSG00000018865 | GSMG0014329 | Sult4a1 | sulfotransferase family 4A, member 1 | 271.50 | 299.57 | -0.14 | 6.58E-01 |
| ENSMUSG00000062760 | GSMG0014331 | 1810041L15Rik | RIKEN cDNA 1810041L15 gene | 734.30 | 445.47 | 0.72 | 1.56E-03 |
| ENSMUSG00000055745 | GSMG0014332 | Ldoc1l | leucine zipper, down-regulated in cancer 1-like | 745.52 | 1016.75 | -0.45 | 4.01E-02 |
| ENSMUSG00000036046 | GSMG0014337 | 5031439G07Rik | RIKEN cDNA 5031439G07 gene | 2647.26 | 977.22 | 1.44 | 8.16E-14 |
| ENSMUSG00000022382 | GSMG0014342 | Wnt7b | wingless-type MMTV integration site family, member 7B | 319.04 | 634.11 | -0.99 | 4.45E-08 |
| ENSMUSG00000064284 | GSMG0014343 | Cdpf1 | cysteine rich, DPF motif domain containing 1 | 335.17 | 294.03 | 0.19 | 4.88E-01 |
| ENSMUSG00000035891 | GSMG0014347 | Cerk | ceramide kinase | 634.25 | 690.54 | -0.12 | 6.87E-01 |
| NULL | GSMG0014349 | NULL | NULL | 555.68 | 497.64 | 0.16 | 6.33E-01 |
| NULL | GSMG0014350 | NULL | NULL | 83.12 | 66.34 | 0.33 | 4.77E-01 |
| NULL | GSMG0014353 | NULL | NULL | 64.68 | 41.89 | 0.63 | 1.56E-01 |
| ENSMUSG00000022387 | GSMG0014358 | Brd1 | bromodomain containing 1 | 296.02 | 238.20 | 0.31 | 3.57E-01 |
| ENSMUSG00000035845 | GSMG0014359 | Alg12 | asparagine-linked glycosylation 12 (alpha-1,6-mannosyltransferase) | 241.95 | 254.60 | -0.07 | 8.47E-01 |
| ENSMUSG00000035805 | GSMG0014362 | Mlc1 | megalencephalic leukoencephalopathy with subcortical cysts 1 homolog | 1330.75 | 1153.00 | 0.21 | 3.07E-01 |
| ENSMUSG00000062906 | GSMG0014365 | Hdac10 | histone deacetylase 10 | 1180.06 | 2020.00 | -0.78 | 9.94E-07 |
| ENSMUSG00000022610 | GSMG0014366 | Mapk12 | mitogen-activated protein kinase 12 | 317.54 | 368.53 | -0.21 | 4.44E-01 |

| | | | | | | | |
|---|---|---|---|---|---|---|---|
| ENSMUSG00000053137 | GSMG0014367 | Mapk11 | mitogen-activated protein kinase 11 | 1204.47 | 1266.78 | -0.07 | 8.13E-01 |
| ENSMUSG00000036606 | GSMG0014368 | Plxnb2 | plexin B2 | 2034.21 | 2620.97 | -0.37 | 1.28E-01 |
| ENSMUSG00000015377 | GSMG0014369 | Dennd6b | DENN/MADD domain containing 6B | 3843.75 | 3198.55 | 0.27 | 2.18E-01 |
| NULL | GSMG0014371 | NULL | NULL | 63.76 | 100.16 | -0.66 | 1.54E-01 |
| ENSMUSG00000036529 | GSMG0014372 | Sbf1 | SET binding factor 1 | 1414.05 | 1348.81 | 0.07 | 8.90E-01 |
| ENSMUSG00000022614 | GSMG0014373 | Lmf2 | lipase maturation factor 2 | 2306.47 | 2370.89 | -0.04 | 8.92E-01 |
| G00000022615,ENSMUSG000 | GSMG0014376 | Sco2 // Tymp | SCO2 cytochrome c oxidase assembly protein // thymidine phosphoryla | 400.35 | 396.86 | 0.01 | 9.71E-01 |
| ENSMUSG00000047394 | GSMG0014377 | Odf3b | outer dense fiber of sperm tails 3B | 153.07 | 75.30 | 1.03 | 4.40E-04 |
| G00000022617,ENSMUSG000 | GSMG0014379 | Gm44502 // Cpt1b | predicted readthrough transcript (NMD candidate), 44502 // carnitine pa | 1253.41 | 1288.15 | -0.04 | 8.95E-01 |
| NULL | GSMG0014380 | NULL | NULL | 154.51 | 257.14 | -0.73 | 2.76E-03 |
| ENSMUSG00000022620 | GSMG0014381 | Arsa | arylsulfatase A | 734.66 | 755.36 | -0.04 | 9.09E-01 |
| ENSMUSG00000022621 | GSMG0014383 | Rabl2 | RAB, member RAS oncogene family-like 2 | 500.94 | 464.49 | 0.11 | 7.51E-01 |
| ENSMUSG00000052560 | GSMG0014388 | Cpne8 | copine VIII | 834.13 | 767.33 | 0.12 | 6.70E-01 |
| ENSMUSG00000022629 | GSMG0014391 | Kif21a | kinesin family member 21A | 877.88 | 976.36 | -0.15 | 5.84E-01 |
| ENSMUSG00000055782 | GSMG0014395 | Abcd2 | ATP-binding cassette, sub-family D (ALD), member 2 | 179.84 | 105.33 | 0.78 | 5.91E-03 |
| ENSMUSG00000036298 | GSMG0014396 | Slc2a13 | solute carrier family 2 (facilitated glucose transporter), member 13 | 281.61 | 206.85 | 0.45 | 1.35E-01 |
| ENSMUSG00000036197 | GSMG0014403 | Gxylt1 | glucoside xylosyltransferase 1 | 299.10 | 277.19 | 0.11 | 7.56E-01 |
| ENSMUSG00000022634 | GSMG0014407 | Yaf2 | YY1 associated factor 2 | 2185.61 | 2455.77 | -0.17 | 4.42E-01 |
| ENSMUSG00000022635 | GSMG0014410 | Zcrb1 | zinc finger CCHC-type and RNA binding motif 1 | 847.01 | 882.90 | -0.06 | 8.55E-01 |
| ENSMUSG00000036158 | GSMG0014411 | Prickle1 | prickle planar cell polarity protein 1 | 1681.23 | 1042.89 | 0.69 | 2.11E-06 |
| NULL | GSMG0014412 | NULL | NULL | 148.43 | 63.35 | 1.23 | 2.68E-05 |
| NULL | GSMG0014413 | NULL | NULL | 422.42 | 157.06 | 1.43 | 1.87E-11 |
| NULL | GSMG0014415 | NULL | NULL | 538.20 | 213.67 | 1.33 | 9.10E-11 |
| ENSMUSG00000022449 | GSMG0014419 | Adamts20 | a disintegrin-like and metallopeptidase (reprolysin type) with thrombos | 536.02 | 633.18 | -0.24 | 4.45E-01 |
| ENSMUSG00000033356 | GSMG0014420 | Pus7l | pseudouridylate synthase 7-like | 167.31 | 215.52 | -0.37 | 2.20E-01 |
| ENSMUSG00000022451 | GSMG0014421 | Twf1 | twinfilin actin binding protein 1 | 386.11 | 497.26 | -0.37 | 1.37E-01 |
| ENSMUSG00000022454 | GSMG0014424 | Nell2 | NEL-like 2 | 1981.56 | 3606.52 | -0.86 | 5.36E-06 |
| NULL | GSMG0014429 | NULL | NULL | 214.55 | 226.24 | -0.07 | 8.87E-01 |
| NULL | GSMG0014441 | E330033B04Rik | RIKEN cDNA E330033B04 gene | 798.36 | 632.91 | 0.33 | 1.53E-01 |
| ENSMUSG00000033228 | GSMG0014443 | Scaf11 | SR-related CTD-associated factor 11 | 320.72 | 401.31 | -0.33 | 1.88E-01 |
| ENSMUSG00000023169 | GSMG0014444 | Slc38a1 | solute carrier family 38, member 1 | 616.44 | 1079.68 | -0.81 | 1.77E-05 |
| ENSMUSG00000022462 | GSMG0014445 | Slc38a2 | solute carrier family 38, member 2 | 11728.93 | 35801.81 | -1.61 | 9.30E-35 |
| ENSMUSG00000048218 | GSMG0014447 | Amigo2 | adhesion molecule with Ig like domain 2 | 385.06 | 234.67 | 0.72 | 6.15E-04 |

| | | | | | | | |
|---|---|---|---|---|---|---|---|
| ENSMUSG00000022466 | GSMG0014449 | Rpap3 | RNA polymerase II associated protein 3 | 855.10 | 1416.80 | -0.73 | 6.90E-05 |
| ENSMUSG00000022469 | GSMG0014451 | Rapgef3 | Rap guanine nucleotide exchange factor (GEF) 3 | 861.57 | 809.07 | 0.09 | 7.66E-01 |
| ENSMUSG00000022475 | GSMG0014452 | Hdac7 | histone deacetylase 7 | 2793.99 | 3440.98 | -0.30 | 2.97E-01 |
| ENSMUSG00000033075 | GSMG0014458 | Senp1 | SUMO1/sentrin specific peptidase 1 | 763.16 | 930.99 | -0.29 | 1.36E-01 |
| ENSMUSG00000048175 | GSMG0014460 | Asb8 | ankyrin repeat and SOCS box-containing 8 | 1550.06 | 1520.21 | 0.03 | 9.22E-01 |
| ENSMUSG00000022987 | GSMG0014464 | Zfp641 | zinc finger protein 641 | 56.80 | 43.18 | 0.39 | 4.88E-01 |
| ENSMUSG00000011960 | GSMG0014470 | Ccnt1 | cyclin T1 | 1654.11 | 1736.10 | -0.07 | 7.83E-01 |
| ENSMUSG00000022994 | GSMG0014471 | Adcy6 | adenylate cyclase 6 | 716.19 | 573.59 | 0.32 | 3.12E-01 |
| NULL | GSMG0014472 | NULL | NULL | 126.46 | 237.02 | -0.90 | 1.77E-03 |
| ENSMUSG00000054855 | GSMG0014474 | Rnd1 | Rho family GTPase 1 | 436.09 | 421.27 | 0.05 | 8.84E-01 |
| ENSMUSG00000003355 | GSMG0014475 | Fkbp11 | FK506 binding protein 11 | 98.60 | 93.23 | 0.08 | 9.05E-01 |
| ENSMUSG00000051853 | GSMG0014476 | Arf3 | ADP-ribosylation factor 3 | 6671.24 | 5589.51 | 0.26 | 1.95E-01 |
| ENSMUSG00000059213 | GSMG0014479 | Ddn | dendrin | 919.90 | 562.51 | 0.71 | 2.73E-04 |
| ENSMUSG00000067713 | GSMG0014480 | Prkag1 | protein kinase, AMP-activated, gamma 1 non-catalytic subunit | 767.54 | 704.34 | 0.13 | 6.33E-01 |
| ENSMUSG00000023755 | GSMG0014485 | Rhebl1 | Ras homolog enriched in brain like 1 | 206.55 | 330.39 | -0.68 | 1.48E-03 |
| ENSMUSG00000022999 | GSMG0014487 | Lmbr1l | limb region 1 like | 566.63 | 666.31 | -0.23 | 3.76E-01 |
| ENSMUSG00000037570 | GSMG0014492 | Mcrs1 | microspherule protein 1 | 804.65 | 1093.61 | -0.44 | 7.44E-03 |
| ENSMUSG00000023008 | GSMG0014493 | Fmnl3 | formin-like 3 | 2966.94 | 3601.44 | -0.28 | 3.65E-01 |
| ENSMUSG00000023009 | GSMG0014494 | Nckap5l | NCK-associated protein 5-like | 495.93 | 605.41 | -0.29 | 2.05E-01 |
| ENSMUSG00000037525 | GSMG0014495 | Bcdin3d | BCDIN3 domain containing | 175.59 | 214.04 | -0.29 | 3.41E-01 |
| ENSMUSG00000023011 | GSMG0014496 | Faim2 | Fas apoptotic inhibitory molecule 2 | 359.71 | 238.31 | 0.59 | 3.41E-02 |
| ENSMUSG00000023015 | GSMG0014497 | Racgap1 | Rac GTPase-activating protein 1 | 556.73 | 1232.31 | -1.14 | 7.24E-13 |
| NULL | GSMG0014498 | NULL | NULL | 119.34 | 54.80 | 1.12 | 3.46E-04 |
| ENSMUSG00000023021 | GSMG0014500 | Cers5 | ceramide synthase 5 | 1262.35 | 1485.95 | -0.23 | 2.50E-01 |
| ENSMUSG00000023022 | GSMG0014501 | Lima1 | LIM domain and actin binding 1 | 686.75 | 304.89 | 1.17 | 2.38E-11 |
| ENSMUSG00000023030 | GSMG0014508 | Slc11a2 | solute carrier family 11 (proton-coupled divalent metal ion transporters) | 1378.00 | 1437.02 | -0.06 | 8.30E-01 |
| ENSMUSG00000044636 | GSMG0014510 | Csrnp2 | cysteine-serine-rich nuclear protein 2 | 184.61 | 195.49 | -0.09 | 8.61E-01 |
| ENSMUSG00000009733 | GSMG0014512 | Tfcp2 | transcription factor CP2 | 376.66 | 370.90 | 0.02 | 9.58E-01 |
| ENSMUSG00000009739 | GSMG0014513 | Pou6f1 | POU domain, class 6, transcription factor 1 | 112.53 | 111.32 | 0.02 | 9.72E-01 |
| ENSMUSG00000053559 | GSMG0014516 | Smagp | small cell adhesion glycoprotein | 720.02 | 439.82 | 0.71 | 4.36E-05 |
| NULL | GSMG0014522 | NULL | NULL | 206.92 | 106.08 | 0.97 | 7.88E-04 |
| NULL | GSMG0014523 | NULL | NULL | 75.16 | 34.50 | 1.13 | 7.73E-03 |
| NULL | GSMG0014524 | NULL | NULL | 174.38 | 127.83 | 0.45 | 2.44E-01 |

| | | | | | | | |
|---|---|---|---|---|---|---|---|
| ENSMUSG00000097915 | GSMG0014525 | A330009N23Rik | RIKEN cDNA A330009N23 gene | 202.69 | 136.60 | 0.57 | 4.56E-02 |
| ENSMUSG00000036966 | GSMG0014550 | Spryd3 | SPRY domain containing 3 | 496.70 | 488.49 | 0.02 | 9.50E-01 |
| ENSMUSG00000023044 | GSMG0014551 | Csad | cysteine sulfinic acid decarboxylase | 763.20 | 628.19 | 0.28 | 2.69E-01 |
| ENSMUSG00000001288 | GSMG0014554 | Rarg | retinoic acid receptor, gamma | 340.32 | 293.64 | 0.21 | 6.54E-01 |
| ENSMUSG00000036678 | GSMG0014556 | Aaas | achalasia, adrenocortical insufficiency, alacrimia | 648.25 | 872.89 | -0.43 | 2.87E-02 |
| ENSMUSG00000023050 | GSMG0014559 | Map3k12 | mitogen-activated protein kinase kinase kinase 12 | 2322.73 | 2228.98 | 0.06 | 8.51E-01 |
| G00000023052,ENSMUSG000 | GSMG0014562 | Npff // Atf7 | neuropeptide FF-amide peptide precursor // activating transcription fact | 492.40 | 562.42 | -0.19 | 5.08E-01 |
| ENSMUSG00000023055 | GSMG0014565 | Calcoco1 | calcium binding and coiled coil domain 1 | 3723.08 | 2952.65 | 0.33 | 4.52E-02 |
| ENSMUSG00000009575 | GSMG0014572 | Cbx5 | chromobox 5 | 1365.72 | 1992.88 | -0.54 | 3.32E-04 |
| ENSMUSG00000036061 | GSMG0014573 | Smug1 | single-strand selective monofunctional uracil DNA glycosylase | 723.08 | 934.99 | -0.37 | 4.44E-02 |
| ENSMUSG00000000552 | GSMG0014581 | Zfp385a | zinc finger protein 385A | 308.24 | 307.78 | 0.00 | 9.97E-01 |
| ENSMUSG00000000555 | GSMG0014582 | Itga5 | integrin alpha 5 (fibronectin receptor alpha) | 2950.97 | 3792.65 | -0.36 | 2.78E-02 |
| ENSMUSG00000022490 | GSMG0014586 | Ppp1r1a | protein phosphatase 1, regulatory (inhibitor) subunit 1A | 783.42 | 1066.78 | -0.45 | 1.46E-02 |
| ENSMUSG00000022529 | GSMG0014589 | Zfp263 | zinc finger protein 263 | 2123.58 | 2273.81 | -0.10 | 6.86E-01 |
| ENSMUSG00000005982 | GSMG0014594 | Naa60 | N(alpha)-acetyltransferase 60, NatF catalytic subunit | 1067.45 | 1013.82 | 0.08 | 7.92E-01 |
| ENSMUSG00000014232 | GSMG0014595 | Cluap1 | clusterin associated protein 1 | 235.95 | 288.22 | -0.29 | 2.97E-01 |
| ENSMUSG00000014303 | GSMG0014599 | Glis2 | GLIS family zinc finger 2 | 173.62 | 154.83 | 0.16 | 7.51E-01 |
| ENSMUSG00000039646 | GSMG0014601 | Vasn | vasorin | 765.04 | 548.75 | 0.48 | 2.02E-02 |
| ENSMUSG00000004069 | GSMG0014602 | Dnaja3 | DnaJ heat shock protein family (Hsp40) member A3 | 1711.87 | 1700.71 | 0.01 | 9.74E-01 |
| ENSMUSG00000004070 | GSMG0014603 | Hmox2 | heme oxygenase 2 | 73.76 | 68.01 | 0.12 | 8.36E-01 |
| ENSMUSG00000022517 | GSMG0014608 | Mgrn1 | mahogunin, ring finger 1 | 4945.52 | 4442.23 | 0.15 | 5.19E-01 |
| ENSMUSG00000022516 | GSMG0014609 | Nudt16l1 | nudix (nucleoside diphosphate linked moiety X)-type motif 16-like 1 | 1814.42 | 1831.81 | -0.01 | 9.64E-01 |
| ENSMUSG00000039473 | GSMG0014612 | Ubn1 | ubinuclein 1 | 1431.12 | 1278.01 | 0.16 | 4.53E-01 |
| ENSMUSG00000039427 | GSMG0014614 | Alg1 | asparagine-linked glycosylation 1 (beta-1,4-mannosyltransferase) | 1027.95 | 1061.57 | -0.05 | 8.71E-01 |
| ENSMUSG00000008658 | GSMG0014618 | Rbfox1 | RNA binding protein, fox-1 homolog (C. elegans) 1 | 634.71 | 555.84 | 0.19 | 4.03E-01 |
| NULL | GSMG0014621 | NULL | NULL | 580.08 | 591.53 | -0.03 | 9.43E-01 |
| NULL | GSMG0014623 | NULL | NULL | 420.06 | 320.79 | 0.39 | 1.14E-01 |
| NULL | GSMG0014624 | NULL | NULL | 82.97 | 70.23 | 0.25 | 6.11E-01 |
| NULL | GSMG0014625 | NULL | NULL | 275.40 | 230.93 | 0.25 | 3.53E-01 |
| NULL | GSMG0014626 | NULL | NULL | 202.14 | 158.22 | 0.36 | 3.95E-01 |
| NULL | GSMG0014627 | NULL | NULL | 638.15 | 558.54 | 0.19 | 5.45E-01 |
| NULL | GSMG0014628 | NULL | NULL | 326.63 | 234.26 | 0.48 | 5.62E-02 |
| NULL | GSMG0014631 | NULL | NULL | 180.21 | 176.84 | 0.03 | 9.47E-01 |

| | | | | | | | |
|---|---|---|---|---|---|---|---|
| NULL | GSMG0014635 | NULL | NULL | 2036.63 | 1370.21 | 0.57 | 8.08E-04 |
| NULL | GSMG0014636 | NULL | NULL | 1269.43 | 883.98 | 0.52 | 5.24E-03 |
| NULL | GSMG0014638 | NULL | NULL | 1719.17 | 1286.02 | 0.42 | 2.92E-02 |
| ENSMUSG00000039345 | GSMG0014644 | Mettl22 | methyltransferase like 22 | 355.60 | 476.47 | -0.42 | 4.89E-02 |
| ENSMUSG00000057880 | GSMG0014645 | Abat | 4-aminobutyrate aminotransferase | 586.68 | 503.00 | 0.22 | 4.05E-01 |
| ENSMUSG00000022711 | GSMG0014647 | Pmm2 | phosphomannomutase 2 | 466.76 | 496.50 | -0.09 | 7.61E-01 |
| ENSMUSG00000022507 | GSMG0014648 | 1810013L24Rik | RIKEN cDNA 1810013L24 gene | 1423.18 | 1203.55 | 0.24 | 2.80E-01 |
| NULL | GSMG0014649 | NULL | NULL | 445.34 | 577.99 | -0.38 | 1.51E-01 |
| ENSMUSG00000022503 | GSMG0014656 | Nubp1 | nucleotide binding protein 1 | 1097.86 | 1403.36 | -0.35 | 4.69E-02 |
| ENSMUSG00000068663 | GSMG0014658 | Clec16a | C-type lectin domain family 16, member A | 792.11 | 731.59 | 0.12 | 6.97E-01 |
| ENSMUSG00000037972 | GSMG0014667 | Snn | stannin | 3035.08 | 3973.03 | -0.39 | 1.69E-01 |
| ENSMUSG00000071669 | GSMG0014674 | Snx29 | sorting nexin 29 | 298.07 | 201.33 | 0.57 | 2.48E-02 |
| ENSMUSG00000022494 | GSMG0014678 | Shisa9 | shisa family member 9 | 528.31 | 707.64 | -0.42 | 3.34E-02 |
| ENSMUSG00000022545 | GSMG0014681 | Ercc4 | excision repair cross-complementing rodent repair deficiency, complem | 455.00 | 457.47 | -0.01 | 9.82E-01 |
| ENSMUSG00000009569 | GSMG0014686 | Mkl2 | MKL/myocardin-like 2 | 2345.07 | 1767.08 | 0.41 | 1.44E-02 |
| NULL | GSMG0014687 | NULL | NULL | 499.41 | 196.48 | 1.34 | 1.37E-09 |
| G00000022684,ENSMUSG000 | GSMG0014691 | Bfar // 3110001I22 | bifunctional apoptosis regulator // RIKEN cDNA 3110001I22 gene | 878.78 | 725.36 | 0.28 | 2.31E-01 |
| ENSMUSG00000022682 | GSMG0014695 | Rrn3 | RRN3 RNA polymerase I transcription factor homolog (yeast) | 736.00 | 866.60 | -0.24 | 2.61E-01 |
| ENSMUSG00000022681 | GSMG0014696 | Ntan1 | N-terminal Asn amidase | 1407.50 | 1511.91 | -0.10 | 6.88E-01 |
| ENSMUSG00000022679 | GSMG0014697 | Mpv17l | Mpv17 transgene, kidney disease mutant-like | 347.63 | 354.49 | -0.03 | 9.41E-01 |
| ENSMUSG00000044117 | GSMG0014700 | 2900011O08Rik | RIKEN cDNA 2900011O08 gene | 1120.29 | 1587.43 | -0.50 | 1.14E-02 |
| ENSMUSG00000022678 | GSMG0014703 | Nde1 | nudE neurodevelopment protein 1 | 882.51 | 838.97 | 0.07 | 8.26E-01 |
| ENSMUSG00000023088 | GSMG0014706 | Abcc1 | ATP-binding cassette, sub-family C (CFTR/MRP), member 1 | 356.18 | 340.44 | 0.07 | 8.72E-01 |
| ENSMUSG00000022676 | GSMG0014709 | Snai2 | snail family zinc finger 2 | 109.56 | 110.51 | -0.01 | 9.79E-01 |
| ENSMUSG00000068617 | GSMG0014710 | Efcab1 | EF hand calcium binding domain 1 | 50.74 | 52.87 | -0.05 | 9.46E-01 |
| ENSMUSG00000022672 | GSMG0014711 | Prkdc | protein kinase, DNA activated, catalytic polypeptide | 197.93 | 140.32 | 0.50 | 7.80E-02 |
| ENSMUSG00000071637 | GSMG0014713 | Cebpd | CCAAT/enhancer binding protein (C/EBP), delta | 2176.71 | 807.97 | 1.43 | 1.94E-13 |
| ENSMUSG00000022792 | GSMG0014716 | Yars2 | tyrosyl-tRNA synthetase 2 (mitochondrial) | 429.47 | 583.08 | -0.44 | 2.84E-02 |
| ENSMUSG00000022779 | GSMG0014718 | Top3b | topoisomerase (DNA) III beta | 2227.48 | 2038.79 | 0.13 | 5.53E-01 |
| ENSMUSG00000026181 | GSMG0014720 | Ppm1f | protein phosphatase 1F (PP2C domain containing) | 1748.45 | 1524.26 | 0.20 | 4.06E-01 |
| ENSMUSG00000063358 | GSMG0014722 | Mapk1 | mitogen-activated protein kinase 1 | 2331.15 | 2290.75 | 0.03 | 9.31E-01 |
| ENSMUSG00000022773 | GSMG0014723 | Ypel1 | yippee-like 1 (Drosophila) | 151.39 | 261.00 | -0.78 | 7.28E-04 |
| ENSMUSG00000041774 | GSMG0014724 | Ydjc | YdjC homolog (bacterial) | 71.85 | 198.11 | -1.47 | 4.45E-08 |

| | | | | | | | |
|---|---|---|---|---|---|---|---|
| NULL | GSMG0014725 | NULL | NULL | 137.74 | 174.04 | -0.34 | 4.18E-01 |
| NULL | GSMG0014726 | NULL | NULL | 272.35 | 306.92 | -0.17 | 5.98E-01 |
| ENSMUSG00000050240 | GSMG0014728 | Hic2 | hypermethylated in cancer 2 | 646.31 | 810.94 | -0.33 | 2.12E-01 |
| ENSMUSG00000055692 | GSMG0014729 | Tmem191c | transmembrane protein 191C | 6721.33 | 5596.72 | 0.26 | 2.03E-01 |
| ENSMUSG00000022765 | GSMG0014731 | Snap29 | synaptosomal-associated protein 29 | 733.91 | 674.94 | 0.12 | 6.82E-01 |
| ENSMUSG00000006134 | GSMG0014732 | Crkl | v-crk avian sarcoma virus CT10 oncogene homolog-like | 2020.52 | 2147.09 | -0.09 | 6.93E-01 |
| ENSMUSG00000022763 | GSMG0014734 | Aifm3 | apoptosis-inducing factor, mitochondrion-associated 3 | 587.65 | 345.46 | 0.77 | 5.31E-03 |
| ENSMUSG00000022761 | GSMG0014735 | Lztr1 | leucine-zipper-like transcriptional regulator, 1 | 1655.61 | 1515.98 | 0.13 | 6.40E-01 |
| ENSMUSG00000005899 | GSMG0014737 | Smpd4 | sphingomyelin phosphodiesterase 4 | 1113.35 | 1101.52 | 0.02 | 9.62E-01 |
| ENSMUSG00000041617 | GSMG0014738 | Ccdc74a | coiled-coil domain containing 74A | 256.78 | 211.08 | 0.28 | 4.02E-01 |
| ENSMUSG00000022750 | GSMG0014739 | Klhl22 | kelch-like 22 | 1727.97 | 1673.82 | 0.05 | 8.81E-01 |
| ENSMUSG00000012017 | GSMG0014740 | Scarf2 | scavenger receptor class F, member 2 | 660.52 | 524.28 | 0.33 | 2.04E-01 |
| ENSMUSG00000003531 | GSMG0014746 | Dgcr6 | DiGeorge syndrome critical region gene 6 | 1627.33 | 1696.19 | -0.06 | 8.15E-01 |
| ENSMUSG00000043811 | GSMG0014748 | Rtn4r | reticulon 4 receptor | 580.22 | 474.55 | 0.29 | 3.38E-01 |
| ENSMUSG00000022721 | GSMG0014750 | Trmt2a | TRM2 tRNA methyltransferase 2A | 1761.73 | 1673.78 | 0.07 | 7.92E-01 |
| ENSMUSG00000000325 | GSMG0014751 | Arvcf | armadillo repeat gene deleted in velo-cardio-facial syndrome | 1309.05 | 1335.58 | -0.03 | 9.42E-01 |
| ENSMUSG00000075704 | GSMG0014753 | Txnrd2 | thioredoxin reductase 2 | 408.29 | 441.13 | -0.11 | 7.40E-01 |
| ENSMUSG00000000884 | GSMG0014755 | Gnb1l | guanine nucleotide binding protein (G protein), beta polypeptide 1-like | 87.99 | 120.29 | -0.45 | 2.11E-01 |
| ENSMUSG00000041378 | GSMG0014759 | Cldn5 | claudin 5 | 52115.28 | 47649.68 | 0.13 | 6.49E-01 |
| ENSMUSG00000005262 | GSMG0014760 | Ufd1l | ubiquitin fusion degradation 1 like | 1260.76 | 1530.72 | -0.28 | 1.22E-01 |
| ENSMUSG00000071632 | GSMG0014762 | 2510002D24Rik | RIKEN cDNA 2510002D24 gene | 989.45 | 955.84 | 0.05 | 8.59E-01 |
| ENSMUSG00000022702 | GSMG0014763 | Hira | histone cell cycle regulator | 64.45 | 101.23 | -0.64 | 1.20E-01 |
| ENSMUSG00000062901 | GSMG0014768 | Klhl24 | kelch-like 24 | 1977.71 | 1704.78 | 0.21 | 3.76E-01 |
| ENSMUSG00000041215 | GSMG0014770 | Yeats2 | YEATS domain containing 2 | 345.93 | 389.94 | -0.17 | 5.76E-01 |
| ENSMUSG00000003235 | GSMG0014772 | Eif2b5 | eukaryotic translation initiation factor 2B, subunit 5 epsilon | 2143.21 | 2323.74 | -0.12 | 5.79E-01 |
| ENSMUSG00000003233 | GSMG0014773 | Dvl3 | dishevelled segment polarity protein 3 | 1028.35 | 1135.52 | -0.14 | 6.20E-01 |
| ENSMUSG00000003234 | GSMG0014775 | Abcf3 | ATP-binding cassette, sub-family F (GCN20), member 3 | 2528.88 | 2314.48 | 0.13 | 5.42E-01 |
| G00000022842,ENSMUSG000 | GSMG0014779 | Ece2 // Eef1akmt4 | endothelin converting enzyme 2 // EEF1A lysine methyltransferase 4 | 477.39 | 487.75 | -0.03 | 9.43E-01 |
| ENSMUSG00000006998 | GSMG0014781 | Psmd2 | proteasome (prosome, macropain) 26S subunit, non-ATPase, 2 | 7206.88 | 8570.90 | -0.25 | 1.67E-01 |
| ENSMUSG00000050821 | GSMG0014783 | Fam131a | family with sequence similarity 131, member A | 528.86 | 328.80 | 0.69 | 1.81E-03 |
| ENSMUSG00000021018 | GSMG0014784 | Polr2h | polymerase (RNA) II (DNA directed) polypeptide H | 150.50 | 164.98 | -0.13 | 7.51E-01 |
| ENSMUSG00000006958 | GSMG0014785 | Chrd | chordin | 1116.67 | 570.56 | 0.97 | 1.80E-05 |
| ENSMUSG00000005958 | GSMG0014786 | Ephb3 | Eph receptor B3 | 400.79 | 299.48 | 0.42 | 1.45E-01 |

| | | | | | | | |
|---|---|---|---|---|---|---|---|
| ENSMUSG00000033653 | GSMG0014789 | Vps8 | VPS8 CORVET complex subunit | 844.16 | 1061.80 | -0.33 | 7.52E-02 |
| ENSMUSG00000097290 | GSMG0014790 | 1300002E11Rik | RIKEN cDNA 1300002E11 gene | 340.99 | 328.10 | 0.06 | 8.72E-01 |
| ENSMUSG00000022855 | GSMG0014792 | Senp2 | SUMO/sentrin specific peptidase 2 | 842.00 | 920.66 | -0.13 | 5.79E-01 |
| ENSMUSG00000004460 | GSMG0014796 | Dnajb11 | DnaJ heat shock protein family (Hsp40) member B11 | 2272.67 | 2683.36 | -0.24 | 2.37E-01 |
| ENSMUSG00000022885 | GSMG0014800 | St6gal1 | beta galactoside alpha 2,6 sialyltransferase 1 | 897.00 | 544.12 | 0.72 | 1.68E-05 |
| ENSMUSG00000022884 | GSMG0014802 | Eif4a2 | eukaryotic translation initiation factor 4A2 | 1638.10 | 1646.85 | -0.01 | 9.83E-01 |
| ENSMUSG00000033355 | GSMG0014818 | Rtp4 | receptor transporter protein 4 | 371.81 | 426.57 | -0.20 | 6.66E-01 |
| ENSMUSG00000033306 | GSMG0014821 | Lpp | LIM domain containing preferred translocation partner in lipoma | 1030.45 | 1023.73 | 0.01 | 9.78E-01 |
| ENSMUSG00000022514 | GSMG0014844 | Il1rap | interleukin 1 receptor accessory protein | 349.04 | 258.04 | 0.44 | 6.31E-02 |
| ENSMUSG00000038127 | GSMG0014847 | Ccdc50 | coiled-coil domain containing 50 | 501.38 | 556.51 | -0.15 | 5.76E-01 |
| NULL | GSMG0014848 | NULL | NULL | 108.62 | 119.82 | -0.15 | 7.59E-01 |
| NULL | GSMG0014851 | NULL | NULL | 709.04 | 506.09 | 0.49 | 1.37E-02 |
| ENSMUSG00000038084 | GSMG0014853 | Opa1 | optic atrophy 1 | 1197.87 | 1620.85 | -0.44 | 6.59E-03 |
| ENSMUSG00000022528 | GSMG0014858 | Hes1 | hairy and enhancer of split 1 (Drosophila) | 1652.13 | 1467.46 | 0.17 | 6.47E-01 |
| NULL | GSMG0014860 | NULL | NULL | 113.11 | 76.92 | 0.55 | 1.39E-01 |
| NULL | GSMG0014861 | NULL | NULL | 1679.69 | 1933.00 | -0.20 | 4.27E-01 |
| ENSMUSG00000046598 | GSMG0014863 | Bdh1 | 3-hydroxybutyrate dehydrogenase, type 1 | 214.21 | 383.24 | -0.84 | 4.15E-04 |
| ENSMUSG00000022770 | GSMG0014867 | Dlg1 | discs, large homolog 1 (Drosophila) | 1254.22 | 915.19 | 0.45 | 9.32E-03 |
| ENSMUSG00000022774 | GSMG0014875 | Ncbp2 | nuclear cap binding protein subunit 2 | 1274.12 | 1410.09 | -0.15 | 5.31E-01 |
| ENSMUSG00000035790 | GSMG0014876 | Cep19 | centrosomal protein 19 | 128.89 | 110.92 | 0.22 | 5.84E-01 |
| ENSMUSG00000022787 | GSMG0014878 | Wdr53 | WD repeat domain 53 | 239.95 | 237.94 | 0.02 | 9.70E-01 |
| ENSMUSG00000014074 | GSMG0014880 | Rnf168 | ring finger protein 168 | 886.16 | 991.97 | -0.16 | 5.73E-01 |
| ENSMUSG00000053774 | GSMG0014882 | Ubxn7 | UBX domain protein 7 | 231.79 | 189.95 | 0.29 | 3.40E-01 |
| ENSMUSG00000014075 | GSMG0014887 | Tctex1d2 | Tctex1 domain containing 2 | 192.07 | 220.67 | -0.20 | 5.81E-01 |
| ENSMUSG00000005615 | GSMG0014888 | Pcyt1a | phosphate cytidylyltransferase 1, choline, alpha isoform | 554.09 | 544.51 | 0.03 | 9.41E-01 |
| ENSMUSG00000022797 | GSMG0014892 | Tfrc | transferrin receptor | 5669.26 | 5519.93 | 0.04 | 8.81E-01 |
| ENSMUSG00000022791 | GSMG0014893 | Tnk2 | tyrosine kinase, non-receptor, 2 | 529.80 | 443.32 | 0.25 | 4.74E-01 |
| ENSMUSG00000022800 | GSMG0014896 | Fyttd1 | forty-two-three domain containing 1 | 728.73 | 793.35 | -0.12 | 6.90E-01 |
| ENSMUSG00000022801 | GSMG0014898 | Lrch3 | leucine-rich repeats and calponin homology (CH) domain containing 3 | 1142.89 | 1437.47 | -0.33 | 4.87E-02 |
| ENSMUSG00000022802 | GSMG0014902 | Lmln | leishmanolysin-like (metallopeptidase M8 family) | 193.34 | 234.05 | -0.28 | 4.12E-01 |
| ENSMUSG00000022807 | GSMG0014903 | Osbpl11 | oxysterol binding protein-like 11 | 1522.02 | 1599.64 | -0.07 | 7.88E-01 |
| ENSMUSG00000022808 | GSMG0014905 | Snx4 | sorting nexin 4 | 1558.52 | 1940.91 | -0.32 | 9.45E-02 |
| ENSMUSG00000022811 | GSMG0014907 | Zfp148 | zinc finger protein 148 | 726.92 | 769.28 | -0.08 | 7.69E-01 |

| | | | | | | | |
|---|---|---|---|---|---|---|---|
| ENSMUSG00000022817 | GSMG0014919 | Itgb5 | integrin beta 5 | 2304.70 | 1620.39 | 0.51 | 8.69E-03 |
| ENSMUSG00000022836 | GSMG0014927 | Mylk | myosin, light polypeptide kinase | 1080.64 | 830.66 | 0.38 | 2.08E-01 |
| ENSMUSG00000035376 | GSMG0014929 | Hacd2 | 3-hydroxyacyl-CoA dehydratase 2 | 240.33 | 236.22 | 0.03 | 9.46E-01 |
| ENSMUSG00000022840 | GSMG0014931 | Adcy5 | adenylate cyclase 5 | 758.57 | 565.43 | 0.42 | 2.05E-01 |
| ENSMUSG00000052133 | GSMG0014933 | Sema5b | sema domain, seven thrombospondin repeats (type 1 and type 1-like), tr | 260.32 | 290.93 | -0.16 | 7.05E-01 |
| ENSMUSG00000022849 | GSMG0014934 | Hspbap1 | Hspb associated protein 1 | 116.90 | 141.15 | -0.27 | 4.81E-01 |
| ENSMUSG00000022906 | GSMG0014937 | Parp9 | poly (ADP-ribose) polymerase family, member 9 | 606.61 | 685.73 | -0.18 | 5.37E-01 |
| ENSMUSG00000022905 | GSMG0014938 | Kpna1 | karyopherin (importin) alpha 1 | 1106.56 | 1155.59 | -0.06 | 8.19E-01 |
| ENSMUSG00000034379 | GSMG0014939 | Wdr5b | WD repeat domain 5B | 284.66 | 372.85 | -0.38 | 1.49E-01 |
| ENSMUSG00000075229 | GSMG0014940 | Ccdc58 | coiled-coil domain containing 58 | 164.86 | 225.26 | -0.45 | 1.38E-01 |
| ENSMUSG00000022837 | GSMG0014947 | Iqcb1 | IQ calmodulin-binding motif containing 1 | 366.65 | 400.15 | -0.13 | 6.82E-01 |
| ENSMUSG00000034243 | GSMG0014949 | Golgb1 | golgi autoantigen, golgin subfamily b, macrogolgin 1 | 671.73 | 653.72 | 0.04 | 8.99E-01 |
| ENSMUSG00000022827 | GSMG0014953 | Rabl3 | RAB, member RAS oncogene family-like 3 | 87.57 | 121.71 | -0.48 | 2.39E-01 |
| ENSMUSG00000022816 | GSMG0014957 | Fstl1 | follistatin-like 1 | 2372.58 | 2445.71 | -0.04 | 8.87E-01 |
| ENSMUSG00000034158 | GSMG0014959 | Lrrc58 | leucine rich repeat containing 58 | 17672.72 | 22447.14 | -0.34 | 2.09E-02 |
| ENSMUSG00000022812 | GSMG0014963 | Gsk3b | glycogen synthase kinase 3 beta | 1020.93 | 1254.99 | -0.30 | 2.17E-01 |
| ENSMUSG00000046516 | GSMG0014969 | Cox17 | cytochrome c oxidase assembly protein 17 | 664.67 | 545.96 | 0.28 | 4.19E-01 |
| ENSMUSG00000002845 | GSMG0014975 | Tmem39a | transmembrane protein 39a | 355.59 | 260.84 | 0.45 | 6.67E-02 |
| ENSMUSG00000022793 | GSMG0014976 | B4galt4 | UDP-Gal:betaGlcNAc beta 1,4-galactosyltransferase, polypeptide 4 | 852.01 | 492.36 | 0.79 | 2.58E-05 |
| ENSMUSG00000022790 | GSMG0014978 | Igsf11 | immunoglobulin superfamily, member 11 | 120.20 | 73.89 | 0.70 | 1.13E-01 |
| ENSMUSG00000061080 | GSMG0014979 | Lsamp | limbic system-associated membrane protein | 518.00 | 511.01 | 0.02 | 9.58E-01 |
| NULL | GSMG0014980 | NULL | NULL | 269.91 | 225.71 | 0.25 | 4.93E-01 |
| NULL | GSMG0014986 | NULL | NULL | 135.85 | 145.72 | -0.11 | 8.52E-01 |
| NULL | GSMG0014989 | NULL | NULL | 195.19 | 176.50 | 0.15 | 7.16E-01 |
| NULL | GSMG0014991 | NULL | NULL | 368.72 | 325.34 | 0.18 | 6.55E-01 |
| NULL | GSMG0014993 | NULL | NULL | 372.48 | 339.53 | 0.13 | 7.34E-01 |
| NULL | GSMG0014994 | NULL | NULL | 303.21 | 257.20 | 0.24 | 4.93E-01 |
| NULL | GSMG0014995 | NULL | NULL | 226.92 | 199.91 | 0.19 | 6.18E-01 |
| NULL | GSMG0015012 | NULL | NULL | 1656.84 | 1452.26 | 0.19 | 3.59E-01 |
| NULL | GSMG0015013 | NULL | NULL | 565.89 | 541.27 | 0.06 | 8.37E-01 |
| NULL | GSMG0015014 | NULL | NULL | 229.21 | 215.47 | 0.09 | 8.37E-01 |
| NULL | GSMG0015018 | NULL | NULL | 340.02 | 352.18 | -0.05 | 9.18E-01 |
| NULL | GSMG0015019 | NULL | NULL | 432.19 | 347.07 | 0.32 | 2.18E-01 |

| | | | | | | | |
|---|---|---|---|---|---|---|---|
| NULL | GSMG0015033 | NULL | NULL | 267.74 | 139.46 | 0.94 | 9.55E-04 |
| NULL | GSMG0015044 | NULL | NULL | 312.23 | 311.33 | 0.01 | 9.89E-01 |
| ENSMUSG00000022708 | GSMG0015045 | Zbtb20 | zinc finger and BTB domain containing 20 | 3801.27 | 2115.99 | 0.85 | 1.82E-08 |
| NULL | GSMG0015054 | NULL | NULL | 271.61 | 169.58 | 0.68 | 2.09E-02 |
| ENSMUSG00000076372 | GSMG0015055 | Mir568 | microRNA 568 | 1339.77 | 1020.43 | 0.39 | 4.97E-02 |
| ENSMUSG00000022698 | GSMG0015065 | Naa50 | N(alpha)-acetyltransferase 50, NatE catalytic subunit | 1460.95 | 1765.15 | -0.27 | 3.02E-01 |
| ENSMUSG00000043065 | GSMG0015069 | Spice1 | spindle and centriole associated protein 1 | 633.85 | 496.23 | 0.35 | 1.48E-01 |
| ENSMUSG00000036208 | GSMG0015072 | Nepro | nucleolus and neural progenitor protein | 295.45 | 342.84 | -0.21 | 4.35E-01 |
| ENSMUSG00000022665 | GSMG0015080 | Ccdc80 | coiled-coil domain containing 80 | 285.14 | 342.09 | -0.26 | 3.49E-01 |
| ENSMUSG00000022663 | GSMG0015083 | Atg3 | autophagy related 3 | 790.77 | 830.60 | -0.07 | 8.21E-01 |
| ENSMUSG00000033031 | GSMG0015099 | C330027C09Rik | RIKEN cDNA C330027C09 gene | 96.15 | 184.75 | -0.95 | 4.77E-04 |
| ENSMUSG00000032965 | GSMG0015101 | Ift57 | intraflagellar transport 57 | 484.26 | 501.05 | -0.05 | 8.89E-01 |
| ENSMUSG00000055447 | GSMG0015104 | Cd47 | CD47 antigen (Rh-related antigen, integrin-associated signal transducer | 1866.02 | 1973.95 | -0.08 | 7.52E-01 |
| NULL | GSMG0015105 | NULL | NULL | 244.08 | 186.07 | 0.39 | 2.96E-01 |
| ENSMUSG00000022637 | GSMG0015115 | Cblb | Casitas B-lineage lymphoma b | 286.62 | 401.86 | -0.49 | 2.83E-02 |
| ENSMUSG00000022601 | GSMG0015124 | Zbtb11 | zinc finger and BTB domain containing 11 | 419.15 | 469.32 | -0.16 | 5.18E-01 |
| ENSMUSG00000022752 | GSMG0015133 | Tomm70a | translocase of outer mitochondrial membrane 70 homolog A (yeast) | 1685.48 | 2236.86 | -0.41 | 7.53E-03 |
| ENSMUSG00000043336 | GSMG0015137 | Filip1l | filamin A interacting protein 1-like | 2466.49 | 1336.19 | 0.88 | 4.80E-07 |
| ENSMUSG00000035107 | GSMG0015139 | Dcbld2 | discoidin, CUB and LCCL domain containing 2 | 326.16 | 319.78 | 0.03 | 9.62E-01 |
| ENSMUSG00000022742 | GSMG0015140 | Cpox | coproporphyrinogen oxidase | 311.97 | 252.04 | 0.31 | 2.84E-01 |
| ENSMUSG00000022744 | GSMG0015141 | Cldnd1 | claudin domain containing 1 | 1078.34 | 1219.16 | -0.18 | 4.50E-01 |
| ENSMUSG00000022724 | GSMG0015145 | Mina | myc induced nuclear antigen | 240.25 | 343.40 | -0.51 | 2.09E-02 |
| NULL | GSMG0015147 | NULL | NULL | 259.12 | 302.92 | -0.23 | 8.19E-01 |
| ENSMUSG00000022912 | GSMG0015153 | Pros1 | protein S (alpha) | 3552.72 | 3348.34 | 0.09 | 7.33E-01 |
| ENSMUSG00000054604 | GSMG0015155 | Cggbp1 | CGG triplet repeat binding protein 1 | 4096.70 | 4451.31 | -0.12 | 6.50E-01 |
| ENSMUSG00000022707 | GSMG0015166 | Gbe1 | glucan (1,4-alpha-), branching enzyme 1 | 866.76 | 768.82 | 0.17 | 4.51E-01 |
| NULL | GSMG0015173 | NULL | NULL | 109.74 | 299.03 | -1.44 | 7.37E-08 |
| NULL | GSMG0015176 | NULL | NULL | 106.55 | 248.62 | -1.22 | 7.84E-07 |
| NULL | GSMG0015178 | NULL | NULL | 180.02 | 462.56 | -1.37 | 2.77E-06 |
| NULL | GSMG0015182 | NULL | NULL | 828.90 | 1146.85 | -0.47 | 5.45E-02 |
| ENSMUSG00000022883 | GSMG0015187 | Robo1 | roundabout guidance receptor 1 | 104.54 | 169.03 | -0.69 | 3.89E-02 |
| NULL | GSMG0015191 | NULL | NULL | 115.15 | 169.96 | -0.56 | 1.29E-01 |
| NULL | GSMG0015193 | NULL | NULL | 169.99 | 315.95 | -0.89 | 1.19E-03 |

| | | | | | | | |
|---|---|---|---|---|---|---|---|
| ENSMUSG00000022867 | GSMG0015204 | Usp25 | ubiquitin specific peptidase 25 | 1761.49 | 1564.73 | 0.17 | 4.56E-01 |
| ENSMUSG00000090386 | GSMG0015206 | Mir99ahg | Mir99a and Mirlet7c-1 host gene (non-protein coding) | 924.40 | 1177.85 | -0.35 | 1.03E-01 |
| NULL | GSMG0015217 | NULL | NULL | 471.30 | 497.06 | -0.08 | 8.17E-01 |
| ENSMUSG00000022865 | GSMG0015224 | Cxadr | coxsackie virus and adenovirus receptor | 138.57 | 583.18 | -2.06 | 3.14E-22 |
| ENSMUSG00000022762 | GSMG0015237 | Ncam2 | neural cell adhesion molecule 2 | 577.57 | 607.77 | -0.07 | 8.24E-01 |
| NULL | GSMG0015240 | NULL | NULL | 278.89 | 246.26 | 0.18 | 5.99E-01 |
| ENSMUSG00000053062 | GSMG0015251 | Jam2 | junction adhesion molecule 2 | 2081.97 | 1852.30 | 0.17 | 4.25E-01 |
| ENSMUSG00000008976 | GSMG0015252 | Gabpa | GA repeat binding protein, alpha | 764.84 | 932.29 | -0.29 | 1.32E-01 |
| ENSMUSG00000044442 | GSMG0015258 | N6amt1 | N-6 adenine-specific DNA methyltransferase 1 (putative) | 394.72 | 553.93 | -0.49 | 1.78E-02 |
| ENSMUSG00000025616 | GSMG0015260 | Usp16 | ubiquitin specific peptidase 16 | 1022.69 | 1036.19 | -0.02 | 9.59E-01 |
| NULL | GSMG0015267 | NULL | NULL | 78.31 | 112.24 | -0.52 | 2.00E-01 |
| ENSMUSG00000022982 | GSMG0015282 | Sod1 | superoxide dismutase 1, soluble | 1388.80 | 1675.34 | -0.27 | 1.32E-01 |
| ENSMUSG00000053414 | GSMG0015283 | Hunk | hormonally upregulated Neu-associated kinase | 227.09 | 269.69 | -0.25 | 4.38E-01 |
| ENSMUSG00000039830 | GSMG0015294 | Olig2 | oligodendrocyte transcription factor 2 | 338.70 | 561.36 | -0.73 | 2.54E-04 |
| ENSMUSG00000046160 | GSMG0015295 | Olig1 | oligodendrocyte transcription factor 1 | 1522.87 | 2251.97 | -0.56 | 2.65E-03 |
| ENSMUSG00000022971 | GSMG0015296 | Ifnar2 | interferon (alpha and beta) receptor 2 | 2675.88 | 2169.77 | 0.30 | 8.76E-02 |
| ENSMUSG00000022969 | GSMG0015297 | Il10rb | interleukin 10 receptor, beta | 1782.26 | 1136.27 | 0.65 | 3.74E-04 |
| ENSMUSG00000084866 | GSMG0015299 | A930006K02Rik | RIKEN cDNA A930006K02 gene | 103.60 | 109.61 | -0.09 | 8.93E-01 |
| ENSMUSG00000022967 | GSMG0015300 | Ifnar1 | interferon (alpha and beta) receptor 1 | 4547.59 | 3442.88 | 0.40 | 1.50E-02 |
| ENSMUSG00000022965 | GSMG0015301 | Ifngr2 | interferon gamma receptor 2 | 1549.55 | 2234.38 | -0.53 | 9.83E-04 |
| NULL | GSMG0015302 | NULL | NULL | 118.73 | 105.11 | 0.18 | 6.86E-01 |
| ENSMUSG00000022961 | GSMG0015304 | Son | Son DNA binding protein | 10514.04 | 9908.44 | 0.09 | 7.70E-01 |
| G00000039680,ENSMUSG000 | GSMG0015311 | Slc5a3 // Mrps6 | solute carrier family 5 (inositol transporters), member 3 // mitochondria | 983.85 | 1210.11 | -0.30 | 1.82E-01 |
| ENSMUSG00000051989 | GSMG0015317 | Smim11 | small integral membrane protein 11 | 557.38 | 579.36 | -0.06 | 8.84E-01 |
| ENSMUSG00000022949 | GSMG0015319 | Clic6 | chloride intracellular channel 6 | 212.50 | 462.34 | -1.12 | 5.34E-05 |
| ENSMUSG00000051483 | GSMG0015327 | Cbr1 | carbonyl reductase 1 | 972.65 | 978.56 | -0.01 | 9.79E-01 |
| ENSMUSG00000022947 | GSMG0015328 | Cbr3 | carbonyl reductase 3 | 139.09 | 179.02 | -0.36 | 2.47E-01 |
| ENSMUSG00000022946 | GSMG0015329 | Dopey2 | dopey family member 2 | 1538.97 | 1982.15 | -0.36 | 1.32E-01 |
| ENSMUSG00000039456 | GSMG0015330 | Morc3 | microrchidia 3 | 942.84 | 978.16 | -0.05 | 8.61E-01 |
| ENSMUSG00000022945 | GSMG0015331 | Chaf1b | chromatin assembly factor 1, subunit B (p60) | 104.64 | 241.93 | -1.21 | 2.10E-06 |
| ENSMUSG00000040785 | GSMG0015335 | Ttc3 | tetratricopeptide repeat domain 3 | 3119.65 | 4329.39 | -0.47 | 1.02E-02 |
| NULL | GSMG0015337 | NULL | NULL | 223.13 | 177.28 | 0.34 | 2.51E-01 |
| ENSMUSG00000022897 | GSMG0015339 | Dyrk1a | dual-specificity tyrosine-(Y)-phosphorylation regulated kinase 1a | 216.83 | 164.53 | 0.40 | 3.10E-01 |

| | | | | | | | |
|---|---|---|---|---|---|---|---|
| ENSMUSG00000022895 | GSMG0015343 | Ets2 | E26 avian leukemia oncogene 2, 3' domain | 1038.71 | 1128.59 | -0.12 | 6.22E-01 |
| ENSMUSG00000023147 | GSMG0015348 | Wrb | tryptophan rich basic protein | 1829.10 | 3134.12 | -0.78 | 4.85E-07 |
| G00000000159,ENSMUSG000 | GSMG0015351 | Igsf5 // Pcp4 | immunoglobulin superfamily, member 5 // Purkinje cell protein 4 | 469.32 | 500.36 | -0.09 | 7.51E-01 |
| ENSMUSG00000040605 | GSMG0015356 | Bace2 | beta-site APP-cleaving enzyme 2 | 186.35 | 167.45 | 0.16 | 6.79E-01 |
| ENSMUSG00000080717 | GSMG0015360 | B230307C23Rik | RIKEN cDNA B230307C23 gene | 173.64 | 132.21 | 0.39 | 2.09E-01 |
| ENSMUSG00000005983 | GSMG0015366 | 1700037C18Rik // | RIKEN cDNA 1700037C18 gene // predicted gene, 41410 | 49.98 | 60.11 | -0.26 | 6.67E-01 |
| ENSMUSG00000049871 | GSMG0015368 | Nlrc3 | NLR family, CARD domain containing 3 | 335.25 | 453.11 | -0.43 | 4.89E-02 |
| ENSMUSG00000039738 | GSMG0015369 | Slx4 | SLX4 structure-specific endonuclease subunit homolog (S. cerevisiae) | 481.01 | 620.83 | -0.37 | 1.12E-01 |
| ENSMUSG00000005981 | GSMG0015370 | Trap1 | TNF receptor-associated protein 1 | 652.00 | 751.40 | -0.20 | 4.76E-01 |
| NULL | GSMG0015371 | NULL | NULL | 871.50 | 972.77 | -0.16 | 5.67E-01 |
| ENSMUSG00000022521 | GSMG0015372 | Crebbp | CREB binding protein | 2221.18 | 1859.77 | 0.26 | 2.73E-01 |
| ENSMUSG00000005580 | GSMG0015378 | Adcy9 | adenylate cyclase 9 | 143.15 | 96.79 | 0.56 | 2.05E-01 |
| NULL | GSMG0015384 | NULL | NULL | 136.30 | 159.99 | -0.23 | 5.10E-01 |
| ENSMUSG00000039637 | GSMG0015386 | Coro7 | coronin 7 | 532.67 | 535.38 | -0.01 | 9.89E-01 |
| ENSMUSG00000063445 | GSMG0015388 | Nmral1 | NmrA-like family domain containing 1 | 114.12 | 247.66 | -1.11 | 8.76E-06 |
| ENSMUSG00000004071 | GSMG0015391 | Cdip1 | cell death inducing Trp53 target 1 | 978.45 | 988.74 | -0.01 | 9.65E-01 |
| ENSMUSG00000039568 | GSMG0015396 | Ubald1 | UBA-like domain containing 1 | 745.34 | 730.32 | 0.03 | 9.41E-01 |
| NULL | GSMG0015398 | NULL | NULL | 519.82 | 425.37 | 0.29 | 3.55E-01 |
| ENSMUSG00000022515 | GSMG0015400 | Anks3 | ankyrin repeat and sterile alpha motif domain containing 3 | 1054.44 | 1036.71 | 0.03 | 9.36E-01 |
| ENSMUSG00000022540 | GSMG0015402 | Rogdi | rogdi homolog | 2466.90 | 1248.50 | 0.98 | 5.26E-09 |
| ENSMUSG00000023143 | GSMG0015406 | Nagpa | N-acetylglucosamine-1-phosphodiester alpha-N-acetylglucosaminidase | 764.43 | 655.76 | 0.22 | 3.44E-01 |
| ENSMUSG00000051669 | GSMG0015407 | AU021092 | expressed sequence AU021092 | 4057.95 | 5361.68 | -0.40 | 2.53E-02 |
| ENSMUSG00000022544 | GSMG0015408 | Eef2kmt | eukaryotic elongation factor 2 lysine methyltransferase | 366.80 | 294.86 | 0.32 | 2.13E-01 |
| ENSMUSG00000043140 | GSMG0015415 | Tmem186 | transmembrane protein 186 | 779.07 | 799.38 | -0.04 | 9.09E-01 |
| NULL | GSMG0015416 | NULL | NULL | 455.91 | 395.63 | 0.21 | 4.13E-01 |
| ENSMUSG00000008393 | GSMG0015417 | Carhsp1 | calcium regulated heat stable protein 1 | 2339.02 | 2611.26 | -0.16 | 4.40E-01 |
| ENSMUSG00000022710 | GSMG0015418 | Usp7 | ubiquitin specific peptidase 7 | 1985.11 | 1370.29 | 0.53 | 5.08E-02 |
| NULL | GSMG0015419 | NULL | NULL | 558.03 | 288.73 | 0.95 | 2.87E-06 |
| NULL | GSMG0015424 | NULL | NULL | 649.73 | 316.08 | 1.04 | 3.51E-07 |
| ENSMUSG00000022505 | GSMG0015426 | Emp2 | epithelial membrane protein 2 | 5221.74 | 2956.57 | 0.82 | 1.54E-08 |
| ENSMUSG00000050908 | GSMG0015429 | Tvp23a | trans-golgi network vesicle protein 23A | 56.95 | 109.87 | -0.94 | 4.18E-03 |
| ENSMUSG00000038055 | GSMG0015431 | Dexi | dexamethasone-induced transcript | 891.48 | 702.38 | 0.34 | 5.55E-02 |
| ENSMUSG00000022500 | GSMG0015439 | Litaf | LPS-induced TN factor | 2271.07 | 2869.98 | -0.34 | 8.23E-02 |

| | | | | | | | |
|---|---|---|---|---|---|---|---|
| ENSMUSG00000022498 | GSMG0015444 | Txndc11 | thioredoxin domain containing 11 | 1413.20 | 1411.84 | 0.00 | 9.97E-01 |
| ENSMUSG00000037965 | GSMG0015445 | Zc3h7a | zinc finger CCCH type containing 7 A | 2920.81 | 2880.52 | 0.02 | 9.47E-01 |
| ENSMUSG00000005846 | GSMG0015446 | Rsl1d1 | ribosomal L1 domain containing 1 | 1129.78 | 1570.57 | -0.48 | 3.94E-02 |
| ENSMUSG00000062203 | GSMG0015448 | Gspt1 | G1 to S phase transition 1 | 1528.60 | 1983.56 | -0.38 | 3.36E-02 |
| ENSMUSG00000065979 | GSMG0015450 | Cpped1 | calcineurin-like phosphoesterase domain containing 1 | 256.39 | 243.32 | 0.08 | 8.52E-01 |
| ENSMUSG00000022685 | GSMG0015458 | Parn | poly(A)-specific ribonuclease (deadenylation nuclease) | 453.69 | 446.42 | 0.02 | 9.54E-01 |
| ENSMUSG00000022680 | GSMG0015463 | Pdxdc1 | pyridoxal-dependent decarboxylase domain containing 1 | 877.84 | 848.77 | 0.05 | 8.78E-01 |
| ENSMUSG00000022677 | GSMG0015468 | Fopnl | Fgfr1op N-terminal like | 1231.12 | 1470.05 | -0.26 | 2.38E-01 |
| ENSMUSG00000018830 | GSMG0015475 | Myh11 | myosin, heavy polypeptide 11, smooth muscle | 283.50 | 46.81 | 2.60 | 1.10E-03 |
| ENSMUSG00000022673 | GSMG0015478 | Mcm4 | minichromosome maintenance complex component 4 | 753.75 | 1571.96 | -1.06 | 5.00E-07 |
| ENSMUSG00000022671 | GSMG0015479 | Mzt2 | mitotic spindle organizing protein 2 | 501.69 | 519.17 | -0.05 | 8.77E-01 |
| ENSMUSG00000041974 | GSMG0015480 | Spidr | scaffolding protein involved in DNA repair | 179.12 | 216.13 | -0.27 | 4.50E-01 |
| ENSMUSG00000022789 | GSMG0015483 | Dnm1l | dynamin 1-like | 3616.60 | 3713.43 | -0.04 | 8.96E-01 |
| ENSMUSG00000022788 | GSMG0015484 | Fgd4 | FYVE, RhoGEF and PH domain containing 4 | 198.21 | 203.82 | -0.03 | 9.41E-01 |
| ENSMUSG00000022771 | GSMG0015494 | Ppil2 | peptidylprolyl isomerase (cyclophilin)-like 2 | 2354.96 | 2459.32 | -0.06 | 7.94E-01 |
| ENSMUSG00000022769 | GSMG0015496 | Sdf2l1 | stromal cell-derived factor 2-like 1 | 443.58 | 446.47 | -0.01 | 9.82E-01 |
| ENSMUSG00000038965 | GSMG0015498 | Ube2l3 | ubiquitin-conjugating enzyme E2L 3 | 2403.41 | 2951.82 | -0.30 | 1.65E-01 |
| ENSMUSG00000041720 | GSMG0015501 | Pi4ka | phosphatidylinositol 4-kinase, catalytic, alpha polypeptide | 1426.31 | 1186.32 | 0.27 | 2.21E-01 |
| NULL | GSMG0015503 | NULL | NULL | 251.45 | 266.55 | -0.08 | 8.30E-01 |
| ENSMUSG00000022760 | GSMG0015504 | Thap7 | THAP domain containing 7 | 704.17 | 666.68 | 0.08 | 8.11E-01 |
| ENSMUSG00000022756 | GSMG0015506 | Slc7a4 | solute carrier family 7 (cationic amino acid transporter, y+ system), mer | 1173.87 | 979.02 | 0.26 | 2.65E-01 |
| ENSMUSG00000012114 | GSMG0015507 | Med15 | mediator complex subunit 15 | 1269.32 | 1371.01 | -0.11 | 6.19E-01 |
| G00000003166,ENSMUSG000 | GSMG0015510 | Car15 // Dgcr2 | carbonic anhydrase 15 // DiGeorge syndrome critical region gene 2 | 1636.30 | 1898.53 | -0.21 | 3.95E-01 |
| ENSMUSG00000003527 | GSMG0015511 | Dgcr14 | DiGeorge syndrome critical region gene 14 | 1268.44 | 1434.85 | -0.18 | 3.65E-01 |
| ENSMUSG00000003528 | GSMG0015513 | Slc25a1 | solute carrier family 25 (mitochondrial carrier, citrate transporter), mem | 1414.72 | 1391.29 | 0.02 | 9.33E-01 |
| ENSMUSG00000003526 | GSMG0015514 | Prodh | proline dehydrogenase | 544.19 | 285.16 | 0.93 | 6.96E-05 |
| ENSMUSG00000060166 | GSMG0015516 | Zdhhc8 | zinc finger, DHHC domain containing 8 | 1461.53 | 1199.47 | 0.28 | 3.27E-01 |
| ENSMUSG00000005732 | GSMG0015517 | Ranbp1 | RAN binding protein 1 | 2775.19 | 4648.15 | -0.74 | 3.06E-04 |
| ENSMUSG00000022718 | GSMG0015518 | Dgcr8 | DiGeorge syndrome critical region gene 8 | 496.93 | 555.61 | -0.16 | 5.58E-01 |
| ENSMUSG00000013539 | GSMG0015519 | Tango2 | transport and golgi organization 2 | 517.86 | 421.03 | 0.30 | 2.36E-01 |
| NULL | GSMG0015520 | NULL | NULL | 373.08 | 193.27 | 0.95 | 8.21E-06 |
| ENSMUSG00000009097 | GSMG0015522 | Tbx1 | T-box 1 | 1215.29 | 850.52 | 0.51 | 3.21E-03 |
| ENSMUSG00000000028 | GSMG0015524 | Cdc45 | cell division cycle 45 | 188.30 | 414.49 | -1.13 | 4.68E-06 |

| | | | | | | | |
|---|---|---|---|---|---|---|---|
| ENSMUSG00000022706 | GSMG0015526 | Mrpl40 | mitochondrial ribosomal protein L40 | 631.26 | 793.65 | -0.33 | 9.65E-02 |
| ENSMUSG00000043008 | GSMG0015538 | Klhl6 | kelch-like 6 | 1153.58 | 1477.64 | -0.36 | 9.54E-02 |
| ENSMUSG00000041205 | GSMG0015539 | Map6d1 | MAP6 domain containing 1 | 470.52 | 129.63 | 1.86 | 5.02E-22 |
| ENSMUSG00000033918 | GSMG0015540 | Parl | presenilin associated, rhomboid-like | 666.40 | 820.12 | -0.30 | 1.30E-01 |
| ENSMUSG00000022822 | GSMG0015543 | Abcc5 | ATP-binding cassette, sub-family C (CFTR/MRP), member 5 | 431.97 | 370.75 | 0.22 | 4.64E-01 |
| ENSMUSG00000033809 | GSMG0015547 | Alg3 | asparagine-linked glycosylation 3 (alpha-1,3-mannosyltransferase) | 583.07 | 837.64 | -0.52 | 3.69E-03 |
| ENSMUSG00000051146 | GSMG0015548 | Camk2n2 | calcium/calmodulin-dependent protein kinase II inhibitor 2 | 385.71 | 522.55 | -0.44 | 1.36E-01 |
| NULL | GSMG0015550 | NULL | NULL | 69.23 | 89.30 | -0.38 | 4.48E-01 |
| ENSMUSG00000022843 | GSMG0015551 | Clcn2 | chloride channel, voltage-sensitive 2 | 3046.56 | 2172.03 | 0.49 | 1.68E-02 |
| NULL | GSMG0015556 | Magefl | melanoma antigen family F, 1 | 437.85 | 436.39 | 0.00 | 9.89E-01 |
| ENSMUSG00000043391 | GSMG0015558 | 2510009E07Rik | RIKEN cDNA 2510009E07 gene | 3641.93 | 3833.09 | -0.07 | 7.58E-01 |
| ENSMUSG00000022856 | GSMG0015564 | Tmem41a | transmembrane protein 41a | 565.85 | 697.31 | -0.30 | 1.51E-01 |
| ENSMUSG00000013089 | GSMG0015569 | Etv5 | ets variant 5 | 304.17 | 324.94 | -0.09 | 7.67E-01 |
| ENSMUSG00000022861 | GSMG0015570 | Dgkg | diacylglycerol kinase, gamma | 655.73 | 319.98 | 1.04 | 1.32E-06 |
| ENSMUSG00000004462 | GSMG0015573 | Tbccd1 | TBCC domain containing 1 | 278.01 | 289.40 | -0.05 | 8.92E-01 |
| ENSMUSG00000022881 | GSMG0015576 | Rfc4 | replication factor C (activator 1) 4 | 224.70 | 516.86 | -1.20 | 1.02E-09 |
| ENSMUSG00000022887 | GSMG0015579 | Masp1 | mannan-binding lectin serine peptidase 1 | 55.04 | 145.15 | -1.39 | 1.42E-05 |
| ENSMUSG00000004366 | GSMG0015581 | Sst | somatostatin | 1548.11 | 1284.46 | 0.27 | 3.59E-01 |
| ENSMUSG00000038168 | GSMG0015593 | P3h2 | prolyl 3-hydroxylase 2 | 107.70 | 57.01 | 0.92 | 6.86E-03 |
| ENSMUSG00000022523 | GSMG0015604 | Fgf12 | fibroblast growth factor 12 | 755.07 | 780.43 | -0.05 | 8.82E-01 |
| NULL | GSMG0015609 | NULL | NULL | 398.97 | 274.81 | 0.54 | 2.96E-02 |
| NULL | GSMG0015610 | NULL | NULL | 213.06 | 213.44 | 0.00 | 9.98E-01 |
| NULL | GSMG0015612 | NULL | NULL | 663.53 | 477.06 | 0.48 | 3.00E-02 |
| NULL | GSMG0015613 | NULL | NULL | 820.47 | 822.72 | 0.00 | 9.92E-01 |
| ENSMUSG00000051065 | GSMG0015614 | Mb21d2 | Mab-21 domain containing 2 | 399.32 | 498.81 | -0.32 | 1.75E-01 |
| ENSMUSG00000048939 | GSMG0015617 | Atp13a5 | ATPase type 13A5 | 10642.94 | 7696.09 | 0.47 | 3.39E-03 |
| ENSMUSG00000038094 | GSMG0015618 | Atp13a4 | ATPase type 13A4 | 75.76 | 32.27 | 1.23 | 3.87E-03 |
| ENSMUSG00000022537 | GSMG0015636 | Tmem44 | transmembrane protein 44 | 3042.03 | 2440.21 | 0.32 | 4.67E-02 |
| NULL | GSMG0015637 | NULL | NULL | 172.30 | 145.00 | 0.25 | 5.11E-01 |
| ENSMUSG00000022538 | GSMG0015638 | Lsg1 | large 60S subunit nuclear export GTPase 1 | 768.85 | 987.18 | -0.36 | 7.92E-02 |
| ENSMUSG00000047434 | GSMG0015640 | Xxylt1 | xyloside xylosyltransferase 1 | 1822.92 | 1578.65 | 0.21 | 3.18E-01 |
| ENSMUSG00000049076 | GSMG0015641 | Acap2 | ArfGAP with coiled-coil, ankyrin repeat and PH domains 2 | 898.33 | 907.97 | -0.02 | 9.58E-01 |
| NULL | GSMG0015643 | NULL | NULL | 53.82 | 35.53 | 0.59 | 3.40E-01 |

| | | | | | | | |
|---|---|---|---|---|---|---|---|
| ENSMUSG00000022548 | GSMG0015646 | Apod | apolipoprotein D | 7395.34 | 6814.46 | 0.12 | 5.92E-01 |
| ENSMUSG00000107002 | GSMG0015653 | 0610012G03Rik | RIKEN cDNA 0610012G03 gene | 1777.97 | 1957.63 | -0.14 | 5.25E-01 |
| ENSMUSG00000022772 | GSMG0015655 | Senp5 | SUMO/sentrin specific peptidase 5 | 831.16 | 910.11 | -0.13 | 6.02E-01 |
| ENSMUSG00000022781 | GSMG0015656 | Pak2 | p21 protein (Cdc42/Rac)-activated kinase 2 | 485.63 | 604.71 | -0.32 | 2.26E-01 |
| ENSMUSG00000023791 | GSMG0015657 | Pigx | phosphatidylinositol glycan anchor biosynthesis, class X | 397.85 | 545.26 | -0.46 | 4.72E-02 |
| ENSMUSG00000052384 | GSMG0015659 | Nrros | negative regulator of reactive oxygen species | 2230.62 | 3242.88 | -0.54 | 3.28E-03 |
| ENSMUSG00000035764 | GSMG0015660 | Fbxo45 | F-box protein 45 | 700.20 | 771.73 | -0.14 | 6.18E-01 |
| ENSMUSG00000035629 | GSMG0015668 | Rubcn | RUN domain and cysteine-rich domain containing, Beclin 1-interacting | 810.29 | 1011.68 | -0.32 | 1.74E-01 |
| ENSMUSG00000035578 | GSMG0015670 | Iqcg | IQ motif containing G | 31.42 | 89.43 | -1.50 | 1.12E-04 |
| NULL | GSMG0015674 | NULL | NULL | 140.83 | 80.63 | 0.80 | 7.45E-02 |
| ENSMUSG00000022814 | GSMG0015675 | Umps | uridine monophosphate synthetase | 300.97 | 508.09 | -0.75 | 6.08E-05 |
| NULL | GSMG0015677 | NULL | NULL | 275.77 | 253.57 | 0.12 | 7.62E-01 |
| NULL | GSMG0015678 | NULL | NULL | 286.54 | 275.54 | 0.06 | 8.82E-01 |
| NULL | GSMG0015681 | NULL | NULL | 92.67 | 71.48 | 0.37 | 4.07E-01 |
| NULL | GSMG0015682 | NULL | NULL | 431.81 | 291.54 | 0.57 | 3.13E-02 |
| NULL | GSMG0015684 | NULL | NULL | 198.30 | 92.41 | 1.11 | 2.72E-05 |
| NULL | GSMG0015686 | NULL | NULL | 403.80 | 304.60 | 0.41 | 1.29E-01 |
| NULL | GSMG0015687 | NULL | NULL | 629.61 | 477.46 | 0.40 | 6.77E-02 |
| ENSMUSG00000034473 | GSMG0015695 | Sec22a | SEC22 homolog A, vesicle trafficking protein | 399.27 | 469.38 | -0.23 | 3.89E-01 |
| ENSMUSG00000022844 | GSMG0015698 | Pdia5 | protein disulfide isomerase associated 5 | 464.01 | 389.63 | 0.25 | 3.66E-01 |
| ENSMUSG00000022848 | GSMG0015701 | Dirc2 | disrupted in renal carcinoma 2 (human) | 1486.47 | 1284.77 | 0.21 | 3.60E-01 |
| ENSMUSG00000049502 | GSMG0015704 | Dtx3l | deltex 3-like, E3 ubiquitin ligase | 489.86 | 574.18 | -0.23 | 3.40E-01 |
| NULL | GSMG0015705 | NULL | NULL | 31770.35 | 36569.31 | -0.20 | 4.70E-01 |
| ENSMUSG00000003955 | GSMG0015706 | Fam162a | family with sequence similarity 162, member A | 957.79 | 930.04 | 0.04 | 8.97E-01 |
| ENSMUSG00000022899 | GSMG0015717 | Slc15a2 | solute carrier family 15 (H+/peptide transporter), member 2 | 92.33 | 146.57 | -0.65 | 7.68E-02 |
| ENSMUSG00000022829 | GSMG0015722 | Stxbp5l | syntaxin binding protein 5-like | 758.70 | 446.35 | 0.77 | 4.11E-05 |
| ENSMUSG00000022828 | GSMG0015723 | Gtf2e1 | general transcription factor II E, polypeptide 1 (alpha subunit) | 165.31 | 191.47 | -0.21 | 5.27E-01 |
| ENSMUSG00000002847 | GSMG0015728 | Pla1a | phospholipase A1 member A | 496.68 | 274.64 | 0.86 | 9.47E-06 |
| ENSMUSG00000002844 | GSMG0015729 | Adprh | ADP-ribosylarginine hydrolase | 1111.56 | 1453.69 | -0.39 | 2.73E-02 |
| ENSMUSG00000002846 | GSMG0015730 | Timmdc1 | translocase of inner mitochondrial membrane domain containing 1 | 262.90 | 232.03 | 0.18 | 5.63E-01 |
| ENSMUSG00000034064 | GSMG0015731 | Poglut1 | protein O-glucosyltransferase 1 | 764.47 | 679.59 | 0.17 | 5.40E-01 |
| NULL | GSMG0015734 | NULL | NULL | 133.78 | 89.02 | 0.58 | 8.77E-02 |
| ENSMUSG00000022799 | GSMG0015736 | Arhgap31 | Rho GTPase activating protein 31 | 1521.13 | 1497.58 | 0.02 | 9.48E-01 |

| | | | | | | | |
|---|---|---|---|---|---|---|---|
| ENSMUSG00000047261 | GSMG0015740 | Gap43 | growth associated protein 43 | 823.40 | 4101.80 | -2.32 | 1.32E-55 |
| NULL | GSMG0015741 | NULL | NULL | 87.36 | 207.00 | -1.24 | 9.51E-07 |
| ENSMUSG00000052459 | GSMG0015749 | Atp6v1a | ATPase, H+ transporting, lysosomal V1 subunit A | 1137.60 | 1181.02 | -0.05 | 8.63E-01 |
| ENSMUSG00000022704 | GSMG0015751 | Qtrtd1 | queuine tRNA-ribosyltransferase domain containing 1 | 154.77 | 193.52 | -0.32 | 3.20E-01 |
| ENSMUSG00000036304 | GSMG0015752 | Zdhhc23 | zinc finger, DHHC domain containing 23 | 126.96 | 121.48 | 0.06 | 9.11E-01 |
| ENSMUSG00000022696 | GSMG0015754 | Sidt1 | SID1 transmembrane family, member 1 | 375.15 | 298.25 | 0.33 | 1.76E-01 |
| ENSMUSG00000022687 | GSMG0015756 | Boc | biregional cell adhesion molecule-related/down-regulated by oncogenes | 272.25 | 200.84 | 0.44 | 1.14E-01 |
| ENSMUSG00000022668 | GSMG0015758 | Gtpbp8 | GTP-binding protein 8 (putative) | 163.89 | 193.77 | -0.25 | 4.87E-01 |
| NULL | GSMG0015759 | NULL | NULL | 460.66 | 473.60 | -0.04 | 9.00E-01 |
| ENSMUSG00000022664 | GSMG0015760 | Slc35a5 | solute carrier family 35, member A5 | 321.18 | 381.76 | -0.25 | 3.46E-01 |
| ENSMUSG00000022661 | GSMG0015762 | Cd200 | CD200 antigen | 1406.03 | 2041.51 | -0.54 | 3.24E-03 |
| ENSMUSG00000022658 | GSMG0015767 | Tagln3 | transgelin 3 | 1410.06 | 1295.98 | 0.12 | 5.98E-01 |
| ENSMUSG00000033157 | GSMG0015768 | Abhd10 | abhydrolase domain containing 10 | 344.54 | 398.45 | -0.21 | 4.13E-01 |
| ENSMUSG00000033149 | GSMG0015769 | Phldb2 | pleckstrin homology like domain, family B, member 2 | 2720.45 | 2571.07 | 0.08 | 7.40E-01 |
| ENSMUSG00000087141 | GSMG0015773 | Plcxd2 | phosphatidylinositol-specific phospholipase C, X domain containing 2 | 25.62 | 69.39 | -1.43 | 7.70E-04 |
| ENSMUSG00000022656 | GSMG0015783 | Nectin3 | nectin cell adhesion molecule 3 | 645.39 | 764.57 | -0.24 | 3.78E-01 |
| NULL | GSMG0015784 | NULL | NULL | 134.55 | 195.07 | -0.54 | 1.00E-01 |
| ENSMUSG00000064061 | GSMG0015789 | Dzip3 | DAZ interacting protein 3, zinc finger | 533.21 | 605.72 | -0.18 | 4.73E-01 |
| ENSMUSG00000022641 | GSMG0015799 | Bbx | bobby sox homolog (Drosophila) | 417.98 | 400.10 | 0.07 | 8.48E-01 |
| ENSMUSG00000022636 | GSMG0015810 | Alcam | activated leukocyte cell adhesion molecule | 439.37 | 649.60 | -0.56 | 3.13E-02 |
| NULL | GSMG0015816 | NULL | NULL | 256.31 | 238.17 | 0.10 | 7.90E-01 |
| ENSMUSG00000035356 | GSMG0015822 | Nfkbiz | nuclear factor of kappa light polypeptide gene enhancer in B cells inhib | 411.94 | 243.86 | 0.76 | 6.52E-04 |
| ENSMUSG00000102101 | GSMG0015825 | Zbtb11os1 | zinc finger and BTB domain containing 11, opposite strand 1 | 187.61 | 181.51 | 0.04 | 9.25E-01 |
| ENSMUSG00000044763 | GSMG0015827 | Trmt10c | tRNA methyltransferase 10C | 392.33 | 454.13 | -0.21 | 4.58E-01 |
| ENSMUSG00000022757 | GSMG0015829 | Tfg | Trk-fused gene | 1032.14 | 1328.19 | -0.36 | 3.83E-02 |
| ENSMUSG00000022754 | GSMG0015831 | Tmem45a | transmembrane protein 45a | 238.59 | 200.39 | 0.25 | 4.66E-01 |
| ENSMUSG00000022751 | GSMG0015834 | Nit2 | nitrilase family, member 2 | 452.94 | 549.56 | -0.28 | 2.46E-01 |
| ENSMUSG00000022749 | GSMG0015835 | Tbc1d23 | TBC1 domain family, member 23 | 451.65 | 465.05 | -0.04 | 9.13E-01 |
| ENSMUSG00000022748 | GSMG0015838 | Cmss1 | cms small ribosomal subunit 1 | 134.46 | 189.73 | -0.50 | 1.40E-01 |
| ENSMUSG00000068196 | GSMG0015844 | Col8a1 | collagen, type VIII, alpha 1 | 1034.70 | 315.58 | 1.71 | 1.14E-15 |
| ENSMUSG00000022747 | GSMG0015847 | St3gal6 | ST3 beta-galactoside alpha-2,3-sialyltransferase 6 | 5283.74 | 3066.88 | 0.78 | 1.61E-06 |
| NULL | GSMG0015848 | NULL | NULL | 244.82 | 153.55 | 0.67 | 1.59E-02 |
| ENSMUSG00000022723 | GSMG0015868 | Crybg3 | beta-gamma crystallin domain containing 3 | 282.99 | 226.19 | 0.33 | 2.67E-01 |

| | | | | | | | |
|---|---|---|---|---|---|---|---|
| ENSMUSG00000022722 | GSMG0015869 | Arl6 | ADP-ribosylation factor-like 6 | 269.49 | 291.10 | -0.11 | 7.49E-01 |
| ENSMUSG00000055540 | GSMG0015872 | Epha6 | Eph receptor A6 | 513.14 | 412.03 | 0.32 | 3.50E-01 |
| NULL | GSMG0015879 | NULL | NULL | 153.27 | 73.13 | 1.06 | 7.39E-04 |
| ENSMUSG00000050312 | GSMG0015881 | Nsun3 | NOL1/NOP2/Sun domain family member 3 | 67.69 | 70.39 | -0.05 | 9.40E-01 |
| ENSMUSG00000022911 | GSMG0015882 | Arl13b | ADP-ribosylation factor-like 13B | 201.60 | 208.74 | -0.05 | 9.13E-01 |
| ENSMUSG00000052504 | GSMG0015883 | Epha3 | Eph receptor A3 | 556.80 | 881.83 | -0.66 | 9.10E-05 |
| ENSMUSG00000059920 | GSMG0015894 | 4930453N24Rik | RIKEN cDNA 4930453N24 gene | 772.34 | 850.94 | -0.14 | 5.96E-01 |
| ENSMUSG00000004843 | GSMG0015900 | Chmp2b | charged multivesicular body protein 2B | 2057.33 | 2317.90 | -0.17 | 4.79E-01 |
| ENSMUSG00000064115 | GSMG0015903 | Cadm2 | cell adhesion molecule 2 | 1046.81 | 1386.18 | -0.40 | 2.53E-02 |
| NULL | GSMG0015905 | NULL | NULL | 97.08 | 70.52 | 0.46 | 3.29E-01 |
| NULL | GSMG0015909 | NULL | NULL | 570.77 | 530.09 | 0.11 | 7.41E-01 |
| NULL | GSMG0015910 | NULL | NULL | 301.28 | 359.75 | -0.26 | 3.89E-01 |
| NULL | GSMG0015914 | NULL | NULL | 308.81 | 243.55 | 0.34 | 3.69E-01 |
| NULL | GSMG0015915 | NULL | NULL | 511.20 | 370.12 | 0.47 | 7.27E-02 |
| NULL | GSMG0015916 | NULL | NULL | 109.71 | 72.32 | 0.60 | 2.40E-01 |
| NULL | GSMG0015920 | NULL | NULL | 533.27 | 383.13 | 0.48 | 7.73E-02 |
| NULL | GSMG0015921 | NULL | NULL | 343.50 | 332.72 | 0.04 | 9.19E-01 |
| NULL | GSMG0015924 | NULL | NULL | 802.46 | 663.30 | 0.28 | 3.20E-01 |
| NULL | GSMG0015925 | NULL | NULL | 588.11 | 421.69 | 0.48 | 3.18E-02 |
| ENSMUSG00000052516 | GSMG0015936 | Robo2 | roundabout guidance receptor 2 | 283.08 | 642.71 | -1.18 | 1.06E-10 |
| NULL | GSMG0015945 | NULL | NULL | 184.90 | 212.65 | -0.20 | 5.84E-01 |
| NULL | GSMG0015947 | NULL | NULL | 311.38 | 295.52 | 0.07 | 8.83E-01 |
| NULL | GSMG0015951 | NULL | NULL | 470.07 | 590.44 | -0.33 | 2.02E-01 |
| NULL | GSMG0015952 | NULL | NULL | 261.79 | 274.89 | -0.07 | 9.03E-01 |
| NULL | GSMG0015953 | NULL | NULL | 929.70 | 1065.68 | -0.20 | 5.41E-01 |
| NULL | GSMG0015954 | NULL | NULL | 110.50 | 124.52 | -0.17 | 7.45E-01 |
| NULL | GSMG0015955 | NULL | NULL | 286.19 | 337.09 | -0.24 | 5.78E-01 |
| NULL | GSMG0015958 | NULL | NULL | 603.48 | 904.35 | -0.58 | 1.64E-02 |
| NULL | GSMG0015960 | NULL | NULL | 288.26 | 517.79 | -0.84 | 2.06E-03 |
| ENSMUSG00000032932 | GSMG0015962 | Hspa13 | heat shock protein 70 family, member 13 | 645.86 | 650.91 | -0.01 | 9.74E-01 |
| ENSMUSG00000022876 | GSMG0015963 | Samsn1 | SAM domain, SH3 domain and nuclear localization signals, 1 | 170.90 | 304.26 | -0.83 | 2.75E-03 |
| ENSMUSG00000048490 | GSMG0015965 | Nrip1 | nuclear receptor interacting protein 1 | 3557.75 | 5498.28 | -0.63 | 2.47E-05 |
| NULL | GSMG0015967 | NULL | NULL | 119.83 | 126.12 | -0.08 | 8.76E-01 |

| | | | | | | | |
|---|---|---|---|---|---|---|---|
| NULL | GSMG0015968 | NULL | NULL | 443.51 | 632.24 | -0.51 | 7.93E-03 |
| NULL | GSMG0015974 | NULL | NULL | 117.60 | 367.88 | -1.65 | 1.33E-09 |
| ENSMUSG00000022864 | GSMG0015981 | D16Ertd472e | DNA segment, Chr 16, ERATO Doi 472, expressed | 139.58 | 191.88 | -0.46 | 1.48E-01 |
| ENSMUSG00000022889 | GSMG0015987 | Mrpl39 | mitochondrial ribosomal protein L39 | 672.53 | 737.37 | -0.13 | 5.93E-01 |
| ENSMUSG00000022890 | GSMG0015988 | Atp5j | ATP synthase, H+ transporting, mitochondrial F0 complex, subunit F | 5076.24 | 5283.33 | -0.06 | 8.30E-01 |
| ENSMUSG00000022892 | GSMG0015993 | App | amyloid beta (A4) precursor protein | 29737.54 | 27225.61 | 0.13 | 5.77E-01 |
| ENSMUSG00000041134 | GSMG0015995 | Cyyr1 | cysteine and tyrosine-rich protein 1 | 470.00 | 387.85 | 0.28 | 2.09E-01 |
| ENSMUSG00000022893 | GSMG0015998 | Adamts1 | a disintegrin-like and metallopeptidase (reprolysin type) with thrombos | 2081.76 | 678.82 | 1.62 | 3.52E-14 |
| ENSMUSG00000052299 | GSMG0016005 | Ltn1 | listerin E3 ubiquitin protein ligase 1 | 935.48 | 1110.12 | -0.25 | 2.16E-01 |
| ENSMUSG00000025613 | GSMG0016008 | Cct8 | chaperonin containing Tcp1, subunit 8 (theta) | 3893.78 | 5028.73 | -0.37 | 5.49E-02 |
| ENSMUSG00000022935 | GSMG0016011 | Grik1 | glutamate receptor, ionotropic, kainate 1 | 349.82 | 539.01 | -0.62 | 3.07E-03 |
| ENSMUSG00000002489 | GSMG0016046 | Tiam1 | T cell lymphoma invasion and metastasis 1 | 833.68 | 942.39 | -0.18 | 4.86E-01 |
| ENSMUSG00000022983 | GSMG0016052 | Scaf4 | SR-related CTD-associated factor 4 | 1359.39 | 1402.32 | -0.04 | 8.70E-01 |
| NULL | GSMG0016053 | NULL | NULL | 265.39 | 281.54 | -0.09 | 8.10E-01 |
| ENSMUSG00000022978 | GSMG0016059 | Mis18a | MIS18 kinetochore protein A | 501.43 | 594.47 | -0.25 | 3.92E-01 |
| ENSMUSG00000022972 | GSMG0016063 | 1110004E09Rik | RIKEN cDNA 1110004E09 gene | 486.17 | 631.01 | -0.38 | 8.16E-02 |
| ENSMUSG00000022973 | GSMG0016065 | Synj1 | synaptojanin 1 | 1253.12 | 1598.39 | -0.35 | 3.23E-02 |
| ENSMUSG00000022974 | GSMG0016067 | Paxbp1 | PAX3 and PAX7 binding protein 1 | 1414.32 | 1535.44 | -0.12 | 6.58E-01 |
| ENSMUSG00000022964 | GSMG0016072 | Tmem50b | transmembrane protein 50B | 2448.75 | 2249.98 | 0.12 | 6.16E-01 |
| ENSMUSG00000022962 | GSMG0016074 | Gart | phosphoribosylglycinamide formyltransferase | 897.78 | 1366.17 | -0.61 | 2.47E-04 |
| ENSMUSG00000022960 | GSMG0016075 | Donson | downstream neighbor of SON | 405.31 | 452.28 | -0.16 | 5.47E-01 |
| ENSMUSG00000058240 | GSMG0016076 | Cryzl1 | crystallin, zeta (quinone reductase)-like 1 | 941.59 | 884.10 | 0.09 | 7.29E-01 |
| ENSMUSG00000022956 | GSMG0016078 | Atp5o | ATP synthase, H+ transporting, mitochondrial F1 complex, O subunit | 3000.96 | 3801.96 | -0.34 | 6.93E-02 |
| ENSMUSG00000022951 | GSMG0016083 | Rcan1 | regulator of calcineurin 1 | 1849.71 | 1641.02 | 0.17 | 4.59E-01 |
| ENSMUSG00000022948 | GSMG0016095 | Setd4 | SET domain containing 4 | 242.01 | 269.28 | -0.16 | 6.16E-01 |
| ENSMUSG00000040820 | GSMG0016100 | Hlcs | holocarboxylase synthetase (biotin- [propriony-Coenzyme A-carboxyla | 358.85 | 301.98 | 0.25 | 4.22E-01 |
| ENSMUSG00000022940 | GSMG0016101 | Pigp | phosphatidylinositol glycan anchor biosynthesis, class P | 730.32 | 738.81 | -0.02 | 9.59E-01 |
| ENSMUSG00000022898 | GSMG0016103 | Dscr3 | Down syndrome critical region gene 3 | 1627.54 | 1565.65 | 0.06 | 8.57E-01 |
| ENSMUSG00000043301 | GSMG0016105 | Kcnj6 | potassium inwardly-rectifying channel, subfamily J, member 6 | 141.80 | 91.07 | 0.64 | 5.66E-02 |
| ENSMUSG00000040732 | GSMG0016110 | Erg | avian erythroblastosis virus E-26 (v-ets) oncogene related | 477.81 | 411.32 | 0.22 | 4.90E-01 |
| ENSMUSG00000022913 | GSMG0016114 | Psmg1 | proteasome (prosome, macropain) assembly chaperone 1 | 360.57 | 528.30 | -0.55 | 2.28E-02 |
| ENSMUSG00000022914 | GSMG0016115 | Brwd1 | bromodomain and WD repeat domain containing 1 | 819.82 | 758.36 | 0.11 | 7.45E-01 |
| ENSMUSG00000040681 | GSMG0016117 | Hmgn1 | high mobility group nucleosomal binding domain 1 | 3198.64 | 4170.04 | -0.38 | 1.05E-01 |

| | | | | | | | |
|---|---|---|---|---|---|---|---|
| NULL | GSMG0016118 | NULL | NULL | 155.09 | 245.30 | -0.66 | 2.02E-02 |
| ENSMUSG00000050272 | GSMG0016122 | Dscam | Down syndrome cell adhesion molecule | 470.09 | 453.55 | 0.05 | 9.16E-01 |
| ENSMUSG00000014039 | GSMG0016135 | Prdm15 | PR domain containing 15 | 429.61 | 419.90 | 0.03 | 9.38E-01 |
| ENSMUSG00000045975 | GSMG0016136 | C2cd2 | C2 calcium-dependent domain containing 2 | 829.90 | 865.02 | -0.06 | 8.24E-01 |
| ENSMUSG00000046201 | GSMG0016144 | Scaf8 | SR-related CTD-associated factor 8 | 1799.10 | 2120.95 | -0.24 | 1.79E-01 |
| NULL | GSMG0016146 | NULL | NULL | 183.14 | 183.99 | -0.01 | 9.91E-01 |
| ENSMUSG00000023800 | GSMG0016148 | Tiam2 | T cell lymphoma invasion and metastasis 2 | 158.80 | 212.34 | -0.42 | 1.66E-01 |
| NULL | GSMG0016150 | NULL | NULL | 86.02 | 305.88 | -1.83 | 6.52E-12 |
| NULL | GSMG0016158 | NULL | NULL | 412.31 | 380.91 | 0.12 | 7.38E-01 |
| NULL | GSMG0016159 | NULL | NULL | 155.38 | 133.12 | 0.22 | 6.33E-01 |
| NULL | GSMG0016160 | NULL | NULL | 232.60 | 261.35 | -0.16 | 6.73E-01 |
| ENSMUSG00000034265 | GSMG0016166 | Zdhhc14 | zinc finger, DHHC domain containing 14 | 231.40 | 112.48 | 1.04 | 4.68E-05 |
| ENSMUSG00000002365 | GSMG0016170 | Snx9 | sorting nexin 9 | 1364.96 | 1524.11 | -0.16 | 4.58E-01 |
| ENSMUSG00000023805 | GSMG0016171 | Synj2 | synaptojanin 2 | 347.67 | 175.45 | 0.99 | 1.86E-04 |
| ENSMUSG00000034345 | GSMG0016173 | Gtf2h5 | general transcription factor IIH, polypeptide 5 | 1184.29 | 1517.58 | -0.36 | 1.39E-01 |
| ENSMUSG00000034377 | GSMG0016175 | Tulp4 | tubby like protein 4 | 2122.76 | 3091.92 | -0.54 | 8.70E-04 |
| ENSMUSG00000023809 | GSMG0016188 | Rps6ka2 | ribosomal protein S6 kinase, polypeptide 2 | 656.18 | 999.86 | -0.61 | 3.37E-03 |
| ENSMUSG00000069135 | GSMG0016193 | Fgfr1op | Fgfr1 oncogene partner | 348.52 | 388.51 | -0.16 | 6.07E-01 |
| ENSMUSG00000055945 | GSMG0016199 | Prr18 | proline rich 18 | 333.91 | 383.05 | -0.20 | 4.80E-01 |
| ENSMUSG00000023868 | GSMG0016202 | Pde10a | phosphodiesterase 10A | 457.90 | 455.42 | 0.01 | 9.88E-01 |
| NULL | GSMG0016204 | NULL | NULL | 248.64 | 9.84 | 4.66 | 5.33E-35 |
| NULL | GSMG0016220 | NULL | NULL | 313.16 | 146.76 | 1.09 | 1.03E-06 |
| ENSMUSG00000023827 | GSMG0016225 | Agpat4 | 1-acylglycerol-3-phosphate O-acyltransferase 4 (lysophosphatidic acid a | 1257.38 | 1045.71 | 0.27 | 1.95E-01 |
| ENSMUSG00000078247 | GSMG0016231 | Airn | antisense Igf2r RNA | 179.25 | 314.08 | -0.81 | 1.21E-04 |
| NULL | GSMG0016234 | NULL | NULL | 170.86 | 172.35 | -0.01 | 9.85E-01 |
| NULL | GSMG0016235 | NULL | NULL | 107.89 | 175.77 | -0.71 | 3.45E-02 |
| ENSMUSG00000068039 | GSMG0016243 | Tcp1 | t-complex protein 1 | 2982.86 | 4046.63 | -0.44 | 4.58E-03 |
| NULL | GSMG0016244 | NULL | NULL | 649.79 | 1717.36 | -1.40 | 5.50E-19 |
| ENSMUSG00000006818 | GSMG0016246 | Sod2 | superoxide dismutase 2, mitochondrial | 1874.08 | 1782.09 | 0.07 | 7.66E-01 |
| ENSMUSG00000068036 | GSMG0016253 | Mllt4 | myeloid/lymphoid or mixed-lineage leukemia%3B translocated to, 4 | 3875.67 | 4392.18 | -0.18 | 4.91E-01 |
| ENSMUSG00000023886 | GSMG0016260 | Smoc2 | SPARC related modular calcium binding 2 | 291.45 | 184.36 | 0.66 | 5.45E-03 |
| ENSMUSG00000050088 | GSMG0016261 | 1600012H06Rik | RIKEN cDNA 1600012H06 gene | 1595.85 | 1520.28 | 0.07 | 8.11E-01 |
| ENSMUSG00000014763 | GSMG0016267 | Fam120b | family with sequence similarity 120, member B | 835.60 | 861.97 | -0.04 | 9.06E-01 |

| | | | | | | | |
|---|---|---|---|---|---|---|---|
| ENSMUSG00000014767 | GSMG0016268 | Tbp | TATA box binding protein | 681.58 | 698.82 | -0.03 | 9.21E-01 |
| ENSMUSG00000047786 | GSMG0016280 | Lix1 | limb and CNS expressed 1 | 465.63 | 358.77 | 0.37 | 8.25E-02 |
| ENSMUSG00000080316 | GSMG0016284 | Spaca6 | sperm acrosome associated 6 | 242.45 | 168.64 | 0.52 | 7.85E-02 |
| ENSMUSG00000007564 | GSMG0016292 | Ppp2r1a | protein phosphatase 2, regulatory subunit A, alpha | 6097.03 | 7335.70 | -0.27 | 1.56E-01 |
| ENSMUSG00000067942 | GSMG0016293 | Zfp160 | zinc finger protein 160 | 627.79 | 583.31 | 0.11 | 7.62E-01 |
| ENSMUSG00000023892 | GSMG0016302 | Zfp51 | zinc finger protein 51 | 380.57 | 404.21 | -0.08 | 8.00E-01 |
| ENSMUSG00000051341 | GSMG0016305 | Zfp52 | zinc finger protein 52 | 381.01 | 217.15 | 0.81 | 1.49E-04 |
| ENSMUSG00000035868 | GSMG0016307 | 3110052M02Rik | RIKEN cDNA 3110052M02 gene | 485.50 | 707.73 | -0.55 | 4.27E-03 |
| ENSMUSG00000067928 | GSMG0016308 | Zfp760 | zinc finger protein 760 | 142.57 | 157.28 | -0.13 | 7.89E-01 |
| ENSMUSG00000053347 | GSMG0016313 | Zfp943 | zinc finger prtoein 943 | 564.28 | 385.93 | 0.55 | 4.43E-03 |
| ENSMUSG00000044501 | GSMG0016315 | Zfp758 | zinc finger protein 758 | 327.04 | 260.36 | 0.33 | 2.33E-01 |
| ENSMUSG00000071266 | GSMG0016317 | Zfp946 | zinc finger protein 946 | 224.74 | 315.75 | -0.49 | 7.56E-02 |
| ENSMUSG00000023904 | GSMG0016324 | Hcfc1r1 | host cell factor C1 regulator 1 (XPO1-dependent) | 165.23 | 190.86 | -0.21 | 5.48E-01 |
| NULL | GSMG0016326 | NULL | NULL | 156.29 | 40.22 | 1.96 | 1.61E-11 |
| ENSMUSG00000023908 | GSMG0016327 | Pkmyt1 | protein kinase, membrane associated tyrosine/threonine 1 | 335.52 | 541.72 | -0.69 | 7.72E-04 |
| ENSMUSG00000097006 | GSMG0016328 | 9530082P21Rik | RIKEN cDNA 9530082P21 gene | 752.05 | 675.18 | 0.15 | 5.76E-01 |
| ENSMUSG00000039218 | GSMG0016331 | Srrm2 | serine/arginine repetitive matrix 2 | 25602.41 | 15476.71 | 0.73 | 4.78E-06 |
| ENSMUSG00000024130 | GSMG0016337 | Abca3 | ATP-binding cassette, sub-family A (ABC1), member 3 | 390.77 | 488.94 | -0.32 | 2.41E-01 |
| ENSMUSG00000024132 | GSMG0016339 | Eci1 | enoyl-Coenzyme A delta isomerase 1 | 571.55 | 513.64 | 0.15 | 6.30E-01 |
| ENSMUSG00000043445 | GSMG0016341 | Pgp | phosphoglycolate phosphatase | 1292.60 | 1384.73 | -0.10 | 7.38E-01 |
| NULL | GSMG0016342 | NULL | NULL | 49.12 | 51.66 | -0.08 | 9.13E-01 |
| ENSMUSG00000045744 | GSMG0016343 | Bricd5 | BRICHOS domain containing 5 | 143.78 | 262.31 | -0.87 | 1.95E-04 |
| ENSMUSG00000033597 | GSMG0016344 | Caskin1 | CASK interacting protein 1 | 980.44 | 1432.94 | -0.55 | 4.11E-01 |
| ENSMUSG00000093565 | GSMG0016345 | Rab26os | RAB26, member RAS oncogene family, opposite strand | 42.96 | 59.64 | -0.49 | 3.71E-01 |
| ENSMUSG00000032855 | GSMG0016346 | Pkd1 | polycystic kidney disease 1 homolog | 981.33 | 732.63 | 0.42 | 1.45E-01 |
| ENSMUSG00000041429 | GSMG0016347 | Nthl1 | nth (endonuclease III)-like 1 (E.coli) | 137.34 | 151.36 | -0.13 | 7.71E-01 |
| ENSMUSG00000041130 | GSMG0016348 | Zfp598 | zinc finger protein 598 | 994.36 | 969.48 | 0.04 | 9.28E-01 |
| ENSMUSG00000019320 | GSMG0016349 | Noxo1 | NADPH oxidase organizer 1 | 162.79 | 218.29 | -0.42 | 1.09E-01 |
| ENSMUSG00000075705 | GSMG0016353 | Msrb1 | methionine sulfoxide reductase B1 | 238.65 | 224.63 | 0.08 | 8.69E-01 |
| ENSMUSG00000024158 | GSMG0016356 | Hagh | hydroxyacyl glutathione hydrolase | 1021.26 | 710.00 | 0.53 | 4.65E-03 |
| ENSMUSG00000024160 | GSMG0016358 | Spsb3 | splA/ryanodine receptor domain and SOCS box containing 3 | 88.64 | 103.94 | -0.23 | 6.30E-01 |
| G00000038880,ENSMUSG000 | GSMG0016359 | Mrps34 // Nme3 | mitochondrial ribosomal protein S34 // NME/NM23 nucleoside diphos | 1214.97 | 1318.63 | -0.12 | 6.68E-01 |
| NULL | GSMG0016360 | NULL | NULL | 86.15 | 109.88 | -0.35 | 3.49E-01 |

| | | | | | | | |
|---|---|---|---|---|---|---|---|
| ENSMUSG00000024169 | GSMG0016362 | Ift140 | intraflagellar transport 140 | 607.67 | 579.93 | 0.07 | 8.52E-01 |
| NULL | GSMG0016363 | NULL | NULL | 665.23 | 490.42 | 0.44 | 2.17E-02 |
| ENSMUSG00000036636 | GSMG0016366 | Clcn7 | chloride channel, voltage-sensitive 7 | 3531.78 | 3258.68 | 0.12 | 6.43E-01 |
| ENSMUSG00000015126 | GSMG0016370 | Tsr3 | TSR3 20S rRNA accumulation | 596.82 | 592.12 | 0.01 | 9.70E-01 |
| NULL | GSMG0016371 | Gm17801 | predicted gene, 17801 | 5799.78 | 8390.25 | -0.53 | 4.76E-04 |
| ENSMUSG00000091475 | GSMG0016378 | 2810468N07Rik | RIKEN cDNA 2810468N07 gene | 339.00 | 134.83 | 1.33 | 8.73E-10 |
| ENSMUSG00000002279 | GSMG0016380 | Lmf1 | lipase maturation factor 1 | 387.80 | 435.53 | -0.17 | 5.34E-01 |
| ENSMUSG00000025739 | GSMG0016382 | Gng13 | guanine nucleotide binding protein (G protein), gamma 13 | 241.52 | 61.58 | 1.97 | 2.01E-11 |
| ENSMUSG00000041199 | GSMG0016383 | Rpusd1 | RNA pseudouridylate synthase domain containing 1 | 563.45 | 442.41 | 0.35 | 1.29E-01 |
| ENSMUSG00000002280 | GSMG0016385 | Narfl | nuclear prelamin A recognition factor-like | 556.12 | 608.75 | -0.13 | 6.21E-01 |
| NULL | GSMG0016387 | NULL | NULL | 294.26 | 237.43 | 0.31 | 3.29E-01 |
| ENSMUSG00000025738 | GSMG0016388 | Fbxl16 | F-box and leucine-rich repeat protein 16 | 1116.13 | 751.66 | 0.57 | 3.53E-02 |
| ENSMUSG00000025737 | GSMG0016389 | Wdr24 | WD repeat domain 24 | 934.79 | 850.69 | 0.14 | 6.45E-01 |
| ENSMUSG00000025736 | GSMG0016390 | Jmjd8 | jumonji domain containing 8 | 520.53 | 508.82 | 0.04 | 9.26E-01 |
| NULL | GSMG0016391 | NULL | NULL | 643.81 | 1020.42 | -0.67 | 8.93E-04 |
| ENSMUSG00000025731 | GSMG0016393 | 0610011F06Rik | RIKEN cDNA 0610011F06 gene | 584.57 | 531.01 | 0.14 | 5.83E-01 |
| NULL | GSMG0016394 | NULL | NULL | 212.50 | 201.51 | 0.08 | 8.65E-01 |
| NULL | GSMG0016395 | NULL | NULL | 292.07 | 255.04 | 0.20 | 5.10E-01 |
| NULL | GSMG0016396 | NULL | NULL | 145.08 | 108.06 | 0.43 | 2.10E-01 |
| NULL | GSMG0016398 | NULL | NULL | 123.00 | 119.34 | 0.04 | 9.37E-01 |
| ENSMUSG00000024180 | GSMG0016400 | Tmem8 | transmembrane protein 8 (five membrane-spanning domains) | 269.94 | 166.84 | 0.70 | 3.67E-02 |
| ENSMUSG00000024181 | GSMG0016401 | Mrpl28 | mitochondrial ribosomal protein L28 | 1174.08 | 1436.45 | -0.29 | 1.29E-01 |
| ENSMUSG00000024182 | GSMG0016402 | Axin1 | axin 1 | 473.54 | 419.80 | 0.17 | 6.20E-01 |
| ENSMUSG00000024186 | GSMG0016404 | Rgs11 | regulator of G-protein signaling 11 | 2224.96 | 1664.89 | 0.42 | 4.00E-02 |
| ENSMUSG00000024188 | GSMG0016405 | Luc7l | Luc7-like | 1285.88 | 1771.21 | -0.46 | 1.14E-02 |
| ENSMUSG00000001576 | GSMG0016409 | Ergic1 | endoplasmic reticulum-golgi intermediate compartment (ERGIC) 1 | 960.22 | 1339.77 | -0.48 | 1.40E-02 |
| ENSMUSG00000015575 | GSMG0016412 | Atp6v0e | ATPase, H+ transporting, lysosomal V0 subunit E | 3360.10 | 2471.13 | 0.44 | 4.81E-03 |
| ENSMUSG00000048249 | GSMG0016413 | Crebrf | CREB3 regulatory factor | 760.58 | 477.78 | 0.67 | 1.48E-04 |
| ENSMUSG00000024191 | GSMG0016414 | Bnip1 | BCL2/adenovirus E1B interacting protein 1 | 416.24 | 539.81 | -0.38 | 9.20E-02 |
| ENSMUSG00000024193 | GSMG0016416 | Phf1 | PHD finger protein 1 | 216.49 | 268.35 | -0.31 | 2.63E-01 |
| ENSMUSG00000042644 | GSMG0016421 | Itpr3 | inositol 1,4,5-triphosphate receptor 3 | 191.14 | 86.07 | 1.15 | 1.44E-04 |
| NULL | GSMG0016422 | NULL | NULL | 207.32 | 206.98 | 0.00 | 9.98E-01 |
| ENSMUSG00000040276 | GSMG0016428 | Pacsin1 | protein kinase C and casein kinase substrate in neurons 1 | 1339.86 | 677.33 | 0.98 | 1.21E-10 |

| | | | | | | | |
|---|---|---|---|---|---|---|---|
| ENSMUSG00000039512 | GSMG0016431 | Uhrf1bp1 | UHRF1 (ICBP90) binding protein 1 | 349.24 | 454.82 | -0.38 | 2.97E-01 |
| ENSMUSG00000024219 | GSMG0016432 | Anks1 | ankyrin repeat and SAM domain containing 1 | 841.15 | 688.55 | 0.29 | 2.76E-01 |
| ENSMUSG00000024220 | GSMG0016434 | Zfp523 | zinc finger protein 523 | 2236.96 | 1710.52 | 0.39 | 1.25E-01 |
| ENSMUSG00000002257 | GSMG0016435 | Def6 | differentially expressed in FDCP 6 | 1001.47 | 1086.01 | -0.12 | 6.42E-01 |
| ENSMUSG00000002250 | GSMG0016437 | Ppard | peroxisome proliferator activator receptor delta | 109.61 | 100.83 | 0.11 | 8.75E-01 |
| ENSMUSG00000007570 | GSMG0016438 | Fance | Fanconi anemia, complementation group E | 723.26 | 741.14 | -0.04 | 9.18E-01 |
| ENSMUSG00000053436 | GSMG0016444 | Mapk14 | mitogen-activated protein kinase 14 | 809.71 | 744.09 | 0.12 | 6.53E-01 |
| ENSMUSG00000063952 | GSMG0016447 | Brpf3 | bromodomain and PHD finger containing, 3 | 172.36 | 118.87 | 0.53 | 1.39E-01 |
| ENSMUSG00000005936 | GSMG0016450 | Kctd20 | potassium channel tetramerisation domain containing 20 | 997.19 | 1153.98 | -0.21 | 3.25E-01 |
| ENSMUSG00000071172 | GSMG0016451 | Srsf3 | serine/arginine-rich splicing factor 3 | 6320.16 | 7892.26 | -0.32 | 1.45E-01 |
| ENSMUSG00000023067 | GSMG0016452 | Cdkn1a | cyclin-dependent kinase inhibitor 1A (P21) | 2551.71 | 2357.88 | 0.11 | 6.70E-01 |
| ENSMUSG00000052712 | GSMG0016454 | BC004004 | cDNA sequence BC004004 | 960.37 | 1004.38 | -0.06 | 8.20E-01 |
| 19,ENSMUSG00000090083,E | GSMG0016456 | Rnf8-cmtr1 // Rnf8 | Rnf8-cmtr1 readthrough // ring finger protein 8 // cap methyltransferase | 2333.23 | 2454.32 | -0.07 | 7.72E-01 |
| ENSMUSG00000024013 | GSMG0016457 | Fgd2 | FYVE, RhoGEF and PH domain containing 2 | 223.47 | 177.91 | 0.33 | 2.73E-01 |
| ENSMUSG00000024014 | GSMG0016458 | Pim1 | proviral integration site 1 | 58.69 | 122.06 | -1.06 | 1.12E-03 |
| ENSMUSG00000042203 | GSMG0016459 | Tbc1d22b | TBC1 domain family, member 22B | 180.38 | 234.75 | -0.38 | 2.09E-01 |
| NULL | GSMG0016461 | NULL | NULL | 161.98 | 91.29 | 0.82 | 2.24E-02 |
| ENSMUSG00000044477 | GSMG0016467 | Zfand3 | zinc finger, AN1-type domain 3 | 1535.53 | 1457.19 | 0.07 | 7.85E-01 |
| NULL | GSMG0016470 | NULL | NULL | 156.90 | 109.42 | 0.52 | 1.10E-01 |
| NULL | GSMG0016473 | NULL | NULL | 422.77 | 275.87 | 0.62 | 4.54E-03 |
| ENSMUSG00000024030 | GSMG0016480 | Abcg1 | ATP-binding cassette, sub-family G (WHITE), member 1 | 744.28 | 767.77 | -0.04 | 9.04E-01 |
| ENSMUSG00000024036 | GSMG0016482 | Slc37a1 | solute carrier family 37 (glycerol-3-phosphate transporter), member 1 | 295.16 | 186.28 | 0.66 | 1.38E-02 |
| ENSMUSG00000041119 | GSMG0016484 | Pde9a | phosphodiesterase 9A | 556.76 | 769.48 | -0.47 | 1.55E-02 |
| ENSMUSG00000024038 | GSMG0016487 | Ndufv3 | NADH dehydrogenase (ubiquinone) flavoprotein 3 | 1536.96 | 1744.10 | -0.18 | 3.72E-01 |
| ENSMUSG00000006705 | GSMG0016489 | Pknox1 | Pbx/knotted 1 homeobox | 336.58 | 385.58 | -0.20 | 4.60E-01 |
| NULL | GSMG0016492 | NULL | NULL | 200.28 | 60.80 | 1.73 | 8.22E-08 |
| ENSMUSG00000058392 | GSMG0016493 | Rrp1b | ribosomal RNA processing 1 homolog B (S. cerevisiae) | 616.70 | 765.02 | -0.31 | 1.27E-01 |
| NULL | GSMG0016495 | NULL | NULL | 228.02 | 190.65 | 0.26 | 4.04E-01 |
| NULL | GSMG0016496 | NULL | NULL | 12565.97 | 7764.86 | 0.69 | 1.55E-05 |
| ENSMUSG00000048440 | GSMG0016500 | Cyp4f16 | cytochrome P450, family 4, subfamily f, polypeptide 16 | 446.68 | 495.20 | -0.15 | 5.88E-01 |
| ENSMUSG00000073424 | GSMG0016503 | Cyp4f15 | cytochrome P450, family 4, subfamily f, polypeptide 15 | 182.31 | 61.11 | 1.58 | 2.54E-07 |
| ENSMUSG00000053600 | GSMG0016506 | Zfp472 | zinc finger protein 472 | 219.98 | 260.76 | -0.24 | 4.46E-01 |
| ENSMUSG00000053390 | GSMG0016507 | Zfp952 | zinc finger protein 952 | 384.78 | 394.45 | -0.04 | 9.20E-01 |

| | | | | | | | |
|---|---|---|---|---|---|---|---|
| ENSMUSG00000024299 | GSMG0016517 | Adamts10 | a disintegrin-like and metallopeptidase (reprolysin type) with thrombos[illegible] | 4479.71 | 2812.35 | 0.67 | 7.37E-04 |
| [illegible]G00000032739,ENSMUSG00[illegible] | GSMG0016519 | Zfp414 // Pram1 | zinc finger protein 414 // PML-RAR alpha-regulated adaptor molecule | 1127.02 | 1366.71 | -0.28 | 1.54E-01 |
| ENSMUSG00000060586 | GSMG0016521 | H2-Eb1 | histocompatibility 2, class II antigen E beta | 68.28 | 27.62 | 1.32 | 3.26E-03 |
| NULL | GSMG0016522 | NULL | NULL | 80.39 | 122.57 | -0.61 | 8.34E-02 |
| ENSMUSG00000042099 | GSMG0016523 | Kank3 | KN motif and ankyrin repeat domains 3 | 11096.38 | 9452.81 | 0.23 | 2.04E-01 |
| ENSMUSG00000041881 | GSMG0016524 | Ndufa7 | NADH dehydrogenase (ubiquinone) 1 alpha subcomplex, 7 (B14.5a) | 2329.83 | 2146.27 | 0.12 | 6.09E-01 |
| ENSMUSG00000002308 | GSMG0016525 | Cd320 | CD320 antigen | 1489.39 | 1529.01 | -0.04 | 9.08E-01 |
| ENSMUSG00000002307 | GSMG0016528 | Daxx | Fas death domain-associated protein | 911.20 | 1097.84 | -0.27 | 1.90E-01 |
| ENSMUSG00000051390 | GSMG0016529 | Zbtb22 | zinc finger and BTB domain containing 22 | 652.03 | 774.30 | -0.25 | 3.42E-01 |
| ENSMUSG00000024308 | GSMG0016530 | Tapbp | TAP binding protein | 2331.04 | 1964.73 | 0.25 | 2.24E-01 |
| ENSMUSG00000041354 | GSMG0016531 | Rgl2 | ral guanine nucleotide dissociation stimulator-like 2 | 3260.53 | 2584.46 | 0.34 | 1.71E-01 |
| ENSMUSG00000024312 | GSMG0016532 | Wdr46 | WD repeat domain 46 | 412.69 | 549.57 | -0.41 | 4.15E-02 |
| ENSMUSG00000024319 | GSMG0016533 | Vps52 | VPS52 GARP complex subunit | 1935.65 | 2026.40 | -0.07 | 8.08E-01 |
| ENSMUSG00000039656 | GSMG0016535 | Rxrb | retinoid X receptor beta | 650.11 | 588.87 | 0.14 | 6.24E-01 |
| ENSMUSG00000024330 | GSMG0016536 | Col11a2 | collagen, type XI, alpha 2 | 720.41 | 230.29 | 1.65 | 6.98E-09 |
| ENSMUSG00000037649 | GSMG0016538 | H2-DMa | histocompatibility 2, class II, locus DMa | 176.80 | 146.47 | 0.27 | 4.90E-01 |
| ENSMUSG00000037321 | GSMG0016541 | Tap1 | transporter 1, ATP-binding cassette, sub-family B (MDR/TAP) | 337.51 | 516.02 | -0.61 | 2.79E-02 |
| ENSMUSG00000024338 | GSMG0016542 | Psmb8 | proteasome (prosome, macropain) subunit, beta type 8 (large multifunct[illegible] | 272.04 | 158.09 | 0.78 | 3.21E-03 |
| ENSMUSG00000024339 | GSMG0016543 | Tap2 | transporter 2, ATP-binding cassette, sub-family B (MDR/TAP) | 1355.36 | 841.96 | 0.69 | 1.07E-04 |
| ENSMUSG00000015468 | GSMG0016544 | Notch4 | notch 4 | 2873.39 | 2301.77 | 0.32 | 2.70E-01 |
| ENSMUSG00000073421 | GSMG0016547 | H2-Ab1 | histocompatibility 2, class II antigen A, beta 1 | 66.93 | 49.81 | 0.43 | 4.18E-01 |
| ENSMUSG00000015452 | GSMG0016553 | Ager | advanced glycosylation end product-specific receptor | 56.87 | 96.63 | -0.76 | 5.49E-02 |
| ENSMUSG00000034254 | GSMG0016554 | Agpat1 | 1-acylglycerol-3-phosphate O-acyltransferase 1 (lysophosphatidic acid [illegible] | 536.93 | 752.32 | -0.49 | 4.08E-02 |
| NULL | GSMG0016555 | NULL | NULL | 130.35 | 108.82 | 0.26 | 5.63E-01 |
| ENSMUSG00000015476 | GSMG0016556 | Prrt1 | proline-rich transmembrane protein 1 | 139.45 | 97.93 | 0.51 | 1.54E-01 |
| ENSMUSG00000033739 | GSMG0016558 | Fkbpl | FK506 binding protein-like | 183.62 | 212.53 | -0.22 | 5.82E-01 |
| ENSMUSG00000015461 | GSMG0016559 | Atf6b | activating transcription factor 6 beta | 3041.95 | 3811.13 | -0.33 | 6.02E-02 |
| ENSMUSG00000040482 | GSMG0016561 | Dxo | decapping exoribonuclease | 720.03 | 602.59 | 0.26 | 2.26E-01 |
| ENSMUSG00000024369 | GSMG0016562 | Nelfe | negative elongation factor complex member E, Rdbp | 1035.33 | 1220.95 | -0.24 | 2.15E-01 |
| ENSMUSG00000013787 | GSMG0016565 | Ehmt2 | euchromatic histone lysine N-methyltransferase 2 | 1672.64 | 2020.61 | -0.27 | 1.90E-01 |
| ENSMUSG00000007038 | GSMG0016567 | Neu1 | neuraminidase 1 | 974.47 | 803.50 | 0.28 | 2.41E-01 |
| ENSMUSG00000007050 | GSMG0016571 | Lsm2 | LSM2 homolog, U6 small nuclear RNA and mRNA degradation associ[illegible] | 415.63 | 726.49 | -0.81 | 7.17E-05 |
| ENSMUSG00000007029 | GSMG0016572 | Vars | valyl-tRNA synthetase | 2598.81 | 5071.75 | -0.96 | 1.03E-07 |

| | | | | | | | |
|---|---|---|---|---|---|---|---|
| ENSMUSG00000007041 | GSMG0016574 | Clic1 | chloride intracellular channel 1 | 4760.73 | 4949.28 | -0.06 | 8.44E-01 |
| ENSMUSG00000007039 | GSMG0016575 | Ddah2 | dimethylarginine dimethylaminohydrolase 2 | 1321.27 | 1602.22 | -0.28 | 1.68E-01 |
| ENSMUSG00000007036 | GSMG0016578 | Abhd16a | abhydrolase domain containing 16A | 3864.40 | 3382.17 | 0.19 | 3.79E-01 |
| ENSMUSG00000092417 | GSMG0016580 | Gpank1 | G patch domain and ankyrin repeats 1 | 120.33 | 108.11 | 0.16 | 7.22E-01 |
| ENSMUSG00000024392 | GSMG0016582 | Bag6 | BCL2-associated athanogene 6 | 3024.92 | 3353.91 | -0.15 | 6.87E-01 |
| ENSMUSG00000024403 | GSMG0016586 | Atp6v1g2 | ATPase, H+ transporting, lysosomal V1 subunit G2 | 2380.11 | 1477.84 | 0.69 | 5.05E-06 |
| NULL | GSMG0016588 | NULL | NULL | 478.51 | 511.30 | -0.10 | 8.22E-01 |
| ENSMUSG00000073411 | GSMG0016589 | H2-D1 | histocompatibility 2, D region locus 1 | 7898.00 | 11949.81 | -0.60 | 3.90E-04 |
| ENSMUSG00000035929 | GSMG0016593 | H2-Q4 | histocompatibility 2, Q region locus 4 | 769.77 | 635.66 | 0.28 | 3.12E-01 |
| ENSMUSG00000040312 | GSMG0016599 | Cchcr1 | coiled-coil alpha-helical rod protein 1 | 284.43 | 353.21 | -0.31 | 2.00E-01 |
| ENSMUSG00000003541 | GSMG0016606 | Ier3 | immediate early response 3 | 1014.76 | 332.95 | 1.61 | 4.38E-21 |
| ENSMUSG00000059714 | GSMG0016607 | Flot1 | flotillin 1 | 2373.52 | 2107.16 | 0.17 | 3.77E-01 |
| ENSMUSG00000059791 | GSMG0016609 | Nrm | nurim (nuclear envelope membrane protein) | 512.67 | 690.05 | -0.43 | 3.74E-02 |
| ENSMUSG00000034595 | GSMG0016610 | Ppp1r18 | protein phosphatase 1, regulatory subunit 18 | 766.05 | 867.55 | -0.18 | 4.65E-01 |
| ENSMUSG00000024422 | GSMG0016611 | Dhx16 | DEAH (Asp-Glu-Ala-His) box polypeptide 16 | 1496.34 | 1422.13 | 0.07 | 7.99E-01 |
| G00000039220,ENSMUSG000 | GSMG0016613 | Ppp1r10 // Gm880 | protein phosphatase 1, regulatory subunit 10 // predicted gene 8801 | 210.12 | 255.57 | -0.29 | 4.82E-01 |
| NULL | GSMG0016614 | NULL | NULL | 342.01 | 287.09 | 0.25 | 4.32E-01 |
| ENSMUSG00000024429 | GSMG0016615 | Gnl1 | guanine nucleotide binding protein-like 1 | 919.84 | 874.01 | 0.07 | 8.19E-01 |
| ENSMUSG00000024457 | GSMG0016626 | Trim26 | tripartite motif-containing 26 | 840.46 | 798.85 | 0.07 | 7.94E-01 |
| ENSMUSG00000036036 | GSMG0016633 | Zfp57 | zinc finger protein 57 | 637.68 | 3224.26 | -2.34 | 3.45E-40 |
| ENSMUSG00000024462 | GSMG0016635 | Gabbr1 | gamma-aminobutyric acid (GABA) B receptor, 1 | 1361.50 | 1620.02 | -0.25 | 3.87E-01 |
| ENSMUSG00000016206 | GSMG0016640 | H2-M3 | histocompatibility 2, M region locus 3 | 181.28 | 179.62 | 0.02 | 9.69E-01 |
| ENSMUSG00000023921 | GSMG0016674 | Mut | methylmalonyl-Coenzyme A mutase | 1207.09 | 1117.06 | 0.11 | 6.70E-01 |
| ENSMUSG00000042256 | GSMG0016676 | Ptchd4 | patched domain containing 4 | 379.27 | 317.99 | 0.25 | 3.23E-01 |
| ENSMUSG00000023915 | GSMG0016682 | Tnfrsf21 | tumor necrosis factor receptor superfamily, member 21 | 3130.29 | 3957.03 | -0.34 | 6.28E-02 |
| ENSMUSG00000056492 | GSMG0016685 | Adgrf5 | adhesion G protein-coupled receptor F5 | 16023.73 | 10242.51 | 0.65 | 7.16E-06 |
| ENSMUSG00000023913 | GSMG0016686 | Pla2g7 | phospholipase A2, group VII (platelet-activating factor acetylhydrolase | 1929.02 | 1590.46 | 0.28 | 1.61E-01 |
| ENSMUSG00000023963 | GSMG0016687 | Cyp39a1 | cytochrome P450, family 39, subfamily a, polypeptide 1 | 108.14 | 127.41 | -0.23 | 5.65E-01 |
| ENSMUSG00000039601 | GSMG0016690 | Rcan2 | regulator of calcineurin 2 | 671.09 | 388.49 | 0.79 | 5.16E-05 |
| ENSMUSG00000023960 | GSMG0016693 | Enpp5 | ectonucleotide pyrophosphatase/phosphodiesterase 5 | 744.49 | 739.29 | 0.01 | 9.75E-01 |
| ENSMUSG00000023959 | GSMG0016696 | Clic5 | chloride intracellular channel 5 | 300.73 | 112.62 | 1.42 | 3.34E-11 |
| ENSMUSG00000038954 | GSMG0016700 | Supt3 | suppressor of Ty 3 | 108.89 | 167.88 | -0.63 | 3.17E-02 |
| ENSMUSG00000097119 | GSMG0016705 | B230354K17Rik | RIKEN cDNA B230354K17 gene | 511.76 | 468.92 | 0.13 | 6.55E-01 |

| | | | | | | | |
|---|---|---|---|---|---|---|---|
| ENSMUSG00000023938 | GSMG0016708 | Aars2 | alanyl-tRNA synthetase 2, mitochondrial | 473.57 | 534.67 | -0.17 | 5.82E-01 |
| ENSMUSG00000037089 | GSMG0016711 | Slc35b2 | solute carrier family 35, member B2 | 2661.45 | 2790.38 | -0.07 | 8.03E-01 |
| NULL | GSMG0016712 | NULL | NULL | 105.60 | 211.07 | -1.00 | 4.48E-05 |
| ENSMUSG00000023939 | GSMG0016713 | Mrpl14 | mitochondrial ribosomal protein L14 | 419.61 | 447.05 | -0.09 | 7.69E-01 |
| ENSMUSG00000023967 | GSMG0016718 | Mrps18a | mitochondrial ribosomal protein S18A | 984.48 | 1212.82 | -0.30 | 1.04E-01 |
| ENSMUSG00000023952 | GSMG0016719 | Gtpbp2 | GTP binding protein 2 | 794.07 | 661.43 | 0.26 | 2.68E-01 |
| ENSMUSG00000067150 | GSMG0016720 | Xpo5 | exportin 5 | 312.48 | 341.07 | -0.12 | 6.99E-01 |
| ENSMUSG00000071074 | GSMG0016721 | Yipf3 | Yip1 domain family, member 3 | 1668.88 | 1425.17 | 0.23 | 2.74E-01 |
| ENSMUSG00000071073 | GSMG0016722 | Lrrc73 | leucine rich repeat containing 73 | 90.33 | 104.73 | -0.22 | 6.22E-01 |
| ENSMUSG00000047428 | GSMG0016723 | Dlk2 | delta-like 2 homolog (Drosophila) | 156.35 | 131.70 | 0.25 | 5.56E-01 |
| ENSMUSG00000015597 | GSMG0016725 | Zfp318 | zinc finger protein 318 | 140.41 | 146.20 | -0.06 | 8.94E-01 |
| ENSMUSG00000002767 | GSMG0016731 | Mrpl2 | mitochondrial ribosomal protein L2 | 956.60 | 1194.71 | -0.32 | 8.56E-02 |
| ENSMUSG00000038545 | GSMG0016732 | Cul7 | cullin 7 | 1278.27 | 1713.38 | -0.42 | 6.33E-02 |
| NULL | GSMG0016734 | NULL | NULL | 82.00 | 74.10 | 0.13 | 8.43E-01 |
| ENSMUSG00000002768 | GSMG0016737 | Mea1 | male enhanced antigen 1 | 1437.31 | 1699.06 | -0.24 | 2.87E-01 |
| ENSMUSG00000002763 | GSMG0016738 | Pex6 | peroxisomal biogenesis factor 6 | 2007.37 | 1849.94 | 0.12 | 6.56E-01 |
| ENSMUSG00000062619 | GSMG0016739 | 2310039H08Rik | RIKEN cDNA 2310039H08 gene | 379.14 | 452.80 | -0.26 | 3.59E-01 |
| ENSMUSG00000036430 | GSMG0016740 | Tbcc | tubulin-specific chaperone C | 1328.38 | 1174.78 | 0.18 | 5.00E-01 |
| ENSMUSG00000064043 | GSMG0016743 | Trerf1 | transcriptional regulating factor 1 | 720.46 | 548.07 | 0.39 | 1.25E-01 |
| ENSMUSG00000034729 | GSMG0016749 | Mrps10 | mitochondrial ribosomal protein S10 | 400.34 | 477.92 | -0.26 | 3.27E-01 |
| ENSMUSG00000034165 | GSMG0016753 | Ccnd3 | cyclin D3 | 2598.91 | 2081.81 | 0.32 | 8.91E-02 |
| G00000090115,ENSMUSG000 | GSMG0016757 | Med20 // Usp49 | mediator complex subunit 20 // ubiquitin specific peptidase 49 | 1715.49 | 1464.60 | 0.23 | 2.39E-01 |
| ENSMUSG00000023266 | GSMG0016760 | Frs3 | fibroblast growth factor receptor substrate 3 | 332.48 | 329.94 | 0.01 | 9.85E-01 |
| ENSMUSG00000023990 | GSMG0016762 | Tfeb | transcription factor EB | 567.16 | 377.91 | 0.58 | 2.05E-03 |
| ENSMUSG00000023992 | GSMG0016770 | Trem2 | triggering receptor expressed on myeloid cells 2 | 101.58 | 67.54 | 0.59 | 1.23E-01 |
| ENSMUSG00000040771 | GSMG0016772 | Oard1 | O-acyl-ADP-ribose deacylase 1 | 467.38 | 515.64 | -0.14 | 6.69E-01 |
| ENSMUSG00000038910 | GSMG0016783 | Plcl2 | phospholipase C-like 2 | 277.12 | 277.66 | -0.01 | 9.90E-01 |
| ENSMUSG00000017831 | GSMG0016795 | Rab5a | RAB5A, member RAS oncogene family | 1238.15 | 1289.20 | -0.06 | 8.52E-01 |
| ENSMUSG00000000708 | GSMG0016797 | Kat2b | K(lysine) acetyltransferase 2B | 505.28 | 341.80 | 0.56 | 4.68E-03 |
| ENSMUSG00000024172 | GSMG0016801 | St6gal2 | beta galactoside alpha 2,6 sialyltransferase 2 | 325.80 | 772.93 | -1.25 | 3.72E-10 |
| ENSMUSG00000003198 | GSMG0016803 | Zfp959 | zinc finger protein 959 | 390.69 | 348.46 | 0.17 | 5.82E-01 |
| ENSMUSG00000003208 | GSMG0016805 | Ccdc94 | coiled-coil domain containing 94 | 342.29 | 436.00 | -0.35 | 1.09E-01 |
| ENSMUSG00000039154 | GSMG0016806 | Shd | src homology 2 domain-containing transforming protein D | 465.91 | 797.79 | -0.78 | 5.45E-06 |

| | | | | | | | |
|---|---|---|---|---|---|---|---|
| ENSMUSG00000011589 | GSMG0016807 | Fsd1 | fibronectin type 3 and SPRY domain-containing protein | 1280.61 | 1681.44 | -0.39 | 2.67E-02 |
| ENSMUSG00000003199 | GSMG0016808 | Mpnd | MPN domain containing | 2452.18 | 2143.50 | 0.19 | 3.15E-01 |
| ENSMUSG00000002835 | GSMG0016809 | Chaf1a | chromatin assembly factor 1, subunit A (p150) | 283.11 | 685.24 | -1.27 | 2.78E-11 |
| ENSMUSG00000002833 | GSMG0016810 | Hdgfrp2 | hepatoma-derived growth factor, related protein 2 | 4284.45 | 4250.92 | 0.01 | 9.67E-01 |
| ENSMUSG00000044469 | GSMG0016812 | Tnfaip8l1 | tumor necrosis factor, alpha-induced protein 8-like 1 | 978.22 | 1289.86 | -0.40 | 3.72E-02 |
| ENSMUSG00000001228 | GSMG0016817 | Uhrf1 | ubiquitin-like, containing PHD and RING finger domains, 1 | 364.58 | 855.44 | -1.23 | 2.37E-08 |
| ENSMUSG00000024201 | GSMG0016818 | Kdm4b | lysine (K)-specific demethylase 4B | 2075.65 | 2224.08 | -0.10 | 7.40E-01 |
| NULL | GSMG0016822 | NULL | NULL | 218.21 | 253.96 | -0.22 | 4.50E-01 |
| ENSMUSG00000002372 | GSMG0016826 | Ranbp3 | RAN binding protein 3 | 1143.45 | 1433.92 | -0.33 | 9.76E-02 |
| ENSMUSG00000002379 | GSMG0016827 | Ndufa11 | NADH dehydrogenase (ubiquinone) 1 alpha subcomplex 11 | 2025.84 | 2145.83 | -0.08 | 7.72E-01 |
| ENSMUSG00000007603 | GSMG0016829 | Dus3l | dihydrouridine synthase 3-like (S. cerevisiae) | 4559.28 | 4890.77 | -0.10 | 6.88E-01 |
| ENSMUSG00000002660 | GSMG0016832 | Clpp | caseinolytic mitochondrial matrix peptidase proteolytic subunit | 1387.69 | 1503.49 | -0.12 | 5.97E-01 |
| ENSMUSG00000002661 | GSMG0016833 | Alkbh7 | alkB homolog 7 | 334.26 | 340.33 | -0.03 | 9.44E-01 |
| ENSMUSG00000019487 | GSMG0016840 | Trip10 | thyroid hormone receptor interactor 10 | 1268.32 | 994.85 | 0.35 | 8.46E-02 |
| ENSMUSG00000000127 | GSMG0016849 | Fer | fer (fms/fps related) protein kinase | 472.08 | 444.44 | 0.09 | 8.07E-01 |
| ENSMUSG00000024085 | GSMG0016854 | Man2a1 | mannosidase 2, alpha 1 | 842.18 | 1001.32 | -0.25 | 4.43E-01 |
| NULL | GSMG0016860 | NULL | NULL | 136.25 | 133.90 | 0.03 | 9.57E-01 |
| ENSMUSG00000061950 | GSMG0016861 | Ppp4r1 | protein phosphatase 4, regulatory subunit 1 | 1112.08 | 1065.57 | 0.06 | 8.50E-01 |
| ENSMUSG00000024101 | GSMG0016865 | Wash1 | WAS protein family homolog 1 | 254.07 | 207.44 | 0.29 | 4.59E-01 |
| ENSMUSG00000035842 | GSMG0016866 | Ddx11 | DEAD/H (Asp-Glu-Ala-Asp/His) box helicase 11 | 390.58 | 584.79 | -0.58 | 1.68E-03 |
| NULL | GSMG0016869 | NULL | NULL | 134.13 | 79.93 | 0.74 | 1.44E-02 |
| ENSMUSG00000032796 | GSMG0016872 | Lama1 | laminin, alpha 1 | 1364.08 | 664.00 | 1.04 | 5.08E-08 |
| ENSMUSG00000024044 | GSMG0016879 | Epb41l3 | erythrocyte membrane protein band 4.1 like 3 | 420.76 | 371.85 | 0.18 | 5.77E-01 |
| ENSMUSG00000049672 | GSMG0016884 | Zbtb14 | zinc finger and BTB domain containing 14 | 1419.75 | 1339.81 | 0.08 | 7.71E-01 |
| ENSMUSG00000003279 | GSMG0016887 | Dlgap1 | discs, large (Drosophila) homolog-associated protein 1 | 2495.48 | 1660.20 | 0.59 | 6.28E-05 |
| NULL | GSMG0016891 | NULL | NULL | 902.55 | 436.35 | 1.05 | 1.26E-08 |
| NULL | GSMG0016892 | NULL | NULL | 613.40 | 515.00 | 0.25 | 4.46E-01 |
| NULL | GSMG0016893 | NULL | NULL | 398.91 | 258.84 | 0.62 | 8.47E-03 |
| NULL | GSMG0016894 | NULL | NULL | 564.88 | 409.80 | 0.46 | 8.59E-02 |
| NULL | GSMG0016895 | NULL | NULL | 471.36 | 259.73 | 0.86 | 4.25E-05 |
| ENSMUSG00000024052 | GSMG0016904 | Lpin2 | lipin 2 | 218.11 | 294.36 | -0.43 | 1.64E-01 |
| ENSMUSG00000041057 | GSMG0016908 | Wdr43 | WD repeat domain 43 | 1488.92 | 2077.38 | -0.48 | 6.49E-03 |
| ENSMUSG00000024059 | GSMG0016911 | Clip4 | CAP-GLY domain containing linker protein family, member 4 | 185.35 | 222.61 | -0.26 | 4.23E-01 |

| | | | | | | | |
|---|---|---|---|---|---|---|---|
| ENSMUSG00000039770 | GSMG0016912 | Ypel5 | yippee-like 5 (Drosophila) | 2224.35 | 1921.51 | 0.21 | 4.36E-01 |
| ENSMUSG00000024063 | GSMG0016914 | Lbh | limb-bud and heart | 461.18 | 519.88 | -0.17 | 5.14E-01 |
| ENSMUSG00000054469 | GSMG0016915 | Lclat1 | lysocardiolipin acyltransferase 1 | 159.25 | 178.03 | -0.16 | 6.89E-01 |
| ENSMUSG00000024065 | GSMG0016918 | Ehd3 | EH-domain containing 3 | 397.52 | 421.38 | -0.08 | 8.32E-01 |
| ENSMUSG00000024068 | GSMG0016919 | Spast | spastin | 734.43 | 853.61 | -0.22 | 4.03E-01 |
| ENSMUSG00000024069 | GSMG0016921 | Slc30a6 | solute carrier family 30 (zinc transporter), member 6 | 443.94 | 407.13 | 0.12 | 6.84E-01 |
| ENSMUSG00000024072 | GSMG0016922 | Yipf4 | Yip1 domain family, member 4 | 960.89 | 865.80 | 0.15 | 6.17E-01 |
| ENSMUSG00000024073 | GSMG0016923 | Birc6 | baculoviral IAP repeat-containing 6 | 2052.73 | 1993.55 | 0.04 | 8.77E-01 |
| ENSMUSG00000024078 | GSMG0016926 | Ttc27 | tetratricopeptide repeat domain 27 | 269.51 | 473.58 | -0.82 | 6.65E-05 |
| ENSMUSG00000001870 | GSMG0016928 | Ltbp1 | latent transforming growth factor beta binding protein 1 | 122.48 | 34.03 | 1.86 | 1.12E-07 |
| ENSMUSG00000071042 | GSMG0016933 | Rasgrp3 | RAS, guanyl releasing protein 3 | 4571.75 | 5042.08 | -0.14 | 5.54E-01 |
| ENSMUSG00000024074 | GSMG0016936 | Crim1 | cysteine rich transmembrane BMP regulator 1 (chordin like) | 152.86 | 79.34 | 0.95 | 3.50E-02 |
| ENSMUSG00000050668 | GSMG0016943 | Gpatch11 | G patch domain containing 11 | 595.05 | 671.83 | -0.18 | 5.05E-01 |
| ENSMUSG00000062691 | GSMG0016945 | Cebpzos | CCAAT/enhancer binding protein (C/EBP), zeta, opposite strand | 342.62 | 453.71 | -0.41 | 1.72E-01 |
| ENSMUSG00000024082 | GSMG0016946 | Ndufaf7 | NADH dehydrogenase (ubiquinone) 1 alpha subcomplex assembly facto | 1231.74 | 1156.87 | 0.09 | 7.36E-01 |
| ENSMUSG00000024084 | GSMG0016947 | Qpct | glutaminyl-peptide cyclotransferase (glutaminyl cyclase) | 135.96 | 146.45 | -0.11 | 8.11E-01 |
| ENSMUSG00000036368 | GSMG0016950 | Rmdn2 | regulator of microtubule dynamics 2 | 259.60 | 214.19 | 0.28 | 3.28E-01 |
| ENSMUSG00000055760 | GSMG0016961 | Gemin6 | gem (nuclear organelle) associated protein 6 | 162.72 | 208.76 | -0.36 | 3.25E-01 |
| ENSMUSG00000024245 | GSMG0016971 | Tmem178 | transmembrane protein 178 | 323.41 | 223.34 | 0.53 | 1.92E-02 |
| ENSMUSG00000024247 | GSMG0016974 | Pkdcc | protein kinase domain containing, cytoplasmic | 146.14 | 103.09 | 0.50 | 2.03E-01 |
| ENSMUSG00000032624 | GSMG0016976 | Eml4 | echinoderm microtubule associated protein like 4 | 595.45 | 753.24 | -0.34 | 1.49E-01 |
| ENSMUSG00000055817 | GSMG0016981 | Mta3 | metastasis associated 3 | 765.14 | 759.18 | 0.01 | 9.75E-01 |
| ENSMUSG00000040852 | GSMG0016992 | Plekhh2 | pleckstrin homology domain containing, family H (with MyTH4 domai | 892.42 | 363.56 | 1.29 | 5.30E-12 |
| ENSMUSG00000024253 | GSMG0016993 | Dync2li1 | dynein cytoplasmic 2 light intermediate chain 1 | 202.40 | 174.16 | 0.22 | 5.35E-01 |
| ENSMUSG00000061130 | GSMG0016995 | Ppm1b | protein phosphatase 1B, magnesium dependent, beta isoform | 1129.50 | 1012.10 | 0.16 | 4.94E-01 |
| ENSMUSG00000071037 | GSMG0016999 | Camkmt | calmodulin-lysine N-methyltransferase | 264.17 | 363.68 | -0.46 | 5.32E-02 |
| ENSMUSG00000045038 | GSMG0017009 | Prkce | protein kinase C, epsilon | 958.05 | 1160.35 | -0.28 | 3.28E-01 |
| NULL | GSMG0017010 | NULL | NULL | 872.40 | 782.46 | 0.16 | 5.41E-01 |
| NULL | GSMG0017011 | NULL | NULL | 568.93 | 407.58 | 0.48 | 3.42E-02 |
| NULL | GSMG0017013 | NULL | NULL | 409.23 | 299.93 | 0.45 | 6.28E-02 |
| NULL | GSMG0017015 | NULL | NULL | 500.39 | 431.31 | 0.22 | 4.56E-01 |
| ENSMUSG00000024140 | GSMG0017022 | Epas1 | endothelial PAS domain protein 1 | 1638.11 | 1657.45 | -0.02 | 9.65E-01 |
| ENSMUSG00000024143 | GSMG0017025 | Rhoq | ras homolog family member Q | 1638.76 | 924.60 | 0.82 | 5.79E-05 |

| | | | | | | | |
|---|---|---|---|---|---|---|---|
| NULL | GSMG0017027 | NULL | NULL | 235.88 | 81.20 | 1.54 | 3.55E-06 |
| ENSMUSG00000024146 | GSMG0017028 | Cript | cysteine-rich PDZ-binding protein | 1433.30 | 1713.38 | -0.26 | 2.09E-01 |
| ENSMUSG00000037104 | GSMG0017029 | Socs5 | suppressor of cytokine signaling 5 | 1487.11 | 1414.74 | 0.07 | 7.97E-01 |
| ENSMUSG00000024151 | GSMG0017032 | Msh2 | mutS homolog 2 | 1047.22 | 1190.48 | -0.18 | 4.04E-01 |
| ENSMUSG00000005370 | GSMG0017034 | Msh6 | mutS homolog 6 | 1044.55 | 1345.53 | -0.37 | 3.62E-02 |
| ENSMUSG00000034998 | GSMG0017036 | Foxn2 | forkhead box N2 | 137.47 | 143.82 | -0.07 | 8.85E-01 |
| ENSMUSG00000034709 | GSMG0017038 | Ppp1r21 | protein phosphatase 1, regulatory subunit 21 | 212.08 | 202.89 | 0.07 | 8.71E-01 |
| NULL | GSMG0017046 | NULL | NULL | 399.19 | 422.72 | -0.08 | 8.43E-01 |
| ENSMUSG00000090160 | GSMG0017047 | 4930480K15Rik | RIKEN cDNA 4930480K15 gene | 168.40 | 208.89 | -0.30 | 5.55E-01 |
| NULL | GSMG0017048 | NULL | NULL | 153.81 | 230.19 | -0.58 | 5.03E-02 |
| ENSMUSG00000098090 | GSMG0017052 | 2700099C18Rik | RIKEN cDNA 2700099C18 gene | 82.24 | 125.82 | -0.61 | 7.63E-02 |
| NULL | GSMG0017057 | NULL | NULL | 207.86 | 202.29 | 0.03 | 9.42E-01 |
| ENSMUSG00000004945 | GSMG0017067 | Tmem242 | transmembrane protein 242 | 686.92 | 686.00 | 0.00 | 9.95E-01 |
| ENSMUSG00000015659 | GSMG0017073 | Serac1 | serine active site containing 1 | 240.18 | 361.96 | -0.59 | 8.68E-03 |
| ENSMUSG00000052397 | GSMG0017077 | Ezr | ezrin | 1721.36 | 1042.39 | 0.72 | 3.01E-04 |
| NULL | GSMG0017099 | NULL | NULL | 736.55 | 486.18 | 0.60 | 1.26E-02 |
| ENSMUSG00000062078 | GSMG0017100 | Qk | quaking | 5726.01 | 2528.50 | 1.18 | 6.58E-02 |
| ENSMUSG00000014426 | GSMG0017115 | Map3k4 | mitogen-activated protein kinase kinase kinase 4 | 955.76 | 1184.91 | -0.31 | 1.29E-01 |
| ENSMUSG00000023830 | GSMG0017119 | Igf2r | insulin-like growth factor 2 receptor | 629.92 | 904.49 | -0.52 | 1.07E-02 |
| ENSMUSG00000057388 | GSMG0017122 | Mrpl18 | mitochondrial ribosomal protein L18 | 1325.78 | 1590.49 | -0.26 | 2.31E-01 |
| ENSMUSG00000023832 | GSMG0017124 | Acat2 | acetyl-Coenzyme A acetyltransferase 2 | 1093.06 | 2944.05 | -1.43 | 1.59E-22 |
| ENSMUSG00000060475 | GSMG0017125 | Wtap | Wilms tumour 1-associating protein | 1477.15 | 1779.08 | -0.27 | 1.61E-01 |
| ENSMUSG00000023885 | GSMG0017137 | Thbs2 | thrombospondin 2 | 959.25 | 788.77 | 0.28 | 2.00E-01 |
| ENSMUSG00000023883 | GSMG0017139 | Phf10 | PHD finger protein 10 | 750.69 | 582.71 | 0.36 | 1.12E-01 |
| ENSMUSG00000014769 | GSMG0017148 | Psmb1 | proteasome (prosome, macropain) subunit, beta type 1 | 3839.57 | 4722.08 | -0.30 | 8.18E-02 |
| ENSMUSG00000014771 | GSMG0017149 | Pdcd2 | programmed cell death 2 | 233.81 | 316.02 | -0.44 | 7.43E-02 |
| NULL | GSMG0017152 | NULL | NULL | 99.85 | 84.28 | 0.24 | 5.95E-01 |
| ENSMUSG00000048027 | GSMG0017153 | Rgmb | repulsive guidance molecule family member B | 225.76 | 320.07 | -0.50 | 5.33E-02 |
| ENSMUSG00000023845 | GSMG0017160 | Lnpep | leucyl/cystinyl aminopeptidase | 278.38 | 303.30 | -0.12 | 7.18E-01 |
| ENSMUSG00000071267 | GSMG0017180 | Zfp942 | zinc finger protein 942 | 563.37 | 564.83 | -0.01 | 9.89E-01 |
| NULL | GSMG0017184 | Gm7072 | predicted gene 7072 | 772.07 | 514.44 | 0.59 | 6.36E-03 |
| ENSMUSG00000033972 | GSMG0017185 | Zfp944 | zinc finger protein 944 | 433.91 | 459.12 | -0.08 | 8.04E-01 |
| ENSMUSG00000059142 | GSMG0017189 | Zfp945 | zinc finger protein 945 | 718.42 | 701.31 | 0.03 | 9.16E-01 |

| | | | | | | | |
|---|---|---|---|---|---|---|---|
| ENSMUSG00000002617 | GSMG0017190 | Zfp40 | zinc finger protein 40 | 748.54 | 681.05 | 0.14 | 5.60E-01 |
| NULL | GSMG0017191 | NULL | NULL | 475.75 | 348.02 | 0.45 | 4.76E-02 |
| ENSMUSG00000062012 | GSMG0017194 | Zfp13 | zinc finger protein 13 | 435.43 | 546.68 | -0.33 | 1.76E-01 |
| ENSMUSG00000023903 | GSMG0017195 | Mmp25 | matrix metallopeptidase 25 | 324.56 | 401.09 | -0.30 | 1.85E-01 |
| ENSMUSG00000041319 | GSMG0017196 | Thoc6 | THO complex 6 | 352.33 | 535.09 | -0.60 | 1.61E-03 |
| ENSMUSG00000023905 | GSMG0017197 | Tnfrsf12a | tumor necrosis factor receptor superfamily, member 12a | 1388.03 | 428.23 | 1.70 | 5.24E-25 |
| ENSMUSG00000023909 | GSMG0017201 | Paqr4 | progestin and adipoQ receptor family member IV | 1947.48 | 1918.95 | 0.02 | 9.46E-01 |
| ENSMUSG00000040097 | GSMG0017203 | Flywch1 | FLYWCH-type zinc finger 1 | 1351.14 | 1263.41 | 0.10 | 6.91E-01 |
| ENSMUSG00000023911 | GSMG0017204 | Flywch2 | FLYWCH family member 2 | 130.56 | 110.62 | 0.23 | 5.61E-01 |
| ENSMUSG00000055839 | GSMG0017205 | Tceb2 | transcription elongation factor B (SIII), polypeptide 2 | 3081.42 | 3926.39 | -0.35 | 2.00E-01 |
| ENSMUSG00000016946 | GSMG0017212 | Kctd5 | potassium channel tetramerisation domain containing 5 | 1530.79 | 1926.70 | -0.33 | 5.13E-02 |
| ENSMUSG00000024122 | GSMG0017213 | Pdpk1 | 3-phosphoinositide dependent protein kinase 1 | 519.30 | 626.26 | -0.27 | 2.78E-01 |
| ENSMUSG00000036820 | GSMG0017216 | Amdhd2 | amidohydrolase domain containing 2 | 487.32 | 409.04 | 0.25 | 3.04E-01 |
| ENSMUSG00000024118 | GSMG0017220 | 1600002H07Rik | RIKEN cDNA 1600002H07 gene | 306.42 | 339.15 | -0.14 | 6.59E-01 |
| ENSMUSG00000072082 | GSMG0017221 | Ccnf | cyclin F | 226.35 | 347.29 | -0.62 | 4.11E-03 |
| ENSMUSG00000073437 | GSMG0017223 | D330041H03Rik | RIKEN cDNA D330041H03 gene | 196.39 | 179.41 | 0.13 | 7.31E-01 |
| ENSMUSG00000024137 | GSMG0017224 | E4f1 | E4F transcription factor 1 | 1781.49 | 1941.48 | -0.12 | 6.73E-01 |
| NULL | GSMG0017225 | NULL | NULL | 51.80 | 54.84 | -0.09 | 8.99E-01 |
| ENSMUSG00000024142 | GSMG0017226 | Mlst8 | MTOR associated protein, LST8 homolog (S. cerevisiae) | 393.72 | 424.75 | -0.11 | 7.42E-01 |
| ENSMUSG00000002496 | GSMG0017228 | Tsc2 | tuberous sclerosis 2 | 4603.12 | 4415.58 | 0.06 | 8.52E-01 |
| ENSMUSG00000002504 | GSMG0017229 | Slc9a3r2 | solute carrier family 9 (sodium/hydrogen exchanger), member 3 regulat | 17664.94 | 10671.52 | 0.73 | 1.73E-07 |
| ENSMUSG00000007021 | GSMG0017231 | Syngr3 | synaptogyrin 3 | 2121.83 | 2919.59 | -0.46 | 2.76E-02 |
| ENSMUSG00000040888 | GSMG0017232 | Gfer | growth factor, erv1 (S. cerevisiae)-like (augmenter of liver regeneration | 354.64 | 533.07 | -0.59 | 1.81E-03 |
| ENSMUSG00000040688 | GSMG0017233 | Tbl3 | transducin (beta)-like 3 | 1429.32 | 1926.22 | -0.43 | 4.69E-02 |
| ENSMUSG00000040048 | GSMG0017235 | Ndufb10 | NADH dehydrogenase (ubiquinone) 1 beta subcomplex, 10 | 1992.21 | 1944.19 | 0.03 | 9.15E-01 |
| ENSMUSG00000045316 | GSMG0017236 | Fahd1 | fumarylacetoacetate hydrolase domain containing 1 | 299.94 | 247.44 | 0.28 | 3.12E-01 |
| ENSMUSG00000039183 | GSMG0017237 | Nubp2 | nucleotide binding protein 2 | 674.29 | 672.15 | 0.01 | 9.85E-01 |
| G00000024163,ENSMUSG000 | GSMG0017238 | Eme2 // Mapk8ip3 | essential meiotic structure-specific endonuclease subunit 2 // mitogen-a | 3265.35 | 3025.14 | 0.11 | 7.37E-01 |
| ENSMUSG00000024165 | GSMG0017239 | Hn1l | hematological and neurological expressed 1-like | 674.23 | 1411.60 | -1.07 | 5.60E-13 |
| ENSMUSG00000024168 | GSMG0017242 | Tmem204 | transmembrane protein 204 | 3723.12 | 3067.14 | 0.28 | 1.24E-01 |
| ENSMUSG00000024170 | GSMG0017243 | Telo2 | telomere maintenance 2 | 167.96 | 215.78 | -0.36 | 2.42E-01 |
| ENSMUSG00000035521 | GSMG0017245 | Gnptg | N-acetylglucosamine-1-phosphotransferase, gamma subunit | 3685.34 | 3174.37 | 0.22 | 2.06E-01 |
| ENSMUSG00000047507 | GSMG0017246 | Baiap3 | BAI1-associated protein 3 | 502.20 | 368.71 | 0.44 | 2.56E-01 |

| | | | | | | | |
|---|---|---|---|---|---|---|---|
| ENSMUSG00000024112 | GSMG0017249 | Cacna1h | calcium channel, voltage-dependent, T type, alpha 1H subunit | 543.42 | 680.31 | -0.32 | 2.72E-01 |
| ENSMUSG00000024176 | GSMG0017251 | Sox8 | SRY (sex determining region Y)-box 8 | 491.60 | 196.30 | 1.32 | 1.15E-08 |
| ENSMUSG00000019214 | GSMG0017253 | Chtf18 | CTF18, chromosome transmission fidelity factor 18 | 271.12 | 468.67 | -0.79 | 1.29E-04 |
| ENSMUSG00000063011 | GSMG0017254 | Msln | mesothelin | 407.22 | 225.74 | 0.85 | 1.99E-03 |
| ENSMUSG00000061046 | GSMG0017255 | Haghl | hydroxyacylglutathione hydrolase-like | 1692.93 | 1527.85 | 0.15 | 5.42E-01 |
| ENSMUSG00000057411 | GSMG0017256 | Fam173a | family with sequence similarity 173, member A | 1132.01 | 817.93 | 0.47 | 9.49E-03 |
| ENSMUSG00000002274 | GSMG0017257 | Metrn | meteorin, glial cell differentiation regulator | 217.65 | 212.95 | 0.03 | 9.48E-01 |
| ENSMUSG00000039615 | GSMG0017259 | Stub1 | STIP1 homology and U-Box containing protein 1 | 1871.36 | 1946.02 | -0.06 | 8.26E-01 |
| ENSMUSG00000025732 | GSMG0017261 | Fam195a | family with sequence similarity 195, member A | 102.51 | 106.81 | -0.06 | 9.07E-01 |
| ENSMUSG00000025730 | GSMG0017263 | Rab40c | Rab40C, member RAS oncogene family | 254.56 | 256.55 | -0.01 | 9.79E-01 |
| ENSMUSG00000025728 | GSMG0017265 | Pigq | phosphatidylinositol glycan anchor biosynthesis, class Q | 1980.20 | 1905.85 | 0.06 | 8.50E-01 |
| ENSMUSG00000037326 | GSMG0017268 | Capn15 | calpain 15 | 1148.16 | 1085.93 | 0.08 | 8.51E-01 |
| ENSMUSG00000037098 | GSMG0017270 | Rab11fip3 | RAB11 family interacting protein 3 (class II) | 707.06 | 745.09 | -0.07 | 8.15E-01 |
| ENSMUSG00000036775 | GSMG0017273 | Decr2 | 2-4-dienoyl-Coenzyme A reductase 2, peroxisomal | 328.15 | 323.35 | 0.02 | 9.56E-01 |
| ENSMUSG00000024177 | GSMG0017274 | Nme4 | NME/NM23 nucleoside diphosphate kinase 4 | 128.48 | 239.55 | -0.90 | 1.12E-04 |
| ENSMUSG00000073433 | GSMG0017276 | Arhgdig | Rho GDP dissociation inhibitor (GDI) gamma | 1690.78 | 2458.99 | -0.54 | 1.93E-03 |
| ENSMUSG00000024187 | GSMG0017277 | Fam234a | family with sequence similarity 234, member A | 1213.76 | 1024.23 | 0.25 | 2.54E-01 |
| ENSMUSG00000024190 | GSMG0017278 | Dusp1 | dual specificity phosphatase 1 | 4407.27 | 1077.41 | 2.03 | 2.74E-45 |
| ENSMUSG00000024194 | GSMG0017282 | Cuta | cutA divalent cation tolerance homolog | 1059.04 | 958.44 | 0.14 | 6.33E-01 |
| ENSMUSG00000024208 | GSMG0017285 | Uqcc2 | ubiquinol-cytochrome c reductase complex assembly factor 2 | 1103.17 | 1244.55 | -0.18 | 5.20E-01 |
| ENSMUSG00000044857 | GSMG0017287 | Lemd2 | LEM domain containing 2 | 781.15 | 626.79 | 0.32 | 3.02E-01 |
| ENSMUSG00000062753 | GSMG0017292 | AI413582 | expressed sequence AI413582 | 703.01 | 638.75 | 0.14 | 6.02E-01 |
| ENSMUSG00000024213 | GSMG0017293 | Nudt3 | nudix (nucleotide diphosphate linked moiety X)-type motif 3 | 2699.26 | 3519.12 | -0.38 | 2.71E-02 |
| ENSMUSG00000056692 | GSMG0017298 | D17Wsu92e | DNA segment, Chr 17, Wayne State University 92, expressed | 150.47 | 155.32 | -0.04 | 9.31E-01 |
| ENSMUSG00000024218 | GSMG0017299 | Taf11 | TATA-box binding protein associated factor 11 | 897.02 | 1115.47 | -0.32 | 1.46E-01 |
| NULL | GSMG0017302 | NULL | NULL | 738.31 | 423.19 | 0.80 | 2.87E-05 |
| NULL | GSMG0017303 | NULL | NULL | 118.04 | 145.37 | -0.31 | 4.15E-01 |
| ENSMUSG00000024222 | GSMG0017306 | Fkbp5 | FK506 binding protein 5 | 219.77 | 349.37 | -0.67 | 2.14E-03 |
| ENSMUSG00000004865 | GSMG0017314 | Srpk1 | serine/arginine-rich protein specific kinase 1 | 710.05 | 802.55 | -0.18 | 4.51E-01 |
| ENSMUSG00000024006 | GSMG0017321 | Stk38 | serine/threonine kinase 38 | 4617.07 | 3600.76 | 0.36 | 2.63E-02 |
| ENSMUSG00000024008 | GSMG0017325 | Cpne5 | copine V | 107.87 | 160.86 | -0.58 | 9.41E-02 |
| ENSMUSG00000024007 | GSMG0017326 | Ppil1 | peptidylprolyl isomerase (cyclophilin)-like 1 | 695.50 | 1112.70 | -0.68 | 7.55E-05 |
| ENSMUSG00000024012 | GSMG0017328 | Mtch1 | mitochondrial carrier 1 | 3367.09 | 3817.72 | -0.18 | 3.89E-01 |

| | | | | | | | |
|---|---|---|---|---|---|---|---|
| ENSMUSG00000024018 | GSMG0017336 | Ccdc167 | coiled-coil domain containing 167 | 613.33 | 512.50 | 0.26 | 2.97E-01 |
| ENSMUSG00000043557 | GSMG0017337 | Mdga1 | MAM domain containing glycosylphosphatidylinositol anchor 1 | 337.64 | 1017.09 | -1.59 | 4.97E-12 |
| NULL | GSMG0017338 | NULL | NULL | 34.62 | 169.15 | -2.29 | 1.60E-12 |
| ENSMUSG00000062202 | GSMG0017339 | Btbd9 | BTB (POZ) domain containing 9 | 873.36 | 1094.40 | -0.32 | 1.49E-01 |
| NULL | GSMG0017341 | NULL | NULL | 203.38 | 178.26 | 0.19 | 6.12E-01 |
| NULL | GSMG0017342 | NULL | NULL | 489.22 | 407.78 | 0.27 | 3.78E-01 |
| ENSMUSG00000024026 | GSMG0017344 | Glo1 | glyoxalase 1 | 1797.32 | 1714.35 | 0.07 | 8.13E-01 |
| ENSMUSG00000024033 | GSMG0017353 | Rsph1 | radial spoke head 1 homolog (Chlamydomonas) | 156.22 | 96.39 | 0.70 | 9.92E-02 |
| ENSMUSG00000024037 | GSMG0017358 | Wdr4 | WD repeat domain 4 | 368.65 | 429.14 | -0.22 | 3.69E-01 |
| ENSMUSG00000024039 | GSMG0017360 | Cbs | cystathionine beta-synthase | 392.89 | 371.55 | 0.08 | 8.24E-01 |
| ENSMUSG00000024042 | GSMG0017362 | Sik1 | salt inducible kinase 1 | 966.25 | 673.75 | 0.52 | 6.38E-02 |
| ENSMUSG00000038146 | GSMG0017366 | Notch3 | notch 3 | 6148.97 | 4610.91 | 0.42 | 5.56E-01 |
| ENSMUSG00000024002 | GSMG0017368 | Brd4 | bromodomain containing 4 | 2353.61 | 2630.87 | -0.16 | 4.39E-01 |
| ENSMUSG00000024045 | GSMG0017370 | Akap8 | A kinase (PRKA) anchor protein 8 | 4058.16 | 4414.65 | -0.12 | 5.68E-01 |
| ENSMUSG00000002625 | GSMG0017371 | Akap8l | A kinase (PRKA) anchor protein 8-like | 7679.77 | 4120.59 | 0.90 | 5.35E-09 |
| ENSMUSG00000024050 | GSMG0017374 | Wiz | widely-interspaced zinc finger motifs | 1028.76 | 1354.28 | -0.40 | 1.56E-01 |
| ENSMUSG00000024298 | GSMG0017380 | Zfp871 | zinc finger protein 871 | 3379.65 | 3779.27 | -0.16 | 4.53E-01 |
| ENSMUSG00000095253 | GSMG0017384 | Zfp799 | zinc finger protein 799 | 816.15 | 755.63 | 0.11 | 6.56E-01 |
| ENSMUSG00000024055 | GSMG0017388 | Cyp4f13 | cytochrome P450, family 4, subfamily f, polypeptide 13 | 385.22 | 253.68 | 0.60 | 4.14E-03 |
| ENSMUSG00000067430 | GSMG0017390 | Zfp763 | zinc finger protein 763 | 344.09 | 363.69 | -0.08 | 8.23E-01 |
| ENSMUSG00000003929 | GSMG0017397 | Zfp81 | zinc finger protein 81 | 243.73 | 229.92 | 0.09 | 8.35E-01 |
| ENSMUSG00000055240 | GSMG0017399 | Zfp101 | zinc finger protein 101 | 367.89 | 382.54 | -0.05 | 8.90E-01 |
| ENSMUSG00000002289 | GSMG0017406 | Angptl4 | angiopoietin-like 4 | 1933.48 | 2669.04 | -0.46 | 9.88E-03 |
| NULL | GSMG0017412 | NULL | NULL | 80.58 | 72.21 | 0.15 | 7.89E-01 |
| ENSMUSG00000024309 | GSMG0017415 | Pfdn6 | prefoldin subunit 6 | 646.64 | 774.93 | -0.26 | 2.89E-01 |
| ENSMUSG00000067370 | GSMG0017416 | B3galt4 | UDP-Gal:betaGlcNAc beta 1,3-galactosyltransferase, polypeptide 4 | 186.20 | 210.16 | -0.18 | 6.12E-01 |
| ENSMUSG00000061232 | GSMG0017418 | H2-K1 | histocompatibility 2, K1, K region | 1909.37 | 2164.19 | -0.18 | 5.36E-01 |
| ENSMUSG00000024325 | GSMG0017419 | Ring1 | ring finger protein 1 | 392.79 | 266.19 | 0.56 | 1.74E-02 |
| G00000024327,ENSMUSG000 | GSMG0017420 | H2-Ke6 // Slc39a7 | H2-K region expressed gene 6 // solute carrier family 39 (zinc transport | 2471.78 | 2728.03 | -0.14 | 5.44E-01 |
| ENSMUSG00000024335 | GSMG0017422 | Brd2 | bromodomain containing 2 | 2333.55 | 2572.67 | -0.14 | 5.79E-01 |
| ENSMUSG00000096727 | GSMG0017423 | Psmb9 | proteasome (prosome, macropain) subunit, beta type 9 (large multifunct | 176.54 | 208.70 | -0.24 | 5.58E-01 |
| NULL | GSMG0017424 | NULL | NULL | 113.49 | 63.93 | 0.83 | 1.24E-02 |
| ENSMUSG00000015478 | GSMG0017429 | Rnf5 | ring finger protein 5 | 1282.87 | 1860.18 | -0.54 | 3.36E-04 |

| | | | | | | | |
|---|---|---|---|---|---|---|---|
| ENSMUSG00000015467 | GSMG0017430 | Egfl8 | EGF-like domain 8 | 1743.57 | 1188.80 | 0.55 | 3.80E-02 |
| ENSMUSG00000015474 | GSMG0017431 | Ppt2 | palmitoyl-protein thioesterase 2 | 945.15 | 917.79 | 0.04 | 9.03E-01 |
| NULL | GSMG0017433 | D17H6S56E-5 | DNA segment, Chr 17, human D6S56E 5 | 725.66 | 902.24 | -0.31 | 3.51E-01 |
| ENSMUSG00000073418 | GSMG0017434 | C4b | complement component 4B (Chido blood group) | 678.51 | 301.79 | 1.17 | 9.25E-06 |
| ENSMUSG00000061207 | GSMG0017437 | Stk19 | serine/threonine kinase 19 | 501.14 | 455.30 | 0.14 | 5.89E-01 |
| ENSMUSG00000040356 | GSMG0017438 | Skiv2l | superkiller viralicidic activity 2-like (S. cerevisiae) | 5244.85 | 5042.27 | 0.06 | 8.48E-01 |
| ENSMUSG00000092203 | GSMG0017444 | 1110038B12Rik | RIKEN cDNA 1110038B12 gene | 116.81 | 195.60 | -0.74 | 3.84E-03 |
| ENSMUSG00000024393 | GSMG0017458 | Prrc2a | proline-rich coiled-coil 2A | 813.10 | 984.54 | -0.28 | 4.88E-01 |
| ENSMUSG00000024397 | GSMG0017459 | Aif1 | allograft inflammatory factor 1 | 186.26 | 145.29 | 0.36 | 3.51E-01 |
| ENSMUSG00000073412 | GSMG0017460 | Lst1 | leukocyte specific transcript 1 | 40.58 | 59.48 | -0.55 | 2.48E-01 |
| ENSMUSG00000042419 | GSMG0017463 | Nfkbil1 | nuclear factor of kappa light polypeptide gene enhancer in B cells inhib | 248.62 | 319.56 | -0.36 | 1.58E-01 |
| NULL | GSMG0017466 | NULL | NULL | 229.34 | 390.23 | -0.77 | 1.33E-03 |
| ENSMUSG00000050410 | GSMG0017468 | Tcf19 | transcription factor 19 | 799.55 | 1319.33 | -0.72 | 5.83E-05 |
| ENSMUSG00000038838 | GSMG0017471 | Vars2 | valyl-tRNA synthetase 2, mitochondrial | 579.35 | 693.03 | -0.26 | 2.72E-01 |
| ENSMUSG00000001524 | GSMG0017472 | Gtf2h4 | general transcription factor II H, polypeptide 4 | 409.77 | 437.47 | -0.09 | 7.72E-01 |
| ENSMUSG00000003534 | GSMG0017473 | Ddr1 | discoidin domain receptor family, member 1 | 810.99 | 675.88 | 0.26 | 4.62E-01 |
| ENSMUSG00000001525 | GSMG0017474 | Tubb5 | tubulin, beta 5 class I | 4823.90 | 11481.59 | -1.25 | 5.40E-16 |
| NULL | GSMG0017476 | NULL | NULL | 258.07 | 353.86 | -0.45 | 6.70E-02 |
| ENSMUSG00000050705 | GSMG0017479 | 2310061I04Rik | RIKEN cDNA 2310061I04 gene | 732.78 | 585.24 | 0.32 | 1.07E-01 |
| ENSMUSG00000024426 | GSMG0017480 | Atat1 | alpha tubulin acetyltransferase 1 | 533.80 | 1270.88 | -1.25 | 3.10E-13 |
| ENSMUSG00000024436 | GSMG0017481 | Mrps18b | mitochondrial ribosomal protein S18B | 222.93 | 263.36 | -0.24 | 4.10E-01 |
| ENSMUSG00000038762 | GSMG0017482 | Abcf1 | ATP-binding cassette, sub-family F (GCN20), member 1 | 3161.88 | 3859.26 | -0.29 | 8.77E-02 |
| ENSMUSG00000053835 | GSMG0017486 | H2-T24 | histocompatibility 2, T region locus 24 | 197.58 | 53.85 | 1.88 | 1.99E-11 |
| ENSMUSG00000024446 | GSMG0017498 | Rpp21 | ribonuclease P 21 subunit | 808.26 | 923.79 | -0.19 | 4.07E-01 |
| ENSMUSG00000045409 | GSMG0017499 | Trim39 | tripartite motif-containing 39 | 1090.45 | 926.25 | 0.24 | 2.70E-01 |
| ENSMUSG00000036398 | GSMG0017509 | Ppp1r11 | protein phosphatase 1, regulatory (inhibitor) subunit 11 | 1284.57 | 1382.05 | -0.11 | 6.88E-01 |
| ENSMUSG00000036315 | GSMG0017510 | Znrd1 | zinc ribbon domain containing, 1 | 438.33 | 439.72 | -0.01 | 9.86E-01 |
| ENSMUSG00000076439 | GSMG0017514 | Mog | myelin oligodendrocyte glycoprotein | 411.70 | 14.33 | 4.84 | 1.16E-57 |
| NULL | GSMG0017538 | NULL | NULL | 866.36 | 938.39 | -0.12 | 7.76E-01 |
| ENSMUSG00000023919 | GSMG0017545 | Cenpq | centromere protein Q | 98.13 | 205.13 | -1.07 | 2.11E-04 |
| ENSMUSG00000061665 | GSMG0017550 | Cd2ap | CD2-associated protein | 1553.93 | 1286.82 | 0.27 | 2.83E-01 |
| ENSMUSG00000023912 | GSMG0017560 | Slc25a27 | solute carrier family 25, member 27 | 165.73 | 178.24 | -0.11 | 7.99E-01 |
| ENSMUSG00000023961 | GSMG0017562 | Enpp4 | ectonucleotide pyrophosphatase/phosphodiesterase 4 | 276.99 | 165.26 | 0.75 | 7.67E-04 |

| | | | | | | | |
|---|---|---|---|---|---|---|---|
| ENSMUSG00000023932 | GSMG0017571 | Cdc5l | cell division cycle 5-like (S. pombe) | 1153.07 | 1405.07 | -0.28 | 2.46E-01 |
| ENSMUSG00000096847 | GSMG0017575 | Tmem151b | transmembrane protein 151B | 782.57 | 883.62 | -0.18 | 5.63E-01 |
| ENSMUSG00000023944 | GSMG0017576 | Hsp90ab1 | heat shock protein 90 alpha (cytosolic), class B member 1 | 19395.12 | 23755.97 | -0.29 | 9.38E-02 |
| ENSMUSG00000023942 | GSMG0017577 | Slc29a1 | solute carrier family 29 (nucleoside transporters), member 1 | 2994.72 | 5952.24 | -0.99 | 4.29E-13 |
| ENSMUSG00000036026 | GSMG0017580 | Tmem63b | transmembrane protein 63b | 1797.56 | 2616.04 | -0.54 | 4.27E-03 |
| ENSMUSG00000023951 | GSMG0017585 | Vegfa | vascular endothelial growth factor A | 1301.58 | 1039.63 | 0.32 | 1.90E-01 |
| ENSMUSG00000023966 | GSMG0017587 | Rsph9 | radial spoke head 9 homolog (Chlamydomonas) | 266.84 | 381.52 | -0.51 | 4.20E-02 |
| ENSMUSG00000034509 | GSMG0017588 | Mad2l1bp | MAD2L1 binding protein | 516.25 | 483.44 | 0.09 | 7.54E-01 |
| ENSMUSG00000023953 | GSMG0017589 | Polh | polymerase (DNA directed), eta (RAD 30 related) | 79.29 | 102.81 | -0.36 | 3.95E-01 |
| ENSMUSG00000067148 | GSMG0017592 | Polr1c | polymerase (RNA) I polypeptide C | 594.24 | 758.23 | -0.35 | 6.12E-02 |
| ENSMUSG00000012296 | GSMG0017593 | Tjap1 | tight junction associated protein 1 | 1227.14 | 1086.27 | 0.18 | 3.93E-01 |
| ENSMUSG00000032842 | GSMG0017594 | Abcc10 | ATP-binding cassette, sub-family C (CFTR/MRP), member 10 | 375.69 | 312.40 | 0.27 | 3.89E-01 |
| ENSMUSG00000040327 | GSMG0017600 | Cul9 | cullin 9 | 538.21 | 411.05 | 0.39 | 2.42E-01 |
| ENSMUSG00000015605 | GSMG0017601 | Srf | serum response factor | 225.29 | 138.85 | 0.70 | 2.59E-02 |
| ENSMUSG00000023972 | GSMG0017602 | Ptk7 | PTK7 protein tyrosine kinase 7 | 233.26 | 330.40 | -0.50 | 1.83E-01 |
| ENSMUSG00000003546 | GSMG0017604 | Klc4 | kinesin light chain 4 | 1212.94 | 1003.19 | 0.27 | 2.37E-01 |
| ENSMUSG00000023971 | GSMG0017606 | Rrp36 | ribosomal RNA processing 36 homolog (S. cerevisiae) | 503.81 | 549.64 | -0.13 | 6.38E-01 |
| ENSMUSG00000063576 | GSMG0017607 | Klhdc3 | kelch domain containing 3 | 2248.51 | 2468.20 | -0.13 | 5.13E-01 |
| ENSMUSG00000059409 | GSMG0017608 | Ppp2r5d | protein phosphatase 2, regulatory subunit B', delta | 1631.22 | 1911.08 | -0.23 | 2.91E-01 |
| ENSMUSG00000023973 | GSMG0017611 | Cnpy3 | canopy FGF signaling regulator 3 | 1926.16 | 2071.77 | -0.10 | 6.53E-01 |
| ENSMUSG00000063888 | GSMG0017613 | Rpl7l1 | ribosomal protein L7-like 1 | 1704.55 | 2085.66 | -0.29 | 1.60E-01 |
| ENSMUSG00000036568 | GSMG0017614 | Gltscr1l | GLTSCR1-like | 140.89 | 111.85 | 0.33 | 3.79E-01 |
| NULL | GSMG0017616 | NULL | NULL | 377.85 | 249.49 | 0.60 | 6.31E-02 |
| ENSMUSG00000023977 | GSMG0017617 | Ubr2 | ubiquitin protein ligase E3 component n-recognin 2 | 1088.41 | 1184.59 | -0.12 | 7.39E-01 |
| ENSMUSG00000023980 | GSMG0017621 | Taf8 | TATA-box binding protein associated factor 8 | 301.12 | 268.10 | 0.17 | 6.07E-01 |
| ENSMUSG00000023988 | GSMG0017622 | Bysl | bystin-like | 235.76 | 337.92 | -0.51 | 7.15E-02 |
| ENSMUSG00000033475 | GSMG0017624 | Tomm6 | translocase of outer mitochondrial membrane 6 homolog (yeast) | 1265.24 | 1608.13 | -0.35 | 2.70E-01 |
| ENSMUSG00000023991 | GSMG0017628 | Foxp4 | forkhead box P4 | 182.76 | 289.88 | -0.67 | 7.03E-02 |
| ENSMUSG00000040260 | GSMG0017638 | Daam2 | dishevelled associated activator of morphogenesis 2 | 895.29 | 649.27 | 0.46 | 9.92E-02 |
| ENSMUSG00000039316 | GSMG0017640 | Rftn1 | raftlin lipid raft linker 1 | 654.84 | 657.44 | 0.00 | 9.88E-01 |
| ENSMUSG00000023923 | GSMG0017646 | Tbc1d5 | TBC1 domain family, member 5 | 425.65 | 786.53 | -0.88 | 1.77E-05 |
| NULL | GSMG0017656 | NULL | NULL | 159.45 | 104.56 | 0.61 | 6.75E-02 |
| ENSMUSG00000023927 | GSMG0017658 | Satb1 | special AT-rich sequence binding protein 1 | 674.10 | 919.00 | -0.45 | 2.93E-02 |

| | | | | | | | |
|---|---|---|---|---|---|---|---|
| NULL | GSMG0017660 | NULL | NULL | 276.76 | 323.54 | -0.23 | 5.14E-01 |
| NULL | GSMG0017661 | NULL | NULL | 167.02 | 255.00 | -0.61 | 4.31E-02 |
| NULL | GSMG0017662 | NULL | NULL | 237.41 | 333.65 | -0.49 | 1.14E-01 |
| ENSMUSG00000057835 | GSMG0017675 | Zfp119a | zinc finger protein 119a | 385.85 | 346.36 | 0.16 | 6.17E-01 |
| ENSMUSG00000062101 | GSMG0017676 | Zfp119b | zinc finger protein 119b | 230.94 | 159.50 | 0.54 | 3.83E-02 |
| ENSMUSG00000038781 | GSMG0017679 | Stap2 | signal transducing adaptor family member 2 | 2863.07 | 2847.52 | 0.01 | 9.79E-01 |
| ENSMUSG00000003200 | GSMG0017681 | Sh3gl1 | SH3-domain GRB2-like 1 | 2513.82 | 2381.31 | 0.08 | 7.37E-01 |
| ENSMUSG00000019578 | GSMG0017682 | Ubxn6 | UBX domain protein 6 | 894.28 | 842.98 | 0.08 | 7.40E-01 |
| NULL | GSMG0017683 | NULL | NULL | 133.50 | 146.24 | -0.13 | 7.41E-01 |
| ENSMUSG00000001227 | GSMG0017687 | Sema6b | sema domain, transmembrane domain (TM), and cytoplasmic domain, ( | 971.31 | 979.51 | -0.01 | 9.79E-01 |
| ENSMUSG00000019579 | GSMG0017688 | Mydgf | myeloid derived growth factor | 2069.13 | 2393.78 | -0.21 | 3.11E-01 |
| ENSMUSG00000001229 | GSMG0017689 | Dpp9 | dipeptidylpeptidase 9 | 2168.35 | 2060.98 | 0.07 | 8.06E-01 |
| ENSMUSG00000024197 | GSMG0017692 | Plin3 | perilipin 3 | 1309.32 | 1090.38 | 0.26 | 1.78E-01 |
| ENSMUSG00000013236 | GSMG0017694 | Ptprs | protein tyrosine phosphatase, receptor type, S | 4197.50 | 5508.67 | -0.39 | 1.26E-01 |
| NULL | GSMG0017695 | NULL | NULL | 2660.48 | 2720.67 | -0.03 | 9.41E-01 |
| ENSMUSG00000042625 | GSMG0017698 | Safb2 | scaffold attachment factor B2 | 2914.21 | 2360.41 | 0.30 | 6.72E-02 |
| ENSMUSG00000049760 | GSMG0017699 | 2410015M20Rik | RIKEN cDNA 2410015M20 gene | 1141.37 | 1292.37 | -0.18 | 4.14E-01 |
| ENSMUSG00000041168 | GSMG0017700 | Lonp1 | lon peptidase 1, mitochondrial | 2615.61 | 2361.54 | 0.15 | 5.75E-01 |
| NULL | GSMG0017701 | NULL | NULL | 231.71 | 278.71 | -0.27 | 3.23E-01 |
| NULL | GSMG0017702 | NULL | NULL | 88.37 | 107.68 | -0.29 | 4.72E-01 |
| ENSMUSG00000054723 | GSMG0017703 | Vmac | vimentin-type intermediate filament associated coiled-coil protein | 831.91 | 808.00 | 0.04 | 8.92E-01 |
| ENSMUSG00000024212 | GSMG0017708 | Mllt1 | myeloid/lymphoid or mixed-lineage leukemia%3B translocated to, 1 | 732.91 | 967.80 | -0.40 | 3.46E-02 |
| ENSMUSG00000002658 | GSMG0017712 | Gtf2f1 | general transcription factor IIF, polypeptide 1 | 863.74 | 957.28 | -0.15 | 5.29E-01 |
| ENSMUSG00000007670 | GSMG0017713 | Khsrp | KH-type splicing regulatory protein | 651.97 | 712.95 | -0.13 | 7.82E-01 |
| ENSMUSG00000046329 | GSMG0017715 | Slc25a23 | solute carrier family 25 (mitochondrial carrier%3B phosphate carrier), r | 2508.71 | 2100.57 | 0.26 | 3.18E-01 |
| ENSMUSG00000062591 | GSMG0017717 | Tubb4a | tubulin, beta 4A class IVA | 3358.08 | 3310.02 | 0.02 | 9.50E-01 |
| ENSMUSG00000005823 | GSMG0017722 | Gpr108 | G protein-coupled receptor 108 | 1639.50 | 1213.48 | 0.43 | 7.77E-03 |
| ENSMUSG00000024228 | GSMG0017726 | Nudt12 | nudix (nucleoside diphosphate linked moiety X)-type motif 12 | 272.78 | 221.70 | 0.30 | 3.18E-01 |
| ENSMUSG00000048915 | GSMG0017728 | Efna5 | ephrin A5 | 414.13 | 735.21 | -0.83 | 1.44E-03 |
| NULL | GSMG0017736 | NULL | NULL | 594.15 | 938.46 | -0.66 | 8.46E-05 |
| NULL | GSMG0017739 | NULL | NULL | 108.67 | 105.68 | 0.04 | 9.46E-01 |
| NULL | GSMG0017741 | NULL | NULL | 468.01 | 520.48 | -0.15 | 6.30E-01 |
| NULL | GSMG0017742 | NULL | NULL | 535.15 | 519.27 | 0.05 | 9.16E-01 |

| | | | | | | | |
|---|---|---|---|---|---|---|---|
| NULL | GSMG0017743 | NULL | NULL | 333.78 | 505.97 | -0.60 | 1.75E-02 |
| ENSMUSG00000023965 | GSMG0017747 | Fbxl17 | F-box and leucine-rich repeat protein 17 | 780.73 | 623.50 | 0.32 | 3.01E-01 |
| ENSMUSG00000024083 | GSMG0017762 | Pja2 | praja 2, RING-H2 motif containing | 3572.88 | 3959.47 | -0.15 | 4.95E-01 |
| ENSMUSG00000024091 | GSMG0017770 | Vapa | vesicle-associated membrane protein, associated protein A | 1664.48 | 1956.72 | -0.23 | 2.93E-01 |
| ENSMUSG00000056515 | GSMG0017773 | Rab31 | RAB31, member RAS oncogene family | 4235.29 | 5248.52 | -0.31 | 4.84E-02 |
| ENSMUSG00000024096 | GSMG0017776 | Ralbp1 | ralA binding protein 1 | 1027.90 | 888.81 | 0.21 | 4.29E-01 |
| ENSMUSG00000024098 | GSMG0017778 | Twsg1 | twisted gastrulation BMP signaling modulator 1 | 830.72 | 755.25 | 0.14 | 6.15E-01 |
| NULL | GSMG0017782 | NULL | NULL | 97.98 | 63.61 | 0.62 | 8.63E-02 |
| ENSMUSG00000024099 | GSMG0017784 | Ndufv2 | NADH dehydrogenase (ubiquinone) flavoprotein 2 | 1977.33 | 2334.83 | -0.24 | 1.71E-01 |
| ENSMUSG00000052105 | GSMG0017788 | Mtcl1 | microtubule crosslinking factor 1 | 999.30 | 1427.55 | -0.52 | 2.21E-02 |
| NULL | GSMG0017790 | NULL | NULL | 333.19 | 294.34 | 0.18 | 5.95E-01 |
| ENSMUSG00000023460 | GSMG0017791 | Rab12 | RAB12, member RAS oncogene family | 3870.57 | 4329.16 | -0.16 | 4.63E-01 |
| ENSMUSG00000033278 | GSMG0017795 | Ptprm | protein tyrosine phosphatase, receptor type, M | 8546.09 | 6886.80 | 0.31 | 8.01E-02 |
| NULL | GSMG0017799 | NULL | NULL | 161.59 | 149.70 | 0.11 | 7.85E-01 |
| ENSMUSG00000024043 | GSMG0017808 | Arhgap28 | Rho GTPase activating protein 28 | 288.26 | 633.58 | -1.14 | 2.21E-09 |
| ENSMUSG00000047407 | GSMG0017822 | Tgif1 | TGFB-induced factor homeobox 1 | 326.37 | 322.03 | 0.02 | 9.58E-01 |
| ENSMUSG00000034868 | GSMG0017827 | Myl12b | myosin, light chain 12B, regulatory | 12641.62 | 9620.63 | 0.39 | 2.42E-02 |
| ENSMUSG00000024048 | GSMG0017828 | Myl12a | myosin, light chain 12A, regulatory, non-sarcomeric | 14218.37 | 12011.33 | 0.24 | 3.21E-01 |
| ENSMUSG00000024056 | GSMG0017835 | Ndc80 | NDC80 kinetochore complex component | 235.92 | 463.93 | -0.97 | 1.13E-04 |
| ENSMUSG00000085492 | GSMG0017836 | Trmt61b | tRNA methyltransferase 61B | 213.28 | 238.50 | -0.16 | 6.37E-01 |
| ENSMUSG00000024064 | GSMG0017847 | Galnt14 | UDP-N-acetyl-alpha-D-galactosamine:polypeptide N-acetylgalactosami | 178.43 | 179.13 | 0.00 | 9.98E-01 |
| NULL | GSMG0017850 | NULL | NULL | 462.78 | 256.26 | 0.85 | 2.89E-05 |
| ENSMUSG00000058704 | GSMG0017855 | Memo1 | mediator of cell motility 1 | 533.95 | 508.31 | 0.07 | 8.19E-01 |
| ENSMUSG00000024067 | GSMG0017858 | Dpy30 | dpy-30, histone methyltransferase complex regulatory subunit | 940.09 | 1179.58 | -0.33 | 1.95E-01 |
| ENSMUSG00000002017 | GSMG0017862 | Fam98a | family with sequence similarity 98, member A | 434.95 | 403.46 | 0.11 | 7.55E-01 |
| ENSMUSG00000056121 | GSMG0017864 | Fez2 | fasciculation and elongation protein zeta 2 (zygin II) | 1044.48 | 1018.44 | 0.04 | 9.03E-01 |
| NULL | GSMG0017865 | NULL | NULL | 500.28 | 636.84 | -0.35 | 2.01E-01 |
| ENSMUSG00000024077 | GSMG0017866 | Strn | striatin, calmodulin binding protein | 247.22 | 225.64 | 0.13 | 7.09E-01 |
| ENSMUSG00000039414 | GSMG0017870 | Heatr5b | HEAT repeat containing 5B | 310.63 | 318.04 | -0.03 | 9.31E-01 |
| ENSMUSG00000024079 | GSMG0017872 | Eif2ak2 | eukaryotic translation initiation factor 2-alpha kinase 2 | 113.99 | 121.82 | -0.10 | 8.51E-01 |
| ENSMUSG00000024081 | GSMG0017874 | Cebpz | CCAAT/enhancer binding protein zeta | 1079.87 | 1203.65 | -0.16 | 5.93E-01 |
| ENSMUSG00000024070 | GSMG0017875 | Prkd3 | protein kinase D3 | 583.55 | 530.79 | 0.14 | 6.74E-01 |
| ENSMUSG00000036533 | GSMG0017876 | Cdc42ep3 | CDC42 effector protein (Rho GTPase binding) 3 | 2474.57 | 2406.83 | 0.04 | 8.99E-01 |

| | | | | | | | |
|---|---|---|---|---|---|---|---|
| ENSMUSG00000059811 | GSMG0017880 | Atl2 | atlastin GTPase 2 | 792.22 | 950.31 | -0.26 | 2.46E-01 |
| ENSMUSG00000024095 | GSMG0017884 | Hnrnpll | heterogeneous nuclear ribonucleoprotein L-like | 1542.80 | 1776.03 | -0.20 | 3.22E-01 |
| ENSMUSG00000024097 | GSMG0017886 | Srsf7 | serine/arginine-rich splicing factor 7 | 5786.60 | 5949.46 | -0.04 | 8.99E-01 |
| ENSMUSG00000035051 | GSMG0017887 | Dhx57 | DEAH (Asp-Glu-Ala-Asp/His) box polypeptide 57 | 641.88 | 847.92 | -0.40 | 2.18E-02 |
| ENSMUSG00000024241 | GSMG0017890 | Sos1 | son of sevenless homolog 1 (Drosophila) | 621.16 | 566.51 | 0.13 | 6.34E-01 |
| ENSMUSG00000024242 | GSMG0017892 | Map4k3 | mitogen-activated protein kinase kinase kinase kinase 3 | 1192.38 | 1013.30 | 0.23 | 2.81E-01 |
| ENSMUSG00000024246 | GSMG0017894 | Thumpd2 | THUMP domain containing 2 | 242.96 | 279.27 | -0.20 | 5.25E-01 |
| ENSMUSG00000054640 | GSMG0017897 | Slc8a1 | solute carrier family 8 (sodium/calcium exchanger), member 1 | 418.43 | 672.42 | -0.68 | 9.50E-04 |
| NULL | GSMG0017899 | NULL | NULL | 63.36 | 125.15 | -0.98 | 2.02E-03 |
| NULL | GSMG0017900 | NULL | NULL | 214.40 | 277.79 | -0.37 | 2.11E-01 |
| NULL | GSMG0017902 | NULL | NULL | 222.20 | 302.21 | -0.45 | 1.04E-01 |
| NULL | GSMG0017903 | NULL | NULL | 338.70 | 528.72 | -0.64 | 2.70E-03 |
| NULL | GSMG0017906 | NULL | NULL | 349.84 | 502.83 | -0.52 | 8.22E-02 |
| NULL | GSMG0017907 | NULL | NULL | 786.60 | 1255.84 | -0.67 | 1.02E-03 |
| NULL | GSMG0017908 | NULL | NULL | 1325.69 | 2080.46 | -0.65 | 1.31E-04 |
| ENSMUSG00000045817 | GSMG0017923 | Zfp36l2 | zinc finger protein 36, C3H type-like 2 | 1425.99 | 1227.39 | 0.21 | 5.42E-01 |
| ENSMUSG00000024251 | GSMG0017924 | Thada | thyroid adenoma associated | 118.66 | 132.51 | -0.15 | 7.27E-01 |
| ENSMUSG00000024120 | GSMG0017930 | Lrpprc | leucine-rich PPR-motif containing | 2346.35 | 2700.61 | -0.20 | 2.59E-01 |
| NULL | GSMG0017933 | NULL | NULL | 799.79 | 635.58 | 0.33 | 8.77E-02 |
| ENSMUSG00000024127 | GSMG0017934 | Prepl | prolyl endopeptidase-like | 883.66 | 883.28 | 0.00 | 9.96E-01 |
| NULL | GSMG0017941 | NULL | NULL | 149.01 | 114.20 | 0.38 | 2.97E-01 |
| ENSMUSG00000024145 | GSMG0017949 | Pigf | phosphatidylinositol glycan anchor biosynthesis, class F | 268.45 | 206.36 | 0.38 | 1.92E-01 |
| ENSMUSG00000024150 | GSMG0017952 | Mcfd2 | multiple coagulation factor deficiency 2 | 1964.47 | 1469.07 | 0.42 | 1.39E-02 |
| ENSMUSG00000036438 | GSMG0017957 | Calm2 | calmodulin 2 | 12616.76 | 16696.77 | -0.40 | 1.62E-02 |
| ENSMUSG00000050138 | GSMG0017958 | Kcnk12 | potassium channel, subfamily K, member 12 | 202.00 | 185.74 | 0.12 | 7.83E-01 |
| ENSMUSG00000005371 | GSMG0017959 | Fbxo11 | F-box protein 11 | 192.89 | 239.84 | -0.31 | 2.53E-01 |
| NULL | GSMG0017964 | NULL | NULL | 155.38 | 97.81 | 0.66 | 9.01E-02 |
| ENSMUSG00000024109 | GSMG0017970 | Nrxn1 | neurexin I | 1383.89 | 1735.07 | -0.33 | 1.20E-01 |
| NULL | GSMG0017972 | NULL | NULL | 178.65 | 192.66 | -0.11 | 7.86E-01 |
| NULL | GSMG0017974 | NULL | NULL | 283.62 | 285.22 | -0.01 | 9.88E-01 |
| NULL | GSMG0017979 | NULL | NULL | 290.59 | 364.33 | -0.32 | 3.47E-01 |
| NULL | GSMG0017980 | NULL | NULL | 945.08 | 1132.51 | -0.26 | 3.18E-01 |
| NULL | GSMG0017986 | NULL | NULL | 467.97 | 475.51 | -0.02 | 9.57E-01 |

| | | | | | | | |
|---|---|---|---|---|---|---|---|
| NULL | GSMG0017989 | NULL | NULL | 597.25 | 776.26 | -0.38 | 1.11E-01 |
| NULL | GSMG0017990 | NULL | NULL | 545.53 | 568.10 | -0.06 | 8.80E-01 |
| NULL | GSMG0017992 | NULL | NULL | 466.75 | 498.83 | -0.10 | 7.93E-01 |
| ENSMUSG00000055660 | GSMG0017995 | Mettl4 | methyltransferase like 4 | 158.66 | 177.37 | -0.16 | 6.62E-01 |
| ENSMUSG00000024231 | GSMG0018003 | Cul2 | cullin 2 | 1386.81 | 1630.89 | -0.23 | 2.28E-01 |
| ENSMUSG00000024232 | GSMG0018005 | Bambi | BMP and activin membrane-bound inhibitor | 2537.60 | 4668.94 | -0.88 | 6.49E-10 |
| ENSMUSG00000024234 | GSMG0018009 | Mtpap | mitochondrial poly(A) polymerase | 1040.25 | 1014.84 | 0.04 | 9.27E-01 |
| ENSMUSG00000033960 | GSMG0018010 | 9430020K01Rik | RIKEN cDNA 9430020K01 gene | 27127.82 | 15943.46 | 0.77 | 1.71E-08 |
| ENSMUSG00000024236 | GSMG0018014 | Svil | supervillin | 1044.85 | 1107.72 | -0.08 | 7.77E-01 |
| ENSMUSG00000024238 | GSMG0018021 | Zeb1 | zinc finger E-box binding homeobox 1 | 431.01 | 488.00 | -0.18 | 4.83E-01 |
| NULL | GSMG0018022 | NULL | NULL | 185.50 | 146.60 | 0.35 | 2.89E-01 |
| NULL | GSMG0018023 | NULL | NULL | 224.41 | 150.96 | 0.57 | 4.80E-02 |
| ENSMUSG00000073639 | GSMG0018031 | Rab18 | RAB18, member RAS oncogene family | 4960.08 | 4405.51 | 0.17 | 4.65E-01 |
| NULL | GSMG0018032 | NULL | NULL | 239.19 | 175.19 | 0.45 | 7.41E-02 |
| NULL | GSMG0018033 | NULL | NULL | 170.87 | 90.33 | 0.91 | 9.73E-03 |
| ENSMUSG00000024283 | GSMG0018036 | Wac | WW domain containing adaptor with coiled-coil | 2136.57 | 2345.36 | -0.13 | 5.11E-01 |
| NULL | GSMG0018037 | NULL | NULL | 232.91 | 205.45 | 0.18 | 6.49E-01 |
| ENSMUSG00000036103 | GSMG0018042 | Colec12 | collectin sub-family member 12 | 2982.59 | 2152.79 | 0.47 | 4.03E-03 |
| ENSMUSG00000024287 | GSMG0018048 | Thoc1 | THO complex 1 | 1204.88 | 1621.17 | -0.43 | 5.27E-02 |
| ENSMUSG00000002477 | GSMG0018057 | Snrpd1 | small nuclear ribonucleoprotein D1 | 543.64 | 968.71 | -0.83 | 1.50E-07 |
| ENSMUSG00000024294 | GSMG0018060 | Mib1 | mindbomb E3 ubiquitin protein ligase 1 | 384.43 | 302.48 | 0.35 | 3.75E-01 |
| ENSMUSG00000041238 | GSMG0018065 | Rbbp8 | retinoblastoma binding protein 8 | 501.13 | 692.34 | -0.47 | 5.78E-02 |
| ENSMUSG00000040957 | GSMG0018068 | Cables1 | CDK5 and Abl enzyme substrate 1 | 201.67 | 108.84 | 0.89 | 6.18E-04 |
| ENSMUSG00000024404 | GSMG0018071 | Riok3 | RIO kinase 3 | 1397.59 | 1379.15 | 0.02 | 9.47E-01 |
| ENSMUSG00000024410 | GSMG0018072 | 3110002H16Rik | RIKEN cDNA 3110002H16 gene | 1077.44 | 1069.59 | 0.01 | 9.71E-01 |
| ENSMUSG00000024424 | GSMG0018075 | Ttc39c | tetratricopeptide repeat domain 39C | 113.30 | 137.82 | -0.28 | 4.81E-01 |
| ENSMUSG00000024430 | GSMG0018076 | Cabyr | calcium-binding tyrosine-(Y)-phosphorylation regulated (fibrousheathi | 72.71 | 83.34 | -0.19 | 7.15E-01 |
| NULL | GSMG0018077 | NULL | NULL | 66.54 | 43.36 | 0.61 | 3.52E-01 |
| ENSMUSG00000024423 | GSMG0018079 | Impact | impact, RWD domain protein | 1547.33 | 1185.15 | 0.38 | 3.11E-02 |
| ENSMUSG00000036743 | GSMG0018085 | Psma8 | proteasome (prosome, macropain) subunit, alpha type, 8 | 123.57 | 116.48 | 0.08 | 8.70E-01 |
| ENSMUSG00000061808 | GSMG0018104 | Ttr | transthyretin | 292.19 | 14897.55 | -5.67 | 5.23E-03 |
| ENSMUSG00000033107 | GSMG0018105 | Rnf125 | ring finger protein 125 | 596.05 | 271.55 | 1.14 | 1.74E-06 |
| ENSMUSG00000024317 | GSMG0018108 | Rnf138 | ring finger protein 138 | 403.22 | 431.09 | -0.10 | 7.69E-01 |

| | | | | | | | |
|---|---|---|---|---|---|---|---|
| ENSMUSG00000024302 | GSMG0018121 | Dtna | dystrobrevin alpha | 712.96 | 864.37 | -0.28 | 2.69E-01 |
| ENSMUSG00000024277 | GSMG0018126 | Mapre2 | microtubule-associated protein, RP/EB family, member 2 | 1076.11 | 1596.10 | -0.57 | 5.11E-04 |
| ENSMUSG00000024276 | GSMG0018130 | Zfp397 | zinc finger protein 397 | 326.58 | 397.80 | -0.29 | 2.29E-01 |
| ENSMUSG00000063281 | GSMG0018132 | Zfp35 | zinc finger protein 35 | 650.71 | 621.90 | 0.07 | 8.58E-01 |
| ENSMUSG00000000420 | GSMG0018133 | Galnt1 | UDP-N-acetyl-alpha-D-galactosamine:polypeptide N-acetylgalactosami | 2906.30 | 2672.33 | 0.12 | 5.87E-01 |
| ENSMUSG00000024273 | GSMG0018135 | 2700062C07Rik | RIKEN cDNA 2700062C07 gene | 348.15 | 439.43 | -0.33 | 1.50E-01 |
| ENSMUSG00000024271 | GSMG0018136 | Elp2 | elongator acetyltransferase complex subunit 2 | 2377.40 | 2775.37 | -0.22 | 2.00E-01 |
| ENSMUSG00000034295 | GSMG0018138 | Fhod3 | formin homology 2 domain containing 3 | 218.68 | 363.65 | -0.74 | 3.33E-03 |
| NULL | GSMG0018144 | NULL | NULL | 328.47 | 307.97 | 0.09 | 7.84E-01 |
| NULL | GSMG0018150 | NULL | NULL | 192.93 | 235.82 | -0.29 | 3.45E-01 |
| NULL | GSMG0018151 | NULL | NULL | 614.04 | 1114.01 | -0.86 | 1.40E-07 |
| ENSMUSG00000033628 | GSMG0018158 | Pik3c3 | phosphoinositide-3-kinase, class 3 | 909.03 | 1021.69 | -0.17 | 4.32E-01 |
| ENSMUSG00000024260 | GSMG0018161 | Sap130 | Sin3A associated protein | 632.06 | 625.11 | 0.02 | 9.61E-01 |
| ENSMUSG00000041915 | GSMG0018162 | Ammecr1l | AMME chromosomal region gene 1-like | 669.86 | 717.89 | -0.10 | 7.34E-01 |
| ENSMUSG00000024258 | GSMG0018163 | Polr2d | polymerase (RNA) II (DNA directed) polypeptide D | 1122.25 | 1472.95 | -0.39 | 5.09E-02 |
| ENSMUSG00000024400 | GSMG0018164 | Wdr33 | WD repeat domain 33 | 851.73 | 920.16 | -0.11 | 6.49E-01 |
| ENSMUSG00000024395 | GSMG0018171 | Lims2 | LIM and senescent cell antigen like domains 2 | 2599.74 | 372.05 | 2.80 | 1.26E-82 |
| ENSMUSG00000024384 | GSMG0018173 | Iws1 | IWS1 homolog (S. cerevisiae) | 284.24 | 330.81 | -0.21 | 4.95E-01 |
| ENSMUSG00000024383 | GSMG0018177 | Map3k2 | mitogen-activated protein kinase kinase kinase 2 | 82.85 | 72.53 | 0.19 | 7.21E-01 |
| ENSMUSG00000024382 | GSMG0018181 | Ercc3 | excision repair cross-complementing rodent repair deficiency, complem | 619.05 | 772.55 | -0.32 | 1.50E-01 |
| ENSMUSG00000024381 | GSMG0018182 | Bin1 | bridging integrator 1 | 1344.22 | 1643.70 | -0.29 | 9.44E-02 |
| ENSMUSG00000038299 | GSMG0018186 | Wdr36 | WD repeat domain 36 | 784.88 | 1108.78 | -0.50 | 6.21E-03 |
| ENSMUSG00000038128 | GSMG0018187 | Camk4 | calcium/calmodulin-dependent protein kinase IV | 196.14 | 232.41 | -0.24 | 4.79E-01 |
| NULL | GSMG0018201 | NULL | NULL | 386.58 | 415.67 | -0.10 | 7.35E-01 |
| ENSMUSG00000005871 | GSMG0018205 | Apc | adenomatosis polyposis coli | 357.67 | 430.27 | -0.26 | 3.75E-01 |
| ENSMUSG00000014504 | GSMG0018209 | Srp19 | signal recognition particle 19 | 1486.66 | 1777.26 | -0.26 | 2.53E-01 |
| ENSMUSG00000003779 | GSMG0018212 | Kif20a | kinesin family member 20A | 603.96 | 1258.07 | -1.06 | 2.35E-07 |
| ENSMUSG00000034300 | GSMG0018214 | Fam53c | family with sequence similarity 53, member C | 364.35 | 390.71 | -0.10 | 7.35E-01 |
| ENSMUSG00000038773 | GSMG0018216 | Kdm3b | KDM3B lysine (K)-specific demethylase 3B | 755.04 | 731.19 | 0.05 | 8.87E-01 |
| NULL | GSMG0018217 | NULL | NULL | 286.13 | 284.24 | 0.01 | 9.89E-01 |
| ENSMUSG00000038555 | GSMG0018219 | Reep2 | receptor accessory protein 2 | 860.48 | 1081.32 | -0.33 | 9.82E-02 |
| ENSMUSG00000038418 | GSMG0018220 | Egr1 | early growth response 1 | 5071.26 | 1146.14 | 2.15 | 8.52E-44 |
| ENSMUSG00000037815 | GSMG0018225 | Ctnna1 | catenin (cadherin associated protein), alpha 1 | 9690.28 | 9164.55 | 0.08 | 7.20E-01 |

| | | | | | | | |
|---|---|---|---|---|---|---|---|
| ENSMUSG00000097769 | GSMG0018229 | Snhg4 | small nucleolar RNA host gene 4 | 184.68 | 238.13 | -0.37 | 1.81E-01 |
| ENSMUSG00000037236 | GSMG0018230 | Matr3 | matrin 3 | 2760.96 | 3814.49 | -0.47 | 1.77E-02 |
| ENSMUSG00000037058 | GSMG0018233 | Paip2 | polyadenylate-binding protein-interacting protein 2 | 1463.31 | 1477.20 | -0.01 | 9.64E-01 |
| NULL | GSMG0018234 | NULL | NULL | 149.30 | 155.61 | -0.06 | 8.92E-01 |
| ENSMUSG00000046668 | GSMG0018238 | Cxxc5 | CXXC finger 5 | 1174.29 | 1250.08 | -0.09 | 7.87E-01 |
| ENSMUSG00000024347 | GSMG0018240 | Psd2 | pleckstrin and Sec7 domain containing 2 | 161.62 | 259.58 | -0.69 | 3.85E-02 |
| ENSMUSG00000043991 | GSMG0018242 | Pura | purine rich element binding protein A | 1363.15 | 729.56 | 0.90 | 8.73E-05 |
| ENSMUSG00000046727 | GSMG0018246 | Cystm1 | cysteine-rich transmembrane module containing 1 | 526.45 | 265.87 | 0.99 | 5.88E-08 |
| ENSMUSG00000033272 | GSMG0018258 | Slc35a4 | solute carrier family 35, member A4 | 3226.83 | 3530.51 | -0.13 | 5.88E-01 |
| ENSMUSG00000006850 | GSMG0018259 | Tmco6 | transmembrane and coiled-coil domains 6 | 256.62 | 230.47 | 0.16 | 6.26E-01 |
| ENSMUSG00000024474 | GSMG0018260 | Ik | IK cytokine | 1359.60 | 1318.26 | 0.05 | 8.83E-01 |
| ENSMUSG00000042660 | GSMG0018261 | Wdr55 | WD repeat domain 55 | 117.48 | 147.66 | -0.32 | 4.01E-01 |
| ENSMUSG00000019143 | GSMG0018262 | Hars2 | histidyl-tRNA synthetase 2 | 810.60 | 951.50 | -0.23 | 2.92E-01 |
| ENSMUSG00000001383 | GSMG0018263 | Zmat2 | zinc finger, matrin type 2 | 934.90 | 1024.05 | -0.13 | 6.13E-01 |
| NULL | GSMG0018272 | NULL | NULL | 171.11 | 219.17 | -0.36 | 2.38E-01 |
| ENSMUSG00000044043 | GSMG0018281 | Pcdhb14 | protocadherin beta 14 | 338.78 | 261.87 | 0.37 | 1.54E-01 |
| ENSMUSG00000047910 | GSMG0018284 | Pcdhb16 | protocadherin beta 16 | 807.49 | 661.36 | 0.29 | 1.76E-01 |
| ENSMUSG00000046387 | GSMG0018285 | Pcdhb17 | protocadherin beta 17 | 645.06 | 725.17 | -0.17 | 4.83E-01 |
| ENSMUSG00000046191 | GSMG0018288 | Pcdhb20 | protocadherin beta 20 | 626.05 | 641.69 | -0.04 | 9.04E-01 |
| G00000103255,ENSMUSG000 | GSMG0018295 | Pcdha1 // Pcdha2 / | protocadherin alpha 1 // protocadherin alpha 2 // protocadherin alpha 3 | 13955.83 | 20620.60 | -0.56 | 2.54E-04 |
| ENSMUSG00000044024 | GSMG0018296 | Rell2 | RELT-like 2 | 332.93 | 274.04 | 0.28 | 3.23E-01 |
| ENSMUSG00000024442 | GSMG0018300 | 0610009O20Rik | RIKEN cDNA 0610009O20 gene | 285.49 | 296.83 | -0.06 | 8.92E-01 |
| ENSMUSG00000060450 | GSMG0018301 | Rnf14 | ring finger protein 14 | 3012.53 | 3278.37 | -0.12 | 5.57E-01 |
| NULL | GSMG0018302 | NULL | NULL | 133.22 | 105.61 | 0.33 | 4.29E-01 |
| ENSMUSG00000024425 | GSMG0018304 | Ndfip1 | Nedd4 family interacting protein 1 | 5136.00 | 7176.42 | -0.48 | 1.73E-02 |
| ENSMUSG00000036452 | GSMG0018311 | Arhgap26 | Rho GTPase activating protein 26 | 301.33 | 233.15 | 0.37 | 1.68E-01 |
| ENSMUSG00000071860 | GSMG0018316 | 2900055J20Rik | RIKEN cDNA 2900055J20 gene | 1105.49 | 697.56 | 0.66 | 5.34E-03 |
| ENSMUSG00000051401 | GSMG0018317 | Kctd16 | potassium channel tetramerisation domain containing 16 | 334.15 | 283.16 | 0.24 | 4.34E-01 |
| NULL | GSMG0018318 | NULL | NULL | 501.46 | 298.78 | 0.74 | 1.47E-03 |
| NULL | GSMG0018319 | NULL | NULL | 95.06 | 44.89 | 1.09 | 9.26E-03 |
| NULL | GSMG0018320 | NULL | NULL | 397.91 | 231.31 | 0.78 | 1.42E-03 |
| NULL | GSMG0018324 | NULL | NULL | 75.47 | 96.39 | -0.35 | 4.06E-01 |
| ENSMUSG00000024491 | GSMG0018327 | Rbm27 | RNA binding motif protein 27 | 254.48 | 293.23 | -0.21 | 5.34E-01 |

| | | | | | | | |
|---|---|---|---|---|---|---|---|
| ENSMUSG00000024498 | GSMG0018329 | Tcerg1 | transcription elongation regulator 1 (CA150) | 1336.11 | 1877.41 | -0.49 | 1.28E-02 |
| NULL | GSMG0018332 | NULL | NULL | 45.99 | 470.74 | -3.36 | 2.79E-52 |
| ENSMUSG00000024472 | GSMG0018341 | Dcp2 | decapping mRNA 2 | 78.39 | 128.76 | -0.72 | 1.99E-02 |
| NULL | GSMG0018343 | NULL | NULL | 315.49 | 476.50 | -0.59 | 8.36E-03 |
| ENSMUSG00000034653 | GSMG0018346 | Ythdc2 | YTH domain containing 2 | 146.09 | 179.68 | -0.29 | 3.53E-01 |
| ENSMUSG00000054477 | GSMG0018350 | Kcnn2 | potassium intermediate/small conductance calcium-activated channel, s | 580.37 | 429.55 | 0.44 | 9.84E-02 |
| ENSMUSG00000057561 | GSMG0018354 | Eif1a | eukaryotic translation initiation factor 1A | 1484.12 | 1698.76 | -0.19 | 4.51E-01 |
| ENSMUSG00000042705 | GSMG0018361 | Commd10 | COMM domain containing 10 | 552.31 | 685.63 | -0.31 | 1.88E-01 |
| ENSMUSG00000037416 | GSMG0018372 | Dmxl1 | Dmx-like 1 | 101.53 | 88.00 | 0.21 | 6.47E-01 |
| ENSMUSG00000024507 | GSMG0018379 | Hsd17b4 | hydroxysteroid (17-beta) dehydrogenase 4 | 2753.98 | 3144.17 | -0.19 | 3.13E-01 |
| ENSMUSG00000024528 | GSMG0018388 | Srfbp1 | serum response factor binding protein 1 | 289.04 | 351.04 | -0.28 | 2.96E-01 |
| ENSMUSG00000024534 | GSMG0018393 | Sncaip | synuclein, alpha interacting protein (synphilin) | 317.77 | 508.14 | -0.67 | 7.19E-04 |
| ENSMUSG00000034484 | GSMG0018398 | Snx2 | sorting nexin 2 | 1685.12 | 1585.38 | 0.09 | 7.55E-01 |
| ENSMUSG00000024535 | GSMG0018399 | Snx24 | sorting nexing 24 | 657.29 | 525.95 | 0.32 | 1.18E-01 |
| NULL | GSMG0018408 | NULL | NULL | 98.06 | 73.84 | 0.40 | 3.47E-01 |
| ENSMUSG00000001700 | GSMG0018419 | Gramd3 | GRAM domain containing 3 | 406.35 | 414.87 | -0.03 | 9.41E-01 |
| ENSMUSG00000008301 | GSMG0018420 | Phax | phosphorylated adaptor for RNA export | 912.06 | 1090.39 | -0.26 | 1.92E-01 |
| ENSMUSG00000024590 | GSMG0018422 | Lmnb1 | lamin B1 | 647.68 | 1545.22 | -1.25 | 1.18E-11 |
| ENSMUSG00000024594 | GSMG0018427 | Prrc1 | proline-rich coiled-coil 1 | 743.69 | 995.59 | -0.42 | 1.88E-02 |
| ENSMUSG00000024597 | GSMG0018432 | Slc12a2 | solute carrier family 12, member 2 | 1337.79 | 1294.44 | 0.05 | 8.90E-01 |
| ENSMUSG00000024601 | GSMG0018437 | Isoc1 | isochorismatase domain containing 1 | 1216.74 | 1649.89 | -0.44 | 1.93E-02 |
| NULL | GSMG0018438 | NULL | NULL | 136.09 | 196.27 | -0.53 | 8.16E-02 |
| ENSMUSG00000050875 | GSMG0018442 | A730017C20Rik | RIKEN cDNA A730017C20 gene | 261.20 | 443.57 | -0.76 | 5.70E-05 |
| ENSMUSG00000058152 | GSMG0018444 | Chsy3 | chondroitin sulfate synthase 3 | 641.19 | 586.63 | 0.13 | 6.46E-01 |
| NULL | GSMG0018446 | NULL | NULL | 2439.33 | 1987.61 | 0.29 | 8.58E-02 |
| ENSMUSG00000024603 | GSMG0018453 | Dctn4 | dynactin 4 | 726.20 | 811.89 | -0.16 | 5.12E-01 |
| ENSMUSG00000024604 | GSMG0018454 | Rbm22 | RNA binding motif protein 22 | 746.60 | 834.75 | -0.16 | 4.97E-01 |
| ENSMUSG00000024608 | GSMG0018458 | Rps14 | ribosomal protein S14 | 2729.00 | 3687.03 | -0.43 | 2.61E-02 |
| ENSMUSG00000024610 | GSMG0018459 | Cd74 | CD74 antigen (invariant polypeptide of major histocompatibility compl | 145.79 | 100.47 | 0.54 | 1.28E-01 |
| ENSMUSG00000024617 | GSMG0018461 | Camk2a | calcium/calmodulin-dependent protein kinase II alpha | 1937.82 | 619.40 | 1.65 | 3.50E-16 |
| ENSMUSG00000024620 | GSMG0018462 | Pdgfrb | platelet derived growth factor receptor, beta polypeptide | 8174.20 | 8887.67 | -0.12 | 6.86E-01 |
| ENSMUSG00000024621 | GSMG0018463 | Csf1r | colony stimulating factor 1 receptor | 1895.26 | 689.68 | 1.46 | 5.75E-10 |
| ENSMUSG00000024576 | GSMG0018467 | Csnk1a1 | casein kinase 1, alpha 1 | 6549.56 | 8820.17 | -0.43 | 1.97E-02 |

| | | | | | | | |
|---|---|---|---|---|---|---|---|
| ENSMUSG00000098098 | GSMG0018470 | Bvht | braveheart long non-coding RNA | 550.20 | 464.55 | 0.24 | 3.18E-01 |
| NULL | GSMG0018472 | NULL | NULL | 132.03 | 115.20 | 0.20 | 6.18E-01 |
| ENSMUSG00000045629 | GSMG0018473 | Sh3tc2 | SH3 domain and tetratricopeptide repeats 2 | 1348.19 | 1418.85 | -0.07 | 7.98E-01 |
| ENSMUSG00000071847 | GSMG0018479 | Apcdd1 | adenomatosis polyposis coli down-regulated 1 | 26754.22 | 26648.41 | 0.01 | 9.86E-01 |
| NULL | GSMG0018481 | NULL | NULL | 942.53 | 730.79 | 0.37 | 6.51E-02 |
| ENSMUSG00000024581 | GSMG0018482 | Napg | N-ethylmaleimide sensitive fusion protein attachment protein gamma | 507.09 | 479.03 | 0.08 | 7.97E-01 |
| ENSMUSG00000040560 | GSMG0018483 | Wdr7 | WD repeat domain 7 | 532.38 | 580.51 | -0.12 | 6.88E-01 |
| ENSMUSG00000056812 | GSMG0018487 | St8sia3 | ST8 alpha-N-acetyl-neuraminide alpha-2,8-sialyltransferase 3 | 914.08 | 1067.51 | -0.22 | 3.51E-01 |
| NULL | GSMG0018495 | NULL | NULL | 247.41 | 213.82 | 0.21 | 5.35E-01 |
| ENSMUSG00000024589 | GSMG0018496 | Nedd4l | neural precursor cell expressed, developmentally down-regulated gene 4 | 2400.73 | 2665.49 | -0.15 | 4.64E-01 |
| ENSMUSG00000032688 | GSMG0018504 | Malt1 | MALT1 paracaspase | 241.51 | 221.94 | 0.12 | 7.32E-01 |
| ENSMUSG00000042439 | GSMG0018508 | Zfp532 | zinc finger protein 532 | 1108.73 | 1145.39 | -0.05 | 9.07E-01 |
| NULL | GSMG0018510 | NULL | NULL | 601.96 | 721.25 | -0.26 | 2.11E-01 |
| ENSMUSG00000024516 | GSMG0018513 | Sec11c | SEC11 homolog C, signal peptidase complex subunit | 1436.53 | 1943.13 | -0.44 | 2.39E-02 |
| ENSMUSG00000024517 | GSMG0018514 | Grp | gastrin releasing peptide | 218.52 | 494.05 | -1.18 | 3.33E-08 |
| ENSMUSG00000024521 | GSMG0018517 | Pmaip1 | phorbol-12-myristate-13-acetate-induced protein 1 | 2516.80 | 1205.05 | 1.06 | 2.91E-06 |
| ENSMUSG00000024524 | GSMG0018520 | Gnal | guanine nucleotide binding protein, alpha stimulating, olfactory type | 134.17 | 309.36 | -1.21 | 1.33E-06 |
| ENSMUSG00000109901 | GSMG0018524 | Chmp1b | charged multivesicular body protein 1B | 981.67 | 837.73 | 0.23 | 4.27E-01 |
| ENSMUSG00000024525 | GSMG0018527 | Impa2 | inositol (myo)-1(or 4)-monophosphatase 2 | 105.45 | 105.25 | 0.00 | 9.94E-01 |
| ENSMUSG00000001473 | GSMG0018529 | Tubb6 | tubulin, beta 6 class V | 2726.89 | 4588.35 | -0.75 | 3.75E-06 |
| NULL | GSMG0018530 | NULL | NULL | 85.57 | 83.48 | 0.03 | 9.69E-01 |
| ENSMUSG00000024530 | GSMG0018532 | Slmo1 | slowmo homolog 1 (Drosophila) | 242.82 | 261.66 | -0.11 | 8.00E-01 |
| ENSMUSG00000024537 | GSMG0018533 | Psmg2 | proteasome (prosome, macropain) assembly chaperone 2 | 870.21 | 973.70 | -0.16 | 5.38E-01 |
| ENSMUSG00000079614 | GSMG0018535 | Seh1l | SEH1-like (S. cerevisiae | 802.03 | 943.21 | -0.23 | 3.03E-01 |
| ENSMUSG00000024544 | GSMG0018539 | Ldlrad4 | low density lipoprotein receptor class A domain containing 4 | 601.69 | 852.16 | -0.50 | 7.15E-03 |
| NULL | GSMG0018540 | NULL | NULL | 387.56 | 416.13 | -0.10 | 7.58E-01 |
| NULL | GSMG0018541 | NULL | NULL | 317.30 | 334.55 | -0.07 | 8.43E-01 |
| NULL | GSMG0018543 | NULL | NULL | 72.58 | 83.98 | -0.21 | 7.27E-01 |
| NULL | GSMG0018548 | NULL | NULL | 122.45 | 154.62 | -0.34 | 2.98E-01 |
| ENSMUSG00000009535 | GSMG0018551 | Rnmt | RNA (guanine-7-) methyltransferase | 696.82 | 954.45 | -0.45 | 5.31E-02 |
| ENSMUSG00000053477 | GSMG0018559 | Tcf4 | transcription factor 4 | 7672.33 | 8722.55 | -0.19 | 3.01E-01 |
| NULL | GSMG0018560 | NULL | NULL | 737.84 | 872.34 | -0.24 | 3.03E-01 |
| NULL | GSMG0018564 | NULL | NULL | 733.38 | 1155.60 | -0.65 | 8.48E-04 |

| | | | | | | | |
|---|---|---|---|---|---|---|---|
| NULL | GSMG0018565 | NULL | NULL | 560.34 | 722.05 | -0.36 | 1.24E-01 |
| NULL | GSMG0018566 | NULL | NULL | 1735.03 | 2318.18 | -0.42 | 3.88E-02 |
| NULL | GSMG0018567 | NULL | NULL | 1352.16 | 1846.15 | -0.45 | 3.64E-02 |
| NULL | GSMG0018568 | NULL | NULL | 1080.56 | 1413.59 | -0.39 | 9.41E-02 |
| NULL | GSMG0018569 | NULL | NULL | 152.27 | 167.63 | -0.14 | 7.63E-01 |
| NULL | GSMG0018571 | NULL | NULL | 548.92 | 661.42 | -0.27 | 3.64E-01 |
| NULL | GSMG0018572 | NULL | NULL | 550.42 | 821.81 | -0.58 | 3.63E-03 |
| ENSMUSG00000024513 | GSMG0018578 | Mbd2 | methyl-CpG binding domain protein 2 | 1303.84 | 717.91 | 0.86 | 1.24E-01 |
| ENSMUSG00000037253 | GSMG0018583 | Mex3c | mex3 RNA binding family member C | 1290.37 | 857.74 | 0.59 | 5.69E-03 |
| NULL | GSMG0018591 | NULL | NULL | 237.22 | 65.64 | 1.85 | 1.47E-11 |
| ENSMUSG00000024560 | GSMG0018592 | Cxxc1 | CXXC finger 1 (PHD domain) | 1712.39 | 1684.11 | 0.02 | 9.34E-01 |
| ENSMUSG00000024561 | GSMG0018593 | Mbd1 | methyl-CpG binding domain protein 1 | 1184.64 | 1421.99 | -0.26 | 2.63E-01 |
| ENSMUSG00000025885 | GSMG0018596 | Myo5b | myosin VB | 149.00 | 171.76 | -0.20 | 5.79E-01 |
| ENSMUSG00000036880 | GSMG0018600 | Acaa2 | acetyl-Coenzyme A acyltransferase 2 (mitochondrial 3-oxoacyl-Coenzy | 2449.04 | 2975.74 | -0.28 | 1.38E-01 |
| ENSMUSG00000036299 | GSMG0018602 | BC031181 | cDNA sequence BC031181 | 2582.82 | 2414.81 | 0.10 | 7.40E-01 |
| ENSMUSG00000035765 | GSMG0018603 | Dym | dymeclin | 397.23 | 516.84 | -0.38 | 8.07E-02 |
| ENSMUSG00000025880 | GSMG0018605 | Smad7 | SMAD family member 7 | 861.88 | 396.24 | 1.12 | 1.06E-04 |
| ENSMUSG00000024563 | GSMG0018615 | Smad2 | SMAD family member 2 | 552.55 | 689.59 | -0.32 | 1.56E-01 |
| ENSMUSG00000090000 | GSMG0018621 | Ier3ip1 | immediate early response 3 interacting protein 1 | 1507.15 | 1485.47 | 0.02 | 9.64E-01 |
| ENSMUSG00000025421 | GSMG0018622 | Hdhd2 | haloacid dehalogenase-like hydrolase domain containing 2 | 579.56 | 569.11 | 0.03 | 9.41E-01 |
| ENSMUSG00000025423 | GSMG0018624 | Pias2 | protein inhibitor of activated STAT 2 | 1283.40 | 1481.27 | -0.21 | 3.69E-01 |
| ENSMUSG00000025425 | GSMG0018626 | St8sia5 | ST8 alpha-N-acetyl-neuraminide alpha-2,8-sialyltransferase 5 | 117.17 | 46.65 | 1.33 | 1.45E-04 |
| ENSMUSG00000025428 | GSMG0018631 | Atp5a1 | ATP synthase, H+ transporting, mitochondrial F1 complex, alpha subur | 11602.43 | 14794.39 | -0.35 | 3.12E-02 |
| ENSMUSG00000039840 | GSMG0018633 | Epg5 | ectopic P-granules autophagy protein 5 homolog (C. elegans) | 270.97 | 258.19 | 0.07 | 8.82E-01 |
| ENSMUSG00000056214 | GSMG0018638 | Pard6g | par-6 family cell polarity regulator gamma | 1765.95 | 2259.66 | -0.36 | 2.52E-02 |
| ENSMUSG00000057130 | GSMG0018640 | Txnl4a | thioredoxin-like 4A | 3713.34 | 4007.31 | -0.11 | 6.08E-01 |
| ENSMUSG00000034006 | GSMG0018641 | Pqlc1 | PQ loop repeat containing 1 | 651.97 | 537.54 | 0.28 | 2.25E-01 |
| ENSMUSG00000041607 | GSMG0018647 | Mbp | myelin basic protein | 42553.59 | 609.27 | 6.13 | 1.42E-221 |
| NULL | GSMG0018648 | NULL | NULL | 314.49 | 159.76 | 0.98 | 5.01E-03 |
| ENSMUSG00000058881 | GSMG0018652 | Zfp516 | zinc finger protein 516 | 250.16 | 195.87 | 0.35 | 3.56E-01 |
| ENSMUSG00000049090 | GSMG0018655 | Zadh2 | zinc binding alcohol dehydrogenase, domain containing 2 | 1495.45 | 1671.44 | -0.16 | 4.61E-01 |
| ENSMUSG00000024646 | GSMG0018658 | Cyb5a | cytochrome b5 type A (microsomal) | 2036.86 | 2005.93 | 0.02 | 9.41E-01 |
| ENSMUSG00000050321 | GSMG0018661 | Neto1 | neuropilin (NRP) and tolloid (TLL)-like 1 | 354.20 | 422.96 | -0.26 | 2.67E-01 |

| | | | | | | | |
|---|---|---|---|---|---|---|---|
| ENSMUSG00000024647 | GSMG0018664 | Cbln2 | cerebellin 2 precursor protein | 627.93 | 850.35 | -0.44 | 2.33E-02 |
| ENSMUSG00000024614 | GSMG0018668 | Tmx3 | thioredoxin-related transmembrane protein 3 | 966.51 | 999.55 | -0.05 | 8.74E-01 |
| ENSMUSG00000063889 | GSMG0018669 | Crem | cAMP responsive element modulator | 164.42 | 151.79 | 0.12 | 8.04E-01 |
| ENSMUSG00000024235 | GSMG0018670 | Map3k8 | mitogen-activated protein kinase kinase kinase 8 | 381.96 | 197.59 | 0.95 | 2.67E-06 |
| ENSMUSG00000041225 | GSMG0018682 | Arhgap12 | Rho GTPase activating protein 12 | 768.59 | 853.48 | -0.15 | 5.12E-01 |
| ENSMUSG00000024240 | GSMG0018689 | Epc1 | enhancer of polycomb homolog 1 (Drosophila) | 468.35 | 638.82 | -0.45 | 5.23E-02 |
| ENSMUSG00000061013 | GSMG0018691 | Mkx | mohawk homeobox | 88.33 | 99.06 | -0.17 | 7.82E-01 |
| ENSMUSG00000057440 | GSMG0018695 | Mpp7 | membrane protein, palmitoylated 7 (MAGUK p55 subfamily member 7 | 356.11 | 316.77 | 0.17 | 5.77E-01 |
| NULL | GSMG0018696 | NULL | NULL | 119.90 | 119.53 | 0.00 | 9.98E-01 |
| ENSMUSG00000024286 | GSMG0018698 | Ccny | cyclin Y | 593.99 | 516.56 | 0.20 | 4.42E-01 |
| ENSMUSG00000047879 | GSMG0018701 | Usp14 | ubiquitin specific peptidase 14 | 1214.16 | 1646.60 | -0.44 | 1.23E-02 |
| ENSMUSG00000024290 | GSMG0018702 | Rock1 | Rho-associated coiled-coil containing protein kinase 1 | 3106.21 | 2769.01 | 0.17 | 5.02E-01 |
| ENSMUSG00000024293 | GSMG0018707 | Esco1 | establishment of sister chromatid cohesion N-acetyltransferase 1 | 337.05 | 314.13 | 0.10 | 7.73E-01 |
| ENSMUSG00000002475 | GSMG0018710 | Abhd3 | abhydrolase domain containing 3 | 825.98 | 295.61 | 1.48 | 1.76E-16 |
| ENSMUSG00000049411 | GSMG0018717 | Tmem241 | transmembrane protein 241 | 247.65 | 160.91 | 0.62 | 1.74E-02 |
| ENSMUSG00000024413 | GSMG0018721 | Npc1 | Niemann-Pick type C1 | 1124.20 | 1145.87 | -0.03 | 9.29E-01 |
| ENSMUSG00000057766 | GSMG0018722 | Ankrd29 | ankyrin repeat domain 29 | 108.63 | 62.23 | 0.80 | 1.92E-02 |
| ENSMUSG00000044252 | GSMG0018724 | Osbpl1a | oxysterol binding protein-like 1A | 2371.52 | 1623.27 | 0.55 | 5.16E-04 |
| NULL | GSMG0018725 | NULL | NULL | 465.46 | 309.70 | 0.59 | 5.42E-03 |
| ENSMUSG00000024420 | GSMG0018727 | Zfp521 | zinc finger protein 521 | 1131.87 | 1624.05 | -0.52 | 1.45E-02 |
| NULL | GSMG0018729 | NULL | NULL | 277.52 | 383.54 | -0.47 | 9.32E-02 |
| ENSMUSG00000037013 | GSMG0018730 | Ss18 | synovial sarcoma translocation, Chromosome 18 | 911.22 | 758.58 | 0.27 | 2.24E-01 |
| ENSMUSG00000036225 | GSMG0018732 | Kctd1 | potassium channel tetramerisation domain containing 1 | 226.88 | 110.68 | 1.03 | 1.46E-05 |
| ENSMUSG00000024411 | GSMG0018737 | Aqp4 | aquaporin 4 | 713.84 | 720.37 | -0.01 | 9.80E-01 |
| ENSMUSG00000024304 | GSMG0018743 | Cdh2 | cadherin 2 | 3015.26 | 5311.93 | -0.82 | 8.06E-10 |
| NULL | GSMG0018747 | NULL | NULL | 354.75 | 434.80 | -0.29 | 3.48E-01 |
| NULL | GSMG0018748 | NULL | NULL | 1106.12 | 1647.62 | -0.58 | 1.28E-03 |
| ENSMUSG00000056124 | GSMG0018758 | B4galt6 | UDP-Gal:betaGlcNAc beta 1,4-galactosyltransferase, polypeptide 6 | 165.10 | 158.04 | 0.06 | 9.07E-01 |
| ENSMUSG00000033382 | GSMG0018759 | Trappc8 | trafficking protein particle complex 8 | 848.03 | 799.48 | 0.09 | 7.69E-01 |
| ENSMUSG00000042680 | GSMG0018761 | Garem | GRB2 associated, regulator of MAPK1 | 362.64 | 436.98 | -0.27 | 3.05E-01 |
| ENSMUSG00000041923 | GSMG0018769 | Nol4 | nucleolar protein 4 | 105.90 | 194.63 | -0.88 | 1.15E-03 |
| ENSMUSG00000051469 | GSMG0018777 | Zfp24 | zinc finger protein 24 | 641.60 | 693.27 | -0.11 | 6.86E-01 |
| ENSMUSG00000047989 | GSMG0018778 | Ino80c | INO80 complex subunit C | 748.46 | 791.46 | -0.08 | 8.13E-01 |

| | | | | | | | |
|---|---|---|---|---|---|---|---|
| ENSMUSG00000040446 | GSMG0018780 | Rprd1a | regulation of nuclear pre-mRNA domain containing 1A | 382.06 | 441.65 | -0.21 | 4.21E-01 |
| NULL | GSMG0018781 | NULL | NULL | 360.60 | 291.60 | 0.30 | 2.96E-01 |
| ENSMUSG00000024270 | GSMG0018782 | Slc39a6 | solute carrier family 39 (metal ion transporter), member 6 | 780.50 | 1664.95 | -1.09 | 1.36E-13 |
| ENSMUSG00000024269 | GSMG0018784 | Tpgs2 | tubulin polyglutamylase complex subunit 2 | 599.61 | 682.01 | -0.18 | 5.01E-01 |
| ENSMUSG00000024268 | GSMG0018785 | Celf4 | CUGBP, Elav-like family member 4 | 982.24 | 1420.83 | -0.53 | 2.76E-03 |
| NULL | GSMG0018786 | NULL | NULL | 265.53 | 457.56 | -0.79 | 3.90E-04 |
| NULL | GSMG0018787 | NULL | NULL | 152.64 | 221.42 | -0.54 | 1.10E-01 |
| NULL | GSMG0018788 | NULL | NULL | 270.12 | 318.66 | -0.24 | 3.85E-01 |
| NULL | GSMG0018789 | NULL | NULL | 247.73 | 265.90 | -0.10 | 7.88E-01 |
| ENSMUSG00000024261 | GSMG0018799 | Syt4 | synaptotagmin IV | 535.67 | 1542.66 | -1.52 | 1.73E-26 |
| ENSMUSG00000024259 | GSMG0018800 | Slc25a46 | solute carrier family 25, member 46 | 1939.88 | 1978.40 | -0.03 | 9.19E-01 |
| NULL | GSMG0018801 | NULL | NULL | 137.36 | 77.43 | 0.82 | 2.19E-02 |
| NULL | GSMG0018803 | NULL | NULL | 526.26 | 487.99 | 0.11 | 6.86E-01 |
| ENSMUSG00000044982 | GSMG0018804 | Sft2d3 | SFT2 domain containing 3 | 1290.99 | 1273.43 | 0.02 | 9.50E-01 |
| ENSMUSG00000052229 | GSMG0018805 | Gpr17 | G protein-coupled receptor 17 | 1617.12 | 993.16 | 0.70 | 2.07E-03 |
| ENSMUSG00000090523 | GSMG0018810 | Gypc | glycophorin C | 519.36 | 637.17 | -0.29 | 1.58E-01 |
| ENSMUSG00000024378 | GSMG0018814 | Stard4 | StAR-related lipid transfer (START) domain containing 4 | 1552.41 | 1997.25 | -0.36 | 5.50E-02 |
| ENSMUSG00000042834 | GSMG0018815 | Nrep | neuronal regeneration related protein | 18076.32 | 29548.91 | -0.71 | 3.93E-04 |
| ENSMUSG00000005873 | GSMG0018820 | Reep5 | receptor accessory protein 5 | 3807.69 | 5190.61 | -0.45 | 1.25E-02 |
| ENSMUSG00000036501 | GSMG0018822 | Fam13b | family with sequence similarity 13, member B | 406.30 | 437.22 | -0.11 | 7.53E-01 |
| ENSMUSG00000035984 | GSMG0018823 | Nme5 | NME/NM23 family member 5 | 101.36 | 95.69 | 0.08 | 8.87E-01 |
| ENSMUSG00000003778 | GSMG0018826 | Brd8 | bromodomain containing 8 | 768.71 | 976.89 | -0.34 | 5.47E-02 |
| ENSMUSG00000024370 | GSMG0018827 | Cdc23 | CDC23 cell division cycle 23 | 2574.69 | 2997.03 | -0.22 | 2.33E-01 |
| ENSMUSG00000044201 | GSMG0018829 | Cdc25c | cell division cycle 25C | 88.84 | 226.31 | -1.35 | 3.79E-07 |
| NULL | GSMG0018832 | NULL | NULL | 189.21 | 155.91 | 0.28 | 4.42E-01 |
| NULL | GSMG0018833 | NULL | NULL | 199.24 | 157.78 | 0.33 | 4.18E-01 |
| ENSMUSG00000024360 | GSMG0018835 | Etf1 | eukaryotic translation termination factor 1 | 881.53 | 1211.68 | -0.46 | 2.52E-02 |
| ENSMUSG00000024359 | GSMG0018836 | Hspa9 | heat shock protein 9 | 3251.62 | 4540.41 | -0.48 | 2.81E-03 |
| ENSMUSG00000071862 | GSMG0018837 | Lrrtm2 | leucine rich repeat transmembrane neuronal 2 | 533.36 | 545.57 | -0.03 | 9.30E-01 |
| ENSMUSG00000024357 | GSMG0018838 | Sil1 | endoplasmic reticulum chaperone SIL1 homolog (S. cerevisiae) | 770.54 | 908.27 | -0.24 | 2.50E-01 |
| ENSMUSG00000024352 | GSMG0018843 | Spata24 | spermatogenesis associated 24 | 92.15 | 97.57 | -0.09 | 8.82E-01 |
| ENSMUSG00000073599 | GSMG0018845 | Ecscr | endothelial cell surface expressed chemotaxis and apoptosis regulator | 1201.91 | 2058.88 | -0.78 | 1.71E-06 |
| ENSMUSG00000024349 | GSMG0018846 | Tmem173 | transmembrane protein 173 | 304.63 | 391.92 | -0.36 | 1.85E-01 |

| | | | | | | | |
|---|---|---|---|---|---|---|---|
| ENSMUSG00000060275 | GSMG0018848 | Nrg2 | neuregulin 2 | 280.08 | 243.34 | 0.20 | 7.08E-01 |
| ENSMUSG00000024346 | GSMG0018851 | Pfdn1 | prefoldin 1 | 1365.09 | 1742.86 | -0.35 | 7.85E-02 |
| ENSMUSG00000024486 | GSMG0018854 | Hbegf | heparin-binding EGF-like growth factor | 1516.91 | 1596.97 | -0.07 | 7.66E-01 |
| ENSMUSG00000006050 | GSMG0018855 | Sra1 | steroid receptor RNA activator 1 | 1440.27 | 1285.23 | 0.16 | 4.23E-01 |
| NULL | GSMG0018856 | Apbb3 | amyloid beta (A4) precursor protein-binding, family B, member 3 | 1162.13 | 969.33 | 0.26 | 2.60E-01 |
| ENSMUSG00000014294 | GSMG0018860 | Ndufa2 | NADH dehydrogenase (ubiquinone) 1 alpha subcomplex, 2 | 2328.26 | 2483.37 | -0.09 | 7.38E-01 |
| ENSMUSG00000001380 | GSMG0018862 | Hars | histidyl-tRNA synthetase | 1809.66 | 2063.31 | -0.19 | 3.92E-01 |
| NULL | GSMG0018875 | NULL | NULL | 1172.04 | 1835.35 | -0.65 | 8.81E-03 |
| NULL | GSMG0018876 | NULL | NULL | 168.28 | 222.10 | -0.40 | 1.78E-01 |
| ENSMUSG00000024456 | GSMG0018878 | Diaph1 | diaphanous related formin 1 | 597.20 | 660.78 | -0.15 | 5.87E-01 |
| ENSMUSG00000024454 | GSMG0018881 | Hdac3 | histone deacetylase 3 | 4363.70 | 3826.49 | 0.19 | 3.85E-01 |
| ENSMUSG00000038524 | GSMG0018882 | Fchsd1 | FCH and double SH3 domains 1 | 242.93 | 451.87 | -0.90 | 3.63E-06 |
| ENSMUSG00000024451 | GSMG0018883 | Arap3 | ArfGAP with RhoGAP domain, ankyrin repeat and PH domain 3 | 3576.99 | 3465.21 | 0.05 | 8.93E-01 |
| ENSMUSG00000024440 | GSMG0018886 | Pcdh12 | protocadherin 12 | 1000.88 | 1763.01 | -0.82 | 5.42E-07 |
| ENSMUSG00000052102 | GSMG0018887 | Gnpda1 | glucosamine-6-phosphate deaminase 1 | 159.57 | 113.03 | 0.49 | 1.59E-01 |
| ENSMUSG00000024427 | GSMG0018889 | Spry4 | sprouty homolog 4 (Drosophila) | 1241.42 | 1114.61 | 0.16 | 4.91E-01 |
| ENSMUSG00000036585 | GSMG0018890 | Fgf1 | fibroblast growth factor 1 | 426.68 | 295.22 | 0.53 | 2.51E-02 |
| ENSMUSG00000024431 | GSMG0018893 | Nr3c1 | nuclear receptor subfamily 3, group C, member 1 | 1043.23 | 829.30 | 0.33 | 6.71E-02 |
| ENSMUSG00000024487 | GSMG0018896 | Yipf5 | Yip1 domain family, member 5 | 721.43 | 777.51 | -0.11 | 6.96E-01 |
| ENSMUSG00000056671 | GSMG0018899 | Prelid2 | PRELI domain containing 2 | 115.85 | 96.30 | 0.27 | 6.09E-01 |
| ENSMUSG00000024493 | GSMG0018903 | Lars | leucyl-tRNA synthetase | 683.06 | 887.27 | -0.38 | 5.68E-02 |
| NULL | GSMG0018904 | NULL | NULL | 311.27 | 439.38 | -0.50 | 2.59E-02 |
| ENSMUSG00000024500 | GSMG0018908 | Ppp2r2b | protein phosphatase 2, regulatory subunit B, beta | 465.92 | 913.23 | -0.97 | 7.99E-09 |
| NULL | GSMG0018909 | NULL | NULL | 303.66 | 296.29 | 0.04 | 9.24E-01 |
| NULL | GSMG0018910 | NULL | NULL | 182.90 | 193.47 | -0.07 | 8.96E-01 |
| NULL | GSMG0018911 | NULL | NULL | 450.79 | 384.43 | 0.23 | 3.66E-01 |
| ENSMUSG00000024501 | GSMG0018916 | Dpysl3 | dihydropyrimidinase-like 3 | 586.46 | 6157.30 | -3.39 | 1.33E-124 |
| ENSMUSG00000024502 | GSMG0018919 | Jakmip2 | janus kinase and microtubule interacting protein 2 | 227.55 | 446.34 | -0.98 | 2.03E-06 |
| ENSMUSG00000071856 | GSMG0018925 | Mcc | mutated in colorectal cancers | 333.11 | 234.81 | 0.51 | 5.41E-02 |
| NULL | GSMG0018927 | NULL | NULL | 267.05 | 200.25 | 0.42 | 1.37E-01 |
| NULL | GSMG0018935 | NULL | NULL | 216.57 | 229.29 | -0.08 | 8.45E-01 |
| ENSMUSG00000024477 | GSMG0018936 | Pggt1b | protein geranylgeranyltransferase type I, beta subunit | 381.01 | 368.68 | 0.05 | 8.96E-01 |
| ENSMUSG00000071855 | GSMG0018938 | Ccdc112 | coiled-coil domain containing 112 | 198.91 | 431.08 | -1.12 | 7.26E-08 |

| | | | | | | | |
|---|---|---|---|---|---|---|---|
| ENSMUSG00000033319 | GSMG0018944 | Fem1c | fem-1 homolog c (C.elegans) | 177.72 | 235.66 | -0.41 | 1.56E-01 |
| ENSMUSG00000033184 | GSMG0018948 | Tmed7 | transmembrane emp24 protein transport domain containing 7 | 4758.78 | 4851.43 | -0.03 | 9.27E-01 |
| ENSMUSG00000033022 | GSMG0018949 | Cdo1 | cysteine dioxygenase 1, cytosolic | 220.29 | 399.89 | -0.86 | 6.30E-05 |
| ENSMUSG00000032905 | GSMG0018950 | Atg12 | autophagy related 12 | 1998.78 | 1963.82 | 0.03 | 9.32E-01 |
| ENSMUSG00000019647 | GSMG0018951 | Sema6a | sema domain, transmembrane domain (TM), and cytoplasmic domain, ( | 2902.87 | 3711.42 | -0.35 | 2.57E-02 |
| NULL | GSMG0018952 | NULL | NULL | 139.09 | 228.91 | -0.72 | 8.73E-03 |
| NULL | GSMG0018954 | NULL | NULL | 86.20 | 171.69 | -1.01 | 3.85E-03 |
| NULL | GSMG0018955 | NULL | NULL | 588.07 | 777.70 | -0.40 | 3.63E-02 |
| ENSMUSG00000024505 | GSMG0018958 | Dtwd2 | DTW domain containing 2 | 59.55 | 51.41 | 0.22 | 7.25E-01 |
| ENSMUSG00000024529 | GSMG0018964 | Lox | lysyl oxidase | 233.92 | 118.09 | 0.99 | 3.30E-03 |
| ENSMUSG00000024538 | GSMG0018967 | Ppic | peptidylprolyl isomerase C | 8704.03 | 10135.36 | -0.22 | 3.75E-01 |
| ENSMUSG00000048799 | GSMG0018969 | Cep120 | centrosomal protein 120 | 719.69 | 809.14 | -0.17 | 4.76E-01 |
| ENSMUSG00000052713 | GSMG0018974 | Zfp608 | zinc finger protein 608 | 578.45 | 734.22 | -0.35 | 1.40E-01 |
| NULL | GSMG0018975 | NULL | NULL | 256.50 | 236.97 | 0.11 | 7.49E-01 |
| ENSMUSG00000053644 | GSMG0018981 | Aldh7a1 | aldehyde dehydrogenase family 7, member A1 | 849.53 | 816.51 | 0.06 | 8.45E-01 |
| ENSMUSG00000032656 | GSMG0018983 | March3 | membrane-associated ring finger (C3HC4) 3 | 116.92 | 105.82 | 0.15 | 7.48E-01 |
| NULL | GSMG0018991 | NULL | NULL | 109.86 | 107.90 | 0.02 | 9.64E-01 |
| ENSMUSG00000024598 | GSMG0018997 | Fbn2 | fibrillin 2 | 90.56 | 332.18 | -1.87 | 3.22E-11 |
| ENSMUSG00000038059 | GSMG0019002 | Smim3 | small integral membrane protein 3 | 612.92 | 348.79 | 0.81 | 4.27E-05 |
| ENSMUSG00000043079 | GSMG0019008 | Synpo | synaptopodin | 668.64 | 254.12 | 1.40 | 2.73E-11 |
| ENSMUSG00000054008 | GSMG0019009 | Ndst1 | N-deacetylase/N-sulfotransferase (heparan glucosaminyl) 1 | 674.19 | 636.28 | 0.08 | 8.63E-01 |
| ENSMUSG00000024613 | GSMG0019013 | Tcof1 | treacle ribosome biogenesis factor 1 | 595.50 | 891.07 | -0.58 | 9.06E-04 |
| ENSMUSG00000052026 | GSMG0019014 | Slc6a7 | solute carrier family 6 (neurotransmitter transporter, L-proline), membe | 1161.62 | 891.70 | 0.38 | 1.21E-01 |
| ENSMUSG00000024622 | GSMG0019016 | Hmgxb3 | HMG box domain containing 3 | 317.56 | 274.61 | 0.21 | 4.77E-01 |
| ENSMUSG00000034320 | GSMG0019017 | Slc26a2 | solute carrier family 26 (sulfate transporter), member 2 | 233.64 | 272.70 | -0.23 | 4.72E-01 |
| ENSMUSG00000033871 | GSMG0019018 | Ppargc1b | peroxisome proliferative activated receptor, gamma, coactivator 1 beta | 540.54 | 331.62 | 0.71 | 2.14E-04 |
| NULL | GSMG0019023 | NULL | NULL | 3495.45 | 3770.35 | -0.11 | 6.11E-01 |
| ENSMUSG00000097324 | GSMG0019024 | Carmn | cardiac mesoderm enhancer-associated non-coding RNA | 132.09 | 166.67 | -0.33 | 3.69E-01 |
| ENSMUSG00000024579 | GSMG0019026 | Pcyox1l | prenylcysteine oxidase 1 like | 233.74 | 311.50 | -0.41 | 8.45E-02 |
| ENSMUSG00000024580 | GSMG0019027 | Grpel2 | GrpE-like 2, mitochondrial | 362.94 | 457.95 | -0.34 | 1.57E-01 |
| ENSMUSG00000033032 | GSMG0019028 | Afap1l1 | actin filament associated protein 1-like 1 | 9225.40 | 6969.80 | 0.40 | 5.31E-03 |
| ENSMUSG00000032735 | GSMG0019029 | Ablim3 | actin binding LIM protein family, member 3 | 196.92 | 479.16 | -1.29 | 5.42E-10 |
| NULL | GSMG0019031 | NULL | NULL | 63.02 | 75.76 | -0.27 | 5.86E-01 |

| | | | | | | | |
|---|---|---|---|---|---|---|---|
| ENSMUSG00000045730 | GSMG0019032 | Adrb2 | adrenergic receptor, beta 2 | 733.19 | 242.69 | 1.59 | 1.78E-18 |
| ENSMUSG00000042211 | GSMG0019035 | Fbxo38 | F-box protein 38 | 1928.65 | 2004.71 | -0.06 | 8.21E-01 |
| NULL | GSMG0019039 | NULL | NULL | 1940.40 | 1871.47 | 0.05 | 8.79E-01 |
| NULL | GSMG0019040 | NULL | NULL | 2578.52 | 2707.91 | -0.07 | 7.93E-01 |
| ENSMUSG00000024583 | GSMG0019045 | Txnl1 | thioredoxin-like 1 | 1658.66 | 2013.00 | -0.28 | 2.31E-01 |
| ENSMUSG00000024588 | GSMG0019049 | Fech | ferrochelatase | 1960.10 | 1936.42 | 0.02 | 9.51E-01 |
| ENSMUSG00000024587 | GSMG0019050 | Nars | asparaginyl-tRNA synthetase | 3017.02 | 3186.89 | -0.08 | 7.31E-01 |
| ENSMUSG00000041891 | GSMG0019060 | Lman1 | lectin, mannose-binding, 1 | 1775.07 | 2012.67 | -0.18 | 4.33E-01 |
| ENSMUSG00000046318 | GSMG0019061 | Ccbe1 | collagen and calcium binding EGF domains 1 | 47.72 | 259.01 | -2.44 | 3.47E-17 |
| ENSMUSG00000062526 | GSMG0019067 | Mppe1 | metallophosphoesterase 1 | 549.91 | 388.44 | 0.50 | 1.19E-02 |
| NULL | GSMG0019069 | NULL | NULL | 1585.83 | 2602.67 | -0.71 | 3.71E-05 |
| ENSMUSG00000024527 | GSMG0019070 | Afg3l2 | AFG3-like AAA ATPase 2 | 758.45 | 944.90 | -0.32 | 1.15E-01 |
| ENSMUSG00000024533 | GSMG0019072 | Spire1 | spire homolog 1 (Drosophila) | 799.36 | 679.21 | 0.24 | 3.69E-01 |
| ENSMUSG00000073542 | GSMG0019075 | Cep76 | centrosomal protein 76 | 238.34 | 213.27 | 0.16 | 6.52E-01 |
| ENSMUSG00000024539 | GSMG0019076 | Ptpn2 | protein tyrosine phosphatase, non-receptor type 2 | 437.82 | 521.03 | -0.25 | 3.16E-01 |
| NULL | GSMG0019077 | NULL | NULL | 293.19 | 167.10 | 0.81 | 1.69E-03 |
| NULL | GSMG0019078 | 4930549G23Rik | RIKEN cDNA 4930549G23 gene | 72.85 | 99.79 | -0.46 | 2.29E-01 |
| ENSMUSG00000038121 | GSMG0019082 | Fam210a | family with sequence similarity 210, member A | 143.37 | 159.48 | -0.15 | 7.19E-01 |
| ENSMUSG00000044906 | GSMG0019092 | 4930503L19Rik | RIKEN cDNA 4930503L19 gene | 233.61 | 243.76 | -0.06 | 8.87E-01 |
| ENSMUSG00000038425 | GSMG0019094 | Poli | polymerase (DNA directed), iota | 159.47 | 172.72 | -0.12 | 7.93E-01 |
| ENSMUSG00000060534 | GSMG0019099 | Dcc | deleted in colorectal carcinoma | 197.15 | 239.04 | -0.28 | 4.00E-01 |
| NULL | GSMG0019108 | NULL | NULL | 224.53 | 251.13 | -0.16 | 6.65E-01 |
| NULL | GSMG0019111 | NULL | NULL | 185.62 | 161.99 | 0.20 | 6.14E-01 |
| NULL | GSMG0019114 | NULL | NULL | 81.46 | 81.32 | 0.01 | 9.87E-01 |
| NULL | GSMG0019120 | NULL | NULL | 190.81 | 199.21 | -0.06 | 8.80E-01 |
| ENSMUSG00000024515 | GSMG0019128 | Smad4 | SMAD family member 4 | 2202.39 | 1282.95 | 0.78 | 8.79E-03 |
| ENSMUSG00000036941 | GSMG0019131 | Elac1 | elaC ribonuclease Z 1 | 155.26 | 135.83 | 0.19 | 5.98E-01 |
| ENSMUSG00000024556 | GSMG0019133 | Me2 | malic enzyme 2, NAD(+)-dependent, mitochondrial | 633.94 | 953.46 | -0.59 | 1.56E-03 |
| ENSMUSG00000024558 | GSMG0019134 | Mapk4 | mitogen-activated protein kinase 4 | 408.99 | 509.80 | -0.31 | 2.24E-01 |
| ENSMUSG00000036223 | GSMG0019136 | Ska1 | spindle and kinetochore associated complex subunit 1 | 41.86 | 146.86 | -1.83 | 6.33E-09 |
| ENSMUSG00000052928 | GSMG0019143 | Ctif | CBP80/20-dependent translation initiation factor | 70.52 | 74.90 | -0.09 | 8.76E-01 |
| NULL | GSMG0019145 | NULL | NULL | 217.28 | 253.33 | -0.22 | 5.35E-01 |
| NULL | GSMG0019146 | NULL | NULL | 96.00 | 129.71 | -0.44 | 2.34E-01 |

| | | | | | | | |
|---|---|---|---|---|---|---|---|
| NULL | GSMG0019163 | NULL | NULL | 367.87 | 321.90 | 0.19 | 5.68E-01 |
| ENSMUSG00000041840 | GSMG0019167 | Haus1 | HAUS augmin-like complex, subunit 1 | 443.68 | 677.45 | -0.61 | 4.03E-03 |
| ENSMUSG00000059336 | GSMG0019170 | Slc14a1 | solute carrier family 14 (urea transporter), member 1 | 83.12 | 92.19 | -0.15 | 7.79E-01 |
| NULL | GSMG0019179 | NULL | NULL | 763.19 | 756.32 | 0.01 | 9.70E-01 |
| ENSMUSG00000053950 | GSMG0019183 | Adnp2 | ADNP homeobox 2 | 819.34 | 730.53 | 0.16 | 5.10E-01 |
| ENSMUSG00000024570 | GSMG0019185 | Rbfa | ribosome binding factor A | 284.99 | 334.25 | -0.23 | 3.84E-01 |
| NULL | GSMG0019186 | NULL | NULL | 78.40 | 93.67 | -0.26 | 5.76E-01 |
| ENSMUSG00000033323 | GSMG0019194 | Ctdp1 | CTD (carboxy-terminal domain, RNA polymerase II, polypeptide A) ph | 1050.02 | 1158.34 | -0.14 | 5.30E-01 |
| ENSMUSG00000033016 | GSMG0019197 | Nfatc1 | nuclear factor of activated T cells, cytoplasmic, calcineurin dependent 1 | 437.40 | 262.07 | 0.74 | 1.04E-03 |
| ENSMUSG00000024566 | GSMG0019198 | Atp9b | ATPase, class II, type 9B | 3359.66 | 3145.45 | 0.10 | 7.28E-01 |
| ENSMUSG00000041258 | GSMG0019207 | Zfp236 | zinc finger protein 236 | 1071.95 | 901.62 | 0.25 | 2.95E-01 |
| ENSMUSG00000046982 | GSMG0019216 | Tshz1 | teashirt zinc finger family member 1 | 71.52 | 23.50 | 1.60 | 1.26E-02 |
| ENSMUSG00000024644 | GSMG0019228 | Cndp2 | CNDP dipeptidase 2 (metallopeptidase M20 family) | 1305.78 | 1445.05 | -0.15 | 5.19E-01 |
| ENSMUSG00000024645 | GSMG0019231 | Timm21 | tranlocase of inner mitochondrial membrane 21 | 511.62 | 527.22 | -0.04 | 9.01E-01 |
| ENSMUSG00000056153 | GSMG0019237 | Socs6 | suppressor of cytokine signaling 6 | 965.29 | 1083.17 | -0.17 | 4.37E-01 |
| ENSMUSG00000073514 | GSMG0019242 | Dok6 | docking protein 6 | 68.30 | 84.56 | -0.30 | 5.03E-01 |
| NULL | GSMG0019244 | NULL | NULL | 132.59 | 143.50 | -0.12 | 8.21E-01 |
| ENSMUSG00000024829 | GSMG0019246 | Mrpl21 | mitochondrial ribosomal protein L21 | 661.50 | 830.39 | -0.33 | 1.35E-01 |
| ENSMUSG00000024900 | GSMG0019247 | Cpt1a | carnitine palmitoyltransferase 1a, liver | 4232.00 | 3804.88 | 0.15 | 5.14E-01 |
| NULL | GSMG0019250 | NULL | NULL | 261.15 | 273.72 | -0.07 | 8.72E-01 |
| ENSMUSG00000035372 | GSMG0019253 | 1810055G02Rik | RIKEN cDNA 1810055G02 gene | 627.26 | 618.82 | 0.02 | 9.59E-01 |
| ENSMUSG00000045098 | GSMG0019254 | Suv420h1 | suppressor of variegation 4-20 homolog 1 (Drosophila) | 608.54 | 652.94 | -0.10 | 7.49E-01 |
| ENSMUSG00000024843 | GSMG0019256 | Chka | choline kinase alpha | 1101.04 | 956.82 | 0.20 | 3.60E-01 |
| ENSMUSG00000036908 | GSMG0019257 | Unc93b1 | unc-93 homolog B1 (C. elegans) | 1419.98 | 996.81 | 0.51 | 1.12E-02 |
| ENSMUSG00000024866 | GSMG0019260 | Acy3 | aspartoacylase (aminoacylase) 3 | 329.08 | 308.55 | 0.10 | 7.82E-01 |
| G00000024871,ENSMUSG000 | GSMG0019262 | Nudt8 // Doc2g | nudix (nucleoside diphosphate linked moiety X)-type motif 8 // double | 119.13 | 163.56 | -0.45 | 1.31E-01 |
| ENSMUSG00000024856 | GSMG0019264 | Cdk2ap2 | CDK2-associated protein 2 | 647.99 | 506.03 | 0.36 | 9.70E-02 |
| ENSMUSG00000024851 | GSMG0019265 | Pitpnm1 | phosphatidylinositol transfer protein, membrane-associated 1 | 5144.53 | 4588.55 | 0.17 | 5.69E-01 |
| ENSMUSG00000024845 | GSMG0019266 | Tmem134 | transmembrane protein 134 | 1343.60 | 1317.41 | 0.03 | 9.21E-01 |
| ENSMUSG00000024835 | GSMG0019268 | Coro1b | coronin, actin binding protein 1B | 10069.11 | 9502.91 | 0.08 | 6.98E-01 |
| ENSMUSG00000040385 | GSMG0019271 | Ppp1ca | protein phosphatase 1, catalytic subunit, alpha isoform | 6508.45 | 8238.94 | -0.34 | 4.24E-02 |
| ENSMUSG00000024892 | GSMG0019277 | Pcx | pyruvate carboxylase | 3722.95 | 5598.64 | -0.59 | 9.74E-04 |
| ENSMUSG00000067889 | GSMG0019280 | Sptbn2 | spectrin beta, non-erythrocytic 2 | 3753.13 | 2400.15 | 0.64 | 3.45E-03 |

| | | | | | | | |
|---|---|---|---|---|---|---|---|
| ENSMUSG00000083282 | GSMG0019283 | Ctsf | cathepsin F | 2360.22 | 1767.11 | 0.42 | 1.97E-02 |
| ENSMUSG00000006463 | GSMG0019284 | Zdhhc24 | zinc finger, DHHC domain containing 24 | 691.54 | 512.90 | 0.43 | 2.96E-02 |
| ENSMUSG00000024902 | GSMG0019286 | Mrpl11 | mitochondrial ribosomal protein L11 | 858.42 | 1151.54 | -0.42 | 7.00E-02 |
| ENSMUSG00000024891 | GSMG0019288 | Slc29a2 | solute carrier family 29 (nucleoside transporters), member 2 | 195.10 | 381.88 | -0.97 | 9.21E-05 |
| G00000047379,ENSMUSG000 | GSMG0019289 | B4gat1 // Brms1 | beta-1,4-glucuronyltransferase 1 // breast cancer metastasis-suppressor | 3738.64 | 3617.36 | 0.05 | 8.53E-01 |
| ENSMUSG00000056481 | GSMG0019292 | Cd248 | CD248 antigen, endosialin | 5654.66 | 6857.89 | -0.28 | 9.01E-02 |
| ENSMUSG00000024875 | GSMG0019293 | Yif1a | Yip1 interacting factor homolog A (S. cerevisiae) | 1001.17 | 1005.39 | 0.00 | 9.89E-01 |
| NULL | GSMG0019295 | NULL | NULL | 69.51 | 84.15 | -0.28 | 5.46E-01 |
| NULL | GSMG0019296 | NULL | NULL | 2112.30 | 2101.37 | 0.01 | 9.83E-01 |
| ENSMUSG00000047658 | GSMG0019297 | Gal3st3 | galactose-3-O-sulfotransferase 3 | 222.12 | 251.89 | -0.18 | 6.99E-01 |
| ENSMUSG00000024841 | GSMG0019299 | Eif1ad | eukaryotic translation initiation factor 1A domain containing | 1196.33 | 1364.09 | -0.19 | 3.29E-01 |
| ENSMUSG00000047423 | GSMG0019302 | AI837181 | expressed sequence AI837181 | 521.42 | 566.06 | -0.12 | 6.84E-01 |
| ENSMUSG00000024911 | GSMG0019304 | Fibp | fibroblast growth factor (acidic) intracellular binding protein | 1425.90 | 1774.89 | -0.32 | 5.52E-02 |
| ENSMUSG00000024909 | GSMG0019306 | Efemp2 | epidermal growth factor-containing fibulin-like extracellular matrix pro | 846.93 | 730.17 | 0.22 | 3.78E-01 |
| NULL | GSMG0019307 | NULL | NULL | 160.61 | 101.39 | 0.67 | 3.66E-02 |
| ENSMUSG00000056201 | GSMG0019308 | Cfl1 | cofilin 1, non-muscle | 17553.58 | 24064.56 | -0.46 | 1.27E-02 |
| ENSMUSG00000049562 | GSMG0019310 | Ap5b1 | adaptor-related protein complex 5, beta 1 subunit | 479.31 | 345.52 | 0.47 | 6.20E-02 |
| ENSMUSG00000024925 | GSMG0019311 | Rnaseh2c | ribonuclease H2, subunit C | 1218.44 | 1511.95 | -0.31 | 2.07E-01 |
| ENSMUSG00000024927 | GSMG0019312 | Rela | v-rel reticuloendotheliosis viral oncogene homolog A (avian) | 1620.58 | 1657.35 | -0.03 | 9.08E-01 |
| ENSMUSG00000004054 | GSMG0019313 | Map3k11 | mitogen-activated protein kinase kinase kinase 11 | 3447.28 | 3393.60 | 0.02 | 9.40E-01 |
| ENSMUSG00000024940 | GSMG0019315 | Ltbp3 | latent transforming growth factor beta binding protein 3 | 664.72 | 477.11 | 0.48 | 8.96E-02 |
| NULL | GSMG0019316 | NULL | NULL | 1891.12 | 586.67 | 1.69 | 1.87E-28 |
| NULL | GSMG0019318 | NULL | NULL | 430.22 | 411.27 | 0.07 | 8.48E-01 |
| ENSMUSG00000024818 | GSMG0019320 | Slc25a45 | solute carrier family 25, member 45 | 215.48 | 182.80 | 0.24 | 5.13E-01 |
| ENSMUSG00000024807 | GSMG0019323 | Syvn1 | synovial apoptosis inhibitor 1, synoviolin | 1009.21 | 947.83 | 0.09 | 7.75E-01 |
| ENSMUSG00000075227 | GSMG0019325 | Znhit2 | zinc finger, HIT domain containing 2 | 735.40 | 800.49 | -0.12 | 6.63E-01 |
| ENSMUSG00000024791 | GSMG0019327 | Cdca5 | cell division cycle associated 5 | 37.95 | 109.97 | -1.53 | 9.15E-06 |
| ENSMUSG00000024773 | GSMG0019333 | Atg2a | autophagy related 2A | 678.14 | 570.05 | 0.25 | 3.84E-01 |
| ENSMUSG00000024772 | GSMG0019334 | Ehd1 | EH-domain containing 1 | 625.89 | 565.76 | 0.15 | 6.13E-01 |
| ENSMUSG00000024947 | GSMG0019336 | Men1 | multiple endocrine neoplasia 1 | 1014.75 | 1025.25 | -0.02 | 9.62E-01 |
| ENSMUSG00000024948 | GSMG0019337 | Map4k2 | mitogen-activated protein kinase kinase kinase kinase 2 | 2459.85 | 2356.07 | 0.06 | 8.28E-01 |
| ENSMUSG00000024949 | GSMG0019338 | Sf1 | splicing factor 1 | 1645.42 | 2082.97 | -0.34 | 7.63E-02 |
| ENSMUSG00000032648 | GSMG0019339 | Pygm | muscle glycogen phosphorylase | 955.60 | 648.52 | 0.56 | 4.77E-03 |

| | | | | | | | |
|---|---|---|---|---|---|---|---|
| ENSMUSG00000032946 | GSMG0019340 | Rasgrp2 | RAS, guanyl releasing protein 2 | 703.71 | 858.79 | -0.29 | 1.86E-01 |
| ENSMUSG00000033768 | GSMG0019341 | Nrxn2 | neurexin II | 208.56 | 346.70 | -0.74 | 1.88E-02 |
| ENSMUSG00000024959 | GSMG0019346 | Bad | BCL2-associated agonist of cell death | 1039.57 | 985.27 | 0.08 | 8.09E-01 |
| ENSMUSG00000047656 | GSMG0019350 | Trpt1 | tRNA phosphotransferase 1 | 202.57 | 180.89 | 0.17 | 6.59E-01 |
| ENSMUSG00000024968 | GSMG0019353 | Rcor2 | REST corepressor 2 | 193.58 | 286.82 | -0.57 | 6.10E-02 |
| ENSMUSG00000053080 | GSMG0019356 | 2700081O15Rik | RIKEN cDNA 2700081O15 gene | 691.52 | 1147.28 | -0.73 | 1.21E-02 |
| ENSMUSG00000024759 | GSMG0019357 | Atl3 | atlastin GTPase 3 | 256.12 | 198.48 | 0.37 | 2.14E-01 |
| ENSMUSG00000060675 | GSMG0019358 | Pla2g16 | phospholipase A2, group XVI | 493.71 | 375.81 | 0.40 | 9.34E-02 |
| ENSMUSG00000063796 | GSMG0019362 | Slc22a8 | solute carrier family 22 (organic anion transporter), member 8 | 21843.02 | 20690.14 | 0.08 | 7.24E-01 |
| ENSMUSG00000024650 | GSMG0019363 | Slc22a6 | solute carrier family 22 (organic anion transporter), member 6 | 566.26 | 566.08 | 0.00 | 9.95E-01 |
| ENSMUSG00000032773 | GSMG0019364 | Chrm1 | cholinergic receptor, muscarinic 1, CNS | 1463.72 | 1197.01 | 0.29 | 2.57E-01 |
| ENSMUSG00000108414 | GSMG0019365 | Snhg1 | small nucleolar RNA host gene 1 | 351.66 | 393.90 | -0.16 | 5.44E-01 |
| ENSMUSG00000042729 | GSMG0019366 | Wdr74 | WD repeat domain 74 | 713.49 | 919.47 | -0.36 | 5.60E-02 |
| ENSMUSG00000010110 | GSMG0019368 | Stx5a | syntaxin 5A | 2310.67 | 2034.24 | 0.18 | 3.24E-01 |
| ENSMUSG00000010097 | GSMG0019370 | Nxf1 // Tmem223 | nuclear RNA export factor 1 // transmembrane protein 223 | 4830.76 | 4073.73 | 0.25 | 1.57E-01 |
| NULL | GSMG0019371 | NULL | NULL | 96.91 | 126.33 | -0.39 | 2.90E-01 |
| ENSMUSG00000071659 | GSMG0019373 | Hnrnpul2 | heterogeneous nuclear ribonucleoprotein U-like 2 | 195.14 | 234.13 | -0.27 | 5.00E-01 |
| ENSMUSG00000071657 | GSMG0019375 | Bscl2 | Berardinelli-Seip congenital lipodystrophy 2 (seipin) | 1948.13 | 1964.27 | -0.01 | 9.70E-01 |
| ENSMUSG00000071655 | GSMG0019377 | Ubxn1 | UBX domain protein 1 | 1778.60 | 1908.98 | -0.10 | 6.94E-01 |
| G00000071653,ENSMUSG000 | GSMG0019378 | Lbhd1 // 1810009A | LBH domain containing 1 // RIKEN cDNA 1810009A15 gene | 669.12 | 768.02 | -0.20 | 4.59E-01 |
| ENSMUSG00000071652 | GSMG0019379 | Ints5 | integrator complex subunit 5 | 859.79 | 1066.68 | -0.31 | 1.85E-01 |
| ENSMUSG00000071650 | GSMG0019380 | Ganab | alpha glucosidase 2 alpha neutral subunit | 4055.22 | 4443.47 | -0.13 | 5.79E-01 |
| ENSMUSG00000071649 | GSMG0019381 | B3gat3 | beta-1,3-glucuronyltransferase 3 (glucuronosyltransferase I) | 1083.78 | 1114.21 | -0.04 | 9.08E-01 |
| ENSMUSG00000071647 | GSMG0019382 | Eml3 | echinoderm microtubule associated protein like 3 | 1015.11 | 853.97 | 0.25 | 1.96E-01 |
| ENSMUSG00000071646 | GSMG0019383 | Mta2 | metastasis-associated gene family, member 2 | 5767.88 | 7113.69 | -0.30 | 5.34E-02 |
| ENSMUSG00000071645 | GSMG0019384 | Tut1 | terminal uridylyl transferase 1, U6 snRNA-specific | 344.44 | 316.94 | 0.12 | 6.93E-01 |
| ENSMUSG00000024661 | GSMG0019394 | Fth1 | ferritin heavy polypeptide 1 | 37507.12 | 50380.07 | -0.43 | 5.67E-03 |
| ENSMUSG00000024663 | GSMG0019395 | Rab3il1 | RAB3A interacting protein (rabin3)-like 1 | 775.84 | 967.03 | -0.32 | 1.64E-01 |
| ENSMUSG00000024664 | GSMG0019396 | Fads3 | fatty acid desaturase 3 | 1627.64 | 2036.90 | -0.32 | 1.23E-01 |
| NULL | GSMG0019397 | NULL | NULL | 33.85 | 56.27 | -0.72 | 1.50E-01 |
| ENSMUSG00000010663 | GSMG0019398 | Fads1 | fatty acid desaturase 1 | 1903.71 | 2368.14 | -0.31 | 6.69E-02 |
| ENSMUSG00000036372 | GSMG0019400 | Tmem258 | transmembrane protein 258 | 1104.47 | 1501.97 | -0.44 | 3.77E-02 |
| ENSMUSG00000024743 | GSMG0019403 | Syt7 | synaptotagmin VII | 370.89 | 206.98 | 0.84 | 1.33E-02 |

| | | | | | | | |
|---|---|---|---|---|---|---|---|
| NULL | GSMG0019404 | NULL | NULL | 396.69 | 163.70 | 1.28 | 3.50E-08 |
| ENSMUSG00000034820 | GSMG0019409 | Cpsf7 | cleavage and polyadenylation specific factor 7 | 319.01 | 417.13 | -0.39 | 1.12E-01 |
| ENSMUSG00000034445 | GSMG0019411 | Cyb561a3 | cytochrome b561 family, member A3 | 549.09 | 483.88 | 0.18 | 4.87E-01 |
| ENSMUSG00000024740 | GSMG0019412 | Ddb1 | damage specific DNA binding protein 1 | 6157.25 | 7847.89 | -0.35 | 8.42E-02 |
| ENSMUSG00000048832 | GSMG0019416 | Vps37c | vacuolar protein sorting 37C | 1063.65 | 1089.94 | -0.03 | 9.13E-01 |
| ENSMUSG00000024735 | GSMG0019420 | Prpf19 | pre-mRNA processing factor 19 | 1211.58 | 1396.34 | -0.20 | 4.69E-01 |
| ENSMUSG00000079419 | GSMG0019428 | Ms4a6c | membrane-spanning 4-domains, subfamily A, member 6C | 65.47 | 22.15 | 1.57 | 4.36E-04 |
| ENSMUSG00000024683 | GSMG0019436 | Mrpl16 | mitochondrial ribosomal protein L16 | 528.86 | 746.36 | -0.50 | 1.20E-02 |
| ENSMUSG00000046139 | GSMG0019439 | Patl1 | protein associated with topoisomerase II homolog 1 (yeast) | 626.31 | 789.13 | -0.34 | 1.95E-01 |
| ENSMUSG00000024687 | GSMG0019440 | Osbp | oxysterol binding protein | 874.40 | 516.80 | 0.76 | 1.43E-02 |
| ENSMUSG00000024691 | GSMG0019448 | Fam111a | family with sequence similarity 111, member A | 564.81 | 992.67 | -0.81 | 4.80E-04 |
| ENSMUSG00000024639 | GSMG0019477 | Gnaq | guanine nucleotide binding protein, alpha q polypeptide | 482.81 | 434.91 | 0.15 | 6.91E-01 |
| NULL | GSMG0019480 | NULL | NULL | 244.74 | 195.77 | 0.32 | 3.02E-01 |
| ENSMUSG00000024712 | GSMG0019491 | Rfk | riboflavin kinase | 2760.11 | 2308.27 | 0.26 | 1.61E-01 |
| NULL | GSMG0019492 | NULL | NULL | 1675.21 | 1538.31 | 0.12 | 6.20E-01 |
| ENSMUSG00000037847 | GSMG0019493 | Nmrk1 | nicotinamide riboside kinase 1 | 99.35 | 72.40 | 0.46 | 2.66E-01 |
| ENSMUSG00000047044 | GSMG0019497 | D030056L22Rik | RIKEN cDNA D030056L22 gene | 363.14 | 389.72 | -0.10 | 7.74E-01 |
| ENSMUSG00000024727 | GSMG0019498 | Trpm6 | transient receptor potential cation channel, subfamily M, member 6 | 582.90 | 468.15 | 0.32 | 2.46E-01 |
| ENSMUSG00000024750 | GSMG0019506 | Zfand5 | zinc finger, AN1-type domain 5 | 3413.25 | 3656.24 | -0.10 | 7.08E-01 |
| ENSMUSG00000047368 | GSMG0019510 | Abhd17b | abhydrolase domain containing 17B | 557.17 | 583.72 | -0.07 | 8.26E-01 |
| ENSMUSG00000024754 | GSMG0019514 | Tmem2 | transmembrane protein 2 | 2250.99 | 3558.61 | -0.66 | 1.82E-05 |
| ENSMUSG00000052387 | GSMG0019518 | Trpm3 | transient receptor potential cation channel, subfamily M, member 3 | 299.70 | 889.34 | -1.57 | 1.04E-09 |
| NULL | GSMG0019524 | NULL | NULL | 269.80 | 521.48 | -0.95 | 4.25E-06 |
| ENSMUSG00000033863 | GSMG0019532 | Klf9 | Kruppel-like factor 9 | 3214.60 | 1011.01 | 1.67 | 7.08E-32 |
| NULL | GSMG0019534 | NULL | NULL | 468.86 | 130.00 | 1.85 | 3.11E-14 |
| ENSMUSG00000074925 | GSMG0019535 | Ptar1 | protein prenyltransferase alpha subunit repeat containing 1 | 180.13 | 164.33 | 0.13 | 7.54E-01 |
| NULL | GSMG0019537 | NULL | NULL | 217.70 | 191.87 | 0.18 | 6.60E-01 |
| ENSMUSG00000024897 | GSMG0019539 | Apba1 | amyloid beta (A4) precursor protein binding, family A, member 1 | 573.38 | 744.25 | -0.38 | 2.87E-01 |
| NULL | GSMG0019544 | NULL | NULL | 57.09 | 47.39 | 0.27 | 6.47E-01 |
| NULL | GSMG0019545 | NULL | NULL | 439.03 | 421.62 | 0.06 | 8.74E-01 |
| ENSMUSG00000048572 | GSMG0019546 | Tmem252 | transmembrane protein 252 | 2953.87 | 1353.36 | 1.13 | 2.04E-06 |
| ENSMUSG00000032702 | GSMG0019549 | Kank1 | KN motif and ankyrin repeat domains 1 | 586.86 | 580.94 | 0.01 | 9.67E-01 |
| ENSMUSG00000024921 | GSMG0019554 | Smarca2 | SWI/SNF related, matrix associated, actin dependent regulator of chron | 1902.21 | 1631.81 | 0.22 | 2.87E-01 |

| | | | | | | | |
|---|---|---|---|---|---|---|---|
| ENSMUSG00000024924 | GSMG0019559 | Vldlr | very low density lipoprotein receptor | 658.24 | 953.47 | -0.53 | 9.12E-03 |
| ENSMUSG00000024935 | GSMG0019565 | Slc1a1 | solute carrier family 1 (neuronal/epithelial high affinity glutamate trans | 322.60 | 212.35 | 0.60 | 9.95E-03 |
| ENSMUSG00000040105 | GSMG0019566 | Plpp6 | phospholipid phosphatase 6 | 3291.97 | 2162.53 | 0.61 | 4.93E-05 |
| ENSMUSG00000024780 | GSMG0019567 | Cdc37l1 | cell division cycle 37-like 1 | 1073.71 | 863.86 | 0.31 | 1.61E-01 |
| ENSMUSG00000024785 | GSMG0019571 | Rcl1 | RNA terminal phosphate cyclase-like 1 | 339.89 | 479.30 | -0.49 | 1.53E-02 |
| ENSMUSG00000024789 | GSMG0019576 | Jak2 | Janus kinase 2 | 3336.65 | 3059.16 | 0.13 | 6.27E-01 |
| ENSMUSG00000038658 | GSMG0019581 | Ric1 | RAB6A GEF complex partner 1 | 147.24 | 108.15 | 0.44 | 1.95E-01 |
| NULL | GSMG0019584 | NULL | NULL | 129.68 | 130.35 | -0.01 | 9.85E-01 |
| ENSMUSG00000024817 | GSMG0019589 | Uhrf2 | ubiquitin-like, containing PHD and RING finger domains 2 | 363.39 | 286.34 | 0.34 | 1.83E-01 |
| ENSMUSG00000053536 | GSMG0019592 | Cstf2t | cleavage stimulation factor, 3' pre-RNA subunit 2, tau | 2102.64 | 2458.69 | -0.23 | 2.27E-01 |
| ENSMUSG00000024896 | GSMG0019598 | Minpp1 | multiple inositol polyphosphate histidine phosphatase 1 | 1982.53 | 2309.39 | -0.22 | 2.89E-01 |
| ENSMUSG00000013663 | GSMG0019601 | Pten | phosphatase and tensin homolog | 1924.87 | 1681.44 | 0.20 | 4.24E-01 |
| NULL | GSMG0019602 | NULL | NULL | 286.02 | 223.45 | 0.36 | 2.75E-01 |
| NULL | GSMG0019604 | NULL | NULL | 348.48 | 392.13 | -0.17 | 5.53E-01 |
| ENSMUSG00000024776 | GSMG0019614 | Stambpl1 | STAM binding protein like 1 | 218.46 | 182.33 | 0.26 | 4.25E-01 |
| ENSMUSG00000024778 | GSMG0019616 | Fas | Fas (TNF receptor superfamily member 6) | 429.26 | 58.63 | 2.88 | 2.62E-29 |
| ENSMUSG00000074896 | GSMG0019620 | Ifit3 | interferon-induced protein with tetratricopeptide repeats 3 | 188.80 | 80.80 | 1.23 | 2.48E-05 |
| ENSMUSG00000024800 | GSMG0019628 | Rpp30 | ribonuclease P/MRP 30 subunit | 280.18 | 378.53 | -0.43 | 8.01E-02 |
| ENSMUSG00000024805 | GSMG0019629 | Pcgf5 | polycomb group ring finger 5 | 153.71 | 191.22 | -0.31 | 3.65E-01 |
| NULL | GSMG0019633 | NULL | NULL | 307.21 | 219.87 | 0.48 | 8.80E-02 |
| ENSMUSG00000040565 | GSMG0019634 | Btaf1 | B-TFIID TATA-box binding protein associated factor 1 | 529.41 | 474.44 | 0.16 | 5.27E-01 |
| ENSMUSG00000023307 | GSMG0019637 | March5 | membrane-associated ring finger (C3HC4) 5 | 823.33 | 986.69 | -0.26 | 1.78E-01 |
| ENSMUSG00000012443 | GSMG0019640 | Kif11 | kinesin family member 11 | 318.31 | 838.70 | -1.39 | 1.63E-08 |
| ENSMUSG00000024986 | GSMG0019642 | Hhex | hematopoietically expressed homeobox | 287.96 | 252.79 | 0.19 | 5.46E-01 |
| ENSMUSG00000053799 | GSMG0019644 | Exoc6 | exocyst complex component 6 | 411.42 | 348.91 | 0.24 | 4.31E-01 |
| ENSMUSG00000024989 | GSMG0019647 | Cep55 | centrosomal protein 55 | 148.03 | 437.69 | -1.56 | 1.38E-08 |
| ENSMUSG00000067242 | GSMG0019650 | Lgi1 | leucine-rich repeat LGI family, member 1 | 62.85 | 61.63 | 0.02 | 9.71E-01 |
| ENSMUSG00000024998 | GSMG0019657 | Plce1 | phospholipase C, epsilon 1 | 244.35 | 231.88 | 0.07 | 8.56E-01 |
| ENSMUSG00000048720 | GSMG0019658 | Tbc1d12 | TBC1D12: TBC1 domain family, member 12 | 418.19 | 416.39 | 0.00 | 9.92E-01 |
| ENSMUSG00000025001 | GSMG0019661 | Hells | helicase, lymphoid specific | 273.85 | 569.50 | -1.06 | 1.18E-04 |
| NULL | GSMG0019670 | NULL | NULL | 54.40 | 66.22 | -0.29 | 5.81E-01 |
| ENSMUSG00000048120 | GSMG0019672 | Entpd1 | ectonucleoside triphosphate diphosphohydrolase 1 | 1947.83 | 640.30 | 1.61 | 1.33E-19 |
| ENSMUSG00000025010 | GSMG0019675 | Ccnj | cyclin J | 148.44 | 171.41 | -0.21 | 5.59E-01 |

| | | | | | | | |
|---|---|---|---|---|---|---|---|
| ENSMUSG00000025019 | GSMG0019679 | Lcor | ligand dependent nuclear receptor corepressor | 224.32 | 305.08 | -0.44 | 8.03E-02 |
| ENSMUSG00000025157 | GSMG0019693 | Zdhhc16 | zinc finger, DHHC domain containing 16 | 1899.64 | 2006.79 | -0.08 | 7.60E-01 |
| ENSMUSG00000025171 | GSMG0019694 | Ubtd1 | ubiquitin domain containing 1 | 1168.82 | 1258.33 | -0.11 | 6.62E-01 |
| ENSMUSG00000025178 | GSMG0019700 | Pi4k2a | phosphatidylinositol 4-kinase type 2 alpha | 391.84 | 345.53 | 0.18 | 5.41E-01 |
| ENSMUSG00000044345 | GSMG0019701 | Marveld1 | MARVEL (membrane-associating) domain containing 1 | 684.29 | 277.04 | 1.30 | 2.89E-08 |
| ENSMUSG00000018820 | GSMG0019703 | Zfyve27 | zinc finger, FYVE domain containing 27 | 1194.39 | 1121.64 | 0.09 | 7.69E-01 |
| ENSMUSG00000042532 | GSMG0019704 | Golga7b | golgi autoantigen, golgin subfamily a, 7B | 248.82 | 239.38 | 0.05 | 9.04E-01 |
| ENSMUSG00000025184 | GSMG0019706 | R3hcc1l | R3H domain and coiled-coil containing 1 like | 706.63 | 736.88 | -0.06 | 8.61E-01 |
| ENSMUSG00000025189 | GSMG0019715 | Cnnm1 | cyclin M1 | 313.46 | 175.24 | 0.84 | 6.57E-04 |
| ENSMUSG00000025192 | GSMG0019719 | Entpd7 | ectonucleoside triphosphate diphosphohydrolase 7 | 135.80 | 138.45 | -0.02 | 9.64E-01 |
| ENSMUSG00000025193 | GSMG0019720 | Cutc | cutC copper transporter | 237.85 | 218.71 | 0.12 | 7.33E-01 |
| ENSMUSG00000025203 | GSMG0019728 | Scd2 | stearoyl-Coenzyme A desaturase 2 | 24676.83 | 29554.45 | -0.26 | 9.53E-02 |
| NULL | GSMG0019730 | NULL | NULL | 94.17 | 121.55 | -0.36 | 3.42E-01 |
| ENSMUSG00000036450 | GSMG0019732 | Hif1an | hypoxia-inducible factor 1, alpha subunit inhibitor | 410.42 | 404.61 | 0.02 | 9.62E-01 |
| ENSMUSG00000036097 | GSMG0019736 | Fam178a | family with sequence similarity 178, member A | 519.30 | 636.97 | -0.30 | 1.78E-01 |
| ENSMUSG00000025207 | GSMG0019737 | Sema4g | sema domain, immunoglobulin domain (Ig), transmembrane domain (T | 126.34 | 333.45 | -1.41 | 6.40E-06 |
| ENSMUSG00000025209 | GSMG0019738 | Peo1 | progressive external ophthalmoplegia 1 (human) | 686.39 | 781.88 | -0.19 | 4.46E-01 |
| ENSMUSG00000035342 | GSMG0019739 | Lzts2 | leucine zipper, putative tumor suppressor 2 | 458.35 | 434.62 | 0.08 | 8.19E-01 |
| ENSMUSG00000025212 | GSMG0019741 | Sfxn3 | sideroflexin 3 | 3861.97 | 3109.13 | 0.31 | 5.84E-02 |
| ENSMUSG00000025217 | GSMG0019747 | Btrc | beta-transducin repeat containing protein | 918.75 | 889.14 | 0.05 | 8.75E-01 |
| NULL | GSMG0019750 | Dpcd | deleted in primary ciliary dyskinesia | 716.15 | 730.11 | -0.03 | 9.27E-01 |
| NULL | GSMG0019751 | NULL | NULL | 373.10 | 500.90 | -0.43 | 5.02E-02 |
| NULL | GSMG0019754 | NULL | NULL | 217.71 | 254.09 | -0.22 | 5.26E-01 |
| ENSMUSG00000055491 | GSMG0019756 | Pprc1 | peroxisome proliferative activated receptor, gamma, coactivator-related | 974.51 | 1615.89 | -0.73 | 2.72E-05 |
| ENSMUSG00000015176 | GSMG0019757 | Nolc1 | nucleolar and coiled-body phosphoprotein 1 | 1213.39 | 1881.21 | -0.63 | 6.31E-04 |
| ENSMUSG00000025224 | GSMG0019759 | Gbf1 | golgi-specific brefeldin A-resistance factor 1 | 2128.41 | 2310.96 | -0.12 | 6.71E-01 |
| ENSMUSG00000025225 | GSMG0019762 | Nfkb2 | nuclear factor of kappa light polypeptide gene enhancer in B cells 2, p4 | 695.54 | 649.46 | 0.10 | 7.17E-01 |
| NULL | GSMG0019763 | NULL | NULL | 282.93 | 251.45 | 0.17 | 7.06E-01 |
| ENSMUSG00000025226 | GSMG0019764 | Fbxl15 | F-box and leucine-rich repeat protein 15 | 397.56 | 350.53 | 0.18 | 5.76E-01 |
| ENSMUSG00000025227 | GSMG0019765 | Tmem180 | transmembrane protein 180 | 267.14 | 246.01 | 0.12 | 7.20E-01 |
| ENSMUSG00000025231 | GSMG0019767 | Sufu | suppressor of fused homolog (Drosophila) | 458.83 | 451.92 | 0.02 | 9.54E-01 |
| ENSMUSG00000025034 | GSMG0019769 | Trim8 | tripartite motif-containing 8 | 3297.89 | 3395.77 | -0.04 | 9.01E-01 |
| ENSMUSG00000025036 | GSMG0019770 | Sfxn2 | sideroflexin 2 | 179.75 | 311.39 | -0.79 | 2.38E-03 |

| | | | | | | | |
|---|---|---|---|---|---|---|---|
| ENSMUSG00000047731 | GSMG0019771 | Wbp1l | WW domain binding protein 1 like | 2497.76 | 2524.45 | -0.02 | 9.62E-01 |
| ENSMUSG00000062376 | GSMG0019773 | Borcs7 | BLOC-1 related complex subunit 7 | 439.09 | 502.35 | -0.20 | 4.52E-01 |
| ENSMUSG00000003559 | GSMG0019774 | As3mt | arsenic (+3 oxidation state) methyltransferase | 745.15 | 703.35 | 0.08 | 8.13E-01 |
| ENSMUSG00000064105 | GSMG0019776 | Cnnm2 | cyclin M2 | 194.36 | 94.35 | 1.05 | 6.25E-05 |
| ENSMUSG00000034336 | GSMG0019779 | Ina | internexin neuronal intermediate filament protein, alpha | 1225.57 | 2002.86 | -0.71 | 4.49E-05 |
| ENSMUSG00000025049 | GSMG0019781 | Taf5 | TATA-box binding protein associated factor 5 | 179.22 | 210.85 | -0.23 | 4.66E-01 |
| ENSMUSG00000025047 | GSMG0019782 | Pdcd11 | programmed cell death 11 | 545.62 | 624.22 | -0.20 | 5.37E-01 |
| ENSMUSG00000006435 | GSMG0019783 | Neurl1a | neuralized E3 ubiquitin protein ligase 1A | 293.78 | 163.90 | 0.84 | 1.34E-03 |
| ENSMUSG00000025060 | GSMG0019787 | Slk | STE20-like kinase | 2653.37 | 1941.16 | 0.45 | 1.79E-01 |
| ENSMUSG00000025066 | GSMG0019788 | Sfr1 | SWI5 dependent recombination repair 1 | 1627.61 | 1823.40 | -0.16 | 5.68E-01 |
| ENSMUSG00000025068 | GSMG0019790 | Gsto1 | glutathione S-transferase omega 1 | 356.08 | 703.73 | -0.98 | 2.87E-08 |
| ENSMUSG00000063434 | GSMG0019793 | Sorcs3 | sortilin-related VPS10 domain containing receptor 3 | 377.12 | 320.05 | 0.24 | 4.59E-01 |
| ENSMUSG00000025026 | GSMG0019803 | Add3 | adducin 3 (gamma) | 3246.70 | 2930.15 | 0.15 | 4.39E-01 |
| ENSMUSG00000025025 | GSMG0019805 | Mxi1 | MAX interactor 1, dimerization protein | 519.65 | 347.66 | 0.58 | 5.15E-03 |
| ENSMUSG00000024974 | GSMG0019811 | Smc3 | structural maintenance of chromosomes 3 | 2277.38 | 3000.00 | -0.40 | 4.01E-02 |
| ENSMUSG00000024975 | GSMG0019815 | Pdcd4 | programmed cell death 4 | 852.30 | 641.57 | 0.41 | 1.91E-02 |
| ENSMUSG00000024976 | GSMG0019817 | Shoc2 | soc-2 (suppressor of clear) homolog (C. elegans) | 1479.64 | 1534.74 | -0.05 | 8.64E-01 |
| NULL | GSMG0019818 | NULL | NULL | 356.54 | 188.34 | 0.92 | 6.72E-05 |
| NULL | GSMG0019820 | NULL | NULL | 775.35 | 931.47 | -0.26 | 1.78E-01 |
| ENSMUSG00000024981 | GSMG0019822 | Acsl5 | acyl-CoA synthetase long-chain family member 5 | 1418.63 | 1780.07 | -0.33 | 1.26E-01 |
| ENSMUSG00000024983 | GSMG0019823 | Vti1a | vesicle transport through interaction with t-SNAREs 1A | 1953.54 | 1918.48 | 0.03 | 9.29E-01 |
| NULL | GSMG0019825 | NULL | NULL | 239.63 | 212.92 | 0.18 | 6.55E-01 |
| NULL | GSMG0019827 | NULL | NULL | 122.69 | 92.00 | 0.42 | 2.65E-01 |
| NULL | GSMG0019830 | NULL | NULL | 327.89 | 225.25 | 0.54 | 2.31E-02 |
| NULL | GSMG0019832 | NULL | NULL | 194.03 | 160.79 | 0.27 | 4.22E-01 |
| ENSMUSG00000024985 | GSMG0019834 | Tcf7l2 | transcription factor 7 like 2, T cell specific, HMG box | 326.93 | 347.91 | -0.09 | 7.97E-01 |
| ENSMUSG00000025076 | GSMG0019845 | Casp7 | caspase 7 | 258.88 | 333.60 | -0.36 | 1.73E-01 |
| ENSMUSG00000025078 | GSMG0019847 | Nhlrc2 | NHL repeat containing 2 | 653.26 | 780.08 | -0.25 | 2.47E-01 |
| ENSMUSG00000035283 | GSMG0019850 | Adrb1 | adrenergic receptor, beta 1 | 863.99 | 259.61 | 1.73 | 1.09E-14 |
| ENSMUSG00000033478 | GSMG0019856 | Fam160b1 | family with sequence similarity 160, member B1 | 789.63 | 1050.11 | -0.41 | 2.53E-02 |
| ENSMUSG00000025086 | GSMG0019857 | Trub1 | TruB pseudouridine (psi) synthase family member 1 | 205.23 | 254.05 | -0.31 | 2.86E-01 |
| ENSMUSG00000054843 | GSMG0019861 | Atrnl1 | attractin like 1 | 844.44 | 810.76 | 0.06 | 8.65E-01 |
| NULL | GSMG0019863 | NULL | NULL | 294.63 | 224.30 | 0.39 | 2.02E-01 |

| | | | | | | | |
|---|---|---|---|---|---|---|---|
| NULL | GSMG0019869 | NULL | NULL | 109.09 | 73.88 | 0.56 | 1.07E-01 |
| ENSMUSG00000025094 | GSMG0019887 | Slc18a2 | solute carrier family 18 (vesicular monoamine), member 2 | 400.72 | 467.11 | -0.23 | 5.14E-01 |
| NULL | GSMG0019889 | NULL | NULL | 73.17 | 49.34 | 0.56 | 2.61E-01 |
| ENSMUSG00000024993 | GSMG0019898 | Fam45a | family with sequence similarity 45, member A | 494.32 | 517.21 | -0.06 | 8.53E-01 |
| ENSMUSG00000003228 | GSMG0019899 | Grk5 | G protein-coupled receptor kinase 5 | 282.74 | 488.01 | -0.78 | 1.42E-04 |
| NULL | GSMG0019904 | NULL | NULL | 198.19 | 355.91 | -0.84 | 2.47E-04 |
| ENSMUSG00000024831 | GSMG0019906 | Ighmbp2 | immunoglobulin mu binding protein 2 | 247.36 | 339.19 | -0.46 | 1.09E-01 |
| ENSMUSG00000024908 | GSMG0019908 | Ppp6r3 | protein phosphatase 6, regulatory subunit 3 | 1385.32 | 1406.63 | -0.02 | 9.41E-01 |
| ENSMUSG00000024913 | GSMG0019909 | Lrp5 | low density lipoprotein receptor-related protein 5 | 592.72 | 484.59 | 0.29 | 4.14E-01 |
| ENSMUSG00000001750 | GSMG0019913 | Tcirg1 | T cell, immune regulator 1, ATPase, H+ transporting, lysosomal V0 pro | 914.69 | 998.71 | -0.13 | 6.36E-01 |
| ENSMUSG00000059734 | GSMG0019914 | Ndufs8 | NADH dehydrogenase (ubiquinone) Fe-S protein 8 | 1950.02 | 1993.73 | -0.03 | 9.07E-01 |
| ENSMUSG00000024885 | GSMG0019915 | Aldh3b1 | aldehyde dehydrogenase 3 family, member B1 | 233.09 | 196.81 | 0.25 | 4.03E-01 |
| ENSMUSG00000037916 | GSMG0019918 | Ndufv1 | NADH dehydrogenase (ubiquinone) flavoprotein 1 | 5174.50 | 6752.27 | -0.38 | 1.09E-02 |
| ENSMUSG00000024847 | GSMG0019923 | Aip | aryl-hydrocarbon receptor-interacting protein | 1095.03 | 926.25 | 0.24 | 2.55E-01 |
| ENSMUSG00000024830 | GSMG0019925 | Rps6kb2 | ribosomal protein S6 kinase, polypeptide 2 | 2434.92 | 2003.47 | 0.28 | 1.15E-01 |
| ENSMUSG00000024824 | GSMG0019927 | Rad9a | RAD9 checkpoint clamp component A | 356.80 | 447.65 | -0.32 | 1.83E-01 |
| ENSMUSG00000005986 | GSMG0019931 | Ankrd13d | ankyrin repeat domain 13 family, member D | 569.24 | 589.61 | -0.05 | 8.77E-01 |
| ENSMUSG00000024858 | GSMG0019932 | Adrbk1 | adrenergic receptor kinase, beta 1 | 1269.26 | 1101.28 | 0.20 | 5.09E-01 |
| ENSMUSG00000054611 | GSMG0019936 | Kdm2a | lysine (K)-specific demethylase 2A | 603.75 | 724.09 | -0.26 | 4.33E-01 |
| ENSMUSG00000041845 | GSMG0019940 | Rhod | ras homolog family member D | 661.01 | 881.77 | -0.42 | 1.66E-02 |
| ENSMUSG00000049303 | GSMG0019941 | Syt12 | synaptotagmin XII | 324.73 | 389.93 | -0.26 | 2.99E-01 |
| ENSMUSG00000045045 | GSMG0019943 | Lrfn4 | leucine rich repeat and fibronectin type III domain containing 4 | 848.07 | 1005.83 | -0.25 | 5.19E-01 |
| ENSMUSG00000024889 | GSMG0019944 | Rce1 | Ras converting CAAX endopeptidase 1 | 627.51 | 699.44 | -0.15 | 5.81E-01 |
| 56,ENSMUSG00000094936,EI | GSMG0019947 | Gm21992 // Rbm4 | predicted gene 21992 // RNA binding motif protein 4 // RNA binding m | 1903.52 | 2198.82 | -0.21 | 2.80E-01 |
| NULL | GSMG0019948 | NULL | NULL | 1624.47 | 1905.75 | -0.23 | 2.37E-01 |
| ENSMUSG00000034108 | GSMG0019952 | Ccs | copper chaperone for superoxide dismutase | 532.74 | 468.93 | 0.19 | 5.30E-01 |
| ENSMUSG00000006464 | GSMG0019956 | Bbs1 | Bardet-Biedl syndrome 1 (human) | 59.68 | 63.84 | -0.09 | 8.96E-01 |
| ENSMUSG00000063904 | GSMG0019957 | Dpp3 | dipeptidylpeptidase 3 | 1994.07 | 2366.84 | -0.25 | 1.58E-01 |
| ENSMUSG00000045903 | GSMG0019959 | Npas4 | neuronal PAS domain protein 4 | 125.61 | 29.18 | 2.10 | 7.77E-08 |
| ENSMUSG00000061451 | GSMG0019961 | Tmem151a | transmembrane protein 151A | 2604.16 | 1485.08 | 0.81 | 2.09E-05 |
| ENSMUSG00000024873 | GSMG0019962 | Cnih2 | cornichon family AMPA receptor auxiliary protein 2 | 1424.61 | 2579.10 | -0.86 | 6.59E-08 |
| ENSMUSG00000024870 | GSMG0019963 | Rab1b | RAB1B, member RAS oncogene family | 6209.09 | 7315.86 | -0.24 | 1.54E-01 |
| ENSMUSG00000024862 | GSMG0019964 | Klc2 | kinesin light chain 2 | 881.22 | 935.01 | -0.09 | 8.64E-01 |

| | | | | | | | |
|---|---|---|---|---|---|---|---|
| ENSMUSG00000024855 | GSMG0019965 | Pacs1 | phosphofurin acidic cluster sorting protein 1 | 977.46 | 1178.94 | -0.27 | 2.47E-01 |
| ENSMUSG00000024853 | GSMG0019967 | Sf3b2 | splicing factor 3b, subunit 2 | 6033.31 | 6774.69 | -0.17 | 4.27E-01 |
| ENSMUSG00000024844 | GSMG0019969 | Banf1 | barrier to autointegration factor 1 | 2141.15 | 3630.41 | -0.76 | 1.64E-05 |
| ENSMUSG00000039148 | GSMG0019971 | Sart1 | squamous cell carcinoma antigen recognized by T cells 1 | 2735.22 | 2934.22 | -0.10 | 6.48E-01 |
| ENSMUSG00000024914 | GSMG0019973 | Drap1 | Dr1 associated protein 1 (negative cofactor 2 alpha) | 753.42 | 1032.76 | -0.46 | 3.37E-02 |
| ENSMUSG00000095098 | GSMG0019975 | Ccdc85b | coiled-coil domain containing 85B | 4012.34 | 4449.96 | -0.15 | 5.31E-01 |
| ENSMUSG00000024910 | GSMG0019976 | Ctsw | cathepsin W | 761.58 | 1380.48 | -0.86 | 3.09E-08 |
| ENSMUSG00000024906 | GSMG0019978 | Mus81 | MUS81 structure-specific endonuclease subunit | 1229.13 | 1010.04 | 0.28 | 1.62E-01 |
| ENSMUSG00000056185 | GSMG0019980 | Snx32 | sorting nexin 32 | 2028.66 | 1760.78 | 0.20 | 2.90E-01 |
| ENSMUSG00000024926 | GSMG0019983 | Kat5 | K(lysine) acetyltransferase 5 | 1256.31 | 1315.43 | -0.07 | 7.93E-01 |
| NULL | GSMG0019984 | NULL | NULL | 646.89 | 547.08 | 0.24 | 3.06E-01 |
| ENSMUSG00000056917 | GSMG0019985 | Sipa1 | signal-induced proliferation associated gene 1 | 7218.35 | 9836.57 | -0.45 | 2.32E-02 |
| ENSMUSG00000054874 | GSMG0019986 | Pcnx3 | pecanex homolog 3 | 1191.47 | 1320.63 | -0.15 | 6.67E-01 |
| ENSMUSG00000024937 | GSMG0019987 | Ehbp1l1 | EH domain binding protein 1-like 1 | 1177.87 | 1063.37 | 0.15 | 6.21E-01 |
| G00000024939,ENSMUSG000 | GSMG0019988 | Fam89b // Sssca1 | family with sequence similarity 89, member B // Sjogren's syndrome/scl | 2200.52 | 2847.90 | -0.37 | 2.52E-02 |
| ENSMUSG00000024941 | GSMG0019991 | Scyl1 | SCY1-like 1 (S. cerevisiae) | 1019.83 | 1073.68 | -0.07 | 7.82E-01 |
| ENSMUSG00000024816 | GSMG0019995 | Frmd8 | FERM domain containing 8 | 862.37 | 885.33 | -0.04 | 9.02E-01 |
| ENSMUSG00000024826 | GSMG0019997 | Dpf2 | D4, zinc and double PHD fingers family 2 | 1635.05 | 1745.92 | -0.09 | 7.04E-01 |
| ENSMUSG00000045664 | GSMG0019998 | Cdc42ep2 | CDC42 effector protein (Rho GTPase binding) 2 | 860.72 | 598.36 | 0.53 | 6.04E-03 |
| ENSMUSG00000024833 | GSMG0020000 | Pola2 | polymerase (DNA directed), alpha 2 | 550.00 | 721.87 | -0.39 | 9.23E-02 |
| ENSMUSG00000024942 | GSMG0020002 | Capn1 | calpain 1 | 313.75 | 290.11 | 0.11 | 8.35E-01 |
| ENSMUSG00000007338 | GSMG0020004 | Mrpl49 | mitochondrial ribosomal protein L49 | 1115.12 | 1122.25 | -0.01 | 9.79E-01 |
| ENSMUSG00000024799 | GSMG0020005 | Tm7sf2 | transmembrane 7 superfamily member 2 | 323.19 | 531.56 | -0.72 | 8.96E-05 |
| ENSMUSG00000024797 | GSMG0020006 | Vps51 | VPS51 GARP complex subunit | 2099.85 | 1993.15 | 0.08 | 8.21E-01 |
| ENSMUSG00000024792 | GSMG0020007 | Zfpl1 | zinc finger like protein 1 | 558.97 | 656.87 | -0.23 | 2.88E-01 |
| ENSMUSG00000024790 | GSMG0020009 | Sac3d1 | SAC3 domain containing 1 | 629.10 | 741.40 | -0.24 | 3.36E-01 |
| ENSMUSG00000024787 | GSMG0020010 | Snx15 | sorting nexin 15 | 529.08 | 527.92 | 0.00 | 9.93E-01 |
| ENSMUSG00000024944 | GSMG0020011 | Arl2 | ADP-ribosylation factor-like 2 | 1225.09 | 1375.40 | -0.17 | 4.23E-01 |
| ENSMUSG00000024777 | GSMG0020012 | Ppp2r5b | protein phosphatase 2, regulatory subunit B', beta | 2029.04 | 2118.99 | -0.06 | 8.31E-01 |
| NULL | GSMG0020016 | NULL | NULL | 269.78 | 233.10 | 0.20 | 6.26E-01 |
| ENSMUSG00000024952 | GSMG0020019 | Rps6ka4 | ribosomal protein S6 kinase, polypeptide 4 | 901.39 | 759.92 | 0.25 | 2.27E-01 |
| ENSMUSG00000024953 | GSMG0020021 | Prdx5 | peroxiredoxin 5 | 2996.13 | 2747.59 | 0.12 | 6.22E-01 |
| ENSMUSG00000024955 | GSMG0020022 | Esrra | estrogen related receptor, alpha | 240.15 | 208.04 | 0.20 | 5.68E-01 |

| | | | | | | | |
|---|---|---|---|---|---|---|---|
| ENSMUSG00000050623 | GSMG0020023 | Tex40 | testis expressed 40 | 161.51 | 100.29 | 0.69 | 1.43E-02 |
| ENSMUSG00000024957 | GSMG0020024 | Kcnk4 | potassium channel, subfamily K, member 4 | 171.14 | 124.46 | 0.46 | 1.25E-01 |
| ENSMUSG00000024958 | GSMG0020025 | Gpr137 | G protein-coupled receptor 137 | 470.42 | 439.97 | 0.10 | 7.84E-01 |
| NULL | GSMG0020026 | NULL | NULL | 245.95 | 244.83 | 0.00 | 9.94E-01 |
| ENSMUSG00000024960 | GSMG0020027 | Plcb3 | phospholipase C, beta 3 | 2605.20 | 2441.96 | 0.09 | 6.79E-01 |
| ENSMUSG00000056629 | GSMG0020028 | Fkbp2 | FK506 binding protein 2 | 878.97 | 757.83 | 0.21 | 3.80E-01 |
| ENSMUSG00000024962 | GSMG0020029 | Vegfb | vascular endothelial growth factor B | 418.89 | 500.86 | -0.26 | 3.11E-01 |
| ENSMUSG00000024963 | GSMG0020030 | Dnajc4 | DnaJ heat shock protein family (Hsp40) member C4 | 477.58 | 410.21 | 0.22 | 4.01E-01 |
| ENSMUSG00000037349 | GSMG0020031 | Nudt22 | nudix (nucleoside diphosphate linked moiety X)-type motif 22 | 295.81 | 239.08 | 0.31 | 2.75E-01 |
| ENSMUSG00000024965 | GSMG0020032 | Fermt3 | fermitin family member 3 | 194.77 | 89.35 | 1.13 | 9.73E-05 |
| ENSMUSG00000024966 | GSMG0020033 | Stip1 | stress-induced phosphoprotein 1 | 1669.12 | 2220.63 | -0.41 | 2.57E-02 |
| NULL | GSMG0020035 | NULL | NULL | 89.07 | 120.11 | -0.43 | 2.54E-01 |
| ENSMUSG00000024767 | GSMG0020039 | Otub1 | OTU domain, ubiquitin aldehyde binding 1 | 1820.76 | 2325.12 | -0.35 | 6.99E-02 |
| ENSMUSG00000035885 | GSMG0020040 | Cox8a | cytochrome c oxidase subunit VIIIa | 6781.90 | 5620.31 | 0.27 | 1.79E-01 |
| ENSMUSG00000024764 | GSMG0020041 | Naa40 | N(alpha)-acetyltransferase 40, NatD catalytic subunit | 555.17 | 524.86 | 0.08 | 7.93E-01 |
| ENSMUSG00000024969 | GSMG0020044 | Mark2 | MAP/microtubule affinity regulating kinase 2 | 673.94 | 651.92 | 0.05 | 8.87E-01 |
| NULL | GSMG0020046 | NULL | NULL | 246.58 | 184.82 | 0.41 | 1.74E-01 |
| ENSMUSG00000024970 | GSMG0020047 | AI846148 | expressed sequence AI846148 | 574.28 | 758.84 | -0.40 | 3.17E-02 |
| ENSMUSG00000024758 | GSMG0020049 | Rtn3 | reticulon 3 | 2457.94 | 3204.71 | -0.38 | 1.51E-02 |
| ENSMUSG00000010095 | GSMG0020062 | Slc3a2 | solute carrier family 3 (activators of dibasic and neutral amino acid tran | 5050.76 | 7707.96 | -0.61 | 4.87E-05 |
| NULL | GSMG0020063 | LOC102640359 // | uncharacterized LOC102640359 // RIKEN cDNA 1700023D09 gene | 119.97 | 123.58 | -0.04 | 9.38E-01 |
| ENSMUSG00000071662 | GSMG0020066 | Polr2g | polymerase (RNA) II (DNA directed) polypeptide G | 1030.30 | 1107.69 | -0.10 | 6.98E-01 |
| ENSMUSG00000071660 | GSMG0020067 | Ttc9c | tetratricopeptide repeat domain 9C | 744.45 | 865.43 | -0.22 | 4.05E-01 |
| ENSMUSG00000071658 | GSMG0020068 | Gng3 | guanine nucleotide binding protein (G protein), gamma 3 | 963.66 | 1612.69 | -0.74 | 3.80E-04 |
| ENSMUSG00000071654 | GSMG0020069 | Uqcc3 | ubiquinol-cytochrome c reductase complex assembly factor 3 | 1182.34 | 1203.22 | -0.03 | 9.41E-01 |
| NULL | GSMG0020070 | NULL | NULL | 49.27 | 53.80 | -0.14 | 8.41E-01 |
| NULL | GSMG0020071 | NULL | NULL | 184.58 | 240.90 | -0.39 | 1.45E-01 |
| ENSMUSG00000071648 | GSMG0020072 | Rom1 | rod outer segment membrane protein 1 | 231.19 | 150.98 | 0.62 | 2.63E-02 |
| NULL | GSMG0020073 | NULL | NULL | 774.50 | 685.18 | 0.18 | 4.38E-01 |
| ENSMUSG00000024654 | GSMG0020075 | Asrgl1 | asparaginase like 1 | 440.24 | 462.22 | -0.07 | 8.35E-01 |
| ENSMUSG00000024660 | GSMG0020079 | Incenp | inner centromere protein | 837.88 | 1551.59 | -0.89 | 1.45E-08 |
| NULL | GSMG0020082 | NULL | NULL | 13721.84 | 17700.28 | -0.37 | 2.01E-02 |
| ENSMUSG00000037418 | GSMG0020083 | Best1 | bestrophin 1 | 245.85 | 88.19 | 1.48 | 1.67E-01 |

| | | | | | | | |
|---|---|---|---|---|---|---|---|
| ENSMUSG00000024665 | GSMG0020084 | Fads2 | fatty acid desaturase 2 | 2765.94 | 4305.99 | -0.64 | 3.36E-05 |
| ENSMUSG00000024742 | GSMG0020086 | Fen1 | flap structure specific endonuclease 1 | 800.67 | 1586.59 | -0.99 | 6.10E-09 |
| ENSMUSG00000035735 | GSMG0020088 | Dagla | diacylglycerol lipase, alpha | 456.95 | 531.51 | -0.22 | 6.36E-01 |
| ENSMUSG00000024668 | GSMG0020091 | Sdhaf2 | succinate dehydrogenase complex assembly factor 2 | 1006.72 | 992.80 | 0.02 | 9.50E-01 |
| ENSMUSG00000024667 | GSMG0020092 | Tmem216 | transmembrane protein 216 | 234.76 | 258.30 | -0.14 | 6.79E-01 |
| NULL | GSMG0020093 | NULL | NULL | 516.24 | 519.39 | -0.01 | 9.80E-01 |
| ENSMUSG00000024666 | GSMG0020094 | Tmem138 | transmembrane protein 138 | 240.85 | 280.29 | -0.22 | 4.64E-01 |
| ENSMUSG00000034371 | GSMG0020096 | Tkfc | triokinase, FMN cyclase | 153.16 | 149.00 | 0.04 | 9.37E-01 |
| ENSMUSG00000024736 | GSMG0020102 | Tmem132a | transmembrane protein 132A | 1668.51 | 1732.65 | -0.05 | 9.03E-01 |
| ENSMUSG00000034659 | GSMG0020103 | Tmem109 | transmembrane protein 109 | 4198.86 | 3691.69 | 0.19 | 3.42E-01 |
| ENSMUSG00000024732 | GSMG0020105 | Ccdc86 | coiled-coil domain containing 86 | 599.28 | 825.96 | -0.46 | 1.84E-02 |
| ENSMUSG00000075010 | GSMG0020109 | AW112010 | expressed sequence AW112010 | 130.96 | 50.87 | 1.37 | 7.29E-05 |
| ENSMUSG00000024672 | GSMG0020116 | Ms4a7 | membrane-spanning 4-domains, subfamily A, member 7 | 99.05 | 16.98 | 2.55 | 1.37E-09 |
| ENSMUSG00000041488 | GSMG0020123 | Stx3 | syntaxin 3 | 568.70 | 391.88 | 0.54 | 5.26E-03 |
| ENSMUSG00000039982 | GSMG0020132 | Dtx4 | deltex 4, E3 ubiquitin ligase | 229.41 | 313.48 | -0.45 | 1.03E-01 |
| NULL | GSMG0020133 | NULL | NULL | 247.83 | 372.05 | -0.58 | 5.03E-03 |
| G00000024695,ENSMUSG000 | GSMG0020136 | Gm44505 // Cntf // | predicted readthrough transcript (NMD candidate), 44505 // ciliary neu | 400.54 | 314.96 | 0.34 | 3.23E-01 |
| ENSMUSG00000024642 | GSMG0020146 | Tle4 | transducin-like enhancer of split 4 | 797.47 | 733.25 | 0.12 | 7.16E-01 |
| ENSMUSG00000024640 | GSMG0020150 | Psat1 | phosphoserine aminotransferase 1 | 795.77 | 1093.65 | -0.46 | 1.32E-02 |
| ENSMUSG00000041491 | GSMG0020152 | Cep78 | centrosomal protein 78 | 503.71 | 684.29 | -0.44 | 2.67E-02 |
| ENSMUSG00000024713 | GSMG0020159 | Pcsk5 | proprotein convertase subtilisin/kexin type 5 | 591.23 | 476.21 | 0.31 | 2.56E-01 |
| ENSMUSG00000024725 | GSMG0020163 | Ostf1 | osteoclast stimulating factor 1 | 1066.32 | 941.43 | 0.18 | 4.54E-01 |
| NULL | GSMG0020164 | NULL | NULL | 47.15 | 56.09 | -0.24 | 7.41E-01 |
| ENSMUSG00000036192 | GSMG0020166 | Rorb | RAR-related orphan receptor beta | 146.16 | 205.29 | -0.48 | 1.69E-01 |
| NULL | GSMG0020170 | NULL | NULL | 159.55 | 282.73 | -0.82 | 1.61E-03 |
| ENSMUSG00000024659 | GSMG0020174 | Anxa1 | annexin A1 | 1964.84 | 1548.65 | 0.34 | 8.56E-02 |
| ENSMUSG00000024747 | GSMG0020177 | Aldh1a7 | aldehyde dehydrogenase family 1, subfamily A7 | 162.69 | 47.63 | 1.77 | 1.34E-10 |
| NULL | GSMG0020182 | NULL | NULL | 486.79 | 512.21 | -0.07 | 8.05E-01 |
| ENSMUSG00000058624 | GSMG0020184 | Gda | guanine deaminase | 128.76 | 115.12 | 0.16 | 7.16E-01 |
| ENSMUSG00000035171 | GSMG0020185 | 1110059E24Rik | RIKEN cDNA 1110059E24 gene | 801.74 | 805.95 | -0.01 | 9.87E-01 |
| NULL | GSMG0020187 | NULL | NULL | 77.27 | 80.12 | -0.06 | 9.27E-01 |
| NULL | GSMG0020192 | NULL | NULL | 579.84 | 182.81 | 1.67 | 8.38E-18 |
| ENSMUSG00000024943 | GSMG0020194 | Smc5 | structural maintenance of chromosomes 5 | 1186.11 | 1401.32 | -0.24 | 2.43E-01 |

| | | | | | | | |
|---|---|---|---|---|---|---|---|
| ENSMUSG00000033207 | GSMG0020195 | Mamdc2 | MAM domain containing 2 | 178.75 | 114.22 | 0.65 | 6.85E-02 |
| ENSMUSG00000071604 | GSMG0020200 | Fam189a2 | family with sequence similarity 189, member A2 | 726.66 | 489.92 | 0.57 | 3.50E-02 |
| ENSMUSG00000024812 | GSMG0020201 | Tjp2 | tight junction protein 2 | 3258.65 | 3286.26 | -0.01 | 9.63E-01 |
| ENSMUSG00000059363 | GSMG0020204 | Fxn | frataxin | 183.34 | 404.22 | -1.14 | 1.58E-07 |
| ENSMUSG00000024867 | GSMG0020206 | Pip5k1b | phosphatidylinositol-4-phosphate 5-kinase, type 1 beta | 445.53 | 285.59 | 0.64 | 5.32E-03 |
| ENSMUSG00000074922 | GSMG0020211 | Fam122a | family with sequence similarity 122, member A | 934.55 | 949.88 | -0.03 | 9.50E-01 |
| NULL | GSMG0020213 | NULL | NULL | 443.54 | 267.83 | 0.73 | 1.31E-04 |
| ENSMUSG00000024878 | GSMG0020219 | Cbwd1 | COBW domain containing 1 | 146.15 | 150.59 | -0.04 | 9.35E-01 |
| ENSMUSG00000041360 | GSMG0020233 | Pum3 | pumilio RNA-binding family member 3 | 885.62 | 1321.07 | -0.58 | 7.19E-04 |
| ENSMUSG00000040929 | GSMG0020236 | Rfx3 | regulatory factor X, 3 (influences HLA class II expression) | 571.57 | 796.23 | -0.48 | 2.28E-02 |
| ENSMUSG00000064202 | GSMG0020252 | 4430402I18Rik | RIKEN cDNA 4430402I18 gene | 93.08 | 70.22 | 0.40 | 3.34E-01 |
| ENSMUSG00000024782 | GSMG0020253 | Ak3 | adenylate kinase 3 | 5805.97 | 4432.65 | 0.39 | 1.24E-02 |
| ENSMUSG00000016495 | GSMG0020257 | Plgrkt | plasminogen receptor, C-terminal lysine transmembrane protein | 679.13 | 661.11 | 0.04 | 9.13E-01 |
| NULL | GSMG0020258 | NULL | NULL | 118.15 | 119.70 | -0.02 | 9.68E-01 |
| ENSMUSG00000046324 | GSMG0020261 | Ermp1 | endoplasmic reticulum metallopeptidase 1 | 1394.69 | 1414.06 | -0.02 | 9.46E-01 |
| ENSMUSG00000046138 | GSMG0020269 | 9930021J03Rik | RIKEN cDNA 9930021J03 gene | 1002.18 | 1138.71 | -0.19 | 4.82E-01 |
| ENSMUSG00000074909 | GSMG0020271 | Ranbp6 | RAN binding protein 6 | 1412.73 | 1351.89 | 0.06 | 8.28E-01 |
| NULL | GSMG0020285 | NULL | NULL | 142.93 | 196.54 | -0.47 | 1.41E-01 |
| NULL | GSMG0020297 | NULL | NULL | 452.50 | 330.94 | 0.45 | 4.00E-02 |
| ENSMUSG00000040451 | GSMG0020307 | Sgms1 | sphingomyelin synthase 1 | 10207.26 | 3557.77 | 1.52 | 4.64E-22 |
| NULL | GSMG0020310 | NULL | NULL | 100.08 | 55.40 | 0.86 | 2.38E-02 |
| ENSMUSG00000013662 | GSMG0020313 | Atad1 | ATPase family, AAA domain containing 1 | 806.63 | 1074.55 | -0.41 | 2.34E-02 |
| ENSMUSG00000024766 | GSMG0020315 | Lipo3 | lipase, member O3 | 554.87 | 563.65 | -0.02 | 9.55E-01 |
| ENSMUSG00000035783 | GSMG0020318 | Acta2 | actin, alpha 2, smooth muscle, aorta | 5744.17 | 935.96 | 2.62 | 2.89E-08 |
| ENSMUSG00000024781 | GSMG0020320 | Lipa | lysosomal acid lipase A | 2171.20 | 2638.59 | -0.28 | 1.92E-01 |
| ENSMUSG00000009378 | GSMG0020325 | Slc16a12 | solute carrier family 16 (monocarboxylic acid transporters), member 12 | 569.10 | 545.22 | 0.06 | 8.46E-01 |
| ENSMUSG00000033610 | GSMG0020328 | Pank1 | pantothenate kinase 1 | 236.17 | 246.18 | -0.06 | 8.97E-01 |
| ENSMUSG00000067279 | GSMG0020335 | Ppp1r3c | protein phosphatase 1, regulatory (inhibitor) subunit 3C | 467.73 | 447.26 | 0.07 | 8.53E-01 |
| ENSMUSG00000047632 | GSMG0020336 | Fgfbp3 | fibroblast growth factor binding protein 3 | 264.01 | 521.01 | -0.98 | 3.95E-06 |
| ENSMUSG00000039652 | GSMG0020337 | Cpeb3 | cytoplasmic polyadenylation element binding protein 3 | 295.18 | 236.72 | 0.32 | 2.98E-01 |
| ENSMUSG00000056999 | GSMG0020342 | Ide | insulin degrading enzyme | 2077.62 | 2381.79 | -0.20 | 3.71E-01 |
| ENSMUSG00000048612 | GSMG0020347 | Myof | myoferlin | 209.14 | 81.69 | 1.36 | 1.39E-07 |
| ENSMUSG00000024990 | GSMG0020350 | Rbp4 | retinol binding protein 4, plasma | 173.34 | 53.03 | 1.71 | 1.44E-10 |

| | | | | | | | |
|---|---|---|---|---|---|---|---|
| ENSMUSG00000054237 | GSMG0020351 | Fra10ac1 | FRA10AC1 homolog (human) | 284.82 | 425.31 | -0.58 | 2.51E-02 |
| ENSMUSG00000024999 | GSMG0020356 | Noc3l | NOC3 like DNA replication regulator | 116.27 | 134.91 | -0.22 | 6.15E-01 |
| ENSMUSG00000055044 | GSMG0020365 | Pdlim1 | PDZ and LIM domain 1 (elfin) | 292.03 | 160.52 | 0.86 | 8.56E-05 |
| ENSMUSG00000025006 | GSMG0020366 | Sorbs1 | sorbin and SH3 domain containing 1 | 4357.96 | 4467.73 | -0.04 | 8.87E-01 |
| NULL | GSMG0020367 | NULL | NULL | 492.87 | 541.15 | -0.14 | 7.11E-01 |
| NULL | GSMG0020370 | NULL | NULL | 365.07 | 253.78 | 0.53 | 1.57E-02 |
| NULL | GSMG0020371 | NULL | NULL | 369.17 | 219.78 | 0.75 | 1.03E-03 |
| ENSMUSG00000025007 | GSMG0020372 | Aldh18a1 | aldehyde dehydrogenase 18 family, member A1 | 935.09 | 1354.61 | -0.53 | 3.19E-03 |
| ENSMUSG00000025008 | GSMG0020373 | Tctn3 | tectonic family member 3 | 1132.16 | 663.66 | 0.77 | 4.13E-06 |
| ENSMUSG00000025016 | GSMG0020383 | Tm9sf3 | transmembrane 9 superfamily member 3 | 6352.66 | 7030.53 | -0.15 | 4.38E-01 |
| ENSMUSG00000025020 | GSMG0020387 | Slit1 | slit homolog 1 (Drosophila) | 890.69 | 955.88 | -0.10 | 7.72E-01 |
| ENSMUSG00000025154 | GSMG0020388 | Arhgap19 | Rho GTPase activating protein 19 | 131.41 | 249.03 | -0.92 | 1.40E-04 |
| ENSMUSG00000035049 | GSMG0020390 | Rrp12 | ribosomal RNA processing 12 homolog (S. cerevisiae) | 716.57 | 1100.71 | -0.62 | 4.72E-03 |
| ENSMUSG00000034321 | GSMG0020392 | Exosc1 | exosome component 1 | 696.57 | 857.77 | -0.30 | 1.80E-01 |
| ENSMUSG00000025159 | GSMG0020394 | Mms19 | MMS19 (MET18 S. cerevisiae) | 1173.00 | 1277.06 | -0.12 | 6.41E-01 |
| ENSMUSG00000049670 | GSMG0020397 | Morn4 | MORN repeat containing 4 | 445.43 | 674.24 | -0.60 | 9.22E-04 |
| ENSMUSG00000018821 | GSMG0020398 | Avpi1 | arginine vasopressin-induced 1 | 489.01 | 281.02 | 0.80 | 2.91E-05 |
| ENSMUSG00000042401 | GSMG0020400 | Crtac1 | cartilage acidic protein 1 | 816.10 | 653.65 | 0.32 | 2.59E-01 |
| ENSMUSG00000025188 | GSMG0020405 | Hps1 | Hermansky-Pudlak syndrome 1 | 307.47 | 292.90 | 0.07 | 8.69E-01 |
| ENSMUSG00000025190 | GSMG0020414 | Got1 | glutamic-oxaloacetic transaminase 1, soluble | 552.30 | 346.06 | 0.67 | 1.84E-04 |
| ENSMUSG00000040414 | GSMG0020416 | Slc25a28 | solute carrier family 25, member 28 | 1210.91 | 1046.18 | 0.21 | 4.98E-01 |
| ENSMUSG00000040018 | GSMG0020417 | Cox15 | cytochrome c oxidase assembly protein 15 | 479.99 | 611.15 | -0.35 | 1.27E-01 |
| ENSMUSG00000025198 | GSMG0020423 | Erlin1 | ER lipid raft associated 1 | 234.83 | 325.45 | -0.47 | 5.26E-02 |
| ENSMUSG00000025199 | GSMG0020424 | Chuk | conserved helix-loop-helix ubiquitous kinase | 3346.92 | 3027.56 | 0.14 | 5.06E-01 |
| ENSMUSG00000025200 | GSMG0020425 | Cwf19l1 | CWF19-like 1, cell cycle control (S. pombe) | 357.45 | 437.01 | -0.29 | 2.40E-01 |
| ENSMUSG00000025204 | GSMG0020432 | Ndufb8 | NADH dehydrogenase (ubiquinone) 1 beta subcomplex 8 | 3913.07 | 3831.87 | 0.03 | 9.11E-01 |
| ENSMUSG00000025208 | GSMG0020436 | Mrpl43 | mitochondrial ribosomal protein L43 | 967.54 | 1092.05 | -0.18 | 4.84E-01 |
| ENSMUSG00000025218 | GSMG0020446 | Poll | polymerase (DNA directed), lambda | 206.58 | 215.55 | -0.07 | 8.76E-01 |
| ENSMUSG00000040913 | GSMG0020447 | Fbxw4 | F-box and WD-40 domain protein 4 | 248.77 | 173.19 | 0.52 | 1.27E-01 |
| ENSMUSG00000025220 | GSMG0020453 | Mgea5 | meningioma expressed antigen 5 (hyaluronidase) | 1879.47 | 2276.74 | -0.28 | 3.27E-01 |
| ENSMUSG00000025221 | GSMG0020454 | Kcnip2 | Kv channel-interacting protein 2 | 223.20 | 157.00 | 0.51 | 7.08E-02 |
| NULL | GSMG0020455 | NULL | NULL | 421.97 | 249.72 | 0.75 | 2.06E-03 |
| ENSMUSG00000039901 | GSMG0020456 | 9130011E15Rik | RIKEN cDNA 9130011E15 gene | 303.92 | 333.26 | -0.13 | 7.06E-01 |

| | | | | | | | |
|---|---|---|---|---|---|---|---|
| ENSMUSG00000025223 | GSMG0020458 | Ldb1 | LIM domain binding 1 | 1613.97 | 1522.91 | 0.08 | 8.16E-01 |
| NULL | GSMG0020461 | 4833438C02Rik | RIKEN cDNA 4833438C02 gene | 209.31 | 180.97 | 0.21 | 5.22E-01 |
| ENSMUSG00000037126 | GSMG0020462 | Psd | pleckstrin and Sec7 domain containing | 1116.69 | 1326.21 | -0.25 | 4.19E-01 |
| ENSMUSG00000036748 | GSMG0020463 | Cuedc2 | CUE domain containing 2 | 1773.76 | 1872.24 | -0.08 | 7.61E-01 |
| ENSMUSG00000025228 | GSMG0020466 | Actr1a | ARP1 actin-related protein 1A, centractin alpha | 1963.90 | 2487.94 | -0.34 | 5.49E-02 |
| NULL | GSMG0020470 | NULL | NULL | 1610.80 | 1536.80 | 0.07 | 8.67E-01 |
| ENSMUSG00000025035 | GSMG0020471 | Arl3 | ADP-ribosylation factor-like 3 | 457.79 | 638.82 | -0.48 | 2.23E-02 |
| NULL | GSMG0020475 | NULL | NULL | 104.16 | 104.58 | 0.00 | 9.96E-01 |
| NULL | GSMG0020476 | NULL | NULL | 64.93 | 70.79 | -0.11 | 8.77E-01 |
| NULL | GSMG0020479 | NULL | NULL | 293.02 | 143.87 | 1.03 | 3.33E-06 |
| ENSMUSG00000025041 | GSMG0020481 | Nt5c2 | 5'-nucleotidase, cytosolic II | 1456.60 | 1161.94 | 0.33 | 6.01E-02 |
| ENSMUSG00000025050 | GSMG0020483 | Pcgf6 | polycomb group ring finger 6 | 606.65 | 777.96 | -0.36 | 1.20E-01 |
| ENSMUSG00000071528 | GSMG0020485 | Usmg5 | upregulated during skeletal muscle growth 5 | 973.47 | 1101.20 | -0.18 | 5.97E-01 |
| ENSMUSG00000033033 | GSMG0020486 | Calhm2 | calcium homeostasis modulator 2 | 193.87 | 123.40 | 0.65 | 6.04E-02 |
| ENSMUSG00000042694 | GSMG0020494 | Obfc1 | oligonucleotide/oligosaccharide-binding fold containing 1 | 186.12 | 167.66 | 0.15 | 6.70E-01 |
| ENSMUSG00000043531 | GSMG0020504 | Sorcs1 | sortilin-related VPS10 domain containing receptor 1 | 260.10 | 340.19 | -0.39 | 2.15E-01 |
| NULL | GSMG0020507 | NULL | NULL | 45.50 | 72.99 | -0.68 | 1.81E-01 |
| NULL | GSMG0020515 | NULL | NULL | 60.11 | 72.20 | -0.28 | 6.34E-01 |
| ENSMUSG00000025027 | GSMG0020516 | Xpnpep1 | X-prolyl aminopeptidase (aminopeptidase P) 1, soluble | 3780.35 | 4786.17 | -0.34 | 2.97E-02 |
| ENSMUSG00000025024 | GSMG0020521 | Smndc1 | survival motor neuron domain containing 1 | 437.74 | 550.36 | -0.33 | 1.75E-01 |
| ENSMUSG00000084957 | GSMG0020527 | Bbip1 | BBSome interacting protein 1 | 1157.50 | 1371.70 | -0.25 | 4.28E-01 |
| ENSMUSG00000024978 | GSMG0020534 | Gpam | glycerol-3-phosphate acyltransferase, mitochondrial | 1463.30 | 1804.91 | -0.30 | 1.62E-01 |
| ENSMUSG00000024982 | GSMG0020537 | Zdhhc6 | zinc finger, DHHC domain containing 6 | 1711.70 | 1820.65 | -0.09 | 7.34E-01 |
| ENSMUSG00000025077 | GSMG0020543 | Dclre1a | DNA cross-link repair 1A | 289.00 | 332.57 | -0.20 | 4.84E-01 |
| ENSMUSG00000035173 | GSMG0020546 | Ccdc186 | coiled-coil domain containing 186 | 779.90 | 898.66 | -0.21 | 3.15E-01 |
| ENSMUSG00000025083 | GSMG0020549 | Afap1l2 | actin filament associated protein 1-like 2 | 2987.71 | 1880.66 | 0.67 | 1.62E-05 |
| ENSMUSG00000025085 | GSMG0020551 | Ablim1 | actin-binding LIM protein 1 | 9106.18 | 7323.38 | 0.31 | 6.45E-02 |
| ENSMUSG00000025089 | GSMG0020557 | Gfra1 | glial cell line derived neurotrophic factor family receptor alpha 1 | 125.00 | 104.21 | 0.26 | 5.17E-01 |
| ENSMUSG00000025092 | GSMG0020560 | Hspa12a | heat shock protein 12A | 300.93 | 317.01 | -0.07 | 8.41E-01 |
| ENSMUSG00000041362 | GSMG0020562 | Shtn1 | shootin 1 | 87.00 | 268.86 | -1.64 | 8.48E-11 |
| ENSMUSG00000074746 | GSMG0020564 | Pdzd8 | PDZ domain containing 8 | 1898.34 | 1737.82 | 0.13 | 5.91E-01 |
| ENSMUSG00000040022 | GSMG0020570 | Rab11fip2 | RAB11 family interacting protein 2 (class I) | 541.02 | 426.36 | 0.34 | 1.59E-01 |
| ENSMUSG00000057858 | GSMG0020573 | Fam204a | family with sequence similarity 204, member A | 535.75 | 643.03 | -0.27 | 3.29E-01 |

| | | | | | | | |
|---|---|---|---|---|---|---|---|
| ENSMUSG00000033417 | GSMG0020576 | Cacul1 | CDK2 associated, cullin domain 1 | 117.22 | 108.52 | 0.11 | 8.25E-01 |
| ENSMUSG00000024991 | GSMG0020579 | Eif3a | eukaryotic translation initiation factor 3, subunit A | 2869.53 | 3062.32 | -0.09 | 6.98E-01 |
| ENSMUSG00000063698 | GSMG0020580 | Sfxn4 | sideroflexin 4 | 201.72 | 172.24 | 0.23 | 4.74E-01 |
| ENSMUSG00000024997 | GSMG0020581 | Prdx3 | peroxiredoxin 3 | 1017.12 | 1223.64 | -0.27 | 2.30E-01 |
| ENSMUSG00000074733 | GSMG0020583 | Zfp950 | zinc finger protein 950 | 466.90 | 285.25 | 0.71 | 2.30E-03 |
| ENSMUSG00000059326 | GSMG0020586 | Csf2ra | colony stimulating factor 2 receptor, alpha, low-affinity (granulocyte-m | 471.31 | 495.49 | -0.08 | 8.74E-01 |
| ENSMUSG00000072258 | GSMG0020587 | Taf1a | TATA-box binding protein associated factor, RNA polymerase I, A | 331.18 | 338.27 | -0.03 | 9.33E-01 |
| ENSMUSG00000030768 | GSMG0020592 | Disp1 | dispatched homolog 1 (Drosophila) | 124.83 | 143.78 | -0.20 | 6.12E-01 |
| ENSMUSG00000046836 | GSMG0020594 | Brox | BRO1 domain and CAAX motif containing | 630.16 | 650.08 | -0.04 | 8.90E-01 |
| ENSMUSG00000056050 | GSMG0020596 | Mia3 | melanoma inhibitory activity 3 | 693.38 | 679.87 | 0.03 | 9.25E-01 |
| ENSMUSG00000050530 | GSMG0020599 | Fam171a1 | family with sequence similarity 171, member A1 | 651.30 | 740.46 | -0.19 | 4.30E-01 |
| ENSMUSG00000026643 | GSMG0020601 | Nmt2 | N-myristoyltransferase 2 | 172.14 | 192.23 | -0.15 | 7.14E-01 |
| ENSMUSG00000026648 | GSMG0020605 | Dclre1c | DNA cross-link repair 1C | 340.45 | 279.59 | 0.29 | 2.73E-01 |
| ENSMUSG00000026655 | GSMG0020609 | Fam107b | family with sequence similarity 107, member B | 1704.78 | 2092.28 | -0.30 | 1.23E-01 |
| NULL | GSMG0020615 | NULL | NULL | 701.91 | 918.71 | -0.39 | 4.39E-02 |
| NULL | GSMG0020617 | NULL | NULL | 715.70 | 1460.11 | -1.03 | 1.95E-09 |
| NULL | GSMG0020618 | NULL | NULL | 301.96 | 472.31 | -0.64 | 4.22E-02 |
| ENSMUSG00000026657 | GSMG0020619 | Frmd4a | FERM domain containing 4A | 288.23 | 492.54 | -0.77 | 5.33E-05 |
| NULL | GSMG0020620 | NULL | NULL | 313.16 | 537.01 | -0.78 | 5.51E-05 |
| NULL | GSMG0020622 | NULL | NULL | 463.49 | 486.78 | -0.07 | 8.86E-01 |
| NULL | GSMG0020623 | NULL | NULL | 201.25 | 197.64 | 0.03 | 9.50E-01 |
| ENSMUSG00000026662 | GSMG0020628 | Sephs1 | selenophosphate synthetase 1 | 755.15 | 1016.00 | -0.43 | 1.59E-02 |
| ENSMUSG00000026664 | GSMG0020629 | Phyh | phytanoyl-CoA hydroxylase | 522.27 | 462.03 | 0.18 | 5.16E-01 |
| ENSMUSG00000026676 | GSMG0020632 | Ccdc3 | coiled-coil domain containing 3 | 1034.56 | 1094.81 | -0.08 | 7.90E-01 |
| ENSMUSG00000025817 | GSMG0020633 | Nudt5 | nudix (nucleoside diphosphate linked moiety X)-type motif 5 | 320.34 | 432.27 | -0.43 | 1.20E-01 |
| NULL | GSMG0020634 | NULL | NULL | 310.99 | 275.15 | 0.18 | 5.55E-01 |
| ENSMUSG00000043241 | GSMG0020635 | Upf2 | UPF2 regulator of nonsense transcripts homolog (yeast) | 430.29 | 403.64 | 0.09 | 8.01E-01 |
| NULL | GSMG0020636 | NULL | NULL | 231.73 | 250.41 | -0.11 | 7.53E-01 |
| ENSMUSG00000039046 | GSMG0020639 | Usp6nl | USP6 N-terminal like | 800.29 | 776.82 | 0.04 | 8.96E-01 |
| ENSMUSG00000037262 | GSMG0020653 | Kin | Kin17 DNA and RNA binding protein | 747.55 | 804.31 | -0.11 | 7.17E-01 |
| ENSMUSG00000025780 | GSMG0020654 | Itih5 | inter-alpha (globulin) inhibitor H5 | 5589.15 | 2147.30 | 1.38 | 5.86E-21 |
| ENSMUSG00000061186 | GSMG0020659 | Sfmbt2 | Scm-like with four mbt domains 2 | 43.94 | 66.55 | -0.58 | 2.14E-01 |
| ENSMUSG00000023206 | GSMG0020671 | Il15ra | interleukin 15 receptor, alpha chain | 110.19 | 78.32 | 0.49 | 1.72E-01 |

| | | | | | | | |
|---|---|---|---|---|---|---|---|
| ENSMUSG00000047909 | GSMG0020672 | Ankrd16 | ankyrin repeat domain 16 | 1625.63 | 1632.88 | -0.01 | 9.85E-01 |
| ENSMUSG00000026730 | GSMG0020674 | Pter | phosphotriesterase related | 55.21 | 113.36 | -1.02 | 7.87E-03 |
| ENSMUSG00000026728 | GSMG0020683 | Vim | vimentin | 8975.74 | 7496.00 | 0.26 | 1.56E-01 |
| ENSMUSG00000026718 | GSMG0020685 | Stam | signal transducing adaptor molecule (SH3 domain and ITAM motif) 1 | 639.42 | 687.69 | -0.10 | 7.42E-01 |
| ENSMUSG00000026712 | GSMG0020688 | Mrc1 | mannose receptor, C type 1 | 741.65 | 311.28 | 1.25 | 8.82E-10 |
| ENSMUSG00000036949 | GSMG0020689 | Slc39a12 | solute carrier family 39 (zinc transporter), member 12 | 404.63 | 136.70 | 1.57 | 6.04E-12 |
| ENSMUSG00000057914 | GSMG0020691 | Cacnb2 | calcium channel, voltage-dependent, beta 2 subunit | 135.05 | 105.36 | 0.35 | 3.50E-01 |
| ENSMUSG00000017418 | GSMG0020699 | Arl5b | ADP-ribosylation factor-like 5B | 423.01 | 378.73 | 0.16 | 5.92E-01 |
| ENSMUSG00000026748 | GSMG0020702 | Plxdc2 | plexin domain containing 2 | 1448.72 | 1267.52 | 0.19 | 3.25E-01 |
| ENSMUSG00000026743 | GSMG0020717 | Mllt10 | myeloid/lymphoid or mixed-lineage leukemia%3B translocated to, 10 | 892.74 | 755.62 | 0.24 | 2.90E-01 |
| ENSMUSG00000051154 | GSMG0020720 | Commd3 | COMM domain containing 3 | 1418.12 | 1578.90 | -0.16 | 5.16E-01 |
| ENSMUSG00000026739 | GSMG0020721 | Bmi1 | Bmi1 polycomb ring finger oncogene | 312.06 | 220.10 | 0.50 | 3.65E-02 |
| ENSMUSG00000023094 | GSMG0020726 | Msrb2 | methionine sulfoxide reductase B2 | 664.21 | 481.52 | 0.47 | 2.11E-02 |
| ENSMUSG00000036617 | GSMG0020730 | Etl4 | enhancer trap locus 4 | 176.32 | 211.70 | -0.26 | 4.72E-01 |
| NULL | GSMG0020733 | NULL | NULL | 164.76 | 55.72 | 1.57 | 5.66E-07 |
| NULL | GSMG0020735 | NULL | NULL | 427.73 | 224.06 | 0.94 | 3.01E-06 |
| NULL | GSMG0020736 | NULL | NULL | 254.84 | 162.96 | 0.65 | 2.35E-02 |
| NULL | GSMG0020737 | NULL | NULL | 248.14 | 150.00 | 0.72 | 2.02E-03 |
| ENSMUSG00000048550 | GSMG0020746 | Thnsl1 | threonine synthase-like 1 (bacterial) | 151.67 | 165.33 | -0.13 | 7.52E-01 |
| ENSMUSG00000045967 | GSMG0020748 | Gpr158 | G protein-coupled receptor 158 | 158.47 | 112.07 | 0.50 | 1.55E-01 |
| ENSMUSG00000026787 | GSMG0020752 | Gad2 | glutamic acid decarboxylase 2 | 338.18 | 347.07 | -0.04 | 9.31E-01 |
| ENSMUSG00000026784 | GSMG0020756 | Pdss1 | prenyl (solanesyl) diphosphate synthase, subunit 1 | 259.35 | 383.85 | -0.56 | 1.18E-02 |
| ENSMUSG00000026781 | GSMG0020760 | Acbd5 | acyl-Coenzyme A binding domain containing 5 | 1225.34 | 1193.21 | 0.04 | 8.90E-01 |
| ENSMUSG00000026775 | GSMG0020762 | Yme1l1 | YME1-like 1 (S. cerevisiae) | 1185.20 | 1438.01 | -0.28 | 2.45E-01 |
| ENSMUSG00000026974 | GSMG0020776 | Zmynd19 | zinc finger, MYND domain containing 19 | 347.06 | 551.88 | -0.67 | 1.03E-03 |
| ENSMUSG00000026975 | GSMG0020777 | Dph7 | diphthamine biosynethesis 7 | 251.45 | 272.18 | -0.11 | 7.42E-01 |
| ENSMUSG00000036833 | GSMG0020778 | Pnpla7 | patatin-like phospholipase domain containing 7 | 231.99 | 194.44 | 0.26 | 4.66E-01 |
| ENSMUSG00000006476 | GSMG0020779 | Nsmf | NMDA receptor synaptonuclear signaling and neuronal migration facto | 6678.08 | 5713.91 | 0.22 | 3.13E-01 |
| NULL | GSMG0020781 | NULL | NULL | 2524.01 | 3282.65 | -0.38 | 2.40E-02 |
| NULL | GSMG0020782 | NULL | NULL | 153.12 | 118.04 | 0.38 | 2.68E-01 |
| NULL | GSMG0020784 | NULL | NULL | 311.06 | 327.25 | -0.08 | 8.74E-01 |
| NULL | GSMG0020785 | NULL | NULL | 271.85 | 291.69 | -0.10 | 7.94E-01 |
| ENSMUSG00000078201 | GSMG0020786 | Tmem203 | transmembrane protein 203 | 540.15 | 592.62 | -0.14 | 6.91E-01 |

| | | | | | | | |
|---|---|---|---|---|---|---|---|
| ENSMUSG00000048707 | GSMG0020787 | Tprn | taperin | 531.88 | 338.27 | 0.65 | 3.12E-03 |
| ENSMUSG00000026965 | GSMG0020788 | Anapc2 | anaphase promoting complex subunit 2 | 2972.98 | 3134.76 | -0.08 | 7.63E-01 |
| ENSMUSG00000085830 | GSMG0020791 | Grin1os | glutamate receptor, ionotropic, NMDA1 (zeta 1), opposite strand | 494.56 | 230.95 | 1.10 | 2.90E-05 |
| ENSMUSG00000036646 | GSMG0020792 | Man1b1 | mannosidase, alpha, class 1B, member 1 | 266.99 | 368.02 | -0.47 | 1.25E-01 |
| ENSMUSG00000026955 | GSMG0020794 | Sapcd2 | suppressor APC domain containing 2 | 66.52 | 128.79 | -0.95 | 5.53E-03 |
| ENSMUSG00000015085 | GSMG0020796 | Entpd2 | ectonucleoside triphosphate diphosphohydrolase 2 | 288.50 | 240.63 | 0.26 | 3.74E-01 |
| ENSMUSG00000015094 | GSMG0020797 | Npdc1 | neural proliferation, differentiation and control 1 | 2923.86 | 3572.81 | -0.29 | 1.45E-01 |
| ENSMUSG00000026944 | GSMG0020799 | Abca2 | ATP-binding cassette, sub-family A (ABC1), member 2 | 1167.89 | 843.21 | 0.47 | 5.56E-02 |
| ENSMUSG00000015095 | GSMG0020801 | Fbxw5 | F-box and WD-40 domain protein 5 | 2670.76 | 2527.15 | 0.08 | 8.17E-01 |
| ENSMUSG00000015092 | GSMG0020802 | Edf1 | endothelial differentiation-related factor 1 | 2512.11 | 2482.83 | 0.02 | 9.59E-01 |
| ENSMUSG00000049086 | GSMG0020810 | Bmyc | brain expressed myelocytomatosis oncogene | 712.99 | 784.96 | -0.14 | 6.54E-01 |
| ENSMUSG00000058740 | GSMG0020815 | Kcnt1 | potassium channel, subfamily T, member 1 | 1583.27 | 960.00 | 0.72 | 9.29E-04 |
| ENSMUSG00000026930 | GSMG0020820 | Gpsm1 | G-protein signalling modulator 1 (AGS3-like, C. elegans) | 454.92 | 792.11 | -0.80 | 7.68E-05 |
| NULL | GSMG0020822 | NULL | NULL | 460.83 | 382.91 | 0.27 | 2.56E-01 |
| ENSMUSG00000026926 | GSMG0020823 | Pmpca | peptidase (mitochondrial processing) alpha | 1449.33 | 1639.31 | -0.18 | 4.54E-01 |
| ENSMUSG00000086714 | GSMG0020824 | 0610009E02Rik | RIKEN cDNA 0610009E02 gene | 76.76 | 72.65 | 0.07 | 9.17E-01 |
| ENSMUSG00000026921 | GSMG0020825 | Egfl7 | EGF-like domain 7 | 7315.97 | 6513.73 | 0.17 | 4.22E-01 |
| ENSMUSG00000036186 | GSMG0020827 | Fam69b | family with sequence similarity 69, member B | 1415.87 | 1721.42 | -0.28 | 1.80E-01 |
| ENSMUSG00000014873 | GSMG0020831 | Surf2 | surfeit gene 2 | 512.14 | 444.75 | 0.20 | 4.02E-01 |
| ENSMUSG00000015488 | GSMG0020834 | Cacfd1 | calcium channel flower domain containing 1 | 1491.34 | 1689.01 | -0.18 | 4.97E-01 |
| ENSMUSG00000036040 | GSMG0020836 | Adamtsl2 | ADAMTS-like 2 | 1086.90 | 1430.60 | -0.40 | 3.12E-02 |
| ENSMUSG00000026917 | GSMG0020837 | Wdr5 | WD repeat domain 5 | 1301.61 | 1598.66 | -0.30 | 9.72E-02 |
| ENSMUSG00000109946 | GSMG0020840 | Brd3os | bromodomain containing 3, opposite strand | 479.66 | 428.30 | 0.16 | 6.11E-01 |
| ENSMUSG00000015846 | GSMG0020842 | Rxra | retinoid X receptor alpha | 238.23 | 133.79 | 0.83 | 3.22E-01 |
| ENSMUSG00000026837 | GSMG0020843 | Col5a1 | collagen, type V, alpha 1 | 3964.19 | 1795.23 | 1.14 | 7.82E-09 |
| ENSMUSG00000026833 | GSMG0020845 | Olfm1 | olfactomedin 1 | 2253.73 | 1195.84 | 0.91 | 3.34E-05 |
| ENSMUSG00000035772 | GSMG0020847 | Mrps2 | mitochondrial ribosomal protein S2 | 935.97 | 1162.98 | -0.31 | 1.10E-01 |
| ENSMUSG00000026821 | GSMG0020850 | Ralgds | ral guanine nucleotide dissociation stimulator | 3651.00 | 4389.58 | -0.27 | 2.61E-01 |
| ENSMUSG00000026812 | GSMG0020852 | Tsc1 | tuberous sclerosis 1 | 1100.83 | 1368.05 | -0.31 | 1.99E-01 |
| ENSMUSG00000026806 | GSMG0020855 | Ddx31 | DEAD/H (Asp-Glu-Ala-Asp/His) box polypeptide 31 | 280.69 | 398.69 | -0.51 | 1.59E-02 |
| ENSMUSG00000026803 | GSMG0020858 | Ttf1 | transcription termination factor, RNA polymerase I | 593.50 | 590.21 | 0.01 | 9.86E-01 |
| ENSMUSG00000043535 | GSMG0020860 | Setx | senataxin | 158.52 | 179.75 | -0.19 | 6.21E-01 |
| ENSMUSG00000026799 | GSMG0020863 | Med27 | mediator complex subunit 27 | 456.38 | 496.19 | -0.12 | 6.93E-01 |

| | | | | | | | |
|---|---|---|---|---|---|---|---|
| ENSMUSG00000019715 | GSMG0020866 | Gle1 | GLE1 RNA export mediator (yeast) | 1567.36 | 1842.69 | -0.23 | 1.92E-01 |
| ENSMUSG00000039844 | GSMG0020867 | Rapgef1 | Rap guanine nucleotide exchange factor (GEF) 1 | 2124.60 | 1660.79 | 0.35 | 2.49E-01 |
| NULL | GSMG0020870 | NULL | NULL | 483.31 | 419.55 | 0.20 | 5.78E-01 |
| ENSMUSG00000026798 | GSMG0020872 | Coq4 | coenzyme Q4 | 176.61 | 170.67 | 0.05 | 9.13E-01 |
| ENSMUSG00000059316 | GSMG0020874 | Slc27a4 | solute carrier family 27 (fatty acid transporter), member 4 | 622.21 | 656.28 | -0.08 | 8.44E-01 |
| ENSMUSG00000069020 | GSMG0020875 | Urm1 | ubiquitin related modifier 1 | 730.25 | 767.44 | -0.07 | 8.29E-01 |
| ENSMUSG00000039787 | GSMG0020876 | Cercam | cerebral endothelial cell adhesion molecule | 189.25 | 296.84 | -0.66 | 5.60E-02 |
| ENSMUSG00000026790 | GSMG0020877 | Odf2 | outer dense fiber of sperm tails 2 | 1601.76 | 1945.04 | -0.28 | 1.68E-01 |
| ENSMUSG00000057738 | GSMG0020879 | Sptan1 | spectrin alpha, non-erythrocytic 1 | 24703.98 | 27188.40 | -0.14 | 5.68E-01 |
| ENSMUSG00000026785 | GSMG0020883 | Pkn3 | protein kinase N3 | 637.82 | 478.54 | 0.42 | 2.93E-02 |
| ENSMUSG00000015337 | GSMG0020886 | Endog | endonuclease G | 500.88 | 373.74 | 0.42 | 7.83E-02 |
| G00000007476,ENSMUSG000 | GSMG0020887 | Lrrc8a // Phyhd1 | leucine rich repeat containing 8A // phytanoyl-CoA dioxygenase domai | 2052.59 | 1827.26 | 0.17 | 5.99E-01 |
| ENSMUSG00000052533 | GSMG0020888 | Nup188 | nucleoporin 188 | 190.77 | 244.94 | -0.36 | 2.17E-01 |
| ENSMUSG00000026858 | GSMG0020889 | Fam73b | family with sequence similarity 73, member B | 1282.55 | 943.20 | 0.44 | 5.09E-02 |
| ENSMUSG00000026856 | GSMG0020890 | Dolpp1 | dolichyl pyrophosphate phosphatase 1 | 336.46 | 380.67 | -0.18 | 5.49E-01 |
| ENSMUSG00000039515 | GSMG0020891 | Ptpa | protein phosphatase 2 protein activator | 2848.22 | 3501.72 | -0.30 | 7.47E-02 |
| ENSMUSG00000047363 | GSMG0020893 | Cstad | CSA-conditional, T cell activation-dependent protein | 74.50 | 27.06 | 1.47 | 4.46E-04 |
| ENSMUSG00000026857 | GSMG0020896 | Ntmt1 | N-terminal Xaa-Pro-Lys N-methyltransferase 1 | 1103.64 | 1389.06 | -0.33 | 5.14E-02 |
| ENSMUSG00000039476 | GSMG0020897 | Prrx2 | paired related homeobox 2 | 157.78 | 200.82 | -0.35 | 2.98E-01 |
| ENSMUSG00000026848 | GSMG0020898 | Tor1b | torsin family 1, member B | 317.33 | 328.31 | -0.05 | 8.94E-01 |
| ENSMUSG00000026854 | GSMG0020899 | Usp20 | ubiquitin specific peptidase 20 | 564.69 | 586.52 | -0.05 | 8.96E-01 |
| ENSMUSG00000000194 | GSMG0020900 | Gpr107 | G protein-coupled receptor 107 | 1158.76 | 1234.50 | -0.09 | 7.18E-01 |
| ENSMUSG00000062661 | GSMG0020902 | Ncs1 | neuronal calcium sensor 1 | 134.74 | 154.27 | -0.19 | 7.18E-01 |
| ENSMUSG00000026843 | GSMG0020905 | Fubp3 | far upstream element (FUSE) binding protein 3 | 1762.34 | 1789.94 | -0.02 | 9.42E-01 |
| ENSMUSG00000039356 | GSMG0020906 | Exosc2 | exosome component 2 | 235.08 | 314.54 | -0.42 | 1.07E-01 |
| ENSMUSG00000026842 | GSMG0020907 | Abl1 | c-abl oncogene 1, non-receptor tyrosine kinase | 620.72 | 811.16 | -0.39 | 1.71E-01 |
| ENSMUSG00000026840 | GSMG0020908 | Lamc3 | laminin gamma 3 | 1371.26 | 859.78 | 0.67 | 2.23E-03 |
| NULL | GSMG0020909 | NULL | NULL | 1040.51 | 567.27 | 0.87 | 4.31E-07 |
| ENSMUSG00000001864 | GSMG0020910 | Aif1l | allograft inflammatory factor 1-like | 2064.21 | 1708.93 | 0.27 | 2.74E-01 |
| ENSMUSG00000001855 | GSMG0020911 | Nup214 | nucleoporin 214 | 1732.99 | 1920.96 | -0.15 | 5.47E-01 |
| ENSMUSG00000051373 | GSMG0020913 | Plpp7 | phospholipid phosphatase 7 (inactive) | 351.95 | 446.52 | -0.34 | 2.01E-01 |
| ENSMUSG00000039262 | GSMG0020914 | Prrc2b | proline-rich coiled-coil 2B | 1835.50 | 2207.39 | -0.27 | 2.56E-01 |
| ENSMUSG00000039254 | GSMG0020916 | Pomt1 | protein-O-mannosyltransferase 1 | 221.13 | 302.83 | -0.45 | 1.14E-01 |

| | | | | | | | |
|---|---|---|---|---|---|---|---|
| ENSMUSG00000002546 | GSMG0020917 | Golga2 | golgi autoantigen, golgin subfamily a, 2 | 1526.34 | 1544.46 | -0.02 | 9.58E-01 |
| ENSMUSG00000039205 | GSMG0020918 | Ciz1 | CDKN1A interacting zinc finger protein 1 | 1554.47 | 1775.91 | -0.19 | 3.89E-01 |
| ENSMUSG00000026820 | GSMG0020919 | Ptges2 | prostaglandin E synthase 2 | 410.26 | 419.27 | -0.03 | 9.32E-01 |
| ENSMUSG00000039164 | GSMG0020921 | Naif1 | nuclear apoptosis inducing factor 1 | 514.94 | 390.27 | 0.40 | 8.59E-02 |
| ENSMUSG00000039157 | GSMG0020923 | Fam102a | family with sequence similarity 102, member A | 270.92 | 305.12 | -0.17 | 5.97E-01 |
| ENSMUSG00000026810 | GSMG0020924 | Dpm2 | dolichol-phosphate (beta-D) mannosyltransferase 2 | 278.57 | 248.40 | 0.16 | 6.20E-01 |
| ENSMUSG00000046854 | GSMG0020925 | Pip5kl1 | phosphatidylinositol-4-phosphate 5-kinase-like 1 | 164.17 | 106.22 | 0.63 | 3.28E-02 |
| ENSMUSG00000079442 | GSMG0020926 | St6galnac4 | ST6 (alpha-N-acetyl-neuraminyl-2,3-beta-galactosyl-1,3)-N-acetylgalac | 898.96 | 967.81 | -0.11 | 7.15E-01 |
| ENSMUSG00000026811 | GSMG0020927 | St6galnac6 | ST6 (alpha-N-acetyl-neuraminyl-2,3-beta-galactosyl-1,3)-N-acetylgalac | 341.13 | 307.97 | 0.15 | 6.31E-01 |
| ENSMUSG00000026817 | GSMG0020928 | Ak1 | adenylate kinase 1 | 166.17 | 174.02 | -0.07 | 8.79E-01 |
| ENSMUSG00000026814 | GSMG0020929 | Eng | endoglin | 17170.93 | 15903.16 | 0.11 | 6.89E-01 |
| ENSMUSG00000059013 | GSMG0020930 | Sh2d3c | SH2 domain containing 3C | 4062.10 | 4111.43 | -0.02 | 9.52E-01 |
| ENSMUSG00000009563 | GSMG0020931 | Tor2a | torsin family 2, member A | 1767.24 | 1983.64 | -0.17 | 3.84E-01 |
| ENSMUSG00000053746 | GSMG0020932 | Ptrh1 | peptidyl-tRNA hydrolase 1 homolog | 243.48 | 214.77 | 0.18 | 6.15E-01 |
| ENSMUSG00000026796 | GSMG0020933 | Fam129b | family with sequence similarity 129, member B | 2745.58 | 2748.53 | 0.00 | 9.96E-01 |
| ENSMUSG00000004105 | GSMG0020937 | Angptl2 | angiopoietin-like 2 | 1321.37 | 1203.60 | 0.14 | 6.55E-01 |
| ENSMUSG00000038696 | GSMG0020942 | Mapkap1 | mitogen-activated protein kinase associated protein 1 | 652.01 | 717.19 | -0.14 | 6.16E-01 |
| ENSMUSG00000026864 | GSMG0020948 | Hspa5 | heat shock protein 5 | 12256.02 | 15192.70 | -0.31 | 1.04E-01 |
| ENSMUSG00000026870 | GSMG0020953 | Cutal | cutA divalent cation tolerance homolog-like | 310.96 | 130.61 | 1.25 | 3.82E-08 |
| ENSMUSG00000026879 | GSMG0020958 | Gsn | gelsolin | 969.44 | 287.77 | 1.75 | 1.66E-15 |
| ENSMUSG00000026883 | GSMG0020960 | Dab2ip | disabled 2 interacting protein | 2117.10 | 1524.10 | 0.47 | 4.44E-01 |
| ENSMUSG00000026887 | GSMG0020966 | Mrrf | mitochondrial ribosome recycling factor | 226.92 | 278.98 | -0.29 | 2.92E-01 |
| ENSMUSG00000047250 | GSMG0020968 | Ptgs1 | prostaglandin-endoperoxide synthase 1 | 200.42 | 97.85 | 1.03 | 1.18E-04 |
| ENSMUSG00000035437 | GSMG0020982 | Rabgap1 | RAB GTPase activating protein 1 | 2700.58 | 2964.82 | -0.13 | 5.04E-01 |
| NULL | GSMG0020986 | NULL | NULL | 345.16 | 461.83 | -0.42 | 8.85E-02 |
| NULL | GSMG0020990 | NULL | NULL | 286.88 | 317.30 | -0.14 | 6.63E-01 |
| ENSMUSG00000000247 | GSMG0020994 | Lhx2 | LIM homeobox protein 2 | 203.39 | 363.85 | -0.84 | 3.28E-04 |
| ENSMUSG00000026749 | GSMG0020997 | Nek6 | NIMA (never in mitosis gene a)-related expressed kinase 6 | 646.21 | 1128.04 | -0.80 | 1.40E-06 |
| ENSMUSG00000046618 | GSMG0021002 | Olfml2a | olfactomedin-like 2A | 517.58 | 618.61 | -0.25 | 4.10E-01 |
| ENSMUSG00000026755 | GSMG0021005 | Arpc5l | actin related protein 2/3 complex, subunit 5-like | 2004.33 | 2320.77 | -0.21 | 2.73E-01 |
| ENSMUSG00000052155 | GSMG0021031 | Acvr2a | activin receptor IIA | 565.23 | 586.30 | -0.05 | 8.71E-01 |
| ENSMUSG00000069495 | GSMG0021047 | Epc2 | enhancer of polycomb homolog 2 (Drosophila) | 152.11 | 150.48 | 0.02 | 9.67E-01 |
| ENSMUSG00000026764 | GSMG0021050 | Kif5c | kinesin family member 5C | 2228.68 | 2865.81 | -0.36 | 5.73E-02 |

| | | | | | | | |
|---|---|---|---|---|---|---|---|
| ENSMUSG00000036202 | GSMG0021062 | Rif1 | replication timing regulatory factor 1 | 94.96 | 88.52 | 0.10 | 8.53E-01 |
| NULL | GSMG0021066 | NULL | NULL | 83.11 | 55.94 | 0.58 | 1.66E-01 |
| NULL | GSMG0021067 | NULL | NULL | 131.96 | 130.42 | 0.01 | 9.92E-01 |
| NULL | GSMG0021070 | NULL | NULL | 587.16 | 585.26 | 0.00 | 9.90E-01 |
| NULL | GSMG0021071 | NULL | NULL | 1475.50 | 1758.74 | -0.25 | 2.95E-01 |
| ENSMUSG00000026960 | GSMG0021079 | Arl6ip6 | ADP-ribosylation factor-like 6 interacting protein 6 | 430.66 | 487.80 | -0.18 | 5.36E-01 |
| NULL | GSMG0021086 | NULL | NULL | 293.57 | 150.08 | 0.97 | 1.04E-03 |
| ENSMUSG00000026827 | GSMG0021091 | Gpd2 | glycerol phosphate dehydrogenase 2, mitochondrial | 4387.61 | 3337.04 | 0.40 | 2.76E-02 |
| NULL | GSMG0021100 | NULL | NULL | 258.87 | 124.38 | 1.05 | 1.00E-05 |
| NULL | GSMG0021101 | NULL | NULL | 307.38 | 170.05 | 0.85 | 5.62E-04 |
| NULL | GSMG0021102 | NULL | NULL | 194.57 | 89.13 | 1.13 | 2.58E-05 |
| ENSMUSG00000026991 | GSMG0021103 | Pkp4 | plakophilin 4 | 354.70 | 270.80 | 0.39 | 9.78E-02 |
| NULL | GSMG0021104 | NULL | NULL | 342.65 | 226.42 | 0.60 | 8.84E-03 |
| ENSMUSG00000026977 | GSMG0021116 | March7 | membrane-associated ring finger (C3HC4) 7 | 2008.37 | 2520.85 | -0.33 | 7.07E-02 |
| ENSMUSG00000064289 | GSMG0021122 | Tank | TRAF family member-associated Nf-kappa B activator | 444.66 | 415.80 | 0.10 | 7.36E-01 |
| ENSMUSG00000026914 | GSMG0021123 | Psmd14 | proteasome (prosome, macropain) 26S subunit, non-ATPase, 14 | 1271.46 | 1601.85 | -0.33 | 6.76E-02 |
| ENSMUSG00000026904 | GSMG0021128 | Slc4a10 | solute carrier family 4, sodium bicarbonate cotransporter-like, member | 946.86 | 908.71 | 0.06 | 8.47E-01 |
| NULL | GSMG0021131 | NULL | NULL | 261.65 | 172.26 | 0.60 | 3.93E-02 |
| ENSMUSG00000044647 | GSMG0021143 | Csrnp3 | cysteine-serine-rich nuclear protein 3 | 192.83 | 343.40 | -0.83 | 1.14E-04 |
| NULL | GSMG0021144 | NULL | NULL | 155.99 | 227.72 | -0.55 | 7.42E-02 |
| ENSMUSG00000087301 | GSMG0021148 | Gm13629 | predicted gene 13629 | 72.42 | 5.88 | 3.65 | 2.10E-10 |
| NULL | GSMG0021152 | NULL | NULL | 741.25 | 896.61 | -0.28 | 2.09E-01 |
| ENSMUSG00000034780 | GSMG0021162 | B3galt1 | UDP-Gal:betaGlcNAc beta 1,3-galactosyltransferase, polypeptide 1 | 380.94 | 733.48 | -0.95 | 1.10E-07 |
| ENSMUSG00000027035 | GSMG0021165 | Cers6 | ceramide synthase 6 | 230.58 | 292.83 | -0.34 | 1.73E-01 |
| NULL | GSMG0021171 | NULL | NULL | 91.87 | 92.52 | -0.02 | 9.78E-01 |
| ENSMUSG00000034738 | GSMG0021175 | Nostrin | nitric oxide synthase trafficker | 3961.74 | 3335.98 | 0.25 | 3.62E-01 |
| ENSMUSG00000063145 | GSMG0021178 | Bbs5 | Bardet-Biedl syndrome 5 (human) | 344.71 | 331.61 | 0.05 | 8.82E-01 |
| ENSMUSG00000042133 | GSMG0021180 | Ppig | peptidyl-prolyl isomerase G (cyclophilin G) | 1470.57 | 1709.38 | -0.22 | 4.88E-01 |
| ENSMUSG00000027088 | GSMG0021181 | Phospho2 | phosphatase, orphan 2 | 843.47 | 938.35 | -0.15 | 5.62E-01 |
| ENSMUSG00000042155 | GSMG0021182 | Klhl23 | kelch-like 23 | 1101.08 | 1114.96 | -0.02 | 9.58E-01 |
| ENSMUSG00000068882 | GSMG0021183 | Ssb | Sjogren syndrome antigen B | 2550.85 | 3569.36 | -0.48 | 5.86E-02 |
| ENSMUSG00000044308 | GSMG0021185 | Ubr3 | ubiquitin protein ligase E3 component n-recognin 3 | 373.88 | 386.85 | -0.05 | 8.89E-01 |
| ENSMUSG00000075304 | GSMG0021190 | Sp5 | trans-acting transcription factor 5 | 102.35 | 203.90 | -1.01 | 5.21E-04 |

| | | | | | | | |
|---|---|---|---|---|---|---|---|
| ENSMUSG00000070880 | GSMG0021193 | Gad1 | glutamate decarboxylase 1 | 93.87 | 75.71 | 0.31 | 4.90E-01 |
| ENSMUSG00000014959 | GSMG0021194 | Gorasp2 | golgi reassembly stacking protein 2 | 1685.16 | 2215.69 | -0.39 | 1.42E-02 |
| ENSMUSG00000041966 | GSMG0021195 | Dcaf17 | DDB1 and CUL4 associated factor 17 | 453.87 | 485.82 | -0.10 | 7.61E-01 |
| ENSMUSG00000027012 | GSMG0021198 | Dync1i2 | dynein cytoplasmic 1 intermediate chain 2 | 2130.46 | 2470.67 | -0.21 | 3.45E-01 |
| ENSMUSG00000027018 | GSMG0021201 | Hat1 | histone aminotransferase 1 | 465.27 | 716.07 | -0.62 | 2.16E-03 |
| ENSMUSG00000041921 | GSMG0021203 | Metap1d | methionyl aminopeptidase type 1D (mitochondrial) | 329.00 | 173.29 | 0.92 | 2.66E-05 |
| ENSMUSG00000041911 | GSMG0021206 | Dlx1 | distal-less homeobox 1 | 211.25 | 544.10 | -1.37 | 3.28E-09 |
| ENSMUSG00000027111 | GSMG0021209 | Itga6 | integrin alpha 6 | 3091.73 | 1677.89 | 0.88 | 2.57E-07 |
| ENSMUSG00000006494 | GSMG0021210 | Pdk1 | pyruvate dehydrogenase kinase, isoenzyme 1 | 524.91 | 428.02 | 0.30 | 2.77E-01 |
| ENSMUSG00000049044 | GSMG0021211 | Rapgef4 | Rap guanine nucleotide exchange factor (GEF) 4 | 2657.88 | 1755.36 | 0.60 | 3.14E-04 |
| ENSMUSG00000004085 | GSMG0021217 | Zak | sterile alpha motif and leucine zipper containing kinase AZK | 903.79 | 687.08 | 0.39 | 3.81E-02 |
| ENSMUSG00000055612 | GSMG0021220 | Cdca7 | cell division cycle associated 7 | 774.19 | 1240.92 | -0.68 | 4.32E-04 |
| ENSMUSG00000063714 | GSMG0021223 | Sp3os | trans-acting transcription factor 3, opposite strand | 151.67 | 163.53 | -0.11 | 8.05E-01 |
| ENSMUSG00000008226 | GSMG0021227 | Scrn3 | secernin 3 | 111.32 | 66.85 | 0.73 | 4.66E-02 |
| ENSMUSG00000027099 | GSMG0021244 | Mtx2 | metaxin 2 | 763.70 | 897.44 | -0.23 | 2.92E-01 |
| ENSMUSG00000042410 | GSMG0021253 | Agps | alkylglycerone phosphate synthase | 725.06 | 819.55 | -0.18 | 4.72E-01 |
| ENSMUSG00000042369 | GSMG0021259 | Rbm45 | RNA binding motif protein 45 | 248.72 | 267.43 | -0.10 | 9.34E-01 |
| ENSMUSG00000042359 | GSMG0021261 | Osbpl6 | oxysterol binding protein-like 6 | 542.94 | 457.17 | 0.25 | 2.93E-01 |
| ENSMUSG00000002733 | GSMG0021267 | Plekha3 | pleckstrin homology domain-containing, family A (phosphoinositide bi | 585.47 | 464.65 | 0.33 | 2.64E-01 |
| ENSMUSG00000027011 | GSMG0021272 | Ube2e3 | ubiquitin-conjugating enzyme E2E 3 | 1401.83 | 1815.32 | -0.37 | 3.60E-02 |
| NULL | GSMG0021279 | NULL | NULL | 279.84 | 166.95 | 0.75 | 1.97E-02 |
| ENSMUSG00000027009 | GSMG0021280 | Itga4 | integrin alpha 4 | 840.44 | 985.71 | -0.23 | 3.37E-01 |
| ENSMUSG00000027007 | GSMG0021284 | Ssfa2 | sperm specific antigen 2 | 2093.00 | 977.50 | 1.10 | 2.10E-11 |
| ENSMUSG00000027006 | GSMG0021288 | Dnajc10 | DnaJ heat shock protein family (Hsp40) member C10 | 6939.13 | 6503.70 | 0.09 | 7.08E-01 |
| NULL | GSMG0021290 | NULL | NULL | 61.96 | 64.43 | -0.05 | 9.36E-01 |
| G00000026999,ENSMUSG000 | GSMG0021291 | Dusp19 // Nup35 | dual specificity phosphatase 19 // nucleoporin 35 | 493.32 | 513.65 | -0.06 | 8.61E-01 |
| ENSMUSG00000027091 | GSMG0021303 | Zc3h15 | zinc finger CCCH-type containing 15 | 994.17 | 1232.59 | -0.31 | 1.33E-01 |
| ENSMUSG00000027087 | GSMG0021305 | Itgav | integrin alpha V | 845.33 | 1060.72 | -0.33 | 2.09E-01 |
| ENSMUSG00000048388 | GSMG0021306 | Fam171b | family with sequence similarity 171, member B | 424.64 | 705.94 | -0.73 | 2.25E-05 |
| ENSMUSG00000027080 | GSMG0021308 | Med19 | mediator complex subunit 19 | 714.49 | 944.91 | -0.41 | 5.21E-02 |
| ENSMUSG00000034059 | GSMG0021310 | Ypel4 | yippee-like 4 (Drosophila) | 166.53 | 42.82 | 1.96 | 3.73E-11 |
| ENSMUSG00000027078 | GSMG0021311 | Ube2l6 | ubiquitin-conjugating enzyme E2L 6 | 1228.63 | 992.65 | 0.31 | 2.07E-01 |
| ENSMUSG00000027076 | GSMG0021312 | Timm10 | translocase of inner mitochondrial membrane 10 | 249.84 | 319.86 | -0.35 | 1.80E-01 |

| | | | | | | | |
|---|---|---|---|---|---|---|---|
| ENSMUSG00000027074 | GSMG0021314 | Slc43a3 | solute carrier family 43, member 3 | 617.18 | 1328.43 | -1.10 | 1.52E-09 |
| ENSMUSG00000027067 | GSMG0021317 | Ssrp1 | structure specific recognition protein 1 | 4198.27 | 5359.50 | -0.35 | 2.19E-02 |
| ENSMUSG00000033955 | GSMG0021319 | Tnks1bp1 | tankyrase 1 binding protein 1 | 717.56 | 722.55 | -0.01 | 9.77E-01 |
| NULL | GSMG0021321 | NULL | NULL | 205.63 | 276.89 | -0.43 | 8.01E-02 |
| NULL | GSMG0021322 | NULL | NULL | 6181.79 | 10481.08 | -0.76 | 1.65E-08 |
| NULL | GSMG0021348 | NULL | NULL | 1332.03 | 1576.65 | -0.24 | 4.04E-01 |
| ENSMUSG00000051329 | GSMG0021366 | Nup160 | nucleoporin 160 | 540.34 | 567.52 | -0.07 | 8.44E-01 |
| ENSMUSG00000008200 | GSMG0021368 | Fnbp4 | formin binding protein 4 | 1174.06 | 1308.61 | -0.16 | 5.15E-01 |
| NULL | GSMG0021369 | NULL | NULL | 504.96 | 542.33 | -0.11 | 7.97E-01 |
| ENSMUSG00000027282 | GSMG0021371 | Mtch2 | mitochondrial carrier 2 | 1271.58 | 1434.75 | -0.17 | 4.65E-01 |
| ENSMUSG00000040794 | GSMG0021372 | C1qtnf4 | C1q and tumor necrosis factor related protein 4 | 887.58 | 905.69 | -0.03 | 9.36E-01 |
| ENSMUSG00000005505 | GSMG0021373 | Kbtbd4 | kelch repeat and BTB (POZ) domain containing 4 | 246.03 | 265.95 | -0.11 | 7.54E-01 |
| ENSMUSG00000005506 | GSMG0021375 | Celf1 | CUGBP, Elav-like family member 1 | 1254.36 | 1309.62 | -0.06 | 8.39E-01 |
| ENSMUSG00000002102 | GSMG0021381 | Psmc3 | proteasome (prosome, macropain) 26S subunit, ATPase 3 | 4775.40 | 5127.68 | -0.10 | 6.75E-01 |
| ENSMUSG00000002103 | GSMG0021385 | Acp2 | acid phosphatase 2, lysosomal | 1405.77 | 1382.38 | 0.03 | 9.36E-01 |
| ENSMUSG00000027257 | GSMG0021387 | Pacsin3 | protein kinase C and casein kinase substrate in neurons 3 | 630.28 | 247.12 | 1.35 | 2.37E-11 |
| ENSMUSG00000027255 | GSMG0021388 | Arfgap2 | ADP-ribosylation factor GTPase activating protein 2 | 624.45 | 690.51 | -0.15 | 5.64E-01 |
| NULL | GSMG0021389 | NULL | NULL | 326.58 | 273.83 | 0.25 | 3.24E-01 |
| ENSMUSG00000027253 | GSMG0021390 | Lrp4 | low density lipoprotein receptor-related protein 4 | 53.15 | 67.42 | -0.34 | 5.67E-01 |
| ENSMUSG00000040549 | GSMG0021391 | Ckap5 | cytoskeleton associated protein 5 | 1650.50 | 2176.20 | -0.40 | 1.24E-02 |
| ENSMUSG00000027247 | GSMG0021393 | Arhgap1 | Rho GTPase activating protein 1 | 1690.64 | 1775.26 | -0.07 | 7.89E-01 |
| ENSMUSG00000027243 | GSMG0021394 | Harbi1 | harbinger transposase derived 1 | 111.44 | 95.31 | 0.22 | 6.32E-01 |
| ENSMUSG00000040506 | GSMG0021395 | Ambra1 | autophagy/beclin 1 regulator 1 | 550.80 | 609.94 | -0.15 | 6.45E-01 |
| NULL | GSMG0021407 | NULL | NULL | 392.10 | 558.54 | -0.51 | 3.21E-02 |
| ENSMUSG00000058318 | GSMG0021408 | Phf21a | PHD finger protein 21A | 410.42 | 469.21 | -0.19 | 4.61E-01 |
| ENSMUSG00000027222 | GSMG0021419 | Pex16 | peroxisomal biogenesis factor 16 | 757.59 | 678.60 | 0.16 | 5.62E-01 |
| ENSMUSG00000044916 | GSMG0021420 | 1700029I15Rik | RIKEN cDNA 1700029I15 gene | 120.33 | 69.80 | 0.78 | 4.13E-02 |
| ENSMUSG00000084781 | GSMG0021421 | D930015M05Rik | RIKEN cDNA D930015M05 gene | 63.99 | 62.56 | 0.03 | 9.64E-01 |
| ENSMUSG00000027221 | GSMG0021425 | Chst1 | carbohydrate (keratan sulfate Gal-6) sulfotransferase 1 | 4098.17 | 2813.31 | 0.54 | 3.50E-03 |
| ENSMUSG00000027220 | GSMG0021426 | Syt13 | synaptotagmin XIII | 295.34 | 172.29 | 0.78 | 2.86E-03 |
| ENSMUSG00000068735 | GSMG0021428 | Trp53i11 | transformation related protein 53 inducible protein 11 | 929.38 | 1549.16 | -0.74 | 3.90E-06 |
| NULL | GSMG0021429 | NULL | NULL | 261.34 | 184.96 | 0.50 | 5.86E-02 |
| ENSMUSG00000045464 | GSMG0021436 | 2810002D19Rik | RIKEN cDNA 2810002D19 gene | 427.75 | 436.22 | -0.03 | 9.41E-01 |

| | | | | | | | |
|---|---|---|---|---|---|---|---|
| NULL | GSMG0021457 | NULL | NULL | 1246.14 | 1227.40 | 0.02 | 9.62E-01 |
| ENSMUSG00000050587 | GSMG0021458 | Lrrc4c | leucine rich repeat containing 4C | 1137.33 | 1398.05 | -0.30 | 1.06E-01 |
| NULL | GSMG0021459 | NULL | NULL | 500.98 | 338.08 | 0.57 | 9.42E-03 |
| NULL | GSMG0021460 | NULL | NULL | 2239.37 | 2425.62 | -0.12 | 6.83E-01 |
| NULL | GSMG0021461 | NULL | NULL | 529.68 | 426.07 | 0.32 | 2.79E-01 |
| NULL | GSMG0021462 | NULL | NULL | 303.83 | 295.38 | 0.04 | 9.15E-01 |
| ENSMUSG00000027164 | GSMG0021467 | Traf6 | TNF receptor-associated factor 6 | 278.02 | 295.82 | -0.09 | 8.27E-01 |
| ENSMUSG00000027163 | GSMG0021470 | Commd9 | COMM domain containing 9 | 762.55 | 943.22 | -0.31 | 1.20E-01 |
| ENSMUSG00000027188 | GSMG0021474 | Pamr1 | peptidase domain containing associated with muscle regeneration 1 | 276.35 | 97.63 | 1.51 | 1.09E-09 |
| ENSMUSG00000005089 | GSMG0021475 | Slc1a2 | solute carrier family 1 (glial high affinity glutamate transporter), membe | 1873.20 | 605.14 | 1.63 | 2.40E-20 |
| ENSMUSG00000010911 | GSMG0021478 | Apip | APAF1 interacting protein | 416.70 | 409.39 | 0.03 | 9.42E-01 |
| ENSMUSG00000032724 | GSMG0021482 | Abtb2 | ankyrin repeat and BTB (POZ) domain containing 2 | 289.71 | 341.23 | -0.24 | 5.84E-01 |
| ENSMUSG00000032698 | GSMG0021490 | Lmo2 | LIM domain only 2 | 7216.30 | 8286.92 | -0.20 | 2.71E-01 |
| ENSMUSG00000027180 | GSMG0021491 | Fbxo3 | F-box protein 3 | 1651.96 | 1403.23 | 0.24 | 2.02E-01 |
| ENSMUSG00000032679 | GSMG0021495 | Cd59a | CD59a antigen | 1891.20 | 1353.51 | 0.48 | 1.49E-02 |
| ENSMUSG00000027176 | GSMG0021501 | Cstf3 | cleavage stimulation factor, 3' pre-RNA, subunit 3 | 1073.95 | 1099.09 | -0.03 | 9.20E-01 |
| NULL | GSMG0021505 | NULL | NULL | 108.44 | 127.37 | -0.24 | 6.07E-01 |
| NULL | GSMG0021506 | NULL | NULL | 272.24 | 391.76 | -0.53 | 2.67E-02 |
| ENSMUSG00000016386 | GSMG0021523 | Mpped2 | metallophosphoesterase domain containing 2 | 749.47 | 1015.46 | -0.44 | 1.19E-02 |
| ENSMUSG00000027115 | GSMG0021531 | Kif18a | kinesin family member 18A | 69.10 | 148.05 | -1.09 | 5.21E-04 |
| ENSMUSG00000027162 | GSMG0021534 | Lin7c | lin-7 homolog C (C. elegans) | 2462.28 | 2857.51 | -0.22 | 4.23E-01 |
| ENSMUSG00000050199 | GSMG0021536 | Lgr4 | leucine-rich repeat-containing G protein-coupled receptor 4 | 493.92 | 355.58 | 0.47 | 4.77E-02 |
| ENSMUSG00000027160 | GSMG0021537 | Ccdc34 | coiled-coil domain containing 34 | 272.86 | 389.30 | -0.52 | 1.67E-01 |
| ENSMUSG00000027134 | GSMG0021551 | Lpcat4 | lysophosphatidylcholine acyltransferase 4 | 2075.94 | 995.50 | 1.06 | 6.04E-12 |
| ENSMUSG00000027133 | GSMG0021552 | Nop10 | NOP10 ribonucleoprotein | 1338.02 | 1807.37 | -0.43 | 4.34E-02 |
| G00000027130,ENSMUSG000 | GSMG0021553 | Slc12a6 // Katnbl1 | solute carrier family 12, member 6 // katanin p80 subunit B like 1 | 4506.67 | 4106.48 | 0.13 | 5.39E-01 |
| ENSMUSG00000055943 | GSMG0021563 | Emc7 | ER membrane protein complex subunit 7 | 2812.54 | 2541.33 | 0.15 | 4.83E-01 |
| ENSMUSG00000003604 | GSMG0021564 | Aven | apoptosis, caspase activation inhibitor | 107.92 | 83.09 | 0.37 | 3.64E-01 |
| ENSMUSG00000040282 | GSMG0021581 | BC052040 | cDNA sequence BC052040 | 51.98 | 59.77 | -0.19 | 7.77E-01 |
| ENSMUSG00000027351 | GSMG0021593 | Spred1 | sprouty protein with EVH-1 domain 1, related sequence | 363.83 | 337.05 | 0.11 | 7.75E-01 |
| ENSMUSG00000005102 | GSMG0021602 | Eif2ak4 | eukaryotic translation initiation factor 2 alpha kinase 4 | 1870.89 | 1987.61 | -0.09 | 7.12E-01 |
| ENSMUSG00000040084 | GSMG0021605 | Bub1b | BUB1B, mitotic checkpoint serine/threonine kinase | 377.59 | 960.32 | -1.34 | 2.40E-09 |
| ENSMUSG00000074923 | GSMG0021606 | Pak6 | p21 protein (Cdc42/Rac)-activated kinase 6 | 576.78 | 929.50 | -0.69 | 6.83E-04 |

| | | | | | | | |
|---|---|---|---|---|---|---|---|
| ENSMUSG00000074918 | GSMG0021607 | Inafm2 | InaF motif containing 2 | 1359.85 | 902.06 | 0.59 | 3.47E-01 |
| ENSMUSG00000040035 | GSMG0021609 | Disp2 | dispatched homolog 2 (Drosophila) | 124.62 | 166.12 | -0.41 | 2.50E-01 |
| ENSMUSG00000027331 | GSMG0021610 | Knstrn | kinetochore-localized astrin/SPAG5 binding | 90.65 | 253.17 | -1.48 | 1.50E-07 |
| ENSMUSG00000027332 | GSMG0021611 | Ivd | isovaleryl coenzyme A dehydrogenase | 459.23 | 490.72 | -0.09 | 7.68E-01 |
| ENSMUSG00000040007 | GSMG0021612 | Bahd1 | bromo adjacent homology domain containing 1 | 339.30 | 241.88 | 0.49 | 2.51E-01 |
| NULL | GSMG0021613 | NULL | NULL | 453.23 | 546.68 | -0.27 | 2.65E-01 |
| ENSMUSG00000027323 | GSMG0021621 | Rad51 | RAD51 recombinase | 81.06 | 212.41 | -1.38 | 4.85E-07 |
| ENSMUSG00000046814 | GSMG0021622 | Gchfr | GTP cyclohydrolase I feedback regulator | 24.30 | 59.59 | -1.30 | 6.95E-03 |
| ENSMUSG00000068580 | GSMG0021624 | Zfyve19 | zinc finger, FYVE domain containing 19 | 424.17 | 444.51 | -0.07 | 8.48E-01 |
| ENSMUSG00000034216 | GSMG0021627 | Vps18 | VPS18 CORVET/HOPS core subunit | 568.68 | 539.57 | 0.07 | 8.69E-01 |
| ENSMUSG00000027314 | GSMG0021628 | Dll4 | delta-like 4 (Drosophila) | 3820.79 | 3409.24 | 0.16 | 5.20E-01 |
| ENSMUSG00000014077 | GSMG0021630 | Chp1 | calcineurin-like EF hand protein 1 | 2064.36 | 1823.62 | 0.18 | 3.56E-01 |
| ENSMUSG00000085438 | GSMG0021632 | 1700020I14Rik | RIKEN cDNA 1700020I14 gene | 6466.44 | 6371.24 | 0.02 | 9.46E-01 |
| ENSMUSG00000027306 | GSMG0021633 | Nusap1 | nucleolar and spindle associated protein 1 | 289.63 | 665.09 | -1.20 | 3.38E-06 |
| ENSMUSG00000027304 | GSMG0021634 | Rtf1 | RTF1, Paf1/RNA polymerase II complex component | 1420.92 | 1691.74 | -0.25 | 2.95E-01 |
| ENSMUSG00000027296 | GSMG0021636 | Itpka | inositol 1,4,5-trisphosphate 3-kinase A | 703.09 | 260.49 | 1.43 | 1.05E-08 |
| ENSMUSG00000027298 | GSMG0021637 | Tyro3 | TYRO3 protein tyrosine kinase 3 | 870.45 | 569.87 | 0.61 | 3.67E-02 |
| ENSMUSG00000033943 | GSMG0021638 | Mga | MAX gene associated | 269.04 | 248.49 | 0.12 | 7.36E-01 |
| ENSMUSG00000033902 | GSMG0021642 | Mapkbp1 | mitogen-activated protein kinase binding protein 1 | 2090.92 | 2466.46 | -0.24 | 4.78E-01 |
| 52,ENSMUSG00000098488,E | GSMG0021643 | Gm28042 // Jmjd7 | predicted gene, 28042 // jumonji domain containing 7 // phospholipase | 71.27 | 119.27 | -0.75 | 4.13E-02 |
| G00000062646,ENSMUSG000 | GSMG0021645 | Ganc // Capn3 | glucosidase, alpha%3B neutral C // calpain 3 | 97.99 | 41.33 | 1.25 | 3.81E-04 |
| ENSMUSG00000027287 | GSMG0021647 | Snap23 | synaptosomal-associated protein 23 | 591.87 | 589.92 | 0.01 | 9.86E-01 |
| ENSMUSG00000027285 | GSMG0021650 | Haus2 | HAUS augmin-like complex, subunit 2 | 780.83 | 954.37 | -0.29 | 1.38E-01 |
| NULL | GSMG0021653 | NULL | NULL | 580.24 | 520.85 | 0.16 | 5.58E-01 |
| ENSMUSG00000054484 | GSMG0021656 | Tmem62 | transmembrane protein 62 | 186.57 | 145.36 | 0.36 | 2.57E-01 |
| ENSMUSG00000023572 | GSMG0021657 | Ccndbp1 | cyclin D-type binding-protein 1 | 809.66 | 848.46 | -0.07 | 8.10E-01 |
| ENSMUSG00000027259 | GSMG0021660 | Adal | adenosine deaminase-like | 194.18 | 214.28 | -0.14 | 7.02E-01 |
| ENSMUSG00000027263 | GSMG0021662 | Tubgcp4 | tubulin, gamma complex associated protein 4 | 800.45 | 819.12 | -0.03 | 9.18E-01 |
| ENSMUSG00000000308 | GSMG0021664 | Ckmt1 | creatine kinase, mitochondrial 1, ubiquitous | 773.75 | 504.81 | 0.62 | 1.23E-03 |
| ENSMUSG00000027248 | GSMG0021665 | Pdia3 | protein disulfide isomerase associated 3 | 11437.21 | 14630.71 | -0.36 | 6.08E-02 |
| ENSMUSG00000027245 | GSMG0021669 | Hypk | huntingtin interacting protein K | 1423.52 | 1958.05 | -0.46 | 4.42E-02 |
| ENSMUSG00000027242 | GSMG0021671 | Wdr76 | WD repeat domain 76 | 125.56 | 154.79 | -0.30 | 3.73E-01 |
| ENSMUSG00000060227 | GSMG0021675 | Casc4 | cancer susceptibility candidate 4 | 372.62 | 461.94 | -0.31 | 2.40E-01 |

| | | | | | | | |
|---|---|---|---|---|---|---|---|
| ENSMUSG00000033411 | GSMG0021678 | Ctdspl2 | CTD (carboxy-terminal domain, RNA polymerase II, polypeptide A) sn | 677.18 | 766.29 | -0.18 | 5.16E-01 |
| ENSMUSG00000060802 | GSMG0021687 | B2m | beta-2 microglobulin | 4369.70 | 4188.93 | 0.06 | 8.51E-01 |
| NULL | GSMG0021699 | NULL | NULL | 1038.54 | 1042.35 | 0.00 | 9.90E-01 |
| ENSMUSG00000005804 | GSMG0021702 | Bloc1s6 | biogenesis of lysosomal organelles complex-1, subunit 6, pallidin | 1034.44 | 1351.66 | -0.39 | 2.90E-02 |
| ENSMUSG00000005803 | GSMG0021703 | Sqrdl | sulfide quinone reductase-like (yeast) | 1044.96 | 933.29 | 0.16 | 4.83E-01 |
| ENSMUSG00000027200 | GSMG0021705 | Sema6d | sema domain, transmembrane domain (TM), and cytoplasmic domain, ( | 2706.94 | 4815.15 | -0.83 | 1.54E-09 |
| NULL | GSMG0021711 | NULL | NULL | 167.73 | 232.58 | -0.47 | 6.78E-02 |
| ENSMUSG00000035183 | GSMG0021713 | Slc24a5 | solute carrier family 24, member 5 | 101.95 | 151.13 | -0.57 | 5.82E-02 |
| ENSMUSG00000027203 | GSMG0021719 | Dut | deoxyuridine triphosphatase | 373.74 | 652.17 | -0.80 | 1.32E-05 |
| ENSMUSG00000091337 | GSMG0021723 | Eid1 | EP300 interacting inhibitor of differentiation 1 | 3219.44 | 4906.27 | -0.61 | 6.03E-04 |
| ENSMUSG00000027207 | GSMG0021726 | Galk2 | galactokinase 2 | 233.11 | 276.44 | -0.24 | 4.58E-01 |
| ENSMUSG00000023330 | GSMG0021731 | Dtwd1 | DTW domain containing 1 | 182.41 | 180.67 | 0.01 | 9.76E-01 |
| ENSMUSG00000027363 | GSMG0021733 | Usp8 | ubiquitin specific peptidase 8 | 1119.44 | 981.38 | 0.19 | 4.16E-01 |
| ENSMUSG00000001998 | GSMG0021735 | Ap4e1 | adaptor-related protein complex AP-4, epsilon 1 | 102.09 | 122.18 | -0.27 | 5.14E-01 |
| ENSMUSG00000001999 | GSMG0021737 | Blvra | biliverdin reductase A | 601.89 | 658.95 | -0.13 | 6.79E-01 |
| ENSMUSG00000003660 | GSMG0021738 | Snrnp200 | small nuclear ribonucleoprotein 200 (U5) | 1015.19 | 1111.96 | -0.13 | 6.44E-01 |
| ENSMUSG00000034850 | GSMG0021739 | Tmem127 | transmembrane protein 127 | 4099.97 | 4375.65 | -0.09 | 6.83E-01 |
| ENSMUSG00000027367 | GSMG0021740 | Stard7 | START domain containing 7 | 791.75 | 931.79 | -0.23 | 2.80E-01 |
| ENSMUSG00000027368 | GSMG0021742 | Dusp2 | dual specificity phosphatase 2 | 616.12 | 1283.46 | -1.06 | 5.20E-08 |
| ENSMUSG00000027374 | GSMG0021747 | Mrps5 | mitochondrial ribosomal protein S5 | 719.41 | 899.33 | -0.32 | 8.72E-02 |
| ENSMUSG00000027381 | GSMG0021754 | Bcl2l11 | BCL2-like 11 (apoptosis facilitator) | 749.22 | 855.03 | -0.19 | 4.67E-01 |
| ENSMUSG00000014361 | GSMG0021758 | Mertk | c-mer proto-oncogene tyrosine kinase | 423.65 | 129.20 | 1.71 | 4.36E-18 |
| ENSMUSG00000014353 | GSMG0021760 | Tmem87b | transmembrane protein 87B | 795.83 | 687.71 | 0.21 | 3.24E-01 |
| ENSMUSG00000027386 | GSMG0021761 | Fbln7 | fibulin 7 | 62.93 | 35.21 | 0.84 | 7.42E-02 |
| ENSMUSG00000027394 | GSMG0021765 | Ttl | tubulin tyrosine ligase | 66.26 | 103.83 | -0.64 | 9.62E-02 |
| ENSMUSG00000027395 | GSMG0021768 | Polr1b | polymerase (RNA) I polypeptide B | 275.40 | 411.74 | -0.58 | 1.87E-02 |
| ENSMUSG00000037938 | GSMG0021769 | Chchd5 | coiled-coil-helix-coiled-coil-helix domain containing 5 | 93.94 | 116.66 | -0.31 | 4.48E-01 |
| ENSMUSG00000027397 | GSMG0021770 | Slc20a1 | solute carrier family 20, member 1 | 363.45 | 462.47 | -0.34 | 1.21E-01 |
| ENSMUSG00000037902 | GSMG0021776 | Sirpa | signal-regulatory protein alpha | 597.61 | 601.16 | -0.01 | 9.86E-01 |
| NULL | GSMG0021777 | NULL | NULL | 82.80 | 41.09 | 1.00 | 1.33E-02 |
| ENSMUSG00000037885 | GSMG0021779 | Stk35 | serine/threonine kinase 35 | 1074.11 | 1024.23 | 0.07 | 8.24E-01 |
| NULL | GSMG0021783 | NULL | NULL | 1272.00 | 1406.34 | -0.14 | 4.87E-01 |
| ENSMUSG00000027405 | GSMG0021785 | Nop56 | NOP56 ribonucleoprotein | 3822.85 | 4659.72 | -0.29 | 9.71E-02 |

| | | | | | | | |
|---|---|---|---|---|---|---|---|
| ENSMUSG00000053552 | GSMG0021787 | Ebf4 | early B cell factor 4 | 248.96 | 159.40 | 0.64 | 6.38E-02 |
| ENSMUSG00000027411 | GSMG0021790 | Vps16 | VSP16 CORVET/HOPS core subunit | 1751.01 | 1895.82 | -0.11 | 6.46E-01 |
| ENSMUSG00000027303 | GSMG0021793 | Ptpra | protein tyrosine phosphatase, receptor type, A | 6735.89 | 7335.65 | -0.12 | 5.56E-01 |
| NULL | GSMG0021795 | NULL | NULL | 183.62 | 136.57 | 0.42 | 3.19E-01 |
| ENSMUSG00000037740 | GSMG0021798 | Mrps26 | mitochondrial ribosomal protein S26 | 508.83 | 536.13 | -0.08 | 8.15E-01 |
| ENSMUSG00000074797 | GSMG0021802 | Itpa | inosine triphosphatase (nucleoside triphosphate pyrophosphatase) | 816.33 | 1018.07 | -0.32 | 1.03E-01 |
| ENSMUSG00000027312 | GSMG0021804 | Atrn | attractin | 879.08 | 902.12 | -0.04 | 9.43E-01 |
| ENSMUSG00000074793 | GSMG0021808 | Hspa12b | heat shock protein 12B | 2160.50 | 1616.33 | 0.42 | 1.02E-01 |
| NULL | GSMG0021809 | NULL | NULL | 545.82 | 470.60 | 0.21 | 4.01E-01 |
| ENSMUSG00000027330 | GSMG0021810 | Cdc25b | cell division cycle 25B | 611.32 | 712.43 | -0.22 | 3.74E-01 |
| ENSMUSG00000068264 | GSMG0021811 | Ap5s1 | adaptor-related protein 5 complex, sigma 1 subunit | 297.80 | 265.95 | 0.16 | 6.02E-01 |
| ENSMUSG00000037523 | GSMG0021812 | Mavs | mitochondrial antiviral signaling protein | 350.44 | 270.25 | 0.37 | 1.55E-01 |
| ENSMUSG00000037514 | GSMG0021813 | Pank2 | pantothenate kinase 2 | 734.34 | 850.61 | -0.21 | 4.04E-01 |
| ENSMUSG00000027333 | GSMG0021815 | Smox | spermine oxidase | 2128.90 | 2462.68 | -0.21 | 2.34E-01 |
| 38,ENSMUSG00000079037,EI | GSMG0021819 | Prn // Prnp // Prnd | prion protein gene complex // prion protein // prion protein dublet | 2085.49 | 5455.60 | -1.39 | 7.51E-19 |
| ENSMUSG00000058793 | GSMG0021823 | Cds2 | CDP-diacylglycerol synthase (phosphatidate cytidylyltransferase) 2 | 1287.64 | 1430.06 | -0.15 | 5.04E-01 |
| ENSMUSG00000044991 | GSMG0021827 | 1110034G24Rik | RIKEN cDNA 1110034G24 gene | 654.31 | 563.53 | 0.22 | 3.44E-01 |
| ENSMUSG00000027350 | GSMG0021828 | Chgb | chromogranin B | 1845.38 | 1776.03 | 0.06 | 8.37E-01 |
| ENSMUSG00000027357 | GSMG0021830 | Crls1 | cardiolipin synthase 1 | 236.40 | 232.60 | 0.02 | 9.62E-01 |
| ENSMUSG00000051177 | GSMG0021836 | Plcb1 | phospholipase C, beta 1 | 1398.99 | 765.22 | 0.87 | 6.93E-09 |
| NULL | GSMG0021841 | NULL | NULL | 905.57 | 512.92 | 0.82 | 1.08E-05 |
| ENSMUSG00000027270 | GSMG0021849 | Lamp5 | lysosomal-associated membrane protein family, member 5 | 489.11 | 78.12 | 2.65 | 4.01E-31 |
| ENSMUSG00000027273 | GSMG0021852 | Snap25 | synaptosomal-associated protein 25 | 1896.95 | 1415.69 | 0.42 | 1.60E-02 |
| ENSMUSG00000062098 | GSMG0021855 | Btbd3 | BTB (POZ) domain containing 3 | 3297.60 | 2381.66 | 0.47 | 2.05E-03 |
| ENSMUSG00000074766 | GSMG0021858 | Ism1 | isthmin 1, angiogenesis inhibitor | 310.89 | 157.11 | 0.98 | 6.25E-06 |
| ENSMUSG00000027384 | GSMG0021860 | Ndufaf5 | NADH dehydrogenase (ubiquinone) complex I, assembly factor 5 | 294.21 | 410.64 | -0.48 | 2.21E-02 |
| NULL | GSMG0021872 | NULL | NULL | 193.91 | 312.56 | -0.69 | 3.26E-03 |
| NULL | GSMG0021873 | NULL | NULL | 165.13 | 134.61 | 0.29 | 4.77E-01 |
| NULL | GSMG0021876 | NULL | NULL | 213.30 | 328.02 | -0.62 | 2.68E-02 |
| NULL | GSMG0021879 | NULL | NULL | 95.81 | 184.06 | -0.93 | 1.34E-03 |
| ENSMUSG00000008333 | GSMG0021882 | Snrpb2 | U2 small nuclear ribonucleoprotein B | 548.56 | 720.13 | -0.39 | 1.90E-01 |
| ENSMUSG00000027419 | GSMG0021884 | Pcsk2 | proprotein convertase subtilisin/kexin type 2 | 654.64 | 700.30 | -0.10 | 7.66E-01 |
| ENSMUSG00000015932 | GSMG0021885 | Dstn | destrin | 7255.84 | 7068.92 | 0.04 | 9.13E-01 |

| | | | | | | | |
|---|---|---|---|---|---|---|---|
| NULL | GSMG0021886 | NULL | NULL | 135.61 | 137.01 | -0.02 | 9.67E-01 |
| ENSMUSG00000027424 | GSMG0021889 | Mgme1 | mitochondrial genome maintainance exonuclease 1 | 155.54 | 186.51 | -0.26 | 4.52E-01 |
| G00000027425,ENSMUSG000 | GSMG0021891 | Pet117 // Csrp2bp | PET117 homolog // cysteine and glycine-rich protein 2 binding protein | 770.06 | 991.40 | -0.37 | 4.50E-02 |
| ENSMUSG00000027427 | GSMG0021895 | Polr3f | polymerase (RNA) III (DNA directed) polypeptide F | 157.08 | 174.61 | -0.15 | 6.81E-01 |
| ENSMUSG00000027429 | GSMG0021896 | Sec23b | SEC23 homolog B, COPII coat complex component | 517.06 | 563.17 | -0.12 | 6.71E-01 |
| ENSMUSG00000074754 | GSMG0021897 | Gm561 | predicted gene 561 | 213.60 | 221.60 | -0.06 | 9.07E-01 |
| ENSMUSG00000027430 | GSMG0021898 | Dtd1 | D-tyrosyl-tRNA deacylase 1 | 358.38 | 490.71 | -0.45 | 2.95E-02 |
| ENSMUSG00000063873 | GSMG0021904 | Slc24a3 | solute carrier family 24 (sodium/potassium/calcium exchanger), membe | 647.30 | 615.64 | 0.07 | 8.67E-01 |
| NULL | GSMG0021911 | NULL | NULL | 256.00 | 302.03 | -0.24 | 4.63E-01 |
| ENSMUSG00000001768 | GSMG0021912 | Rin2 | Ras and Rab interactor 2 | 312.40 | 194.17 | 0.69 | 3.66E-03 |
| ENSMUSG00000002728 | GSMG0021914 | Naa20 | N(alpha)-acetyltransferase 20, NatB catalytic subunit | 1639.55 | 1808.23 | -0.14 | 5.05E-01 |
| ENSMUSG00000074749 | GSMG0021920 | Kiz | kizuna centrosomal protein | 658.42 | 527.31 | 0.32 | 1.96E-01 |
| ENSMUSG00000027433 | GSMG0021921 | Xrn2 | 5'-3' exoribonuclease 2 | 3396.11 | 4773.29 | -0.49 | 4.83E-03 |
| NULL | GSMG0021928 | NULL | NULL | 602.22 | 541.54 | 0.15 | 6.07E-01 |
| ENSMUSG00000036992 | GSMG0021930 | Nxt1 | NTF2-related export protein 1 | 535.53 | 732.17 | -0.45 | 1.84E-02 |
| ENSMUSG00000027439 | GSMG0021931 | Gzf1 | GDNF-inducible zinc finger protein 1 | 628.00 | 629.31 | 0.00 | 9.90E-01 |
| ENSMUSG00000074736 | GSMG0021940 | Syndig1 | synapse differentiation inducing 1 | 262.53 | 491.89 | -0.90 | 7.21E-06 |
| ENSMUSG00000033068 | GSMG0021945 | Entpd6 | ectonucleoside triphosphate diphosphohydrolase 6 | 701.47 | 981.88 | -0.48 | 5.50E-03 |
| ENSMUSG00000033059 | GSMG0021946 | Pygb | brain glycogen phosphorylase | 1709.21 | 2419.05 | -0.50 | 1.01E-02 |
| ENSMUSG00000027454 | GSMG0021949 | Gins1 | GINS complex subunit 1 (Psf1 homolog) | 40.36 | 136.60 | -1.76 | 6.59E-08 |
| ENSMUSG00000027455 | GSMG0021954 | Nsfl1c | NSFL1 (p97) cofactor (p47) | 1491.33 | 1741.21 | -0.22 | 2.93E-01 |
| ENSMUSG00000032966 | GSMG0021956 | Fkbp1a | FK506 binding protein 1a | 3298.95 | 4226.66 | -0.36 | 3.47E-02 |
| ENSMUSG00000044364 | GSMG0021959 | Tmem74b | transmembrane protein 74B | 42.07 | 105.88 | -1.34 | 7.73E-04 |
| ENSMUSG00000027463 | GSMG0021964 | Slc52a3 | solute carrier protein family 52, member 3 | 2015.60 | 836.52 | 1.27 | 8.48E-13 |
| ENSMUSG00000032802 | GSMG0021967 | Srxn1 | sulfiredoxin 1 homolog (S. cerevisiae) | 658.92 | 378.29 | 0.80 | 4.68E-06 |
| ENSMUSG00000068079 | GSMG0021968 | Tcf15 | transcription factor 15 | 89.85 | 90.81 | -0.02 | 9.68E-01 |
| ENSMUSG00000027465 | GSMG0021970 | Tbc1d20 | TBC1 domain family, member 20 | 531.69 | 495.81 | 0.10 | 7.12E-01 |
| ENSMUSG00000000359 | GSMG0021979 | Rem1 | rad and gem related GTP binding protein 1 | 348.28 | 213.43 | 0.71 | 2.55E-03 |
| ENSMUSG00000019188 | GSMG0021980 | H13 | histocompatibility 13 | 2137.40 | 2794.59 | -0.39 | 1.37E-02 |
| ENSMUSG00000042814 | GSMG0021981 | Mcts2 | malignant T cell amplified sequence 2 | 310.30 | 394.72 | -0.35 | 1.21E-01 |
| ENSMUSG00000042745 | GSMG0021983 | Id1 | inhibitor of DNA binding 1 | 4767.09 | 2549.94 | 0.90 | 4.00E-06 |
| ENSMUSG00000009876 | GSMG0021984 | Cox4i2 | cytochrome c oxidase subunit IV isoform 2 | 655.84 | 519.23 | 0.34 | 1.05E-01 |
| ENSMUSG00000027469 | GSMG0021986 | Tpx2 | TPX2, microtubule-associated | 653.42 | 1734.35 | -1.41 | 1.46E-12 |

| | | | | | | | |
|---|---|---|---|---|---|---|---|
| ENSMUSG00000027474 | GSMG0021990 | Ccm2l | cerebral cavernous malformation 2-like | 1443.57 | 841.46 | 0.78 | 5.89E-04 |
| ENSMUSG00000068040 | GSMG0021992 | Tm9sf4 | transmembrane 9 superfamily protein member 4 | 1862.77 | 2217.90 | -0.25 | 1.61E-01 |
| ENSMUSG00000046020 | GSMG0021994 | Pofut1 | protein O-fucosyltransferase 1 | 1108.50 | 947.61 | 0.23 | 2.45E-01 |
| ENSMUSG00000027475 | GSMG0021995 | Kif3b | kinesin family member 3B | 498.29 | 494.43 | 0.01 | 9.79E-01 |
| ENSMUSG00000042548 | GSMG0021996 | Asxl1 | additional sex combs like 1 | 741.57 | 882.96 | -0.25 | 4.67E-01 |
| NULL | GSMG0021998 | NULL | NULL | 141.16 | 164.86 | -0.23 | 5.24E-01 |
| ENSMUSG00000027478 | GSMG0021999 | Dnmt3b | DNA methyltransferase 3B | 260.57 | 526.32 | -1.02 | 3.57E-07 |
| ENSMUSG00000027479 | GSMG0022001 | Mapre1 | microtubule-associated protein, RP/EB family, member 1 | 2329.60 | 2673.81 | -0.20 | 4.20E-01 |
| ENSMUSG00000038533 | GSMG0022016 | Cbfa2t2 | core-binding factor, runt domain, alpha subunit 2, translocated to, 2 (hu | 279.61 | 358.36 | -0.36 | 1.52E-01 |
| ENSMUSG00000038467 | GSMG0022022 | Chmp4b | charged multivesicular body protein 4B | 122.28 | 131.55 | -0.11 | 8.57E-01 |
| ENSMUSG00000027593 | GSMG0022030 | Raly | hnRNP-associated with lethal yellow | 746.90 | 998.80 | -0.42 | 9.02E-02 |
| ENSMUSG00000027598 | GSMG0022033 | Itch | itchy, E3 ubiquitin protein ligase | 450.31 | 450.77 | 0.00 | 9.98E-01 |
| ENSMUSG00000047459 | GSMG0022035 | Dynlrb1 | dynein light chain roadblock-type 1 | 3003.61 | 3472.26 | -0.21 | 3.10E-01 |
| ENSMUSG00000027602 | GSMG0022036 | Map1lc3a | microtubule-associated protein 1 light chain 3 alpha | 607.35 | 590.45 | 0.04 | 9.03E-01 |
| ENSMUSG00000038375 | GSMG0022037 | Trp53inp2 | transformation related protein 53 inducible nuclear protein 2 | 8031.45 | 5391.29 | 0.58 | 4.48E-05 |
| ENSMUSG00000027605 | GSMG0022038 | Acss2 | acyl-CoA synthetase short-chain family member 2 | 570.12 | 415.43 | 0.46 | 3.34E-02 |
| ENSMUSG00000027611 | GSMG0022042 | Procr | protein C receptor, endothelial | 702.35 | 639.13 | 0.14 | 6.43E-01 |
| ENSMUSG00000027612 | GSMG0022043 | Mmp24 | matrix metallopeptidase 24 | 148.56 | 195.96 | -0.40 | 7.28E-01 |
| ENSMUSG00000005881 | GSMG0022048 | Ergic3 | ERGIC and golgi 3 | 3468.01 | 4126.70 | -0.25 | 1.95E-01 |
| ENSMUSG00000038180 | GSMG0022049 | Spag4 | sperm associated antigen 4 | 118.41 | 135.71 | -0.19 | 6.41E-01 |
| ENSMUSG00000067847 | GSMG0022050 | Romo1 | reactive oxygen species modulator 1 | 1174.46 | 1548.50 | -0.40 | 8.23E-02 |
| ENSMUSG00000038116 | GSMG0022052 | Phf20 | PHD finger protein 20 | 1476.63 | 1969.57 | -0.41 | 2.14E-02 |
| ENSMUSG00000027624 | GSMG0022057 | Epb41l1 | erythrocyte membrane protein band 4.1 like 1 | 1724.06 | 1865.94 | -0.11 | 7.66E-01 |
| ENSMUSG00000027628 | GSMG0022058 | Aar2 | AAR2 splicing factor homolog | 696.11 | 748.83 | -0.11 | 6.96E-01 |
| ENSMUSG00000061689 | GSMG0022060 | Dlgap4 | discs, large homolog-associated protein 4 (Drosophila) | 1676.16 | 2712.70 | -0.70 | 3.93E-06 |
| ENSMUSG00000067818 | GSMG0022062 | Myl9 | myosin, light polypeptide 9, regulatory | 8171.24 | 4500.86 | 0.86 | 1.31E-07 |
| ENSMUSG00000062175 | GSMG0022063 | Tgif2 | TGFB-induced factor homeobox 2 | 174.97 | 327.60 | -0.91 | 3.49E-05 |
| ENSMUSG00000027637 | GSMG0022064 | 1110008F13Rik | RIKEN cDNA 1110008F13 gene | 308.42 | 331.18 | -0.10 | 7.99E-01 |
| ENSMUSG00000027642 | GSMG0022066 | Rpn2 | ribophorin II | 6555.36 | 8415.46 | -0.36 | 3.83E-02 |
| ENSMUSG00000063019 | GSMG0022068 | Manbal | mannosidase, beta A, lysosomal-like | 328.46 | 296.49 | 0.15 | 6.31E-01 |
| ENSMUSG00000027646 | GSMG0022069 | Src | Rous sarcoma oncogene | 791.35 | 819.85 | -0.05 | 8.99E-01 |
| ENSMUSG00000067786 | GSMG0022070 | Nnat | neuronatin | 3081.12 | 7328.55 | -1.25 | 6.01E-12 |
| ENSMUSG00000027649 | GSMG0022071 | Ctnnbl1 | catenin, beta like 1 | 1312.16 | 1917.53 | -0.55 | 1.10E-03 |

| | | | | | | | |
|---|---|---|---|---|---|---|---|
| ENSMUSG00000037843 | GSMG0022074 | Vstm2l | V-set and transmembrane domain containing 2-like | 325.42 | 475.68 | -0.55 | 6.35E-02 |
| ENSMUSG00000027651 | GSMG0022076 | Rprd1b | regulation of nuclear pre-mRNA domain containing 1B | 334.63 | 418.82 | -0.32 | 2.05E-01 |
| ENSMUSG00000016024 | GSMG0022080 | Lbp | lipopolysaccharide binding protein | 738.55 | 368.77 | 1.00 | 6.25E-04 |
| ENSMUSG00000044349 | GSMG0022081 | Snhg11 | small nucleolar RNA host gene 11 | 119405.13 | 52717.17 | 1.18 | 2.69E-17 |
| ENSMUSG00000027652 | GSMG0022083 | Ralgapb | Ral GTPase activating protein, beta subunit (non-catalytic) | 329.77 | 314.32 | 0.07 | 8.52E-01 |
| ENSMUSG00000037761 | GSMG0022090 | Actr5 | ARP5 actin-related protein 5 | 669.61 | 748.03 | -0.16 | 5.30E-01 |
| ENSMUSG00000037754 | GSMG0022091 | Ppp1r16b | protein phosphatase 1, regulatory (inhibitor) subunit 16B | 1404.95 | 953.51 | 0.56 | 4.64E-03 |
| ENSMUSG00000027654 | GSMG0022094 | Fam83d | family with sequence similarity 83, member D | 185.94 | 396.48 | -1.10 | 2.84E-07 |
| ENSMUSG00000027655 | GSMG0022095 | Dhx35 | DEAH (Asp-Glu-Ala-His) box polypeptide 35 | 398.86 | 488.07 | -0.29 | 1.91E-01 |
| ENSMUSG00000070544 | GSMG0022099 | Top1 | topoisomerase (DNA) I | 2022.97 | 2343.84 | -0.21 | 5.18E-01 |
| ENSMUSG00000016933 | GSMG0022100 | Plcg1 | phospholipase C, gamma 1 | 1089.48 | 1096.50 | -0.01 | 9.81E-01 |
| NULL | GSMG0022101 | NULL | NULL | 307.18 | 325.11 | -0.08 | 8.51E-01 |
| NULL | GSMG0022102 | NULL | NULL | 667.13 | 597.26 | 0.16 | 5.11E-01 |
| ENSMUSG00000016921 | GSMG0022109 | Srsf6 | serine/arginine-rich splicing factor 6 | 1095.19 | 1480.33 | -0.43 | 9.02E-03 |
| ENSMUSG00000035576 | GSMG0022110 | L3mbtl1 | l(3)mbt-like (Drosophila) | 892.37 | 1060.56 | -0.25 | 3.65E-01 |
| ENSMUSG00000017858 | GSMG0022112 | Ift52 | intraflagellar transport 52 | 616.76 | 681.48 | -0.14 | 5.68E-01 |
| ENSMUSG00000017861 | GSMG0022113 | Mybl2 | myeloblastosis oncogene-like 2 | 126.18 | 388.09 | -1.61 | 1.19E-14 |
| ENSMUSG00000099632 | GSMG0022116 | 2900093K20Rik | RIKEN cDNA 2900093K20 gene | 187.08 | 159.19 | 0.23 | 6.68E-01 |
| ENSMUSG00000017943 | GSMG0022117 | Gdap1l1 | ganglioside-induced differentiation-associated protein 1-like 1 | 143.77 | 236.59 | -0.71 | 5.98E-03 |
| ENSMUSG00000017679 | GSMG0022119 | Ttpal | tocopherol (alpha) transfer protein-like | 297.29 | 287.09 | 0.05 | 8.94E-01 |
| ENSMUSG00000035268 | GSMG0022121 | Pkig | protein kinase inhibitor, gamma | 1673.23 | 1624.90 | 0.04 | 9.19E-01 |
| ENSMUSG00000018326 | GSMG0022126 | Ywhab | tyrosine 3-monooxygenase/tryptophan 5-monooxygenase activation pro | 4959.20 | 5523.29 | -0.16 | 5.08E-01 |
| ENSMUSG00000018209 | GSMG0022129 | Stk4 | serine/threonine kinase 4 | 421.72 | 440.79 | -0.06 | 8.53E-01 |
| ENSMUSG00000045503 | GSMG0022140 | Sys1 | SYS1 Golgi-localized integral membrane protein homolog (S. cerevisia | 2154.68 | 2222.26 | -0.04 | 8.72E-01 |
| ENSMUSG00000017734 | GSMG0022141 | Dbndd2 | dysbindin (dystrobrevin binding protein 1) domain containing 2 | 1877.61 | 1100.77 | 0.77 | 3.13E-07 |
| ENSMUSG00000017721 | GSMG0022143 | Pigt | phosphatidylinositol glycan anchor biosynthesis, class T | 647.55 | 807.34 | -0.32 | 2.89E-01 |
| ENSMUSG00000017299 | GSMG0022153 | Dnttip1 | deoxynucleotidyltransferase, terminal, interacting protein 1 | 1416.41 | 1477.87 | -0.06 | 8.18E-01 |
| ENSMUSG00000001403 | GSMG0022154 | Ube2c | ubiquitin-conjugating enzyme E2C | 397.49 | 930.66 | -1.23 | 1.23E-11 |
| ENSMUSG00000050373 | GSMG0022155 | Snx21 | sorting nexin family member 21 | 924.22 | 981.97 | -0.09 | 7.37E-01 |
| ENSMUSG00000017764 | GSMG0022157 | Zswim1 | zinc finger SWIM-type containing 1 | 365.87 | 437.08 | -0.26 | 3.02E-01 |
| ENSMUSG00000017760 | GSMG0022158 | Ctsa | cathepsin A | 5451.51 | 5091.78 | 0.10 | 6.86E-01 |
| ENSMUSG00000039849 | GSMG0022159 | Pcif1 | PDX1 C-terminal inhibiting factor 1 | 1367.42 | 1718.27 | -0.33 | 5.94E-02 |
| NULL | GSMG0022160 | NULL | NULL | 165.15 | 153.48 | 0.10 | 8.45E-01 |

| | | | | | | | |
|---|---|---|---|---|---|---|---|
| NULL | GSMG0022162 | NULL | NULL | 382.17 | 321.15 | 0.25 | 4.35E-01 |
| ENSMUSG00000017737 | GSMG0022164 | Mmp9 | matrix metallopeptidase 9 | 45.14 | 1046.64 | -4.54 | 1.25E-88 |
| ENSMUSG00000017740 | GSMG0022165 | Slc12a5 | solute carrier family 12, member 5 | 899.33 | 808.06 | 0.15 | 6.84E-01 |
| ENSMUSG00000017652 | GSMG0022166 | Cd40 | CD40 antigen | 808.90 | 1098.07 | -0.44 | 3.47E-02 |
| ENSMUSG00000017897 | GSMG0022170 | Eya2 | EYA transcriptional coactivator and phosphatase 2 | 117.07 | 139.63 | -0.25 | 5.83E-01 |
| ENSMUSG00000027678 | GSMG0022178 | Ncoa3 | nuclear receptor coactivator 3 | 615.91 | 446.42 | 0.46 | 6.59E-02 |
| NULL | GSMG0022182 | NULL | NULL | 168.74 | 146.95 | 0.19 | 6.51E-01 |
| ENSMUSG00000074582 | GSMG0022184 | Arfgef2 | ADP-ribosylation factor guanine nucleotide-exchange factor 2 (brefeldi | 458.19 | 457.65 | 0.00 | 9.99E-01 |
| ENSMUSG00000002718 | GSMG0022186 | Cse1l | chromosome segregation 1-like (S. cerevisiae) | 664.37 | 878.69 | -0.40 | 4.69E-02 |
| ENSMUSG00000017999 | GSMG0022187 | Ddx27 | DEAD (Asp-Glu-Ala-Asp) box polypeptide 27 | 1487.00 | 1655.42 | -0.15 | 4.67E-01 |
| ENSMUSG00000074578 | GSMG0022189 | Zfas1 | zinc finger, NFX1-type containing 1, antisense RNA 1 | 268.70 | 301.29 | -0.16 | 5.82E-01 |
| ENSMUSG00000006418 | GSMG0022193 | Rnf114 | ring finger protein 114 | 2537.46 | 2107.05 | 0.27 | 1.50E-01 |
| ENSMUSG00000056501 | GSMG0022198 | Cebpb | CCAAT/enhancer binding protein (C/EBP), beta | 408.42 | 228.43 | 0.84 | 2.17E-04 |
| ENSMUSG00000027540 | GSMG0022201 | Ptpn1 | protein tyrosine phosphatase, non-receptor type 1 | 1218.72 | 1312.51 | -0.11 | 6.49E-01 |
| NULL | GSMG0022205 | NULL | NULL | 192.67 | 173.58 | 0.15 | 7.03E-01 |
| ENSMUSG00000047907 | GSMG0022215 | Tshz2 | teashirt zinc finger family member 2 | 1038.92 | 1416.11 | -0.45 | 2.35E-02 |
| NULL | GSMG0022218 | NULL | NULL | 422.34 | 652.07 | -0.63 | 1.50E-03 |
| NULL | GSMG0022219 | NULL | NULL | 356.63 | 518.68 | -0.54 | 1.83E-02 |
| ENSMUSG00000052033 | GSMG0022226 | Pfdn4 | prefoldin 4 | 258.81 | 421.07 | -0.71 | 2.11E-02 |
| ENSMUSG00000027560 | GSMG0022227 | Dok5 | docking protein 5 | 199.97 | 635.86 | -1.67 | 3.66E-20 |
| ENSMUSG00000027495 | GSMG0022234 | Fam210b | family with sequence similarity 210, member B | 1893.70 | 2191.01 | -0.21 | 2.74E-01 |
| ENSMUSG00000027498 | GSMG0022235 | Cstf1 | cleavage stimulation factor, 3' pre-RNA, subunit 1 | 415.69 | 577.05 | -0.47 | 1.84E-02 |
| ENSMUSG00000027502 | GSMG0022237 | Rtfdc1 | replication termination factor 2 domain containing 1 | 1055.06 | 1010.62 | 0.06 | 8.23E-01 |
| ENSMUSG00000027509 | GSMG0022243 | Rae1 | ribonucleic acid export 1 | 1267.69 | 1654.68 | -0.38 | 1.68E-02 |
| ENSMUSG00000027519 | GSMG0022252 | Rab22a | RAB22A, member RAS oncogene family | 2779.57 | 2532.05 | 0.13 | 5.28E-01 |
| ENSMUSG00000054455 | GSMG0022255 | Vapb | vesicle-associated membrane protein, associated protein B and C | 226.26 | 197.23 | 0.20 | 5.92E-01 |
| ENSMUSG00000027522 | GSMG0022256 | Stx16 | syntaxin 16 | 193.97 | 161.29 | 0.27 | 4.85E-01 |
| ENSMUSG00000039263 | GSMG0022258 | Npepl1 | aminopeptidase-like 1 | 3252.39 | 2692.66 | 0.27 | 1.61E-01 |
| ENSMUSG00000027523 | GSMG0022260 | Gnas | GNAS (guanine nucleotide binding protein, alpha stimulating) complex | 23658.12 | 24539.63 | -0.05 | 8.25E-01 |
| ENSMUSG00000016253 | GSMG0022262 | Nelfcd | negative elongation factor complex member C/D, Th1l | 2002.92 | 2372.49 | -0.24 | 1.81E-01 |
| ENSMUSG00000027524 | GSMG0022266 | Edn3 | endothelin 3 | 1231.29 | 236.45 | 2.38 | 5.22E-34 |
| ENSMUSG00000094786 | GSMG0022272 | Gm14403 | predicted gene 14403 | 1127.27 | 371.17 | 1.60 | 7.46E-21 |
| ENSMUSG00000074519 | GSMG0022275 | Zfp971 | zinc finger protein 971 | 366.14 | 318.65 | 0.20 | 4.97E-01 |

| | | | | | | | |
|---|---|---|---|---|---|---|---|
| ENSMUSG00000027525 | GSMG0022276 | Phactr3 | phosphatase and actin regulator 3 | 393.60 | 431.09 | -0.13 | 6.45E-01 |
| ENSMUSG00000000305 | GSMG0022284 | Cdh4 | cadherin 4 | 75.04 | 110.35 | -0.56 | 1.39E-01 |
| ENSMUSG00000085704 | GSMG0022287 | 4921531C22Rik | RIKEN cDNA 4921531C22 gene | 264.82 | 214.67 | 0.30 | 3.09E-01 |
| ENSMUSG00000039108 | GSMG0022288 | Lsm14b | LSM family member 14B | 90.84 | 121.61 | -0.42 | 4.12E-01 |
| ENSMUSG00000039086 | GSMG0022289 | Ss18l1 | synovial sarcoma translocation gene on chromosome 18-like 1 | 190.43 | 271.94 | -0.51 | 1.22E-01 |
| ENSMUSG00000039069 | GSMG0022291 | Mtg2 | mitochondrial ribosome associated GTPase 2 | 338.33 | 362.86 | -0.10 | 7.68E-01 |
| ENSMUSG00000039050 | GSMG0022292 | Osbpl2 | oxysterol binding protein-like 2 | 587.62 | 503.92 | 0.22 | 4.55E-01 |
| ENSMUSG00000039001 | GSMG0022294 | Rps21 | ribosomal protein S21 | 557.34 | 718.03 | -0.37 | 3.21E-01 |
| ENSMUSG00000038963 | GSMG0022297 | Slco4a1 | solute carrier organic anion transporter family, member 4a1 | 103.80 | 261.95 | -1.34 | 3.23E-08 |
| ENSMUSG00000027568 | GSMG0022298 | Ntsr1 | neurotensin receptor 1 | 77.37 | 418.49 | -2.43 | 9.14E-27 |
| ENSMUSG00000027569 | GSMG0022299 | Mrgbp | MRG/MORF4L binding protein | 1134.42 | 1260.39 | -0.15 | 4.65E-01 |
| ENSMUSG00000049401 | GSMG0022301 | Ogfr | opioid growth factor receptor | 2222.55 | 2447.68 | -0.14 | 5.28E-01 |
| ENSMUSG00000027570 | GSMG0022302 | Col9a3 | collagen, type IX, alpha 3 | 1434.65 | 2084.43 | -0.54 | 9.95E-03 |
| NULL | GSMG0022304 | NULL | NULL | 268.74 | 273.87 | -0.02 | 9.55E-01 |
| ENSMUSG00000027573 | GSMG0022305 | Gid8 | GID complex subunit 8 | 1111.25 | 1346.90 | -0.28 | 1.54E-01 |
| ENSMUSG00000027575 | GSMG0022312 | Arfgap1 | ADP-ribosylation factor GTPase activating protein 1 | 1178.28 | 1181.52 | 0.00 | 9.93E-01 |
| ENSMUSG00000016344 | GSMG0022315 | Ppdpf | pancreatic progenitor cell differentiation and proliferation factor | 1415.36 | 1713.04 | -0.28 | 2.12E-01 |
| ENSMUSG00000038685 | GSMG0022318 | Rtel1 | regulator of telomere elongation helicase 1 | 1098.39 | 1677.75 | -0.61 | 5.25E-03 |
| G00000027582,ENSMUSG000 | GSMG0022319 | Zgpat // Lime1 | zinc finger, CCCH-type with G patch domain // Lck interacting transme | 2852.66 | 2621.08 | 0.12 | 6.72E-01 |
| ENSMUSG00000085028 | GSMG0022320 | Slc2a4rg-ps | Slc2a4 regulator, pseudogene | 206.09 | 170.11 | 0.27 | 4.56E-01 |
| ENSMUSG00000000827 | GSMG0022323 | Tpd52l2 | tumor protein D52-like 2 | 738.99 | 771.31 | -0.06 | 8.58E-01 |
| ENSMUSG00000000826 | GSMG0022324 | Dnajc5 | DnaJ heat shock protein family (Hsp40) member C5 | 688.04 | 911.26 | -0.41 | 6.42E-02 |
| ENSMUSG00000002455 | GSMG0022327 | Prpf6 | pre-mRNA splicing factor 6 | 595.69 | 600.15 | -0.01 | 9.80E-01 |
| ENSMUSG00000059540 | GSMG0022329 | Tcea2 | transcription elongation factor A (SII), 2 | 1858.60 | 1168.29 | 0.67 | 8.20E-05 |
| NULL | GSMG0022330 | NULL | NULL | 124.26 | 151.22 | -0.28 | 5.14E-01 |
| ENSMUSG00000027589 | GSMG0022333 | Pcmtd2 | protein-L-isoaspartate (D-aspartate) O-methyltransferase domain contai | 915.56 | 782.34 | 0.23 | 2.97E-01 |
| ENSMUSG00000038628 | GSMG0022334 | Polr3k | polymerase (RNA) III (DNA directed) polypeptide K | 765.81 | 927.14 | -0.27 | 2.65E-01 |
| ENSMUSG00000049950 | GSMG0022335 | Rpp38 | ribonuclease P/MRP 38 subunit | 221.45 | 210.40 | 0.07 | 8.67E-01 |
| ENSMUSG00000026646 | GSMG0022340 | Suv39h2 | suppressor of variegation 3-9 homolog 2 (Drosophila) | 65.55 | 103.60 | -0.67 | 8.22E-02 |
| ENSMUSG00000109865 | GSMG0022341 | Hspa14 | heat shock protein 14 | 1347.82 | 1525.94 | -0.18 | 4.17E-01 |
| ENSMUSG00000039449 | GSMG0022345 | Prpf18 | pre-mRNA processing factor 18 | 566.60 | 543.03 | 0.06 | 8.62E-01 |
| ENSMUSG00000026669 | GSMG0022346 | Mcm10 | minichromosome maintenance 10 replication initiation factor | 49.37 | 195.18 | -1.97 | 6.87E-11 |
| ENSMUSG00000026672 | GSMG0022348 | Optn | optineurin | 322.29 | 341.28 | -0.08 | 7.97E-01 |

| | | | | | | | |
|---|---|---|---|---|---|---|---|
| ENSMUSG00000039145 | GSMG0022351 | Camk1d | calcium/calmodulin-dependent protein kinase ID | 167.92 | 165.42 | 0.02 | 9.65E-01 |
| ENSMUSG00000039128 | GSMG0022354 | Cdc123 | cell division cycle 123 | 938.69 | 1157.03 | -0.30 | 1.51E-01 |
| ENSMUSG00000025816 | GSMG0022358 | Sec61a2 | Sec61, alpha subunit 2 (S. cerevisiae) | 992.67 | 928.18 | 0.10 | 6.91E-01 |
| ENSMUSG00000045319 | GSMG0022360 | Proser2 | proline and serine rich 2 | 705.64 | 683.48 | 0.05 | 8.91E-01 |
| ENSMUSG00000002107 | GSMG0022366 | Celf2 | CUGBP, Elav-like family member 2 | 1660.42 | 2031.69 | -0.29 | 1.33E-01 |
| NULL | GSMG0022367 | NULL | NULL | 366.91 | 489.41 | -0.41 | 1.32E-01 |
| NULL | GSMG0022368 | NULL | NULL | 373.52 | 503.72 | -0.43 | 4.07E-02 |
| NULL | GSMG0022369 | NULL | NULL | 80.01 | 119.72 | -0.57 | 1.67E-01 |
| NULL | GSMG0022370 | NULL | NULL | 132.22 | 227.63 | -0.78 | 3.01E-03 |
| NULL | GSMG0022371 | NULL | NULL | 391.78 | 419.24 | -0.10 | 8.15E-01 |
| NULL | GSMG0022373 | NULL | NULL | 722.40 | 864.77 | -0.26 | 3.59E-01 |
| NULL | GSMG0022374 | NULL | NULL | 960.42 | 1123.47 | -0.23 | 3.90E-01 |
| NULL | GSMG0022375 | NULL | NULL | 99.62 | 145.90 | -0.55 | 1.37E-01 |
| NULL | GSMG0022377 | NULL | NULL | 172.92 | 255.62 | -0.56 | 4.58E-02 |
| NULL | GSMG0022379 | NULL | NULL | 897.78 | 1434.60 | -0.68 | 1.20E-05 |
| NULL | GSMG0022380 | NULL | NULL | 1924.04 | 1182.58 | 0.70 | 1.53E-04 |
| NULL | GSMG0022381 | NULL | NULL | 567.61 | 439.05 | 0.37 | 6.55E-02 |
| NULL | GSMG0022382 | 5031426D15Rik | RIKEN cDNA 5031426D15 gene | 2798.66 | 2007.29 | 0.48 | 8.44E-03 |
| NULL | GSMG0022384 | NULL | NULL | 956.72 | 617.07 | 0.63 | 2.94E-03 |
| NULL | GSMG0022385 | NULL | NULL | 546.04 | 384.86 | 0.51 | 1.05E-02 |
| NULL | GSMG0022386 | NULL | NULL | 592.65 | 353.47 | 0.75 | 3.91E-04 |
| NULL | GSMG0022387 | NULL | NULL | 557.95 | 377.26 | 0.56 | 3.99E-02 |
| NULL | GSMG0022388 | NULL | NULL | 482.40 | 380.92 | 0.34 | 1.26E-01 |
| NULL | GSMG0022389 | NULL | NULL | 278.46 | 181.46 | 0.62 | 2.79E-02 |
| NULL | GSMG0022390 | NULL | NULL | 911.27 | 605.23 | 0.59 | 2.93E-03 |
| NULL | GSMG0022391 | NULL | NULL | 526.68 | 399.88 | 0.40 | 9.02E-02 |
| ENSMUSG00000025782 | GSMG0022400 | Taf3 | TATA-box binding protein associated factor 3 | 154.70 | 211.25 | -0.45 | 1.01E-01 |
| ENSMUSG00000025781 | GSMG0022405 | Atp5c1 | ATP synthase, H+ transporting, mitochondrial F1 complex, gamma poly | 4517.32 | 5155.91 | -0.19 | 3.07E-01 |
| ENSMUSG00000037254 | GSMG0022407 | Itih2 | inter-alpha trypsin inhibitor, heavy chain 2 | 288.16 | 265.64 | 0.12 | 7.33E-01 |
| ENSMUSG00000026773 | GSMG0022413 | Pfkfb3 | 6-phosphofructo-2-kinase/fructose-2,6-biphosphatase 3 | 360.04 | 654.02 | -0.86 | 6.40E-06 |
| ENSMUSG00000037197 | GSMG0022415 | Rbm17 | RNA binding motif protein 17 | 1552.26 | 2001.74 | -0.37 | 5.55E-02 |
| ENSMUSG00000058594 | GSMG0022416 | Fbxo18 | F-box protein 18 | 1569.54 | 1401.45 | 0.16 | 4.29E-01 |
| ENSMUSG00000026768 | GSMG0022417 | Itga8 | integrin alpha 8 | 342.83 | 258.53 | 0.41 | 1.37E-01 |

| | | | | | | | |
|---|---|---|---|---|---|---|---|
| ENSMUSG00000026767 | GSMG0022419 | Fam188a | family with sequence similarity 188, member A | 430.33 | 404.42 | 0.09 | 7.89E-01 |
| ENSMUSG00000026727 | GSMG0022424 | Rsu1 | Ras suppressor protein 1 | 2958.00 | 3115.46 | -0.07 | 8.08E-01 |
| ENSMUSG00000026723 | GSMG0022430 | Trdmt1 | tRNA aspartic acid methyltransferase 1 | 142.36 | 153.22 | -0.10 | 8.24E-01 |
| ENSMUSG00000003418 | GSMG0022431 | St8sia6 | ST8 alpha-N-acetyl-neuraminide alpha-2,8-sialyltransferase 6 | 147.39 | 23.23 | 2.67 | 6.82E-15 |
| ENSMUSG00000063275 | GSMG0022432 | Hacd1 | 3-hydroxyacyl-CoA dehydratase 1 | 273.10 | 291.06 | -0.09 | 7.99E-01 |
| ENSMUSG00000026707 | GSMG0022441 | Nsun6 | NOL1/NOP2/Sun domain family member 6 | 103.00 | 117.56 | -0.19 | 6.67E-01 |
| NULL | GSMG0022445 | NULL | NULL | 1549.16 | 590.34 | 1.39 | 3.48E-20 |
| ENSMUSG00000053702 | GSMG0022446 | Nebl | nebulette | 3611.55 | 1556.32 | 1.21 | 2.94E-17 |
| NULL | GSMG0022448 | NULL | NULL | 197.40 | 97.65 | 1.01 | 7.48E-05 |
| ENSMUSG00000026740 | GSMG0022459 | Dnajc1 | DnaJ heat shock protein family (Hsp40) member C1 | 1854.09 | 1882.16 | -0.02 | 9.49E-01 |
| ENSMUSG00000026737 | GSMG0022472 | Pip4k2a | phosphatidylinositol-5-phosphate 4-kinase, type II, alpha | 292.38 | 259.33 | 0.18 | 5.62E-01 |
| ENSMUSG00000036591 | GSMG0022486 | Arhgap21 | Rho GTPase activating protein 21 | 1640.46 | 1862.36 | -0.18 | 5.69E-01 |
| NULL | GSMG0022487 | NULL | NULL | 222.50 | 214.54 | 0.05 | 8.99E-01 |
| ENSMUSG00000058835 | GSMG0022492 | Abi1 | abl-interactor 1 | 872.32 | 937.99 | -0.10 | 6.94E-01 |
| ENSMUSG00000026779 | GSMG0022496 | Mastl | microtubule associated serine/threonine kinase-like | 78.28 | 158.97 | -1.01 | 1.78E-03 |
| ENSMUSG00000026771 | GSMG0022500 | Spopl | speckle-type POZ protein-like | 253.85 | 285.45 | -0.17 | 6.24E-01 |
| ENSMUSG00000026986 | GSMG0022501 | Hnmt | histamine N-methyltransferase | 57.70 | 69.17 | -0.27 | 6.55E-01 |
| ENSMUSG00000004113 | GSMG0022506 | Cacna1b | calcium channel, voltage-dependent, N type, alpha 1B subunit | 130.59 | 184.57 | -0.50 | 1.35E-01 |
| ENSMUSG00000036893 | GSMG0022508 | Ehmt1 | euchromatic histone methyltransferase 1 | 645.26 | 681.20 | -0.08 | 7.96E-01 |
| ENSMUSG00000026972 | GSMG0022511 | Arrdc1 | arrestin domain containing 1 | 1562.59 | 1729.08 | -0.15 | 5.07E-01 |
| ENSMUSG00000036850 | GSMG0022512 | Mrpl41 | mitochondrial ribosomal protein L41 | 2466.11 | 1944.18 | 0.34 | 6.86E-02 |
| ENSMUSG00000059555 | GSMG0022515 | Tor4a | torsin family 4, member A | 994.96 | 819.03 | 0.28 | 1.53E-01 |
| ENSMUSG00000036752 | GSMG0022518 | Tubb4b | tubulin, beta 4B class IVB | 5584.11 | 6345.69 | -0.18 | 3.53E-01 |
| ENSMUSG00000006471 | GSMG0022523 | Ndor1 | NADPH dependent diflavin oxidoreductase 1 | 970.31 | 909.29 | 0.09 | 7.66E-01 |
| ENSMUSG00000026966 | GSMG0022524 | Ssna1 | Sjogren's syndrome nuclear autoantigen 1 | 2190.93 | 2293.65 | -0.07 | 8.02E-01 |
| ENSMUSG00000026959 | GSMG0022525 | Grin1 | glutamate receptor, ionotropic, NMDA1 (zeta 1) | 2258.24 | 1073.17 | 1.07 | 4.85E-07 |
| ENSMUSG00000026958 | GSMG0022527 | Dpp7 | dipeptidylpeptidase 7 | 1245.75 | 1017.68 | 0.29 | 2.10E-01 |
| ENSMUSG00000026956 | GSMG0022528 | Uap1l1 | UDP-N-acteylglucosamine pyrophosphorylase 1-like 1 | 174.72 | 145.51 | 0.27 | 4.80E-01 |
| ENSMUSG00000047617 | GSMG0022529 | BC029214 | cDNA sequence BC029214 | 773.01 | 828.65 | -0.10 | 6.98E-01 |
| ENSMUSG00000015090 | GSMG0022531 | Ptgds | prostaglandin D2 synthase (brain) | 39186.73 | 21274.47 | 0.88 | 5.04E-04 |
| ENSMUSG00000026942 | GSMG0022534 | Traf2 | TNF receptor-associated factor 2 | 525.49 | 542.56 | -0.04 | 8.97E-01 |
| ENSMUSG00000036504 | GSMG0022537 | Phpt1 | phosphohistidine phosphatase 1 | 987.95 | 1221.53 | -0.31 | 1.30E-01 |
| NULL | GSMG0022538 | NULL | NULL | 1732.31 | 971.24 | 0.83 | 5.22E-04 |

| | | | | | | | |
|---|---|---|---|---|---|---|---|
| ENSMUSG00000015087 | GSMG0022539 | Rabl6 | RAB, member RAS oncogene family-like 6 | 1454.38 | 1495.24 | -0.04 | 8.94E-01 |
| ENSMUSG00000026939 | GSMG0022541 | Tmem141 | transmembrane protein 141 | 276.82 | 212.95 | 0.38 | 1.66E-01 |
| NULL | GSMG0022543 | NULL | NULL | 452.59 | 553.05 | -0.29 | 2.48E-01 |
| ENSMUSG00000026933 | GSMG0022547 | Camsap1 | calmodulin regulated spectrin-associated protein 1 | 142.06 | 157.55 | -0.15 | 7.56E-01 |
| ENSMUSG00000036352 | GSMG0022548 | Ubac1 | ubiquitin associated domain containing 1 | 2376.96 | 2721.75 | -0.20 | 2.85E-01 |
| ENSMUSG00000026932 | GSMG0022550 | Nacc2 | nucleus accumbens associated 2, BEN and BTB (POZ) domain containi | 675.40 | 518.97 | 0.38 | 8.39E-02 |
| ENSMUSG00000036327 | GSMG0022556 | Qsox2 | quiescin Q6 sulfhydryl oxidase 2 | 540.63 | 508.29 | 0.09 | 7.98E-01 |
| ENSMUSG00000075467 | GSMG0022559 | Dnlz | DNL-type zinc finger | 1011.03 | 1105.37 | -0.13 | 5.98E-01 |
| ENSMUSG00000036281 | GSMG0022561 | Snapc4 | small nuclear RNA activating complex, polypeptide 4 | 706.96 | 807.18 | -0.19 | 4.75E-01 |
| ENSMUSG00000026927 | GSMG0022562 | Sdccag3 | serologically defined colon cancer antigen 3 | 2312.63 | 2446.73 | -0.08 | 7.29E-01 |
| ENSMUSG00000026925 | GSMG0022563 | Inpp5e | inositol polyphosphate-5-phosphatase E | 513.48 | 486.90 | 0.08 | 8.13E-01 |
| ENSMUSG00000026923 | GSMG0022566 | Notch1 | notch 1 | 6036.14 | 5619.40 | 0.10 | 7.91E-01 |
| ENSMUSG00000026922 | GSMG0022567 | Agpat2 | 1-acylglycerol-3-phosphate O-acyltransferase 2 (lysophosphatidic acid a | 270.35 | 260.89 | 0.05 | 8.89E-01 |
| NULL | GSMG0022568 | Snhg7 | small nucleolar RNA host gene 7 | 137.08 | 181.84 | -0.41 | 1.76E-01 |
| ENSMUSG00000015776 | GSMG0022572 | Med22 | mediator complex subunit 22 | 2241.27 | 1953.94 | 0.20 | 4.48E-01 |
| ENSMUSG00000015790 | GSMG0022573 | Surf1 | surfeit gene 1 | 1048.67 | 915.14 | 0.20 | 3.87E-01 |
| ENSMUSG00000014867 | GSMG0022574 | Surf4 | surfeit gene 4 | 8618.29 | 10230.76 | -0.25 | 1.43E-01 |
| ENSMUSG00000052406 | GSMG0022575 | Rexo4 | REX4, 3'-5' exonuclease | 438.86 | 471.95 | -0.10 | 7.39E-01 |
| ENSMUSG00000036067 | GSMG0022576 | Slc2a6 | solute carrier family 2 (facilitated glucose transporter), member 6 | 527.81 | 399.14 | 0.40 | 1.19E-01 |
| ENSMUSG00000009216 | GSMG0022578 | Fam163b | family with sequence similarity 163, member B | 1154.90 | 939.97 | 0.30 | 1.93E-01 |
| ENSMUSG00000009614 | GSMG0022580 | Sardh | sarcosine dehydrogenase | 385.34 | 393.75 | -0.03 | 9.34E-01 |
| ENSMUSG00000009621 | GSMG0022581 | Vav2 | vav 2 oncogene | 334.15 | 460.97 | -0.46 | 1.30E-01 |
| ENSMUSG00000026918 | GSMG0022583 | Brd3 | bromodomain containing 3 | 1223.56 | 1570.29 | -0.36 | 5.18E-02 |
| ENSMUSG00000026816 | GSMG0022590 | Gtf3c5 | general transcription factor IIIC, polypeptide 5 | 738.07 | 903.44 | -0.29 | 1.48E-01 |
| ENSMUSG00000035666 | GSMG0022593 | Gtf3c4 | general transcription factor IIIC, polypeptide 4 | 470.90 | 524.83 | -0.16 | 5.77E-01 |
| ENSMUSG00000039826 | GSMG0022598 | Trub2 | TruB pseudouridine (psi) synthase family member 2 | 291.55 | 316.63 | -0.12 | 7.28E-01 |
| NULL | GSMG0022600 | NULL | NULL | 380.45 | 38.41 | 3.31 | 8.54E-30 |
| NULL | GSMG0022602 | NULL | NULL | 155.78 | 133.51 | 0.22 | 5.83E-01 |
| ENSMUSG00000039715 | GSMG0022605 | Wdr34 | WD repeat domain 34 | 362.52 | 491.26 | -0.44 | 5.35E-02 |
| ENSMUSG00000015335 | GSMG0022606 | Zdhhc12 | zinc finger, DHHC domain containing 12 | 213.05 | 189.12 | 0.17 | 6.51E-01 |
| ENSMUSG00000039686 | GSMG0022607 | Zer1 | zyg-11 related, cell cycle regulator | 1316.52 | 1076.71 | 0.29 | 4.28E-01 |
| ENSMUSG00000039660 | GSMG0022608 | D2Wsu81e | DNA segment, Chr 2, Wayne State University 81, expressed | 857.55 | 836.94 | 0.04 | 9.11E-01 |
| ENSMUSG00000039648 | GSMG0022609 | Kyat1 | kynurenine aminotransferase 1 | 228.80 | 182.11 | 0.33 | 2.39E-01 |

| | | | | | | | |
|---|---|---|---|---|---|---|---|
| ENSMUSG00000026860 | GSMG0022613 | Sh3glb2 | SH3-domain GRB2-like endophilin B2 | 3078.65 | 2351.89 | 0.39 | 5.34E-02 |
| ENSMUSG00000026853 | GSMG0022614 | Crat | carnitine acetyltransferase | 987.99 | 868.55 | 0.19 | 4.29E-01 |
| ENSMUSG00000089762 | GSMG0022616 | Ier5l | immediate early response 5-like | 778.96 | 283.92 | 1.45 | 1.02E-06 |
| ENSMUSG00000039483 | GSMG0022619 | Asb6 | ankyrin repeat and SOCS box-containing 6 | 1298.30 | 1226.29 | 0.08 | 7.79E-01 |
| ENSMUSG00000050737 | GSMG0022620 | Ptges | prostaglandin E synthase | 2539.42 | 3501.13 | -0.46 | 4.24E-03 |
| ENSMUSG00000026849 | GSMG0022621 | Tor1a | torsin family 1, member A (torsin A) | 693.11 | 789.41 | -0.19 | 4.15E-01 |
| ENSMUSG00000026851 | GSMG0022622 | BC005624 | cDNA sequence BC005624 | 759.64 | 798.64 | -0.07 | 7.97E-01 |
| ENSMUSG00000075415 | GSMG0022623 | Fnbp1 | formin binding protein 1 | 423.63 | 432.82 | -0.03 | 9.27E-01 |
| NULL | GSMG0022624 | NULL | NULL | 382.76 | 363.93 | 0.07 | 8.90E-01 |
| NULL | GSMG0022625 | NULL | NULL | 152.84 | 116.21 | 0.39 | 2.30E-01 |
| ENSMUSG00000087269 | GSMG0022626 | D330023K18Rik | RIKEN cDNA D330023K18 gene | 86.77 | 103.35 | -0.25 | 5.49E-01 |
| NULL | GSMG0022633 | NULL | NULL | 122.79 | 136.50 | -0.15 | 7.78E-01 |
| ENSMUSG00000002550 | GSMG0022634 | Uck1 | uridine-cytidine kinase 1 | 325.83 | 334.68 | -0.03 | 9.35E-01 |
| ENSMUSG00000044627 | GSMG0022635 | Swi5 | SWI5 recombination repair homolog (yeast) | 1582.17 | 2015.04 | -0.35 | 4.03E-02 |
| ENSMUSG00000026825 | GSMG0022637 | Dnm1 | dynamin 1 | 2480.08 | 1498.81 | 0.73 | 1.18E-03 |
| NULL | GSMG0022638 | NULL | NULL | 2565.86 | 1715.50 | 0.58 | 2.57E-03 |
| ENSMUSG00000039195 | GSMG0022640 | 1110008P14Rik | RIKEN cDNA 1110008P14 gene | 1286.48 | 1041.01 | 0.30 | 1.47E-01 |
| ENSMUSG00000026822 | GSMG0022641 | Lcn2 | lipocalin 2 | 113.95 | 51.83 | 1.13 | 4.19E-04 |
| ENSMUSG00000026819 | GSMG0022642 | Slc25a25 | solute carrier family 25 (mitochondrial carrier, phosphate carrier), mem | 583.06 | 429.98 | 0.44 | 7.02E-02 |
| ENSMUSG00000009566 | GSMG0022645 | Fpgs | folylpolyglutamyl synthetase | 445.47 | 815.53 | -0.87 | 3.14E-06 |
| ENSMUSG00000009555 | GSMG0022646 | Cdk9 | cyclin-dependent kinase 9 (CDC2-related kinase) | 1257.15 | 995.89 | 0.34 | 8.58E-02 |
| ENSMUSG00000026797 | GSMG0022650 | Stxbp1 | syntaxin binding protein 1 | 2867.32 | 3416.96 | -0.25 | 2.51E-01 |
| ENSMUSG00000026792 | GSMG0022652 | Lrsam1 | leucine rich repeat and sterile alpha motif containing 1 | 436.50 | 658.52 | -0.59 | 8.22E-03 |
| ENSMUSG00000026791 | GSMG0022653 | Slc2a8 | solute carrier family 2, (facilitated glucose transporter), member 8 | 1508.78 | 1146.64 | 0.40 | 4.12E-02 |
| ENSMUSG00000038860 | GSMG0022654 | Garnl3 | GTPase activating RANGAP domain-like 3 | 2271.56 | 2481.96 | -0.13 | 5.93E-01 |
| NULL | GSMG0022655 | NULL | NULL | 359.45 | 367.23 | -0.03 | 9.43E-01 |
| ENSMUSG00000038831 | GSMG0022656 | Ralgps1 | Ral GEF with PH domain and SH3 binding motif 1 | 497.70 | 646.22 | -0.38 | 9.52E-02 |
| ENSMUSG00000068966 | GSMG0022661 | Zbtb34 | zinc finger and BTB domain containing 34 | 1250.68 | 1588.02 | -0.35 | 1.98E-01 |
| ENSMUSG00000026788 | GSMG0022662 | Zbtb43 | zinc finger and BTB domain containing 43 | 1391.01 | 1109.83 | 0.33 | 6.74E-02 |
| ENSMUSG00000038740 | GSMG0022666 | Mvb12b | multivesicular body subunit 12B | 240.37 | 338.07 | -0.50 | 3.69E-02 |
| ENSMUSG00000038718 | GSMG0022669 | Pbx3 | pre B cell leukemia homeobox 3 | 210.21 | 263.29 | -0.33 | 2.52E-01 |
| ENSMUSG00000026867 | GSMG0022675 | Gapvd1 | GTPase activating protein and VPS9 domains 1 | 3465.98 | 3876.08 | -0.16 | 4.18E-01 |
| ENSMUSG00000070953 | GSMG0022679 | Rabepk | Rab9 effector protein with kelch motifs | 360.35 | 300.11 | 0.26 | 3.05E-01 |

| | | | | | | | |
|---|---|---|---|---|---|---|---|
| NULL | GSMG0022680 | NULL | NULL | 154.35 | 139.05 | 0.14 | 7.39E-01 |
| ENSMUSG00000035949 | GSMG0022681 | Fbxw2 | F-box and WD-40 domain protein 2 | 903.69 | 789.03 | 0.20 | 3.79E-01 |
| ENSMUSG00000026869 | GSMG0022682 | Psmd5 | proteasome (prosome, macropain) 26S subunit, non-ATPase, 5 | 1051.14 | 1105.74 | -0.07 | 8.04E-01 |
| ENSMUSG00000026878 | GSMG0022687 | Rab14 | RAB14, member RAS oncogene family | 6230.52 | 6442.41 | -0.05 | 8.70E-01 |
| ENSMUSG00000026880 | GSMG0022688 | Stom | stomatin | 1687.65 | 875.73 | 0.95 | 3.46E-09 |
| ENSMUSG00000035778 | GSMG0022691 | Ggta1 | glycoprotein galactosyltransferase alpha 1, 3 | 433.08 | 428.58 | 0.02 | 9.71E-01 |
| ENSMUSG00000026895 | GSMG0022698 | Ndufa8 | NADH dehydrogenase (ubiquinone) 1 alpha subcomplex, 8 | 3248.63 | 3381.69 | -0.06 | 8.42E-01 |
| ENSMUSG00000026890 | GSMG0022700 | Lhx6 | LIM homeobox protein 6 | 399.73 | 463.24 | -0.21 | 4.69E-01 |
| ENSMUSG00000026889 | GSMG0022701 | Rbm18 | RNA binding motif protein 18 | 433.20 | 444.38 | -0.04 | 9.16E-01 |
| ENSMUSG00000009030 | GSMG0022708 | Pdcl | phosducin-like | 1662.48 | 1779.84 | -0.10 | 7.18E-01 |
| ENSMUSG00000075376 | GSMG0022711 | Rc3h2 | ring finger and CCCH-type zinc finger domains 2 | 1627.95 | 1541.48 | 0.08 | 7.60E-01 |
| ENSMUSG00000026915 | GSMG0022715 | Strbp | spermatid perinuclear RNA binding protein | 1081.81 | 1109.25 | -0.04 | 8.99E-01 |
| NULL | GSMG0022719 | NULL | NULL | 803.96 | 447.43 | 0.84 | 2.30E-04 |
| ENSMUSG00000035392 | GSMG0022720 | Dennd1a | DENN/MADD domain containing 1A | 561.82 | 700.82 | -0.32 | 1.65E-01 |
| NULL | GSMG0022729 | NULL | NULL | 199.71 | 171.55 | 0.22 | 4.92E-01 |
| ENSMUSG00000026750 | GSMG0022733 | Psmb7 | proteasome (prosome, macropain) subunit, beta type 7 | 1274.03 | 1625.96 | -0.35 | 5.18E-02 |
| ENSMUSG00000026754 | GSMG0022744 | Golga1 | golgi autoantigen, golgin subfamily a, 1 | 573.08 | 667.32 | -0.22 | 4.36E-01 |
| ENSMUSG00000035236 | GSMG0022746 | Scai | suppressor of cancer cell invasion | 296.89 | 301.20 | -0.02 | 9.63E-01 |
| NULL | GSMG0022748 | NULL | NULL | 388.08 | 312.88 | 0.31 | 2.60E-01 |
| ENSMUSG00000026753 | GSMG0022749 | Ppp6c | protein phosphatase 6, catalytic subunit | 697.62 | 720.74 | -0.05 | 8.78E-01 |
| ENSMUSG00000036890 | GSMG0022770 | Gtdc1 | glycosyltransferase-like domain containing 1 | 230.24 | 258.77 | -0.16 | 6.53E-01 |
| ENSMUSG00000026872 | GSMG0022776 | Zeb2 | zinc finger E-box binding homeobox 2 | 1418.94 | 1642.78 | -0.21 | 3.67E-01 |
| NULL | GSMG0022778 | NULL | NULL | 405.38 | 602.27 | -0.57 | 1.20E-02 |
| NULL | GSMG0022779 | NULL | NULL | 135.12 | 173.82 | -0.36 | 3.77E-01 |
| NULL | GSMG0022780 | NULL | NULL | 725.48 | 890.68 | -0.30 | 1.42E-01 |
| NULL | GSMG0022782 | NULL | NULL | 1315.92 | 1794.57 | -0.45 | 1.59E-02 |
| NULL | GSMG0022783 | NULL | NULL | 301.41 | 376.19 | -0.32 | 2.50E-01 |
| ENSMUSG00000026761 | GSMG0022789 | Orc4 | origin recognition complex, subunit 4 | 657.14 | 705.80 | -0.10 | 7.84E-01 |
| ENSMUSG00000026766 | GSMG0022794 | Mmadhc | methylmalonic aciduria (cobalamin deficiency) cblD type, with homocy | 1372.90 | 1607.47 | -0.23 | 2.26E-01 |
| ENSMUSG00000017144 | GSMG0022796 | Rnd3 | Rho family GTPase 3 | 1329.53 | 1536.64 | -0.21 | 3.16E-01 |
| ENSMUSG00000036249 | GSMG0022803 | Rbm43 | RNA binding motif protein 43 | 489.82 | 552.01 | -0.17 | 5.44E-01 |
| ENSMUSG00000026946 | GSMG0022804 | Nmi | N-myc (and STAT) interactor | 605.85 | 632.26 | -0.06 | 8.74E-01 |
| ENSMUSG00000036093 | GSMG0022810 | Arl5a | ADP-ribosylation factor-like 5A | 619.96 | 681.99 | -0.14 | 5.76E-01 |

| | | | | | | | |
|---|---|---|---|---|---|---|---|
| ENSMUSG00000017412 | GSMG0022811 | Cacnb4 | calcium channel, voltage-dependent, beta 4 subunit | 296.20 | 244.01 | 0.28 | 3.80E-01 |
| NULL | GSMG0022813 | NULL | NULL | 553.13 | 360.55 | 0.61 | 5.07E-02 |
| NULL | GSMG0022816 | NULL | NULL | 598.14 | 393.09 | 0.61 | 9.68E-04 |
| ENSMUSG00000055371 | GSMG0022818 | Stam2 | signal transducing adaptor molecule (SH3 domain and ITAM motif) 2 | 684.34 | 683.57 | 0.00 | 9.96E-01 |
| NULL | GSMG0022819 | NULL | NULL | 84.52 | 104.51 | -0.30 | 5.32E-01 |
| ENSMUSG00000061136 | GSMG0022823 | Prpf40a | pre-mRNA processing factor 40A | 1452.54 | 1939.58 | -0.42 | 1.92E-02 |
| ENSMUSG00000075334 | GSMG0022825 | Rprm | reprimo, TP53 dependent G2 arrest mediator candidate | 418.23 | 1007.65 | -1.27 | 2.77E-13 |
| ENSMUSG00000026826 | GSMG0022829 | Nr4a2 | nuclear receptor subfamily 4, group A, member 2 | 473.56 | 451.62 | 0.07 | 8.61E-01 |
| ENSMUSG00000026836 | GSMG0022840 | Acvr1 | activin A receptor, type 1 | 595.84 | 431.60 | 0.47 | 1.30E-01 |
| ENSMUSG00000026988 | GSMG0022847 | Wdsub1 | WD repeat, SAM and U-box domain containing 1 | 592.40 | 493.38 | 0.26 | 3.39E-01 |
| ENSMUSG00000026987 | GSMG0022849 | Baz2b | bromodomain adjacent to zinc finger domain, 2B | 1533.95 | 1506.19 | 0.03 | 9.41E-01 |
| ENSMUSG00000060703 | GSMG0022854 | Cd302 | CD302 antigen | 611.02 | 576.09 | 0.08 | 7.89E-01 |
| ENSMUSG00000026980 | GSMG0022855 | Ly75 | lymphocyte antigen 75 | 3723.54 | 4220.16 | -0.18 | 4.12E-01 |
| ENSMUSG00000054580 | GSMG0022857 | Pla2r1 | phospholipase A2 receptor 1 | 290.68 | 177.46 | 0.71 | 2.82E-03 |
| NULL | GSMG0022861 | NULL | NULL | 173.57 | 179.15 | -0.05 | 9.34E-01 |
| ENSMUSG00000026970 | GSMG0022862 | Rbms1 | RNA binding motif, single stranded interacting protein 1 | 1765.17 | 1556.71 | 0.18 | 3.60E-01 |
| ENSMUSG00000035000 | GSMG0022873 | Dpp4 | dipeptidylpeptidase 4 | 1294.45 | 957.44 | 0.44 | 9.42E-02 |
| ENSMUSG00000059742 | GSMG0022880 | Kcnh7 | potassium voltage-gated channel, subfamily H (eag-related), member 7 | 978.40 | 656.93 | 0.58 | 1.76E-03 |
| NULL | GSMG0022884 | NULL | NULL | 730.66 | 746.81 | -0.03 | 9.45E-01 |
| ENSMUSG00000026888 | GSMG0022898 | Grb14 | growth factor receptor bound protein 14 | 940.40 | 707.87 | 0.41 | 2.80E-02 |
| ENSMUSG00000034903 | GSMG0022900 | Cobll1 | Cobl-like 1 | 1866.40 | 1327.01 | 0.49 | 3.34E-03 |
| ENSMUSG00000061171 | GSMG0022904 | Slc38a11 | solute carrier family 38, member 11 | 408.69 | 139.01 | 1.56 | 6.83E-14 |
| ENSMUSG00000034848 | GSMG0022911 | Ttc21b | tetratricopeptide repeat domain 21B | 305.88 | 468.50 | -0.61 | 4.87E-03 |
| NULL | GSMG0022916 | NULL | NULL | 71.26 | 37.40 | 0.92 | 7.03E-02 |
| NULL | GSMG0022919 | NULL | NULL | 79.63 | 64.05 | 0.32 | 5.30E-01 |
| ENSMUSG00000075316 | GSMG0022921 | Scn9a | sodium channel, voltage-gated, type IX, alpha | 19.95 | 258.44 | -3.69 | 3.57E-36 |
| ENSMUSG00000027030 | GSMG0022927 | Stk39 | serine/threonine kinase 39 | 664.60 | 674.45 | -0.02 | 9.48E-01 |
| NULL | GSMG0022931 | NULL | NULL | 74.92 | 45.44 | 0.73 | 9.28E-02 |
| ENSMUSG00000005233 | GSMG0022932 | Spc25 | SPC25, NDC80 kinetochore complex component, homolog (S. cerevisi | 303.51 | 662.69 | -1.13 | 3.96E-06 |
| ENSMUSG00000027086 | GSMG0022937 | Fastkd1 | FAST kinase domains 1 | 59.04 | 70.03 | -0.25 | 6.44E-01 |
| ENSMUSG00000070883 | GSMG0022938 | Ccdc173 | coiled-coil domain containing 173 | 117.77 | 89.77 | 0.39 | 3.39E-01 |
| ENSMUSG00000051730 | GSMG0022939 | Mettl5 | methyltransferase like 5 | 230.14 | 239.72 | -0.06 | 8.88E-01 |
| ENSMUSG00000041997 | GSMG0022944 | Tlk1 | tousled-like kinase 1 | 501.77 | 499.72 | 0.00 | 9.91E-01 |

| | | | | | | | |
|---|---|---|---|---|---|---|---|
| NULL | GSMG0022946 | NULL | NULL | 593.79 | 581.31 | 0.03 | 9.37E-01 |
| ENSMUSG00000041975 | GSMG0022948 | Mettl8 | methyltransferase like 8 | 113.33 | 123.36 | -0.12 | 8.09E-01 |
| ENSMUSG00000027010 | GSMG0022952 | Slc25a12 | solute carrier family 25 (mitochondrial carrier, Aralar), member 12 | 1299.18 | 1574.07 | -0.28 | 1.44E-01 |
| ENSMUSG00000084946 | GSMG0022957 | Dlx1as | distal-less homeobox 1, antisense | 110.11 | 262.62 | -1.26 | 1.54E-07 |
| ENSMUSG00000027109 | GSMG0022967 | Sp3 | trans-acting transcription factor 3 | 2019.02 | 1830.08 | 0.14 | 4.90E-01 |
| ENSMUSG00000027108 | GSMG0022970 | Ola1 | Obg-like ATPase 1 | 349.30 | 488.08 | -0.48 | 1.58E-02 |
| ENSMUSG00000041762 | GSMG0022973 | Gpr155 | G protein-coupled receptor 155 | 1100.34 | 828.46 | 0.41 | 2.20E-02 |
| ENSMUSG00000056486 | GSMG0022983 | Chn1 | chimerin 1 | 1267.57 | 660.36 | 0.94 | 3.34E-08 |
| ENSMUSG00000027104 | GSMG0022984 | Atf2 | activating transcription factor 2 | 530.98 | 637.63 | -0.26 | 2.45E-01 |
| ENSMUSG00000018770 | GSMG0022985 | Atp5g3 | ATP synthase, H+ transporting, mitochondrial F0 complex, subunit C3 | 2778.05 | 3215.78 | -0.21 | 3.17E-01 |
| ENSMUSG00000009207 | GSMG0022990 | Lnp | limb and neural patterns | 180.37 | 282.25 | -0.65 | 5.15E-03 |
| ENSMUSG00000015839 | GSMG0022999 | Nfe2l2 | nuclear factor, erythroid derived 2, like 2 | 2838.04 | 2384.41 | 0.25 | 2.42E-01 |
| ENSMUSG00000075273 | GSMG0023002 | Ttc30b | tetratricopeptide repeat domain 30B | 398.50 | 415.78 | -0.06 | 8.57E-01 |
| ENSMUSG00000002731 | GSMG0023008 | Prkra | protein kinase, interferon inducible double stranded RNA dependent ac | 306.23 | 450.30 | -0.55 | 4.14E-02 |
| ENSMUSG00000002732 | GSMG0023009 | Fkbp7 | FK506 binding protein 7 | 716.28 | 653.37 | 0.13 | 6.81E-01 |
| NULL | GSMG0023025 | NULL | NULL | 295.05 | 310.29 | -0.08 | 8.75E-01 |
| ENSMUSG00000044033 | GSMG0023026 | Ccdc141 | coiled-coil domain containing 141 | 5071.58 | 4072.58 | 0.32 | 1.50E-01 |
| ENSMUSG00000042272 | GSMG0023029 | Sestd1 | SEC14 and spectrin domains 1 | 1155.60 | 1341.90 | -0.21 | 4.45E-01 |
| ENSMUSG00000027016 | GSMG0023030 | Zfp385b | zinc finger protein 385B | 102.67 | 94.79 | 0.12 | 8.43E-01 |
| ENSMUSG00000027014 | GSMG0023032 | Cwc22 | CWC22 spliceosome-associated protein | 2919.94 | 6073.72 | -1.06 | 3.90E-07 |
| ENSMUSG00000059173 | GSMG0023037 | Pde1a | phosphodiesterase 1A, calmodulin-dependent | 472.64 | 362.42 | 0.39 | 1.42E-01 |
| NULL | GSMG0023038 | NULL | NULL | 285.01 | 130.75 | 1.12 | 4.72E-07 |
| NULL | GSMG0023039 | NULL | NULL | 319.33 | 231.40 | 0.47 | 5.27E-02 |
| ENSMUSG00000027004 | GSMG0023040 | Frzb | frizzled-related protein | 248.20 | 417.35 | -0.74 | 6.60E-04 |
| ENSMUSG00000027002 | GSMG0023043 | Nckap1 | NCK-associated protein 1 | 11013.46 | 11107.41 | -0.01 | 9.64E-01 |
| ENSMUSG00000059588 | GSMG0023050 | Calcrl | calcitonin receptor-like | 698.44 | 278.15 | 1.33 | 6.41E-09 |
| ENSMUSG00000027082 | GSMG0023051 | Tfpi | tissue factor pathway inhibitor | 861.74 | 1019.99 | -0.24 | 3.49E-01 |
| NULL | GSMG0023053 | NULL | NULL | 320.83 | 245.71 | 0.39 | 1.25E-01 |
| ENSMUSG00000034101 | GSMG0023054 | Ctnnd1 | catenin (cadherin associated protein), delta 1 | 3013.63 | 3227.60 | -0.10 | 7.05E-01 |
| NULL | GSMG0023056 | NULL | NULL | 439.38 | 316.74 | 0.48 | 8.77E-02 |
| NULL | GSMG0023057 | NULL | NULL | 201.92 | 207.99 | -0.04 | 9.24E-01 |
| ENSMUSG00000076437 | GSMG0023058 | 2700094K13Rik | RIKEN cDNA 2700094K13 gene | 824.41 | 1653.65 | -1.00 | 1.76E-08 |
| ENSMUSG00000050043 | GSMG0023059 | Tmx2 | thioredoxin-related transmembrane protein 2 | 679.80 | 739.44 | -0.12 | 6.49E-01 |

| | | | | | | | |
|---|---|---|---|---|---|---|---|
| ENSMUSG00000034075 | GSMG0023060 | Zdhhc5 | zinc finger, DHHC domain containing 5 | 1815.85 | 1873.21 | -0.04 | 8.92E-01 |
| ENSMUSG00000027079 | GSMG0023061 | Clp1 | CLP1, cleavage and polyadenylation factor I subunit | 191.03 | 178.47 | 0.10 | 8.15E-01 |
| ENSMUSG00000023224 | GSMG0023064 | Serping1 | serine (or cysteine) peptidase inhibitor, clade G, member 1 | 1186.71 | 472.29 | 1.33 | 9.90E-11 |
| ENSMUSG00000050896 | GSMG0023067 | Rtn4rl2 | reticulon 4 receptor-like 2 | 122.02 | 161.88 | -0.41 | 3.14E-01 |
| ENSMUSG00000075224 | GSMG0023071 | Lrrc55 | leucine rich repeat containing 55 | 1239.52 | 1820.58 | -0.55 | 2.25E-03 |
| ENSMUSG00000025314 | GSMG0023152 | Ptprj | protein tyrosine phosphatase, receptor type, J | 838.15 | 357.87 | 1.23 | 1.28E-07 |
| NULL | GSMG0023159 | NULL | NULL | 1485.15 | 1839.76 | -0.31 | 3.13E-01 |
| ENSMUSG00000063235 | GSMG0023162 | Ptpmt1 | protein tyrosine phosphatase, mitochondrial 1 | 1417.31 | 1385.58 | 0.03 | 9.09E-01 |
| ENSMUSG00000002105 | GSMG0023165 | Slc39a13 | solute carrier family 39 (metal ion transporter), member 13 | 351.89 | 293.13 | 0.26 | 3.25E-01 |
| ENSMUSG00000040687 | GSMG0023167 | Madd | MAP-kinase activating death domain | 2185.26 | 1829.17 | 0.26 | 3.22E-01 |
| ENSMUSG00000002108 | GSMG0023168 | Nr1h3 | nuclear receptor subfamily 1, group H, member 3 | 417.45 | 325.88 | 0.36 | 1.73E-01 |
| ENSMUSG00000002109 | GSMG0023171 | Ddb2 | damage specific DNA binding protein 2 | 117.65 | 85.32 | 0.47 | 2.27E-01 |
| ENSMUSG00000040591 | GSMG0023174 | 1110051M20Rik | RIKEN cDNA 1110051M20 gene | 413.27 | 524.98 | -0.34 | 1.07E-01 |
| ENSMUSG00000075040 | GSMG0023182 | Zfp408 | zinc finger protein 408 | 387.22 | 415.50 | -0.10 | 7.74E-01 |
| ENSMUSG00000027244 | GSMG0023183 | Atg13 | autophagy related 13 | 1035.69 | 843.41 | 0.30 | 1.11E-01 |
| ENSMUSG00000027239 | GSMG0023185 | Mdk | midkine | 227.63 | 224.84 | 0.02 | 9.66E-01 |
| ENSMUSG00000040479 | GSMG0023186 | Dgkz | diacylglycerol kinase zeta | 1529.42 | 1182.12 | 0.37 | 1.71E-01 |
| ENSMUSG00000027230 | GSMG0023188 | Creb3l1 | cAMP responsive element binding protein 3-like 1 | 211.04 | 355.06 | -0.75 | 5.56E-04 |
| ENSMUSG00000027223 | GSMG0023193 | Mapk8ip1 | mitogen-activated protein kinase 8 interacting protein 1 | 1511.50 | 1380.32 | 0.13 | 7.54E-01 |
| ENSMUSG00000068742 | GSMG0023194 | Cry2 | cryptochrome 2 (photolyase-like) | 1373.05 | 1036.07 | 0.41 | 3.47E-02 |
| ENSMUSG00000049922 | GSMG0023195 | Slc35c1 | solute carrier family 35, member C1 | 301.36 | 237.93 | 0.34 | 2.92E-01 |
| ENSMUSG00000027217 | GSMG0023200 | Tspan18 | tetraspanin 18 | 1060.24 | 2561.84 | -1.27 | 3.06E-19 |
| NULL | GSMG0023203 | NULL | NULL | 103.86 | 212.65 | -1.03 | 5.83E-05 |
| ENSMUSG00000027215 | GSMG0023204 | Cd82 | CD82 antigen | 1404.95 | 2395.26 | -0.77 | 5.36E-06 |
| ENSMUSG00000027198 | GSMG0023205 | Ext2 | exostoses (multiple) 2 | 4482.39 | 4796.50 | -0.10 | 6.90E-01 |
| ENSMUSG00000040272 | GSMG0023206 | Accs | 1-aminocyclopropane-1-carboxylate synthase (non-functional) | 130.70 | 193.90 | -0.57 | 4.62E-02 |
| ENSMUSG00000040174 | GSMG0023210 | Alkbh3 | alkB homolog 3, alpha-ketoglutarate-dependent dioxygenase | 573.73 | 660.48 | -0.20 | 4.46E-01 |
| ENSMUSG00000027195 | GSMG0023211 | Hsd17b12 | hydroxysteroid (17-beta) dehydrogenase 12 | 773.57 | 1022.19 | -0.40 | 4.87E-02 |
| ENSMUSG00000027194 | GSMG0023216 | Ttc17 | tetratricopeptide repeat domain 17 | 475.97 | 460.46 | 0.05 | 9.13E-01 |
| G00000027165,ENSMUSG000 | GSMG0023225 | B230118H07Rik // | RIKEN cDNA B230118H07 gene // recombination activating gene 1 | 291.30 | 308.66 | -0.08 | 8.47E-01 |
| ENSMUSG00000032841 | GSMG0023227 | Prr5l | proline rich 5 like | 600.79 | 444.66 | 0.44 | 3.12E-02 |
| NULL | GSMG0023231 | NULL | NULL | 190.47 | 162.87 | 0.23 | 5.47E-01 |
| ENSMUSG00000048058 | GSMG0023232 | Ldlrad3 | low density lipoprotein receptor class A domain containing 3 | 1335.86 | 1318.66 | 0.02 | 9.49E-01 |

| | | | | | | | |
|---|---|---|---|---|---|---|---|
| ENSMUSG00000027189 | GSMG0023234 | Trim44 | tripartite motif-containing 44 | 4425.74 | 4314.12 | 0.04 | 9.20E-01 |
| ENSMUSG00000075012 | GSMG0023237 | Fjx1 | four jointed box 1 (Drosophila) | 1264.94 | 1094.13 | 0.21 | 3.87E-01 |
| ENSMUSG00000087185 | GSMG0023238 | Gm13872 | predicted gene 13872 | 159.44 | 55.92 | 1.51 | 2.09E-06 |
| ENSMUSG00000010914 | GSMG0023242 | Pdhx | pyruvate dehydrogenase complex, component X | 685.00 | 811.20 | -0.24 | 3.25E-01 |
| ENSMUSG00000027187 | GSMG0023247 | Cat | catalase | 3712.99 | 5279.92 | -0.51 | 7.75E-04 |
| ENSMUSG00000027185 | GSMG0023249 | Nat10 | N-acetyltransferase 10 | 894.46 | 879.66 | 0.02 | 9.36E-01 |
| ENSMUSG00000027184 | GSMG0023250 | Caprin1 | cell cycle associated protein 1 | 4449.01 | 5200.60 | -0.22 | 2.51E-01 |
| ENSMUSG00000068373 | GSMG0023255 | D430041D05Rik | RIKEN cDNA D430041D05 gene | 133.93 | 189.41 | -0.50 | 1.56E-01 |
| ENSMUSG00000027177 | GSMG0023259 | Hipk3 | homeodomain interacting protein kinase 3 | 2725.44 | 2491.55 | 0.13 | 7.25E-01 |
| NULL | GSMG0023262 | NULL | NULL | 126.95 | 174.74 | -0.47 | 1.54E-01 |
| ENSMUSG00000027175 | GSMG0023264 | Tcp11l1 | t-complex 11 like 1 | 289.93 | 403.74 | -0.48 | 2.86E-02 |
| ENSMUSG00000074994 | GSMG0023267 | Qser1 | glutamine and serine rich 1 | 136.06 | 151.34 | -0.16 | 7.09E-01 |
| ENSMUSG00000027170 | GSMG0023271 | Eif3m | eukaryotic translation initiation factor 3, subunit M | 1289.88 | 1677.03 | -0.38 | 3.45E-02 |
| ENSMUSG00000005973 | GSMG0023278 | Rcn1 | reticulocalbin 1 | 1486.31 | 2050.94 | -0.46 | 1.70E-02 |
| ENSMUSG00000027167 | GSMG0023282 | Elp4 | elongator acetyltransferase complex subunit 4 | 260.97 | 326.77 | -0.32 | 2.28E-01 |
| ENSMUSG00000027166 | GSMG0023290 | Dnajc24 | DnaJ heat shock protein family (Hsp40) member C24 | 327.91 | 328.78 | -0.01 | 9.86E-01 |
| ENSMUSG00000027122 | GSMG0023294 | Arl14ep | ADP-ribosylation factor-like 14 effector protein | 546.46 | 639.80 | -0.23 | 5.10E-01 |
| ENSMUSG00000057234 | GSMG0023297 | Mettl15 | methyltransferase like 15 | 104.52 | 123.01 | -0.23 | 5.97E-01 |
| ENSMUSG00000074968 | GSMG0023306 | Ano3 | anoctamin 3 | 367.04 | 144.56 | 1.34 | 5.39E-12 |
| ENSMUSG00000027131 | GSMG0023329 | Emc4 | ER membrane protein complex subunit 4 | 2284.24 | 2347.58 | -0.04 | 8.89E-01 |
| ENSMUSG00000023236 | GSMG0023341 | Scg5 | secretogranin V | 1812.33 | 2013.53 | -0.15 | 5.13E-01 |
| ENSMUSG00000041219 | GSMG0023343 | Arhgap11a | Rho GTPase activating protein 11A | 312.94 | 706.33 | -1.17 | 6.69E-09 |
| ENSMUSG00000040383 | GSMG0023346 | Aqr | aquarius | 197.78 | 279.28 | -0.50 | 7.55E-02 |
| ENSMUSG00000057147 | GSMG0023351 | Dph6 | diphthamine biosynthesis 6 | 544.12 | 608.69 | -0.16 | 5.78E-01 |
| NULL | GSMG0023359 | NULL | NULL | 122.70 | 173.88 | -0.51 | 1.57E-01 |
| ENSMUSG00000027347 | GSMG0023366 | Rasgrp1 | RAS guanyl releasing protein 1 | 784.45 | 275.54 | 1.51 | 1.48E-19 |
| ENSMUSG00000009549 | GSMG0023379 | Srp14 | signal recognition particle 14 | 3173.92 | 3036.36 | 0.06 | 8.10E-01 |
| ENSMUSG00000045838 | GSMG0023384 | A430105I19Rik | RIKEN cDNA A430105I19 gene | 219.61 | 166.31 | 0.40 | 1.40E-01 |
| ENSMUSG00000039983 | GSMG0023385 | Ccdc32 | coiled-coil domain containing 32 | 831.70 | 858.99 | -0.05 | 8.70E-01 |
| ENSMUSG00000070730 | GSMG0023386 | Rmdn3 | regulator of microtubule dynamics 3 | 1207.96 | 1354.43 | -0.16 | 4.26E-01 |
| ENSMUSG00000034278 | GSMG0023387 | Dnajc17 | DnaJ heat shock protein family (Hsp40) member C17 | 181.31 | 251.99 | -0.48 | 6.45E-02 |
| ENSMUSG00000034226 | GSMG0023390 | Rhov | ras homolog family member V | 59.71 | 157.96 | -1.40 | 4.85E-07 |
| ENSMUSG00000034154 | GSMG0023392 | Ino80 | INO80 complex subunit | 226.62 | 287.67 | -0.34 | 2.02E-01 |

| | | | | | | | |
|---|---|---|---|---|---|---|---|
| ENSMUSG00000027305 | GSMG0023399 | Ndufaf1 | NADH dehydrogenase (ubiquinone) 1 alpha subcomplex, assembly fact | 346.36 | 362.87 | -0.07 | 8.69E-01 |
| ENSMUSG00000027297 | GSMG0023401 | Ltk | leukocyte tyrosine kinase | 304.91 | 74.57 | 2.03 | 5.01E-10 |
| ENSMUSG00000034032 | GSMG0023402 | Rpap1 | RNA polymerase II associated protein 1 | 1326.45 | 1349.95 | -0.02 | 9.44E-01 |
| NULL | GSMG0023405 | NULL | NULL | 125.93 | 140.74 | -0.17 | 6.99E-01 |
| ENSMUSG00000027293 | GSMG0023406 | Ehd4 | EH-domain containing 4 | 849.91 | 686.52 | 0.31 | 1.69E-01 |
| ENSMUSG00000050211 | GSMG0023407 | Pla2g4e | phospholipase A2, group IVE | 890.00 | 1093.95 | -0.30 | 1.96E-01 |
| ENSMUSG00000027291 | GSMG0023410 | Vps39 | VPS39 HOPS complex subunit | 1028.11 | 953.67 | 0.11 | 6.75E-01 |
| ENSMUSG00000033808 | GSMG0023411 | Tmem87a | transmembrane protein 87A | 1275.24 | 1272.50 | 0.00 | 9.90E-01 |
| ENSMUSG00000027286 | GSMG0023418 | Lrrc57 | leucine rich repeat containing 57 | 1012.41 | 768.43 | 0.40 | 4.69E-02 |
| G00000027284,ENSMUSG000 | GSMG0023419 | Cdan1 // Ttbk2 | congenital dyserythropoietic anemia, type I (human) // tau tubulin kinas | 342.35 | 335.83 | 0.03 | 9.41E-01 |
| ENSMUSG00000027272 | GSMG0023421 | Ubr1 | ubiquitin protein ligase E3 component n-recognin 1 | 1290.80 | 1287.13 | 0.01 | 9.88E-01 |
| NULL | GSMG0023429 | NULL | NULL | 411.05 | 481.47 | -0.23 | 3.85E-01 |
| ENSMUSG00000050619 | GSMG0023430 | Zscan29 | zinc finger SCAN domains 29 | 147.56 | 123.03 | 0.27 | 4.84E-01 |
| ENSMUSG00000043909 | GSMG0023431 | Trp53bp1 | transformation related protein 53 binding protein 1 | 819.51 | 975.38 | -0.25 | 2.94E-01 |
| ENSMUSG00000033526 | GSMG0023434 | Ppip5k1 | diphosphoinositol pentakisphosphate kinase 1 | 473.34 | 290.93 | 0.70 | 1.15E-03 |
| ENSMUSG00000033486 | GSMG0023436 | Catsper2 | cation channel, sperm associated 2 | 166.64 | 179.75 | -0.11 | 7.79E-01 |
| ENSMUSG00000046110 | GSMG0023438 | Serinc4 | serine incorporator 4 | 613.96 | 586.72 | 0.07 | 8.22E-01 |
| ENSMUSG00000027238 | GSMG0023441 | Frmd5 | FERM domain containing 5 | 361.77 | 445.92 | -0.30 | 2.72E-01 |
| ENSMUSG00000033396 | GSMG0023448 | Spg11 | spastic paraplegia 11 | 395.81 | 368.74 | 0.10 | 8.01E-01 |
| ENSMUSG00000033256 | GSMG0023453 | Shf | Src homology 2 domain containing F | 313.95 | 271.42 | 0.21 | 5.88E-01 |
| ENSMUSG00000027199 | GSMG0023455 | Gatm | glycine amidinotransferase (L-arginine:glycine amidinotransferase) | 6996.23 | 5588.71 | 0.32 | 4.52E-02 |
| ENSMUSG00000005802 | GSMG0023456 | Slc30a4 | solute carrier family 30 (zinc transporter), member 4 | 255.64 | 226.79 | 0.17 | 6.40E-01 |
| ENSMUSG00000027204 | GSMG0023466 | Fbn1 | fibrillin 1 | 233.31 | 263.30 | -0.17 | 6.94E-01 |
| ENSMUSG00000068394 | GSMG0023470 | Cep152 | centrosomal protein 152 | 167.53 | 253.44 | -0.59 | 2.58E-02 |
| ENSMUSG00000035093 | GSMG0023472 | Secisbp2l | SECIS binding protein 2-like | 1116.19 | 1127.82 | -0.02 | 9.59E-01 |
| ENSMUSG00000027206 | GSMG0023473 | Cops2 | COP9 signalosome subunit 2 | 567.85 | 632.29 | -0.16 | 6.28E-01 |
| ENSMUSG00000027361 | GSMG0023479 | Gabpb1 | GA repeat binding protein, beta 1 | 369.59 | 410.62 | -0.15 | 6.15E-01 |
| ENSMUSG00000027365 | GSMG0023483 | Trpm7 | transient receptor potential cation channel, subfamily M, member 7 | 2734.03 | 2629.13 | 0.06 | 8.20E-01 |
| ENSMUSG00000027366 | GSMG0023484 | Sppl2a | signal peptide peptidase like 2A | 1457.72 | 1587.16 | -0.12 | 6.30E-01 |
| ENSMUSG00000034906 | GSMG0023485 | Ncaph | non-SMC condensin I complex, subunit H | 956.40 | 1926.99 | -1.01 | 2.09E-08 |
| ENSMUSG00000074825 | GSMG0023486 | Itpripl1 | inositol 1,4,5-triphosphate receptor interacting protein-like 1 | 420.79 | 477.01 | -0.18 | 5.46E-01 |
| ENSMUSG00000003662 | GSMG0023488 | Ciao1 | cytosolic iron-sulfur protein assembly 1 | 598.67 | 543.85 | 0.14 | 6.03E-01 |
| ENSMUSG00000027371 | GSMG0023491 | Fahd2a | fumarylacetoacetate hydrolase domain containing 2A | 808.33 | 762.80 | 0.08 | 7.55E-01 |

| | | | | | | | |
|---|---|---|---|---|---|---|---|
| ENSMUSG00000079056 | GSMG0023492 | Kcnip3 | Kv channel interacting protein 3, calsenilin | 203.87 | 181.25 | 0.17 | 6.65E-01 |
| ENSMUSG00000027375 | GSMG0023496 | Mal | myelin and lymphocyte protein, T cell differentiation protein | 3626.21 | 741.04 | 2.29 | 2.02E-44 |
| ENSMUSG00000027378 | GSMG0023498 | Nphp1 | nephronophthisis 1 (juvenile) homolog (human) | 749.40 | 702.56 | 0.09 | 7.21E-01 |
| ENSMUSG00000051319 | GSMG0023500 | 1500011K16Rik | RIKEN cDNA 1500011K16 gene | 437.38 | 437.68 | 0.00 | 9.95E-01 |
| ENSMUSG00000027379 | GSMG0023501 | Bub1 | BUB1, mitotic checkpoint serine/threonine kinase | 191.56 | 526.11 | -1.45 | 8.14E-08 |
| ENSMUSG00000074813 | GSMG0023504 | Gm14005 | predicted gene 14005 | 475.62 | 579.56 | -0.29 | 3.03E-01 |
| ENSMUSG00000014355 | GSMG0023509 | Anapc1 | anaphase promoting complex subunit 1 | 390.63 | 502.79 | -0.36 | 1.16E-01 |
| ENSMUSG00000027387 | GSMG0023512 | Zc3h8 | zinc finger CCCH type containing 8 | 177.93 | 274.64 | -0.63 | 6.02E-03 |
| NULL | GSMG0023514 | NULL | NULL | 271.18 | 412.43 | -0.61 | 2.20E-02 |
| NULL | GSMG0023518 | NULL | NULL | 477.96 | 605.10 | -0.34 | 1.24E-01 |
| ENSMUSG00000048327 | GSMG0023520 | Ckap2l | cytoskeleton associated protein 2-like | 202.50 | 550.19 | -1.44 | 2.51E-12 |
| ENSMUSG00000027400 | GSMG0023525 | Pdyn | prodynorphin | 265.98 | 129.00 | 1.05 | 1.52E-04 |
| ENSMUSG00000027404 | GSMG0023530 | Snrpb | small nuclear ribonucleoprotein B | 4583.05 | 5365.88 | -0.23 | 1.95E-01 |
| ENSMUSG00000027406 | GSMG0023533 | Idh3b | isocitrate dehydrogenase 3 (NAD+) beta | 3207.69 | 2338.53 | 0.46 | 9.75E-03 |
| ENSMUSG00000060029 | GSMG0023538 | 4930473A02Rik | RIKEN cDNA 4930473A02 gene | 112.49 | 122.32 | -0.12 | 7.94E-01 |
| G00000027300,ENSMUSG000 | GSMG0023540 | Ubox5 // Fastkd5 | U box domain containing 5 // FAST kinase domains 5 | 793.74 | 765.49 | 0.05 | 8.59E-01 |
| ENSMUSG00000037703 | GSMG0023542 | Lzts3 | leucine zipper, putative tumor suppressor family member 3 | 1216.12 | 489.37 | 1.31 | 2.12E-09 |
| ENSMUSG00000068290 | GSMG0023543 | Ddrgk1 | DDRGK domain containing 1 | 2461.53 | 2556.96 | -0.05 | 8.60E-01 |
| ENSMUSG00000027309 | GSMG0023546 | 4930402H24Rik | RIKEN cDNA 4930402H24 gene | 653.36 | 826.71 | -0.34 | 9.41E-02 |
| ENSMUSG00000085566 | GSMG0023551 | A730017L22Rik | RIKEN cDNA A730017L22 gene | 1231.61 | 1468.53 | -0.25 | 2.25E-01 |
| ENSMUSG00000027316 | GSMG0023556 | Gfra4 | glial cell line derived neurotrophic factor family receptor alpha 4 | 208.75 | 187.69 | 0.15 | 6.94E-01 |
| ENSMUSG00000027327 | GSMG0023559 | 1700037H04Rik | RIKEN cDNA 1700037H04 gene | 106.35 | 134.74 | -0.35 | 3.49E-01 |
| ENSMUSG00000027329 | GSMG0023560 | Spef1 | sperm flagellar 1 | 119.64 | 115.31 | 0.04 | 9.41E-01 |
| NULL | GSMG0023561 | NULL | NULL | 2324.81 | 1687.21 | 0.46 | 1.24E-01 |
| NULL | GSMG0023564 | NULL | NULL | 294.29 | 403.12 | -0.45 | 4.19E-02 |
| ENSMUSG00000027339 | GSMG0023572 | Rassf2 | Ras association (RalGDS/AF-6) domain family member 2 | 296.48 | 566.99 | -0.94 | 4.67E-06 |
| ENSMUSG00000027340 | GSMG0023574 | Slc23a2 | solute carrier family 23 (nucleobase transporters), member 2 | 593.71 | 667.86 | -0.17 | 5.30E-01 |
| ENSMUSG00000027341 | GSMG0023576 | Tmem230 | transmembrane protein 230 | 522.62 | 659.30 | -0.33 | 1.11E-01 |
| NULL | GSMG0023578 | NULL | NULL | 62.47 | 89.59 | -0.52 | 1.94E-01 |
| NULL | GSMG0023582 | NULL | NULL | 919.55 | 511.87 | 0.85 | 2.39E-03 |
| ENSMUSG00000027346 | GSMG0023583 | Gpcpd1 | glycerophosphocholine phosphodiesterase 1 | 11672.52 | 9064.82 | 0.36 | 4.97E-02 |
| NULL | GSMG0023584 | NULL | NULL | 1097.54 | 543.25 | 1.01 | 7.80E-07 |
| ENSMUSG00000037376 | GSMG0023585 | Trmt6 | tRNA methyltransferase 6 | 332.47 | 395.37 | -0.25 | 3.89E-01 |

| | | | | | | | |
|---|---|---|---|---|---|---|---|
| ENSMUSG00000034723 | GSMG0023591 | Tmx4 | thioredoxin-related transmembrane protein 4 | 3200.81 | 3409.94 | -0.09 | 6.74E-01 |
| ENSMUSG00000039913 | GSMG0023595 | Pak7 | p21 protein (Cdc42/Rac)-activated kinase 7 | 148.26 | 245.79 | -0.73 | 6.01E-03 |
| NULL | GSMG0023599 | NULL | NULL | 83.82 | 110.85 | -0.41 | 3.04E-01 |
| ENSMUSG00000027274 | GSMG0023601 | Mkks | McKusick-Kaufman syndrome | 193.05 | 195.90 | -0.02 | 9.68E-01 |
| ENSMUSG00000027276 | GSMG0023603 | Jag1 | jagged 1 | 838.80 | 769.34 | 0.12 | 7.07E-01 |
| ENSMUSG00000039033 | GSMG0023607 | Tasp1 | taspase, threonine aspartase 1 | 264.51 | 227.32 | 0.22 | 4.88E-01 |
| ENSMUSG00000045624 | GSMG0023609 | Esf1 | ESF1 nucleolar pre-rRNA processing protein homolog | 855.18 | 1039.62 | -0.28 | 3.23E-01 |
| ENSMUSG00000051379 | GSMG0023612 | Flrt3 | fibronectin leucine rich transmembrane protein 3 | 636.87 | 845.52 | -0.41 | 2.71E-02 |
| ENSMUSG00000038844 | GSMG0023614 | Kif16b | kinesin family member 16B | 353.11 | 307.98 | 0.20 | 5.50E-01 |
| ENSMUSG00000027422 | GSMG0023623 | Rrbp1 | ribosome binding protein 1 | 706.70 | 897.39 | -0.35 | 1.90E-01 |
| ENSMUSG00000027423 | GSMG0023626 | Snx5 | sorting nexin 5 | 846.38 | 951.01 | -0.17 | 4.91E-01 |
| ENSMUSG00000037259 | GSMG0023629 | Dzank1 | double zinc ribbon and ankyrin repeat domains 1 | 319.45 | 177.54 | 0.85 | 1.88E-04 |
| ENSMUSG00000027428 | GSMG0023630 | Rbbp9 | retinoblastoma binding protein 9 | 1379.32 | 1249.77 | 0.14 | 5.95E-01 |
| NULL | GSMG0023636 | NULL | NULL | 60.51 | 69.07 | -0.18 | 7.40E-01 |
| ENSMUSG00000001767 | GSMG0023637 | Crnkl1 | Crn, crooked neck-like 1 (Drosophila) | 975.68 | 1086.91 | -0.16 | 5.28E-01 |
| ENSMUSG00000074743 | GSMG0023658 | Thbd | thrombomodulin | 3219.70 | 2149.70 | 0.58 | 2.63E-05 |
| ENSMUSG00000027435 | GSMG0023660 | Cd93 | CD93 antigen | 3855.06 | 5463.56 | -0.50 | 6.56E-02 |
| ENSMUSG00000027438 | GSMG0023661 | Napb | N-ethylmaleimide sensitive fusion protein attachment protein beta | 320.39 | 233.79 | 0.45 | 6.34E-02 |
| ENSMUSG00000027447 | GSMG0023664 | Cst3 | cystatin C | 37250.53 | 23892.48 | 0.64 | 2.06E-06 |
| ENSMUSG00000033096 | GSMG0023678 | Apmap | adipocyte plasma membrane associated protein | 860.46 | 867.39 | -0.01 | 9.74E-01 |
| ENSMUSG00000027452 | GSMG0023679 | Acss1 | acyl-CoA synthetase short-chain family member 1 | 285.81 | 326.86 | -0.19 | 6.24E-01 |
| ENSMUSG00000032046 | GSMG0023683 | Abhd12 | abhydrolase domain containing 12 | 1151.61 | 950.26 | 0.28 | 1.96E-01 |
| ENSMUSG00000027457 | GSMG0023693 | Snph | syntaphilin | 2120.42 | 944.53 | 1.17 | 9.08E-11 |
| ENSMUSG00000032869 | GSMG0023695 | Psmf1 | proteasome (prosome, macropain) inhibitor subunit 1 | 404.91 | 456.01 | -0.17 | 5.54E-01 |
| ENSMUSG00000027459 | GSMG0023698 | Fam110a | family with sequence similarity 110, member A | 287.27 | 371.56 | -0.37 | 1.11E-01 |
| ENSMUSG00000027466 | GSMG0023701 | Rbck1 | RanBP-type and C3HC4-type zinc finger containing 1 | 3018.24 | 2938.77 | 0.04 | 8.92E-01 |
| ENSMUSG00000059361 | GSMG0023703 | Nrsn2 | neurensin 2 | 239.22 | 169.51 | 0.50 | 1.31E-01 |
| NULL | GSMG0023704 | NULL | NULL | 716.73 | 1068.86 | -0.58 | 6.45E-02 |
| ENSMUSG00000074682 | GSMG0023705 | Zcchc3 | zinc finger, CCHC domain containing 3 | 585.49 | 733.91 | -0.33 | 1.50E-01 |
| NULL | GSMG0023714 | NULL | NULL | 164.18 | 200.35 | -0.28 | 4.19E-01 |
| ENSMUSG00000007659 | GSMG0023716 | Bcl2l1 | BCL2-like 1 | 1508.12 | 1183.59 | 0.35 | 9.46E-02 |
| NULL | GSMG0023717 | NULL | NULL | 445.34 | 123.70 | 1.85 | 1.20E-13 |
| NULL | GSMG0023719 | NULL | NULL | 3309.73 | 1608.20 | 1.04 | 6.39E-12 |

| | | | | | | | |
|---|---|---|---|---|---|---|---|
| ENSMUSG00000074676 | GSMG0023720 | Foxs1 | forkhead box S1 | 671.92 | 620.06 | 0.11 | 6.66E-01 |
| ENSMUSG00000042662 | GSMG0023722 | Dusp15 | dual specificity phosphatase-like 15 | 528.37 | 300.94 | 0.81 | 1.75E-03 |
| ENSMUSG00000027472 | GSMG0023723 | Pdrg1 | p53 and DNA damage regulated 1 | 1875.46 | 1895.17 | -0.01 | 9.59E-01 |
| ENSMUSG00000074671 | GSMG0023724 | Tspyl3 | TSPY-like 3 | 552.48 | 700.87 | -0.34 | 8.95E-02 |
| ENSMUSG00000073236 | GSMG0023726 | 2500004C02Rik | RIKEN cDNA 2500004C02 gene | 189.86 | 150.68 | 0.33 | 3.73E-01 |
| ENSMUSG00000061411 | GSMG0023728 | Nol4l | nucleolar protein 4-like | 211.13 | 320.17 | -0.60 | 2.06E-02 |
| ENSMUSG00000056941 | GSMG0023730 | Commd7 | COMM domain containing 7 | 1420.45 | 1873.70 | -0.40 | 1.88E-02 |
| NULL | GSMG0023732 | NULL | NULL | 195.26 | 251.63 | -0.37 | 2.20E-01 |
| ENSMUSG00000027487 | GSMG0023734 | Cdk5rap1 | CDK5 regulatory subunit associated protein 1 | 261.14 | 260.58 | 0.00 | 9.92E-01 |
| ENSMUSG00000027488 | GSMG0023735 | Snta1 | syntrophin, acidic 1 | 270.97 | 221.77 | 0.29 | 3.59E-01 |
| ENSMUSG00000027489 | GSMG0023736 | Necab3 | N-terminal EF-hand calcium binding protein 3 | 866.53 | 375.82 | 1.21 | 3.02E-10 |
| ENSMUSG00000027490 | GSMG0023737 | E2f1 | E2F transcription factor 1 | 117.19 | 225.59 | -0.94 | 6.11E-04 |
| ENSMUSG00000000876 | GSMG0023738 | Pxmp4 | peroxisomal membrane protein 4 | 781.38 | 707.24 | 0.14 | 5.73E-01 |
| ENSMUSG00000074656 | GSMG0023742 | Eif2s2 | eukaryotic translation initiation factor 2, subunit 2 (beta) | 887.56 | 1238.57 | -0.48 | 1.04E-01 |
| ENSMUSG00000038383 | GSMG0023745 | Pigu | phosphatidylinositol glycan anchor biosynthesis, class U | 611.82 | 744.08 | -0.28 | 2.45E-01 |
| NULL | GSMG0023746 | NULL | NULL | 308.76 | 244.45 | 0.33 | 2.12E-01 |
| ENSMUSG00000038369 | GSMG0023747 | Ncoa6 | nuclear receptor coactivator 6 | 573.63 | 689.87 | -0.27 | 3.29E-01 |
| ENSMUSG00000027603 | GSMG0023749 | Ggt7 | gamma-glutamyltransferase 7 | 1920.13 | 2007.61 | -0.06 | 8.47E-01 |
| ENSMUSG00000027610 | GSMG0023751 | Gss | glutathione synthetase | 446.55 | 432.10 | 0.05 | 8.86E-01 |
| ENSMUSG00000038324 | GSMG0023752 | Trpc4ap | transient receptor potential cation channel, subfamily C, member 4 asso | 6095.01 | 7044.96 | -0.21 | 2.15E-01 |
| ENSMUSG00000038312 | GSMG0023753 | Edem2 | ER degradation enhancer, mannosidase alpha-like 2 | 1150.49 | 1153.25 | 0.00 | 9.93E-01 |
| ENSMUSG00000074649 | GSMG0023754 | BC029722 | cDNA sequence BC029722 | 705.10 | 617.20 | 0.19 | 3.93E-01 |
| ENSMUSG00000027613 | GSMG0023755 | Eif6 | eukaryotic translation initiation factor 6 | 1209.81 | 1449.58 | -0.26 | 1.99E-01 |
| ENSMUSG00000005882 | GSMG0023757 | Uqcc1 | ubiquinol-cytochrome c reductase complex assembly factor 1 | 258.60 | 282.24 | -0.12 | 7.37E-01 |
| NULL | GSMG0023765 | NULL | NULL | 352.24 | 194.97 | 0.85 | 2.77E-04 |
| G00000074643,ENSMUSG000 | GSMG0023766 | Cpne1 // Rbm12 | copine I // RNA binding motif protein 12 | 1153.87 | 1122.58 | 0.04 | 8.97E-01 |
| ENSMUSG00000027618 | GSMG0023767 | Nfs1 | nitrogen fixation gene 1 (S. cerevisiae) | 1217.72 | 1155.35 | 0.08 | 7.94E-01 |
| ENSMUSG00000027620 | GSMG0023768 | Rbm39 | RNA binding motif protein 39 | 7219.90 | 6589.35 | 0.13 | 5.79E-01 |
| ENSMUSG00000046229 | GSMG0023771 | Scand1 | SCAN domain-containing 1 | 663.11 | 836.20 | -0.34 | 2.52E-01 |
| NULL | GSMG0023777 | NULL | NULL | 60.23 | 112.06 | -0.91 | 1.15E-02 |
| NULL | GSMG0023780 | NULL | NULL | 177.62 | 188.17 | -0.09 | 8.45E-01 |
| ENSMUSG00000027634 | GSMG0023782 | Ndrg3 | N-myc downstream regulated gene 3 | 1531.11 | 1399.53 | 0.13 | 6.22E-01 |
| ENSMUSG00000027635 | GSMG0023784 | Dsn1 | DSN1 homolog, MIS12 kinetochore complex component | 387.23 | 431.14 | -0.16 | 5.84E-01 |

| | | | | | | | |
|---|---|---|---|---|---|---|---|
| ENSMUSG00000055485 | GSMG0023787 | Soga1 | suppressor of glucose, autophagy associated 1 | 143.23 | 193.95 | -0.44 | 3.84E-01 |
| ENSMUSG00000027639 | GSMG0023788 | Samhd1 | SAM domain and HD domain, 1 | 651.04 | 455.83 | 0.51 | 2.36E-02 |
| ENSMUSG00000027641 | GSMG0023790 | Rbl1 | retinoblastoma-like 1 (p107) | 72.31 | 131.20 | -0.86 | 9.68E-03 |
| ENSMUSG00000067787 | GSMG0023794 | Blcap | bladder cancer associated protein | 1762.29 | 2179.32 | -0.31 | 1.39E-01 |
| ENSMUSG00000027650 | GSMG0023798 | Tti1 | TELO2 interacting protein 1 | 232.43 | 234.59 | -0.01 | 9.87E-01 |
| ENSMUSG00000037820 | GSMG0023800 | Tgm2 | transglutaminase 2, C polypeptide | 3555.71 | 2015.97 | 0.82 | 5.68E-06 |
| ENSMUSG00000085385 | GSMG0023802 | Snhg17 | small nucleolar RNA host gene 17 | 334.96 | 550.53 | -0.72 | 1.45E-04 |
| ENSMUSG00000074622 | GSMG0023812 | Mafb | v-maf musculoaponeurotic fibrosarcoma oncogene family, protein B (av | 489.91 | 807.17 | -0.72 | 6.45E-04 |
| ENSMUSG00000035877 | GSMG0023813 | Zhx3 | zinc fingers and homeoboxes 3 | 1303.28 | 1024.92 | 0.35 | 9.95E-02 |
| NULL | GSMG0023814 | NULL | NULL | 294.43 | 206.08 | 0.52 | 2.93E-02 |
| ENSMUSG00000057133 | GSMG0023823 | Chd6 | chromodomain helicase DNA binding protein 6 | 1344.74 | 1243.40 | 0.11 | 6.40E-01 |
| ENSMUSG00000053141 | GSMG0023826 | Ptprt | protein tyrosine phosphatase, receptor type, T | 552.50 | 566.74 | -0.04 | 9.40E-01 |
| ENSMUSG00000048486 | GSMG0023838 | Fitm2 | fat storage-inducing transmembrane protein 2 | 309.53 | 340.08 | -0.13 | 6.88E-01 |
| NULL | GSMG0023840 | NULL | NULL | 665.58 | 491.11 | 0.44 | 2.33E-02 |
| ENSMUSG00000017707 | GSMG0023841 | Serinc3 | serine incorporator 3 | 24597.83 | 15195.34 | 0.69 | 9.89E-05 |
| NULL | GSMG0023845 | NULL | NULL | 373.00 | 378.04 | -0.02 | 9.66E-01 |
| ENSMUSG00000035226 | GSMG0023846 | Rims4 | regulating synaptic membrane exocytosis 4 | 50.43 | 79.01 | -0.64 | 2.30E-01 |
| ENSMUSG00000018322 | GSMG0023847 | Tomm34 | translocase of outer mitochondrial membrane 34 | 424.82 | 605.69 | -0.51 | 3.11E-02 |
| ENSMUSG00000016995 | GSMG0023857 | Matn4 | matrilin 4 | 341.76 | 267.62 | 0.35 | 3.55E-01 |
| ENSMUSG00000017009 | GSMG0023858 | Sdc4 | syndecan 4 | 530.94 | 268.56 | 0.98 | 1.42E-05 |
| NULL | GSMG0023873 | NULL | NULL | 95.26 | 128.37 | -0.42 | 2.33E-01 |
| ENSMUSG00000017307 | GSMG0023874 | Acot8 | acyl-CoA thioesterase 8 | 229.32 | 209.33 | 0.13 | 7.20E-01 |
| ENSMUSG00000039873 | GSMG0023877 | Neurl2 | neuralized E3 ubiquitin protein ligase 2 | 137.83 | 130.35 | 0.08 | 8.57E-01 |
| ENSMUSG00000017754 | GSMG0023878 | Pltp | phospholipid transfer protein | 44228.66 | 23803.56 | 0.89 | 5.20E-11 |
| NULL | GSMG0023880 | NULL | NULL | 991.50 | 821.98 | 0.27 | 2.19E-01 |
| ENSMUSG00000039834 | GSMG0023881 | Zfp335 | zinc finger protein 335 | 2600.83 | 2437.73 | 0.09 | 7.75E-01 |
| ENSMUSG00000039804 | GSMG0023882 | Ncoa5 | nuclear receptor coactivator 5 | 628.87 | 805.70 | -0.36 | 6.73E-02 |
| ENSMUSG00000053166 | GSMG0023885 | Cdh22 | cadherin 22 | 238.92 | 98.17 | 1.28 | 3.98E-06 |
| ENSMUSG00000017664 | GSMG0023886 | Slc35c2 | solute carrier family 35, member C2 | 626.58 | 717.23 | -0.19 | 4.05E-01 |
| ENSMUSG00000017670 | GSMG0023887 | Elmo2 | engulfment and cell motility 2 | 1227.73 | 1012.73 | 0.28 | 1.59E-01 |
| ENSMUSG00000017667 | GSMG0023889 | Zfp334 | zinc finger protein 334 | 545.07 | 827.46 | -0.60 | 8.06E-04 |
| ENSMUSG00000018459 | GSMG0023891 | Slc13a3 | solute carrier family 13 (sodium-dependent dicarboxylate transporter), r | 444.56 | 597.92 | -0.43 | 8.07E-02 |
| ENSMUSG00000039725 | GSMG0023892 | Trp53rka | transformation related protein 53 regulating kinase A | 433.60 | 479.34 | -0.14 | 6.06E-01 |

| | | | | | | | |
|---|---|---|---|---|---|---|---|
| ENSMUSG00000039671 | GSMG0023894 | Zmynd8 | zinc finger, MYND-type containing 8 | 1246.39 | 1280.40 | -0.04 | 8.97E-01 |
| NULL | GSMG0023895 | NULL | NULL | 184.30 | 104.05 | 0.83 | 1.81E-03 |
| ENSMUSG00000006800 | GSMG0023897 | Sulf2 | sulfatase 2 | 1176.20 | 1598.55 | -0.44 | 6.90E-02 |
| NULL | GSMG0023898 | NULL | NULL | 285.51 | 191.23 | 0.58 | 5.68E-02 |
| ENSMUSG00000039621 | GSMG0023901 | Prex1 | phosphatidylinositol-3,4,5-trisphosphate-dependent Rac exchange facto | 1578.10 | 1928.57 | -0.29 | 2.84E-01 |
| ENSMUSG00000039536 | GSMG0023902 | Stau1 | staufen (RNA binding protein) homolog 1 (Drosophila) | 1416.09 | 1578.78 | -0.16 | 4.30E-01 |
| ENSMUSG00000039501 | GSMG0023903 | Znfx1 | zinc finger, NFX1-type containing 1 | 2069.37 | 1528.96 | 0.44 | 3.93E-02 |
| ENSMUSG00000050556 | GSMG0023905 | Kcnb1 | potassium voltage gated channel, Shab-related subfamily, member 1 | 1343.95 | 883.57 | 0.60 | 7.63E-03 |
| NULL | GSMG0023906 | NULL | NULL | 295.30 | 103.28 | 1.52 | 8.37E-11 |
| ENSMUSG00000017969 | GSMG0023907 | Ptgis | prostaglandin I2 (prostacyclin) synthase | 1193.57 | 1075.31 | 0.15 | 6.11E-01 |
| ENSMUSG00000017929 | GSMG0023908 | B4galt5 | UDP-Gal:betaGlcNAc beta 1,4-galactosyltransferase, polypeptide 5 | 222.67 | 199.21 | 0.16 | 6.70E-01 |
| ENSMUSG00000047030 | GSMG0023911 | Spata2 | spermatogenesis associated 2 | 444.97 | 492.12 | -0.14 | 6.56E-01 |
| ENSMUSG00000090213 | GSMG0023913 | Tmem189 | transmembrane protein 189 | 682.08 | 585.29 | 0.22 | 3.54E-01 |
| 49,ENSMUSG00000078919,EI | GSMG0023924 | Dpm1-adnp // Adn | Dpm1-adnp readthrough // activity-dependent neuroprotective protein / | 748.25 | 748.33 | 0.00 | 9.96E-01 |
| NULL | GSMG0023925 | NULL | NULL | 75.20 | 117.34 | -0.66 | 1.02E-01 |
| ENSMUSG00000074575 | GSMG0023927 | Kcng1 | potassium voltage-gated channel, subfamily G, member 1 | 349.03 | 451.36 | -0.37 | 8.56E-02 |
| ENSMUSG00000027546 | GSMG0023930 | Atp9a | ATPase, class II, type 9A | 5344.43 | 6302.86 | -0.24 | 5.19E-01 |
| ENSMUSG00000027551 | GSMG0023933 | Zfp64 | zinc finger protein 64 | 227.04 | 328.42 | -0.53 | 2.09E-02 |
| ENSMUSG00000013523 | GSMG0023950 | Bcas1 | breast carcinoma amplified sequence 1 | 302.92 | 102.56 | 1.57 | 4.97E-11 |
| ENSMUSG00000027496 | GSMG0023958 | Aurka | aurora kinase A | 190.98 | 440.08 | -1.20 | 1.18E-08 |
| NULL | GSMG0023961 | NULL | NULL | 159.94 | 122.56 | 0.39 | 2.67E-01 |
| ENSMUSG00000008999 | GSMG0023963 | Bmp7 | bone morphogenetic protein 7 | 268.02 | 282.30 | -0.07 | 8.76E-01 |
| ENSMUSG00000038400 | GSMG0023967 | Pmepa1 | prostate transmembrane protein, androgen induced 1 | 2700.53 | 2504.64 | 0.11 | 7.80E-01 |
| NULL | GSMG0023968 | NULL | NULL | 575.12 | 634.32 | -0.14 | 7.32E-01 |
| ENSMUSG00000055897 | GSMG0023970 | Ppp4r1l-ps | protein phosphatase 4, regulatory subunit 1-like, pseudogene | 156.42 | 135.07 | 0.21 | 5.49E-01 |
| ENSMUSG00000016256 | GSMG0023977 | Ctsz | cathepsin Z | 1540.13 | 1498.86 | 0.04 | 8.98E-01 |
| ENSMUSG00000016252 | GSMG0023978 | Atp5e | ATP synthase, H+ transporting, mitochondrial F1 complex, epsilon sub | 1741.62 | 2217.33 | -0.35 | 1.80E-01 |
| ENSMUSG00000016257 | GSMG0023979 | Slmo2 | slowmo homolog 2 (Drosophila) | 959.33 | 1210.08 | -0.33 | 1.46E-01 |
| NULL | GSMG0023997 | NULL | NULL | 352.14 | 200.40 | 0.81 | 5.80E-04 |
| ENSMUSG00000039117 | GSMG0024003 | Taf4 | TATA-box binding protein associated factor 4 | 81.38 | 95.76 | -0.23 | 6.33E-01 |
| ENSMUSG00000027566 | GSMG0024004 | Psma7 | proteasome (prosome, macropain) subunit, alpha type 7 | 3588.22 | 4167.91 | -0.22 | 2.59E-01 |
| ENSMUSG00000039059 | GSMG0024005 | Hrh3 | histamine receptor H3 | 373.35 | 136.03 | 1.46 | 3.82E-09 |
| ENSMUSG00000015647 | GSMG0024007 | Lama5 | laminin, alpha 5 | 1062.24 | 426.63 | 1.32 | 4.04E-09 |

| | | | | | | | |
|---|---|---|---|---|---|---|---|
| ENSMUSG00000038990 | GSMG0024008 | Cables2 | CDK5 and Abl enzyme substrate 2 | 681.83 | 623.41 | 0.13 | 6.04E-01 |
| ENSMUSG00000038914 | GSMG0024018 | Dido1 | death inducer-obliterator 1 | 1294.16 | 1294.30 | 0.00 | 1.00E+00 |
| ENSMUSG00000038848 | GSMG0024023 | Ythdf1 | YTH domain family 1 | 1155.93 | 1398.68 | -0.27 | 1.89E-01 |
| ENSMUSG00000027574 | GSMG0024024 | Nkain4 | Na+/K+ transporting ATPase interacting 4 | 521.73 | 394.04 | 0.41 | 7.22E-02 |
| ENSMUSG00000027577 | GSMG0024026 | Chrna4 | cholinergic receptor, nicotinic, alpha polypeptide 4 | 143.21 | 179.17 | -0.32 | 3.54E-01 |
| ENSMUSG00000016346 | GSMG0024027 | Kcnq2 | potassium voltage-gated channel, subfamily Q, member 2 | 709.59 | 866.87 | -0.29 | 3.25E-01 |
| ENSMUSG00000016349 | GSMG0024030 | Eef1a2 | eukaryotic translation elongation factor 1 alpha 2 | 597.82 | 329.13 | 0.86 | 2.63E-03 |
| NULL | GSMG0024031 | NULL | NULL | 228.12 | 258.73 | -0.19 | 5.94E-01 |
| ENSMUSG00000027579 | GSMG0024034 | Srms | src-related kinase lacking C-terminal regulatory tyrosine and N-termina | 129.23 | 63.18 | 1.04 | 1.93E-03 |
| ENSMUSG00000027580 | GSMG0024035 | Helz2 | helicase with zinc finger 2, transcriptional coactivator | 149.88 | 105.38 | 0.51 | 1.97E-01 |
| ENSMUSG00000038705 | GSMG0024036 | Gmeb2 | glucocorticoid modulatory element binding protein 2 | 115.06 | 135.74 | -0.24 | 5.79E-01 |
| ENSMUSG00000027581 | GSMG0024039 | Stmn3 | stathmin-like 3 | 3627.64 | 6392.71 | -0.82 | 1.23E-08 |
| ENSMUSG00000038671 | GSMG0024040 | Arfrp1 | ADP-ribosylation factor related protein 1 | 716.71 | 745.19 | -0.06 | 8.51E-01 |
| ENSMUSG00000027583 | GSMG0024042 | Zbtb46 | zinc finger and BTB domain containing 46 | 966.45 | 774.31 | 0.32 | 9.84E-02 |
| ENSMUSG00000089917 | GSMG0024045 | Uckl1 | uridine-cytidine kinase 1-like 1 | 850.65 | 688.14 | 0.31 | 1.31E-01 |
| ENSMUSG00000000823 | GSMG0024046 | Zfp512b | zinc finger protein 512B | 1891.96 | 2308.83 | -0.29 | 3.87E-01 |
| ENSMUSG00000038605 | GSMG0024047 | Samd10 | sterile alpha motif domain containing 10 | 1085.68 | 1364.69 | -0.33 | 7.49E-02 |
| ENSMUSG00000046470 | GSMG0024048 | Sox18 | SRY (sex determining region Y)-box 18 | 4584.49 | 4300.99 | 0.09 | 7.07E-01 |
| ENSMUSG00000002458 | GSMG0024049 | Rgs19 | regulator of G-protein signaling 19 | 665.60 | 905.61 | -0.44 | 1.90E-02 |
| NULL | GSMG0024053 | NULL | NULL | 150.32 | 151.96 | -0.02 | 9.74E-01 |
| ENSMUSG00000027499 | GSMG0024065 | Pkia | protein kinase inhibitor, alpha | 2096.16 | 3865.78 | -0.88 | 1.63E-08 |
| ENSMUSG00000043542 | GSMG0024070 | Zc2hc1a | zinc finger, C2HC-type containing 1A | 334.71 | 468.44 | -0.49 | 3.55E-02 |
| ENSMUSG00000027500 | GSMG0024075 | Stmn2 | stathmin-like 2 | 3324.40 | 10045.33 | -1.60 | 1.97E-35 |
| NULL | GSMG0024079 | NULL | NULL | 330.18 | 300.78 | 0.13 | 6.94E-01 |
| ENSMUSG00000027533 | GSMG0024086 | Fabp5 | fatty acid binding protein 5, epidermal | 1637.50 | 4039.09 | -1.30 | 1.47E-24 |
| ENSMUSG00000039717 | GSMG0024094 | Ralyl | RALY RNA binding protein-like | 347.26 | 542.17 | -0.65 | 2.08E-03 |
| NULL | GSMG0024096 | NULL | NULL | 433.47 | 407.44 | 0.09 | 8.00E-01 |
| NULL | GSMG0024101 | NULL | NULL | 372.01 | 391.71 | -0.08 | 8.66E-01 |
| ENSMUSG00000027550 | GSMG0024107 | Lrrcc1 | leucine rich repeat and coiled-coil domain containing 1 | 746.39 | 850.26 | -0.19 | 4.78E-01 |
| ENSMUSG00000027552 | GSMG0024108 | E2f5 | E2F transcription factor 5 | 357.42 | 332.15 | 0.11 | 7.63E-01 |
| ENSMUSG00000027562 | GSMG0024111 | Car2 | carbonic anhydrase 2 | 3707.62 | 5666.43 | -0.61 | 5.97E-04 |
| ENSMUSG00000047213 | GSMG0024115 | Ythdf3 | YTH domain family 3 | 1448.16 | 1446.44 | 0.00 | 9.95E-01 |
| ENSMUSG00000025128 | GSMG0024124 | Bhlhe22 | basic helix-loop-helix family, member e22 | 375.15 | 687.72 | -0.88 | 1.38E-04 |

| | | | | | | | |
|---|---|---|---|---|---|---|---|
| ENSMUSG00000027601 | GSMG0024127 | Mtfr1 | mitochondrial fission regulator 1 | 358.78 | 365.81 | -0.03 | 9.47E-01 |
| ENSMUSG00000048106 | GSMG0024132 | 4632415L05Rik | RIKEN cDNA 4632415L05 gene | 853.47 | 661.10 | 0.37 | 7.03E-02 |
| ENSMUSG00000003617 | GSMG0024133 | Cp | ceruloplasmin | 1265.96 | 553.57 | 1.19 | 1.20E-08 |
| ENSMUSG00000002428 | GSMG0024135 | Hltf | helicase-like transcription factor | 370.67 | 294.33 | 0.33 | 2.75E-01 |
| ENSMUSG00000027630 | GSMG0024139 | Tbl1xr1 | transducin (beta)-like 1X-linked receptor 1 | 439.22 | 400.29 | 0.14 | 6.21E-01 |
| ENSMUSG00000027698 | GSMG0024146 | Nceh1 | neutral cholesterol ester hydrolase 1 | 1797.40 | 1319.88 | 0.45 | 4.84E-03 |
| NULL | GSMG0024147 | NULL | NULL | 381.78 | 208.80 | 0.87 | 1.80E-05 |
| ENSMUSG00000039304 | GSMG0024148 | Tnfsf10 | tumor necrosis factor (ligand) superfamily, member 10 | 250.89 | 230.05 | 0.13 | 7.29E-01 |
| ENSMUSG00000027695 | GSMG0024151 | Pld1 | phospholipase D1 | 226.36 | 150.69 | 0.59 | 5.62E-02 |
| NULL | GSMG0024152 | NULL | NULL | 242.53 | 141.17 | 0.78 | 9.70E-03 |
| ENSMUSG00000027692 | GSMG0024155 | Tnik | TRAF2 and NCK interacting kinase | 2249.38 | 2088.79 | 0.11 | 7.33E-01 |
| ENSMUSG00000039221 | GSMG0024162 | Rpl22l1 | ribosomal protein L22 like 1 | 373.40 | 437.13 | -0.23 | 5.53E-01 |
| ENSMUSG00000063600 | GSMG0024166 | Egfem1 | EGF-like and EMI domain containing 1 | 386.76 | 383.47 | 0.02 | 9.70E-01 |
| NULL | GSMG0024170 | NULL | NULL | 180.90 | 236.96 | -0.39 | 3.44E-01 |
| NULL | GSMG0024171 | NULL | NULL | 331.25 | 441.69 | -0.41 | 1.02E-01 |
| ENSMUSG00000037730 | GSMG0024175 | Mynn | myoneurin | 670.90 | 597.07 | 0.17 | 4.78E-01 |
| ENSMUSG00000027706 | GSMG0024179 | Sec62 | SEC62 homolog (S. cerevisiae) | 1796.13 | 1531.40 | 0.23 | 7.79E-01 |
| ENSMUSG00000037661 | GSMG0024181 | Gpr160 | G protein-coupled receptor 160 | 245.66 | 77.89 | 1.66 | 7.49E-09 |
| ENSMUSG00000037643 | GSMG0024182 | Prkci | protein kinase C, iota | 711.79 | 785.58 | -0.14 | 5.80E-01 |
| ENSMUSG00000027660 | GSMG0024184 | Skil | SKI-like | 2264.02 | 2571.90 | -0.18 | 3.87E-01 |
| ENSMUSG00000037625 | GSMG0024186 | Cldn11 | claudin 11 | 299.53 | 133.95 | 1.16 | 4.01E-05 |
| ENSMUSG00000027665 | GSMG0024193 | Pik3ca | phosphatidylinositol 3-kinase, catalytic, alpha polypeptide | 1542.84 | 948.17 | 0.70 | 8.64E-03 |
| ENSMUSG00000027667 | GSMG0024195 | Zfp639 | zinc finger protein 639 | 365.43 | 393.01 | -0.11 | 7.64E-01 |
| ENSMUSG00000027668 | GSMG0024196 | Mfn1 | mitofusin 1 | 944.02 | 956.25 | -0.02 | 9.49E-01 |
| NULL | GSMG0024197 | NULL | NULL | 263.85 | 257.45 | 0.04 | 9.28E-01 |
| ENSMUSG00000027671 | GSMG0024198 | Actl6a | actin-like 6A | 1506.00 | 1924.01 | -0.35 | 6.53E-02 |
| ENSMUSG00000027673 | GSMG0024199 | Ndufb5 | NADH dehydrogenase (ubiquinone) 1 beta subcomplex, 5 | 1778.24 | 2147.35 | -0.27 | 1.74E-01 |
| ENSMUSG00000056900 | GSMG0024200 | Usp13 | ubiquitin specific peptidase 13 (isopeptidase T-3) | 292.14 | 334.44 | -0.19 | 4.71E-01 |
| ENSMUSG00000087625 | GSMG0024202 | 4930419G24Rik | RIKEN cDNA 4930419G24 gene | 110.64 | 59.90 | 0.88 | 7.22E-03 |
| ENSMUSG00000027677 | GSMG0024203 | Ttc14 | tetratricopeptide repeat domain 14 | 2923.19 | 2275.47 | 0.36 | 2.93E-02 |
| ENSMUSG00000027680 | GSMG0024205 | Fxr1 | fragile X mental retardation gene 1, autosomal homolog | 502.05 | 587.21 | -0.22 | 4.52E-01 |
| NULL | GSMG0024209 | NULL | NULL | 196.04 | 249.17 | -0.35 | 2.52E-01 |
| ENSMUSG00000037400 | GSMG0024213 | Atp11b | ATPase, class VI, type 11B | 2200.10 | 1759.60 | 0.32 | 1.04E-01 |

| | | | | | | | |
|---|---|---|---|---|---|---|---|
| ENSMUSG00000027710 | GSMG0024216 | Acad9 | acyl-Coenzyme A dehydrogenase family, member 9 | 1708.31 | 1846.35 | -0.11 | 6.23E-01 |
| ENSMUSG00000033883 | GSMG0024218 | D3Ertd254e | DNA segment, Chr 3, ERATO Doi 254, expressed | 891.34 | 960.21 | -0.11 | 6.68E-01 |
| ENSMUSG00000027714 | GSMG0024222 | Exosc9 | exosome component 9 | 616.17 | 771.83 | -0.32 | 1.01E-01 |
| ENSMUSG00000027722 | GSMG0024232 | Spata5 | spermatogenesis associated 5 | 353.20 | 505.34 | -0.51 | 4.64E-02 |
| ENSMUSG00000037211 | GSMG0024238 | Spry1 | sprouty homolog 1 (Drosophila) | 1436.05 | 777.51 | 0.89 | 1.29E-03 |
| ENSMUSG00000046743 | GSMG0024244 | Fat4 | FAT atypical cadherin 4 | 168.65 | 215.34 | -0.35 | 2.92E-01 |
| ENSMUSG00000025757 | GSMG0024248 | Hspa4l | heat shock protein 4 like | 473.60 | 441.31 | 0.10 | 7.25E-01 |
| ENSMUSG00000025758 | GSMG0024250 | Plk4 | polo-like kinase 4 | 245.00 | 399.50 | -0.71 | 6.33E-04 |
| ENSMUSG00000037818 | GSMG0024252 | Abhd18 | abhydrolase domain containing 18 | 277.00 | 215.39 | 0.36 | 2.21E-01 |
| ENSMUSG00000025762 | GSMG0024254 | Larp1b | La ribonucleoprotein domain family, member 1B | 260.01 | 178.80 | 0.54 | 6.33E-02 |
| ENSMUSG00000025764 | GSMG0024259 | Jade1 | jade family PHD finger 1 | 1000.90 | 870.20 | 0.20 | 3.59E-01 |
| ENSMUSG00000023087 | GSMG0024273 | Noct | nocturnin | 729.13 | 770.74 | -0.08 | 8.02E-01 |
| ENSMUSG00000063273 | GSMG0024277 | Naa15 | N(alpha)-acetyltransferase 15, NatA auxiliary subunit | 598.64 | 957.76 | -0.68 | 3.09E-04 |
| ENSMUSG00000027739 | GSMG0024279 | Rab33b | RAB33B, member RAS oncogene family | 1092.47 | 1180.37 | -0.11 | 6.74E-01 |
| ENSMUSG00000097885 | GSMG0024281 | 5031434O11Rik | RIKEN cDNA 5031434O11 gene | 159.97 | 89.02 | 0.84 | 2.35E-02 |
| ENSMUSG00000044167 | GSMG0024290 | Foxo1 | forkhead box O1 | 2780.74 | 2555.52 | 0.12 | 5.68E-01 |
| ENSMUSG00000048332 | GSMG0024294 | Lhfp | lipoma HMGIC fusion partner | 2249.95 | 1908.02 | 0.24 | 2.48E-01 |
| NULL | GSMG0024295 | NULL | NULL | 452.80 | 632.38 | -0.48 | 3.03E-02 |
| NULL | GSMG0024297 | NULL | NULL | 308.85 | 392.43 | -0.34 | 2.51E-01 |
| ENSMUSG00000049504 | GSMG0024301 | Proser1 | proline and serine rich 1 | 692.53 | 868.32 | -0.33 | 7.33E-02 |
| ENSMUSG00000027748 | GSMG0024307 | Trpc4 | transient receptor potential cation channel, subfamily C, member 4 | 329.96 | 555.08 | -0.75 | 3.66E-05 |
| ENSMUSG00000027750 | GSMG0024308 | Postn | periostin, osteoblast specific factor | 612.17 | 1314.81 | -1.10 | 2.77E-08 |
| ENSMUSG00000027751 | GSMG0024309 | Supt20 | suppressor of Ty 20 | 670.80 | 726.50 | -0.12 | 6.74E-01 |
| ENSMUSG00000036632 | GSMG0024310 | Alg5 | asparagine-linked glycosylation 5 (dolichyl-phosphate beta-glucosyltra | 863.12 | 1003.26 | -0.22 | 3.36E-01 |
| ENSMUSG00000036580 | GSMG0024313 | Spg20 | spastic paraplegia 20, spartin (Troyer syndrome) homolog (human) | 734.12 | 679.81 | 0.11 | 6.93E-01 |
| ENSMUSG00000027797 | GSMG0024316 | Dclk1 | doublecortin-like kinase 1 | 1408.94 | 1805.36 | -0.36 | 7.03E-02 |
| NULL | GSMG0024317 | NULL | NULL | 446.84 | 475.01 | -0.09 | 8.19E-01 |
| NULL | GSMG0024318 | NULL | NULL | 612.55 | 534.35 | 0.20 | 4.30E-01 |
| NULL | GSMG0024319 | NULL | NULL | 226.74 | 209.35 | 0.12 | 7.60E-01 |
| NULL | GSMG0024320 | NULL | NULL | 454.50 | 431.99 | 0.08 | 8.33E-01 |
| NULL | GSMG0024324 | NULL | NULL | 317.49 | 158.30 | 1.01 | 1.60E-05 |
| NULL | GSMG0024325 | NULL | NULL | 261.64 | 172.64 | 0.60 | 3.48E-02 |
| ENSMUSG00000036503 | GSMG0024332 | Rnf13 | ring finger protein 13 | 1501.51 | 1314.14 | 0.19 | 4.29E-01 |

| | | | | | | | |
|---|---|---|---|---|---|---|---|
| ENSMUSG00000027806 | GSMG0024335 | Tsc22d2 | TSC22 domain family, member 2 | 2457.43 | 2140.22 | 0.20 | 4.01E-01 |
| ENSMUSG00000027810 | GSMG0024339 | Eif2a | eukaryotic translation initiation factor 2A | 373.28 | 480.66 | -0.36 | 1.04E-01 |
| ENSMUSG00000075700 | GSMG0024341 | Selt | selenoprotein T | 921.13 | 884.20 | 0.06 | 8.60E-01 |
| ENSMUSG00000027763 | GSMG0024356 | Mbnl1 | muscleblind-like 1 (Drosophila) | 738.99 | 794.12 | -0.10 | 7.30E-01 |
| ENSMUSG00000036885 | GSMG0024366 | Arhgef26 | Rho guanine nucleotide exchange factor (GEF) 26 | 140.23 | 285.80 | -1.03 | 2.29E-04 |
| ENSMUSG00000027823 | GSMG0024371 | Gmps | guanine monophosphate synthetase | 1261.58 | 1721.52 | -0.45 | 2.59E-02 |
| ENSMUSG00000034640 | GSMG0024379 | Tiparp | TCDD-inducible poly(ADP-ribose) polymerase | 976.24 | 844.58 | 0.21 | 3.02E-01 |
| ENSMUSG00000034544 | GSMG0024387 | Rsrc1 | arginine/serine-rich coiled-coil 1 | 419.63 | 479.64 | -0.19 | 5.16E-01 |
| ENSMUSG00000048416 | GSMG0024399 | Mlf1 | myeloid leukemia factor 1 | 53.42 | 106.30 | -0.98 | 2.25E-02 |
| ENSMUSG00000027774 | GSMG0024401 | Gfm1 | G elongation factor, mitochondrial 1 | 625.11 | 711.56 | -0.19 | 4.11E-01 |
| ENSMUSG00000027775 | GSMG0024404 | Mfsd1 | major facilitator superfamily domain containing 1 | 1456.35 | 1446.21 | 0.01 | 9.75E-01 |
| 77,ENSMUSG00000051777,EI | GSMG0024405 | Gm21949 // Iqcj // | predicted gene, 21949 // IQ motif containing J // schwannomin interacti | 1065.74 | 1323.16 | -0.31 | 1.10E-01 |
| NULL | GSMG0024412 | NULL | NULL | 460.16 | 325.81 | 0.50 | 5.55E-02 |
| NULL | GSMG0024413 | NULL | NULL | 501.34 | 343.44 | 0.55 | 5.16E-03 |
| ENSMUSG00000034349 | GSMG0024416 | Smc4 | structural maintenance of chromosomes 4 | 1092.22 | 1339.61 | -0.29 | 2.34E-01 |
| ENSMUSG00000027784 | GSMG0024419 | Ppm1l | protein phosphatase 1 (formerly 2C)-like | 3427.19 | 3627.99 | -0.08 | 7.14E-01 |
| ENSMUSG00000027787 | GSMG0024424 | Nmd3 | NMD3 ribosome export adaptor | 757.87 | 977.27 | -0.36 | 1.04E-01 |
| ENSMUSG00000027834 | GSMG0024430 | Serpini1 | serine (or cysteine) peptidase inhibitor, clade I, member 1 | 711.85 | 680.89 | 0.06 | 8.52E-01 |
| ENSMUSG00000034098 | GSMG0024433 | Fstl5 | follistatin-like 5 | 103.55 | 223.04 | -1.10 | 2.54E-05 |
| ENSMUSG00000027804 | GSMG0024441 | Ppid | peptidylprolyl isomerase D (cyclophilin D) | 793.93 | 865.84 | -0.13 | 7.33E-01 |
| ENSMUSG00000027811 | GSMG0024443 | 4930579G24Rik | RIKEN cDNA 4930579G24 gene | 250.53 | 443.00 | -0.82 | 1.60E-04 |
| ENSMUSG00000028015 | GSMG0024451 | Ctso | cathepsin O | 921.63 | 630.93 | 0.55 | 1.53E-03 |
| ENSMUSG00000033900 | GSMG0024453 | Map9 | microtubule-associated protein 9 | 379.20 | 465.62 | -0.30 | 1.73E-01 |
| ENSMUSG00000027998 | GSMG0024457 | Plrg1 | pleiotropic regulator 1 | 1193.61 | 1290.83 | -0.11 | 6.74E-01 |
| ENSMUSG00000027996 | GSMG0024459 | Sfrp2 | secreted frizzled-related protein 2 | 863.48 | 826.32 | 0.06 | 8.62E-01 |
| ENSMUSG00000056498 | GSMG0024464 | Tmem154 | transmembrane protein 154 | 992.21 | 1988.32 | -1.00 | 5.39E-10 |
| ENSMUSG00000028086 | GSMG0024466 | Fbxw7 | F-box and WD-40 domain protein 7 | 1140.98 | 945.40 | 0.27 | 2.73E-01 |
| ENSMUSG00000028085 | GSMG0024473 | Gatb | glutamyl-tRNA(Gln) amidotransferase, subunit B | 279.40 | 276.92 | 0.02 | 9.67E-01 |
| ENSMUSG00000028082 | GSMG0024479 | Sh3d19 | SH3 domain protein D19 | 205.77 | 212.36 | -0.05 | 9.20E-01 |
| ENSMUSG00000003382 | GSMG0024494 | Etv3 | ets variant 3 | 1239.48 | 1050.42 | 0.24 | 3.49E-01 |
| ENSMUSG00000041977 | GSMG0024495 | Arhgef11 | Rho guanine nucleotide exchange factor (GEF) 11 | 496.29 | 500.24 | -0.01 | 9.80E-01 |
| ENSMUSG00000004897 | GSMG0024501 | Hdgf | hepatoma-derived growth factor | 1418.25 | 1706.65 | -0.27 | 1.72E-01 |
| ENSMUSG00000019710 | GSMG0024502 | Mrpl24 | mitochondrial ribosomal protein L24 | 454.58 | 438.37 | 0.05 | 8.88E-01 |

| | | | | | | | |
|---|---|---|---|---|---|---|---|
| ENSMUSG00000048039 | GSMG0024503 | Isg20l2 | interferon stimulated exonuclease gene 20-like 2 | 384.10 | 481.44 | -0.33 | 1.63E-01 |
| ENSMUSG00000004885 | GSMG0024504 | Crabp2 | cellular retinoic acid binding protein II | 102.42 | 137.56 | -0.42 | 2.37E-01 |
| ENSMUSG00000004891 | GSMG0024505 | Nes | nestin | 9490.88 | 8645.15 | 0.13 | 6.08E-01 |
| ENSMUSG00000028069 | GSMG0024507 | Gpatch4 | G patch domain containing 4 | 406.95 | 589.66 | -0.54 | 7.37E-03 |
| ENSMUSG00000028068 | GSMG0024508 | Iqgap3 | IQ motif containing GTPase activating protein 3 | 443.96 | 1159.93 | -1.38 | 1.37E-13 |
| ENSMUSG00000001419 | GSMG0024509 | Mef2d | myocyte enhancer factor 2D | 1281.26 | 1318.31 | -0.04 | 8.93E-01 |
| NULL | GSMG0024510 | NULL | NULL | 364.08 | 617.49 | -0.76 | 4.65E-05 |
| NULL | GSMG0024511 | NULL | NULL | 2960.66 | 3640.90 | -0.30 | 1.42E-01 |
| NULL | GSMG0024512 | NULL | NULL | 224.99 | 408.35 | -0.86 | 2.05E-03 |
| ENSMUSG00000001416 | GSMG0024513 | Cct3 | chaperonin containing Tcp1, subunit 3 (gamma) | 3214.34 | 4399.29 | -0.45 | 1.84E-02 |
| ENSMUSG00000001418 | GSMG0024514 | Glmp | glycosylated lysosomal membrane protein | 2807.01 | 2481.83 | 0.18 | 3.65E-01 |
| ENSMUSG00000001415 | GSMG0024515 | Smg5 | Smg-5 homolog, nonsense mediated mRNA decay factor (C. elegans) | 2848.52 | 2820.69 | 0.01 | 9.70E-01 |
| ENSMUSG00000008604 | GSMG0024520 | Ubqln4 | ubiquilin 4 | 405.58 | 479.03 | -0.24 | 3.32E-01 |
| ENSMUSG00000041355 | GSMG0024521 | Ssr2 | signal sequence receptor, beta | 4745.61 | 5451.32 | -0.20 | 3.35E-01 |
| ENSMUSG00000028059 | GSMG0024522 | Arhgef2 | rho/rac guanine nucleotide exchange factor (GEF) 2 | 7074.57 | 4702.45 | 0.59 | 1.73E-04 |
| ENSMUSG00000028060 | GSMG0024523 | 2810403A07Rik | RIKEN cDNA 2810403A07 gene | 1449.77 | 1828.30 | -0.33 | 5.63E-02 |
| ENSMUSG00000028057 | GSMG0024524 | Rit1 | Ras-like without CAAX 1 | 1847.19 | 1400.05 | 0.40 | 1.32E-02 |
| G00000028049,ENSMUSG000 | GSMG0024533 | Clk2-scamp3 // Clk | Clk2-scamp3 readthrough // CDC-like kinase 2 // secretory carrier mem | 3091.38 | 3134.56 | -0.02 | 9.43E-01 |
| ENSMUSG00000032657 | GSMG0024534 | Fam189b | family with sequence similarity 189, member B | 706.64 | 721.36 | -0.03 | 9.41E-01 |
| ENSMUSG00000028048 | GSMG0024535 | Gba | glucosidase, beta, acid | 1750.59 | 1614.25 | 0.12 | 6.32E-01 |
| NULL | GSMG0024536 | NULL | NULL | 524.94 | 531.75 | -0.02 | 9.62E-01 |
| ENSMUSG00000028047 | GSMG0024537 | Thbs3 | thrombospondin 3 | 541.85 | 1155.85 | -1.09 | 1.13E-06 |
| ENSMUSG00000042747 | GSMG0024539 | Krtcap2 | keratinocyte associated protein 2 | 2259.71 | 2572.34 | -0.19 | 4.33E-01 |
| ENSMUSG00000042737 | GSMG0024540 | Dpm3 | dolichyl-phosphate mannosyltransferase polypeptide 3 | 469.64 | 476.75 | -0.02 | 9.54E-01 |
| NULL | GSMG0024542 | NULL | NULL | 544.61 | 117.16 | 2.22 | 2.13E-27 |
| NULL | GSMG0024543 | NULL | NULL | 811.42 | 446.86 | 0.86 | 1.15E-06 |
| ENSMUSG00000042626 | GSMG0024546 | Shc1 | src homology 2 domain-containing transforming protein C1 | 1282.87 | 1217.04 | 0.08 | 7.62E-01 |
| ENSMUSG00000047824 | GSMG0024547 | Pygo2 | pygopus 2 | 1901.55 | 1874.76 | 0.02 | 9.42E-01 |
| ENSMUSG00000042613 | GSMG0024548 | Pbxip1 | pre B cell leukemia transcription factor interacting protein 1 | 2111.33 | 1419.26 | 0.57 | 2.30E-04 |
| ENSMUSG00000027952 | GSMG0024550 | Pmvk | phosphomevalonate kinase | 1030.72 | 1166.68 | -0.18 | 4.22E-01 |
| ENSMUSG00000027951 | GSMG0024553 | Adar | adenosine deaminase, RNA-specific | 1103.54 | 1166.72 | -0.08 | 7.87E-01 |
| ENSMUSG00000042572 | GSMG0024555 | Ube2q1 | ubiquitin-conjugating enzyme E2Q family member 1 | 662.72 | 427.29 | 0.63 | 2.84E-02 |
| ENSMUSG00000046280 | GSMG0024556 | She | src homology 2 domain-containing transforming protein E | 936.63 | 642.22 | 0.54 | 5.56E-03 |

| | | | | | | | |
|---|---|---|---|---|---|---|---|
| ENSMUSG00000027942 | GSMG0024563 | 4933434E20Rik // | RIKEN cDNA 4933434E20 gene // RIKEN cDNA 1700094D03 gene | 942.84 | 974.26 | -0.05 | 8.85E-01 |
| NULL | GSMG0024565 | NULL | NULL | 487.08 | 436.44 | 0.16 | 5.81E-01 |
| ENSMUSG00000027935 | GSMG0024568 | Rab13 | RAB13, member RAS oncogene family | 528.68 | 604.35 | -0.19 | 4.65E-01 |
| ENSMUSG00000027937 | GSMG0024569 | Jtb | jumping translocation breakpoint | 1813.15 | 1903.91 | -0.07 | 7.93E-01 |
| ENSMUSG00000052310 | GSMG0024570 | Slc39a1 | solute carrier family 39 (zinc transporter), member 1 | 238.73 | 319.46 | -0.42 | 1.64E-01 |
| ENSMUSG00000027936 | GSMG0024571 | Crtc2 | CREB regulated transcription coactivator 2 | 726.28 | 734.19 | -0.01 | 9.69E-01 |
| ENSMUSG00000042404 | GSMG0024572 | Dennd4b | DENN/MADD domain containing 4B | 318.89 | 335.02 | -0.07 | 8.77E-01 |
| ENSMUSG00000042390 | GSMG0024576 | Gatad2b | GATA zinc finger domain containing 2B | 164.54 | 200.61 | -0.28 | 4.13E-01 |
| ENSMUSG00000001016 | GSMG0024579 | Ilf2 | interleukin enhancer binding factor 2 | 742.10 | 1013.04 | -0.45 | 3.06E-02 |
| ENSMUSG00000042312 | GSMG0024580 | S100a13 | S100 calcium binding protein A13 | 1090.55 | 881.25 | 0.31 | 1.97E-01 |
| ENSMUSG00000074457 | GSMG0024582 | S100a16 | S100 calcium binding protein A16 | 640.93 | 937.62 | -0.55 | 9.75E-03 |
| ENSMUSG00000001025 | GSMG0024587 | S100a6 | S100 calcium binding protein A6 (calcyclin) | 282.22 | 167.43 | 0.75 | 5.70E-03 |
| ENSMUSG00000056054 | GSMG0024589 | S100a8 | S100 calcium binding protein A8 (calgranulin A) | 12.73 | 56.05 | -2.19 | 6.12E-05 |
| ENSMUSG00000041959 | GSMG0024624 | S100a10 | S100 calcium binding protein A10 (calpactin) | 277.44 | 409.87 | -0.56 | 1.06E-02 |
| ENSMUSG00000028145 | GSMG0024626 | Them4 | thioesterase superfamily member 4 | 270.80 | 403.80 | -0.58 | 6.68E-03 |
| ENSMUSG00000041912 | GSMG0024630 | Tdrkh | tudor and KH domain containing protein | 170.43 | 272.65 | -0.68 | 3.81E-03 |
| ENSMUSG00000028140 | GSMG0024633 | Mrpl9 | mitochondrial ribosomal protein L9 | 300.45 | 347.38 | -0.21 | 4.63E-01 |
| ENSMUSG00000028137 | GSMG0024634 | Celf3 | CUGBP, Elav-like family member 3 | 948.01 | 1017.31 | -0.10 | 7.10E-01 |
| ENSMUSG00000038902 | GSMG0024637 | Pogz | pogo transposable element with ZNF domain | 541.61 | 548.52 | -0.02 | 9.63E-01 |
| ENSMUSG00000005774 | GSMG0024639 | Rfx5 | regulatory factor X, 5 (influences HLA class II expression) | 155.23 | 167.36 | -0.11 | 7.97E-01 |
| ENSMUSG00000038861 | GSMG0024640 | Pi4kb | phosphatidylinositol 4-kinase, catalytic, beta polypeptide | 1089.74 | 1387.91 | -0.35 | 6.15E-02 |
| NULL | GSMG0024643 | NULL | NULL | 975.71 | 1006.88 | -0.05 | 8.68E-01 |
| ENSMUSG00000008958 | GSMG0024644 | Vps72 | vacuolar protein sorting 72 | 878.44 | 1272.76 | -0.53 | 1.85E-03 |
| ENSMUSG00000038777 | GSMG0024647 | Sema6c | sema domain, transmembrane domain (TM), and cytoplasmic domain, ( | 603.29 | 1229.77 | -1.03 | 1.93E-05 |
| ENSMUSG00000046722 | GSMG0024649 | Cdc42se1 | CDC42 small effector 1 | 1135.60 | 1244.86 | -0.13 | 5.79E-01 |
| ENSMUSG00000038712 | GSMG0024651 | Fam63a | family with sequence similarity 63, member A | 6704.94 | 3645.19 | 0.88 | 2.56E-10 |
| ENSMUSG00000015714 | GSMG0024652 | Cers2 | ceramide synthase 2 | 3490.44 | 3537.61 | -0.02 | 9.52E-01 |
| ENSMUSG00000028111 | GSMG0024657 | Ctsk | cathepsin K | 462.18 | 389.42 | 0.25 | 3.64E-01 |
| ENSMUSG00000038642 | GSMG0024658 | Ctss | cathepsin S | 466.94 | 175.15 | 1.42 | 1.45E-10 |
| ENSMUSG00000046519 | GSMG0024660 | Golph3l | golgi phosphoprotein 3-like | 231.25 | 194.40 | 0.25 | 4.33E-01 |
| ENSMUSG00000038619 | GSMG0024661 | Ensa | endosulfine alpha | 1013.98 | 690.67 | 0.55 | 7.08E-03 |
| ENSMUSG00000038612 | GSMG0024663 | Mcl1 | myeloid cell leukemia sequence 1 | 8821.15 | 6446.33 | 0.45 | 4.85E-03 |
| ENSMUSG00000080727 | GSMG0024666 | C920021L13Rik | RIKEN cDNA C920021L13 gene | 136.27 | 119.38 | 0.19 | 6.59E-01 |

| | | | | | | | |
|---|---|---|---|---|---|---|---|
| ENSMUSG00000015749 | GSMG0024669 | Anp32e | acidic (leucine-rich) nuclear phosphoprotein 32 family, member E | 767.07 | 1328.67 | -0.79 | 3.08E-05 |
| ENSMUSG00000038495 | GSMG0024675 | Otud7b | OTU domain containing 7B | 395.39 | 321.60 | 0.30 | 2.40E-01 |
| ENSMUSG00000045934 | GSMG0024678 | Mtmr11 | myotubularin related protein 11 | 516.99 | 346.55 | 0.58 | 6.45E-03 |
| ENSMUSG00000038486 | GSMG0024680 | Sv2a | synaptic vesicle glycoprotein 2 a | 1124.01 | 1241.64 | -0.14 | 6.65E-01 |
| ENSMUSG00000038393 | GSMG0024690 | Txnip | thioredoxin interacting protein | 1075.90 | 1070.91 | 0.01 | 9.85E-01 |
| ENSMUSG00000049097 | GSMG0024692 | Ankrd34a | ankyrin repeat domain 34A | 355.02 | 230.60 | 0.62 | 4.35E-02 |
| ENSMUSG00000038374 | GSMG0024694 | Rbm8a | RNA binding motif protein 8a | 1549.81 | 1902.02 | -0.30 | 1.93E-01 |
| G00000028102,ENSMUSG000 | GSMG0024695 | Pex11b // Itga10 | peroxisomal biogenesis factor 11 beta // integrin, alpha 10 | 466.44 | 517.79 | -0.15 | 5.83E-01 |
| ENSMUSG00000028101 | GSMG0024697 | Pias3 | protein inhibitor of activated STAT 3 | 635.27 | 653.11 | -0.04 | 9.04E-01 |
| ENSMUSG00000028098 | GSMG0024698 | Rnf115 | ring finger protein 115 | 763.08 | 708.50 | 0.11 | 7.02E-01 |
| ENSMUSG00000057123 | GSMG0024700 | Gja5 | gap junction protein, alpha 5 | 729.18 | 226.37 | 1.69 | 6.90E-12 |
| ENSMUSG00000028093 | GSMG0024701 | Acp6 | acid phosphatase 6, lysophosphatidic | 741.57 | 932.68 | -0.33 | 9.54E-02 |
| ENSMUSG00000028088 | GSMG0024702 | Fmo5 | flavin containing monooxygenase 5 | 203.39 | 87.20 | 1.22 | 1.63E-05 |
| ENSMUSG00000038205 | GSMG0024703 | Prkab2 | protein kinase, AMP-activated, beta 2 non-catalytic subunit | 134.37 | 159.09 | -0.24 | 4.87E-01 |
| ENSMUSG00000027879 | GSMG0024706 | Sec22b | SEC22 homolog B, vesicle trafficking protein | 1931.52 | 2009.66 | -0.06 | 8.44E-01 |
| NULL | GSMG0024707 | NULL | NULL | 373.32 | 396.74 | -0.09 | 7.94E-01 |
| ENSMUSG00000027878 | GSMG0024709 | Notch2 | notch 2 | 220.61 | 221.18 | 0.00 | 1.00E+00 |
| ENSMUSG00000027875 | GSMG0024713 | Hmgcs2 | 3-hydroxy-3-methylglutaryl-Coenzyme A synthase 2 | 12085.07 | 12893.58 | -0.09 | 7.13E-01 |
| ENSMUSG00000004233 | GSMG0024716 | Wars2 | tryptophanyl tRNA synthetase 2 (mitochondrial) | 63.78 | 67.81 | -0.08 | 9.05E-01 |
| ENSMUSG00000027868 | GSMG0024719 | Tbx15 | T-box 15 | 103.89 | 79.35 | 0.39 | 3.28E-01 |
| ENSMUSG00000027865 | GSMG0024722 | Gdap2 | ganglioside-induced differentiation-associated-protein 2 | 1108.91 | 1160.22 | -0.06 | 8.09E-01 |
| ENSMUSG00000042035 | GSMG0024730 | Igsf3 | immunoglobulin superfamily, member 3 | 156.51 | 388.76 | -1.31 | 2.15E-09 |
| ENSMUSG00000027861 | GSMG0024736 | Casq2 | calsequestrin 2 | 563.83 | 106.79 | 2.40 | 2.51E-27 |
| ENSMUSG00000027858 | GSMG0024744 | Tspan2 | tetraspanin 2 | 356.63 | 376.00 | -0.07 | 8.25E-01 |
| ENSMUSG00000027854 | GSMG0024747 | Sike1 | suppressor of IKBKE 1 | 1582.57 | 1766.40 | -0.16 | 5.06E-01 |
| ENSMUSG00000068823 | GSMG0024749 | Csde1 | cold shock domain containing E1, RNA binding | 5813.15 | 6521.51 | -0.17 | 4.35E-01 |
| ENSMUSG00000027852 | GSMG0024750 | Nras | neuroblastoma ras oncogene | 1346.34 | 1505.24 | -0.16 | 4.75E-01 |
| ENSMUSG00000007379 | GSMG0024754 | Dennd2c | DENN/MADD domain containing 2C | 231.94 | 377.34 | -0.70 | 1.29E-03 |
| ENSMUSG00000005687 | GSMG0024755 | Bcas2 | breast carcinoma amplified sequence 2 | 1882.33 | 2467.27 | -0.39 | 8.11E-02 |
| ENSMUSG00000033014 | GSMG0024756 | Trim33 | tripartite motif-containing 33 | 99.52 | 120.92 | -0.29 | 4.83E-01 |
| ENSMUSG00000027849 | GSMG0024759 | Syt6 | synaptotagmin VI | 90.34 | 116.12 | -0.37 | 3.47E-01 |
| G00000032952,ENSMUSG000 | GSMG0024763 | Ap4b1 // Bcl2l15 | adaptor-related protein complex AP-4, beta 1 // BCLl2-like 15 | 528.72 | 534.70 | -0.01 | 9.65E-01 |
| ENSMUSG00000044098 | GSMG0024766 | Rsbn1 | rosbin, round spermatid basic protein 1 | 213.68 | 299.20 | -0.49 | 1.16E-01 |

| | | | | | | | |
|---|---|---|---|---|---|---|---|
| ENSMUSG00000058388 | GSMG0024769 | Phtf1 | putative homeodomain transcription factor 1 | 1332.60 | 1374.05 | -0.04 | 8.91E-01 |
| ENSMUSG00000032902 | GSMG0024772 | Slc16a1 | solute carrier family 16 (monocarboxylic acid transporters), member 1 | 135016.61 | 79908.91 | 0.76 | 5.88E-06 |
| ENSMUSG00000002228 | GSMG0024775 | Ppm1j | protein phosphatase 1J | 278.26 | 277.57 | 0.01 | 9.90E-01 |
| ENSMUSG00000002233 | GSMG0024776 | Rhoc | ras homolog family member C | 3604.34 | 3655.95 | -0.02 | 9.48E-01 |
| ENSMUSG00000045576 | GSMG0024777 | St7l | suppression of tumorigenicity 7-like | 576.20 | 503.01 | 0.20 | 4.07E-01 |
| ENSMUSG00000048458 | GSMG0024783 | Fam212b | family with sequence similarity 212, member B | 444.78 | 246.50 | 0.85 | 4.05E-06 |
| G00000000561,ENSMUSG000 | GSMG0024787 | Wdr77 // Ovgp1 | WD repeat domain 77 // oviductal glycoprotein 1 | 850.05 | 1181.00 | -0.47 | 1.94E-02 |
| ENSMUSG00000027900 | GSMG0024791 | Dram2 | DNA-damage regulated autophagy modulator 2 | 1226.25 | 1139.19 | 0.11 | 6.87E-01 |
| ENSMUSG00000056260 | GSMG0024794 | Lrif1 | ligand dependent nuclear receptor interacting factor 1 | 444.66 | 442.20 | 0.01 | 9.89E-01 |
| ENSMUSG00000047959 | GSMG0024795 | Kcna3 | potassium voltage-gated channel, shaker-related subfamily, member 3 | 463.31 | 711.36 | -0.62 | 7.67E-03 |
| ENSMUSG00000040724 | GSMG0024798 | Kcna2 | potassium voltage-gated channel, shaker-related subfamily, member 2 | 1316.44 | 656.11 | 1.00 | 4.00E-08 |
| ENSMUSG00000087260 | GSMG0024803 | Lamtor5 | late endosomal/lysosomal adaptor, MAPK and MTOR activator 5 | 1200.49 | 1026.28 | 0.23 | 3.25E-01 |
| ENSMUSG00000027896 | GSMG0024804 | Slc16a4 | solute carrier family 16 (monocarboxylic acid transporters), member 4 | 3682.23 | 2508.89 | 0.55 | 1.10E-02 |
| ENSMUSG00000004032 | GSMG0024809 | Gstm5 | glutathione S-transferase, mu 5 | 2554.55 | 2582.81 | -0.02 | 9.54E-01 |
| ENSMUSG00000050947 | GSMG0024813 | Amigo1 | adhesion molecule with Ig like domain 1 | 170.91 | 146.30 | 0.22 | 6.20E-01 |
| ENSMUSG00000068747 | GSMG0024816 | Sort1 | sortilin 1 | 1828.22 | 1609.50 | 0.18 | 6.06E-01 |
| ENSMUSG00000068744 | GSMG0024820 | Psrc1 | proline/serine-rich coiled-coil 1 | 169.92 | 139.06 | 0.29 | 3.92E-01 |
| ENSMUSG00000048100 | GSMG0024823 | Taf13 | TATA-box binding protein associated factor 13 | 723.12 | 616.15 | 0.23 | 4.05E-01 |
| ENSMUSG00000040389 | GSMG0024824 | Wdr47 | WD repeat domain 47 | 882.95 | 1254.75 | -0.51 | 2.68E-03 |
| ENSMUSG00000027884 | GSMG0024825 | Clcc1 | chloride channel CLIC-like 1 | 1480.88 | 1459.55 | 0.02 | 9.45E-01 |
| ENSMUSG00000040322 | GSMG0024830 | Slc25a24 | solute carrier family 25 (mitochondrial carrier, phosphate carrier), mem | 297.72 | 260.16 | 0.20 | 4.97E-01 |
| NULL | GSMG0024836 | NULL | NULL | 219.56 | 202.48 | 0.11 | 7.68E-01 |
| ENSMUSG00000027966 | GSMG0024837 | Col11a1 | collagen, type XI, alpha 1 | 482.88 | 441.82 | 0.13 | 6.71E-01 |
| ENSMUSG00000033554 | GSMG0024841 | Dph5 | diphthamide biosynthesis 5 | 159.73 | 225.53 | -0.50 | 5.77E-02 |
| ENSMUSG00000027963 | GSMG0024843 | Extl2 | exostoses (multiple)-like 2 | 218.66 | 245.89 | -0.17 | 6.01E-01 |
| ENSMUSG00000000340 | GSMG0024846 | Dbt | dihydrolipoamide branched chain transacylase E2 | 182.36 | 197.05 | -0.11 | 8.13E-01 |
| ENSMUSG00000027961 | GSMG0024847 | Lrrc39 | leucine rich repeat containing 39 | 573.21 | 487.14 | 0.23 | 3.36E-01 |
| ENSMUSG00000027959 | GSMG0024848 | Sass6 | SAS-6 centriolar assembly protein | 296.87 | 436.40 | -0.56 | 1.78E-02 |
| ENSMUSG00000033386 | GSMG0024850 | Frrs1 | ferric-chelate reductase 1 | 618.44 | 692.32 | -0.16 | 5.77E-01 |
| ENSMUSG00000033342 | GSMG0024854 | Plppr5 | phospholipid phosphatase related 5 | 153.74 | 215.48 | -0.48 | 1.41E-01 |
| NULL | GSMG0024857 | NULL | NULL | 1027.81 | 1230.02 | -0.26 | 4.62E-01 |
| NULL | GSMG0024859 | NULL | NULL | 795.77 | 1214.68 | -0.61 | 5.59E-03 |
| NULL | GSMG0024860 | NULL | NULL | 632.44 | 782.98 | -0.31 | 2.07E-01 |

| | | | | | | | |
|---|---|---|---|---|---|---|---|
| NULL | GSMG0024862 | NULL | NULL | 309.84 | 456.43 | -0.56 | 9.33E-03 |
| ENSMUSG00000039887 | GSMG0024869 | Alg14 | asparagine-linked glycosylation 14 | 1367.76 | 1250.31 | 0.13 | 5.96E-01 |
| ENSMUSG00000053931 | GSMG0024870 | Cnn3 | calponin 3, acidic | 4107.43 | 4511.31 | -0.14 | 5.85E-01 |
| ENSMUSG00000028128 | GSMG0024874 | F3 | coagulation factor III | 471.16 | 161.74 | 1.55 | 6.31E-14 |
| ENSMUSG00000039831 | GSMG0024875 | Arhgap29 | Rho GTPase activating protein 29 | 5819.59 | 4932.90 | 0.24 | 2.40E-01 |
| ENSMUSG00000028124 | GSMG0024878 | Gclm | glutamate-cysteine ligase, modifier subunit | 908.01 | 826.38 | 0.14 | 6.35E-01 |
| ENSMUSG00000039756 | GSMG0024879 | Dnttip2 | deoxynucleotidyltransferase, terminal, interacting protein 2 | 1615.43 | 2197.57 | -0.44 | 3.36E-02 |
| ENSMUSG00000028121 | GSMG0024880 | Bcar3 | breast cancer anti-estrogen resistance 3 | 795.04 | 571.28 | 0.48 | 7.75E-03 |
| ENSMUSG00000053965 | GSMG0024882 | Pde5a | phosphodiesterase 5A, cGMP-specific | 310.69 | 342.81 | -0.14 | 6.77E-01 |
| ENSMUSG00000054091 | GSMG0024884 | 1810037I17Rik | RIKEN cDNA 1810037I17 gene | 2221.71 | 1925.62 | 0.21 | 4.80E-01 |
| ENSMUSG00000039234 | GSMG0024886 | Sec24d | Sec24 related gene family, member D (S. cerevisiae) | 630.71 | 716.81 | -0.19 | 4.31E-01 |
| ENSMUSG00000027978 | GSMG0024887 | Prss12 | protease, serine 12 neurotrypsin (motopsin) | 147.26 | 373.75 | -1.33 | 2.03E-06 |
| ENSMUSG00000044528 | GSMG0024889 | Tram1l1 | translocation associated membrane protein 1-like 1 | 588.51 | 697.59 | -0.25 | 2.95E-01 |
| ENSMUSG00000053819 | GSMG0024894 | Camk2d | calcium/calmodulin-dependent protein kinase II, delta | 656.78 | 518.44 | 0.34 | 1.45E-01 |
| ENSMUSG00000051278 | GSMG0024903 | Zgrf1 | zinc finger, GRF-type containing 1 | 199.00 | 216.13 | -0.12 | 7.36E-01 |
| ENSMUSG00000046688 | GSMG0024905 | Tifa | TRAF-interacting protein with forkhead-associated domain | 503.76 | 602.55 | -0.26 | 2.19E-01 |
| ENSMUSG00000041220 | GSMG0024916 | Elovl6 | ELOVL family member 6, elongation of long chain fatty acids (yeast) | 1032.61 | 1488.25 | -0.53 | 5.91E-04 |
| ENSMUSG00000027999 | GSMG0024923 | Pla2g12a | phospholipase A2, group XIIA | 1243.52 | 1582.98 | -0.35 | 5.20E-02 |
| ENSMUSG00000027997 | GSMG0024924 | Casp6 | caspase 6 | 1409.55 | 2032.19 | -0.53 | 7.71E-03 |
| ENSMUSG00000027985 | GSMG0024937 | Lef1 | lymphoid enhancer binding factor 1 | 1959.42 | 1583.82 | 0.31 | 6.95E-02 |
| NULL | GSMG0024939 | NULL | NULL | 627.49 | 339.36 | 0.89 | 3.51E-05 |
| NULL | GSMG0024940 | NULL | NULL | 207.43 | 139.45 | 0.57 | 9.16E-02 |
| NULL | GSMG0024943 | NULL | NULL | 187.17 | 190.64 | -0.03 | 9.55E-01 |
| NULL | GSMG0024945 | NULL | NULL | 429.03 | 490.25 | -0.19 | 5.63E-01 |
| ENSMUSG00000028032 | GSMG0024948 | Papss1 | 3'-phosphoadenosine 5'-phosphosulfate synthase 1 | 1581.95 | 2040.88 | -0.37 | 3.56E-02 |
| ENSMUSG00000028031 | GSMG0024950 | Dkk2 | dickkopf WNT signaling pathway inhibitor 2 | 355.85 | 231.94 | 0.62 | 1.82E-02 |
| ENSMUSG00000028016 | GSMG0024958 | Ints12 | integrator complex subunit 12 | 225.59 | 275.38 | -0.29 | 3.66E-01 |
| ENSMUSG00000028013 | GSMG0024959 | Ppa2 | pyrophosphatase (inorganic) 2 | 570.72 | 667.08 | -0.22 | 3.30E-01 |
| ENSMUSG00000044365 | GSMG0024962 | Cxxc4 | CXXC finger 4 | 65.49 | 156.61 | -1.26 | 7.78E-06 |
| ENSMUSG00000045328 | GSMG0024965 | Cenpe | centromere protein E | 171.81 | 441.18 | -1.35 | 1.25E-07 |
| ENSMUSG00000078578 | GSMG0024972 | Ube2d3 | ubiquitin-conjugating enzyme E2D 3 | 1028.84 | 1272.15 | -0.31 | 1.70E-01 |
| ENSMUSG00000028164 | GSMG0024975 | Manba | mannosidase, beta A, lysosomal | 635.94 | 464.07 | 0.46 | 3.36E-02 |
| ENSMUSG00000053897 | GSMG0024979 | Slc39a8 | solute carrier family 39 (metal ion transporter), member 8 | 5332.79 | 9492.75 | -0.83 | 1.23E-07 |

| | | | | | | | |
|---|---|---|---|---|---|---|---|
| ENSMUSG00000028161 | GSMG0024983 | Ppp3ca | protein phosphatase 3, catalytic subunit, alpha isoform | 6151.33 | 5714.73 | 0.11 | 6.49E-01 |
| NULL | GSMG0024986 | NULL | NULL | 611.49 | 342.67 | 0.84 | 1.60E-06 |
| NULL | GSMG0024987 | NULL | NULL | 414.39 | 245.73 | 0.76 | 1.42E-04 |
| NULL | GSMG0024988 | NULL | NULL | 397.20 | 260.15 | 0.61 | 1.54E-02 |
| NULL | GSMG0024989 | NULL | NULL | 537.00 | 438.50 | 0.29 | 1.85E-01 |
| NULL | GSMG0024990 | NULL | NULL | 416.53 | 340.83 | 0.29 | 3.13E-01 |
| NULL | GSMG0024992 | NULL | NULL | 308.40 | 196.05 | 0.66 | 3.94E-03 |
| ENSMUSG00000054690 | GSMG0024993 | Emcn | endomucin | 1943.92 | 1154.19 | 0.75 | 1.33E-04 |
| ENSMUSG00000046818 | GSMG0024995 | Ddit4l | DNA-damage-inducible transcript 4-like | 1691.90 | 680.63 | 1.31 | 3.68E-15 |
| ENSMUSG00000074212 | GSMG0024997 | Dnajb14 | DnaJ heat shock protein family (Hsp40) member B14 | 176.83 | 185.60 | -0.07 | 8.80E-01 |
| NULL | GSMG0025001 | NULL | NULL | 127.77 | 109.42 | 0.22 | 6.03E-01 |
| ENSMUSG00000091512 | GSMG0025002 | Lamtor3 | late endosomal/lysosomal adaptor, MAPK and MTOR activator 3 | 714.43 | 645.95 | 0.14 | 6.49E-01 |
| ENSMUSG00000004127 | GSMG0025004 | Trmt10a | tRNA methyltransferase 10A | 106.51 | 148.56 | -0.48 | 1.32E-01 |
| ENSMUSG00000074207 | GSMG0025007 | Adh1 | alcohol dehydrogenase 1 (class I) | 375.68 | 457.69 | -0.29 | 2.79E-01 |
| ENSMUSG00000028138 | GSMG0025013 | Adh5 | alcohol dehydrogenase 5 (class III), chi polypeptide | 2140.42 | 2600.68 | -0.28 | 1.54E-01 |
| ENSMUSG00000028156 | GSMG0025015 | Eif4e | eukaryotic translation initiation factor 4E | 590.56 | 739.89 | -0.32 | 1.60E-01 |
| ENSMUSG00000028152 | GSMG0025020 | Tspan5 | tetraspanin 5 | 2619.85 | 2524.89 | 0.05 | 8.50E-01 |
| NULL | GSMG0025023 | NULL | NULL | 299.93 | 268.63 | 0.16 | 6.09E-01 |
| NULL | GSMG0025024 | NULL | NULL | 302.59 | 253.41 | 0.26 | 3.89E-01 |
| ENSMUSG00000059921 | GSMG0025028 | Unc5c | unc-5 netrin receptor C | 293.27 | 432.19 | -0.56 | 1.97E-02 |
| NULL | GSMG0025031 | NULL | NULL | 141.10 | 148.07 | -0.08 | 8.94E-01 |
| ENSMUSG00000105504 | GSMG0025042 | Gbp5 | guanylate binding protein 5 | 170.28 | 98.65 | 0.79 | 3.53E-02 |
| ENSMUSG00000040253 | GSMG0025044 | Gbp7 | guanylate binding protein 7 | 1062.54 | 1034.83 | 0.04 | 9.28E-01 |
| ENSMUSG00000028268 | GSMG0025045 | Gbp3 | guanylate binding protein 3 | 1044.63 | 1051.11 | -0.01 | 9.86E-01 |
| ENSMUSG00000028270 | GSMG0025046 | Gbp2 | guanylate binding protein 2 | 410.97 | 270.37 | 0.61 | 8.34E-02 |
| ENSMUSG00000028271 | GSMG0025048 | Gtf2b | general transcription factor IIB | 641.11 | 659.79 | -0.04 | 9.02E-01 |
| ENSMUSG00000037072 | GSMG0025059 | Sept15 | selenoprotein 15 | 10510.97 | 11031.69 | -0.07 | 8.04E-01 |
| ENSMUSG00000028256 | GSMG0025063 | Odf2l | outer dense fiber of sperm tails 2-like | 277.59 | 264.56 | 0.07 | 8.72E-01 |
| ENSMUSG00000028197 | GSMG0025064 | Col24a1 | collagen, type XXIV, alpha 1 | 918.52 | 324.02 | 1.50 | 1.27E-10 |
| ENSMUSG00000074182 | GSMG0025065 | Znhit6 | zinc finger, HIT type 6 | 271.84 | 386.74 | -0.51 | 3.03E-02 |
| ENSMUSG00000028194 | GSMG0025068 | Ddah1 | dimethylarginine dimethylaminohydrolase 1 | 5319.16 | 6544.60 | -0.30 | 8.06E-02 |
| ENSMUSG00000028191 | GSMG0025070 | Bcl10 | B cell leukemia/lymphoma 10 | 1431.38 | 1413.02 | 0.02 | 9.58E-01 |
| ENSMUSG00000036873 | GSMG0025071 | 2410004B18Rik | RIKEN cDNA 2410004B18 gene | 715.27 | 726.88 | -0.02 | 9.42E-01 |

| | | | | | | | |
|---|---|---|---|---|---|---|---|
| ENSMUSG00000036825 | GSMG0025077 | Ssx2ip | synovial sarcoma, X breakpoint 2 interacting protein | 1633.63 | 2210.33 | -0.44 | 5.22E-03 |
| ENSMUSG00000028189 | GSMG0025078 | Ctbs | chitobiase, di-N-acetyl- | 174.57 | 143.55 | 0.29 | 4.01E-01 |
| ENSMUSG00000036745 | GSMG0025083 | Ttll7 | tubulin tyrosine ligase-like family, member 7 | 455.58 | 244.92 | 0.89 | 1.09E-05 |
| ENSMUSG00000039167 | GSMG0025094 | Adgrl4 | adhesion G protein-coupled receptor L4 | 18507.84 | 14021.64 | 0.40 | 1.15E-02 |
| ENSMUSG00000028034 | GSMG0025099 | Fubp1 | far upstream element (FUSE) binding protein 1 | 3355.48 | 3786.18 | -0.17 | 3.92E-01 |
| ENSMUSG00000025437 | GSMG0025100 | Usp33 | ubiquitin specific peptidase 33 | 2046.86 | 2057.15 | -0.01 | 9.80E-01 |
| ENSMUSG00000039068 | GSMG0025102 | Zzz3 | zinc finger, ZZ domain containing 3 | 495.81 | 533.26 | -0.10 | 7.02E-01 |
| ENSMUSG00000097745 | GSMG0025105 | AI115009 | expressed sequence AI115009 | 259.07 | 108.35 | 1.26 | 1.80E-06 |
| ENSMUSG00000039047 | GSMG0025106 | Pigk | phosphatidylinositol glycan anchor biosynthesis, class K | 1031.45 | 1137.45 | -0.14 | 5.30E-01 |
| ENSMUSG00000028360 | GSMG0025109 | Slc44a5 | solute carrier family 44, member 5 | 313.14 | 394.29 | -0.33 | 1.72E-01 |
| ENSMUSG00000028199 | GSMG0025112 | Cryz | crystallin, zeta | 215.89 | 266.71 | -0.30 | 3.28E-01 |
| ENSMUSG00000040037 | GSMG0025119 | Negr1 | neuronal growth regulator 1 | 1018.50 | 1043.41 | -0.03 | 9.04E-01 |
| NULL | GSMG0025120 | NULL | NULL | 403.59 | 305.29 | 0.40 | 1.15E-01 |
| NULL | GSMG0025123 | NULL | NULL | 174.42 | 122.65 | 0.51 | 7.57E-02 |
| NULL | GSMG0025124 | NULL | NULL | 237.19 | 156.31 | 0.60 | 9.53E-02 |
| NULL | GSMG0025126 | NULL | NULL | 212.15 | 142.78 | 0.57 | 7.17E-02 |
| NULL | GSMG0025129 | NULL | NULL | 195.81 | 166.23 | 0.24 | 5.11E-01 |
| ENSMUSG00000028180 | GSMG0025130 | Zranb2 | zinc finger, RAN-binding domain containing 2 | 4175.83 | 3599.48 | 0.21 | 4.04E-01 |
| ENSMUSG00000040016 | GSMG0025131 | Ptger3 | prostaglandin E receptor 3 (subtype EP3) | 76.65 | 133.17 | -0.79 | 8.60E-03 |
| ENSMUSG00000039988 | GSMG0025133 | Ankrd13c | ankyrin repeat domain 13c | 747.33 | 693.31 | 0.11 | 6.81E-01 |
| ENSMUSG00000063052 | GSMG0025135 | Lrrc40 | leucine rich repeat containing 40 | 430.67 | 616.89 | -0.52 | 2.46E-02 |
| NULL | GSMG0025136 | NULL | NULL | 585.09 | 474.59 | 0.30 | 2.37E-01 |
| ENSMUSG00000028173 | GSMG0025141 | Wls | wntless homolog (Drosophila) | 1943.17 | 1741.90 | 0.16 | 5.18E-01 |
| ENSMUSG00000040374 | GSMG0025145 | Pex2 | peroxisomal biogenesis factor 2 | 1259.85 | 1402.88 | -0.16 | 4.83E-01 |
| ENSMUSG00000040289 | GSMG0025148 | Hey1 | hairy/enhancer-of-split related with YRPW motif 1 | 3394.39 | 1994.75 | 0.77 | 1.37E-04 |
| ENSMUSG00000040269 | GSMG0025150 | Mrps28 | mitochondrial ribosomal protein S28 | 518.18 | 763.41 | -0.56 | 2.25E-03 |
| ENSMUSG00000027506 | GSMG0025152 | Tpd52 | tumor protein D52 | 2825.74 | 2167.07 | 0.38 | 9.34E-02 |
| ENSMUSG00000040209 | GSMG0025153 | Zfp704 | zinc finger protein 704 | 212.35 | 218.23 | -0.04 | 9.42E-01 |
| ENSMUSG00000027531 | GSMG0025165 | Impa1 | inositol (myo)-1(or 4)-monophosphatase 1 | 642.42 | 635.17 | 0.02 | 9.63E-01 |
| ENSMUSG00000027534 | GSMG0025168 | Snx16 | sorting nexin 16 | 208.20 | 248.96 | -0.26 | 3.72E-01 |
| NULL | GSMG0025175 | 4930539N22Rik | RIKEN cDNA 4930539N22 gene | 105.71 | 105.66 | 0.01 | 9.89E-01 |
| ENSMUSG00000039519 | GSMG0025190 | Cyp7b1 | cytochrome P450, family 7, subfamily b, polypeptide 1 | 180.15 | 102.79 | 0.81 | 2.75E-03 |
| ENSMUSG00000027599 | GSMG0025194 | Armc1 | armadillo repeat containing 1 | 522.61 | 593.98 | -0.19 | 5.23E-01 |

| | | | | | | | |
|---|---|---|---|---|---|---|---|
| ENSMUSG00000069094 | GSMG0025197 | Pde7a | phosphodiesterase 7A | 905.26 | 1060.36 | -0.23 | 2.79E-01 |
| ENSMUSG00000049796 | GSMG0025199 | Crh | corticotropin releasing hormone | 139.57 | 101.68 | 0.46 | 2.32E-01 |
| ENSMUSG00000027615 | GSMG0025200 | Hps3 | Hermansky-Pudlak syndrome 3 | 395.50 | 469.71 | -0.25 | 3.14E-01 |
| ENSMUSG00000019528 | GSMG0025202 | Gyg | glycogenin | 842.35 | 953.95 | -0.18 | 4.30E-01 |
| NULL | GSMG0025208 | NULL | NULL | 72.93 | 81.07 | -0.16 | 8.02E-01 |
| ENSMUSG00000063887 | GSMG0025214 | Nlgn1 | neuroligin 1 | 248.00 | 397.63 | -0.68 | 3.67E-03 |
| NULL | GSMG0025218 | NULL | NULL | 206.96 | 140.11 | 0.57 | 1.51E-01 |
| NULL | GSMG0025220 | NULL | NULL | 456.35 | 354.00 | 0.37 | 1.66E-01 |
| NULL | GSMG0025222 | NULL | NULL | 845.28 | 733.67 | 0.20 | 5.12E-01 |
| NULL | GSMG0025225 | NULL | NULL | 457.83 | 374.46 | 0.29 | 3.29E-01 |
| NULL | GSMG0025226 | NULL | NULL | 463.88 | 456.36 | 0.02 | 9.61E-01 |
| NULL | GSMG0025227 | NULL | NULL | 686.28 | 606.19 | 0.18 | 5.27E-01 |
| NULL | GSMG0025230 | NULL | NULL | 1149.32 | 963.51 | 0.25 | 2.84E-01 |
| NULL | GSMG0025231 | NULL | NULL | 629.81 | 511.61 | 0.30 | 2.66E-01 |
| NULL | GSMG0025232 | NULL | NULL | 256.19 | 199.40 | 0.36 | 2.98E-01 |
| NULL | GSMG0025233 | NULL | NULL | 206.64 | 216.04 | -0.07 | 8.74E-01 |
| NULL | GSMG0025240 | NULL | NULL | 205.76 | 148.01 | 0.47 | 1.55E-01 |
| NULL | GSMG0025241 | NULL | NULL | 82.48 | 90.06 | -0.13 | 8.04E-01 |
| ENSMUSG00000027699 | GSMG0025248 | Ect2 | ect2 oncogene | 291.11 | 620.07 | -1.09 | 2.38E-06 |
| ENSMUSG00000039286 | GSMG0025250 | Fndc3b | fibronectin type III domain containing 3B | 694.64 | 619.13 | 0.17 | 5.84E-01 |
| NULL | GSMG0025254 | NULL | NULL | 334.69 | 226.56 | 0.56 | 4.10E-02 |
| ENSMUSG00000043164 | GSMG0025256 | Tmem212 | transmembrane protein 212 | 65.11 | 27.88 | 1.22 | 4.72E-03 |
| ENSMUSG00000027684 | GSMG0025264 | Mecom | MDS1 and EVI1 complex locus | 519.82 | 342.22 | 0.61 | 1.97E-03 |
| NULL | GSMG0025271 | NULL | NULL | 178.20 | 122.19 | 0.55 | 1.24E-01 |
| ENSMUSG00000037652 | GSMG0025280 | Phc3 | polyhomeotic-like 3 (Drosophila) | 322.26 | 387.11 | -0.26 | 3.72E-01 |
| ENSMUSG00000069072 | GSMG0025283 | Slc7a14 | solute carrier family 7 (cationic amino acid transporter, y+ system), mer | 212.70 | 272.17 | -0.35 | 2.52E-01 |
| ENSMUSG00000027669 | GSMG0025289 | Gnb4 | guanine nucleotide binding protein (G protein), beta 4 | 2811.95 | 3463.42 | -0.30 | 7.43E-02 |
| ENSMUSG00000037531 | GSMG0025290 | Mrpl47 | mitochondrial ribosomal protein L47 | 202.87 | 234.57 | -0.21 | 5.68E-01 |
| ENSMUSG00000027674 | GSMG0025291 | Pex5l | peroxisomal biogenesis factor 5-like | 407.57 | 214.36 | 0.92 | 8.23E-06 |
| NULL | GSMG0025294 | NULL | NULL | 878.14 | 489.56 | 0.84 | 5.56E-05 |
| NULL | GSMG0025295 | NULL | NULL | 976.55 | 562.50 | 0.80 | 4.22E-06 |
| ENSMUSG00000027676 | GSMG0025298 | Ccdc39 | coiled-coil domain containing 39 | 473.45 | 554.32 | -0.23 | 4.45E-01 |
| ENSMUSG00000027679 | GSMG0025300 | Dnajc19 | DnaJ heat shock protein family (Hsp40) member C19 | 621.09 | 544.24 | 0.19 | 4.25E-01 |

| | | | | | | | |
|---|---|---|---|---|---|---|---|
| NULL | GSMG0025312 | NULL | NULL | 104.52 | 60.81 | 0.78 | 1.91E-02 |
| ENSMUSG00000027709 | GSMG0025313 | Mccc1 | methylcrotonoyl-Coenzyme A carboxylase 1 (alpha) | 1498.93 | 1450.06 | 0.05 | 8.63E-01 |
| ENSMUSG00000027712 | GSMG0025317 | Anxa5 | annexin A5 | 5838.30 | 4195.47 | 0.48 | 1.17E-02 |
| ENSMUSG00000027715 | GSMG0025321 | Ccna2 | cyclin A2 | 498.01 | 933.36 | -0.90 | 8.94E-07 |
| ENSMUSG00000037325 | GSMG0025322 | Bbs7 | Bardet-Biedl syndrome 7 (human) | 379.05 | 386.53 | -0.03 | 9.42E-01 |
| ENSMUSG00000027716 | GSMG0025323 | Trpc3 | transient receptor potential cation channel, subfamily C, member 3 | 1068.67 | 911.79 | 0.23 | 2.60E-01 |
| NULL | GSMG0025332 | NULL | NULL | 931.27 | 445.70 | 1.06 | 2.14E-04 |
| ENSMUSG00000044864 | GSMG0025338 | Ankrd50 | ankyrin repeat domain 50 | 2959.20 | 2379.20 | 0.31 | 1.72E-01 |
| ENSMUSG00000025759 | GSMG0025343 | Mfsd8 | major facilitator superfamily domain containing 8 | 231.44 | 252.39 | -0.12 | 7.15E-01 |
| NULL | GSMG0025346 | NULL | NULL | 699.21 | 780.73 | -0.16 | 5.53E-01 |
| ENSMUSG00000049940 | GSMG0025350 | Pgrmc2 | progesterone receptor membrane component 2 | 1318.11 | 1493.97 | -0.18 | 4.28E-01 |
| ENSMUSG00000059834 | GSMG0025356 | Sclt1 | sodium channel and clathrin linker 1 | 567.81 | 535.81 | 0.08 | 7.96E-01 |
| ENSMUSG00000097040 | GSMG0025359 | 2610316D01Rik | RIKEN cDNA 2610316D01 gene | 153.11 | 147.80 | 0.05 | 9.08E-01 |
| ENSMUSG00000037892 | GSMG0025364 | Pcdh18 | protocadherin 18 | 4003.70 | 5070.92 | -0.34 | 3.45E-02 |
| ENSMUSG00000027737 | GSMG0025365 | Slc7a11 | solute carrier family 7 (cationic amino acid transporter, y+ system), mer | 174.54 | 94.51 | 0.89 | 7.11E-03 |
| ENSMUSG00000037174 | GSMG0025375 | Elf2 | E74-like factor 2 | 626.42 | 666.76 | -0.09 | 7.69E-01 |
| ENSMUSG00000037152 | GSMG0025378 | Ndufc1 | NADH dehydrogenase (ubiquinone) 1, subcomplex unknown, 1 | 1214.57 | 1432.28 | -0.24 | 4.90E-01 |
| ENSMUSG00000037111 | GSMG0025380 | Setd7 | SET domain containing (lysine methyltransferase) 7 | 89.92 | 91.73 | -0.03 | 9.63E-01 |
| NULL | GSMG0025384 | NULL | NULL | 104.89 | 82.98 | 0.34 | 4.20E-01 |
| NULL | GSMG0025391 | NULL | NULL | 1025.22 | 635.89 | 0.69 | 8.03E-04 |
| ENSMUSG00000027742 | GSMG0025397 | Cog6 | component of oligomeric golgi complex 6 | 897.91 | 872.03 | 0.04 | 8.88E-01 |
| ENSMUSG00000042997 | GSMG0025400 | Nhlrc3 | NHL repeat containing 3 | 189.94 | 201.54 | -0.09 | 8.31E-01 |
| ENSMUSG00000027746 | GSMG0025404 | Ufm1 | ubiquitin-fold modifier 1 | 1325.72 | 1422.75 | -0.10 | 7.15E-01 |
| ENSMUSG00000027752 | GSMG0025407 | Exosc8 | exosome component 8 | 566.29 | 799.07 | -0.50 | 1.50E-02 |
| ENSMUSG00000036615 | GSMG0025409 | Rfxap | regulatory factor X-associated protein | 1055.90 | 1031.11 | 0.03 | 9.15E-01 |
| NULL | GSMG0025414 | NULL | NULL | 349.18 | 356.97 | -0.03 | 9.41E-01 |
| ENSMUSG00000027799 | GSMG0025421 | Nbea | neurobeachin | 141.99 | 245.37 | -0.78 | 1.08E-03 |
| NULL | GSMG0025426 | NULL | NULL | 98.69 | 126.04 | -0.36 | 3.95E-01 |
| ENSMUSG00000027800 | GSMG0025433 | Tm4sf1 | transmembrane 4 superfamily member 1 | 8951.08 | 2902.99 | 1.62 | 2.12E-24 |
| ENSMUSG00000027803 | GSMG0025434 | Wwtr1 | WW domain containing transcription regulator 1 | 7384.60 | 8442.02 | -0.19 | 3.04E-01 |
| ENSMUSG00000036513 | GSMG0025438 | Commd2 | COMM domain containing 2 | 1031.76 | 1292.95 | -0.33 | 1.53E-01 |
| ENSMUSG00000027805 | GSMG0025442 | Pfn2 | profilin 2 | 1528.78 | 2391.15 | -0.65 | 6.92E-05 |
| ENSMUSG00000027808 | GSMG0025446 | Serp1 | stress-associated endoplasmic reticulum protein 1 | 4609.84 | 5113.21 | -0.15 | 5.48E-01 |

| | | | | | | | |
|---|---|---|---|---|---|---|---|
| ENSMUSG00000036432 | GSMG0025448 | Siah2 | seven in absentia 2 | 526.98 | 462.49 | 0.19 | 5.12E-01 |
| ENSMUSG00000036381 | GSMG0025453 | P2ry14 | purinergic receptor P2Y, G-protein coupled, 14 | 2018.66 | 759.58 | 1.41 | 3.83E-13 |
| ENSMUSG00000036362 | GSMG0025455 | P2ry13 | purinergic receptor P2Y, G-protein coupled 13 | 419.53 | 136.98 | 1.62 | 2.72E-11 |
| ENSMUSG00000036353 | GSMG0025456 | P2ry12 | purinergic receptor P2Y, G-protein coupled 12 | 1104.66 | 312.33 | 1.82 | 2.18E-16 |
| NULL | GSMG0025461 | NULL | NULL | 232.91 | 212.34 | 0.13 | 8.32E-01 |
| ENSMUSG00000027770 | GSMG0025466 | Dhx36 | DEAH (Asp-Glu-Ala-His) box polypeptide 36 | 497.58 | 567.02 | -0.19 | 5.30E-01 |
| ENSMUSG00000048581 | GSMG0025471 | E130311K13Rik | RIKEN cDNA E130311K13 gene | 80.29 | 73.90 | 0.12 | 8.53E-01 |
| ENSMUSG00000027822 | GSMG0025472 | Slc33a1 | solute carrier family 33 (acetyl-CoA transporter), member 1 | 260.36 | 271.89 | -0.06 | 8.88E-01 |
| ENSMUSG00000027828 | GSMG0025478 | Ssr3 | signal sequence receptor, gamma | 6586.51 | 8346.93 | -0.34 | 6.98E-02 |
| ENSMUSG00000027829 | GSMG0025483 | Ccnl1 | cyclin L1 | 2035.11 | 1722.27 | 0.24 | 1.98E-01 |
| ENSMUSG00000047557 | GSMG0025488 | Lxn | latexin | 1239.04 | 2998.20 | -1.28 | 1.57E-15 |
| ENSMUSG00000027778 | GSMG0025494 | Ift80 | intraflagellar transport 80 | 153.45 | 134.15 | 0.20 | 6.29E-01 |
| ENSMUSG00000034317 | GSMG0025495 | Trim59 | tripartite motif-containing 59 | 349.14 | 580.19 | -0.73 | 1.15E-03 |
| ENSMUSG00000027782 | GSMG0025498 | Kpna4 | karyopherin (importin) alpha 4 | 868.01 | 840.32 | 0.05 | 8.89E-01 |
| ENSMUSG00000043300 | GSMG0025503 | B3galnt1 | UDP-GalNAc:betaGlcNAc beta 1,3-galactosaminyltransferase, polypep | 679.17 | 817.04 | -0.27 | 2.53E-01 |
| ENSMUSG00000048304 | GSMG0025510 | Slitrk3 | SLIT and NTRK-like family, member 3 | 1194.77 | 1272.48 | -0.09 | 7.72E-01 |
| ENSMUSG00000027792 | GSMG0025512 | Bche | butyrylcholinesterase | 849.59 | 338.29 | 1.33 | 8.39E-12 |
| ENSMUSG00000034109 | GSMG0025523 | Golim4 | golgi integral membrane protein 4 | 3792.22 | 2582.45 | 0.55 | 1.11E-04 |
| ENSMUSG00000062232 | GSMG0025529 | Rapgef2 | Rap guanine nucleotide exchange factor (GEF) 2 | 2310.29 | 1801.29 | 0.36 | 1.62E-01 |
| NULL | GSMG0025533 | NULL | NULL | 966.06 | 962.81 | 0.00 | 9.88E-01 |
| NULL | GSMG0025536 | NULL | NULL | 846.11 | 758.66 | 0.16 | 6.96E-01 |
| ENSMUSG00000061175 | GSMG0025537 | Fnip2 | folliculin interacting protein 2 | 192.45 | 244.29 | -0.34 | 3.66E-01 |
| ENSMUSG00000027809 | GSMG0025539 | Etfdh | electron transferring flavoprotein, dehydrogenase | 871.85 | 963.72 | -0.14 | 5.83E-01 |
| ENSMUSG00000034009 | GSMG0025542 | Rxfp1 | relaxin/insulin-like family peptide receptor 1 | 279.12 | 594.34 | -1.09 | 4.20E-09 |
| ENSMUSG00000027956 | GSMG0025544 | Tmem144 | transmembrane protein 144 | 122.65 | 104.59 | 0.23 | 5.89E-01 |
| ENSMUSG00000033981 | GSMG0025547 | Gria2 | glutamate receptor, ionotropic, AMPA2 (alpha 2) | 12764.26 | 13241.96 | -0.05 | 8.46E-01 |
| NULL | GSMG0025548 | NULL | NULL | 196.41 | 199.22 | -0.02 | 9.61E-01 |
| NULL | GSMG0025549 | NULL | NULL | 642.48 | 542.03 | 0.25 | 3.44E-01 |
| ENSMUSG00000028020 | GSMG0025552 | Glrb | glycine receptor, beta subunit | 580.09 | 551.14 | 0.07 | 8.12E-01 |
| ENSMUSG00000028005 | GSMG0025556 | Gucy1b3 | guanylate cyclase 1, soluble, beta 3 | 1695.86 | 1673.38 | 0.02 | 9.56E-01 |
| ENSMUSG00000033910 | GSMG0025557 | Gucy1a3 | guanylate cyclase 1, soluble, alpha 3 | 2068.04 | 2993.44 | -0.53 | 3.19E-04 |
| ENSMUSG00000033767 | GSMG0025568 | D930015E06Rik | RIKEN cDNA D930015E06 gene | 547.72 | 1290.71 | -1.24 | 9.24E-16 |
| ENSMUSG00000027993 | GSMG0025572 | Trim2 | tripartite motif-containing 2 | 967.99 | 1757.76 | -0.86 | 9.96E-08 |

| | | | | | | | |
|---|---|---|---|---|---|---|---|
| NULL | GSMG0025581 | NULL | NULL | 56.64 | 36.16 | 0.64 | 4.37E-01 |
| ENSMUSG00000028078 | GSMG0025594 | Dclk2 | doublecortin-like kinase 2 | 705.29 | 1233.24 | -0.81 | 3.88E-04 |
| ENSMUSG00000028076 | GSMG0025597 | Cd1d1 | CD1d1 antigen | 174.41 | 328.70 | -0.91 | 3.79E-05 |
| ENSMUSG00000041734 | GSMG0025600 | Kirrel | kin of IRRE like (Drosophila) | 252.49 | 271.60 | -0.10 | 8.17E-01 |
| ENSMUSG00000015852 | GSMG0025604 | Fcrls | Fc receptor-like S, scavenger receptor | 497.18 | 190.20 | 1.39 | 1.29E-08 |
| ENSMUSG00000004895 | GSMG0025607 | Prcc | papillary renal cell carcinoma (translocation-associated) | 106.26 | 147.91 | -0.47 | 1.82E-01 |
| ENSMUSG00000004896 | GSMG0025608 | Rrnad1 | ribosomal RNA adenine dimethylase domain containing 1 | 252.28 | 216.34 | 0.22 | 4.74E-01 |
| ENSMUSG00000004892 | GSMG0025609 | Bcan | brevican | 2709.51 | 2157.87 | 0.33 | 2.04E-01 |
| ENSMUSG00000004894 | GSMG0025610 | Hapln2 | hyaluronan and proteoglycan link protein 2 | 91.80 | 12.59 | 2.85 | 3.67E-11 |
| ENSMUSG00000028070 | GSMG0025611 | Apoa1bp | apolipoprotein A-I binding protein | 1459.07 | 1576.44 | -0.11 | 6.57E-01 |
| NULL | GSMG0025616 | NULL | NULL | 112.30 | 223.08 | -0.99 | 4.89E-05 |
| ENSMUSG00000028066 | GSMG0025623 | Pmf1 | polyamine-modulated factor 1 | 278.03 | 486.32 | -0.80 | 9.17E-05 |
| ENSMUSG00000050144 | GSMG0025624 | Slc25a44 | solute carrier family 25, member 44 | 410.70 | 432.51 | -0.07 | 8.47E-01 |
| ENSMUSG00000028064 | GSMG0025625 | Sema4a | sema domain, immunoglobulin domain (Ig), transmembrane domain (T | 1645.63 | 1314.21 | 0.32 | 1.66E-01 |
| ENSMUSG00000028063 | GSMG0025626 | Lmna | lamin A | 3001.66 | 1986.06 | 0.60 | 2.68E-04 |
| ENSMUSG00000028062 | GSMG0025630 | Lamtor2 | late endosomal/lysosomal adaptor, MAPK and MTOR activator 2 | 1045.42 | 1132.91 | -0.12 | 6.32E-01 |
| NULL | GSMG0025631 | NULL | NULL | 65.93 | 74.14 | -0.18 | 7.52E-01 |
| ENSMUSG00000027947 | GSMG0025634 | Il6ra | interleukin 6 receptor, alpha | 137.07 | 56.69 | 1.28 | 9.84E-04 |
| G00000068923,ENSMUSG000 | GSMG0025638 | Syt11 // 1500004A | synaptotagmin XI // RIKEN cDNA 1500004A13 gene | 6785.73 | 7027.24 | -0.05 | 8.47E-01 |
| ENSMUSG00000068922 | GSMG0025639 | Msto1 | misato 1, mitochondrial distribution and morphology regulator | 1057.68 | 1300.93 | -0.30 | 1.48E-01 |
| ENSMUSG00000068921 | GSMG0025640 | Dap3 | death associated protein 3 | 905.10 | 1056.40 | -0.22 | 3.61E-01 |
| ENSMUSG00000041263 | GSMG0025642 | Rusc1 | RUN and SH3 domain containing 1 | 2108.01 | 2147.63 | -0.03 | 9.37E-01 |
| ENSMUSG00000059743 | GSMG0025643 | Fdps | farnesyl diphosphate synthetase | 604.35 | 1379.61 | -1.19 | 7.59E-14 |
| ENSMUSG00000028051 | GSMG0025644 | Hcn3 | hyperpolarization-activated, cyclic nucleotide-gated K+ 3 | 153.11 | 566.35 | -1.89 | 8.60E-26 |
| ENSMUSG00000064068 | GSMG0025645 | Mtx1 | metaxin 1 | 286.43 | 344.94 | -0.27 | 3.16E-01 |
| ENSMUSG00000042766 | GSMG0025646 | Trim46 | tripartite motif-containing 46 | 1087.34 | 1074.62 | 0.02 | 9.61E-01 |
| ENSMUSG00000027953 | GSMG0025647 | Slc50a1 | solute carrier family 50 (sugar transporter), member 1 | 2332.98 | 1554.46 | 0.59 | 9.89E-05 |
| ENSMUSG00000027954 | GSMG0025648 | Efna1 | ephrin A1 | 2753.87 | 3539.98 | -0.36 | 1.84E-02 |
| ENSMUSG00000028039 | GSMG0025649 | Efna3 | ephrin A3 | 216.27 | 399.96 | -0.89 | 1.44E-04 |
| ENSMUSG00000028040 | GSMG0025650 | Efna4 | ephrin A4 | 133.90 | 159.67 | -0.26 | 5.24E-01 |
| ENSMUSG00000028041 | GSMG0025651 | Adam15 | a disintegrin and metallopeptidase domain 15 (metargidin) | 10556.12 | 5888.80 | 0.84 | 1.73E-08 |
| G00000042642,ENSMUSG000 | GSMG0025654 | Lenep // Flad1 | lens epithelial protein // flavin adenine dinucleotide synthetase 1 | 1809.61 | 1589.42 | 0.19 | 3.96E-01 |
| ENSMUSG00000028044 | GSMG0025655 | Cks1b | CDC28 protein kinase 1b | 331.95 | 785.57 | -1.24 | 9.74E-11 |

| | | | | | | | |
|---|---|---|---|---|---|---|---|
| ENSMUSG00000060671 | GSMG0025658 | Atp8b2 | ATPase, class I, type 8B, member 2 | 2316.49 | 1752.24 | 0.40 | 1.10E-01 |
| ENSMUSG00000042520 | GSMG0025660 | Ubap2l | ubiquitin-associated protein 2-like | 4683.88 | 5631.56 | -0.27 | 1.36E-01 |
| ENSMUSG00000027932 | GSMG0025665 | Slc27a3 | solute carrier family 27 (fatty acid transporter), member 3 | 121.04 | 284.39 | -1.24 | 3.48E-08 |
| ENSMUSG00000027933 | GSMG0025666 | Ints3 | integrator complex subunit 3 | 756.84 | 867.71 | -0.20 | 3.86E-01 |
| ENSMUSG00000027931 | GSMG0025667 | Npr1 | natriuretic peptide receptor 1 | 460.72 | 378.80 | 0.28 | 3.29E-01 |
| ENSMUSG00000001018 | GSMG0025668 | Snapin | SNAP-associated protein | 2559.36 | 2197.30 | 0.22 | 2.33E-01 |
| ENSMUSG00000001017 | GSMG0025669 | Chtop | chromatin target of PRMT1 | 643.51 | 739.84 | -0.20 | 4.26E-01 |
| ENSMUSG00000044080 | GSMG0025670 | S100a1 | S100 calcium binding protein A1 | 792.76 | 589.09 | 0.43 | 2.50E-02 |
| ENSMUSG00000056071 | GSMG0025671 | S100a9 | S100 calcium binding protein A9 (calgranulin B) | 46.24 | 108.20 | -1.25 | 1.73E-03 |
| ENSMUSG00000043165 | GSMG0025673 | Lor | loricrin | 121.48 | 222.32 | -0.88 | 3.06E-03 |
| ENSMUSG00000028136 | GSMG0025714 | Snx27 | sorting nexin family member 27 | 588.10 | 632.28 | -0.10 | 7.25E-01 |
| ENSMUSG00000005779 | GSMG0025721 | Psmb4 | proteasome (prosome, macropain) subunit, beta type 4 | 1891.62 | 2284.33 | -0.27 | 1.49E-01 |
| NULL | GSMG0025725 | NULL | NULL | 232.08 | 274.17 | -0.24 | 4.15E-01 |
| ENSMUSG00000019338 | GSMG0025727 | Zfp687 | zinc finger protein 687 | 190.09 | 219.64 | -0.21 | 5.68E-01 |
| ENSMUSG00000005625 | GSMG0025728 | Psmd4 | proteasome (prosome, macropain) 26S subunit, non-ATPase, 4 | 3262.78 | 3504.63 | -0.10 | 6.45E-01 |
| ENSMUSG00000028126 | GSMG0025729 | Pip5k1a | phosphatidylinositol-4-phosphate 5-kinase, type 1 alpha | 628.96 | 628.50 | 0.00 | 9.92E-01 |
| NULL | GSMG0025730 | NULL | NULL | 270.42 | 355.48 | -0.39 | 8.83E-02 |
| ENSMUSG00000092607 | GSMG0025731 | Scnm1 | sodium channel modifier 1 | 401.89 | 355.61 | 0.17 | 5.36E-01 |
| ENSMUSG00000038766 | GSMG0025735 | Gabpb2 | GA repeat binding protein, beta 2 | 120.60 | 83.05 | 0.54 | 1.34E-01 |
| ENSMUSG00000053192 | GSMG0025736 | Mllt11 | myeloid/lymphoid or mixed-lineage leukemia%3B translocated to, 11 | 2053.06 | 4335.72 | -1.08 | 1.08E-10 |
| ENSMUSG00000015711 | GSMG0025739 | Prune | prune exopolyphosphatase | 779.32 | 881.66 | -0.18 | 4.86E-01 |
| ENSMUSG00000015697 | GSMG0025742 | Setdb1 | SET domain, bifurcated 1 | 272.86 | 324.45 | -0.25 | 3.86E-01 |
| ENSMUSG00000028108 | GSMG0025746 | Ecm1 | extracellular matrix protein 1 | 4113.56 | 3374.77 | 0.29 | 1.54E-01 |
| ENSMUSG00000028107 | GSMG0025747 | Tars2 | threonyl-tRNA synthetase 2, mitochondrial (putative) | 719.59 | 709.83 | 0.02 | 9.50E-01 |
| ENSMUSG00000028106 | GSMG0025748 | Rprd2 | regulation of nuclear pre-mRNA domain containing 2 | 216.53 | 263.42 | -0.29 | 3.24E-01 |
| ENSMUSG00000015748 | GSMG0025749 | Prpf3 | pre-mRNA processing factor 3 | 1116.07 | 1194.49 | -0.10 | 6.88E-01 |
| ENSMUSG00000054312 | GSMG0025750 | Mrps21 | mitochondrial ribosomal protein S21 | 1098.90 | 1235.55 | -0.17 | 5.82E-01 |
| ENSMUSG00000038550 | GSMG0025751 | Ciart | circadian associated repressor of transcription | 250.73 | 131.13 | 0.94 | 5.23E-05 |
| ENSMUSG00000038543 | GSMG0025752 | BC028528 | cDNA sequence BC028528 | 2244.09 | 2106.80 | 0.09 | 7.37E-01 |
| ENSMUSG00000038526 | GSMG0025753 | Car14 | carbonic anhydrase 14 | 526.79 | 344.56 | 0.62 | 2.96E-02 |
| ENSMUSG00000015745 | GSMG0025754 | Plekho1 | pleckstrin homology domain containing, family O member 1 | 292.75 | 309.68 | -0.09 | 8.15E-01 |
| ENSMUSG00000015747 | GSMG0025755 | Vps45 | vacuolar protein sorting 45 | 538.94 | 687.74 | -0.35 | 7.60E-02 |
| ENSMUSG00000015943 | GSMG0025759 | Bola1 | bolA-like 1 (E. coli) | 420.72 | 536.54 | -0.35 | 1.86E-01 |

| | | | | | | | |
|---|---|---|---|---|---|---|---|
| ENSMUSG00000074398 | GSMG0025771 | Gm15441 | predicted gene 15441 | 140.88 | 169.44 | -0.26 | 4.57E-01 |
| ENSMUSG00000028104 | GSMG0025772 | Polr3gl | polymerase (RNA) III (DNA directed) polypeptide G like | 180.49 | 219.60 | -0.28 | 3.60E-01 |
| ENSMUSG00000045327 | GSMG0025773 | 6330549D23Rik | RIKEN cDNA 6330549D23 gene | 213.98 | 283.43 | -0.40 | 1.28E-01 |
| ENSMUSG00000028100 | GSMG0025775 | Nudt17 | nudix (nucleoside diphosphate linked moiety X)-type motif 17 | 151.16 | 124.24 | 0.29 | 4.50E-01 |
| ENSMUSG00000028099 | GSMG0025776 | Polr3c | polymerase (RNA) III (DNA directed) polypeptide C | 349.37 | 356.93 | -0.03 | 9.30E-01 |
| ENSMUSG00000028096 | GSMG0025780 | Gpr89 | G protein-coupled receptor 89 | 564.32 | 641.24 | -0.18 | 4.64E-01 |
| ENSMUSG00000038256 | GSMG0025784 | Bcl9 | B cell CLL/lymphoma 9 | 619.00 | 969.90 | -0.65 | 1.81E-04 |
| ENSMUSG00000028089 | GSMG0025789 | Chd1l | chromodomain helicase DNA binding protein 1-like | 263.92 | 285.25 | -0.11 | 7.40E-01 |
| ENSMUSG00000038170 | GSMG0025790 | Pde4dip | phosphodiesterase 4D interacting protein (myomegalin) | 526.60 | 581.88 | -0.14 | 6.64E-01 |
| NULL | GSMG0025791 | NULL | NULL | 510.14 | 386.47 | 0.40 | 1.08E-01 |
| ENSMUSG00000033285 | GSMG0025806 | Wdr3 | WD repeat domain 3 | 1154.94 | 1431.54 | -0.31 | 7.44E-02 |
| ENSMUSG00000044468 | GSMG0025807 | Fam46c | family with sequence similarity 46, member C | 829.96 | 795.23 | 0.06 | 8.24E-01 |
| ENSMUSG00000008763 | GSMG0025808 | Man1a2 | mannosidase, alpha, class 1A, member 2 | 828.46 | 870.35 | -0.07 | 8.08E-01 |
| ENSMUSG00000033222 | GSMG0025813 | Ttf2 | transcription termination factor, RNA polymerase II | 356.73 | 548.74 | -0.62 | 6.64E-04 |
| ENSMUSG00000033161 | GSMG0025819 | Atp1a1 | ATPase, Na+/K+ transporting, alpha 1 polypeptide | 5412.62 | 4947.70 | 0.13 | 7.15E-01 |
| ENSMUSG00000027860 | GSMG0025825 | Vangl1 | vang-like 1 (van gogh, Drosophila) | 501.37 | 463.60 | 0.11 | 6.94E-01 |
| ENSMUSG00000027848 | GSMG0025833 | Olfml3 | olfactomedin-like 3 | 635.12 | 619.14 | 0.04 | 9.16E-01 |
| ENSMUSG00000008730 | GSMG0025834 | Hipk1 | homeodomain interacting protein kinase 1 | 957.24 | 674.83 | 0.50 | 1.38E-01 |
| NULL | GSMG0025840 | NULL | NULL | 78.22 | 49.63 | 0.66 | 1.10E-01 |
| ENSMUSG00000052539 | GSMG0025841 | Magi3 | membrane associated guanylate kinase, WW and PDZ domain containi | 1193.70 | 848.79 | 0.49 | 1.35E-01 |
| ENSMUSG00000032913 | GSMG0025854 | Lrig2 | leucine-rich repeats and immunoglobulin-like domains 2 | 825.18 | 701.78 | 0.23 | 3.74E-01 |
| NULL | GSMG0025855 | NULL | NULL | 715.31 | 776.37 | -0.12 | 6.31E-01 |
| NULL | GSMG0025856 | NULL | NULL | 621.49 | 610.67 | 0.02 | 9.49E-01 |
| NULL | GSMG0025857 | NULL | NULL | 177.40 | 126.37 | 0.49 | 2.42E-01 |
| ENSMUSG00000002227 | GSMG0025859 | Mov10 | Moloney leukemia virus 10 | 1116.17 | 1200.69 | -0.11 | 6.69E-01 |
| NULL | GSMG0025861 | NULL | NULL | 186.86 | 208.62 | -0.17 | 6.79E-01 |
| ENSMUSG00000062127 | GSMG0025864 | Cttnbp2nl | CTTNBP2 N-terminal like | 4506.48 | 3584.88 | 0.33 | 1.50E-01 |
| ENSMUSG00000027905 | GSMG0025866 | Ddx20 | DEAD (Asp-Glu-Ala-Asp) box polypeptide 20 | 396.86 | 588.77 | -0.57 | 1.37E-03 |
| ENSMUSG00000068798 | GSMG0025868 | Rap1a | RAS-related protein-1a | 851.45 | 888.06 | -0.06 | 8.57E-01 |
| ENSMUSG00000000563 | GSMG0025874 | Atp5f1 | ATP synthase, H+ transporting, mitochondrial F0 complex, subunit B1 | 2377.37 | 2759.40 | -0.21 | 2.89E-01 |
| ENSMUSG00000040774 | GSMG0025882 | Cept1 | choline/ethanolaminephosphotransferase 1 | 1073.19 | 804.24 | 0.42 | 3.81E-02 |
| ENSMUSG00000040747 | GSMG0025888 | Cd53 | CD53 antigen | 109.40 | 49.06 | 1.17 | 3.96E-03 |
| ENSMUSG00000027895 | GSMG0025897 | Kcnc4 | potassium voltage gated channel, Shaw-related subfamily, member 4 | 231.30 | 107.90 | 1.10 | 4.52E-04 |

| | | | | | | | |
|---|---|---|---|---|---|---|---|
| ENSMUSG00000027894 | GSMG0025900 | Slc6a17 | solute carrier family 6 (neurotransmitter transporter), member 17 | 119.05 | 106.01 | 0.17 | 7.47E-01 |
| ENSMUSG00000014601 | GSMG0025902 | Strip1 | striatin interacting protein 1 | 4508.51 | 5478.87 | -0.28 | 8.44E-02 |
| ENSMUSG00000027893 | GSMG0025905 | Ahcyl1 | S-adenosylhomocysteine hydrolase-like 1 | 4148.03 | 3603.13 | 0.20 | 3.80E-01 |
| ENSMUSG00000014599 | GSMG0025907 | Csf1 | colony stimulating factor 1 (macrophage) | 1059.26 | 1077.52 | -0.03 | 9.47E-01 |
| ENSMUSG00000086968 | GSMG0025910 | 4933431E20Rik | RIKEN cDNA 4933431E20 gene | 1608.83 | 1683.88 | -0.07 | 8.32E-01 |
| ENSMUSG00000004035 | GSMG0025911 | Gstm7 | glutathione S-transferase, mu 7 | 1336.83 | 1226.99 | 0.12 | 6.39E-01 |
| ENSMUSG00000040562 | GSMG0025914 | Gstm2 | glutathione S-transferase, mu 2 | 208.46 | 102.07 | 1.03 | 4.65E-04 |
| ENSMUSG00000058135 | GSMG0025915 | Gstm1 | glutathione S-transferase, mu 1 | 2496.70 | 1493.80 | 0.74 | 5.01E-07 |
| ENSMUSG00000027890 | GSMG0025917 | Gstm4 | glutathione S-transferase, mu 4 | 88.04 | 103.60 | -0.24 | 5.94E-01 |
| ENSMUSG00000027889 | GSMG0025918 | Ampd2 | adenosine monophosphate deaminase 2 | 1922.34 | 2027.98 | -0.08 | 8.09E-01 |
| ENSMUSG00000000001 | GSMG0025919 | Gnai3 | guanine nucleotide binding protein (G protein), alpha inhibiting 3 | 1368.47 | 1735.45 | -0.34 | 7.73E-02 |
| ENSMUSG00000048796 | GSMG0025922 | Cyb561d1 | cytochrome b-561 domain containing 1 | 652.24 | 632.29 | 0.04 | 9.01E-01 |
| ENSMUSG00000048997 | GSMG0025923 | Atxn7l2 | ataxin 7-like 2 | 739.53 | 889.62 | -0.27 | 2.17E-01 |
| ENSMUSG00000068740 | GSMG0025928 | Celsr2 | cadherin, EGF LAG seven-pass G-type receptor 2 | 1460.58 | 2052.89 | -0.49 | 4.59E-02 |
| ENSMUSG00000068739 | GSMG0025929 | Sars | seryl-aminoacyl-tRNA synthetase | 3432.00 | 3413.81 | 0.01 | 9.77E-01 |
| ENSMUSG00000040412 | GSMG0025930 | 5330417C22Rik | RIKEN cDNA 5330417C22 gene | 320.96 | 370.09 | -0.21 | 4.84E-01 |
| NULL | GSMG0025931 | NULL | NULL | 266.14 | 171.44 | 0.64 | 5.59E-03 |
| ENSMUSG00000068732 | GSMG0025932 | Tmem167b | transmembrane protein 167B | 1687.76 | 1549.23 | 0.12 | 5.87E-01 |
| ENSMUSG00000027883 | GSMG0025933 | Gpsm2 | G-protein signalling modulator 2 (AGS3-like, C. elegans) | 543.05 | 844.44 | -0.63 | 6.03E-04 |
| ENSMUSG00000027882 | GSMG0025935 | Stxbp3 | syntaxin binding protein 3 | 788.63 | 635.57 | 0.31 | 1.62E-01 |
| ENSMUSG00000027881 | GSMG0025939 | Prpf38b | PRP38 pre-mRNA processing factor 38 (yeast) domain containing B | 1612.97 | 1664.97 | -0.05 | 8.71E-01 |
| ENSMUSG00000040339 | GSMG0025943 | Fam102b | family with sequence similarity 102, member B | 193.91 | 409.30 | -1.08 | 4.26E-05 |
| ENSMUSG00000059857 | GSMG0025947 | Ntng1 | netrin G1 | 108.32 | 102.93 | 0.08 | 9.00E-01 |
| ENSMUSG00000049300 | GSMG0025952 | Prmt6 | protein arginine N-methyltransferase 6 | 486.92 | 607.17 | -0.32 | 1.31E-01 |
| ENSMUSG00000045092 | GSMG0025962 | S1pr1 | sphingosine-1-phosphate receptor 1 | 12870.51 | 7341.54 | 0.81 | 1.40E-05 |
| ENSMUSG00000054426 | GSMG0025963 | A930005H10Rik | RIKEN cDNA A930005H10 gene | 267.70 | 324.55 | -0.28 | 2.85E-01 |
| ENSMUSG00000054414 | GSMG0025966 | Slc30a7 | solute carrier family 30 (zinc transporter), member 7 | 452.97 | 486.12 | -0.10 | 7.23E-01 |
| ENSMUSG00000027962 | GSMG0025967 | Vcam1 | vascular cell adhesion molecule 1 | 2481.05 | 998.76 | 1.31 | 3.31E-11 |
| ENSMUSG00000068696 | GSMG0025968 | Gpr88 | G-protein coupled receptor 88 | 613.90 | 530.28 | 0.21 | 4.70E-01 |
| ENSMUSG00000033502 | GSMG0025969 | Cdc14a | CDC14 cell division cycle 14A | 242.49 | 180.74 | 0.42 | 1.77E-01 |
| ENSMUSG00000000339 | GSMG0025970 | Rtca | RNA 3'-terminal phosphate cyclase | 1304.02 | 1578.19 | -0.28 | 1.44E-01 |
| ENSMUSG00000089911 | GSMG0025973 | Mfsd14a | major facilitator superfamily domain containing 14A | 2716.77 | 2223.09 | 0.29 | 1.01E-01 |
| ENSMUSG00000027957 | GSMG0025974 | Slc35a3 | solute carrier family 35 (UDP-N-acetylglucosamine (UDP-GlcNAc) tra | 386.80 | 387.32 | 0.00 | 9.98E-01 |

| | | | | | | | |
|---|---|---|---|---|---|---|---|
| ENSMUSG00000033400 | GSMG0025975 | Agl | amylo-1,6-glucosidase, 4-alpha-glucanotransferase | 782.12 | 794.23 | -0.02 | 9.50E-01 |
| NULL | GSMG0025977 | NULL | NULL | 798.24 | 518.29 | 0.62 | 1.41E-03 |
| ENSMUSG00000033377 | GSMG0025978 | Palmd | palmdelphin | 4469.59 | 4117.46 | 0.12 | 6.80E-01 |
| ENSMUSG00000044667 | GSMG0025980 | Plppr4 | phospholipid phosphatase related 4 | 322.74 | 381.29 | -0.24 | 5.18E-01 |
| NULL | GSMG0025981 | NULL | NULL | 1357.85 | 1084.15 | 0.32 | 1.81E-01 |
| ENSMUSG00000028007 | GSMG0025982 | Snx7 | sorting nexin 7 | 665.91 | 729.55 | -0.13 | 6.55E-01 |
| ENSMUSG00000028134 | GSMG0025983 | Ptbp2 | polypyrimidine tract binding protein 2 | 120.78 | 221.89 | -0.87 | 1.79E-03 |
| ENSMUSG00000028133 | GSMG0025989 | Rwdd3 | RWD domain containing 3 | 105.88 | 132.70 | -0.33 | 3.62E-01 |
| ENSMUSG00000028127 | GSMG0025996 | Abcd3 | ATP-binding cassette, sub-family D (ALD), member 3 | 1360.75 | 1503.44 | -0.14 | 5.28E-01 |
| ENSMUSG00000039701 | GSMG0026007 | Usp53 | ubiquitin specific peptidase 53 | 363.89 | 288.80 | 0.33 | 1.90E-01 |
| ENSMUSG00000050315 | GSMG0026009 | Synpo2 | synaptopodin 2 | 312.18 | 226.42 | 0.46 | 5.16E-02 |
| NULL | GSMG0026010 | NULL | NULL | 124.22 | 90.26 | 0.45 | 2.18E-01 |
| ENSMUSG00000028114 | GSMG0026014 | Mettl14 | methyltransferase like 14 | 211.00 | 241.93 | -0.20 | 5.37E-01 |
| NULL | GSMG0026015 | NULL | NULL | 76.09 | 80.39 | -0.09 | 8.83E-01 |
| ENSMUSG00000027977 | GSMG0026016 | Ndst3 | N-deacetylase/N-sulfotransferase (heparan glucosaminyl) 3 | 255.20 | 182.54 | 0.48 | 6.61E-02 |
| ENSMUSG00000032854 | GSMG0026022 | Ugt8a | UDP galactosyltransferase 8A | 176.83 | 42.95 | 2.04 | 5.08E-10 |
| ENSMUSG00000032826 | GSMG0026025 | Ank2 | ankyrin 2, brain | 7449.79 | 9622.51 | -0.37 | 2.74E-02 |
| NULL | GSMG0026026 | NULL | NULL | 452.70 | 368.50 | 0.30 | 2.26E-01 |
| NULL | GSMG0026028 | NULL | NULL | 407.60 | 456.21 | -0.16 | 6.04E-01 |
| NULL | GSMG0026032 | NULL | NULL | 211.28 | 318.56 | -0.59 | 4.57E-02 |
| ENSMUSG00000027968 | GSMG0026033 | Larp7 | La ribonucleoprotein domain family, member 7 | 360.74 | 456.35 | -0.34 | 2.36E-01 |
| NULL | GSMG0026034 | NULL | NULL | 369.90 | 578.58 | -0.64 | 1.09E-03 |
| NULL | GSMG0026035 | NULL | NULL | 287.49 | 398.57 | -0.47 | 1.01E-01 |
| NULL | GSMG0026036 | NULL | NULL | 630.15 | 722.98 | -0.20 | 4.23E-01 |
| NULL | GSMG0026037 | NULL | NULL | 416.93 | 383.99 | 0.12 | 7.44E-01 |
| NULL | GSMG0026038 | NULL | NULL | 365.00 | 368.82 | -0.01 | 9.72E-01 |
| ENSMUSG00000028028 | GSMG0026040 | Alpk1 | alpha-kinase 1 | 100.58 | 51.55 | 0.96 | 7.38E-03 |
| ENSMUSG00000074238 | GSMG0026044 | Ap1ar | adaptor-related protein complex 1 associated regulatory protein | 1459.10 | 1681.97 | -0.21 | 3.22E-01 |
| ENSMUSG00000050549 | GSMG0026048 | 5730508B09Rik | RIKEN cDNA 5730508B09 gene | 495.71 | 465.42 | 0.09 | 8.24E-01 |
| ENSMUSG00000028024 | GSMG0026052 | Enpep | glutamyl aminopeptidase | 1586.13 | 891.85 | 0.83 | 7.33E-06 |
| ENSMUSG00000028010 | GSMG0026057 | Gar1 | GAR1 ribonucleoprotein | 208.55 | 323.95 | -0.64 | 9.00E-03 |
| ENSMUSG00000027994 | GSMG0026058 | Ccdc109b | coiled-coil domain containing 109B | 88.91 | 31.17 | 1.52 | 1.18E-04 |
| ENSMUSG00000001052 | GSMG0026059 | Sec24b | Sec24 related gene family, member B (S. cerevisiae) | 1286.50 | 1419.06 | -0.14 | 5.90E-01 |

| | | | | | | | |
|---|---|---|---|---|---|---|---|
| NULL | GSMG0026061 | NULL | NULL | 409.72 | 398.23 | 0.04 | 9.08E-01 |
| ENSMUSG00000041084 | GSMG0026062 | Ostc | oligosaccharyltransferase complex subunit (non-catalytic) | 2717.52 | 3652.42 | -0.43 | 1.85E-02 |
| NULL | GSMG0026067 | NULL | NULL | 1227.59 | 841.41 | 0.54 | 5.20E-03 |
| ENSMUSG00000027984 | GSMG0026068 | Hadh | hydroxyacyl-Coenzyme A dehydrogenase | 1472.24 | 1566.11 | -0.09 | 7.64E-01 |
| ENSMUSG00000027983 | GSMG0026070 | Cyp2u1 | cytochrome P450, family 2, subfamily u, polypeptide 1 | 66.92 | 51.78 | 0.37 | 4.89E-01 |
| ENSMUSG00000050931 | GSMG0026072 | Sgms2 | sphingomyelin synthase 2 | 445.87 | 220.77 | 1.01 | 8.18E-08 |
| ENSMUSG00000028029 | GSMG0026076 | Aimp1 | aminoacyl tRNA synthetase complex-interacting multifunctional protei | 526.67 | 638.58 | -0.28 | 2.60E-01 |
| ENSMUSG00000040998 | GSMG0026079 | Npnt | nephronectin | 1061.36 | 563.58 | 0.91 | 1.10E-04 |
| ENSMUSG00000040943 | GSMG0026088 | Tet2 | tet methylcytosine dioxygenase 2 | 3089.20 | 2793.17 | 0.15 | 5.46E-01 |
| ENSMUSG00000028165 | GSMG0026094 | Cisd2 | CDGSH iron sulfur domain 2 | 1858.28 | 1825.27 | 0.03 | 9.38E-01 |
| ENSMUSG00000028163 | GSMG0026099 | Nfkb1 | nuclear factor of kappa light polypeptide gene enhancer in B cells 1, p1 | 739.53 | 1000.62 | -0.43 | 1.84E-02 |
| ENSMUSG00000005813 | GSMG0026116 | Metap1 | methionyl aminopeptidase 1 | 403.91 | 485.26 | -0.26 | 2.58E-01 |
| NULL | GSMG0026118 | NULL | NULL | 768.66 | 740.81 | 0.05 | 8.70E-01 |
| ENSMUSG00000028149 | GSMG0026119 | Rap1gds1 | RAP1, GTP-GDP dissociation stimulator 1 | 2238.53 | 2081.80 | 0.10 | 6.63E-01 |
| NULL | GSMG0026123 | NULL | NULL | 263.33 | 205.34 | 0.36 | 3.46E-01 |
| ENSMUSG00000028273 | GSMG0026134 | Pdlim5 | PDZ and LIM domain 5 | 1415.06 | 1328.24 | 0.09 | 7.53E-01 |
| ENSMUSG00000004591 | GSMG0026139 | Pkn2 | protein kinase N2 | 1008.66 | 985.36 | 0.03 | 9.17E-01 |
| ENSMUSG00000028266 | GSMG0026145 | Lmo4 | LIM domain only 4 | 1318.18 | 1518.44 | -0.21 | 3.80E-01 |
| NULL | GSMG0026146 | NULL | NULL | 234.26 | 206.24 | 0.18 | 6.82E-01 |
| ENSMUSG00000040151 | GSMG0026148 | Hs2st1 | heparan sulfate 2-O-sulfotransferase 1 | 415.37 | 476.84 | -0.20 | 4.60E-01 |
| NULL | GSMG0026150 | NULL | NULL | 218.22 | 135.15 | 0.70 | 1.12E-02 |
| ENSMUSG00000037062 | GSMG0026152 | Sh3glb1 | SH3-domain GRB2-like B1 (endophilin) | 3695.52 | 4131.29 | -0.16 | 4.00E-01 |
| ENSMUSG00000028195 | GSMG0026162 | Cyr61 | cysteine rich protein 61 | 2075.40 | 671.73 | 1.63 | 9.96E-16 |
| ENSMUSG00000028187 | GSMG0026167 | Rpf1 | ribosome production factor 1 homolog | 614.63 | 682.57 | -0.15 | 5.97E-01 |
| ENSMUSG00000005034 | GSMG0026171 | Prkacb | protein kinase, cAMP dependent, catalytic, beta | 1096.04 | 1147.26 | -0.07 | 8.07E-01 |
| ENSMUSG00000028184 | GSMG0026174 | Adgrl2 | adhesion G protein-coupled receptor L2 | 3125.16 | 4288.54 | -0.46 | 4.17E-03 |
| NULL | GSMG0026176 | NULL | NULL | 128.42 | 168.04 | -0.38 | 2.84E-01 |
| NULL | GSMG0026179 | NULL | NULL | 80.97 | 295.32 | -1.87 | 4.27E-15 |
| NULL | GSMG0026183 | NULL | NULL | 96.36 | 260.45 | -1.44 | 1.19E-08 |
| ENSMUSG00000028037 | GSMG0026188 | Ifi44 | interferon-induced protein 44 | 62.59 | 106.66 | -0.78 | 4.44E-02 |
| ENSMUSG00000054942 | GSMG0026194 | Fam73a | family with sequence similarity 73, member A | 581.90 | 675.42 | -0.22 | 3.33E-01 |
| ENSMUSG00000039058 | GSMG0026199 | Ak5 | adenylate kinase 5 | 735.09 | 521.40 | 0.50 | 1.34E-02 |
| NULL | GSMG0026201 | NULL | NULL | 206.45 | 343.37 | -0.73 | 9.56E-04 |

| | | | | | | | |
|---|---|---|---|---|---|---|---|
| ENSMUSG00000039037 | GSMG0026202 | St6galnac5 | ST6 (alpha-N-acetyl-neuraminyl-2,3-beta-galactosyl-1,3)-N-acetylgalac | 359.65 | 607.49 | -0.76 | 3.46E-05 |
| NULL | GSMG0026203 | NULL | NULL | 274.64 | 272.28 | 0.01 | 9.71E-01 |
| ENSMUSG00000052544 | GSMG0026207 | St6galnac3 | ST6 (alpha-N-acetyl-neuraminyl-2,3-beta-galactosyl-1,3)-N-acetylgalac | 458.72 | 340.79 | 0.43 | 4.11E-02 |
| NULL | GSMG0026210 | NULL | NULL | 239.57 | 251.86 | -0.07 | 8.72E-01 |
| ENSMUSG00000038975 | GSMG0026216 | Rabggtb | Rab geranylgeranyl transferase, b subunit | 489.45 | 487.92 | 0.00 | 9.89E-01 |
| ENSMUSG00000062908 | GSMG0026217 | Acadm | acyl-Coenzyme A dehydrogenase, medium chain | 1626.40 | 1521.90 | 0.10 | 7.65E-01 |
| ENSMUSG00000047583 | GSMG0026220 | Tyw3 | tRNA-yW synthesizing protein 3 homolog (S. cerevisiae) | 351.74 | 418.68 | -0.25 | 3.01E-01 |
| NULL | GSMG0026221 | NULL | NULL | 57.89 | 77.49 | -0.42 | 3.26E-01 |
| ENSMUSG00000053870 | GSMG0026225 | Fpgt | fucose-1-phosphate guanylyltransferase | 349.97 | 262.51 | 0.41 | 6.65E-02 |
| ENSMUSG00000055436 | GSMG0026234 | Srsf11 | serine/arginine-rich splicing factor 11 | 7010.44 | 7376.04 | -0.07 | 7.61E-01 |
| NULL | GSMG0026237 | NULL | NULL | 757.63 | 654.75 | 0.21 | 3.59E-01 |
| NULL | GSMG0026238 | NULL | NULL | 625.11 | 483.08 | 0.37 | 1.28E-01 |
| ENSMUSG00000028176 | GSMG0026239 | Lrrc7 | leucine rich repeat containing 7 | 722.88 | 665.20 | 0.12 | 6.39E-01 |
| NULL | GSMG0026241 | NULL | NULL | 420.83 | 413.83 | 0.02 | 9.61E-01 |
| NULL | GSMG0026242 | NULL | NULL | 224.41 | 155.97 | 0.53 | 5.10E-02 |
| NULL | GSMG0026243 | NULL | NULL | 379.76 | 267.59 | 0.51 | 2.40E-02 |
| ENSMUSG00000028233 | GSMG0026244 | Tgs1 | trimethylguanosine synthase 1 | 1094.42 | 1461.52 | -0.42 | 3.82E-02 |
| ENSMUSG00000042228 | GSMG0026247 | Lyn | LYN proto-oncogene, Src family tyrosine kinase | 2063.13 | 2601.77 | -0.33 | 7.49E-02 |
| ENSMUSG00000042198 | GSMG0026249 | Chchd7 | coiled-coil-helix-coiled-coil-helix domain containing 7 | 578.37 | 641.72 | -0.15 | 6.57E-01 |
| ENSMUSG00000049119 | GSMG0026255 | Fam110b | family with sequence similarity 110, member B | 777.04 | 1273.49 | -0.71 | 9.75E-06 |
| ENSMUSG00000028243 | GSMG0026258 | Ubxn2b | UBX domain protein 2B | 172.73 | 216.55 | -0.32 | 3.28E-01 |
| ENSMUSG00000028249 | GSMG0026259 | Sdcbp | syndecan binding protein | 5476.28 | 5860.96 | -0.10 | 6.88E-01 |
| ENSMUSG00000047187 | GSMG0026263 | Rab2a | RAB2A, member RAS oncogene family | 7463.12 | 8283.18 | -0.15 | 5.29E-01 |
| ENSMUSG00000041216 | GSMG0026273 | Clvs1 | clavesin 1 | 62.07 | 89.12 | -0.52 | 1.79E-01 |
| ENSMUSG00000028211 | GSMG0026279 | Trp53inp1 | transformation related protein 53 inducible nuclear protein 1 | 223.93 | 136.11 | 0.72 | 4.09E-03 |
| ENSMUSG00000028212 | GSMG0026280 | Ccne2 | cyclin E2 | 109.81 | 241.28 | -1.14 | 4.58E-04 |
| ENSMUSG00000040720 | GSMG0026283 | 1110037F02Rik | RIKEN cDNA 1110037F02 gene | 1479.96 | 1526.00 | -0.04 | 8.70E-01 |
| ENSMUSG00000028214 | GSMG0026285 | Gem | GTP binding protein (gene overexpressed in skeletal muscle) | 127.64 | 63.41 | 1.01 | 1.01E-03 |
| ENSMUSG00000055963 | GSMG0026293 | Triqk | triple QxxK/R motif containing | 197.47 | 241.29 | -0.29 | 4.27E-01 |
| ENSMUSG00000006586 | GSMG0026296 | Runx1t1 | runt-related transcription factor 1%3B translocated to, 1 (cyclin D-relat | 294.60 | 641.29 | -1.12 | 1.31E-07 |
| NULL | GSMG0026298 | NULL | NULL | 58.72 | 92.24 | -0.66 | 1.50E-01 |
| NULL | GSMG0026299 | NULL | NULL | 37.78 | 129.64 | -1.80 | 6.59E-08 |
| ENSMUSG00000028221 | GSMG0026300 | Tmem55a | transmembrane protein 55A | 279.62 | 334.27 | -0.26 | 3.39E-01 |

| | | | | | | | |
|---|---|---|---|---|---|---|---|
| NULL | GSMG0026303 | NULL | NULL | 355.83 | 493.10 | -0.48 | 1.46E-01 |
| ENSMUSG00000043252 | GSMG0026305 | Tmem64 | transmembrane protein 64 | 2690.22 | 1596.62 | 0.75 | 2.17E-06 |
| ENSMUSG00000028222 | GSMG0026306 | Calb1 | calbindin 1 | 282.39 | 101.17 | 1.48 | 1.51E-06 |
| ENSMUSG00000028224 | GSMG0026308 | Nbn | nibrin | 568.48 | 626.27 | -0.14 | 7.11E-01 |
| ENSMUSG00000028226 | GSMG0026312 | Mmp16 | matrix metallopeptidase 16 | 505.20 | 779.01 | -0.62 | 8.91E-04 |
| NULL | GSMG0026313 | NULL | NULL | 945.94 | 1223.21 | -0.37 | 2.81E-02 |
| ENSMUSG00000028229 | GSMG0026320 | Rmdn1 | regulator of microtubule dynamics 1 | 510.46 | 475.17 | 0.11 | 7.11E-01 |
| ENSMUSG00000073987 | GSMG0026325 | Ggh | gamma-glutamyl hydrolase | 447.29 | 473.06 | -0.08 | 8.44E-01 |
| ENSMUSG00000028252 | GSMG0026328 | Ccnc | cyclin C | 340.81 | 361.87 | -0.09 | 8.08E-01 |
| ENSMUSG00000040455 | GSMG0026330 | Usp45 | ubiquitin specific petidase 45 | 187.97 | 203.53 | -0.11 | 7.74E-01 |
| ENSMUSG00000028248 | GSMG0026334 | Pnisr | PNN interacting serine/arginine-rich | 6211.11 | 5617.99 | 0.15 | 5.29E-01 |
| ENSMUSG00000028247 | GSMG0026336 | Coq3 | coenzyme Q3 methyltransferase | 190.84 | 211.48 | -0.15 | 7.33E-01 |
| ENSMUSG00000028246 | GSMG0026339 | Faxc | failed axon connections homolog | 518.91 | 897.12 | -0.79 | 1.84E-06 |
| ENSMUSG00000040410 | GSMG0026341 | Fbxl4 | F-box and leucine-rich repeat protein 4 | 525.08 | 502.48 | 0.06 | 8.31E-01 |
| NULL | GSMG0026343 | NULL | NULL | 28.60 | 90.31 | -1.65 | 1.33E-05 |
| ENSMUSG00000045751 | GSMG0026345 | Mms22l | MMS22-like, DNA repair protein | 169.76 | 368.60 | -1.11 | 8.02E-07 |
| ENSMUSG00000028261 | GSMG0026349 | Ndufaf4 | NADH dehydrogenase (ubiquinone) 1 alpha subcomplex, assembly fact | 1037.04 | 996.90 | 0.06 | 8.93E-01 |
| ENSMUSG00000028289 | GSMG0026356 | Epha7 | Eph receptor A7 | 656.06 | 1153.33 | -0.81 | 6.56E-07 |
| NULL | GSMG0026357 | NULL | NULL | 198.37 | 205.83 | -0.06 | 9.12E-01 |
| ENSMUSG00000028284 | GSMG0026362 | Map3k7 | mitogen-activated protein kinase kinase kinase 7 | 689.32 | 722.19 | -0.07 | 8.49E-01 |
| NULL | GSMG0026374 | NULL | NULL | 404.53 | 750.10 | -0.90 | 2.91E-04 |
| NULL | GSMG0026375 | NULL | NULL | 159.04 | 259.01 | -0.71 | 2.18E-02 |
| NULL | GSMG0026376 | NULL | NULL | 154.78 | 274.85 | -0.83 | 4.23E-03 |
| ENSMUSG00000028282 | GSMG0026379 | Casp8ap2 | caspase 8 associated protein 2 | 106.08 | 128.26 | -0.27 | 4.60E-01 |
| ENSMUSG00000045854 | GSMG0026385 | Lyrm2 | LYR motif containing 2 | 777.71 | 830.32 | -0.09 | 7.63E-01 |
| ENSMUSG00000028278 | GSMG0026386 | Rragd | Ras-related GTP binding D | 134.70 | 234.01 | -0.79 | 1.45E-03 |
| ENSMUSG00000028277 | GSMG0026389 | Ube2j1 | ubiquitin-conjugating enzyme E2J 1 | 644.30 | 819.90 | -0.35 | 7.18E-02 |
| ENSMUSG00000054679 | GSMG0026393 | Srsf12 | serine/arginine-rich splicing factor 12 | 455.75 | 869.72 | -0.94 | 6.39E-08 |
| NULL | GSMG0026394 | NULL | NULL | 104.15 | 101.04 | 0.04 | 9.54E-01 |
| ENSMUSG00000028274 | GSMG0026395 | Rngtt | RNA guanylyltransferase and 5'-phosphatase | 442.43 | 574.40 | -0.38 | 8.50E-02 |
| ENSMUSG00000044288 | GSMG0026404 | Cnr1 | cannabinoid receptor 1 (brain) | 736.28 | 1403.24 | -0.93 | 1.21E-09 |
| ENSMUSG00000028291 | GSMG0026405 | Akirin2 | akirin 2 | 425.05 | 524.75 | -0.31 | 1.65E-01 |
| ENSMUSG00000028292 | GSMG0026409 | Rars2 | arginyl-tRNA synthetase 2, mitochondrial | 755.06 | 868.83 | -0.20 | 4.38E-01 |

| | | | | | | | |
|---|---|---|---|---|---|---|---|
| ENSMUSG00000028405 | GSMG0026419 | Aco1 | aconitase 1 | 1965.47 | 2595.35 | -0.40 | 9.32E-03 |
| ENSMUSG00000028407 | GSMG0026420 | Smim27 | small integral membrane protein 27 | 264.23 | 251.31 | 0.07 | 8.92E-01 |
| ENSMUSG00000028419 | GSMG0026427 | Chmp5 | charged multivesicular body protein 5 | 1241.50 | 1153.65 | 0.11 | 7.33E-01 |
| ENSMUSG00000028423 | GSMG0026428 | Nfx1 | nuclear transcription factor, X-box binding 1 | 2690.14 | 3021.67 | -0.17 | 4.04E-01 |
| ENSMUSG00000036241 | GSMG0026430 | Ube2r2 | ubiquitin-conjugating enzyme E2R 2 | 1781.06 | 1809.82 | -0.02 | 9.33E-01 |
| ENSMUSG00000028437 | GSMG0026436 | Ubap1 | ubiquitin-associated protein 1 | 718.73 | 761.67 | -0.08 | 7.59E-01 |
| ENSMUSG00000028443 | GSMG0026442 | Nudt2 | nudix (nucleoside diphosphate linked moiety X)-type motif 2 | 251.95 | 440.20 | -0.81 | 2.31E-04 |
| ENSMUSG00000036073 | GSMG0026447 | Galt | galactose-1-phosphate uridyl transferase | 1127.57 | 942.26 | 0.26 | 1.87E-01 |
| ENSMUSG00000036062 | GSMG0026457 | Phf24 | PHD finger protein 24 | 464.51 | 414.28 | 0.17 | 6.11E-01 |
| ENSMUSG00000036052 | GSMG0026458 | Dnajb5 | DnaJ heat shock protein family (Hsp40) member B5 | 1445.04 | 1252.46 | 0.21 | 4.66E-01 |
| ENSMUSG00000028456 | GSMG0026463 | Unc13b | unc-13 homolog B (C. elegans) | 2963.90 | 2387.37 | 0.31 | 2.35E-01 |
| NULL | GSMG0026467 | NULL | NULL | 475.21 | 376.36 | 0.33 | 1.98E-01 |
| ENSMUSG00000035969 | GSMG0026468 | Rusc2 | RUN and SH3 domain containing 2 | 1936.12 | 1829.15 | 0.08 | 8.26E-01 |
| ENSMUSG00000028458 | GSMG0026469 | Tesk1 | testis specific protein kinase 1 | 423.80 | 303.44 | 0.48 | 5.88E-01 |
| ENSMUSG00000028461 | GSMG0026470 | Ccdc107 | coiled-coil domain containing 107 | 601.89 | 740.17 | -0.30 | 1.89E-01 |
| ENSMUSG00000028466 | GSMG0026472 | Creb3 | cAMP responsive element binding protein 3 | 1471.08 | 1310.72 | 0.17 | 4.39E-01 |
| ENSMUSG00000028468 | GSMG0026475 | Rgp1 | RAB6A GEF compex partner 1 | 199.63 | 157.97 | 0.34 | 2.93E-01 |
| ENSMUSG00000028469 | GSMG0026477 | Npr2 | natriuretic peptide receptor 2 | 911.75 | 637.25 | 0.52 | 5.61E-02 |
| ENSMUSG00000078716 | GSMG0026478 | Tmem8b | transmembrane protein 8B | 369.19 | 392.69 | -0.09 | 8.40E-01 |
| ENSMUSG00000071001 | GSMG0026479 | Hrct1 | histidine rich carboxyl terminus 1 | 652.83 | 505.77 | 0.37 | 6.73E-02 |
| ENSMUSG00000028475 | GSMG0026480 | Spaar | small regulatory polypeptide of amino acid response | 404.73 | 227.79 | 0.83 | 2.66E-05 |
| ENSMUSG00000028476 | GSMG0026483 | Reck | reversion-inducing-cysteine-rich protein with kazal motifs | 1309.24 | 847.13 | 0.63 | 9.95E-05 |
| ENSMUSG00000028480 | GSMG0026485 | Glipr2 | GLI pathogenesis-related 2 | 539.10 | 1081.52 | -1.00 | 4.32E-10 |
| ENSMUSG00000028478 | GSMG0026487 | Clta | clathrin, light polypeptide (Lca) | 2316.47 | 2730.50 | -0.24 | 2.29E-01 |
| ENSMUSG00000035683 | GSMG0026491 | Melk | maternal embryonic leucine zipper kinase | 180.39 | 436.22 | -1.27 | 1.10E-06 |
| ENSMUSG00000035649 | GSMG0026495 | Zcchc7 | zinc finger, CCHC domain containing 7 | 634.89 | 484.00 | 0.39 | 4.96E-02 |
| ENSMUSG00000035637 | GSMG0026506 | Grhpr | glyoxylate reductase/hydroxypyruvate reductase | 376.76 | 413.26 | -0.13 | 6.65E-01 |
| ENSMUSG00000028318 | GSMG0026508 | Polr1e | polymerase (RNA) I polypeptide E | 231.23 | 326.61 | -0.49 | 4.96E-02 |
| ENSMUSG00000035601 | GSMG0026510 | Trmt10b | tRNA methyltransferase 10B | 278.20 | 311.49 | -0.16 | 5.71E-01 |
| ENSMUSG00000035572 | GSMG0026513 | Dcaf10 | DDB1 and CUL4 associated factor 10 | 349.88 | 351.66 | -0.01 | 9.78E-01 |
| ENSMUSG00000035561 | GSMG0026517 | Aldh1b1 | aldehyde dehydrogenase 1 family, member B1 | 53.62 | 266.04 | -2.31 | 5.21E-21 |
| ENSMUSG00000035517 | GSMG0026521 | Tdrd7 | tudor domain containing 7 | 1775.06 | 2222.38 | -0.32 | 8.21E-02 |
| ENSMUSG00000028328 | GSMG0026523 | Tmod1 | tropomodulin 1 | 89.18 | 19.84 | 2.15 | 4.88E-07 |

| | | | | | | | |
|---|---|---|---|---|---|---|---|
| ENSMUSG00000028330 | GSMG0026525 | Ncbp1 | nuclear cap binding protein subunit 1 | 756.83 | 927.92 | -0.29 | 2.08E-01 |
| ENSMUSG00000028333 | GSMG0026530 | Anp32b | acidic (leucine-rich) nuclear phosphoprotein 32 family, member B | 1202.78 | 1687.51 | -0.49 | 9.84E-03 |
| ENSMUSG00000028334 | GSMG0026531 | Nans | N-acetylneuraminic acid synthase (sialic acid synthase) | 535.70 | 800.03 | -0.58 | 3.34E-03 |
| ENSMUSG00000028339 | GSMG0026538 | Col15a1 | collagen, type XV, alpha 1 | 106.72 | 75.24 | 0.50 | 3.38E-01 |
| ENSMUSG00000007613 | GSMG0026539 | Tgfbr1 | transforming growth factor, beta receptor I | 303.61 | 330.45 | -0.12 | 7.00E-01 |
| ENSMUSG00000053317 | GSMG0026541 | Sec61b | Sec61 beta subunit | 1072.76 | 1180.14 | -0.14 | 6.30E-01 |
| ENSMUSG00000061455 | GSMG0026544 | Stx17 | syntaxin 17 | 345.31 | 355.09 | -0.04 | 9.06E-01 |
| ENSMUSG00000028344 | GSMG0026545 | Invs | inversin | 269.21 | 314.47 | -0.22 | 4.83E-01 |
| ENSMUSG00000039693 | GSMG0026546 | Msantd3 | Myb/SANT-like DNA-binding domain containing 3 | 174.57 | 187.33 | -0.11 | 7.97E-01 |
| ENSMUSG00000028347 | GSMG0026549 | Tmeff1 | transmembrane protein with EGF-like and two follistatin-like domains | 610.85 | 575.31 | 0.09 | 7.91E-01 |
| ENSMUSG00000063446 | GSMG0026552 | Plppr1 | phospholipid phosphatase related 1 | 105.47 | 679.79 | -2.69 | 2.34E-45 |
| ENSMUSG00000039634 | GSMG0026553 | Zfp189 | zinc finger protein 189 | 356.92 | 398.03 | -0.16 | 6.39E-01 |
| ENSMUSG00000028309 | GSMG0026555 | Rnf20 | ring finger protein 20 | 1051.32 | 1148.98 | -0.13 | 5.93E-01 |
| ENSMUSG00000028312 | GSMG0026557 | Smc2 | structural maintenance of chromosomes 2 | 219.72 | 460.48 | -1.07 | 9.51E-06 |
| ENSMUSG00000015247 | GSMG0026563 | Nipsnap3b | nipsnap homolog 3B (C. elegans) | 438.72 | 535.95 | -0.29 | 2.68E-01 |
| ENSMUSG00000028412 | GSMG0026565 | Slc44a1 | solute carrier family 44, member 1 | 3331.57 | 2425.27 | 0.46 | 1.71E-02 |
| ENSMUSG00000028414 | GSMG0026573 | Fktn | fukutin | 354.43 | 368.29 | -0.05 | 8.91E-01 |
| ENSMUSG00000028420 | GSMG0026575 | Tmem38b | transmembrane protein 38B | 101.99 | 74.49 | 0.46 | 2.21E-01 |
| ENSMUSG00000028426 | GSMG0026584 | Rad23b | RAD23 homolog B, nucleotide excision repair protein | 357.82 | 509.24 | -0.51 | 4.89E-02 |
| ENSMUSG00000038827 | GSMG0026589 | Fam206a | family with sequence similarity 206, member A | 896.97 | 881.70 | 0.03 | 9.43E-01 |
| NULL | GSMG0026591 | NULL | NULL | 282.56 | 271.18 | 0.06 | 8.77E-01 |
| NULL | GSMG0026600 | NULL | NULL | 258.55 | 363.22 | -0.49 | 7.47E-02 |
| ENSMUSG00000070972 | GSMG0026605 | Dnajc25 | DnaJ heat shock protein family (Hsp40) member C25 | 170.63 | 159.57 | 0.10 | 8.24E-01 |
| ENSMUSG00000038607 | GSMG0026607 | Gng10 | guanine nucleotide binding protein (G protein), gamma 10 | 705.62 | 1090.42 | -0.63 | 1.75E-03 |
| ENSMUSG00000028381 | GSMG0026608 | Ugcg | UDP-glucose ceramide glucosyltransferase | 317.19 | 566.67 | -0.84 | 1.60E-06 |
| ENSMUSG00000028383 | GSMG0026612 | Hsdl2 | hydroxysteroid dehydrogenase like 2 | 534.38 | 586.43 | -0.13 | 6.64E-01 |
| ENSMUSG00000045071 | GSMG0026613 | E130308A19Rik | RIKEN cDNA E130308A19 gene | 258.26 | 353.99 | -0.45 | 6.80E-02 |
| ENSMUSG00000028385 | GSMG0026616 | Snx30 | sorting nexin family member 30 | 198.34 | 193.66 | 0.04 | 9.33E-01 |
| ENSMUSG00000066152 | GSMG0026619 | Slc31a2 | solute carrier family 31, member 2 | 236.02 | 219.47 | 0.10 | 7.83E-01 |
| ENSMUSG00000066150 | GSMG0026620 | Slc31a1 | solute carrier family 31, member 1 | 3702.21 | 3014.40 | 0.30 | 1.05E-01 |
| ENSMUSG00000066148 | GSMG0026622 | Prpf4 | pre-mRNA processing factor 4 | 740.64 | 863.97 | -0.22 | 3.71E-01 |
| ENSMUSG00000028392 | GSMG0026624 | Bspry | B-box and SPRY domain containing | 136.76 | 120.79 | 0.17 | 7.00E-01 |
| ENSMUSG00000059810 | GSMG0026627 | Rgs3 | regulator of G-protein signaling 3 | 804.91 | 813.26 | -0.02 | 9.61E-01 |

| | | | | | | | |
|---|---|---|---|---|---|---|---|
| ENSMUSG00000028358 | GSMG0026629 | Zfp618 | zinc finger protein 618 | 113.90 | 157.77 | -0.48 | 1.80E-01 |
| NULL | GSMG0026630 | NULL | NULL | 169.59 | 257.13 | -0.60 | 5.92E-01 |
| ENSMUSG00000045672 | GSMG0026635 | Col27a1 | collagen, type XXVII, alpha 1 | 123.78 | 71.16 | 0.80 | 3.57E-02 |
| ENSMUSG00000045917 | GSMG0026641 | 6330416G13Rik | RIKEN cDNA 6330416G13 gene | 99.59 | 94.19 | 0.09 | 8.83E-01 |
| NULL | GSMG0026645 | NULL | NULL | 106.81 | 509.50 | -2.24 | 1.86E-26 |
| ENSMUSG00000051675 | GSMG0026650 | Trim32 | tripartite motif-containing 32 | 1119.59 | 1497.87 | -0.42 | 2.42E-02 |
| NULL | GSMG0026660 | NULL | NULL | 321.59 | 303.99 | 0.08 | 8.66E-01 |
| ENSMUSG00000049122 | GSMG0026665 | Frmd3 | FERM domain containing 3 | 213.35 | 490.46 | -1.20 | 8.62E-10 |
| ENSMUSG00000028397 | GSMG0026666 | Kdm4c | lysine (K)-specific demethylase 4C | 773.85 | 666.44 | 0.22 | 3.40E-01 |
| NULL | GSMG0026672 | NULL | NULL | 200.38 | 268.97 | -0.43 | 2.12E-01 |
| ENSMUSG00000048706 | GSMG0026676 | Lurap1l | leucine rich adaptor protein 1-like | 1322.83 | 1435.28 | -0.12 | 6.89E-01 |
| ENSMUSG00000028483 | GSMG0026688 | Snapc3 | small nuclear RNA activating complex, polypeptide 3 | 599.05 | 559.51 | 0.10 | 7.53E-01 |
| NULL | GSMG0026690 | NULL | NULL | 236.27 | 289.17 | -0.29 | 2.71E-01 |
| ENSMUSG00000028488 | GSMG0026701 | Sh3gl2 | SH3-domain GRB2-like 2 | 450.82 | 485.78 | -0.11 | 7.26E-01 |
| ENSMUSG00000070934 | GSMG0026711 | Rraga | Ras-related GTP binding A | 1700.04 | 2068.59 | -0.28 | 1.72E-01 |
| ENSMUSG00000038024 | GSMG0026713 | Dennd4c | DENN/MADD domain containing 4C | 891.65 | 639.46 | 0.48 | 8.03E-03 |
| ENSMUSG00000038007 | GSMG0026717 | Acer2 | alkaline ceramidase 2 | 4564.31 | 567.34 | 3.01 | 6.72E-59 |
| ENSMUSG00000038368 | GSMG0026720 | Focad | focadhesin | 1233.01 | 900.55 | 0.45 | 8.05E-03 |
| ENSMUSG00000062937 | GSMG0026734 | Mtap | methylthioadenosine phosphorylase | 476.73 | 748.72 | -0.65 | 5.90E-04 |
| ENSMUSG00000028576 | GSMG0026744 | Ift74 | intraflagellar transport 74 | 264.41 | 311.34 | -0.24 | 5.35E-01 |
| ENSMUSG00000006386 | GSMG0026745 | Tek | endothelial-specific receptor tyrosine kinase | 15114.45 | 11667.74 | 0.37 | 4.27E-02 |
| ENSMUSG00000028573 | GSMG0026750 | Fggy | FGGY carbohydrate kinase domain containing | 136.85 | 14.40 | 3.25 | 1.06E-02 |
| ENSMUSG00000028572 | GSMG0026751 | Hook1 | hook microtubule tethering protein 1 | 786.01 | 894.23 | -0.19 | 4.12E-01 |
| ENSMUSG00000028565 | GSMG0026755 | Nfia | nuclear factor I/A | 1417.44 | 1531.84 | -0.11 | 6.34E-01 |
| NULL | GSMG0026757 | NULL | NULL | 269.73 | 374.56 | -0.47 | 1.36E-01 |
| ENSMUSG00000028560 | GSMG0026770 | Usp1 | ubiquitin specific peptidase 1 | 953.12 | 1485.36 | -0.64 | 1.24E-03 |
| ENSMUSG00000028550 | GSMG0026772 | Atg4c | autophagy related 4C, cysteine peptidase | 244.95 | 237.31 | 0.04 | 9.23E-01 |
| ENSMUSG00000073792 | GSMG0026777 | Alg6 | asparagine-linked glycosylation 6 (alpha-1,3,-glucosyltransferase) | 492.25 | 491.93 | 0.00 | 9.99E-01 |
| ENSMUSG00000025791 | GSMG0026780 | Pgm2 | phosphoglucomutase 2 | 222.80 | 265.98 | -0.26 | 3.75E-01 |
| NULL | GSMG0026781 | NULL | NULL | 143.77 | 84.32 | 0.77 | 9.05E-03 |
| ENSMUSG00000035305 | GSMG0026782 | Ror1 | receptor tyrosine kinase-like orphan receptor 1 | 199.81 | 141.97 | 0.50 | 1.25E-01 |
| ENSMUSG00000028532 | GSMG0026788 | Cachd1 | cache domain containing 1 | 569.32 | 676.31 | -0.25 | 4.09E-01 |
| ENSMUSG00000035275 | GSMG0026792 | Raver2 | ribonucleoprotein, PTB-binding 2 | 130.68 | 142.29 | -0.12 | 8.12E-01 |

| | | | | | | | |
|---|---|---|---|---|---|---|---|
| NULL | GSMG0026794 | NULL | NULL | 320.98 | 361.18 | -0.17 | 6.08E-01 |
| ENSMUSG00000028527 | GSMG0026798 | Ak4 | adenylate kinase 4 | 82.86 | 46.92 | 0.82 | 3.85E-02 |
| ENSMUSG00000028528 | GSMG0026800 | Dnajc6 | DnaJ heat shock protein family (Hsp40) member C6 | 320.66 | 321.24 | 0.00 | 1.00E+00 |
| ENSMUSG00000035212 | GSMG0026801 | Leprot | leptin receptor overlapping transcript | 5270.20 | 5432.42 | -0.04 | 8.80E-01 |
| ENSMUSG00000057722 | GSMG0026802 | Lepr | leptin receptor | 122.06 | 119.76 | 0.03 | 9.49E-01 |
| NULL | GSMG0026807 | NULL | NULL | 451.12 | 200.40 | 1.17 | 1.07E-09 |
| ENSMUSG00000028525 | GSMG0026812 | Pde4b | phosphodiesterase 4B, cAMP specific | 1348.92 | 1467.07 | -0.12 | 6.63E-01 |
| ENSMUSG00000028524 | GSMG0026814 | Sgip1 | SH3-domain GRB2-like (endophilin) interacting protein 1 | 817.99 | 932.75 | -0.19 | 3.93E-01 |
| ENSMUSG00000028522 | GSMG0026820 | Mier1 | MEIR1 treanscription regulator | 189.30 | 173.11 | 0.13 | 7.72E-01 |
| NULL | GSMG0026823 | NULL | NULL | 51.16 | 38.21 | 0.43 | 4.60E-01 |
| ENSMUSG00000035069 | GSMG0026826 | Oma1 | OMA1 zinc metallopeptidase | 541.88 | 528.76 | 0.04 | 9.29E-01 |
| NULL | GSMG0026830 | NULL | NULL | 167.97 | 229.16 | -0.44 | 1.15E-01 |
| ENSMUSG00000028519 | GSMG0026831 | Dab1 | disabled 1 | 220.54 | 463.94 | -1.07 | 3.66E-08 |
| NULL | GSMG0026832 | NULL | NULL | 431.84 | 562.27 | -0.38 | 6.88E-02 |
| NULL | GSMG0026833 | NULL | NULL | 480.74 | 336.43 | 0.52 | 1.19E-01 |
| NULL | GSMG0026835 | NULL | NULL | 282.99 | 313.97 | -0.15 | 6.57E-01 |
| NULL | GSMG0026836 | NULL | NULL | 232.68 | 225.44 | 0.05 | 9.09E-01 |
| NULL | GSMG0026838 | NULL | NULL | 386.85 | 539.07 | -0.48 | 2.56E-02 |
| NULL | GSMG0026839 | NULL | NULL | 481.82 | 490.06 | -0.02 | 9.52E-01 |
| ENSMUSG00000028517 | GSMG0026845 | Plpp3 | phospholipid phosphatase 3 | 1728.86 | 1270.40 | 0.45 | 1.73E-02 |
| ENSMUSG00000034926 | GSMG0026856 | Dhcr24 | 24-dehydrocholesterol reductase | 784.81 | 1333.61 | -0.77 | 9.45E-03 |
| ENSMUSG00000043572 | GSMG0026859 | Pars2 | prolyl-tRNA synthetase (mitochondrial)(putative) | 213.49 | 262.74 | -0.30 | 2.54E-01 |
| ENSMUSG00000061887 | GSMG0026862 | Ssbp3 | single-stranded DNA binding protein 3 | 569.01 | 657.44 | -0.21 | 4.53E-01 |
| NULL | GSMG0026863 | NULL | NULL | 191.30 | 134.26 | 0.52 | 1.56E-01 |
| NULL | GSMG0026864 | NULL | NULL | 121.98 | 173.55 | -0.51 | 1.01E-01 |
| ENSMUSG00000028618 | GSMG0026869 | Tmem59 | transmembrane protein 59 | 7013.42 | 6920.47 | 0.02 | 9.48E-01 |
| ENSMUSG00000057375 | GSMG0026871 | Yipf1 | Yip1 domain family, member 1 | 759.96 | 705.49 | 0.11 | 6.70E-01 |
| ENSMUSG00000028614 | GSMG0026872 | Ndc1 | NDC1 transmembrane nucleoporin | 190.48 | 335.41 | -0.82 | 9.22E-05 |
| ENSMUSG00000028613 | GSMG0026876 | Lrp8 | low density lipoprotein receptor-related protein 8, apolipoprotein e rece | 7783.41 | 10617.09 | -0.45 | 4.76E-02 |
| ENSMUSG00000028609 | GSMG0026877 | Magoh | mago homolog, exon junction complex core component | 682.42 | 849.42 | -0.32 | 2.02E-01 |
| ENSMUSG00000028608 | GSMG0026878 | 0610037L13Rik | RIKEN cDNA 0610037L13 gene | 1993.04 | 1702.19 | 0.23 | 2.44E-01 |
| ENSMUSG00000028601 | GSMG0026881 | Echdc2 | enoyl Coenzyme A hydratase domain containing 2 | 486.54 | 509.11 | -0.06 | 8.60E-01 |
| ENSMUSG00000048351 | GSMG0026882 | Coa7 | cytochrome c oxidase assembly factor 7 | 1018.31 | 1209.22 | -0.25 | 2.33E-01 |

| | | | | | | | |
|---|---|---|---|---|---|---|---|
| ENSMUSG00000034610 | GSMG0026886 | Zcchc11 | zinc finger, CCHC domain containing 11 | 808.06 | 872.22 | -0.11 | 6.89E-01 |
| ENSMUSG00000028582 | GSMG0026889 | Cc2d1b | coiled-coil and C2 domain containing 1B | 1234.91 | 1173.84 | 0.07 | 8.08E-01 |
| ENSMUSG00000028567 | GSMG0026892 | Txndc12 | thioredoxin domain containing 12 (endoplasmic reticulum) | 627.90 | 902.74 | -0.53 | 3.23E-03 |
| ENSMUSG00000003411 | GSMG0026895 | Rab3b | RAB3B, member RAS oncogene family | 338.71 | 697.02 | -1.04 | 2.27E-08 |
| ENSMUSG00000053510 | GSMG0026896 | Nrd1 | nardilysin, N-arginine dibasic convertase, NRD convertase 1 | 3451.80 | 3357.07 | 0.04 | 8.72E-01 |
| NULL | GSMG0026897 | NULL | NULL | 558.63 | 582.32 | -0.06 | 8.61E-01 |
| ENSMUSG00000028552 | GSMG0026899 | Eps15 | epidermal growth factor receptor pathway substrate 15 | 1886.71 | 1975.12 | -0.07 | 7.97E-01 |
| ENSMUSG00000010517 | GSMG0026902 | Faf1 | Fas-associated factor 1 | 1953.11 | 2018.97 | -0.05 | 8.57E-01 |
| ENSMUSG00000028545 | GSMG0026914 | Bend5 | BEN domain containing 5 | 641.89 | 717.73 | -0.16 | 4.98E-01 |
| ENSMUSG00000034401 | GSMG0026916 | Spata6 | spermatogenesis associated 6 | 612.13 | 536.72 | 0.19 | 4.25E-01 |
| ENSMUSG00000028718 | GSMG0026931 | Stil | Scl/Tal1 interrupting locus | 98.82 | 290.87 | -1.55 | 9.43E-10 |
| ENSMUSG00000034210 | GSMG0026939 | Efcab14 | EF-hand calcium binding domain 14 | 709.57 | 656.46 | 0.11 | 6.71E-01 |
| ENSMUSG00000028709 | GSMG0026942 | Mob3c | MOB kinase activator 3C | 460.59 | 405.91 | 0.18 | 4.98E-01 |
| ENSMUSG00000028708 | GSMG0026943 | Mknk1 | MAP kinase-interacting serine/threonine kinase 1 | 898.88 | 810.07 | 0.15 | 5.79E-01 |
| ENSMUSG00000028703 | GSMG0026946 | Lrrc41 | leucine rich repeat containing 41 | 243.82 | 257.26 | -0.08 | 8.36E-01 |
| ENSMUSG00000028700 | GSMG0026950 | Pomgnt1 | protein O-linked mannose beta 1,2-N-acetylglucosaminyltransferase | 922.76 | 921.03 | 0.00 | 9.91E-01 |
| ENSMUSG00000028698 | GSMG0026952 | Pik3r3 | phosphatidylinositol 3 kinase, regulatory subunit, polypeptide 3 (p55) | 960.83 | 638.94 | 0.59 | 1.15E-03 |
| ENSMUSG00000028696 | GSMG0026955 | Ipp | IAP promoted placental gene | 234.81 | 274.13 | -0.22 | 4.27E-01 |
| ENSMUSG00000034042 | GSMG0026956 | Gpbp1l1 | GC-rich promoter binding protein 1-like 1 | 775.49 | 783.07 | -0.02 | 9.64E-01 |
| ENSMUSG00000034035 | GSMG0026957 | Ccdc17 | coiled-coil domain containing 17 | 201.42 | 117.04 | 0.79 | 7.37E-03 |
| ENSMUSG00000028691 | GSMG0026960 | Prdx1 | peroxiredoxin 1 | 5038.33 | 5923.55 | -0.23 | 2.70E-01 |
| ENSMUSG00000028689 | GSMG0026964 | Ccdc163 | coiled-coil domain containing 163 | 128.16 | 133.12 | -0.05 | 9.23E-01 |
| ENSMUSG00000028687 | GSMG0026967 | Mutyh | mutY DNA glycosylase | 198.66 | 190.63 | 0.06 | 8.99E-01 |
| ENSMUSG00000046861 | GSMG0026971 | Hectd3 | HECT domain containing 3 | 1723.99 | 1657.18 | 0.06 | 8.71E-01 |
| ENSMUSG00000028683 | GSMG0026972 | Eif2b3 | eukaryotic translation initiation factor 2B, subunit 3 | 704.68 | 783.39 | -0.15 | 6.12E-01 |
| ENSMUSG00000048772 | GSMG0026977 | Tmem53 | transmembrane protein 53 | 77.71 | 94.42 | -0.28 | 5.17E-01 |
| ENSMUSG00000033423 | GSMG0026978 | Eri3 | exoribonuclease 3 | 1201.08 | 1728.95 | -0.53 | 7.67E-04 |
| NULL | GSMG0026979 | NULL | NULL | 125.51 | 108.55 | 0.22 | 6.30E-01 |
| ENSMUSG00000028542 | GSMG0026982 | Slc6a9 | solute carrier family 6 (neurotransmitter transporter, glycine), member 9 | 429.31 | 537.11 | -0.32 | 2.89E-01 |
| ENSMUSG00000006395 | GSMG0026985 | Hyi | hydroxypyruvate isomerase (putative) | 107.98 | 111.40 | -0.05 | 9.28E-01 |
| ENSMUSG00000006392 | GSMG0026986 | Med8 | mediator complex subunit 8 | 680.88 | 906.03 | -0.41 | 3.83E-02 |
| ENSMUSG00000006390 | GSMG0026987 | Elovl1 | elongation of very long chain fatty acids (FEN1/Elo2, SUR4/Elo3, yeas | 1473.38 | 1112.72 | 0.41 | 3.71E-02 |
| ENSMUSG00000028645 | GSMG0026997 | Slc2a1 | solute carrier family 2 (facilitated glucose transporter), member 1 | 31397.21 | 14860.71 | 1.08 | 5.15E-11 |

| | | | | | | | |
|---|---|---|---|---|---|---|---|
| ENSMUSG00000028643 | GSMG0026998 | Svbp | small vasohibin binding protein | 203.49 | 353.86 | -0.80 | 3.51E-04 |
| ENSMUSG00000028641 | GSMG0026999 | P3h1 | prolyl 3-hydroxylase 1 | 917.81 | 762.37 | 0.27 | 2.03E-01 |
| ENSMUSG00000032998 | GSMG0027005 | Foxj3 | forkhead box J3 | 404.69 | 435.72 | -0.11 | 7.18E-01 |
| ENSMUSG00000000085 | GSMG0027013 | Scmh1 | sex comb on midleg homolog 1 | 453.56 | 484.26 | -0.09 | 7.58E-01 |
| ENSMUSG00000032890 | GSMG0027021 | Rims3 | regulating synaptic membrane exocytosis 3 | 49.94 | 122.92 | -1.31 | 8.12E-05 |
| ENSMUSG00000028626 | GSMG0027022 | Col9a2 | collagen, type IX, alpha 2 | 383.93 | 503.54 | -0.39 | 2.16E-01 |
| ENSMUSG00000028657 | GSMG0027024 | Ppt1 | palmitoyl-protein thioesterase 1 | 1783.37 | 1537.50 | 0.21 | 2.92E-01 |
| ENSMUSG00000028654 | GSMG0027027 | Mycl | v-myc avian myelocytomatosis viral oncogene lung carcinoma derived | 758.81 | 1299.48 | -0.78 | 1.82E-07 |
| ENSMUSG00000028653 | GSMG0027028 | Trit1 | tRNA isopentenyltransferase 1 | 286.07 | 307.82 | -0.10 | 7.44E-01 |
| ENSMUSG00000046093 | GSMG0027032 | Hpcal4 | hippocalcin-like 4 | 3288.12 | 1414.75 | 1.22 | 1.03E-13 |
| ENSMUSG00000032744 | GSMG0027034 | Heyl | hairy/enhancer-of-split related with YRPW motif-like | 3937.24 | 4096.90 | -0.06 | 8.31E-01 |
| ENSMUSG00000028647 | GSMG0027039 | Mycbp | MYC binding protein | 1177.49 | 1150.69 | 0.03 | 9.12E-01 |
| ENSMUSG00000028646 | GSMG0027040 | Rragc | Ras-related GTP binding C | 696.38 | 643.57 | 0.11 | 7.07E-01 |
| ENSMUSG00000090125 | GSMG0027043 | Pou3f1 | POU domain, class 3, transcription factor 1 | 631.85 | 1501.23 | -1.25 | 3.80E-10 |
| ENSMUSG00000032643 | GSMG0027044 | Fhl3 | four and a half LIM domains 3 | 36.84 | 204.51 | -2.48 | 9.67E-10 |
| ENSMUSG00000028902 | GSMG0027046 | Sf3a3 | splicing factor 3a, subunit 3 | 1683.24 | 2081.78 | -0.31 | 1.15E-01 |
| ENSMUSG00000028894 | GSMG0027047 | Inpp5b | inositol polyphosphate-5-phosphatase B | 1834.39 | 1484.81 | 0.31 | 7.56E-02 |
| NULL | GSMG0027048 | NULL | NULL | 276.06 | 195.08 | 0.50 | 1.03E-01 |
| ENSMUSG00000028890 | GSMG0027049 | Mtf1 | metal response element binding transcription factor 1 | 552.27 | 495.26 | 0.16 | 5.67E-01 |
| ENSMUSG00000028889 | GSMG0027052 | Yrdc | yrdC domain containing (E.coli) | 888.99 | 1110.48 | -0.32 | 9.09E-02 |
| ENSMUSG00000044730 | GSMG0027055 | 9930104L06Rik | RIKEN cDNA 9930104L06 gene | 151.25 | 159.56 | -0.08 | 8.67E-01 |
| ENSMUSG00000028869 | GSMG0027057 | Gnl2 | guanine nucleotide binding protein-like 2 (nucleolar) | 3476.53 | 3462.04 | 0.01 | 9.82E-01 |
| ENSMUSG00000050213 | GSMG0027058 | Snip1 | Smad nuclear interacting protein 1 | 195.54 | 178.66 | 0.13 | 7.24E-01 |
| ENSMUSG00000028863 | GSMG0027059 | Meaf6 | MYST/Esa1-associated factor 6 | 1099.53 | 1523.18 | -0.47 | 4.85E-03 |
| ENSMUSG00000028861 | GSMG0027067 | Mrps15 | mitochondrial ribosomal protein S15 | 776.95 | 997.53 | -0.36 | 1.06E-01 |
| ENSMUSG00000042616 | GSMG0027068 | Oscp1 | organic solute carrier partner 1 | 238.22 | 228.21 | 0.06 | 9.03E-01 |
| ENSMUSG00000050188 | GSMG0027069 | Lsm10 | U7 snRNP-specific Sm-like protein LSM10 | 417.90 | 457.01 | -0.13 | 6.55E-01 |
| ENSMUSG00000042608 | GSMG0027070 | Stk40 | serine/threonine kinase 40 | 570.56 | 519.39 | 0.13 | 6.89E-01 |
| ENSMUSG00000050212 | GSMG0027071 | Eva1b | eva-1 homolog B (C. elegans) | 2365.99 | 2330.80 | 0.02 | 9.42E-01 |
| ENSMUSG00000028847 | GSMG0027074 | Trappc3 | trafficking protein particle complex 3 | 1837.06 | 2131.69 | -0.21 | 2.88E-01 |
| ENSMUSG00000028837 | GSMG0027083 | Psmb2 | proteasome (prosome, macropain) subunit, beta type 2 | 3551.31 | 4428.35 | -0.32 | 7.53E-02 |
| ENSMUSG00000028830 | GSMG0027086 | AU040320 | expressed sequence AU040320 | 748.04 | 794.85 | -0.09 | 7.70E-01 |
| ENSMUSG00000042408 | GSMG0027092 | Zmym6 | zinc finger, MYM-type 6 | 856.92 | 854.35 | 0.01 | 9.87E-01 |

| | | | | | | | |
|---|---|---|---|---|---|---|---|
| ENSMUSG00000070737 | GSMG0027093 | Gm12942 | predicted gene 12942 | 148.60 | 179.18 | -0.27 | 4.21E-01 |
| ENSMUSG00000042388 | GSMG0027097 | Dlgap3 | discs, large (Drosophila) homolog-associated protein 3 | 677.78 | 371.36 | 0.87 | 1.35E-04 |
| ENSMUSG00000042380 | GSMG0027098 | Smim12 | small integral membrane protein 12 | 1170.08 | 1210.47 | -0.05 | 8.67E-01 |
| ENSMUSG00000028804 | GSMG0027103 | Csmd2 | CUB and Sushi multiple domains 2 | 623.24 | 761.68 | -0.29 | 2.96E-01 |
| NULL | GSMG0027107 | NULL | NULL | 86.82 | 105.93 | -0.29 | 5.35E-01 |
| ENSMUSG00000028796 | GSMG0027109 | Phc2 | polyhomeotic-like 2 (Drosophila) | 1922.93 | 1633.70 | 0.23 | 4.88E-01 |
| NULL | GSMG0027110 | NULL | NULL | 476.40 | 437.43 | 0.12 | 6.69E-01 |
| ENSMUSG00000041000 | GSMG0027113 | Trim62 | tripartite motif-containing 62 | 1350.74 | 1420.99 | -0.07 | 7.84E-01 |
| ENSMUSG00000028792 | GSMG0027114 | Ak2 | adenylate kinase 2 | 982.81 | 1176.23 | -0.26 | 1.91E-01 |
| ENSMUSG00000028793 | GSMG0027115 | Rnf19b | ring finger protein 19B | 322.30 | 396.70 | -0.30 | 3.16E-01 |
| ENSMUSG00000001334 | GSMG0027118 | Fndc5 | fibronectin type III domain containing 5 | 189.14 | 100.78 | 0.91 | 6.07E-03 |
| ENSMUSG00000028811 | GSMG0027119 | Yars | tyrosyl-tRNA synthetase | 1061.73 | 1427.69 | -0.43 | 8.50E-03 |
| NULL | GSMG0027123 | NULL | NULL | 156.58 | 194.50 | -0.32 | 5.76E-01 |
| ENSMUSG00000040859 | GSMG0027126 | Bsdc1 | BSD domain containing 1 | 459.18 | 456.12 | 0.01 | 9.75E-01 |
| ENSMUSG00000047945 | GSMG0027128 | Marcksl1 | MARCKS-like 1 | 2352.69 | 7511.85 | -1.68 | 4.13E-28 |
| ENSMUSG00000028797 | GSMG0027129 | Tmem234 | transmembrane protein 234 | 3296.96 | 3119.01 | 0.08 | 7.27E-01 |
| ENSMUSG00000028788 | GSMG0027133 | Ptp4a2 | protein tyrosine phosphatase 4a2 | 6103.39 | 8004.98 | -0.39 | 1.07E-01 |
| ENSMUSG00000028782 | GSMG0027139 | Adgrb2 | adhesion G protein-coupled receptor B2 | 2462.88 | 1948.40 | 0.34 | 1.54E-01 |
| ENSMUSG00000040690 | GSMG0027140 | Col16a1 | collagen, type XVI, alpha 1 | 1098.14 | 1048.65 | 0.07 | 8.61E-01 |
| ENSMUSG00000028779 | GSMG0027141 | Pef1 | penta-EF hand domain containing 1 | 561.91 | 511.83 | 0.14 | 6.29E-01 |
| ENSMUSG00000074088 | GSMG0027143 | Snrnp40 | small nuclear ribonucleoprotein 40 (U5) | 988.04 | 1319.50 | -0.42 | 3.08E-02 |
| ENSMUSG00000078532 | GSMG0027145 | Nkain1 | Na+/K+ transporting ATPase interacting 1 | 75.71 | 293.37 | -1.94 | 1.64E-13 |
| ENSMUSG00000028580 | GSMG0027147 | Pum1 | pumilio RNA-binding family member 1 | 2576.10 | 3651.26 | -0.50 | 1.54E-03 |
| NULL | GSMG0027149 | NULL | NULL | 130.36 | 204.36 | -0.66 | 4.01E-02 |
| ENSMUSG00000025743 | GSMG0027151 | Sdc3 | syndecan 3 | 7501.87 | 9632.39 | -0.36 | 1.13E-01 |
| ENSMUSG00000028581 | GSMG0027152 | Laptm5 | lysosomal-associated protein transmembrane 5 | 1063.92 | 706.55 | 0.59 | 9.44E-04 |
| ENSMUSG00000028910 | GSMG0027156 | Mecr | mitochondrial trans-2-enoyl-CoA reductase | 470.68 | 497.36 | -0.08 | 7.96E-01 |
| ENSMUSG00000028911 | GSMG0027157 | Srsf4 | serine/arginine-rich splicing factor 4 | 595.32 | 706.55 | -0.25 | 2.29E-01 |
| NULL | GSMG0027158 | NULL | NULL | 136.53 | 114.91 | 0.25 | 5.29E-01 |
| NULL | GSMG0027159 | NULL | NULL | 225.44 | 171.63 | 0.40 | 3.05E-01 |
| NULL | GSMG0027161 | NULL | NULL | 47.52 | 58.88 | -0.33 | 7.14E-01 |
| ENSMUSG00000028899 | GSMG0027163 | Taf12 | TATA-box binding protein associated factor 12 | 312.34 | 325.32 | -0.06 | 8.89E-01 |
| ENSMUSG00000086290 | GSMG0027165 | Snhg12 | small nucleolar RNA host gene 12 | 120.64 | 117.76 | 0.03 | 9.66E-01 |

| | | | | | | | |
|---|---|---|---|---|---|---|---|
| NULL | GSMG0027167 | NULL | NULL | 208.23 | 178.45 | 0.22 | 4.95E-01 |
| ENSMUSG00000054405 | GSMG0027170 | Dnajc8 | DnaJ heat shock protein family (Hsp40) member C8 | 3337.07 | 3814.24 | -0.19 | 4.93E-01 |
| ENSMUSG00000028886 | GSMG0027173 | Eya3 | EYA transcriptional coactivator and phosphatase 3 | 253.05 | 287.51 | -0.18 | 5.86E-01 |
| ENSMUSG00000028884 | GSMG0027175 | Rpa2 | replication protein A2 | 286.18 | 483.31 | -0.75 | 4.75E-05 |
| NULL | GSMG0027177 | NULL | NULL | 67.02 | 86.19 | -0.37 | 4.45E-01 |
| ENSMUSG00000037692 | GSMG0027179 | Ahdc1 | AT hook, DNA binding motif, containing 1 | 690.95 | 755.37 | -0.13 | 7.11E-01 |
| ENSMUSG00000028868 | GSMG0027181 | Wasf2 | WAS protein family, member 2 | 819.24 | 984.29 | -0.27 | 3.27E-01 |
| ENSMUSG00000028862 | GSMG0027185 | Map3k6 | mitogen-activated protein kinase kinase kinase 6 | 280.10 | 105.09 | 1.41 | 2.00E-06 |
| ENSMUSG00000028854 | GSMG0027186 | Slc9a1 | solute carrier family 9 (sodium/hydrogen exchanger), member 1 | 971.70 | 1078.09 | -0.15 | 4.94E-01 |
| ENSMUSG00000028850 | GSMG0027191 | Gpatch3 | G patch domain containing 3 | 191.59 | 203.77 | -0.09 | 8.43E-01 |
| ENSMUSG00000028848 | GSMG0027192 | Gpn2 | GPN-loop GTPase 2 | 316.21 | 332.40 | -0.07 | 8.31E-01 |
| NULL | GSMG0027194 | NULL | NULL | 44.84 | 53.97 | -0.28 | 6.73E-01 |
| ENSMUSG00000012126 | GSMG0027201 | Ubxn11 | UBX domain protein 11 | 200.23 | 123.20 | 0.70 | 1.01E-02 |
| ENSMUSG00000037366 | GSMG0027207 | Pafah2 | platelet-activating factor acetylhydrolase 2 | 649.80 | 558.70 | 0.22 | 3.75E-01 |
| ENSMUSG00000037348 | GSMG0027209 | Paqr7 | progestin and adipoQ receptor family member VII | 3206.77 | 2308.78 | 0.47 | 1.41E-03 |
| NULL | GSMG0027211 | NULL | NULL | 248.56 | 154.76 | 0.68 | 3.26E-02 |
| NULL | GSMG0027215 | NULL | NULL | 873.09 | 898.81 | -0.04 | 9.03E-01 |
| ENSMUSG00000037266 | GSMG0027216 | Rsrp1 | arginine/serine rich protein 1 | 20288.70 | 14756.75 | 0.46 | 8.45E-03 |
| ENSMUSG00000028821 | GSMG0027217 | Syf2 | SYF2 homolog, RNA splicing factor (S. cerevisiae) | 1394.13 | 1392.49 | 0.00 | 9.98E-01 |
| ENSMUSG00000028676 | GSMG0027231 | Srsf10 | serine/arginine-rich splicing factor 10 | 2651.55 | 3075.74 | -0.21 | 3.80E-01 |
| ENSMUSG00000028673 | GSMG0027233 | Fuca1 | fucosidase, alpha-L- 1, tissue | 1918.88 | 2288.80 | -0.25 | 1.62E-01 |
| ENSMUSG00000028672 | GSMG0027234 | Hmgcl | 3-hydroxy-3-methylglutaryl-Coenzyme A lyase | 853.05 | 948.45 | -0.15 | 5.01E-01 |
| ENSMUSG00000028671 | GSMG0027235 | Gale | galactose-4-epimerase, UDP | 596.80 | 787.96 | -0.40 | 6.57E-02 |
| ENSMUSG00000007872 | GSMG0027236 | Id3 | inhibitor of DNA binding 3 | 11740.84 | 8938.07 | 0.39 | 2.93E-02 |
| ENSMUSG00000018983 | GSMG0027237 | E2f2 | E2F transcription factor 2 | 54.42 | 113.23 | -1.08 | 2.83E-03 |
| ENSMUSG00000036995 | GSMG0027238 | Asap3 | ArfGAP with SH3 domain, ankyrin repeat and PH domain 3 | 188.91 | 231.89 | -0.30 | 3.40E-01 |
| ENSMUSG00000051351 | GSMG0027241 | Zfp46 | zinc finger protein 46 | 718.00 | 765.88 | -0.09 | 7.19E-01 |
| ENSMUSG00000066037 | GSMG0027242 | Hnrnpr | heterogeneous nuclear ribonucleoprotein R | 758.87 | 1042.18 | -0.46 | 6.95E-03 |
| NULL | GSMG0027245 | NULL | NULL | 401.36 | 410.02 | -0.04 | 9.49E-01 |
| NULL | GSMG0027246 | NULL | NULL | 674.35 | 672.46 | 0.00 | 9.99E-01 |
| NULL | GSMG0027249 | NULL | NULL | 146.77 | 120.42 | 0.29 | 4.20E-01 |
| ENSMUSG00000036856 | GSMG0027258 | Wnt4 | wingless-type MMTV integration site family, member 4 | 175.33 | 126.40 | 0.47 | 1.58E-01 |
| ENSMUSG00000028763 | GSMG0027261 | Hspg2 | perlecan (heparan sulfate proteoglycan 2) | 2265.50 | 3040.18 | -0.42 | 7.70E-02 |

| | | | | | | | |
|---|---|---|---|---|---|---|---|
| ENSMUSG00000043411 | GSMG0027262 | Usp48 | ubiquitin specific peptidase 48 | 1075.92 | 1174.16 | -0.13 | 6.33E-01 |
| NULL | GSMG0027263 | NULL | NULL | 327.77 | 444.23 | -0.44 | 9.71E-02 |
| ENSMUSG00000041351 | GSMG0027264 | Rap1gap | Rap1 GTPase-activating protein | 1478.60 | 1202.78 | 0.30 | 2.89E-01 |
| ENSMUSG00000057530 | GSMG0027265 | Ece1 | endothelin converting enzyme 1 | 7087.12 | 6096.02 | 0.22 | 2.61E-01 |
| NULL | GSMG0027266 | NULL | NULL | 515.40 | 570.25 | -0.14 | 7.05E-01 |
| ENSMUSG00000028760 | GSMG0027267 | Eif4g3 | eukaryotic translation initiation factor 4 gamma, 3 | 1518.84 | 1798.47 | -0.24 | 1.90E-01 |
| ENSMUSG00000028759 | GSMG0027271 | Hp1bp3 | heterochromatin protein 1, binding protein 3 | 2066.63 | 2286.22 | -0.15 | 4.76E-01 |
| G00000028758,ENSMUSG000 | GSMG0027272 | Sh2d5 // Kif17 | SH2 domain containing 5 // kinesin family member 17 | 235.84 | 93.25 | 1.34 | 3.89E-07 |
| ENSMUSG00000028757 | GSMG0027273 | Ddost | dolichyl-di-phosphooligosaccharide-protein glycotransferase | 5481.39 | 6759.11 | -0.30 | 8.37E-02 |
| ENSMUSG00000041241 | GSMG0027275 | Mul1 | mitochondrial ubiquitin ligase activator of NFKB 1 | 754.74 | 682.99 | 0.14 | 6.26E-01 |
| ENSMUSG00000046447 | GSMG0027277 | Camk2n1 | calcium/calmodulin-dependent protein kinase II inhibitor 1 | 2614.59 | 810.65 | 1.69 | 1.12E-24 |
| ENSMUSG00000028745 | GSMG0027286 | Capzb | capping protein (actin filament) muscle Z-line, beta | 3689.82 | 4002.40 | -0.12 | 5.72E-01 |
| ENSMUSG00000028743 | GSMG0027288 | Akr7a5 | aldo-keto reductase family 7, member A5 (aflatoxin aldehyde reductase | 395.79 | 491.82 | -0.32 | 1.87E-01 |
| ENSMUSG00000028737 | GSMG0027294 | Aldh4a1 | aldehyde dehydrogenase 4 family, member A1 | 821.89 | 787.12 | 0.06 | 8.43E-01 |
| ENSMUSG00000040945 | GSMG0027301 | Rcc2 | regulator of chromosome condensation 2 | 964.80 | 984.48 | -0.03 | 9.52E-01 |
| ENSMUSG00000009863 | GSMG0027305 | Sdhb | succinate dehydrogenase complex, subunit B, iron sulfur (Ip) | 3050.56 | 3602.48 | -0.24 | 1.82E-01 |
| ENSMUSG00000036622 | GSMG0027306 | Atp13a2 | ATPase type 13A2 | 8601.28 | 11029.34 | -0.36 | 1.16E-01 |
| ENSMUSG00000060572 | GSMG0027307 | Mfap2 | microfibrillar-associated protein 2 | 947.81 | 1382.49 | -0.54 | 6.11E-03 |
| NULL | GSMG0027311 | NULL | NULL | 76.49 | 62.11 | 0.30 | 6.45E-01 |
| ENSMUSG00000028920 | GSMG0027313 | Fbxo42 | F-box protein 42 | 386.00 | 398.37 | -0.05 | 9.11E-01 |
| ENSMUSG00000006445 | GSMG0027317 | Epha2 | Eph receptor A2 | 281.61 | 253.37 | 0.15 | 6.28E-01 |
| ENSMUSG00000006218 | GSMG0027319 | Fam131c | family with sequence similarity 131, member C | 12.15 | 102.27 | -3.03 | 3.33E-13 |
| ENSMUSG00000006215 | GSMG0027321 | Zbtb17 | zinc finger and BTB domain containing 17 | 1505.94 | 1595.10 | -0.08 | 7.90E-01 |
| ENSMUSG00000028914 | GSMG0027330 | Casp9 | caspase 9 | 300.57 | 314.34 | -0.06 | 8.61E-01 |
| ENSMUSG00000046862 | GSMG0027340 | Pramef8 | PRAME family member 8 | 603.62 | 544.54 | 0.15 | 5.84E-01 |
| ENSMUSG00000066026 | GSMG0027354 | Dhrs3 | dehydrogenase/reductase (SDR family) member 3 | 681.54 | 659.66 | 0.05 | 8.94E-01 |
| ENSMUSG00000044496 | GSMG0027373 | 2510039O18Rik | RIKEN cDNA 2510039O18 gene | 1638.71 | 1467.57 | 0.16 | 5.25E-01 |
| ENSMUSG00000029009 | GSMG0027376 | Mthfr | 5,10-methylenetetrahydrofolate reductase | 818.41 | 659.24 | 0.31 | 3.40E-01 |
| ENSMUSG00000029003 | GSMG0027377 | Mad2l2 | MAD2 mitotic arrest deficient-like 2 | 567.70 | 785.24 | -0.47 | 7.53E-03 |
| ENSMUSG00000041556 | GSMG0027378 | Fbxo2 | F-box protein 2 | 326.10 | 119.95 | 1.45 | 1.87E-07 |
| ENSMUSG00000028991 | GSMG0027382 | Mtor | mechanistic target of rapamycin (serine/threonine kinase) | 2231.31 | 2764.62 | -0.31 | 1.79E-01 |
| ENSMUSG00000017264 | GSMG0027384 | Exosc10 | exosome component 10 | 3306.54 | 3833.95 | -0.21 | 2.20E-01 |
| ENSMUSG00000006442 | GSMG0027385 | Srm | spermidine synthase | 941.50 | 1622.74 | -0.78 | 1.05E-07 |

| | | | | | | | |
|---|---|---|---|---|---|---|---|
| ENSMUSG00000028979 | GSMG0027386 | Masp2 | mannan-binding lectin serine peptidase 2 | 60.17 | 103.30 | -0.79 | 1.19E-01 |
| ENSMUSG00000028974 | GSMG0027391 | Dffa | DNA fragmentation factor, alpha subunit | 400.82 | 408.01 | -0.02 | 9.53E-01 |
| ENSMUSG00000028990 | GSMG0027395 | Lzic | leucine zipper and CTNNBIP1 domain containing | 568.48 | 822.91 | -0.53 | 3.08E-03 |
| ENSMUSG00000028988 | GSMG0027397 | Ctnnbip1 | catenin beta interacting protein 1 | 2355.36 | 3345.97 | -0.51 | 1.27E-02 |
| ENSMUSG00000039953 | GSMG0027400 | Clstn1 | calsyntenin 1 | 5876.16 | 4813.50 | 0.29 | 3.13E-01 |
| NULL | GSMG0027407 | NULL | NULL | 64.48 | 61.89 | 0.06 | 9.30E-01 |
| ENSMUSG00000039852 | GSMG0027409 | Rere | arginine glutamic acid dipeptide (RE) repeats | 980.00 | 1065.41 | -0.12 | 7.04E-01 |
| NULL | GSMG0027410 | NULL | NULL | 82.57 | 106.49 | -0.38 | 4.06E-01 |
| NULL | GSMG0027412 | NULL | NULL | 262.61 | 310.18 | -0.24 | 5.02E-01 |
| NULL | GSMG0027413 | NULL | NULL | 362.23 | 370.28 | -0.03 | 9.43E-01 |
| NULL | GSMG0027416 | NULL | NULL | 210.71 | 211.57 | -0.01 | 9.90E-01 |
| ENSMUSG00000028967 | GSMG0027419 | Errfi1 | ERBB receptor feedback inhibitor 1 | 1323.92 | 673.16 | 0.98 | 3.95E-07 |
| NULL | GSMG0027425 | NULL | NULL | 1151.12 | 949.70 | 0.28 | 1.62E-01 |
| ENSMUSG00000039768 | GSMG0027430 | Dnajc11 | DnaJ heat shock protein family (Hsp40) member C11 | 1119.49 | 1427.33 | -0.35 | 6.44E-02 |
| ENSMUSG00000073700 | GSMG0027431 | Klhl21 | kelch-like 21 | 532.73 | 392.07 | 0.44 | 1.42E-01 |
| ENSMUSG00000028948 | GSMG0027433 | Nol9 | nucleolar protein 9 | 1035.02 | 1333.48 | -0.37 | 6.10E-02 |
| ENSMUSG00000039713 | GSMG0027434 | Plekhg5 | pleckstrin homology domain containing, family G (with RhoGef domai | 2355.12 | 1949.23 | 0.27 | 2.62E-01 |
| ENSMUSG00000024793 | GSMG0027435 | Tnfrsf25 | tumor necrosis factor receptor superfamily, member 25 | 109.48 | 64.68 | 0.75 | 4.80E-02 |
| ENSMUSG00000028937 | GSMG0027437 | Acot7 | acyl-CoA thioesterase 7 | 927.48 | 1612.91 | -0.80 | 9.63E-07 |
| ENSMUSG00000042804 | GSMG0027439 | Gpr153 | G protein-coupled receptor 153 | 175.03 | 133.80 | 0.39 | 3.64E-01 |
| ENSMUSG00000039662 | GSMG0027440 | Icmt | isoprenylcysteine carboxyl methyltransferase | 2754.96 | 2537.89 | 0.12 | 6.70E-01 |
| ENSMUSG00000028936 | GSMG0027441 | Rpl22 | ribosomal protein L22 | 992.49 | 1580.51 | -0.67 | 6.40E-05 |
| ENSMUSG00000005045 | GSMG0027442 | Chd5 | chromodomain helicase DNA binding protein 5 | 554.16 | 635.19 | -0.20 | 5.61E-01 |
| ENSMUSG00000039577 | GSMG0027443 | Nphp4 | nephronophthisis 4 (juvenile) homolog (human) | 519.28 | 316.28 | 0.71 | 4.51E-03 |
| ENSMUSG00000047613 | GSMG0027447 | A430005L14Rik | RIKEN cDNA A430005L14 gene | 600.65 | 630.75 | -0.07 | 8.24E-01 |
| ENSMUSG00000039523 | GSMG0027448 | Cep104 | centrosomal protein 104 | 976.83 | 1022.55 | -0.07 | 8.13E-01 |
| ENSMUSG00000029028 | GSMG0027449 | Lrrc47 | leucine rich repeat containing 47 | 515.96 | 513.37 | 0.01 | 9.88E-01 |
| ENSMUSG00000029029 | GSMG0027452 | Wrap73 | WD repeat containing, antisense to Trp73 | 632.50 | 601.19 | 0.07 | 8.26E-01 |
| ENSMUSG00000048001 | GSMG0027464 | Hes5 | hairy and enhancer of split 5 (Drosophila) | 507.19 | 820.69 | -0.70 | 9.59E-04 |
| ENSMUSG00000029056 | GSMG0027465 | Pank4 | pantothenate kinase 4 | 782.41 | 764.66 | 0.03 | 9.28E-01 |
| ENSMUSG00000029047 | GSMG0027468 | Pex10 | peroxisomal biogenesis factor 10 | 217.03 | 216.87 | 0.00 | 9.98E-01 |
| ENSMUSG00000029049 | GSMG0027470 | Morn1 | MORN repeat containing 1 | 255.92 | 189.70 | 0.43 | 1.24E-01 |
| ENSMUSG00000029070 | GSMG0027472 | Mxra8 | matrix-remodelling associated 8 | 1878.86 | 1794.65 | 0.07 | 8.03E-01 |

| | | | | | | | |
|---|---|---|---|---|---|---|---|
| ENSMUSG00000073684 | GSMG0027473 | Faap20 | Fanconi anemia core complex associated protein 20 | 569.07 | 640.37 | -0.17 | 5.03E-01 |
| ENSMUSG00000078490 | GSMG0027477 | Cfap74 | cilia and flagella associated protein 74 | 104.76 | 67.48 | 0.63 | 8.97E-02 |
| ENSMUSG00000029064 | GSMG0027481 | Gnb1 | guanine nucleotide binding protein (G protein), beta 1 | 20511.14 | 24248.81 | -0.24 | 1.63E-01 |
| ENSMUSG00000029063 | GSMG0027484 | Nadk | NAD kinase | 1479.83 | 1655.47 | -0.16 | 4.68E-01 |
| ENSMUSG00000042202 | GSMG0027486 | Slc35e2 | solute carrier family 35, member E2 | 141.66 | 168.47 | -0.25 | 5.22E-01 |
| ENSMUSG00000029062 | GSMG0027487 | Cdk11b | cyclin-dependent kinase 11B | 4568.94 | 4756.85 | -0.06 | 8.13E-01 |
| ENSMUSG00000074738 | GSMG0027488 | B930041F14Rik | RIKEN cDNA B930041F14 gene | 920.04 | 868.16 | 0.08 | 8.19E-01 |
| ENSMUSG00000029038 | GSMG0027489 | Ssu72 | Ssu72 RNA polymerase II CTD phosphatase homolog (yeast) | 2435.01 | 2663.26 | -0.13 | 5.77E-01 |
| ENSMUSG00000084845 | GSMG0027491 | Tmem240 | transmembrane protein 240 | 234.30 | 151.94 | 0.62 | 5.67E-01 |
| NULL | GSMG0027494 | NULL | NULL | 1251.01 | 1265.18 | -0.02 | 9.56E-01 |
| ENSMUSG00000029066 | GSMG0027496 | Mrpl20 | mitochondrial ribosomal protein L20 | 1376.79 | 1643.53 | -0.26 | 2.42E-01 |
| ENSMUSG00000029068 | GSMG0027497 | Ccnl2 | cyclin L2 | 4693.38 | 4028.64 | 0.22 | 1.88E-01 |
| ENSMUSG00000065990 | GSMG0027498 | Aurkaip1 | aurora kinase A interacting protein 1 | 1034.82 | 1124.89 | -0.12 | 6.07E-01 |
| ENSMUSG00000029071 | GSMG0027499 | Dvl1 | dishevelled segment polarity protein 1 | 2872.05 | 1756.90 | 0.71 | 2.91E-01 |
| ENSMUSG00000029034 | GSMG0027500 | Cpsf3l | cleavage and polyadenylation specific factor 3-like | 1982.43 | 2218.65 | -0.16 | 4.16E-01 |
| ENSMUSG00000029033 | GSMG0027501 | Acap3 | ArfGAP with coiled-coil, ankyrin repeat and PH domains 3 | 4752.91 | 4908.75 | -0.05 | 9.01E-01 |
| ENSMUSG00000023286 | GSMG0027502 | Ube2j2 | ubiquitin-conjugating enzyme E2J 2 | 799.20 | 1143.89 | -0.52 | 6.38E-03 |
| ENSMUSG00000023571 | GSMG0027503 | Fam132a | family with sequence similarity 132, member A | 324.42 | 378.30 | -0.22 | 4.04E-01 |
| ENSMUSG00000029076 | GSMG0027504 | Sdf4 | stromal cell derived factor 4 | 3897.36 | 3936.51 | -0.01 | 9.59E-01 |
| ENSMUSG00000041954 | GSMG0027506 | Tnfrsf18 | tumor necrosis factor receptor superfamily, member 18 | 70.87 | 85.27 | -0.26 | 5.98E-01 |
| ENSMUSG00000059939 | GSMG0027507 | 9430015G10Rik | RIKEN cDNA 9430015G10 gene | 320.25 | 376.79 | -0.23 | 3.91E-01 |
| ENSMUSG00000028232 | GSMG0027514 | Tmem68 | transmembrane protein 68 | 354.99 | 444.18 | -0.32 | 2.06E-01 |
| ENSMUSG00000028234 | GSMG0027518 | Rps20 | ribosomal protein S20 | 1305.87 | 2022.19 | -0.63 | 2.57E-03 |
| ENSMUSG00000045573 | GSMG0027524 | Penk | preproenkephalin | 139.51 | 60.02 | 1.21 | 1.11E-02 |
| ENSMUSG00000066324 | GSMG0027525 | Impad1 | inositol monophosphatase domain containing 1 | 2062.01 | 2577.08 | -0.32 | 6.93E-02 |
| ENSMUSG00000028245 | GSMG0027529 | Nsmaf | neutral sphingomyelinase (N-SMase) activation associated factor | 452.47 | 434.98 | 0.06 | 8.67E-01 |
| ENSMUSG00000041272 | GSMG0027531 | Tox | thymocyte selection-associated high mobility group box | 166.16 | 293.45 | -0.81 | 7.94E-04 |
| ENSMUSG00000041261 | GSMG0027542 | Car8 | carbonic anhydrase 8 | 82.58 | 253.36 | -1.62 | 3.14E-12 |
| ENSMUSG00000028207 | GSMG0027548 | Asph | aspartate-beta-hydroxylase | 533.09 | 718.65 | -0.43 | 2.21E-02 |
| ENSMUSG00000049969 | GSMG0027551 | Plekhf2 | pleckstrin homology domain containing, family F (with FYVE domain) | 1521.55 | 1537.74 | -0.02 | 9.66E-01 |
| ENSMUSG00000050323 | GSMG0027552 | Ndufaf6 | NADH dehydrogenase (ubiquinone) complex I, assembly factor 6 | 165.60 | 258.30 | -0.64 | 6.83E-03 |
| ENSMUSG00000040738 | GSMG0027554 | Ints8 | integrator complex subunit 8 | 1643.40 | 1433.52 | 0.20 | 3.74E-01 |
| NULL | GSMG0027555 | NULL | NULL | 84.86 | 88.80 | -0.08 | 9.08E-01 |

| | | | | | | | |
|---|---|---|---|---|---|---|---|
| ENSMUSG00000045205 | GSMG0027556 | Dpy19l4 | dpy-19-like 4 (C. elegans) | 327.48 | 406.78 | -0.31 | 1.79E-01 |
| ENSMUSG00000049225 | GSMG0027561 | Pdp1 | pyruvate dehyrogenase phosphatase catalytic subunit 1 | 1456.55 | 1080.52 | 0.43 | 9.72E-03 |
| ENSMUSG00000049488 | GSMG0027565 | Tmem67 | transmembrane protein 67 | 232.34 | 235.86 | -0.02 | 9.56E-01 |
| ENSMUSG00000028218 | GSMG0027566 | Fam92a | family with sequence similarity 92, member A | 1042.44 | 1383.03 | -0.41 | 6.44E-02 |
| ENSMUSG00000040550 | GSMG0027575 | Otud6b | OTU domain containing 6B | 433.29 | 447.63 | -0.05 | 9.02E-01 |
| ENSMUSG00000040536 | GSMG0027577 | Necab1 | N-terminal EF-hand calcium binding protein 1 | 338.39 | 585.38 | -0.79 | 1.51E-05 |
| ENSMUSG00000028223 | GSMG0027578 | Decr1 | 2,4-dienoyl CoA reductase 1, mitochondrial | 601.19 | 635.18 | -0.08 | 8.26E-01 |
| ENSMUSG00000041135 | GSMG0027581 | Ripk2 | receptor (TNFRSF)-interacting serine-threonine kinase 2 | 343.55 | 279.16 | 0.30 | 2.76E-01 |
| NULL | GSMG0027583 | NULL | NULL | 564.18 | 579.02 | -0.04 | 9.12E-01 |
| ENSMUSG00000028228 | GSMG0027584 | Cpne3 | copine III | 670.26 | 631.87 | 0.09 | 8.04E-01 |
| ENSMUSG00000041058 | GSMG0027587 | Wwp1 | WW domain containing E3 ubiquitin protein ligase 1 | 135.68 | 117.49 | 0.20 | 6.66E-01 |
| ENSMUSG00000040387 | GSMG0027614 | Klhl32 | kelch-like 32 | 182.47 | 250.86 | -0.46 | 8.96E-02 |
| ENSMUSG00000028259 | GSMG0027620 | Fhl5 | four and a half LIM domains 5 | 92.39 | 18.12 | 2.35 | 1.40E-09 |
| ENSMUSG00000040359 | GSMG0027621 | Ufl1 | UFM1 specific ligase 1 | 532.07 | 585.81 | -0.14 | 6.86E-01 |
| ENSMUSG00000040520 | GSMG0027626 | Manea | mannosidase, endo-alpha | 224.93 | 223.92 | 0.00 | 9.96E-01 |
| ENSMUSG00000040183 | GSMG0027638 | Ankrd6 | ankyrin repeat domain 6 | 327.55 | 311.78 | 0.07 | 8.40E-01 |
| ENSMUSG00000040128 | GSMG0027643 | Pnrc1 | proline-rich nuclear receptor coactivator 1 | 2431.37 | 1989.36 | 0.29 | 2.34E-01 |
| ENSMUSG00000040044 | GSMG0027649 | Orc3 | origin recognition complex, subunit 3 | 953.95 | 1156.25 | -0.28 | 1.31E-01 |
| ENSMUSG00000028293 | GSMG0027651 | Slc35a1 | solute carrier family 35 (CMP-sialic acid transporter), member 1 | 721.19 | 642.59 | 0.17 | 4.92E-01 |
| ENSMUSG00000028295 | GSMG0027654 | Smim8 | small integral membrane protein 8 | 272.03 | 304.19 | -0.16 | 6.16E-01 |
| ENSMUSG00000039967 | GSMG0027655 | Zfp292 | zinc finger protein 292 | 152.48 | 189.97 | -0.31 | 4.10E-01 |
| ENSMUSG00000028300 | GSMG0027665 | 3110043O21Rik | RIKEN cDNA 3110043O21 gene | 210.40 | 205.29 | 0.04 | 9.28E-01 |
| NULL | GSMG0027670 | NULL | NULL | 110.30 | 75.26 | 0.55 | 1.56E-01 |
| NULL | GSMG0027676 | NULL | NULL | 763.72 | 729.45 | 0.06 | 8.76E-01 |
| ENSMUSG00000040296 | GSMG0027679 | Ddx58 | DEAD (Asp-Glu-Ala-Asp) box polypeptide 58 | 745.92 | 844.40 | -0.18 | 5.00E-01 |
| ENSMUSG00000036822 | GSMG0027681 | Topors | topoisomerase I binding, arginine/serine-rich | 1878.03 | 2274.09 | -0.28 | 1.10E-01 |
| ENSMUSG00000071014 | GSMG0027682 | Ndufb6 | NADH dehydrogenase (ubiquinone) 1 beta subcomplex, 6 | 1506.66 | 1846.71 | -0.29 | 1.40E-01 |
| ENSMUSG00000028411 | GSMG0027685 | Aptx | aprataxin | 351.51 | 398.87 | -0.18 | 5.23E-01 |
| ENSMUSG00000028409 | GSMG0027687 | Smu1 | smu-1 suppressor of mec-8 and unc-52 homolog (C. elegans) | 2255.54 | 2688.81 | -0.25 | 1.95E-01 |
| ENSMUSG00000028413 | GSMG0027688 | B4galt1 | UDP-Gal:betaGlcNAc beta 1,4- galactosyltransferase, polypeptide 1 | 361.49 | 357.71 | 0.02 | 9.65E-01 |
| ENSMUSG00000028416 | GSMG0027689 | Bag1 | BCL2-associated athanogene 1 | 2088.39 | 2710.87 | -0.38 | 6.47E-02 |
| ENSMUSG00000028430 | GSMG0027692 | Nol6 | nucleolar protein family 6 (RNA-associated) | 1127.86 | 1431.86 | -0.34 | 1.01E-01 |
| NULL | GSMG0027693 | NULL | NULL | 472.21 | 482.89 | -0.03 | 9.29E-01 |

| | | | | | | | |
|---|---|---|---|---|---|---|---|
| ENSMUSG00000028433 | GSMG0027694 | Ubap2 | ubiquitin-associated protein 2 | 1357.38 | 1701.92 | -0.33 | 9.79E-02 |
| ENSMUSG00000028436 | GSMG0027697 | Dcaf12 | DDB1 and CUL4 associated factor 12 | 1293.07 | 1319.41 | -0.03 | 9.24E-01 |
| ENSMUSG00000046312 | GSMG0027700 | AI464131 | expressed sequence AI464131 | 689.23 | 594.83 | 0.21 | 5.25E-01 |
| ENSMUSG00000028441 | GSMG0027701 | 1110017D15Rik | RIKEN cDNA 1110017D15 gene | 118.59 | 124.85 | -0.08 | 8.89E-01 |
| ENSMUSG00000028439 | GSMG0027702 | Fam219a | family with sequence similarity 219, member A | 366.33 | 317.64 | 0.20 | 4.86E-01 |
| NULL | GSMG0027703 | NULL | NULL | 345.82 | 278.57 | 0.32 | 3.55E-01 |
| NULL | GSMG0027704 | NULL | NULL | 316.22 | 288.40 | 0.13 | 7.04E-01 |
| ENSMUSG00000028445 | GSMG0027705 | Enho | energy homeostasis associated | 566.49 | 739.66 | -0.38 | 4.22E-02 |
| ENSMUSG00000028444 | GSMG0027706 | Cntfr | ciliary neurotrophic factor receptor | 178.54 | 137.96 | 0.37 | 4.37E-01 |
| ENSMUSG00000036114 | GSMG0027707 | Rpp25l | ribonuclease P/MRP 25 subunit-like | 425.85 | 474.42 | -0.16 | 5.99E-01 |
| ENSMUSG00000028447 | GSMG0027708 | Dctn3 | dynactin 3 | 1837.06 | 1982.10 | -0.11 | 6.58E-01 |
| ENSMUSG00000036078 | GSMG0027710 | Sigmar1 | sigma non-opioid intracellular receptor 1 | 1778.84 | 1821.82 | -0.03 | 9.05E-01 |
| ENSMUSG00000028453 | GSMG0027721 | Fancg | Fanconi anemia, complementation group G | 283.49 | 304.48 | -0.10 | 7.80E-01 |
| ENSMUSG00000028454 | GSMG0027722 | Pigo | phosphatidylinositol glycan anchor biosynthesis, class O | 1425.13 | 1439.65 | -0.01 | 9.64E-01 |
| ENSMUSG00000028455 | GSMG0027723 | Stoml2 | stomatin (Epb7.2)-like 2 | 1460.87 | 1878.09 | -0.36 | 3.75E-02 |
| ENSMUSG00000036002 | GSMG0027724 | Fam214b | family with sequence similarity 214, member B | 831.78 | 806.48 | 0.04 | 9.03E-01 |
| ENSMUSG00000051517 | GSMG0027728 | Arhgef39 | Rho guanine nucleotide exchange factor (GEF) 39 | 136.32 | 235.97 | -0.79 | 2.65E-03 |
| ENSMUSG00000028464 | GSMG0027729 | Tpm2 | tropomyosin 2, beta | 968.88 | 295.58 | 1.71 | 6.25E-15 |
| ENSMUSG00000028465 | GSMG0027730 | Tln1 | talin 1 | 5535.38 | 5217.93 | 0.09 | 8.19E-01 |
| ENSMUSG00000028467 | GSMG0027731 | Gba2 | glucosidase beta 2 | 2585.33 | 1969.31 | 0.39 | 7.07E-02 |
| ENSMUSG00000066196 | GSMG0027733 | Spag8 | sperm associated antigen 8 | 114.54 | 129.47 | -0.17 | 7.57E-01 |
| ENSMUSG00000028470 | GSMG0027734 | Hint2 | histidine triad nucleotide binding protein 2 | 580.60 | 539.94 | 0.10 | 7.71E-01 |
| ENSMUSG00000028479 | GSMG0027745 | Gne | glucosamine (UDP-N-acetyl)-2-epimerase/N-acetylmannosamine kinase | 1155.72 | 1276.64 | -0.14 | 5.47E-01 |
| ENSMUSG00000035696 | GSMG0027746 | Rnf38 | ring finger protein 38 | 314.31 | 393.02 | -0.32 | 2.58E-01 |
| ENSMUSG00000049657 | GSMG0027756 | Zbtb5 | zinc finger and BTB domain containing 5 | 741.74 | 892.23 | -0.27 | 1.72E-01 |
| ENSMUSG00000048232 | GSMG0027758 | Fbxo10 | F-box protein 10 | 415.82 | 340.76 | 0.29 | 4.10E-01 |
| ENSMUSG00000028322 | GSMG0027762 | Exosc3 | exosome component 3 | 467.27 | 564.58 | -0.27 | 2.33E-01 |
| ENSMUSG00000045973 | GSMG0027763 | Slc25a51 | solute carrier family 25, member 51 | 1318.47 | 1659.42 | -0.33 | 1.85E-01 |
| ENSMUSG00000044813 | GSMG0027764 | Shb | src homology 2 domain-containing transforming protein B | 162.08 | 128.61 | 0.33 | 4.95E-01 |
| ENSMUSG00000035551 | GSMG0027767 | Igfbpl1 | insulin-like growth factor binding protein-like 1 | 40.87 | 284.84 | -2.81 | 2.07E-05 |
| ENSMUSG00000035495 | GSMG0027768 | Tstd2 | thiosulfate sulfurtransferase (rhodanese)-like domain containing 2 | 416.76 | 535.16 | -0.36 | 6.85E-02 |
| ENSMUSG00000028329 | GSMG0027769 | Xpa | xeroderma pigmentosum, complementation group A | 368.12 | 452.49 | -0.30 | 1.98E-01 |
| ENSMUSG00000028331 | GSMG0027772 | Trmo | tRNA methyltransferase O | 149.11 | 133.01 | 0.17 | 6.85E-01 |

| | | | | | | | |
|---|---|---|---|---|---|---|---|
| ENSMUSG00000028337 | GSMG0027775 | Coro2a | coronin, actin binding protein 2A | 24.20 | 54.48 | -1.17 | 1.17E-02 |
| ENSMUSG00000039813 | GSMG0027777 | Tbc1d2 | TBC1 domain family, member 2 | 283.18 | 238.59 | 0.24 | 5.13E-01 |
| ENSMUSG00000039809 | GSMG0027778 | Gabbr2 | gamma-aminobutyric acid (GABA) B receptor, 2 | 391.07 | 339.79 | 0.20 | 5.91E-01 |
| NULL | GSMG0027779 | NULL | NULL | 816.72 | 583.13 | 0.49 | 1.61E-02 |
| ENSMUSG00000066191 | GSMG0027781 | Anks6 | ankyrin repeat and sterile alpha motif domain containing 6 | 810.78 | 655.38 | 0.31 | 2.97E-01 |
| ENSMUSG00000039740 | GSMG0027782 | Alg2 | asparagine-linked glycosylation 2 (alpha-1,3-mannosyltransferase) | 1919.86 | 3354.67 | -0.80 | 3.10E-07 |
| ENSMUSG00000028343 | GSMG0027788 | Erp44 | endoplasmic reticulum protein 44 | 994.11 | 979.25 | 0.02 | 9.54E-01 |
| ENSMUSG00000028345 | GSMG0027790 | Tex10 | testis expressed gene 10 | 927.29 | 1126.70 | -0.28 | 1.49E-01 |
| ENSMUSG00000044018 | GSMG0027797 | Mrpl50 | mitochondrial ribosomal protein L50 | 1005.18 | 1341.66 | -0.42 | 7.47E-02 |
| ENSMUSG00000039611 | GSMG0027800 | Tmem246 | transmembrane protein 246 | 1702.24 | 2251.76 | -0.40 | 2.31E-02 |
| ENSMUSG00000039579 | GSMG0027801 | Grin3a | glutamate receptor ionotropic, NMDA3A | 312.27 | 614.87 | -0.97 | 6.21E-05 |
| ENSMUSG00000015243 | GSMG0027811 | Abca1 | ATP-binding cassette, sub-family A (ABC1), member 1 | 5789.72 | 9933.83 | -0.78 | 1.79E-03 |
| ENSMUSG00000003032 | GSMG0027822 | Klf4 | Kruppel-like factor 4 (gut) | 1204.72 | 585.64 | 1.04 | 4.14E-05 |
| ENSMUSG00000028431 | GSMG0027826 | Ikbkap | inhibitor of kappa light polypeptide enhancer in B cells, kinase complex | 1803.85 | 2312.66 | -0.36 | 9.33E-02 |
| ENSMUSG00000038816 | GSMG0027829 | Ctnnal1 | catenin (cadherin associated protein), alpha-like 1 | 283.40 | 283.00 | 0.00 | 9.99E-01 |
| ENSMUSG00000055296 | GSMG0027832 | Tmem245 | transmembrane protein 245 | 717.57 | 551.75 | 0.38 | 1.12E-01 |
| ENSMUSG00000045589 | GSMG0027834 | Frrs1l | ferric-chelate reductase 1 like | 945.36 | 854.15 | 0.15 | 5.66E-01 |
| ENSMUSG00000028434 | GSMG0027837 | Epb41l4b | erythrocyte membrane protein band 4.1 like 4b | 93.67 | 28.81 | 1.69 | 6.66E-04 |
| ENSMUSG00000028367 | GSMG0027848 | Txn1 | thioredoxin 1 | 2404.99 | 3603.19 | -0.58 | 1.81E-03 |
| ENSMUSG00000038668 | GSMG0027852 | Lpar1 | lysophosphatidic acid receptor 1 | 468.17 | 305.62 | 0.62 | 1.96E-02 |
| ENSMUSG00000050812 | GSMG0027856 | AI314180 | expressed sequence AI314180 | 2354.88 | 2186.56 | 0.11 | 6.50E-01 |
| ENSMUSG00000028378 | GSMG0027858 | Ptgr1 | prostaglandin reductase 1 | 72.51 | 59.34 | 0.30 | 5.79E-01 |
| ENSMUSG00000028382 | GSMG0027863 | Ptbp3 | polypyrimidine tract binding protein 3 | 1043.97 | 1081.94 | -0.05 | 8.83E-01 |
| ENSMUSG00000038544 | GSMG0027870 | Inip | INTS3 and NABP interacting protein | 717.55 | 787.00 | -0.13 | 5.96E-01 |
| ENSMUSG00000028389 | GSMG0027885 | Zfp37 | zinc finger protein 37 | 309.10 | 345.24 | -0.16 | 6.16E-01 |
| ENSMUSG00000066151 | GSMG0027888 | Fkbp15 | FK506 binding protein 15 | 534.86 | 598.18 | -0.16 | 5.56E-01 |
| ENSMUSG00000066149 | GSMG0027889 | Cdc26 | cell division cycle 26 | 1518.46 | 1598.95 | -0.07 | 8.11E-01 |
| ENSMUSG00000028391 | GSMG0027891 | Wdr31 | WD repeat domain 31 | 86.23 | 101.11 | -0.23 | 6.33E-01 |
| ENSMUSG00000038422 | GSMG0027892 | Hdhd3 | haloacid dehalogenase-like hydrolase domain containing 3 | 632.21 | 476.41 | 0.41 | 3.72E-02 |
| ENSMUSG00000028393 | GSMG0027893 | Alad | aminolevulinate, delta-, dehydratase | 3047.68 | 3639.40 | -0.26 | 1.42E-01 |
| ENSMUSG00000028394 | GSMG0027894 | Pole3 | polymerase (DNA directed), epsilon 3 (p17 subunit) | 1045.54 | 1323.33 | -0.34 | 1.08E-01 |
| NULL | GSMG0027895 | NULL | NULL | 97.51 | 81.01 | 0.26 | 5.60E-01 |
| ENSMUSG00000039158 | GSMG0027898 | Akna | AT-hook transcription factor | 510.25 | 470.71 | 0.12 | 7.39E-01 |

| | | | | | | | |
|---|---|---|---|---|---|---|---|
| ENSMUSG00000039137 | GSMG0027899 | Whrn | whirlin | 472.35 | 264.49 | 0.84 | 4.00E-04 |
| NULL | GSMG0027901 | NULL | NULL | 68.63 | 46.42 | 0.57 | 2.30E-01 |
| ENSMUSG00000028364 | GSMG0027906 | Tnc | tenascin C | 127.27 | 734.07 | -2.53 | 6.58E-31 |
| ENSMUSG00000028373 | GSMG0027913 | Astn2 | astrotactin 2 | 341.63 | 220.23 | 0.63 | 3.47E-02 |
| ENSMUSG00000028351 | GSMG0027925 | Brinp1 | bone morphogenic protein/retinoic acid inducible neural specific 1 | 669.39 | 712.66 | -0.09 | 7.63E-01 |
| NULL | GSMG0027926 | NULL | NULL | 229.29 | 153.47 | 0.58 | 3.77E-02 |
| NULL | GSMG0027927 | NULL | NULL | 340.95 | 168.43 | 1.02 | 2.06E-05 |
| ENSMUSG00000039298 | GSMG0027931 | Cdk5rap2 | CDK5 regulatory subunit associated protein 2 | 609.61 | 1117.13 | -0.87 | 1.55E-07 |
| ENSMUSG00000039270 | GSMG0027932 | Megf9 | multiple EGF-like-domains 9 | 145.54 | 228.51 | -0.65 | 3.95E-02 |
| ENSMUSG00000008305 | GSMG0027935 | Tle1 | transducin-like enhancer of split 1 | 1814.33 | 1619.61 | 0.16 | 4.42E-01 |
| ENSMUSG00000059343 | GSMG0027936 | Aldoart1 | aldolase 1 A, retrogene 1 | 250.70 | 192.68 | 0.38 | 1.73E-01 |
| ENSMUSG00000028398 | GSMG0027940 | Tmem261 | transmembrane protein 261 | 294.98 | 366.27 | -0.31 | 2.36E-01 |
| NULL | GSMG0027944 | NULL | NULL | 77.04 | 82.84 | -0.11 | 8.51E-01 |
| NULL | GSMG0027945 | NULL | NULL | 464.13 | 441.65 | 0.07 | 8.78E-01 |
| NULL | GSMG0027946 | NULL | NULL | 355.08 | 320.41 | 0.15 | 7.01E-01 |
| NULL | GSMG0027947 | NULL | NULL | 396.71 | 399.26 | -0.01 | 9.89E-01 |
| NULL | GSMG0027949 | NULL | NULL | 284.35 | 274.05 | 0.06 | 8.97E-01 |
| NULL | GSMG0027950 | NULL | NULL | 132.86 | 130.19 | 0.03 | 9.56E-01 |
| NULL | GSMG0027951 | NULL | NULL | 341.00 | 303.67 | 0.17 | 6.47E-01 |
| NULL | GSMG0027952 | NULL | NULL | 308.89 | 289.07 | 0.10 | 7.94E-01 |
| NULL | GSMG0027954 | NULL | NULL | 346.03 | 371.58 | -0.10 | 8.49E-01 |
| NULL | GSMG0027955 | NULL | NULL | 375.54 | 358.64 | 0.07 | 8.77E-01 |
| NULL | GSMG0027956 | NULL | NULL | 696.49 | 554.65 | 0.33 | 2.30E-01 |
| NULL | GSMG0027960 | NULL | NULL | 401.25 | 374.36 | 0.10 | 8.13E-01 |
| NULL | GSMG0027963 | NULL | NULL | 346.49 | 326.19 | 0.09 | 8.67E-01 |
| NULL | GSMG0027964 | NULL | NULL | 234.21 | 217.54 | 0.11 | 8.16E-01 |
| NULL | GSMG0027967 | NULL | NULL | 324.64 | 309.61 | 0.07 | 8.60E-01 |
| NULL | GSMG0027968 | NULL | NULL | 408.96 | 353.57 | 0.21 | 5.31E-01 |
| NULL | GSMG0027969 | NULL | NULL | 386.07 | 319.21 | 0.27 | 4.01E-01 |
| NULL | GSMG0027970 | NULL | NULL | 800.66 | 654.88 | 0.29 | 3.29E-01 |
| NULL | GSMG0027973 | NULL | NULL | 347.38 | 292.55 | 0.25 | 4.87E-01 |
| ENSMUSG00000028402 | GSMG0027976 | Mpdz | multiple PDZ domain protein | 1292.50 | 1600.61 | -0.31 | 1.48E-01 |
| ENSMUSG00000008575 | GSMG0027989 | Nfib | nuclear factor I/B | 2498.73 | 3013.60 | -0.27 | 1.14E-01 |

| | | | | | | | |
|---|---|---|---|---|---|---|---|
| ENSMUSG00000028403 | GSMG0027996 | Zdhhc21 | zinc finger, DHHC domain containing 21 | 100.31 | 108.79 | -0.12 | 8.05E-01 |
| ENSMUSG00000059049 | GSMG0027998 | Frem1 | Fras1 related extracellular matrix protein 1 | 21.66 | 80.32 | -1.89 | 3.46E-06 |
| ENSMUSG00000038172 | GSMG0027999 | Ttc39b | tetratricopeptide repeat domain 39B | 536.49 | 848.38 | -0.66 | 1.38E-04 |
| ENSMUSG00000028484 | GSMG0028002 | Psip1 | PC4 and SFRS1 interacting protein 1 | 748.83 | 1346.94 | -0.85 | 4.13E-05 |
| ENSMUSG00000038047 | GSMG0028016 | Haus6 | HAUS augmin-like complex, subunit 6 | 139.60 | 204.16 | -0.55 | 3.77E-02 |
| ENSMUSG00000028494 | GSMG0028017 | Plin2 | perilipin 2 | 1120.52 | 1425.26 | -0.35 | 8.57E-02 |
| ENSMUSG00000037996 | GSMG0028023 | Slc24a2 | solute carrier family 24 (sodium/potassium/calcium exchanger), membe | 450.70 | 162.03 | 1.48 | 1.07E-11 |
| NULL | GSMG0028027 | NULL | NULL | 87.11 | 31.93 | 1.46 | 3.47E-04 |
| ENSMUSG00000028496 | GSMG0028030 | Mllt3 | myeloid/lymphoid or mixed-lineage leukemia%3B translocated to, 3 | 1362.94 | 1394.01 | -0.03 | 9.27E-01 |
| ENSMUSG00000028497 | GSMG0028035 | Hacd4 | 3-hydroxyacyl-CoA dehydratase 4 | 215.98 | 148.28 | 0.55 | 6.64E-02 |
| ENSMUSG00000070923 | GSMG0028048 | Klhl9 | kelch-like 9 | 1565.62 | 1742.80 | -0.16 | 5.28E-01 |
| ENSMUSG00000073802 | GSMG0028057 | Cdkn2b | cyclin-dependent kinase inhibitor 2B (p15, inhibits CDK4) | 3011.22 | 1903.45 | 0.66 | 4.67E-05 |
| ENSMUSG00000008489 | GSMG0028061 | Elavl2 | ELAV (embryonic lethal, abnormal vision, Drosophila)-like 2 (Hu antig | 259.13 | 374.29 | -0.53 | 1.74E-02 |
| ENSMUSG00000054000 | GSMG0028067 | Tusc1 | tumor suppressor candidate 1 | 146.23 | 178.78 | -0.30 | 4.26E-01 |
| ENSMUSG00000028578 | GSMG0028071 | Caap1 | caspase activity and apoptosis inhibitor 1 | 505.14 | 457.66 | 0.14 | 6.88E-01 |
| ENSMUSG00000028577 | GSMG0028075 | Plaa | phospholipase A2, activating protein | 446.44 | 463.43 | -0.05 | 8.75E-01 |
| ENSMUSG00000062627 | GSMG0028078 | Mysm1 | myb-like, SWIRM and MPN domains 1 | 419.44 | 395.10 | 0.08 | 8.26E-01 |
| NULL | GSMG0028080 | NULL | NULL | 5130.35 | 2203.34 | 1.22 | 1.85E-15 |
| ENSMUSG00000052914 | GSMG0028088 | Cyp2j6 | cytochrome P450, family 2, subfamily j, polypeptide 6 | 584.53 | 549.86 | 0.09 | 7.77E-01 |
| ENSMUSG00000015224 | GSMG0028090 | Cyp2j9 | cytochrome P450, family 2, subfamily j, polypeptide 9 | 196.31 | 97.75 | 1.01 | 1.40E-04 |
| ENSMUSG00000028563 | GSMG0028099 | Tm2d1 | TM2 domain containing 1 | 653.60 | 713.32 | -0.13 | 7.19E-01 |
| ENSMUSG00000028556 | GSMG0028104 | Dock7 | dedicator of cytokinesis 7 | 620.54 | 1014.18 | -0.71 | 1.03E-05 |
| ENSMUSG00000028549 | GSMG0028109 | Itgb3bp | integrin beta 3 binding protein (beta3-endonexin) | 99.65 | 68.62 | 0.53 | 1.94E-01 |
| ENSMUSG00000028530 | GSMG0028118 | Jak1 | Janus kinase 1 | 3347.91 | 4059.03 | -0.28 | 1.10E-01 |
| ENSMUSG00000086782 | GSMG0028119 | E130102H24Rik | RIKEN cDNA E130102H24 gene | 124.50 | 71.91 | 0.79 | 1.33E-02 |
| NULL | GSMG0028130 | NULL | NULL | 73.52 | 69.19 | 0.09 | 8.84E-01 |
| ENSMUSG00000028521 | GSMG0028131 | Slc35d1 | solute carrier family 35 (UDP-glucuronic acid/UDP-N-acetylgalactosan | 200.91 | 145.94 | 0.46 | 9.73E-02 |
| ENSMUSG00000028518 | GSMG0028137 | Prkaa2 | protein kinase, AMP-activated, alpha 2 catalytic subunit | 163.65 | 220.00 | -0.42 | 1.69E-01 |
| NULL | GSMG0028139 | NULL | NULL | 161.43 | 133.62 | 0.27 | 4.57E-01 |
| ENSMUSG00000025413 | GSMG0028147 | Ttc4 | tetratricopeptide repeat domain 4 | 1240.26 | 1265.28 | -0.03 | 9.28E-01 |
| ENSMUSG00000028622 | GSMG0028151 | Mrpl37 | mitochondrial ribosomal protein L37 | 1035.74 | 1176.31 | -0.18 | 4.06E-01 |
| ENSMUSG00000028617 | GSMG0028154 | Lrrc42 | leucine rich repeat containing 42 | 346.69 | 392.35 | -0.18 | 5.23E-01 |
| ENSMUSG00000028607 | GSMG0028162 | Cpt2 | carnitine palmitoyltransferase 2 | 967.97 | 758.10 | 0.35 | 1.02E-01 |

| | | | | | | | |
|---|---|---|---|---|---|---|---|
| ENSMUSG00000028603 | GSMG0028165 | Scp2 | sterol carrier protein 2, liver | 1092.82 | 1106.55 | -0.02 | 9.61E-01 |
| ENSMUSG00000034636 | GSMG0028170 | Zyg11b | zyg-ll family member B, cell cycle regulator | 705.13 | 885.59 | -0.33 | 1.08E-01 |
| ENSMUSG00000028597 | GSMG0028172 | Gpx7 | glutathione peroxidase 7 | 1384.45 | 1148.49 | 0.27 | 2.20E-01 |
| ENSMUSG00000063800 | GSMG0028174 | Prpf38a | PRP38 pre-mRNA processing factor 38 (yeast) domain containing A | 903.57 | 1016.67 | -0.17 | 5.01E-01 |
| NULL | GSMG0028181 | NULL | NULL | 620.68 | 838.59 | -0.43 | 2.36E-02 |
| ENSMUSG00000028559 | GSMG0028183 | Osbpl9 | oxysterol binding protein-like 9 | 2911.60 | 2008.42 | 0.54 | 2.12E-03 |
| NULL | GSMG0028184 | NULL | NULL | 244.82 | 284.45 | -0.22 | 5.00E-01 |
| NULL | GSMG0028185 | NULL | NULL | 58.02 | 59.26 | -0.03 | 9.67E-01 |
| ENSMUSG00000028557 | GSMG0028188 | Rnf11 | ring finger protein 11 | 3759.76 | 3806.48 | -0.02 | 9.51E-01 |
| ENSMUSG00000028551 | GSMG0028192 | Cdkn2c | cyclin-dependent kinase inhibitor 2C (p18, inhibits CDK4) | 150.94 | 186.54 | -0.30 | 3.74E-01 |
| ENSMUSG00000028546 | GSMG0028195 | Elavl4 | ELAV (embryonic lethal, abnormal vision, Drosophila)-like 4 (Hu antig | 179.65 | 301.68 | -0.75 | 5.24E-04 |
| NULL | GSMG0028196 | NULL | NULL | 269.74 | 209.74 | 0.36 | 2.85E-01 |
| ENSMUSG00000028719 | GSMG0028211 | Cmpk1 | cytidine monophosphate (UMP-CMP) kinase 1 | 1042.69 | 1251.05 | -0.26 | 3.18E-01 |
| ENSMUSG00000047155 | GSMG0028214 | Cyp4x1 | cytochrome P450, family 4, subfamily x, polypeptide 1 | 141.58 | 277.31 | -0.97 | 9.38E-05 |
| ENSMUSG00000034171 | GSMG0028222 | Faah | fatty acid amide hydrolase | 1206.44 | 852.64 | 0.50 | 3.87E-02 |
| ENSMUSG00000028706 | GSMG0028224 | Nsun4 | NOL1/NOP2/Sun domain family, member 4 | 208.49 | 211.25 | -0.02 | 9.64E-01 |
| ENSMUSG00000063882 | GSMG0028225 | Uqcrh | ubiquinol-cytochrome c reductase hinge protein | 1968.54 | 1995.86 | -0.02 | 9.50E-01 |
| ENSMUSG00000028702 | GSMG0028226 | Rad54l | RAD54 like (S. cerevisiae) | 84.97 | 213.95 | -1.33 | 6.37E-08 |
| ENSMUSG00000028701 | GSMG0028229 | Lurap1 | leucine rich adaptor protein 1 | 204.06 | 145.64 | 0.49 | 7.25E-02 |
| ENSMUSG00000003810 | GSMG0028231 | Mast2 | microtubule associated serine/threonine kinase 2 | 1095.67 | 1447.65 | -0.40 | 1.39E-01 |
| NULL | GSMG0028234 | NULL | NULL | 337.60 | 418.69 | -0.31 | 2.58E-01 |
| ENSMUSG00000055900 | GSMG0028235 | Tmem69 | transmembrane protein 69 | 114.41 | 138.00 | -0.27 | 4.85E-01 |
| ENSMUSG00000028693 | GSMG0028237 | Nasp | nuclear autoantigenic sperm protein (histone-binding) | 873.06 | 1520.21 | -0.80 | 5.16E-04 |
| ENSMUSG00000028692 | GSMG0028238 | Akr1a1 | aldo-keto reductase family 1, member A1 (aldehyde reductase) | 7298.12 | 7592.41 | -0.06 | 8.26E-01 |
| ENSMUSG00000028688 | GSMG0028243 | Toe1 | target of EGR1, member 1 (nuclear) | 118.61 | 97.88 | 0.28 | 5.21E-01 |
| ENSMUSG00000028684 | GSMG0028245 | Urod | uroporphyrinogen decarboxylase | 1115.38 | 1125.02 | -0.01 | 9.68E-01 |
| NULL | GSMG0028246 | NULL | NULL | 111.91 | 118.23 | -0.08 | 8.77E-01 |
| ENSMUSG00000028680 | GSMG0028248 | Plk3 | polo-like kinase 3 | 2075.70 | 1367.31 | 0.60 | 6.73E-04 |
| ENSMUSG00000028678 | GSMG0028250 | Kif2c | kinesin family member 2C | 184.21 | 427.47 | -1.21 | 1.19E-08 |
| ENSMUSG00000028677 | GSMG0028253 | Rnf220 | ring finger protein 220 | 889.16 | 949.94 | -0.10 | 7.64E-01 |
| ENSMUSG00000009640 | GSMG0028256 | Dmap1 | DNA methyltransferase 1-associated protein 1 | 1145.41 | 1336.85 | -0.22 | 2.62E-01 |
| G00000028541,ENSMUSG000 | GSMG0028258 | Ccdc24 // B4galt2 | coiled-coil domain containing 24 // UDP-Gal:betaGlcNAc beta 1,4- gal | 642.49 | 985.59 | -0.62 | 2.78E-03 |
| ENSMUSG00000033379 | GSMG0028259 | Atp6v0b | ATPase, H+ transporting, lysosomal V0 subunit B | 7057.82 | 5174.81 | 0.45 | 2.55E-03 |

| | | | | | | | |
|---|---|---|---|---|---|---|---|
| ENSMUSG00000028540 | GSMG0028260 | Dph2 | DPH2 homolog | 213.90 | 286.11 | -0.41 | 1.50E-01 |
| ENSMUSG00000033365 | GSMG0028261 | Ipo13 | importin 13 | 2456.56 | 2346.76 | 0.07 | 8.42E-01 |
| ENSMUSG00000028538 | GSMG0028263 | St3gal3 | ST3 beta-galactoside alpha-2,3-sialyltransferase 3 | 197.68 | 227.41 | -0.20 | 6.18E-01 |
| ENSMUSG00000033326 | GSMG0028265 | Kdm4a | lysine (K)-specific demethylase 4A | 842.76 | 1079.83 | -0.36 | 1.38E-01 |
| ENSMUSG00000033295 | GSMG0028266 | Ptprf | protein tyrosine phosphatase, receptor type, F | 542.76 | 876.46 | -0.69 | 5.19E-03 |
| NULL | GSMG0028267 | NULL | NULL | 311.87 | 472.96 | -0.60 | 3.12E-02 |
| ENSMUSG00000033253 | GSMG0028268 | Szt2 | seizure threshold 2 | 755.58 | 661.19 | 0.19 | 5.00E-01 |
| ENSMUSG00000006398 | GSMG0028269 | Cdc20 | cell division cycle 20 | 678.28 | 1284.49 | -0.92 | 1.14E-06 |
| ENSMUSG00000033191 | GSMG0028271 | Tie1 | tyrosine kinase with immunoglobulin-like and EGF-like domains 1 | 14161.11 | 14780.08 | -0.06 | 8.13E-01 |
| NULL | GSMG0028274 | NULL | NULL | 87.43 | 100.92 | -0.21 | 6.89E-01 |
| ENSMUSG00000045268 | GSMG0028284 | Zfp691 | zinc finger protein 691 | 413.84 | 402.45 | 0.04 | 9.23E-01 |
| G00000028642,ENSMUSG000 | GSMG0028286 | 4930538K18Rik // | RIKEN cDNA 4930538K18 gene // expressed sequence AU022252 | 476.36 | 505.79 | -0.09 | 7.79E-01 |
| ENSMUSG00000028639 | GSMG0028288 | Ybx1 | Y box protein 1 | 4245.21 | 6894.02 | -0.70 | 7.50E-07 |
| ENSMUSG00000060288 | GSMG0028289 | Ppih | peptidyl prolyl isomerase H | 168.52 | 346.51 | -1.04 | 6.70E-06 |
| ENSMUSG00000028637 | GSMG0028290 | Ccdc30 | coiled-coil domain containing 30 | 105.39 | 117.28 | -0.16 | 7.36E-01 |
| ENSMUSG00000028636 | GSMG0028291 | Ppcs | phosphopantothenoylcysteine synthetase | 129.47 | 141.59 | -0.13 | 7.95E-01 |
| ENSMUSG00000048899 | GSMG0028292 | Rimkla | ribosomal modification protein rimK-like family member A | 188.73 | 354.12 | -0.90 | 2.17E-04 |
| ENSMUSG00000028633 | GSMG0028297 | Ctps | cytidine 5'-triphosphate synthase | 1103.19 | 1246.39 | -0.17 | 5.35E-01 |
| ENSMUSG00000028631 | GSMG0028298 | Kcnq4 | potassium voltage-gated channel, subfamily Q, member 4 | 175.83 | 274.70 | -0.65 | 5.47E-03 |
| ENSMUSG00000028629 | GSMG0028302 | Exo5 | exonuclease 5 | 100.05 | 106.37 | -0.09 | 8.63E-01 |
| ENSMUSG00000064141 | GSMG0028303 | Zfp69 | zinc finger protein 69 | 450.18 | 799.99 | -0.83 | 5.47E-07 |
| ENSMUSG00000032870 | GSMG0028304 | Smap2 | small ArfGAP 2 | 533.27 | 629.27 | -0.24 | 4.35E-01 |
| ENSMUSG00000043207 | GSMG0028306 | Zmpste24 | zinc metallopeptidase, STE24 | 1094.23 | 1134.82 | -0.05 | 8.76E-01 |
| ENSMUSG00000049878 | GSMG0028308 | Rlf | rearranged L-myc fusion sequence | 273.48 | 347.01 | -0.34 | 2.06E-01 |
| NULL | GSMG0028309 | NULL | NULL | 298.37 | 325.19 | -0.13 | 7.53E-01 |
| ENSMUSG00000028656 | GSMG0028312 | Cap1 | CAP, adenylate cyclase-associated protein 1 (yeast) | 1595.38 | 3593.03 | -1.17 | 3.12E-15 |
| ENSMUSG00000028655 | GSMG0028313 | Mfsd2a | major facilitator superfamily domain containing 2A | 21371.45 | 25547.00 | -0.26 | 1.12E-01 |
| ENSMUSG00000028651 | GSMG0028315 | Ppie | peptidylprolyl isomerase E (cyclophilin E) | 497.73 | 559.90 | -0.17 | 5.12E-01 |
| ENSMUSG00000028649 | GSMG0028316 | Macf1 | microtubule-actin crosslinking factor 1 | 11495.19 | 11958.04 | -0.06 | 8.04E-01 |
| NULL | GSMG0028321 | D830031N03Rik | RIKEN cDNA D830031N03 gene | 1057.76 | 1148.06 | -0.12 | 7.21E-01 |
| NULL | GSMG0028323 | NULL | NULL | 486.26 | 549.29 | -0.17 | 5.51E-01 |
| ENSMUSG00000023075 | GSMG0028325 | Akirin1 | akirin 1 | 2053.60 | 1945.22 | 0.08 | 7.56E-01 |
| NULL | GSMG0028330 | NULL | NULL | 1040.07 | 918.43 | 0.18 | 4.05E-01 |

| | | | | | | | |
|---|---|---|---|---|---|---|---|
| ENSMUSG00000028907 | GSMG0028336 | Utp11l | UTP11-like, U3 small nucleolar ribonucleoprotein, (yeast) | 781.89 | 878.09 | -0.17 | 5.82E-01 |
| ENSMUSG00000078570 | GSMG0028340 | 1110065P20Rik | RIKEN cDNA 1110065P20 gene | 141.04 | 140.75 | 0.00 | 1.00E+00 |
| ENSMUSG00000042763 | GSMG0028341 | Maneal | mannosidase, endo-alpha-like | 845.29 | 868.88 | -0.04 | 9.31E-01 |
| ENSMUSG00000028873 | GSMG0028343 | Cdca8 | cell division cycle associated 8 | 351.46 | 928.80 | -1.40 | 1.31E-13 |
| ENSMUSG00000073758 | GSMG0028348 | Sh3d21 | SH3 domain containing 21 | 344.36 | 261.98 | 0.39 | 1.06E-01 |
| ENSMUSG00000043962 | GSMG0028349 | Thrap3 | thyroid hormone receptor associated protein 3 | 3366.67 | 4363.40 | -0.37 | 1.94E-02 |
| ENSMUSG00000028849 | GSMG0028350 | Map7d1 | MAP7 domain containing 1 | 5137.01 | 4713.88 | 0.12 | 6.78E-01 |
| ENSMUSG00000042558 | GSMG0028352 | Adprhl2 | ADP-ribosylhydrolase like 2 | 610.06 | 696.45 | -0.19 | 4.93E-01 |
| ENSMUSG00000028845 | GSMG0028353 | Tekt2 | tektin 2 | 104.44 | 205.23 | -0.97 | 7.77E-04 |
| ENSMUSG00000028842 | GSMG0028355 | Ago3 | argonaute RISC catalytic subunit 3 | 480.98 | 322.94 | 0.58 | 8.04E-03 |
| ENSMUSG00000041530 | GSMG0028360 | Ago1 | argonaute RISC catalytic subunit 1 | 251.01 | 315.71 | -0.33 | 2.81E-01 |
| ENSMUSG00000042500 | GSMG0028361 | Ago4 | argonaute RISC catalytic subunit 4 | 180.51 | 185.37 | -0.04 | 9.45E-01 |
| ENSMUSG00000028833 | GSMG0028363 | Ncdn | neurochondrin | 718.01 | 664.67 | 0.11 | 7.65E-01 |
| ENSMUSG00000042446 | GSMG0028364 | Zmym4 | zinc finger, MYM-type 4 | 951.33 | 1039.35 | -0.13 | 6.40E-01 |
| ENSMUSG00000043872 | GSMG0028368 | Zmym1 | zinc finger, MYM domain containing 1 | 87.05 | 91.68 | -0.07 | 9.00E-01 |
| ENSMUSG00000050234 | GSMG0028369 | Gja4 | gap junction protein, alpha 4 | 5741.96 | 4214.75 | 0.45 | 6.35E-03 |
| ENSMUSG00000062545 | GSMG0028381 | Tlr12 | toll-like receptor 12 | 664.56 | 1535.00 | -1.21 | 1.49E-12 |
| ENSMUSG00000028799 | GSMG0028382 | Zfp362 | zinc finger protein 362 | 935.97 | 950.16 | -0.02 | 9.56E-01 |
| ENSMUSG00000028789 | GSMG0028384 | Azin2 | antizyme inhibitor 2 | 254.57 | 216.22 | 0.24 | 4.24E-01 |
| ENSMUSG00000028785 | GSMG0028387 | Hpca | hippocalcin | 451.83 | 407.46 | 0.15 | 7.40E-01 |
| ENSMUSG00000040928 | GSMG0028389 | S100pbp | S100P binding protein | 131.33 | 134.53 | -0.04 | 9.34E-01 |
| ENSMUSG00000050390 | GSMG0028390 | C77080 | expressed sequence C77080 | 603.70 | 446.15 | 0.43 | 1.46E-01 |
| ENSMUSG00000057236 | GSMG0028394 | Rbbp4 | retinoblastoma binding protein 4 | 3352.70 | 4319.03 | -0.37 | 1.54E-01 |
| ENSMUSG00000028807 | GSMG0028395 | Zbtb8a | zinc finger and BTB domain containing 8a | 234.66 | 240.13 | -0.03 | 9.40E-01 |
| ENSMUSG00000028798 | GSMG0028404 | Eif3i | eukaryotic translation initiation factor 3, subunit I | 1520.98 | 1911.75 | -0.33 | 6.96E-02 |
| ENSMUSG00000040795 | GSMG0028406 | Iqcc | IQ motif containing C | 350.14 | 342.88 | 0.03 | 9.38E-01 |
| ENSMUSG00000028795 | GSMG0028407 | Ccdc28b | coiled coil domain containing 28B | 1059.76 | 1241.71 | -0.23 | 2.62E-01 |
| ENSMUSG00000053841 | GSMG0028409 | Txlna | taxilin alpha | 4062.15 | 4124.35 | -0.02 | 9.37E-01 |
| ENSMUSG00000003731 | GSMG0028410 | Kpna6 | karyopherin (importin) alpha 6 | 712.90 | 781.92 | -0.14 | 6.22E-01 |
| ENSMUSG00000053730 | GSMG0028411 | Tmem39b | transmembrane protein 39b | 254.83 | 346.64 | -0.44 | 1.08E-01 |
| ENSMUSG00000028790 | GSMG0028412 | Khdrbs1 | KH domain containing, RNA binding, signal transduction associated 1 | 1099.13 | 1450.95 | -0.40 | 2.22E-02 |
| NULL | GSMG0028413 | NULL | NULL | 523.98 | 635.72 | -0.28 | 4.27E-01 |
| ENSMUSG00000028778 | GSMG0028417 | Hcrtr1 | hypocretin (orexin) receptor 1 | 154.69 | 134.79 | 0.20 | 6.89E-01 |

| | | | | | | | |
|---|---|---|---|---|---|---|---|
| ENSMUSG00000028776 | GSMG0028418 | Tinagl1 | tubulointerstitial nephritis antigen-like 1 | 1320.00 | 413.31 | 1.68 | 8.18E-19 |
| ENSMUSG00000023232 | GSMG0028420 | Serinc2 | serine incorporator 2 | 72.86 | 158.76 | -1.12 | 1.31E-04 |
| ENSMUSG00000028772 | GSMG0028423 | Zcchc17 | zinc finger, CCHC domain containing 17 | 930.11 | 1100.10 | -0.24 | 2.53E-01 |
| ENSMUSG00000028909 | GSMG0028431 | Ptpru | protein tyrosine phosphatase, receptor type, U | 419.72 | 413.63 | 0.02 | 9.69E-01 |
| ENSMUSG00000085667 | GSMG0028434 | Gm12992 | predicted gene 12992 | 673.68 | 703.16 | -0.06 | 8.37E-01 |
| ENSMUSG00000028906 | GSMG0028435 | Epb41 | erythrocyte membrane protein band 4.1 | 2945.89 | 3669.09 | -0.32 | 1.13E-01 |
| ENSMUSG00000040025 | GSMG0028438 | Ythdf2 | YTH domain family 2 | 1722.79 | 2313.42 | -0.42 | 1.06E-02 |
| ENSMUSG00000028901 | GSMG0028441 | Gmeb1 | glucocorticoid modulatory element binding protein 1 | 96.65 | 113.93 | -0.24 | 5.97E-01 |
| ENSMUSG00000028898 | GSMG0028443 | Trnau1ap | tRNA selenocysteine 1 associated protein 1 | 884.34 | 924.64 | -0.06 | 8.46E-01 |
| ENSMUSG00000028896 | GSMG0028444 | Rcc1 | regulator of chromosome condensation 1 | 322.44 | 408.23 | -0.34 | 1.38E-01 |
| ENSMUSG00000085241 | GSMG0028445 | Snhg3 | small nucleolar RNA host gene 3 | 167.45 | 136.99 | 0.29 | 3.75E-01 |
| ENSMUSG00000066043 | GSMG0028446 | Phactr4 | phosphatase and actin regulator 4 | 827.03 | 684.92 | 0.27 | 3.09E-01 |
| ENSMUSG00000066042 | GSMG0028448 | Med18 | mediator complex subunit 18 | 182.33 | 232.38 | -0.34 | 2.33E-01 |
| ENSMUSG00000028893 | GSMG0028450 | Sesn2 | sestrin 2 | 219.96 | 240.10 | -0.13 | 7.94E-01 |
| ENSMUSG00000054428 | GSMG0028451 | Atpif1 | ATPase inhibitory factor 1 | 2796.51 | 4026.04 | -0.53 | 6.34E-03 |
| ENSMUSG00000028885 | GSMG0028455 | Smpdl3b | sphingomyelin phosphodiesterase, acid-like 3B | 127.30 | 129.65 | -0.02 | 9.70E-01 |
| ENSMUSG00000028882 | GSMG0028457 | Ppp1r8 | protein phosphatase 1, regulatory (inhibitor) subunit 8 | 308.21 | 386.92 | -0.33 | 2.44E-01 |
| ENSMUSG00000028879 | GSMG0028458 | Stx12 | syntaxin 12 | 1358.88 | 1490.96 | -0.13 | 6.38E-01 |
| ENSMUSG00000028857 | GSMG0028459 | Tmem222 | transmembrane protein 222 | 1586.28 | 1792.83 | -0.18 | 4.26E-01 |
| ENSMUSG00000028878 | GSMG0028461 | Fam76a | family with sequence similarity 76, member A | 975.70 | 838.65 | 0.22 | 3.11E-01 |
| ENSMUSG00000049649 | GSMG0028463 | Gpr3 | G-protein coupled receptor 3 | 186.71 | 44.92 | 2.05 | 2.47E-12 |
| ENSMUSG00000056596 | GSMG0028468 | Trnp1 | TMF1-regulated nuclear protein 1 | 559.28 | 226.14 | 1.30 | 3.41E-09 |
| ENSMUSG00000028851 | GSMG0028470 | Nudc | nudC nuclear distribution protein | 1964.63 | 2702.56 | -0.46 | 3.91E-03 |
| ENSMUSG00000037553 | GSMG0028472 | Zdhhc18 | zinc finger, DHHC domain containing 18 | 579.41 | 367.74 | 0.65 | 1.61E-02 |
| ENSMUSG00000043257 | GSMG0028473 | Pigv | phosphatidylinositol glycan anchor biosynthesis, class V | 149.72 | 176.59 | -0.24 | 5.09E-01 |
| ENSMUSG00000007880 | GSMG0028474 | Arid1a | AT rich interactive domain 1A (SWI-like) | 815.70 | 870.69 | -0.10 | 8.03E-01 |
| ENSMUSG00000003644 | GSMG0028477 | Rps6ka1 | ribosomal protein S6 kinase polypeptide 1 | 718.13 | 1005.40 | -0.49 | 9.16E-03 |
| ENSMUSG00000012117 | GSMG0028479 | Dhdds | dehydrodolichyl diphosphate synthase | 681.59 | 687.44 | -0.01 | 9.68E-01 |
| ENSMUSG00000000682 | GSMG0028481 | Cd52 | CD52 antigen | 26.64 | 54.92 | -1.03 | 3.52E-02 |
| ENSMUSG00000028843 | GSMG0028482 | Sh3bgrl3 | SH3 domain binding glutamic acid-rich protein-like 3 | 3646.60 | 4464.89 | -0.29 | 9.77E-02 |
| ENSMUSG00000037443 | GSMG0028483 | Cep85 | centrosomal protein 85 | 505.06 | 612.67 | -0.28 | 2.20E-01 |
| ENSMUSG00000028840 | GSMG0028487 | Zfp593 | zinc finger protein 593 | 260.18 | 309.39 | -0.25 | 3.74E-01 |
| ENSMUSG00000050105 | GSMG0028488 | Grrp1 | glycine/arginine rich protein 1 | 1153.62 | 960.63 | 0.26 | 2.96E-01 |

| | | | | | | | |
|---|---|---|---|---|---|---|---|
| ENSMUSG00000050890 | GSMG0028489 | Pdik1l | PDLIM1 interacting kinase 1 like | 789.35 | 954.19 | -0.28 | 2.23E-01 |
| ENSMUSG00000028838 | GSMG0028490 | Extl1 | exostoses (multiple)-like 1 | 245.97 | 50.75 | 2.28 | 1.95E-16 |
| ENSMUSG00000046671 | GSMG0028492 | Mtfr1l | mitochondrial fission regulator 1-like | 2828.82 | 3206.77 | -0.18 | 3.67E-01 |
| ENSMUSG00000050989 | GSMG0028493 | Sepn1 | selenoprotein N, 1 | 1278.56 | 825.64 | 0.63 | 3.33E-02 |
| ENSMUSG00000037306 | GSMG0028494 | Man1c1 | mannosidase, alpha, class 1C, member 1 | 1112.43 | 1020.00 | 0.12 | 7.76E-01 |
| ENSMUSG00000037295 | GSMG0028496 | Ldlrap1 | low density lipoprotein receptor adaptor protein 1 | 588.59 | 571.31 | 0.04 | 8.90E-01 |
| ENSMUSG00000028826 | GSMG0028497 | Tmem57 | transmembrane protein 57 | 1319.65 | 1759.43 | -0.42 | 1.27E-02 |
| ENSMUSG00000028822 | GSMG0028499 | Tmem50a | transmembrane protein 50A | 4847.72 | 5052.61 | -0.06 | 8.47E-01 |
| ENSMUSG00000037242 | GSMG0028502 | Clic4 | chloride intracellular channel 4 (mitochondrial) | 3411.64 | 2238.37 | 0.61 | 3.06E-04 |
| ENSMUSG00000028809 | GSMG0028503 | Srrm1 | serine/arginine repetitive matrix 1 | 3002.19 | 3653.96 | -0.28 | 1.51E-01 |
| ENSMUSG00000059713 | GSMG0028507 | Rcan3 | regulator of calcineurin 3 | 221.36 | 313.50 | -0.51 | 4.58E-02 |
| ENSMUSG00000028803 | GSMG0028508 | Nipal3 | NIPA-like domain containing 3 | 286.66 | 266.31 | 0.10 | 7.90E-01 |
| ENSMUSG00000028675 | GSMG0028511 | Pnrc2 | proline-rich nuclear receptor coactivator 2 | 3006.84 | 3385.63 | -0.17 | 4.95E-01 |
| ENSMUSG00000028670 | GSMG0028512 | Lypla2 | lysophospholipase 2 | 1226.08 | 1392.71 | -0.18 | 3.43E-01 |
| ENSMUSG00000028669 | GSMG0028513 | Pithd1 | PITH (C-terminal proteasome-interacting domain of thioredoxin-like) d | 286.93 | 272.75 | 0.07 | 8.59E-01 |
| ENSMUSG00000028668 | GSMG0028514 | Tceb3 | transcription elongation factor B (SIII), polypeptide 3 | 1030.72 | 968.73 | 0.09 | 7.72E-01 |
| ENSMUSG00000036940 | GSMG0028519 | Kdm1a | lysine (K)-specific demethylase 1A | 2077.91 | 2447.25 | -0.24 | 2.55E-01 |
| ENSMUSG00000028664 | GSMG0028521 | Ephb2 | Eph receptor B2 | 259.51 | 491.46 | -0.92 | 6.64E-04 |
| ENSMUSG00000028661 | GSMG0028522 | Epha8 | Eph receptor A8 | 598.03 | 984.90 | -0.72 | 8.68E-03 |
| NULL | GSMG0028524 | NULL | NULL | 81.91 | 109.39 | -0.41 | 4.90E-01 |
| ENSMUSG00000036905 | GSMG0028526 | C1qb | complement component 1, q subcomponent, beta polypeptide | 1000.60 | 308.46 | 1.70 | 7.26E-17 |
| ENSMUSG00000036896 | GSMG0028527 | C1qc | complement component 1, q subcomponent, C chain | 1109.75 | 409.60 | 1.44 | 2.04E-17 |
| ENSMUSG00000036887 | GSMG0028528 | C1qa | complement component 1, q subcomponent, alpha polypeptide | 755.77 | 272.34 | 1.48 | 3.74E-14 |
| ENSMUSG00000006699 | GSMG0028532 | Cdc42 | cell division cycle 42 | 4762.90 | 5565.84 | -0.22 | 2.57E-01 |
| NULL | GSMG0028535 | NULL | NULL | 104.15 | 89.03 | 0.22 | 6.72E-01 |
| ENSMUSG00000028766 | GSMG0028539 | Alpl | alkaline phosphatase, liver/bone/kidney | 1158.14 | 369.53 | 1.65 | 9.39E-23 |
| ENSMUSG00000028756 | GSMG0028544 | Pink1 | PTEN induced putative kinase 1 | 2869.14 | 2261.24 | 0.34 | 5.04E-02 |
| ENSMUSG00000078235 | GSMG0028546 | Fam43b | family with sequence similarity 43, member B | 238.72 | 275.49 | -0.21 | 5.68E-01 |
| ENSMUSG00000041120 | GSMG0028555 | Nbl1 | neuroblastoma, suppression of tumorigenicity 1 | 727.26 | 534.61 | 0.44 | 3.76E-02 |
| ENSMUSG00000050608 | GSMG0028556 | Minos1 | mitochondrial inner membrane organizing system 1 | 3361.15 | 4034.38 | -0.26 | 2.26E-01 |
| NULL | GSMG0028558 | NULL | NULL | 84.32 | 79.88 | 0.07 | 9.12E-01 |
| ENSMUSG00000028744 | GSMG0028559 | Pqlc2 | PQ loop repeat containing 2 | 533.46 | 494.14 | 0.11 | 7.22E-01 |
| ENSMUSG00000028741 | GSMG0028560 | Mrto4 | mRNA turnover 4, ribosome maturation factor | 315.71 | 456.71 | -0.53 | 6.72E-03 |

| | | | | | | | |
|---|---|---|---|---|---|---|---|
| ENSMUSG00000040972 | GSMG0028568 | Igsf21 | immunoglobulin superfamily, member 21 | 884.31 | 1196.72 | -0.44 | 2.27E-02 |
| ENSMUSG00000040964 | GSMG0028572 | Arhgef10l | Rho guanine nucleotide exchange factor (GEF) 10-like | 652.59 | 693.23 | -0.09 | 8.32E-01 |
| ENSMUSG00000040860 | GSMG0028579 | Crocc | ciliary rootlet coiled-coil, rootletin | 983.96 | 392.98 | 1.33 | 9.16E-15 |
| ENSMUSG00000028923 | GSMG0028580 | Necap2 | NECAP endocytosis associated 2 | 1473.02 | 1458.17 | 0.01 | 9.62E-01 |
| ENSMUSG00000040842 | GSMG0028582 | Szrd1 | SUZ RNA binding domain containing 1 | 773.95 | 875.49 | -0.18 | 6.19E-01 |
| ENSMUSG00000070637 | GSMG0028587 | Gm694 | predicted gene 694 | 371.56 | 54.24 | 2.78 | 5.22E-34 |
| ENSMUSG00000006219 | GSMG0028589 | Fblim1 | filamin binding LIM protein 1 | 160.21 | 199.07 | -0.32 | 3.08E-01 |
| ENSMUSG00000028917 | GSMG0028592 | Plekhm2 | pleckstrin homology domain containing, family M (with RUN domain) | 1376.42 | 956.90 | 0.52 | 6.01E-02 |
| NULL | GSMG0028594 | NULL | NULL | 413.31 | 537.83 | -0.38 | 2.44E-01 |
| ENSMUSG00000078515 | GSMG0028595 | Ddi2 | DNA-damage inducible protein 2 | 540.15 | 817.87 | -0.60 | 2.78E-03 |
| ENSMUSG00000040697 | GSMG0028597 | Dnajc16 | DnaJ heat shock protein family (Hsp40) member C16 | 454.37 | 562.08 | -0.31 | 2.21E-01 |
| ENSMUSG00000040659 | GSMG0028600 | Efhd2 | EF hand domain containing 2 | 4365.00 | 4503.80 | -0.05 | 8.59E-01 |
| ENSMUSG00000040616 | GSMG0028603 | Tmem51 | transmembrane protein 51 | 2666.22 | 2360.05 | 0.18 | 4.12E-01 |
| ENSMUSG00000040606 | GSMG0028605 | Kazn | kazrin, periplakin interacting protein | 236.25 | 273.41 | -0.22 | 5.43E-01 |
| NULL | GSMG0028607 | NULL | NULL | 425.96 | 337.13 | 0.34 | 1.51E-01 |
| NULL | GSMG0028609 | NULL | NULL | 195.96 | 141.63 | 0.47 | 8.16E-02 |
| ENSMUSG00000057637 | GSMG0028616 | Prdm2 | PR domain containing 2, with ZNF domain | 854.21 | 1019.85 | -0.26 | 3.17E-01 |
| NULL | GSMG0028630 | NULL | NULL | 49.22 | 90.13 | -0.88 | 2.49E-02 |
| ENSMUSG00000028599 | GSMG0028634 | Tnfrsf1b | tumor necrosis factor receptor superfamily, member 1b | 198.33 | 195.04 | 0.03 | 9.49E-01 |
| ENSMUSG00000029022 | GSMG0028645 | Miip | migration and invasion inhibitory protein | 718.07 | 499.90 | 0.52 | 2.10E-02 |
| ENSMUSG00000029020 | GSMG0028646 | Mfn2 | mitofusin 2 | 1809.46 | 1963.13 | -0.12 | 5.98E-01 |
| ENSMUSG00000019055 | GSMG0028647 | Plod1 | procollagen-lysine, 2-oxoglutarate 5-dioxygenase 1 | 6855.75 | 7441.85 | -0.12 | 5.84E-01 |
| ENSMUSG00000029016 | GSMG0028649 | Clcn6 | chloride channel, voltage-sensitive 6 | 685.71 | 1011.35 | -0.56 | 2.18E-02 |
| ENSMUSG00000055401 | GSMG0028652 | Fbxo6 | F-box protein 6 | 986.60 | 728.91 | 0.44 | 1.53E-02 |
| ENSMUSG00000029001 | GSMG0028653 | Fbxo44 | F-box protein 44 | 216.21 | 165.17 | 0.39 | 1.77E-01 |
| ENSMUSG00000041544 | GSMG0028654 | Ptchd2 | patched domain containing 2 | 389.35 | 230.39 | 0.76 | 4.89E-03 |
| ENSMUSG00000047719 | GSMG0028656 | Ubiad1 | UbiA prenyltransferase domain containing 1 | 1075.13 | 1100.45 | -0.03 | 9.26E-01 |
| ENSMUSG00000041459 | GSMG0028659 | Tardbp | TAR DNA binding protein | 1183.14 | 1344.74 | -0.18 | 4.08E-01 |
| ENSMUSG00000028971 | GSMG0028661 | Cort | cortistatin | 76.99 | 85.17 | -0.15 | 7.87E-01 |
| ENSMUSG00000028975 | GSMG0028662 | Pex14 | peroxisomal biogenesis factor 14 | 178.23 | 216.61 | -0.29 | 4.13E-01 |
| ENSMUSG00000073705 | GSMG0028664 | Apitd1 | apoptosis-inducing, TAF9-like domain 1 | 104.72 | 252.14 | -1.27 | 6.19E-07 |
| ENSMUSG00000028961 | GSMG0028665 | Pgd | phosphogluconate dehydrogenase | 1290.36 | 1627.81 | -0.34 | 8.52E-02 |
| ENSMUSG00000063077 | GSMG0028666 | Kif1b | kinesin family member 1B | 3884.34 | 4476.13 | -0.20 | 4.04E-01 |

| | | | | | | | |
|---|---|---|---|---|---|---|---|
| NULL | GSMG0028667 | NULL | NULL | 263.15 | 241.07 | 0.13 | 7.02E-01 |
| NULL | GSMG0028668 | NULL | NULL | 727.41 | 722.45 | 0.01 | 9.72E-01 |
| ENSMUSG00000028960 | GSMG0028669 | Ube4b | ubiquitination factor E4B | 1428.45 | 1760.01 | -0.30 | 1.20E-01 |
| ENSMUSG00000028992 | GSMG0028673 | Nmnat1 | nicotinamide nucleotide adenylyltransferase 1 | 384.76 | 392.11 | -0.03 | 9.43E-01 |
| NULL | GSMG0028674 | NULL | NULL | 548.68 | 622.63 | -0.19 | 6.40E-01 |
| NULL | GSMG0028675 | NULL | NULL | 218.95 | 150.06 | 0.54 | 9.92E-02 |
| ENSMUSG00000039936 | GSMG0028676 | Pik3cd | phosphatidylinositol 3-kinase catalytic delta polypeptide | 488.15 | 666.85 | -0.45 | 7.26E-02 |
| ENSMUSG00000044700 | GSMG0028677 | Tmem201 | transmembrane protein 201 | 447.36 | 595.47 | -0.41 | 1.20E-01 |
| ENSMUSG00000028982 | GSMG0028678 | Slc25a33 | solute carrier family 25, member 33 | 393.12 | 237.30 | 0.73 | 3.51E-04 |
| ENSMUSG00000039911 | GSMG0028681 | Spsb1 | splA/ryanodine receptor domain and SOCS box containing 1 | 317.85 | 675.07 | -1.09 | 4.51E-10 |
| ENSMUSG00000028980 | GSMG0028683 | H6pd | hexose-6-phosphate dehydrogenase (glucose 1-dehydrogenase) | 111.56 | 148.32 | -0.41 | 2.10E-01 |
| ENSMUSG00000039838 | GSMG0028687 | Slc45a1 | solute carrier family 45, member 1 | 400.25 | 416.12 | -0.06 | 9.00E-01 |
| ENSMUSG00000028964 | GSMG0028689 | Park7 | Parkinson disease (autosomal recessive, early onset) 7 | 2419.47 | 3099.98 | -0.36 | 4.52E-02 |
| ENSMUSG00000028957 | GSMG0028691 | Per3 | period circadian clock 3 | 1133.32 | 474.92 | 1.25 | 8.78E-15 |
| ENSMUSG00000028955 | GSMG0028692 | Vamp3 | vesicle-associated membrane protein 3 | 3570.03 | 2960.86 | 0.27 | 1.24E-01 |
| NULL | GSMG0028697 | NULL | NULL | 177.83 | 102.20 | 0.80 | 2.90E-03 |
| ENSMUSG00000039759 | GSMG0028700 | Thap3 | THAP domain containing, apoptosis associated protein 3 | 379.84 | 395.65 | -0.06 | 8.87E-01 |
| ENSMUSG00000047777 | GSMG0028701 | Phf13 | PHD finger protein 13 | 141.63 | 162.92 | -0.21 | 6.05E-01 |
| ENSMUSG00000028952 | GSMG0028707 | Zbtb48 | zinc finger and BTB domain containing 48 | 653.22 | 624.03 | 0.07 | 8.30E-01 |
| ENSMUSG00000058498 | GSMG0028712 | Rnf207 | ring finger protein 207 | 141.26 | 40.02 | 1.81 | 2.23E-05 |
| ENSMUSG00000028931 | GSMG0028715 | Kcnab2 | potassium voltage-gated channel, shaker-related subfamily, beta membe | 250.03 | 110.30 | 1.18 | 3.28E-04 |
| ENSMUSG00000039546 | GSMG0028719 | Ajap1 | adherens junction associated protein 1 | 503.26 | 226.15 | 1.15 | 3.75E-04 |
| ENSMUSG00000029027 | GSMG0028722 | Dffb | DNA fragmentation factor, beta subunit | 196.61 | 189.71 | 0.05 | 8.99E-01 |
| ENSMUSG00000029030 | GSMG0028727 | Tprgl | transformation related protein 63 regulated like | 8091.17 | 6569.92 | 0.30 | 5.73E-02 |
| ENSMUSG00000039410 | GSMG0028730 | Prdm16 | PR domain containing 16 | 77.63 | 98.26 | -0.34 | 4.11E-01 |
| ENSMUSG00000029059 | GSMG0028736 | Fam213b | family with sequence similarity 213, member B | 357.79 | 315.91 | 0.18 | 5.43E-01 |
| ENSMUSG00000029055 | GSMG0028740 | Plch2 | phospholipase C, eta 2 | 622.26 | 621.77 | 0.00 | 9.99E-01 |
| ENSMUSG00000029048 | GSMG0028741 | Rer1 | retention in endoplasmic reticulum sorting receptor 1 | 1563.97 | 1731.77 | -0.15 | 5.59E-01 |
| ENSMUSG00000029050 | GSMG0028742 | Ski | ski sarcoma viral oncogene homolog (avian) | 604.78 | 314.31 | 0.94 | 2.97E-01 |
| NULL | GSMG0028743 | NULL | NULL | 1550.37 | 1070.80 | 0.53 | 5.86E-03 |
| ENSMUSG00000029053 | GSMG0028744 | Prkcz | protein kinase C, zeta | 432.67 | 484.65 | -0.16 | 5.41E-01 |
| ENSMUSG00000029054 | GSMG0028745 | Gabrd | gamma-aminobutyric acid (GABA) A receptor, subunit delta | 455.76 | 116.68 | 1.97 | 1.58E-21 |
| NULL | GSMG0028747 | NULL | NULL | 252.99 | 422.41 | -0.74 | 1.38E-02 |

| | | | | | | | |
|---|---|---|---|---|---|---|---|
| ENSMUSG00000029061 | GSMG0028751 | Mmp23 | matrix metallopeptidase 23 | 47.83 | 70.76 | -0.55 | 3.11E-01 |
| ENSMUSG00000029060 | GSMG0028752 | Mib2 | mindbomb E3 ubiquitin protein ligase 2 | 3244.63 | 2512.99 | 0.37 | 1.13E-01 |
| ENSMUSG00000029036 | GSMG0028754 | Atad3a | ATPase family, AAA domain containing 3A | 1267.80 | 1440.72 | -0.18 | 3.88E-01 |
| ENSMUSG00000042116 | GSMG0028755 | Vwa1 | von Willebrand factor A domain containing 1 | 14232.16 | 13132.35 | 0.12 | 5.77E-01 |
| NULL | GSMG0028757 | NULL | NULL | 145.09 | 154.33 | -0.10 | 8.39E-01 |
| ENSMUSG00000029073 | GSMG0028760 | Cptp | ceramide-1-phosphate transfer protein | 982.92 | 1151.93 | -0.23 | 3.06E-01 |
| ENSMUSG00000051557 | GSMG0028762 | Pusl1 | pseudouridylate synthase-like 1 | 1113.31 | 877.77 | 0.34 | 8.48E-02 |
| ENSMUSG00000050796 | GSMG0028764 | B3galt6 | UDP-Gal:betaGal beta 1,3-galactosyltransferase, polypeptide 6 | 562.72 | 741.17 | -0.40 | 5.40E-02 |
| ENSMUSG00000041936 | GSMG0028768 | Agrn | agrin | 3624.26 | 4194.45 | -0.21 | 3.99E-01 |
| ENSMUSG00000035692 | GSMG0028769 | Isg15 | ISG15 ubiquitin-like modifier | 136.05 | 101.23 | 0.42 | 2.68E-01 |
| NULL | GSMG0028771 | Dhrsx | dehydrogenase/reductase (SDR family) X chromosome | 311.18 | 371.51 | -0.26 | 3.34E-01 |
| ENSMUSG00000040274 | GSMG0028779 | Cdk6 | cyclin-dependent kinase 6 | 183.34 | 114.88 | 0.67 | 1.23E-02 |
| ENSMUSG00000058503 | GSMG0028782 | Fam133b | family with sequence similarity 133, member B | 457.83 | 514.11 | -0.17 | 6.90E-01 |
| ENSMUSG00000005907 | GSMG0028785 | Pex1 | peroxisomal biogenesis factor 1 | 713.49 | 768.35 | -0.11 | 6.81E-01 |
| ENSMUSG00000000600 | GSMG0028787 | Krit1 | KRIT1, ankyrin repeat containing | 1463.09 | 1553.13 | -0.09 | 7.52E-01 |
| ENSMUSG00000040407 | GSMG0028789 | Akap9 | A kinase (PRKA) anchor protein (yotiao) 9 | 656.98 | 866.22 | -0.40 | 4.13E-02 |
| ENSMUSG00000003161 | GSMG0028799 | Sri | sorcin | 2962.15 | 2939.26 | 0.01 | 9.72E-01 |
| ENSMUSG00000054099 | GSMG0028800 | Slc25a40 | solute carrier family 25, member 40 | 200.68 | 188.26 | 0.09 | 8.53E-01 |
| ENSMUSG00000040584 | GSMG0028802 | Abcb1a | ATP-binding cassette, sub-family B (MDR/TAP), member 1A | 3746.35 | 1612.23 | 1.22 | 7.19E-15 |
| ENSMUSG00000079659 | GSMG0028806 | Tmem243 | transmembrane protein 243, mitochondrial | 594.52 | 423.75 | 0.49 | 5.86E-02 |
| ENSMUSG00000056004 | GSMG0028808 | 9330182L06Rik | RIKEN cDNA 9330182L06 gene | 210.82 | 336.78 | -0.67 | 3.02E-03 |
| ENSMUSG00000028883 | GSMG0028820 | Sema3a | sema domain, immunoglobulin domain (Ig), short basic domain, secrete | 92.83 | 185.86 | -1.00 | 4.36E-03 |
| NULL | GSMG0028829 | NULL | NULL | 186.11 | 122.37 | 0.60 | 1.28E-01 |
| ENSMUSG00000061601 | GSMG0028831 | Pclo | piccolo (presynaptic cytomatrix protein) | 1507.97 | 1643.86 | -0.12 | 6.50E-01 |
| ENSMUSG00000040118 | GSMG0028843 | Cacna2d1 | calcium channel, voltage-dependent, alpha2/delta subunit 1 | 563.08 | 766.70 | -0.44 | 1.49E-02 |
| NULL | GSMG0028845 | NULL | NULL | 120.86 | 93.01 | 0.38 | 4.27E-01 |
| ENSMUSG00000028780 | GSMG0028856 | Sema3c | sema domain, immunoglobulin domain (Ig), short basic domain, secrete | 4287.35 | 547.18 | 2.97 | 2.03E-75 |
| NULL | GSMG0028873 | NULL | NULL | 147.60 | 286.60 | -0.97 | 2.25E-04 |
| NULL | GSMG0028874 | NULL | NULL | 258.52 | 206.96 | 0.32 | 3.28E-01 |
| NULL | GSMG0028876 | NULL | NULL | 280.05 | 230.78 | 0.28 | 3.46E-01 |
| NULL | GSMG0028877 | NULL | NULL | 216.51 | 227.45 | -0.07 | 8.72E-01 |
| ENSMUSG00000045435 | GSMG0028883 | Tmem60 | transmembrane protein 60 | 570.88 | 575.69 | -0.01 | 9.69E-01 |
| ENSMUSG00000101013 | GSMG0028884 | A630072M18Rik | RIKEN cDNA A630072M18 gene | 551.00 | 499.75 | 0.14 | 6.81E-01 |

| | | | | | | | |
|---|---|---|---|---|---|---|---|
| ENSMUSG00000039934 | GSMG0028885 | Gsap | gamma-secretase activating protein | 213.29 | 354.16 | -0.73 | 1.57E-03 |
| ENSMUSG00000047221 | GSMG0028887 | Fam185a | family with sequence similarity 185, member A | 50.67 | 66.61 | -0.39 | 4.53E-01 |
| ENSMUSG00000038525 | GSMG0028889 | Armc10 | armadillo repeat containing 10 | 291.66 | 385.56 | -0.40 | 1.07E-01 |
| ENSMUSG00000029017 | GSMG0028890 | Pmpcb | peptidase (mitochondrial processing) beta | 974.49 | 1164.51 | -0.26 | 1.96E-01 |
| ENSMUSG00000028932 | GSMG0028891 | Psmc2 | proteasome (prosome, macropain) 26S subunit, ATPase 2 | 2507.63 | 2915.81 | -0.22 | 3.18E-01 |
| ENSMUSG00000106379 | GSMG0028897 | Lhfpl3 | lipoma HMGIC fusion partner-like 3 | 252.51 | 442.37 | -0.81 | 5.86E-05 |
| ENSMUSG00000029004 | GSMG0028904 | Kmt2e | lysine (K)-specific methyltransferase 2E | 5025.32 | 4132.19 | 0.28 | 9.26E-02 |
| NULL | GSMG0028907 | NULL | NULL | 115.15 | 76.49 | 0.58 | 1.70E-01 |
| ENSMUSG00000028999 | GSMG0028908 | Rint1 | RAD50 interactor 1 | 835.66 | 718.32 | 0.22 | 4.07E-01 |
| ENSMUSG00000097180 | GSMG0028910 | 2700038G22Rik | RIKEN cDNA 2700038G22 gene | 188.52 | 271.10 | -0.52 | 2.54E-02 |
| ENSMUSG00000028986 | GSMG0028911 | Klhl7 | kelch-like 7 | 707.15 | 972.87 | -0.46 | 3.41E-02 |
| NULL | GSMG0028912 | NULL | NULL | 166.50 | 222.43 | -0.42 | 1.72E-01 |
| ENSMUSG00000048439 | GSMG0028916 | Nupl2 | nucleoporin like 2 | 157.56 | 162.42 | -0.05 | 9.16E-01 |
| ENSMUSG00000028978 | GSMG0028917 | Nos3 | nitric oxide synthase 3, endothelial cell | 14213.40 | 12349.97 | 0.20 | 3.02E-01 |
| ENSMUSG00000028973 | GSMG0028918 | Abcb8 | ATP-binding cassette, sub-family B (MDR/TAP), member 8 | 1274.25 | 2402.64 | -0.91 | 1.17E-06 |
| ENSMUSG00000028962 | GSMG0028920 | Slc4a2 | solute carrier family 4 (anion exchanger), member 2 | 1274.65 | 1202.18 | 0.08 | 7.84E-01 |
| ENSMUSG00000023353 | GSMG0028922 | Agap3 | ArfGAP with GTPase domain, ankyrin repeat and PH domain 3 | 2070.63 | 2031.39 | 0.03 | 9.52E-01 |
| ENSMUSG00000038181 | GSMG0028924 | Chpf2 | chondroitin polymerizing factor 2 | 1665.06 | 1103.09 | 0.59 | 1.04E-02 |
| ENSMUSG00000028954 | GSMG0028926 | Nub1 | negative regulator of ubiquitin-like proteins 1 | 1578.65 | 1836.31 | -0.22 | 2.20E-01 |
| ENSMUSG00000086446 | GSMG0028928 | Prkag2os1 | protein kinase, AMP-activated, gamma 2 non-catalytic subunit, opposit | 104.93 | 96.90 | 0.11 | 8.27E-01 |
| NULL | GSMG0028929 | NULL | NULL | 54.03 | 164.57 | -1.61 | 1.38E-08 |
| ENSMUSG00000038072 | GSMG0028932 | Galnt11 | UDP-N-acetyl-alpha-D-galactosamine:polypeptide N-acetylgalactosami | 1857.15 | 1604.34 | 0.21 | 3.00E-01 |
| ENSMUSG00000056367 | GSMG0028938 | Actr3b | ARP3 actin-related protein 3B | 574.49 | 705.74 | -0.30 | 2.53E-01 |
| ENSMUSG00000061576 | GSMG0028944 | Dpp6 | dipeptidylpeptidase 6 | 641.21 | 1067.56 | -0.74 | 3.34E-04 |
| ENSMUSG00000045294 | GSMG0028956 | Insig1 | insulin induced gene 1 | 1113.92 | 1482.34 | -0.41 | 5.32E-02 |
| ENSMUSG00000029130 | GSMG0028966 | Rnf32 | ring finger protein 32 | 167.19 | 134.19 | 0.32 | 4.07E-01 |
| ENSMUSG00000001569 | GSMG0028971 | Nom1 | nucleolar protein with MIF4G domain 1 | 790.49 | 1053.68 | -0.41 | 1.71E-02 |
| ENSMUSG00000039000 | GSMG0028973 | Ube3c | ubiquitin protein ligase E3C | 2797.42 | 3062.04 | -0.13 | 5.33E-01 |
| ENSMUSG00000075703 | GSMG0028986 | Ept1 | ethanolaminephosphotransferase 1 (CDP-ethanolamine-specific) | 112.20 | 135.13 | -0.26 | 4.83E-01 |
| ENSMUSG00000073102 | GSMG0028987 | Drc1 | dynein regulatory complex subunit 1 | 407.05 | 364.91 | 0.16 | 6.04E-01 |
| ENSMUSG00000049265 | GSMG0028991 | Kcnk3 | potassium channel, subfamily K, member 3 | 443.13 | 259.33 | 0.77 | 1.87E-03 |
| ENSMUSG00000029175 | GSMG0028993 | Slc35f6 | solute carrier family 35, member F6 | 368.46 | 277.67 | 0.41 | 1.44E-01 |
| ENSMUSG00000029177 | GSMG0028994 | Cenpa | centromere protein A | 589.18 | 1036.22 | -0.81 | 9.09E-06 |

| | | | | | | | |
|---|---|---|---|---|---|---|---|
| ENSMUSG00000029168 | GSMG0028995 | Dpysl5 | dihydropyrimidinase-like 5 | 86.90 | 643.31 | -2.90 | 4.36E-45 |
| ENSMUSG00000029166 | GSMG0028996 | Mapre3 | microtubule-associated protein, RP/EB family, member 3 | 711.12 | 770.07 | -0.12 | 7.34E-01 |
| NULL | GSMG0028997 | NULL | NULL | 797.70 | 585.80 | 0.44 | 1.14E-01 |
| ENSMUSG00000038828 | GSMG0028998 | Tmem214 | transmembrane protein 214 | 2307.90 | 2156.12 | 0.10 | 7.30E-01 |
| ENSMUSG00000029165 | GSMG0028999 | Agbl5 | ATP/GTP binding protein-like 5 | 558.35 | 776.21 | -0.48 | 6.70E-02 |
| ENSMUSG00000029163 | GSMG0029001 | Emilin1 | elastin microfibril interfacer 1 | 775.45 | 942.15 | -0.28 | 3.11E-01 |
| ENSMUSG00000029162 | GSMG0029002 | Khk | ketohexokinase | 236.51 | 184.96 | 0.35 | 2.15E-01 |
| ENSMUSG00000006638 | GSMG0029003 | Abhd1 | abhydrolase domain containing 1 | 386.09 | 310.33 | 0.31 | 2.80E-01 |
| ENSMUSG00000013622 | GSMG0029005 | Atraid | all-trans retinoic acid induced differentiation factor | 3417.84 | 3538.83 | -0.05 | 8.51E-01 |
| ENSMUSG00000013629 | GSMG0029006 | Cad | carbamoyl-phosphate synthetase 2, aspartate transcarbamylase, and dihy | 1128.30 | 1304.08 | -0.21 | 5.23E-01 |
| ENSMUSG00000029146 | GSMG0029010 | Snx17 | sorting nexin 17 | 2530.64 | 2678.03 | -0.08 | 7.34E-01 |
| ENSMUSG00000029148 | GSMG0029011 | Nrbp1 | nuclear receptor binding protein 1 | 4019.18 | 4078.66 | -0.02 | 9.42E-01 |
| ENSMUSG00000062761 | GSMG0029014 | Zfp512 | zinc finger protein 512 | 848.01 | 1146.78 | -0.43 | 2.95E-02 |
| ENSMUSG00000064037 | GSMG0029017 | Gpn1 | GPN-loop GTPase 1 | 618.93 | 826.22 | -0.42 | 2.69E-02 |
| ENSMUSG00000029141 | GSMG0029018 | Slc4a1ap | solute carrier family 4 (anion exchanger), member 1, adaptor protein | 386.12 | 378.09 | 0.03 | 9.33E-01 |
| ENSMUSG00000106918 | GSMG0029021 | Mrpl33 | mitochondrial ribosomal protein L33 | 1132.81 | 1378.32 | -0.28 | 2.59E-01 |
| ENSMUSG00000052139 | GSMG0029023 | Bre | brain and reproductive organ-expressed protein | 498.36 | 552.71 | -0.15 | 5.63E-01 |
| ENSMUSG00000029135 | GSMG0029026 | Fosl2 | fos-like antigen 2 | 636.17 | 441.37 | 0.53 | 4.19E-02 |
| ENSMUSG00000014956 | GSMG0029030 | Ppp1cb | protein phosphatase 1, catalytic subunit, beta isoform | 2480.58 | 2077.31 | 0.26 | 2.18E-01 |
| NULL | GSMG0029031 | NULL | NULL | 217.80 | 160.49 | 0.44 | 1.58E-01 |
| ENSMUSG00000014932 | GSMG0029034 | Yes1 | YES proto-oncogene 1, Src family tyrosine kinase | 1487.20 | 1604.32 | -0.11 | 6.74E-01 |
| ENSMUSG00000037426 | GSMG0029036 | Depdc5 | DEP domain containing 5 | 526.94 | 593.21 | -0.17 | 5.28E-01 |
| ENSMUSG00000079562 | GSMG0029041 | Maea | macrophage erythroblast attacher | 859.47 | 832.50 | 0.05 | 8.76E-01 |
| ENSMUSG00000037355 | GSMG0029042 | Uvssa | UV stimulated scaffold protein A | 547.66 | 538.96 | 0.02 | 9.53E-01 |
| ENSMUSG00000037313 | GSMG0029045 | Tacc3 | transforming, acidic coiled-coil containing protein 3 | 704.05 | 1362.61 | -0.95 | 6.02E-08 |
| ENSMUSG00000054252 | GSMG0029046 | Fgfr3 | fibroblast growth factor receptor 3 | 1390.98 | 918.40 | 0.60 | 1.87E-02 |
| ENSMUSG00000057406 | GSMG0029048 | Whsc1 | Wolf-Hirschhorn syndrome candidate 1 (human) | 783.29 | 1080.95 | -0.46 | 9.31E-03 |
| ENSMUSG00000070858 | GSMG0029053 | Gm1673 | predicted gene 1673 | 472.39 | 995.88 | -1.08 | 2.73E-05 |
| ENSMUSG00000048142 | GSMG0029054 | Nat8l | N-acetyltransferase 8-like | 864.25 | 398.95 | 1.11 | 7.10E-05 |
| ENSMUSG00000029110 | GSMG0029058 | Rnf4 | ring finger protein 4 | 867.15 | 1125.00 | -0.37 | 3.63E-02 |
| ENSMUSG00000037210 | GSMG0029060 | Fam193a | family with sequence similarity 193, member A | 872.83 | 702.38 | 0.31 | 1.80E-01 |
| ENSMUSG00000054520 | GSMG0029063 | Sh3bp2 | SH3-domain binding protein 2 | 84.47 | 347.88 | -2.04 | 6.71E-21 |
| ENSMUSG00000029106 | GSMG0029065 | Add1 | adducin 1 (alpha) | 4279.32 | 4251.18 | 0.01 | 9.74E-01 |

| | | | | | | | |
|---|---|---|---|---|---|---|---|
| NULL | GSMG0029066 | NULL | NULL | 281.55 | 262.14 | 0.10 | 8.11E-01 |
| NULL | GSMG0029068 | NULL | NULL | 838.30 | 645.80 | 0.38 | 3.43E-02 |
| ENSMUSG00000052783 | GSMG0029070 | Grk4 | G protein-coupled receptor kinase 4 | 176.45 | 157.46 | 0.16 | 6.77E-01 |
| ENSMUSG00000029104 | GSMG0029071 | Htt | huntingtin | 291.64 | 250.08 | 0.22 | 4.90E-01 |
| ENSMUSG00000029101 | GSMG0029073 | Rgs12 | regulator of G-protein signaling 12 | 3383.56 | 3055.71 | 0.15 | 5.09E-01 |
| ENSMUSG00000029098 | GSMG0029080 | Acox3 | acyl-Coenzyme A oxidase 3, pristanoyl | 880.89 | 900.60 | -0.03 | 9.27E-01 |
| ENSMUSG00000029095 | GSMG0029081 | Ablim2 | actin-binding LIM protein 2 | 1144.14 | 999.12 | 0.20 | 4.62E-01 |
| ENSMUSG00000029094 | GSMG0029082 | Afap1 | actin filament associated protein 1 | 268.96 | 325.33 | -0.28 | 4.04E-01 |
| NULL | GSMG0029083 | NULL | NULL | 192.61 | 234.10 | -0.29 | 4.59E-01 |
| ENSMUSG00000029198 | GSMG0029085 | Grpel1 | GrpE-like 1, mitochondrial | 1813.12 | 2417.49 | -0.41 | 9.25E-03 |
| NULL | GSMG0029089 | NULL | NULL | 350.51 | 479.31 | -0.45 | 1.32E-01 |
| ENSMUSG00000029120 | GSMG0029090 | Ppp2r2c | protein phosphatase 2, regulatory subunit B, gamma | 386.38 | 385.85 | 0.00 | 9.96E-01 |
| ENSMUSG00000063646 | GSMG0029091 | Jakmip1 | janus kinase and microtubule interacting protein 1 | 649.26 | 857.25 | -0.40 | 4.99E-02 |
| ENSMUSG00000029121 | GSMG0029094 | Crmp1 | collapsin response mediator protein 1 | 874.77 | 4690.34 | -2.42 | 2.96E-48 |
| ENSMUSG00000062329 | GSMG0029097 | Cytl1 | cytokine-like 1 | 92.50 | 8.69 | 3.41 | 2.55E-12 |
| ENSMUSG00000029125 | GSMG0029100 | Stx18 | syntaxin 18 | 457.45 | 493.77 | -0.11 | 7.31E-01 |
| ENSMUSG00000067367 | GSMG0029102 | Lyar | Ly1 antibody reactive clone | 544.13 | 688.57 | -0.34 | 2.47E-01 |
| ENSMUSG00000067365 | GSMG0029103 | Tmem128 | transmembrane protein 128 | 1401.10 | 1392.95 | 0.01 | 9.80E-01 |
| ENSMUSG00000039782 | GSMG0029114 | Cpeb2 | cytoplasmic polyadenylation element binding protein 2 | 191.52 | 204.52 | -0.10 | 8.05E-01 |
| ENSMUSG00000039765 | GSMG0029120 | Cc2d2a | coiled-coil and C2 domain containing 2A | 448.50 | 438.07 | 0.03 | 9.17E-01 |
| ENSMUSG00000029084 | GSMG0029124 | Cd38 | CD38 antigen | 237.51 | 200.07 | 0.25 | 4.59E-01 |
| ENSMUSG00000039682 | GSMG0029136 | Lap3 | leucine aminopeptidase 3 | 495.70 | 464.51 | 0.10 | 7.77E-01 |
| ENSMUSG00000015804 | GSMG0029137 | Med28 | mediator complex subunit 28 | 1678.94 | 1615.62 | 0.05 | 8.68E-01 |
| ENSMUSG00000015880 | GSMG0029142 | Ncapg | non-SMC condensin I complex, subunit G | 105.16 | 202.01 | -0.95 | 1.95E-03 |
| NULL | GSMG0029151 | Pacrgl | PARK2 co-regulated-like | 184.49 | 213.67 | -0.21 | 5.13E-01 |
| ENSMUSG00000072941 | GSMG0029161 | Sod3 | superoxide dismutase 3, extracellular | 5285.57 | 2541.36 | 1.06 | 1.11E-13 |
| ENSMUSG00000029186 | GSMG0029163 | Pi4k2b | phosphatidylinositol 4-kinase type 2 beta | 148.66 | 213.54 | -0.53 | 4.93E-02 |
| ENSMUSG00000029179 | GSMG0029164 | Zcchc4 | zinc finger, CCHC domain containing 4 | 174.97 | 200.84 | -0.20 | 5.53E-01 |
| ENSMUSG00000029176 | GSMG0029165 | Anapc4 | anaphase promoting complex subunit 4 | 1852.38 | 2141.15 | -0.21 | 3.16E-01 |
| ENSMUSG00000061461 | GSMG0029167 | Smim20 | small integral membrane protein 20 | 659.07 | 685.17 | -0.06 | 8.51E-01 |
| ENSMUSG00000039178 | GSMG0029171 | Tbc1d19 | TBC1 domain family, member 19 | 395.63 | 483.43 | -0.29 | 2.25E-01 |
| ENSMUSG00000039156 | GSMG0029174 | Stim2 | stromal interaction molecule 2 | 572.24 | 794.67 | -0.48 | 1.67E-02 |
| ENSMUSG00000029108 | GSMG0029181 | Pcdh7 | protocadherin 7 | 2751.27 | 2682.43 | 0.04 | 9.29E-01 |

| | | | | | | | |
|---|---|---|---|---|---|---|---|
| NULL | GSMG0029189 | NULL | NULL | 297.62 | 272.11 | 0.13 | 7.64E-01 |
| NULL | GSMG0029190 | NULL | NULL | 244.99 | 246.83 | -0.01 | 9.75E-01 |
| ENSMUSG00000090061 | GSMG0029198 | Nwd2 | NACHT and WD repeat domain containing 2 | 114.32 | 265.49 | -1.21 | 1.16E-06 |
| ENSMUSG00000029171 | GSMG0029201 | Pgm1 | phosphoglucomutase 1 | 1757.67 | 2395.75 | -0.45 | 3.49E-02 |
| ENSMUSG00000029174 | GSMG0029202 | Tbc1d1 | TBC1 domain family, member 1 | 847.40 | 746.75 | 0.18 | 4.57E-01 |
| ENSMUSG00000029178 | GSMG0029206 | Klf3 | Kruppel-like factor 3 (basic) | 1355.05 | 1307.53 | 0.05 | 8.53E-01 |
| ENSMUSG00000029185 | GSMG0029208 | Fam114a1 | family with sequence similarity 114, member A1 | 631.84 | 518.79 | 0.29 | 1.78E-01 |
| ENSMUSG00000054920 | GSMG0029210 | Klhl5 | kelch-like 5 | 1739.17 | 1593.49 | 0.13 | 5.95E-01 |
| ENSMUSG00000037890 | GSMG0029211 | Wdr19 | WD repeat domain 19 | 602.81 | 493.40 | 0.29 | 1.91E-01 |
| ENSMUSG00000029199 | GSMG0029213 | Lias | lipoic acid synthetase | 827.41 | 1010.33 | -0.29 | 1.23E-01 |
| NULL | GSMG0029215 | NULL | NULL | 689.37 | 560.42 | 0.30 | 2.16E-01 |
| ENSMUSG00000029203 | GSMG0029217 | Ube2k | ubiquitin-conjugating enzyme E2K | 728.75 | 1005.23 | -0.46 | 1.12E-02 |
| NULL | GSMG0029218 | NULL | NULL | 473.62 | 543.37 | -0.20 | 4.67E-01 |
| ENSMUSG00000029223 | GSMG0029238 | Uchl1 | ubiquitin carboxy-terminal hydrolase L1 | 2330.59 | 3701.77 | -0.67 | 1.85E-05 |
| ENSMUSG00000037736 | GSMG0029241 | Limch1 | LIM and calponin homology domains 1 | 2356.37 | 3174.80 | -0.43 | 1.19E-02 |
| ENSMUSG00000037720 | GSMG0029245 | Tmem33 | transmembrane protein 33 | 682.55 | 674.07 | 0.02 | 9.61E-01 |
| ENSMUSG00000029221 | GSMG0029247 | Slc30a9 | solute carrier family 30 (zinc transporter), member 9 | 942.30 | 914.93 | 0.04 | 8.91E-01 |
| ENSMUSG00000029208 | GSMG0029256 | Guf1 | GUF1 homolog, GTPase | 2830.91 | 2515.39 | 0.17 | 3.82E-01 |
| ENSMUSG00000029212 | GSMG0029257 | Gabrb1 | gamma-aminobutyric acid (GABA) A receptor, subunit beta 1 | 649.80 | 719.48 | -0.14 | 5.77E-01 |
| NULL | GSMG0029259 | NULL | NULL | 207.96 | 155.80 | 0.41 | 2.45E-01 |
| ENSMUSG00000046808 | GSMG0029263 | Atp10d | ATPase, class V, type 10D | 179.18 | 148.89 | 0.27 | 5.10E-01 |
| ENSMUSG00000036087 | GSMG0029269 | Slain2 | SLAIN motif family, member 2 | 889.76 | 740.87 | 0.26 | 1.86E-01 |
| ENSMUSG00000029152 | GSMG0029274 | Ociad1 | OCIA domain containing 1 | 1642.43 | 1924.05 | -0.23 | 3.67E-01 |
| ENSMUSG00000051674 | GSMG0029277 | Dcun1d4 | DCN1, defective in cullin neddylation 1, domain containing 4 (S. cerev | 347.50 | 331.52 | 0.07 | 8.42E-01 |
| ENSMUSG00000053656 | GSMG0029285 | Dancr | differentiation antagonizing non-protein coding RNA | 255.15 | 273.54 | -0.10 | 7.91E-01 |
| ENSMUSG00000049907 | GSMG0029286 | Rasl11b | RAS-like, family 11, member B | 767.73 | 485.50 | 0.66 | 5.53E-04 |
| ENSMUSG00000029227 | GSMG0029287 | Fip1l1 | FIP1 like 1 (S. cerevisiae) | 915.60 | 1013.85 | -0.15 | 5.22E-01 |
| ENSMUSG00000029231 | GSMG0029295 | Pdgfra | platelet derived growth factor receptor, alpha polypeptide | 248.61 | 237.13 | 0.07 | 8.78E-01 |
| ENSMUSG00000005672 | GSMG0029296 | Kit | kit oncogene | 1333.94 | 2818.84 | -1.08 | 2.31E-14 |
| ENSMUSG00000029233 | GSMG0029299 | Srd5a3 | steroid 5 alpha-reductase 3 | 318.33 | 306.29 | 0.06 | 8.79E-01 |
| ENSMUSG00000029234 | GSMG0029300 | Tmem165 | transmembrane protein 165 | 2925.91 | 2813.36 | 0.06 | 8.30E-01 |
| ENSMUSG00000098449 | GSMG0029301 | Gm7467 | predicted gene 7467 | 95.94 | 70.95 | 0.43 | 3.51E-01 |
| ENSMUSG00000036435 | GSMG0029304 | Exoc1 | exocyst complex component 1 | 1402.59 | 1886.77 | -0.43 | 7.63E-03 |

| | | | | | | | |
|---|---|---|---|---|---|---|---|
| ENSMUSG00000036403 | GSMG0029305 | Cep135 | centrosomal protein 135 | 444.52 | 555.99 | -0.32 | 1.80E-01 |
| NULL | GSMG0029311 | NULL | NULL | 106.90 | 199.20 | -0.90 | 4.03E-04 |
| ENSMUSG00000036377 | GSMG0029312 | C530008M17Rik | RIKEN cDNA C530008M17 gene | 2010.89 | 3272.63 | -0.70 | 6.54E-07 |
| ENSMUSG00000029247 | GSMG0029314 | Paics | phosphoribosylaminoimidazole carboxylase, phosphoribosylaminoribos | 1605.40 | 1952.84 | -0.28 | 2.04E-01 |
| ENSMUSG00000036323 | GSMG0029315 | Srp72 | signal recognition particle 72 | 1806.00 | 2093.58 | -0.21 | 3.41E-01 |
| ENSMUSG00000029249 | GSMG0029318 | Rest | RE1-silencing transcription factor | 1247.67 | 1506.14 | -0.27 | 1.47E-01 |
| NULL | GSMG0029319 | NULL | NULL | 368.45 | 397.04 | -0.11 | 7.69E-01 |
| ENSMUSG00000029250 | GSMG0029320 | Polr2b | polymerase (RNA) II (DNA directed) polypeptide B | 2367.96 | 2640.58 | -0.16 | 4.35E-01 |
| ENSMUSG00000037605 | GSMG0029323 | Adgrl3 | adhesion G protein-coupled receptor L3 | 2054.34 | 2111.02 | -0.04 | 9.11E-01 |
| NULL | GSMG0029324 | NULL | NULL | 356.13 | 223.89 | 0.67 | 1.06E-02 |
| NULL | GSMG0029327 | NULL | NULL | 377.77 | 338.45 | 0.16 | 5.95E-01 |
| NULL | GSMG0029328 | NULL | NULL | 272.87 | 306.97 | -0.16 | 7.06E-01 |
| NULL | GSMG0029331 | NULL | NULL | 241.17 | 264.30 | -0.13 | 7.52E-01 |
| NULL | GSMG0029343 | NULL | NULL | 475.97 | 597.21 | -0.33 | 2.52E-01 |
| NULL | GSMG0029345 | NULL | NULL | 367.94 | 455.10 | -0.31 | 2.70E-01 |
| ENSMUSG00000035851 | GSMG0029357 | Ythdc1 | YTH domain containing 1 | 457.19 | 412.51 | 0.15 | 5.96E-01 |
| NULL | GSMG0029378 | NULL | NULL | 1238.27 | 1571.24 | -0.34 | 1.24E-01 |
| ENSMUSG00000029291 | GSMG0029379 | Rufy3 | RUN and FYVE domain containing 3 | 805.41 | 1344.67 | -0.74 | 2.93E-06 |
| ENSMUSG00000006262 | GSMG0029380 | Mob1b | MOB kinase activator 1B | 72.07 | 79.95 | -0.15 | 7.79E-01 |
| ENSMUSG00000029366 | GSMG0029381 | Dck | deoxycytidine kinase | 164.65 | 341.29 | -1.05 | 3.63E-06 |
| ENSMUSG00000060961 | GSMG0029384 | Slc4a4 | solute carrier family 4 (anion exchanger), member 4 | 1021.79 | 917.95 | 0.15 | 6.51E-01 |
| NULL | GSMG0029392 | NULL | NULL | 423.53 | 297.55 | 0.51 | 1.08E-02 |
| ENSMUSG00000029373 | GSMG0029399 | Pf4 | platelet factor 4 | 403.54 | 181.83 | 1.15 | 2.57E-07 |
| ENSMUSG00000034981 | GSMG0029415 | Parm1 | prostate androgen-regulated mucin-like protein 1 | 333.06 | 160.54 | 1.06 | 1.11E-05 |
| NULL | GSMG0029420 | NULL | NULL | 1101.58 | 1430.48 | -0.38 | 1.28E-01 |
| ENSMUSG00000029407 | GSMG0029423 | Uso1 | USO1 vesicle docking factor | 1657.07 | 1847.03 | -0.16 | 4.47E-01 |
| ENSMUSG00000034842 | GSMG0029424 | Art3 | ADP-ribosyltransferase 3 | 2434.32 | 1732.49 | 0.49 | 1.60E-02 |
| ENSMUSG00000058013 | GSMG0029433 | Sept11 | septin 11 | 5061.45 | 4933.91 | 0.04 | 8.85E-01 |
| ENSMUSG00000029385 | GSMG0029440 | Ccng2 | cyclin G2 | 1562.01 | 1296.75 | 0.27 | 2.36E-01 |
| ENSMUSG00000029486 | GSMG0029448 | Mrpl1 | mitochondrial ribosomal protein L1 | 278.61 | 296.71 | -0.09 | 8.08E-01 |
| ENSMUSG00000029484 | GSMG0029452 | Anxa3 | annexin A3 | 6091.88 | 4468.90 | 0.45 | 6.41E-03 |
| ENSMUSG00000034663 | GSMG0029457 | Bmp2k | BMP2 inducible kinase | 215.91 | 350.04 | -0.70 | 4.14E-03 |
| ENSMUSG00000029326 | GSMG0029475 | Enoph1 | enolase-phosphatase 1 | 339.34 | 441.89 | -0.38 | 7.21E-02 |

| | | | | | | | |
|---|---|---|---|---|---|---|---|
| ENSMUSG00000035297 | GSMG0029480 | Cops4 | COP9 signalosome subunit 4 | 1466.99 | 1969.28 | -0.42 | 2.03E-02 |
| ENSMUSG00000016833 | GSMG0029484 | Mrps18c | mitochondrial ribosomal protein S18C | 723.87 | 945.93 | -0.39 | 6.20E-02 |
| ENSMUSG00000029330 | GSMG0029488 | Cds1 | CDP-diacylglycerol synthase 1 | 230.26 | 164.66 | 0.48 | 1.21E-01 |
| ENSMUSG00000057315 | GSMG0029491 | Arhgap24 | Rho GTPase activating protein 24 | 70.79 | 59.00 | 0.26 | 6.07E-01 |
| ENSMUSG00000029313 | GSMG0029501 | Aff1 | AF4/FMR2 family, member 1 | 552.14 | 441.52 | 0.32 | 1.54E-01 |
| ENSMUSG00000029310 | GSMG0029504 | Nudt9 | nudix (nucleoside diphosphate linked moiety X)-type motif 9 | 1215.38 | 1273.05 | -0.07 | 8.03E-01 |
| ENSMUSG00000029304 | GSMG0029510 | Spp1 | secreted phosphoprotein 1 | 212.74 | 482.23 | -1.17 | 5.24E-05 |
| ENSMUSG00000034462 | GSMG0029511 | Pkd2 | polycystic kidney disease 2 | 3003.96 | 2249.21 | 0.42 | 1.14E-02 |
| ENSMUSG00000079065 | GSMG0029513 | BC005561 | cDNA sequence BC005561 | 6456.46 | 3750.64 | 0.78 | 2.72E-05 |
| ENSMUSG00000097392 | GSMG0029514 | D930016D06Rik | RIKEN cDNA D930016D06 gene | 896.70 | 460.32 | 0.96 | 6.56E-08 |
| ENSMUSG00000070639 | GSMG0029520 | Lrrc8b | leucine rich repeat containing 8 family, member B | 2390.90 | 2395.89 | 0.00 | 9.93E-01 |
| ENSMUSG00000054720 | GSMG0029524 | Lrrc8c | leucine rich repeat containing 8 family, member C | 2583.19 | 2381.30 | 0.12 | 6.21E-01 |
| ENSMUSG00000046079 | GSMG0029527 | Lrrc8d | leucine rich repeat containing 8D | 1332.14 | 2589.72 | -0.96 | 2.34E-10 |
| ENSMUSG00000029290 | GSMG0029530 | Zfp326 | zinc finger protein 326 | 329.68 | 446.38 | -0.44 | 3.96E-02 |
| ENSMUSG00000029283 | GSMG0029538 | Cdc7 | cell division cycle 7 (S. cerevisiae) | 437.54 | 613.20 | -0.49 | 2.15E-02 |
| ENSMUSG00000033805 | GSMG0029544 | Ephx4 | epoxide hydrolase 4 | 175.01 | 65.73 | 1.41 | 8.99E-07 |
| ENSMUSG00000033773 | GSMG0029547 | Rpap2 | RNA polymerase II associated protein 2 | 356.91 | 325.35 | 0.13 | 6.78E-01 |
| ENSMUSG00000029267 | GSMG0029556 | Mtf2 | metal response element binding transcription factor 2 | 1045.91 | 1063.83 | -0.02 | 9.37E-01 |
| ENSMUSG00000029265 | GSMG0029561 | Dr1 | down-regulator of transcription 1 | 1090.05 | 1193.86 | -0.13 | 6.54E-01 |
| NULL | GSMG0029564 | NULL | NULL | 867.77 | 311.97 | 1.47 | 2.92E-12 |
| ENSMUSG00000033623 | GSMG0029568 | Pcgf3 | polycomb group ring finger 3 | 398.17 | 476.12 | -0.26 | 3.05E-01 |
| ENSMUSG00000013495 | GSMG0029570 | Tmem175 | transmembrane protein 175 | 640.21 | 522.58 | 0.29 | 1.83E-01 |
| NULL | GSMG0029571 | NULL | NULL | 61.28 | 44.55 | 0.46 | 3.49E-01 |
| ENSMUSG00000033540 | GSMG0029572 | Idua | iduronidase, alpha-L- | 270.13 | 198.40 | 0.45 | 1.14E-01 |
| ENSMUSG00000008090 | GSMG0029573 | Fgfrl1 | fibroblast growth factor receptor-like 1 | 151.98 | 126.68 | 0.26 | 5.61E-01 |
| ENSMUSG00000090015 | GSMG0029579 | Gm15446 | predicted gene 15446 | 211.57 | 108.12 | 0.97 | 1.86E-04 |
| ENSMUSG00000064247 | GSMG0029581 | Plcxd1 | phosphatidylinositol-specific phospholipase C, X domain containing 1 | 457.79 | 388.58 | 0.24 | 3.89E-01 |
| ENSMUSG00000023284 | GSMG0029582 | Zfp605 | zinc finger protein 605 | 242.80 | 421.98 | -0.79 | 6.33E-05 |
| ENSMUSG00000014668 | GSMG0029584 | Chfr | checkpoint with forkhead and ring finger domains | 1814.17 | 1734.21 | 0.07 | 8.13E-01 |
| ENSMUSG00000029502 | GSMG0029586 | Golga3 | golgi autoantigen, golgin subfamily a, 3 | 190.23 | 183.16 | 0.05 | 9.18E-01 |
| ENSMUSG00000029501 | GSMG0029589 | Ankle2 | ankyrin repeat and LEM domain containing 2 | 1115.21 | 1295.74 | -0.22 | 2.68E-01 |
| ENSMUSG00000007080 | GSMG0029590 | Pole | polymerase (DNA directed), epsilon | 50.14 | 93.43 | -0.89 | 1.56E-02 |
| ENSMUSG00000033316 | GSMG0029595 | Galnt9 | UDP-N-acetyl-alpha-D-galactosamine:polypeptide N-acetylgalactosami | 278.68 | 194.73 | 0.52 | 9.08E-02 |

| | | | | | | | |
|---|---|---|---|---|---|---|---|
| ENSMUSG00000029504 | GSMG0029596 | Ddx51 | DEAD (Asp-Glu-Ala-Asp) box polypeptide 51 | 303.45 | 294.94 | 0.04 | 9.28E-01 |
| ENSMUSG00000029521 | GSMG0029600 | Chek2 | checkpoint kinase 2 | 186.50 | 362.93 | -0.96 | 1.65E-05 |
| ENSMUSG00000050017 | GSMG0029610 | Pitpnb | phosphatidylinositol transfer protein, beta | 2627.18 | 2882.36 | -0.13 | 5.52E-01 |
| ENSMUSG00000029344 | GSMG0029619 | Tpst2 | protein-tyrosine sulfotransferase 2 | 2370.67 | 2309.68 | 0.04 | 8.96E-01 |
| ENSMUSG00000029345 | GSMG0029621 | Tfip11 | tuftelin interacting protein 11 | 406.13 | 472.82 | -0.22 | 3.46E-01 |
| ENSMUSG00000042328 | GSMG0029622 | Hps4 | Hermansky-Pudlak syndrome 4 | 423.66 | 359.94 | 0.23 | 3.32E-01 |
| ENSMUSG00000053334 | GSMG0029639 | Ficd | FIC domain containing | 309.92 | 261.35 | 0.25 | 4.46E-01 |
| ENSMUSG00000025825 | GSMG0029640 | Iscu | iron-sulfur cluster assembly enzyme | 2121.99 | 1814.40 | 0.23 | 2.77E-01 |
| NULL | GSMG0029641 | NULL | NULL | 182.10 | 254.29 | -0.48 | 5.82E-02 |
| NULL | GSMG0029642 | NULL | NULL | 524.74 | 721.28 | -0.46 | 4.89E-02 |
| ENSMUSG00000029592 | GSMG0029648 | Usp30 | ubiquitin specific peptidase 30 | 150.88 | 128.58 | 0.23 | 5.74E-01 |
| ENSMUSG00000042010 | GSMG0029651 | Acacb | acetyl-Coenzyme A carboxylase beta | 186.39 | 116.61 | 0.67 | 4.05E-02 |
| ENSMUSG00000066952 | GSMG0029652 | Myo1h | myosin 1H | 662.09 | 488.41 | 0.44 | 2.14E-02 |
| ENSMUSG00000029577 | GSMG0029654 | Ube3b | ubiquitin protein ligase E3B | 2454.27 | 2751.90 | -0.17 | 4.85E-01 |
| ENSMUSG00000041939 | GSMG0029656 | Mvk | mevalonate kinase | 360.35 | 563.50 | -0.65 | 5.89E-04 |
| ENSMUSG00000002486 | GSMG0029659 | Tchp | trichoplein, keratin filament binding | 289.78 | 328.45 | -0.18 | 5.21E-01 |
| ENSMUSG00000041870 | GSMG0029660 | Ankrd13a | ankyrin repeat domain 13a | 6056.48 | 5118.40 | 0.24 | 1.97E-01 |
| NULL | GSMG0029661 | NULL | NULL | 351.11 | 356.01 | -0.02 | 9.62E-01 |
| ENSMUSG00000029561 | GSMG0029663 | Oasl2 | 2'-5' oligoadenylate synthetase-like 2 | 109.45 | 161.55 | -0.56 | 9.85E-02 |
| ENSMUSG00000029559 | GSMG0029666 | 2210016L21Rik | RIKEN cDNA 2210016L21 gene | 1849.94 | 1972.63 | -0.09 | 7.08E-01 |
| ENSMUSG00000029550 | GSMG0029669 | Sppl3 | signal peptide peptidase 3 | 795.01 | 700.76 | 0.18 | 6.21E-01 |
| ENSMUSG00000060152 | GSMG0029672 | Pop5 | processing of precursor 5, ribonuclease P/MRP family (S. cerevisiae) | 1706.94 | 1761.13 | -0.05 | 8.84E-01 |
| ENSMUSG00000041733 | GSMG0029673 | Coq5 | coenzyme Q5 methyltransferase | 360.96 | 328.76 | 0.14 | 6.83E-01 |
| ENSMUSG00000029538 | GSMG0029675 | Srsf9 | serine/arginine-rich splicing factor 9 | 1964.49 | 2089.99 | -0.09 | 7.18E-01 |
| ENSMUSG00000029535 | GSMG0029677 | Triap1 | TP53 regulated inhibitor of apoptosis 1 | 640.04 | 730.09 | -0.19 | 4.38E-01 |
| ENSMUSG00000029528 | GSMG0029681 | Pxn | paxillin | 1265.90 | 1315.30 | -0.06 | 8.60E-01 |
| ENSMUSG00000067274 | GSMG0029682 | Rplp0 | ribosomal protein, large, P0 | 4232.41 | 5923.35 | -0.48 | 9.50E-04 |
| ENSMUSG00000041638 | GSMG0029683 | Gcn1l1 | GCN1 general control of amino-acid synthesis 1-like 1 (yeast) | 1812.05 | 2134.03 | -0.24 | 4.20E-01 |
| ENSMUSG00000029518 | GSMG0029684 | Rab35 | RAB35, member RAS oncogene family | 617.87 | 611.05 | 0.01 | 9.79E-01 |
| ENSMUSG00000029516 | GSMG0029686 | Cit | citron | 577.34 | 411.65 | 0.49 | 2.84E-02 |
| ENSMUSG00000061288 | GSMG0029695 | Taok3 | TAO kinase 3 | 718.62 | 563.18 | 0.35 | 6.58E-02 |
| ENSMUSG00000066894 | GSMG0029701 | Vsig10 | V-set and immunoglobulin domain containing 10 | 392.66 | 369.23 | 0.09 | 8.03E-01 |
| ENSMUSG00000029364 | GSMG0029702 | Wsb2 | WD repeat and SOCS box-containing 2 | 2327.93 | 2848.82 | -0.29 | 1.06E-01 |

| | | | | | | | |
|---|---|---|---|---|---|---|---|
| ENSMUSG00000061578 | GSMG0029704 | Ksr2 | kinase suppressor of ras 2 | 53.04 | 40.46 | 0.39 | 4.83E-01 |
| ENSMUSG00000032898 | GSMG0029709 | Fbxo21 | F-box protein 21 | 1081.17 | 1338.79 | -0.31 | 1.40E-01 |
| ENSMUSG00000032840 | GSMG0029714 | 2410131K14Rik | RIKEN cDNA 2410131K14 gene | 450.15 | 545.39 | -0.28 | 2.56E-01 |
| ENSMUSG00000018076 | GSMG0029715 | Med13l | mediator complex subunit 13-like | 180.43 | 182.21 | -0.01 | 9.81E-01 |
| ENSMUSG00000018604 | GSMG0029724 | Tbx3 | T-box 3 | 1325.33 | 1006.93 | 0.39 | 1.78E-01 |
| ENSMUSG00000029594 | GSMG0029726 | Rbm19 | RNA binding motif protein 19 | 371.24 | 498.73 | -0.43 | 7.80E-02 |
| ENSMUSG00000032754 | GSMG0029731 | Slc8b1 | solute carrier family 8 (sodium/lithium/calcium exchanger), member B1 | 412.10 | 329.52 | 0.32 | 1.99E-01 |
| ENSMUSG00000029599 | GSMG0029734 | Ddx54 | DEAD (Asp-Glu-Ala-Asp) box polypeptide 54 | 1476.13 | 1582.41 | -0.10 | 7.04E-01 |
| ENSMUSG00000029602 | GSMG0029735 | Rasal1 | RAS protein activator like 1 (GAP1 like) | 1227.25 | 775.78 | 0.66 | 2.42E-04 |
| NULL | GSMG0029736 | NULL | NULL | 220.60 | 318.41 | -0.53 | 1.91E-02 |
| ENSMUSG00000042719 | GSMG0029748 | Naa25 | N(alpha)-acetyltransferase 25, NatB auxiliary subunit | 1093.17 | 1387.82 | -0.34 | 4.14E-02 |
| NULL | GSMG0029754 | NULL | NULL | 473.59 | 169.32 | 1.48 | 9.69E-10 |
| ENSMUSG00000029458 | GSMG0029756 | Brap | BRCA1 associated protein | 572.64 | 459.36 | 0.32 | 2.30E-01 |
| ENSMUSG00000042605 | GSMG0029757 | Atxn2 | ataxin 2 | 1768.15 | 1523.70 | 0.21 | 4.15E-01 |
| ENSMUSG00000044134 | GSMG0029761 | Fam109a | family with sequence similarity 109, member A | 424.23 | 443.43 | -0.06 | 8.68E-01 |
| ENSMUSG00000038582 | GSMG0029767 | Pptc7 | PTC7 protein phosphatase homolog | 356.91 | 261.83 | 0.45 | 6.03E-02 |
| ENSMUSG00000029462 | GSMG0029769 | Vps29 | VPS29 retromer complex component | 2696.07 | 3091.13 | -0.20 | 4.31E-01 |
| ENSMUSG00000029464 | GSMG0029770 | Gpn3 | GPN-loop GTPase 3 | 392.68 | 448.26 | -0.19 | 4.77E-01 |
| ENSMUSG00000029465 | GSMG0029771 | Arpc3 | actin related protein 2/3 complex, subunit 3 | 4582.50 | 4993.06 | -0.12 | 6.08E-01 |
| ENSMUSG00000029466 | GSMG0029773 | Anapc7 | anaphase promoting complex subunit 7 | 899.53 | 976.76 | -0.12 | 6.24E-01 |
| ENSMUSG00000029468 | GSMG0029777 | P2rx7 | purinergic receptor P2X, ligand-gated ion channel, 7 | 144.87 | 63.26 | 1.20 | 1.07E-04 |
| ENSMUSG00000029470 | GSMG0029779 | P2rx4 | purinergic receptor P2X, ligand-gated ion channel 4 | 399.93 | 289.80 | 0.47 | 3.48E-02 |
| ENSMUSG00000029474 | GSMG0029781 | Rnf34 | ring finger protein 34 | 694.38 | 615.99 | 0.17 | 4.84E-01 |
| ENSMUSG00000049686 | GSMG0029783 | Orai1 | ORAI calcium release-activated calcium modulator 1 | 1354.61 | 1384.55 | -0.03 | 9.21E-01 |
| ENSMUSG00000054434 | GSMG0029785 | Tmem120b | transmembrane protein 120B | 173.17 | 282.64 | -0.70 | 4.70E-03 |
| NULL | GSMG0029786 | NULL | NULL | 200.63 | 180.34 | 0.15 | 6.98E-01 |
| ENSMUSG00000029440 | GSMG0029791 | Psmd9 | proteasome (prosome, macropain) 26S subunit, non-ATPase, 9 | 782.44 | 958.77 | -0.30 | 1.49E-01 |
| ENSMUSG00000029438 | GSMG0029793 | Bcl7a | B cell CLL/lymphoma 7A | 281.06 | 415.66 | -0.57 | 2.90E-02 |
| ENSMUSG00000038342 | GSMG0029795 | Mlxip | MLX interacting protein | 396.40 | 510.70 | -0.36 | 1.86E-01 |
| ENSMUSG00000023106 | GSMG0029803 | Denr | density-regulated protein | 966.37 | 1268.39 | -0.39 | 7.38E-02 |
| ENSMUSG00000000915 | GSMG0029806 | Hip1r | huntingtin interacting protein 1 related | 1351.08 | 1161.59 | 0.22 | 4.88E-01 |
| NULL | GSMG0029807 | NULL | NULL | 339.87 | 282.51 | 0.27 | 3.14E-01 |
| ENSMUSG00000023707 | GSMG0029809 | Ogfod2 | 2-oxoglutarate and iron-dependent oxygenase domain containing 2 | 268.41 | 233.54 | 0.20 | 5.58E-01 |

| | | | | | | | |
|---|---|---|---|---|---|---|---|
| ENSMUSG00000029404 | GSMG0029810 | Arl6ip4 | ADP-ribosylation factor-like 6 interacting protein 4 | 1039.68 | 1258.61 | -0.28 | 1.81E-01 |
| ENSMUSG00000047635 | GSMG0029814 | 2810006K23Rik | RIKEN cDNA 2810006K23 gene | 270.82 | 313.81 | -0.21 | 5.06E-01 |
| ENSMUSG00000049327 | GSMG0029815 | Kmt5a | lysine methyltransferase 5A | 495.30 | 485.38 | 0.03 | 9.46E-01 |
| ENSMUSG00000029402 | GSMG0029816 | Snrnp35 | small nuclear ribonucleoprotein 35 (U11/U12) | 187.89 | 252.17 | -0.42 | 1.25E-01 |
| ENSMUSG00000029389 | GSMG0029819 | Ddx55 | DEAD (Asp-Glu-Ala-Asp) box polypeptide 55 | 615.90 | 725.95 | -0.24 | 2.46E-01 |
| ENSMUSG00000029387 | GSMG0029820 | Gtf2h3 | general transcription factor IIH, polypeptide 3 | 482.85 | 523.04 | -0.11 | 6.86E-01 |
| ENSMUSG00000029386 | GSMG0029821 | Tctn2 | tectonic family member 2 | 599.83 | 712.05 | -0.25 | 2.71E-01 |
| ENSMUSG00000079215 | GSMG0029824 | Zfp664 | zinc finger protein 664 | 3505.28 | 3655.48 | -0.06 | 8.22E-01 |
| ENSMUSG00000037962 | GSMG0029826 | Fam101a | family with sequence similarity 101, member A | 445.06 | 477.32 | -0.10 | 7.27E-01 |
| ENSMUSG00000037905 | GSMG0029829 | Bri3bp | Bri3 binding protein | 580.11 | 857.27 | -0.56 | 7.47E-04 |
| ENSMUSG00000029482 | GSMG0029831 | Aacs | acetoacetyl-CoA synthetase | 884.62 | 1613.13 | -0.87 | 3.45E-07 |
| ENSMUSG00000070498 | GSMG0029833 | Tmem132b | transmembrane protein 132B | 1178.30 | 804.82 | 0.55 | 3.36E-02 |
| ENSMUSG00000029439 | GSMG0029849 | Sfswap | splicing factor, suppressor of white-apricot homolog (Drosophila) | 1558.64 | 2098.04 | -0.43 | 1.26E-02 |
| NULL | GSMG0029850 | NULL | NULL | 644.92 | 521.98 | 0.31 | 1.62E-01 |
| ENSMUSG00000029436 | GSMG0029851 | Mmp17 | matrix metallopeptidase 17 | 710.99 | 490.91 | 0.54 | 2.78E-02 |
| ENSMUSG00000034211 | GSMG0029852 | Mrps17 | mitochondrial ribosomal protein S17 | 1356.61 | 1498.66 | -0.14 | 5.72E-01 |
| ENSMUSG00000029432 | GSMG0029854 | Gbas | glioblastoma amplified sequence | 689.56 | 710.28 | -0.04 | 8.83E-01 |
| ENSMUSG00000025538 | GSMG0029859 | Sumf2 | sulfatase modifying factor 2 | 642.18 | 607.75 | 0.08 | 7.74E-01 |
| ENSMUSG00000066735 | GSMG0029862 | Vkorc1l1 | vitamin K epoxide reductase complex, subunit 1-like 1 | 931.64 | 1096.53 | -0.24 | 3.57E-01 |
| ENSMUSG00000025532 | GSMG0029866 | Crcp | calcitonin gene-related peptide-receptor component protein | 635.94 | 843.18 | -0.41 | 4.89E-02 |
| ENSMUSG00000034118 | GSMG0029867 | Tpst1 | protein-tyrosine sulfotransferase 1 | 2521.28 | 2577.48 | -0.03 | 9.29E-01 |
| ENSMUSG00000034110 | GSMG0029870 | Kctd7 | potassium channel tetramerisation domain containing 7 | 585.74 | 534.05 | 0.13 | 6.87E-01 |
| ENSMUSG00000025340 | GSMG0029871 | Rabgef1 | RAB guanine nucleotide exchange factor (GEF) 1 | 505.02 | 547.28 | -0.11 | 7.02E-01 |
| ENSMUSG00000053094 | GSMG0029872 | Tmem248 | transmembrane protein 248 | 957.20 | 1074.95 | -0.17 | 4.49E-01 |
| ENSMUSG00000056310 | GSMG0029874 | Tyw1 | tRNA-yW synthesizing protein 1 homolog (S. cerevisiae) | 375.63 | 457.30 | -0.28 | 2.44E-01 |
| ENSMUSG00000060371 | GSMG0029875 | Caln1 | calneuron 1 | 396.16 | 690.17 | -0.80 | 1.87E-05 |
| NULL | GSMG0029876 | NULL | NULL | 117.35 | 176.77 | -0.60 | 8.24E-02 |
| ENSMUSG00000015944 | GSMG0029884 | Gatsl2 | GATS protein-like 2 | 167.76 | 200.20 | -0.26 | 4.81E-01 |
| NULL | GSMG0029885 | NULL | NULL | 267.39 | 234.61 | 0.19 | 5.58E-01 |
| ENSMUSG00000015942 | GSMG0029887 | Gtf2ird2 | GTF2I repeat domain containing 2 | 139.28 | 156.24 | -0.16 | 7.01E-01 |
| ENSMUSG00000023104 | GSMG0029888 | Rfc2 | replication factor C (activator 1) 2 | 1010.76 | 1352.25 | -0.42 | 1.15E-02 |
| ENSMUSG00000040557 | GSMG0029893 | Wbscr27 | Williams Beuren syndrome chromosome region 27 (human) | 330.41 | 357.44 | -0.12 | 7.42E-01 |
| ENSMUSG00000040532 | GSMG0029895 | Abhd11 | abhydrolase domain containing 11 | 1044.31 | 1352.92 | -0.37 | 3.35E-02 |

| | | | | | | | |
|---|---|---|---|---|---|---|---|
| ENSMUSG00000007207 | GSMG0029896 | Stx1a | syntaxin 1A (brain) | 1538.43 | 1973.11 | -0.36 | 2.70E-02 |
| ENSMUSG00000061118 | GSMG0029897 | Dnajc30 | DnaJ heat shock protein family (Hsp40) member C30 | 613.69 | 574.54 | 0.10 | 7.63E-01 |
| ENSMUSG00000005374 | GSMG0029899 | Tbl2 | transducin (beta)-like 2 | 162.81 | 183.56 | -0.17 | 6.22E-01 |
| ENSMUSG00000029681 | GSMG0029900 | Bcl7b | B cell CLL/lymphoma 7B | 1368.57 | 1210.31 | 0.18 | 3.76E-01 |
| ENSMUSG00000002748 | GSMG0029901 | Baz1b | bromodomain adjacent to zinc finger domain, 1B | 315.21 | 349.66 | -0.15 | 6.11E-01 |
| ENSMUSG00000000916 | GSMG0029904 | Nsun5 | NOL1/NOP2/Sun domain family, member 5 | 464.65 | 535.03 | -0.20 | 4.19E-01 |
| ENSMUSG00000039917 | GSMG0029905 | Rhbdd2 | rhomboid domain containing 2 | 811.44 | 1066.26 | -0.39 | 4.15E-02 |
| ENSMUSG00000005514 | GSMG0029906 | Por | P450 (cytochrome) oxidoreductase | 2515.38 | 2452.74 | 0.04 | 9.13E-01 |
| ENSMUSG00000019179 | GSMG0029908 | Mdh2 | malate dehydrogenase 2, NAD (mitochondrial) | 5656.46 | 6868.30 | -0.28 | 1.19E-01 |
| ENSMUSG00000039860 | GSMG0029909 | Srrm3 | serine/arginine repetitive matrix 3 | 629.29 | 601.58 | 0.06 | 8.45E-01 |
| NULL | GSMG0029910 | NULL | NULL | 732.75 | 655.83 | 0.16 | 6.02E-01 |
| ENSMUSG00000004947 | GSMG0029914 | Dtx2 | deltex 2, E3 ubiquitin ligase | 316.36 | 402.82 | -0.35 | 1.34E-01 |
| ENSMUSG00000004952 | GSMG0029917 | Rasa4 | RAS p21 protein activator 4 | 286.70 | 526.94 | -0.88 | 9.15E-06 |
| ENSMUSG00000039771 | GSMG0029918 | Polr2j | polymerase (RNA) II (DNA directed) polypeptide J | 932.98 | 1102.61 | -0.24 | 2.45E-01 |
| ENSMUSG00000039754 | GSMG0029919 | Alkbh4 | alkB homolog 4, lysine demethylase | 409.89 | 428.16 | -0.06 | 8.53E-01 |
| ENSMUSG00000007987 | GSMG0029922 | Ift22 | intraflagellar transport 22 | 267.31 | 294.62 | -0.14 | 6.67E-01 |
| ENSMUSG00000019054 | GSMG0029925 | Fis1 | fission, mitochondrial 1 | 2950.13 | 2885.00 | 0.03 | 9.04E-01 |
| ENSMUSG00000004846 | GSMG0029927 | Plod3 | procollagen-lysine, 2-oxoglutarate 5-dioxygenase 3 | 2192.72 | 2307.45 | -0.07 | 7.77E-01 |
| ENSMUSG00000037428 | GSMG0029928 | Vgf | VGF nerve growth factor inducible | 1012.47 | 472.87 | 1.10 | 1.27E-04 |
| ENSMUSG00000023328 | GSMG0029929 | Ache | acetylcholinesterase | 558.11 | 857.92 | -0.62 | 4.21E-04 |
| ENSMUSG00000051502 | GSMG0029930 | Ufsp1 | UFM1-specific peptidase 1 | 81.86 | 156.41 | -0.93 | 1.49E-03 |
| ENSMUSG00000029710 | GSMG0029931 | Ephb4 | Eph receptor B4 | 4576.80 | 4393.58 | 0.06 | 8.34E-01 |
| ENSMUSG00000029714 | GSMG0029933 | Gigyf1 | GRB10 interacting GYF protein 1 | 684.45 | 670.79 | 0.03 | 9.39E-01 |
| NULL | GSMG0029934 | NULL | NULL | 1324.70 | 1695.70 | -0.36 | 1.04E-01 |
| ENSMUSG00000029712 | GSMG0029935 | Actl6b | actin-like 6B | 892.96 | 1003.40 | -0.17 | 5.19E-01 |
| ENSMUSG00000029716 | GSMG0029936 | Tfr2 | transferrin receptor 2 | 355.60 | 453.55 | -0.35 | 1.38E-01 |
| 20,ENSMUSG00000079165,E | GSMG0029938 | Gm20605 // Lrch4 | predicted gene 20605 // leucine-rich repeats and calponin homology (C | 1457.66 | 1374.69 | 0.08 | 7.58E-01 |
| ENSMUSG00000029723 | GSMG0029939 | Tsc22d4 | TSC22 domain family, member 4 | 291.48 | 396.45 | -0.45 | 5.57E-02 |
| ENSMUSG00000029725 | GSMG0029940 | Ppp1r35 | protein phosphatase 1, regulatory subunit 35 | 167.67 | 180.64 | -0.11 | 8.08E-01 |
| ENSMUSG00000037108 | GSMG0029941 | Zcwpw1 | zinc finger, CW type with PWWP domain 1 | 202.78 | 260.31 | -0.36 | 2.30E-01 |
| ENSMUSG00000029729 | GSMG0029947 | Zkscan1 | zinc finger with KRAB and SCAN domains 1 | 186.60 | 227.44 | -0.28 | 4.12E-01 |
| ENSMUSG00000037017 | GSMG0029948 | Zscan21 | zinc finger and SCAN domain containing 21 | 386.73 | 465.43 | -0.26 | 2.71E-01 |
| ENSMUSG00000019494 | GSMG0029949 | Cops6 | COP9 signalosome subunit 6 | 2225.81 | 2604.01 | -0.23 | 2.26E-01 |

| | | | | | | | |
|---|---|---|---|---|---|---|---|
| ENSMUSG00000019518 | GSMG0029950 | Ap4m1 | adaptor-related protein complex AP-4, mu 1 | 158.66 | 197.83 | -0.31 | 3.50E-01 |
| ENSMUSG00000036968 | GSMG0029952 | Cnpy4 | canopy FGF signaling regulator 4 | 935.18 | 1059.31 | -0.18 | 4.63E-01 |
| ENSMUSG00000049285 | GSMG0029953 | Mblac1 | metallo-beta-lactamase domain containing 1 | 176.10 | 188.55 | -0.10 | 8.26E-01 |
| ENSMUSG00000050552 | GSMG0029955 | Lamtor4 | late endosomal/lysosomal adaptor, MAPK and MTOR activator 4 | 545.73 | 528.51 | 0.04 | 8.95E-01 |
| ENSMUSG00000036928 | GSMG0029957 | Stag3 | stromal antigen 3 | 372.33 | 1083.23 | -1.54 | 8.45E-22 |
| ENSMUSG00000036898 | GSMG0029959 | Zfp157 | zinc finger protein 157 | 779.04 | 891.45 | -0.19 | 3.54E-01 |
| ENSMUSG00000056014 | GSMG0029961 | A430033K04Rik | RIKEN cDNA A430033K04 gene | 272.04 | 376.34 | -0.46 | 4.83E-02 |
| ENSMUSG00000025854 | GSMG0029962 | Fam20c | family with sequence similarity 20, member C | 590.97 | 834.84 | -0.50 | 5.88E-03 |
| ENSMUSG00000025857 | GSMG0029965 | Dnaaf5 | dynein, axonemal assembly factor 5 | 422.01 | 390.88 | 0.11 | 7.64E-01 |
| ENSMUSG00000036817 | GSMG0029966 | Sun1 | Sad1 and UNC84 domain containing 1 | 4728.23 | 3617.25 | 0.39 | 1.32E-02 |
| ENSMUSG00000025858 | GSMG0029967 | Get4 | golgi to ER traffic protein 4 | 2672.79 | 3220.09 | -0.27 | 1.31E-01 |
| ENSMUSG00000044197 | GSMG0029969 | Gpr146 | G protein-coupled receptor 146 | 3480.24 | 1645.09 | 1.08 | 1.15E-13 |
| ENSMUSG00000053647 | GSMG0029971 | Gper1 | G protein-coupled estrogen receptor 1 | 3048.25 | 2103.54 | 0.54 | 5.82E-04 |
| ENSMUSG00000018143 | GSMG0029975 | Mafk | v-maf musculoaponeurotic fibrosarcoma oncogene family, protein K (av | 1572.27 | 758.45 | 1.05 | 6.52E-12 |
| ENSMUSG00000036639 | GSMG0029982 | Nudt1 | nudix (nucleoside diphosphate linked moiety X)-type motif 1 | 125.45 | 154.23 | -0.29 | 4.18E-01 |
| ENSMUSG00000056076 | GSMG0029985 | Eif3b | eukaryotic translation initiation factor 3, subunit B | 6219.67 | 8116.09 | -0.38 | 9.41E-03 |
| ENSMUSG00000036599 | GSMG0029986 | Chst12 | carbohydrate sulfotransferase 12 | 603.30 | 761.77 | -0.33 | 1.55E-01 |
| ENSMUSG00000029570 | GSMG0029988 | Lfng | LFNG O-fucosylpeptide 3-beta-N-acetylglucosaminyltransferase | 726.41 | 472.12 | 0.62 | 1.41E-02 |
| ENSMUSG00000000148 | GSMG0029990 | Brat1 | BRCA1-associated ATM activator 1 | 678.15 | 573.42 | 0.24 | 4.24E-01 |
| ENSMUSG00000056493 | GSMG0030003 | Foxk1 | forkhead box K1 | 126.60 | 129.75 | -0.04 | 9.50E-01 |
| ENSMUSG00000039623 | GSMG0030004 | Ap5z1 | adaptor-related protein complex 5, zeta 1 subunit | 236.93 | 195.57 | 0.28 | 3.59E-01 |
| ENSMUSG00000029578 | GSMG0030005 | Wipi2 | WD repeat domain, phosphoinositide interacting 2 | 1189.58 | 1215.03 | -0.03 | 9.28E-01 |
| ENSMUSG00000029581 | GSMG0030007 | Fscn1 | fascin actin-bundling protein 1 | 5593.59 | 12426.24 | -1.15 | 1.64E-02 |
| ENSMUSG00000029587 | GSMG0030012 | Zfp12 | zinc finger protein 12 | 1089.12 | 971.24 | 0.17 | 5.31E-01 |
| ENSMUSG00000079111 | GSMG0030018 | Kdelr2 | KDEL (Lys-Asp-Glu-Leu) endoplasmic reticulum protein retention rece | 794.72 | 998.74 | -0.33 | 1.45E-01 |
| ENSMUSG00000039206 | GSMG0030020 | Daglb | diacylglycerol lipase, beta | 406.36 | 336.56 | 0.27 | 3.60E-01 |
| ENSMUSG00000083012 | GSMG0030021 | Fam220a | family with sequence similarity 220, member A | 510.04 | 628.77 | -0.30 | 1.55E-01 |
| ENSMUSG00000018001 | GSMG0030023 | Cyth3 | cytohesin 3 | 1371.44 | 1077.88 | 0.35 | 3.96E-02 |
| ENSMUSG00000029613 | GSMG0030026 | Eif2ak1 | eukaryotic translation initiation factor 2 alpha kinase 1 | 1746.87 | 2049.96 | -0.23 | 2.35E-01 |
| ENSMUSG00000038970 | GSMG0030029 | Lmtk2 | lemur tyrosine kinase 2 | 820.53 | 898.03 | -0.13 | 7.36E-01 |
| NULL | GSMG0030031 | NULL | NULL | 321.06 | 294.60 | 0.13 | 7.03E-01 |
| ENSMUSG00000059991 | GSMG0030037 | Nptx2 | neuronal pentraxin 2 | 113.19 | 46.41 | 1.29 | 7.19E-03 |
| ENSMUSG00000045482 | GSMG0030038 | Trrap | transformation/transcription domain-associated protein | 1584.21 | 1612.19 | -0.03 | 9.32E-01 |

| | | | | | | | |
|---|---|---|---|---|---|---|---|
| ENSMUSG00000029621 | GSMG0030040 | Arpc1a | actin related protein 2/3 complex, subunit 1A | 2281.87 | 2033.07 | 0.17 | 4.34E-01 |
| ENSMUSG00000029622 | GSMG0030041 | Arpc1b | actin related protein 2/3 complex, subunit 1B | 7320.50 | 6160.72 | 0.25 | 1.53E-01 |
| ENSMUSG00000038722 | GSMG0030043 | Bud31 | BUD31 homolog | 1087.47 | 1345.37 | -0.31 | 2.04E-01 |
| ENSMUSG00000029625 | GSMG0030044 | Cpsf4 | cleavage and polyadenylation specific factor 4 | 363.47 | 388.23 | -0.10 | 8.00E-01 |
| ENSMUSG00000055991 | GSMG0030045 | Zkscan5 | zinc finger with KRAB and SCAN domains 5 | 107.17 | 122.21 | -0.19 | 6.63E-01 |
| ENSMUSG00000007812 | GSMG0030046 | Zfp655 | zinc finger protein 655 | 934.13 | 880.85 | 0.08 | 7.69E-01 |
| ENSMUSG00000029635 | GSMG0030049 | Cdk8 | cyclin-dependent kinase 8 | 225.37 | 238.89 | -0.09 | 8.19E-01 |
| ENSMUSG00000029636 | GSMG0030050 | Wasf3 | WAS protein family, member 3 | 160.68 | 149.77 | 0.10 | 8.18E-01 |
| ENSMUSG00000041453 | GSMG0030056 | Rpl21 | ribosomal protein L21 | 141.32 | 410.90 | -1.54 | 1.44E-15 |
| ENSMUSG00000029641 | GSMG0030057 | Rasl11a | RAS-like, family 11, member A | 349.13 | 211.26 | 0.73 | 1.86E-03 |
| ENSMUSG00000016503 | GSMG0030058 | Gtf3a | general transcription factor III A | 202.05 | 243.74 | -0.27 | 4.13E-01 |
| ENSMUSG00000029642 | GSMG0030059 | Polr1d | polymerase (RNA) I polypeptide D | 958.62 | 1099.06 | -0.20 | 4.08E-01 |
| ENSMUSG00000029647 | GSMG0030066 | Pan3 | PAN3 poly(A) specific ribonuclease subunit | 516.09 | 557.69 | -0.11 | 7.28E-01 |
| ENSMUSG00000029649 | GSMG0030072 | Pomp | proteasome maturation protein | 4991.43 | 6007.84 | -0.27 | 2.44E-01 |
| ENSMUSG00000029651 | GSMG0030073 | Mtus2 | microtubule associated tumor suppressor candidate 2 | 104.78 | 136.03 | -0.37 | 2.69E-01 |
| ENSMUSG00000041264 | GSMG0030085 | Uspl1 | ubiquitin specific peptidase like 1 | 348.84 | 439.50 | -0.33 | 1.32E-01 |
| ENSMUSG00000051950 | GSMG0030094 | B3glct | beta-3-glucosyltransferase | 418.58 | 596.44 | -0.51 | 5.91E-03 |
| NULL | GSMG0030103 | NULL | NULL | 77.04 | 56.14 | 0.46 | 2.92E-01 |
| NULL | GSMG0030106 | NULL | NULL | 398.38 | 272.29 | 0.55 | 3.72E-02 |
| ENSMUSG00000034021 | GSMG0030110 | Pds5b | PDS5 cohesin associated factor B | 945.71 | 1129.89 | -0.26 | 1.80E-01 |
| NULL | GSMG0030111 | NULL | NULL | 292.40 | 276.38 | 0.08 | 8.15E-01 |
| ENSMUSG00000040302 | GSMG0030123 | Rbm48 | RNA binding motif protein 48 | 199.84 | 215.46 | -0.11 | 7.62E-01 |
| ENSMUSG00000040351 | GSMG0030126 | Ankib1 | ankyrin repeat and IBR domain containing 1 | 788.80 | 1008.49 | -0.35 | 7.00E-02 |
| ENSMUSG00000001467 | GSMG0030130 | Cyp51 | cytochrome P450, family 51 | 855.38 | 1355.81 | -0.66 | 3.44E-05 |
| ENSMUSG00000044674 | GSMG0030132 | Fzd1 | frizzled class receptor 1 | 1103.48 | 950.02 | 0.21 | 4.87E-01 |
| ENSMUSG00000028926 | GSMG0030134 | Cdk14 | cyclin-dependent kinase 14 | 532.41 | 552.74 | -0.05 | 8.91E-01 |
| ENSMUSG00000046798 | GSMG0030145 | Cldn12 | claudin 12 | 1318.72 | 923.76 | 0.51 | 8.77E-03 |
| NULL | GSMG0030153 | NULL | NULL | 1187.13 | 1344.95 | -0.18 | 5.45E-01 |
| ENSMUSG00000040537 | GSMG0030161 | Adam22 | a disintegrin and metallopeptidase domain 22 | 175.66 | 157.45 | 0.16 | 6.70E-01 |
| ENSMUSG00000002297 | GSMG0030164 | Dbf4 | DBF4 zinc finger | 142.80 | 307.86 | -1.11 | 3.30E-06 |
| ENSMUSG00000040570 | GSMG0030165 | Rundc3b | RUN domain containing 3B | 277.02 | 548.33 | -0.99 | 2.94E-07 |
| ENSMUSG00000042508 | GSMG0030168 | Dmtf1 | cyclin D binding myb-like transcription factor 1 | 541.77 | 576.45 | -0.09 | 7.70E-01 |
| ENSMUSG00000003623 | GSMG0030170 | Crot | carnitine O-octanoyltransferase | 1231.29 | 1095.20 | 0.17 | 4.50E-01 |

| | | | | | | | |
|---|---|---|---|---|---|---|---|
| NULL | GSMG0030173 | NULL | NULL | 318.90 | 667.51 | -1.07 | 1.61E-05 |
| ENSMUSG00000003974 | GSMG0030175 | Grm3 | glutamate receptor, metabotropic 3 | 855.48 | 1247.27 | -0.54 | 2.68E-03 |
| NULL | GSMG0030178 | NULL | NULL | 218.15 | 215.88 | 0.01 | 9.81E-01 |
| ENSMUSG00000057614 | GSMG0030199 | Gnai1 | guanine nucleotide binding protein (G protein), alpha inhibiting 1 | 487.73 | 805.75 | -0.73 | 4.36E-05 |
| ENSMUSG00000039987 | GSMG0030205 | Phtf2 | putative homeodomain transcription factor 2 | 170.94 | 226.11 | -0.40 | 1.58E-01 |
| ENSMUSG00000039968 | GSMG0030209 | Rsbn1l | round spermatid basic protein 1-like | 167.29 | 173.87 | -0.06 | 8.99E-01 |
| ENSMUSG00000028771 | GSMG0030211 | Ptpn12 | protein tyrosine phosphatase, non-receptor type 12 | 915.67 | 945.20 | -0.05 | 8.80E-01 |
| ENSMUSG00000044968 | GSMG0030220 | Napepld | N-acyl phosphatidylethanolamine phospholipase D | 245.40 | 167.29 | 0.55 | 3.25E-02 |
| ENSMUSG00000029014 | GSMG0030222 | Dnajc2 | DnaJ heat shock protein family (Hsp40) member C2 | 380.90 | 557.36 | -0.55 | 2.70E-02 |
| ENSMUSG00000042453 | GSMG0030224 | Reln | reelin | 117.79 | 252.07 | -1.09 | 8.89E-06 |
| ENSMUSG00000029012 | GSMG0030229 | Orc5 | origin recognition complex, subunit 5 | 530.72 | 570.89 | -0.10 | 7.36E-01 |
| NULL | GSMG0030232 | NULL | NULL | 82.67 | 70.02 | 0.23 | 6.66E-01 |
| ENSMUSG00000062604 | GSMG0030237 | Srpk2 | serine/arginine-rich protein specific kinase 2 | 498.29 | 553.90 | -0.15 | 5.87E-01 |
| NULL | GSMG0030239 | NULL | NULL | 93.70 | 225.71 | -1.27 | 1.39E-06 |
| ENSMUSG00000057541 | GSMG0030240 | Pus7 | pseudouridylate synthase 7 | 645.32 | 844.19 | -0.39 | 4.25E-02 |
| ENSMUSG00000028998 | GSMG0030243 | Tomm7 | translocase of outer mitochondrial membrane 7 homolog (yeast) | 1016.71 | 1085.34 | -0.10 | 7.75E-01 |
| ENSMUSG00000028995 | GSMG0030245 | Fam126a | family with sequence similarity 126, member A | 417.66 | 565.60 | -0.44 | 4.74E-02 |
| NULL | GSMG0030250 | NULL | NULL | 670.31 | 561.38 | 0.25 | 3.72E-01 |
| NULL | GSMG0030252 | NULL | NULL | 35.72 | 51.94 | -0.54 | 3.31E-01 |
| ENSMUSG00000028969 | GSMG0030253 | Cdk5 | cyclin-dependent kinase 5 | 1846.57 | 1430.72 | 0.37 | 4.66E-02 |
| ENSMUSG00000028959 | GSMG0030254 | Fastk | Fas-activated serine/threonine kinase | 3740.31 | 2837.10 | 0.40 | 9.47E-03 |
| ENSMUSG00000028958 | GSMG0030255 | Tmub1 | transmembrane and ubiquitin-like domain containing 1 | 122.91 | 155.18 | -0.33 | 3.70E-01 |
| ENSMUSG00000028953 | GSMG0030261 | Abcf2 | ATP-binding cassette, sub-family F (GCN20), member 2 | 2292.93 | 2447.10 | -0.09 | 7.14E-01 |
| ENSMUSG00000028949 | GSMG0030262 | Smarcd3 | SWI/SNF related, matrix associated, actin dependent regulator of chron | 735.29 | 971.15 | -0.40 | 3.82E-02 |
| ENSMUSG00000028945 | GSMG0030267 | Rheb | Ras homolog enriched in brain | 1149.44 | 1313.89 | -0.19 | 5.07E-01 |
| ENSMUSG00000028944 | GSMG0030269 | Prkag2 | protein kinase, AMP-activated, gamma 2 non-catalytic subunit | 552.86 | 614.57 | -0.15 | 5.78E-01 |
| NULL | GSMG0030270 | NULL | NULL | 168.62 | 121.33 | 0.47 | 1.84E-01 |
| ENSMUSG00000038056 | GSMG0030275 | Kmt2c | lysine (K)-specific methyltransferase 2C | 150.21 | 169.71 | -0.17 | 6.91E-01 |
| ENSMUSG00000002221 | GSMG0030289 | Paxip1 | PAX interacting (with transcription-activation domain) protein 1 | 1232.11 | 1390.96 | -0.17 | 4.72E-01 |
| ENSMUSG00000010721 | GSMG0030297 | Lmbr1 | limb region 1 | 975.76 | 1260.48 | -0.37 | 4.71E-02 |
| ENSMUSG00000025747 | GSMG0030307 | Tyms | thymidylate synthase | 260.32 | 660.50 | -1.34 | 1.60E-11 |
| ENSMUSG00000025745 | GSMG0030309 | Hadha | hydroxyacyl-Coenzyme A dehydrogenase/3-ketoacyl-Coenzyme A thiol | 3032.38 | 3761.13 | -0.31 | 5.73E-02 |
| ENSMUSG00000062372 | GSMG0030311 | Otof | otoferlin | 820.37 | 659.49 | 0.31 | 3.59E-01 |

| | | | | | | | |
|---|---|---|---|---|---|---|---|
| NULL | GSMG0030314 | NULL | NULL | 309.92 | 282.20 | 0.13 | 6.93E-01 |
| ENSMUSG00000038803 | GSMG0030315 | Ost4 | oligosaccharyltransferase complex subunit 4 (non-catalytic) | 1746.01 | 1856.77 | -0.09 | 7.75E-01 |
| ENSMUSG00000029161 | GSMG0030316 | Cgref1 | cell growth regulator with EF hand domain 1 | 398.31 | 793.88 | -1.00 | 9.36E-09 |
| ENSMUSG00000045302 | GSMG0030317 | Preb | prolactin regulatory element binding | 1607.45 | 1550.37 | 0.05 | 8.47E-01 |
| ENSMUSG00000029151 | GSMG0030319 | Slc30a3 | solute carrier family 30 (zinc transporter), member 3 | 1030.15 | 189.37 | 2.44 | 2.25E-30 |
| G00000106864,ENSMUSG000 | GSMG0030323 | Mpv17 // Gtf3c2 | MpV17 mitochondrial inner membrane protein // general transcription f | 1755.09 | 2027.13 | -0.21 | 3.09E-01 |
| ENSMUSG00000029145 | GSMG0030324 | Eif2b4 | eukaryotic translation initiation factor 2B, subunit 4 delta | 1481.08 | 1565.95 | -0.08 | 7.44E-01 |
| ENSMUSG00000043059 | GSMG0030325 | Zfp513 | zinc finger protein 513 | 1012.87 | 741.03 | 0.45 | 7.55E-02 |
| ENSMUSG00000029147 | GSMG0030326 | Ppm1g | protein phosphatase 1G (formerly 2C), magnesium-dependent, gamma i | 2072.99 | 2740.86 | -0.40 | 1.07E-02 |
| NULL | GSMG0030329 | NULL | NULL | 818.31 | 780.55 | 0.07 | 8.09E-01 |
| ENSMUSG00000038564 | GSMG0030330 | Ift172 | intraflagellar transport 172 | 2849.17 | 2277.14 | 0.32 | 1.33E-01 |
| ENSMUSG00000038552 | GSMG0030331 | Fndc4 | fibronectin type III domain containing 4 | 1079.72 | 1406.29 | -0.38 | 8.95E-02 |
| ENSMUSG00000053134 | GSMG0030334 | Supt7l | suppressor of Ty 7-like | 256.22 | 259.45 | -0.02 | 9.71E-01 |
| ENSMUSG00000029136 | GSMG0030337 | Rbks | ribokinase | 99.15 | 69.02 | 0.53 | 1.92E-01 |
| ENSMUSG00000023452 | GSMG0030346 | Pisd | phosphatidylserine decarboxylase | 1575.64 | 1510.84 | 0.06 | 8.53E-01 |
| ENSMUSG00000037379 | GSMG0030354 | Spon2 | spondin 2, extracellular matrix protein | 100.35 | 181.57 | -0.85 | 6.01E-03 |
| ENSMUSG00000037373 | GSMG0030355 | Ctbp1 | C-terminal binding protein 1 | 3299.81 | 3753.32 | -0.19 | 4.80E-01 |
| ENSMUSG00000037339 | GSMG0030357 | Fam53a | family with sequence similarity 53, member A | 580.97 | 618.35 | -0.09 | 7.94E-01 |
| ENSMUSG00000005299 | GSMG0030358 | Letm1 | leucine zipper-EF-hand containing transmembrane protein 1 | 1531.92 | 1779.03 | -0.22 | 2.78E-01 |
| ENSMUSG00000004642 | GSMG0030359 | Slbp | stem-loop binding protein | 680.87 | 946.82 | -0.48 | 7.63E-02 |
| ENSMUSG00000019295 | GSMG0030360 | Tmem129 | transmembrane protein 129 | 386.51 | 306.07 | 0.34 | 2.03E-01 |
| ENSMUSG00000029111 | GSMG0030363 | Nelfa | negative elongation factor complex member A, Whsc2 | 923.23 | 1018.01 | -0.14 | 5.83E-01 |
| G00000045102,ENSMUSG000 | GSMG0030366 | Poln // Haus3 | DNA polymerase N // HAUS augmin-like complex, subunit 3 | 559.79 | 573.70 | -0.03 | 9.29E-01 |
| NULL | GSMG0030368 | NULL | NULL | 843.94 | 685.61 | 0.30 | 1.17E-01 |
| NULL | GSMG0030369 | NULL | NULL | 52.15 | 49.93 | 0.07 | 9.21E-01 |
| ENSMUSG00000037235 | GSMG0030370 | Mxd4 | Max dimerization protein 4 | 1093.47 | 749.17 | 0.54 | 7.09E-04 |
| ENSMUSG00000037224 | GSMG0030371 | Zfyve28 | zinc finger, FYVE domain containing 28 | 325.90 | 147.37 | 1.15 | 8.93E-05 |
| ENSMUSG00000059866 | GSMG0030373 | Tnip2 | TNFAIP3 interacting protein 2 | 654.05 | 485.08 | 0.43 | 2.11E-02 |
| ENSMUSG00000001082 | GSMG0030374 | Mfsd10 | major facilitator superfamily domain containing 10 | 2070.83 | 2777.97 | -0.42 | 1.11E-02 |
| ENSMUSG00000036693 | GSMG0030375 | Nop14 | NOP14 nucleolar protein | 259.52 | 364.88 | -0.49 | 3.96E-02 |
| ENSMUSG00000029103 | GSMG0030378 | Lrpap1 | low density lipoprotein receptor-related protein associated protein 1 | 3532.50 | 3135.81 | 0.17 | 3.94E-01 |
| ENSMUSG00000029097 | GSMG0030386 | Trmt44 | tRNA methyltransferase 44 | 181.39 | 212.14 | -0.22 | 5.10E-01 |
| ENSMUSG00000029096 | GSMG0030388 | Htra3 | HtrA serine peptidase 3 | 2528.54 | 4959.76 | -0.97 | 1.34E-13 |

| | | | | | | | |
|---|---|---|---|---|---|---|---|
| ENSMUSG00000036553 | GSMG0030389 | Sh3tc1 | SH3 domain and tetratricopeptide repeats 1 | 1615.32 | 1802.26 | -0.16 | 5.78E-01 |
| ENSMUSG00000029093 | GSMG0030390 | Sorcs2 | sortilin-related VPS10 domain containing receptor 2 | 69.89 | 95.74 | -0.44 | 5.14E-01 |
| ENSMUSG00000029192 | GSMG0030393 | Tbc1d14 | TBC1 domain family, member 14 | 2075.51 | 2006.49 | 0.05 | 8.65E-01 |
| ENSMUSG00000060708 | GSMG0030396 | Bloc1s4 | biogenesis of lysosomal organelles complex-1, subunit 4, cappuccino | 608.72 | 543.98 | 0.16 | 5.67E-01 |
| ENSMUSG00000055302 | GSMG0030397 | Mrfap1 | Morf4 family associated protein 1 | 7908.69 | 10017.94 | -0.34 | 1.40E-01 |
| ENSMUSG00000029119 | GSMG0030398 | Man2b2 | mannosidase 2, alpha B2 | 536.40 | 451.15 | 0.25 | 4.76E-01 |
| ENSMUSG00000039474 | GSMG0030400 | Wfs1 | Wolfram syndrome 1 homolog (human) | 272.21 | 193.52 | 0.49 | 1.28E-01 |
| NULL | GSMG0030401 | NULL | NULL | 109.93 | 85.85 | 0.36 | 3.76E-01 |
| ENSMUSG00000048450 | GSMG0030404 | Msx1 | msh homeobox 1 | 676.62 | 445.48 | 0.60 | 2.06E-02 |
| ENSMUSG00000029126 | GSMG0030405 | Nsg1 | neuron specific gene family member 1 | 655.66 | 2306.21 | -1.81 | 2.34E-34 |
| ENSMUSG00000029127 | GSMG0030408 | Zbtb49 | zinc finger and BTB domain containing 49 | 316.25 | 315.86 | 0.00 | 9.92E-01 |
| ENSMUSG00000005103 | GSMG0030411 | Wdr1 | WD repeat domain 1 | 7485.44 | 8094.67 | -0.11 | 6.29E-01 |
| ENSMUSG00000051022 | GSMG0030417 | Hs3st1 | heparan sulfate (glucosamine) 3-O-sulfotransferase 1 | 1417.60 | 774.89 | 0.87 | 6.75E-08 |
| ENSMUSG00000029128 | GSMG0030420 | Rab28 | RAB28, member RAS oncogene family | 1701.15 | 1535.98 | 0.15 | 6.03E-01 |
| ENSMUSG00000039753 | GSMG0030430 | Fbxl5 | F-box and leucine-rich repeat protein 5 | 1546.93 | 1440.93 | 0.10 | 6.93E-01 |
| ENSMUSG00000048373 | GSMG0030435 | Fgfbp1 | fibroblast growth factor binding protein 1 | 1069.23 | 654.63 | 0.71 | 6.22E-04 |
| ENSMUSG00000029086 | GSMG0030436 | Prom1 | prominin 1 | 13892.31 | 4974.61 | 1.48 | 4.63E-28 |
| ENSMUSG00000046985 | GSMG0030438 | Tapt1 | transmembrane anterior posterior transformation 1 | 2189.88 | 1849.14 | 0.24 | 1.65E-01 |
| ENSMUSG00000039706 | GSMG0030443 | Ldb2 | LIM domain binding 2 | 182.21 | 421.94 | -1.20 | 8.72E-09 |
| NULL | GSMG0030445 | NULL | NULL | 69.55 | 139.69 | -1.01 | 8.26E-03 |
| NULL | GSMG0030446 | NULL | NULL | 255.83 | 471.95 | -0.89 | 3.09E-05 |
| NULL | GSMG0030451 | NULL | NULL | 248.26 | 496.42 | -1.00 | 2.24E-06 |
| ENSMUSG00000015806 | GSMG0030454 | Qdpr | quinoid dihydropteridine reductase | 949.62 | 888.06 | 0.10 | 7.66E-01 |
| ENSMUSG00000015879 | GSMG0030455 | Fam184b | family with sequence similarity 184, member B | 313.26 | 254.82 | 0.30 | 2.77E-01 |
| ENSMUSG00000015882 | GSMG0030457 | Lcorl | ligand dependent nuclear receptor corepressor-like | 323.10 | 372.49 | -0.20 | 4.48E-01 |
| ENSMUSG00000029088 | GSMG0030464 | Kcnip4 | Kv channel interacting protein 4 | 130.35 | 95.07 | 0.45 | 1.76E-01 |
| NULL | GSMG0030469 | NULL | NULL | 298.40 | 169.00 | 0.82 | 2.95E-03 |
| NULL | GSMG0030470 | NULL | NULL | 198.36 | 126.21 | 0.65 | 7.75E-02 |
| NULL | GSMG0030474 | NULL | NULL | 901.33 | 581.30 | 0.63 | 5.64E-03 |
| NULL | GSMG0030475 | NULL | NULL | 534.70 | 311.55 | 0.78 | 2.68E-03 |
| NULL | GSMG0030476 | NULL | NULL | 298.87 | 140.67 | 1.08 | 1.46E-04 |
| NULL | GSMG0030478 | NULL | NULL | 223.13 | 115.78 | 0.95 | 8.07E-04 |
| NULL | GSMG0030480 | NULL | NULL | 261.14 | 166.02 | 0.65 | 1.39E-02 |

| | | | | | | | |
|---|---|---|---|---|---|---|---|
| NULL | GSMG0030481 | NULL | NULL | 75.97 | 34.96 | 1.12 | 3.79E-03 |
| NULL | GSMG0030483 | NULL | NULL | 273.17 | 187.88 | 0.54 | 4.50E-02 |
| NULL | GSMG0030484 | NULL | NULL | 1109.04 | 717.48 | 0.63 | 1.81E-03 |
| NULL | GSMG0030485 | NULL | NULL | 271.68 | 147.87 | 0.87 | 6.70E-04 |
| NULL | GSMG0030486 | NULL | NULL | 294.05 | 179.44 | 0.71 | 5.67E-03 |
| NULL | GSMG0030488 | NULL | NULL | 222.96 | 119.12 | 0.91 | 1.38E-03 |
| NULL | GSMG0030489 | NULL | NULL | 816.31 | 445.77 | 0.87 | 5.83E-05 |
| NULL | GSMG0030490 | NULL | NULL | 276.18 | 144.71 | 0.93 | 3.50E-04 |
| ENSMUSG00000029090 | GSMG0030492 | Adgra3 | adhesion G protein-coupled receptor A3 | 409.81 | 478.56 | -0.22 | 4.70E-01 |
| ENSMUSG00000029167 | GSMG0030496 | Ppargc1a | peroxisome proliferative activated receptor, gamma, coactivator 1 alpha | 263.71 | 224.55 | 0.23 | 4.85E-01 |
| NULL | GSMG0030504 | NULL | NULL | 301.74 | 145.83 | 1.05 | 8.63E-06 |
| ENSMUSG00000029169 | GSMG0030506 | Dhx15 | DEAH (Asp-Glu-Ala-His) box polypeptide 15 | 2124.71 | 2537.97 | -0.26 | 1.52E-01 |
| ENSMUSG00000039252 | GSMG0030510 | Lgi2 | leucine-rich repeat LGI family, member 2 | 64.88 | 69.36 | -0.11 | 8.83E-01 |
| ENSMUSG00000029173 | GSMG0030513 | Sepsecs | Sep (O-phosphoserine) tRNA:Sec (selenocysteine) tRNA synthase | 177.82 | 156.05 | 0.19 | 6.21E-01 |
| ENSMUSG00000029189 | GSMG0030515 | Sel1l3 | sel-1 suppressor of lin-12-like 3 (C. elegans) | 722.43 | 648.46 | 0.16 | 6.33E-01 |
| ENSMUSG00000037999 | GSMG0030523 | Arap2 | ArfGAP with RhoGAP domain, ankyrin repeat and PH domain 2 | 3331.38 | 2695.70 | 0.31 | 1.40E-01 |
| ENSMUSG00000047881 | GSMG0030527 | Rell1 | RELT-like 1 | 540.02 | 531.51 | 0.02 | 9.42E-01 |
| NULL | GSMG0030532 | NULL | NULL | 98.78 | 186.92 | -0.93 | 1.58E-02 |
| ENSMUSG00000029191 | GSMG0030537 | Rfc1 | replication factor C (activator 1) 1 | 752.67 | 999.59 | -0.41 | 4.19E-02 |
| ENSMUSG00000029201 | GSMG0030541 | Ugdh | UDP-glucose dehydrogenase | 912.40 | 1211.17 | -0.41 | 1.44E-02 |
| NULL | GSMG0030543 | NULL | NULL | 547.67 | 396.25 | 0.46 | 2.49E-02 |
| ENSMUSG00000037822 | GSMG0030544 | Smim14 | small integral membrane protein 14 | 3075.20 | 2231.15 | 0.46 | 1.10E-02 |
| NULL | GSMG0030548 | NULL | NULL | 263.60 | 297.41 | -0.18 | 6.03E-01 |
| ENSMUSG00000029207 | GSMG0030563 | Apbb2 | amyloid beta (A4) precursor protein-binding, family B, member 2 | 4056.26 | 5027.30 | -0.31 | 5.65E-02 |
| NULL | GSMG0030564 | NULL | NULL | 117.99 | 208.53 | -0.82 | 5.86E-03 |
| NULL | GSMG0030565 | NULL | NULL | 215.50 | 213.67 | 0.01 | 9.79E-01 |
| NULL | GSMG0030569 | NULL | NULL | 1028.63 | 513.56 | 1.00 | 1.99E-07 |
| ENSMUSG00000037685 | GSMG0030578 | Atp8a1 | ATPase, aminophospholipid transporter (APLT), class I, type 8A, mem | 1750.18 | 1442.31 | 0.28 | 1.46E-01 |
| NULL | GSMG0030579 | NULL | NULL | 214.67 | 221.25 | -0.05 | 9.18E-01 |
| NULL | GSMG0030582 | NULL | NULL | 384.37 | 266.48 | 0.53 | 1.40E-01 |
| ENSMUSG00000029209 | GSMG0030597 | Gnpda2 | glucosamine-6-phosphate deaminase 2 | 872.80 | 735.19 | 0.25 | 2.90E-01 |
| ENSMUSG00000001260 | GSMG0030598 | Gabrg1 | gamma-aminobutyric acid (GABA) A receptor, subunit gamma 1 | 118.06 | 150.24 | -0.35 | 2.87E-01 |
| ENSMUSG00000000560 | GSMG0030599 | Gabra2 | gamma-aminobutyric acid (GABA) A receptor, subunit alpha 2 | 490.12 | 569.66 | -0.21 | 5.14E-01 |

| | | | | | | | |
|---|---|---|---|---|---|---|---|
| ENSMUSG00000029211 | GSMG0030602 | Gabra4 | gamma-aminobutyric acid (GABA) A receptor, subunit alpha 4 | 282.71 | 186.79 | 0.60 | 1.80E-02 |
| ENSMUSG00000029213 | GSMG0030604 | Commd8 | COMM domain containing 8 | 1494.78 | 1710.51 | -0.20 | 4.07E-01 |
| ENSMUSG00000072889 | GSMG0030607 | Nfxl1 | nuclear transcription factor, X-box binding-like 1 | 138.30 | 155.27 | -0.16 | 6.81E-01 |
| ENSMUSG00000029217 | GSMG0030614 | Tec | tec protein tyrosine kinase | 632.30 | 483.65 | 0.39 | 6.34E-02 |
| ENSMUSG00000070733 | GSMG0030619 | Fryl | FRY like transcription coactivator | 545.84 | 506.58 | 0.11 | 7.00E-01 |
| ENSMUSG00000029153 | GSMG0030623 | Ociad2 | OCIA domain containing 2 | 391.01 | 216.04 | 0.85 | 9.46E-05 |
| ENSMUSG00000029156 | GSMG0030625 | Sgcb | sarcoglycan, beta (dystrophin-associated glycoprotein) | 806.45 | 859.25 | -0.09 | 7.45E-01 |
| ENSMUSG00000054814 | GSMG0030627 | Usp46 | ubiquitin specific peptidase 46 | 580.89 | 799.98 | -0.46 | 2.93E-02 |
| ENSMUSG00000062110 | GSMG0030632 | Scfd2 | Sec1 family domain containing 2 | 134.53 | 130.89 | 0.04 | 9.37E-01 |
| ENSMUSG00000029228 | GSMG0030635 | Lnx1 | ligand of numb-protein X 1 | 434.29 | 151.66 | 1.52 | 2.37E-12 |
| ENSMUSG00000029229 | GSMG0030638 | Chic2 | cysteine-rich hydrophobic domain 2 | 1392.89 | 1594.24 | -0.20 | 3.29E-01 |
| ENSMUSG00000062960 | GSMG0030642 | Kdr | kinase insert domain protein receptor | 13159.66 | 14052.19 | -0.09 | 7.18E-01 |
| ENSMUSG00000029238 | GSMG0030646 | Clock | circadian locomotor output cycles kaput | 890.89 | 749.89 | 0.25 | 2.51E-01 |
| ENSMUSG00000055923 | GSMG0030657 | Aasdh | aminoadipate-semialdehyde dehydrogenase | 379.94 | 372.37 | 0.03 | 9.46E-01 |
| ENSMUSG00000029246 | GSMG0030658 | Ppat | phosphoribosyl pyrophosphate amidotransferase | 278.33 | 427.88 | -0.62 | 3.31E-03 |
| NULL | GSMG0030659 | NULL | NULL | 402.19 | 430.26 | -0.10 | 7.74E-01 |
| ENSMUSG00000059325 | GSMG0030661 | Hopx | HOP homeobox | 683.27 | 818.34 | -0.26 | 1.90E-01 |
| ENSMUSG00000036285 | GSMG0030664 | Noa1 | nitric oxide associated 1 | 244.65 | 274.37 | -0.16 | 6.48E-01 |
| ENSMUSG00000036256 | GSMG0030665 | Igfbp7 | insulin-like growth factor binding protein 7 | 36907.87 | 31965.70 | 0.21 | 2.84E-01 |
| NULL | GSMG0030668 | NULL | NULL | 206.98 | 217.08 | -0.07 | 8.91E-01 |
| ENSMUSG00000029245 | GSMG0030673 | Epha5 | Eph receptor A5 | 489.45 | 819.41 | -0.74 | 1.14E-04 |
| ENSMUSG00000029253 | GSMG0030676 | Cenpc1 | centromere protein C1 | 233.36 | 299.39 | -0.36 | 2.06E-01 |
| ENSMUSG00000035898 | GSMG0030677 | Uba6 | ubiquitin-like modifier activating enzyme 6 | 351.79 | 517.38 | -0.55 | 1.47E-02 |
| ENSMUSG00000044221 | GSMG0030704 | Grsf1 | G-rich RNA sequence binding factor 1 | 751.65 | 757.45 | -0.01 | 9.75E-01 |
| ENSMUSG00000043635 | GSMG0030709 | Adamts3 | a disintegrin-like and metallopeptidase (reprolysin type) with thrombos | 314.45 | 424.15 | -0.43 | 8.84E-02 |
| ENSMUSG00000035505 | GSMG0030711 | Cox18 | cytochrome c oxidase assembly protein 18 | 326.58 | 386.30 | -0.24 | 3.72E-01 |
| ENSMUSG00000029397 | GSMG0030727 | Rchy1 | ring finger and CHY zinc finger domain containing 1 | 402.99 | 437.49 | -0.12 | 7.08E-01 |
| ENSMUSG00000029403 | GSMG0030729 | Cdkl2 | cyclin-dependent kinase-like 2 (CDC2-related kinase) | 249.09 | 439.81 | -0.82 | 3.91E-05 |
| ENSMUSG00000029405 | GSMG0030730 | G3bp2 | GTPase activating protein (SH3 domain) binding protein 2 | 2065.59 | 2671.23 | -0.37 | 6.31E-02 |
| ENSMUSG00000029413 | GSMG0030735 | Naaa | N-acylethanolamine acid amidase | 199.47 | 143.85 | 0.48 | 8.99E-02 |
| ENSMUSG00000029415 | GSMG0030736 | Sdad1 | SDA1 domain containing 1 | 1014.52 | 1414.65 | -0.48 | 3.75E-03 |
| ENSMUSG00000034826 | GSMG0030740 | Nup54 | nucleoporin 54 | 391.76 | 535.45 | -0.45 | 2.65E-02 |
| ENSMUSG00000029426 | GSMG0030741 | Scarb2 | scavenger receptor class B, member 2 | 954.25 | 1101.76 | -0.21 | 3.24E-01 |

| | | | | | | | |
|---|---|---|---|---|---|---|---|
| NULL | GSMG0030748 | NULL | NULL | 1833.59 | 2286.79 | -0.32 | 8.14E-02 |
| ENSMUSG00000063015 | GSMG0030749 | Ccni | cyclin I | 4069.16 | 3963.26 | 0.04 | 9.09E-01 |
| ENSMUSG00000034724 | GSMG0030758 | Cnot6l | CCR4-NOT transcription complex, subunit 6-like | 389.02 | 340.00 | 0.19 | 4.72E-01 |
| ENSMUSG00000055725 | GSMG0030763 | Paqr3 | progestin and adipoQ receptor family member III | 91.93 | 123.44 | -0.43 | 2.63E-01 |
| ENSMUSG00000029338 | GSMG0030767 | Antxr2 | anthrax toxin receptor 2 | 210.44 | 261.74 | -0.31 | 3.39E-01 |
| ENSMUSG00000089809 | GSMG0030777 | Rasgef1b | RasGEF domain family, member 1B | 277.18 | 255.11 | 0.12 | 7.37E-01 |
| ENSMUSG00000000568 | GSMG0030783 | Hnrnpd | heterogeneous nuclear ribonucleoprotein D | 1646.69 | 2344.27 | -0.51 | 1.82E-02 |
| ENSMUSG00000029328 | GSMG0030784 | Hnrnpdl | heterogeneous nuclear ribonucleoprotein D-like | 1360.21 | 1911.42 | -0.49 | 2.37E-03 |
| ENSMUSG00000050640 | GSMG0030785 | Tmem150c | transmembrane protein 150C | 72.67 | 262.35 | -1.86 | 4.93E-16 |
| ENSMUSG00000035325 | GSMG0030786 | Sec31a | Sec31 homolog A (S. cerevisiae) | 3825.08 | 4577.12 | -0.26 | 1.22E-01 |
| ENSMUSG00000097772 | GSMG0030788 | 5430416N02Rik | RIKEN cDNA 5430416N02 gene | 111.17 | 141.30 | -0.35 | 3.80E-01 |
| ENSMUSG00000035310 | GSMG0030789 | Lin54 | lin-54 homolog (C. elegans) | 227.06 | 294.48 | -0.37 | 1.28E-01 |
| ENSMUSG00000029319 | GSMG0030793 | Coq2 | coenzyme Q2 4-hydroxybenzoate polyprenyltransferase | 494.60 | 434.47 | 0.19 | 5.30E-01 |
| ENSMUSG00000035266 | GSMG0030795 | Helq | helicase, POLQ-like | 537.74 | 685.64 | -0.35 | 9.40E-02 |
| ENSMUSG00000035234 | GSMG0030796 | Fam175a | family with sequence similarity 175, member A | 211.92 | 251.97 | -0.25 | 3.93E-01 |
| NULL | GSMG0030808 | NULL | NULL | 77.00 | 53.27 | 0.54 | 2.44E-01 |
| ENSMUSG00000046709 | GSMG0030810 | Mapk10 | mitogen-activated protein kinase 10 | 595.09 | 602.30 | -0.02 | 9.61E-01 |
| NULL | GSMG0030813 | NULL | NULL | 231.65 | 187.89 | 0.30 | 3.32E-01 |
| NULL | GSMG0030816 | NULL | NULL | 559.14 | 344.12 | 0.70 | 6.06E-04 |
| NULL | GSMG0030817 | NULL | NULL | 494.09 | 308.85 | 0.68 | 1.61E-03 |
| ENSMUSG00000029312 | GSMG0030825 | Klhl8 | kelch-like 8 | 523.10 | 530.26 | -0.02 | 9.58E-01 |
| ENSMUSG00000029309 | GSMG0030830 | Sparcl1 | SPARC-like 1 | 49449.30 | 41501.79 | 0.25 | 1.88E-01 |
| ENSMUSG00000029298 | GSMG0030838 | Gbp9 | guanylate-binding protein 9 | 1289.89 | 691.64 | 0.90 | 3.30E-05 |
| ENSMUSG00000079363 | GSMG0030839 | Gbp4 | guanylate binding protein 4 | 503.44 | 416.11 | 0.28 | 5.04E-01 |
| ENSMUSG00000049606 | GSMG0030854 | Zfp644 | zinc finger protein 644 | 2277.59 | 2506.42 | -0.14 | 5.22E-01 |
| ENSMUSG00000029287 | GSMG0030857 | Tgfbr3 | transforming growth factor, beta receptor III | 599.46 | 386.63 | 0.63 | 1.31E-03 |
| ENSMUSG00000029276 | GSMG0030863 | Glmn | glomulin, FKBP associated protein | 401.29 | 439.05 | -0.13 | 7.13E-01 |
| ENSMUSG00000011831 | GSMG0030869 | Evi5 | ecotropic viral integration site 5 | 1237.42 | 1163.55 | 0.09 | 7.55E-01 |
| ENSMUSG00000029270 | GSMG0030873 | Fam69a | family with sequence similarity 69, member A | 2044.89 | 1916.26 | 0.09 | 6.97E-01 |
| ENSMUSG00000063406 | GSMG0030876 | Tmed5 | transmembrane emp24 protein transport domain containing 5 | 1340.05 | 1502.25 | -0.16 | 5.26E-01 |
| ENSMUSG00000050856 | GSMG0030879 | Atp5k | ATP synthase, H+ transporting, mitochondrial F1F0 complex, subunit E | 1145.52 | 1258.10 | -0.14 | 7.59E-01 |
| ENSMUSG00000033615 | GSMG0030881 | Cplx1 | complexin 1 | 1618.01 | 1146.63 | 0.50 | 7.89E-03 |
| ENSMUSG00000062234 | GSMG0030884 | Gak | cyclin G associated kinase | 1031.76 | 1076.03 | -0.06 | 8.57E-01 |

| | | | | | | | |
|---|---|---|---|---|---|---|---|
| ENSMUSG00000004815 | GSMG0030886 | Dgkq | diacylglycerol kinase, theta | 478.13 | 472.97 | 0.02 | 9.75E-01 |
| ENSMUSG00000033467 | GSMG0030891 | Crlf2 | cytokine receptor-like factor 2 | 154.54 | 172.55 | -0.17 | 7.10E-01 |
| NULL | GSMG0030897 | NULL | NULL | 87.05 | 40.39 | 1.11 | 5.30E-03 |
| ENSMUSG00000033434 | GSMG0030898 | Gtpbp6 | GTP binding protein 6 (putative) | 628.71 | 584.29 | 0.11 | 7.03E-01 |
| ENSMUSG00000029500 | GSMG0030901 | Pgam5 | phosphoglycerate mutase family member 5 | 1227.79 | 1649.84 | -0.43 | 6.64E-03 |
| ENSMUSG00000029503 | GSMG0030903 | P2rx2 | purinergic receptor P2X, ligand-gated ion channel, 2 | 5.41 | 226.08 | -5.39 | 4.10E-34 |
| ENSMUSG00000043323 | GSMG0030904 | Fbrsl1 | fibrosin-like 1 | 300.96 | 303.02 | -0.01 | 9.81E-01 |
| ENSMUSG00000033294 | GSMG0030908 | Noc4l | NOC4 like | 754.93 | 827.97 | -0.13 | 6.48E-01 |
| ENSMUSG00000029505 | GSMG0030909 | Ep400 | E1A binding protein p400 | 1290.09 | 1745.32 | -0.43 | 5.79E-02 |
| ENSMUSG00000029507 | GSMG0030911 | Pus1 | pseudouridine synthase 1 | 295.18 | 373.49 | -0.34 | 1.62E-01 |
| ENSMUSG00000029512 | GSMG0030912 | Ulk1 | unc-51 like kinase 1 | 1703.21 | 1736.13 | -0.03 | 9.48E-01 |
| ENSMUSG00000043510 | GSMG0030913 | Hscb | HscB iron-sulfur cluster co-chaperone | 141.30 | 146.22 | -0.05 | 9.20E-01 |
| ENSMUSG00000097767 | GSMG0030919 | Miat | myocardial infarction associated transcript (non-protein coding) | 15054.87 | 15657.32 | -0.06 | 8.69E-01 |
| ENSMUSG00000029346 | GSMG0030921 | Srrd | SRR1 domain containing | 470.32 | 444.99 | 0.08 | 7.99E-01 |
| ENSMUSG00000029348 | GSMG0030924 | Asphd2 | aspartate beta-hydroxylase domain containing 2 | 1590.04 | 1781.42 | -0.16 | 5.15E-01 |
| ENSMUSG00000058153 | GSMG0030925 | Sez6l | seizure related 6 homolog like | 380.80 | 434.91 | -0.19 | 6.02E-01 |
| ENSMUSG00000042249 | GSMG0030930 | Adrbk2 | adrenergic receptor kinase, beta 2 | 324.78 | 379.79 | -0.23 | 4.68E-01 |
| ENSMUSG00000051339 | GSMG0030933 | 2900026A02Rik | RIKEN cDNA 2900026A02 gene | 619.23 | 612.81 | 0.01 | 9.75E-01 |
| NULL | GSMG0030935 | NULL | NULL | 258.27 | 133.86 | 0.95 | 3.29E-04 |
| ENSMUSG00000042216 | GSMG0030936 | Sgsm1 | small G protein signaling modulator 1 | 392.93 | 466.53 | -0.25 | 4.53E-01 |
| ENSMUSG00000018974 | GSMG0030940 | Sart3 | squamous cell carcinoma antigen recognized by T cells 3 | 1191.68 | 1465.51 | -0.30 | 1.74E-01 |
| ENSMUSG00000054675 | GSMG0030941 | Tmem119 | transmembrane protein 119 | 746.45 | 271.95 | 1.46 | 1.63E-11 |
| ENSMUSG00000048163 | GSMG0030942 | Selplg | selectin, platelet (p-selectin) ligand | 612.57 | 147.78 | 2.05 | 1.82E-19 |
| ENSMUSG00000004530 | GSMG0030943 | Coro1c | coronin, actin binding protein 1C | 1297.77 | 2038.91 | -0.65 | 6.69E-06 |
| ENSMUSG00000042121 | GSMG0030945 | Ssh1 | slingshot homolog 1 (Drosophila) | 288.50 | 367.08 | -0.35 | 1.93E-01 |
| ENSMUSG00000042078 | GSMG0030946 | Svop | SV2 related protein | 872.67 | 841.00 | 0.05 | 8.87E-01 |
| ENSMUSG00000044339 | GSMG0030947 | Alkbh2 | alkB homolog 2, alpha-ketoglutarate-dependent dioxygenase | 149.83 | 125.36 | 0.26 | 5.32E-01 |
| ENSMUSG00000001098 | GSMG0030950 | Kctd10 | potassium channel tetramerisation domain containing 10 | 2849.45 | 2373.70 | 0.26 | 1.58E-01 |
| NULL | GSMG0030951 | NULL | NULL | 191.89 | 158.42 | 0.28 | 4.43E-01 |
| NULL | GSMG0030952 | NULL | NULL | 133.86 | 130.41 | 0.03 | 9.46E-01 |
| ENSMUSG00000029575 | GSMG0030953 | Mmab | methylmalonic aciduria (cobalamin deficiency) cblB type homolog (hur | 189.74 | 242.73 | -0.36 | 2.16E-01 |
| ENSMUSG00000014158 | GSMG0030956 | Trpv4 | transient receptor potential cation channel, subfamily V, member 4 | 305.98 | 81.23 | 1.91 | 4.42E-11 |
| ENSMUSG00000011884 | GSMG0030957 | Gltp | glycolipid transfer protein | 2595.84 | 1749.34 | 0.57 | 1.32E-03 |

| | | | | | | | |
|---|---|---|---|---|---|---|---|
| ENSMUSG00000041890 | GSMG0030958 | Git2 | G protein-coupled receptor kinase-interactor 2 | 1569.01 | 1705.67 | -0.12 | 6.05E-01 |
| ENSMUSG00000072694 | GSMG0030960 | 1500011B03Rik | RIKEN cDNA 1500011B03 gene | 1418.80 | 1392.33 | 0.03 | 9.31E-01 |
| ENSMUSG00000092486 | GSMG0030961 | 2610524H06Rik | RIKEN cDNA 2610524H06 gene | 732.75 | 949.26 | -0.37 | 8.54E-02 |
| NULL | GSMG0030965 | NULL | NULL | 179.67 | 149.45 | 0.26 | 4.18E-01 |
| ENSMUSG00000029545 | GSMG0030966 | Acads | acyl-Coenzyme A dehydrogenase, short chain | 463.65 | 623.04 | -0.43 | 7.11E-02 |
| ENSMUSG00000046562 | GSMG0030967 | Unc119b | unc-119 lipid binding chaperone B | 3293.32 | 2533.12 | 0.38 | 1.37E-02 |
| ENSMUSG00000048578 | GSMG0030968 | Mlec | malectin | 2917.95 | 2987.79 | -0.03 | 9.21E-01 |
| ENSMUSG00000029544 | GSMG0030970 | Cabp1 | calcium binding protein 1 | 479.81 | 284.41 | 0.75 | 1.44E-02 |
| ENSMUSG00000041740 | GSMG0030971 | Rnf10 | ring finger protein 10 | 5373.32 | 6603.37 | -0.30 | 9.86E-02 |
| ENSMUSG00000009013 | GSMG0030972 | Dynll1 | dynein light chain LC8-type 1 | 6057.83 | 6073.54 | 0.00 | 9.91E-01 |
| ENSMUSG00000029536 | GSMG0030973 | Gatc | glutamyl-tRNA(Gln) amidotransferase, subunit C | 1191.39 | 1466.78 | -0.30 | 9.78E-02 |
| ENSMUSG00000041697 | GSMG0030974 | Cox6a1 | cytochrome c oxidase subunit VIa polypeptide 1 | 6632.67 | 7381.25 | -0.15 | 5.00E-01 |
| ENSMUSG00000029524 | GSMG0030979 | Sirt4 | sirtuin 4 | 502.34 | 421.83 | 0.25 | 3.19E-01 |
| ENSMUSG00000107121 | GSMG0030981 | 1110006O24Rik | RIKEN cDNA 1110006O24 gene | 117.40 | 102.89 | 0.19 | 6.79E-01 |
| ENSMUSG00000041609 | GSMG0030982 | Ccdc64 | coiled-coil domain containing 64 | 292.91 | 259.54 | 0.17 | 7.49E-01 |
| NULL | GSMG0030983 | NULL | NULL | 919.22 | 502.38 | 0.87 | 4.70E-04 |
| ENSMUSG00000029513 | GSMG0030984 | Prkab1 | protein kinase, AMP-activated, beta 1 non-catalytic subunit | 656.47 | 675.12 | -0.04 | 9.00E-01 |
| ENSMUSG00000041548 | GSMG0030989 | Hspb8 | heat shock protein 8 | 552.89 | 318.97 | 0.80 | 1.19E-03 |
| ENSMUSG00000063919 | GSMG0030990 | Srrm4 | serine/arginine repetitive matrix 4 | 332.19 | 535.81 | -0.69 | 2.99E-03 |
| NULL | GSMG0030991 | NULL | NULL | 133.19 | 128.22 | 0.06 | 9.12E-01 |
| NULL | GSMG0030992 | NULL | NULL | 346.83 | 454.35 | -0.39 | 7.66E-02 |
| ENSMUSG00000066900 | GSMG0030995 | Suds3 | suppressor of defective silencing 3 homolog (S. cerevisiae) | 855.12 | 916.92 | -0.10 | 7.14E-01 |
| NULL | GSMG0030998 | NULL | NULL | 156.04 | 125.21 | 0.31 | 4.67E-01 |
| ENSMUSG00000029363 | GSMG0030999 | Rfc5 | replication factor C (activator 1) 5 | 296.35 | 448.59 | -0.60 | 3.37E-03 |
| ENSMUSG00000032867 | GSMG0031002 | Fbxw8 | F-box and WD-40 domain protein 8 | 1284.65 | 1288.90 | -0.01 | 9.89E-01 |
| ENSMUSG00000032850 | GSMG0031003 | Rnft2 | ring finger protein, transmembrane 2 | 210.98 | 297.02 | -0.49 | 1.15E-01 |
| NULL | GSMG0031004 | NULL | NULL | 266.07 | 358.01 | -0.43 | 1.37E-01 |
| NULL | GSMG0031009 | AW549542 | expressed sequence AW549542 | 174.05 | 86.04 | 1.02 | 3.09E-04 |
| ENSMUSG00000029598 | GSMG0031018 | Plbd2 | phospholipase B domain containing 2 | 4368.86 | 3175.12 | 0.46 | 1.73E-02 |
| ENSMUSG00000032741 | GSMG0031019 | Tpcn1 | two pore channel 1 | 664.10 | 576.91 | 0.20 | 4.15E-01 |
| ENSMUSG00000029600 | GSMG0031021 | Rita1 | RBPJ interacting and tubulin associated 1 | 137.50 | 148.05 | -0.11 | 8.13E-01 |
| NULL | GSMG0031022 | NULL | NULL | 265.71 | 305.73 | -0.21 | 5.09E-01 |
| ENSMUSG00000029603 | GSMG0031023 | Dtx1 | deltex 1, E3 ubiquitin ligase | 234.75 | 425.32 | -0.86 | 9.37E-04 |

| | | | | | | | |
|---|---|---|---|---|---|---|---|
| ENSMUSG00000029608 | GSMG0031030 | Rph3a | rabphilin 3A | 358.16 | 282.60 | 0.34 | 2.38E-01 |
| NULL | GSMG0031031 | NULL | NULL | 338.64 | 185.66 | 0.87 | 4.30E-05 |
| ENSMUSG00000043733 | GSMG0031032 | Ptpn11 | protein tyrosine phosphatase, non-receptor type 11 | 1163.33 | 1364.76 | -0.23 | 3.40E-01 |
| ENSMUSG00000042726 | GSMG0031034 | Trafd1 | TRAF type zinc finger domain containing 1 | 1666.03 | 2182.64 | -0.39 | 3.26E-02 |
| ENSMUSG00000029616 | GSMG0031035 | Erp29 | endoplasmic reticulum protein 29 | 2571.60 | 2898.19 | -0.17 | 3.66E-01 |
| ENSMUSG00000029454 | GSMG0031037 | Mapkapk5 | MAP kinase-activated protein kinase 5 | 1055.70 | 1008.76 | 0.07 | 8.08E-01 |
| ENSMUSG00000029455 | GSMG0031038 | Aldh2 | aldehyde dehydrogenase 2, mitochondrial | 3690.50 | 2426.35 | 0.60 | 2.26E-05 |
| ENSMUSG00000042647 | GSMG0031039 | Acad12 | acyl-Coenzyme A dehydrogenase family, member 12 | 210.40 | 84.70 | 1.32 | 6.41E-07 |
| ENSMUSG00000042594 | GSMG0031042 | Sh2b3 | SH2B adaptor protein 3 | 1589.83 | 1429.27 | 0.15 | 5.87E-01 |
| NULL | GSMG0031044 | NULL | NULL | 245.81 | 450.52 | -0.88 | 1.98E-01 |
| ENSMUSG00000042589 | GSMG0031045 | Cux2 | cut-like homeobox 2 | 178.04 | 222.83 | -0.32 | 3.61E-01 |
| ENSMUSG00000038593 | GSMG0031048 | Tctn1 | tectonic family member 1 | 149.08 | 125.29 | 0.26 | 5.30E-01 |
| ENSMUSG00000038569 | GSMG0031049 | Rad9b | RAD9 checkpoint clamp component B | 143.26 | 147.99 | -0.05 | 9.21E-01 |
| ENSMUSG00000029463 | GSMG0031051 | Fam216a | family with sequence similarity 216, member A | 676.47 | 757.82 | -0.16 | 5.40E-01 |
| ENSMUSG00000029467 | GSMG0031053 | Atp2a2 | ATPase, Ca++ transporting, cardiac muscle, slow twitch 2 | 6261.53 | 5011.18 | 0.32 | 2.20E-01 |
| ENSMUSG00000029469 | GSMG0031054 | Ift81 | intraflagellar transport 81 | 370.95 | 437.71 | -0.24 | 3.18E-01 |
| ENSMUSG00000029471 | GSMG0031056 | Camkk2 | calcium/calmodulin-dependent protein kinase kinase 2, beta | 1213.34 | 1020.59 | 0.25 | 2.93E-01 |
| ENSMUSG00000029472 | GSMG0031057 | Anapc5 | anaphase-promoting complex subunit 5 | 10156.95 | 12213.41 | -0.27 | 8.74E-02 |
| ENSMUSG00000029475 | GSMG0031058 | Kdm2b | lysine (K)-specific demethylase 2B | 479.40 | 971.30 | -1.02 | 3.49E-07 |
| ENSMUSG00000029449 | GSMG0031061 | Rhof | ras homolog family member F (in filopodia) | 340.36 | 473.06 | -0.47 | 3.10E-02 |
| ENSMUSG00000029433 | GSMG0031067 | Diablo | diablo, IAP-binding mitochondrial protein | 1752.88 | 1663.07 | 0.08 | 7.69E-01 |
| ENSMUSG00000029434 | GSMG0031068 | Vps33a | VPS33A CORVET/HOPS core subunit | 930.13 | 1034.87 | -0.15 | 5.05E-01 |
| ENSMUSG00000049550 | GSMG0031070 | Clip1 | CAP-GLY domain containing linker protein 1 | 3716.93 | 4393.03 | -0.24 | 2.08E-01 |
| ENSMUSG00000029427 | GSMG0031072 | Zcchc8 | zinc finger, CCHC domain containing 8 | 537.05 | 639.30 | -0.25 | 3.38E-01 |
| ENSMUSG00000029422 | GSMG0031073 | Rsrc2 | arginine/serine-rich coiled-coil 2 | 1283.98 | 1420.44 | -0.15 | 6.20E-01 |
| ENSMUSG00000029408 | GSMG0031078 | Abcb9 | ATP-binding cassette, sub-family B (MDR/TAP), member 9 | 736.48 | 554.75 | 0.41 | 3.95E-02 |
| ENSMUSG00000029406 | GSMG0031079 | Pitpnm2 | phosphatidylinositol transfer protein, membrane-associated 2 | 550.36 | 457.89 | 0.26 | 7.85E-01 |
| ENSMUSG00000038126 | GSMG0031082 | Mphosph9 | M-phase phosphoprotein 9 | 304.06 | 332.13 | -0.13 | 7.51E-01 |
| ENSMUSG00000038095 | GSMG0031087 | Sbno1 | strawberry notch homolog 1 (Drosophila) | 1735.44 | 1731.78 | 0.00 | 9.92E-01 |
| ENSMUSG00000029401 | GSMG0031088 | Rilpl2 | Rab interacting lysosomal protein-like 2 | 764.65 | 609.33 | 0.33 | 9.33E-02 |
| ENSMUSG00000029392 | GSMG0031090 | Rilpl1 | Rab interacting lysosomal protein-like 1 | 771.00 | 665.91 | 0.21 | 3.74E-01 |
| NULL | GSMG0031091 | NULL | NULL | 151.17 | 166.50 | -0.13 | 7.77E-01 |
| ENSMUSG00000029388 | GSMG0031092 | Eif2b1 | eukaryotic translation initiation factor 2B, subunit 1 (alpha) | 932.19 | 1106.00 | -0.25 | 2.44E-01 |

| | | | | | | | |
|---|---|---|---|---|---|---|---|
| NULL | GSMG0031093 | NULL | NULL | 147.62 | 139.09 | 0.08 | 8.77E-01 |
| ENSMUSG00000037979 | GSMG0031094 | Ccdc92 | coiled-coil domain containing 92 | 408.02 | 428.35 | -0.07 | 8.28E-01 |
| ENSMUSG00000029478 | GSMG0031097 | Ncor2 | nuclear receptor co-repressor 2 | 885.69 | 666.13 | 0.41 | 6.45E-01 |
| ENSMUSG00000037936 | GSMG0031099 | Scarb1 | scavenger receptor class B, member 1 | 7639.81 | 9104.48 | -0.25 | 1.19E-01 |
| ENSMUSG00000029480 | GSMG0031103 | Dhx37 | DEAH (Asp-Glu-Ala-His) box polypeptide 37 | 273.61 | 375.89 | -0.46 | 6.60E-02 |
| ENSMUSG00000029416 | GSMG0031105 | Slc15a4 | solute carrier family 15, member 4 | 976.30 | 873.66 | 0.16 | 4.68E-01 |
| ENSMUSG00000034310 | GSMG0031106 | Tmem132d | transmembrane protein 132D | 880.23 | 157.67 | 2.48 | 3.44E-37 |
| ENSMUSG00000029420 | GSMG0031113 | Rimbp2 | RIMS binding protein 2 | 574.12 | 346.28 | 0.73 | 8.52E-03 |
| ENSMUSG00000029428 | GSMG0031114 | Stx2 | syntaxin 2 | 481.69 | 469.36 | 0.04 | 9.14E-01 |
| ENSMUSG00000029446 | GSMG0031119 | Psph | phosphoserine phosphatase | 430.21 | 487.22 | -0.18 | 4.94E-01 |
| NULL | GSMG0031120 | NULL | NULL | 307.24 | 428.98 | -0.48 | 1.34E-01 |
| ENSMUSG00000025537 | GSMG0031121 | Phkg1 | phosphorylase kinase gamma 1 | 114.68 | 31.78 | 1.86 | 3.72E-08 |
| ENSMUSG00000025534 | GSMG0031124 | Gusb | glucuronidase, beta | 1044.86 | 1132.62 | -0.11 | 6.75E-01 |
| ENSMUSG00000025533 | GSMG0031125 | Asl | argininosuccinate lyase | 956.10 | 1155.76 | -0.27 | 1.61E-01 |
| NULL | GSMG0031128 | NULL | NULL | 168.07 | 150.10 | 0.16 | 6.83E-01 |
| ENSMUSG00000025337 | GSMG0031129 | Sbds | Shwachman-Bodian-Diamond syndrome homolog (human) | 1743.32 | 1435.69 | 0.28 | 1.87E-01 |
| ENSMUSG00000034040 | GSMG0031132 | Wbscr17 | Williams-Beuren syndrome chromosome region 17 homolog (human) | 181.69 | 140.14 | 0.37 | 2.93E-01 |
| NULL | GSMG0031138 | NULL | NULL | 74.66 | 80.97 | -0.13 | 8.38E-01 |
| NULL | GSMG0031139 | NULL | NULL | 568.33 | 869.44 | -0.62 | 5.02E-03 |
| NULL | GSMG0031140 | NULL | NULL | 236.12 | 340.98 | -0.53 | 6.95E-02 |
| NULL | GSMG0031142 | NULL | NULL | 154.48 | 153.44 | 0.01 | 9.86E-01 |
| NULL | GSMG0031144 | NULL | NULL | 358.83 | 318.94 | 0.17 | 5.57E-01 |
| NULL | GSMG0031146 | NULL | NULL | 786.71 | 810.20 | -0.04 | 8.97E-01 |
| NULL | GSMG0031147 | NULL | NULL | 282.40 | 269.15 | 0.07 | 8.67E-01 |
| NULL | GSMG0031148 | NULL | NULL | 803.36 | 989.85 | -0.30 | 1.38E-01 |
| NULL | GSMG0031149 | NULL | NULL | 126.56 | 134.92 | -0.09 | 8.47E-01 |
| NULL | GSMG0031152 | NULL | NULL | 292.15 | 369.64 | -0.34 | 2.34E-01 |
| NULL | GSMG0031153 | NULL | NULL | 1931.72 | 2798.06 | -0.54 | 1.25E-03 |
| NULL | GSMG0031154 | NULL | NULL | 170.23 | 214.42 | -0.33 | 2.80E-01 |
| NULL | GSMG0031155 | NULL | NULL | 570.12 | 727.11 | -0.35 | 1.36E-01 |
| ENSMUSG00000029673 | GSMG0031156 | Auts2 | autism susceptibility candidate 2 | 437.13 | 492.72 | -0.17 | 6.10E-01 |
| NULL | GSMG0031157 | NULL | NULL | 357.66 | 401.85 | -0.17 | 5.74E-01 |
| NULL | GSMG0031160 | NULL | NULL | 458.66 | 507.65 | -0.15 | 6.15E-01 |

| | | | | | | | |
|---|---|---|---|---|---|---|---|
| NULL | GSMG0031161 | NULL | NULL | 337.39 | 354.37 | -0.07 | 8.71E-01 |
| NULL | GSMG0031164 | NULL | NULL | 109.38 | 114.07 | -0.05 | 9.28E-01 |
| NULL | GSMG0031165 | NULL | NULL | 570.85 | 626.57 | -0.13 | 6.10E-01 |
| NULL | GSMG0031166 | NULL | NULL | 102.17 | 178.36 | -0.81 | 1.46E-02 |
| NULL | GSMG0031167 | NULL | NULL | 187.28 | 58.66 | 1.68 | 8.47E-08 |
| ENSMUSG00000061979 | GSMG0031171 | Wbscr16 | Williams-Beuren syndrome chromosome region 16 homolog (human) | 300.31 | 417.92 | -0.48 | 1.88E-02 |
| ENSMUSG00000060261 | GSMG0031173 | Gtf2i | general transcription factor II I | 3913.77 | 4883.52 | -0.32 | 2.36E-01 |
| NULL | GSMG0031174 | NULL | NULL | 200.98 | 154.55 | 0.38 | 2.13E-01 |
| NULL | GSMG0031175 | NULL | NULL | 1097.47 | 900.83 | 0.28 | 2.76E-01 |
| ENSMUSG00000023079 | GSMG0031176 | Gtf2ird1 | general transcription factor II I repeat domain-containing 1 | 1681.18 | 1874.86 | -0.16 | 5.64E-01 |
| ENSMUSG00000063146 | GSMG0031178 | Clip2 | CAP-GLY domain containing linker protein 2 | 384.61 | 633.45 | -0.72 | 3.25E-03 |
| ENSMUSG00000040731 | GSMG0031182 | Eif4h | eukaryotic translation initiation factor 4H | 10378.82 | 11163.39 | -0.11 | 6.21E-01 |
| ENSMUSG00000029674 | GSMG0031183 | Limk1 | LIM-domain containing, protein kinase | 599.03 | 677.54 | -0.18 | 5.27E-01 |
| ENSMUSG00000029675 | GSMG0031184 | Eln | elastin | 8565.24 | 2335.19 | 1.87 | 2.49E-03 |
| ENSMUSG00000005378 | GSMG0031193 | Wbscr22 | Williams Beuren syndrome chromosome region 22 | 825.22 | 898.74 | -0.12 | 6.48E-01 |
| ENSMUSG00000043614 | GSMG0031194 | Vps37d | vacuolar protein sorting 37D | 185.07 | 277.17 | -0.59 | 4.45E-02 |
| ENSMUSG00000049551 | GSMG0031196 | Fzd9 | frizzled class receptor 9 | 145.28 | 330.50 | -1.19 | 1.42E-08 |
| ENSMUSG00000053293 | GSMG0031198 | Pom121 | nuclear pore membrane protein 121 | 938.46 | 1135.88 | -0.28 | 1.90E-01 |
| ENSMUSG00000039959 | GSMG0031199 | Hip1 | huntingtin interacting protein 1 | 4147.26 | 3574.07 | 0.21 | 4.85E-01 |
| ENSMUSG00000039886 | GSMG0031206 | Tmem120a | transmembrane protein 120A | 545.16 | 932.39 | -0.77 | 1.71E-06 |
| ENSMUSG00000051391 | GSMG0031209 | Ywhag | tyrosine 3-monooxygenase/tryptophan 5-monooxygenase activation pro | 2228.84 | 3879.18 | -0.80 | 9.67E-08 |
| ENSMUSG00000029703 | GSMG0031212 | Lrwd1 | leucine-rich repeats and WD repeat domain containing 1 | 573.82 | 649.40 | -0.18 | 5.22E-01 |
| ENSMUSG00000039747 | GSMG0031213 | Orai2 | ORAI calcium release-activated calcium modulator 2 | 826.02 | 1234.88 | -0.58 | 9.50E-04 |
| ENSMUSG00000039737 | GSMG0031214 | Prkrip1 | Prkr interacting protein 1 (IL11 inducible) | 663.35 | 709.67 | -0.10 | 7.51E-01 |
| ENSMUSG00000005057 | GSMG0031215 | Sh2b2 | SH2B adaptor protein 2 | 120.70 | 214.87 | -0.83 | 5.87E-03 |
| ENSMUSG00000029705 | GSMG0031218 | Cux1 | cut-like homeobox 1 | 1249.74 | 1504.87 | -0.27 | 1.79E-01 |
| NULL | GSMG0031220 | NULL | NULL | 164.26 | 154.03 | 0.09 | 8.32E-01 |
| ENSMUSG00000004415 | GSMG0031225 | Col26a1 | collagen, type XXVI, alpha 1 | 360.75 | 340.60 | 0.08 | 8.24E-01 |
| ENSMUSG00000059518 | GSMG0031226 | Znhit1 | zinc finger, HIT domain containing 1 | 722.20 | 883.96 | -0.29 | 2.31E-01 |
| ENSMUSG00000004849 | GSMG0031227 | Ap1s1 | adaptor protein complex AP-1, sigma 1 | 2095.21 | 2987.00 | -0.51 | 1.73E-03 |
| ENSMUSG00000037364 | GSMG0031232 | Srrt | serrate RNA effector molecule homolog (Arabidopsis) | 1707.10 | 1720.12 | -0.01 | 9.71E-01 |
| ENSMUSG00000023348 | GSMG0031233 | Trip6 | thyroid hormone receptor interactor 6 | 585.78 | 620.88 | -0.08 | 8.23E-01 |
| ENSMUSG00000037344 | GSMG0031234 | Slc12a9 | solute carrier family 12 (potassium/chloride transporters), member 9 | 591.25 | 581.50 | 0.02 | 9.48E-01 |

| | | | | | | | |
|---|---|---|---|---|---|---|---|
| ENSMUSG00000029715 | GSMG0031238 | Pop7 | processing of precursor 7, ribonuclease P family, (S. cerevisiae) | 436.17 | 613.08 | -0.49 | 1.72E-02 |
| ENSMUSG00000037221 | GSMG0031239 | Mospd3 | motile sperm domain containing 3 | 657.22 | 621.32 | 0.08 | 7.93E-01 |
| ENSMUSG00000029718 | GSMG0031240 | Pcolce | procollagen C-endopeptidase enhancer protein | 1753.63 | 1272.50 | 0.46 | 1.21E-02 |
| NULL | GSMG0031242 | NULL | NULL | 857.35 | 682.71 | 0.33 | 2.16E-01 |
| ENSMUSG00000029722 | GSMG0031244 | Agfg2 | ArfGAP with FG repeats 2 | 345.33 | 444.17 | -0.37 | 1.35E-01 |
| ENSMUSG00000045348 | GSMG0031247 | Nyap1 | neuronal tyrosine-phosphorylated phosphoinositide 3-kinase adaptor 1 | 175.09 | 350.34 | -1.00 | 2.14E-05 |
| ENSMUSG00000029726 | GSMG0031248 | Mepce | methylphosphate capping enzyme | 359.23 | 283.28 | 0.34 | 2.53E-01 |
| ENSMUSG00000029730 | GSMG0031256 | Mcm7 | minichromosome maintenance complex component 7 | 852.38 | 1675.42 | -0.97 | 7.33E-09 |
| ENSMUSG00000036980 | GSMG0031257 | Taf6 | TATA-box binding protein associated factor 6 | 1216.68 | 1164.96 | 0.06 | 8.31E-01 |
| ENSMUSG00000036948 | GSMG0031260 | BC037034 | cDNA sequence BC037034 | 554.89 | 486.21 | 0.19 | 5.76E-01 |
| ENSMUSG00000029510 | GSMG0031262 | Gpc2 | glypican 2 (cerebroglycan) | 75.61 | 417.47 | -2.46 | 2.04E-33 |
| ENSMUSG00000058291 | GSMG0031264 | Zfp68 | zinc finger protein 68 | 1946.34 | 1840.76 | 0.08 | 7.40E-01 |
| ENSMUSG00000025856 | GSMG0031265 | Pdgfa | platelet derived growth factor, alpha | 700.39 | 478.55 | 0.55 | 1.76E-02 |
| ENSMUSG00000025855 | GSMG0031266 | Prkar1b | protein kinase, cAMP dependent regulatory, type I beta | 455.61 | 668.13 | -0.55 | 8.04E-03 |
| ENSMUSG00000056413 | GSMG0031271 | Adap1 | ArfGAP with dual PH domains 1 | 805.80 | 738.52 | 0.13 | 7.00E-01 |
| ENSMUSG00000045438 | GSMG0031273 | Cox19 | cytochrome c oxidase assembly protein 19 | 645.41 | 698.84 | -0.12 | 6.72E-01 |
| ENSMUSG00000053553 | GSMG0031274 | 3110082I17Rik | RIKEN cDNA 3110082I17 gene | 347.95 | 511.06 | -0.56 | 3.30E-03 |
| ENSMUSG00000053581 | GSMG0031278 | Zfand2a | zinc finger, AN1-type domain 2A | 404.53 | 351.34 | 0.20 | 4.51E-01 |
| ENSMUSG00000036718 | GSMG0031279 | Micall2 | MICAL-like 2 | 272.23 | 392.47 | -0.53 | 2.82E-02 |
| ENSMUSG00000029547 | GSMG0031280 | Ints1 | integrator complex subunit 1 | 519.74 | 531.17 | -0.03 | 9.37E-01 |
| ENSMUSG00000029551 | GSMG0031282 | Psmg3 | proteasome (prosome, macropain) assembly chaperone 3 | 573.74 | 668.22 | -0.22 | 3.21E-01 |
| ENSMUSG00000029554 | GSMG0031283 | Mad1l1 | MAD1 mitotic arrest deficient 1-like 1 | 250.45 | 340.74 | -0.44 | 4.92E-02 |
| ENSMUSG00000029557 | GSMG0031287 | Mrm2 | mitochondrial rRNA methyltransferase 2 | 235.30 | 191.95 | 0.30 | 3.28E-01 |
| ENSMUSG00000029560 | GSMG0031288 | Snx8 | sorting nexin 8 | 224.12 | 306.19 | -0.45 | 5.29E-02 |
| ENSMUSG00000036565 | GSMG0031292 | Ttyh3 | tweety family member 3 | 1170.25 | 1871.72 | -0.68 | 3.11E-01 |
| ENSMUSG00000036555 | GSMG0031293 | Iqce | IQ motif containing E | 285.22 | 297.64 | -0.07 | 8.74E-01 |
| ENSMUSG00000000149 | GSMG0031296 | Gna12 | guanine nucleotide binding protein, alpha 12 | 893.34 | 757.13 | 0.24 | 5.79E-01 |
| ENSMUSG00000029576 | GSMG0031301 | Radil | Ras association and DIL domains | 993.89 | 1124.76 | -0.18 | 5.30E-01 |
| ENSMUSG00000039533 | GSMG0031304 | Mmd2 | monocyte to macrophage differentiation-associated 2 | 3812.14 | 3760.33 | 0.02 | 9.46E-01 |
| ENSMUSG00000039477 | GSMG0031308 | Tnrc18 | trinucleotide repeat containing 18 | 193.59 | 270.68 | -0.49 | 2.21E-01 |
| ENSMUSG00000066640 | GSMG0031311 | Fbxl18 | F-box and leucine-rich repeat protein 18 | 232.41 | 323.65 | -0.48 | 3.70E-02 |
| ENSMUSG00000029580 | GSMG0031313 | Actb | actin, beta | 96587.76 | 123032.93 | -0.35 | 6.90E-02 |
| ENSMUSG00000045078 | GSMG0031314 | Rnf216 | ring finger protein 216 | 1195.80 | 1301.66 | -0.12 | 6.23E-01 |

| | | | | | | | |
|---|---|---|---|---|---|---|---|
| ENSMUSG00000061898 | GSMG0031321 | Rbak | RB-associated KRAB zinc finger | 988.08 | 1101.01 | -0.16 | 5.59E-01 |
| ENSMUSG00000046658 | GSMG0031323 | Zfp316 | zinc finger protein 316 | 76.21 | 68.01 | 0.16 | 8.09E-01 |
| ENSMUSG00000039244 | GSMG0031324 | E130309D02Rik | RIKEN cDNA E130309D02 gene | 446.06 | 697.25 | -0.65 | 1.99E-04 |
| ENSMUSG00000001844 | GSMG0031325 | Zdhhc4 | zinc finger, DHHC domain containing 4 | 1276.81 | 1319.81 | -0.05 | 8.63E-01 |
| ENSMUSG00000001847 | GSMG0031329 | Rac1 | RAS-related C3 botulinum substrate 1 | 14836.18 | 15797.63 | -0.09 | 7.14E-01 |
| NULL | GSMG0031330 | NULL | NULL | 280.08 | 246.22 | 0.19 | 5.53E-01 |
| NULL | GSMG0031331 | NULL | NULL | 115.89 | 73.61 | 0.66 | 9.87E-02 |
| ENSMUSG00000051306 | GSMG0031332 | Usp42 | ubiquitin specific peptidase 42 | 305.93 | 328.35 | -0.10 | 7.93E-01 |
| ENSMUSG00000029610 | GSMG0031335 | Aimp2 | aminoacyl tRNA synthetase complex-interacting multifunctional protei | 336.12 | 471.38 | -0.49 | 2.66E-02 |
| ENSMUSG00000029617 | GSMG0031336 | Ccz1 | CCZ1 vacuolar protein trafficking and biogenesis associated | 613.07 | 791.58 | -0.37 | 4.07E-02 |
| ENSMUSG00000066621 | GSMG0031339 | Tecpr1 | tectonin beta-propeller repeat containing 1 | 8008.47 | 7608.70 | 0.07 | 8.07E-01 |
| ENSMUSG00000043388 | GSMG0031344 | Tmem130 | transmembrane protein 130 | 459.48 | 1058.59 | -1.20 | 1.65E-09 |
| ENSMUSG00000038780 | GSMG0031347 | Smurf1 | SMAD specific E3 ubiquitin protein ligase 1 | 695.17 | 627.60 | 0.15 | 7.11E-01 |
| NULL | GSMG0031348 | NULL | NULL | 278.16 | 249.46 | 0.15 | 6.96E-01 |
| ENSMUSG00000029623 | GSMG0031350 | Pdap1 | PDGFA associated protein 1 | 925.53 | 1437.16 | -0.64 | 6.85E-04 |
| ENSMUSG00000029624 | GSMG0031351 | Ptcd1 | pentatricopeptide repeat domain 1 | 570.10 | 578.11 | -0.02 | 9.52E-01 |
| ENSMUSG00000038690 | GSMG0031352 | Atp5j2 | ATP synthase, H+ transporting, mitochondrial F0 complex, subunit F2 | 2497.56 | 2851.88 | -0.19 | 4.78E-01 |
| ENSMUSG00000029627 | GSMG0031353 | Zkscan14 | zinc finger with KRAB and SCAN domains 14 | 368.09 | 284.84 | 0.37 | 1.31E-01 |
| ENSMUSG00000029634 | GSMG0031365 | Rnf6 | ring finger protein (C3H2C3 type) 6 | 2732.89 | 2485.42 | 0.14 | 5.37E-01 |
| NULL | GSMG0031366 | NULL | NULL | 141.58 | 146.54 | -0.05 | 9.15E-01 |
| ENSMUSG00000041468 | GSMG0031367 | Gpr12 | G-protein coupled receptor 12 | 271.96 | 516.80 | -0.93 | 7.95E-07 |
| ENSMUSG00000029640 | GSMG0031368 | Usp12 | ubiquitin specific peptidase 12 | 411.20 | 497.00 | -0.28 | 2.67E-01 |
| ENSMUSG00000016510 | GSMG0031371 | Mtif3 | mitochondrial translational initiation factor 3 | 255.64 | 224.69 | 0.18 | 5.58E-01 |
| ENSMUSG00000016520 | GSMG0031372 | Lnx2 | ligand of numb-protein X 2 | 1459.56 | 1111.37 | 0.39 | 1.57E-02 |
| ENSMUSG00000042817 | GSMG0031376 | Flt3 | FMS-like tyrosine kinase 3 | 205.91 | 47.91 | 2.10 | 3.76E-16 |
| NULL | GSMG0031377 | NULL | NULL | 130.99 | 98.02 | 0.42 | 2.07E-01 |
| NULL | GSMG0031378 | NULL | NULL | 128.27 | 160.42 | -0.33 | 3.78E-01 |
| ENSMUSG00000029648 | GSMG0031379 | Flt1 | FMS-like tyrosine kinase 1 | 23539.41 | 17107.83 | 0.46 | 2.99E-03 |
| ENSMUSG00000029650 | GSMG0031380 | Slc46a3 | solute carrier family 46, member 3 | 1679.89 | 1556.03 | 0.11 | 6.95E-01 |
| ENSMUSG00000041313 | GSMG0031384 | Slc7a1 | solute carrier family 7 (cationic amino acid transporter, y+ system), mer | 8572.12 | 14438.85 | -0.75 | 5.54E-08 |
| NULL | GSMG0031385 | NULL | NULL | 100.14 | 159.71 | -0.68 | 3.10E-02 |
| ENSMUSG00000001687 | GSMG0031386 | Ubl3 | ubiquitin-like 3 | 4815.82 | 3916.62 | 0.30 | 7.59E-02 |
| ENSMUSG00000041298 | GSMG0031390 | Katnal1 | katanin p60 subunit A-like 1 | 260.34 | 317.97 | -0.28 | 3.44E-01 |

| | | | | | | | |
|---|---|---|---|---|---|---|---|
| NULL | GSMG0031396 | NULL | NULL | 404.62 | 568.86 | -0.50 | 2.75E-02 |
| ENSMUSG00000029657 | GSMG0031401 | Hsph1 | heat shock 105kDa/110kDa protein 1 | 1128.34 | 1024.37 | 0.14 | 5.60E-01 |
| ENSMUSG00000041132 | GSMG0031411 | N4bp2l1 | NEDD4 binding protein 2-like 1 | 476.74 | 281.87 | 0.76 | 4.83E-05 |
| NULL | GSMG0031413 | NULL | NULL | 432.13 | 255.44 | 0.76 | 9.46E-05 |
| NULL | GSMG0031414 | NULL | NULL | 125.79 | 73.89 | 0.77 | 2.90E-02 |
| ENSMUSG00000029655 | GSMG0031416 | N4bp2l2 | NEDD4 binding protein 2-like 2 | 1308.02 | 1195.77 | 0.13 | 6.39E-01 |
| NULL | GSMG0031417 | NULL | NULL | 299.86 | 262.18 | 0.19 | 6.46E-01 |
| ENSMUSG00000016128 | GSMG0031419 | Stard13 | StAR-related lipid transfer (START) domain containing 13 | 1482.02 | 1529.90 | -0.05 | 8.89E-01 |
| NULL | GSMG0031422 | NULL | NULL | 1655.79 | 1382.59 | 0.26 | 2.22E-01 |
| ENSMUSG00000033970 | GSMG0031429 | Rfc3 | replication factor C (activator 1) 3 | 247.78 | 476.36 | -0.94 | 1.21E-06 |
| ENSMUSG00000001376 | GSMG0031430 | Vps50 | VPS50 EARP/GARPII complex subunit | 424.41 | 440.71 | -0.05 | 8.72E-01 |
| ENSMUSG00000032766 | GSMG0031434 | Gng11 | guanine nucleotide binding protein (G protein), gamma 11 | 4728.67 | 4784.31 | -0.02 | 9.64E-01 |
| ENSMUSG00000029661 | GSMG0031435 | Col1a2 | collagen, type I, alpha 2 | 9144.70 | 6472.31 | 0.50 | 1.11E-03 |
| ENSMUSG00000015189 | GSMG0031437 | Casd1 | CAS1 domain containing 1 | 423.49 | 535.69 | -0.34 | 1.28E-01 |
| ENSMUSG00000042607 | GSMG0031447 | Asb4 | ankyrin repeat and SOCS box-containing 4 | 192.81 | 258.17 | -0.41 | 1.99E-01 |
| ENSMUSG00000029757 | GSMG0031448 | Dync1i1 | dynein cytoplasmic 1 intermediate chain 1 | 660.14 | 1221.74 | -0.89 | 1.73E-06 |
| ENSMUSG00000042460 | GSMG0031459 | C1galt1 | core 1 synthase, glycoprotein-N-acetylgalactosamine 3-beta-galactosylt | 198.89 | 194.91 | 0.03 | 9.63E-01 |
| ENSMUSG00000042447 | GSMG0031462 | Mios | missing oocyte, meiosis regulator, homolog (Drosophila) | 249.28 | 329.69 | -0.40 | 7.31E-02 |
| NULL | GSMG0031463 | NULL | NULL | 222.78 | 323.42 | -0.54 | 7.13E-02 |
| G00000029638,ENSMUSG000 | GSMG0031469 | Umad1 // Glcci1 | UMAP1-MVP12 associated (UMA) domain containing 1 // glucocortic | 199.62 | 172.75 | 0.21 | 5.44E-01 |
| ENSMUSG00000046178 | GSMG0031474 | Nxph1 | neurexophilin 1 | 497.90 | 764.29 | -0.62 | 1.91E-03 |
| ENSMUSG00000029629 | GSMG0031484 | Phf14 | PHD finger protein 14 | 315.58 | 367.08 | -0.22 | 4.46E-01 |
| NULL | GSMG0031488 | NULL | NULL | 358.51 | 305.53 | 0.23 | 4.73E-01 |
| ENSMUSG00000029571 | GSMG0031492 | Tmem106b | transmembrane protein 106B | 606.67 | 626.97 | -0.05 | 8.79E-01 |
| ENSMUSG00000041390 | GSMG0031510 | Mdfic | MyoD family inhibitor domain containing | 70.61 | 75.56 | -0.10 | 8.61E-01 |
| ENSMUSG00000029552 | GSMG0031516 | Tes | testis derived transcript | 1252.61 | 1687.36 | -0.43 | 3.04E-02 |
| ENSMUSG00000000058 | GSMG0031519 | Cav2 | caveolin 2 | 4238.78 | 2226.02 | 0.93 | 3.45E-07 |
| ENSMUSG00000007655 | GSMG0031520 | Cav1 | caveolin 1, caveolae protein | 9270.36 | 7472.12 | 0.31 | 2.20E-01 |
| ENSMUSG00000009376 | GSMG0031521 | Met | met proto-oncogene | 273.10 | 796.92 | -1.54 | 8.73E-15 |
| ENSMUSG00000015733 | GSMG0031525 | Capza2 | capping protein (actin filament) muscle Z-line, alpha 2 | 2690.62 | 3041.59 | -0.18 | 4.54E-01 |
| ENSMUSG00000029534 | GSMG0031526 | St7 | suppression of tumorigenicity 7 | 681.92 | 1023.24 | -0.59 | 2.37E-04 |
| NULL | GSMG0031530 | NULL | NULL | 213.43 | 217.35 | -0.03 | 9.49E-01 |
| ENSMUSG00000044155 | GSMG0031531 | Lsm8 | LSM8 homolog, U6 small nuclear RNA associated | 431.36 | 476.69 | -0.15 | 5.93E-01 |

| | | | | | | | |
|---|---|---|---|---|---|---|---|
| ENSMUSG00000060882 | GSMG0031534 | Kcnd2 | potassium voltage-gated channel, Shal-related family, member 2 | 96.78 | 167.14 | -0.79 | 1.42E-02 |
| NULL | GSMG0031535 | NULL | NULL | 348.87 | 252.49 | 0.47 | 8.55E-02 |
| NULL | GSMG0031537 | NULL | NULL | 373.24 | 269.21 | 0.47 | 7.61E-02 |
| NULL | GSMG0031538 | NULL | NULL | 1079.58 | 756.46 | 0.51 | 2.66E-03 |
| NULL | GSMG0031539 | NULL | NULL | 266.46 | 205.10 | 0.38 | 2.92E-01 |
| NULL | GSMG0031544 | NULL | NULL | 403.18 | 270.64 | 0.57 | 1.93E-02 |
| NULL | GSMG0031548 | NULL | NULL | 597.68 | 450.10 | 0.41 | 1.61E-01 |
| NULL | GSMG0031549 | NULL | NULL | 77.48 | 39.40 | 0.97 | 6.87E-02 |
| NULL | GSMG0031551 | NULL | NULL | 381.76 | 332.88 | 0.20 | 4.79E-01 |
| ENSMUSG00000029670 | GSMG0031556 | Ing3 | inhibitor of growth family, member 3 | 402.42 | 430.58 | -0.10 | 7.44E-01 |
| ENSMUSG00000062980 | GSMG0031558 | Cped1 | cadherin-like and PC-esterase domain containing 1 | 483.19 | 375.14 | 0.37 | 1.31E-01 |
| ENSMUSG00000068748 | GSMG0031567 | Ptprz1 | protein tyrosine phosphatase, receptor type Z, polypeptide 1 | 830.01 | 2047.95 | -1.30 | 4.90E-17 |
| ENSMUSG00000020440 | GSMG0031579 | Arf5 | ADP-ribosylation factor 5 | 4491.07 | 5026.10 | -0.16 | 4.68E-01 |
| ENSMUSG00000001424 | GSMG0031581 | Snd1 | staphylococcal nuclease and tudor domain containing 1 | 3742.94 | 5094.08 | -0.44 | 1.55E-03 |
| ENSMUSG00000043421 | GSMG0031589 | Hilpda | hypoxia inducible lipid droplet associated | 654.93 | 671.32 | -0.03 | 9.26E-01 |
| ENSMUSG00000029767 | GSMG0031592 | Calu | calumenin | 3286.83 | 3148.99 | 0.06 | 8.28E-01 |
| ENSMUSG00000029769 | GSMG0031594 | Ccdc136 | coiled-coil domain containing 136 | 400.06 | 684.24 | -0.77 | 1.04E-04 |
| ENSMUSG00000004285 | GSMG0031596 | Atp6v1f | ATPase, H+ transporting, lysosomal V1 subunit F | 1975.97 | 1914.34 | 0.05 | 8.61E-01 |
| ENSMUSG00000029771 | GSMG0031599 | Irf5 | interferon regulatory factor 5 | 254.44 | 203.00 | 0.33 | 2.48E-01 |
| ENSMUSG00000001763 | GSMG0031601 | Tspan33 | tetraspanin 33 | 133.75 | 72.45 | 0.88 | 1.02E-02 |
| ENSMUSG00000001761 | GSMG0031603 | Smo | smoothened, frizzled class receptor | 577.30 | 580.99 | -0.01 | 9.85E-01 |
| ENSMUSG00000029772 | GSMG0031605 | Ahcyl2 | S-adenosylhomocysteine hydrolase-like 2 | 190.12 | 152.10 | 0.32 | 3.83E-01 |
| ENSMUSG00000058440 | GSMG0031614 | Nrf1 | nuclear respiratory factor 1 | 738.26 | 766.05 | -0.05 | 8.67E-01 |
| ENSMUSG00000029775 | GSMG0031620 | Klhdc10 | kelch domain containing 10 | 356.88 | 377.38 | -0.08 | 8.42E-01 |
| ENSMUSG00000054446 | GSMG0031628 | Cpa1 | carboxypeptidase A1, pancreatic | 46.81 | 233.55 | -2.30 | 4.08E-18 |
| ENSMUSG00000051855 | GSMG0031630 | Mest | mesoderm specific transcript | 825.48 | 2254.74 | -1.45 | 6.88E-13 |
| NULL | GSMG0031631 | NULL | NULL | 1038.83 | 1538.96 | -0.57 | 1.02E-03 |
| ENSMUSG00000087380 | GSMG0031635 | 2210408F21Rik | RIKEN cDNA 2210408F21 gene | 137.32 | 26.58 | 2.37 | 1.27E-13 |
| ENSMUSG00000025609 | GSMG0031643 | Mkln1 | muskelin 1, intracellular mediator containing kelch motifs | 606.66 | 592.50 | 0.03 | 9.22E-01 |
| ENSMUSG00000029763 | GSMG0031652 | Exoc4 | exocyst complex component 4 | 578.00 | 603.92 | -0.06 | 8.77E-01 |
| ENSMUSG00000061758 | GSMG0031668 | Akr1b10 | aldo-keto reductase family 1, member B10 (aldose reductase) | 234.69 | 196.86 | 0.26 | 4.32E-01 |
| ENSMUSG00000038871 | GSMG0031670 | Bpgm | 2,3-bisphosphoglycerate mutase | 1126.97 | 1472.09 | -0.38 | 3.85E-02 |
| ENSMUSG00000029761 | GSMG0031672 | Cald1 | caldesmon 1 | 4392.00 | 3820.69 | 0.20 | 4.76E-01 |

| | | | | | | | |
|---|---|---|---|---|---|---|---|
| NULL | GSMG0031677 | NULL | NULL | 204.54 | 122.22 | 0.74 | 4.18E-02 |
| ENSMUSG00000038836 | GSMG0031678 | Agbl3 | ATP/GTP binding protein-like 3 | 105.99 | 108.91 | -0.04 | 9.45E-01 |
| ENSMUSG00000038759 | GSMG0031682 | Nup205 | nucleoporin 205 | 1481.74 | 2311.85 | -0.64 | 4.46E-05 |
| ENSMUSG00000073155 | GSMG0031683 | 1810058I24Rik | RIKEN cDNA 1810058I24 gene | 1386.75 | 1413.97 | -0.03 | 9.28E-01 |
| ENSMUSG00000029833 | GSMG0031694 | Trim24 | tripartite motif-containing 24 | 779.78 | 799.78 | -0.04 | 9.06E-01 |
| ENSMUSG00000038538 | GSMG0031706 | Ubn2 | ubinuclein 2 | 1451.50 | 1160.40 | 0.32 | 8.84E-02 |
| ENSMUSG00000019689 | GSMG0031712 | 1110001J03Rik | RIKEN cDNA 1110001J03 gene | 413.56 | 527.45 | -0.35 | 2.30E-01 |
| ENSMUSG00000029823 | GSMG0031714 | Luc7l2 | LUC7-like 2 (S. cerevisiae) | 562.88 | 557.36 | 0.01 | 9.72E-01 |
| ENSMUSG00000073144 | GSMG0031720 | 4930599N23Rik | RIKEN cDNA 4930599N23 gene | 90.86 | 60.51 | 0.58 | 1.48E-01 |
| ENSMUSG00000046947 | GSMG0031726 | Adck2 | aarF domain containing kinase 2 | 467.91 | 429.19 | 0.12 | 6.66E-01 |
| ENSMUSG00000002416 | GSMG0031727 | Ndufb2 | NADH dehydrogenase (ubiquinone) 1 beta subcomplex, 2 | 1740.37 | 2039.80 | -0.23 | 3.03E-01 |
| NULL | GSMG0031728 | NULL | NULL | 336.89 | 269.69 | 0.32 | 2.09E-01 |
| NULL | GSMG0031730 | NULL | NULL | 492.11 | 375.09 | 0.39 | 1.26E-01 |
| NULL | GSMG0031731 | NULL | NULL | 394.89 | 345.26 | 0.20 | 5.58E-01 |
| ENSMUSG00000057716 | GSMG0031733 | Tmem178b | transmembrane protein 178B | 686.48 | 963.84 | -0.49 | 9.10E-03 |
| ENSMUSG00000029916 | GSMG0031737 | Agk | acylglycerol kinase | 162.25 | 262.63 | -0.69 | 3.09E-03 |
| ENSMUSG00000029911 | GSMG0031741 | Ssbp1 | single-stranded DNA binding protein 1 | 820.57 | 733.57 | 0.16 | 4.94E-01 |
| ENSMUSG00000029869 | GSMG0031755 | Ephb6 | Eph receptor B6 | 3265.20 | 3477.79 | -0.09 | 7.86E-01 |
| ENSMUSG00000029864 | GSMG0031763 | Gstk1 | glutathione S-transferase kappa 1 | 461.80 | 412.82 | 0.16 | 5.71E-01 |
| ENSMUSG00000029863 | GSMG0031765 | Casp2 | caspase 2 | 247.39 | 299.22 | -0.27 | 3.64E-01 |
| ENSMUSG00000029860 | GSMG0031767 | Zyx | zyxin | 2305.78 | 2372.46 | -0.04 | 8.96E-01 |
| ENSMUSG00000039419 | GSMG0031785 | Cntnap2 | contactin associated protein-like 2 | 603.04 | 712.61 | -0.24 | 3.30E-01 |
| NULL | GSMG0031788 | NULL | NULL | 464.06 | 361.48 | 0.36 | 2.20E-01 |
| NULL | GSMG0031796 | NULL | NULL | 162.60 | 122.87 | 0.41 | 3.07E-01 |
| ENSMUSG00000029686 | GSMG0031820 | Cul1 | cullin 1 | 4023.87 | 4409.86 | -0.13 | 5.24E-01 |
| ENSMUSG00000062519 | GSMG0031824 | Zfp398 | zinc finger protein 398 | 200.43 | 249.70 | -0.31 | 2.57E-01 |
| ENSMUSG00000025821 | GSMG0031826 | Zfp282 | zinc finger protein 282 | 276.82 | 336.87 | -0.29 | 2.97E-01 |
| ENSMUSG00000052763 | GSMG0031827 | Zfp212 | Zinc finger protein 212 | 192.94 | 241.80 | -0.33 | 2.29E-01 |
| ENSMUSG00000042810 | GSMG0031833 | Krba1 | KRAB-A domain containing 1 | 1236.88 | 1115.93 | 0.15 | 5.50E-01 |
| ENSMUSG00000107476 | GSMG0031835 | Zfp862-ps | zinc finger protein 862, pseudogene | 397.99 | 424.19 | -0.09 | 8.20E-01 |
| ENSMUSG00000039347 | GSMG0031836 | Atp6v0e2 | ATPase, H+ transporting, lysosomal V0 subunit E2 | 5018.59 | 4380.75 | 0.20 | 3.28E-01 |
| ENSMUSG00000073096 | GSMG0031838 | Lrrc61 | leucine rich repeat containing 61 | 324.63 | 309.66 | 0.07 | 8.57E-01 |
| ENSMUSG00000052751 | GSMG0031840 | Repin1 | replication initiator 1 | 1905.32 | 1402.13 | 0.44 | 2.70E-02 |

| | | | | | | | |
|---|---|---|---|---|---|---|---|
| ENSMUSG00000064262 | GSMG0031843 | Gimap8 | GTPase, IMAP family member 8 | 279.11 | 224.99 | 0.31 | 2.45E-01 |
| ENSMUSG00000051124 | GSMG0031845 | Gimap9 | GTPase, IMAP family member 9 | 431.86 | 571.72 | -0.40 | 1.13E-01 |
| ENSMUSG00000054435 | GSMG0031846 | Gimap4 | GTPase, IMAP family member 4 | 497.34 | 656.97 | -0.40 | 1.77E-01 |
| ENSMUSG00000090019 | GSMG0031849 | Gimap1 | GTPase, IMAP family member 1 | 143.43 | 127.45 | 0.17 | 6.99E-01 |
| ENSMUSG00000043505 | GSMG0031850 | Gimap5 | GTPase, IMAP family member 5 | 977.57 | 816.12 | 0.26 | 3.23E-01 |
| ENSMUSG00000023367 | GSMG0031852 | Tmem176a | transmembrane protein 176A | 555.31 | 436.62 | 0.35 | 1.07E-01 |
| ENSMUSG00000029815 | GSMG0031859 | Malsu1 | mitochondrial assembly of ribosomal large subunit 1 | 579.34 | 811.46 | -0.49 | 6.88E-03 |
| NULL | GSMG0031860 | NULL | NULL | 177.51 | 204.89 | -0.21 | 5.82E-01 |
| ENSMUSG00000050786 | GSMG0031863 | Ccdc126 | coiled-coil domain containing 126 | 673.33 | 651.00 | 0.05 | 8.99E-01 |
| ENSMUSG00000029819 | GSMG0031866 | Npy | neuropeptide Y | 868.32 | 530.12 | 0.71 | 5.74E-04 |
| ENSMUSG00000038388 | GSMG0031868 | Mpp6 | membrane protein, palmitoylated 6 (MAGUK p55 subfamily member 6 | 675.84 | 1195.42 | -0.82 | 9.97E-08 |
| NULL | GSMG0031870 | NULL | NULL | 268.94 | 394.56 | -0.55 | 1.27E-02 |
| NULL | GSMG0031871 | NULL | NULL | 161.30 | 308.18 | -0.94 | 1.10E-03 |
| ENSMUSG00000029832 | GSMG0031881 | Nfe2l3 | nuclear factor, erythroid derived 2, like 3 | 74.10 | 277.84 | -1.90 | 3.52E-17 |
| ENSMUSG00000038301 | GSMG0031885 | Snx10 | sorting nexin 10 | 762.74 | 995.55 | -0.39 | 1.05E-01 |
| ENSMUSG00000004535 | GSMG0031900 | Tax1bp1 | Tax1 (human T cell leukemia virus type I) binding protein 1 | 4710.11 | 4761.19 | -0.02 | 9.62E-01 |
| ENSMUSG00000004633 | GSMG0031911 | Chn2 | chimerin 2 | 535.57 | 377.72 | 0.51 | 2.75E-02 |
| ENSMUSG00000005225 | GSMG0031917 | Plekha8 | pleckstrin homology domain containing, family A (phosphoinositide bi | 232.08 | 295.18 | -0.35 | 2.22E-01 |
| ENSMUSG00000038065 | GSMG0031919 | Mturn | maturin, neural progenitor differentiation regulator homolog (Xenopus) | 234.36 | 468.80 | -1.00 | 1.68E-07 |
| ENSMUSG00000058446 | GSMG0031921 | Znrf2 | zinc and ring finger 2 | 614.03 | 346.18 | 0.83 | 2.36E-01 |
| ENSMUSG00000029777 | GSMG0031925 | Gars | glycyl-tRNA synthetase | 3143.14 | 3566.99 | -0.18 | 3.31E-01 |
| ENSMUSG00000038022 | GSMG0031926 | Fam188b | family with sequence similarity 188, member B | 172.14 | 191.01 | -0.15 | 6.88E-01 |
| ENSMUSG00000029778 | GSMG0031930 | Adcyap1r1 | adenylate cyclase activating polypeptide 1 receptor 1 | 395.60 | 524.29 | -0.41 | 2.19E-01 |
| ENSMUSG00000029787 | GSMG0031935 | Avl9 | AVL9 homolog (S. cerevisiase) | 345.33 | 372.96 | -0.11 | 7.74E-01 |
| ENSMUSG00000029781 | GSMG0031939 | Fkbp9 | FK506 binding protein 9 | 6159.61 | 6162.78 | 0.00 | 9.98E-01 |
| NULL | GSMG0031940 | NULL | NULL | 34.88 | 55.98 | -0.70 | 2.69E-01 |
| ENSMUSG00000062190 | GSMG0031953 | Lancl2 | LanC (bacterial lantibiotic synthetase component C)-like 2 | 383.20 | 530.80 | -0.47 | 2.50E-02 |
| ENSMUSG00000029802 | GSMG0031959 | Abcg2 | ATP-binding cassette, sub-family G (WHITE), member 2 | 9315.40 | 7014.99 | 0.41 | 2.99E-02 |
| ENSMUSG00000029804 | GSMG0031962 | Herc3 | hect domain and RLD 3 | 673.68 | 620.68 | 0.12 | 7.17E-01 |
| ENSMUSG00000049232 | GSMG0031965 | Tigd2 | tigger transposable element derived 2 | 1230.90 | 1057.74 | 0.22 | 3.71E-01 |
| NULL | GSMG0031976 | NULL | NULL | 160.94 | 240.91 | -0.59 | 3.41E-02 |
| ENSMUSG00000071424 | GSMG0031979 | Grid2 | glutamate receptor, ionotropic, delta 2 | 303.91 | 279.17 | 0.12 | 7.08E-01 |
| ENSMUSG00000029920 | GSMG0032001 | Smarcad1 | SWI/SNF-related, matrix-associated actin-dependent regulator of chron | 1029.95 | 1258.58 | -0.29 | 1.61E-01 |

| | | | | | | | |
|---|---|---|---|---|---|---|---|
| ENSMUSG00000049001 | GSMG0032007 | Ndnf | neuron-derived neurotrophic factor | 2465.19 | 762.57 | 1.69 | 5.24E-30 |
| NULL | GSMG0032008 | NULL | NULL | 368.08 | 137.23 | 1.42 | 2.25E-09 |
| ENSMUSG00000029913 | GSMG0032009 | Prdm5 | PR domain containing 5 | 152.64 | 112.91 | 0.44 | 1.96E-01 |
| ENSMUSG00000029910 | GSMG0032012 | Mad2l1 | MAD2 mitotic arrest deficient-like 1 | 224.67 | 427.06 | -0.92 | 4.04E-05 |
| ENSMUSG00000036402 | GSMG0032014 | Gng12 | guanine nucleotide binding protein (G protein), gamma 12 | 4480.15 | 5156.35 | -0.20 | 2.84E-01 |
| ENSMUSG00000087231 | GSMG0032017 | E230016M11Rik | RIKEN cDNA E230016M11 gene | 99.17 | 73.06 | 0.44 | 2.52E-01 |
| ENSMUSG00000036371 | GSMG0032022 | Serbp1 | serpine1 mRNA binding protein 1 | 2613.79 | 3799.56 | -0.54 | 9.35E-03 |
| ENSMUSG00000031668 | GSMG0032040 | Eif2ak3 | eukaryotic translation initiation factor 2 alpha kinase 3 | 1156.83 | 1138.66 | 0.02 | 9.42E-01 |
| ENSMUSG00000053012 | GSMG0032044 | Krcc1 | lysine-rich coiled-coil 1 | 1367.96 | 1090.30 | 0.33 | 1.31E-01 |
| ENSMUSG00000052656 | GSMG0032050 | Rnf103 | ring finger protein 103 | 1219.86 | 970.60 | 0.33 | 8.23E-02 |
| ENSMUSG00000053119 | GSMG0032051 | Chmp3 | charged multivesicular body protein 3 | 1554.28 | 1270.53 | 0.29 | 1.04E-01 |
| ENSMUSG00000052852 | GSMG0032055 | Reep1 | receptor accessory protein 1 | 1041.55 | 1777.21 | -0.77 | 3.58E-07 |
| ENSMUSG00000052337 | GSMG0032058 | Immt | inner membrane protein, mitochondrial | 1124.72 | 1385.47 | -0.30 | 1.66E-01 |
| ENSMUSG00000049553 | GSMG0032061 | Polr1a | polymerase (RNA) I polypeptide A | 1200.49 | 1889.20 | -0.65 | 8.48E-05 |
| ENSMUSG00000056091 | GSMG0032062 | St3gal5 | ST3 beta-galactoside alpha-2,3-sialyltransferase 5 | 1440.82 | 1325.15 | 0.12 | 6.02E-01 |
| ENSMUSG00000058706 | GSMG0032066 | 0610030E20Rik | RIKEN cDNA 0610030E20 gene | 1153.37 | 1073.49 | 0.10 | 6.89E-01 |
| ENSMUSG00000055912 | GSMG0032067 | Tmem150a | transmembrane protein 150A | 133.46 | 210.15 | -0.65 | 2.46E-02 |
| ENSMUSG00000053460 | GSMG0032068 | Ggcx | gamma-glutamyl carboxylase | 837.72 | 853.43 | -0.03 | 9.50E-01 |
| ENSMUSG00000056737 | GSMG0032070 | Capg | capping protein (actin filament), gelsolin-like | 1715.75 | 1019.12 | 0.75 | 2.17E-07 |
| ENSMUSG00000056666 | GSMG0032071 | Retsat | retinol saturase (all trans retinol 13,14 reductase) | 342.63 | 480.28 | -0.48 | 3.15E-02 |
| ENSMUSG00000052738 | GSMG0032078 | Suclg1 | succinate-CoA ligase, GDP-forming, alpha subunit | 2158.07 | 2320.92 | -0.10 | 6.69E-01 |
| ENSMUSG00000060780 | GSMG0032079 | Lrrtm1 | leucine rich repeat transmembrane neuronal 1 | 946.67 | 1205.96 | -0.35 | 1.54E-01 |
| ENSMUSG00000052581 | GSMG0032087 | Lrrtm4 | leucine rich repeat transmembrane neuronal 4 | 164.76 | 175.81 | -0.09 | 8.42E-01 |
| NULL | GSMG0032090 | NULL | NULL | 656.14 | 307.09 | 1.10 | 2.24E-10 |
| NULL | GSMG0032091 | NULL | NULL | 315.08 | 184.23 | 0.78 | 4.55E-03 |
| NULL | GSMG0032093 | NULL | NULL | 279.76 | 147.43 | 0.92 | 1.63E-03 |
| NULL | GSMG0032095 | NULL | NULL | 397.99 | 208.09 | 0.94 | 1.04E-04 |
| ENSMUSG00000035125 | GSMG0032098 | Gcfc2 | GC-rich sequence DNA binding factor 2 | 346.38 | 533.29 | -0.62 | 6.02E-03 |
| ENSMUSG00000035104 | GSMG0032100 | Eva1a | eva-1 homolog A (C. elegans) | 504.74 | 670.65 | -0.41 | 3.28E-02 |
| ENSMUSG00000000693 | GSMG0032106 | Loxl3 | lysyl oxidase-like 3 | 787.35 | 866.30 | -0.14 | 6.11E-01 |
| ENSMUSG00000068328 | GSMG0032107 | Aup1 | ancient ubiquitous protein 1 | 2808.68 | 2581.42 | 0.12 | 5.91E-01 |
| ENSMUSG00000069678 | GSMG0032110 | Pcgf1 | polycomb group ring finger 1 | 309.03 | 280.37 | 0.14 | 6.62E-01 |
| G00000030037,ENSMUSG000 | GSMG0032112 | Ccdc142 // Mrpl53 | coiled-coil domain containing 142 // mitochondrial ribosomal protein L | 2063.63 | 2188.65 | -0.09 | 7.15E-01 |

| | | | | | | | |
|---|---|---|---|---|---|---|---|
| ENSMUSG00000030036 | GSMG0032113 | Mogs | mannosyl-oligosaccharide glucosidase | 757.76 | 697.26 | 0.12 | 7.49E-01 |
| ENSMUSG00000034930 | GSMG0032114 | Rtkn | rhotekin | 326.18 | 161.12 | 1.02 | 7.83E-06 |
| ENSMUSG00000031865 | GSMG0032116 | Dctn1 | dynactin 1 | 1322.38 | 1488.85 | -0.17 | 4.75E-01 |
| ENSMUSG00000043131 | GSMG0032118 | Mob1a | MOB kinase activator 1A | 480.23 | 430.38 | 0.16 | 5.68E-01 |
| ENSMUSG00000045160 | GSMG0032119 | Bola3 | bolA-like 3 (E. coli) | 636.73 | 647.54 | -0.03 | 9.50E-01 |
| NULL | GSMG0032126 | NULL | NULL | 315.65 | 347.59 | -0.14 | 6.60E-01 |
| ENSMUSG00000034744 | GSMG0032127 | Nagk | N-acetylglucosamine kinase | 880.58 | 951.40 | -0.11 | 6.70E-01 |
| ENSMUSG00000030016 | GSMG0032129 | Zfp638 | zinc finger protein 638 | 660.14 | 663.58 | -0.01 | 9.84E-01 |
| ENSMUSG00000033788 | GSMG0032133 | Dysf | dysferlin | 704.57 | 922.34 | -0.39 | 5.21E-02 |
| ENSMUSG00000033726 | GSMG0032136 | Emx1 | empty spiracles homeobox 1 | 80.39 | 141.01 | -0.82 | 1.24E-02 |
| ENSMUSG00000033706 | GSMG0032139 | Smyd5 | SET and MYND domain containing 5 | 195.52 | 339.62 | -0.80 | 1.36E-04 |
| ENSMUSG00000030007 | GSMG0032140 | Cct7 | chaperonin containing Tcp1, subunit 7 (eta) | 4805.82 | 5807.15 | -0.27 | 1.83E-01 |
| ENSMUSG00000054226 | GSMG0032146 | Tprkb | Tp53rk binding protein | 425.70 | 323.01 | 0.40 | 7.40E-02 |
| ENSMUSG00000030000 | GSMG0032149 | Add2 | adducin 2 (beta) | 470.48 | 1094.20 | -1.22 | 1.43E-13 |
| ENSMUSG00000029999 | GSMG0032151 | Tgfa | transforming growth factor alpha | 3672.32 | 3128.80 | 0.23 | 1.86E-01 |
| ENSMUSG00000057497 | GSMG0032154 | Fam136a | family with sequence similarity 136, member A | 668.74 | 821.88 | -0.30 | 1.78E-01 |
| ENSMUSG00000071337 | GSMG0032156 | Tia1 | cytotoxic granule-associated RNA binding protein 1 | 1843.88 | 2337.81 | -0.34 | 3.99E-02 |
| ENSMUSG00000046679 | GSMG0032157 | C87436 | expressed sequence C87436 | 257.78 | 216.71 | 0.25 | 4.10E-01 |
| ENSMUSG00000097048 | GSMG0032159 | 1600020E01Rik | RIKEN cDNA 1600020E01 gene | 545.90 | 551.36 | -0.01 | 9.68E-01 |
| ENSMUSG00000057230 | GSMG0032163 | Aak1 | AP2 associated kinase 1 | 393.32 | 292.86 | 0.43 | 1.09E-01 |
| NULL | GSMG0032165 | NULL | NULL | 177.07 | 103.95 | 0.77 | 5.96E-03 |
| ENSMUSG00000029993 | GSMG0032168 | Nfu1 | NFU1 iron-sulfur cluster scaffold | 613.16 | 632.17 | -0.04 | 8.97E-01 |
| ENSMUSG00000029992 | GSMG0032169 | Gfpt1 | glutamine fructose-6-phosphate transaminase 1 | 713.48 | 835.71 | -0.23 | 3.50E-01 |
| ENSMUSG00000068263 | GSMG0032177 | Efcc1 | EF hand and coiled-coil domain containing 1 | 128.13 | 74.68 | 0.77 | 2.57E-02 |
| ENSMUSG00000030058 | GSMG0032183 | Copg1 | coatomer protein complex, subunit gamma 1 | 3794.23 | 4197.75 | -0.15 | 5.61E-01 |
| ENSMUSG00000030060 | GSMG0032184 | Hmces | 5-hydroxymethylcytosine (hmC) binding, ES cell specific | 335.75 | 374.82 | -0.15 | 6.39E-01 |
| NULL | GSMG0032187 | NULL | NULL | 149.55 | 134.38 | 0.15 | 8.04E-01 |
| NULL | GSMG0032188 | NULL | NULL | 595.25 | 500.16 | 0.25 | 2.94E-01 |
| ENSMUSG00000030062 | GSMG0032189 | Rpn1 | ribophorin I | 7865.91 | 10404.09 | -0.40 | 1.27E-02 |
| ENSMUSG00000015053 | GSMG0032190 | Gata2 | GATA binding protein 2 | 2503.71 | 1220.15 | 1.04 | 8.89E-12 |
| ENSMUSG00000033174 | GSMG0032194 | Mgll | monoglyceride lipase | 5836.31 | 2813.35 | 1.05 | 1.71E-15 |
| NULL | GSMG0032195 | NULL | NULL | 851.81 | 331.25 | 1.36 | 2.59E-16 |
| NULL | GSMG0032197 | NULL | NULL | 357.67 | 140.01 | 1.35 | 7.06E-08 |

| | | | | | | | |
|---|---|---|---|---|---|---|---|
| ENSMUSG00000085828 | GSMG0032198 | Gm15612 | predicted gene 15612 | 442.05 | 821.23 | -0.90 | 1.79E-05 |
| ENSMUSG00000002871 | GSMG0032199 | Tpra1 | transmembrane protein, adipocyte asscociated 1 | 2250.16 | 1980.96 | 0.18 | 3.96E-01 |
| ENSMUSG00000000811 | GSMG0032205 | Txnrd3 | thioredoxin reductase 3 | 246.98 | 270.97 | -0.14 | 6.96E-01 |
| ENSMUSG00000034430 | GSMG0032215 | Zxdc | ZXD family zinc finger C | 448.06 | 446.07 | 0.01 | 9.89E-01 |
| ENSMUSG00000030087 | GSMG0032217 | Klf15 | Kruppel-like factor 15 | 189.73 | 212.52 | -0.16 | 6.91E-01 |
| ENSMUSG00000030088 | GSMG0032218 | Aldh1l1 | aldehyde dehydrogenase 1 family, member L1 | 604.33 | 1112.46 | -0.88 | 1.07E-05 |
| ENSMUSG00000030089 | GSMG0032219 | Slc41a3 | solute carrier family 41, member 3 | 152.39 | 168.45 | -0.14 | 7.57E-01 |
| ENSMUSG00000034245 | GSMG0032221 | Hdac11 | histone deacetylase 11 | 728.54 | 589.46 | 0.30 | 1.87E-01 |
| ENSMUSG00000064080 | GSMG0032222 | Fbln2 | fibulin 2 | 7092.96 | 2188.70 | 1.70 | 1.59E-16 |
| ENSMUSG00000030095 | GSMG0032225 | Tmem43 | transmembrane protein 43 | 905.17 | 1045.16 | -0.21 | 3.36E-01 |
| ENSMUSG00000034192 | GSMG0032226 | Lsm3 | LSM3 homolog, U6 small nuclear RNA and mRNA degradation associa | 461.53 | 686.73 | -0.57 | 7.15E-03 |
| ENSMUSG00000030096 | GSMG0032228 | Slc6a6 | solute carrier family 6 (neurotransmitter transporter, taurine), member 6 | 33069.37 | 15284.51 | 1.11 | 1.08E-14 |
| ENSMUSG00000034083 | GSMG0032230 | Ccdc174 | coiled-coil domain containing 174 | 425.49 | 453.06 | -0.09 | 8.20E-01 |
| ENSMUSG00000034037 | GSMG0032232 | Fgd5 | FYVE, RhoGEF and PH domain containing 5 | 1322.79 | 1135.53 | 0.22 | 4.82E-01 |
| ENSMUSG00000005893 | GSMG0032234 | Nr2c2 | nuclear receptor subfamily 2, group C, member 2 | 612.61 | 668.55 | -0.12 | 6.95E-01 |
| ENSMUSG00000045100 | GSMG0032243 | Slc25a26 | solute carrier family 25 (mitochondrial carrier, phosphate carrier), mem | 101.75 | 100.20 | 0.03 | 9.63E-01 |
| ENSMUSG00000059187 | GSMG0032249 | Fam19a1 | family with sequence similarity 19, member A1 | 1067.71 | 970.83 | 0.14 | 6.89E-01 |
| NULL | GSMG0032251 | NULL | NULL | 1805.54 | 967.44 | 0.90 | 3.26E-07 |
| ENSMUSG00000035199 | GSMG0032252 | Arl6ip5 | ADP-ribosylation factor-like 6 interacting protein 5 | 4387.17 | 4907.84 | -0.16 | 4.32E-01 |
| ENSMUSG00000072875 | GSMG0032264 | Gpr27 | G protein-coupled receptor 27 | 621.90 | 1077.75 | -0.80 | 2.29E-03 |
| ENSMUSG00000030077 | GSMG0032277 | Chl1 | cell adhesion molecule L1-like | 229.77 | 1211.75 | -2.40 | 3.61E-49 |
| ENSMUSG00000030092 | GSMG0032282 | Cntn6 | contactin 6 | 504.57 | 918.17 | -0.86 | 4.38E-06 |
| ENSMUSG00000064293 | GSMG0032284 | Cntn4 | contactin 4 | 500.02 | 435.95 | 0.20 | 4.89E-01 |
| ENSMUSG00000013736 | GSMG0032291 | Trnt1 | tRNA nucleotidyl transferase, CCA-adding, 1 | 348.10 | 415.58 | -0.26 | 4.20E-01 |
| ENSMUSG00000034648 | GSMG0032292 | Lrrn1 | leucine rich repeat protein 1, neuronal | 503.50 | 485.66 | 0.05 | 8.92E-01 |
| ENSMUSG00000030102 | GSMG0032295 | Itpr1 | inositol 1,4,5-trisphosphate receptor 1 | 1089.91 | 1136.28 | -0.06 | 8.53E-01 |
| ENSMUSG00000030103 | GSMG0032301 | Bhlhe40 | basic helix-loop-helix family, member e40 | 3319.11 | 1668.51 | 0.99 | 2.07E-07 |
| ENSMUSG00000030105 | GSMG0032302 | Arl8b | ADP-ribosylation factor-like 8B | 2103.25 | 2216.29 | -0.08 | 7.83E-01 |
| ENSMUSG00000030104 | GSMG0032305 | Edem1 | ER degradation enhancer, mannosidase alpha-like 1 | 463.40 | 522.95 | -0.17 | 5.09E-01 |
| ENSMUSG00000056755 | GSMG0032308 | Grm7 | glutamate receptor, metabotropic 7 | 661.31 | 547.44 | 0.27 | 2.77E-01 |
| NULL | GSMG0032310 | NULL | NULL | 505.17 | 382.55 | 0.40 | 1.49E-01 |
| ENSMUSG00000057604 | GSMG0032318 | Lmcd1 | LIM and cysteine-rich domains 1 | 336.53 | 205.62 | 0.71 | 5.08E-03 |
| ENSMUSG00000030264 | GSMG0032322 | Thumpd3 | THUMP domain containing 3 | 583.85 | 702.26 | -0.27 | 2.38E-01 |

| | | | | | | | |
|---|---|---|---|---|---|---|---|
| ENSMUSG00000034269 | GSMG0032323 | Setd5 | SET domain containing 5 | 604.84 | 597.72 | 0.02 | 9.55E-01 |
| ENSMUSG00000030269 | GSMG0032327 | Mtmr14 | myotubularin related protein 14 | 544.52 | 645.07 | -0.24 | 2.83E-01 |
| ENSMUSG00000030270 | GSMG0032328 | Cpne9 | copine family member IX | 292.54 | 98.35 | 1.58 | 2.38E-11 |
| ENSMUSG00000001632 | GSMG0032329 | Brpf1 | bromodomain and PHD finger containing, 1 | 728.14 | 711.39 | 0.03 | 9.29E-01 |
| ENSMUSG00000030271 | GSMG0032330 | Ogg1 | 8-oxoguanine DNA-glycosylase 1 | 130.01 | 148.62 | -0.20 | 6.58E-01 |
| ENSMUSG00000051256 | GSMG0032333 | Jagn1 | jagunal homolog 1 | 1143.32 | 1237.36 | -0.11 | 6.58E-01 |
| ENSMUSG00000030281 | GSMG0032335 | Il17rc | interleukin 17 receptor C | 449.72 | 285.46 | 0.66 | 4.74E-03 |
| ENSMUSG00000030284 | GSMG0032336 | Creld1 | cysteine-rich with EGF-like domains 1 | 877.87 | 772.05 | 0.19 | 4.21E-01 |
| ENSMUSG00000034023 | GSMG0032337 | Fancd2 | Fanconi anemia, complementation group D2 | 146.23 | 315.40 | -1.10 | 1.31E-05 |
| ENSMUSG00000033940 | GSMG0032338 | Brk1 | BRICK1, SCAR/WAVE actin-nucleating complex subunit | 4069.52 | 4180.26 | -0.04 | 8.96E-01 |
| ENSMUSG00000033933 | GSMG0032340 | Vhl | von Hippel-Lindau tumor suppressor | 1691.31 | 1747.64 | -0.05 | 8.66E-01 |
| ENSMUSG00000060477 | GSMG0032341 | Irak2 | interleukin-1 receptor-associated kinase 2 | 438.55 | 283.39 | 0.63 | 4.40E-03 |
| ENSMUSG00000030307 | GSMG0032345 | Slc6a11 | solute carrier family 6 (neurotransmitter transporter, GABA), member 1 | 517.91 | 916.94 | -0.82 | 1.03E-03 |
| ENSMUSG00000030310 | GSMG0032346 | Slc6a1 | solute carrier family 6 (neurotransmitter transporter, GABA), member 1 | 1252.35 | 1373.84 | -0.13 | 7.32E-01 |
| ENSMUSG00000030314 | GSMG0032348 | Atg7 | autophagy related 7 | 809.43 | 881.56 | -0.12 | 6.26E-01 |
| ENSMUSG00000009394 | GSMG0032350 | Syn2 | synapsin II | 1028.49 | 699.77 | 0.56 | 1.89E-03 |
| NULL | GSMG0032352 | NULL | NULL | 305.47 | 151.21 | 1.01 | 6.91E-05 |
| ENSMUSG00000000440 | GSMG0032353 | Pparg | peroxisome proliferator activated receptor gamma | 118.73 | 23.66 | 2.33 | 4.20E-12 |
| ENSMUSG00000042389 | GSMG0032355 | Tsen2 | tRNA splicing endonuclease subunit 2 | 392.37 | 561.34 | -0.51 | 6.76E-03 |
| ENSMUSG00000000439 | GSMG0032357 | Mkrn2 | makorin, ring finger protein, 2 | 538.14 | 567.22 | -0.08 | 7.91E-01 |
| NULL | GSMG0032358 | NULL | NULL | 953.93 | 1131.50 | -0.25 | 2.28E-01 |
| ENSMUSG00000030319 | GSMG0032360 | Cand2 | cullin-associated and neddylation-dissociated 2 (putative) | 34.73 | 92.17 | -1.41 | 6.87E-04 |
| ENSMUSG00000030323 | GSMG0032361 | Ift122 | intraflagellar transport 122 | 1006.59 | 1077.55 | -0.10 | 7.34E-01 |
| ENSMUSG00000024104 | GSMG0032366 | Fam21 | family with sequence similarity 21 | 2584.64 | 2901.97 | -0.17 | 3.73E-01 |
| ENSMUSG00000025702 | GSMG0032369 | March8 | membrane-associated ring finger (C3HC4) 8 | 1784.02 | 2311.36 | -0.37 | 6.77E-02 |
| ENSMUSG00000048489 | GSMG0032378 | 8430408G22Rik | RIKEN cDNA 8430408G22 gene | 4788.01 | 528.31 | 3.18 | 5.03E-11 |
| ENSMUSG00000061353 | GSMG0032380 | Cxcl12 | chemokine (C-X-C motif) ligand 12 | 8863.02 | 2224.59 | 1.99 | 5.88E-53 |
| ENSMUSG00000059689 | GSMG0032385 | Zfp637 | zinc finger protein 637 | 1171.97 | 1242.47 | -0.08 | 7.48E-01 |
| ENSMUSG00000030134 | GSMG0032389 | Rasgef1a | RasGEF domain family, member 1A | 862.18 | 484.60 | 0.83 | 1.82E-04 |
| ENSMUSG00000041477 | GSMG0032393 | Dcp1b | decapping mRNA 1B | 572.95 | 564.23 | 0.02 | 9.50E-01 |
| ENSMUSG00000030019 | GSMG0032395 | Fbxl14 | F-box and leucine-rich repeat protein 14 | 757.46 | 488.92 | 0.63 | 3.03E-02 |
| ENSMUSG00000030166 | GSMG0032398 | Rad52 | RAD52 homolog, DNA repair protein | 431.43 | 274.77 | 0.65 | 2.75E-03 |
| ENSMUSG00000030180 | GSMG0032402 | Kdm5a | lysine (K)-specific demethylase 5A | 928.85 | 1047.08 | -0.17 | 4.65E-01 |

| | | | | | | | |
|---|---|---|---|---|---|---|---|
| ENSMUSG00000002897 | GSMG0032408 | Il17ra | interleukin 17 receptor A | 129.62 | 88.47 | 0.55 | 1.25E-01 |
| ENSMUSG00000004902 | GSMG0032414 | Slc25a18 | solute carrier family 25 (mitochondrial carrier), member 18 | 222.28 | 313.86 | -0.49 | 3.37E-02 |
| ENSMUSG00000009112 | GSMG0032415 | Bcl2l13 | BCL2-like 13 (apoptosis facilitator) | 199.89 | 240.03 | -0.27 | 4.19E-01 |
| NULL | GSMG0032419 | NULL | NULL | 75.05 | 59.28 | 0.34 | 5.03E-01 |
| ENSMUSG00000067825 | GSMG0032426 | Pex26 | peroxisomal biogenesis factor 26 | 124.52 | 121.73 | 0.03 | 9.56E-01 |
| ENSMUSG00000030107 | GSMG0032428 | Usp18 | ubiquitin specific peptidase 18 | 154.91 | 146.03 | 0.09 | 8.76E-01 |
| ENSMUSG00000030108 | GSMG0032429 | Slc6a13 | solute carrier family 6 (neurotransmitter transporter, GABA), member 1 | 1967.23 | 1419.08 | 0.47 | 5.60E-02 |
| ENSMUSG00000007458 | GSMG0032436 | M6pr | mannose-6-phosphate receptor, cation dependent | 2470.62 | 2668.18 | -0.11 | 6.76E-01 |
| ENSMUSG00000003154 | GSMG0032444 | Foxj2 | forkhead box J2 | 494.38 | 403.87 | 0.29 | 3.39E-01 |
| ENSMUSG00000030327 | GSMG0032445 | Necap1 | NECAP endocytosis associated 1 | 446.54 | 507.81 | -0.19 | 5.33E-01 |
| ENSMUSG00000004270 | GSMG0032463 | Lpcat3 | lysophosphatidylcholine acyltransferase 3 | 3081.51 | 3083.76 | 0.00 | 9.98E-01 |
| ENSMUSG00000004264 | GSMG0032465 | Phb2 | prohibitin 2 | 2067.71 | 2904.40 | -0.49 | 4.40E-03 |
| ENSMUSG00000038451 | GSMG0032468 | Spsb2 | splA/ryanodine receptor domain and SOCS box containing 2 | 209.10 | 100.69 | 1.05 | 3.40E-05 |
| ENSMUSG00000023505 | GSMG0032469 | Cdca3 | cell division cycle associated 3 | 147.93 | 332.28 | -1.16 | 1.64E-06 |
| ENSMUSG00000030120 | GSMG0032471 | Mlf2 | myeloid leukemia factor 2 | 5687.04 | 5885.99 | -0.05 | 8.40E-01 |
| ENSMUSG00000030329 | GSMG0032473 | Pianp | PILR alpha associated neural protein | 164.65 | 151.61 | 0.11 | 8.16E-01 |
| ENSMUSG00000038346 | GSMG0032474 | Zfp384 | zinc finger protein 384 | 833.49 | 824.07 | 0.02 | 9.63E-01 |
| G00000030330,ENSMUSG000 | GSMG0032476 | Ing4 // Acrbp | inhibitor of growth family, member 4 // proacrosin binding protein | 1044.89 | 963.62 | 0.12 | 6.50E-01 |
| ENSMUSG00000063870 | GSMG0032480 | Chd4 | chromodomain helicase DNA binding protein 4 | 3948.36 | 4310.29 | -0.13 | 6.40E-01 |
| ENSMUSG00000038279 | GSMG0032482 | Nop2 | NOP2 nucleolar protein | 605.95 | 950.69 | -0.65 | 1.76E-04 |
| ENSMUSG00000038271 | GSMG0032483 | Iffo1 | intermediate filament family orphan 1 | 365.25 | 207.92 | 0.81 | 3.34E-03 |
| ENSMUSG00000030335 | GSMG0032484 | Mrpl51 | mitochondrial ribosomal protein L51 | 785.23 | 902.21 | -0.20 | 4.66E-01 |
| ENSMUSG00000030337 | GSMG0032485 | Vamp1 // E130112 | vesicle-associated membrane protein 1 // RIKEN cDNA E130112N10 g | 1169.82 | 742.30 | 0.66 | 1.25E-04 |
| ENSMUSG00000030341 | GSMG0032490 | Tnfrsf1a | tumor necrosis factor receptor superfamily, member 1a | 969.95 | 983.82 | -0.02 | 9.46E-01 |
| ENSMUSG00000001930 | GSMG0032492 | Vwf | Von Willebrand factor | 4345.14 | 5090.49 | -0.23 | 7.79E-01 |
| ENSMUSG00000030347 | GSMG0032498 | D6Wsu163e | DNA segment, Chr 6, Wayne State University 163, expressed | 180.48 | 220.59 | -0.29 | 4.25E-01 |
| ENSMUSG00000037997 | GSMG0032505 | Parp11 | poly (ADP-ribose) polymerase family, member 11 | 559.96 | 830.54 | -0.57 | 2.65E-03 |
| NULL | GSMG0032509 | NULL | NULL | 356.23 | 388.48 | -0.12 | 6.93E-01 |
| ENSMUSG00000001520 | GSMG0032514 | Nrip2 | nuclear receptor interacting protein 2 | 335.44 | 87.81 | 1.94 | 3.79E-17 |
| ENSMUSG00000030161 | GSMG0032535 | Gabarapl1 | gamma-aminobutyric acid (GABA) A receptor-associated protein-like 1 | 488.01 | 494.64 | -0.02 | 9.50E-01 |
| ENSMUSG00000030199 | GSMG0032554 | Etv6 | ets variant 6 | 560.28 | 678.88 | -0.28 | 2.53E-01 |
| NULL | GSMG0032555 | NULL | NULL | 592.34 | 550.09 | 0.10 | 7.26E-01 |
| NULL | GSMG0032557 | NULL | NULL | 388.98 | 427.37 | -0.13 | 7.33E-01 |

| | | | | | | | |
|---|---|---|---|---|---|---|---|
| ENSMUSG00000042992 | GSMG0032566 | Borcs5 | BLOC-1 related complex subunit 5 | 325.61 | 464.16 | -0.51 | 1.83E-02 |
| ENSMUSG00000032652 | GSMG0032568 | Crebl2 | cAMP responsive element binding protein-like 2 | 365.52 | 178.01 | 1.04 | 4.96E-07 |
| ENSMUSG00000098318 | GSMG0032571 | Lockd | lncRNA downstream of Cdkn1b | 41.75 | 74.85 | -0.85 | 5.80E-02 |
| ENSMUSG00000030204 | GSMG0032576 | Ddx47 | DEAD (Asp-Glu-Ala-Asp) box polypeptide 47 | 1749.23 | 1743.08 | 0.01 | 9.85E-01 |
| ENSMUSG00000030207 | GSMG0032578 | Fam234b | family with sequence similarity 234, member B | 1091.77 | 1321.67 | -0.28 | 2.09E-01 |
| ENSMUSG00000030208 | GSMG0032580 | Emp1 | epithelial membrane protein 1 | 544.51 | 304.52 | 0.84 | 5.88E-06 |
| ENSMUSG00000030213 | GSMG0032585 | Atf7ip | activating transcription factor 7 interacting protein | 1461.89 | 1913.75 | -0.39 | 3.71E-02 |
| ENSMUSG00000060032 | GSMG0032588 | H2afj | H2A histone family, member J | 940.01 | 885.79 | 0.08 | 8.15E-01 |
| ENSMUSG00000030223 | GSMG0032591 | Ptpro | protein tyrosine phosphatase, receptor type, O | 184.57 | 520.77 | -1.49 | 3.66E-14 |
| ENSMUSG00000030224 | GSMG0032593 | Strap | serine/threonine kinase receptor associated protein | 942.64 | 1148.36 | -0.28 | 1.26E-01 |
| ENSMUSG00000030225 | GSMG0032594 | Dera | deoxyribose-phosphate aldolase (putative) | 279.29 | 575.05 | -1.04 | 7.43E-08 |
| ENSMUSG00000008540 | GSMG0032595 | Mgst1 | microsomal glutathione S-transferase 1 | 334.30 | 303.44 | 0.14 | 7.21E-01 |
| ENSMUSG00000030231 | GSMG0032604 | Plekha5 | pleckstrin homology domain containing, family A member 5 | 988.49 | 951.86 | 0.05 | 8.67E-01 |
| NULL | GSMG0032605 | NULL | NULL | 425.53 | 326.37 | 0.38 | 1.72E-01 |
| ENSMUSG00000030232 | GSMG0032607 | Aebp2 | AE binding protein 2 | 454.07 | 314.24 | 0.53 | 2.92E-02 |
| ENSMUSG00000041741 | GSMG0032612 | Pde3a | phosphodiesterase 3A, cGMP inhibited | 462.88 | 310.14 | 0.58 | 2.57E-02 |
| ENSMUSG00000030235 | GSMG0032619 | Slco1c1 | solute carrier organic anion transporter family, member 1c1 | 52079.32 | 49002.66 | 0.09 | 7.52E-01 |
| ENSMUSG00000041671 | GSMG0032622 | Pyroxd1 | pyridine nucleotide-disulphide oxidoreductase domain 1 | 397.63 | 347.22 | 0.20 | 4.84E-01 |
| ENSMUSG00000030245 | GSMG0032623 | Golt1b | golgi transport 1B | 1757.31 | 2724.78 | -0.63 | 9.48E-05 |
| ENSMUSG00000030282 | GSMG0032626 | Cmas | cytidine monophospho-N-acetylneuraminic acid synthetase | 861.77 | 1133.69 | -0.40 | 3.29E-02 |
| ENSMUSG00000030275 | GSMG0032630 | Etnk1 | ethanolamine kinase 1 | 878.66 | 839.34 | 0.07 | 8.04E-01 |
| ENSMUSG00000040370 | GSMG0032638 | Lyrm5 | LYR motif containing 5 | 469.52 | 438.28 | 0.10 | 7.86E-01 |
| ENSMUSG00000086013 | GSMG0032640 | Gm15706 | predicted gene 15706 | 300.39 | 218.78 | 0.45 | 1.39E-01 |
| ENSMUSG00000030259 | GSMG0032647 | Rassf8 | Ras association (RalGDS/AF-6) domain family (N-terminal) member 8 | 247.43 | 262.24 | -0.08 | 8.25E-01 |
| NULL | GSMG0032648 | NULL | NULL | 498.40 | 466.44 | 0.10 | 7.71E-01 |
| ENSMUSG00000030255 | GSMG0032650 | Sspn | sarcospan | 690.51 | 449.80 | 0.62 | 6.31E-03 |
| ENSMUSG00000040242 | GSMG0032653 | Fgfr1op2 | FGFR1 oncogene partner 2 | 626.19 | 615.86 | 0.03 | 9.40E-01 |
| ENSMUSG00000030291 | GSMG0032654 | Med21 | mediator complex subunit 21 | 924.98 | 904.47 | 0.03 | 9.16E-01 |
| ENSMUSG00000001630 | GSMG0032656 | Stk38l | serine/threonine kinase 38 like | 100.41 | 80.95 | 0.31 | 4.84E-01 |
| ENSMUSG00000016487 | GSMG0032660 | Ppfibp1 | PTPRF interacting protein, binding protein 1 (liprin beta 1) | 3506.35 | 3421.75 | 0.04 | 8.94E-01 |
| ENSMUSG00000040121 | GSMG0032661 | Rep15 | RAB15 effector protein | 75.08 | 111.21 | -0.56 | 1.79E-01 |
| ENSMUSG00000040112 | GSMG0032662 | Mrps35 | mitochondrial ribosomal protein S35 | 455.76 | 558.26 | -0.29 | 1.84E-01 |
| ENSMUSG00000040102 | GSMG0032665 | Klhl42 | kelch-like 42 | 1207.82 | 1197.40 | 0.01 | 9.75E-01 |

| | | | | | | | |
|---|---|---|---|---|---|---|---|
| ENSMUSG00000030301 | GSMG0032667 | Ccdc91 | coiled-coil domain containing 91 | 663.57 | 530.81 | 0.32 | 1.64E-01 |
| ENSMUSG00000030303 | GSMG0032670 | Far2 | fatty acyl CoA reductase 2 | 2406.97 | 2662.73 | -0.15 | 5.83E-01 |
| ENSMUSG00000063171 | GSMG0032672 | Rps4l | ribosomal protein S4-like | 642.46 | 920.94 | -0.52 | 2.30E-02 |
| ENSMUSG00000039958 | GSMG0032676 | Mettl20 | methyltransferase like 20 | 56.26 | 66.71 | -0.25 | 6.69E-01 |
| ENSMUSG00000032712 | GSMG0032677 | 2810474O19Rik | RIKEN cDNA 2810474O19 gene | 711.90 | 733.57 | -0.04 | 8.87E-01 |
| ENSMUSG00000003452 | GSMG0032681 | Bicd1 | bicaudal D homolog 1 (Drosophila) | 1490.88 | 1205.13 | 0.31 | 1.54E-01 |
| ENSMUSG00000047735 | GSMG0032686 | Samd9l | sterile alpha motif domain containing 9-like | 1555.66 | 1210.17 | 0.36 | 1.28E-01 |
| ENSMUSG00000032757 | GSMG0032691 | Bet1 | Bet1 golgi vesicular membrane trafficking protein | 993.68 | 838.90 | 0.24 | 3.78E-01 |
| ENSMUSG00000004631 | GSMG0032695 | Sgce | sarcoglycan, epsilon | 731.00 | 1087.80 | -0.57 | 2.64E-03 |
| NULL | GSMG0032696 | NULL | NULL | 205.26 | 279.16 | -0.45 | 9.93E-02 |
| ENSMUSG00000029759 | GSMG0032700 | Pon3 | paraoxonase 3 | 137.14 | 196.70 | -0.52 | 1.05E-01 |
| ENSMUSG00000032667 | GSMG0032701 | Pon2 | paraoxonase 2 | 3300.55 | 3529.48 | -0.10 | 7.13E-01 |
| ENSMUSG00000019577 | GSMG0032703 | Pdk4 | pyruvate dehydrogenase kinase, isoenzyme 4 | 362.01 | 450.88 | -0.32 | 2.20E-01 |
| ENSMUSG00000042541 | GSMG0032709 | Shfm1 | split hand/foot malformation (ectrodactyly) type 1 | 2678.47 | 3124.39 | -0.22 | 4.01E-01 |
| NULL | GSMG0032711 | NULL | NULL | 252.67 | 278.00 | -0.14 | 6.86E-01 |
| NULL | GSMG0032714 | NULL | NULL | 100.95 | 134.08 | -0.41 | 2.27E-01 |
| ENSMUSG00000029752 | GSMG0032719 | Asns | asparagine synthetase | 748.70 | 1035.96 | -0.47 | 7.45E-03 |
| ENSMUSG00000068794 | GSMG0032720 | Col28a1 | collagen, type XXVIII, alpha 1 | 354.22 | 259.39 | 0.45 | 6.68E-02 |
| ENSMUSG00000012483 | GSMG0032721 | Rpa3 | replication protein A3 | 398.61 | 447.84 | -0.17 | 5.97E-01 |
| ENSMUSG00000062995 | GSMG0032723 | Ica1 | islet cell autoantigen 1 | 874.78 | 1109.42 | -0.34 | 7.46E-02 |
| ENSMUSG00000029632 | GSMG0032729 | Ndufa4 | NADH dehydrogenase (ubiquinone) 1 alpha subcomplex, 4 | 4044.59 | 4575.16 | -0.18 | 4.49E-01 |
| ENSMUSG00000029569 | GSMG0032739 | Tmem168 | transmembrane protein 168 | 1039.26 | 1074.12 | -0.05 | 8.80E-01 |
| ENSMUSG00000042742 | GSMG0032740 | Bmt2 | base methyltransferase of 25S rRNA 2 | 719.93 | 649.47 | 0.15 | 6.03E-01 |
| ENSMUSG00000048216 | GSMG0032742 | Gpr85 | G protein-coupled receptor 85 | 1378.03 | 2458.48 | -0.84 | 3.08E-08 |
| ENSMUSG00000052419 | GSMG0032744 | 2610001J05Rik | RIKEN cDNA 2610001J05 gene | 1858.14 | 2250.55 | -0.28 | 2.27E-01 |
| ENSMUSG00000000416 | GSMG0032759 | Cttnbp2 | cortactin binding protein 2 | 2014.84 | 1729.75 | 0.22 | 2.82E-01 |
| NULL | GSMG0032760 | NULL | NULL | 166.82 | 146.54 | 0.18 | 7.07E-01 |
| ENSMUSG00000029669 | GSMG0032764 | Tspan12 | tetraspanin 12 | 1275.89 | 1307.69 | -0.04 | 9.10E-01 |
| ENSMUSG00000029672 | GSMG0032766 | Fam3c | family with sequence similarity 3, member C | 799.93 | 961.60 | -0.26 | 2.12E-01 |
| ENSMUSG00000017978 | GSMG0032770 | Cadps2 | Ca2+-dependent activator protein for secretion 2 | 370.96 | 208.66 | 0.83 | 4.89E-04 |
| ENSMUSG00000023089 | GSMG0032780 | Ndufa5 | NADH dehydrogenase (ubiquinone) 1 alpha subcomplex, 5 | 1325.88 | 1433.81 | -0.11 | 6.97E-01 |
| ENSMUSG00000029684 | GSMG0032781 | Wasl | Wiskott-Aldrich syndrome-like (human) | 538.69 | 478.54 | 0.17 | 6.23E-01 |
| ENSMUSG00000048022 | GSMG0032785 | Tmem229a | transmembrane protein 229A | 581.68 | 551.95 | 0.08 | 8.15E-01 |

| | | | | | | | |
|---|---|---|---|---|---|---|---|
| ENSMUSG00000029676 | GSMG0032788 | Pot1a | protection of telomeres 1A | 769.44 | 839.69 | -0.12 | 6.57E-01 |
| ENSMUSG00000024211 | GSMG0032789 | Grm8 | glutamate receptor, metabotropic 8 | 365.91 | 333.76 | 0.14 | 6.69E-01 |
| ENSMUSG00000039841 | GSMG0032796 | Zfp800 | zinc finger protein 800 | 646.96 | 557.74 | 0.21 | 4.02E-01 |
| ENSMUSG00000029708 | GSMG0032797 | Gcc1 | golgi coiled coil 1 | 275.95 | 205.75 | 0.42 | 1.83E-01 |
| ENSMUSG00000049939 | GSMG0032799 | Lrrc4 | leucine rich repeat containing 4 | 396.24 | 297.55 | 0.41 | 2.20E-01 |
| ENSMUSG00000029701 | GSMG0032800 | Rbm28 | RNA binding motif protein 28 | 1437.00 | 1671.47 | -0.22 | 2.32E-01 |
| ENSMUSG00000079654 | GSMG0032801 | Prrt4 | proline-rich transmembrane protein 4 | 231.21 | 137.16 | 0.75 | 1.12E-02 |
| ENSMUSG00000003500 | GSMG0032802 | Impdh1 | inosine 5'-phosphate dehydrogenase 1 | 4019.53 | 3498.69 | 0.20 | 3.23E-01 |
| ENSMUSG00000059022 | GSMG0032806 | Kcp | kielin/chordin-like protein | 3867.19 | 4183.31 | -0.11 | 7.00E-01 |
| ENSMUSG00000012535 | GSMG0032807 | Tnpo3 | transportin 3 | 2205.76 | 2584.79 | -0.23 | 2.58E-01 |
| ENSMUSG00000039159 | GSMG0032811 | Ube2h | ubiquitin-conjugating enzyme E2H | 3991.34 | 3658.16 | 0.13 | 5.95E-01 |
| ENSMUSG00000039130 | GSMG0032816 | Zc3hc1 | zinc finger, C3HC type 1 | 578.23 | 850.02 | -0.55 | 3.58E-03 |
| ENSMUSG00000029782 | GSMG0032817 | Tmem209 | transmembrane protein 209 | 625.65 | 797.53 | -0.35 | 1.03E-01 |
| ENSMUSG00000029790 | GSMG0032818 | Cep41 | centrosomal protein 41 | 56.61 | 75.27 | -0.41 | 4.00E-01 |
| ENSMUSG00000025607 | GSMG0032819 | Copg2 | coatomer protein complex, subunit gamma 2 | 741.41 | 789.61 | -0.09 | 7.89E-01 |
| NULL | GSMG0032824 | NULL | NULL | 526.81 | 60.88 | 3.12 | 1.49E-36 |
| NULL | GSMG0032825 | NULL | NULL | 4332.14 | 3825.00 | 0.18 | 3.40E-01 |
| ENSMUSG00000025608 | GSMG0032830 | Podxl | podocalyxin-like | 15769.00 | 10639.05 | 0.57 | 1.74E-03 |
| ENSMUSG00000029765 | GSMG0032834 | Plxna4 | plexin A4 | 456.18 | 585.51 | -0.36 | 1.72E-01 |
| ENSMUSG00000053768 | GSMG0032836 | Chchd3 | coiled-coil-helix-coiled-coil-helix domain containing 3 | 817.49 | 886.37 | -0.12 | 6.63E-01 |
| ENSMUSG00000018999 | GSMG0032847 | Slc35b4 | solute carrier family 35, member B4 | 371.77 | 373.63 | 0.00 | 9.93E-01 |
| NULL | GSMG0032850 | NULL | NULL | 134.01 | 41.39 | 1.69 | 1.64E-07 |
| NULL | GSMG0032852 | NULL | NULL | 152.97 | 135.17 | 0.18 | 7.89E-01 |
| ENSMUSG00000046806 | GSMG0032853 | 3110062M04Rik | RIKEN cDNA 3110062M04 gene | 701.90 | 473.41 | 0.57 | 1.82E-03 |
| ENSMUSG00000058486 | GSMG0032854 | Wdr91 | WD repeat domain 91 | 617.89 | 657.96 | -0.09 | 7.73E-01 |
| ENSMUSG00000038784 | GSMG0032858 | Cnot4 | CCR4-NOT transcription complex, subunit 4 | 987.24 | 889.41 | 0.15 | 5.81E-01 |
| ENSMUSG00000029843 | GSMG0032867 | Slc13a4 | solute carrier family 13 (sodium/sulfate symporters), member 4 | 671.19 | 676.42 | -0.01 | 9.86E-01 |
| ENSMUSG00000047420 | GSMG0032868 | Fam180a | family with sequence similarity 180, member A | 721.63 | 340.41 | 1.09 | 2.06E-07 |
| ENSMUSG00000029840 | GSMG0032871 | Mtpn | myotrophin | 4475.52 | 5395.69 | -0.27 | 1.72E-01 |
| ENSMUSG00000029838 | GSMG0032877 | Ptn | pleiotrophin | 28837.23 | 25727.91 | 0.16 | 5.99E-01 |
| NULL | GSMG0032879 | NULL | NULL | 444.02 | 444.44 | 0.00 | 9.97E-01 |
| ENSMUSG00000038665 | GSMG0032884 | Dgki | diacylglycerol kinase, iota | 132.00 | 160.48 | -0.27 | 4.79E-01 |
| NULL | GSMG0032885 | NULL | NULL | 311.51 | 236.63 | 0.40 | 2.02E-01 |

| | | | | | | | |
|---|---|---|---|---|---|---|---|
| NULL | GSMG0032888 | NULL | NULL | 399.56 | 234.27 | 0.77 | 1.81E-03 |
| ENSMUSG00000038648 | GSMG0032889 | Creb3l2 | cAMP responsive element binding protein 3-like 2 | 646.08 | 611.18 | 0.08 | 7.79E-01 |
| ENSMUSG00000063455 | GSMG0032896 | D630045J12Rik | RIKEN cDNA D630045J12 gene | 70.96 | 98.47 | -0.47 | 2.67E-01 |
| ENSMUSG00000029826 | GSMG0032901 | Zc3hav1 | zinc finger CCCH type, antiviral 1 | 309.24 | 279.42 | 0.15 | 6.93E-01 |
| ENSMUSG00000061436 | GSMG0032910 | Hipk2 | homeodomain interacting protein kinase 2 | 1045.99 | 927.05 | 0.17 | 6.79E-01 |
| ENSMUSG00000038507 | GSMG0032915 | Parp12 | poly (ADP-ribose) polymerase family, member 12 | 449.92 | 418.36 | 0.11 | 7.55E-01 |
| ENSMUSG00000042599 | GSMG0032918 | Kdm7a | lysine (K)-specific demethylase 7A | 201.47 | 140.70 | 0.52 | 6.31E-02 |
| ENSMUSG00000029924 | GSMG0032920 | Slc37a3 | solute carrier family 37 (glycerol-3-phosphate transporter), member 3 | 757.88 | 828.90 | -0.13 | 6.14E-01 |
| ENSMUSG00000029922 | GSMG0032921 | Mkrn1 | makorin, ring finger protein, 1 | 1731.65 | 1776.11 | -0.04 | 8.92E-01 |
| ENSMUSG00000038456 | GSMG0032922 | Dennd2a | DENN/MADD domain containing 2A | 770.69 | 978.51 | -0.35 | 1.91E-01 |
| NULL | GSMG0032925 | NULL | NULL | 164.01 | 174.88 | -0.10 | 8.24E-01 |
| ENSMUSG00000002413 | GSMG0032926 | Braf | Braf transforming gene | 375.28 | 300.00 | 0.32 | 1.87E-01 |
| NULL | GSMG0032927 | NULL | NULL | 335.16 | 300.59 | 0.16 | 6.23E-01 |
| ENSMUSG00000037172 | GSMG0032934 | E330009J07Rik | RIKEN cDNA E330009J07 gene | 407.75 | 360.58 | 0.18 | 6.83E-01 |
| ENSMUSG00000029861 | GSMG0032956 | Fam131b | family with sequence similarity 131, member B | 67.81 | 45.37 | 0.58 | 2.08E-01 |
| NULL | GSMG0032958 | NULL | NULL | 91.09 | 97.15 | -0.09 | 8.73E-01 |
| ENSMUSG00000029851 | GSMG0032963 | Tcaf2 | TRPM8 channel-associated factor 2 | 387.78 | 378.73 | 0.03 | 9.38E-01 |
| ENSMUSG00000036667 | GSMG0032964 | Tcaf1 | TRPM8 channel-associated factor 1 | 3237.45 | 3340.57 | -0.04 | 8.83E-01 |
| ENSMUSG00000029735 | GSMG0032970 | Tpk1 | thiamine pyrophosphokinase | 205.26 | 215.19 | -0.06 | 8.76E-01 |
| NULL | GSMG0032973 | NULL | NULL | 79.64 | 75.13 | 0.09 | 8.83E-01 |
| NULL | GSMG0032974 | NULL | NULL | 88.50 | 105.58 | -0.25 | 5.94E-01 |
| ENSMUSG00000029687 | GSMG0032977 | Ezh2 | enhancer of zeste 2 polycomb repressive complex 2 subunit | 581.08 | 982.12 | -0.76 | 1.30E-04 |
| ENSMUSG00000025823 | GSMG0032983 | Pdia4 | protein disulfide isomerase associated 4 | 4679.58 | 5747.89 | -0.30 | 1.25E-01 |
| ENSMUSG00000029814 | GSMG0032985 | Igf2bp3 | insulin-like growth factor 2 mRNA binding protein 3 | 37.45 | 224.81 | -2.57 | 3.68E-20 |
| ENSMUSG00000071477 | GSMG0032986 | Zfp777 | zinc finger protein 777 | 462.51 | 290.65 | 0.67 | 5.09E-02 |
| ENSMUSG00000057691 | GSMG0032987 | Zfp746 | zinc finger protein 746 | 1396.49 | 1341.93 | 0.06 | 8.44E-01 |
| ENSMUSG00000068551 | GSMG0032990 | Zfp467 | zinc finger protein 467 | 520.38 | 564.82 | -0.12 | 6.69E-01 |
| ENSMUSG00000009281 | GSMG0032992 | Rarres2 | retinoic acid receptor responder (tazarotene induced) 2 | 990.91 | 614.95 | 0.69 | 2.31E-04 |
| ENSMUSG00000047867 | GSMG0032993 | Gimap6 | GTPase, IMAP family member 6 | 5133.28 | 4284.84 | 0.26 | 2.55E-01 |
| ENSMUSG00000029810 | GSMG0032996 | Tmem176b | transmembrane protein 176B | 2175.28 | 2114.50 | 0.04 | 8.95E-01 |
| ENSMUSG00000029817 | GSMG0033004 | Tra2a | transformer 2 alpha homolog (Drosophila) | 646.68 | 562.78 | 0.20 | 3.90E-01 |
| ENSMUSG00000029821 | GSMG0033006 | Dfna5 | deafness, autosomal dominant 5 (human) | 57.87 | 132.16 | -1.20 | 1.62E-04 |
| ENSMUSG00000029822 | GSMG0033007 | Osbpl3 | oxysterol binding protein-like 3 | 597.99 | 708.91 | -0.25 | 2.75E-01 |

| | | | | | | | |
|---|---|---|---|---|---|---|---|
| ENSMUSG00000004980 | GSMG0033016 | Hnrnpa2b1 | heterogeneous nuclear ribonucleoprotein A2/B1 | 10346.25 | 13718.68 | -0.41 | 1.86E-02 |
| ENSMUSG00000059182 | GSMG0033019 | Skap2 | src family associated phosphoprotein 2 | 300.00 | 182.41 | 0.72 | 2.70E-03 |
| ENSMUSG00000029776 | GSMG0033042 | Hibadh | 3-hydroxyisobutyrate dehydrogenase | 583.74 | 609.29 | -0.06 | 8.58E-01 |
| ENSMUSG00000063568 | GSMG0033044 | Jazf1 | JAZF zinc finger 1 | 378.85 | 732.78 | -0.95 | 1.79E-07 |
| NULL | GSMG0033046 | NULL | NULL | 199.30 | 157.68 | 0.34 | 3.12E-01 |
| NULL | GSMG0033047 | NULL | NULL | 465.75 | 379.74 | 0.30 | 2.01E-01 |
| ENSMUSG00000043496 | GSMG0033049 | Tril | TLR4 interactor with leucine-rich repeats | 2360.33 | 2255.01 | 0.07 | 8.31E-01 |
| ENSMUSG00000019124 | GSMG0033056 | Scrn1 | secernin 1 | 443.94 | 605.53 | -0.45 | 6.26E-02 |
| ENSMUSG00000038074 | GSMG0033058 | Fkbp14 | FK506 binding protein 14 | 518.68 | 490.95 | 0.08 | 8.40E-01 |
| ENSMUSG00000038058 | GSMG0033064 | Nod1 | nucleotide-binding oligomerization domain containing 1 | 469.70 | 262.63 | 0.84 | 6.57E-05 |
| ENSMUSG00000037984 | GSMG0033068 | Neurod6 | neurogenic differentiation 6 | 397.64 | 1408.36 | -1.83 | 2.02E-27 |
| NULL | GSMG0033072 | NULL | NULL | 287.90 | 406.90 | -0.50 | 5.25E-02 |
| ENSMUSG00000029780 | GSMG0033075 | Nt5c3 | 5'-nucleotidase, cytosolic III | 531.79 | 774.88 | -0.55 | 2.28E-03 |
| ENSMUSG00000037826 | GSMG0033080 | Ppm1k | protein phosphatase 1K (PP2C domain containing) | 52.61 | 41.49 | 0.34 | 5.90E-01 |
| ENSMUSG00000043162 | GSMG0033081 | Pyurf | Pigy upstream reading frame | 445.64 | 442.04 | 0.01 | 9.72E-01 |
| ENSMUSG00000037788 | GSMG0033082 | Vopp1 | vesicular, overexpressed in cancer, prosurvival protein 1 | 1696.25 | 1974.74 | -0.22 | 2.64E-01 |
| ENSMUSG00000055430 | GSMG0033088 | Nap1l5 | nucleosome assembly protein 1-like 5 | 2036.95 | 955.09 | 1.09 | 4.55E-07 |
| ENSMUSG00000037709 | GSMG0033089 | Fam13a | family with sequence similarity 13, member A | 974.33 | 1111.75 | -0.19 | 5.03E-01 |
| ENSMUSG00000025889 | GSMG0033094 | Snca | synuclein, alpha | 2537.60 | 2972.15 | -0.23 | 3.04E-01 |
| ENSMUSG00000036390 | GSMG0033107 | Gadd45a | growth arrest and DNA-damage-inducible 45 alpha | 205.26 | 161.92 | 0.34 | 2.89E-01 |
| NULL | GSMG0033151 | NULL | NULL | 125.57 | 217.59 | -0.80 | 1.38E-02 |
| ENSMUSG00000054474 | GSMG0033152 | Thnsl2 | threonine synthase-like 2 (bacterial) | 364.38 | 335.68 | 0.12 | 7.00E-01 |
| ENSMUSG00000002222 | GSMG0033156 | Rmnd5a | required for meiotic nuclear division 5 homolog A | 231.38 | 152.61 | 0.60 | 6.34E-02 |
| ENSMUSG00000053470 | GSMG0033161 | Kdm3a | lysine (K)-specific demethylase 3A | 480.60 | 503.78 | -0.07 | 8.20E-01 |
| ENSMUSG00000052962 | GSMG0033162 | Mrpl35 | mitochondrial ribosomal protein L35 | 385.95 | 593.11 | -0.62 | 1.28E-03 |
| ENSMUSG00000063884 | GSMG0033164 | Ptcd3 | pentatricopeptide repeat domain 3 | 1382.09 | 1584.93 | -0.20 | 3.16E-01 |
| ENSMUSG00000056305 | GSMG0033169 | Usp39 | ubiquitin specific peptidase 39 | 693.46 | 909.43 | -0.39 | 4.24E-02 |
| NULL | GSMG0033170 | NULL | NULL | 204.04 | 220.01 | -0.12 | 8.03E-01 |
| ENSMUSG00000055850 | GSMG0033171 | Rnf181 | ring finger protein 181 | 2065.29 | 1811.32 | 0.19 | 3.18E-01 |
| ENSMUSG00000073002 | GSMG0033173 | Vamp5 | vesicle-associated membrane protein 5 | 602.29 | 618.08 | -0.04 | 9.16E-01 |
| NULL | GSMG0033174 | NULL | NULL | 250.49 | 413.23 | -0.72 | 2.26E-03 |
| ENSMUSG00000050732 | GSMG0033176 | Vamp8 | vesicle-associated membrane protein 8 | 1433.46 | 1105.42 | 0.37 | 3.03E-02 |
| ENSMUSG00000053907 | GSMG0033177 | Mat2a | methionine adenosyltransferase II, alpha | 6533.52 | 6080.67 | 0.10 | 6.60E-01 |

| | | | | | | | |
|---|---|---|---|---|---|---|---|
| ENSMUSG00000056698 | GSMG0033179 | Elmod3 | ELMO/CED-12 domain containing 3 | 458.77 | 414.47 | 0.15 | 5.96E-01 |
| ENSMUSG00000056429 | GSMG0033181 | Tgoln1 | trans-golgi network protein | 1588.09 | 1501.54 | 0.08 | 7.90E-01 |
| ENSMUSG00000055799 | GSMG0033182 | Tcf7l1 | transcription factor 7 like 1 (T cell specific, HMG box) | 148.29 | 145.98 | 0.02 | 9.74E-01 |
| ENSMUSG00000055239 | GSMG0033186 | Kcmf1 | potassium channel modulatory factor 1 | 1102.62 | 981.74 | 0.17 | 5.53E-01 |
| NULL | GSMG0033189 | NULL | NULL | 469.28 | 527.38 | -0.17 | 5.69E-01 |
| ENSMUSG00000063063 | GSMG0033200 | Ctnna2 | catenin (cadherin associated protein), alpha 2 | 570.93 | 915.85 | -0.68 | 1.73E-04 |
| NULL | GSMG0033204 | NULL | NULL | 399.97 | 550.99 | -0.47 | 8.34E-02 |
| ENSMUSG00000108077 | GSMG0033209 | 6330415B21Rik | RIKEN cDNA 6330415B21 gene | 360.14 | 508.72 | -0.50 | 1.88E-02 |
| NULL | GSMG0033211 | NULL | NULL | 740.74 | 972.48 | -0.39 | 4.42E-02 |
| NULL | GSMG0033215 | NULL | NULL | 145.48 | 142.83 | 0.03 | 9.58E-01 |
| NULL | GSMG0033221 | NULL | NULL | 192.50 | 194.63 | -0.01 | 9.82E-01 |
| ENSMUSG00000030045 | GSMG0033224 | Mrpl19 | mitochondrial ribosomal protein L19 | 551.90 | 795.06 | -0.53 | 1.96E-02 |
| ENSMUSG00000030042 | GSMG0033228 | Pole4 | polymerase (DNA-directed), epsilon 4 (p12 subunit) | 1096.76 | 1361.12 | -0.31 | 1.60E-01 |
| ENSMUSG00000000627 | GSMG0033231 | Sema4f | sema domain, immunoglobulin domain (Ig), TM domain, and short cyto | 189.65 | 257.55 | -0.44 | 2.10E-01 |
| ENSMUSG00000068329 | GSMG0033233 | Htra2 | HtrA serine peptidase 2 | 740.80 | 805.71 | -0.12 | 6.44E-01 |
| ENSMUSG00000030035 | GSMG0033237 | Wbp1 | WW domain binding protein 1 | 461.88 | 446.10 | 0.05 | 8.77E-01 |
| ENSMUSG00000030034 | GSMG0033238 | Ino80b | INO80 complex subunit B | 270.39 | 306.09 | -0.18 | 5.45E-01 |
| ENSMUSG00000030032 | GSMG0033239 | Wdr54 | WD repeat domain 54 | 245.79 | 233.77 | 0.07 | 8.57E-01 |
| ENSMUSG00000005667 | GSMG0033242 | Mthfd2 | methylenetetrahydrofolate dehydrogenase (NAD+ dependent), metheny | 111.86 | 225.50 | -1.01 | 2.70E-04 |
| NULL | GSMG0033244 | NULL | NULL | 155.66 | 135.61 | 0.20 | 5.97E-01 |
| NULL | GSMG0033245 | NULL | NULL | 151.27 | 133.53 | 0.18 | 6.34E-01 |
| NULL | GSMG0033248 | NULL | NULL | 404.58 | 374.60 | 0.11 | 7.97E-01 |
| NULL | GSMG0033250 | NULL | NULL | 738.37 | 586.67 | 0.33 | 3.58E-01 |
| ENSMUSG00000014554 | GSMG0033251 | Dguok | deoxyguanosine kinase | 543.10 | 446.92 | 0.28 | 1.98E-01 |
| ENSMUSG00000059430 | GSMG0033252 | Actg2 | actin, gamma 2, smooth muscle, enteric | 107.34 | 77.66 | 0.47 | 4.43E-01 |
| ENSMUSG00000006906 | GSMG0033253 | Stambp | STAM binding protein | 511.25 | 662.19 | -0.37 | 5.82E-02 |
| ENSMUSG00000014748 | GSMG0033258 | Tex261 | testis expressed gene 261 | 1806.09 | 2254.74 | -0.32 | 6.73E-02 |
| ENSMUSG00000045896 | GSMG0033260 | Paip2b | poly(A) binding protein interacting protein 2B | 538.78 | 684.57 | -0.35 | 1.25E-01 |
| ENSMUSG00000033769 | GSMG0033266 | Exoc6b | exocyst complex component 6B | 1661.45 | 1722.22 | -0.05 | 8.48E-01 |
| ENSMUSG00000033735 | GSMG0033277 | Spr | sepiapterin reductase | 739.30 | 761.30 | -0.04 | 9.03E-01 |
| ENSMUSG00000033720 | GSMG0033280 | Sfxn5 | sideroflexin 5 | 242.39 | 159.52 | 0.61 | 1.33E-02 |
| NULL | GSMG0033281 | NULL | NULL | 239.68 | 120.34 | 0.99 | 4.69E-04 |
| ENSMUSG00000051343 | GSMG0033283 | Rab11fip5 | RAB11 family interacting protein 5 (class I) | 275.80 | 182.98 | 0.59 | 7.94E-02 |

| | | | | | | | |
|---|---|---|---|---|---|---|---|
| ENSMUSG00000030008 | GSMG0033285 | Pradc1 | protease-associated domain containing 1 | 646.29 | 412.78 | 0.65 | 7.61E-04 |
| ENSMUSG00000047013 | GSMG0033286 | Fbxo41 | F-box protein 41 | 375.15 | 452.51 | -0.27 | 4.05E-01 |
| ENSMUSG00000071341 | GSMG0033287 | Egr4 | early growth response 4 | 369.82 | 53.81 | 2.78 | 1.38E-18 |
| ENSMUSG00000068299 | GSMG0033294 | Nat8f4 | N-acetyltransferase 8 (GCN5-related) family member 4 | 129.95 | 44.35 | 1.55 | 5.93E-06 |
| ENSMUSG00000057103 | GSMG0033295 | Nat8f1 | N-acetyltransferase 8 (GCN5-related) family member 1 | 428.10 | 252.90 | 0.76 | 8.97E-05 |
| ENSMUSG00000030002 | GSMG0033297 | Dusp11 | dual specificity phosphatase 11 (RNA/RNP complex 1-interacting) | 2637.99 | 2587.76 | 0.03 | 9.20E-01 |
| ENSMUSG00000029998 | GSMG0033300 | Pcyox1 | prenylcysteine oxidase 1 | 3689.71 | 4266.74 | -0.21 | 2.30E-01 |
| ENSMUSG00000051695 | GSMG0033305 | Pcbp1 | poly(rC) binding protein 1 | 5820.12 | 6146.67 | -0.08 | 7.41E-01 |
| ENSMUSG00000001156 | GSMG0033306 | Mxd1 | MAX dimerization protein 1 | 959.07 | 737.43 | 0.38 | 5.34E-02 |
| ENSMUSG00000001158 | GSMG0033308 | Snrnp27 | small nuclear ribonucleoprotein 27 (U4/U6.U5) | 769.24 | 1000.49 | -0.38 | 1.96E-01 |
| ENSMUSG00000001157 | GSMG0033309 | Gmcl1 | germ cell-less, spermatogenesis associated 1 | 1690.78 | 1539.85 | 0.13 | 5.54E-01 |
| ENSMUSG00000029994 | GSMG0033310 | Anxa4 | annexin A4 | 777.87 | 536.89 | 0.54 | 5.94E-03 |
| ENSMUSG00000100164 | GSMG0033312 | 2610306M01Rik | RIKEN cDNA 2610306M01 gene | 246.14 | 212.26 | 0.21 | 5.09E-01 |
| ENSMUSG00000033420 | GSMG0033314 | Antxr1 | anthrax toxin receptor 1 | 130.33 | 140.59 | -0.11 | 8.07E-01 |
| ENSMUSG00000030048 | GSMG0033321 | Gkn3 | gastrokine 3 | 1130.68 | 173.95 | 2.70 | 1.19E-26 |
| ENSMUSG00000030047 | GSMG0033322 | Arhgap25 | Rho GTPase activating protein 25 | 844.60 | 856.59 | -0.02 | 9.53E-01 |
| ENSMUSG00000030051 | GSMG0033324 | Aplf | aprataxin and PNKP like factor | 57.74 | 65.31 | -0.17 | 7.66E-01 |
| ENSMUSG00000030055 | GSMG0033326 | Rab43 | RAB43, member RAS oncogene family | 1671.43 | 1358.78 | 0.30 | 2.77E-01 |
| ENSMUSG00000030056 | GSMG0033329 | Isy1 | ISY1 splicing factor homolog | 373.14 | 526.02 | -0.49 | 7.16E-02 |
| ENSMUSG00000030057 | GSMG0033331 | Cnbp | cellular nucleic acid binding protein | 9332.29 | 10789.62 | -0.21 | 3.66E-01 |
| ENSMUSG00000079477 | GSMG0033334 | Rab7 | RAB7, member RAS oncogene family | 1043.02 | 1111.74 | -0.09 | 7.21E-01 |
| NULL | GSMG0033336 | NULL | NULL | 320.89 | 549.99 | -0.78 | 3.20E-01 |
| ENSMUSG00000033216 | GSMG0033339 | Eefsec | eukaryotic elongation factor, selenocysteine-tRNA-specific | 530.87 | 709.70 | -0.42 | 5.17E-02 |
| ENSMUSG00000030082 | GSMG0033342 | Sec61a1 | Sec61 alpha 1 subunit (S. cerevisiae) | 7265.43 | 7915.60 | -0.12 | 5.53E-01 |
| ENSMUSG00000030083 | GSMG0033347 | Abtb1 | ankyrin repeat and BTB (POZ) domain containing 1 | 1555.25 | 1103.43 | 0.49 | 9.72E-03 |
| ENSMUSG00000033152 | GSMG0033348 | Podxl2 | podocalyxin-like 2 | 3813.29 | 7836.03 | -1.04 | 1.49E-12 |
| ENSMUSG00000002870 | GSMG0033349 | Mcm2 | minichromosome maintenance complex component 2 | 839.18 | 2024.72 | -1.27 | 3.25E-11 |
| ENSMUSG00000030084 | GSMG0033353 | Plxna1 | plexin A1 | 968.83 | 1470.34 | -0.60 | 3.96E-02 |
| ENSMUSG00000030086 | GSMG0033354 | Chchd6 | coiled-coil-helix-coiled-coil-helix domain containing 6 | 464.79 | 691.31 | -0.57 | 3.78E-03 |
| NULL | GSMG0033368 | NULL | NULL | 1099.27 | 792.35 | 0.47 | 4.16E-02 |
| ENSMUSG00000030091 | GSMG0033369 | Nup210 | nucleoporin 210 | 1044.45 | 2083.85 | -1.00 | 1.23E-05 |
| ENSMUSG00000034203 | GSMG0033373 | Chchd4 | coiled-coil-helix-coiled-coil-helix domain containing 4 | 469.42 | 575.52 | -0.29 | 1.91E-01 |
| ENSMUSG00000030094 | GSMG0033374 | Xpc | xeroderma pigmentosum, complementation group C | 1509.74 | 1536.89 | -0.03 | 9.33E-01 |

| | | | | | | | |
|---|---|---|---|---|---|---|---|
| ENSMUSG00000030098 | GSMG0033378 | Grip2 | glutamate receptor interacting protein 2 | 210.68 | 493.83 | -1.23 | 4.02E-08 |
| ENSMUSG00000014551 | GSMG0033386 | Mrps25 | mitochondrial ribosomal protein S25 | 1045.56 | 1107.82 | -0.08 | 7.39E-01 |
| ENSMUSG00000014550 | GSMG0033387 | Rbsn | rabenosyn, RAB effector | 614.00 | 627.06 | -0.03 | 9.29E-01 |
| ENSMUSG00000030020 | GSMG0033389 | Prickle2 | prickle planar cell polarity protein 2 | 1277.62 | 1021.63 | 0.32 | 1.36E-01 |
| NULL | GSMG0033393 | NULL | NULL | 355.56 | 366.97 | -0.05 | 9.12E-01 |
| NULL | GSMG0033394 | NULL | NULL | 62.04 | 66.91 | -0.10 | 9.09E-01 |
| NULL | GSMG0033395 | NULL | NULL | 783.14 | 803.19 | -0.04 | 9.10E-01 |
| NULL | GSMG0033396 | NULL | NULL | 1047.31 | 1276.77 | -0.29 | 1.72E-01 |
| NULL | GSMG0033397 | NULL | NULL | 520.55 | 570.43 | -0.13 | 6.77E-01 |
| ENSMUSG00000030022 | GSMG0033399 | Adamts9 | a disintegrin-like and metallopeptidase (reprolysin type) with thrombos | 183.36 | 173.99 | 0.08 | 8.56E-01 |
| NULL | GSMG0033400 | NULL | NULL | 161.50 | 109.38 | 0.57 | 7.61E-02 |
| ENSMUSG00000045095 | GSMG0033403 | Magi1 | membrane associated guanylate kinase, WW and PDZ domain containi | 1557.10 | 2002.33 | -0.36 | 6.61E-02 |
| NULL | GSMG0033406 | NULL | NULL | 408.21 | 485.78 | -0.25 | 4.41E-01 |
| ENSMUSG00000030029 | GSMG0033413 | Lrig1 | leucine-rich repeats and immunoglobulin-like domains 1 | 1274.82 | 1146.93 | 0.15 | 6.21E-01 |
| ENSMUSG00000035245 | GSMG0033430 | Eogt | EGF domain-specific O-linked N-acetylglucosamine (GlcNAc) transfer | 22615.40 | 13557.67 | 0.74 | 1.74E-08 |
| ENSMUSG00000030059 | GSMG0033432 | Tmf1 | TATA element modulatory factor 1 | 1514.78 | 1125.07 | 0.43 | 2.70E-02 |
| ENSMUSG00000030061 | GSMG0033433 | Uba3 | ubiquitin-like modifier activating enzyme 3 | 802.45 | 795.46 | 0.01 | 9.69E-01 |
| ENSMUSG00000030064 | GSMG0033436 | Frmd4b | FERM domain containing 4B | 2770.77 | 2444.47 | 0.18 | 3.88E-01 |
| NULL | GSMG0033437 | NULL | NULL | 231.64 | 183.25 | 0.34 | 3.12E-01 |
| G00000030067,ENSMUSG000 | GSMG0033446 | Foxp1 // Eif4e3 | forkhead box P1 // eukaryotic translation initiation factor 4E member 3 | 2002.85 | 2098.43 | -0.07 | 7.90E-01 |
| NULL | GSMG0033447 | NULL | NULL | 395.13 | 452.36 | -0.19 | 5.11E-01 |
| NULL | GSMG0033449 | NULL | NULL | 55.57 | 61.43 | -0.15 | 8.04E-01 |
| NULL | GSMG0033451 | NULL | NULL | 151.97 | 160.07 | -0.08 | 8.66E-01 |
| NULL | GSMG0033458 | NULL | NULL | 260.63 | 332.21 | -0.35 | 1.78E-01 |
| NULL | GSMG0033463 | NULL | NULL | 104.21 | 101.87 | 0.03 | 9.57E-01 |
| NULL | GSMG0033466 | NULL | NULL | 324.10 | 305.69 | 0.09 | 8.34E-01 |
| NULL | GSMG0033470 | NULL | NULL | 422.20 | 601.94 | -0.51 | 1.22E-02 |
| ENSMUSG00000035378 | GSMG0033483 | Shq1 | SHQ1 homolog (S. cerevisiae) | 89.65 | 142.42 | -0.67 | 2.68E-02 |
| ENSMUSG00000035357 | GSMG0033486 | Pdzrn3 | PDZ domain containing RING finger 3 | 168.48 | 172.13 | -0.03 | 9.50E-01 |
| NULL | GSMG0033487 | NULL | NULL | 485.09 | 414.41 | 0.23 | 3.78E-01 |
| ENSMUSG00000030075 | GSMG0033492 | Cntn3 | contactin 3 | 368.10 | 456.39 | -0.31 | 2.88E-01 |
| NULL | GSMG0033495 | NULL | NULL | 159.35 | 132.85 | 0.26 | 5.29E-01 |
| ENSMUSG00000005362 | GSMG0033500 | Crbn | cereblon | 1021.59 | 901.67 | 0.18 | 4.12E-01 |

| | | | | | | | |
|---|---|---|---|---|---|---|---|
| ENSMUSG00000030101 | GSMG0033501 | Sumf1 | sulfatase modifying factor 1 | 666.30 | 551.70 | 0.27 | 2.08E-01 |
| NULL | GSMG0033508 | NULL | NULL | 303.66 | 292.98 | 0.05 | 8.97E-01 |
| NULL | GSMG0033509 | NULL | NULL | 422.22 | 418.97 | 0.01 | 9.72E-01 |
| ENSMUSG00000030254 | GSMG0033517 | Rad18 | RAD18 E3 ubiquitin protein ligase | 235.41 | 458.27 | -0.96 | 1.73E-06 |
| ENSMUSG00000030257 | GSMG0033518 | Srgap3 | SLIT-ROBO Rho GTPase activating protein 3 | 304.62 | 419.00 | -0.46 | 3.32E-02 |
| NULL | GSMG0033519 | NULL | NULL | 121.63 | 142.73 | -0.24 | 6.72E-01 |
| NULL | GSMG0033520 | NULL | NULL | 187.51 | 238.17 | -0.35 | 2.36E-01 |
| NULL | GSMG0033521 | NULL | NULL | 687.42 | 734.09 | -0.10 | 8.06E-01 |
| ENSMUSG00000086429 | GSMG0033524 | Gt(ROSA)26Sor | gene trap ROSA 26, Philippe Soriano | 119.32 | 113.53 | 0.08 | 8.81E-01 |
| ENSMUSG00000042873 | GSMG0033525 | Lhfpl4 | lipoma HMGIC fusion partner-like protein 4 | 108.94 | 231.87 | -1.09 | 8.99E-05 |
| ENSMUSG00000030272 | GSMG0033530 | Camk1 | calcium/calmodulin-dependent protein kinase I | 2673.19 | 1869.09 | 0.52 | 1.31E-03 |
| ENSMUSG00000048930 | GSMG0033531 | Tada3 | transcriptional adaptor 3 | 847.40 | 862.19 | -0.02 | 9.37E-01 |
| ENSMUSG00000051169 | GSMG0033532 | Rpusd3 | RNA pseudouridylate synthase domain containing 3 | 267.15 | 292.64 | -0.13 | 6.78E-01 |
| ENSMUSG00000045009 | GSMG0033535 | Prrt3 | proline-rich transmembrane protein 3 | 542.01 | 326.16 | 0.73 | 2.00E-04 |
| ENSMUSG00000030286 | GSMG0033536 | Emc3 | ER membrane protein complex subunit 3 | 5098.38 | 5295.58 | -0.05 | 8.49E-01 |
| ENSMUSG00000030298 | GSMG0033541 | Sec13 | SEC13 homolog, nuclear pore and COPII coat complex component | 2824.49 | 3022.92 | -0.10 | 6.84E-01 |
| ENSMUSG00000030302 | GSMG0033542 | Atp2b2 | ATPase, Ca++ transporting, plasma membrane 2 | 976.90 | 829.84 | 0.24 | 5.75E-01 |
| ENSMUSG00000030315 | GSMG0033544 | Vgll4 | vestigial like family member 4 | 571.21 | 648.27 | -0.18 | 4.56E-01 |
| NULL | GSMG0033545 | NULL | NULL | 688.09 | 774.04 | -0.17 | 5.55E-01 |
| ENSMUSG00000030316 | GSMG0033548 | Tamm41 | TAM41 mitochondrial translocator assembly and maintenance homolog | 282.69 | 323.46 | -0.19 | 5.29E-01 |
| ENSMUSG00000030317 | GSMG0033550 | Timp4 | tissue inhibitor of metalloproteinase 4 | 908.73 | 526.51 | 0.79 | 1.07E-05 |
| NULL | GSMG0033551 | NULL | NULL | 500.40 | 278.25 | 0.85 | 6.89E-06 |
| ENSMUSG00000000441 | GSMG0033554 | Raf1 | v-raf-leukemia viral oncogene 1 | 2892.26 | 3264.53 | -0.17 | 4.20E-01 |
| ENSMUSG00000057841 | GSMG0033556 | Rpl32 | ribosomal protein L32 | 2572.36 | 4018.53 | -0.64 | 1.58E-03 |
| ENSMUSG00000030322 | GSMG0033558 | Mbd4 | methyl-CpG binding domain protein 4 | 221.18 | 228.31 | -0.05 | 9.19E-01 |
| ENSMUSG00000030123 | GSMG0033560 | Plxnd1 | plexin D1 | 1821.15 | 2240.31 | -0.30 | 4.01E-01 |
| ENSMUSG00000030126 | GSMG0033562 | Tmcc1 | transmembrane and coiled coil domains 1 | 193.29 | 215.72 | -0.16 | 6.58E-01 |
| NULL | GSMG0033566 | NULL | NULL | 261.44 | 458.86 | -0.81 | 2.82E-04 |
| NULL | GSMG0033567 | NULL | NULL | 45.87 | 51.23 | -0.16 | 8.09E-01 |
| ENSMUSG00000059878 | GSMG0033571 | Zfp422 | zinc finger protein 422 | 2720.54 | 3215.03 | -0.24 | 3.46E-01 |
| ENSMUSG00000042129 | GSMG0033572 | Rassf4 | Ras association (RalGDS/AF-6) domain family member 4 | 133.06 | 103.85 | 0.36 | 3.03E-01 |
| NULL | GSMG0033579 | NULL | NULL | 368.60 | 431.31 | -0.23 | 5.14E-01 |
| ENSMUSG00000042042 | GSMG0033581 | Csgalnact2 | chondroitin sulfate N-acetylgalactosaminyltransferase 2 | 650.73 | 541.86 | 0.26 | 2.62E-01 |

| | | | | | | | |
|---|---|---|---|---|---|---|---|
| ENSMUSG00000030110 | GSMG0033582 | Ret | ret proto-oncogene | 1451.90 | 947.65 | 0.62 | 1.00E-03 |
| ENSMUSG00000030138 | GSMG0033584 | Bms1 | BMS1, ribosome biogenesis factor | 1365.89 | 1551.57 | -0.18 | 3.79E-01 |
| ENSMUSG00000030145 | GSMG0033585 | Zfp248 | zinc finger protein 248 | 79.66 | 87.56 | -0.14 | 7.91E-01 |
| ENSMUSG00000007827 | GSMG0033587 | Ankrd26 | ankyrin repeat domain 26 | 122.19 | 148.72 | -0.28 | 4.87E-01 |
| ENSMUSG00000051331 | GSMG0033589 | Cacna1c | calcium channel, voltage-dependent, L type, alpha 1C subunit | 939.91 | 922.42 | 0.03 | 9.42E-01 |
| NULL | GSMG0033593 | NULL | NULL | 252.46 | 222.84 | 0.18 | 6.83E-01 |
| NULL | GSMG0033594 | NULL | NULL | 1036.06 | 896.90 | 0.21 | 3.28E-01 |
| NULL | GSMG0033595 | NULL | NULL | 214.57 | 249.82 | -0.22 | 4.85E-01 |
| NULL | GSMG0033597 | NULL | NULL | 746.33 | 661.26 | 0.17 | 5.54E-01 |
| NULL | GSMG0033599 | NULL | NULL | 136.01 | 106.44 | 0.36 | 3.92E-01 |
| ENSMUSG00000055003 | GSMG0033600 | Lrtm2 | leucine-rich repeats and transmembrane domains 2 | 498.51 | 418.38 | 0.25 | 3.90E-01 |
| ENSMUSG00000030168 | GSMG0033601 | Adipor2 | adiponectin receptor 2 | 258.75 | 179.67 | 0.53 | 4.86E-02 |
| ENSMUSG00000030170 | GSMG0033603 | Wnt5b | wingless-type MMTV integration site family, member 5B | 203.73 | 87.03 | 1.23 | 8.94E-07 |
| ENSMUSG00000030172 | GSMG0033605 | Erc1 | ELKS/RAB6-interacting/CAST family member 1 | 1021.86 | 1217.30 | -0.25 | 1.80E-01 |
| ENSMUSG00000045962 | GSMG0033615 | Wnk1 | WNK lysine deficient protein kinase 1 | 592.39 | 548.47 | 0.11 | 7.27E-01 |
| ENSMUSG00000030177 | GSMG0033623 | Ccdc77 | coiled-coil domain containing 77 | 155.03 | 175.93 | -0.19 | 5.98E-01 |
| NULL | GSMG0033625 | NULL | NULL | 802.65 | 338.78 | 1.24 | 4.47E-07 |
| ENSMUSG00000058979 | GSMG0033626 | Cecr5 | cat eye syndrome chromosome region, candidate 5 | 1180.03 | 1006.46 | 0.23 | 3.08E-01 |
| ENSMUSG00000019210 | GSMG0033628 | Atp6v1e1 | ATPase, H+ transporting, lysosomal V1 subunit E1 | 1998.82 | 1900.17 | 0.07 | 7.83E-01 |
| ENSMUSG00000004446 | GSMG0033630 | Bid | BH3 interacting domain death agonist | 179.60 | 314.76 | -0.80 | 5.50E-04 |
| ENSMUSG00000051586 | GSMG0033632 | Mical3 | microtubule associated monooxygenase, calponin and LIM domain con | 1051.96 | 763.83 | 0.46 | 7.53E-02 |
| NULL | GSMG0033633 | NULL | NULL | 278.14 | 239.44 | 0.22 | 5.24E-01 |
| ENSMUSG00000040797 | GSMG0033638 | Iqsec3 | IQ motif and Sec7 domain 3 | 321.07 | 194.10 | 0.73 | 1.03E-02 |
| ENSMUSG00000040669 | GSMG0033640 | Phc1 | polyhomeotic-like 1 (Drosophila) | 1616.49 | 1501.49 | 0.11 | 7.26E-01 |
| ENSMUSG00000040649 | GSMG0033641 | Rimklb | ribosomal modification protein rimK-like family member B | 113.50 | 394.34 | -1.79 | 1.86E-16 |
| ENSMUSG00000003153 | GSMG0033645 | Slc2a3 | solute carrier family 2 (facilitated glucose transporter), member 3 | 1570.15 | 1021.98 | 0.62 | 4.36E-04 |
| ENSMUSG00000005069 | GSMG0033656 | Pex5 | peroxisomal biogenesis factor 5 | 1048.80 | 1034.37 | 0.02 | 9.46E-01 |
| ENSMUSG00000008153 | GSMG0033657 | Clstn3 | calsyntenin 3 | 2343.18 | 2197.26 | 0.09 | 7.97E-01 |
| ENSMUSG00000004268 | GSMG0033660 | Emg1 | EMG1 N1-specific pseudouridine methyltransferase | 1116.23 | 1443.59 | -0.37 | 4.03E-02 |
| ENSMUSG00000004266 | GSMG0033661 | Ptpn6 | protein tyrosine phosphatase, non-receptor type 6 | 222.36 | 200.72 | 0.15 | 6.91E-01 |
| G00000004263,ENSMUSG000 | GSMG0033662 | Grcc10 // Atn1 | gene rich cluster, C10 gene // atrophin 1 | 3259.81 | 4104.22 | -0.33 | 7.44E-02 |
| ENSMUSG00000004267 | GSMG0033663 | Eno2 | enolase 2, gamma neuronal | 2639.10 | 1500.80 | 0.81 | 9.62E-07 |
| ENSMUSG00000023456 | GSMG0033665 | Tpi1 | triosephosphate isomerase 1 | 2546.09 | 2209.35 | 0.21 | 3.85E-01 |

| | | | | | | | |
|---|---|---|---|---|---|---|---|
| ENSMUSG00000038429 | GSMG0033666 | Usp5 | ubiquitin specific peptidase 5 (isopeptidase T) | 4073.01 | 5118.04 | -0.33 | 8.51E-02 |
| ENSMUSG00000023191 | GSMG0033668 | P3h3 | prolyl 3-hydroxylase 3 | 1121.87 | 1117.65 | 0.01 | 9.87E-01 |
| ENSMUSG00000038390 | GSMG0033669 | Gpr162 | G protein-coupled receptor 162 | 780.91 | 597.40 | 0.38 | 2.85E-01 |
| ENSMUSG00000030122 | GSMG0033672 | Ptms | parathymosin | 1362.39 | 1807.04 | -0.41 | 3.94E-02 |
| ENSMUSG00000030127 | GSMG0033673 | Cops7a | COP9 signalosome subunit 7A | 1689.60 | 1906.31 | -0.17 | 3.73E-01 |
| ENSMUSG00000038252 | GSMG0033679 | Ncapd2 | non-SMC condensin I complex, subunit D2 | 1326.70 | 1769.80 | -0.41 | 1.81E-02 |
| ENSMUSG00000038213 | GSMG0033680 | Tapbpl | TAP binding protein-like | 370.62 | 298.76 | 0.31 | 2.85E-01 |
| ENSMUSG00000030339 | GSMG0033684 | Ltbr | lymphotoxin B receptor | 907.71 | 1038.70 | -0.20 | 3.44E-01 |
| ENSMUSG00000030342 | GSMG0033686 | Cd9 | CD9 antigen | 2340.45 | 2236.05 | 0.07 | 8.01E-01 |
| ENSMUSG00000049107 | GSMG0033691 | Ntf3 | neurotrophin 3 | 24.88 | 130.92 | -2.39 | 1.23E-11 |
| ENSMUSG00000047976 | GSMG0033698 | Kcna1 | potassium voltage-gated channel, shaker-related subfamily, member 1 | 1851.77 | 312.51 | 2.57 | 5.01E-37 |
| ENSMUSG00000038077 | GSMG0033699 | Kcna6 | potassium voltage-gated channel, shaker-related, subfamily, member 6 | 756.07 | 1454.79 | -0.95 | 7.55E-08 |
| ENSMUSG00000000399 | GSMG0033700 | Ndufa9 | NADH dehydrogenase (ubiquinone) 1 alpha subcomplex, 9 | 2237.89 | 2875.38 | -0.36 | 4.24E-02 |
| ENSMUSG00000030346 | GSMG0033702 | Rad51ap1 | RAD51 associated protein 1 | 184.00 | 381.38 | -1.05 | 1.82E-06 |
| ENSMUSG00000000184 | GSMG0033704 | Ccnd2 | cyclin D2 | 1626.84 | 1717.30 | -0.08 | 7.93E-01 |
| ENSMUSG00000030350 | GSMG0033707 | Prmt8 | protein arginine N-methyltransferase 8 | 196.72 | 199.63 | -0.02 | 9.64E-01 |
| ENSMUSG00000030352 | GSMG0033714 | Tspan9 | tetraspanin 9 | 843.58 | 1025.37 | -0.28 | 1.55E-01 |
| ENSMUSG00000030353 | GSMG0033722 | Tead4 | TEA domain family member 4 | 98.86 | 90.28 | 0.14 | 8.08E-01 |
| ENSMUSG00000048668 | GSMG0033725 | Rhno1 | RAD9-HUS1-RAD1 interacting nuclear orphan 1 | 488.01 | 749.82 | -0.62 | 2.33E-03 |
| ENSMUSG00000001518 | GSMG0033726 | Itfg2 | integrin alpha FG-GAP repeat containing 2 | 383.51 | 425.72 | -0.15 | 5.83E-01 |
| ENSMUSG00000030357 | GSMG0033727 | Fkbp4 | FK506 binding protein 4 | 3186.08 | 3367.83 | -0.08 | 7.55E-01 |
| ENSMUSG00000033082 | GSMG0033745 | Clec1a | C-type lectin domain family 1, member a | 218.67 | 295.65 | -0.43 | 1.15E-01 |
| ENSMUSG00000030188 | GSMG0033771 | Magohb | mago homolog B, exon junction complex core component | 141.28 | 213.75 | -0.59 | 3.48E-02 |
| ENSMUSG00000030189 | GSMG0033774 | Ybx3 | Y box protein 3 | 3015.56 | 2438.38 | 0.31 | 3.61E-01 |
| NULL | GSMG0033791 | NULL | NULL | 94.49 | 89.26 | 0.08 | 8.82E-01 |
| ENSMUSG00000030201 | GSMG0033799 | Lrp6 | low density lipoprotein receptor-related protein 6 | 712.36 | 759.55 | -0.09 | 8.30E-01 |
| ENSMUSG00000032718 | GSMG0033803 | Mansc1 | MANSC domain containing 1 | 314.43 | 288.51 | 0.13 | 7.13E-01 |
| ENSMUSG00000030203 | GSMG0033804 | Dusp16 | dual specificity phosphatase 16 | 115.33 | 93.15 | 0.31 | 4.10E-01 |
| ENSMUSG00000032641 | GSMG0033806 | Gpr19 | G protein-coupled receptor 19 | 563.79 | 476.24 | 0.24 | 2.77E-01 |
| ENSMUSG00000042770 | GSMG0033809 | Hebp1 | heme binding protein 1 | 852.99 | 686.91 | 0.31 | 2.23E-01 |
| NULL | GSMG0033813 | NULL | NULL | 148.83 | 181.34 | -0.28 | 5.34E-01 |
| ENSMUSG00000030209 | GSMG0033814 | Grin2b | glutamate receptor, ionotropic, NMDA2B (epsilon 2) | 145.88 | 153.25 | -0.06 | 9.19E-01 |
| NULL | GSMG0033815 | NULL | NULL | 159.54 | 204.56 | -0.35 | 3.65E-01 |

| | | | | | | | |
|---|---|---|---|---|---|---|---|
| NULL | GSMG0033816 | NULL | NULL | 414.86 | 436.27 | -0.08 | 8.34E-01 |
| ENSMUSG00000030216 | GSMG0033822 | Wbp11 | WW domain binding protein 11 | 524.48 | 633.93 | -0.28 | 2.38E-01 |
| ENSMUSG00000030218 | GSMG0033825 | Mgp | matrix Gla protein | 3738.80 | 826.03 | 2.18 | 7.61E-51 |
| ENSMUSG00000030220 | GSMG0033827 | Arhgdib | Rho, GDP dissociation inhibitor (GDI) beta | 2455.73 | 2211.57 | 0.15 | 5.79E-01 |
| ENSMUSG00000030222 | GSMG0033828 | Rerg | RAS-like, estrogen-regulated, growth-inhibitor | 223.73 | 299.56 | -0.42 | 7.77E-02 |
| ENSMUSG00000015766 | GSMG0033831 | Eps8 | epidermal growth factor receptor pathway substrate 8 | 606.25 | 745.31 | -0.30 | 1.57E-01 |
| ENSMUSG00000030226 | GSMG0033836 | Lmo3 | LIM domain only 3 | 506.83 | 1026.68 | -1.02 | 1.61E-06 |
| ENSMUSG00000030237 | GSMG0033846 | Slco1a4 | solute carrier organic anion transporter family, member 1a4 | 18971.56 | 3566.88 | 2.41 | 1.03E-62 |
| ENSMUSG00000030243 | GSMG0033850 | Recql | RecQ protein-like | 480.34 | 526.29 | -0.13 | 6.69E-01 |
| ENSMUSG00000030247 | GSMG0033854 | Kcnj8 | potassium inwardly-rectifying channel, subfamily J, member 8 | 1753.16 | 2219.89 | -0.34 | 6.57E-02 |
| ENSMUSG00000030249 | GSMG0033855 | Abcc9 | ATP-binding cassette, sub-family C (CFTR/MRP), member 9 | 2204.11 | 3439.74 | -0.64 | 1.09E-03 |
| ENSMUSG00000030279 | GSMG0033860 | C2cd5 | C2 calcium-dependent domain containing 5 | 1156.40 | 1249.60 | -0.11 | 6.40E-01 |
| ENSMUSG00000041540 | GSMG0033867 | Sox5 | SRY (sex determining region Y)-box 5 | 131.60 | 220.26 | -0.74 | 2.08E-03 |
| NULL | GSMG0033869 | NULL | NULL | 138.01 | 229.02 | -0.73 | 1.59E-02 |
| ENSMUSG00000030268 | GSMG0033877 | Bcat1 | branched chain aminotransferase 1, cytosolic | 133.93 | 226.15 | -0.75 | 2.19E-03 |
| ENSMUSG00000030265 | GSMG0033880 | Kras | Kirsten rat sarcoma viral oncogene homolog | 3955.38 | 3870.24 | 0.03 | 9.34E-01 |
| NULL | GSMG0033881 | NULL | NULL | 262.35 | 224.58 | 0.23 | 5.34E-01 |
| ENSMUSG00000054966 | GSMG0033882 | Lmntd1 | lamin tail domain containing 1 | 103.56 | 82.08 | 0.33 | 4.80E-01 |
| ENSMUSG00000030256 | GSMG0033886 | Bhlhe41 | basic helix-loop-helix family, member e41 | 483.26 | 664.95 | -0.46 | 1.08E-01 |
| ENSMUSG00000030287 | GSMG0033889 | Itpr2 | inositol 1,4,5-triphosphate receptor 2 | 711.88 | 576.41 | 0.30 | 1.30E-01 |
| ENSMUSG00000040250 | GSMG0033893 | Asun | asunder, spermatogenesis regulator | 1548.02 | 1497.73 | 0.05 | 8.79E-01 |
| ENSMUSG00000040234 | GSMG0033894 | Tm7sf3 | transmembrane 7 superfamily member 3 | 1652.30 | 1233.28 | 0.42 | 1.22E-02 |
| NULL | GSMG0033895 | NULL | NULL | 174.49 | 118.86 | 0.55 | 5.60E-02 |
| ENSMUSG00000048776 | GSMG0033900 | Pthlh | parathyroid hormone-like peptide | 68.20 | 37.03 | 0.89 | 7.36E-02 |
| ENSMUSG00000030304 | GSMG0033905 | Ergic2 | ERGIC and golgi 2 | 1459.75 | 1623.97 | -0.15 | 5.02E-01 |
| NULL | GSMG0033906 | NULL | NULL | 219.07 | 234.06 | -0.10 | 7.97E-01 |
| ENSMUSG00000030306 | GSMG0033908 | Tmtc1 | transmembrane and tetratricopeptide repeat containing 1 | 657.36 | 334.10 | 0.98 | 1.90E-04 |
| ENSMUSG00000040029 | GSMG0033913 | Ipo8 | importin 8 | 746.63 | 836.53 | -0.16 | 4.87E-01 |
| ENSMUSG00000030309 | GSMG0033914 | Caprin2 | caprin family member 2 | 237.26 | 261.77 | -0.14 | 6.93E-01 |
| ENSMUSG00000030313 | GSMG0033920 | Dennd5b | DENN/MADD domain containing 5B | 698.60 | 697.55 | 0.00 | 9.93E-01 |
| ENSMUSG00000068250 | GSMG0033926 | Amn1 | antagonist of mitotic exit network 1 | 215.93 | 261.35 | -0.28 | 3.16E-01 |
| ENSMUSG00000068566 | GSMG0033931 | Myadm | myeloid-associated differentiation marker | 5191.15 | 4170.71 | 0.32 | 1.10E-01 |
| ENSMUSG00000078816 | GSMG0033932 | Prkcg | protein kinase C, gamma | 1261.45 | 1413.95 | -0.16 | 5.52E-01 |

| | | | | | | | |
|---|---|---|---|---|---|---|---|
| ENSMUSG00000035674 | GSMG0033937 | Ndufa3 | NADH dehydrogenase (ubiquinone) 1 alpha subcomplex, 3 | 1290.81 | 1277.69 | 0.01 | 9.75E-01 |
| ENSMUSG00000008373 | GSMG0033938 | Prpf31 | pre-mRNA processing factor 31 | 1140.59 | 1418.01 | -0.31 | 1.08E-01 |
| ENSMUSG00000035632 | GSMG0033939 | Cnot3 | CCR4-NOT transcription complex, subunit 3 | 1169.08 | 1102.09 | 0.08 | 7.90E-01 |
| ENSMUSG00000035585 | GSMG0033941 | Tsen34 | tRNA splicing endonuclease subunit 34 | 2407.66 | 3039.75 | -0.34 | 3.57E-02 |
| ENSMUSG00000006333 | GSMG0033942 | Rps9 | ribosomal protein S9 | 2839.06 | 4030.51 | -0.51 | 1.53E-02 |
| ENSMUSG00000030428 | GSMG0033944 | Ttyh1 | tweety family member 1 | 4275.91 | 2280.83 | 0.91 | 2.18E-08 |
| ENSMUSG00000035545 | GSMG0033945 | Leng8 | leukocyte receptor cluster (LRC) member 8 | 9654.29 | 7541.31 | 0.36 | 2.29E-01 |
| NULL | GSMG0033946 | NULL | NULL | 120.13 | 75.74 | 0.66 | 5.51E-02 |
| ENSMUSG00000059851 | GSMG0033956 | Suv420h2 | suppressor of variegation 4-20 homolog 2 (Drosophila) | 802.30 | 908.18 | -0.18 | 4.63E-01 |
| ENSMUSG00000035285 | GSMG0033961 | Nat14 | N-acetyltransferase 14 | 656.76 | 493.73 | 0.41 | 6.77E-02 |
| ENSMUSG00000051184 | GSMG0033964 | Zfp524 | zinc finger protein 524 | 225.92 | 184.05 | 0.29 | 3.32E-01 |
| ENSMUSG00000055633 | GSMG0033967 | Zfp580 | zinc finger protein 580 | 365.80 | 518.21 | -0.51 | 1.55E-02 |
| ENSMUSG00000035228 | GSMG0033968 | Ccdc106 | coiled-coil domain containing 106 | 578.84 | 611.84 | -0.08 | 7.94E-01 |
| ENSMUSG00000030435 | GSMG0033969 | U2af2 | U2 small nuclear ribonucleoprotein auxiliary factor (U2AF) 2 | 1039.27 | 1406.46 | -0.44 | 4.12E-02 |
| ENSMUSG00000035203 | GSMG0033970 | Epn1 | epsin 1 | 1412.64 | 1261.51 | 0.16 | 6.85E-01 |
| NULL | GSMG0033974 | NULL | NULL | 106.66 | 119.55 | -0.17 | 6.97E-01 |
| ENSMUSG00000044876 | GSMG0033976 | Zfp444 | zinc finger protein 444 | 815.41 | 826.66 | -0.02 | 9.51E-01 |
| ENSMUSG00000054893 | GSMG0033979 | Zfp667 | zinc finger protein 667 | 401.35 | 375.99 | 0.10 | 7.89E-01 |
| ENSMUSG00000062861 | GSMG0033982 | Zfp28 | zinc finger protein 28 | 70.09 | 110.36 | -0.65 | 1.04E-01 |
| ENSMUSG00000051527 | GSMG0033987 | Usp29 | ubiquitin specific peptidase 29 | 188.45 | 326.39 | -0.79 | 5.80E-04 |
| NULL | GSMG0033999 | NULL | NULL | 131.40 | 150.55 | -0.19 | 6.32E-01 |
| ENSMUSG00000030386 | GSMG0034018 | Zfp606 | zinc finger protein 606 | 705.88 | 728.23 | -0.04 | 8.89E-01 |
| ENSMUSG00000058638 | GSMG0034024 | Zfp110 | zinc finger protein 110 | 399.38 | 348.90 | 0.19 | 4.63E-01 |
| ENSMUSG00000060397 | GSMG0034026 | Zfp128 | zinc finger protein 128 | 402.37 | 450.90 | -0.16 | 6.06E-01 |
| ENSMUSG00000054715 | GSMG0034027 | Zscan22 | zinc finger and SCAN domain containing 22 | 565.39 | 499.26 | 0.18 | 4.99E-01 |
| ENSMUSG00000012848 | GSMG0034028 | Rps5 | ribosomal protein S5 | 1804.73 | 2626.05 | -0.54 | 1.55E-03 |
| ENSMUSG00000004500 | GSMG0034030 | Zfp324 | zinc finger protein 324 | 650.67 | 494.32 | 0.39 | 1.35E-01 |
| ENSMUSG00000033961 | GSMG0034031 | Zfp446 | zinc finger protein 446 | 109.31 | 96.53 | 0.18 | 7.04E-01 |
| ENSMUSG00000005566 | GSMG0034032 | Trim28 | tripartite motif-containing 28 | 3923.87 | 5069.08 | -0.37 | 5.12E-02 |
| NULL | GSMG0034033 | NULL | NULL | 1905.65 | 1615.07 | 0.24 | 2.33E-01 |
| ENSMUSG00000056394 | GSMG0034034 | Lig1 | ligase I, DNA, ATP-dependent | 695.05 | 1403.45 | -1.01 | 8.61E-11 |
| ENSMUSG00000006024 | GSMG0034045 | Napa | N-ethylmaleimide sensitive fusion protein attachment protein alpha | 2586.47 | 2272.97 | 0.19 | 3.24E-01 |
| ENSMUSG00000006021 | GSMG0034046 | Kptn | kaptin | 582.62 | 507.64 | 0.20 | 4.25E-01 |

| | | | | | | | |
|---|---|---|---|---|---|---|---|
| ENSMUSG00000041420 | GSMG0034049 | Meis3 | Meis homeobox 3 | 1036.08 | 1452.58 | -0.49 | 1.31E-02 |
| ENSMUSG00000002083 | GSMG0034050 | Bbc3 | BCL2 binding component 3 | 214.89 | 274.60 | -0.36 | 3.33E-01 |
| ENSMUSG00000059273 | GSMG0034051 | Zc3h4 | zinc finger CCCH-type containing 4 | 127.54 | 136.10 | -0.09 | 8.47E-01 |
| ENSMUSG00000019158 | GSMG0034055 | Tmem160 | transmembrane protein 160 | 1258.45 | 1145.13 | 0.13 | 6.55E-01 |
| ENSMUSG00000008036 | GSMG0034059 | Ap2s1 | adaptor-related protein complex 2, sigma 1 subunit | 2252.06 | 2725.34 | -0.28 | 1.67E-01 |
| ENSMUSG00000001918 | GSMG0034061 | Slc1a5 | solute carrier family 1 (neutral amino acid transporter), member 5 | 433.77 | 398.89 | 0.12 | 6.95E-01 |
| NULL | GSMG0034063 | NULL | NULL | 49.35 | 57.61 | -0.22 | 7.18E-01 |
| ENSMUSG00000030374 | GSMG0034064 | Strn4 | striatin, calmodulin binding protein 4 | 4611.88 | 4431.25 | 0.06 | 8.41E-01 |
| ENSMUSG00000041187 | GSMG0034065 | Prkd2 | protein kinase D2 | 1892.54 | 2080.08 | -0.14 | 5.70E-01 |
| ENSMUSG00000078794 | GSMG0034066 | Dact3 | dishevelled-binding antagonist of beta-catenin 3 | 697.86 | 361.35 | 0.95 | 1.85E-04 |
| ENSMUSG00000070802 | GSMG0034069 | Pnmal2 | PNMA-like 2 | 954.88 | 1029.06 | -0.11 | 8.10E-01 |
| ENSMUSG00000041141 | GSMG0034070 | Pnmal1 | PNMA-like 1 | 278.49 | 275.52 | 0.01 | 9.77E-01 |
| ENSMUSG00000066760 | GSMG0034074 | Psg16 | pregnancy specific glycoprotein 16 | 110.24 | 150.76 | -0.45 | 2.27E-01 |
| ENSMUSG00000040987 | GSMG0034088 | Mill2 | MHC I like leukocyte 2 | 116.33 | 118.69 | -0.02 | 9.77E-01 |
| NULL | GSMG0034089 | NULL | NULL | 83.59 | 143.04 | -0.78 | 1.31E-02 |
| ENSMUSG00000030413 | GSMG0034090 | Pglyrp1 | peptidoglycan recognition protein 1 | 10947.52 | 7229.68 | 0.60 | 1.89E-04 |
| ENSMUSG00000044030 | GSMG0034097 | Irf2bp1 | interferon regulatory factor 2 binding protein 1 | 1282.46 | 1267.18 | 0.02 | 9.64E-01 |
| ENSMUSG00000023118 | GSMG0034098 | Sympk | symplekin | 2907.34 | 2479.80 | 0.23 | 4.13E-01 |
| ENSMUSG00000030410 | GSMG0034100 | Dmwd | dystrophia myotonica-containing WD repeat motif | 1164.46 | 1159.87 | 0.01 | 9.91E-01 |
| ENSMUSG00000030409 | GSMG0034101 | Dmpk | dystrophia myotonica-protein kinase | 429.13 | 312.85 | 0.45 | 8.54E-02 |
| ENSMUSG00000050428 | GSMG0034103 | Fbxo46 | F-box protein 46 | 474.91 | 418.58 | 0.18 | 5.68E-01 |
| ENSMUSG00000040824 | GSMG0034104 | Snrpd2 | small nuclear ribonucleoprotein D2 | 778.65 | 1224.30 | -0.65 | 7.60E-04 |
| ENSMUSG00000040811 | GSMG0034105 | Eml2 | echinoderm microtubule associated protein like 2 | 1242.01 | 864.85 | 0.52 | 5.55E-01 |
| ENSMUSG00000044317 | GSMG0034106 | Gpr4 | G protein-coupled receptor 4 | 3614.56 | 3573.33 | 0.02 | 9.51E-01 |
| ENSMUSG00000052214 | GSMG0034107 | Opa3 | optic atrophy 3 | 1451.53 | 1774.03 | -0.29 | 1.39E-01 |
| ENSMUSG00000030401 | GSMG0034109 | Rtn2 | reticulon 2 (Z-band associated protein) | 356.86 | 592.91 | -0.73 | 9.72E-04 |
| ENSMUSG00000003549 | GSMG0034111 | Ercc1 | excision repair cross-complementing rodent repair deficiency, complem | 556.62 | 668.58 | -0.26 | 2.74E-01 |
| ENSMUSG00000030400 | GSMG0034113 | Ercc2 | excision repair cross-complementing rodent repair deficiency, complem | 302.55 | 356.92 | -0.24 | 4.96E-01 |
| NULL | GSMG0034115 | NULL | NULL | 146.52 | 251.54 | -0.79 | 1.65E-03 |
| ENSMUSG00000002043 | GSMG0034117 | Trappc6a | trafficking protein particle complex 6A | 500.67 | 456.25 | 0.13 | 6.28E-01 |
| ENSMUSG00000057101 | GSMG0034137 | Zfp180 | zinc finger protein 180 | 1203.81 | 1186.47 | 0.02 | 9.48E-01 |
| ENSMUSG00000047603 | GSMG0034139 | Zfp235 | zinc finger protein 235 | 875.22 | 683.09 | 0.36 | 6.01E-02 |
| ENSMUSG00000002210 | GSMG0034149 | Smg9 | smg-9 homolog, nonsense mediated mRNA decay factor (C. elegans) | 810.35 | 849.88 | -0.07 | 8.32E-01 |

| | | | | | | | |
|---|---|---|---|---|---|---|---|
| ENSMUSG00000046223 | GSMG0034150 | Plaur | plasminogen activator, urokinase receptor | 460.65 | 340.10 | 0.44 | 6.51E-02 |
| ENSMUSG00000054793 | GSMG0034151 | Cadm4 | cell adhesion molecule 4 | 725.54 | 1003.69 | -0.47 | 5.33E-01 |
| ENSMUSG00000064264 | GSMG0034152 | Zfp428 | zinc finger protein 428 | 218.49 | 430.59 | -0.98 | 2.56E-05 |
| ENSMUSG00000041037 | GSMG0034153 | Irgq | immunity-related GTPase family, Q | 92.04 | 88.10 | 0.06 | 9.36E-01 |
| ENSMUSG00000051768 | GSMG0034154 | Xrcc1 | X-ray repair complementing defective repair in Chinese hamster cells 1 | 1380.33 | 1450.36 | -0.07 | 7.90E-01 |
| ENSMUSG00000064254 | GSMG0034155 | Ethe1 | ethylmalonic encephalopathy 1 | 298.28 | 258.13 | 0.21 | 4.61E-01 |
| ENSMUSG00000040940 | GSMG0034167 | Arhgef1 | Rho guanine nucleotide exchange factor (GEF) 1 | 7135.61 | 7546.93 | -0.08 | 7.75E-01 |
| ENSMUSG00000045252 | GSMG0034168 | Zfp574 | zinc finger protein 574 | 637.46 | 558.23 | 0.19 | 5.50E-01 |
| ENSMUSG00000043843 | GSMG0034175 | Tmem145 | transmembrane protein 145 | 3536.06 | 4204.22 | -0.25 | 3.84E-01 |
| ENSMUSG00000045039 | GSMG0034176 | Megf8 | multiple EGF-like-domains 8 | 442.56 | 417.94 | 0.08 | 8.74E-01 |
| ENSMUSG00000057229 | GSMG0034180 | Atp5sl | ATP5S-like | 419.44 | 369.81 | 0.18 | 5.01E-01 |
| ENSMUSG00000061286 | GSMG0034182 | Exosc5 | exosome component 5 | 324.38 | 397.22 | -0.29 | 2.63E-01 |
| ENSMUSG00000063439 | GSMG0034183 | B9d2 | B9 protein domain 2 | 346.83 | 368.89 | -0.09 | 7.97E-01 |
| ENSMUSG00000002603 | GSMG0034184 | Tgfb1 | transforming growth factor, beta 1 | 674.45 | 605.14 | 0.15 | 7.16E-01 |
| ENSMUSG00000003762 | GSMG0034206 | Adck4 | aarF domain containing kinase 4 | 461.76 | 364.86 | 0.34 | 1.63E-01 |
| ENSMUSG00000063160 | GSMG0034208 | Numbl | numb-like | 345.39 | 348.38 | -0.01 | 9.71E-01 |
| ENSMUSG00000040466 | GSMG0034211 | Blvrb | biliverdin reductase B (flavin reductase (NADPH)) | 762.27 | 723.57 | 0.08 | 8.00E-01 |
| ENSMUSG00000055200 | GSMG0034212 | Sertad3 | SERTA domain containing 3 | 487.08 | 403.45 | 0.27 | 3.27E-01 |
| ENSMUSG00000008384 | GSMG0034213 | Sertad1 | SERTA domain containing 1 | 1362.64 | 697.55 | 0.97 | 2.85E-08 |
| ENSMUSG00000049643 | GSMG0034216 | 2310022A10Rik | RIKEN cDNA 2310022A10 gene | 293.38 | 348.89 | -0.25 | 4.04E-01 |
| ENSMUSG00000004056 | GSMG0034218 | Akt2 | thymoma viral proto-oncogene 2 | 1036.70 | 907.57 | 0.19 | 4.24E-01 |
| ENSMUSG00000007944 | GSMG0034219 | Ttc9b | tetratricopeptide repeat domain 9B | 670.88 | 575.61 | 0.22 | 4.43E-01 |
| ENSMUSG00000037640 | GSMG0034223 | Zfp60 | zinc finger protein 60 | 881.18 | 835.87 | 0.08 | 7.75E-01 |
| ENSMUSG00000002409 | GSMG0034238 | Dyrk1b | dual-specificity tyrosine-(Y)-phosphorylation regulated kinase 1b | 541.72 | 397.56 | 0.45 | 1.02E-01 |
| NULL | GSMG0034239 | NULL | NULL | 208.66 | 197.34 | 0.08 | 8.57E-01 |
| ENSMUSG00000046058 | GSMG0034242 | Eid2 | EP300 interacting inhibitor of differentiation 2 | 749.11 | 660.09 | 0.18 | 6.32E-01 |
| ENSMUSG00000070705 | GSMG0034243 | Eid2b | EP300 interacting inhibitor of differentiation 2B | 376.54 | 469.62 | -0.32 | 2.18E-01 |
| NULL | GSMG0034245 | NULL | NULL | 437.06 | 187.57 | 1.22 | 3.38E-09 |
| ENSMUSG00000003437 | GSMG0034246 | Paf1 | Paf1, RNA polymerase II complex component | 1982.77 | 1736.30 | 0.19 | 3.60E-01 |
| ENSMUSG00000030600 | GSMG0034249 | Lrfn1 | leucine rich repeat and fibronectin type III domain containing 1 | 202.97 | 387.13 | -0.94 | 4.76E-05 |
| ENSMUSG00000037463 | GSMG0034254 | Fbxo27 | F-box protein 27 | 107.49 | 131.49 | -0.29 | 4.38E-01 |
| ENSMUSG00000070699 | GSMG0034256 | Sars2 | seryl-aminoacyl-tRNA synthetase 2 | 460.27 | 502.57 | -0.13 | 6.72E-01 |
| ENSMUSG00000015149 | GSMG0034257 | Sirt2 | sirtuin 2 | 1489.98 | 896.08 | 0.73 | 2.53E-05 |

| | | | | | | | |
|---|---|---|---|---|---|---|---|
| ENSMUSG00000051735 | GSMG0034258 | Rinl | Ras and Rab interactor-like | 76.00 | 78.16 | -0.04 | 9.53E-01 |
| ENSMUSG00000015165 | GSMG0034259 | Hnrnpl | heterogeneous nuclear ribonucleoprotein L | 2517.93 | 2986.89 | -0.25 | 3.21E-01 |
| ENSMUSG00000053898 | GSMG0034260 | Ech1 | enoyl coenzyme A hydratase 1, peroxisomal | 4274.15 | 4015.38 | 0.09 | 7.21E-01 |
| ENSMUSG00000053964 | GSMG0034261 | Lgals4 | lectin, galactose binding, soluble 4 | 248.78 | 313.13 | -0.34 | 2.37E-01 |
| NULL | GSMG0034264 | NULL | NULL | 650.65 | 664.07 | -0.03 | 9.30E-01 |
| ENSMUSG00000037337 | GSMG0034265 | Map4k1 | mitogen-activated protein kinase kinase kinase kinase 1 | 69.90 | 155.93 | -1.15 | 1.09E-04 |
| ENSMUSG00000030589 | GSMG0034268 | Rasgrp4 | RAS guanyl releasing protein 4 | 380.95 | 403.89 | -0.09 | 8.07E-01 |
| NULL | GSMG0034269 | NULL | NULL | 127.03 | 72.99 | 0.79 | 3.10E-02 |
| ENSMUSG00000030588 | GSMG0034273 | Yif1b | Yip1 interacting factor homolog B (S. cerevisiae) | 1180.93 | 1284.44 | -0.12 | 6.21E-01 |
| ENSMUSG00000037166 | GSMG0034274 | Ppp1r14a | protein phosphatase 1, regulatory (inhibitor) subunit 14A | 383.17 | 269.79 | 0.50 | 9.95E-02 |
| ENSMUSG00000030584 | GSMG0034275 | Dpf1 | D4, zinc and double PHD fingers family 1 | 392.29 | 842.96 | -1.11 | 9.49E-11 |
| ENSMUSG00000046185 | GSMG0034281 | Zfp84 | zinc finger protein 84 | 929.95 | 832.51 | 0.16 | 5.46E-01 |
| ENSMUSG00000047473 | GSMG0034283 | Zfp30 | zinc finger protein 30 | 535.90 | 455.94 | 0.23 | 3.99E-01 |
| ENSMUSG00000011427 | GSMG0034284 | Zfp790 | zinc finger protein 790 | 997.48 | 789.79 | 0.34 | 6.50E-02 |
| ENSMUSG00000074221 | GSMG0034290 | Zfp568 | zinc finger protein 568 | 104.16 | 119.87 | -0.20 | 6.22E-01 |
| ENSMUSG00000049421 | GSMG0034292 | Zfp260 | zinc finger protein 260 | 2397.69 | 2227.91 | 0.11 | 6.37E-01 |
| ENSMUSG00000019738 | GSMG0034299 | Polr2i | polymerase (RNA) II (DNA directed) polypeptide I | 507.77 | 600.25 | -0.24 | 3.10E-01 |
| ENSMUSG00000013921 | GSMG0034301 | Clip3 | CAP-GLY domain containing linker protein 3 | 301.20 | 340.81 | -0.18 | 5.87E-01 |
| ENSMUSG00000042831 | GSMG0034302 | Alkbh6 | alkB homolog 6 | 1174.95 | 970.79 | 0.28 | 1.43E-01 |
| NULL | GSMG0034304 | NULL | NULL | 216.27 | 232.25 | -0.10 | 7.83E-01 |
| ENSMUSG00000030579 | GSMG0034305 | Tyrobp | TYRO protein tyrosine kinase binding protein | 399.12 | 217.54 | 0.88 | 7.82E-05 |
| ENSMUSG00000036931 | GSMG0034306 | Nfkbid | nuclear factor of kappa light polypeptide gene enhancer in B cells inhib | 245.55 | 198.58 | 0.31 | 2.79E-01 |
| G00000036826,ENSMUSG000 | GSMG0034312 | U2af1l4 // Igflr1 | U2 small nuclear RNA auxiliary factor 1-like 4 // IGF-like family recep | 1169.37 | 1164.77 | 0.01 | 9.83E-01 |
| NULL | GSMG0034317 | NULL | NULL | 136.40 | 150.42 | -0.14 | 7.77E-01 |
| NULL | GSMG0034322 | NULL | NULL | 1789.76 | 1301.77 | 0.46 | 8.91E-03 |
| ENSMUSG00000036560 | GSMG0034324 | Lgi4 | leucine-rich repeat LGI family, member 4 | 116.85 | 18.46 | 2.66 | 8.95E-12 |
| NULL | GSMG0034335 | NULL | NULL | 1404.46 | 1297.13 | 0.11 | 6.39E-01 |
| ENSMUSG00000063931 | GSMG0034341 | Pepd | peptidase D | 891.67 | 1033.57 | -0.21 | 3.28E-01 |
| NULL | GSMG0034343 | NULL | NULL | 1898.20 | 1568.89 | 0.27 | 1.23E-01 |
| ENSMUSG00000030495 | GSMG0034345 | Slc7a10 | solute carrier family 7 (cationic amino acid transporter, y+ system), mer | 767.18 | 522.11 | 0.56 | 8.96E-03 |
| ENSMUSG00000030494 | GSMG0034347 | Rhpn2 | rhophilin, Rho GTPase binding protein 2 | 300.63 | 91.54 | 1.72 | 9.54E-13 |
| ENSMUSG00000023072 | GSMG0034348 | Cep89 | centrosomal protein 89 | 878.64 | 1048.47 | -0.26 | 1.93E-01 |
| ENSMUSG00000034867 | GSMG0034351 | Ankrd27 | ankyrin repeat domain 27 (VPS9 domain) | 1432.76 | 1231.37 | 0.22 | 2.99E-01 |

| | | | | | | | |
|---|---|---|---|---|---|---|---|
| ENSMUSG00000021217 | GSMG0034355 | Tshz3 | teashirt zinc finger family member 3 | 484.89 | 700.64 | -0.53 | 1.86E-02 |
| NULL | GSMG0034356 | NULL | NULL | 242.81 | 232.12 | 0.06 | 8.95E-01 |
| NULL | GSMG0034357 | NULL | NULL | 639.21 | 608.21 | 0.07 | 8.23E-01 |
| ENSMUSG00000054676 | GSMG0034362 | 1600014C10Rik | RIKEN cDNA 1600014C10 gene | 448.18 | 383.62 | 0.22 | 4.27E-01 |
| ENSMUSG00000030424 | GSMG0034364 | Zfp939 | zinc finger protein 939 | 312.70 | 304.27 | 0.04 | 9.15E-01 |
| ENSMUSG00000039257 | GSMG0034374 | Vstm2b | V-set and transmembrane domain containing 2B | 369.26 | 399.80 | -0.12 | 7.18E-01 |
| NULL | GSMG0034379 | NULL | NULL | 192.68 | 179.38 | 0.11 | 8.09E-01 |
| ENSMUSG00000074165 | GSMG0034380 | Zfp788 | zinc finger protein 788 | 384.00 | 377.07 | 0.03 | 9.39E-01 |
| ENSMUSG00000004610 | GSMG0034390 | Etfb | electron transferring flavoprotein, beta polypeptide | 2078.16 | 2044.48 | 0.02 | 9.40E-01 |
| ENSMUSG00000030469 | GSMG0034394 | Zfp719 | zinc finger protein 719 | 975.82 | 877.75 | 0.15 | 5.47E-01 |
| ENSMUSG00000038888 | GSMG0034397 | Ctu1 | cytosolic thiouridylase subunit 1 | 170.85 | 227.45 | -0.41 | 1.33E-01 |
| ENSMUSG00000064023 | GSMG0034404 | Klk8 | kallikrein related-peptidase 8 | 186.06 | 237.53 | -0.36 | 2.00E-01 |
| G00000045411,ENSMUSG000 | GSMG0034425 | 2410002F23Rik // | RIKEN cDNA 2410002F23 gene // predicted gene 15517 | 2253.10 | 3549.45 | -0.66 | 2.12E-05 |
| ENSMUSG00000030731 | GSMG0034428 | Syt3 | synaptotagmin III | 1116.44 | 1148.09 | -0.04 | 9.00E-01 |
| ENSMUSG00000047085 | GSMG0034430 | Lrrc4b | leucine rich repeat containing 4B | 564.69 | 846.65 | -0.59 | 3.20E-02 |
| ENSMUSG00000038695 | GSMG0034432 | Josd2 | Josephin domain containing 2 | 438.50 | 234.09 | 0.91 | 1.53E-06 |
| ENSMUSG00000062785 | GSMG0034439 | Kcnc3 | potassium voltage gated channel, Shaw-related subfamily, member 3 | 192.74 | 124.56 | 0.63 | 5.02E-02 |
| ENSMUSG00000002205 | GSMG0034442 | Vrk3 | vaccinia related kinase 3 | 1117.01 | 1011.10 | 0.14 | 5.22E-01 |
| ENSMUSG00000011658 | GSMG0034443 | Fuz | fuzzy planar cell polarity protein | 327.56 | 227.10 | 0.53 | 3.40E-02 |
| G00000074141,ENSMUSG000 | GSMG0034445 | Nup62 // Gm21948 | nucleoporin 62 // predicted gene, 21948 // interleukin 4 induced 1 | 1106.57 | 1745.22 | -0.66 | 5.29E-06 |
| ENSMUSG00000011096 | GSMG0034446 | Akt1s1 | AKT1 substrate 1 (proline-rich) | 300.08 | 316.31 | -0.08 | 8.25E-01 |
| ENSMUSG00000002963 | GSMG0034447 | Pnkp | polynucleotide kinase 3'- phosphatase | 1502.60 | 1210.46 | 0.31 | 7.26E-02 |
| NULL | GSMG0034448 | NULL | NULL | 93.20 | 117.16 | -0.33 | 3.71E-01 |
| NULL | GSMG0034452 | NULL | NULL | 101.54 | 92.72 | 0.13 | 7.95E-01 |
| ENSMUSG00000003184 | GSMG0034454 | Irf3 | interferon regulatory factor 3 | 1117.26 | 977.62 | 0.19 | 3.32E-01 |
| ENSMUSG00000038387 | GSMG0034455 | Rras | related RAS viral (r-ras) oncogene | 3078.50 | 2048.15 | 0.59 | 8.25E-05 |
| ENSMUSG00000003421 | GSMG0034456 | Nosip | nitric oxide synthase interacting protein | 1137.81 | 1080.74 | 0.07 | 7.78E-01 |
| ENSMUSG00000003423 | GSMG0034458 | Pih1d1 | PIH1 domain containing 1 | 603.43 | 924.39 | -0.61 | 6.37E-04 |
| ENSMUSG00000070570 | GSMG0034459 | Slc17a7 | solute carrier family 17 (sodium-dependent inorganic phosphate cotrans | 7821.70 | 2639.02 | 1.57 | 2.50E-17 |
| ENSMUSG00000030796 | GSMG0034461 | Tead2 | TEA domain family member 2 | 842.02 | 1267.53 | -0.59 | 5.03E-03 |
| ENSMUSG00000003865 | GSMG0034470 | Gys1 | glycogen synthase 1, muscle | 318.78 | 371.78 | -0.22 | 4.88E-01 |
| ENSMUSG00000030826 | GSMG0034476 | Bcat2 | branched chain aminotransferase 2, mitochondrial | 388.77 | 511.54 | -0.39 | 6.69E-02 |
| G00000044562,ENSMUSG000 | GSMG0034478 | Izumo1 // Rasip1 | izumo sperm-egg fusion 1 // Ras interacting protein 1 | 2478.52 | 2243.34 | 0.14 | 6.70E-01 |

| | | | | | | | |
|---|---|---|---|---|---|---|---|
| ENSMUSG00000003273 | GSMG0034481 | Car11 | carbonic anhydrase 11 | 2102.20 | 2002.80 | 0.07 | 8.13E-01 |
| ENSMUSG00000059824 | GSMG0034482 | Dbp | D site albumin promoter binding protein | 555.91 | 206.18 | 1.43 | 5.09E-14 |
| NULL | GSMG0034487 | NULL | NULL | 125.27 | 73.02 | 0.78 | 1.63E-02 |
| ENSMUSG00000002778 | GSMG0034488 | Kdelr1 | KDEL (Lys-Asp-Glu-Leu) endoplasmic reticulum protein retention rece | 886.01 | 880.34 | 0.01 | 9.76E-01 |
| ENSMUSG00000002781 | GSMG0034489 | Tmem143 | transmembrane protein 143 | 80.26 | 66.26 | 0.27 | 5.96E-01 |
| ENSMUSG00000030835 | GSMG0034491 | Nomo1 | nodal modulator 1 | 16779.57 | 17179.36 | -0.03 | 8.92E-01 |
| ENSMUSG00000058975 | GSMG0034496 | Kcnc1 | potassium voltage gated channel, Shaw-related subfamily, member 1 | 554.35 | 370.10 | 0.58 | 3.10E-03 |
| ENSMUSG00000006599 | GSMG0034498 | Gtf2h1 | general transcription factor II H, polypeptide 1 | 922.89 | 1009.69 | -0.13 | 5.80E-01 |
| ENSMUSG00000063229 | GSMG0034499 | Ldha | lactate dehydrogenase A | 14569.29 | 11737.36 | 0.31 | 1.43E-01 |
| ENSMUSG00000010307 | GSMG0034502 | Tmem86a | transmembrane protein 86A | 1728.19 | 2066.19 | -0.26 | 1.29E-01 |
| ENSMUSG00000030471 | GSMG0034507 | Zdhhc13 | zinc finger, DHHC domain containing 13 | 446.36 | 592.59 | -0.41 | 7.39E-02 |
| NULL | GSMG0034514 | NULL | NULL | 2179.70 | 3207.91 | -0.56 | 1.01E-04 |
| NULL | GSMG0034515 | NULL | NULL | 519.78 | 473.94 | 0.14 | 6.65E-01 |
| ENSMUSG00000030505 | GSMG0034526 | Prmt3 | protein arginine N-methyltransferase 3 | 424.44 | 569.90 | -0.43 | 3.51E-02 |
| ENSMUSG00000055409 | GSMG0034530 | Nell1 | NEL-like 1 | 166.46 | 222.64 | -0.42 | 2.61E-01 |
| ENSMUSG00000030500 | GSMG0034535 | Slc17a6 | solute carrier family 17 (sodium-dependent inorganic phosphate cotrans | 68.95 | 300.02 | -2.11 | 5.50E-19 |
| ENSMUSG00000030498 | GSMG0034538 | Gas2 | growth arrest specific 2 | 78.07 | 109.01 | -0.48 | 1.86E-01 |
| ENSMUSG00000063297 | GSMG0034539 | Luzp2 | leucine zipper protein 2 | 444.07 | 610.70 | -0.46 | 2.56E-02 |
| ENSMUSG00000033790 | GSMG0034547 | Tubgcp5 | tubulin, gamma complex associated protein 5 | 1163.93 | 1169.11 | -0.01 | 9.86E-01 |
| ENSMUSG00000030447 | GSMG0034548 | Cyfip1 | cytoplasmic FMR1 interacting protein 1 | 3913.31 | 4198.80 | -0.10 | 6.22E-01 |
| ENSMUSG00000030451 | GSMG0034550 | Herc2 | HECT and RLD domain containing E3 ubiquitin protein ligase 2 | 811.84 | 759.19 | 0.10 | 7.34E-01 |
| ENSMUSG00000033676 | GSMG0034554 | Gabrb3 | gamma-aminobutyric acid (GABA) A receptor, subunit beta 3 | 350.72 | 580.44 | -0.73 | 2.54E-04 |
| ENSMUSG00000025324 | GSMG0034558 | Atp10a | ATPase, class V, type 10A | 1598.69 | 337.61 | 2.24 | 1.70E-33 |
| ENSMUSG00000025326 | GSMG0034561 | Ube3a | ubiquitin protein ligase E3A | 1313.22 | 1172.86 | 0.16 | 5.33E-01 |
| NULL | GSMG0034563 | NULL | NULL | 505.75 | 341.20 | 0.57 | 4.32E-02 |
| NULL | GSMG0034566 | NULL | NULL | 190.09 | 553.99 | -1.55 | 8.13E-08 |
| ENSMUSG00000033510 | GSMG0034567 | Otud7a | OTU domain containing 7A | 350.36 | 264.97 | 0.40 | 3.04E-01 |
| NULL | GSMG0034570 | NULL | NULL | 898.50 | 1236.17 | -0.46 | 9.40E-02 |
| ENSMUSG00000030522 | GSMG0034575 | Mtmr10 | myotubularin related protein 10 | 355.99 | 296.84 | 0.26 | 3.02E-01 |
| ENSMUSG00000033429 | GSMG0034576 | Mcee | methylmalonyl CoA epimerase | 519.33 | 490.96 | 0.08 | 8.36E-01 |
| ENSMUSG00000030519 | GSMG0034577 | Apba2 | amyloid beta (A4) precursor protein-binding, family A, member 2 | 793.42 | 932.51 | -0.23 | 3.83E-01 |
| ENSMUSG00000030515 | GSMG0034579 | Tarsl2 | threonyl-tRNA synthetase-like 2 | 392.82 | 395.84 | -0.01 | 9.73E-01 |
| ENSMUSG00000078681 | GSMG0034580 | Tm2d3 | TM2 domain containing 3 | 1443.93 | 1642.94 | -0.19 | 3.60E-01 |

| | | | | | | | |
|---|---|---|---|---|---|---|---|
| ENSMUSG00000030512 | GSMG0034583 | Snrpa1 | small nuclear ribonucleoprotein polypeptide A' | 986.33 | 1512.17 | -0.62 | 8.04E-04 |
| ENSMUSG00000075701 | GSMG0034584 | Vimp | VCP-interacting membrane protein | 2239.47 | 2089.52 | 0.10 | 6.59E-01 |
| ENSMUSG00000032640 | GSMG0034585 | Chsy1 | chondroitin sulfate synthase 1 | 606.04 | 353.48 | 0.78 | 1.86E-02 |
| ENSMUSG00000053091 | GSMG0034588 | Lins1 | lines homolog 1 | 303.94 | 384.71 | -0.34 | 1.76E-01 |
| ENSMUSG00000058145 | GSMG0034592 | Adamts17 | a disintegrin-like and metallopeptidase (reprolysin type) with thrombos | 296.97 | 216.53 | 0.46 | 1.06E-01 |
| ENSMUSG00000043831 | GSMG0034596 | Lysmd4 | LysM, putative peptidoglycan-binding, domain containing 4 | 778.49 | 835.55 | -0.10 | 6.84E-01 |
| ENSMUSG00000030555 | GSMG0034598 | Ttc23 | tetratricopeptide repeat domain 23 | 257.03 | 232.89 | 0.15 | 6.70E-01 |
| ENSMUSG00000005533 | GSMG0034600 | Igf1r | insulin-like growth factor I receptor | 6514.56 | 5425.01 | 0.26 | 3.25E-01 |
| ENSMUSG00000100005 | GSMG0034615 | B130024G19Rik | RIKEN cDNA B130024G19 gene | 261.77 | 202.35 | 0.37 | 1.80E-01 |
| ENSMUSG00000070509 | GSMG0034627 | Rgma | repulsive guidance molecule family member A | 114.44 | 183.45 | -0.68 | 5.18E-01 |
| NULL | GSMG0034629 | NULL | NULL | 40.11 | 178.81 | -2.15 | 8.48E-13 |
| ENSMUSG00000078670 | GSMG0034633 | Fam174b | family with sequence similarity 174, member B | 731.88 | 1150.24 | -0.65 | 7.08E-04 |
| ENSMUSG00000055652 | GSMG0034652 | Klhl25 | kelch-like 25 | 197.27 | 282.29 | -0.51 | 5.49E-02 |
| ENSMUSG00000030611 | GSMG0034660 | Mrps11 | mitochondrial ribosomal protein S11 | 654.20 | 781.82 | -0.26 | 2.56E-01 |
| ENSMUSG00000030609 | GSMG0034663 | Aen | apoptosis enhancing nuclease | 223.26 | 352.40 | -0.66 | 2.79E-03 |
| ENSMUSG00000039236 | GSMG0034664 | Isg20 | interferon-stimulated protein | 186.06 | 173.07 | 0.10 | 7.91E-01 |
| ENSMUSG00000039202 | GSMG0034666 | Abhd2 | abhydrolase domain containing 2 | 1355.21 | 936.50 | 0.53 | 3.85E-02 |
| NULL | GSMG0034669 | NULL | NULL | 879.02 | 719.75 | 0.29 | 2.86E-01 |
| ENSMUSG00000097023 | GSMG0034672 | Mir9-3hg | Mir9-3 host gene | 684.01 | 703.42 | -0.04 | 9.16E-01 |
| NULL | GSMG0034673 | NULL | NULL | 145.20 | 158.36 | -0.12 | 7.92E-01 |
| NULL | GSMG0034674 | NULL | NULL | 88.85 | 85.25 | 0.06 | 9.14E-01 |
| NULL | GSMG0034683 | NULL | NULL | 233.15 | 175.69 | 0.40 | 1.86E-01 |
| ENSMUSG00000048897 | GSMG0034684 | Zfp710 | zinc finger protein 710 | 329.68 | 428.39 | -0.38 | 1.36E-01 |
| ENSMUSG00000047084 | GSMG0034689 | Ngrn | neugrin, neurite outgrowth associated | 1088.76 | 1100.97 | -0.02 | 9.61E-01 |
| ENSMUSG00000030534 | GSMG0034690 | Vps33b | vacuolar protein sorting 33B | 1216.27 | 1531.87 | -0.33 | 6.60E-02 |
| ENSMUSG00000038943 | GSMG0034691 | Prc1 | protein regulator of cytokinesis 1 | 663.30 | 1512.67 | -1.19 | 6.38E-10 |
| ENSMUSG00000030532 | GSMG0034692 | Hddc3 | HD domain containing 3 | 569.76 | 647.93 | -0.18 | 4.28E-01 |
| NULL | GSMG0034697 | NULL | NULL | 63.47 | 81.85 | -0.37 | 3.98E-01 |
| ENSMUSG00000005621 | GSMG0034699 | Zfp592 | zinc finger protein 592 | 997.73 | 1044.52 | -0.07 | 8.53E-01 |
| ENSMUSG00000025584 | GSMG0034703 | Pde8a | phosphodiesterase 8A | 1723.96 | 1704.52 | 0.02 | 9.62E-01 |
| NULL | GSMG0034705 | NULL | NULL | 51.68 | 25.67 | 1.01 | 6.23E-02 |
| NULL | GSMG0034706 | NULL | NULL | 86.42 | 38.32 | 1.17 | 1.76E-03 |
| ENSMUSG00000097277 | GSMG0034708 | 2900076A07Rik | RIKEN cDNA 2900076A07 gene | 199.15 | 186.51 | 0.09 | 8.34E-01 |

| | | | | | | | |
|---|---|---|---|---|---|---|---|
| ENSMUSG00000045795 | GSMG0034709 | Whamm | WAS protein homolog associated with actin, golgi membranes and micı | 517.40 | 376.02 | 0.46 | 7.51E-02 |
| ENSMUSG00000038623 | GSMG0034712 | Tm6sf1 | transmembrane 6 superfamily member 1 | 1270.37 | 884.04 | 0.52 | 7.87E-03 |
| NULL | GSMG0034713 | NULL | NULL | 109.83 | 131.74 | -0.26 | 4.98E-01 |
| ENSMUSG00000030638 | GSMG0034716 | Sh3gl3 | SH3-domain GRB2-like 3 | 185.42 | 239.54 | -0.37 | 1.60E-01 |
| ENSMUSG00000038563 | GSMG0034719 | Efl1 | elongation factor like GPTase 1 | 838.35 | 933.60 | -0.15 | 5.40E-01 |
| ENSMUSG00000046027 | GSMG0034728 | Stard5 | StAR-related lipid transfer (START) domain containing 5 | 494.67 | 786.57 | -0.67 | 1.38E-04 |
| ENSMUSG00000038503 | GSMG0034733 | Mesdc2 | mesoderm development candidate 2 | 1692.52 | 2029.04 | -0.26 | 2.21E-01 |
| ENSMUSG00000030562 | GSMG0034746 | Nox4 | NADPH oxidase 4 | 231.84 | 376.79 | -0.70 | 4.83E-04 |
| ENSMUSG00000049583 | GSMG0034747 | Grm5 | glutamate receptor, metabotropic 5 | 466.89 | 726.14 | -0.64 | 3.88E-03 |
| NULL | GSMG0034749 | NULL | NULL | 104.93 | 88.86 | 0.25 | 6.33E-01 |
| NULL | GSMG0034750 | NULL | NULL | 264.98 | 325.55 | -0.30 | 3.62E-01 |
| NULL | GSMG0034752 | NULL | NULL | 335.78 | 375.84 | -0.16 | 6.19E-01 |
| ENSMUSG00000030560 | GSMG0034753 | Ctsc | cathepsin C | 1118.30 | 881.63 | 0.34 | 1.87E-01 |
| ENSMUSG00000049791 | GSMG0034757 | Fzd4 | frizzled class receptor 4 | 3775.59 | 2619.39 | 0.53 | 8.47E-03 |
| NULL | GSMG0034758 | NULL | NULL | 2389.78 | 1342.68 | 0.83 | 1.80E-06 |
| NULL | GSMG0034760 | NULL | NULL | 420.35 | 98.50 | 2.10 | 5.48E-24 |
| ENSMUSG00000030621 | GSMG0034761 | Me3 | malic enzyme 3, NADP(+)-dependent, mitochondrial | 388.02 | 189.28 | 1.04 | 4.67E-06 |
| ENSMUSG00000039361 | GSMG0034767 | Picalm | phosphatidylinositol binding clathrin assembly protein | 1244.22 | 792.60 | 0.65 | 1.40E-04 |
| ENSMUSG00000030616 | GSMG0034772 | Sytl2 | synaptotagmin-like 2 | 545.71 | 227.02 | 1.27 | 1.77E-09 |
| NULL | GSMG0034779 | NULL | NULL | 724.88 | 594.81 | 0.28 | 2.13E-01 |
| NULL | GSMG0034782 | NULL | NULL | 410.30 | 386.49 | 0.09 | 8.38E-01 |
| NULL | GSMG0034784 | NULL | NULL | 1143.02 | 871.74 | 0.39 | 4.89E-02 |
| NULL | GSMG0034785 | NULL | NULL | 696.38 | 649.55 | 0.10 | 8.08E-01 |
| NULL | GSMG0034786 | NULL | NULL | 388.56 | 285.95 | 0.44 | 8.11E-02 |
| NULL | GSMG0034788 | NULL | NULL | 273.88 | 198.73 | 0.46 | 5.57E-02 |
| NULL | GSMG0034789 | NULL | NULL | 792.71 | 688.31 | 0.20 | 4.30E-01 |
| NULL | GSMG0034790 | NULL | NULL | 751.36 | 654.43 | 0.20 | 4.63E-01 |
| NULL | GSMG0034791 | NULL | NULL | 972.36 | 1060.13 | -0.13 | 7.14E-01 |
| NULL | GSMG0034792 | NULL | NULL | 611.95 | 473.54 | 0.37 | 1.44E-01 |
| NULL | GSMG0034801 | NULL | NULL | 387.93 | 449.53 | -0.21 | 6.12E-01 |
| ENSMUSG00000030613 | GSMG0034802 | Ccdc90b | coiled-coil domain containing 90B | 454.12 | 494.56 | -0.12 | 6.84E-01 |
| NULL | GSMG0034804 | NULL | NULL | 376.65 | 261.87 | 0.53 | 4.84E-02 |
| ENSMUSG00000030643 | GSMG0034806 | Rab30 | RAB30, member RAS oncogene family | 203.81 | 187.38 | 0.12 | 7.69E-01 |

| | | | | | | | |
|---|---|---|---|---|---|---|---|
| NULL | GSMG0034807 | NULL | NULL | 204.17 | 153.02 | 0.41 | 1.77E-01 |
| ENSMUSG00000061119 | GSMG0034810 | Prcp | prolylcarboxypeptidase (angiotensinase C) | 904.21 | 1250.84 | -0.47 | 1.47E-02 |
| ENSMUSG00000051515 | GSMG0034814 | Fam181b | family with sequence similarity 181, member B | 670.73 | 550.88 | 0.28 | 1.89E-01 |
| ENSMUSG00000048078 | GSMG0034819 | Tenm4 | teneurin transmembrane protein 4 | 849.84 | 883.85 | -0.06 | 9.01E-01 |
| ENSMUSG00000018995 | GSMG0034830 | Nars2 | asparaginyl-tRNA synthetase 2 (mitochondrial)(putative) | 472.94 | 483.40 | -0.03 | 9.34E-01 |
| ENSMUSG00000004508 | GSMG0034834 | Gab2 | growth factor receptor bound protein 2-associated protein 2 | 423.85 | 400.11 | 0.09 | 8.44E-01 |
| ENSMUSG00000035704 | GSMG0034838 | Alg8 | asparagine-linked glycosylation 8 (alpha-1,3-glucosyltransferase) | 314.38 | 400.91 | -0.35 | 1.58E-01 |
| ENSMUSG00000030647 | GSMG0034839 | Ndufc2 | NADH dehydrogenase (ubiquinone) 1, subcomplex unknown, 2 | 2320.75 | 3214.55 | -0.47 | 7.04E-02 |
| ENSMUSG00000025133 | GSMG0034842 | Ints4 | integrator complex subunit 4 | 372.87 | 433.24 | -0.21 | 4.18E-01 |
| ENSMUSG00000035623 | GSMG0034844 | Rsf1 | remodeling and spacing factor 1 | 370.23 | 387.38 | -0.06 | 8.72E-01 |
| ENSMUSG00000025439 | GSMG0034849 | Clns1a | chloride channel, nucleotide-sensitive, 1A | 830.50 | 914.42 | -0.14 | 6.10E-01 |
| ENSMUSG00000030774 | GSMG0034852 | Pak1 | p21 protein (Cdc42/Rac)-activated kinase 1 | 458.22 | 587.18 | -0.36 | 7.41E-02 |
| NULL | GSMG0034857 | NULL | NULL | 364.78 | 350.99 | 0.05 | 8.89E-01 |
| ENSMUSG00000030753 | GSMG0034858 | Prkrir | protein-kinase, interferon-inducible double stranded RNA dependent in | 2331.78 | 2608.89 | -0.16 | 4.24E-01 |
| ENSMUSG00000055407 | GSMG0034864 | Map6 | microtubule-associated protein 6 | 229.64 | 332.28 | -0.53 | 2.07E-02 |
| ENSMUSG00000035314 | GSMG0034866 | Gdpd5 | glycerophosphodiester phosphodiesterase domain containing 5 | 1578.75 | 2216.02 | -0.49 | 1.32E-02 |
| NULL | GSMG0034867 | NULL | NULL | 114.42 | 99.69 | 0.21 | 6.61E-01 |
| ENSMUSG00000018909 | GSMG0034869 | Arrb1 | arrestin, beta 1 | 221.34 | 260.04 | -0.23 | 5.51E-01 |
| ENSMUSG00000030725 | GSMG0034875 | Lipt2 | lipoyl(octanoyl) transferase 2 (putative) | 154.87 | 256.76 | -0.73 | 4.80E-03 |
| ENSMUSG00000030729 | GSMG0034877 | Pgm2l1 | phosphoglucomutase 2-like 1 | 537.96 | 459.94 | 0.23 | 3.28E-01 |
| ENSMUSG00000047248 | GSMG0034879 | C2cd3 | C2 calcium-dependent domain containing 3 | 594.27 | 583.87 | 0.03 | 9.43E-01 |
| ENSMUSG00000033685 | GSMG0034882 | Ucp2 | uncoupling protein 2 (mitochondrial, proton carrier) | 6750.52 | 4765.75 | 0.50 | 2.97E-03 |
| ENSMUSG00000030704 | GSMG0034885 | Rab6a | RAB6A, member RAS oncogene family | 6400.93 | 6491.57 | -0.02 | 9.50E-01 |
| ENSMUSG00000029461 | GSMG0034887 | Fam168a | family with sequence similarity 168, member A | 366.17 | 420.51 | -0.20 | 5.26E-01 |
| ENSMUSG00000030691 | GSMG0034893 | Fchsd2 | FCH and double SH3 domains 2 | 283.83 | 249.44 | 0.19 | 5.67E-01 |
| ENSMUSG00000030688 | GSMG0034898 | Stard10 | START domain containing 10 | 187.23 | 83.15 | 1.17 | 9.44E-05 |
| ENSMUSG00000032812 | GSMG0034899 | Arap1 | ArfGAP with RhoGAP domain, ankyrin repeat and PH domain 1 | 787.69 | 591.86 | 0.41 | 9.62E-02 |
| ENSMUSG00000110195 | GSMG0034900 | Pde2a | phosphodiesterase 2A, cGMP-stimulated | 7276.82 | 6274.25 | 0.21 | 3.40E-01 |
| ENSMUSG00000001829 | GSMG0034901 | Clpb | ClpB caseinolytic peptidase B | 1111.91 | 1327.48 | -0.25 | 2.03E-01 |
| ENSMUSG00000030649 | GSMG0034905 | Anapc15 | anaphase prompoting complex C subunit 15 | 76.20 | 111.99 | -0.55 | 1.08E-01 |
| ENSMUSG00000030842 | GSMG0034906 | Lamtor1 | late endosomal/lysosomal adaptor, MAPK and MTOR activator 1 | 1780.70 | 1812.66 | -0.03 | 9.28E-01 |
| ENSMUSG00000066306 | GSMG0034907 | Numa1 | nuclear mitotic apparatus protein 1 | 2501.44 | 2809.50 | -0.17 | 5.08E-01 |
| 25,ENSMUSG00000099481,EI | GSMG0034909 | Xntrpc // Xndc1 // | Xndc1-transient receptor potential cation channel, subfamily C, membe | 275.64 | 310.50 | -0.17 | 6.14E-01 |

| | | | | | | | |
|---|---|---|---|---|---|---|---|
| ENSMUSG00000030990 | GSMG0034911 | Pgap2 | post-GPI attachment to proteins 2 | 188.24 | 263.93 | -0.48 | 1.03E-01 |
| ENSMUSG00000030987 | GSMG0034914 | Stim1 | stromal interaction molecule 1 | 1217.21 | 953.92 | 0.35 | 1.31E-01 |
| ENSMUSG00000030978 | GSMG0034918 | Rrm1 | ribonucleotide reductase M1 | 1483.96 | 2726.21 | -0.88 | 5.74E-06 |
| NULL | GSMG0034941 | NULL | NULL | 1888.95 | 2002.10 | -0.08 | 7.75E-01 |
| NULL | GSMG0034942 | NULL | NULL | 7104.09 | 7245.72 | -0.03 | 9.22E-01 |
| ENSMUSG00000030898 | GSMG0034975 | Cckbr | cholecystokinin B receptor | 274.51 | 39.14 | 2.82 | 1.95E-23 |
| NULL | GSMG0034976 | NULL | NULL | 694.23 | 457.68 | 0.60 | 1.93E-03 |
| ENSMUSG00000037049 | GSMG0034977 | Smpd1 | sphingomyelin phosphodiesterase 1, acid lysosomal | 2968.96 | 2685.60 | 0.15 | 5.18E-01 |
| ENSMUSG00000089847 | GSMG0034978 | Timm10b | translocase of inner mitochondrial membrane 10B | 1620.44 | 1824.81 | -0.17 | 5.07E-01 |
| ENSMUSG00000030890 | GSMG0034979 | Ilk | integrin linked kinase | 7597.08 | 6727.91 | 0.18 | 3.62E-01 |
| ENSMUSG00000051041 | GSMG0034994 | Olfml1 | olfactomedin-like 1 | 189.35 | 154.56 | 0.30 | 4.74E-01 |
| ENSMUSG00000036528 | GSMG0034996 | Ppfibp2 | PTPRF interacting protein, binding protein 2 (liprin beta 2) | 412.76 | 283.95 | 0.54 | 9.42E-03 |
| ENSMUSG00000031029 | GSMG0035013 | Eif3f | eukaryotic translation initiation factor 3, subunit F | 997.84 | 1202.19 | -0.27 | 1.99E-01 |
| ENSMUSG00000031023 | GSMG0035018 | Akip1 | A kinase (PRKA) interacting protein 1 | 327.83 | 358.40 | -0.13 | 6.79E-01 |
| NULL | GSMG0035019 | NULL | NULL | 335.38 | 314.48 | 0.09 | 7.91E-01 |
| ENSMUSG00000066232 | GSMG0035021 | Ipo7 | importin 7 | 979.96 | 1172.92 | -0.26 | 2.99E-01 |
| ENSMUSG00000061079 | GSMG0035023 | Zfp143 | zinc finger protein 143 | 456.71 | 438.21 | 0.06 | 8.52E-01 |
| ENSMUSG00000031016 | GSMG0035024 | Wee1 | WEE 1 homolog 1 (S. pombe) | 434.24 | 541.57 | -0.32 | 1.80E-01 |
| ENSMUSG00000031015 | GSMG0035026 | Swap70 | SWA-70 protein | 3229.99 | 3825.86 | -0.24 | 1.76E-01 |
| ENSMUSG00000030790 | GSMG0035027 | Adm | adrenomedullin | 217.62 | 141.89 | 0.62 | 5.31E-02 |
| ENSMUSG00000005686 | GSMG0035028 | Ampd3 | adenosine monophosphate deaminase 3 | 327.51 | 250.39 | 0.38 | 1.70E-01 |
| ENSMUSG00000005609 | GSMG0035029 | Ctr9 | CTR9 homolog, Paf1/RNA polymerase II complex component | 1291.40 | 1499.45 | -0.22 | 3.05E-01 |
| ENSMUSG00000059263 | GSMG0035032 | Usp47 | ubiquitin specific peptidase 47 | 1068.60 | 1136.29 | -0.09 | 7.18E-01 |
| ENSMUSG00000038244 | GSMG0035034 | Mical2 | microtubule associated monooxygenase, calponin and LIM domain con | 2552.51 | 2627.80 | -0.04 | 9.03E-01 |
| ENSMUSG00000030770 | GSMG0035036 | Parva | parvin, alpha | 1012.22 | 1196.52 | -0.24 | 2.35E-01 |
| ENSMUSG00000055116 | GSMG0035049 | Arntl | aryl hydrocarbon receptor nuclear translocator-like | 296.59 | 371.79 | -0.32 | 2.17E-01 |
| NULL | GSMG0035051 | NULL | NULL | 418.46 | 418.61 | 0.00 | 9.99E-01 |
| ENSMUSG00000030759 | GSMG0035054 | Far1 | fatty acyl CoA reductase 1 | 757.98 | 794.68 | -0.07 | 8.48E-01 |
| ENSMUSG00000038156 | GSMG0035057 | Spon1 | spondin 1, (f-spondin) extracellular matrix protein | 553.73 | 690.15 | -0.32 | 2.23E-01 |
| ENSMUSG00000048782 | GSMG0035062 | Insc | inscuteable homolog (Drosophila) | 101.48 | 20.57 | 2.31 | 2.36E-09 |
| ENSMUSG00000030663 | GSMG0035067 | 1110004F10Rik | RIKEN cDNA 1110004F10 gene | 2648.60 | 3583.93 | -0.44 | 1.43E-02 |
| NULL | GSMG0035069 | NULL | NULL | 213.08 | 198.15 | 0.10 | 8.04E-01 |
| ENSMUSG00000030659 | GSMG0035070 | Nucb2 | nucleobindin 2 | 710.52 | 758.32 | -0.09 | 7.67E-01 |

| | | | | | | | |
|---|---|---|---|---|---|---|---|
| NULL | GSMG0035071 | NULL | NULL | 276.02 | 430.84 | -0.65 | 9.44E-03 |
| ENSMUSG00000030657 | GSMG0035072 | Xylt1 | xylosyltransferase 1 | 572.53 | 530.41 | 0.11 | 7.89E-01 |
| NULL | GSMG0035073 | NULL | NULL | 592.06 | 465.18 | 0.35 | 2.08E-01 |
| NULL | GSMG0035080 | NULL | NULL | 206.10 | 164.64 | 0.32 | 2.88E-01 |
| ENSMUSG00000033904 | GSMG0035081 | Ccp110 | centriolar coiled coil protein 110 | 94.67 | 93.85 | 0.00 | 9.96E-01 |
| ENSMUSG00000030982 | GSMG0035082 | 9030624J02Rik | RIKEN cDNA 9030624J02 gene | 2231.87 | 2841.18 | -0.35 | 4.16E-02 |
| ENSMUSG00000030924 | GSMG0035092 | 2610020H08Rik | RIKEN cDNA 2610020H08 gene | 100.42 | 112.96 | -0.18 | 7.06E-01 |
| ENSMUSG00000030917 | GSMG0035096 | Tmem159 | transmembrane protein 159 | 405.92 | 307.63 | 0.40 | 1.28E-01 |
| ENSMUSG00000030884 | GSMG0035104 | Uqcrc2 | ubiquinol cytochrome c reductase core protein 2 | 2782.52 | 3164.92 | -0.19 | 3.46E-01 |
| ENSMUSG00000046096 | GSMG0035106 | BC030336 | cDNA sequence BC030336 | 182.99 | 168.66 | 0.11 | 8.07E-01 |
| ENSMUSG00000035064 | GSMG0035108 | Eef2k | eukaryotic elongation factor-2 kinase | 826.55 | 1116.02 | -0.43 | 3.50E-02 |
| ENSMUSG00000030876 | GSMG0035113 | Mettl9 | methyltransferase like 9 | 1390.45 | 1925.25 | -0.47 | 5.59E-03 |
| ENSMUSG00000046321 | GSMG0035117 | Hs3st2 | heparan sulfate (glucosamine) 3-O-sulfotransferase 2 | 222.54 | 302.51 | -0.44 | 1.00E-01 |
| ENSMUSG00000030870 | GSMG0035122 | Ubfd1 | ubiquitin family domain containing 1 | 1643.85 | 1819.82 | -0.15 | 4.78E-01 |
| G00000030867,ENSMUSG000 | GSMG0035125 | Dctn5 // Plk1 | dynactin 5 // polo-like kinase 1 | 3083.46 | 3660.02 | -0.25 | 1.98E-01 |
| ENSMUSG00000030865 | GSMG0035126 | Chp2 | calcineurin-like EF hand protein 2 | 399.02 | 563.38 | -0.50 | 1.07E-02 |
| ENSMUSG00000052889 | GSMG0035127 | Prkcb | protein kinase C, beta | 3617.84 | 3076.37 | 0.23 | 2.26E-01 |
| NULL | GSMG0035132 | NULL | NULL | 405.95 | 227.88 | 0.83 | 3.94E-05 |
| ENSMUSG00000066189 | GSMG0035135 | Cacng3 | calcium channel, voltage-dependent, gamma subunit 3 | 897.16 | 238.92 | 1.91 | 2.68E-24 |
| ENSMUSG00000052707 | GSMG0035139 | Tnrc6a | trinucleotide repeat containing 6a | 345.68 | 282.86 | 0.29 | 2.81E-01 |
| ENSMUSG00000030763 | GSMG0035143 | Lcmt1 | leucine carboxyl methyltransferase 1 | 694.17 | 713.91 | -0.04 | 8.94E-01 |
| ENSMUSG00000030752 | GSMG0035151 | Kdm8 | lysine (K)-specific demethylase 8 | 108.97 | 177.23 | -0.70 | 8.38E-03 |
| ENSMUSG00000030748 | GSMG0035153 | Il4ra | interleukin 4 receptor, alpha | 384.29 | 509.53 | -0.41 | 1.42E-01 |
| ENSMUSG00000042978 | GSMG0035158 | Sbk1 | SH3-binding kinase 1 | 742.80 | 3403.55 | -2.19 | 2.59E-28 |
| ENSMUSG00000030727 | GSMG0035161 | Rabep2 | rabaptin, RAB GTPase binding effector protein 2 | 982.48 | 1194.99 | -0.28 | 1.15E-01 |
| NULL | GSMG0035165 | NULL | NULL | 256.97 | 197.96 | 0.38 | 1.58E-01 |
| ENSMUSG00000030714 | GSMG0035168 | Sgf29 | SAGA complex associated factor 29 | 535.66 | 599.58 | -0.16 | 5.18E-01 |
| ENSMUSG00000047721 | GSMG0035170 | Bola2 | bolA-like 2 (E. coli) | 516.68 | 734.07 | -0.51 | 3.73E-02 |
| ENSMUSG00000063065 | GSMG0035172 | Mapk3 | mitogen-activated protein kinase 3 | 2729.47 | 3018.43 | -0.15 | 5.04E-01 |
| ENSMUSG00000030703 | GSMG0035173 | Gdpd3 | glycerophosphodiester phosphodiesterase domain containing 3 | 186.62 | 19.57 | 3.25 | 1.50E-20 |
| ENSMUSG00000042675 | GSMG0035174 | Ypel3 | yippee-like 3 (Drosophila) | 2607.24 | 1650.83 | 0.66 | 5.80E-06 |
| ENSMUSG00000058966 | GSMG0035176 | Fam57b | family with sequence similarity 57, member B | 1127.05 | 1100.13 | 0.03 | 9.47E-01 |
| ENSMUSG00000052301 | GSMG0035179 | Doc2a | double C2, alpha | 242.18 | 429.61 | -0.83 | 5.04E-05 |

| | | | | | | | |
|---|---|---|---|---|---|---|---|
| ENSMUSG00000042606 | GSMG0035180 | Hirip3 | HIRA interacting protein 3 | 327.41 | 320.19 | 0.03 | 9.35E-01 |
| ENSMUSG00000030685 | GSMG0035181 | Kctd13 | potassium channel tetramerisation domain containing 13 | 252.66 | 465.40 | -0.88 | 1.64E-05 |
| ENSMUSG00000030683 | GSMG0035183 | Sez6l2 | seizure related 6 homolog like 2 | 672.90 | 893.44 | -0.41 | 8.33E-02 |
| ENSMUSG00000030682 | GSMG0035184 | Cdipt | CDP-diacylglycerol--inositol 3-phosphatidyltransferase (phosphatidylin | 4000.02 | 4462.70 | -0.16 | 4.16E-01 |
| ENSMUSG00000045165 | GSMG0035185 | AI467606 | expressed sequence AI467606 | 875.48 | 551.42 | 0.67 | 1.42E-03 |
| ENSMUSG00000030672 | GSMG0035186 | Mylpf | myosin light chain, phosphorylatable, fast skeletal muscle | 112.92 | 74.93 | 0.59 | 1.07E-01 |
| ENSMUSG00000045598 | GSMG0035187 | Zfp553 | zinc finger protein 553 | 442.33 | 571.22 | -0.37 | 7.15E-02 |
| ENSMUSG00000054716 | GSMG0035188 | Zfp771 | zinc finger protein 771 | 801.60 | 683.99 | 0.23 | 2.89E-01 |
| NULL | GSMG0035189 | NULL | NULL | 428.10 | 628.98 | -0.56 | 2.32E-03 |
| NULL | GSMG0035192 | NULL | NULL | 528.51 | 393.22 | 0.43 | 3.45E-02 |
| ENSMUSG00000030822 | GSMG0035193 | Prr14 | proline rich 14 | 1890.13 | 1696.56 | 0.16 | 4.48E-01 |
| ENSMUSG00000042423 | GSMG0035195 | Fbrs | fibrosin | 492.56 | 415.96 | 0.24 | 4.67E-01 |
| ENSMUSG00000053877 | GSMG0035198 | Srcap | Snf2-related CREBBP activator protein | 243.04 | 239.27 | 0.02 | 9.64E-01 |
| ENSMUSG00000030815 | GSMG0035201 | Phkg2 | phosphorylase kinase, gamma 2 (testis) | 637.17 | 525.27 | 0.28 | 1.71E-01 |
| ENSMUSG00000030816 | GSMG0035202 | Rnf40 | ring finger protein 40 | 1601.54 | 1769.07 | -0.14 | 5.74E-01 |
| ENSMUSG00000030811 | GSMG0035206 | Fbxl19 | F-box and leucine-rich repeat protein 19 | 156.67 | 202.94 | -0.38 | 3.46E-01 |
| ENSMUSG00000043964 | GSMG0035207 | Orai3 | ORAI calcium release-activated calcium modulator 3 | 903.23 | 866.85 | 0.06 | 8.52E-01 |
| G00000042289,ENSMUSG000 | GSMG0035208 | Setd1a // Hsd3b7 | SET domain containing 1A // hydroxy-delta-5-steroid dehydrogenase, 3 | 481.20 | 350.45 | 0.46 | 4.20E-02 |
| ENSMUSG00000030805 | GSMG0035209 | Stx4a | syntaxin 4A (placental) | 869.54 | 702.42 | 0.31 | 1.71E-01 |
| ENSMUSG00000049739 | GSMG0035210 | Zfp646 | zinc finger protein 646 | 1110.44 | 1060.48 | 0.07 | 8.66E-01 |
| ENSMUSG00000030802 | GSMG0035211 | Bckdk | branched chain ketoacid dehydrogenase kinase | 1769.71 | 2000.41 | -0.18 | 3.80E-01 |
| ENSMUSG00000030801 | GSMG0035212 | Kat8 | K(lysine) acetyltransferase 8 | 526.52 | 555.03 | -0.08 | 8.02E-01 |
| ENSMUSG00000030795 | GSMG0035213 | Fus | fused in sarcoma | 692.81 | 873.45 | -0.33 | 2.11E-01 |
| ENSMUSG00000042178 | GSMG0035220 | Armc5 | armadillo repeat containing 5 | 831.48 | 874.56 | -0.07 | 8.30E-01 |
| ENSMUSG00000030782 | GSMG0035221 | Tgfb1i1 | transforming growth factor beta 1 induced transcript 1 | 1355.63 | 1205.90 | 0.17 | 5.16E-01 |
| ENSMUSG00000030847 | GSMG0035224 | Bag3 | BCL2-associated athanogene 3 | 315.89 | 321.80 | -0.03 | 9.43E-01 |
| ENSMUSG00000042105 | GSMG0035226 | Inpp5f | inositol polyphosphate-5-phosphatase F | 258.34 | 340.05 | -0.40 | 1.42E-01 |
| ENSMUSG00000055319 | GSMG0035229 | Sec23ip | Sec23 interacting protein | 1034.34 | 1155.21 | -0.16 | 5.51E-01 |
| ENSMUSG00000042055 | GSMG0035231 | Wdr11 | WD repeat domain 11 | 1269.00 | 1434.53 | -0.18 | 4.90E-01 |
| ENSMUSG00000030852 | GSMG0035236 | Tacc2 | transforming, acidic coiled-coil containing protein 2 | 1044.28 | 1342.87 | -0.36 | 6.89E-02 |
| ENSMUSG00000040268 | GSMG0035239 | Plekha1 | pleckstrin homology domain containing, family A (phosphoinositide bi | 1430.36 | 1700.41 | -0.25 | 1.87E-01 |
| ENSMUSG00000006205 | GSMG0035240 | Htra1 | HtrA serine peptidase 1 | 4023.26 | 1793.18 | 1.17 | 3.91E-13 |
| ENSMUSG00000063179 | GSMG0035245 | Pstk | phosphoseryl-tRNA kinase | 260.13 | 266.62 | -0.04 | 9.28E-01 |

| | | | | | | | |
|---|---|---|---|---|---|---|---|
| NULL | GSMG0035246 | NULL | NULL | 553.96 | 549.38 | 0.01 | 9.73E-01 |
| ENSMUSG00000030861 | GSMG0035248 | Acadsb | acyl-Coenzyme A dehydrogenase, short/branched chain | 1080.36 | 1268.82 | -0.23 | 2.52E-01 |
| ENSMUSG00000066979 | GSMG0035253 | Bub3 | BUB3 mitotic checkpoint protein | 1627.42 | 1905.35 | -0.23 | 2.82E-01 |
| NULL | GSMG0035256 | NULL | NULL | 600.74 | 442.91 | 0.44 | 2.12E-02 |
| ENSMUSG00000030946 | GSMG0035261 | Lhpp | phospholysine phosphohistidine inorganic pyrophosphate phosphatase | 465.36 | 655.38 | -0.49 | 7.41E-03 |
| ENSMUSG00000030965 | GSMG0035263 | Fam175b | family with sequence similarity 175, member B | 262.09 | 284.75 | -0.12 | 7.34E-01 |
| ENSMUSG00000039990 | GSMG0035269 | Edrf1 | erythroid differentiation regulatory factor 1 | 729.85 | 649.13 | 0.17 | 4.67E-01 |
| ENSMUSG00000030983 | GSMG0035270 | Bccip | BRCA2 and CDKN1A interacting protein | 1180.17 | 1325.02 | -0.17 | 4.71E-01 |
| ENSMUSG00000058325 | GSMG0035273 | Dock1 | dedicator of cytokinesis 1 | 512.93 | 642.18 | -0.32 | 1.29E-01 |
| ENSMUSG00000041836 | GSMG0035283 | Ptpre | protein tyrosine phosphatase, receptor type, E | 684.01 | 808.74 | -0.24 | 2.54E-01 |
| NULL | GSMG0035284 | NULL | NULL | 225.56 | 209.83 | 0.10 | 8.13E-01 |
| ENSMUSG00000041769 | GSMG0035298 | Ppp2r2d | protein phosphatase 2, regulatory subunit B, delta | 877.88 | 876.13 | 0.00 | 9.92E-01 |
| ENSMUSG00000025478 | GSMG0035302 | Dpysl4 | dihydropyrimidinase-like 4 | 788.04 | 1099.73 | -0.48 | 3.01E-02 |
| ENSMUSG00000060260 | GSMG0035305 | Pwwp2b | PWWP domain containing 2B | 533.54 | 360.25 | 0.56 | 9.40E-03 |
| ENSMUSG00000025477 | GSMG0035306 | Inpp5a | inositol polyphosphate-5-phosphatase A | 1092.45 | 888.54 | 0.30 | 1.13E-01 |
| ENSMUSG00000025475 | GSMG0035307 | Adgra1 | adhesion G protein-coupled receptor A1 | 342.90 | 135.07 | 1.34 | 1.19E-08 |
| ENSMUSG00000066129 | GSMG0035308 | Kndc1 | kinase non-catalytic C-lobe domain (KIND) containing 1 | 574.42 | 311.97 | 0.88 | 1.71E-01 |
| ENSMUSG00000025470 | GSMG0035311 | Zfp511 | zinc finger protein 511 | 700.44 | 726.91 | -0.05 | 8.72E-01 |
| ENSMUSG00000039018 | GSMG0035314 | Mtg1 | mitochondrial ribosome-associated GTPase 1 | 437.40 | 417.87 | 0.07 | 8.34E-01 |
| ENSMUSG00000025485 | GSMG0035332 | Ric8a | RIC8 guanine nucleotide exchange factor A | 1965.69 | 1953.63 | 0.01 | 9.74E-01 |
| ENSMUSG00000025487 | GSMG0035333 | Psmd13 | proteasome (prosome, macropain) 26S subunit, non-ATPase, 13 | 1999.92 | 2374.05 | -0.25 | 1.71E-01 |
| ENSMUSG00000062031 | GSMG0035336 | Athl1 | ATH1, acid trehalase-like 1 (yeast) | 411.59 | 292.68 | 0.50 | 5.57E-02 |
| ENSMUSG00000025491 | GSMG0035337 | Ifitm1 | interferon induced transmembrane protein 1 | 619.37 | 438.22 | 0.50 | 4.72E-02 |
| ENSMUSG00000055629 | GSMG0035338 | B4galnt4 | beta-1,4-N-acetyl-galactosaminyl transferase 4 | 1913.54 | 2483.33 | -0.38 | 2.34E-01 |
| ENSMUSG00000025495 | GSMG0035341 | Ptdss2 | phosphatidylserine synthase 2 | 778.76 | 878.93 | -0.17 | 5.01E-01 |
| ENSMUSG00000038611 | GSMG0035347 | Phrf1 | PHD and ring finger domains 1 | 429.33 | 466.23 | -0.12 | 7.11E-01 |
| ENSMUSG00000002957 | GSMG0035349 | Ap2a2 | adaptor-related protein complex 2, alpha 2 subunit | 3889.49 | 4451.27 | -0.19 | 2.87E-01 |
| ENSMUSG00000025505 | GSMG0035351 | Tmem80 | transmembrane protein 80 | 807.47 | 622.46 | 0.37 | 4.75E-02 |
| ENSMUSG00000025504 | GSMG0035352 | Eps8l2 | EPS8-like 2 | 663.50 | 60.15 | 3.47 | 2.79E-53 |
| ENSMUSG00000025503 | GSMG0035354 | Taldo1 | transaldolase 1 | 3331.21 | 3556.44 | -0.09 | 7.42E-01 |
| ENSMUSG00000025509 | GSMG0035356 | Pnpla2 | patatin-like phospholipase domain containing 2 | 2375.00 | 1475.97 | 0.69 | 1.08E-06 |
| ENSMUSG00000048200 | GSMG0035357 | Cracr2b | calcium release activated channel regulator 2B | 694.51 | 517.10 | 0.42 | 2.32E-02 |
| ENSMUSG00000025510 | GSMG0035358 | Cd151 | CD151 antigen | 840.40 | 863.62 | -0.04 | 9.65E-01 |

| | | | | | | | |
|---|---|---|---|---|---|---|---|
| ENSMUSG00000025511 | GSMG0035359 | Tspan4 | tetraspanin 4 | 1801.98 | 1680.03 | 0.10 | 6.91E-01 |
| ENSMUSG00000053046 | GSMG0035366 | Brsk2 | BR serine/threonine kinase 2 | 547.50 | 439.63 | 0.32 | 6.99E-01 |
| ENSMUSG00000018819 | GSMG0035374 | Lsp1 | lymphocyte specific 1 | 88.25 | 83.48 | 0.08 | 8.87E-01 |
| ENSMUSG00000037706 | GSMG0035380 | Cd81 | CD81 antigen | 11485.71 | 10912.17 | 0.07 | 7.84E-01 |
| ENSMUSG00000045752 | GSMG0035381 | Tssc4 | tumor-suppressing subchromosomal transferable fragment 4 | 827.53 | 907.49 | -0.13 | 5.65E-01 |
| ENSMUSG00000009545 | GSMG0035382 | Kcnq1 | potassium voltage-gated channel, subfamily Q, member 1 | 792.80 | 1113.30 | -0.49 | 2.54E-03 |
| ENSMUSG00000058454 | GSMG0035391 | Dhcr7 | 7-dehydrocholesterol reductase | 613.37 | 840.59 | -0.45 | 3.28E-02 |
| ENSMUSG00000037541 | GSMG0035392 | Shank2 | SH3/ankyrin domain gene 2 | 419.24 | 543.82 | -0.38 | 1.34E-01 |
| NULL | GSMG0035393 | NULL | NULL | 583.60 | 414.62 | 0.50 | 2.57E-02 |
| NULL | GSMG0035395 | NULL | NULL | 368.66 | 374.36 | -0.03 | 9.50E-01 |
| ENSMUSG00000031072 | GSMG0035404 | Oraov1 | oral cancer overexpressed 1 | 167.61 | 200.04 | -0.26 | 4.08E-01 |
| ENSMUSG00000107585 | GSMG0035412 | 3300002P13Rik | RIKEN cDNA 3300002P13 gene | 135.46 | 111.81 | 0.27 | 5.13E-01 |
| ENSMUSG00000006335 | GSMG0035416 | Tfpt | TCF3 (E2A) fusion partner | 466.77 | 431.75 | 0.11 | 7.18E-01 |
| ENSMUSG00000035596 | GSMG0035419 | Mboat7 | membrane bound O-acyltransferase domain containing 7 | 281.43 | 358.25 | -0.35 | 2.12E-01 |
| ENSMUSG00000063838 | GSMG0035428 | Cdc42ep5 | CDC42 effector protein (Rho GTPase binding) 5 | 73.98 | 84.55 | -0.19 | 6.91E-01 |
| ENSMUSG00000008435 | GSMG0035432 | Rdh13 | retinol dehydrogenase 13 (all-trans and 9-cis) | 276.07 | 271.75 | 0.02 | 9.49E-01 |
| ENSMUSG00000019254 | GSMG0035434 | Ppp1r12c | protein phosphatase 1, regulatory (inhibitor) subunit 12C | 2723.87 | 1782.20 | 0.61 | 6.57E-03 |
| ENSMUSG00000004961 | GSMG0035438 | Syt5 | synaptotagmin V | 1264.90 | 2304.01 | -0.87 | 6.61E-08 |
| ENSMUSG00000045282 | GSMG0035440 | Tmem86b | transmembrane protein 86B | 259.11 | 138.28 | 0.90 | 3.05E-04 |
| ENSMUSG00000052296 | GSMG0035441 | Ppp6r1 | protein phosphatase 6, regulatory subunit 1 | 4395.82 | 3431.60 | 0.36 | 1.10E-01 |
| ENSMUSG00000063802 | GSMG0035442 | Hspbp1 | HSPA (heat shock 70kDa) binding protein, cytoplasmic cochaperone 1 | 1096.76 | 1438.29 | -0.39 | 1.73E-02 |
| ENSMUSG00000051811 | GSMG0035445 | Cox6b2 | cytochrome c oxidase subunit VIb polypeptide 2 | 113.36 | 106.03 | 0.09 | 8.61E-01 |
| ENSMUSG00000030431 | GSMG0035448 | Tmem238 | transmembrane protein 238 | 290.92 | 264.10 | 0.13 | 7.64E-01 |
| ENSMUSG00000053550 | GSMG0035450 | Shisa7 | shisa family member 7 | 771.39 | 537.59 | 0.52 | 2.59E-02 |
| ENSMUSG00000051550 | GSMG0035453 | Zfp579 | zinc finger protein 579 | 209.84 | 295.16 | -0.50 | 5.08E-02 |
| ENSMUSG00000061374 | GSMG0035454 | Fiz1 | Flt3 interacting zinc finger protein 1 | 897.99 | 1113.29 | -0.31 | 1.04E-01 |
| ENSMUSG00000043290 | GSMG0035456 | Zfp784 | zinc finger protein 784 | 232.67 | 177.12 | 0.39 | 1.84E-01 |
| ENSMUSG00000046792 | GSMG0035468 | Zfp787 | zinc finger protein 787 | 464.65 | 602.07 | -0.38 | 7.08E-02 |
| NULL | GSMG0035470 | NULL | NULL | 114.38 | 101.92 | 0.17 | 7.16E-01 |
| ENSMUSG00000002265 | GSMG0035480 | Peg3 | paternally expressed 3 | 212.28 | 204.67 | 0.05 | 9.00E-01 |
| ENSMUSG00000062116 | GSMG0035482 | Zfp954 | zinc finger protein 954 | 508.57 | 405.50 | 0.33 | 1.46E-01 |
| ENSMUSG00000066838 | GSMG0035484 | Zfp772 | zinc finger protein 772 | 387.62 | 400.57 | -0.05 | 8.99E-01 |
| ENSMUSG00000000605 | GSMG0035488 | Clcn4 | chloride channel, voltage-sensitive 4 | 4061.40 | 4879.37 | -0.26 | 1.29E-01 |

| | | | | | | | |
|---|---|---|---|---|---|---|---|
| ENSMUSG00000030393 | GSMG0035497 | Zik1 | zinc finger protein interacting with K protein 1 | 502.30 | 645.48 | -0.36 | 6.35E-02 |
| ENSMUSG00000034071 | GSMG0035505 | Zfp551 | zinc fingr protein 551 | 172.52 | 252.01 | -0.54 | 3.32E-02 |
| ENSMUSG00000070822 | GSMG0035506 | Zscan18 | zinc finger and SCAN domain containing 18 | 612.09 | 1185.64 | -0.95 | 3.70E-10 |
| ENSMUSG00000057894 | GSMG0035508 | Zfp329 | zinc finger protein 329 | 310.08 | 403.34 | -0.38 | 1.17E-01 |
| ENSMUSG00000049600 | GSMG0035513 | Zbtb45 | zinc finger and BTB domain containing 45 | 221.17 | 257.11 | -0.22 | 5.22E-01 |
| NULL | GSMG0035515 | NULL | NULL | 121.75 | 138.85 | -0.18 | 7.03E-01 |
| ENSMUSG00000033916 | GSMG0035516 | Chmp2a | charged multivesicular body protein 2A | 3121.73 | 3145.46 | -0.01 | 9.70E-01 |
| ENSMUSG00000005575 | GSMG0035517 | Ube2m | ubiquitin-conjugating enzyme E2M | 1177.40 | 950.94 | 0.31 | 3.74E-01 |
| ENSMUSG00000030380 | GSMG0035518 | Mzf1 | myeloid zinc finger 1 | 192.08 | 185.25 | 0.06 | 9.15E-01 |
| ENSMUSG00000041571 | GSMG0035530 | Sepw1 | selenoprotein W, muscle 1 | 6155.39 | 6759.99 | -0.14 | 6.81E-01 |
| ENSMUSG00000041560 | GSMG0035531 | Gltscr2 | glioma tumor suppressor candidate region gene 2 | 2029.24 | 2235.63 | -0.14 | 5.32E-01 |
| ENSMUSG00000074364 | GSMG0035532 | Ehd2 | EH-domain containing 2 | 1883.07 | 1117.01 | 0.75 | 1.83E-05 |
| ENSMUSG00000006019 | GSMG0035535 | Dhx34 | DEAH (Asp-Glu-Ala-His) box polypeptide 34 | 1070.90 | 1010.30 | 0.08 | 7.94E-01 |
| ENSMUSG00000091811 | GSMG0035538 | Inafm1 | InaF motif containing 1 | 673.81 | 833.05 | -0.31 | 1.41E-01 |
| ENSMUSG00000041375 | GSMG0035539 | Ccdc9 | coiled-coil domain containing 9 | 683.76 | 618.77 | 0.14 | 6.81E-01 |
| ENSMUSG00000052833 | GSMG0035540 | Sae1 | SUMO1 activating enzyme subunit 1 | 1744.70 | 2402.09 | -0.46 | 4.76E-03 |
| ENSMUSG00000001988 | GSMG0035542 | Npas1 | neuronal PAS domain protein 1 | 149.30 | 246.90 | -0.72 | 1.40E-02 |
| ENSMUSG00000058230 | GSMG0035543 | Arhgap35 | Rho GTPase activating protein 35 | 1122.71 | 1263.30 | -0.17 | 6.14E-01 |
| NULL | GSMG0035545 | NULL | NULL | 229.05 | 235.46 | -0.05 | 9.20E-01 |
| ENSMUSG00000048920 | GSMG0035548 | Fkrp | fukutin related protein | 1021.56 | 922.30 | 0.15 | 5.53E-01 |
| ENSMUSG00000019370 | GSMG0035551 | Calm3 | calmodulin 3 | 3852.36 | 5846.73 | -0.60 | 2.85E-03 |
| NULL | GSMG0035552 | NULL | NULL | 214.55 | 288.42 | -0.43 | 8.99E-02 |
| ENSMUSG00000003099 | GSMG0035553 | Ppp5c | protein phosphatase 5, catalytic subunit | 2566.16 | 2657.11 | -0.05 | 8.45E-01 |
| ENSMUSG00000074358 | GSMG0035565 | Ccdc61 | coiled-coil domain containing 61 | 521.80 | 551.32 | -0.08 | 8.01E-01 |
| ENSMUSG00000030407 | GSMG0035571 | Qpctl | glutaminyl-peptide cyclotransferase-like | 342.76 | 421.32 | -0.30 | 2.70E-01 |
| ENSMUSG00000030406 | GSMG0035572 | Gipr | gastric inhibitory polypeptide receptor | 56.24 | 241.99 | -2.10 | 4.49E-14 |
| ENSMUSG00000030403 | GSMG0035575 | Vasp | vasodilator-stimulated phosphoprotein | 1394.03 | 1561.86 | -0.17 | 5.79E-01 |
| ENSMUSG00000047649 | GSMG0035580 | Cd3eap | CD3E antigen, epsilon polypeptide associated protein | 436.28 | 410.67 | 0.08 | 8.03E-01 |
| ENSMUSG00000030397 | GSMG0035583 | Mark4 | MAP/microtubule affinity regulating kinase 4 | 105.81 | 163.87 | -0.64 | 4.29E-02 |
| ENSMUSG00000051403 | GSMG0035585 | Ppp1r37 | protein phosphatase 1, regulatory subunit 37 | 1689.98 | 1287.40 | 0.39 | 2.16E-01 |
| ENSMUSG00000044709 | GSMG0035586 | Gemin7 | gem (nuclear organelle) associated protein 7 | 1363.43 | 1562.25 | -0.20 | 3.15E-01 |
| ENSMUSG00000061028 | GSMG0035587 | Clasrp | CLK4-associating serine/arginine rich protein | 4297.99 | 4055.07 | 0.08 | 7.58E-01 |
| NULL | GSMG0035588 | NULL | NULL | 339.28 | 337.71 | 0.01 | 9.85E-01 |

| | | | | | | | |
|---|---|---|---|---|---|---|---|
| ENSMUSG00000002983 | GSMG0035589 | Relb | avian reticuloendotheliosis viral (v-rel) oncogene related B | 352.35 | 417.44 | -0.24 | 4.08E-01 |
| ENSMUSG00000002981 | GSMG0035590 | Clptm1 | cleft lip and palate associated transmembrane protein 1 | 5898.66 | 6353.05 | -0.11 | 6.31E-01 |
| ENSMUSG00000002985 | GSMG0035595 | Apoe | apolipoprotein E | 38317.54 | 20178.67 | 0.93 | 1.73E-08 |
| ENSMUSG00000002984 | GSMG0035596 | Tomm40 | translocase of outer mitochondrial membrane 40 homolog (yeast) | 1500.25 | 1726.06 | -0.20 | 3.07E-01 |
| ENSMUSG00000062300 | GSMG0035597 | Nectin2 | nectin cell adhesion molecule 2 | 1013.26 | 1886.51 | -0.90 | 3.80E-10 |
| ENSMUSG00000002980 | GSMG0035598 | Bcam | basal cell adhesion molecule | 4475.20 | 2086.22 | 1.10 | 3.53E-12 |
| ENSMUSG00000053175 | GSMG0035601 | Bcl3 | B cell leukemia/lymphoma 3 | 245.93 | 199.44 | 0.30 | 3.77E-01 |
| ENSMUSG00000040511 | GSMG0035604 | Pvr | poliovirus receptor | 501.21 | 319.70 | 0.65 | 2.75E-02 |
| ENSMUSG00000050605 | GSMG0035614 | Zfp61 | zinc finger protein 61 | 237.63 | 240.25 | -0.01 | 9.75E-01 |
| ENSMUSG00000003380 | GSMG0035630 | Rabac1 | Rab acceptor 1 (prenylated) | 4401.47 | 3845.81 | 0.19 | 2.99E-01 |
| ENSMUSG00000040907 | GSMG0035631 | Atp1a3 | ATPase, Na+/K+ transporting, alpha 3 polypeptide | 8205.91 | 12919.43 | -0.65 | 3.27E-04 |
| ENSMUSG00000003378 | GSMG0035632 | Grik5 | glutamate receptor, ionotropic, kainate 5 (gamma 2) | 2133.12 | 1531.45 | 0.48 | 3.03E-02 |
| ENSMUSG00000008496 | GSMG0035636 | Pou2f2 | POU domain, class 2, transcription factor 2 | 159.57 | 175.86 | -0.15 | 7.62E-01 |
| ENSMUSG00000054499 | GSMG0035638 | Dedd2 | death effector domain-containing DNA binding protein 2 | 143.08 | 150.09 | -0.07 | 8.77E-01 |
| ENSMUSG00000057177 | GSMG0035639 | Gsk3a | glycogen synthase kinase 3 alpha | 531.41 | 490.08 | 0.12 | 7.80E-01 |
| ENSMUSG00000005447 | GSMG0035642 | Pafah1b3 | platelet-activating factor acetylhydrolase, isoform 1b, subunit 3 | 175.45 | 739.28 | -2.08 | 9.22E-32 |
| ENSMUSG00000003123 | GSMG0035644 | Lipe | lipase, hormone sensitive | 613.38 | 551.80 | 0.15 | 5.46E-01 |
| ENSMUSG00000074272 | GSMG0035647 | Ceacam1 | carcinoembryonic antigen-related cell adhesion molecule 1 | 598.54 | 253.75 | 1.24 | 1.60E-11 |
| ENSMUSG00000060376 | GSMG0035650 | Bckdha | branched chain ketoacid dehydrogenase E1, alpha polypeptide | 975.22 | 833.98 | 0.23 | 3.24E-01 |
| ENSMUSG00000061702 | GSMG0035653 | Tmem91 | transmembrane protein 91 | 318.48 | 211.33 | 0.59 | 4.99E-02 |
| ENSMUSG00000002608 | GSMG0035654 | Ccdc97 | coiled-coil domain containing 97 | 431.40 | 474.44 | -0.14 | 6.20E-01 |
| ENSMUSG00000040725 | GSMG0035655 | Hnrnpul1 | heterogeneous nuclear ribonucleoprotein U-like 1 | 1167.52 | 1499.27 | -0.36 | 8.73E-02 |
| ENSMUSG00000002602 | GSMG0035656 | Axl | AXL receptor tyrosine kinase | 5095.29 | 4202.53 | 0.28 | 1.07E-01 |
| ENSMUSG00000058709 | GSMG0035664 | Egln2 | egl-9 family hypoxia-inducible factor 2 | 2038.11 | 1981.42 | 0.04 | 9.02E-01 |
| G00000053291,ENSMUSG000 | GSMG0035665 | Rab4b // Mia | RAB4B, member RAS oncogene family // melanoma inhibitory activity | 1077.24 | 1239.48 | -0.20 | 3.42E-01 |
| ENSMUSG00000061479 | GSMG0035666 | Snrpa | small nuclear ribonucleoprotein polypeptide A | 1153.78 | 1507.08 | -0.39 | 2.75E-02 |
| ENSMUSG00000003752 | GSMG0035667 | Itpkc | inositol 1,4,5-trisphosphate 3-kinase C | 66.36 | 58.75 | 0.17 | 7.74E-01 |
| ENSMUSG00000040488 | GSMG0035669 | Ltbp4 | latent transforming growth factor beta binding protein 4 | 24076.43 | 5088.36 | 2.24 | 7.88E-29 |
| ENSMUSG00000089832 | GSMG0035670 | Shkbp1 | Sh3kbp1 binding protein 1 | 615.44 | 773.87 | -0.33 | 1.40E-01 |
| ENSMUSG00000011751 | GSMG0035671 | Sptbn4 | spectrin beta, non-erythrocytic 4 | 1114.78 | 556.38 | 1.00 | 1.17E-04 |
| NULL | GSMG0035673 | NULL | NULL | 164.27 | 19.77 | 3.06 | 2.21E-21 |
| ENSMUSG00000003363 | GSMG0035674 | Pld3 | phospholipase D family, member 3 | 4655.04 | 3597.79 | 0.37 | 4.35E-02 |
| ENSMUSG00000030603 | GSMG0035687 | Psmc4 | proteasome (prosome, macropain) 26S subunit, ATPase, 4 | 3955.87 | 4358.56 | -0.14 | 5.01E-01 |

| | | | | | | | |
|---|---|---|---|---|---|---|---|
| ENSMUSG00000003438 | GSMG0035692 | Timm50 | translocase of inner mitochondrial membrane 50 | 794.06 | 990.98 | -0.32 | 1.00E-01 |
| ENSMUSG00000003435 | GSMG0035693 | Supt5 | suppressor of Ty 5 | 4584.44 | 4919.66 | -0.10 | 6.90E-01 |
| ENSMUSG00000037552 | GSMG0035694 | Plekhg2 | pleckstrin homology domain containing, family G (with RhoGef domai | 1493.80 | 1517.28 | -0.02 | 9.51E-01 |
| ENSMUSG00000044786 | GSMG0035695 | Zfp36 | zinc finger protein 36 | 463.08 | 195.56 | 1.24 | 2.02E-10 |
| ENSMUSG00000003444 | GSMG0035696 | Med29 | mediator complex subunit 29 | 356.07 | 447.15 | -0.33 | 1.38E-01 |
| ENSMUSG00000109336 | GSMG0035697 | Samd4b | sterile alpha motif domain containing 4B | 1209.38 | 1217.21 | -0.01 | 9.83E-01 |
| ENSMUSG00000030602 | GSMG0035700 | Pak4 | p21 protein (Cdc42/Rac)-activated kinase 4 | 295.86 | 265.00 | 0.16 | 6.00E-01 |
| ENSMUSG00000045948 | GSMG0035703 | Mrps12 | mitochondrial ribosomal protein S12 | 695.54 | 634.90 | 0.13 | 6.30E-01 |
| ENSMUSG00000030595 | GSMG0035704 | Nfkbib | nuclear factor of kappa light polypeptide gene enhancer in B cells inhib | 431.10 | 530.79 | -0.30 | 1.73E-01 |
| NULL | GSMG0035705 | Gm19897 | predicted gene, 19897 | 285.14 | 155.34 | 0.87 | 1.46E-02 |
| ENSMUSG00000054808 | GSMG0035707 | Actn4 | actinin alpha 4 | 27776.54 | 29564.18 | -0.09 | 6.80E-01 |
| NULL | GSMG0035709 | NULL | NULL | 367.47 | 276.99 | 0.41 | 1.76E-01 |
| ENSMUSG00000053565 | GSMG0035710 | Eif3k | eukaryotic translation initiation factor 3, subunit K | 1465.89 | 1771.87 | -0.27 | 1.59E-01 |
| ENSMUSG00000030592 | GSMG0035711 | Ryr1 | ryanodine receptor 1, skeletal muscle | 172.58 | 254.34 | -0.56 | 7.44E-02 |
| ENSMUSG00000030590 | GSMG0035712 | Fam98c | family with sequence similarity 98, member C | 523.78 | 381.35 | 0.46 | 1.92E-02 |
| ENSMUSG00000037239 | GSMG0035713 | Spred3 | sprouty-related, EVH1 domain containing 3 | 78.11 | 52.21 | 0.58 | 1.88E-01 |
| ENSMUSG00000030591 | GSMG0035714 | Psmd8 | proteasome (prosome, macropain) 26S subunit, non-ATPase, 8 | 2376.71 | 2626.08 | -0.14 | 5.20E-01 |
| ENSMUSG00000074227 | GSMG0035718 | Spint2 | serine protease inhibitor, Kunitz type 2 | 286.35 | 487.46 | -0.77 | 9.55E-03 |
| ENSMUSG00000030583 | GSMG0035720 | Sipa1l3 | signal-induced proliferation-associated 1 like 3 | 107.53 | 164.67 | -0.62 | 1.14E-01 |
| ENSMUSG00000050855 | GSMG0035724 | Zfp940 | zinc finger protein 940 | 602.01 | 476.96 | 0.34 | 1.15E-01 |
| ENSMUSG00000062040 | GSMG0035727 | Zfp27 | zinc finger protein 27 | 486.13 | 462.35 | 0.07 | 8.12E-01 |
| ENSMUSG00000059975 | GSMG0035728 | Zfp74 | zinc finger protein 74 | 90.15 | 98.24 | -0.13 | 8.17E-01 |
| ENSMUSG00000053985 | GSMG0035729 | Zfp14 | zinc finger protein 14 | 627.86 | 541.82 | 0.21 | 3.95E-01 |
| ENSMUSG00000078768 | GSMG0035731 | Zfp566 | zinc finger protein 566 | 139.88 | 223.74 | -0.67 | 1.83E-02 |
| ENSMUSG00000001794 | GSMG0035734 | Capns1 | calpain, small subunit 1 | 1589.36 | 1335.72 | 0.25 | 3.81E-01 |
| ENSMUSG00000006095 | GSMG0035735 | Tbcb | tubulin folding cofactor B | 1104.70 | 1373.35 | -0.31 | 1.35E-01 |
| ENSMUSG00000037020 | GSMG0035736 | Wdr62 | WD repeat domain 62 | 176.95 | 280.11 | -0.66 | 4.52E-03 |
| ENSMUSG00000074211 | GSMG0035738 | Sdhaf1 | succinate dehydrogenase complex assembly factor 1 | 461.67 | 518.18 | -0.17 | 5.31E-01 |
| ENSMUSG00000036957 | GSMG0035740 | Lrfn3 | leucine rich repeat and fibronectin type III domain containing 3 | 100.00 | 151.24 | -0.61 | 2.20E-01 |
| ENSMUSG00000006651 | GSMG0035742 | Aplp1 | amyloid beta (A4) precursor-like protein 1 | 9346.19 | 14131.75 | -0.60 | 2.72E-04 |
| ENSMUSG00000036882 | GSMG0035747 | Arhgap33 | Rho GTPase activating protein 33 | 2812.17 | 3896.77 | -0.47 | 7.38E-02 |
| ENSMUSG00000036864 | GSMG0035748 | Proser3 | proline and serine rich 3 | 63.16 | 87.67 | -0.48 | 2.86E-01 |
| ENSMUSG00000036845 | GSMG0035749 | Lin37 | lin-37 homolog (C. elegans) | 575.50 | 457.38 | 0.33 | 1.12E-01 |

| | | | | | | | |
|---|---|---|---|---|---|---|---|
| ENSMUSG00000006307 | GSMG0035751 | Kmt2b | lysine (K)-specific methyltransferase 2B | 529.95 | 546.89 | -0.04 | 9.02E-01 |
| ENSMUSG00000036751 | GSMG0035754 | Cox6b1 | cytochrome c oxidase, subunit VIb polypeptide 1 | 4479.44 | 5075.01 | -0.18 | 4.43E-01 |
| G00000036733,ENSMUSG000 | GSMG0035755 | Rbm42 // Haus5 | RNA binding motif protein 42 // HAUS augmin-like complex, subunit 5 | 3169.87 | 3607.82 | -0.19 | 3.03E-01 |
| ENSMUSG00000006315 | GSMG0035757 | Tmem147 | transmembrane protein 147 | 2600.43 | 3096.60 | -0.25 | 1.72E-01 |
| ENSMUSG00000036634 | GSMG0035762 | Mag | myelin-associated glycoprotein | 1399.99 | 43.52 | 5.01 | 2.07E-93 |
| ENSMUSG00000058239 | GSMG0035765 | Usf2 | upstream transcription factor 2 | 269.19 | 202.58 | 0.41 | 1.07E-01 |
| ENSMUSG00000001247 | GSMG0035766 | Lsr | lipolysis stimulated lipoprotein receptor | 10058.57 | 8149.35 | 0.30 | 6.39E-02 |
| NULL | GSMG0035767 | NULL | NULL | 563.27 | 366.67 | 0.62 | 7.33E-03 |
| ENSMUSG00000009687 | GSMG0035768 | Fxyd5 | FXYD domain-containing ion transport regulator 5 | 3297.31 | 2281.80 | 0.53 | 4.61E-04 |
| ENSMUSG00000036578 | GSMG0035769 | Fxyd7 | FXYD domain-containing ion transport regulator 7 | 606.73 | 381.41 | 0.67 | 3.80E-03 |
| ENSMUSG00000036570 | GSMG0035770 | Fxyd1 | FXYD domain-containing ion transport regulator 1 | 455.01 | 188.78 | 1.27 | 5.23E-02 |
| ENSMUSG00000019194 | GSMG0035773 | Scn1b | sodium channel, voltage-gated, type I, beta | 858.31 | 262.49 | 1.71 | 2.31E-14 |
| ENSMUSG00000001248 | GSMG0035774 | Gramd1a | GRAM domain containing 1A | 2361.00 | 2823.22 | -0.26 | 2.28E-01 |
| ENSMUSG00000036459 | GSMG0035782 | Wtip | WT1-interacting protein | 898.92 | 318.52 | 1.50 | 2.43E-02 |
| ENSMUSG00000052997 | GSMG0035783 | Uba2 | ubiquitin-like modifier activating enzyme 2 | 4993.41 | 4625.95 | 0.11 | 6.38E-01 |
| ENSMUSG00000002635 | GSMG0035784 | Pdcd2l | programmed cell death 2-like | 547.55 | 630.30 | -0.20 | 4.21E-01 |
| ENSMUSG00000036427 | GSMG0035785 | Gpi1 | glucose phosphate isomerase 1 | 7084.33 | 6378.58 | 0.15 | 4.65E-01 |
| ENSMUSG00000066571 | GSMG0035786 | 4931406P16Rik | RIKEN cDNA 4931406P16 gene | 5347.06 | 5429.26 | -0.02 | 9.42E-01 |
| ENSMUSG00000066568 | GSMG0035789 | Lsm14a | LSM14A mRNA processing body assembly factor | 1552.94 | 1679.07 | -0.11 | 6.46E-01 |
| NULL | GSMG0035798 | NULL | NULL | 77.58 | 108.09 | -0.49 | 2.45E-01 |
| ENSMUSG00000056216 | GSMG0035799 | Cebpg | CCAAT/enhancer binding protein (C/EBP), gamma | 1126.63 | 564.20 | 1.00 | 4.60E-07 |
| ENSMUSG00000001802 | GSMG0035801 | Lrp3 | low density lipoprotein receptor-related protein 3 | 1102.39 | 760.06 | 0.54 | 4.76E-01 |
| ENSMUSG00000063808 | GSMG0035802 | Gpatch1 | G patch domain containing 1 | 1021.82 | 1059.65 | -0.05 | 8.55E-01 |
| ENSMUSG00000030493 | GSMG0035803 | Faap24 | Fanconi anemia core complex associated protein 24 | 225.49 | 323.20 | -0.52 | 4.01E-02 |
| ENSMUSG00000034875 | GSMG0035805 | Nudt19 | nudix (nucleoside diphosphate linked moiety X)-type motif 19 | 741.44 | 756.55 | -0.03 | 9.41E-01 |
| ENSMUSG00000030417 | GSMG0035808 | Pdcd5 | programmed cell death 5 | 613.43 | 817.77 | -0.42 | 2.68E-02 |
| ENSMUSG00000043671 | GSMG0035810 | Dpy19l3 | dpy-19-like 3 (C. elegans) | 261.74 | 418.22 | -0.68 | 8.89E-04 |
| ENSMUSG00000044452 | GSMG0035811 | Zfp507 | zinc finger protein 507 | 661.12 | 796.92 | -0.27 | 2.42E-01 |
| ENSMUSG00000043456 | GSMG0035813 | Zfp536 | zinc finger protein 536 | 466.94 | 372.07 | 0.33 | 2.13E-01 |
| ENSMUSG00000030421 | GSMG0035819 | Uri1 | URI1, prefoldin-like chaperone | 682.84 | 790.79 | -0.21 | 4.12E-01 |
| ENSMUSG00000002068 | GSMG0035820 | Ccne1 | cyclin E1 | 143.23 | 240.81 | -0.75 | 4.74E-03 |
| ENSMUSG00000030423 | GSMG0035823 | Pop4 | processing of precursor 4, ribonuclease P/MRP family, (S. cerevisiae) | 471.26 | 510.71 | -0.12 | 7.17E-01 |
| ENSMUSG00000056383 | GSMG0035837 | AI987944 | expressed sequence AI987944 | 496.97 | 381.94 | 0.38 | 7.47E-02 |

| | | | | | | | |
|---|---|---|---|---|---|---|---|
| ENSMUSG00000092416 | GSMG0035842 | Zfp141 | zinc finger protein 141 | 410.45 | 411.61 | 0.00 | 9.96E-01 |
| ENSMUSG00000012640 | GSMG0035851 | Zfp715 | zinc finger protein 715 | 1007.69 | 939.47 | 0.10 | 7.28E-01 |
| ENSMUSG00000008140 | GSMG0035864 | Emc10 | ER membrane protein complex subunit 10 | 1507.40 | 1828.37 | -0.28 | 1.60E-01 |
| ENSMUSG00000038644 | GSMG0035868 | Pold1 | polymerase (DNA directed), delta 1, catalytic subunit | 617.30 | 1024.45 | -0.73 | 2.13E-05 |
| ENSMUSG00000060601 | GSMG0035869 | Nr1h2 | nuclear receptor subfamily 1, group H, member 2 | 2066.77 | 1871.43 | 0.14 | 4.86E-01 |
| ENSMUSG00000038520 | GSMG0035875 | Tbc1d17 | TBC1 domain family, member 17 | 2196.00 | 2011.46 | 0.13 | 5.96E-01 |
| ENSMUSG00000038502 | GSMG0035876 | Ptov1 | prostate tumor over expressed gene 1 | 4864.98 | 6148.76 | -0.34 | 7.33E-02 |
| ENSMUSG00000002968 | GSMG0035877 | Med25 | mediator complex subunit 25 | 2331.56 | 2761.39 | -0.24 | 2.11E-01 |
| ENSMUSG00000060279 | GSMG0035879 | Ap2a1 | adaptor-related protein complex 2, alpha 1 subunit | 3506.65 | 3536.52 | -0.01 | 9.72E-01 |
| ENSMUSG00000007783 | GSMG0035881 | Cpt1c | carnitine palmitoyltransferase 1c | 3649.34 | 3020.19 | 0.27 | 2.07E-01 |
| ENSMUSG00000109324 | GSMG0035882 | Prmt1 | protein arginine N-methyltransferase 1 | 888.68 | 1710.03 | -0.94 | 1.15E-08 |
| ENSMUSG00000003190 | GSMG0035883 | Bcl2l12 | BCL2-like 12 (proline rich) | 179.45 | 230.03 | -0.36 | 2.24E-01 |
| ENSMUSG00000038406 | GSMG0035884 | Scaf1 | SR-related CTD-associated factor 1 | 1708.85 | 1962.77 | -0.20 | 5.17E-01 |
| ENSMUSG00000007837 | GSMG0035888 | Prrg2 | proline-rich Gla (G-carboxyglutamic acid) polypeptide 2 | 633.62 | 482.74 | 0.39 | 5.43E-02 |
| NULL | GSMG0035889 | NULL | NULL | 73.72 | 57.48 | 0.36 | 4.35E-01 |
| ENSMUSG00000019539 | GSMG0035890 | Rcn3 | reticulocalbin 3, EF-hand calcium binding domain | 1342.97 | 1393.30 | -0.05 | 8.63E-01 |
| ENSMUSG00000003420 | GSMG0035891 | Fcgrt | Fc receptor, IgG, alpha chain transporter | 9927.75 | 6002.69 | 0.73 | 1.85E-06 |
| ENSMUSG00000003429 | GSMG0035892 | Rps11 | ribosomal protein S11 | 2517.88 | 4014.66 | -0.67 | 3.66E-05 |
| ENSMUSG00000007833 | GSMG0035895 | Aldh16a1 | aldehyde dehydrogenase 16 family, member A1 | 262.89 | 246.91 | 0.09 | 8.04E-01 |
| ENSMUSG00000030792 | GSMG0035898 | Dkkl1 | dickkopf-like 1 | 52.15 | 84.80 | -0.70 | 9.33E-02 |
| ENSMUSG00000038260 | GSMG0035901 | Trpm4 | transient receptor potential cation channel, subfamily M, member 4 | 483.08 | 439.78 | 0.14 | 6.59E-01 |
| ENSMUSG00000003863 | GSMG0035902 | Ppfia3 | protein tyrosine phosphatase, receptor type, f polypeptide (PTPRF), inte | 1384.22 | 1281.70 | 0.11 | 7.47E-01 |
| ENSMUSG00000003872 | GSMG0035903 | Lin7b | lin-7 homolog B (C. elegans) | 1656.64 | 779.59 | 1.09 | 3.04E-11 |
| ENSMUSG00000063511 | GSMG0035904 | Snrnp70 | small nuclear ribonucleoprotein 70 (U1) | 5225.81 | 4045.57 | 0.37 | 1.14E-01 |
| ENSMUSG00000003868 | GSMG0035906 | Ruvbl2 | RuvB-like protein 2 | 1328.31 | 1844.17 | -0.47 | 9.16E-03 |
| ENSMUSG00000003873 | GSMG0035908 | Bax | BCL2-associated X protein | 2330.91 | 2740.47 | -0.23 | 1.91E-01 |
| ENSMUSG00000030824 | GSMG0035911 | Nucb1 | nucleobindin 1 | 4910.07 | 5116.44 | -0.06 | 8.24E-01 |
| ENSMUSG00000040435 | GSMG0035912 | Ppp1r15a | protein phosphatase 1, regulatory (inhibitor) subunit 15A | 903.52 | 586.42 | 0.62 | 5.57E-04 |
| ENSMUSG00000110357 | GSMG0035915 | A030001D20Rik | RIKEN cDNA A030001D20 gene | 748.56 | 684.36 | 0.13 | 6.96E-01 |
| ENSMUSG00000057342 | GSMG0035918 | Sphk2 | sphingosine kinase 2 | 323.02 | 311.00 | 0.06 | 8.85E-01 |
| ENSMUSG00000003269 | GSMG0035922 | Cyth2 | cytohesin 2 | 1889.02 | 1959.73 | -0.05 | 8.43E-01 |
| ENSMUSG00000058743 | GSMG0035923 | Kcnj14 | potassium inwardly-rectifying channel, subfamily J, member 14 | 84.81 | 50.50 | 0.75 | 6.32E-02 |
| ENSMUSG00000053801 | GSMG0035924 | Grwd1 | glutamate-rich WD repeat containing 1 | 208.39 | 415.23 | -0.99 | 2.35E-07 |

| | | | | | | | |
|---|---|---|---|---|---|---|---|
| ENSMUSG00000002771 | GSMG0035925 | Grin2d | glutamate receptor, ionotropic, NMDA2D (epsilon 4) | 233.61 | 466.84 | -1.00 | 2.86E-03 |
| ENSMUSG00000040212 | GSMG0035928 | Emp3 | epithelial membrane protein 3 | 654.61 | 482.37 | 0.44 | 1.82E-02 |
| ENSMUSG00000030834 | GSMG0035929 | Abcc6 | ATP-binding cassette, sub-family C (CFTR/MRP), member 6 | 3817.48 | 2061.14 | 0.89 | 2.98E-06 |
| ENSMUSG00000040136 | GSMG0035931 | Abcc8 | ATP-binding cassette, sub-family C (CFTR/MRP), member 8 | 387.23 | 248.91 | 0.64 | 1.21E-02 |
| ENSMUSG00000030839 | GSMG0035935 | Sergef | secretion regulating guanine nucleotide exchange factor | 351.65 | 320.05 | 0.14 | 6.55E-01 |
| ENSMUSG00000006763 | GSMG0035939 | Saal1 | serum amyloid A-like 1 | 291.01 | 341.21 | -0.23 | 3.83E-01 |
| ENSMUSG00000014418 | GSMG0035943 | Hps5 | Hermansky-Pudlak syndrome 5 | 948.88 | 1036.31 | -0.13 | 5.85E-01 |
| ENSMUSG00000014402 | GSMG0035946 | Tsg101 | tumor susceptibility gene 101 | 1059.72 | 1204.23 | -0.18 | 4.13E-01 |
| ENSMUSG00000049516 | GSMG0035950 | Spty2d1 | SPT2, Suppressor of Ty, domain containing 1 (S. cerevisiae) | 458.49 | 412.93 | 0.15 | 5.79E-01 |
| ENSMUSG00000030854 | GSMG0035951 | Ptpn5 | protein tyrosine phosphatase, non-receptor type 5 | 1716.98 | 1988.90 | -0.21 | 3.91E-01 |
| NULL | GSMG0035970 | NULL | NULL | 133.56 | 114.38 | 0.22 | 7.48E-01 |
| ENSMUSG00000092118 | GSMG0035978 | Fancf | Fanconi anemia, complementation group F | 184.52 | 228.28 | -0.30 | 3.09E-01 |
| ENSMUSG00000074093 | GSMG0035979 | Svip | small VCP/p97-interacting protein | 329.89 | 414.93 | -0.33 | 1.52E-01 |
| ENSMUSG00000030452 | GSMG0035985 | Nipa2 | non imprinted in Prader-Willi/Angelman syndrome 2 homolog (human) | 565.10 | 647.10 | -0.20 | 4.59E-01 |
| NULL | GSMG0035987 | NULL | NULL | 139.69 | 175.76 | -0.34 | 3.25E-01 |
| ENSMUSG00000047037 | GSMG0035988 | Nipa1 | non imprinted in Prader-Willi/Angelman syndrome 1 homolog (human) | 445.43 | 676.26 | -0.60 | 8.86E-04 |
| ENSMUSG00000055026 | GSMG0035993 | Gabrg3 | gamma-aminobutyric acid (GABA) A receptor, subunit gamma 3 | 547.73 | 439.65 | 0.32 | 1.37E-01 |
| NULL | GSMG0035995 | NULL | NULL | 339.66 | 246.88 | 0.46 | 1.03E-01 |
| ENSMUSG00000055078 | GSMG0035996 | Gabra5 | gamma-aminobutyric acid (GABA) A receptor, subunit alpha 5 | 289.39 | 737.79 | -1.35 | 9.12E-14 |
| NULL | GSMG0035997 | NULL | NULL | 225.04 | 50.81 | 2.14 | 6.68E-15 |
| NULL | GSMG0036014 | NULL | NULL | 568.80 | 429.50 | 0.40 | 1.09E-01 |
| NULL | GSMG0036026 | NULL | NULL | 189.86 | 106.14 | 0.84 | 6.68E-03 |
| ENSMUSG00000109006 | GSMG0036028 | B230209E15Rik | RIKEN cDNA B230209E15 gene | 259.51 | 221.13 | 0.23 | 5.64E-01 |
| ENSMUSG00000109554 | GSMG0036031 | A230057D06Rik | RIKEN cDNA A230057D06 gene | 237.60 | 206.65 | 0.20 | 5.38E-01 |
| NULL | GSMG0036032 | NULL | NULL | 833.50 | 716.02 | 0.22 | 3.18E-01 |
| NULL | GSMG0036035 | NULL | NULL | 250.51 | 202.52 | 0.30 | 4.21E-01 |
| NULL | GSMG0036037 | NULL | NULL | 274.58 | 232.27 | 0.24 | 4.76E-01 |
| ENSMUSG00000109321 | GSMG0036039 | A330076H08Rik | RIKEN cDNA A330076H08 gene | 473.96 | 314.31 | 0.59 | 8.60E-03 |
| NULL | GSMG0036044 | NULL | NULL | 468.71 | 620.81 | -0.40 | 8.67E-02 |
| NULL | GSMG0036046 | NULL | NULL | 1747.48 | 3119.50 | -0.84 | 6.99E-07 |
| ENSMUSG00000070527 | GSMG0036047 | Mkrn3 | makorin, ring finger protein, 3 | 250.37 | 319.61 | -0.36 | 2.07E-01 |
| ENSMUSG00000052040 | GSMG0036054 | Klf13 | Kruppel-like factor 13 | 2724.64 | 2688.18 | 0.02 | 9.61E-01 |
| ENSMUSG00000033458 | GSMG0036061 | Fan1 | FANCD2/FANCI-associated nuclease 1 | 398.04 | 333.99 | 0.25 | 3.33E-01 |

| | | | | | | | |
|---|---|---|---|---|---|---|---|
| ENSMUSG00000030521 | GSMG0036062 | Mphosph10 | M-phase phosphoprotein 10 (U3 small nucleolar ribonucleoprotein) | 745.50 | 988.56 | -0.41 | 1.87E-01 |
| ENSMUSG00000030518 | GSMG0036063 | Fam189a1 | family with sequence similarity 189, member A1 | 100.17 | 79.39 | 0.34 | 5.04E-01 |
| ENSMUSG00000070520 | GSMG0036064 | Ndnl2 | necdin-like 2 | 945.53 | 933.42 | 0.02 | 9.50E-01 |
| ENSMUSG00000030516 | GSMG0036068 | Tjp1 | tight junction protein 1 | 2596.58 | 2711.58 | -0.06 | 8.18E-01 |
| ENSMUSG00000015133 | GSMG0036075 | Lrrk1 | leucine-rich repeat kinase 1 | 240.00 | 184.44 | 0.38 | 2.41E-01 |
| ENSMUSG00000030509 | GSMG0036078 | Asb7 | ankyrin repeat and SOCS box-containing 7 | 63.00 | 85.55 | -0.44 | 3.54E-01 |
| ENSMUSG00000030557 | GSMG0036081 | Mef2a | myocyte enhancer factor 2A | 1754.75 | 1881.28 | -0.10 | 6.64E-01 |
| ENSMUSG00000030556 | GSMG0036087 | Lrrc28 | leucine rich repeat containing 28 | 285.49 | 341.55 | -0.26 | 3.03E-01 |
| ENSMUSG00000030554 | GSMG0036090 | Synm | synemin, intermediate filament protein | 2824.51 | 1989.47 | 0.51 | 2.44E-02 |
| ENSMUSG00000042659 | GSMG0036096 | Arrdc4 | arrestin domain containing 4 | 378.57 | 890.61 | -1.24 | 4.02E-14 |
| ENSMUSG00000030551 | GSMG0036099 | Nr2f2 | nuclear receptor subfamily 2, group F, member 2 | 405.96 | 347.55 | 0.22 | 4.43E-01 |
| NULL | GSMG0036113 | NULL | NULL | 295.69 | 278.54 | 0.08 | 8.46E-01 |
| NULL | GSMG0036116 | NULL | NULL | 429.59 | 520.10 | -0.28 | 2.13E-01 |
| ENSMUSG00000025789 | GSMG0036120 | St8sia2 | ST8 alpha-N-acetyl-neuraminide alpha-2,8-sialyltransferase 2 | 25.32 | 400.49 | -3.98 | 1.22E-47 |
| ENSMUSG00000025790 | GSMG0036124 | Slco3a1 | solute carrier organic anion transporter family, member 3a1 | 1997.45 | 1575.73 | 0.34 | 3.32E-02 |
| NULL | GSMG0036127 | NULL | NULL | 242.74 | 246.73 | -0.02 | 9.55E-01 |
| ENSMUSG00000053025 | GSMG0036132 | Sv2b | synaptic vesicle glycoprotein 2 b | 541.57 | 471.63 | 0.20 | 5.11E-01 |
| NULL | GSMG0036142 | NULL | NULL | 572.94 | 522.01 | 0.13 | 6.72E-01 |
| ENSMUSG00000030612 | GSMG0036145 | Mrpl46 | mitochondrial ribosomal protein L46 | 703.18 | 926.87 | -0.40 | 4.12E-02 |
| ENSMUSG00000030610 | GSMG0036146 | Det1 | de-etiolated homolog 1 (Arabidopsis) | 235.99 | 273.62 | -0.22 | 4.96E-01 |
| ENSMUSG00000030606 | GSMG0036149 | Hapln3 | hyaluronan and proteoglycan link protein 3 | 451.85 | 545.71 | -0.27 | 3.30E-01 |
| ENSMUSG00000030605 | GSMG0036150 | Mfge8 | milk fat globule-EGF factor 8 protein | 9849.82 | 9638.69 | 0.03 | 9.05E-01 |
| ENSMUSG00000039194 | GSMG0036151 | Rlbp1 | retinaldehyde binding protein 1 | 75.11 | 314.36 | -2.05 | 6.93E-20 |
| ENSMUSG00000039176 | GSMG0036153 | Polg | polymerase (DNA directed), gamma | 3193.02 | 3085.14 | 0.05 | 8.94E-01 |
| ENSMUSG00000050382 | GSMG0036157 | Kif7 | kinesin family member 7 | 148.01 | 132.07 | 0.16 | 7.07E-01 |
| ENSMUSG00000030545 | GSMG0036159 | Pex11a | peroxisomal biogenesis factor 11 alpha | 218.79 | 153.33 | 0.51 | 7.07E-02 |
| ENSMUSG00000039062 | GSMG0036161 | Anpep | alanyl (membrane) aminopeptidase | 843.26 | 850.81 | -0.01 | 9.73E-01 |
| ENSMUSG00000063801 | GSMG0036163 | Ap3s2 | adaptor-related protein complex 3, sigma 2 subunit | 935.45 | 1284.54 | -0.46 | 5.13E-03 |
| ENSMUSG00000039043 | GSMG0036165 | Arpin | actin-related protein 2/3 complex inhibitor | 257.19 | 264.97 | -0.04 | 9.20E-01 |
| ENSMUSG00000030541 | GSMG0036169 | Idh2 | isocitrate dehydrogenase 2 (NADP+), mitochondrial | 2077.79 | 2815.37 | -0.44 | 1.67E-02 |
| ENSMUSG00000030528 | GSMG0036171 | Blm | Bloom syndrome, RecQ helicase-like | 151.76 | 322.05 | -1.09 | 1.00E-06 |
| ENSMUSG00000030538 | GSMG0036172 | Cib1 | calcium and integrin binding 1 (calmyrin) | 713.50 | 642.65 | 0.15 | 5.48E-01 |
| ENSMUSG00000038930 | GSMG0036173 | Rccd1 | RCC1 domain containing 1 | 577.85 | 625.83 | -0.11 | 7.05E-01 |

| | | | | | | | |
|---|---|---|---|---|---|---|---|
| ENSMUSG00000030533 | GSMG0036174 | Unc45a | unc-45 myosin chaperone A | 2134.94 | 2077.41 | 0.04 | 8.94E-01 |
| ENSMUSG00000038886 | GSMG0036176 | Man2a2 | mannosidase 2, alpha 2 | 548.71 | 472.80 | 0.21 | 6.14E-01 |
| ENSMUSG00000053158 | GSMG0036177 | Fes | feline sarcoma oncogene | 1392.98 | 1098.31 | 0.34 | 4.36E-02 |
| ENSMUSG00000030530 | GSMG0036178 | Furin | furin (paired basic amino acid cleaving enzyme) | 2064.41 | 1548.47 | 0.41 | 1.95E-01 |
| ENSMUSG00000030527 | GSMG0036183 | Crtc3 | CREB regulated transcription coactivator 3 | 371.58 | 413.60 | -0.15 | 6.23E-01 |
| ENSMUSG00000030536 | GSMG0036184 | Iqgap1 | IQ motif containing GTPase activating protein 1 | 7007.13 | 6694.70 | 0.07 | 7.97E-01 |
| ENSMUSG00000025722 | GSMG0036185 | Wdr73 | WD repeat domain 73 | 322.61 | 429.72 | -0.41 | 5.56E-02 |
| ENSMUSG00000025723 | GSMG0036186 | Nmb | neuromedin B | 179.35 | 178.38 | 0.01 | 9.86E-01 |
| ENSMUSG00000025724 | GSMG0036188 | Sec11a | SEC11 homolog A, signal peptidase complex subunit | 1974.86 | 2210.58 | -0.16 | 4.94E-01 |
| ENSMUSG00000025586 | GSMG0036191 | Cpeb1 | cytoplasmic polyadenylation element binding protein 1 | 497.30 | 154.97 | 1.68 | 3.96E-13 |
| ENSMUSG00000062444 | GSMG0036192 | Ap3b2 | adaptor-related protein complex 3, beta 2 subunit | 2337.53 | 3153.97 | -0.43 | 6.36E-02 |
| ENSMUSG00000025813 | GSMG0036198 | Homer2 | homer scaffolding protein 2 | 200.30 | 145.43 | 0.47 | 1.08E-01 |
| ENSMUSG00000025102 | GSMG0036202 | 3110040N11Rik | RIKEN cDNA 3110040N11 gene | 243.54 | 330.73 | -0.44 | 7.49E-02 |
| ENSMUSG00000025104 | GSMG0036208 | Hdgfrp3 | hepatoma-derived growth factor, related protein 3 | 204.81 | 240.88 | -0.24 | 4.24E-01 |
| ENSMUSG00000070462 | GSMG0036226 | Mesdc1 | mesoderm development candidate 1 | 1175.10 | 1241.84 | -0.08 | 8.57E-01 |
| ENSMUSG00000038459 | GSMG0036228 | Abhd17c | abhydrolase domain containing 17C | 858.23 | 1527.64 | -0.83 | 5.88E-06 |
| ENSMUSG00000015709 | GSMG0036230 | Arnt2 | aryl hydrocarbon receptor nuclear translocator 2 | 405.33 | 305.36 | 0.41 | 8.38E-02 |
| ENSMUSG00000030630 | GSMG0036231 | Fah | fumarylacetoacetate hydrolase | 560.79 | 477.00 | 0.24 | 3.51E-01 |
| ENSMUSG00000030629 | GSMG0036232 | Zfand6 | zinc finger, AN1-type domain 6 | 2545.26 | 2602.81 | -0.03 | 9.25E-01 |
| ENSMUSG00000039428 | GSMG0036243 | Tmem135 | transmembrane protein 135 | 624.16 | 611.86 | 0.03 | 9.30E-01 |
| ENSMUSG00000039405 | GSMG0036247 | Prss23 | protease, serine 23 | 2140.93 | 545.29 | 1.97 | 3.36E-33 |
| NULL | GSMG0036248 | NULL | NULL | 129.99 | 52.13 | 1.32 | 8.62E-05 |
| ENSMUSG00000062797 | GSMG0036250 | l7Rn6 | lethal, Chr 7, Rinchik 6 | 507.21 | 571.37 | -0.17 | 6.11E-01 |
| ENSMUSG00000030619 | GSMG0036251 | Eed | embryonic ectoderm development | 520.52 | 572.08 | -0.14 | 6.35E-01 |
| ENSMUSG00000030615 | GSMG0036256 | Tmem126a | transmembrane protein 126A | 443.63 | 581.95 | -0.39 | 7.86E-02 |
| ENSMUSG00000030614 | GSMG0036257 | Tmem126b | transmembrane protein 126B | 320.78 | 280.52 | 0.19 | 5.93E-01 |
| ENSMUSG00000041343 | GSMG0036261 | Ankrd42 | ankyrin repeat domain 42 | 140.90 | 191.96 | -0.44 | 1.30E-01 |
| ENSMUSG00000041328 | GSMG0036264 | Pcf11 | PCF11 cleavage and polyadenylation factor subunit | 319.01 | 236.29 | 0.44 | 9.03E-02 |
| ENSMUSG00000074024 | GSMG0036266 | 4632427E13Rik | RIKEN cDNA 4632427E13 gene | 3564.25 | 3188.61 | 0.16 | 5.31E-01 |
| ENSMUSG00000035686 | GSMG0036279 | Thrsp | thyroid hormone responsive | 170.23 | 91.54 | 0.90 | 4.27E-03 |
| ENSMUSG00000035642 | GSMG0036280 | Aamdc | adipogenesis associated Mth938 domain containing | 513.41 | 438.36 | 0.23 | 3.37E-01 |
| ENSMUSG00000042797 | GSMG0036285 | Aqp11 | aquaporin 11 | 1370.58 | 609.89 | 1.17 | 3.65E-09 |
| ENSMUSG00000035547 | GSMG0036287 | Capn5 | calpain 5 | 197.81 | 264.26 | -0.42 | 1.21E-01 |

| | | | | | | | |
|---|---|---|---|---|---|---|---|
| ENSMUSG00000030760 | GSMG0036291 | Acer3 | alkaline ceramidase 3 | 201.93 | 152.91 | 0.40 | 1.54E-01 |
| ENSMUSG00000035401 | GSMG0036296 | Emsy | EMSY, BRCA2-interacting transcriptional repressor | 1073.77 | 994.18 | 0.11 | 6.39E-01 |
| ENSMUSG00000035354 | GSMG0036301 | Uvrag | UV radiation resistance associated gene | 711.87 | 622.33 | 0.19 | 4.30E-01 |
| ENSMUSG00000030747 | GSMG0036305 | Dgat2 | diacylglycerol O-acyltransferase 2 | 394.38 | 338.82 | 0.22 | 5.02E-01 |
| ENSMUSG00000070436 | GSMG0036309 | Serpinh1 | serine (or cysteine) peptidase inhibitor, clade H, member 1 | 11082.78 | 14239.31 | -0.36 | 1.08E-01 |
| NULL | GSMG0036310 | NULL | NULL | 28.88 | 50.75 | -0.79 | 1.42E-01 |
| ENSMUSG00000030744 | GSMG0036311 | Rps3 | ribosomal protein S3 | 3882.03 | 5293.79 | -0.45 | 3.54E-03 |
| ENSMUSG00000030737 | GSMG0036314 | Slco2b1 | solute carrier organic anion transporter family, member 2b1 | 5573.07 | 5656.46 | -0.02 | 9.43E-01 |
| ENSMUSG00000035227 | GSMG0036317 | Spcs2 | signal peptidase complex subunit 2 homolog (S. cerevisiae) | 4076.65 | 4861.57 | -0.25 | 2.18E-01 |
| ENSMUSG00000058761 | GSMG0036320 | Rnf169 | ring finger protein 169 | 1396.76 | 1207.18 | 0.21 | 4.16E-01 |
| ENSMUSG00000030726 | GSMG0036324 | Pold3 | polymerase (DNA-directed), delta 3, accessory subunit | 959.33 | 1295.26 | -0.43 | 2.02E-02 |
| ENSMUSG00000030718 | GSMG0036325 | Ppme1 | protein phosphatase methylesterase 1 | 763.58 | 753.09 | 0.02 | 9.57E-01 |
| ENSMUSG00000030701 | GSMG0036331 | Plekhb1 | pleckstrin homology domain containing, family B (evectins) member 1 | 748.56 | 365.55 | 1.03 | 1.60E-07 |
| ENSMUSG00000032875 | GSMG0036333 | Arhgef17 | Rho guanine nucleotide exchange factor (GEF) 17 | 385.98 | 394.94 | -0.04 | 9.43E-01 |
| ENSMUSG00000047767 | GSMG0036338 | Atg16l2 | autophagy related 16-like 2 (S. cerevisiae) | 1033.28 | 579.10 | 0.84 | 2.23E-06 |
| ENSMUSG00000032737 | GSMG0036344 | Inppl1 | inositol polyphosphate phosphatase-like 1 | 1497.05 | 1207.26 | 0.31 | 2.61E-01 |
| ENSMUSG00000078630 | GSMG0036347 | Tomt | transmembrane O-methyltransferase | 205.14 | 296.78 | -0.53 | 3.32E-02 |
| ENSMUSG00000064307 | GSMG0036349 | Lrrc51 | leucine rich repeat containing 51 | 117.97 | 110.24 | 0.10 | 8.32E-01 |
| ENSMUSG00000070427 | GSMG0036351 | Il18bp | interleukin 18 binding protein | 69.80 | 100.12 | -0.52 | 2.03E-01 |
| ENSMUSG00000070426 | GSMG0036352 | Rnf121 | ring finger protein 121 | 343.58 | 423.52 | -0.30 | 2.06E-01 |
| ENSMUSG00000063550 | GSMG0036355 | Nup98 | nucleoporin 98 | 320.92 | 346.37 | -0.11 | 7.52E-01 |
| ENSMUSG00000073982 | GSMG0036358 | Rhog | ras homolog family member G | 1445.54 | 1260.57 | 0.20 | 3.75E-01 |
| NULL | GSMG0036359 | NULL | NULL | 49.14 | 103.93 | -1.08 | 3.39E-03 |
| ENSMUSG00000030966 | GSMG0036363 | Trim21 | tripartite motif-containing 21 | 168.11 | 170.07 | -0.01 | 9.77E-01 |
| ENSMUSG00000073968 | GSMG0036365 | Trim68 | tripartite motif-containing 68 | 134.85 | 136.65 | -0.01 | 9.78E-01 |
| ENSMUSG00000043366 | GSMG0036368 | Olfr78 | olfactory receptor 78 | 461.88 | 74.00 | 2.65 | 9.35E-24 |
| ENSMUSG00000057143 | GSMG0036413 | Trim12c | tripartite motif-containing 12C | 131.96 | 140.14 | -0.08 | 8.87E-01 |
| ENSMUSG00000030921 | GSMG0036416 | Trim30a | tripartite motif-containing 30A | 82.65 | 86.06 | -0.05 | 9.41E-01 |
| ENSMUSG00000044465 | GSMG0036432 | Fam160a2 | family with sequence similarity 160, member A2 | 1531.11 | 1255.00 | 0.29 | 1.66E-01 |
| ENSMUSG00000037060 | GSMG0036433 | Prkcdbp | protein kinase C, delta binding protein | 1615.41 | 1088.40 | 0.57 | 1.10E-03 |
| NULL | GSMG0036434 | NULL | NULL | 100.87 | 73.83 | 0.45 | 2.41E-01 |
| ENSMUSG00000037032 | GSMG0036435 | Apbb1 | amyloid beta (A4) precursor protein-binding, family B, member 1 | 2215.11 | 2362.71 | -0.09 | 7.05E-01 |
| ENSMUSG00000030881 | GSMG0036437 | Arfip2 | ADP-ribosylation factor interacting protein 2 | 1015.91 | 1003.88 | 0.02 | 9.57E-01 |

| | | | | | | | |
|---|---|---|---|---|---|---|---|
| ENSMUSG00000036989 | GSMG0036438 | Trim3 | tripartite motif-containing 3 | 3654.78 | 4022.63 | -0.14 | 5.79E-01 |
| ENSMUSG00000030888 | GSMG0036440 | Rrp8 | ribosomal RNA processing 8, methyltransferase, homolog (yeast) | 406.29 | 507.13 | -0.32 | 1.36E-01 |
| ENSMUSG00000030894 | GSMG0036442 | Tpp1 | tripeptidyl peptidase I | 3509.38 | 3340.15 | 0.07 | 7.92E-01 |
| ENSMUSG00000030879 | GSMG0036446 | Mrpl17 | mitochondrial ribosomal protein L17 | 1018.60 | 1268.00 | -0.32 | 9.20E-02 |
| NULL | GSMG0036453 | NULL | NULL | 224.12 | 112.03 | 1.00 | 2.26E-04 |
| ENSMUSG00000048330 | GSMG0036492 | Ric3 | RIC3 acetylcholine receptor chaperone | 87.96 | 118.00 | -0.43 | 2.25E-01 |
| ENSMUSG00000036111 | GSMG0036493 | Lmo1 | LIM domain only 1 | 88.51 | 240.12 | -1.45 | 2.33E-06 |
| ENSMUSG00000031024 | GSMG0036496 | St5 | suppression of tumorigenicity 5 | 2163.83 | 2586.15 | -0.26 | 1.53E-01 |
| ENSMUSG00000031021 | GSMG0036500 | Tmem9b | TMEM9 domain family, member B | 3482.28 | 3146.08 | 0.15 | 5.56E-01 |
| ENSMUSG00000034825 | GSMG0036501 | Nrip3 | nuclear receptor interacting protein 3 | 231.11 | 170.39 | 0.44 | 1.59E-01 |
| ENSMUSG00000007279 | GSMG0036503 | Scube2 | signal peptide, CUB domain, EGF-like 2 | 312.65 | 120.48 | 1.38 | 1.61E-05 |
| ENSMUSG00000035901 | GSMG0036504 | Dennd5a | DENN/MADD domain containing 5A | 3968.31 | 2676.30 | 0.57 | 2.39E-02 |
| ENSMUSG00000030788 | GSMG0036522 | Rnf141 | ring finger protein 141 | 243.71 | 247.21 | -0.02 | 9.56E-01 |
| ENSMUSG00000030787 | GSMG0036523 | Lyve1 | lymphatic vessel endothelial hyaluronan receptor 1 | 106.35 | 84.74 | 0.33 | 5.69E-01 |
| ENSMUSG00000005611 | GSMG0036524 | Mrvi1 | MRV integration site 1 | 934.98 | 698.49 | 0.42 | 3.09E-02 |
| ENSMUSG00000005610 | GSMG0036528 | Eif4g2 | eukaryotic translation initiation factor 4, gamma 2 | 27986.35 | 36735.10 | -0.39 | 6.60E-03 |
| ENSMUSG00000038296 | GSMG0036531 | Galnt18 | UDP-N-acetyl-alpha-D-galactosamine:polypeptide N-acetylgalactosami | 1480.95 | 1466.77 | 0.01 | 9.63E-01 |
| NULL | GSMG0036533 | NULL | NULL | 222.99 | 163.51 | 0.45 | 1.68E-01 |
| ENSMUSG00000030772 | GSMG0036535 | Dkk3 | dickkopf WNT signaling pathway inhibitor 3 | 461.34 | 582.21 | -0.33 | 1.28E-01 |
| ENSMUSG00000038187 | GSMG0036540 | Btbd10 | BTB (POZ) domain containing 10 | 1232.77 | 1232.43 | 0.00 | 9.97E-01 |
| ENSMUSG00000055723 | GSMG0036546 | Rras2 | related RAS viral (r-ras) oncogene 2 | 1787.00 | 1897.38 | -0.09 | 7.59E-01 |
| ENSMUSG00000030754 | GSMG0036548 | Copb1 | coatomer protein complex, subunit beta 1 | 1955.80 | 2093.78 | -0.10 | 7.08E-01 |
| ENSMUSG00000030751 | GSMG0036549 | Psma1 | proteasome (prosome, macropain) subunit, alpha type 1 | 2198.33 | 2471.56 | -0.17 | 4.26E-01 |
| ENSMUSG00000051910 | GSMG0036557 | Sox6 | SRY (sex determining region Y)-box 6 | 268.61 | 385.49 | -0.52 | 1.55E-02 |
| ENSMUSG00000045659 | GSMG0036561 | Plekha7 | pleckstrin homology domain containing, family A member 7 | 437.66 | 310.68 | 0.49 | 9.68E-02 |
| ENSMUSG00000030660 | GSMG0036563 | Pik3c2a | phosphatidylinositol 3-kinase, C2 domain containing, alpha polypeptide | 1987.16 | 1704.92 | 0.22 | 2.50E-01 |
| ENSMUSG00000008683 | GSMG0036566 | Rps15a | ribosomal protein S15A | 631.20 | 1034.90 | -0.71 | 1.66E-04 |
| ENSMUSG00000030654 | GSMG0036567 | Arl6ip1 | ADP-ribosylation factor-like 6 interacting protein 1 | 8045.70 | 11037.81 | -0.46 | 1.37E-02 |
| ENSMUSG00000058420 | GSMG0036572 | Syt17 | synaptotagmin XVII | 217.40 | 89.79 | 1.28 | 3.14E-07 |
| ENSMUSG00000095115 | GSMG0036573 | Itpripl2 | inositol 1,4,5-triphosphate receptor interacting protein-like 2 | 749.31 | 399.71 | 0.91 | 1.08E-05 |
| ENSMUSG00000030652 | GSMG0036574 | Coq7 | demethyl-Q 7 | 692.99 | 858.86 | -0.31 | 1.09E-01 |
| ENSMUSG00000042246 | GSMG0036575 | Tmc7 | transmembrane channel-like gene family 7 | 2127.90 | 1481.78 | 0.52 | 1.82E-03 |
| ENSMUSG00000033917 | GSMG0036577 | Gde1 | glycerophosphodiester phosphodiesterase 1 | 2191.82 | 1840.27 | 0.25 | 2.35E-01 |

| | | | | | | | |
|---|---|---|---|---|---|---|---|
| ENSMUSG00000030980 | GSMG0036579 | Knop1 | lysine rich nucleolar protein 1 | 637.14 | 671.30 | -0.07 | 7.96E-01 |
| ENSMUSG00000008734 | GSMG0036581 | Gprc5b | G protein-coupled receptor, family C, group 5, member B | 254.86 | 148.57 | 0.78 | 1.73E-03 |
| ENSMUSG00000030942 | GSMG0036590 | Thumpd1 | THUMP domain containing 1 | 1515.90 | 2122.76 | -0.49 | 5.54E-03 |
| ENSMUSG00000030929 | GSMG0036591 | Eri2 | exoribonuclease 2 | 102.70 | 119.77 | -0.23 | 6.06E-01 |
| ENSMUSG00000048787 | GSMG0036593 | Dcun1d3 | DCN1, defective in cullin neddylation 1, domain containing 3 (S. cerev | 312.16 | 209.70 | 0.57 | 1.24E-02 |
| ENSMUSG00000030905 | GSMG0036597 | Crym | crystallin, mu | 189.97 | 486.91 | -1.36 | 7.35E-06 |
| ENSMUSG00000030878 | GSMG0036604 | Cdr2 | cerebellar degeneration-related 2 | 387.60 | 497.18 | -0.36 | 1.58E-01 |
| ENSMUSG00000030872 | GSMG0036612 | Gga2 | golgi associated, gamma adaptin ear containing, ARF binding protein 2 | 743.53 | 1277.36 | -0.78 | 6.84E-07 |
| ENSMUSG00000034951 | GSMG0036614 | Cog7 | component of oligomeric golgi complex 7 | 736.21 | 697.27 | 0.08 | 8.07E-01 |
| ENSMUSG00000030871 | GSMG0036616 | Ears2 | glutamyl-tRNA synthetase 2, mitochondrial | 104.25 | 172.89 | -0.72 | 8.29E-03 |
| ENSMUSG00000108452 | GSMG0036625 | 4930413G21Rik | RIKEN cDNA 4930413G21 gene | 124.99 | 121.90 | 0.03 | 9.56E-01 |
| ENSMUSG00000030766 | GSMG0036626 | Arhgap17 | Rho GTPase activating protein 17 | 1215.37 | 1033.65 | 0.23 | 2.17E-01 |
| NULL | GSMG0036627 | NULL | NULL | 284.55 | 385.48 | -0.44 | 7.76E-02 |
| ENSMUSG00000030757 | GSMG0036628 | Zkscan2 | zinc finger with KRAB and SCAN domains 2 | 295.53 | 382.39 | -0.37 | 2.56E-01 |
| ENSMUSG00000030750 | GSMG0036634 | Nsmce1 | NSE1 homolog, SMC5-SMC6 complex component | 1281.63 | 1897.69 | -0.57 | 1.51E-03 |
| ENSMUSG00000032777 | GSMG0036636 | Gtf3c1 | general transcription factor III C 1 | 1253.21 | 1611.27 | -0.36 | 9.54E-02 |
| ENSMUSG00000046182 | GSMG0036638 | Gsg1l | GSG1-like | 93.05 | 79.98 | 0.21 | 6.79E-01 |
| ENSMUSG00000000131 | GSMG0036641 | Xpo6 | exportin 6 | 2870.52 | 3457.82 | -0.27 | 1.93E-01 |
| ENSMUSG00000030741 | GSMG0036647 | Spns1 | spinster homolog 1 | 1379.99 | 1428.82 | -0.05 | 8.67E-01 |
| ENSMUSG00000030733 | GSMG0036652 | Sh2b1 | SH2B adaptor protein 1 | 1289.07 | 1395.47 | -0.11 | 6.50E-01 |
| ENSMUSG00000032637 | GSMG0036653 | Atxn2l | ataxin 2-like | 4439.81 | 4704.40 | -0.08 | 7.83E-01 |
| ENSMUSG00000030738 | GSMG0036654 | Eif3c | eukaryotic translation initiation factor 3, subunit C | 3435.73 | 4195.38 | -0.29 | 1.02E-01 |
| ENSMUSG00000030720 | GSMG0036655 | Cln3 | ceroid lipofuscinosis, neuronal 3, juvenile (Batten, Spielmeyer-Vogt dis | 490.38 | 594.35 | -0.28 | 2.15E-01 |
| ENSMUSG00000030717 | GSMG0036657 | Nupr1 | nuclear protein transcription regulator 1 | 596.41 | 340.11 | 0.81 | 9.75E-06 |
| NULL | GSMG0036658 | NULL | NULL | 97.03 | 139.45 | -0.53 | 1.10E-01 |
| ENSMUSG00000030711 | GSMG0036659 | Sult1a1 | sulfotransferase family 1A, phenol-preferring, member 1 | 630.29 | 223.06 | 1.50 | 3.48E-15 |
| ENSMUSG00000059772 | GSMG0036661 | Slx1b | SLX1 structure-specific endonuclease subunit homolog B (S. cerevisiae | 188.07 | 280.42 | -0.58 | 1.02E-02 |
| ENSMUSG00000030707 | GSMG0036662 | Coro1a | coronin, actin binding protein 1A | 1639.21 | 1481.62 | 0.15 | 5.99E-01 |
| NULL | GSMG0036663 | NULL | NULL | 219.08 | 255.50 | -0.22 | 4.51E-01 |
| NULL | GSMG0036665 | NULL | NULL | 352.49 | 228.15 | 0.63 | 3.00E-03 |
| ENSMUSG00000030697 | GSMG0036666 | Ppp4c | protein phosphatase 4, catalytic subunit | 1661.54 | 2155.20 | -0.37 | 4.12E-02 |
| ENSMUSG00000030695 | GSMG0036667 | Aldoa | aldolase A, fructose-bisphosphate | 8617.09 | 5907.57 | 0.54 | 5.92E-04 |
| ENSMUSG00000030689 | GSMG0036670 | Ino80e | INO80 complex subunit E | 791.40 | 1165.20 | -0.56 | 3.08E-04 |

| | | | | | | | |
|---|---|---|---|---|---|---|---|
| ENSMUSG00000059981 | GSMG0036671 | Taok2 | TAO kinase 2 | 3182.54 | 2917.35 | 0.13 | 6.18E-01 |
| ENSMUSG00000060538 | GSMG0036672 | Tmem219 | transmembrane protein 219 | 442.85 | 388.44 | 0.19 | 4.51E-01 |
| ENSMUSG00000046378 | GSMG0036674 | Asphd1 | aspartate beta-hydroxylase domain containing 1 | 134.91 | 113.60 | 0.25 | 5.08E-01 |
| ENSMUSG00000030681 | GSMG0036676 | Mvp | major vault protein | 1107.73 | 750.72 | 0.56 | 2.14E-03 |
| G00000030680,ENSMUSG000 | GSMG0036677 | Pagr1a // Prrt2 | PAXIP1 associated glutamate rich protein 1A // proline-rich transmemb | 1717.53 | 1823.83 | -0.09 | 7.32E-01 |
| ENSMUSG00000030678 | GSMG0036678 | Maz | MYC-associated zinc finger protein (purine-binding transcription factor | 1453.57 | 1502.65 | -0.05 | 9.04E-01 |
| ENSMUSG00000030677 | GSMG0036679 | Kif22 | kinesin family member 22 | 501.67 | 1226.34 | -1.29 | 5.45E-09 |
| ENSMUSG00000042502 | GSMG0036684 | Cd2bp2 | CD2 antigen (cytoplasmic tail) binding protein 2 | 586.69 | 685.19 | -0.22 | 2.93E-01 |
| ENSMUSG00000042492 | GSMG0036685 | Tbc1d10b | TBC1 domain family, member 10b | 647.58 | 504.33 | 0.36 | 2.35E-01 |
| ENSMUSG00000000486 | GSMG0036687 | Sept1 | septin 1 | 117.53 | 206.11 | -0.82 | 2.22E-03 |
| NULL | GSMG0036688 | Gm4532 | predicted gene 4532 | 121.28 | 145.47 | -0.27 | 4.93E-01 |
| ENSMUSG00000042462 | GSMG0036689 | Dctpp1 | dCTP pyrophosphatase 1 | 272.10 | 508.45 | -0.90 | 9.81E-06 |
| ENSMUSG00000049091 | GSMG0036690 | Sephs2 | selenophosphate synthetase 2 | 636.43 | 941.28 | -0.57 | 8.29E-04 |
| ENSMUSG00000047371 | GSMG0036691 | Zfp768 | zinc finger protein 768 | 401.98 | 387.92 | 0.05 | 8.83E-01 |
| ENSMUSG00000054381 | GSMG0036692 | Zfp747 | zinc finger protein 747 | 352.42 | 252.94 | 0.48 | 3.52E-02 |
| ENSMUSG00000045757 | GSMG0036695 | Zfp764 | zinc finger protein 764 | 315.59 | 252.27 | 0.32 | 2.80E-01 |
| ENSMUSG00000045251 | GSMG0036696 | Zfp688 | zinc finger protein 688 | 323.57 | 308.21 | 0.07 | 8.57E-01 |
| ENSMUSG00000048921 | GSMG0036697 | Zfp689 | zinc finger protein 689 | 288.18 | 334.79 | -0.22 | 4.44E-01 |
| NULL | GSMG0036700 | NULL | NULL | 81.62 | 134.45 | -0.73 | 2.34E-02 |
| ENSMUSG00000057176 | GSMG0036701 | Ccdc189 | coiled-coil domain containing 189 | 125.15 | 71.90 | 0.80 | 2.78E-02 |
| ENSMUSG00000045639 | GSMG0036703 | Zfp629 | zinc finger protein 629 | 1228.58 | 1228.38 | 0.00 | 1.00E+00 |
| ENSMUSG00000030814 | GSMG0036705 | Bcl7c | B cell CLL/lymphoma 7C | 306.52 | 357.60 | -0.23 | 4.47E-01 |
| NULL | GSMG0036708 | NULL | NULL | 569.27 | 372.01 | 0.62 | 4.42E-03 |
| ENSMUSG00000030806 | GSMG0036709 | Stx1b | syntaxin 1B | 290.70 | 174.62 | 0.74 | 1.65E-03 |
| NULL | GSMG0036710 | NULL | NULL | 161.34 | 103.65 | 0.63 | 4.47E-02 |
| ENSMUSG00000049728 | GSMG0036713 | Zfp668 | zinc finger protein 668 | 129.40 | 162.29 | -0.32 | 3.27E-01 |
| ENSMUSG00000096145 | GSMG0036715 | Vkorc1 | vitamin K epoxide reductase complex, subunit 1 | 623.31 | 606.88 | 0.04 | 9.01E-01 |
| ENSMUSG00000062944 | GSMG0036725 | 9130023H24Rik | RIKEN cDNA 9130023H24 gene | 980.75 | 954.22 | 0.04 | 8.99E-01 |
| ENSMUSG00000030780 | GSMG0036726 | BC017158 | cDNA sequence BC017158 | 503.93 | 474.51 | 0.09 | 7.95E-01 |
| ENSMUSG00000030844 | GSMG0036727 | Rgs10 | regulator of G-protein signalling 10 | 254.27 | 196.74 | 0.37 | 2.34E-01 |
| ENSMUSG00000030846 | GSMG0036730 | Tial1 | Tia1 cytotoxic granule-associated RNA binding protein-like 1 | 741.91 | 753.06 | -0.02 | 9.46E-01 |
| ENSMUSG00000048170 | GSMG0036732 | Mcmbp | minichromosome maintenance complex binding protein | 1403.56 | 1510.60 | -0.11 | 6.58E-01 |
| ENSMUSG00000030849 | GSMG0036740 | Fgfr2 | fibroblast growth factor receptor 2 | 261.56 | 354.21 | -0.43 | 1.96E-01 |

| | | | | | | | |
|---|---|---|---|---|---|---|---|
| ENSMUSG00000030850 | GSMG0036742 | Ate1 | arginyltransferase 1 | 1975.28 | 2093.94 | -0.08 | 7.34E-01 |
| ENSMUSG00000040331 | GSMG0036745 | Nsmce4a | NSE4 homolog A, SMC5-SMC6 complex component | 1505.56 | 1663.84 | -0.14 | 5.38E-01 |
| ENSMUSG00000040177 | GSMG0036755 | 2310057M21Rik | RIKEN cDNA 2310057M21 gene | 219.31 | 263.85 | -0.27 | 3.24E-01 |
| ENSMUSG00000040167 | GSMG0036756 | Ikzf5 | IKAROS family zinc finger 5 | 145.31 | 142.22 | 0.03 | 9.46E-01 |
| NULL | GSMG0036758 | NULL | NULL | 157.48 | 213.27 | -0.44 | 1.60E-01 |
| ENSMUSG00000030930 | GSMG0036762 | Chst15 | carbohydrate (N-acetylgalactosamine 4-sulfate 6-O) sulfotransferase 15 | 282.97 | 396.58 | -0.49 | 6.64E-02 |
| ENSMUSG00000030934 | GSMG0036767 | Oat | ornithine aminotransferase | 1371.52 | 1350.44 | 0.02 | 9.41E-01 |
| ENSMUSG00000030956 | GSMG0036771 | Fam53b | family with sequence similarity 53, member B | 1080.47 | 548.09 | 0.98 | 4.19E-09 |
| NULL | GSMG0036772 | NULL | NULL | 653.57 | 360.49 | 0.86 | 8.91E-05 |
| ENSMUSG00000030960 | GSMG0036774 | Mettl10 | methyltransferase like 10 | 304.89 | 368.40 | -0.27 | 3.89E-01 |
| ENSMUSG00000030970 | GSMG0036777 | Ctbp2 | C-terminal binding protein 2 | 1120.52 | 941.84 | 0.25 | 2.33E-01 |
| NULL | GSMG0036778 | NULL | NULL | 343.30 | 325.14 | 0.07 | 8.70E-01 |
| ENSMUSG00000030979 | GSMG0036782 | Uros | uroporphyrinogen III synthase | 432.50 | 313.69 | 0.47 | 6.06E-02 |
| ENSMUSG00000031004 | GSMG0036802 | Mki67 | antigen identified by monoclonal antibody Ki 67 | 255.11 | 583.25 | -1.19 | 4.68E-09 |
| ENSMUSG00000040139 | GSMG0036811 | 9430038I01Rik | RIKEN cDNA 9430038I01 gene | 269.13 | 287.31 | -0.10 | 8.08E-01 |
| ENSMUSG00000091002 | GSMG0036815 | Tcerg1l | transcription elongation regulator 1-like | 76.51 | 167.68 | -1.13 | 5.90E-05 |
| ENSMUSG00000041775 | GSMG0036817 | Mapk1ip1 | mitogen-activated protein kinase 1 interacting protein 1 | 712.67 | 578.64 | 0.30 | 1.82E-01 |
| ENSMUSG00000078566 | GSMG0036818 | Bnip3 | BCL2/adenovirus E1B interacting protein 3 | 921.49 | 611.15 | 0.59 | 9.59E-04 |
| ENSMUSG00000025474 | GSMG0036830 | Tubgcp2 | tubulin, gamma complex associated protein 2 | 553.47 | 590.82 | -0.09 | 7.61E-01 |
| ENSMUSG00000025468 | GSMG0036832 | Caly | calcyon neuron-specific vesicular protein | 981.00 | 1453.64 | -0.57 | 1.86E-03 |
| ENSMUSG00000025465 | GSMG0036834 | Echs1 | enoyl Coenzyme A hydratase, short chain, 1, mitochondrial | 962.34 | 1173.47 | -0.29 | 1.73E-01 |
| ENSMUSG00000045733 | GSMG0036835 | Sprn | shadow of prion protein | 1816.85 | 1324.81 | 0.46 | 5.08E-02 |
| ENSMUSG00000060314 | GSMG0036844 | Zfp941 | zinc finger protein 941 | 509.07 | 240.34 | 1.08 | 1.13E-07 |
| ENSMUSG00000025484 | GSMG0036846 | Bet1l | Bet1 golgi vesicular membrane trafficking protein like | 1210.51 | 967.38 | 0.32 | 7.36E-02 |
| ENSMUSG00000025486 | GSMG0036847 | Sirt3 | sirtuin 3 | 631.74 | 455.04 | 0.47 | 3.14E-02 |
| ENSMUSG00000060591 | GSMG0036850 | Ifitm2 | interferon induced transmembrane protein 2 | 2191.35 | 1744.01 | 0.33 | 1.08E-01 |
| ENSMUSG00000025492 | GSMG0036851 | Ifitm3 | interferon induced transmembrane protein 3 | 2693.67 | 1764.85 | 0.61 | 1.17E-03 |
| ENSMUSG00000025494 | GSMG0036853 | Sigirr | single immunoglobulin and toll-interleukin 1 receptor (TIR) domain | 1994.10 | 1287.05 | 0.63 | 1.31E-04 |
| ENSMUSG00000025507 | GSMG0036855 | Pidd1 | p53 induced death domain protein 1 | 85.70 | 212.08 | -1.30 | 1.24E-06 |
| ENSMUSG00000038650 | GSMG0036856 | Rnh1 | ribonuclease/angiogenin inhibitor 1 | 1937.15 | 1436.46 | 0.43 | 2.60E-02 |
| ENSMUSG00000025499 | GSMG0036858 | Hras | Harvey rat sarcoma virus oncogene | 1256.02 | 1214.01 | 0.05 | 8.71E-01 |
| ENSMUSG00000025498 | GSMG0036860 | Irf7 | interferon regulatory factor 7 | 336.50 | 289.00 | 0.22 | 4.85E-01 |
| ENSMUSG00000058886 | GSMG0036863 | Deaf1 | deformed epidermal autoregulatory factor 1 (Drosophila) | 1886.87 | 2517.81 | -0.42 | 3.66E-02 |

| | | | | | | | |
|---|---|---|---|---|---|---|---|
| ENSMUSG00000051007 | GSMG0036865 | Pddc1 | Parkinson disease 7 domain containing 1 | 300.22 | 281.73 | 0.09 | 8.06E-01 |
| ENSMUSG00000060240 | GSMG0036866 | Cend1 | cell cycle exit and neuronal differentiation 1 | 1110.93 | 766.48 | 0.54 | 8.20E-03 |
| ENSMUSG00000019082 | GSMG0036867 | Slc25a22 | solute carrier family 25 (mitochondrial carrier, glutamate), member 22 | 808.48 | 857.06 | -0.08 | 7.79E-01 |
| ENSMUSG00000025512 | GSMG0036874 | Chid1 | chitinase domain containing 1 | 376.51 | 373.11 | 0.02 | 9.68E-01 |
| NULL | GSMG0036877 | NULL | NULL | 264.21 | 278.08 | -0.07 | 8.55E-01 |
| ENSMUSG00000025139 | GSMG0036879 | Tollip | toll interacting protein | 862.34 | 829.43 | 0.06 | 8.73E-01 |
| ENSMUSG00000025147 | GSMG0036881 | Mob2 | MOB kinase activator 2 | 2066.26 | 1617.27 | 0.35 | 3.39E-02 |
| ENSMUSG00000037887 | GSMG0036884 | Dusp8 | dual specificity phosphatase 8 | 544.55 | 461.32 | 0.24 | 4.09E-01 |
| ENSMUSG00000007891 | GSMG0036890 | Ctsd | cathepsin D | 10889.17 | 7714.25 | 0.50 | 7.79E-03 |
| ENSMUSG00000048583 | GSMG0036894 | Igf2 | insulin-like growth factor 2 | 2639.69 | 2599.32 | 0.02 | 9.56E-01 |
| ENSMUSG00000059277 | GSMG0036899 | R74862 | expressed sequence R74862 | 134.53 | 153.40 | -0.18 | 6.43E-01 |
| ENSMUSG00000037664 | GSMG0036912 | Cdkn1c | cyclin-dependent kinase inhibitor 1C (P57) | 2540.66 | 1057.70 | 1.26 | 2.47E-12 |
| ENSMUSG00000059119 | GSMG0036914 | Nap1l4 | nucleosome assembly protein 1-like 4 | 5926.83 | 7437.81 | -0.33 | 5.47E-02 |
| ENSMUSG00000010755 | GSMG0036915 | Cars | cysteinyl-tRNA synthetase | 2313.37 | 2970.08 | -0.36 | 2.06E-02 |
| ENSMUSG00000010751 | GSMG0036917 | Tnfrsf22 | tumor necrosis factor receptor superfamily, member 22 | 538.32 | 579.69 | -0.11 | 7.18E-01 |
| ENSMUSG00000037606 | GSMG0036919 | Osbpl5 | oxysterol binding protein-like 5 | 331.56 | 488.67 | -0.56 | 1.97E-02 |
| ENSMUSG00000048965 | GSMG0036922 | Mrgpre | MAS-related GPR, member E | 567.43 | 650.59 | -0.20 | 4.49E-01 |
| ENSMUSG00000031090 | GSMG0036923 | Nadsyn1 | NAD synthetase 1 | 372.27 | 311.34 | 0.26 | 3.33E-01 |
| NULL | GSMG0036925 | NULL | NULL | 82.47 | 90.17 | -0.14 | 8.09E-01 |
| ENSMUSG00000031078 | GSMG0036928 | Cttn | cortactin | 2003.57 | 2218.74 | -0.15 | 4.50E-01 |
| ENSMUSG00000037519 | GSMG0036929 | Ppfia1 | protein tyrosine phosphatase, receptor type, f polypeptide (PTPRF), inte | 651.34 | 577.23 | 0.17 | 5.96E-01 |
| ENSMUSG00000031075 | GSMG0036931 | Ano1 | anoctamin 1, calcium activated chloride channel | 842.24 | 474.18 | 0.83 | 1.32E-06 |
| ENSMUSG00000048677 | GSMG0036937 | Tpcn2 | two pore segment channel 2 | 338.96 | 271.92 | 0.32 | 3.30E-01 |
| ENSMUSG00000004568 | GSMG0036940 | Arhgef18 | rho/rac guanine nucleotide exchange factor (GEF) 18 | 690.64 | 652.78 | 0.08 | 8.25E-01 |
| ENSMUSG00000047264 | GSMG0036944 | Zfp358 | zinc finger protein 358 | 1027.05 | 1120.44 | -0.12 | 5.87E-01 |
| ENSMUSG00000004567 | GSMG0036945 | Mcoln1 | mucolipin 1 | 3059.74 | 3010.16 | 0.02 | 9.32E-01 |
| ENSMUSG00000004565 | GSMG0036946 | Pnpla6 | patatin-like phospholipase domain containing 6 | 6706.26 | 6338.06 | 0.08 | 7.69E-01 |
| ENSMUSG00000044433 | GSMG0036949 | Camsap3 | calmodulin regulated spectrin-associated protein family, member 3 | 289.99 | 697.53 | -1.27 | 2.48E-09 |
| ENSMUSG00000087687 | GSMG0036950 | Pet100 | PET100 homolog | 540.66 | 540.88 | 0.00 | 9.96E-01 |
| ENSMUSG00000004626 | GSMG0036951 | Stxbp2 | syntaxin binding protein 2 | 2217.17 | 2432.36 | -0.13 | 6.03E-01 |
| ENSMUSG00000040236 | GSMG0036955 | Trappc5 | trafficking protein particle complex 5 | 1160.06 | 1404.58 | -0.28 | 1.71E-01 |
| ENSMUSG00000011832 | GSMG0036957 | Evi5l | ecotropic viral integration site 5 like | 1086.87 | 1432.92 | -0.40 | 7.00E-02 |
| G00000002948,ENSMUSG000 | GSMG0036960 | Map2k7 // Tgfbr3l | mitogen-activated protein kinase kinase 7 // transforming growth factor, | 932.06 | 885.26 | 0.07 | 8.15E-01 |

| | | | | | | | |
|---|---|---|---|---|---|---|---|
| ENSMUSG00000011837 | GSMG0036961 | Snapc2 | small nuclear RNA activating complex, polypeptide 2 | 701.42 | 795.70 | -0.18 | 4.76E-01 |
| ENSMUSG00000008206 | GSMG0036965 | Cers4 | ceramide synthase 4 | 2045.37 | 1708.11 | 0.26 | 1.77E-01 |
| ENSMUSG00000058748 | GSMG0036968 | Zfp958 | zinc finger protein 958 | 295.62 | 252.36 | 0.23 | 4.18E-01 |
| ENSMUSG00000040396 | GSMG0036984 | Abhd13 | abhydrolase domain containing 13 | 1468.92 | 1428.60 | 0.04 | 8.84E-01 |
| ENSMUSG00000031503 | GSMG0036995 | Col4a2 | collagen, type IV, alpha 2 | 54042.65 | 54958.37 | -0.02 | 9.48E-01 |
| ENSMUSG00000031505 | GSMG0036998 | Carkd | carbohydrate kinase domain containing | 975.67 | 972.84 | 0.01 | 9.86E-01 |
| ENSMUSG00000045969 | GSMG0036999 | Ing1 | inhibitor of growth family, member 1 | 1279.47 | 1201.43 | 0.09 | 7.44E-01 |
| ENSMUSG00000031511 | GSMG0037004 | Arhgef7 | Rho guanine nucleotide exchange factor (GEF7) | 772.31 | 907.06 | -0.23 | 3.86E-01 |
| ENSMUSG00000031441 | GSMG0037014 | Atp11a | ATPase, class VI, type 11A | 1462.35 | 1118.40 | 0.39 | 2.54E-01 |
| ENSMUSG00000031442 | GSMG0037015 | Mcf2l | mcf.2 transforming sequence-like | 3637.40 | 3533.12 | 0.04 | 8.80E-01 |
| ENSMUSG00000031446 | GSMG0037020 | Cul4a | cullin 4A | 3609.56 | 3383.46 | 0.09 | 6.73E-01 |
| ENSMUSG00000031447 | GSMG0037022 | Lamp1 | lysosomal-associated membrane protein 1 | 4116.97 | 3729.06 | 0.14 | 5.58E-01 |
| ENSMUSG00000038497 | GSMG0037025 | Tmco3 | transmembrane and coiled-coil domains 3 | 712.39 | 721.66 | -0.02 | 9.58E-01 |
| ENSMUSG00000038482 | GSMG0037026 | Tfdp1 | transcription factor Dp 1 | 1022.55 | 1205.92 | -0.24 | 2.60E-01 |
| ENSMUSG00000038457 | GSMG0037028 | Tmem255b | transmembrane protein 255B | 115.65 | 62.45 | 0.90 | 1.27E-02 |
| ENSMUSG00000038416 | GSMG0037030 | Cdc16 | CDC16 cell division cycle 16 | 2848.98 | 3999.34 | -0.49 | 8.53E-04 |
| ENSMUSG00000038398 | GSMG0037032 | Upf3a | UPF3 regulator of nonsense transcripts homolog A (yeast) | 848.59 | 845.78 | 0.00 | 9.91E-01 |
| ENSMUSG00000047710 | GSMG0037033 | Champ1 | chromosome alignment maintaining phosphoprotein 1 | 554.59 | 769.04 | -0.47 | 7.11E-03 |
| ENSMUSG00000038365 | GSMG0037035 | Fbxo25 | F-box protein 25 | 384.32 | 450.89 | -0.23 | 3.32E-01 |
| NULL | GSMG0037038 | NULL | NULL | 504.40 | 144.21 | 1.81 | 1.01E-18 |
| ENSMUSG00000047495 | GSMG0037039 | Dlgap2 | discs, large (Drosophila) homolog-associated protein 2 | 340.49 | 237.43 | 0.52 | 4.96E-02 |
| NULL | GSMG0037040 | NULL | NULL | 337.80 | 161.42 | 1.07 | 5.24E-05 |
| ENSMUSG00000026317 | GSMG0037042 | Cln8 | ceroid-lipofuscinosis, neuronal 8 | 274.99 | 340.08 | -0.31 | 2.22E-01 |
| NULL | GSMG0037043 | NULL | NULL | 199.86 | 29.61 | 2.76 | 1.97E-22 |
| ENSMUSG00000071176 | GSMG0037044 | Arhgef10 | Rho guanine nucleotide exchange factor (GEF) 10 | 1401.41 | 1106.17 | 0.34 | 1.79E-01 |
| ENSMUSG00000055675 | GSMG0037046 | Kbtbd11 | kelch repeat and BTB (POZ) domain containing 11 | 1447.94 | 1465.34 | -0.02 | 9.65E-01 |
| ENSMUSG00000039842 | GSMG0037052 | Mcph1 | microcephaly, primary autosomal recessive 1 | 365.82 | 428.61 | -0.23 | 3.54E-01 |
| ENSMUSG00000031467 | GSMG0037054 | Agpat5 | 1-acylglycerol-3-phosphate O-acyltransferase 5 (lysophosphatidic acid a | 1710.64 | 1656.69 | 0.05 | 8.77E-01 |
| ENSMUSG00000063362 | GSMG0037082 | Alg11 | asparagine-linked glycosylation 11 (alpha-1,2-mannosyltransferase) | 786.17 | 952.29 | -0.28 | 2.34E-01 |
| ENSMUSG00000031479 | GSMG0037083 | Vps36 | vacuolar protein sorting 36 | 1305.99 | 1362.94 | -0.06 | 8.44E-01 |
| ENSMUSG00000031480 | GSMG0037084 | Thsd1 | thrombospondin, type I, domain 1 | 2661.16 | 2564.03 | 0.05 | 8.47E-01 |
| ENSMUSG00000031533 | GSMG0037089 | Mrps31 | mitochondrial ribosomal protein S31 | 340.99 | 454.88 | -0.42 | 6.86E-02 |
| ENSMUSG00000037656 | GSMG0037090 | Slc20a2 | solute carrier family 20, member 2 | 1947.36 | 1923.31 | 0.02 | 9.51E-01 |

| | | | | | | | |
|---|---|---|---|---|---|---|---|
| ENSMUSG00000031538 | GSMG0037095 | Plat | plasminogen activator, tissue | 8989.43 | 7478.15 | 0.27 | 1.32E-01 |
| ENSMUSG00000031540 | GSMG0037097 | Kat6a | K(lysine) acetyltransferase 6A | 456.67 | 395.24 | 0.21 | 4.63E-01 |
| ENSMUSG00000031548 | GSMG0037107 | Sfrp1 | secreted frizzled-related protein 1 | 315.09 | 1136.80 | -1.85 | 1.13E-24 |
| ENSMUSG00000031556 | GSMG0037114 | Tm2d2 | TM2 domain containing 2 | 565.16 | 562.25 | 0.01 | 9.78E-01 |
| ENSMUSG00000031565 | GSMG0037120 | Fgfr1 | fibroblast growth factor receptor 1 | 1103.09 | 1091.37 | 0.02 | 9.69E-01 |
| ENSMUSG00000054823 | GSMG0037122 | Whsc1l1 | Wolf-Hirschhorn syndrome candidate 1-like 1 (human) | 454.14 | 503.59 | -0.15 | 6.05E-01 |
| ENSMUSG00000031570 | GSMG0037126 | Plpp5 | phospholipid phosphatase 5 | 525.43 | 550.57 | -0.07 | 8.30E-01 |
| ENSMUSG00000037296 | GSMG0037128 | Lsm1 | LSM1 homolog, mRNA degradation associated | 858.50 | 878.48 | -0.03 | 9.29E-01 |
| ENSMUSG00000013878 | GSMG0037133 | Rnf170 | ring finger protein 170 | 169.61 | 157.07 | 0.11 | 7.86E-01 |
| ENSMUSG00000031483 | GSMG0037138 | Erlin2 | ER lipid raft associated 2 | 743.60 | 898.07 | -0.27 | 1.75E-01 |
| ENSMUSG00000031485 | GSMG0037139 | Prosc | proline synthetase co-transcribed | 1571.46 | 1458.36 | 0.11 | 6.61E-01 |
| ENSMUSG00000031486 | GSMG0037141 | Adgra2 | adhesion G protein-coupled receptor A2 | 225.74 | 202.57 | 0.15 | 7.70E-01 |
| ENSMUSG00000031490 | GSMG0037145 | Eif4ebp1 | eukaryotic translation initiation factor 4E binding protein 1 | 397.70 | 485.36 | -0.29 | 2.09E-01 |
| ENSMUSG00000039661 | GSMG0037154 | Dusp26 | dual specificity phosphatase 26 (putative) | 191.81 | 327.05 | -0.77 | 9.36E-04 |
| ENSMUSG00000039328 | GSMG0037156 | Rnf122 | ring finger protein 122 | 707.18 | 730.87 | -0.05 | 8.84E-01 |
| ENSMUSG00000031577 | GSMG0037157 | Tti2 | TELO2 interacting protein 2 | 239.52 | 230.84 | 0.06 | 8.96E-01 |
| ENSMUSG00000046152 | GSMG0037158 | Fut10 | fucosyltransferase 10 | 186.44 | 199.38 | -0.10 | 8.09E-01 |
| NULL | GSMG0037165 | NULL | NULL | 348.65 | 267.56 | 0.38 | 1.52E-01 |
| ENSMUSG00000009630 | GSMG0037173 | Ppp2cb | protein phosphatase 2 (formerly 2A), catalytic subunit, beta isoform | 997.52 | 612.35 | 0.70 | 1.90E-02 |
| NULL | GSMG0037174 | NULL | NULL | 255.27 | 150.52 | 0.76 | 1.36E-03 |
| ENSMUSG00000031584 | GSMG0037175 | Gsr | glutathione reductase | 1356.88 | 1478.05 | -0.12 | 6.80E-01 |
| ENSMUSG00000031585 | GSMG0037176 | Gtf2e2 | general transcription factor II E, polypeptide 2 (beta subunit) | 614.16 | 712.33 | -0.21 | 4.54E-01 |
| ENSMUSG00000031532 | GSMG0037181 | Saraf | store-operated calcium entry-associated regulatory factor | 4509.71 | 4361.70 | 0.05 | 8.65E-01 |
| ENSMUSG00000070056 | GSMG0037189 | Mfhas1 | malignant fibrous histiocytoma amplified sequence 1 | 3435.03 | 3472.17 | -0.02 | 9.63E-01 |
| NULL | GSMG0037190 | NULL | NULL | 200.32 | 206.39 | -0.04 | 9.37E-01 |
| ENSMUSG00000050271 | GSMG0037197 | D8Ertd82e | DNA segment, Chr 8, ERATO Doi 82, expressed | 230.68 | 351.34 | -0.60 | 2.83E-02 |
| ENSMUSG00000039620 | GSMG0037202 | 6430573F11Rik | RIKEN cDNA 6430573F11 gene | 166.81 | 176.38 | -0.08 | 8.45E-01 |
| ENSMUSG00000074384 | GSMG0037204 | AI429214 | expressed sequence AI429214 | 263.39 | 246.48 | 0.10 | 8.25E-01 |
| ENSMUSG00000039470 | GSMG0037215 | Zdhhc2 | zinc finger, DHHC domain containing 2 | 336.29 | 728.84 | -1.11 | 1.85E-11 |
| ENSMUSG00000031600 | GSMG0037216 | Vps37a | vacuolar protein sorting 37A | 630.73 | 530.53 | 0.25 | 2.74E-01 |
| ENSMUSG00000031596 | GSMG0037222 | Slc7a2 | solute carrier family 7 (cationic amino acid transporter, y+ system), mer | 195.65 | 261.59 | -0.42 | 1.40E-01 |
| ENSMUSG00000031595 | GSMG0037223 | Pdgfrl | platelet-derived growth factor receptor-like | 191.19 | 139.16 | 0.46 | 1.60E-01 |
| NULL | GSMG0037225 | NULL | NULL | 174.02 | 121.19 | 0.52 | 8.95E-02 |

| | | | | | | | |
|---|---|---|---|---|---|---|---|
| ENSMUSG00000031592 | GSMG0037227 | Pcm1 | pericentriolar material 1 | 627.13 | 755.17 | -0.27 | 2.23E-01 |
| ENSMUSG00000070047 | GSMG0037237 | Fat1 | FAT atypical cadherin 1 | 156.93 | 236.97 | -0.59 | 1.44E-02 |
| ENSMUSG00000031636 | GSMG0037245 | Pdlim3 | PDZ and LIM domain 3 | 68.54 | 4.84 | 3.84 | 9.61E-12 |
| ENSMUSG00000031634 | GSMG0037248 | Ufsp2 | UFM1-specific peptidase 2 | 814.23 | 831.87 | -0.03 | 9.30E-01 |
| ENSMUSG00000031631 | GSMG0037252 | Cfap97 | cilia and flagella associated protein 97 | 344.10 | 351.84 | -0.03 | 9.34E-01 |
| ENSMUSG00000018796 | GSMG0037255 | Acsl1 | acyl-CoA synthetase long-chain family member 1 | 1739.06 | 1783.62 | -0.04 | 8.94E-01 |
| ENSMUSG00000031629 | GSMG0037256 | Cenpu | centromere protein U | 70.38 | 123.11 | -0.81 | 1.56E-02 |
| ENSMUSG00000031628 | GSMG0037258 | Casp3 | caspase 3 | 153.58 | 428.64 | -1.48 | 4.83E-14 |
| ENSMUSG00000031627 | GSMG0037259 | Irf2 | interferon regulatory factor 2 | 645.64 | 537.54 | 0.26 | 2.33E-01 |
| ENSMUSG00000038173 | GSMG0037260 | Enpp6 | ectonucleotide pyrophosphatase/phosphodiesterase 6 | 115.28 | 22.69 | 2.36 | 9.01E-09 |
| ENSMUSG00000031568 | GSMG0037264 | Rwdd4a | RWD domain containing 4A | 186.04 | 236.54 | -0.35 | 2.11E-01 |
| ENSMUSG00000031562 | GSMG0037268 | Dctd | dCMP deaminase | 180.92 | 497.14 | -1.45 | 3.59E-12 |
| ENSMUSG00000031521 | GSMG0037276 | Aga | aspartylglucosaminidase | 486.97 | 430.90 | 0.18 | 5.01E-01 |
| ENSMUSG00000031520 | GSMG0037277 | Vegfc | vascular endothelial growth factor C | 874.68 | 242.41 | 1.85 | 2.57E-23 |
| ENSMUSG00000031517 | GSMG0037280 | Gpm6a | glycoprotein m6a | 2211.21 | 2511.25 | -0.18 | 4.94E-01 |
| NULL | GSMG0037281 | NULL | NULL | 278.37 | 320.87 | -0.20 | 6.16E-01 |
| NULL | GSMG0037284 | NULL | NULL | 241.29 | 283.17 | -0.23 | 5.62E-01 |
| ENSMUSG00000031613 | GSMG0037292 | Hpgd | hydroxyprostaglandin dehydrogenase 15 (NAD) | 58.36 | 28.76 | 1.03 | 4.34E-02 |
| ENSMUSG00000038206 | GSMG0037294 | Fbxo8 | F-box protein 8 | 476.28 | 456.93 | 0.06 | 8.49E-01 |
| ENSMUSG00000031610 | GSMG0037297 | Scrg1 | scrapie responsive gene 1 | 314.94 | 197.97 | 0.67 | 7.50E-03 |
| ENSMUSG00000049946 | GSMG0037302 | BC030500 | cDNA sequence BC030500 | 124.69 | 192.89 | -0.63 | 1.82E-02 |
| ENSMUSG00000031647 | GSMG0037306 | Mfap3l | microfibrillar-associated protein 3-like | 786.43 | 575.80 | 0.45 | 5.03E-02 |
| ENSMUSG00000038005 | GSMG0037307 | Hpf1 | histone PARylation factor 1 | 547.00 | 691.55 | -0.34 | 2.61E-01 |
| ENSMUSG00000031644 | GSMG0037309 | Nek1 | NIMA (never in mitosis gene a)-related expressed kinase 1 | 335.02 | 309.68 | 0.11 | 7.56E-01 |
| ENSMUSG00000031642 | GSMG0037314 | Sh3rf1 | SH3 domain containing ring finger 1 | 633.07 | 569.21 | 0.15 | 5.37E-01 |
| ENSMUSG00000031641 | GSMG0037317 | Cbr4 | carbonyl reductase 4 | 317.84 | 305.31 | 0.06 | 8.76E-01 |
| ENSMUSG00000054162 | GSMG0037325 | Spock3 | sparc/osteonectin, cwcv and kazal-like domains proteoglycan 3 | 252.57 | 264.12 | -0.06 | 8.90E-01 |
| ENSMUSG00000025521 | GSMG0037331 | Tmem192 | transmembrane protein 192 | 288.15 | 275.10 | 0.07 | 8.59E-01 |
| ENSMUSG00000036469 | GSMG0037332 | March1 | membrane-associated ring finger (C3HC4) 1 | 418.92 | 787.84 | -0.91 | 2.60E-08 |
| ENSMUSG00000036437 | GSMG0037342 | Npy1r | neuropeptide Y receptor Y1 | 281.69 | 179.73 | 0.65 | 3.29E-02 |
| ENSMUSG00000031864 | GSMG0037351 | Ints10 | integrator complex subunit 10 | 1897.67 | 2188.12 | -0.21 | 2.89E-01 |
| ENSMUSG00000015568 | GSMG0037352 | Lpl | lipoprotein lipase | 572.91 | 1320.74 | -1.20 | 6.65E-14 |
| ENSMUSG00000006273 | GSMG0037353 | Atp6v1b2 | ATPase, H+ transporting, lysosomal V1 subunit B2 | 1001.79 | 975.40 | 0.04 | 9.00E-01 |

| | | | | | | | |
|---|---|---|---|---|---|---|---|
| ENSMUSG00000059897 | GSMG0037355 | Zfp930 | zinc finger protein 930 | 279.50 | 302.61 | -0.11 | 7.28E-01 |
| ENSMUSG00000031862 | GSMG0037360 | Atp13a1 | ATPase type 13A1 | 5183.78 | 5958.33 | -0.20 | 4.51E-01 |
| ENSMUSG00000036246 | GSMG0037361 | Gmip | Gem-interacting protein | 234.11 | 733.72 | -1.65 | 5.05E-17 |
| ENSMUSG00000031860 | GSMG0037363 | Pbx4 | pre B cell leukemia homeobox 4 | 267.00 | 234.07 | 0.19 | 6.03E-01 |
| ENSMUSG00000011306 | GSMG0037366 | Sugp1 | SURP and G patch domain containing 1 | 897.05 | 956.69 | -0.09 | 7.26E-01 |
| ENSMUSG00000071078 | GSMG0037371 | Nr2c2ap | nuclear receptor 2C2-associated protein | 317.70 | 430.39 | -0.44 | 3.71E-02 |
| ENSMUSG00000002342 | GSMG0037375 | Tmem161a | transmembrane protein 161A | 1391.00 | 1280.74 | 0.12 | 6.11E-01 |
| ENSMUSG00000036054 | GSMG0037376 | Sugp2 | SURP and G patch domain containing 2 | 991.39 | 1018.45 | -0.04 | 9.21E-01 |
| ENSMUSG00000003573 | GSMG0037378 | Homer3 | homer scaffolding protein 3 | 546.23 | 721.08 | -0.40 | 5.97E-02 |
| ENSMUSG00000055681 | GSMG0037380 | Cope | coatomer protein complex, subunit epsilon | 3622.05 | 3121.83 | 0.21 | 3.03E-01 |
| G00000087408,ENSMUSG000 | GSMG0037381 | Gdf1 // Cers1 | growth differentiation factor 1 // ceramide synthase 1 | 361.94 | 307.84 | 0.23 | 4.40E-01 |
| ENSMUSG00000007888 | GSMG0037385 | Crlf1 | cytokine receptor-like factor 1 | 187.67 | 525.40 | -1.49 | 1.13E-12 |
| NULL | GSMG0037388 | NULL | NULL | 254.78 | 205.71 | 0.31 | 2.83E-01 |
| ENSMUSG00000019428 | GSMG0037389 | Fkbp8 | FK506 binding protein 8 | 4747.54 | 4725.78 | 0.01 | 9.81E-01 |
| ENSMUSG00000070002 | GSMG0037390 | Ell | elongation factor RNA polymerase II | 579.11 | 600.19 | -0.05 | 8.77E-01 |
| ENSMUSG00000019139 | GSMG0037392 | Isyna1 | myo-inositol 1-phosphate synthase A1 | 5520.69 | 4056.64 | 0.44 | 9.68E-03 |
| ENSMUSG00000031848 | GSMG0037395 | Lsm4 | LSM4 homolog, U6 small nuclear RNA and mRNA degradation associa | 907.95 | 1319.52 | -0.54 | 5.04E-03 |
| NULL | GSMG0037396 | NULL | NULL | 2806.71 | 2114.29 | 0.41 | 1.26E-01 |
| ENSMUSG00000031840 | GSMG0037398 | Rab3a | RAB3A, member RAS oncogene family | 2405.31 | 2074.64 | 0.21 | 3.21E-01 |
| NULL | GSMG0037400 | NULL | NULL | 74.93 | 172.21 | -1.21 | 1.83E-03 |
| NULL | GSMG0037401 | NULL | NULL | 64.57 | 82.58 | -0.35 | 4.77E-01 |
| ENSMUSG00000100691 | GSMG0037402 | 2010320M18Rik | RIKEN cDNA 2010320M18 gene | 423.21 | 509.24 | -0.27 | 3.82E-01 |
| ENSMUSG00000019261 | GSMG0037406 | Map1s | microtubule-associated protein 1S | 1532.36 | 1790.84 | -0.23 | 3.09E-01 |
| NULL | GSMG0037407 | NULL | NULL | 181.04 | 171.27 | 0.08 | 8.49E-01 |
| ENSMUSG00000004677 | GSMG0037412 | Myo9b | myosin IXb | 581.19 | 708.39 | -0.29 | 2.81E-01 |
| ENSMUSG00000002395 | GSMG0037413 | Use1 | unconventional SNARE in the ER 1 homolog (S. cerevisiae) | 863.19 | 818.68 | 0.08 | 8.02E-01 |
| ENSMUSG00000002396 | GSMG0037414 | Ocel1 | occludin/ELL domain containing 1 | 1903.49 | 1433.18 | 0.41 | 2.23E-02 |
| NULL | GSMG0037415 | NULL | NULL | 66.33 | 73.33 | -0.16 | 8.19E-01 |
| ENSMUSG00000031820 | GSMG0037416 | Babam1 | BRISC and BRCA1 A complex member 1 | 1371.81 | 1706.17 | -0.31 | 8.15E-02 |
| ENSMUSG00000046295 | GSMG0037417 | Ankle1 | ankyrin repeat and LEM domain containing 1 | 143.29 | 189.81 | -0.40 | 2.43E-01 |
| ENSMUSG00000034880 | GSMG0037420 | Mrpl34 | mitochondrial ribosomal protein L34 | 418.01 | 573.74 | -0.46 | 7.93E-02 |
| ENSMUSG00000074247 | GSMG0037421 | Dda1 | DET1 and DDB1 associated 1 | 2409.31 | 2971.36 | -0.30 | 7.25E-02 |
| ENSMUSG00000007610 | GSMG0037422 | Gtpbp3 | GTP binding protein 3 | 541.19 | 392.99 | 0.46 | 1.08E-01 |

| | | | | | | | |
|---|---|---|---|---|---|---|---|
| ENSMUSG00000031813 | GSMG0037424 | Mvb12a | multivesicular body subunit 12A | 808.97 | 832.06 | -0.04 | 8.91E-01 |
| ENSMUSG00000031808 | GSMG0037425 | Slc27a1 | solute carrier family 27 (fatty acid transporter), member 1 | 1529.60 | 1452.39 | 0.07 | 8.30E-01 |
| G00000031807,ENSMUSG000 | GSMG0037426 | Pgls // Fam129c | 6-phosphogluconolactonase // family with sequence similarity 129, mer | 1077.84 | 1361.05 | -0.34 | 7.40E-02 |
| ENSMUSG00000034807 | GSMG0037427 | Colgalt1 | collagen beta(1-O)galactosyltransferase 1 | 2115.72 | 2114.15 | 0.00 | 9.97E-01 |
| NULL | GSMG0037428 | NULL | NULL | 337.48 | 259.53 | 0.38 | 1.34E-01 |
| G00000031805,ENSMUSG000 | GSMG0037429 | Jak3 // Insl3 | Janus kinase 3 // insulin-like 3 | 2968.51 | 3648.25 | -0.30 | 1.31E-01 |
| ENSMUSG00000052446 | GSMG0037434 | Zfp961 | zinc finger protein 961 | 593.65 | 939.81 | -0.66 | 4.25E-05 |
| NULL | GSMG0037435 | NULL | NULL | 232.00 | 244.13 | -0.08 | 9.01E-01 |
| ENSMUSG00000031799 | GSMG0037441 | Tpm4 | tropomyosin 4 | 3900.47 | 4841.38 | -0.31 | 2.09E-01 |
| ENSMUSG00000003037 | GSMG0037442 | Rab8a | RAB8A, member RAS oncogene family | 2718.64 | 3242.79 | -0.25 | 2.06E-01 |
| ENSMUSG00000003039 | GSMG0037444 | Fam32a | family with sequence similarity 32, member A | 3102.64 | 3171.66 | -0.03 | 9.11E-01 |
| ENSMUSG00000003033 | GSMG0037445 | Ap1m1 | adaptor-related protein complex AP-1, mu subunit 1 | 2160.14 | 2868.16 | -0.41 | 2.09E-02 |
| ENSMUSG00000055148 | GSMG0037447 | Klf2 | Kruppel-like factor 2 (lung) | 2411.88 | 1463.07 | 0.72 | 5.12E-04 |
| ENSMUSG00000052794 | GSMG0037448 | 1700030K09Rik | RIKEN cDNA 1700030K09 gene | 146.76 | 173.62 | -0.24 | 4.91E-01 |
| ENSMUSG00000031622 | GSMG0037457 | Sin3b | transcriptional regulator, SIN3B (yeast) | 1429.91 | 1826.41 | -0.35 | 4.07E-02 |
| ENSMUSG00000034518 | GSMG0037463 | Hmgxb4 | HMG box domain containing 4 | 696.68 | 880.99 | -0.34 | 6.66E-02 |
| ENSMUSG00000005413 | GSMG0037465 | Hmox1 | heme oxygenase 1 | 666.09 | 661.55 | 0.01 | 9.75E-01 |
| ENSMUSG00000005410 | GSMG0037466 | Mcm5 | minichromosome maintenance complex component 5 | 605.16 | 1548.63 | -1.35 | 8.54E-16 |
| ENSMUSG00000034472 | GSMG0037467 | Rasd2 | RASD family, member 2 | 531.30 | 138.63 | 1.94 | 9.62E-18 |
| ENSMUSG00000031618 | GSMG0037470 | Nr3c2 | nuclear receptor subfamily 3, group C, member 2 | 582.02 | 415.66 | 0.49 | 4.55E-02 |
| NULL | GSMG0037471 | NULL | NULL | 217.53 | 157.16 | 0.46 | 1.54E-01 |
| ENSMUSG00000037134 | GSMG0037479 | Prmt9 | protein arginine methyltransferase 9 | 216.49 | 255.78 | -0.24 | 4.51E-01 |
| ENSMUSG00000031684 | GSMG0037484 | Slc10a7 | solute carrier family 10 (sodium/bile acid cotransporter family), membe | 257.14 | 251.83 | 0.03 | 9.44E-01 |
| ENSMUSG00000036990 | GSMG0037490 | Otud4 | OTU domain containing 4 | 574.49 | 647.30 | -0.17 | 5.85E-01 |
| ENSMUSG00000036977 | GSMG0037492 | Anapc10 | anaphase promoting complex subunit 10 | 431.05 | 478.27 | -0.15 | 6.34E-01 |
| ENSMUSG00000037940 | GSMG0037499 | Inpp4b | inositol polyphosphate-4-phosphatase, type II | 1880.37 | 974.80 | 0.95 | 2.84E-06 |
| ENSMUSG00000033938 | GSMG0037518 | Ndufb7 | NADH dehydrogenase (ubiquinone) 1 beta subcomplex, 7 | 2329.35 | 2638.78 | -0.18 | 5.08E-01 |
| ENSMUSG00000005483 | GSMG0037521 | Dnajb1 | DnaJ heat shock protein family (Hsp40) member B1 | 818.09 | 841.54 | -0.04 | 8.94E-01 |
| ENSMUSG00000019433 | GSMG0037522 | Gipc1 | GIPC PDZ domain containing family, member 1 | 2576.21 | 3056.86 | -0.25 | 1.48E-01 |
| ENSMUSG00000019464 | GSMG0037523 | Ptger1 | prostaglandin E receptor 1 (subtype EP1) | 1808.55 | 1674.00 | 0.11 | 7.25E-01 |
| ENSMUSG00000005481 | GSMG0037524 | Ddx39 | DEAD (Asp-Glu-Ala-Asp) box polypeptide 39 | 2593.51 | 3489.88 | -0.43 | 7.67E-03 |
| ENSMUSG00000013033 | GSMG0037525 | Adgrl1 | adhesion G protein-coupled receptor L1 | 840.39 | 816.94 | 0.04 | 9.28E-01 |
| ENSMUSG00000005470 | GSMG0037526 | Asf1b | anti-silencing function 1B histone chaperone | 164.29 | 397.04 | -1.27 | 7.95E-07 |

| | | | | | | | |
|---|---|---|---|---|---|---|---|
| ENSMUSG00000005469 | GSMG0037527 | Prkaca | protein kinase, cAMP dependent, catalytic, alpha | 1292.75 | 1655.66 | -0.36 | 3.92E-02 |
| NULL | GSMG0037528 | NULL | NULL | 223.85 | 187.28 | 0.25 | 5.27E-01 |
| ENSMUSG00000031706 | GSMG0037532 | Rfx1 | regulatory factor X, 1 (influences HLA class II expression) | 914.68 | 807.13 | 0.18 | 6.20E-01 |
| ENSMUSG00000034656 | GSMG0037536 | Cacna1a | calcium channel, voltage-dependent, P/Q type, alpha 1A subunit | 2726.43 | 1286.41 | 1.08 | 1.43E-07 |
| ENSMUSG00000001909 | GSMG0037538 | Trmt1 | tRNA methyltransferase 1 | 1756.07 | 1920.73 | -0.13 | 6.10E-01 |
| NULL | GSMG0037540 | NULL | NULL | 79.97 | 139.34 | -0.80 | 1.44E-02 |
| ENSMUSG00000033751 | GSMG0037542 | Gadd45gip1 | growth arrest and DNA-damage-inducible, gamma interacting protein 1 | 636.55 | 719.88 | -0.18 | 4.77E-01 |
| NULL | GSMG0037543 | NULL | NULL | 155.77 | 170.03 | -0.12 | 7.85E-01 |
| ENSMUSG00000003808 | GSMG0037545 | Farsa | phenylalanyl-tRNA synthetase, alpha subunit | 932.82 | 1101.94 | -0.24 | 2.38E-01 |
| ENSMUSG00000003824 | GSMG0037546 | Syce2 | synaptonemal complex central element protein 2 | 65.12 | 180.38 | -1.47 | 8.38E-08 |
| ENSMUSG00000003812 | GSMG0037549 | Dnase2a // Gm384 | deoxyribonuclease II alpha // predicted gene, 38426 | 116.13 | 86.35 | 0.42 | 2.84E-01 |
| ENSMUSG00000048617 | GSMG0037550 | Rtbdn | retbindin | 55.33 | 112.97 | -1.03 | 2.75E-03 |
| ENSMUSG00000005161 | GSMG0037551 | Prdx2 | peroxiredoxin 2 | 2443.61 | 3835.82 | -0.65 | 1.62E-04 |
| ENSMUSG00000052566 | GSMG0037552 | Hook2 | hook microtubule tethering protein 2 | 769.03 | 1559.23 | -1.02 | 8.07E-08 |
| ENSMUSG00000041203 | GSMG0037553 | 2310036O22Rik | RIKEN cDNA 2310036O22 gene | 1230.95 | 1443.35 | -0.23 | 3.02E-01 |
| ENSMUSG00000031691 | GSMG0037554 | Tnpo2 | transportin 2 (importin 3, karyopherin beta 2b) | 924.09 | 1027.72 | -0.15 | 7.13E-01 |
| ENSMUSG00000008167 | GSMG0037555 | Fbxw9 | F-box and WD-40 domain protein 9 | 973.67 | 1071.32 | -0.14 | 5.86E-01 |
| ENSMUSG00000060038 | GSMG0037556 | Dhps | deoxyhypusine synthase | 1053.69 | 1210.74 | -0.20 | 3.35E-01 |
| ENSMUSG00000059355 | GSMG0037557 | Wdr83os | WD repeat domain 83 opposite strand | 221.54 | 261.52 | -0.24 | 4.52E-01 |
| ENSMUSG00000005142 | GSMG0037558 | Man2b1 | mannosidase 2, alpha B1 | 1551.83 | 1763.42 | -0.18 | 4.47E-01 |
| ENSMUSG00000031697 | GSMG0037561 | Orc6 | origin recognition complex, subunit 6 | 309.13 | 541.75 | -0.81 | 3.44E-05 |
| ENSMUSG00000031700 | GSMG0037563 | Gpt2 | glutamic pyruvate transaminase (alanine aminotransferase) 2 | 364.41 | 332.45 | 0.14 | 7.26E-01 |
| ENSMUSG00000036879 | GSMG0037568 | Phkb | phosphorylase kinase beta | 1001.27 | 844.22 | 0.25 | 2.45E-01 |
| ENSMUSG00000047866 | GSMG0037572 | Lonp2 | lon peptidase 2, peroxisomal | 1233.59 | 1341.96 | -0.12 | 6.45E-01 |
| ENSMUSG00000036810 | GSMG0037584 | Cnep1r1 | CTD nuclear envelope phosphatase 1 regulatory subunit 1 | 887.55 | 861.34 | 0.04 | 9.04E-01 |
| ENSMUSG00000031657 | GSMG0037586 | Heatr3 | HEAT repeat containing 3 | 782.53 | 899.67 | -0.20 | 4.23E-01 |
| ENSMUSG00000036779 | GSMG0037589 | Papd5 | PAP associated domain containing 5 | 565.51 | 446.38 | 0.34 | 3.18E-01 |
| ENSMUSG00000031659 | GSMG0037591 | Adcy7 | adenylate cyclase 7 | 254.26 | 379.50 | -0.57 | 3.17E-02 |
| ENSMUSG00000031661 | GSMG0037592 | Nkd1 | naked cuticle 1 homolog (Drosophila) | 3283.14 | 2342.34 | 0.49 | 1.44E-03 |
| ENSMUSG00000036712 | GSMG0037595 | Cyld | cylindromatosis (turban tumor syndrome) | 187.42 | 155.84 | 0.26 | 3.99E-01 |
| ENSMUSG00000056608 | GSMG0037598 | Chd9 | chromodomain helicase DNA binding protein 9 | 368.66 | 358.34 | 0.04 | 9.26E-01 |
| ENSMUSG00000031666 | GSMG0037603 | Rbl2 | retinoblastoma-like 2 | 887.56 | 755.03 | 0.23 | 2.64E-01 |
| ENSMUSG00000055932 | GSMG0037605 | Fto | fat mass and obesity associated | 1932.83 | 2077.79 | -0.10 | 6.99E-01 |

| | | | | | | | |
|---|---|---|---|---|---|---|---|
| NULL | GSMG0037610 | NULL | NULL | 219.91 | 161.86 | 0.44 | 1.09E-01 |
| ENSMUSG00000031740 | GSMG0037619 | Mmp2 | matrix metallopeptidase 2 | 654.26 | 662.07 | -0.02 | 9.66E-01 |
| ENSMUSG00000033192 | GSMG0037620 | Lpcat2 | lysophosphatidylcholine acyltransferase 2 | 225.35 | 169.81 | 0.41 | 2.28E-01 |
| ENSMUSG00000031748 | GSMG0037625 | Gnao1 | guanine nucleotide binding protein, alpha O | 782.46 | 1112.89 | -0.51 | 9.08E-03 |
| NULL | GSMG0037626 | NULL | NULL | 889.45 | 718.29 | 0.31 | 1.19E-01 |
| ENSMUSG00000033009 | GSMG0037628 | Ogfod1 | 2-oxoglutarate and iron-dependent oxygenase domain containing 1 | 170.97 | 166.65 | 0.03 | 9.41E-01 |
| ENSMUSG00000031760 | GSMG0037631 | Mt3 | metallothionein 3 | 1154.63 | 707.95 | 0.70 | 2.29E-05 |
| ENSMUSG00000031762 | GSMG0037632 | Mt2 | metallothionein 2 | 521.63 | 204.19 | 1.35 | 3.96E-11 |
| ENSMUSG00000031765 | GSMG0037633 | Mt1 | metallothionein 1 | 2266.31 | 930.85 | 1.28 | 1.43E-20 |
| ENSMUSG00000032939 | GSMG0037634 | Nup93 | nucleoporin 93 | 1401.45 | 1691.04 | -0.27 | 1.22E-01 |
| ENSMUSG00000031770 | GSMG0037640 | Herpud1 | homocysteine-inducible, endoplasmic reticulum stress-inducible, ubiqu | 1399.58 | 914.28 | 0.61 | 1.11E-03 |
| ENSMUSG00000034361 | GSMG0037644 | Cpne2 | copine II | 675.35 | 631.90 | 0.10 | 7.32E-01 |
| ENSMUSG00000050079 | GSMG0037645 | Rspry1 | ring finger and SPRY domain containing 1 | 462.25 | 509.46 | -0.14 | 6.34E-01 |
| ENSMUSG00000031776 | GSMG0037648 | Arl2bp | ADP-ribosylation factor-like 2 binding protein | 6281.19 | 5642.89 | 0.15 | 4.46E-01 |
| NULL | GSMG0037649 | NULL | NULL | 489.43 | 192.13 | 1.35 | 8.63E-12 |
| ENSMUSG00000031778 | GSMG0037651 | Cx3cl1 | chemokine (C-X3-C motif) ligand 1 | 7504.11 | 3181.22 | 1.24 | 6.01E-12 |
| ENSMUSG00000031780 | GSMG0037652 | Ccl17 | chemokine (C-C motif) ligand 17 | 54.82 | 24.84 | 1.13 | 1.57E-02 |
| ENSMUSG00000031782 | GSMG0037653 | Coq9 | coenzyme Q9 | 753.46 | 600.03 | 0.33 | 9.83E-02 |
| ENSMUSG00000031785 | GSMG0037656 | Adgrg1 | adhesion G protein-coupled receptor G1 | 2452.65 | 2300.35 | 0.09 | 7.80E-01 |
| ENSMUSG00000060470 | GSMG0037657 | Adgrg3 | adhesion G protein-coupled receptor G3 | 165.00 | 369.15 | -1.16 | 4.70E-07 |
| ENSMUSG00000031787 | GSMG0037660 | Katnb1 | katanin p80 (WD40-containing) subunit B 1 | 1395.86 | 1630.07 | -0.22 | 3.39E-01 |
| ENSMUSG00000031792 | GSMG0037663 | Usb1 | U6 snRNA biogenesis 1 | 357.22 | 391.95 | -0.13 | 6.73E-01 |
| ENSMUSG00000031790 | GSMG0037664 | Mmp15 | matrix metallopeptidase 15 | 1067.30 | 1371.18 | -0.36 | 6.88E-01 |
| ENSMUSG00000031669 | GSMG0037669 | Gins3 | GINS complex subunit 3 (Psf3 homolog) | 197.59 | 299.73 | -0.61 | 1.75E-02 |
| ENSMUSG00000036564 | GSMG0037670 | Ndrg4 | N-myc downstream regulated gene 4 | 2637.82 | 1702.72 | 0.63 | 5.55E-03 |
| ENSMUSG00000031671 | GSMG0037671 | Setd6 | SET domain containing 6 | 374.28 | 476.47 | -0.35 | 1.09E-01 |
| NULL | GSMG0037681 | NULL | NULL | 39.42 | 50.47 | -0.37 | 5.96E-01 |
| ENSMUSG00000031871 | GSMG0037685 | Cdh5 | cadherin 5 | 2589.02 | 2504.42 | 0.05 | 9.03E-01 |
| ENSMUSG00000031872 | GSMG0037686 | Bean1 | brain expressed, associated with Nedd4, 1 | 48.14 | 114.66 | -1.25 | 1.16E-03 |
| ENSMUSG00000054400 | GSMG0037688 | Cklf | chemokine-like factor | 243.67 | 458.34 | -0.91 | 6.43E-07 |
| ENSMUSG00000031875 | GSMG0037690 | Cmtm3 | CKLF-like MARVEL transmembrane domain containing 3 | 488.01 | 889.89 | -0.87 | 5.71E-06 |
| NULL | GSMG0037691 | NULL | NULL | 123.92 | 91.06 | 0.44 | 3.84E-01 |
| NULL | GSMG0037692 | NULL | NULL | 176.45 | 189.60 | -0.10 | 8.05E-01 |

| | | | | | | | |
|---|---|---|---|---|---|---|---|
| ENSMUSG00000031883 | GSMG0037694 | Car7 | carbonic anhydrase 7 | 261.79 | 55.72 | 2.23 | 1.69E-19 |
| ENSMUSG00000050097 | GSMG0037700 | Ces2b | carboxyesterase 2B | 310.48 | 201.26 | 0.63 | 1.85E-02 |
| ENSMUSG00000031886 | GSMG0037702 | Ces2e | carboxylesterase 2E | 146.00 | 72.82 | 1.00 | 8.99E-04 |
| ENSMUSG00000031885 | GSMG0037709 | Cbfb | core binding factor beta | 1209.56 | 1266.54 | -0.07 | 8.58E-01 |
| ENSMUSG00000031889 | GSMG0037711 | D230025D16Rik | RIKEN cDNA D230025D16 gene | 1627.65 | 1643.03 | -0.01 | 9.62E-01 |
| ENSMUSG00000014776 | GSMG0037714 | Nol3 | nucleolar protein 3 (apoptosis repressor with CARD domain) | 150.08 | 334.22 | -1.15 | 4.52E-07 |
| NULL | GSMG0037715 | NULL | NULL | 150.07 | 321.64 | -1.10 | 1.64E-06 |
| ENSMUSG00000014859 | GSMG0037716 | E2f4 | E2F transcription factor 4 | 949.71 | 1149.77 | -0.28 | 1.47E-01 |
| ENSMUSG00000014856 | GSMG0037718 | Tmem208 | transmembrane protein 208 | 1583.32 | 1800.08 | -0.18 | 3.48E-01 |
| ENSMUSG00000014786 | GSMG0037719 | Slc9a5 | solute carrier family 9 (sodium/hydrogen exchanger), member 5 | 642.94 | 813.77 | -0.34 | 2.02E-01 |
| ENSMUSG00000038604 | GSMG0037723 | Fam65a | family with sequence similarity 65, member A | 6137.08 | 5882.68 | 0.06 | 8.59E-01 |
| ENSMUSG00000005698 | GSMG0037724 | Ctcf | CCCTC-binding factor | 1666.25 | 1879.77 | -0.17 | 4.32E-01 |
| ENSMUSG00000050357 | GSMG0037728 | Carmil2 | capping protein regulator and myosin 1 linker 2 | 493.82 | 555.84 | -0.17 | 5.30E-01 |
| ENSMUSG00000005699 | GSMG0037730 | Pard6a | par-6 family cell polarity regulator alpha | 622.40 | 523.02 | 0.25 | 2.84E-01 |
| ENSMUSG00000036442 | GSMG0037733 | Thap11 | THAP domain containing 11 | 1092.86 | 1058.16 | 0.05 | 8.77E-01 |
| ENSMUSG00000036270 | GSMG0037735 | Edc4 | enhancer of mRNA decapping 4 | 843.34 | 800.38 | 0.08 | 8.07E-01 |
| ENSMUSG00000048310 | GSMG0037737 | Pskh1 | protein serine kinase H1 | 243.68 | 288.15 | -0.24 | 5.07E-01 |
| ENSMUSG00000031901 | GSMG0037739 | Dus2 | dihydrouridine synthase 2 | 378.70 | 383.04 | -0.01 | 9.70E-01 |
| ENSMUSG00000031902 | GSMG0037740 | Nfatc3 | nuclear factor of activated T cells, cytoplasmic, calcineurin dependent 3 | 525.99 | 516.46 | 0.03 | 9.44E-01 |
| ENSMUSG00000031903 | GSMG0037746 | Pla2g15 | phospholipase A2, group XV | 736.04 | 632.41 | 0.22 | 3.74E-01 |
| ENSMUSG00000031904 | GSMG0037747 | Slc7a6 | solute carrier family 7 (cationic amino acid transporter, y+ system), mer | 520.50 | 520.20 | 0.00 | 9.98E-01 |
| ENSMUSG00000060098 | GSMG0037749 | Prmt7 | protein arginine N-methyltransferase 7 | 1125.73 | 1481.17 | -0.39 | 3.47E-02 |
| ENSMUSG00000031907 | GSMG0037753 | Zfp90 | zinc finger protein 90 | 522.17 | 874.25 | -0.74 | 6.41E-06 |
| ENSMUSG00000041949 | GSMG0037758 | Tango6 | transport and golgi organization 6 | 358.97 | 412.91 | -0.20 | 4.71E-01 |
| ENSMUSG00000041438 | GSMG0037760 | Cirh1a | cirrhosis, autosomal recessive 1A (human) | 1079.30 | 1101.98 | -0.03 | 9.32E-01 |
| ENSMUSG00000041308 | GSMG0037761 | Sntb2 | syntrophin, basic 2 | 529.74 | 453.36 | 0.23 | 3.32E-01 |
| ENSMUSG00000031913 | GSMG0037764 | Vps4a | vacuolar protein sorting 4A | 2728.73 | 2989.80 | -0.13 | 5.23E-01 |
| NULL | GSMG0037765 | NULL | NULL | 240.91 | 250.01 | -0.06 | 8.83E-01 |
| ENSMUSG00000031917 | GSMG0037766 | Nip7 | NIP7, nucleolar pre-rRNA processing protein | 821.61 | 1093.97 | -0.41 | 1.42E-02 |
| ENSMUSG00000031924 | GSMG0037769 | Cyb5b | cytochrome b5 type B | 5837.97 | 6599.18 | -0.18 | 3.95E-01 |
| ENSMUSG00000003847 | GSMG0037777 | Nfat5 | nuclear factor of activated T cells 5 | 477.24 | 391.72 | 0.29 | 2.43E-01 |
| ENSMUSG00000031930 | GSMG0037779 | Wwp2 | WW domain containing E3 ubiquitin protein ligase 2 | 909.79 | 906.56 | 0.01 | 9.86E-01 |
| ENSMUSG00000031723 | GSMG0037798 | Txnl4b | thioredoxin-like 4B | 261.66 | 321.20 | -0.30 | 3.38E-01 |

| | | | | | | | |
|---|---|---|---|---|---|---|---|
| ENSMUSG00000031728 | GSMG0037801 | Zfp821 | zinc finger protein 821 | 436.35 | 543.34 | -0.32 | 1.25E-01 |
| ENSMUSG00000031731 | GSMG0037802 | Ap1g1 | adaptor protein complex AP-1, gamma 1 subunit | 787.91 | 881.36 | -0.16 | 5.36E-01 |
| ENSMUSG00000044676 | GSMG0037809 | Zfp612 | zinc finger protein 612 | 1418.02 | 702.78 | 1.01 | 1.13E-09 |
| ENSMUSG00000010936 | GSMG0037812 | Vac14 | Vac14 homolog (S. cerevisiae) | 1169.22 | 1762.95 | -0.59 | 1.36E-04 |
| ENSMUSG00000033763 | GSMG0037813 | Mtss1l | metastasis suppressor 1-like | 599.65 | 777.84 | -0.38 | 6.54E-01 |
| ENSMUSG00000031753 | GSMG0037815 | Cog4 | component of oligomeric golgi complex 4 | 698.17 | 673.20 | 0.05 | 8.66E-01 |
| ENSMUSG00000031749 | GSMG0037816 | St3gal2 | ST3 beta-galactoside alpha-2,3-sialyltransferase 2 | 578.99 | 531.08 | 0.13 | 6.52E-01 |
| ENSMUSG00000031960 | GSMG0037818 | Aars | alanyl-tRNA synthetase | 4014.92 | 5122.28 | -0.35 | 2.10E-02 |
| ENSMUSG00000033624 | GSMG0037819 | Pdpr | pyruvate dehydrogenase phosphatase regulatory subunit | 542.73 | 596.78 | -0.14 | 6.82E-01 |
| ENSMUSG00000033545 | GSMG0037821 | Znrf1 | zinc and ring finger 1 | 3483.45 | 3933.04 | -0.18 | 4.35E-01 |
| ENSMUSG00000055835 | GSMG0037825 | Zfp1 | zinc finger protein 1 | 541.86 | 602.16 | -0.15 | 5.96E-01 |
| ENSMUSG00000033430 | GSMG0037829 | Terf2ip | telomeric repeat binding factor 2, interacting protein | 1211.27 | 1301.10 | -0.10 | 6.52E-01 |
| ENSMUSG00000031772 | GSMG0037831 | Cntnap4 | contactin associated protein-like 4 | 416.67 | 406.42 | 0.04 | 9.30E-01 |
| ENSMUSG00000031767 | GSMG0037839 | Nudt7 | nudix (nucleoside diphosphate linked moiety X)-type motif 7 | 171.62 | 108.51 | 0.66 | 4.06E-02 |
| ENSMUSG00000004637 | GSMG0037844 | Wwox | WW domain-containing oxidoreductase | 154.61 | 216.08 | -0.48 | 8.96E-02 |
| ENSMUSG00000031756 | GSMG0037863 | Cenpn | centromere protein N | 49.37 | 146.86 | -1.57 | 2.73E-06 |
| ENSMUSG00000047388 | GSMG0037865 | Atmin | ATM interactor | 937.71 | 1157.90 | -0.30 | 1.42E-01 |
| ENSMUSG00000034390 | GSMG0037870 | Cmip | c-Maf inducing protein | 1338.43 | 1956.66 | -0.55 | 9.01E-03 |
| NULL | GSMG0037871 | NULL | NULL | 1827.35 | 1649.88 | 0.15 | 5.75E-01 |
| NULL | GSMG0037873 | NULL | NULL | 638.12 | 484.93 | 0.40 | 6.67E-02 |
| ENSMUSG00000034330 | GSMG0037876 | Plcg2 | phospholipase C, gamma 2 | 300.43 | 102.14 | 1.56 | 1.21E-11 |
| ENSMUSG00000031839 | GSMG0037884 | Hsbp1 | heat shock factor binding protein 1 | 4180.83 | 4761.75 | -0.19 | 4.21E-01 |
| ENSMUSG00000074064 | GSMG0037885 | Mlycd | malonyl-CoA decarboxylase | 1319.04 | 1749.03 | -0.41 | 1.80E-02 |
| ENSMUSG00000074063 | GSMG0037886 | Osgin1 | oxidative stress induced growth inhibitor 1 | 82.62 | 45.26 | 0.87 | 3.63E-02 |
| ENSMUSG00000031837 | GSMG0037887 | Necab2 | N-terminal EF-hand calcium binding protein 2 | 376.17 | 262.32 | 0.52 | 1.53E-01 |
| ENSMUSG00000023336 | GSMG0037890 | Wfdc1 | WAP four-disulfide core domain 1 | 3219.33 | 1137.65 | 1.50 | 2.69E-24 |
| NULL | GSMG0037892 | NULL | NULL | 76.41 | 204.96 | -1.42 | 2.21E-04 |
| ENSMUSG00000031828 | GSMG0037893 | Klhl36 | kelch-like 36 | 208.18 | 243.37 | -0.22 | 4.90E-01 |
| ENSMUSG00000031826 | GSMG0037895 | Usp10 | ubiquitin specific peptidase 10 | 1632.06 | 2354.35 | -0.53 | 7.11E-04 |
| ENSMUSG00000031825 | GSMG0037896 | Crispld2 | cysteine-rich secretory protein LCCL domain containing 2 | 393.10 | 54.12 | 2.87 | 3.37E-23 |
| NULL | GSMG0037897 | NULL | NULL | 181.42 | 144.36 | 0.33 | 2.97E-01 |
| NULL | GSMG0037898 | NULL | NULL | 84.41 | 57.96 | 0.54 | 1.95E-01 |
| NULL | GSMG0037899 | NULL | NULL | 388.41 | 254.70 | 0.61 | 5.51E-03 |

| | | | | | | | |
|---|---|---|---|---|---|---|---|
| ENSMUSG00000031824 | GSMG0037901 | 6430548M08Rik | RIKEN cDNA 6430548M08 gene | 422.86 | 357.55 | 0.24 | 3.63E-01 |
| ENSMUSG00000031822 | GSMG0037904 | Gse1 | genetic suppressor element 1, coiled-coil protein | 314.23 | 233.14 | 0.43 | 2.86E-01 |
| ENSMUSG00000031818 | GSMG0037908 | Cox4i1 | cytochrome c oxidase subunit IV isoform 1 | 10176.56 | 10571.71 | -0.06 | 8.40E-01 |
| ENSMUSG00000041515 | GSMG0037911 | Irf8 | interferon regulatory factor 8 | 132.24 | 109.14 | 0.27 | 4.67E-01 |
| ENSMUSG00000042812 | GSMG0037913 | Foxf1 | forkhead box F1 | 440.00 | 316.78 | 0.47 | 1.22E-01 |
| ENSMUSG00000031812 | GSMG0037921 | Map1lc3b | microtubule-associated protein 1 light chain 3 beta | 4803.17 | 4069.79 | 0.24 | 1.58E-01 |
| NULL | GSMG0037922 | NULL | NULL | 149.71 | 198.01 | -0.41 | 1.56E-01 |
| ENSMUSG00000025318 | GSMG0037923 | Jph3 | junctophilin 3 | 672.99 | 799.95 | -0.25 | 3.89E-01 |
| ENSMUSG00000025316 | GSMG0037926 | Banp | BTG3 associated nuclear protein | 699.45 | 551.06 | 0.35 | 9.68E-02 |
| ENSMUSG00000049577 | GSMG0037928 | Zfpm1 | zinc finger protein, multitype 1 | 324.00 | 155.56 | 1.06 | 1.93E-01 |
| ENSMUSG00000017478 | GSMG0037930 | Zc3h18 | zinc finger CCCH-type containing 18 | 1052.75 | 1234.50 | -0.23 | 3.17E-01 |
| NULL | GSMG0037932 | 9330133O14Rik | RIKEN cDNA 9330133O14 gene | 268.33 | 370.49 | -0.47 | 2.92E-02 |
| ENSMUSG00000049482 | GSMG0037936 | Ctu2 | cytosolic thiouridylase subunit 2 | 1148.82 | 1325.76 | -0.21 | 4.40E-01 |
| ENSMUSG00000006585 | GSMG0037940 | Cdt1 | chromatin licensing and DNA replication factor 1 | 78.75 | 195.63 | -1.32 | 5.50E-07 |
| ENSMUSG00000015013 | GSMG0037941 | Trappc2l | trafficking protein particle complex 2-like | 832.61 | 797.11 | 0.06 | 8.33E-01 |
| ENSMUSG00000015016 | GSMG0037944 | Acsf3 | acyl-CoA synthetase family member 3 | 166.31 | 170.85 | -0.04 | 9.44E-01 |
| ENSMUSG00000001482 | GSMG0037950 | Def8 | differentially expressed in FDCP 8 | 557.77 | 578.69 | -0.05 | 8.72E-01 |
| ENSMUSG00000099881 | GSMG0037951 | 2810013P06Rik | RIKEN cDNA 2810013P06 gene | 901.28 | 806.97 | 0.16 | 5.58E-01 |
| ENSMUSG00000000738 | GSMG0037952 | Spg7 | spastic paraplegia 7 homolog (human) | 1124.78 | 1207.36 | -0.10 | 6.88E-01 |
| ENSMUSG00000034796 | GSMG0037954 | Cpne7 | copine VII | 1632.53 | 923.42 | 0.82 | 4.81E-04 |
| ENSMUSG00000033862 | GSMG0037958 | Cdk10 | cyclin-dependent kinase 10 | 343.65 | 344.43 | 0.00 | 9.99E-01 |
| ENSMUSG00000097637 | GSMG0037959 | 4933417D19Rik | RIKEN cDNA 4933417D19 gene | 111.35 | 38.20 | 1.54 | 1.93E-05 |
| ENSMUSG00000001065 | GSMG0037960 | Zfp276 | zinc finger protein (C2H2 type) 276 | 749.14 | 582.26 | 0.36 | 1.59E-01 |
| ENSMUSG00000010154 | GSMG0037962 | Spire2 | spire homolog 2 (Drosophila) | 253.05 | 458.02 | -0.86 | 1.10E-05 |
| ENSMUSG00000001472 | GSMG0037964 | Tcf25 | transcription factor 25 (basic helix-loop-helix) | 6050.58 | 5441.70 | 0.15 | 5.35E-01 |
| ENSMUSG00000062380 | GSMG0037966 | Tubb3 | tubulin, beta 3 class III | 1239.64 | 7036.62 | -2.51 | 1.10E-64 |
| ENSMUSG00000031967 | GSMG0037970 | Afg3l1 | AFG3-like AAA ATPase 1 | 1643.54 | 1905.32 | -0.21 | 2.52E-01 |
| ENSMUSG00000040220 | GSMG0037972 | Gas8 | growth arrest specific 8 | 648.78 | 714.68 | -0.14 | 5.58E-01 |
| ENSMUSG00000039960 | GSMG0037974 | Rhou | ras homolog family member U | 371.90 | 604.35 | -0.70 | 6.01E-03 |
| ENSMUSG00000019478 | GSMG0037976 | Rab4a | RAB4A, member RAS oncogene family | 619.34 | 641.85 | -0.05 | 8.64E-01 |
| ENSMUSG00000031976 | GSMG0037982 | Urb2 | URB2 ribosome biogenesis 2 homolog (S. cerevisiae) | 271.87 | 309.27 | -0.18 | 5.78E-01 |
| ENSMUSG00000089704 | GSMG0037983 | Galnt2 | UDP-N-acetyl-alpha-D-galactosamine:polypeptide N-acetylgalactosami | 912.85 | 1009.72 | -0.15 | 5.97E-01 |
| ENSMUSG00000031979 | GSMG0037985 | Cog2 | component of oligomeric golgi complex 2 | 1219.64 | 1211.69 | 0.01 | 9.74E-01 |

| | | | | | | | |
|---|---|---|---|---|---|---|---|
| ENSMUSG00000031982 | GSMG0037987 | Arv1 | ARV1 homolog, fatty acid homeostasis modulator | 126.56 | 207.82 | -0.72 | 9.03E-03 |
| ENSMUSG00000036913 | GSMG0037989 | Trim67 | tripartite motif-containing 67 | 44.42 | 1085.54 | -4.64 | 2.08E-94 |
| ENSMUSG00000043051 | GSMG0037992 | Disc1 | disrupted in schizophrenia 1 | 182.91 | 179.61 | 0.03 | 9.46E-01 |
| NULL | GSMG0037993 | NULL | NULL | 286.69 | 301.83 | -0.07 | 8.47E-01 |
| ENSMUSG00000031851 | GSMG0037998 | Ntpcr | nucleoside-triphosphatase, cancer-related | 499.60 | 440.69 | 0.18 | 4.89E-01 |
| ENSMUSG00000033998 | GSMG0038001 | Kcnk1 | potassium channel, subfamily K, member 1 | 781.40 | 332.58 | 1.23 | 1.96E-08 |
| ENSMUSG00000057060 | GSMG0038002 | Slc35f3 | solute carrier family 35, member F3 | 148.37 | 147.67 | 0.01 | 9.91E-01 |
| ENSMUSG00000051671 | GSMG0038006 | Coa6 | cytochrome c oxidase assembly factor 6 | 299.88 | 337.01 | -0.17 | 6.58E-01 |
| ENSMUSG00000025812 | GSMG0038013 | Pard3 | par-3 family cell polarity regulator | 501.79 | 516.14 | -0.04 | 9.17E-01 |
| NULL | GSMG0038020 | NULL | NULL | 443.01 | 616.47 | -0.48 | 4.62E-02 |
| ENSMUSG00000025810 | GSMG0038022 | Nrp1 | neuropilin 1 | 6258.10 | 7020.98 | -0.17 | 3.83E-01 |
| NULL | GSMG0038023 | NULL | NULL | 377.79 | 346.55 | 0.12 | 6.94E-01 |
| ENSMUSG00000025809 | GSMG0038026 | Itgb1 | integrin beta 1 (fibronectin receptor beta) | 23403.35 | 20152.36 | 0.22 | 2.89E-01 |
| ENSMUSG00000005534 | GSMG0038028 | Insr | insulin receptor | 2347.96 | 1774.14 | 0.41 | 6.11E-02 |
| ENSMUSG00000019470 | GSMG0038032 | Xab2 | XPA binding protein 2 | 2936.98 | 2857.73 | 0.04 | 8.94E-01 |
| NULL | GSMG0038044 | NULL | NULL | 191.89 | 258.41 | -0.43 | 1.22E-01 |
| ENSMUSG00000064125 | GSMG0038045 | Prr36 | proline rich 36 | 266.97 | 662.27 | -1.31 | 6.60E-12 |
| ENSMUSG00000048644 | GSMG0038046 | Ctxn1 | cortexin 1 | 1390.73 | 1642.81 | -0.24 | 3.04E-01 |
| ENSMUSG00000002949 | GSMG0038047 | Timm44 | translocase of inner mitochondrial membrane 44 | 2435.43 | 1967.59 | 0.31 | 8.88E-02 |
| ENSMUSG00000040028 | GSMG0038049 | Elavl1 | ELAV (embryonic lethal, abnormal vision)-like 1 (Hu antigen R) | 3195.74 | 3888.74 | -0.28 | 2.08E-01 |
| NULL | GSMG0038050 | NULL | NULL | 100.98 | 76.88 | 0.40 | 4.51E-01 |
| ENSMUSG00000022322 | GSMG0038052 | Shcbp1 | Shc SH2-domain binding protein 1 | 191.40 | 473.23 | -1.31 | 7.64E-08 |
| ENSMUSG00000001300 | GSMG0038057 | Efnb2 | ephrin B2 | 2531.07 | 2430.65 | 0.06 | 8.51E-01 |
| ENSMUSG00000040459 | GSMG0038060 | Arglu1 | arginine and glutamate rich 1 | 2297.76 | 2355.08 | -0.04 | 9.74E-01 |
| ENSMUSG00000079157 | GSMG0038064 | Fam155a | family with sequence similarity 155, member A | 390.80 | 426.29 | -0.13 | 6.64E-01 |
| NULL | GSMG0038066 | NULL | NULL | 380.22 | 311.73 | 0.29 | 2.51E-01 |
| NULL | GSMG0038068 | NULL | NULL | 853.75 | 666.16 | 0.36 | 6.50E-02 |
| NULL | GSMG0038070 | NULL | NULL | 644.54 | 527.72 | 0.29 | 2.12E-01 |
| NULL | GSMG0038071 | NULL | NULL | 561.99 | 470.95 | 0.25 | 3.62E-01 |
| NULL | GSMG0038074 | NULL | NULL | 510.65 | 373.54 | 0.45 | 3.32E-02 |
| NULL | GSMG0038075 | NULL | NULL | 129.65 | 84.55 | 0.61 | 8.06E-02 |
| NULL | GSMG0038076 | NULL | NULL | 451.56 | 296.57 | 0.61 | 3.55E-03 |
| ENSMUSG00000038894 | GSMG0038088 | Irs2 | insulin receptor substrate 2 | 979.07 | 654.98 | 0.58 | 8.45E-02 |

| | | | | | | | |
|---|---|---|---|---|---|---|---|
| ENSMUSG00000031502 | GSMG0038089 | Col4a1 | collagen, type IV, alpha 1 | 20712.14 | 21622.90 | -0.06 | 8.65E-01 |
| ENSMUSG00000056228 | GSMG0038093 | Cars2 | cysteinyl-tRNA synthetase 2 (mitochondrial)(putative) | 242.65 | 261.70 | -0.11 | 7.68E-01 |
| ENSMUSG00000031508 | GSMG0038094 | Ankrd10 | ankyrin repeat domain 10 | 617.33 | 647.68 | -0.07 | 8.15E-01 |
| ENSMUSG00000000759 | GSMG0038104 | Tubgcp3 | tubulin, gamma complex associated protein 3 | 1566.50 | 1626.06 | -0.05 | 8.32E-01 |
| ENSMUSG00000038542 | GSMG0038107 | Pcid2 | PCI domain containing 2 | 747.57 | 771.63 | -0.05 | 8.79E-01 |
| ENSMUSG00000038515 | GSMG0038108 | Grtp1 | GH regulated TBC protein 1 | 146.51 | 154.59 | -0.08 | 8.58E-01 |
| ENSMUSG00000038506 | GSMG0038110 | Dcun1d2 | DCN1, defective in cullin neddylation 1, domain containing 2 (S. cerev | 171.86 | 176.39 | -0.03 | 9.50E-01 |
| ENSMUSG00000031451 | GSMG0038114 | Gas6 | growth arrest specific 6 | 5705.24 | 2843.95 | 1.00 | 2.95E-11 |
| ENSMUSG00000031453 | GSMG0038117 | Rasa3 | RAS p21 protein activator 3 | 1010.81 | 1479.96 | -0.55 | 1.40E-03 |
| NULL | GSMG0038118 | NULL | NULL | 422.24 | 454.23 | -0.10 | 7.82E-01 |
| ENSMUSG00000031458 | GSMG0038121 | Coprs | coordinator of PRMT5, differentiation stimulator | 263.51 | 354.89 | -0.43 | 6.46E-02 |
| ENSMUSG00000050052 | GSMG0038124 | Tdrp | testis development related protein | 281.14 | 299.60 | -0.09 | 7.85E-01 |
| ENSMUSG00000051978 | GSMG0038125 | Erich1 | glutamate rich 1 | 118.09 | 110.07 | 0.11 | 8.41E-01 |
| ENSMUSG00000060924 | GSMG0038134 | Csmd1 | CUB and Sushi multiple domains 1 | 2405.79 | 1893.58 | 0.35 | 1.16E-01 |
| NULL | GSMG0038137 | NULL | NULL | 305.12 | 207.82 | 0.55 | 3.09E-02 |
| NULL | GSMG0038139 | NULL | NULL | 578.35 | 365.16 | 0.66 | 8.10E-04 |
| NULL | GSMG0038142 | NULL | NULL | 347.06 | 211.15 | 0.72 | 9.83E-04 |
| NULL | GSMG0038143 | NULL | NULL | 286.25 | 186.62 | 0.62 | 2.51E-02 |
| NULL | GSMG0038146 | NULL | NULL | 761.44 | 537.38 | 0.50 | 1.44E-02 |
| NULL | GSMG0038148 | NULL | NULL | 535.41 | 323.08 | 0.73 | 7.12E-04 |
| NULL | GSMG0038149 | NULL | NULL | 246.24 | 146.79 | 0.74 | 5.58E-03 |
| NULL | GSMG0038151 | NULL | NULL | 1616.33 | 931.51 | 0.80 | 7.47E-08 |
| NULL | GSMG0038152 | NULL | NULL | 2046.57 | 1202.60 | 0.77 | 8.37E-07 |
| ENSMUSG00000031465 | GSMG0038155 | Angpt2 | angiopoietin 2 | 1250.63 | 938.53 | 0.42 | 2.65E-02 |
| ENSMUSG00000031478 | GSMG0038179 | Nek3 | NIMA (never in mitosis gene a)-related expressed kinase 3 | 131.60 | 157.61 | -0.26 | 5.06E-01 |
| ENSMUSG00000037725 | GSMG0038180 | Ckap2 | cytoskeleton associated protein 2 | 243.50 | 636.18 | -1.38 | 8.79E-09 |
| ENSMUSG00000031482 | GSMG0038182 | Slc25a15 | solute carrier family 25 (mitochondrial carrier ornithine transporter), me | 271.02 | 297.00 | -0.13 | 7.28E-01 |
| ENSMUSG00000031534 | GSMG0038184 | Smim19 | small integral membrane protein 19 | 508.56 | 522.61 | -0.04 | 9.03E-01 |
| ENSMUSG00000031536 | GSMG0038186 | Polb | polymerase (DNA directed), beta | 1028.38 | 1310.87 | -0.35 | 1.02E-01 |
| NULL | GSMG0038187 | NULL | NULL | 106.14 | 99.11 | 0.09 | 8.65E-01 |
| ENSMUSG00000031537 | GSMG0038189 | Ikbkb | inhibitor of kappaB kinase beta | 2597.30 | 2567.02 | 0.02 | 9.50E-01 |
| ENSMUSG00000031539 | GSMG0038190 | Ap3m2 | adaptor-related protein complex 3, mu 2 subunit | 470.64 | 416.22 | 0.18 | 4.71E-01 |
| ENSMUSG00000031545 | GSMG0038193 | Gpat4 | glycerol-3-phosphate acyltransferase 4 | 1444.71 | 1824.99 | -0.34 | 5.82E-02 |

| | | | | | | | |
|---|---|---|---|---|---|---|---|
| ENSMUSG00000031546 | GSMG0038194 | Gins4 | GINS complex subunit 4 (Sld5 homolog) | 526.50 | 613.98 | -0.22 | 3.49E-01 |
| ENSMUSG00000015341 | GSMG0038195 | Golga7 | golgi autoantigen, golgin subfamily a, 7 | 2781.04 | 2909.19 | -0.07 | 8.31E-01 |
| ENSMUSG00000056313 | GSMG0038197 | 1810011O10Rik | RIKEN cDNA 1810011O10 gene | 332.54 | 220.17 | 0.60 | 2.93E-02 |
| ENSMUSG00000031555 | GSMG0038205 | Adam9 | a disintegrin and metallopeptidase domain 9 (meltrin gamma) | 1792.28 | 1458.67 | 0.30 | 1.12E-01 |
| ENSMUSG00000031557 | GSMG0038207 | Plekha2 | pleckstrin homology domain-containing, family A (phosphoinositide bi | 324.13 | 169.22 | 0.94 | 3.12E-05 |
| ENSMUSG00000065954 | GSMG0038209 | Tacc1 | transforming, acidic coiled-coil containing protein 1 | 865.66 | 700.21 | 0.31 | 1.45E-01 |
| ENSMUSG00000037363 | GSMG0038213 | Letm2 | leucine zipper-EF-hand containing transmembrane protein 2 | 97.54 | 81.11 | 0.26 | 5.84E-01 |
| ENSMUSG00000061313 | GSMG0038215 | Ddhd2 | DDHD domain containing 2 | 855.94 | 884.60 | -0.05 | 8.76E-01 |
| ENSMUSG00000037316 | GSMG0038217 | Bag4 | BCL2-associated athanogene 4 | 148.82 | 157.57 | -0.08 | 8.58E-01 |
| ENSMUSG00000031575 | GSMG0038219 | Ash2l | ash2 (absent, small, or homeotic)-like (Drosophila) | 1081.33 | 1453.51 | -0.43 | 5.15E-02 |
| NULL | GSMG0038220 | NULL | NULL | 194.66 | 168.35 | 0.21 | 5.47E-01 |
| ENSMUSG00000037260 | GSMG0038221 | Hgsnat | heparan-alpha-glucosaminide N-acetyltransferase | 1639.94 | 1659.97 | -0.02 | 9.50E-01 |
| ENSMUSG00000037251 | GSMG0038222 | Pomk | protein-O-mannose kinase | 155.38 | 186.38 | -0.26 | 4.33E-01 |
| ENSMUSG00000015994 | GSMG0038223 | Fnta | farnesyltransferase, CAAX box, alpha | 1567.52 | 1350.92 | 0.22 | 3.34E-01 |
| ENSMUSG00000037234 | GSMG0038226 | Hook3 | hook microtubule tethering protein 3 | 406.99 | 428.68 | -0.07 | 8.59E-01 |
| ENSMUSG00000031487 | GSMG0038234 | Brf2 | BRF2, RNA polymerase III transcription initiation factor 50kDa subuni | 352.75 | 376.63 | -0.09 | 7.74E-01 |
| ENSMUSG00000031488 | GSMG0038235 | Rab11fip1 | RAB11 family interacting protein 1 (class I) | 820.11 | 520.25 | 0.66 | 2.55E-03 |
| ENSMUSG00000063626 | GSMG0038241 | Unc5d | unc-5 netrin receptor D | 601.27 | 1077.84 | -0.84 | 6.68E-07 |
| NULL | GSMG0038242 | NULL | NULL | 567.37 | 469.22 | 0.28 | 2.13E-01 |
| NULL | GSMG0038244 | NULL | NULL | 431.75 | 419.23 | 0.05 | 9.11E-01 |
| NULL | GSMG0038246 | NULL | NULL | 554.31 | 335.72 | 0.73 | 3.21E-04 |
| ENSMUSG00000031578 | GSMG0038248 | Mak16 | MAK16 homolog | 657.97 | 804.42 | -0.29 | 1.76E-01 |
| ENSMUSG00000062991 | GSMG0038251 | Nrg1 | neuregulin 1 | 79.70 | 107.47 | -0.42 | 2.88E-01 |
| NULL | GSMG0038257 | NULL | NULL | 202.89 | 125.39 | 0.69 | 7.04E-02 |
| NULL | GSMG0038258 | NULL | NULL | 276.59 | 205.32 | 0.43 | 1.07E-01 |
| ENSMUSG00000031583 | GSMG0038265 | Wrn | Werner syndrome homolog (human) | 291.98 | 320.64 | -0.14 | 6.64E-01 |
| ENSMUSG00000052906 | GSMG0038268 | Ubxn8 | UBX domain protein 8 | 248.60 | 318.36 | -0.36 | 1.64E-01 |
| ENSMUSG00000094500 | GSMG0038270 | Smim18 | small integral membrane protein 18 | 69.10 | 151.55 | -1.14 | 8.90E-05 |
| ENSMUSG00000031586 | GSMG0038271 | Rbpms | RNA binding protein gene with multiple splicing | 1248.11 | 1354.79 | -0.12 | 6.13E-01 |
| NULL | GSMG0038272 | NULL | NULL | 847.05 | 488.32 | 0.79 | 5.20E-06 |
| NULL | GSMG0038275 | NULL | NULL | 436.36 | 160.19 | 1.44 | 2.41E-10 |
| ENSMUSG00000031516 | GSMG0038277 | Dctn6 | dynactin 6 | 809.39 | 840.81 | -0.05 | 8.51E-01 |
| ENSMUSG00000031513 | GSMG0038278 | Leprotl1 | leptin receptor overlapping transcript-like 1 | 1941.78 | 2436.66 | -0.33 | 1.04E-01 |

| | | | | | | | |
|---|---|---|---|---|---|---|---|
| ENSMUSG00000031529 | GSMG0038285 | Tnks | tankyrase, TRF1-interacting ankyrin-related ADP-ribose polymerase | 841.90 | 1003.38 | -0.25 | 4.22E-01 |
| ENSMUSG00000031527 | GSMG0038290 | Eri1 | exoribonuclease 1 | 267.33 | 392.93 | -0.55 | 2.61E-02 |
| ENSMUSG00000039633 | GSMG0038297 | Lonrf1 | LON peptidase N-terminal domain and ring finger 1 | 189.61 | 224.88 | -0.24 | 4.66E-01 |
| ENSMUSG00000031523 | GSMG0038299 | Dlc1 | deleted in liver cancer 1 | 2322.79 | 2007.81 | 0.21 | 3.17E-01 |
| NULL | GSMG0038301 | NULL | NULL | 1546.14 | 1330.01 | 0.22 | 4.70E-01 |
| NULL | GSMG0038302 | NULL | NULL | 468.56 | 330.50 | 0.50 | 5.29E-02 |
| NULL | GSMG0038323 | NULL | NULL | 151.75 | 74.19 | 1.03 | 2.72E-04 |
| NULL | GSMG0038327 | NULL | NULL | 57.35 | 28.24 | 1.01 | 5.36E-02 |
| ENSMUSG00000098431 | GSMG0038332 | Mir7666 | microRNA 7666 | 143.14 | 117.68 | 0.27 | 5.23E-01 |
| ENSMUSG00000031601 | GSMG0038334 | Cnot7 | CCR4-NOT transcription complex, subunit 7 | 669.60 | 982.97 | -0.55 | 2.11E-03 |
| ENSMUSG00000039431 | GSMG0038335 | Mtmr7 | myotubularin related protein 7 | 298.37 | 433.69 | -0.54 | 6.19E-03 |
| ENSMUSG00000045636 | GSMG0038337 | Mtus1 | mitochondrial tumor suppressor 1 | 1640.69 | 1254.53 | 0.39 | 2.98E-02 |
| ENSMUSG00000031591 | GSMG0038344 | Asah1 | N-acylsphingosine amidohydrolase 1 | 1317.58 | 1241.25 | 0.09 | 7.37E-01 |
| ENSMUSG00000031590 | GSMG0038346 | Frg1 | FSHD region gene 1 | 950.22 | 1170.17 | -0.30 | 2.56E-01 |
| G00000079057,ENSMUSG000 | GSMG0038360 | Klkb1 // Cyp4v3 | kallikrein B, plasma 1 // cytochrome P450, family 4, subfamily v, polyp | 163.32 | 152.45 | 0.11 | 8.09E-01 |
| ENSMUSG00000070044 | GSMG0038361 | Fam149a | family with sequence similarity 149, member A | 544.24 | 754.68 | -0.47 | 1.44E-02 |
| ENSMUSG00000031639 | GSMG0038362 | Tlr3 | toll-like receptor 3 | 656.14 | 758.03 | -0.21 | 4.78E-01 |
| ENSMUSG00000085440 | GSMG0038364 | Sorbs2os | sorbin and SH3 domain containing 2, opposite strand | 243.83 | 113.16 | 1.11 | 3.13E-05 |
| ENSMUSG00000071103 | GSMG0038365 | 1700029J07Rik | RIKEN cDNA 1700029J07 gene | 157.45 | 105.55 | 0.58 | 5.72E-02 |
| ENSMUSG00000050914 | GSMG0038366 | Ankrd37 | ankyrin repeat domain 37 | 963.31 | 282.97 | 1.77 | 3.54E-16 |
| ENSMUSG00000038291 | GSMG0038367 | Snx25 | sorting nexin 25 | 735.50 | 558.25 | 0.40 | 2.16E-01 |
| ENSMUSG00000031633 | GSMG0038369 | Slc25a4 | solute carrier family 25 (mitochondrial carrier, adenine nucleotide trans | 5064.35 | 5111.00 | -0.01 | 9.63E-01 |
| NULL | GSMG0038371 | NULL | NULL | 640.35 | 536.11 | 0.26 | 3.16E-01 |
| ENSMUSG00000038225 | GSMG0038372 | Primpol | primase and polymerase (DNA-directed) | 247.01 | 294.18 | -0.25 | 4.01E-01 |
| ENSMUSG00000038143 | GSMG0038377 | Stox2 | storkhead box 2 | 1135.20 | 1234.84 | -0.12 | 5.99E-01 |
| NULL | GSMG0038380 | NULL | NULL | 174.86 | 144.26 | 0.28 | 4.07E-01 |
| NULL | GSMG0038381 | NULL | NULL | 375.50 | 380.84 | -0.02 | 9.50E-01 |
| NULL | GSMG0038383 | NULL | NULL | 151.48 | 116.39 | 0.38 | 2.78E-01 |
| ENSMUSG00000038102 | GSMG0038384 | Trappc11 | trafficking protein particle complex 11 | 2330.21 | 2061.93 | 0.18 | 3.66E-01 |
| ENSMUSG00000063049 | GSMG0038385 | Ing2 | inhibitor of growth family, member 2 | 755.10 | 602.21 | 0.33 | 9.27E-02 |
| ENSMUSG00000038069 | GSMG0038386 | Cdkn2aip | CDKN2A interacting protein | 1099.83 | 1053.22 | 0.06 | 8.32E-01 |
| ENSMUSG00000031563 | GSMG0038389 | Wwc2 | WW, C2 and coiled-coil domain containing 2 | 722.22 | 651.37 | 0.15 | 5.99E-01 |
| NULL | GSMG0038391 | NULL | NULL | 156.97 | 210.49 | -0.43 | 2.05E-01 |

| | | | | | | | |
|---|---|---|---|---|---|---|---|
| NULL | GSMG0038398 | NULL | NULL | 165.95 | 221.29 | -0.42 | 1.78E-01 |
| ENSMUSG00000031561 | GSMG0038400 | Tenm3 | teneurin transmembrane protein 3 | 60.60 | 154.76 | -1.35 | 1.09E-04 |
| NULL | GSMG0038407 | NULL | NULL | 144.49 | 246.34 | -0.77 | 4.58E-03 |
| ENSMUSG00000039396 | GSMG0038431 | Neil3 | nei like 3 (E. coli) | 174.33 | 402.37 | -1.20 | 1.09E-06 |
| ENSMUSG00000054408 | GSMG0038433 | Spcs3 | signal peptidase complex subunit 3 homolog (S. cerevisiae) | 2733.23 | 3110.68 | -0.19 | 4.62E-01 |
| ENSMUSG00000039375 | GSMG0038435 | Wdr17 | WD repeat domain 17 | 109.24 | 157.04 | -0.52 | 7.26E-02 |
| ENSMUSG00000038215 | GSMG0038439 | Cep44 | centrosomal protein 44 | 285.94 | 252.83 | 0.18 | 5.97E-01 |
| ENSMUSG00000031609 | GSMG0038443 | Sap30 | sin3 associated polypeptide | 322.67 | 387.88 | -0.27 | 3.32E-01 |
| ENSMUSG00000031608 | GSMG0038444 | Galnt7 | UDP-N-acetyl-alpha-D-galactosamine: polypeptide N-acetylgalactosam | 304.07 | 329.86 | -0.12 | 7.08E-01 |
| NULL | GSMG0038456 | NULL | NULL | 143.12 | 191.11 | -0.42 | 2.04E-01 |
| NULL | GSMG0038472 | NULL | NULL | 278.67 | 381.40 | -0.45 | 3.72E-02 |
| ENSMUSG00000004319 | GSMG0038473 | Clcn3 | chloride channel, voltage-sensitive 3 | 4268.97 | 4744.07 | -0.15 | 4.39E-01 |
| ENSMUSG00000058056 | GSMG0038479 | Palld | palladin, cytoskeletal associated protein | 227.11 | 154.83 | 0.55 | 6.07E-02 |
| ENSMUSG00000037852 | GSMG0038486 | Cpe | carboxypeptidase E | 50248.05 | 34912.33 | 0.53 | 1.91E-04 |
| ENSMUSG00000031604 | GSMG0038489 | Msmo1 | methylsterol monoxygenase 1 | 720.06 | 1105.76 | -0.62 | 5.77E-03 |
| ENSMUSG00000031605 | GSMG0038491 | Klhl2 | kelch-like 2, Mayven | 1057.49 | 830.57 | 0.35 | 1.66E-01 |
| ENSMUSG00000030465 | GSMG0038506 | Psd3 | pleckstrin and Sec7 domain containing 3 | 136.55 | 100.24 | 0.45 | 2.06E-01 |
| NULL | GSMG0038507 | NULL | NULL | 228.59 | 152.61 | 0.59 | 2.11E-02 |
| ENSMUSG00000036356 | GSMG0038513 | Csgalnact1 | chondroitin sulfate N-acetylgalactosaminyltransferase 1 | 259.10 | 236.78 | 0.13 | 6.92E-01 |
| NULL | GSMG0038521 | NULL | NULL | 614.57 | 530.31 | 0.21 | 3.82E-01 |
| NULL | GSMG0038522 | NULL | NULL | 1478.59 | 1683.16 | -0.19 | 4.66E-01 |
| ENSMUSG00000060427 | GSMG0038529 | Zfp868 | zinc finger protein 868 | 537.62 | 585.81 | -0.12 | 6.29E-01 |
| G00000043090,ENSMUSG000 | GSMG0038532 | Zfp963 // Zfp866 | zinc finger protein 963 // zinc finger protein 866 | 740.89 | 649.29 | 0.19 | 4.86E-01 |
| ENSMUSG00000036199 | GSMG0038536 | Ndufa13 | NADH dehydrogenase (ubiquinone) 1 alpha subcomplex, 13 | 3473.39 | 3015.48 | 0.20 | 3.84E-01 |
| ENSMUSG00000036180 | GSMG0038537 | Gatad2a | GATA zinc finger domain containing 2A | 879.93 | 852.98 | 0.04 | 8.99E-01 |
| ENSMUSG00000031858 | GSMG0038541 | Mau2 | MAU2 sister chromatid cohesion factor | 4219.11 | 3762.45 | 0.17 | 4.35E-01 |
| ENSMUSG00000002341 | GSMG0038542 | Ncan | neurocan | 4370.89 | 11117.27 | -1.35 | 1.41E-18 |
| ENSMUSG00000036120 | GSMG0038543 | Rfxank | regulatory factor X-associated ankyrin-containing protein | 244.65 | 194.59 | 0.33 | 2.90E-01 |
| ENSMUSG00000002346 | GSMG0038544 | Slc25a42 | solute carrier family 25, member 42 | 512.75 | 526.27 | -0.04 | 9.16E-01 |
| ENSMUSG00000002343 | GSMG0038546 | Armc6 | armadillo repeat containing 6 | 536.49 | 726.86 | -0.44 | 2.30E-02 |
| ENSMUSG00000057788 | GSMG0038547 | Ddx49 | DEAD (Asp-Glu-Ala-Asp) box polypeptide 49 | 566.63 | 618.82 | -0.12 | 6.68E-01 |
| NULL | GSMG0038548 | NULL | NULL | 69.61 | 59.00 | 0.24 | 6.72E-01 |
| ENSMUSG00000058301 | GSMG0038549 | Upf1 | UPF1 regulator of nonsense transcripts homolog (yeast) | 1189.50 | 963.07 | 0.30 | 3.28E-01 |

| | | | | | | | |
|---|---|---|---|---|---|---|---|
| ENSMUSG00000003575 | GSMG0038550 | Crtc1 | CREB regulated transcription coactivator 1 | 534.86 | 570.15 | -0.09 | 7.94E-01 |
| ENSMUSG00000055707 | GSMG0038551 | Klhl26 | kelch-like 26 | 1421.80 | 1105.41 | 0.36 | 9.95E-02 |
| ENSMUSG00000035964 | GSMG0038552 | Tmem59l | transmembrane protein 59-like | 981.08 | 810.90 | 0.27 | 2.04E-01 |
| ENSMUSG00000058833 | GSMG0038553 | 2810428I15Rik | RIKEN cDNA 2810428I15 gene | 1123.72 | 1371.46 | -0.29 | 1.76E-01 |
| ENSMUSG00000055553 | GSMG0038556 | Kxd1 | KxDL motif containing 1 | 515.08 | 492.48 | 0.07 | 8.34E-01 |
| ENSMUSG00000070003 | GSMG0038557 | Ssbp4 | single stranded DNA binding protein 4 | 6551.40 | 5373.37 | 0.29 | 1.32E-01 |
| NULL | GSMG0038561 | NULL | NULL | 906.84 | 609.74 | 0.57 | 3.18E-02 |
| ENSMUSG00000035559 | GSMG0038562 | Mpv17l2 | MPV17 mitochondrial membrane protein-like 2 | 739.51 | 673.73 | 0.14 | 5.98E-01 |
| ENSMUSG00000031838 | GSMG0038563 | Ifi30 | interferon gamma inducible protein 30 | 150.03 | 191.88 | -0.35 | 2.98E-01 |
| ENSMUSG00000031834 | GSMG0038564 | Pik3r2 | phosphatidylinositol 3-kinase, regulatory subunit, polypeptide 2 (p85 be | 2740.79 | 3108.06 | -0.18 | 5.36E-01 |
| ENSMUSG00000031833 | GSMG0038565 | Mast3 | microtubule associated serine/threonine kinase 3 | 267.96 | 193.13 | 0.47 | 1.89E-01 |
| ENSMUSG00000002910 | GSMG0038566 | Arrdc2 | arrestin domain containing 2 | 373.25 | 342.85 | 0.12 | 6.96E-01 |
| ENSMUSG00000002908 | GSMG0038567 | Kcnn1 | potassium intermediate/small conductance calcium-activated channel, s | 113.45 | 206.61 | -0.86 | 3.70E-03 |
| ENSMUSG00000007721 | GSMG0038568 | Ccdc124 | coiled-coil domain containing 124 | 1451.28 | 1951.94 | -0.43 | 2.12E-02 |
| ENSMUSG00000000792 | GSMG0038569 | Slc5a5 | solute carrier family 5 (sodium iodide symporter), member 5 | 813.63 | 452.90 | 0.84 | 1.88E-05 |
| ENSMUSG00000035439 | GSMG0038572 | Haus8 | 4HAUS augmin-like complex, subunit 8 | 151.97 | 277.53 | -0.87 | 1.26E-04 |
| ENSMUSG00000002393 | GSMG0038573 | Nr2f6 | nuclear receptor subfamily 2, group F, member 6 | 2041.39 | 2213.05 | -0.12 | 6.29E-01 |
| ENSMUSG00000034911 | GSMG0038574 | Ushbp1 | Usher syndrome 1C binding protein 1 | 7868.25 | 7676.01 | 0.04 | 8.92E-01 |
| ENSMUSG00000007950 | GSMG0038576 | Abhd8 | abhydrolase domain containing 8 | 1262.14 | 1111.08 | 0.18 | 5.06E-01 |
| NULL | GSMG0038577 | NULL | NULL | 326.82 | 442.01 | -0.44 | 1.09E-01 |
| ENSMUSG00000034863 | GSMG0038579 | Ano8 | anoctamin 8 | 1006.36 | 1046.27 | -0.06 | 9.10E-01 |
| ENSMUSG00000034845 | GSMG0038580 | Plvap | plasmalemma vesicle associated protein | 19.24 | 117.69 | -2.64 | 1.21E-06 |
| ENSMUSG00000046718 | GSMG0038582 | Bst2 | bone marrow stromal cell antigen 2 | 390.19 | 452.63 | -0.21 | 5.07E-01 |
| NULL | GSMG0038586 | NULL | NULL | 206.16 | 212.36 | -0.04 | 9.15E-01 |
| ENSMUSG00000034799 | GSMG0038587 | Unc13a | unc-13 homolog A (C. elegans) | 288.71 | 275.99 | 0.06 | 9.11E-01 |
| ENSMUSG00000070000 | GSMG0038590 | Fcho1 | FCH domain only 1 | 3864.81 | 3735.29 | 0.05 | 8.90E-01 |
| NULL | GSMG0038594 | NULL | NULL | 114.75 | 72.65 | 0.66 | 8.28E-02 |
| ENSMUSG00000006276 | GSMG0038596 | Eps15l1 | epidermal growth factor receptor pathway substrate 15-like 1 | 1901.20 | 2209.46 | -0.22 | 3.32E-01 |
| ENSMUSG00000019732 | GSMG0038600 | Calr3 | calreticulin 3 | 79.10 | 66.23 | 0.26 | 6.35E-01 |
| ENSMUSG00000052488 | GSMG0038601 | Cherp | calcium homeostasis endoplasmic reticulum protein | 246.31 | 341.01 | -0.47 | 3.99E-02 |
| ENSMUSG00000019731 | GSMG0038602 | Slc35e1 | solute carrier family 35, member E1 | 299.75 | 301.24 | -0.01 | 9.87E-01 |
| ENSMUSG00000045248 | GSMG0038603 | Med26 | mediator complex subunit 26 | 495.14 | 558.18 | -0.17 | 4.94E-01 |
| ENSMUSG00000044600 | GSMG0038605 | Smim7 | small integral membrane protein 7 | 1800.95 | 2154.17 | -0.26 | 1.73E-01 |

| | | | | | | | |
|---|---|---|---|---|---|---|---|
| ENSMUSG00000004383 | GSMG0038606 | Large | like-glycosyltransferase | 500.92 | 569.03 | -0.18 | 5.74E-01 |
| ENSMUSG00000037148 | GSMG0038621 | Arhgap10 | Rho GTPase activating protein 10 | 326.74 | 202.34 | 0.69 | 1.39E-03 |
| ENSMUSG00000031617 | GSMG0038627 | Tmem184c | transmembrane protein 184C | 483.92 | 473.18 | 0.03 | 9.21E-01 |
| ENSMUSG00000031616 | GSMG0038629 | Ednra | endothelin receptor type A | 1096.99 | 1525.50 | -0.47 | 1.81E-02 |
| ENSMUSG00000031683 | GSMG0038633 | Lsm6 | LSM6 homolog, U6 small nuclear RNA and mRNA degradation associa | 1044.61 | 1428.24 | -0.45 | 5.76E-03 |
| ENSMUSG00000037022 | GSMG0038635 | Mmaa | methylmalonic aciduria (cobalamin deficiency) type A | 229.97 | 179.60 | 0.36 | 2.07E-01 |
| ENSMUSG00000031681 | GSMG0038636 | Smad1 | SMAD family member 1 | 1783.55 | 2539.95 | -0.51 | 8.58E-04 |
| ENSMUSG00000058355 | GSMG0038639 | Abce1 | ATP-binding cassette, sub-family E (OABP), member 1 | 2325.38 | 3033.39 | -0.38 | 6.36E-02 |
| ENSMUSG00000031715 | GSMG0038643 | Smarca5 | SWI/SNF related, matrix associated, actin dependent regulator of chrom | 849.21 | 1009.89 | -0.25 | 2.98E-01 |
| ENSMUSG00000031714 | GSMG0038644 | Gab1 | growth factor receptor bound protein 2-associated protein 1 | 1766.89 | 2281.66 | -0.37 | 2.79E-02 |
| ENSMUSG00000038250 | GSMG0038649 | Usp38 | ubiquitin specific peptidase 38 | 727.98 | 880.72 | -0.27 | 1.94E-01 |
| ENSMUSG00000031712 | GSMG0038651 | Il15 | interleukin 15 | 59.25 | 50.89 | 0.22 | 7.03E-01 |
| ENSMUSG00000031711 | GSMG0038653 | Zfp330 | zinc finger protein 330 | 446.40 | 495.77 | -0.15 | 5.82E-01 |
| ENSMUSG00000035151 | GSMG0038657 | Elmod2 | ELMO/CED-12 domain containing 2 | 195.05 | 202.65 | -0.05 | 8.96E-01 |
| ENSMUSG00000031708 | GSMG0038660 | Tecr | trans-2,3-enoyl-CoA reductase | 4203.75 | 5891.33 | -0.49 | 1.69E-03 |
| NULL | GSMG0038662 | NULL | NULL | 1144.29 | 1055.42 | 0.12 | 6.96E-01 |
| ENSMUSG00000057672 | GSMG0038663 | Pkn1 | protein kinase N1 | 2860.07 | 2915.21 | -0.03 | 9.41E-01 |
| ENSMUSG00000002885 | GSMG0038664 | Adgre5 | adhesion G protein-coupled receptor E5 | 4596.31 | 3233.56 | 0.51 | 7.54E-04 |
| ENSMUSG00000005465 | GSMG0038670 | Il27ra | interleukin 27 receptor, alpha | 228.20 | 317.01 | -0.47 | 8.70E-02 |
| ENSMUSG00000037103 | GSMG0038674 | Dcaf15 | DDB1 and CUL4 associated factor 15 | 714.90 | 995.88 | -0.48 | 5.05E-02 |
| ENSMUSG00000036686 | GSMG0038675 | Cc2d1a | coiled-coil and C2 domain containing 1A | 1272.28 | 1933.33 | -0.60 | 4.05E-03 |
| ENSMUSG00000056155 | GSMG0038676 | Nanos3 | nanos homolog 3 (Drosophila) | 47.33 | 90.29 | -0.94 | 1.39E-02 |
| ENSMUSG00000035671 | GSMG0038678 | Zswim4 | zinc finger SWIM-type containing 4 | 721.85 | 819.16 | -0.18 | 5.51E-01 |
| ENSMUSG00000004996 | GSMG0038680 | Mri1 | methylthioribose-1-phosphate isomerase 1 | 330.08 | 436.22 | -0.40 | 5.37E-02 |
| ENSMUSG00000004994 | GSMG0038681 | Ccdc130 | coiled-coil domain containing 130 | 311.39 | 267.07 | 0.22 | 4.23E-01 |
| ENSMUSG00000053560 | GSMG0038682 | Ier2 | immediate early response 2 | 4670.97 | 3319.47 | 0.49 | 2.55E-03 |
| ENSMUSG00000001910 | GSMG0038684 | Nacc1 | nucleus accumbens associated 1, BEN and BTB (POZ) domain contain | 898.06 | 1021.98 | -0.19 | 5.98E-01 |
| ENSMUSG00000001911 | GSMG0038687 | Nfix | nuclear factor I/X | 1344.50 | 2322.50 | -0.79 | 4.31E-05 |
| ENSMUSG00000053226 | GSMG0038689 | Dand5 | DAN domain family member 5, BMP antagonist | 130.13 | 86.28 | 0.60 | 5.76E-02 |
| ENSMUSG00000003813 | GSMG0038690 | Rad23a | RAD23 homolog A, nucleotide excision repair protein | 987.95 | 1072.61 | -0.12 | 6.24E-01 |
| ENSMUSG00000003814 | GSMG0038691 | Calr | calreticulin | 14963.42 | 19123.80 | -0.35 | 2.55E-02 |
| ENSMUSG00000003809 | GSMG0038692 | Gcdh | glutaryl-Coenzyme A dehydrogenase | 637.72 | 452.63 | 0.50 | 1.89E-02 |
| ENSMUSG00000053693 | GSMG0038693 | Mast1 | microtubule associated serine/threonine kinase 1 | 665.79 | 995.24 | -0.58 | 1.08E-02 |

| | | | | | | | |
|---|---|---|---|---|---|---|---|
| ENSMUSG00000052926 | GSMG0038694 | Rnaseh2a | ribonuclease H2, large subunit | 353.97 | 491.52 | -0.47 | 3.18E-02 |
| NULL | GSMG0038695 | NULL | NULL | 1773.52 | 1212.82 | 0.55 | 2.38E-03 |
| ENSMUSG00000052456 | GSMG0038697 | Asna1 | arsA arsenite transporter, ATP-binding, homolog 1 (bacterial) | 887.13 | 1067.47 | -0.27 | 2.32E-01 |
| NULL | GSMG0038700 | NULL | NULL | 216.29 | 233.99 | -0.11 | 7.56E-01 |
| ENSMUSG00000005150 | GSMG0038701 | Wdr83 | WD repeat domain containing 83 | 728.01 | 666.90 | 0.13 | 6.30E-01 |
| ENSMUSG00000031696 | GSMG0038704 | Vps35 | VPS35 retromer complex component | 3463.76 | 4428.50 | -0.35 | 5.27E-02 |
| ENSMUSG00000036934 | GSMG0038707 | 4921524J17Rik | RIKEN cDNA 4921524J17 gene | 655.25 | 740.63 | -0.18 | 6.44E-01 |
| ENSMUSG00000031701 | GSMG0038711 | Dnaja2 | DnaJ heat shock protein family (Hsp40) member A2 | 2962.54 | 3240.01 | -0.13 | 5.99E-01 |
| ENSMUSG00000036902 | GSMG0038712 | Neto2 | neuropilin (NRP) and tolloid (TLL)-like 2 | 392.04 | 1074.99 | -1.46 | 8.40E-17 |
| ENSMUSG00000031703 | GSMG0038714 | Itfg1 | integrin alpha FG-GAP repeat containing 1 | 1615.17 | 2018.39 | -0.32 | 6.80E-02 |
| ENSMUSG00000036840 | GSMG0038721 | Siah1a | seven in absentia 1A | 1299.59 | 1512.28 | -0.22 | 3.10E-01 |
| ENSMUSG00000045333 | GSMG0038730 | Zfp423 | zinc finger protein 423 | 1848.02 | 2382.46 | -0.37 | 1.84E-01 |
| ENSMUSG00000031660 | GSMG0038737 | Brd7 | bromodomain containing 7 | 2811.41 | 3276.16 | -0.22 | 2.53E-01 |
| ENSMUSG00000043668 | GSMG0038744 | Tox3 | TOX high mobility group box family member 3 | 81.73 | 151.15 | -0.89 | 2.38E-03 |
| ENSMUSG00000031667 | GSMG0038749 | Aktip | thymoma viral proto-oncogene 1 interacting protein | 1095.09 | 926.36 | 0.24 | 2.53E-01 |
| ENSMUSG00000031751 | GSMG0038776 | Amfr | autocrine motility factor receptor | 9201.18 | 9072.46 | 0.02 | 9.44E-01 |
| ENSMUSG00000031755 | GSMG0038778 | Bbs2 | Bardet-Biedl syndrome 2 (human) | 764.25 | 716.34 | 0.09 | 7.49E-01 |
| ENSMUSG00000031774 | GSMG0038781 | Fam192a | family with sequence similarity 192, member A | 524.77 | 619.56 | -0.24 | 3.19E-01 |
| ENSMUSG00000031775 | GSMG0038782 | Pllp | plasma membrane proteolipid | 1579.66 | 406.26 | 1.96 | 1.56E-25 |
| ENSMUSG00000031781 | GSMG0038784 | Ciapin1 | cytokine induced apoptosis inhibitor 1 | 1712.70 | 2101.17 | -0.30 | 9.94E-02 |
| ENSMUSG00000040631 | GSMG0038785 | Dok4 | docking protein 4 | 2834.31 | 4213.14 | -0.57 | 6.82E-04 |
| ENSMUSG00000063605 | GSMG0038786 | Ccdc102a | coiled-coil domain containing 102A | 566.97 | 545.63 | 0.06 | 8.55E-01 |
| NULL | GSMG0038788 | NULL | NULL | 137.33 | 153.30 | -0.15 | 7.78E-01 |
| ENSMUSG00000031788 | GSMG0038789 | Kifc3 | kinesin family member C3 | 1278.69 | 1013.88 | 0.33 | 1.43E-01 |
| ENSMUSG00000046556 | GSMG0038792 | Zfp319 | zinc finger protein 319 | 1662.94 | 1833.73 | -0.14 | 6.20E-01 |
| ENSMUSG00000031796 | GSMG0038793 | Cfap20 | cilia and flagella associated protein 20 | 1742.70 | 2075.11 | -0.25 | 1.62E-01 |
| ENSMUSG00000046707 | GSMG0038794 | Csnk2a2 | casein kinase 2, alpha prime polypeptide | 387.47 | 274.96 | 0.49 | 1.22E-01 |
| ENSMUSG00000036550 | GSMG0038797 | Cnot1 | CCR4-NOT transcription complex, subunit 1 | 2257.11 | 2773.89 | -0.30 | 1.48E-01 |
| ENSMUSG00000036534 | GSMG0038799 | Slc38a7 | solute carrier family 38, member 7 | 226.04 | 307.71 | -0.44 | 1.36E-01 |
| ENSMUSG00000036510 | GSMG0038803 | Cdh8 | cadherin 8 | 797.15 | 1463.12 | -0.87 | 3.09E-05 |
| ENSMUSG00000031673 | GSMG0038810 | Cdh11 | cadherin 11 | 2052.43 | 3228.90 | -0.65 | 3.29E-05 |
| NULL | GSMG0038813 | NULL | NULL | 418.14 | 438.62 | -0.07 | 8.36E-01 |
| ENSMUSG00000035824 | GSMG0038819 | Tk2 | thymidine kinase 2, mitochondrial | 204.36 | 208.32 | -0.03 | 9.52E-01 |

| | | | | | | | |
|---|---|---|---|---|---|---|---|
| ENSMUSG00000035770 | GSMG0038825 | Dync1li2 | dynein, cytoplasmic 1 light intermediate chain 2 | 2820.31 | 2451.86 | 0.20 | 2.70E-01 |
| ENSMUSG00000031878 | GSMG0038827 | Nae1 | NEDD8 activating enzyme E1 subunit 1 | 1217.01 | 1338.66 | -0.14 | 6.13E-01 |
| ENSMUSG00000031880 | GSMG0038829 | Rrad | Ras-related associated with diabetes | 347.55 | 128.67 | 1.44 | 7.51E-12 |
| ENSMUSG00000031879 | GSMG0038830 | Fam96b | family with sequence similarity 96, member B | 674.69 | 695.96 | -0.05 | 9.03E-01 |
| ENSMUSG00000031887 | GSMG0038833 | Tradd | TNFRSF1A-associated via death domain | 211.08 | 187.20 | 0.17 | 6.18E-01 |
| ENSMUSG00000014837 | GSMG0038835 | 4931428F04Rik | RIKEN cDNA 4931428F04 gene | 166.83 | 582.99 | -1.80 | 8.40E-25 |
| ENSMUSG00000043251 | GSMG0038836 | Exoc3l | exocyst complex component 3-like | 1067.96 | 2060.10 | -0.95 | 8.20E-11 |
| ENSMUSG00000014846 | GSMG0038842 | Tppp3 | tubulin polymerization-promoting protein family member 3 | 508.07 | 775.98 | -0.61 | 1.32E-03 |
| ENSMUSG00000039199 | GSMG0038843 | Zdhhc1 | zinc finger, DHHC domain containing 1 | 1488.21 | 1026.05 | 0.54 | 3.47E-03 |
| ENSMUSG00000013160 | GSMG0038844 | Atp6v0d1 | ATPase, H+ transporting, lysosomal V0 subunit D1 | 2695.11 | 2832.04 | -0.07 | 7.93E-01 |
| NULL | GSMG0038845 | NULL | NULL | 166.05 | 113.48 | 0.55 | 6.73E-02 |
| ENSMUSG00000038000 | GSMG0038849 | Acd | adrenocortical dysplasia | 1289.42 | 1339.78 | -0.05 | 8.37E-01 |
| ENSMUSG00000013155 | GSMG0038850 | Enkd1 | enkurin domain containing 1 | 261.13 | 309.37 | -0.25 | 3.87E-01 |
| ENSMUSG00000013150 | GSMG0038851 | Gfod2 | glucose-fructose oxidoreductase domain containing 2 | 1077.23 | 1466.81 | -0.44 | 6.57E-03 |
| ENSMUSG00000037415 | GSMG0038853 | Ranbp10 | RAN binding protein 10 | 728.65 | 905.00 | -0.31 | 1.24E-01 |
| ENSMUSG00000036672 | GSMG0038855 | Cenpt | centromere protein T | 593.55 | 1015.55 | -0.77 | 4.42E-06 |
| ENSMUSG00000031897 | GSMG0038858 | Psmb10 | proteasome (prosome, macropain) subunit, beta type 10 | 567.51 | 551.58 | 0.04 | 9.11E-01 |
| ENSMUSG00000035237 | GSMG0038859 | Lcat | lecithin cholesterol acyltransferase | 267.37 | 84.32 | 1.67 | 2.26E-10 |
| ENSMUSG00000017765 | GSMG0038860 | Slc12a4 | solute carrier family 12, member 4 | 2457.33 | 1729.33 | 0.51 | 9.10E-03 |
| ENSMUSG00000045538 | GSMG0038863 | Ddx28 | DEAD (Asp-Glu-Ala-Asp) box polypeptide 28 | 459.10 | 516.00 | -0.17 | 4.99E-01 |
| ENSMUSG00000033106 | GSMG0038869 | Slc7a6os | solute carrier family 7, member 6 opposite strand | 485.26 | 535.54 | -0.14 | 5.84E-01 |
| ENSMUSG00000031906 | GSMG0038871 | Smpd3 | sphingomyelin phosphodiesterase 3, neutral | 131.34 | 318.87 | -1.28 | 1.76E-05 |
| ENSMUSG00000046691 | GSMG0038875 | Chtf8 | CTF8, chromosome transmission fidelity factor 8 | 972.23 | 1113.13 | -0.19 | 5.39E-01 |
| ENSMUSG00000031921 | GSMG0038884 | Terf2 | telomeric repeat binding factor 2 | 156.33 | 198.27 | -0.35 | 3.05E-01 |
| ENSMUSG00000003849 | GSMG0038886 | Nqo1 | NAD(P)H dehydrogenase, quinone 1 | 62.95 | 53.44 | 0.24 | 7.07E-01 |
| ENSMUSG00000003848 | GSMG0038887 | Nob1 | NIN1/RPN12 binding protein 1 homolog | 326.98 | 438.68 | -0.43 | 5.23E-02 |
| ENSMUSG00000039067 | GSMG0038889 | Psmd7 | proteasome (prosome, macropain) 26S subunit, non-ATPase, 7 | 2527.90 | 2996.60 | -0.25 | 2.68E-01 |
| ENSMUSG00000110528 | GSMG0038897 | Gm1943 | predicted gene 1943 | 216.88 | 290.48 | -0.42 | 1.54E-01 |
| ENSMUSG00000037993 | GSMG0038899 | Dhx38 | DEAH (Asp-Glu-Ala-His) box polypeptide 38 | 1094.28 | 1207.37 | -0.14 | 6.12E-01 |
| ENSMUSG00000031730 | GSMG0038901 | Dhodh | dihydroorotate dehydrogenase | 520.46 | 628.91 | -0.27 | 2.20E-01 |
| ENSMUSG00000031729 | GSMG0038902 | Ist1 | increased sodium tolerance 1 homolog (yeast) | 2103.74 | 2151.95 | -0.03 | 9.07E-01 |
| NULL | GSMG0038906 | NULL | NULL | 199.22 | 265.88 | -0.42 | 9.97E-02 |
| ENSMUSG00000003657 | GSMG0038910 | Calb2 | calbindin 2 | 67.28 | 83.90 | -0.32 | 5.33E-01 |

| | | | | | | | |
|---|---|---|---|---|---|---|---|
| ENSMUSG00000031750 | GSMG0038913 | Il34 | interleukin 34 | 1004.69 | 405.06 | 1.31 | 1.64E-12 |
| ENSMUSG00000033732 | GSMG0038915 | Sf3b3 | splicing factor 3b, subunit 3 | 2846.87 | 3452.78 | -0.28 | 1.96E-01 |
| ENSMUSG00000033703 | GSMG0038916 | Fuk | fucokinase | 725.55 | 647.95 | 0.16 | 6.15E-01 |
| ENSMUSG00000003316 | GSMG0038923 | Glg1 | golgi apparatus protein 1 | 2074.29 | 2174.88 | -0.07 | 8.21E-01 |
| ENSMUSG00000012519 | GSMG0038929 | Mlkl | mixed lineage kinase domain-like | 235.13 | 212.20 | 0.15 | 7.42E-01 |
| ENSMUSG00000033579 | GSMG0038930 | Fa2h | fatty acid 2-hydroxylase | 117.21 | 13.35 | 3.13 | 1.34E-13 |
| ENSMUSG00000031959 | GSMG0038931 | Wdr59 | WD repeat domain 59 | 353.99 | 335.81 | 0.07 | 8.43E-01 |
| ENSMUSG00000031958 | GSMG0038932 | Ldhd | lactate dehydrogenase D | 214.39 | 62.43 | 1.78 | 3.03E-09 |
| ENSMUSG00000031955 | GSMG0038934 | Bcar1 | breast cancer anti-estrogen resistance 1 | 2144.77 | 2502.74 | -0.22 | 2.41E-01 |
| ENSMUSG00000031954 | GSMG0038935 | Cfdp1 | craniofacial development protein 1 | 1393.86 | 1823.42 | -0.39 | 5.64E-02 |
| ENSMUSG00000031951 | GSMG0038939 | Tmem231 | transmembrane protein 231 | 54.53 | 88.65 | -0.70 | 6.18E-02 |
| ENSMUSG00000031949 | GSMG0038940 | Adat1 | adenosine deaminase, tRNA-specific 1 | 244.45 | 299.99 | -0.29 | 3.24E-01 |
| ENSMUSG00000031948 | GSMG0038941 | Kars | lysyl-tRNA synthetase | 1647.04 | 2518.47 | -0.61 | 3.91E-05 |
| ENSMUSG00000031758 | GSMG0038961 | Cdyl2 | chromodomain protein, Y chromosome-like 2 | 250.48 | 346.28 | -0.47 | 7.12E-02 |
| ENSMUSG00000034424 | GSMG0038968 | Gcsh | glycine cleavage system protein H (aminomethyl carrier) | 794.72 | 920.77 | -0.21 | 3.07E-01 |
| ENSMUSG00000031843 | GSMG0038973 | Mphosph6 | M phase phosphoprotein 6 | 483.60 | 661.35 | -0.45 | 2.33E-02 |
| NULL | GSMG0038974 | NULL | NULL | 1422.01 | 1593.78 | -0.17 | 5.23E-01 |
| ENSMUSG00000031835 | GSMG0038976 | Mbtps1 | membrane-bound transcription factor peptidase, site 1 | 5253.80 | 4969.66 | 0.08 | 7.50E-01 |
| ENSMUSG00000034189 | GSMG0038978 | Hsdl1 | hydroxysteroid dehydrogenase like 1 | 417.37 | 503.29 | -0.27 | 2.43E-01 |
| ENSMUSG00000031832 | GSMG0038979 | Taf1c | TATA-box binding protein associated factor, RNA polymerase I, C | 463.23 | 470.27 | -0.02 | 9.64E-01 |
| ENSMUSG00000034105 | GSMG0038983 | Tldc1 | TBC/LysM associated domain containing 1 | 127.01 | 110.98 | 0.19 | 6.74E-01 |
| ENSMUSG00000031827 | GSMG0038984 | Cotl1 | coactosin-like 1 (Dictyostelium) | 672.95 | 1922.68 | -1.51 | 1.05E-02 |
| ENSMUSG00000031823 | GSMG0038987 | Zdhhc7 | zinc finger, DHHC domain containing 7 | 737.24 | 643.04 | 0.20 | 4.43E-01 |
| ENSMUSG00000031821 | GSMG0038991 | Gins2 | GINS complex subunit 2 (Psf2 homolog) | 348.05 | 602.96 | -0.79 | 4.00E-05 |
| ENSMUSG00000097336 | GSMG0038994 | Fendrr | Foxf1 adjacent non-coding developmental regulatory RNA | 241.58 | 143.14 | 0.75 | 1.35E-03 |
| ENSMUSG00000031816 | GSMG0038997 | Mthfsd | methenyltetrahydrofolate synthetase domain containing | 855.92 | 729.28 | 0.23 | 3.17E-01 |
| ENSMUSG00000052934 | GSMG0039003 | Fbxo31 | F-box protein 31 | 543.23 | 572.59 | -0.08 | 8.23E-01 |
| NULL | GSMG0039004 | NULL | NULL | 3000.06 | 2471.56 | 0.28 | 1.14E-01 |
| ENSMUSG00000061410 | GSMG0039005 | Zcchc14 | zinc finger, CCHC domain containing 14 | 309.90 | 385.09 | -0.31 | 1.75E-01 |
| NULL | GSMG0039006 | NULL | NULL | 326.91 | 303.50 | 0.11 | 7.70E-01 |
| ENSMUSG00000040263 | GSMG0039008 | Klhdc4 | kelch domain containing 4 | 588.42 | 603.63 | -0.03 | 9.20E-01 |
| ENSMUSG00000040010 | GSMG0039010 | Slc7a5 | solute carrier family 7 (cationic amino acid transporter, y+ system), mer | 75622.81 | 100133.82 | -0.41 | 2.76E-02 |
| ENSMUSG00000006519 | GSMG0039015 | Cyba | cytochrome b-245, alpha polypeptide | 1969.16 | 1994.89 | -0.02 | 9.55E-01 |

| | | | | | | | |
|---|---|---|---|---|---|---|---|
| ENSMUSG00000006517 | GSMG0039016 | Mvd | mevalonate (diphospho) decarboxylase | 286.23 | 613.59 | -1.10 | 5.32E-09 |
| ENSMUSG00000014470 | GSMG0039019 | Rnf166 | ring finger protein 166 | 854.18 | 643.20 | 0.41 | 3.52E-02 |
| ENSMUSG00000014444 | GSMG0039020 | Piezo1 | piezo-type mechanosensitive ion channel component 1 | 2185.03 | 2222.07 | -0.02 | 9.56E-01 |
| ENSMUSG00000006589 | GSMG0039022 | Aprt | adenine phosphoribosyl transferase | 1064.37 | 1775.54 | -0.74 | 3.37E-06 |
| ENSMUSG00000015027 | GSMG0039023 | Galns | galactosamine (N-acetyl)-6-sulfate sulfatase | 150.14 | 201.47 | -0.42 | 1.74E-01 |
| ENSMUSG00000006362 | GSMG0039026 | Cbfa2t3 | core-binding factor, runt domain, alpha subunit 2, translocated to, 3 (hu | 678.67 | 481.90 | 0.49 | 6.26E-02 |
| NULL | GSMG0039027 | NULL | NULL | 599.79 | 394.52 | 0.61 | 6.87E-04 |
| ENSMUSG00000035569 | GSMG0039029 | Ankrd11 | ankyrin repeat domain 11 | 2387.66 | 2024.37 | 0.24 | 3.87E-01 |
| NULL | GSMG0039031 | NULL | NULL | 367.64 | 341.51 | 0.11 | 8.41E-01 |
| ENSMUSG00000000743 | GSMG0039037 | Chmp1a | charged multivesicular body protein 1A | 875.78 | 881.52 | -0.01 | 9.74E-01 |
| ENSMUSG00000001062 | GSMG0039039 | Vps9d1 | VPS9 domain containing 1 | 1445.37 | 1343.70 | 0.11 | 7.12E-01 |
| NULL | GSMG0039040 | NULL | NULL | 169.31 | 164.28 | 0.04 | 9.24E-01 |
| ENSMUSG00000032815 | GSMG0039041 | Fanca | Fanconi anemia, complementation group A | 93.58 | 202.38 | -1.12 | 8.67E-06 |
| ENSMUSG00000031971 | GSMG0039050 | Ccsap | centriole, cilia and spindle associated protein | 213.47 | 154.00 | 0.47 | 8.19E-02 |
| ENSMUSG00000039509 | GSMG0039052 | Nup133 | nucleoporin 133 | 840.57 | 1045.60 | -0.31 | 1.07E-01 |
| ENSMUSG00000031974 | GSMG0039053 | Abcb10 | ATP-binding cassette, sub-family B (MDR/TAP), member 10 | 971.27 | 1280.80 | -0.40 | 4.03E-02 |
| ENSMUSG00000038697 | GSMG0039054 | Taf5l | TATA-box binding protein associated factor 5 like | 548.12 | 682.49 | -0.31 | 1.82E-01 |
| ENSMUSG00000050751 | GSMG0039057 | Pgbd5 | piggyBac transposable element derived 5 | 1270.33 | 624.95 | 1.02 | 1.40E-06 |
| ENSMUSG00000031980 | GSMG0039058 | Agt | angiotensinogen (serpin peptidase inhibitor, clade A, member 8) | 644.99 | 254.26 | 1.34 | 4.89E-07 |
| ENSMUSG00000031983 | GSMG0039059 | 2310022B05Rik | RIKEN cDNA 2310022B05 gene | 623.84 | 504.97 | 0.30 | 2.48E-01 |
| ENSMUSG00000037300 | GSMG0039060 | Ttc13 | tetratricopeptide repeat domain 13 | 1663.00 | 2186.25 | -0.40 | 1.82E-02 |
| ENSMUSG00000031984 | GSMG0039062 | 2810004N23Rik | RIKEN cDNA 2810004N23 gene | 312.75 | 387.90 | -0.31 | 2.67E-01 |
| ENSMUSG00000031987 | GSMG0039065 | Egln1 | egl-9 family hypoxia-inducible factor 1 | 2510.31 | 1366.81 | 0.88 | 1.43E-03 |
| ENSMUSG00000001995 | GSMG0039067 | Sipa1l2 | signal-induced proliferation-associated 1 like 2 | 984.14 | 1051.09 | -0.10 | 7.96E-01 |
| ENSMUSG00000060212 | GSMG0039069 | Pcnx2 | pecanex homolog 2 | 560.77 | 385.76 | 0.54 | 3.99E-02 |
| ENSMUSG00000093904 | GSMG0039077 | Tomm20 | translocase of outer mitochondrial membrane 20 homolog (yeast) | 2099.02 | 2471.40 | -0.24 | 1.80E-01 |
| ENSMUSG00000033931 | GSMG0039078 | Rbm34 | RNA binding motif protein 34 | 185.82 | 264.57 | -0.51 | 7.78E-02 |
| ENSMUSG00000025899 | GSMG0039100 | Alkbh8 | alkB homolog 8, tRNA methyltransferase | 163.77 | 240.12 | -0.55 | 3.20E-02 |
| ENSMUSG00000025898 | GSMG0039102 | Cwf19l2 | CWF19-like 2, cell cycle control (S. pombe) | 603.85 | 661.20 | -0.13 | 6.92E-01 |
| ENSMUSG00000041624 | GSMG0039104 | Gucy1a2 | guanylate cyclase 1, soluble, alpha 2 | 339.94 | 308.49 | 0.14 | 6.59E-01 |
| ENSMUSG00000025893 | GSMG0039107 | Kbtbd3 | kelch repeat and BTB (POZ) domain containing 3 | 377.80 | 395.32 | -0.07 | 8.72E-01 |
| ENSMUSG00000041124 | GSMG0039108 | Msantd4 | Myb/SANT-like DNA-binding domain containing 4 with coiled-coils | 218.17 | 205.21 | 0.09 | 8.35E-01 |
| ENSMUSG00000025887 | GSMG0039111 | Casp12 | caspase 12 | 176.74 | 74.70 | 1.25 | 1.97E-04 |

| | | | | | | | |
|---|---|---|---|---|---|---|---|
| ENSMUSG00000032006 | GSMG0039112 | Pdgfd | platelet-derived growth factor, D polypeptide | 680.46 | 262.55 | 1.37 | 8.46E-12 |
| ENSMUSG00000032002 | GSMG0039116 | Dcun1d5 | DCN1, defective in cullin neddylation 1, domain containing 5 (S. cerev | 1254.79 | 1282.33 | -0.03 | 9.30E-01 |
| ENSMUSG00000050912 | GSMG0039128 | Tmem123 | transmembrane protein 123 | 5866.57 | 4917.50 | 0.25 | 2.46E-01 |
| ENSMUSG00000031997 | GSMG0039135 | Trpc6 | transient receptor potential cation channel, subfamily C, member 6 | 175.29 | 185.55 | -0.08 | 8.74E-01 |
| ENSMUSG00000031802 | GSMG0039144 | Phxr4 | per-hexamer repeat gene 4 | 327.23 | 339.32 | -0.05 | 8.93E-01 |
| ENSMUSG00000031925 | GSMG0039147 | Maml2 | mastermind like 2 (Drosophila) | 503.72 | 625.40 | -0.31 | 1.55E-01 |
| ENSMUSG00000031918 | GSMG0039152 | Mtmr2 | myotubularin related protein 2 | 2229.16 | 2128.45 | 0.07 | 7.85E-01 |
| ENSMUSG00000037808 | GSMG0039153 | Fam76b | family with sequence similarity 76, member B | 530.09 | 527.50 | 0.01 | 9.81E-01 |
| ENSMUSG00000032009 | GSMG0039155 | Sesn3 | sestrin 3 | 266.98 | 301.67 | -0.18 | 5.53E-01 |
| NULL | GSMG0039158 | NULL | NULL | 428.52 | 362.95 | 0.24 | 3.42E-01 |
| ENSMUSG00000004096 | GSMG0039159 | Cwc15 | CWC15 spliceosome-associated protein | 1107.78 | 1275.90 | -0.20 | 4.48E-01 |
| ENSMUSG00000031928 | GSMG0039162 | Mre11a | MRE11A homolog A, double strand break repair nuclease | 255.05 | 370.54 | -0.54 | 1.12E-02 |
| ENSMUSG00000031937 | GSMG0039164 | Vstm5 | V-set and transmembrane domain containing 5 | 257.93 | 388.99 | -0.59 | 1.69E-02 |
| ENSMUSG00000031939 | GSMG0039165 | Taf1d | TATA-box binding protein associated factor, RNA polymerase I, D | 625.17 | 817.48 | -0.39 | 1.22E-01 |
| NULL | GSMG0039166 | NULL | NULL | 137.29 | 182.85 | -0.41 | 2.14E-01 |
| NULL | GSMG0039167 | NULL | NULL | 1074.96 | 1163.32 | -0.11 | 7.19E-01 |
| ENSMUSG00000043885 | GSMG0039170 | Slc36a4 | solute carrier family 36 (proton/amino acid symporter), member 4 | 438.57 | 432.77 | 0.02 | 9.55E-01 |
| ENSMUSG00000001774 | GSMG0039175 | Chordc1 | cysteine and histidine-rich domain (CHORD)-containing, zinc-binding | 668.83 | 643.84 | 0.06 | 8.61E-01 |
| ENSMUSG00000057551 | GSMG0039185 | Zfp317 | zinc finger protein 317 | 233.05 | 259.30 | -0.16 | 6.31E-01 |
| NULL | GSMG0039194 | NULL | NULL | 475.28 | 632.27 | -0.41 | 4.26E-02 |
| ENSMUSG00000058192 | GSMG0039195 | Zfp846 | zinc finger protein 846 | 394.43 | 609.44 | -0.63 | 6.26E-04 |
| ENSMUSG00000084786 | GSMG0039199 | Ubl5 | ubiquitin-like 5 | 1686.89 | 1952.76 | -0.21 | 4.53E-01 |
| ENSMUSG00000032171 | GSMG0039200 | Pin1 | protein (peptidyl-prolyl cis/trans isomerase) NIMA-interacting 1 | 1322.63 | 1408.01 | -0.09 | 7.34E-01 |
| ENSMUSG00000038884 | GSMG0039203 | A230050P20Rik | RIKEN cDNA A230050P20 gene | 915.74 | 689.66 | 0.41 | 6.89E-02 |
| ENSMUSG00000004100 | GSMG0039204 | Ppan | peter pan homolog (Drosophila) | 664.71 | 915.80 | -0.46 | 8.72E-03 |
| ENSMUSG00000003299 | GSMG0039206 | Mrpl4 | mitochondrial ribosomal protein L4 | 2297.75 | 2242.81 | 0.04 | 8.93E-01 |
| ENSMUSG00000037405 | GSMG0039207 | Icam1 | intercellular adhesion molecule 1 | 2146.20 | 1591.71 | 0.43 | 1.59E-02 |
| ENSMUSG00000032174 | GSMG0039209 | Icam5 | intercellular adhesion molecule 5, telencephalin | 2484.18 | 737.55 | 1.75 | 2.69E-17 |
| ENSMUSG00000032177 | GSMG0039211 | Pde4a | phosphodiesterase 4A, cAMP specific | 875.72 | 578.87 | 0.60 | 1.05E-02 |
| ENSMUSG00000002820 | GSMG0039214 | Atg4d | autophagy related 4D, cysteine peptidase | 780.94 | 601.20 | 0.38 | 1.03E-01 |
| ENSMUSG00000032185 | GSMG0039215 | Carm1 | coactivator-associated arginine methyltransferase 1 | 1767.20 | 1572.27 | 0.17 | 6.51E-01 |
| ENSMUSG00000057193 | GSMG0039217 | Slc44a2 | solute carrier family 44, member 2 | 1455.03 | 1639.52 | -0.17 | 4.43E-01 |
| ENSMUSG00000032178 | GSMG0039218 | Ilf3 | interleukin enhancer binding factor 3 | 1575.95 | 2758.56 | -0.81 | 3.52E-06 |

| | | | | | | | |
|---|---|---|---|---|---|---|---|
| ENSMUSG00000002825 | GSMG0039220 | Qtrt1 | queuine tRNA-ribosyltransferase 1 | 481.32 | 572.07 | -0.25 | 2.77E-01 |
| ENSMUSG00000033335 | GSMG0039221 | Dnm2 | dynamin 2 | 6130.31 | 6033.99 | 0.02 | 9.29E-01 |
| ENSMUSG00000048429 | GSMG0039225 | 1810026J23Rik | RIKEN cDNA 1810026J23 gene | 2074.07 | 2384.29 | -0.20 | 3.11E-01 |
| ENSMUSG00000032187 | GSMG0039226 | Smarca4 | SWI/SNF related, matrix associated, actin dependent regulator of chron | 3488.00 | 4481.21 | -0.36 | 2.06E-02 |
| ENSMUSG00000032193 | GSMG0039227 | Ldlr | low density lipoprotein receptor | 634.52 | 1189.03 | -0.91 | 3.80E-06 |
| ENSMUSG00000006241 | GSMG0039229 | Ccdc159 | coiled-coil domain containing 159 | 309.88 | 348.92 | -0.17 | 6.55E-01 |
| ENSMUSG00000040563 | GSMG0039230 | Plppr2 | phospholipid phosphatase related 2 | 1873.47 | 2385.71 | -0.35 | 8.12E-02 |
| ENSMUSG00000051238 | GSMG0039231 | Swsap1 | SWIM type zinc finger 7 associated protein 1 | 430.40 | 412.00 | 0.06 | 8.44E-01 |
| ENSMUSG00000003402 | GSMG0039232 | Prkcsh | protein kinase C substrate 80K-H | 2284.29 | 2482.21 | -0.12 | 5.77E-01 |
| NULL | GSMG0039233 | NULL | NULL | 125.72 | 186.85 | -0.57 | 3.82E-02 |
| ENSMUSG00000096981 | GSMG0039234 | Gm16845 | predicted gene, 16845 | 250.20 | 237.93 | 0.07 | 8.42E-01 |
| ENSMUSG00000010607 | GSMG0039237 | Pigyl | phosphatidylinositol glycan anchor biosynthesis, class Y-like | 777.26 | 797.92 | -0.04 | 9.17E-01 |
| ENSMUSG00000057982 | GSMG0039239 | Zfp809 | zinc finger protein 809 | 258.95 | 200.33 | 0.37 | 2.00E-01 |
| ENSMUSG00000036411 | GSMG0039241 | 9530077C05Rik | RIKEN cDNA 9530077C05 gene | 85.78 | 110.53 | -0.36 | 4.14E-01 |
| ENSMUSG00000035919 | GSMG0039243 | Bbs9 | Bardet-Biedl syndrome 9 (human) | 549.40 | 577.56 | -0.07 | 8.34E-01 |
| ENSMUSG00000001833 | GSMG0039257 | Sept7 | septin 7 | 7899.49 | 7250.55 | 0.12 | 6.72E-01 |
| ENSMUSG00000036611 | GSMG0039260 | Eepd1 | endonuclease/exonuclease/phosphatase family domain containing 1 | 378.16 | 330.97 | 0.19 | 5.74E-01 |
| ENSMUSG00000045994 | GSMG0039267 | B3gat1 | beta-1,3-glucuronyltransferase 1 (glucuronosyltransferase P) | 274.08 | 308.90 | -0.17 | 7.04E-01 |
| NULL | GSMG0039268 | NULL | NULL | 88.29 | 50.92 | 0.80 | 4.48E-02 |
| ENSMUSG00000035443 | GSMG0039269 | Thyn1 | thymocyte nuclear protein 1 | 307.70 | 420.56 | -0.45 | 4.35E-02 |
| ENSMUSG00000035024 | GSMG0039270 | Ncapd3 | non-SMC condensin II complex, subunit D3 | 148.48 | 199.83 | -0.43 | 1.42E-01 |
| NULL | GSMG0039273 | NULL | NULL | 477.61 | 383.64 | 0.32 | 1.63E-01 |
| ENSMUSG00000062257 | GSMG0039280 | Opcml | opioid binding protein/cell adhesion molecule-like | 289.06 | 315.51 | -0.12 | 7.97E-01 |
| NULL | GSMG0039281 | NULL | NULL | 183.96 | 187.89 | -0.02 | 9.63E-01 |
| ENSMUSG00000031993 | GSMG0039288 | Snx19 | sorting nexin 19 | 447.90 | 471.68 | -0.07 | 8.30E-01 |
| ENSMUSG00000031994 | GSMG0039289 | Adamts8 | a disintegrin-like and metallopeptidase (reprolysin type) with thrombos | 183.62 | 227.81 | -0.31 | 3.75E-01 |
| ENSMUSG00000047412 | GSMG0039290 | Zbtb44 | zinc finger and BTB domain containing 44 | 206.26 | 188.54 | 0.13 | 7.25E-01 |
| ENSMUSG00000042496 | GSMG0039295 | Prdm10 | PR domain containing 10 | 87.27 | 95.92 | -0.14 | 8.06E-01 |
| NULL | GSMG0039297 | NULL | NULL | 191.60 | 293.74 | -0.62 | 1.26E-02 |
| ENSMUSG00000042185 | GSMG0039298 | Nfrkb | nuclear factor related to kappa B binding protein | 619.84 | 803.27 | -0.37 | 1.44E-01 |
| ENSMUSG00000041444 | GSMG0039301 | Arhgap32 | Rho GTPase activating protein 32 | 1057.75 | 537.45 | 0.98 | 2.03E-06 |
| ENSMUSG00000032035 | GSMG0039304 | Ets1 | E26 avian leukemia oncogene 1, 5' domain | 5559.55 | 6568.99 | -0.24 | 1.94E-01 |
| ENSMUSG00000032036 | GSMG0039310 | Kirrel3 | kin of IRRE like 3 (Drosophila) | 442.82 | 545.55 | -0.30 | 3.50E-01 |

| | | | | | | | |
|---|---|---|---|---|---|---|---|
| NULL | GSMG0039313 | NULL | NULL | 170.69 | 117.11 | 0.54 | 8.18E-02 |
| NULL | GSMG0039314 | NULL | NULL | 627.70 | 623.41 | 0.01 | 9.83E-01 |
| NULL | GSMG0039315 | NULL | NULL | 682.51 | 671.26 | 0.02 | 9.41E-01 |
| NULL | GSMG0039316 | NULL | NULL | 164.68 | 148.74 | 0.15 | 6.94E-01 |
| NULL | GSMG0039318 | NULL | NULL | 463.77 | 416.13 | 0.16 | 5.75E-01 |
| NULL | GSMG0039319 | NULL | NULL | 293.26 | 305.05 | -0.05 | 8.90E-01 |
| ENSMUSG00000032042 | GSMG0039325 | Srpr | signal recognition particle receptor ('docking protein') | 3665.18 | 3638.99 | 0.01 | 9.70E-01 |
| ENSMUSG00000032044 | GSMG0039327 | Rpusd4 | RNA pseudouridylate synthase domain containing 4 | 184.75 | 188.61 | -0.02 | 9.59E-01 |
| NULL | GSMG0039330 | NULL | NULL | 299.07 | 324.39 | -0.12 | 7.37E-01 |
| ENSMUSG00000032103 | GSMG0039332 | Pus3 | pseudouridine synthase 3 | 291.24 | 352.66 | -0.27 | 2.95E-01 |
| ENSMUSG00000032118 | GSMG0039340 | Fez1 | fasciculation and elongation protein zeta 1 (zygin I) | 1426.42 | 2457.63 | -0.78 | 2.10E-07 |
| NULL | GSMG0039341 | NULL | NULL | 49.82 | 59.04 | -0.25 | 6.87E-01 |
| ENSMUSG00000032121 | GSMG0039343 | Tmem218 | transmembrane protein 218 | 693.46 | 560.22 | 0.31 | 1.65E-01 |
| ENSMUSG00000046240 | GSMG0039347 | Hepacam | hepatocyte cell adhesion molecule | 401.16 | 290.65 | 0.47 | 1.31E-01 |
| ENSMUSG00000032125 | GSMG0039349 | Robo4 | roundabout guidance receptor 4 | 6437.25 | 5693.63 | 0.18 | 4.07E-01 |
| ENSMUSG00000042138 | GSMG0039350 | Msantd2 | Myb/SANT-like DNA-binding domain containing 2 | 775.10 | 969.43 | -0.32 | 1.10E-01 |
| NULL | GSMG0039352 | NULL | NULL | 159.77 | 166.07 | -0.06 | 9.02E-01 |
| ENSMUSG00000001946 | GSMG0039353 | Esam | endothelial cell-specific adhesion molecule | 14967.15 | 11339.45 | 0.40 | 1.28E-02 |
| ENSMUSG00000001943 | GSMG0039354 | Vsig2 | V-set and immunoglobulin domain containing 2 | 404.53 | 387.92 | 0.06 | 8.67E-01 |
| ENSMUSG00000001942 | GSMG0039355 | Siae | sialic acid acetylesterase | 197.96 | 164.51 | 0.27 | 4.18E-01 |
| ENSMUSG00000023186 | GSMG0039371 | Vwa5a | von Willebrand factor A domain containing 5A | 1148.06 | 918.51 | 0.32 | 1.01E-01 |
| ENSMUSG00000025602 | GSMG0039396 | Zfp202 | zinc finger protein 202 | 222.84 | 397.85 | -0.83 | 8.91E-04 |
| ENSMUSG00000049281 | GSMG0039397 | Scn3b | sodium channel, voltage-gated, type III, beta | 1389.94 | 2372.24 | -0.77 | 7.74E-06 |
| ENSMUSG00000032024 | GSMG0039400 | Clmp | CXADR-like membrane protein | 182.91 | 537.64 | -1.55 | 1.45E-11 |
| NULL | GSMG0039408 | NULL | NULL | 157.89 | 204.80 | -0.38 | 2.59E-01 |
| NULL | GSMG0039413 | NULL | NULL | 192.44 | 347.83 | -0.86 | 1.77E-03 |
| ENSMUSG00000032012 | GSMG0039421 | Nectin1 | nectin cell adhesion molecule 1 | 58.33 | 93.06 | -0.67 | 1.16E-01 |
| ENSMUSG00000032011 | GSMG0039424 | Thy1 | thymus cell antigen 1, theta | 868.61 | 363.57 | 1.25 | 2.10E-11 |
| ENSMUSG00000032010 | GSMG0039425 | Usp2 | ubiquitin specific peptidase 2 | 617.48 | 275.95 | 1.16 | 4.68E-10 |
| G00000034739,ENSMUSG000 | GSMG0039426 | Mfrp // C1qtnf5 | membrane frizzled-related protein // C1q and tumor necrosis factor relat | 766.24 | 518.39 | 0.56 | 1.47E-03 |
| ENSMUSG00000032135 | GSMG0039428 | Mcam | melanoma cell adhesion molecule | 5256.12 | 5483.22 | -0.06 | 8.24E-01 |
| NULL | GSMG0039429 | NULL | NULL | 311.88 | 373.28 | -0.26 | 3.39E-01 |
| ENSMUSG00000032123 | GSMG0039432 | Dpagt1 | dolichyl-phosphate (UDP-N-acetylglucosamine) acetylglucosaminephos | 1076.48 | 1101.53 | -0.03 | 9.21E-01 |

| | | | | | | | |
|---|---|---|---|---|---|---|---|
| ENSMUSG00000049932 | GSMG0039433 | H2afx | H2A histone family, member X | 1257.33 | 2269.56 | -0.85 | 8.01E-06 |
| NULL | GSMG0039434 | NULL | NULL | 425.89 | 530.47 | -0.32 | 1.52E-01 |
| ENSMUSG00000032115 | GSMG0039435 | Hyou1 | hypoxia up-regulated 1 | 1175.46 | 1722.03 | -0.55 | 3.02E-02 |
| ENSMUSG00000032114 | GSMG0039436 | Slc37a4 | solute carrier family 37 (glucose-6-phosphate transporter), member 4 | 599.90 | 607.35 | -0.02 | 9.62E-01 |
| NULL | GSMG0039438 | NULL | NULL | 87.10 | 44.32 | 0.97 | 8.29E-03 |
| ENSMUSG00000032097 | GSMG0039441 | Ddx6 | DEAD (Asp-Glu-Ala-Asp) box polypeptide 6 | 753.47 | 882.06 | -0.23 | 3.61E-01 |
| ENSMUSG00000002031 | GSMG0039447 | Ift46 | intraflagellar transport 46 | 531.23 | 592.10 | -0.16 | 5.58E-01 |
| NULL | GSMG0039448 | NULL | NULL | 107.33 | 108.54 | -0.01 | 9.86E-01 |
| ENSMUSG00000070304 | GSMG0039454 | Scn2b | sodium channel, voltage-gated, type II, beta | 464.82 | 162.90 | 1.51 | 2.21E-09 |
| ENSMUSG00000066705 | GSMG0039458 | Fxyd6 | FXYD domain-containing ion transport regulator 6 | 1923.02 | 4299.11 | -1.16 | 3.01E-13 |
| ENSMUSG00000032087 | GSMG0039460 | Dscaml1 | Down syndrome cell adhesion molecule like 1 | 245.03 | 326.18 | -0.41 | 2.09E-01 |
| ENSMUSG00000032086 | GSMG0039462 | Bace1 | beta-site APP cleaving enzyme 1 | 265.40 | 227.18 | 0.22 | 5.14E-01 |
| ENSMUSG00000035382 | GSMG0039464 | Pcsk7 | proprotein convertase subtilisin/kexin type 7 | 1567.92 | 1345.82 | 0.22 | 4.09E-01 |
| NULL | GSMG0039465 | NULL | NULL | 785.03 | 181.70 | 2.11 | 1.56E-25 |
| ENSMUSG00000034135 | GSMG0039467 | Sik3 | SIK family kinase 3 | 734.52 | 580.49 | 0.34 | 1.80E-01 |
| ENSMUSG00000032078 | GSMG0039478 | Zpr1 | ZPR1 zinc finger | 1327.22 | 1480.20 | -0.16 | 4.52E-01 |
| ENSMUSG00000032077 | GSMG0039479 | Bud13 | BUD13 homolog | 320.72 | 364.90 | -0.18 | 5.11E-01 |
| ENSMUSG00000032076 | GSMG0039483 | Cadm1 | cell adhesion molecule 1 | 1438.29 | 2095.06 | -0.54 | 1.78E-03 |
| ENSMUSG00000042293 | GSMG0039493 | Gm5617 | predicted gene 5617 | 205.17 | 272.69 | -0.41 | 1.45E-01 |
| ENSMUSG00000032267 | GSMG0039495 | Usp28 | ubiquitin specific peptidase 28 | 1431.99 | 1070.56 | 0.42 | 1.32E-02 |
| ENSMUSG00000032264 | GSMG0039496 | Zw10 | zw10 kinetochore protein | 984.23 | 1133.92 | -0.20 | 3.01E-01 |
| ENSMUSG00000039217 | GSMG0039506 | Il18 | interleukin 18 | 385.33 | 270.17 | 0.51 | 3.66E-02 |
| ENSMUSG00000039016 | GSMG0039508 | Timm8b | translocase of inner mitochondrial membrane 8B | 970.71 | 1060.32 | -0.13 | 6.82E-01 |
| ENSMUSG00000032060 | GSMG0039512 | Cryab | crystallin, alpha B | 1385.48 | 1133.39 | 0.29 | 2.05E-01 |
| ENSMUSG00000037845 | GSMG0039513 | Fdxacb1 | ferredoxin-fold anticodon binding domain containing 1 | 118.42 | 115.88 | 0.03 | 9.49E-01 |
| ENSMUSG00000032059 | GSMG0039514 | Alg9 | asparagine-linked glycosylation 9 (alpha 1,2 mannosyltransferase) | 442.81 | 628.39 | -0.50 | 1.23E-02 |
| ENSMUSG00000032058 | GSMG0039517 | Ppp2r1b | protein phosphatase 2, regulatory subunit A, beta | 1075.04 | 1260.12 | -0.23 | 3.18E-01 |
| ENSMUSG00000053199 | GSMG0039522 | Arhgap20 | Rho GTPase activating protein 20 | 260.72 | 318.17 | -0.29 | 3.23E-01 |
| ENSMUSG00000032050 | GSMG0039525 | Rdx | radixin | 1713.97 | 1817.43 | -0.08 | 7.82E-01 |
| ENSMUSG00000034487 | GSMG0039533 | Kdelc2 | KDEL (Lys-Asp-Glu-Leu) containing 2 | 1549.59 | 1380.58 | 0.17 | 5.46E-01 |
| ENSMUSG00000033054 | GSMG0039535 | Npat | nuclear protein in the AT region | 904.31 | 1042.42 | -0.21 | 3.23E-01 |
| ENSMUSG00000042195 | GSMG0039538 | Slc35f2 | solute carrier family 35, member F2 | 1653.01 | 1819.09 | -0.14 | 5.12E-01 |
| ENSMUSG00000032279 | GSMG0039546 | Idh3a | isocitrate dehydrogenase 3 (NAD+) alpha | 1531.17 | 1866.85 | -0.29 | 1.47E-01 |

| | | | | | | | |
|---|---|---|---|---|---|---|---|
| ENSMUSG00000032285 | GSMG0039547 | Dnaja4 | DnaJ heat shock protein family (Hsp40) member A4 | 404.68 | 298.81 | 0.44 | 4.40E-02 |
| ENSMUSG00000032293 | GSMG0039549 | Ireb2 | iron responsive element binding protein 2 | 1062.86 | 1116.55 | -0.07 | 7.90E-01 |
| ENSMUSG00000035878 | GSMG0039551 | Hykk | hydroxylysine kinase 1 | 61.04 | 63.34 | -0.06 | 9.28E-01 |
| ENSMUSG00000032301 | GSMG0039552 | Psma4 | proteasome (prosome, macropain) subunit, alpha type 4 | 1300.24 | 1523.54 | -0.23 | 3.01E-01 |
| ENSMUSG00000032309 | GSMG0039556 | Fbxo22 | F-box protein 22 | 1327.15 | 1353.66 | -0.03 | 9.28E-01 |
| ENSMUSG00000032320 | GSMG0039560 | Rcn2 | reticulocalbin 2 | 4204.76 | 3963.60 | 0.09 | 7.46E-01 |
| ENSMUSG00000032329 | GSMG0039564 | Hmg20a | high mobility group 20A | 636.60 | 711.86 | -0.16 | 4.73E-01 |
| ENSMUSG00000032911 | GSMG0039568 | Cspg4 | chondroitin sulfate proteoglycan 4 | 3383.00 | 3692.32 | -0.13 | 7.44E-01 |
| ENSMUSG00000032288 | GSMG0039571 | Imp3 | IMP3, U3 small nucleolar ribonucleoprotein | 908.06 | 913.57 | -0.01 | 9.79E-01 |
| ENSMUSG00000055334 | GSMG0039572 | Snupn | snurportin 1 | 260.51 | 286.22 | -0.13 | 7.07E-01 |
| ENSMUSG00000032290 | GSMG0039574 | Ptpn9 | protein tyrosine phosphatase, non-receptor type 9 | 1105.24 | 1096.17 | 0.01 | 9.69E-01 |
| ENSMUSG00000042557 | GSMG0039576 | Sin3a | transcriptional regulator, SIN3A (yeast) | 787.45 | 857.00 | -0.12 | 6.15E-01 |
| ENSMUSG00000032295 | GSMG0039577 | Man2c1 | mannosidase, alpha, class 2C, member 1 | 2387.61 | 2720.31 | -0.19 | 4.38E-01 |
| ENSMUSG00000062309 | GSMG0039582 | Rpp25 | ribonuclease P/MRP 25 subunit | 54.15 | 138.18 | -1.35 | 2.24E-06 |
| ENSMUSG00000000088 | GSMG0039583 | Cox5a | cytochrome c oxidase subunit Va | 3120.79 | 3400.48 | -0.12 | 6.63E-01 |
| ENSMUSG00000032305 | GSMG0039585 | Fam219b | family with sequence similarity 219, member B | 1265.61 | 1310.68 | -0.05 | 8.80E-01 |
| NULL | GSMG0039586 | NULL | NULL | 202.39 | 186.61 | 0.11 | 7.74E-01 |
| NULL | GSMG0039587 | NULL | NULL | 584.34 | 610.18 | -0.06 | 8.62E-01 |
| ENSMUSG00000040188 | GSMG0039588 | Scamp2 | secretory carrier membrane protein 2 | 866.53 | 932.59 | -0.11 | 6.99E-01 |
| ENSMUSG00000032308 | GSMG0039590 | Ulk3 | unc-51-like kinase 3 | 515.66 | 465.43 | 0.15 | 6.52E-01 |
| ENSMUSG00000038957 | GSMG0039592 | Edc3 | enhancer of mRNA decapping 3 | 478.65 | 583.43 | -0.29 | 1.81E-01 |
| ENSMUSG00000055720 | GSMG0039596 | Ubl7 | ubiquitin-like 7 (bone marrow stromal cell-derived) | 828.60 | 735.94 | 0.17 | 5.12E-01 |
| ENSMUSG00000038264 | GSMG0039597 | Sema7a | sema domain, immunoglobulin domain (Ig), and GPI membrane anchor | 525.36 | 537.86 | -0.03 | 9.40E-01 |
| ENSMUSG00000032327 | GSMG0039601 | Stra6 | stimulated by retinoic acid gene 6 | 2025.93 | 3389.19 | -0.74 | 6.34E-06 |
| ENSMUSG00000032333 | GSMG0039605 | Stoml1 | stomatin-like 1 | 773.16 | 1050.57 | -0.44 | 1.63E-02 |
| ENSMUSG00000032336 | GSMG0039611 | Nptn | neuroplastin | 2828.64 | 2559.64 | 0.14 | 4.88E-01 |
| NULL | GSMG0039613 | NULL | NULL | 1015.08 | 356.58 | 1.51 | 5.90E-18 |
| NULL | GSMG0039614 | NULL | NULL | 631.93 | 194.90 | 1.70 | 2.79E-15 |
| ENSMUSG00000025236 | GSMG0039618 | Adpgk | ADP-dependent glucokinase | 280.85 | 362.39 | -0.36 | 1.64E-01 |
| ENSMUSG00000025232 | GSMG0039620 | Hexa | hexosaminidase A | 1656.04 | 1457.90 | 0.18 | 4.01E-01 |
| ENSMUSG00000032297 | GSMG0039621 | Celf6 | CUGBP, Elav-like family member 6 | 197.62 | 457.46 | -1.21 | 5.59E-10 |
| ENSMUSG00000025237 | GSMG0039623 | Parp6 | poly (ADP-ribose) polymerase family, member 6 | 1114.26 | 1349.53 | -0.28 | 2.24E-01 |
| ENSMUSG00000032294 | GSMG0039624 | Pkm | pyruvate kinase, muscle | 21797.25 | 23787.99 | -0.13 | 5.35E-01 |

| | | | | | | | |
|---|---|---|---|---|---|---|---|
| ENSMUSG00000074259 | GSMG0039626 | Gramd2 | GRAM domain containing 2 | 40.15 | 102.85 | -1.37 | 2.29E-04 |
| ENSMUSG00000034839 | GSMG0039637 | Larp6 | La ribonucleoprotein domain family, member 6 | 275.68 | 291.52 | -0.08 | 8.26E-01 |
| ENSMUSG00000034485 | GSMG0039638 | Uaca | uveal autoantigen with coiled-coil domains and ankyrin repeats | 1978.51 | 2450.26 | -0.31 | 6.78E-02 |
| ENSMUSG00000032280 | GSMG0039641 | Tle3 | transducin-like enhancer of split 3 | 370.73 | 415.17 | -0.16 | 7.32E-01 |
| ENSMUSG00000032249 | GSMG0039645 | Anp32a | acidic (leucine-rich) nuclear phosphoprotein 32 family, member A | 438.12 | 548.55 | -0.32 | 2.05E-01 |
| ENSMUSG00000037257 | GSMG0039656 | Aagab | alpha- and gamma-adaptin binding protein | 2117.26 | 1866.99 | 0.18 | 3.80E-01 |
| ENSMUSG00000032399 | GSMG0039660 | Rpl4 | ribosomal protein L4 | 5142.09 | 7203.83 | -0.49 | 1.00E-02 |
| ENSMUSG00000032398 | GSMG0039661 | Snapc5 | small nuclear RNA activating complex, polypeptide 5 | 397.26 | 319.74 | 0.32 | 2.49E-01 |
| ENSMUSG00000035337 | GSMG0039663 | Uchl4 | ubiquitin carboxyl-terminal esterase L4 | 229.14 | 328.22 | -0.52 | 3.06E-02 |
| ENSMUSG00000032397 | GSMG0039664 | Tipin | timeless interacting protein | 530.51 | 833.30 | -0.65 | 1.89E-04 |
| ENSMUSG00000036466 | GSMG0039665 | Megf11 | multiple EGF-like-domains 11 | 80.06 | 25.34 | 1.66 | 1.19E-04 |
| ENSMUSG00000034263 | GSMG0039669 | Vwa9 | von Willebrand factor A domain containing 9 | 794.62 | 995.45 | -0.33 | 7.90E-02 |
| ENSMUSG00000032393 | GSMG0039670 | Dpp8 | dipeptidylpeptidase 8 | 2914.35 | 2961.58 | -0.02 | 9.42E-01 |
| NULL | GSMG0039672 | NULL | NULL | 604.95 | 523.13 | 0.21 | 4.13E-01 |
| ENSMUSG00000032816 | GSMG0039673 | Igdcc4 | immunoglobulin superfamily, DCC subclass, member 4 | 374.02 | 567.06 | -0.60 | 3.43E-02 |
| ENSMUSG00000032394 | GSMG0039674 | Igdcc3 | immunoglobulin superfamily, DCC subclass, member 3 | 273.53 | 310.83 | -0.19 | 5.74E-01 |
| ENSMUSG00000032392 | GSMG0039677 | Parp16 | poly (ADP-ribose) polymerase family, member 16 | 167.18 | 223.34 | -0.42 | 1.40E-01 |
| ENSMUSG00000015357 | GSMG0039679 | Clpx | caseinolytic mitochondrial matrix peptidase chaperone subunit | 1342.35 | 1560.74 | -0.22 | 3.12E-01 |
| ENSMUSG00000041837 | GSMG0039680 | Pdcd7 | programmed cell death 7 | 416.19 | 399.67 | 0.06 | 8.65E-01 |
| ENSMUSG00000041696 | GSMG0039682 | Rasl12 | RAS-like, family 12 | 1167.18 | 721.70 | 0.69 | 1.21E-05 |
| ENSMUSG00000059183 | GSMG0039684 | Mtfmt | mitochondrial methionyl-tRNA formyltransferase | 255.73 | 261.99 | -0.03 | 9.28E-01 |
| ENSMUSG00000032388 | GSMG0039685 | Spg21 | spastic paraplegia 21 homolog (human) | 634.83 | 742.61 | -0.23 | 3.37E-01 |
| ENSMUSG00000032384 | GSMG0039695 | Csnk1g1 | casein kinase 1, gamma 1 | 340.77 | 427.70 | -0.32 | 2.83E-01 |
| ENSMUSG00000032383 | GSMG0039700 | Ppib | peptidylprolyl isomerase B | 7067.49 | 7866.13 | -0.15 | 4.49E-01 |
| ENSMUSG00000032381 | GSMG0039701 | Fam96a | family with sequence similarity 96, member A | 1017.47 | 1386.68 | -0.45 | 4.16E-02 |
| ENSMUSG00000032380 | GSMG0039702 | Dapk2 | death-associated protein kinase 2 | 84.76 | 137.43 | -0.69 | 3.15E-02 |
| ENSMUSG00000036781 | GSMG0039712 | Rps27l | ribosomal protein S27-like | 1411.07 | 1621.04 | -0.20 | 4.83E-01 |
| ENSMUSG00000032238 | GSMG0039719 | Rora | RAR-related orphan receptor alpha | 658.39 | 455.41 | 0.53 | 1.71E-02 |
| NULL | GSMG0039724 | NULL | NULL | 234.50 | 191.34 | 0.30 | 3.92E-01 |
| NULL | GSMG0039726 | NULL | NULL | 264.72 | 235.62 | 0.17 | 5.79E-01 |
| NULL | GSMG0039727 | NULL | NULL | 541.40 | 416.39 | 0.38 | 9.53E-02 |
| NULL | GSMG0039732 | NULL | NULL | 503.80 | 437.52 | 0.21 | 5.20E-01 |
| NULL | GSMG0039733 | NULL | NULL | 521.18 | 414.73 | 0.33 | 1.80E-01 |

| | | | | | | | |
|---|---|---|---|---|---|---|---|
| ENSMUSG00000032235 | GSMG0039745 | Ice2 | interactor of little elongation complex ELL subunit 2 | 496.53 | 536.36 | -0.11 | 7.34E-01 |
| ENSMUSG00000032231 | GSMG0039746 | Anxa2 | annexin A2 | 5934.83 | 5304.84 | 0.16 | 5.07E-01 |
| ENSMUSG00000011958 | GSMG0039749 | Bnip2 | BCL2/adenovirus E1B interacting protein 2 | 3843.19 | 3161.29 | 0.28 | 1.48E-01 |
| ENSMUSG00000033543 | GSMG0039750 | Gtf2a2 | general transcription factor II A, 2 | 1100.78 | 1292.75 | -0.23 | 2.74E-01 |
| ENSMUSG00000032220 | GSMG0039754 | Myo1e | myosin IE | 992.48 | 904.79 | 0.13 | 5.64E-01 |
| ENSMUSG00000032212 | GSMG0039761 | Sltm | SAFB-like, transcription modulator | 2091.29 | 2314.23 | -0.15 | 5.57E-01 |
| NULL | GSMG0039762 | NULL | NULL | 350.11 | 333.03 | 0.07 | 8.67E-01 |
| NULL | GSMG0039763 | NULL | NULL | 170.24 | 155.19 | 0.14 | 7.31E-01 |
| NULL | GSMG0039764 | NULL | NULL | 118.68 | 153.18 | -0.36 | 2.98E-01 |
| NULL | GSMG0039765 | NULL | NULL | 498.44 | 359.54 | 0.47 | 4.34E-02 |
| ENSMUSG00000054693 | GSMG0039766 | Adam10 | a disintegrin and metallopeptidase domain 10 | 4493.31 | 4901.80 | -0.13 | 5.77E-01 |
| ENSMUSG00000013584 | GSMG0039772 | Aldh1a2 | aldehyde dehydrogenase family 1, subfamily A2 | 491.25 | 374.41 | 0.40 | 1.80E-01 |
| ENSMUSG00000038535 | GSMG0039773 | Zfp280d | zinc finger protein 280D | 872.77 | 999.52 | -0.20 | 3.93E-01 |
| ENSMUSG00000037674 | GSMG0039782 | Rfx7 | regulatory factor X, 7 | 114.50 | 130.69 | -0.19 | 6.66E-01 |
| ENSMUSG00000032216 | GSMG0039785 | Nedd4 | neural precursor cell expressed, developmentally down-regulated 4 | 11250.28 | 13644.66 | -0.28 | 9.91E-02 |
| ENSMUSG00000034563 | GSMG0039794 | Ccpg1 | cell cycle progression 1 | 2740.12 | 1816.08 | 0.59 | 1.08E-04 |
| ENSMUSG00000098332 | GSMG0039796 | 2310009A05Rik | RIKEN cDNA 2310009A05 gene | 255.17 | 262.58 | -0.05 | 9.34E-01 |
| ENSMUSG00000032215 | GSMG0039801 | Rsl24d1 | ribosomal L24 domain containing 1 | 995.16 | 1266.60 | -0.35 | 1.13E-01 |
| ENSMUSG00000034858 | GSMG0039806 | Fam214a | family with sequence similarity 214, member A | 1243.76 | 933.56 | 0.41 | 2.93E-02 |
| ENSMUSG00000034593 | GSMG0039809 | Myo5a | myosin VA | 1339.07 | 1410.48 | -0.07 | 8.08E-01 |
| ENSMUSG00000032192 | GSMG0039815 | Gnb5 | guanine nucleotide binding protein (G protein), beta 5 | 667.50 | 596.81 | 0.16 | 5.50E-01 |
| ENSMUSG00000042487 | GSMG0039818 | Leo1 | Leo1, Paf1/RNA polymerase II complex component | 1011.97 | 1196.12 | -0.24 | 2.17E-01 |
| ENSMUSG00000032184 | GSMG0039819 | Lysmd2 | LysM, putative peptidoglycan-binding, domain containing 2 | 895.83 | 605.87 | 0.56 | 3.29E-03 |
| ENSMUSG00000032179 | GSMG0039820 | Bmp5 | bone morphogenetic protein 5 | 125.62 | 131.11 | -0.06 | 8.94E-01 |
| ENSMUSG00000007908 | GSMG0039822 | Hmgcll1 | 3-hydroxymethyl-3-methylglutaryl-Coenzyme A lyase-like 1 | 49.68 | 75.51 | -0.61 | 1.75E-01 |
| ENSMUSG00000032350 | GSMG0039830 | Gclc | glutamate-cysteine ligase, catalytic subunit | 1063.43 | 916.68 | 0.21 | 4.62E-01 |
| ENSMUSG00000032349 | GSMG0039833 | Elovl5 | ELOVL family member 5, elongation of long chain fatty acids (yeast) | 1795.56 | 1616.38 | 0.15 | 5.10E-01 |
| ENSMUSG00000009828 | GSMG0039836 | Ick | intestinal cell kinase | 466.20 | 464.08 | 0.01 | 9.85E-01 |
| ENSMUSG00000032348 | GSMG0039838 | Gsta4 | glutathione S-transferase, alpha 4 | 1234.01 | 731.50 | 0.76 | 2.14E-05 |
| ENSMUSG00000032342 | GSMG0039843 | Mto1 | mitochondrial tRNA translation optimization 1 | 163.17 | 157.17 | 0.06 | 8.99E-01 |
| ENSMUSG00000046186 | GSMG0039844 | Cd109 | CD109 antigen | 101.50 | 382.63 | -1.92 | 1.40E-16 |
| ENSMUSG00000034252 | GSMG0039850 | Senp6 | SUMO/sentrin specific peptidase 6 | 531.22 | 560.85 | -0.08 | 7.96E-01 |
| ENSMUSG00000033577 | GSMG0039853 | Myo6 | myosin VI | 3098.79 | 3322.12 | -0.10 | 6.62E-01 |

| | | | | | | | |
|---|---|---|---|---|---|---|---|
| ENSMUSG00000032251 | GSMG0039860 | Irak1bp1 | interleukin-1 receptor-associated kinase 1 binding protein 1 | 209.53 | 245.48 | -0.23 | 5.11E-01 |
| ENSMUSG00000038379 | GSMG0039871 | Ttk | Ttk protein kinase | 186.58 | 428.12 | -1.19 | 4.61E-05 |
| ENSMUSG00000032263 | GSMG0039873 | Bckdhb | branched chain ketoacid dehydrogenase E1, beta polypeptide | 345.37 | 440.27 | -0.35 | 1.43E-01 |
| ENSMUSG00000035274 | GSMG0039877 | Tpbg | trophoblast glycoprotein | 805.60 | 1188.93 | -0.56 | 3.12E-03 |
| ENSMUSG00000032417 | GSMG0039880 | Rwdd2a | RWD domain containing 2A | 181.90 | 190.01 | -0.07 | 8.86E-01 |
| ENSMUSG00000032425 | GSMG0039891 | Zfp949 | zinc finger protein 949 | 494.53 | 415.10 | 0.25 | 3.11E-01 |
| ENSMUSG00000032356 | GSMG0039904 | Rasgrf1 | RAS protein-specific guanine nucleotide-releasing factor 1 | 1179.09 | 174.80 | 2.76 | 2.95E-46 |
| ENSMUSG00000032359 | GSMG0039905 | Ctsh | cathepsin H | 9736.89 | 5521.74 | 0.82 | 3.72E-08 |
| ENSMUSG00000032369 | GSMG0039911 | Plscr1 | phospholipid scramblase 1 | 130.83 | 119.39 | 0.13 | 7.99E-01 |
| ENSMUSG00000032372 | GSMG0039912 | Plscr2 | phospholipid scramblase 2 | 100.55 | 44.45 | 1.18 | 1.04E-03 |
| ENSMUSG00000032377 | GSMG0039915 | Plscr4 | phospholipid scramblase 4 | 261.86 | 56.87 | 2.21 | 6.15E-18 |
| ENSMUSG00000032374 | GSMG0039916 | Plod2 | procollagen lysine, 2-oxoglutarate 5-dioxygenase 2 | 173.57 | 225.18 | -0.37 | 2.13E-01 |
| ENSMUSG00000031129 | GSMG0039920 | Slc9a9 | solute carrier family 9 (sodium/hydrogen exchanger), member 9 | 360.16 | 272.56 | 0.41 | 1.99E-01 |
| ENSMUSG00000032410 | GSMG0039938 | Xrn1 | 5'-3' exoribonuclease 1 | 349.12 | 363.92 | -0.06 | 8.79E-01 |
| ENSMUSG00000041440 | GSMG0039942 | Gk5 | glycerol kinase 5 (putative) | 140.92 | 127.42 | 0.14 | 7.31E-01 |
| ENSMUSG00000032411 | GSMG0039944 | Tfdp2 | transcription factor Dp 2 | 493.58 | 468.89 | 0.07 | 8.47E-01 |
| ENSMUSG00000046402 | GSMG0039956 | Rbp1 | retinol binding protein 1, cellular | 1736.11 | 2164.72 | -0.32 | 1.76E-01 |
| ENSMUSG00000032458 | GSMG0039959 | Copb2 | coatomer protein complex, subunit beta 2 (beta prime) | 4394.30 | 5160.22 | -0.23 | 3.03E-01 |
| ENSMUSG00000032463 | GSMG0039964 | Faim | Fas apoptotic inhibitory molecule | 1607.39 | 1554.01 | 0.05 | 8.72E-01 |
| ENSMUSG00000056267 | GSMG0039966 | Cep70 | centrosomal protein 70 | 287.34 | 321.61 | -0.16 | 5.90E-01 |
| ENSMUSG00000032469 | GSMG0039971 | Dbr1 | debranching RNA lariats 1 | 130.10 | 152.64 | -0.24 | 5.28E-01 |
| ENSMUSG00000037784 | GSMG0039973 | Dzip1l | DAZ interacting protein 1-like | 221.77 | 180.51 | 0.29 | 3.33E-01 |
| NULL | GSMG0039976 | NULL | NULL | 353.43 | 291.85 | 0.27 | 3.16E-01 |
| ENSMUSG00000037286 | GSMG0039978 | Stag1 | stromal antigen 1 | 445.68 | 473.80 | -0.09 | 7.75E-01 |
| ENSMUSG00000032531 | GSMG0040002 | Amotl2 | angiomotin-like 2 | 1716.80 | 1400.22 | 0.29 | 2.28E-01 |
| ENSMUSG00000032547 | GSMG0040004 | Ryk | receptor-like tyrosine kinase | 986.83 | 661.75 | 0.57 | 2.85E-02 |
| ENSMUSG00000032548 | GSMG0040006 | Slco2a1 | solute carrier organic anion transporter family, member 2a1 | 757.99 | 1396.59 | -0.88 | 2.01E-06 |
| ENSMUSG00000032555 | GSMG0040010 | Topbp1 | topoisomerase (DNA) II binding protein 1 | 726.42 | 1201.09 | -0.73 | 5.48E-06 |
| G00000032558,ENSMUSG000 | GSMG0040019 | Nphp3 // Acad11 | nephronophthisis 3 (adolescent) // acyl-Coenzyme A dehydrogenase far | 2074.90 | 1622.41 | 0.36 | 3.83E-02 |
| ENSMUSG00000032564 | GSMG0040023 | Cpne4 | copine IV | 41.83 | 66.86 | -0.69 | 1.80E-01 |
| ENSMUSG00000032563 | GSMG0040025 | Mrpl3 | mitochondrial ribosomal protein L3 | 612.35 | 724.31 | -0.24 | 3.57E-01 |
| ENSMUSG00000032567 | GSMG0040029 | Aste1 | asteroid homolog 1 (Drosophila) | 387.24 | 365.96 | 0.08 | 7.97E-01 |
| ENSMUSG00000032571 | GSMG0040032 | Pik3r4 | phosphatidylinositol 3 kinase, regulatory subunit, polypeptide 4, p150 | 1377.85 | 1359.79 | 0.02 | 9.50E-01 |

| | | | | | | | |
|---|---|---|---|---|---|---|---|
| ENSMUSG00000020257 | GSMG0040034 | Wdr82 | WD repeat domain containing 82 | 1335.95 | 1309.26 | 0.03 | 9.21E-01 |
| ENSMUSG00000023277 | GSMG0040035 | Twf2 | twinfilin actin binding protein 2 | 1663.31 | 1555.75 | 0.10 | 7.14E-01 |
| ENSMUSG00000023345 | GSMG0040038 | Poc1a | POC1 centriolar protein A | 124.42 | 173.94 | -0.48 | 1.23E-01 |
| ENSMUSG00000053716 | GSMG0040039 | Dusp7 | dual specificity phosphatase 7 | 2251.20 | 1489.94 | 0.59 | 3.03E-02 |
| ENSMUSG00000042073 | GSMG0040041 | Abhd14b | abhydrolase domain containing 14b | 197.59 | 206.50 | -0.07 | 9.08E-01 |
| ENSMUSG00000023495 | GSMG0040042 | Pcbp4 | poly(rC) binding protein 4 | 1712.09 | 3111.65 | -0.86 | 3.34E-07 |
| ENSMUSG00000041506 | GSMG0040043 | Rrp9 | RRP9, small subunit (SSU) processome component, homolog (yeast) | 886.30 | 1069.95 | -0.27 | 1.45E-01 |
| ENSMUSG00000040325 | GSMG0040046 | Vprbp | Vpr (HIV-1) binding protein | 304.41 | 355.17 | -0.22 | 4.90E-01 |
| NULL | GSMG0040047 | NULL | NULL | 233.00 | 279.48 | -0.27 | 4.71E-01 |
| ENSMUSG00000010045 | GSMG0040054 | Tmem115 | transmembrane protein 115 | 1367.22 | 1363.47 | 0.00 | 9.88E-01 |
| ENSMUSG00000010057 | GSMG0040055 | Nprl2 | nitrogen permease regulator-like 2 | 1306.04 | 1221.19 | 0.10 | 7.13E-01 |
| ENSMUSG00000010067 | GSMG0040057 | Rassf1 | Ras association (RalGDS/AF-6) domain family member 1 | 1101.43 | 795.29 | 0.47 | 7.02E-03 |
| ENSMUSG00000010054 | GSMG0040058 | Tusc2 | tumor suppressor candidate 2 | 1817.84 | 1806.76 | 0.01 | 9.77E-01 |
| ENSMUSG00000010047 | GSMG0040059 | Hyal2 | hyaluronoglucosaminidase 2 | 1609.15 | 1626.55 | -0.02 | 9.61E-01 |
| ENSMUSG00000010048 | GSMG0040061 | Ifrd2 | interferon-related developmental regulator 2 | 279.07 | 416.63 | -0.58 | 8.66E-03 |
| ENSMUSG00000032583 | GSMG0040065 | Mon1a | MON1 homolog A, secretory traffciking associated | 445.94 | 328.20 | 0.44 | 3.95E-02 |
| ENSMUSG00000032936 | GSMG0040069 | Camkv | CaM kinase-like vesicle-associated | 3926.99 | 6005.32 | -0.61 | 6.95E-04 |
| ENSMUSG00000032586 | GSMG0040070 | Traip | TRAF-interacting protein | 288.24 | 399.73 | -0.47 | 3.65E-02 |
| ENSMUSG00000032596 | GSMG0040071 | Uba7 | ubiquitin-like modifier activating enzyme 7 | 699.15 | 467.09 | 0.58 | 1.27E-03 |
| ENSMUSG00000032594 | GSMG0040072 | Ip6k1 | inositol hexaphosphate kinase 1 | 1371.59 | 1035.18 | 0.41 | 7.09E-02 |
| ENSMUSG00000070284 | GSMG0040074 | Gmppb | GDP-mannose pyrophosphorylase B | 370.06 | 307.60 | 0.27 | 3.72E-01 |
| NULL | GSMG0040076 | NULL | NULL | 58.39 | 59.43 | -0.02 | 9.77E-01 |
| NULL | GSMG0040079 | NULL | NULL | 173.77 | 128.14 | 0.43 | 3.07E-01 |
| ENSMUSG00000032606 | GSMG0040080 | Nicn1 | nicolin 1 | 1541.02 | 1843.07 | -0.26 | 1.75E-01 |
| ENSMUSG00000007815 | GSMG0040082 | Rhoa | ras homolog family member A | 13239.55 | 12976.83 | 0.03 | 9.28E-01 |
| ENSMUSG00000063856 | GSMG0040083 | Gpx1 | glutathione peroxidase 1 | 5049.20 | 5128.74 | -0.02 | 9.42E-01 |
| ENSMUSG00000032612 | GSMG0040084 | Usp4 | ubiquitin specific peptidase 4 (proto-oncogene) | 4523.84 | 5066.96 | -0.16 | 3.91E-01 |
| ENSMUSG00000049305 | GSMG0040087 | Ccdc71 | coiled-coil domain containing 71 | 990.40 | 1107.14 | -0.16 | 5.23E-01 |
| ENSMUSG00000052911 | GSMG0040089 | Lamb2 | laminin, beta 2 | 19759.95 | 9027.94 | 1.13 | 6.73E-09 |
| ENSMUSG00000006676 | GSMG0040090 | Usp19 | ubiquitin specific peptidase 19 | 4497.66 | 3733.63 | 0.27 | 2.43E-01 |
| ENSMUSG00000006673 | GSMG0040092 | Qrich1 | glutamine-rich 1 | 2234.05 | 2540.44 | -0.19 | 3.66E-01 |
| NULL | GSMG0040093 | NULL | NULL | 298.30 | 201.95 | 0.56 | 4.62E-02 |
| ENSMUSG00000019039 | GSMG0040098 | Dalrd3 | DALR anticodon binding domain containing 3 | 2335.93 | 2124.66 | 0.14 | 6.07E-01 |

| | | | | | | | |
|---|---|---|---|---|---|---|---|
| ENSMUSG00000032602 | GSMG0040100 | Slc25a20 | solute carrier family 25 (mitochondrial carnitine/acylcarnitine transloca | 962.95 | 1174.05 | -0.28 | 1.77E-01 |
| ENSMUSG00000032601 | GSMG0040101 | Prkar2a | protein kinase, cAMP dependent regulatory, type II alpha | 463.53 | 519.65 | -0.16 | 5.47E-01 |
| ENSMUSG00000032599 | GSMG0040104 | Ip6k2 | inositol hexaphosphate kinase 2 | 326.76 | 622.09 | -0.93 | 2.22E-07 |
| ENSMUSG00000032598 | GSMG0040105 | Nckipsd | NCK interacting protein with SH3 domain | 1123.51 | 1202.53 | -0.10 | 7.52E-01 |
| ENSMUSG00000023473 | GSMG0040106 | Celsr3 | cadherin, EGF LAG seven-pass G-type receptor 3 | 922.79 | 1157.62 | -0.33 | 3.28E-01 |
| ENSMUSG00000023259 | GSMG0040107 | Slc26a6 | solute carrier family 26, member 6 | 491.24 | 373.99 | 0.40 | 1.22E-01 |
| ENSMUSG00000025651 | GSMG0040109 | Uqcrc1 | ubiquinol-cytochrome c reductase core protein 1 | 5479.32 | 5521.78 | -0.01 | 9.73E-01 |
| ENSMUSG00000025650 | GSMG0040110 | Col7a1 | collagen, type VII, alpha 1 | 803.87 | 543.02 | 0.57 | 3.99E-01 |
| ENSMUSG00000025648 | GSMG0040112 | Pfkfb4 | 6-phosphofructo-2-kinase/fructose-2,6-biphosphatase 4 | 164.11 | 226.33 | -0.46 | 1.42E-01 |
| ENSMUSG00000025647 | GSMG0040113 | Shisa5 | shisa family member 5 | 3732.55 | 3579.49 | 0.06 | 8.03E-01 |
| ENSMUSG00000025645 | GSMG0040114 | Ccdc51 | coiled-coil domain containing 51 | 307.47 | 292.42 | 0.07 | 8.39E-01 |
| ENSMUSG00000053646 | GSMG0040115 | Plxnb1 | plexin B1 | 807.99 | 911.28 | -0.17 | 6.15E-01 |
| ENSMUSG00000032478 | GSMG0040121 | Nme6 | NME/NM23 nucleoside diphosphate kinase 6 | 213.02 | 362.86 | -0.77 | 3.03E-04 |
| ENSMUSG00000032477 | GSMG0040122 | Cdc25a | cell division cycle 25A | 562.85 | 822.21 | -0.55 | 2.47E-03 |
| ENSMUSG00000032479 | GSMG0040123 | Map4 | microtubule-associated protein 4 | 3511.65 | 2771.43 | 0.34 | 3.77E-02 |
| NULL | GSMG0040129 | NULL | NULL | 301.72 | 174.41 | 0.79 | 6.08E-04 |
| ENSMUSG00000032481 | GSMG0040130 | Smarcc1 | SWI/SNF related, matrix associated, actin dependent regulator of chron | 1334.01 | 2277.93 | -0.77 | 2.40E-08 |
| ENSMUSG00000032482 | GSMG0040132 | Cspg5 | chondroitin sulfate proteoglycan 5 | 4634.38 | 7940.35 | -0.78 | 1.12E-07 |
| ENSMUSG00000054836 | GSMG0040133 | Elp6 | elongator acetyltransferase complex subunit 6 | 139.68 | 158.15 | -0.18 | 6.62E-01 |
| ENSMUSG00000032485 | GSMG0040134 | Scap | SREBF chaperone | 1538.93 | 1513.97 | 0.02 | 9.46E-01 |
| ENSMUSG00000032484 | GSMG0040136 | Ngp | neutrophilic granule protein | 23.69 | 129.12 | -2.44 | 1.91E-12 |
| ENSMUSG00000019659 | GSMG0040144 | Ccdc12 | coiled-coil domain containing 12 | 1296.35 | 1435.14 | -0.15 | 5.53E-01 |
| ENSMUSG00000044037 | GSMG0040152 | Als2cl | ALS2 C-terminal like | 831.73 | 365.93 | 1.18 | 2.45E-08 |
| ENSMUSG00000057802 | GSMG0040155 | Gm10030 | predicted gene 10030 | 452.41 | 609.54 | -0.43 | 1.16E-01 |
| ENSMUSG00000032497 | GSMG0040157 | Lrrfip2 | leucine rich repeat (in FLII) interacting protein 2 | 569.09 | 537.96 | 0.08 | 7.80E-01 |
| NULL | GSMG0040159 | NULL | NULL | 1566.19 | 1617.86 | -0.05 | 8.57E-01 |
| ENSMUSG00000032500 | GSMG0040163 | Dclk3 | doublecortin-like kinase 3 | 230.53 | 121.99 | 0.92 | 1.92E-04 |
| ENSMUSG00000089718 | GSMG0040165 | 2310075C17Rik | RIKEN cDNA 2310075C17 gene | 39.80 | 75.29 | -0.92 | 2.17E-02 |
| ENSMUSG00000033392 | GSMG0040171 | Clasp2 | CLIP associating protein 2 | 3566.84 | 3999.61 | -0.17 | 3.81E-01 |
| NULL | GSMG0040172 | NULL | NULL | 248.96 | 231.86 | 0.10 | 7.93E-01 |
| ENSMUSG00000009741 | GSMG0040174 | Ubp1 | upstream binding protein 1 | 925.50 | 908.86 | 0.02 | 9.40E-01 |
| G00000045594,ENSMUSG000 | GSMG0040178 | Glb1 // Tmppe | galactosidase, beta 1 // transmembrane protein with metallophosphoeste | 513.31 | 589.37 | -0.20 | 4.57E-01 |
| ENSMUSG00000032435 | GSMG0040181 | Dync1li1 | dynein cytoplasmic 1 light intermediate chain 1 | 2161.37 | 2722.10 | -0.33 | 3.89E-02 |

| | | | | | | | |
|---|---|---|---|---|---|---|---|
| ENSMUSG00000032434 | GSMG0040183 | Cmtm6 | CKLF-like MARVEL transmembrane domain containing 6 | 11922.24 | 12746.77 | -0.10 | 6.57E-01 |
| ENSMUSG00000040875 | GSMG0040189 | Osbpl10 | oxysterol binding protein-like 10 | 412.59 | 410.02 | 0.01 | 9.86E-01 |
| NULL | GSMG0040191 | NULL | NULL | 146.19 | 140.89 | 0.05 | 9.21E-01 |
| ENSMUSG00000056880 | GSMG0040196 | Gadl1 | glutamate decarboxylase-like 1 | 423.05 | 625.75 | -0.57 | 2.53E-02 |
| ENSMUSG00000039285 | GSMG0040203 | Azi2 | 5-azacytidine induced gene 2 | 788.50 | 812.36 | -0.04 | 8.93E-01 |
| ENSMUSG00000038708 | GSMG0040207 | Golga4 | golgi autoantigen, golgin subfamily a, 4 | 539.52 | 536.76 | 0.01 | 9.83E-01 |
| ENSMUSG00000039115 | GSMG0040209 | Itga9 | integrin alpha 9 | 184.77 | 105.40 | 0.81 | 2.18E-02 |
| ENSMUSG00000047409 | GSMG0040214 | Ctdspl | CTD (carboxy-terminal domain, RNA polymerase II, polypeptide A) sm | 609.23 | 254.99 | 1.26 | 2.33E-07 |
| ENSMUSG00000036138 | GSMG0040219 | Acaa1a | acetyl-Coenzyme A acyltransferase 1A | 1828.13 | 1560.22 | 0.23 | 2.44E-01 |
| ENSMUSG00000061393 | GSMG0040222 | Acvr2b | activin receptor IIB | 524.44 | 517.11 | 0.02 | 9.57E-01 |
| ENSMUSG00000042787 | GSMG0040224 | Exog | endo/exonuclease (5'-3'), endonuclease G-like | 631.11 | 739.72 | -0.23 | 3.37E-01 |
| ENSMUSG00000032512 | GSMG0040227 | Wdr48 | WD repeat domain 48 | 1572.38 | 1416.38 | 0.15 | 5.44E-01 |
| ENSMUSG00000032519 | GSMG0040233 | Slc25a38 | solute carrier family 25, member 38 | 347.96 | 331.40 | 0.07 | 8.32E-01 |
| ENSMUSG00000032517 | GSMG0040235 | Mobp | myelin-associated oligodendrocytic basic protein | 1794.45 | 0.66 | 11.46 | 3.89E-26 |
| ENSMUSG00000041794 | GSMG0040236 | Myrip | myosin VIIA and Rab interacting protein | 608.30 | 253.32 | 1.26 | 2.75E-07 |
| ENSMUSG00000006941 | GSMG0040237 | Eif1b | eukaryotic translation initiation factor 1B | 1601.16 | 2248.80 | -0.49 | 1.10E-02 |
| ENSMUSG00000006932 | GSMG0040243 | Ctnnb1 | catenin (cadherin associated protein), beta 1 | 7284.70 | 6415.52 | 0.18 | 4.79E-01 |
| ENSMUSG00000032536 | GSMG0040244 | Trak1 | trafficking protein, kinesin binding 1 | 1636.77 | 1569.94 | 0.06 | 8.66E-01 |
| ENSMUSG00000032528 | GSMG0040248 | Vipr1 | vasoactive intestinal peptide receptor 1 | 135.57 | 31.88 | 2.09 | 1.22E-08 |
| ENSMUSG00000032525 | GSMG0040249 | Nktr | natural killer tumor recognition sequence | 345.81 | 425.05 | -0.30 | 2.05E-01 |
| ENSMUSG00000032526 | GSMG0040252 | Deb1 | differentially expressed in B16F10 1 | 385.20 | 319.03 | 0.27 | 2.94E-01 |
| ENSMUSG00000013419 | GSMG0040253 | Zfp651 | zinc finger protein 651 | 1724.86 | 1284.91 | 0.42 | 6.97E-02 |
| ENSMUSG00000044534 | GSMG0040257 | Ackr2 | atypical chemokine receptor 2 | 1684.64 | 3054.36 | -0.86 | 1.00E-07 |
| ENSMUSG00000043773 | GSMG0040258 | 1700048O20Rik | RIKEN cDNA 1700048O20 gene | 225.96 | 320.83 | -0.51 | 6.93E-02 |
| ENSMUSG00000038145 | GSMG0040260 | Snrk | SNF related kinase | 2712.30 | 1956.89 | 0.47 | 5.05E-03 |
| ENSMUSG00000032540 | GSMG0040263 | Abhd5 | abhydrolase domain containing 5 | 1391.92 | 1592.25 | -0.19 | 4.21E-01 |
| ENSMUSG00000097736 | GSMG0040264 | 9530059O14Rik | RIKEN cDNA 9530059O14 gene | 3091.98 | 1362.30 | 1.18 | 1.05E-12 |
| ENSMUSG00000046603 | GSMG0040265 | Tcaim | T cell activation inhibitor, mitochondrial | 246.12 | 191.41 | 0.36 | 3.22E-01 |
| ENSMUSG00000057895 | GSMG0040268 | Zfp105 | zinc finger protein 105 | 431.94 | 469.93 | -0.12 | 6.93E-01 |
| ENSMUSG00000036768 | GSMG0040269 | Kif15 | kinesin family member 15 | 158.68 | 415.40 | -1.38 | 9.18E-07 |
| ENSMUSG00000066233 | GSMG0040271 | Tmem42 | transmembrane protein 42 | 469.61 | 428.79 | 0.13 | 6.59E-01 |
| ENSMUSG00000025785 | GSMG0040273 | Exosc7 | exosome component 7 | 707.25 | 791.76 | -0.16 | 5.15E-01 |
| ENSMUSG00000025784 | GSMG0040275 | Clec3b | C-type lectin domain family 3, member b | 155.65 | 29.06 | 2.43 | 1.09E-10 |

| | | | | | | | |
|---|---|---|---|---|---|---|---|
| ENSMUSG00000035202 | GSMG0040276 | Lars2 | leucyl-tRNA synthetase, mitochondrial | 470.46 | 637.12 | -0.44 | 3.65E-02 |
| ENSMUSG00000025239 | GSMG0040278 | Limd1 | LIM domains containing 1 | 2785.35 | 2638.66 | 0.08 | 7.69E-01 |
| ENSMUSG00000025240 | GSMG0040279 | Sacm1l | SAC1 suppressor of actin mutations 1-like (yeast) | 1380.61 | 1165.67 | 0.24 | 3.03E-01 |
| NULL | GSMG0040280 | NULL | NULL | 848.59 | 822.96 | 0.04 | 8.97E-01 |
| ENSMUSG00000029530 | GSMG0040282 | Ccr9 | chemokine (C-C motif) receptor 9 | 762.94 | 349.42 | 1.13 | 1.23E-11 |
| ENSMUSG00000025894 | GSMG0040293 | Aasdhppt | aminoadipate-semialdehyde dehydrogenase-phosphopantetheinyl transf | 483.56 | 657.51 | -0.44 | 6.92E-02 |
| ENSMUSG00000025892 | GSMG0040294 | Gria4 | glutamate receptor, ionotropic, AMPA4 (alpha 4) | 842.02 | 873.75 | -0.05 | 8.72E-01 |
| NULL | GSMG0040295 | NULL | NULL | 293.12 | 253.54 | 0.21 | 6.24E-01 |
| NULL | GSMG0040297 | NULL | NULL | 288.41 | 214.14 | 0.43 | 3.59E-01 |
| NULL | GSMG0040300 | NULL | NULL | 251.32 | 262.81 | -0.07 | 8.90E-01 |
| ENSMUSG00000047193 | GSMG0040309 | Dync2h1 | dynein cytoplasmic 2 heavy chain 1 | 609.91 | 615.61 | -0.01 | 9.72E-01 |
| NULL | GSMG0040314 | NULL | NULL | 697.23 | 650.37 | 0.10 | 7.56E-01 |
| ENSMUSG00000032000 | GSMG0040317 | Birc3 | baculoviral IAP repeat-containing 3 | 509.71 | 437.31 | 0.22 | 4.29E-01 |
| NULL | GSMG0040319 | NULL | NULL | 188.08 | 103.49 | 0.87 | 5.81E-03 |
| ENSMUSG00000053110 | GSMG0040320 | Yap1 | yes-associated protein 1 | 424.95 | 459.52 | -0.11 | 7.70E-01 |
| ENSMUSG00000050730 | GSMG0040328 | Arhgap42 | Rho GTPase activating protein 42 | 685.40 | 650.09 | 0.08 | 7.87E-01 |
| NULL | GSMG0040334 | NULL | NULL | 204.54 | 160.14 | 0.36 | 3.37E-01 |
| NULL | GSMG0040341 | NULL | NULL | 352.49 | 509.21 | -0.53 | 3.12E-02 |
| NULL | GSMG0040344 | NULL | NULL | 84.13 | 132.71 | -0.66 | 7.73E-02 |
| NULL | GSMG0040348 | NULL | NULL | 138.27 | 150.15 | -0.12 | 8.09E-01 |
| NULL | GSMG0040352 | NULL | NULL | 104.31 | 171.00 | -0.71 | 2.76E-02 |
| ENSMUSG00000031922 | GSMG0040358 | Cep57 | centrosomal protein 57 | 613.18 | 796.70 | -0.38 | 6.96E-02 |
| ENSMUSG00000037419 | GSMG0040360 | Endod1 | endonuclease domain containing 1 | 2659.37 | 2283.80 | 0.22 | 2.14E-01 |
| ENSMUSG00000013076 | GSMG0040363 | Amotl1 | angiomotin-like 1 | 391.05 | 396.68 | -0.02 | 9.58E-01 |
| ENSMUSG00000031931 | GSMG0040369 | Ankrd49 | ankyrin repeat domain 49 | 538.16 | 712.71 | -0.41 | 8.44E-02 |
| ENSMUSG00000031934 | GSMG0040371 | Panx1 | pannexin 1 | 222.24 | 609.39 | -1.45 | 9.18E-15 |
| ENSMUSG00000031935 | GSMG0040374 | Med17 | mediator complex subunit 17 | 657.84 | 681.54 | -0.05 | 8.65E-01 |
| ENSMUSG00000031938 | GSMG0040375 | 4931406C07Rik | RIKEN cDNA 4931406C07 gene | 126.15 | 160.03 | -0.35 | 4.44E-01 |
| NULL | GSMG0040386 | NULL | NULL | 208.98 | 219.79 | -0.08 | 8.57E-01 |
| NULL | GSMG0040387 | NULL | NULL | 333.73 | 350.66 | -0.07 | 8.30E-01 |
| ENSMUSG00000043943 | GSMG0040392 | Naalad2 | N-acetylated alpha-linked acidic dipeptidase 2 | 618.97 | 607.71 | 0.03 | 9.37E-01 |
| ENSMUSG00000045519 | GSMG0040409 | Zfp560 | zinc finger protein 560 | 365.39 | 445.82 | -0.29 | 2.31E-01 |
| ENSMUSG00000063108 | GSMG0040412 | Zfp26 | zinc finger protein 26 | 153.46 | 206.36 | -0.43 | 1.56E-01 |

| | | | | | | | |
|---|---|---|---|---|---|---|---|
| ENSMUSG00000059475 | GSMG0040414 | Zfp426 | zinc finger protein 426 | 135.73 | 148.76 | -0.13 | 7.60E-01 |
| ENSMUSG00000060510 | GSMG0040417 | Zfp266 | zinc finger protein 266 | 278.77 | 272.83 | 0.03 | 9.43E-01 |
| ENSMUSG00000066892 | GSMG0040422 | Fbxl12 | F-box and leucine-rich repeat protein 12 | 580.87 | 677.52 | -0.22 | 3.51E-01 |
| ENSMUSG00000032172 | GSMG0040423 | Olfm2 | olfactomedin 2 | 447.42 | 701.43 | -0.65 | 1.87E-03 |
| ENSMUSG00000004098 | GSMG0040424 | Col5a3 | collagen, type V, alpha 3 | 929.06 | 373.27 | 1.31 | 3.96E-09 |
| ENSMUSG00000038742 | GSMG0040425 | Angptl6 | angiopoietin-like 6 | 194.06 | 173.42 | 0.16 | 6.72E-01 |
| ENSMUSG00000070319 | GSMG0040426 | Eif3g | eukaryotic translation initiation factor 3, subunit G | 2631.61 | 3140.36 | -0.25 | 1.24E-01 |
| ENSMUSG00000004099 | GSMG0040427 | Dnmt1 | DNA methyltransferase (cytosine-5) 1 | 2806.62 | 4761.67 | -0.76 | 5.40E-08 |
| ENSMUSG00000043895 | GSMG0040428 | S1pr2 | sphingosine-1-phosphate receptor 2 | 334.35 | 359.16 | -0.10 | 7.95E-01 |
| NULL | GSMG0040431 | NULL | NULL | 294.94 | 131.05 | 1.17 | 5.18E-05 |
| G00000079677,ENSMUSG000 | GSMG0040432 | Zglp1 // Gm38431 | zinc finger, GATA-like protein 1 // predicted gene, 38431 // ferredoxin | 1008.37 | 1172.78 | -0.22 | 4.24E-01 |
| ENSMUSG00000032175 | GSMG0040433 | Tyk2 | tyrosine kinase 2 | 1299.02 | 1052.19 | 0.30 | 1.75E-01 |
| ENSMUSG00000019471 | GSMG0040434 | Cdc37 | cell division cycle 37 | 3293.02 | 3829.86 | -0.22 | 2.91E-01 |
| ENSMUSG00000003308 | GSMG0040437 | Keap1 | kelch-like ECH-associated protein 1 | 1046.89 | 1066.06 | -0.03 | 9.33E-01 |
| ENSMUSG00000035047 | GSMG0040440 | Kri1 | KRI1 homolog | 456.56 | 496.33 | -0.12 | 6.82E-01 |
| ENSMUSG00000096472 | GSMG0040441 | Cdkn2d | cyclin-dependent kinase inhibitor 2D (p19, inhibits CDK4) | 931.18 | 1189.58 | -0.35 | 7.65E-02 |
| ENSMUSG00000074476 | GSMG0040444 | Spc24 | SPC24, NDC80 kinetochore complex component, homolog (S. cerevisi | 167.84 | 420.52 | -1.32 | 1.11E-07 |
| ENSMUSG00000032180 | GSMG0040447 | Tmed1 | transmembrane emp24 domain containing 1 | 1323.80 | 1519.28 | -0.20 | 3.66E-01 |
| ENSMUSG00000032182 | GSMG0040451 | Yipf2 | Yip1 domain family, member 2 | 541.52 | 367.01 | 0.56 | 1.28E-02 |
| ENSMUSG00000032194 | GSMG0040454 | Kank2 | KN motif and ankyrin repeat domains 2 | 1066.36 | 929.56 | 0.20 | 4.61E-01 |
| ENSMUSG00000032198 | GSMG0040455 | Dock6 | dedicator of cytokinesis 6 | 3396.47 | 3131.74 | 0.12 | 7.02E-01 |
| ENSMUSG00000019066 | GSMG0040456 | Rab3d | RAB3D, member RAS oncogene family | 476.88 | 624.83 | -0.39 | 8.72E-02 |
| ENSMUSG00000040883 | GSMG0040457 | Tmem205 | transmembrane protein 205 | 687.72 | 626.96 | 0.13 | 6.33E-01 |
| ENSMUSG00000006235 | GSMG0040459 | Epor | erythropoietin receptor | 331.57 | 512.69 | -0.63 | 1.45E-03 |
| ENSMUSG00000003410 | GSMG0040462 | Elavl3 | ELAV (embryonic lethal, abnormal vision, Drosophila)-like 3 (Hu antig | 104.41 | 222.45 | -1.10 | 1.47E-04 |
| ENSMUSG00000038895 | GSMG0040465 | Zfp653 | zinc finger protein 653 | 1277.24 | 1292.53 | -0.02 | 9.65E-01 |
| ENSMUSG00000066839 | GSMG0040467 | Ecsit | ECSIT signalling integrator | 996.58 | 1089.96 | -0.13 | 6.08E-01 |
| ENSMUSG00000013822 | GSMG0040468 | Elof1 | ELF1 homolog, elongation factor 1 | 962.26 | 1043.75 | -0.12 | 6.81E-01 |
| ENSMUSG00000001348 | GSMG0040469 | Acp5 | acid phosphatase 5, tartrate resistant | 132.22 | 159.26 | -0.26 | 5.10E-01 |
| ENSMUSG00000066829 | GSMG0040472 | Zfp810 | zinc finger protein 810 | 222.20 | 292.63 | -0.40 | 1.32E-01 |
| ENSMUSG00000036777 | GSMG0040475 | Anln | anillin, actin binding protein | 370.32 | 754.22 | -1.02 | 2.92E-07 |
| ENSMUSG00000032239 | GSMG0040477 | Rp9 | retinitis pigmentosa 9 (human) | 1168.46 | 1346.70 | -0.21 | 3.64E-01 |
| ENSMUSG00000043067 | GSMG0040485 | Dpy19l1 | dpy-19-like 1 (C. elegans) | 552.24 | 381.57 | 0.53 | 1.76E-02 |

| | | | | | | | |
|---|---|---|---|---|---|---|---|
| ENSMUSG00000008429 | GSMG0040492 | Herpud2 | HERPUD family member 2 | 1992.74 | 1714.28 | 0.22 | 2.77E-01 |
| ENSMUSG00000031969 | GSMG0040498 | Acad8 | acyl-Coenzyme A dehydrogenase family, member 8 | 661.51 | 665.41 | -0.01 | 9.85E-01 |
| ENSMUSG00000031988 | GSMG0040500 | Vps26b | VPS26 retromer complex component B | 396.42 | 445.83 | -0.17 | 5.79E-01 |
| ENSMUSG00000031990 | GSMG0040502 | Jam3 | junction adhesion molecule 3 | 656.46 | 565.53 | 0.22 | 3.58E-01 |
| NULL | GSMG0040504 | NULL | NULL | 253.99 | 245.03 | 0.06 | 8.94E-01 |
| ENSMUSG00000059974 | GSMG0040508 | Ntm | neurotrimin | 1492.36 | 2040.28 | -0.45 | 1.16E-02 |
| NULL | GSMG0040511 | NULL | NULL | 195.70 | 174.91 | 0.17 | 7.10E-01 |
| NULL | GSMG0040512 | NULL | NULL | 390.68 | 413.82 | -0.08 | 8.54E-01 |
| NULL | GSMG0040515 | NULL | NULL | 335.54 | 374.60 | -0.16 | 7.26E-01 |
| NULL | GSMG0040516 | NULL | NULL | 697.60 | 624.64 | 0.16 | 6.46E-01 |
| NULL | GSMG0040517 | NULL | NULL | 155.77 | 176.44 | -0.17 | 7.34E-01 |
| NULL | GSMG0040518 | NULL | NULL | 640.58 | 827.09 | -0.37 | 1.10E-01 |
| ENSMUSG00000031996 | GSMG0040528 | Aplp2 | amyloid beta (A4) precursor-like protein 2 | 23678.11 | 22604.55 | 0.07 | 7.78E-01 |
| ENSMUSG00000016087 | GSMG0040536 | Fli1 | Friend leukemia integration 1 | 1788.84 | 2266.82 | -0.34 | 9.94E-02 |
| NULL | GSMG0040540 | NULL | NULL | 735.27 | 609.71 | 0.27 | 1.89E-01 |
| NULL | GSMG0040541 | NULL | NULL | 554.69 | 500.24 | 0.15 | 5.69E-01 |
| NULL | GSMG0040543 | NULL | NULL | 526.51 | 665.30 | -0.34 | 2.21E-01 |
| ENSMUSG00000032038 | GSMG0040547 | St3gal4 | ST3 beta-galactoside alpha-2,3-sialyltransferase 4 | 276.23 | 250.22 | 0.14 | 6.64E-01 |
| ENSMUSG00000032040 | GSMG0040549 | Dcps | decapping enzyme, scavenger | 299.16 | 382.29 | -0.35 | 1.35E-01 |
| ENSMUSG00000032041 | GSMG0040551 | Tirap | toll-interleukin 1 receptor (TIR) domain-containing adaptor protein | 458.28 | 476.05 | -0.06 | 8.73E-01 |
| ENSMUSG00000039048 | GSMG0040552 | Foxred1 | FAD-dependent oxidoreductase domain containing 1 | 1400.47 | 1099.10 | 0.35 | 7.19E-02 |
| ENSMUSG00000050471 | GSMG0040553 | Fam118b | family with sequence similarity 118, member B | 824.78 | 724.51 | 0.19 | 4.66E-01 |
| ENSMUSG00000032101 | GSMG0040557 | Ddx25 | DEAD (Asp-Glu-Ala-Asp) box polypeptide 25 | 333.32 | 530.00 | -0.67 | 5.89E-04 |
| ENSMUSG00000032113 | GSMG0040570 | Chek1 | checkpoint kinase 1 | 52.69 | 145.21 | -1.45 | 1.45E-05 |
| ENSMUSG00000032116 | GSMG0040571 | Stt3a | STT3, subunit of the oligosaccharyltransferase complex, homolog A (S. | 2283.06 | 2786.95 | -0.29 | 1.50E-01 |
| ENSMUSG00000062762 | GSMG0040572 | Ei24 | etoposide induced 2.4 mRNA | 1679.41 | 1858.76 | -0.15 | 4.85E-01 |
| ENSMUSG00000035934 | GSMG0040573 | Pknox2 | Pbx/knotted 1 homeobox 2 | 182.72 | 178.72 | 0.03 | 9.47E-01 |
| NULL | GSMG0040574 | NULL | NULL | 102.21 | 38.03 | 1.42 | 1.09E-04 |
| ENSMUSG00000032128 | GSMG0040581 | Robo3 | roundabout guidance receptor 3 | 1076.35 | 194.05 | 2.47 | 2.49E-30 |
| ENSMUSG00000053310 | GSMG0040582 | Nrgn | neurogranin | 3063.93 | 1443.61 | 1.08 | 1.94E-05 |
| ENSMUSG00000011114 | GSMG0040584 | Tbrg1 | transforming growth factor beta regulated gene 1 | 1875.18 | 2197.38 | -0.23 | 2.41E-01 |
| ENSMUSG00000040111 | GSMG0040611 | Gramd1b | GRAM domain containing 1B | 447.53 | 469.30 | -0.07 | 8.37E-01 |
| NULL | GSMG0040616 | NULL | NULL | 46.52 | 93.67 | -1.00 | 5.26E-03 |

| | | | | | | | |
|---|---|---|---|---|---|---|---|
| ENSMUSG00000032020 | GSMG0040619 | Ubash3b | ubiquitin associated and SH3 domain containing, B | 176.83 | 280.21 | -0.66 | 2.12E-02 |
| ENSMUSG00000049313 | GSMG0040624 | Sorl1 | sortilin-related receptor, LDLR class A repeats-containing | 390.36 | 261.79 | 0.57 | 1.03E-02 |
| ENSMUSG00000032018 | GSMG0040626 | Sc5d | sterol-C5-desaturase | 187.40 | 277.08 | -0.56 | 4.11E-02 |
| ENSMUSG00000037287 | GSMG0040628 | Tbcel | tubulin folding cofactor E-like | 431.12 | 363.26 | 0.25 | 3.80E-01 |
| ENSMUSG00000032017 | GSMG0040629 | Grik4 | glutamate receptor, ionotropic, kainate 4 | 360.58 | 309.65 | 0.22 | 5.56E-01 |
| ENSMUSG00000059495 | GSMG0040631 | Arhgef12 | Rho guanine nucleotide exchange factor (GEF) 12 | 2373.36 | 2518.71 | -0.09 | 8.02E-01 |
| ENSMUSG00000048503 | GSMG0040634 | Tmem136 | transmembrane protein 136 | 1088.96 | 1202.56 | -0.14 | 5.10E-01 |
| ENSMUSG00000032014 | GSMG0040636 | Oaf | out at first homolog | 648.56 | 488.51 | 0.41 | 4.00E-02 |
| NULL | GSMG0040644 | NULL | NULL | 880.00 | 1328.88 | -0.60 | 6.13E-04 |
| ENSMUSG00000034342 | GSMG0040646 | Cbl | Casitas B-lineage lymphoma | 383.32 | 462.23 | -0.27 | 2.42E-01 |
| ENSMUSG00000032109 | GSMG0040649 | Nlrx1 | NLR family member X1 | 266.20 | 257.03 | 0.05 | 8.95E-01 |
| ENSMUSG00000032131 | GSMG0040650 | Abcg4 | ATP-binding cassette, sub-family G (WHITE), member 4 | 423.19 | 594.54 | -0.49 | 4.37E-02 |
| ENSMUSG00000032119 | GSMG0040651 | Hinfp | histone H4 transcription factor | 312.20 | 361.13 | -0.21 | 4.92E-01 |
| ENSMUSG00000032120 | GSMG0040652 | C2cd2l | C2 calcium-dependent domain containing 2-like | 1229.06 | 967.78 | 0.34 | 1.55E-01 |
| ENSMUSG00000032126 | GSMG0040653 | Hmbs | hydroxymethylbilane synthase | 460.58 | 614.06 | -0.41 | 2.96E-02 |
| ENSMUSG00000032127 | GSMG0040654 | Vps11 | VPS11, CORVET/HOPS core subunit | 1842.16 | 1907.47 | -0.05 | 8.64E-01 |
| ENSMUSG00000032112 | GSMG0040656 | Trappc4 | trafficking protein particle complex 4 | 1610.03 | 2017.61 | -0.33 | 9.54E-02 |
| ENSMUSG00000043923 | GSMG0040657 | Ccdc84 | coiled-coil domain containing 84 | 611.88 | 608.06 | 0.01 | 9.79E-01 |
| ENSMUSG00000048537 | GSMG0040667 | Phldb1 | pleckstrin homology like domain, family B, member 1 | 379.16 | 404.59 | -0.10 | 8.47E-01 |
| NULL | GSMG0040668 | NULL | NULL | 96.21 | 81.92 | 0.22 | 6.72E-01 |
| ENSMUSG00000032096 | GSMG0040669 | Arcn1 | archain 1 | 2463.05 | 2885.57 | -0.23 | 2.78E-01 |
| ENSMUSG00000002032 | GSMG0040670 | Tmem25 | transmembrane protein 25 | 1310.95 | 610.64 | 1.10 | 6.19E-11 |
| ENSMUSG00000002028 | GSMG0040672 | Kmt2a | lysine (K)-specific methyltransferase 2A | 267.60 | 312.52 | -0.23 | 4.72E-01 |
| NULL | GSMG0040674 | NULL | NULL | 315.78 | 258.58 | 0.29 | 2.84E-01 |
| ENSMUSG00000059890 | GSMG0040676 | Ube4a | ubiquitination factor E4A | 561.53 | 624.72 | -0.15 | 5.44E-01 |
| NULL | GSMG0040686 | NULL | NULL | 496.71 | 1106.31 | -1.15 | 2.41E-11 |
| NULL | GSMG0040692 | NULL | NULL | 447.14 | 378.43 | 0.24 | 3.21E-01 |
| ENSMUSG00000042790 | GSMG0040693 | Rnf214 | ring finger protein 214 | 575.26 | 654.78 | -0.19 | 4.06E-01 |
| ENSMUSG00000032085 | GSMG0040695 | Tagln | transgelin | 1182.91 | 201.83 | 2.55 | 8.26E-32 |
| ENSMUSG00000034908 | GSMG0040696 | Sidt2 | SID1 transmembrane family, member 2 | 4664.48 | 3872.44 | 0.27 | 2.13E-01 |
| ENSMUSG00000003131 | GSMG0040698 | Pafah1b2 | platelet-activating factor acetylhydrolase, isoform 1b, subunit 2 | 4874.90 | 5369.52 | -0.14 | 5.24E-01 |
| ENSMUSG00000032026 | GSMG0040710 | Rexo2 | RNA exonuclease 2 | 2082.90 | 2090.04 | -0.01 | 9.88E-01 |
| ENSMUSG00000042396 | GSMG0040711 | Rbm7 | RNA binding motif protein 7 | 2111.90 | 2033.14 | 0.05 | 8.66E-01 |

| | | | | | | | |
|---|---|---|---|---|---|---|---|
| ENSMUSG00000066687 | GSMG0040714 | Zbtb16 | zinc finger and BTB domain containing 16 | 74.24 | 60.00 | 0.31 | 5.88E-01 |
| ENSMUSG00000040219 | GSMG0040724 | Ttc12 | tetratricopeptide repeat domain 12 | 125.19 | 83.94 | 0.58 | 7.30E-02 |
| ENSMUSG00000039542 | GSMG0040726 | Ncam1 | neural cell adhesion molecule 1 | 1541.03 | 3794.74 | -1.30 | 3.03E-12 |
| ENSMUSG00000032067 | GSMG0040733 | Pts | 6-pyruvoyl-tetrahydropterin synthase | 610.99 | 506.51 | 0.27 | 2.68E-01 |
| ENSMUSG00000000171 | GSMG0040736 | Sdhd | succinate dehydrogenase complex, subunit D, integral membrane protei | 3777.73 | 4378.59 | -0.21 | 2.86E-01 |
| ENSMUSG00000059820 | GSMG0040737 | AU019823 | expressed sequence AU019823 | 1092.73 | 1276.35 | -0.23 | 3.66E-01 |
| ENSMUSG00000000168 | GSMG0040738 | Dlat | dihydrolipoamide S-acetyltransferase (E2 component of pyruvate dehyc | 1156.07 | 1214.32 | -0.07 | 7.84E-01 |
| ENSMUSG00000032064 | GSMG0040739 | Dixdc1 | DIX domain containing 1 | 425.42 | 371.01 | 0.20 | 4.55E-01 |
| ENSMUSG00000032062 | GSMG0040741 | 2310030G06Rik | RIKEN cDNA 2310030G06 gene | 122.01 | 90.96 | 0.43 | 2.20E-01 |
| ENSMUSG00000038086 | GSMG0040742 | Hspb2 | heat shock protein 2 | 160.92 | 87.85 | 0.88 | 1.26E-03 |
| ENSMUSG00000037971 | GSMG0040743 | 1110032A03Rik | RIKEN cDNA 1110032A03 gene | 529.16 | 560.91 | -0.08 | 7.69E-01 |
| ENSMUSG00000037112 | GSMG0040747 | Sik2 | salt inducible kinase 2 | 730.55 | 639.27 | 0.19 | 4.76E-01 |
| ENSMUSG00000060594 | GSMG0040749 | Layn | layilin | 291.52 | 121.08 | 1.27 | 1.18E-08 |
| ENSMUSG00000032051 | GSMG0040755 | Fdx1 | ferredoxin 1 | 534.34 | 532.88 | 0.00 | 9.93E-01 |
| ENSMUSG00000078307 | GSMG0040761 | AI593442 | expressed sequence AI593442 | 1893.37 | 1967.71 | -0.06 | 8.48E-01 |
| ENSMUSG00000053289 | GSMG0040762 | Ddx10 | DEAD (Asp-Glu-Ala-Asp) box polypeptide 10 | 434.76 | 525.50 | -0.27 | 3.18E-01 |
| ENSMUSG00000034218 | GSMG0040765 | Atm | ataxia telangiectasia mutated | 249.46 | 295.27 | -0.24 | 4.00E-01 |
| ENSMUSG00000032047 | GSMG0040768 | Acat1 | acetyl-Coenzyme A acetyltransferase 1 | 2344.49 | 3178.56 | -0.44 | 3.33E-02 |
| ENSMUSG00000032030 | GSMG0040769 | Cul5 | cullin 5 | 762.78 | 857.99 | -0.17 | 5.06E-01 |
| ENSMUSG00000041986 | GSMG0040772 | Elmod1 | ELMO/CED-12 domain containing 1 | 580.65 | 876.17 | -0.60 | 8.65E-03 |
| ENSMUSG00000037493 | GSMG0040778 | Cib2 | calcium and integrin binding family member 2 | 121.32 | 447.15 | -1.88 | 7.80E-22 |
| ENSMUSG00000032281 | GSMG0040779 | Acsbg1 | acyl-CoA synthetase bubblegum family member 1 | 1699.72 | 1447.77 | 0.23 | 3.25E-01 |
| ENSMUSG00000061559 | GSMG0040782 | Wdr61 | WD repeat domain 61 | 1329.51 | 1492.33 | -0.17 | 4.27E-01 |
| ENSMUSG00000032314 | GSMG0040790 | Etfa | electron transferring flavoprotein, alpha polypeptide | 1784.36 | 2044.64 | -0.20 | 3.93E-01 |
| ENSMUSG00000034007 | GSMG0040795 | Scaper | S phase cyclin A-associated protein in the ER | 750.45 | 840.94 | -0.16 | 5.44E-01 |
| ENSMUSG00000032324 | GSMG0040802 | Tspan3 | tetraspanin 3 | 3948.40 | 4226.70 | -0.10 | 7.07E-01 |
| ENSMUSG00000074305 | GSMG0040803 | Peak1 | pseudopodium-enriched atypical kinase 1 | 575.04 | 473.77 | 0.28 | 2.84E-01 |
| ENSMUSG00000049556 | GSMG0040808 | Lingo1 | leucine rich repeat and Ig domain containing 1 | 2541.46 | 2436.72 | 0.06 | 8.37E-01 |
| ENSMUSG00000032733 | GSMG0040811 | Snx33 | sorting nexin 33 | 485.15 | 319.49 | 0.60 | 1.57E-02 |
| ENSMUSG00000032298 | GSMG0040816 | Neil1 | nei endonuclease VIII-like 1 (E. coli) | 404.83 | 392.33 | 0.05 | 8.90E-01 |
| ENSMUSG00000032299 | GSMG0040817 | Commd4 | COMM domain containing 4 | 1683.85 | 1561.94 | 0.11 | 6.50E-01 |
| ENSMUSG00000032300 | GSMG0040818 | 1700017B05Rik | RIKEN cDNA 1700017B05 gene | 472.71 | 395.06 | 0.26 | 3.01E-01 |
| ENSMUSG00000063849 | GSMG0040819 | Ppcdc | phosphopantothenoylcysteine decarboxylase | 360.33 | 291.38 | 0.31 | 2.21E-01 |

| | | | | | | | |
|---|---|---|---|---|---|---|---|
| ENSMUSG00000040722 | GSMG0040820 | Scamp5 | secretory carrier membrane protein 5 | 2879.21 | 3346.50 | -0.22 | 3.28E-01 |
| NULL | GSMG0040822 | NULL | NULL | 310.38 | 251.32 | 0.30 | 4.44E-01 |
| ENSMUSG00000032306 | GSMG0040824 | Mpi | mannose phosphate isomerase | 597.58 | 595.76 | 0.01 | 9.86E-01 |
| ENSMUSG00000032312 | GSMG0040827 | Csk | c-src tyrosine kinase | 688.80 | 783.47 | -0.19 | 5.31E-01 |
| ENSMUSG00000032316 | GSMG0040830 | Clk3 | CDC-like kinase 3 | 1608.18 | 1406.17 | 0.19 | 3.55E-01 |
| ENSMUSG00000004661 | GSMG0040832 | Arid3b | AT rich interactive domain 3B (BRIGHT-like) | 283.74 | 288.70 | -0.02 | 9.53E-01 |
| ENSMUSG00000037206 | GSMG0040836 | Islr | immunoglobulin superfamily containing leucine-rich repeat | 1135.42 | 934.21 | 0.28 | 3.30E-01 |
| ENSMUSG00000051243 | GSMG0040837 | Islr2 | immunoglobulin superfamily containing leucine-rich repeat 2 | 127.60 | 356.00 | -1.49 | 9.81E-09 |
| ENSMUSG00000036986 | GSMG0040838 | Pml | promyelocytic leukemia | 108.81 | 94.31 | 0.20 | 6.61E-01 |
| ENSMUSG00000035914 | GSMG0040842 | Cd276 | CD276 antigen | 269.62 | 651.71 | -1.27 | 3.95E-07 |
| ENSMUSG00000032340 | GSMG0040846 | Neo1 | neogenin | 1610.03 | 2676.16 | -0.73 | 1.29E-05 |
| ENSMUSG00000025235 | GSMG0040849 | Bbs4 | Bardet-Biedl syndrome 4 (human) | 344.79 | 341.93 | 0.02 | 9.69E-01 |
| ENSMUSG00000025234 | GSMG0040850 | Arih1 | ariadne RBR E3 ubiquitin protein ligase 1 | 408.90 | 386.63 | 0.08 | 8.38E-01 |
| NULL | GSMG0040854 | NULL | NULL | 139.76 | 84.51 | 0.73 | 4.39E-02 |
| ENSMUSG00000047766 | GSMG0040871 | Lrrc49 | leucine rich repeat containing 49 | 1647.74 | 1048.05 | 0.65 | 5.21E-04 |
| ENSMUSG00000007892 | GSMG0040878 | Rplp1 | ribosomal protein, large, P1 | 2266.56 | 2612.86 | -0.21 | 4.33E-01 |
| ENSMUSG00000032254 | GSMG0040879 | Kif23 | kinesin family member 23 | 304.32 | 544.37 | -0.84 | 2.15E-05 |
| ENSMUSG00000032278 | GSMG0040880 | Paqr5 | progestin and adipoQ receptor family member V | 685.05 | 86.46 | 2.99 | 1.74E-55 |
| ENSMUSG00000032252 | GSMG0040883 | Glce | glucuronyl C5-epimerase | 1586.30 | 1370.48 | 0.21 | 2.95E-01 |
| ENSMUSG00000041729 | GSMG0040889 | Coro2b | coronin, actin binding protein, 2B | 390.32 | 613.52 | -0.65 | 3.94E-03 |
| ENSMUSG00000032244 | GSMG0040891 | Fem1b | feminization 1 homolog b (C. elegans) | 2273.96 | 2359.30 | -0.05 | 8.53E-01 |
| ENSMUSG00000032405 | GSMG0040896 | Pias1 | protein inhibitor of activated STAT 1 | 1687.88 | 1461.40 | 0.21 | 3.20E-01 |
| ENSMUSG00000058444 | GSMG0040899 | Map2k5 | mitogen-activated protein kinase kinase 5 | 669.06 | 659.08 | 0.02 | 9.47E-01 |
| NULL | GSMG0040902 | NULL | NULL | 211.11 | 125.29 | 0.75 | 2.38E-03 |
| ENSMUSG00000032403 | GSMG0040904 | 2300009A05Rik | RIKEN cDNA 2300009A05 gene | 336.84 | 301.86 | 0.15 | 6.22E-01 |
| ENSMUSG00000032402 | GSMG0040906 | Smad3 | SMAD family member 3 | 267.08 | 351.09 | -0.40 | 1.72E-01 |
| ENSMUSG00000036867 | GSMG0040910 | Smad6 | SMAD family member 6 | 1104.22 | 382.82 | 1.53 | 1.57E-17 |
| ENSMUSG00000032400 | GSMG0040912 | Zwilch | zwilch kinetochore protein | 72.76 | 198.93 | -1.44 | 1.26E-05 |
| ENSMUSG00000004936 | GSMG0040913 | Map2k1 | mitogen-activated protein kinase kinase 1 | 1367.48 | 1457.94 | -0.09 | 7.32E-01 |
| ENSMUSG00000032396 | GSMG0040915 | Dis3l | DIS3 like exosome 3'-5' exoribonuclease | 696.38 | 778.25 | -0.16 | 4.91E-01 |
| NULL | GSMG0040917 | NULL | NULL | 525.51 | 667.64 | -0.35 | 1.38E-01 |
| ENSMUSG00000004771 | GSMG0040918 | Rab11a | RAB11A, member RAS oncogene family | 6766.37 | 6722.36 | 0.01 | 9.74E-01 |
| ENSMUSG00000033629 | GSMG0040922 | Hacd3 | 3-hydroxyacyl-CoA dehydratase 3 | 1525.34 | 2011.69 | -0.40 | 2.53E-02 |

| | | | | | | | |
|---|---|---|---|---|---|---|---|
| ENSMUSG00000040524 | GSMG0040931 | Zfp609 | zinc finger protein 609 | 368.31 | 385.06 | -0.07 | 8.77E-01 |
| NULL | GSMG0040933 | NULL | NULL | 161.36 | 178.12 | -0.15 | 7.28E-01 |
| ENSMUSG00000032386 | GSMG0040936 | Trip4 | thyroid hormone receptor interactor 4 | 597.39 | 561.40 | 0.09 | 7.96E-01 |
| ENSMUSG00000039452 | GSMG0040938 | Snx22 | sorting nexin 22 | 1365.62 | 1380.26 | -0.02 | 9.56E-01 |
| ENSMUSG00000032382 | GSMG0040939 | Snx1 | sorting nexin 1 | 2136.62 | 2002.38 | 0.09 | 6.91E-01 |
| ENSMUSG00000032376 | GSMG0040941 | Usp3 | ubiquitin specific peptidase 3 | 652.82 | 1057.04 | -0.69 | 2.23E-05 |
| ENSMUSG00000032375 | GSMG0040946 | Aph1b | aph1 homolog B, gamma secretase subunit | 739.38 | 819.50 | -0.15 | 5.18E-01 |
| ENSMUSG00000036943 | GSMG0040949 | Rab8b | RAB8B, member RAS oncogene family | 822.26 | 894.75 | -0.12 | 6.66E-01 |
| ENSMUSG00000032370 | GSMG0040952 | Lactb | lactamase, beta | 413.38 | 485.67 | -0.23 | 3.91E-01 |
| ENSMUSG00000032366 | GSMG0040953 | Tpm1 | tropomyosin 1, alpha | 4309.49 | 2723.21 | 0.66 | 3.88E-04 |
| ENSMUSG00000052698 | GSMG0040954 | Tln2 | talin 2 | 192.08 | 219.68 | -0.19 | 6.08E-01 |
| NULL | GSMG0040955 | NULL | NULL | 144.65 | 121.13 | 0.25 | 5.94E-01 |
| NULL | GSMG0040962 | NULL | NULL | 661.62 | 616.42 | 0.10 | 7.25E-01 |
| ENSMUSG00000032224 | GSMG0040968 | Fam81a | family with sequence similarity 81, member A | 781.36 | 639.81 | 0.29 | 1.95E-01 |
| ENSMUSG00000032218 | GSMG0040970 | Ccnb2 | cyclin B2 | 280.23 | 586.19 | -1.06 | 2.17E-06 |
| ENSMUSG00000032217 | GSMG0040971 | Rnf111 | ring finger 111 | 231.64 | 304.46 | -0.40 | 1.17E-01 |
| ENSMUSG00000042444 | GSMG0040975 | Fam63b | family with sequence similarity 63, member B | 1794.54 | 1231.05 | 0.54 | 1.15E-03 |
| NULL | GSMG0040976 | NULL | NULL | 828.24 | 976.72 | -0.24 | 3.90E-01 |
| ENSMUSG00000032207 | GSMG0040977 | Lipc | lipase, hepatic | 193.76 | 440.97 | -1.19 | 1.97E-09 |
| ENSMUSG00000032199 | GSMG0040980 | Polr2m | polymerase (RNA) II (DNA directed) polypeptide M | 4196.35 | 5024.97 | -0.26 | 1.42E-01 |
| ENSMUSG00000041361 | GSMG0040981 | Myzap | myocardial zonula adherens protein | 117.14 | 404.97 | -1.79 | 1.88E-17 |
| ENSMUSG00000032232 | GSMG0040982 | Cgnl1 | cingulin-like 1 | 4715.83 | 4339.81 | 0.12 | 6.21E-01 |
| NULL | GSMG0040998 | NULL | NULL | 358.63 | 300.69 | 0.25 | 4.50E-01 |
| ENSMUSG00000032228 | GSMG0041000 | Tcf12 | transcription factor 12 | 1015.82 | 1133.18 | -0.16 | 6.13E-01 |
| ENSMUSG00000090626 | GSMG0041004 | Tex9 | testis expressed gene 9 | 131.09 | 166.51 | -0.35 | 3.49E-01 |
| ENSMUSG00000086158 | GSMG0041007 | Ccpg1os | cell cycle progression 1, opposite strand | 83.52 | 87.71 | -0.08 | 9.02E-01 |
| NULL | GSMG0041017 | NULL | NULL | 313.33 | 221.39 | 0.50 | 3.58E-02 |
| ENSMUSG00000042688 | GSMG0041020 | Mapk6 | mitogen-activated protein kinase 6 | 703.25 | 685.91 | 0.04 | 9.03E-01 |
| ENSMUSG00000058587 | GSMG0041022 | Tmod3 | tropomodulin 3 | 2500.91 | 2186.58 | 0.19 | 5.05E-01 |
| ENSMUSG00000032186 | GSMG0041024 | Tmod2 | tropomodulin 2 | 1984.68 | 3291.49 | -0.73 | 2.44E-06 |
| ENSMUSG00000032181 | GSMG0041026 | Scg3 | secretogranin III | 1514.90 | 1279.99 | 0.24 | 2.96E-01 |
| ENSMUSG00000001366 | GSMG0041042 | Fbxo9 | f-box protein 9 | 1495.87 | 1311.01 | 0.19 | 3.58E-01 |
| ENSMUSG00000049624 | GSMG0041053 | Slc17a5 | solute carrier family 17 (anion/sugar transporter), member 5 | 463.55 | 522.64 | -0.17 | 5.03E-01 |

| | | | | | | | |
|---|---|---|---|---|---|---|---|
| ENSMUSG00000032330 | GSMG0041060 | Cox7a2 | cytochrome c oxidase subunit VIIa 2 | 3038.25 | 3480.31 | -0.20 | 4.50E-01 |
| ENSMUSG00000032328 | GSMG0041061 | Tmem30a | transmembrane protein 30A | 2126.01 | 1946.31 | 0.13 | 5.86E-01 |
| ENSMUSG00000034898 | GSMG0041064 | Filip1 | filamin A interacting protein 1 | 1210.79 | 886.32 | 0.45 | 2.21E-02 |
| ENSMUSG00000032253 | GSMG0041073 | Phip | pleckstrin homology domain interacting protein | 2087.74 | 2031.05 | 0.04 | 8.94E-01 |
| ENSMUSG00000032258 | GSMG0041077 | Lca5 | Leber congenital amaurosis 5 (human) | 215.44 | 324.78 | -0.59 | 1.58E-02 |
| ENSMUSG00000032262 | GSMG0041078 | Elovl4 | elongation of very long chain fatty acids (FEN1/Elo2, SUR4/Elo3, yeas | 185.60 | 304.76 | -0.71 | 4.31E-03 |
| ENSMUSG00000035941 | GSMG0041081 | Ibtk | inhibitor of Bruton agammaglobulinemia tyrosine kinase | 235.72 | 258.34 | -0.13 | 7.26E-01 |
| ENSMUSG00000056131 | GSMG0041085 | Pgm3 | phosphoglucomutase 3 | 365.68 | 615.79 | -0.75 | 4.06E-05 |
| ENSMUSG00000032418 | GSMG0041086 | Me1 | malic enzyme 1, NADP(+)-dependent, cytosolic | 190.02 | 335.51 | -0.82 | 8.77E-05 |
| ENSMUSG00000033419 | GSMG0041088 | Snap91 | synaptosomal-associated protein 91 | 1000.61 | 1591.05 | -0.67 | 6.30E-05 |
| ENSMUSG00000056919 | GSMG0041092 | Cep162 | centrosomal protein 162 | 181.36 | 207.84 | -0.19 | 5.53E-01 |
| ENSMUSG00000032419 | GSMG0041093 | Tbx18 | T-box18 | 250.62 | 154.20 | 0.70 | 5.94E-03 |
| ENSMUSG00000032422 | GSMG0041096 | Snx14 | sorting nexin 14 | 1110.55 | 1207.58 | -0.12 | 6.48E-01 |
| ENSMUSG00000032423 | GSMG0041099 | Syncrip | synaptotagmin binding, cytoplasmic RNA interacting protein | 835.11 | 1389.33 | -0.73 | 8.58E-05 |
| ENSMUSG00000097195 | GSMG0041100 | Snhg5 | small nucleolar RNA host gene 5 | 230.84 | 337.79 | -0.55 | 1.01E-02 |
| ENSMUSG00000032353 | GSMG0041114 | Tmed3 | transmembrane emp24 domain containing 3 | 1353.67 | 1278.91 | 0.08 | 7.79E-01 |
| ENSMUSG00000037410 | GSMG0041119 | Tbc1d2b | TBC1 domain family, member 2B | 232.97 | 297.56 | -0.35 | 2.74E-01 |
| NULL | GSMG0041130 | NULL | NULL | 152.59 | 104.58 | 0.54 | 1.05E-01 |
| ENSMUSG00000045414 | GSMG0041136 | 1190002N15Rik | RIKEN cDNA 1190002N15 gene | 760.34 | 725.46 | 0.07 | 8.37E-01 |
| NULL | GSMG0041141 | NULL | NULL | 323.93 | 275.72 | 0.23 | 3.99E-01 |
| ENSMUSG00000033350 | GSMG0041142 | Chst2 | carbohydrate sulfotransferase 2 | 3204.45 | 3309.80 | -0.05 | 8.74E-01 |
| ENSMUSG00000032407 | GSMG0041144 | U2surp | U2 snRNP-associated SURP domain containing | 1979.96 | 2535.17 | -0.36 | 1.15E-01 |
| ENSMUSG00000032839 | GSMG0041146 | Trpc1 | transient receptor potential cation channel, subfamily C, member 1 | 286.79 | 293.74 | -0.03 | 9.28E-01 |
| ENSMUSG00000032412 | GSMG0041152 | Atp1b3 | ATPase, Na+/K+ transporting, beta 3 polypeptide | 2190.62 | 2346.91 | -0.10 | 7.24E-01 |
| ENSMUSG00000051234 | GSMG0041155 | Rnf7 | ring finger protein 7 | 1562.21 | 1911.23 | -0.29 | 2.14E-01 |
| ENSMUSG00000032413 | GSMG0041156 | Rasa2 | RAS p21 protein activator 2 | 876.40 | 927.12 | -0.08 | 7.99E-01 |
| NULL | GSMG0041159 | NULL | NULL | 1272.36 | 1254.91 | 0.02 | 9.58E-01 |
| ENSMUSG00000040433 | GSMG0041160 | Zbtb38 // E030011 | zinc finger and BTB domain containing 38 // RIKEN cDNA E030011O | 656.14 | 695.45 | -0.08 | 8.15E-01 |
| ENSMUSG00000043587 | GSMG0041163 | Pxylp1 | 2-phosphoxylose phosphatase 1 | 37.64 | 60.10 | -0.66 | 2.02E-01 |
| ENSMUSG00000046997 | GSMG0041164 | Spsb4 | splA/ryanodine receptor domain and SOCS box containing 4 | 37.60 | 93.12 | -1.31 | 1.18E-03 |
| ENSMUSG00000032449 | GSMG0041167 | Slc25a36 | solute carrier family 25, member 36 | 1664.32 | 1911.12 | -0.20 | 3.73E-01 |
| ENSMUSG00000032452 | GSMG0041170 | Clstn2 | calsyntenin 2 | 567.54 | 806.95 | -0.51 | 4.30E-02 |
| NULL | GSMG0041173 | NULL | NULL | 194.76 | 242.55 | -0.32 | 3.49E-01 |

| | | | | | | | |
|---|---|---|---|---|---|---|---|
| NULL | GSMG0041174 | NULL | NULL | 283.35 | 371.73 | -0.39 | 1.76E-01 |
| ENSMUSG00000032459 | GSMG0041179 | Mrps22 | mitochondrial ribosomal protein S22 | 255.84 | 348.42 | -0.44 | 5.89E-02 |
| ENSMUSG00000097072 | GSMG0041183 | Foxl2os | forkhead box L2, opposite strand | 509.80 | 492.72 | 0.05 | 8.90E-01 |
| ENSMUSG00000032462 | GSMG0041185 | Pik3cb | phosphatidylinositol 3-kinase, catalytic, beta polypeptide | 237.09 | 180.37 | 0.39 | 1.74E-01 |
| ENSMUSG00000032470 | GSMG0041190 | Mras | muscle and microspikes RAS | 212.03 | 176.75 | 0.26 | 4.63E-01 |
| ENSMUSG00000032468 | GSMG0041191 | Armc8 | armadillo repeat containing 8 | 533.53 | 605.96 | -0.18 | 5.30E-01 |
| ENSMUSG00000032475 | GSMG0041200 | Nck1 | non-catalytic region of tyrosine kinase adaptor protein 1 | 826.08 | 713.02 | 0.21 | 4.09E-01 |
| ENSMUSG00000032527 | GSMG0041203 | Pccb | propionyl Coenzyme A carboxylase, beta polypeptide | 600.17 | 694.32 | -0.21 | 3.78E-01 |
| ENSMUSG00000032537 | GSMG0041214 | Ephb1 | Eph receptor B1 | 1048.48 | 1249.65 | -0.25 | 2.35E-01 |
| ENSMUSG00000032534 | GSMG0041216 | Cep63 | centrosomal protein 63 | 579.01 | 640.90 | -0.15 | 5.76E-01 |
| ENSMUSG00000032553 | GSMG0041219 | Srprb | signal recognition particle receptor, B subunit | 1510.82 | 1539.60 | -0.03 | 9.38E-01 |
| ENSMUSG00000032554 | GSMG0041221 | Trf | transferrin | 3850.40 | 9017.20 | -1.23 | 5.69E-20 |
| ENSMUSG00000032803 | GSMG0041223 | Cdv3 | carnitine deficiency-associated gene expressed in ventricle 3 | 1211.23 | 1596.46 | -0.40 | 4.40E-02 |
| NULL | GSMG0041229 | NULL | NULL | 70.02 | 104.05 | -0.56 | 1.44E-01 |
| ENSMUSG00000032557 | GSMG0041232 | Uba5 | ubiquitin-like modifier activating enzyme 5 | 1289.84 | 1398.04 | -0.12 | 6.71E-01 |
| ENSMUSG00000032565 | GSMG0041239 | Nudt16 | nudix (nucleoside diphosphate linked moiety X)-type motif 16 | 723.58 | 1001.68 | -0.47 | 4.62E-03 |
| ENSMUSG00000032570 | GSMG0041243 | Atp2c1 | ATPase, Ca++-sequestering | 2856.51 | 3076.33 | -0.11 | 6.54E-01 |
| ENSMUSG00000020253 | GSMG0041249 | Ppm1m | protein phosphatase 1M | 425.53 | 319.59 | 0.42 | 6.28E-02 |
| ENSMUSG00000032786 | GSMG0041250 | Alas1 | aminolevulinic acid synthase 1 | 5785.52 | 4068.84 | 0.51 | 1.05E-03 |
| ENSMUSG00000023262 | GSMG0041252 | Acy1 | aminoacylase 1 | 255.49 | 237.01 | 0.11 | 7.70E-01 |
| ENSMUSG00000042210 | GSMG0041253 | Abhd14a | abhydrolase domain containing 14A | 250.41 | 302.23 | -0.27 | 3.43E-01 |
| NULL | GSMG0041255 | NULL | NULL | 189.71 | 370.68 | -0.97 | 7.94E-06 |
| ENSMUSG00000023249 | GSMG0041258 | Parp3 | poly (ADP-ribose) polymerase family, member 3 | 337.88 | 241.20 | 0.49 | 2.79E-02 |
| ENSMUSG00000023192 | GSMG0041261 | Grm2 | glutamate receptor, metabotropic 2 | 499.62 | 450.85 | 0.15 | 6.78E-01 |
| ENSMUSG00000040813 | GSMG0041262 | Tex264 | testis expressed gene 264 | 1775.34 | 1589.42 | 0.16 | 4.57E-01 |
| NULL | GSMG0041268 | NULL | NULL | 914.37 | 961.41 | -0.07 | 8.58E-01 |
| ENSMUSG00000032575 | GSMG0041269 | Manf | mesencephalic astrocyte-derived neurotrophic factor | 1481.18 | 2076.52 | -0.49 | 2.74E-02 |
| ENSMUSG00000039716 | GSMG0041270 | Dock3 | dedicator of cyto-kinesis 3 | 220.54 | 348.07 | -0.66 | 1.59E-02 |
| ENSMUSG00000032579 | GSMG0041273 | Hemk1 | HemK methyltransferase family member 1 | 377.66 | 377.78 | 0.00 | 9.98E-01 |
| ENSMUSG00000037190 | GSMG0041276 | Cyb561d2 | cytochrome b-561 domain containing 2 | 351.09 | 426.63 | -0.28 | 3.11E-01 |
| NULL | GSMG0041277 | NULL | NULL | 969.39 | 753.16 | 0.36 | 4.53E-02 |
| ENSMUSG00000032562 | GSMG0041279 | Gnai2 | guanine nucleotide binding protein (G protein), alpha inhibiting 2 | 28784.01 | 32776.55 | -0.19 | 2.75E-01 |
| ENSMUSG00000010064 | GSMG0041281 | Slc38a3 | solute carrier family 38, member 3 | 11868.24 | 11390.56 | 0.06 | 8.34E-01 |

| | | | | | | | |
|---|---|---|---|---|---|---|---|
| ENSMUSG00000034684 | GSMG0041283 | Sema3f | sema domain, immunoglobulin domain (Ig), short basic domain, secrete | 411.42 | 459.46 | -0.16 | 6.97E-01 |
| ENSMUSG00000032580 | GSMG0041284 | Rbm5 | RNA binding motif protein 5 | 12291.04 | 12567.59 | -0.03 | 9.03E-01 |
| ENSMUSG00000032582 | GSMG0041285 | Rbm6 | RNA binding motif protein 6 | 2370.68 | 2556.86 | -0.11 | 6.76E-01 |
| ENSMUSG00000042106 | GSMG0041288 | Fam212a | family with sequence similarity 212, member A | 1041.53 | 1293.02 | -0.31 | 9.09E-02 |
| ENSMUSG00000041528 | GSMG0041290 | Rnf123 | ring finger protein 123 | 3519.88 | 3308.03 | 0.09 | 7.67E-01 |
| ENSMUSG00000032590 | GSMG0041291 | Apeh | acylpeptide hydrolase | 799.96 | 818.35 | -0.03 | 9.29E-01 |
| ENSMUSG00000039952 | GSMG0041295 | Dag1 | dystroglycan 1 | 8069.46 | 8148.77 | -0.01 | 9.59E-01 |
| ENSMUSG00000039461 | GSMG0041297 | Tcta | T cell leukemia translocation altered gene | 387.02 | 318.64 | 0.28 | 2.75E-01 |
| ENSMUSG00000032609 | GSMG0041299 | Klhdc8b | kelch domain containing 8B | 2143.20 | 1456.54 | 0.56 | 1.99E-03 |
| NULL | GSMG0041302 | NULL | NULL | 184.46 | 189.64 | -0.03 | 9.46E-01 |
| ENSMUSG00000070283 | GSMG0041303 | Ndufaf3 | NADH dehydrogenase (ubiquinone) 1 alpha subcomplex, assembly fact | 868.26 | 885.01 | -0.03 | 9.28E-01 |
| ENSMUSG00000066357 | GSMG0041304 | Wdr6 | WD repeat domain 6 | 2685.01 | 3644.81 | -0.44 | 3.34E-02 |
| ENSMUSG00000006675 | GSMG0041305 | P4htm | prolyl 4-hydroxylase, transmembrane (endoplasmic reticulum) | 508.83 | 348.98 | 0.54 | 1.69E-02 |
| ENSMUSG00000064145 | GSMG0041306 | Arih2 | ariadne RBR E3 ubiquitin protein ligase 2 | 715.94 | 759.26 | -0.08 | 8.18E-01 |
| NULL | GSMG0041309 | NULL | NULL | 84.80 | 90.35 | -0.09 | 8.65E-01 |
| G00000025646,ENSMUSG000 | GSMG0041312 | Trex1 // Atrip | three prime repair exonuclease 1 // ATR interacting protein | 1273.67 | 1346.38 | -0.08 | 7.56E-01 |
| ENSMUSG00000032480 | GSMG0041331 | Dhx30 | DEAH (Asp-Glu-Ala-His) box polypeptide 30 | 3245.19 | 3433.52 | -0.08 | 7.79E-01 |
| ENSMUSG00000036057 | GSMG0041336 | Ptpn23 | protein tyrosine phosphatase, non-receptor type 23 | 188.31 | 156.77 | 0.26 | 4.71E-01 |
| ENSMUSG00000054792 | GSMG0041338 | Klhl18 | kelch-like 18 | 119.76 | 141.38 | -0.24 | 5.53E-01 |
| ENSMUSG00000056724 | GSMG0041341 | Nbeal2 | neurobeachin-like 2 | 367.79 | 326.26 | 0.17 | 7.42E-01 |
| ENSMUSG00000032492 | GSMG0041343 | Pth1r | parathyroid hormone 1 receptor | 1963.60 | 1345.49 | 0.55 | 2.05E-03 |
| ENSMUSG00000043953 | GSMG0041349 | Ccrl2 | chemokine (C-C motif) receptor-like 2 | 780.71 | 474.17 | 0.72 | 9.10E-04 |
| ENSMUSG00000032498 | GSMG0041350 | Mlh1 | mutL homolog 1 | 649.89 | 795.27 | -0.29 | 2.17E-01 |
| NULL | GSMG0041357 | NULL | NULL | 312.94 | 299.06 | 0.07 | 8.66E-01 |
| NULL | GSMG0041358 | NULL | NULL | 1849.61 | 1699.73 | 0.12 | 6.61E-01 |
| NULL | GSMG0041359 | NULL | NULL | 1976.77 | 3501.38 | -0.83 | 3.00E-05 |
| NULL | GSMG0041360 | NULL | NULL | 97.29 | 119.12 | -0.28 | 5.31E-01 |
| ENSMUSG00000032503 | GSMG0041361 | Arpp21 | cyclic AMP-regulated phosphoprotein, 21 | 5740.24 | 4878.40 | 0.23 | 1.74E-01 |
| NULL | GSMG0041362 | NULL | NULL | 246.52 | 399.80 | -0.69 | 4.32E-03 |
| NULL | GSMG0041363 | NULL | NULL | 157.30 | 229.50 | -0.55 | 5.42E-02 |
| NULL | GSMG0041364 | NULL | NULL | 1552.41 | 1910.29 | -0.30 | 1.15E-01 |
| NULL | GSMG0041365 | NULL | NULL | 550.91 | 865.91 | -0.65 | 1.50E-04 |
| ENSMUSG00000032504 | GSMG0041371 | Pdcd6ip | programmed cell death 6 interacting protein | 2907.23 | 3296.81 | -0.18 | 3.40E-01 |

| | | | | | | | |
|---|---|---|---|---|---|---|---|
| ENSMUSG00000032507 | GSMG0041373 | Fbxl2 | F-box and leucine-rich repeat protein 2 | 825.62 | 767.07 | 0.11 | 6.80E-01 |
| ENSMUSG00000032431 | GSMG0041378 | Crtap | cartilage associated protein | 1764.45 | 1377.41 | 0.36 | 4.45E-02 |
| ENSMUSG00000056167 | GSMG0041382 | Cnot10 | CCR4-NOT transcription complex, subunit 10 | 3347.93 | 2919.40 | 0.20 | 2.93E-01 |
| NULL | GSMG0041383 | NULL | NULL | 143.69 | 165.65 | -0.21 | 5.84E-01 |
| ENSMUSG00000032436 | GSMG0041385 | Cmtm7 | CKLF-like MARVEL transmembrane domain containing 7 | 258.89 | 315.21 | -0.28 | 3.26E-01 |
| ENSMUSG00000041012 | GSMG0041386 | Cmtm8 | CKLF-like MARVEL transmembrane domain containing 8 | 697.43 | 550.84 | 0.34 | 1.31E-01 |
| ENSMUSG00000050627 | GSMG0041387 | Gpd1l | glycerol-3-phosphate dehydrogenase 1-like | 524.39 | 565.95 | -0.11 | 6.95E-01 |
| ENSMUSG00000032437 | GSMG0041389 | Stt3b | STT3, subunit of the oligosaccharyltransferase complex, homolog B (S. | 3756.53 | 4591.64 | -0.29 | 1.07E-01 |
| ENSMUSG00000032440 | GSMG0041396 | Tgfbr2 | transforming growth factor, beta receptor II | 4857.00 | 4253.15 | 0.19 | 3.21E-01 |
| ENSMUSG00000039163 | GSMG0041423 | Cmc1 | COX assembly mitochondrial protein 1 | 265.08 | 297.75 | -0.17 | 6.99E-01 |
| ENSMUSG00000010660 | GSMG0041430 | Plcd1 | phospholipase C, delta 1 | 647.43 | 1014.00 | -0.65 | 6.18E-04 |
| ENSMUSG00000036737 | GSMG0041435 | Oxsr1 | oxidative-stress responsive 1 | 570.48 | 647.66 | -0.18 | 4.33E-01 |
| ENSMUSG00000032508 | GSMG0041437 | Myd88 | myeloid differentiation primary response gene 88 | 340.24 | 242.23 | 0.49 | 2.66E-02 |
| ENSMUSG00000032513 | GSMG0041443 | Gorasp1 | golgi reassembly stacking protein 1 | 271.03 | 286.23 | -0.07 | 8.58E-01 |
| ENSMUSG00000052336 | GSMG0041447 | Cx3cr1 | chemokine (C-X3-C motif) receptor 1 | 640.04 | 292.41 | 1.13 | 3.10E-06 |
| ENSMUSG00000040936 | GSMG0041455 | Ulk4 | unc-51-like kinase 4 | 197.95 | 100.63 | 0.98 | 1.66E-03 |
| ENSMUSG00000032532 | GSMG0041458 | Cck | cholecystokinin | 597.90 | 306.12 | 0.96 | 2.76E-06 |
| ENSMUSG00000038412 | GSMG0041459 | Higd1a | HIG1 domain family, member 1A | 1263.00 | 1085.81 | 0.22 | 2.92E-01 |
| ENSMUSG00000032523 | GSMG0041467 | Hhatl | hedgehog acyltransferase-like | 144.30 | 20.83 | 2.80 | 2.16E-13 |
| ENSMUSG00000066235 | GSMG0041473 | Pomgnt2 | protein O-linked mannose beta 1,4-N-acetylglucosaminyltransferase 2 | 515.58 | 604.75 | -0.23 | 3.60E-01 |
| ENSMUSG00000037949 | GSMG0041475 | Ano10 | anoctamin 10 | 1049.36 | 1030.74 | 0.03 | 9.32E-01 |
| ENSMUSG00000047036 | GSMG0041481 | Zfp445 | zinc finger protein 445 | 5088.44 | 4794.53 | 0.09 | 7.16E-01 |
| ENSMUSG00000032551 | GSMG0041483 | 1110059G10Rik | RIKEN cDNA 1110059G10 gene | 547.51 | 585.02 | -0.10 | 7.76E-01 |
| NULL | GSMG0041485 | NULL | NULL | 434.49 | 418.21 | 0.05 | 8.84E-01 |
| ENSMUSG00000025786 | GSMG0041486 | Zdhhc3 | zinc finger, DHHC domain containing 3 | 1127.61 | 1386.89 | -0.30 | 9.32E-02 |
| ENSMUSG00000054871 | GSMG0041488 | Tmem158 | transmembrane protein 158 | 1176.32 | 2777.98 | -1.24 | 7.77E-14 |
| ENSMUSG00000036814 | GSMG0041491 | Slc6a20a | solute carrier family 6 (neurotransmitter transporter), member 20A | 4062.36 | 1978.47 | 1.04 | 4.58E-09 |
| ENSMUSG00000025245 | GSMG0041493 | Lztfl1 | leucine zipper transcription factor-like 1 | 303.37 | 259.49 | 0.22 | 5.02E-01 |
| NULL | GSMG0041499 | 4930526I15Rik | RIKEN cDNA 4930526I15 gene | 185.92 | 239.20 | -0.37 | 2.68E-01 |
| ENSMUSG00000079084 | GSMG0041501 | Ccdc82 | coiled-coil domain containing 82 | 412.27 | 384.05 | 0.10 | 7.87E-01 |
| NULL | GSMG0041505 | NULL | NULL | 254.00 | 192.62 | 0.40 | 1.33E-01 |
| ENSMUSG00000074829 | GSMG0041507 | 2010315B03Rik | RIKEN cDNA 2010315B03 gene | 395.34 | 300.41 | 0.40 | 6.67E-02 |
| NULL | GSMG0041510 | NULL | NULL | 1062.28 | 732.37 | 0.54 | 8.23E-02 |

| | | | | | | | |
|---|---|---|---|---|---|---|---|
| ENSMUSG00000079083 | GSMG0041511 | Jrkl | Jrk-like | 310.33 | 276.75 | 0.17 | 6.62E-01 |
| NULL | GSMG0041512 | NULL | NULL | 36396.46 | 31524.21 | 0.21 | 5.24E-01 |
| NULL | GSMG0041513 | NULL | NULL | 5005.78 | 3874.83 | 0.37 | 1.06E-01 |
| NULL | GSMG0041517 | NULL | NULL | 413.36 | 275.30 | 0.59 | 5.27E-01 |
| NULL | GSMG0041518 | NULL | NULL | 34639.72 | 30711.61 | 0.17 | 5.17E-01 |
| NULL | GSMG0041519 | NULL | NULL | 373.63 | 252.75 | 0.57 | 1.55E-02 |
| NULL | GSMG0041520 | NULL | NULL | 1506.31 | 1344.21 | 0.17 | 6.44E-01 |
| NULL | GSMG0041522 | NULL | NULL | 1427.79 | 1340.81 | 0.09 | 7.58E-01 |
| ENSMUSG00000079685 | GSMG0041529 | Ulbp1 | UL16 binding protein 1 | 26.27 | 56.37 | -1.09 | 1.88E-02 |
| ENSMUSG00000068270 | GSMG0041549 | Shroom4 | shroom family member 4 | 1523.04 | 1362.52 | 0.16 | 5.86E-01 |
| ENSMUSG00000031144 | GSMG0041557 | Syp | synaptophysin | 1472.63 | 1362.35 | 0.11 | 6.83E-01 |
| ENSMUSG00000031145 | GSMG0041558 | Prickle3 | prickle planar cell polarity protein 3 | 252.05 | 335.82 | -0.41 | 1.08E-01 |
| ENSMUSG00000031148 | GSMG0041559 | Gpkow | G patch domain and KOW motifs | 1604.90 | 1733.50 | -0.11 | 6.53E-01 |
| ENSMUSG00000039382 | GSMG0041560 | Wdr45 | WD repeat domain 45 | 752.71 | 637.70 | 0.24 | 3.07E-01 |
| ENSMUSG00000031149 | GSMG0041561 | Praf2 | PRA1 domain family 2 | 178.18 | 233.15 | -0.39 | 1.59E-01 |
| ENSMUSG00000000134 | GSMG0041562 | Tfe3 | transcription factor E3 | 912.38 | 838.06 | 0.12 | 6.29E-01 |
| ENSMUSG00000031153 | GSMG0041563 | Gripap1 | GRIP1 associated protein 1 | 2655.20 | 2811.88 | -0.08 | 7.24E-01 |
| ENSMUSG00000009731 | GSMG0041565 | Kcnd1 | potassium voltage-gated channel, Shal-related family, member 1 | 40.08 | 69.79 | -0.79 | 1.27E-01 |
| ENSMUSG00000031154 | GSMG0041566 | Otud5 | OTU domain containing 5 | 1309.88 | 1097.58 | 0.26 | 3.48E-01 |
| ENSMUSG00000031155 | GSMG0041567 | Pim2 | proviral integration site 2 | 254.48 | 159.69 | 0.67 | 1.39E-02 |
| ENSMUSG00000031156 | GSMG0041568 | Slc35a2 | solute carrier family 35 (UDP-galactose transporter), member A2 | 793.13 | 734.60 | 0.11 | 6.81E-01 |
| ENSMUSG00000031158 | GSMG0041569 | Timm17b | translocase of inner mitochondrial membrane 17b | 739.02 | 943.09 | -0.35 | 9.84E-02 |
| ENSMUSG00000039278 | GSMG0041570 | Pcsk1n | proprotein convertase subtilisin/kexin type 1 inhibitor | 2648.45 | 2518.90 | 0.07 | 8.60E-01 |
| ENSMUSG00000055188 | GSMG0041572 | Rbm3os | RNA binding motif protein 3, opposite strand | 62.17 | 88.26 | -0.51 | 2.04E-01 |
| ENSMUSG00000031170 | GSMG0041575 | Slc38a5 | solute carrier family 38, member 5 | 14266.52 | 25444.55 | -0.83 | 2.16E-06 |
| ENSMUSG00000052364 | GSMG0041581 | B630019K06Rik | RIKEN cDNA B630019K06 gene | 246.78 | 317.31 | -0.37 | 2.17E-01 |
| ENSMUSG00000047344 | GSMG0041585 | Lancl3 | LanC lantibiotic synthetase component C-like 3 (bacterial) | 52.75 | 43.09 | 0.31 | 6.35E-01 |
| ENSMUSG00000058254 | GSMG0041595 | Tspan7 | tetraspanin 7 | 4448.09 | 3951.87 | 0.17 | 3.81E-01 |
| NULL | GSMG0041596 | NULL | NULL | 169.31 | 95.26 | 0.83 | 2.29E-03 |
| ENSMUSG00000008035 | GSMG0041597 | Mid1ip1 | Mid1 interacting protein 1 (gastrulation specific G12-like (zebrafish)) | 1603.32 | 1547.43 | 0.05 | 8.56E-01 |
| ENSMUSG00000031007 | GSMG0041599 | Atp6ap2 | ATPase, H+ transporting, lysosomal accessory protein 2 | 1586.58 | 1744.73 | -0.14 | 5.76E-01 |
| ENSMUSG00000031010 | GSMG0041605 | Usp9x | ubiquitin specific peptidase 9, X chromosome | 745.38 | 860.53 | -0.21 | 3.90E-01 |
| ENSMUSG00000000787 | GSMG0041607 | Ddx3x | DEAD/H (Asp-Glu-Ala-Asp/His) box polypeptide 3, X-linked | 2342.60 | 2377.46 | -0.02 | 9.52E-01 |

| | | | | | | | |
|---|---|---|---|---|---|---|---|
| ENSMUSG00000040229 | GSMG0041609 | Gpr34 | G protein-coupled receptor 34 | 301.19 | 160.47 | 0.91 | 1.54E-04 |
| ENSMUSG00000025037 | GSMG0041612 | Maoa | monoamine oxidase A | 5460.05 | 4943.00 | 0.14 | 5.94E-01 |
| ENSMUSG00000037369 | GSMG0041616 | Kdm6a | lysine (K)-specific demethylase 6A | 1472.09 | 1527.78 | -0.05 | 8.32E-01 |
| ENSMUSG00000037347 | GSMG0041621 | Chst7 | carbohydrate (N-acetylglucosamino) sulfotransferase 7 | 106.98 | 176.49 | -0.72 | 8.68E-03 |
| ENSMUSG00000060090 | GSMG0041622 | Rp2 | retinitis pigmentosa 2 homolog | 246.61 | 259.26 | -0.07 | 8.68E-01 |
| ENSMUSG00000037315 | GSMG0041624 | Jade3 | jade family PHD finger 3 | 176.17 | 214.20 | -0.29 | 3.54E-01 |
| ENSMUSG00000031060 | GSMG0041626 | Rbm10 | RNA binding motif protein 10 | 1129.75 | 969.33 | 0.22 | 2.98E-01 |
| ENSMUSG00000001924 | GSMG0041628 | Uba1 | ubiquitin-like modifier activating enzyme 1 | 5957.38 | 8201.43 | -0.46 | 1.55E-03 |
| ENSMUSG00000031065 | GSMG0041629 | Cdk16 | cyclin-dependent kinase 16 | 5299.57 | 4987.79 | 0.09 | 7.67E-01 |
| ENSMUSG00000031066 | GSMG0041630 | Usp11 | ubiquitin specific peptidase 11 | 1395.50 | 1241.32 | 0.17 | 5.15E-01 |
| ENSMUSG00000001127 | GSMG0041631 | Araf | Araf proto-oncogene, serine/threonine kinase | 1607.57 | 1713.95 | -0.09 | 7.11E-01 |
| ENSMUSG00000001131 | GSMG0041633 | Timp1 | tissue inhibitor of metalloproteinase 1 | 168.73 | 290.31 | -0.78 | 2.50E-03 |
| ENSMUSG00000086877 | GSMG0041635 | A230072C01Rik | RIKEN cDNA A230072C01 gene | 238.32 | 215.86 | 0.14 | 7.51E-01 |
| ENSMUSG00000036769 | GSMG0041641 | Wdr44 | WD repeat domain 44 | 403.81 | 360.36 | 0.17 | 5.93E-01 |
| ENSMUSG00000031093 | GSMG0041658 | Dock11 | dedicator of cytokinesis 11 | 220.52 | 240.46 | -0.13 | 7.13E-01 |
| ENSMUSG00000017057 | GSMG0041661 | Il13ra1 | interleukin 13 receptor, alpha 1 | 878.21 | 722.48 | 0.28 | 1.51E-01 |
| ENSMUSG00000036699 | GSMG0041663 | Zcchc12 | zinc finger, CCHC domain containing 12 | 327.09 | 602.95 | -0.88 | 3.52E-03 |
| ENSMUSG00000006373 | GSMG0041666 | Pgrmc1 | progesterone receptor membrane component 1 | 5616.46 | 6840.72 | -0.28 | 1.21E-01 |
| ENSMUSG00000016319 | GSMG0041668 | Slc25a5 | solute carrier family 25 (mitochondrial carrier, adenine nucleotide trans | 4143.44 | 5119.72 | -0.30 | 9.07E-02 |
| ENSMUSG00000016308 | GSMG0041669 | Ube2a | ubiquitin-conjugating enzyme E2A | 1298.95 | 1474.50 | -0.18 | 4.50E-01 |
| ENSMUSG00000016409 | GSMG0041672 | Nkap | NFKB activating protein | 258.93 | 328.48 | -0.35 | 2.60E-01 |
| ENSMUSG00000036537 | GSMG0041674 | Rnf113a1 | ring finger protein 113A1 | 224.44 | 232.67 | -0.05 | 9.06E-01 |
| ENSMUSG00000000355 | GSMG0041695 | Mcts1 | malignant T cell amplified sequence 1 | 643.24 | 760.79 | -0.24 | 3.02E-01 |
| ENSMUSG00000001986 | GSMG0041699 | Gria3 | glutamate receptor, ionotropic, AMPA3 (alpha 3) | 704.98 | 368.08 | 0.94 | 6.16E-07 |
| NULL | GSMG0041707 | NULL | NULL | 745.27 | 210.67 | 1.82 | 6.01E-21 |
| ENSMUSG00000025860 | GSMG0041708 | Xiap | X-linked inhibitor of apoptosis | 254.12 | 346.05 | -0.44 | 4.35E-02 |
| NULL | GSMG0041709 | NULL | NULL | 476.05 | 420.79 | 0.17 | 7.08E-01 |
| ENSMUSG00000025862 | GSMG0041710 | Stag2 | stromal antigen 2 | 918.15 | 957.78 | -0.06 | 8.63E-01 |
| ENSMUSG00000001173 | GSMG0041723 | Ocrl | oculocerebrorenal syndrome of Lowe | 1411.74 | 1183.48 | 0.25 | 1.94E-01 |
| ENSMUSG00000063785 | GSMG0041726 | Utp14a | UTP14A small subunit processome component | 191.97 | 237.70 | -0.31 | 3.57E-01 |
| ENSMUSG00000031104 | GSMG0041732 | Rab33a | RAB33A, member RAS oncogene family | 227.12 | 368.26 | -0.70 | 3.76E-03 |
| ENSMUSG00000031105 | GSMG0041733 | Slc25a14 | solute carrier family 25 (mitochondrial carrier, brain), member 14 | 306.96 | 325.03 | -0.08 | 8.01E-01 |
| ENSMUSG00000031107 | GSMG0041734 | Rbmx2 | RNA binding motif protein, X-linked 2 | 182.07 | 225.55 | -0.31 | 3.30E-01 |

| | | | | | | | |
|---|---|---|---|---|---|---|---|
| ENSMUSG00000025626 | GSMG0041749 | Phf6 | PHD finger protein 6 | 336.83 | 474.02 | -0.49 | 1.77E-02 |
| ENSMUSG00000025630 | GSMG0041750 | Hprt | hypoxanthine guanine phosphoribosyl transferase | 3068.19 | 3038.05 | 0.01 | 9.65E-01 |
| ENSMUSG00000073176 | GSMG0041760 | Zfp449 | zinc finger protein 449 | 85.03 | 94.55 | -0.15 | 7.95E-01 |
| ENSMUSG00000054850 | GSMG0041761 | Smim10l2a | small integral membrane protein 10 like 2A | 252.24 | 361.43 | -0.52 | 6.93E-02 |
| ENSMUSG00000035967 | GSMG0041763 | Ddx26b | DEAD/H (Asp-Glu-Ala-Asp/His) box polypeptide 26B | 859.65 | 795.27 | 0.11 | 6.56E-01 |
| ENSMUSG00000060681 | GSMG0041765 | Slc9a6 | solute carrier family 9 (sodium/hydrogen exchanger), member 6 | 374.20 | 549.84 | -0.56 | 6.79E-03 |
| ENSMUSG00000023092 | GSMG0041766 | Fhl1 | four and a half LIM domains 1 | 993.25 | 1124.34 | -0.18 | 5.29E-01 |
| ENSMUSG00000067873 | GSMG0041770 | Htatsf1 | HIV TAT specific factor 1 | 1144.10 | 1233.04 | -0.11 | 6.39E-01 |
| ENSMUSG00000067860 | GSMG0041775 | Zic3 | zinc finger protein of the cerebellum 3 | 4349.68 | 4760.74 | -0.13 | 5.39E-01 |
| NULL | GSMG0041784 | NULL | NULL | 521.68 | 257.55 | 1.02 | 4.96E-07 |
| ENSMUSG00000000838 | GSMG0041799 | Fmr1 | fragile X mental retardation syndrome 1 | 556.91 | 717.44 | -0.37 | 9.05E-02 |
| ENSMUSG00000045237 | GSMG0041808 | 1110012L19Rik | RIKEN cDNA 1110012L19 gene | 133.20 | 123.05 | 0.12 | 8.18E-01 |
| ENSMUSG00000059401 | GSMG0041809 | Mamld1 | mastermind-like domain containing 1 | 759.24 | 944.88 | -0.31 | 1.06E-01 |
| ENSMUSG00000031337 | GSMG0041813 | Mtm1 | X-linked myotubular myopathy gene 1 | 163.62 | 145.19 | 0.17 | 6.92E-01 |
| ENSMUSG00000015214 | GSMG0041814 | Mtmr1 | myotubularin related protein 1 | 435.52 | 486.78 | -0.16 | 5.67E-01 |
| ENSMUSG00000073131 | GSMG0041818 | Vma21 | VMA21 vacuolar H+-ATPase homolog (S. cerevisiae) | 1093.34 | 1057.02 | 0.05 | 9.08E-01 |
| ENSMUSG00000033361 | GSMG0041820 | Prrg3 | proline rich Gla (G-carboxyglutamic acid) 3 (transmembrane) | 476.29 | 924.93 | -0.96 | 3.82E-09 |
| ENSMUSG00000031349 | GSMG0041826 | Nsdhl | NAD(P) dependent steroid dehydrogenase-like | 279.00 | 415.56 | -0.58 | 8.54E-03 |
| ENSMUSG00000078317 | GSMG0041833 | F8a | factor 8-associated gene A | 359.33 | 423.19 | -0.24 | 3.99E-01 |
| ENSMUSG00000031365 | GSMG0041834 | Zfp275 | zinc finger protein 275 | 943.10 | 976.87 | -0.05 | 8.63E-01 |
| ENSMUSG00000031375 | GSMG0041837 | Bgn | biglycan | 3468.93 | 5242.74 | -0.60 | 1.90E-04 |
| ENSMUSG00000031376 | GSMG0041838 | Atp2b3 | ATPase, Ca++ transporting, plasma membrane 3 | 676.34 | 462.23 | 0.55 | 1.14E-02 |
| ENSMUSG00000019558 | GSMG0041842 | Slc6a8 | solute carrier family 6 (neurotransmitter transporter, creatine), member | 642.31 | 399.13 | 0.69 | 3.21E-04 |
| ENSMUSG00000031378 | GSMG0041843 | Abcd1 | ATP-binding cassette, sub-family D (ALD), member 1 | 389.77 | 254.37 | 0.62 | 9.23E-03 |
| ENSMUSG00000002007 | GSMG0041846 | Srpk3 | serine/arginine-rich protein specific kinase 3 | 204.68 | 49.09 | 2.07 | 5.19E-12 |
| ENSMUSG00000002014 | GSMG0041847 | Ssr4 | signal sequence receptor, delta | 1672.47 | 1657.93 | 0.01 | 9.64E-01 |
| ENSMUSG00000009995 | GSMG0041854 | Taz | tafazzin | 1279.06 | 1272.60 | 0.01 | 9.79E-01 |
| ENSMUSG00000019087 | GSMG0041856 | Atp6ap1 | ATPase, H+ transporting, lysosomal accessory protein 1 | 4091.40 | 4076.20 | 0.01 | 9.86E-01 |
| ENSMUSG00000015291 | GSMG0041857 | Gdi1 | guanosine diphosphate (GDP) dissociation inhibitor 1 | 6685.69 | 6520.63 | 0.04 | 9.01E-01 |
| ENSMUSG00000001962 | GSMG0041858 | Fam50a | family with sequence similarity 50, member A | 1203.95 | 1078.67 | 0.16 | 5.32E-01 |
| ENSMUSG00000031398 | GSMG0041859 | Plxna3 | plexin A3 | 556.46 | 1650.97 | -1.57 | 1.00E-13 |
| ENSMUSG00000004221 | GSMG0041861 | Ikbkg | inhibitor of kappaB kinase gamma | 455.93 | 380.25 | 0.26 | 2.80E-01 |
| ENSMUSG00000031403 | GSMG0041862 | Dkc1 | dyskeratosis congenita 1, dyskerin | 615.97 | 855.65 | -0.47 | 8.35E-03 |

| | | | | | | | |
|---|---|---|---|---|---|---|---|
| ENSMUSG00000031198 | GSMG0041863 | Fundc2 | FUN14 domain containing 2 | 598.33 | 636.47 | -0.09 | 7.89E-01 |
| ENSMUSG00000031201 | GSMG0041864 | Brcc3 | BRCA1/BRCA2-containing complex, subunit 3 | 363.47 | 421.52 | -0.21 | 4.52E-01 |
| ENSMUSG00000031197 | GSMG0041865 | Vbp1 | von Hippel-Lindau binding protein 1 | 1714.76 | 1956.51 | -0.19 | 4.70E-01 |
| ENSMUSG00000025246 | GSMG0041872 | Tbl1x | transducin (beta)-like 1 X-linked | 1017.58 | 1165.01 | -0.19 | 4.55E-01 |
| ENSMUSG00000025666 | GSMG0041885 | Tmem47 | transmembrane protein 47 | 1914.21 | 1325.94 | 0.53 | 3.75E-03 |
| NULL | GSMG0041890 | NULL | NULL | 172.78 | 74.03 | 1.22 | 1.41E-05 |
| ENSMUSG00000045103 | GSMG0041898 | Dmd | dystrophin, muscular dystrophy | 2335.75 | 1817.46 | 0.36 | 4.90E-02 |
| ENSMUSG00000035277 | GSMG0041912 | Arx | aristaless related homeobox | 163.94 | 346.04 | -1.08 | 3.23E-07 |
| ENSMUSG00000035246 | GSMG0041914 | Pcyt1b | phosphate cytidylyltransferase 1, choline, beta isoform | 269.97 | 557.69 | -1.05 | 2.98E-09 |
| NULL | GSMG0041922 | NULL | NULL | 391.98 | 324.05 | 0.27 | 2.71E-01 |
| ENSMUSG00000031207 | GSMG0041931 | Msn | moesin | 19945.20 | 21070.91 | -0.08 | 7.24E-01 |
| ENSMUSG00000031209 | GSMG0041933 | Heph | hephaestin | 373.39 | 496.22 | -0.41 | 1.08E-01 |
| ENSMUSG00000047694 | GSMG0041940 | Yipf6 | Yip1 domain family, member 6 | 510.90 | 526.77 | -0.04 | 9.01E-01 |
| ENSMUSG00000031216 | GSMG0041941 | Stard8 | START domain containing 8 | 849.03 | 670.28 | 0.34 | 1.33E-01 |
| ENSMUSG00000031217 | GSMG0041942 | Efnb1 | ephrin B1 | 220.00 | 264.43 | -0.27 | 4.70E-01 |
| ENSMUSG00000031221 | GSMG0041949 | Igbp1 | immunoglobulin (CD79A) binding protein 1 | 283.76 | 312.40 | -0.14 | 6.79E-01 |
| ENSMUSG00000034311 | GSMG0041954 | Kif4 | kinesin family member 4 | 213.67 | 548.19 | -1.36 | 2.16E-08 |
| ENSMUSG00000019359 | GSMG0041956 | Gdpd2 | glycerophosphodiester phosphodiesterase domain containing 2 | 347.31 | 367.26 | -0.08 | 8.51E-01 |
| ENSMUSG00000000881 | GSMG0041958 | Dlg3 | discs, large homolog 3 (Drosophila) | 1005.65 | 1134.63 | -0.17 | 4.17E-01 |
| NULL | GSMG0041959 | NULL | NULL | 110.27 | 117.28 | -0.09 | 8.58E-01 |
| G00000031302,ENSMUSG000 | GSMG0041961 | Med12 // Nlgn3 | mediator complex subunit 12 // neuroligin 3 | 1610.86 | 1624.21 | -0.01 | 9.76E-01 |
| NULL | GSMG0041962 | NULL | NULL | 663.81 | 1172.28 | -0.82 | 4.24E-06 |
| ENSMUSG00000031311 | GSMG0041964 | Nono | non-POU-domain-containing, octamer binding protein | 3665.43 | 4902.87 | -0.42 | 2.48E-02 |
| ENSMUSG00000031314 | GSMG0041966 | Taf1 | TATA-box binding protein associated factor 1 | 677.13 | 784.15 | -0.21 | 3.87E-01 |
| ENSMUSG00000034160 | GSMG0041968 | Ogt | O-linked N-acetylglucosamine (GlcNAc) transferase (UDP-N-acetylglu | 4602.50 | 4464.62 | 0.04 | 8.72E-01 |
| ENSMUSG00000085715 | GSMG0041990 | Tsix | X (inactive)-specific transcript, opposite strand | 1463.66 | 893.74 | 0.71 | 5.95E-01 |
| ENSMUSG00000031226 | GSMG0042000 | Pbdc1 | polysaccharide biosynthesis domain containing 1 | 570.60 | 771.37 | -0.44 | 8.64E-02 |
| ENSMUSG00000031227 | GSMG0042002 | Magee1 | melanoma antigen, family E, 1 | 1050.79 | 1387.47 | -0.40 | 4.16E-02 |
| ENSMUSG00000031231 | GSMG0042006 | Cox7b | cytochrome c oxidase subunit VIIb | 1443.11 | 2093.26 | -0.54 | 1.83E-03 |
| ENSMUSG00000033792 | GSMG0042007 | Atp7a | ATPase, Cu++ transporting, alpha polypeptide | 283.60 | 350.62 | -0.30 | 2.79E-01 |
| ENSMUSG00000049929 | GSMG0042015 | Lpar4 | lysophosphatidic acid receptor 4 | 527.50 | 620.27 | -0.23 | 3.69E-01 |
| ENSMUSG00000031246 | GSMG0042026 | Sh3bgrl | SH3-binding domain glutamic acid-rich protein like | 5124.00 | 4260.83 | 0.27 | 3.92E-01 |
| ENSMUSG00000025525 | GSMG0042039 | Apool | apolipoprotein O-like | 322.55 | 370.91 | -0.20 | 4.65E-01 |

| | | | | | | | |
|---|---|---|---|---|---|---|---|
| ENSMUSG00000025529 | GSMG0042042 | Zfp711 | zinc finger protein 711 | 77.57 | 163.47 | -1.07 | 1.07E-04 |
| ENSMUSG00000025597 | GSMG0042049 | Klhl4 | kelch-like 4 | 1465.08 | 1502.93 | -0.04 | 9.27E-01 |
| ENSMUSG00000034755 | GSMG0042058 | Pcdh11x | protocadherin 11 X-linked | 973.16 | 1034.44 | -0.09 | 8.05E-01 |
| ENSMUSG00000034480 | GSMG0042074 | Diaph2 | diaphanous related formin 2 | 299.45 | 364.23 | -0.28 | 2.68E-01 |
| ENSMUSG00000031256 | GSMG0042089 | Cstf2 | cleavage stimulation factor, 3' pre-RNA subunit 2 | 3642.86 | 2760.11 | 0.40 | 1.21E-02 |
| ENSMUSG00000033578 | GSMG0042092 | Tmem35 | transmembrane protein 35 | 571.15 | 1473.16 | -1.36 | 1.85E-15 |
| ENSMUSG00000031262 | GSMG0042093 | Cenpi | centromere protein I | 108.20 | 254.86 | -1.23 | 2.36E-04 |
| ENSMUSG00000000223 | GSMG0042094 | Drp2 | dystrophin related protein 2 | 497.01 | 402.13 | 0.31 | 2.46E-01 |
| ENSMUSG00000045427 | GSMG0042097 | Hnrnph2 | heterogeneous nuclear ribonucleoprotein H2 | 3819.77 | 3769.96 | 0.02 | 9.47E-01 |
| ENSMUSG00000049804 | GSMG0042098 | Armcx4 | armadillo repeat containing, X-linked 4 | 362.34 | 586.26 | -0.69 | 1.40E-04 |
| ENSMUSG00000033460 | GSMG0042100 | Armcx1 | armadillo repeat containing, X-linked 1 | 979.17 | 1015.99 | -0.05 | 8.54E-01 |
| NULL | GSMG0042106 | NULL | NULL | 589.36 | 525.99 | 0.16 | 5.35E-01 |
| NULL | GSMG0042108 | NULL | NULL | 235.08 | 127.31 | 0.89 | 1.62E-03 |
| ENSMUSG00000042712 | GSMG0042118 | Wbp5 | WW domain binding protein 5 | 1576.42 | 1806.20 | -0.20 | 4.75E-01 |
| ENSMUSG00000046432 | GSMG0042119 | Ngfrap1 | nerve growth factor receptor (TNFRSF16) associated protein 1 | 1659.84 | 2501.34 | -0.59 | 3.47E-04 |
| ENSMUSG00000049536 | GSMG0042123 | Tceal1 | transcription elongation factor A (SII)-like 1 | 182.03 | 201.46 | -0.14 | 7.09E-01 |
| ENSMUSG00000031425 | GSMG0042127 | Plp1 | proteolipid protein (myelin) 1 | 2913.50 | 131.01 | 4.48 | 1.03E-114 |
| ENSMUSG00000031428 | GSMG0042129 | Zcchc18 | zinc finger, CCHC domain containing 18 | 6447.07 | 6763.64 | -0.07 | 7.98E-01 |
| ENSMUSG00000042595 | GSMG0042130 | Fam199x | family with sequence similarity 199, X-linked | 129.01 | 81.75 | 0.66 | 5.07E-02 |
| ENSMUSG00000059203 | GSMG0042132 | Il1rapl2 | interleukin 1 receptor accessory protein-like 2 | 100.93 | 105.95 | -0.07 | 9.00E-01 |
| ENSMUSG00000031432 | GSMG0042152 | Prps1 | phosphoribosyl pyrophosphate synthetase 1 | 724.26 | 921.35 | -0.35 | 8.25E-02 |
| ENSMUSG00000000266 | GSMG0042153 | Mid2 | midline 2 | 99.71 | 113.03 | -0.17 | 7.21E-01 |
| ENSMUSG00000031274 | GSMG0042159 | Col4a5 | collagen, type IV, alpha 5 | 300.06 | 150.86 | 0.99 | 2.76E-04 |
| ENSMUSG00000042271 | GSMG0042160 | Nxt2 | nuclear transport factor 2-like export factor 2 | 460.99 | 440.88 | 0.06 | 8.63E-01 |
| ENSMUSG00000047045 | GSMG0042161 | Tmem164 | transmembrane protein 164 | 128.33 | 189.76 | -0.56 | 4.47E-02 |
| ENSMUSG00000031284 | GSMG0042167 | Pak3 | p21 protein (Cdc42/Rac)-activated kinase 3 | 212.31 | 528.80 | -1.32 | 8.85E-13 |
| ENSMUSG00000025270 | GSMG0042186 | Alas2 | aminolevulinic acid synthase 2, erythroid | 5440.63 | 5246.62 | 0.05 | 8.31E-01 |
| ENSMUSG00000025265 | GSMG0042189 | Fgd1 | FYVE, RhoGEF and PH domain containing 1 | 551.08 | 719.96 | -0.39 | 1.88E-01 |
| ENSMUSG00000025262 | GSMG0042193 | Fam120c | family with sequence similarity 120, member C | 184.04 | 260.36 | -0.50 | 1.27E-01 |
| ENSMUSG00000041229 | GSMG0042197 | Phf8 | PHD finger protein 8 | 339.00 | 338.31 | 0.00 | 9.92E-01 |
| ENSMUSG00000025261 | GSMG0042200 | Huwe1 | HECT, UBA and WWE domain containing 1 | 3387.59 | 3783.61 | -0.16 | 4.61E-01 |
| ENSMUSG00000025260 | GSMG0042202 | Hsd17b10 | hydroxysteroid (17-beta) dehydrogenase 10 | 632.28 | 795.11 | -0.33 | 1.35E-01 |
| ENSMUSG00000041133 | GSMG0042203 | Smc1a | structural maintenance of chromosomes 1A | 2300.67 | 2735.69 | -0.25 | 2.17E-01 |

| | | | | | | | |
|---|---|---|---|---|---|---|---|
| NULL | GSMG0042206 | NULL | NULL | 833.06 | 507.19 | 0.72 | 1.62E-03 |
| NULL | GSMG0042208 | NULL | NULL | 54.41 | 41.61 | 0.38 | 4.84E-01 |
| ENSMUSG00000025332 | GSMG0042209 | Kdm5c | lysine (K)-specific demethylase 5C | 2738.73 | 2867.70 | -0.07 | 8.60E-01 |
| ENSMUSG00000041658 | GSMG0042218 | Rragb | Ras-related GTP binding B | 280.68 | 267.66 | 0.06 | 8.87E-01 |
| ENSMUSG00000086316 | GSMG0042222 | 2210013O21Rik | RIKEN cDNA 2210013O21 gene | 927.40 | 1001.69 | -0.11 | 6.70E-01 |
| ENSMUSG00000025287 | GSMG0042225 | Acot9 | acyl-CoA thioesterase 9 | 1134.60 | 1244.62 | -0.13 | 5.97E-01 |
| ENSMUSG00000031309 | GSMG0042235 | Rps6ka3 | ribosomal protein S6 kinase polypeptide 3 | 486.42 | 574.13 | -0.24 | 4.08E-01 |
| ENSMUSG00000067194 | GSMG0042243 | Eif1ax | eukaryotic translation initiation factor 1A, X-linked | 1825.08 | 2634.24 | -0.53 | 5.33E-03 |
| ENSMUSG00000041020 | GSMG0042245 | Map7d2 | MAP7 domain containing 2 | 243.65 | 194.72 | 0.33 | 3.16E-01 |
| ENSMUSG00000044150 | GSMG0042246 | A830080D01Rik | RIKEN cDNA A830080D01 gene | 498.83 | 656.88 | -0.40 | 4.34E-02 |
| ENSMUSG00000040990 | GSMG0042249 | Sh3kbp1 | SH3-domain kinase binding protein 1 | 1050.03 | 2267.22 | -1.11 | 4.73E-14 |
| ENSMUSG00000031295 | GSMG0042255 | Phka2 | phosphorylase kinase alpha 2 | 1636.46 | 969.97 | 0.76 | 1.52E-05 |
| ENSMUSG00000031353 | GSMG0042259 | Rbbp7 | retinoblastoma binding protein 7 | 4066.86 | 5071.68 | -0.32 | 1.10E-01 |
| ENSMUSG00000031360 | GSMG0042260 | Ctps2 | cytidine 5'-triphosphate synthase 2 | 1308.41 | 1539.14 | -0.23 | 2.56E-01 |
| ENSMUSG00000031367 | GSMG0042264 | Ap1s2 | adaptor-related protein complex 1, sigma 2 subunit | 1412.07 | 1177.15 | 0.26 | 3.12E-01 |
| ENSMUSG00000015405 | GSMG0042268 | Ace2 | angiotensin I converting enzyme (peptidyl-dipeptidase A) 2 | 2194.44 | 927.85 | 1.24 | 1.49E-12 |
| ENSMUSG00000031379 | GSMG0042269 | Pir | pirin | 617.53 | 279.74 | 1.14 | 8.85E-09 |
| ENSMUSG00000031381 | GSMG0042271 | Piga | phosphatidylinositol glycan anchor biosynthesis, class A | 80.97 | 85.80 | -0.08 | 9.01E-01 |
| ENSMUSG00000031342 | GSMG0042279 | Gpm6b | glycoprotein m6b | 4805.03 | 5287.92 | -0.14 | 5.40E-01 |
| ENSMUSG00000031355 | GSMG0042285 | Arhgap6 | Rho GTPase activating protein 6 | 216.26 | 104.37 | 1.05 | 1.43E-05 |
| ENSMUSG00000087159 | GSMG0042292 | Gm15246 | predicted gene 15246 | 466.10 | 634.23 | -0.44 | 3.04E-02 |
| ENSMUSG00000035299 | GSMG0042301 | Mid1 | midline 1 | 365.02 | 839.25 | -1.20 | 5.75E-13 |
| ENSMUSG00000073293 | GSMG0042308 | Nudt10 | nudix (nucleoside diphosphate linked moiety X)-type motif 10 | 139.24 | 172.82 | -0.32 | 3.88E-01 |
| NULL | GSMG0042314 | NULL | NULL | 120.57 | 172.25 | -0.51 | 8.36E-02 |
| ENSMUSG00000004317 | GSMG0042315 | Clcn5 | chloride channel, voltage-sensitive 5 | 159.58 | 276.23 | -0.79 | 1.06E-03 |
| G00000031143,ENSMUSG000 | GSMG0042318 | Ppp1r3f // Ccdc22 | protein phosphatase 1, regulatory (inhibitor) subunit 3F // coiled-coil do | 1827.76 | 1662.51 | 0.14 | 5.77E-01 |
| NULL | GSMG0042319 | NULL | NULL | 523.72 | 444.89 | 0.24 | 3.55E-01 |
| ENSMUSG00000031147 | GSMG0042321 | Magix | MAGI family member, X-linked | 67.59 | 26.96 | 1.33 | 1.85E-03 |
| ENSMUSG00000031150 | GSMG0042322 | Ccdc120 | coiled-coil domain containing 120 | 86.72 | 157.12 | -0.86 | 3.10E-03 |
| ENSMUSG00000031157 | GSMG0042323 | Pqbp1 | polyglutamine binding protein 1 | 1013.44 | 1206.21 | -0.25 | 3.03E-01 |
| ENSMUSG00000031161 | GSMG0042325 | Hdac6 | histone deacetylase 6 | 1311.19 | 1582.94 | -0.27 | 2.49E-01 |
| ENSMUSG00000039231 | GSMG0042329 | Suv39h1 | suppressor of variegation 3-9 homolog 1 (Drosophila) | 305.08 | 487.80 | -0.67 | 5.82E-04 |
| ENSMUSG00000031166 | GSMG0042332 | Wdr13 | WD repeat domain 13 | 2035.84 | 2067.13 | -0.02 | 9.37E-01 |

| | | | | | | | |
|---|---|---|---|---|---|---|---|
| ENSMUSG00000039201 | GSMG0042334 | Tbc1d25 | TBC1 domain family, member 25 | 521.21 | 534.04 | -0.03 | 9.14E-01 |
| ENSMUSG00000031168 | GSMG0042335 | Ebp | phenylalkylamine Ca2+ antagonist (emopamil) binding protein | 756.54 | 722.38 | 0.07 | 8.32E-01 |
| ENSMUSG00000031169 | GSMG0042336 | Porcn | porcupine homolog (Drosophila) | 1169.71 | 859.55 | 0.44 | 2.11E-02 |
| ENSMUSG00000031171 | GSMG0042337 | Ftsj1 | FtsJ RNA methyltransferase homolog 1 (E. coli) | 515.30 | 581.64 | -0.17 | 4.69E-01 |
| ENSMUSG00000031176 | GSMG0042349 | Dynlt3 | dynein light chain Tctex-type 3 | 365.76 | 307.61 | 0.25 | 3.70E-01 |
| G00000031174,ENSMUSG000 | GSMG0042350 | Srpx // Rpgr | sushi-repeat-containing protein // retinitis pigmentosa GTPase regulator | 219.92 | 278.39 | -0.34 | 1.84E-01 |
| ENSMUSG00000040363 | GSMG0042356 | Bcor | BCL6 interacting corepressor | 761.07 | 851.13 | -0.16 | 5.66E-01 |
| ENSMUSG00000044148 | GSMG0042359 | 1810030O07Rik | RIKEN cDNA 1810030O07 gene | 441.22 | 463.21 | -0.07 | 8.46E-01 |
| ENSMUSG00000064127 | GSMG0042361 | Med14 | mediator complex subunit 14 | 577.85 | 583.42 | -0.01 | 9.75E-01 |
| ENSMUSG00000031012 | GSMG0042367 | Cask | calcium/calmodulin-dependent serine protein kinase (MAGUK family) | 1160.05 | 1046.80 | 0.15 | 7.09E-01 |
| ENSMUSG00000040147 | GSMG0042376 | Maob | monoamine oxidase B | 309.73 | 160.99 | 0.95 | 3.52E-04 |
| ENSMUSG00000025040 | GSMG0042382 | Fundc1 | FUN14 domain containing 1 | 697.35 | 700.48 | -0.01 | 9.86E-01 |
| ENSMUSG00000037358 | GSMG0042385 | 4930578C19Rik | RIKEN cDNA 4930578C19 gene | 341.04 | 340.52 | 0.00 | 9.95E-01 |
| ENSMUSG00000037341 | GSMG0042390 | Slc9a7 | solute carrier family 9 (sodium/hydrogen exchanger), member 7 | 82.19 | 103.74 | -0.33 | 4.34E-01 |
| ENSMUSG00000031059 | GSMG0042391 | Ndufb11 | NADH dehydrogenase (ubiquinone) 1 beta subcomplex, 11 | 2924.14 | 3233.43 | -0.15 | 5.71E-01 |
| ENSMUSG00000037217 | GSMG0042392 | Syn1 | synapsin I | 706.27 | 582.58 | 0.28 | 2.99E-01 |
| ENSMUSG00000001128 | GSMG0042394 | Cfp | complement factor properdin | 280.16 | 137.09 | 1.04 | 3.54E-05 |
| ENSMUSG00000009406 | GSMG0042396 | Elk1 | ELK1, member of ETS oncogene family | 450.64 | 400.27 | 0.17 | 5.58E-01 |
| ENSMUSG00000001134 | GSMG0042397 | Uxt | ubiquitously expressed transcript | 154.44 | 168.40 | -0.12 | 7.71E-01 |
| ENSMUSG00000036782 | GSMG0042403 | Klhl13 | kelch-like 13 | 481.22 | 657.15 | -0.45 | 4.32E-02 |
| ENSMUSG00000059708 | GSMG0042415 | Akap17b | A kinase (PRKA) anchor protein 17B | 133.07 | 127.09 | 0.07 | 8.83E-01 |
| ENSMUSG00000006423 | GSMG0042417 | C330007P06Rik | RIKEN cDNA C330007P06 gene | 974.98 | 1038.19 | -0.09 | 7.93E-01 |
| ENSMUSG00000050379 | GSMG0042420 | Sept6 | septin 6 | 345.99 | 592.51 | -0.77 | 1.36E-05 |
| ENSMUSG00000036572 | GSMG0042421 | Upf3b | UPF3 regulator of nonsense transcripts homolog B (yeast) | 424.59 | 477.97 | -0.17 | 5.66E-01 |
| ENSMUSG00000016427 | GSMG0042423 | Ndufa1 | NADH dehydrogenase (ubiquinone) 1 alpha subcomplex, 1 | 1840.31 | 1722.67 | 0.10 | 6.81E-01 |
| ENSMUSG00000016534 | GSMG0042436 | Lamp2 | lysosomal-associated membrane protein 2 | 2420.17 | 2485.85 | -0.04 | 9.03E-01 |
| G00000031095,ENSMUSG000 | GSMG0042438 | Cul4b // C1galt1c1 | cullin 4B // C1GALT1-specific chaperone 1 | 563.53 | 746.62 | -0.41 | 6.50E-02 |
| ENSMUSG00000016150 | GSMG0042447 | Tenm1 | teneurin transmembrane protein 1 | 801.92 | 1073.23 | -0.42 | 4.42E-02 |
| ENSMUSG00000031099 | GSMG0042462 | Smarca1 | SWI/SNF related, matrix associated, actin dependent regulator of chron | 263.04 | 413.82 | -0.66 | 4.94E-03 |
| ENSMUSG00000037010 | GSMG0042464 | Apln | apelin | 8346.53 | 11781.35 | -0.50 | 5.81E-03 |
| ENSMUSG00000036985 | GSMG0042466 | Zdhhc9 | zinc finger, DHHC domain containing 9 | 785.12 | 828.97 | -0.08 | 7.76E-01 |
| ENSMUSG00000036932 | GSMG0042473 | Aifm1 | apoptosis-inducing factor, mitochondrion-associated 1 | 810.60 | 927.84 | -0.19 | 3.79E-01 |
| ENSMUSG00000036916 | GSMG0042474 | Zfp280c | zinc finger protein 280C | 684.04 | 689.54 | -0.01 | 9.78E-01 |

| | | | | | | | |
|---|---|---|---|---|---|---|---|
| ENSMUSG00000031109 | GSMG0042478 | Enox2 | ecto-NOX disulfide-thiol exchanger 2 | 344.96 | 341.20 | 0.02 | 9.65E-01 |
| ENSMUSG00000050029 | GSMG0042489 | Rap2c | RAP2C, member of RAS oncogene family | 354.78 | 438.59 | -0.31 | 2.77E-01 |
| ENSMUSG00000062184 | GSMG0042496 | Hs6st2 | heparan sulfate 6-O-sulfotransferase 2 | 58.93 | 160.82 | -1.45 | 2.22E-06 |
| ENSMUSG00000031119 | GSMG0042501 | Gpc4 | glypican 4 | 354.49 | 281.37 | 0.34 | 2.32E-01 |
| ENSMUSG00000055653 | GSMG0042504 | Gpc3 | glypican 3 | 1272.91 | 1334.79 | -0.07 | 8.36E-01 |
| ENSMUSG00000023074 | GSMG0042520 | Mospd1 | motile sperm domain containing 1 | 2481.71 | 2308.25 | 0.10 | 6.91E-01 |
| ENSMUSG00000061273 | GSMG0042533 | Mmgt1 | membrane magnesium transporter 1 | 718.12 | 910.91 | -0.34 | 1.08E-01 |
| NULL | GSMG0042536 | NULL | NULL | 317.82 | 317.95 | 0.00 | 9.93E-01 |
| ENSMUSG00000031133 | GSMG0042537 | Arhgef6 | Rac/Cdc42 guanine nucleotide exchange factor (GEF) 6 | 558.11 | 394.75 | 0.50 | 2.62E-02 |
| ENSMUSG00000031134 | GSMG0042540 | Rbmx | RNA binding motif protein, X chromosome | 1714.40 | 2189.85 | -0.35 | 4.53E-02 |
| ENSMUSG00000031137 | GSMG0042543 | Fgf13 | fibroblast growth factor 13 | 893.85 | 1531.57 | -0.78 | 1.97E-05 |
| ENSMUSG00000062949 | GSMG0042554 | Atp11c | ATPase, class VI, type 11C | 501.18 | 492.72 | 0.03 | 9.42E-01 |
| ENSMUSG00000046699 | GSMG0042563 | Slitrk4 | SLIT and NTRK-like family, member 4 | 361.90 | 400.17 | -0.15 | 6.11E-01 |
| ENSMUSG00000035847 | GSMG0042570 | Ids | iduronate 2-sulfatase | 2130.42 | 1988.47 | 0.10 | 6.68E-01 |
| ENSMUSG00000073139 | GSMG0042572 | BC023829 | cDNA sequence BC023829 | 310.33 | 339.99 | -0.13 | 7.07E-01 |
| ENSMUSG00000035776 | GSMG0042574 | Cd99l2 | CD99 antigen-like 2 | 256.38 | 313.65 | -0.29 | 2.60E-01 |
| ENSMUSG00000031343 | GSMG0042578 | Gabra3 | gamma-aminobutyric acid (GABA) A receptor, subunit alpha 3 | 73.25 | 175.71 | -1.26 | 7.18E-06 |
| ENSMUSG00000031347 | GSMG0042581 | Cetn2 | centrin 2 | 1143.99 | 1355.43 | -0.25 | 3.83E-01 |
| ENSMUSG00000031371 | GSMG0042593 | Haus7 | HAUS augmin-like complex, subunit 7 | 355.61 | 534.90 | -0.59 | 2.43E-03 |
| ENSMUSG00000002012 | GSMG0042594 | Pnck | pregnancy upregulated non-ubiquitously expressed CaM kinase | 1603.50 | 1845.04 | -0.20 | 4.53E-01 |
| ENSMUSG00000002015 | GSMG0042595 | Bcap31 | B cell receptor associated protein 31 | 3141.17 | 3053.15 | 0.04 | 8.89E-01 |
| ENSMUSG00000002010 | GSMG0042596 | Idh3g | isocitrate dehydrogenase 3 (NAD+), gamma | 3644.86 | 2821.38 | 0.37 | 6.46E-02 |
| NULL | GSMG0042598 | NULL | NULL | 209.28 | 171.76 | 0.29 | 3.51E-01 |
| ENSMUSG00000031391 | GSMG0042600 | L1cam | L1 cell adhesion molecule | 318.19 | 986.25 | -1.63 | 1.22E-10 |
| G00000031388,ENSMUSG000 | GSMG0042601 | Arhgap4 // Naa10 | Rho GTPase activating protein 4 // N(alpha)-acetyltransferase 10, NatA | 290.39 | 207.74 | 0.48 | 1.60E-01 |
| ENSMUSG00000031387 | GSMG0042603 | Renbp | renin binding protein | 510.87 | 513.91 | -0.01 | 9.86E-01 |
| ENSMUSG00000031386 | GSMG0042604 | Hcfc1 | host cell factor C1 | 1149.40 | 1528.02 | -0.41 | 1.64E-01 |
| ENSMUSG00000031392 | GSMG0042606 | Irak1 | interleukin-1 receptor-associated kinase 1 | 2325.66 | 1879.43 | 0.31 | 1.91E-01 |
| ENSMUSG00000031393 | GSMG0042609 | Mecp2 | methyl CpG binding protein 2 | 1251.24 | 1223.91 | 0.03 | 9.18E-01 |
| ENSMUSG00000031328 | GSMG0042610 | Flna | filamin, alpha | 24130.24 | 33585.82 | -0.48 | 5.67E-02 |
| ENSMUSG00000019088 | GSMG0042611 | Dnase1l1 | deoxyribonuclease 1-like 1 | 387.92 | 276.32 | 0.49 | 3.20E-02 |
| ENSMUSG00000015289 | GSMG0042613 | Lage3 | L antigen family, member 3 | 822.44 | 857.58 | -0.06 | 8.57E-01 |
| G00000015290,ENSMUSG000 | GSMG0042614 | Ubl4a // Gm44504 | ubiquitin-like 4A // predicted readthrough transcript (NMD candidate), | 318.20 | 367.31 | -0.21 | 4.32E-01 |

| | | | | | | | |
|---|---|---|---|---|---|---|---|
| ENSMUSG00000031399 | GSMG0042615 | Fam3a | family with sequence similarity 3, member A | 921.82 | 929.51 | -0.01 | 9.69E-01 |
| ENSMUSG00000031400 | GSMG0042616 | G6pdx | glucose-6-phosphate dehydrogenase X-linked | 567.26 | 649.82 | -0.19 | 5.20E-01 |
| ENSMUSG00000032750 | GSMG0042617 | Gm38425 // Gab3 | predicted gene, 38425 // growth factor receptor bound protein 2-associa | 67.48 | 60.99 | 0.15 | 8.00E-01 |
| ENSMUSG00000031402 | GSMG0042618 | Mpp1 | membrane protein, palmitoylated | 1293.61 | 1550.55 | -0.26 | 2.41E-01 |
| G00000031200,ENSMUSG000 | GSMG0042623 | Cmc4 // Mtcp1 | C-x(9)-C motif containing 4 // mature T cell proliferation 1 | 362.87 | 367.25 | -0.02 | 9.58E-01 |
| ENSMUSG00000031202 | GSMG0042625 | Rab39b | RAB39B, member RAS oncogene family | 269.40 | 325.21 | -0.27 | 2.90E-01 |
| ENSMUSG00000016382 | GSMG0042627 | Pls3 | plastin 3 (T-isoform) | 8441.71 | 6839.88 | 0.30 | 1.28E-01 |
| ENSMUSG00000035725 | GSMG0042633 | Prkx | protein kinase, X-linked | 498.61 | 530.99 | -0.09 | 7.84E-01 |
| ENSMUSG00000025059 | GSMG0042646 | Gk | glycerol kinase | 257.33 | 261.64 | -0.02 | 9.60E-01 |
| ENSMUSG00000052372 | GSMG0042655 | Il1rapl1 | interleukin 1 receptor accessory protein-like 1 | 521.26 | 398.79 | 0.39 | 8.79E-02 |
| NULL | GSMG0042663 | NULL | NULL | 349.75 | 189.66 | 0.89 | 3.28E-04 |
| ENSMUSG00000006678 | GSMG0042676 | Pola1 | polymerase (DNA directed), alpha 1 | 167.21 | 272.81 | -0.70 | 2.54E-03 |
| ENSMUSG00000035232 | GSMG0042680 | Pdk3 | pyruvate dehydrogenase kinase, isoenzyme 3 | 654.01 | 751.07 | -0.20 | 4.92E-01 |
| ENSMUSG00000079509 | GSMG0042681 | Zfx | zinc finger protein X-linked | 236.67 | 229.02 | 0.05 | 8.98E-01 |
| ENSMUSG00000025151 | GSMG0042685 | Maged1 | melanoma antigen, family D, 1 | 9340.57 | 11421.53 | -0.29 | 9.21E-02 |
| ENSMUSG00000025656 | GSMG0042689 | Arhgef9 | CDC42 guanine nucleotide exchange factor (GEF) 9 | 880.17 | 652.96 | 0.43 | 4.38E-02 |
| ENSMUSG00000050332 | GSMG0042692 | Amer1 | APC membrane recruitment 1 | 535.19 | 1031.59 | -0.95 | 4.25E-08 |
| ENSMUSG00000035062 | GSMG0042694 | Zc4h2 | zinc finger, C4H2 domain containing | 85.84 | 124.58 | -0.53 | 1.66E-01 |
| ENSMUSG00000057421 | GSMG0042696 | Las1l | LAS1-like (S. cerevisiae) | 2894.04 | 3471.48 | -0.26 | 1.44E-01 |
| ENSMUSG00000031214 | GSMG0042702 | Ophn1 | oligophrenin 1 | 1957.42 | 2266.15 | -0.21 | 2.86E-01 |
| ENSMUSG00000034403 | GSMG0042709 | Pja1 | praja ring finger 1, E3 ubiquitin protein ligase | 3938.63 | 3983.90 | -0.02 | 9.52E-01 |
| ENSMUSG00000015668 | GSMG0042712 | Pdzd11 | PDZ domain containing 11 | 1144.68 | 1033.95 | 0.15 | 5.08E-01 |
| ENSMUSG00000031297 | GSMG0042716 | Slc7a3 | solute carrier family 7 (cationic amino acid transporter, y+ system), mer | 932.12 | 924.66 | 0.01 | 9.73E-01 |
| ENSMUSG00000046032 | GSMG0042722 | Snx12 | sorting nexin 12 | 210.48 | 261.56 | -0.32 | 2.34E-01 |
| ENSMUSG00000031304 | GSMG0042724 | Il2rg | interleukin 2 receptor, gamma chain | 2306.65 | 2242.28 | 0.04 | 8.98E-01 |
| ENSMUSG00000031310 | GSMG0042725 | Zmym3 | zinc finger, MYM-type 3 | 820.46 | 904.40 | -0.14 | 6.69E-01 |
| ENSMUSG00000067567 | GSMG0042732 | Hdac8 | histone deacetylase 8 | 173.48 | 143.44 | 0.28 | 4.61E-01 |
| ENSMUSG00000034055 | GSMG0042743 | Phka1 | phosphorylase kinase alpha 1 | 393.33 | 267.53 | 0.56 | 1.43E-02 |
| ENSMUSG00000086370 | GSMG0042755 | Ftx | Ftx transcript, Xist regulator (non-protein coding) | 586.69 | 265.54 | 1.14 | 3.86E-09 |
| ENSMUSG00000033965 | GSMG0042756 | Slc16a2 | solute carrier family 16 (monocarboxylic acid transporters), member 2 | 1793.87 | 1290.53 | 0.48 | 1.63E-02 |
| ENSMUSG00000056537 | GSMG0042760 | Rlim | ring finger protein, LIM domain interacting | 1301.65 | 1321.56 | -0.02 | 9.49E-01 |
| ENSMUSG00000046449 | GSMG0042762 | C77370 | expressed sequence C77370 | 251.19 | 845.10 | -1.75 | 3.45E-26 |
| ENSMUSG00000031333 | GSMG0042764 | Abcb7 | ATP-binding cassette, sub-family B (MDR/TAP), member 7 | 194.64 | 235.36 | -0.27 | 3.92E-01 |

| | | | | | | | |
|---|---|---|---|---|---|---|---|
| NULL | GSMG0042765 | NULL | NULL | 651.66 | 829.39 | -0.35 | 6.01E-02 |
| ENSMUSG00000033906 | GSMG0042769 | Zdhhc15 | zinc finger, DHHC domain containing 15 | 93.69 | 155.23 | -0.73 | 2.37E-02 |
| ENSMUSG00000087174 | GSMG0042772 | 5530601H04Rik | RIKEN cDNA 5530601H04 gene | 174.39 | 161.04 | 0.12 | 7.93E-01 |
| ENSMUSG00000087620 | GSMG0042779 | 5330434G04Rik | RIKEN cDNA 5330434G04 gene | 1841.37 | 733.07 | 1.33 | 1.53E-09 |
| ENSMUSG00000031229 | GSMG0042783 | Atrx | alpha thalassemia/mental retardation syndrome X-linked | 1066.25 | 1309.42 | -0.30 | 9.04E-02 |
| ENSMUSG00000031232 | GSMG0042788 | Magt1 | magnesium transporter 1 | 678.15 | 1007.97 | -0.57 | 1.46E-02 |
| ENSMUSG00000047242 | GSMG0042789 | Taf9b | TATA-box binding protein associated factor 9B | 323.29 | 272.97 | 0.25 | 4.11E-01 |
| ENSMUSG00000031239 | GSMG0042796 | Itm2a | integral membrane protein 2A | 38243.17 | 27442.14 | 0.48 | 1.85E-02 |
| ENSMUSG00000031242 | GSMG0042798 | 2610002M06Rik | RIKEN cDNA 2610002M06 gene | 136.58 | 158.18 | -0.21 | 5.65E-01 |
| ENSMUSG00000031245 | GSMG0042805 | Hmgn5 | high-mobility group nucleosome binding domain 5 | 531.59 | 619.72 | -0.22 | 4.64E-01 |
| ENSMUSG00000034551 | GSMG0042810 | Hdx | highly divergent homeobox | 143.08 | 182.01 | -0.34 | 2.97E-01 |
| ENSMUSG00000025531 | GSMG0042814 | Chm | choroidermia (RAB escort protein 1) | 574.45 | 562.70 | 0.03 | 9.34E-01 |
| ENSMUSG00000055733 | GSMG0042821 | Nap1l3 | nucleosome assembly protein 1-like 3 | 380.54 | 386.23 | -0.03 | 9.52E-01 |
| ENSMUSG00000051323 | GSMG0042831 | Pcdh19 | protocadherin 19 | 254.57 | 332.58 | -0.39 | 1.10E-01 |
| ENSMUSG00000067377 | GSMG0042832 | Tspan6 | tetraspanin 6 | 1509.84 | 3104.29 | -1.04 | 5.11E-11 |
| ENSMUSG00000067369 | GSMG0042838 | Trmt2b | TRM2 tRNA methyltransferase 2B | 264.28 | 188.91 | 0.49 | 8.35E-02 |
| ENSMUSG00000031266 | GSMG0042843 | Gla | galactosidase, alpha | 85.75 | 103.67 | -0.27 | 5.33E-01 |
| ENSMUSG00000033436 | GSMG0042847 | Armcx2 | armadillo repeat containing, X-linked 2 | 1751.43 | 2599.92 | -0.57 | 1.18E-04 |
| ENSMUSG00000042750 | GSMG0042856 | Bex2 | brain expressed X-linked 2 | 63.26 | 84.52 | -0.43 | 3.25E-01 |
| ENSMUSG00000051579 | GSMG0042858 | Tceal8 | transcription elongation factor A (SII)-like 8 | 745.90 | 779.37 | -0.06 | 8.89E-01 |
| ENSMUSG00000054034 | GSMG0042859 | Tceal5 | transcription elongation factor A (SII)-like 5 | 365.76 | 225.87 | 0.69 | 6.62E-03 |
| NULL | GSMG0042861 | NULL | NULL | 200.28 | 248.65 | -0.31 | 2.61E-01 |
| ENSMUSG00000031422 | GSMG0042863 | Morf4l2 | mortality factor 4 like 2 | 5855.29 | 7376.25 | -0.33 | 5.42E-02 |
| ENSMUSG00000044348 | GSMG0042867 | Slc25a53 | solute carrier family 25, member 53 | 109.87 | 114.94 | -0.07 | 8.90E-01 |
| ENSMUSG00000031434 | GSMG0042873 | Morc4 | microrchidia 4 | 83.29 | 114.51 | -0.45 | 2.42E-01 |
| ENSMUSG00000031433 | GSMG0042874 | Rbm41 | RNA binding motif protein 41 | 96.71 | 112.72 | -0.22 | 6.03E-01 |
| ENSMUSG00000031431 | GSMG0042876 | Tsc22d3 | TSC22 domain family, member 3 | 8339.73 | 3871.71 | 1.11 | 9.26E-11 |
| ENSMUSG00000031429 | GSMG0042880 | Psmd10 | proteasome (prosome, macropain) 26S subunit, non-ATPase, 10 | 487.16 | 447.15 | 0.12 | 7.13E-01 |
| ENSMUSG00000090122 | GSMG0042889 | Kcne1l | potassium voltage-gated channel, Isk-related family, member 1-like, ps | 228.35 | 382.01 | -0.75 | 8.07E-04 |
| ENSMUSG00000031278 | GSMG0042890 | Acsl4 | acyl-CoA synthetase long-chain family member 4 | 250.79 | 342.06 | -0.45 | 3.96E-02 |
| ENSMUSG00000042225 | GSMG0042894 | Ammecr1 | Alport syndrome, mental retardation, midface hypoplasia and elliptocyt | 315.44 | 412.04 | -0.38 | 1.58E-01 |
| ENSMUSG00000031283 | GSMG0042897 | Chrdl1 | chordin-like 1 | 64.79 | 85.54 | -0.40 | 3.44E-01 |
| ENSMUSG00000031285 | GSMG0042900 | Dcx | doublecortin | 278.74 | 786.29 | -1.50 | 3.58E-18 |

| | | | | | | | |
|---|---|---|---|---|---|---|---|
| NULL | GSMG0042901 | NULL | NULL | 198.80 | 357.95 | -0.85 | 4.30E-04 |
| ENSMUSG00000041688 | GSMG0042908 | Amot | angiomotin | 98.82 | 69.91 | 0.50 | 1.76E-01 |
| ENSMUSG00000041353 | GSMG0042922 | Tmem29 | transmembrane protein 29 | 433.36 | 472.78 | -0.12 | 6.91E-01 |
| ENSMUSG00000025269 | GSMG0042926 | Apex2 | apurinic/apyrimidinic endonuclease 2 | 254.42 | 374.29 | -0.55 | 1.52E-02 |
| ENSMUSG00000025272 | GSMG0042928 | Tro | trophinin | 1084.77 | 1806.94 | -0.74 | 2.86E-04 |
| ENSMUSG00000025268 | GSMG0042929 | Maged2 | melanoma antigen, family D, 2 | 4381.70 | 7637.54 | -0.80 | 1.96E-07 |
| ENSMUSG00000025266 | GSMG0042931 | Gnl3l | guanine nucleotide binding protein-like 3 (nucleolar)-like | 2104.41 | 2151.61 | -0.03 | 9.11E-01 |
| ENSMUSG00000025264 | GSMG0042934 | Tsr2 | TSR2 20S rRNA accumulation | 649.00 | 683.94 | -0.07 | 8.02E-01 |
| ENSMUSG00000041096 | GSMG0042942 | Tspyl2 | TSPY-like 2 | 1719.50 | 1301.64 | 0.40 | 1.06E-01 |
| ENSMUSG00000045180 | GSMG0042944 | Shroom2 | shroom family member 2 | 365.25 | 587.95 | -0.69 | 4.77E-04 |
| ENSMUSG00000047238 | GSMG0042948 | Mageh1 | melanoma antigen, family H, 1 | 1041.45 | 1440.13 | -0.47 | 1.55E-02 |
| ENSMUSG00000041633 | GSMG0042949 | Kctd12b | potassium channel tetramerisation domain containing 12b | 1233.01 | 1343.88 | -0.12 | 6.84E-01 |
| ENSMUSG00000025283 | GSMG0042958 | Sat1 | spermidine/spermine N1-acetyl transferase 1 | 4917.66 | 5593.58 | -0.19 | 2.99E-01 |
| ENSMUSG00000025289 | GSMG0042960 | Prdx4 | peroxiredoxin 4 | 976.58 | 1565.85 | -0.68 | 2.85E-04 |
| ENSMUSG00000041552 | GSMG0042962 | Ptchd1 | patched domain containing 1 | 74.43 | 72.36 | 0.04 | 9.50E-01 |
| ENSMUSG00000046873 | GSMG0042966 | Mbtps2 | membrane-bound transcription factor peptidase, site 2 | 113.04 | 148.62 | -0.39 | 2.84E-01 |
| NULL | GSMG0042968 | NULL | NULL | 654.88 | 604.07 | 0.12 | 7.06E-01 |
| ENSMUSG00000025658 | GSMG0042969 | Cnksr2 | connector enhancer of kinase suppressor of Ras 2 | 392.15 | 382.77 | 0.03 | 9.30E-01 |
| ENSMUSG00000031299 | GSMG0042971 | Pdha1 | pyruvate dehydrogenase E1 alpha 1 | 1767.24 | 1861.60 | -0.07 | 7.98E-01 |
| NULL | GSMG0042974 | NULL | NULL | 137.71 | 136.15 | 0.01 | 9.83E-01 |
| ENSMUSG00000038344 | GSMG0042995 | Txlng | taxilin gamma | 163.07 | 147.14 | 0.14 | 7.22E-01 |
| ENSMUSG00000031370 | GSMG0043001 | Zrsr2 | zinc finger (CCCH type), RNA binding motif and serine/arginine rich 2 | 616.59 | 672.40 | -0.13 | 6.59E-01 |
| ENSMUSG00000040749 | GSMG0043005 | Siah1b | seven in absentia 1B | 273.60 | 375.75 | -0.46 | 4.88E-02 |
| ENSMUSG00000031377 | GSMG0043007 | Bmx | BMX non-receptor tyrosine kinase | 945.72 | 115.97 | 3.03 | 4.26E-32 |
| ENSMUSG00000061778 | GSMG0043010 | Mospd2 | motile sperm domain containing 2 | 152.09 | 147.41 | 0.05 | 9.24E-01 |
| ENSMUSG00000018589 | GSMG0043012 | Glra2 | glycine receptor, alpha 2 subunit | 37.03 | 146.69 | -2.00 | 1.66E-10 |
| ENSMUSG00000040586 | GSMG0043014 | Ofd1 | oral-facial-digital syndrome 1 gene homolog (human) | 130.49 | 182.22 | -0.48 | 1.22E-01 |
| ENSMUSG00000079316 | GSMG0043015 | Rab9 | RAB9, member RAS oncogene family | 601.89 | 660.47 | -0.14 | 6.89E-01 |
| ENSMUSG00000049775 | GSMG0043021 | Tmsb4x | thymosin, beta 4, X chromosome | 74943.22 | 100669.77 | -0.43 | 1.37E-01 |
| ENSMUSG00000049176 | GSMG0043025 | Frmpd4 | FERM and PDZ domain containing 4 | 329.22 | 368.29 | -0.16 | 5.93E-01 |
| ENSMUSG00000031358 | GSMG0043030 | Msl3 | male-specific lethal 3 homolog (Drosophila) | 621.81 | 649.71 | -0.06 | 8.31E-01 |
| ENSMUSG00000031352 | GSMG0043034 | Hccs | holocytochrome c synthetase | 863.64 | 865.34 | 0.00 | 9.91E-01 |
| NULL | GSMG0043037 | Vamp7 | vesicle-associated membrane protein 7 | 344.74 | 400.95 | -0.22 | 4.46E-01 |

| | | | | | | | |
|---|---|---|---|---|---|---|---|
| NULL | GSMG0043039 | Tmlhe | trimethyllysine hydroxylase, epsilon | 312.77 | 340.40 | -0.12 | 7.12E-01 |
| ENSMUSG00000056673 | GSMG0043044 | Kdm5d | lysine (K)-specific demethylase 5D | 175.03 | 244.84 | -0.48 | 2.21E-01 |
| NULL | GSMG0043045 | NULL | NULL | 525.81 | 266.97 | 0.98 | 5.42E-06 |
| ENSMUSG00000069049 | GSMG0043046 | Eif2s3y | eukaryotic translation initiation factor 2, subunit 3, structural gene Y-lir | 1667.73 | 1657.47 | 0.01 | 9.83E-01 |
| ENSMUSG00000068457 | GSMG0043052 | Uty | ubiquitously transcribed tetratricopeptide repeat gene, Y chromosome | 415.02 | 405.01 | 0.04 | 9.35E-01 |
| ENSMUSG00000069045 | GSMG0043055 | Ddx3y | DEAD (Asp-Glu-Ala-Asp) box polypeptide 3, Y-linked | 607.14 | 644.00 | -0.08 | 8.58E-01 |
| ENSMUSG00000032560 | GSMG0043102 | Dnajc13 | DnaJ heat shock protein family (Hsp40) member C13 | 1340.99 | 1794.15 | -0.42 | 2.39E-02 |
| NULL | GSMG0043174 | NULL | NULL | 349.38 | 379.68 | -0.12 | 6.89E-01 |
| ENSMUSG00000095930 | GSMG0043184 | Nim1k | NIM1 serine/threonine protein kinase | 262.45 | 309.06 | -0.24 | 3.79E-01 |
| NULL | GSMG0043186 | NULL | NULL | 142.74 | 338.06 | -1.25 | 6.53E-05 |
| NULL | GSMG0043188 | NULL | NULL | 380.80 | 368.42 | 0.04 | 9.13E-01 |
| ENSMUSG00000093930 | GSMG0043189 | Hmgcs1 | 3-hydroxy-3-methylglutaryl-Coenzyme A synthase 1 | 1588.31 | 3256.53 | -1.04 | 5.12E-14 |
| NULL | GSMG0043226 | NULL | NULL | 224.27 | 87.95 | 1.35 | 3.69E-06 |
| ENSMUSG00000071640 | GSMG0043287 | Stxbp3-ps | syntaxin-binding protein 3, pseudogene | 230.43 | 186.82 | 0.31 | 3.07E-01 |
| NULL | GSMG0043374 | NULL | NULL | 3005.73 | 2616.39 | 0.20 | 2.89E-01 |
| NULL | GSMG0043389 | NULL | NULL | 774.37 | 1267.07 | -0.71 | 4.20E-05 |
| ENSMUSG00000056820 | GSMG0043399 | Tsnax | translin-associated factor X | 4044.04 | 4018.41 | 0.01 | 9.73E-01 |
| ENSMUSG00000040481 | GSMG0043470 | Bptf | bromodomain PHD finger transcription factor | 291.81 | 298.93 | -0.04 | 9.30E-01 |
| ENSMUSG00000059486 | GSMG0043483 | Kbtbd2 | kelch repeat and BTB (POZ) domain containing 2 | 2059.68 | 1963.27 | 0.07 | 7.92E-01 |
| NULL | GSMG0043484 | NULL | NULL | 556.81 | 536.73 | 0.05 | 8.77E-01 |
| ENSMUSG00000001517 | GSMG0043487 | Foxm1 | forkhead box M1 | 96.47 | 213.24 | -1.15 | 9.89E-05 |
| ENSMUSG00000078813 | GSMG0043495 | Leng1 | leukocyte receptor cluster (LRC) member 1 | 192.38 | 149.46 | 0.36 | 2.71E-01 |
| NULL | GSMG0043508 | A830010M20Rik | RIKEN cDNA A830010M20 gene | 132.88 | 109.87 | 0.27 | 5.46E-01 |
| ENSMUSG00000040003 | GSMG0043512 | Magi2 | membrane associated guanylate kinase, WW and PDZ domain containin | 2097.70 | 3041.61 | -0.54 | 3.37E-03 |
| ENSMUSG00000006641 | GSMG0043514 | Slc5a6 | solute carrier family 5 (sodium-dependent vitamin transporter), member | 2997.89 | 1938.56 | 0.63 | 1.39E-04 |
| ENSMUSG00000055204 | GSMG0043515 | Ankrd17 | ankyrin repeat domain 17 | 2569.86 | 2454.17 | 0.07 | 8.02E-01 |
| ENSMUSG00000079109 | GSMG0043519 | Pms2 | postmeiotic segregation increased 2 (S. cerevisiae) | 820.11 | 816.35 | 0.01 | 9.83E-01 |
| ENSMUSG00000007415 | GSMG0043520 | Gatad1 | GATA zinc finger domain containing 1 | 661.13 | 691.35 | -0.07 | 8.48E-01 |
| ENSMUSG00000054641 | GSMG0043525 | Mmrn1 | multimerin 1 | 242.90 | 697.52 | -1.52 | 2.08E-15 |
| ENSMUSG00000079426 | GSMG0043530 | Arpc4 | actin related protein 2/3 complex, subunit 4 | 4910.03 | 5895.62 | -0.26 | 2.35E-01 |
| ENSMUSG00000030276 | GSMG0043531 | Ttll3 | tubulin tyrosine ligase-like family, member 3 | 346.95 | 277.20 | 0.32 | 2.16E-01 |
| ENSMUSG00000032827 | GSMG0043532 | Ppp1r9a | protein phosphatase 1, regulatory (inhibitor) subunit 9A | 525.08 | 699.03 | -0.41 | 1.04E-01 |
| ENSMUSG00000061762 | GSMG0043535 | Tac1 | tachykinin 1 | 370.84 | 200.01 | 0.89 | 2.14E-01 |

| | | | | | | | |
|---|---|---|---|---|---|---|---|
| ENSMUSG00000045466 | GSMG0043536 | Zfp956 | zinc finger protein 956 | 251.14 | 276.30 | -0.14 | 7.16E-01 |
| ENSMUSG00000072653 | GSMG0043537 | Zfp783 | zinc finger protein 783 | 61.87 | 101.53 | -0.71 | 7.29E-02 |
| ENSMUSG00000057137 | GSMG0043541 | Tmem140 | transmembrane protein 140 | 466.71 | 293.71 | 0.67 | 8.00E-04 |
| ENSMUSG00000033594 | GSMG0043542 | Spata2l | spermatogenesis associated 2-like | 1873.65 | 566.91 | 1.72 | 2.20E-23 |
| ENSMUSG00000007656 | GSMG0043550 | Arpp19 | cAMP-regulated phosphoprotein 19 | 3580.29 | 3309.75 | 0.11 | 6.69E-01 |
| G00000010051,ENSMUSG000 | GSMG0043555 | Hyal1 // Nat6 | hyaluronoglucosaminidase 1 // N-acetyltransferase 6 | 871.55 | 628.85 | 0.47 | 5.59E-03 |
| ENSMUSG00000066415 | GSMG0043563 | Msl2 | male-specific lethal 2 homolog (Drosophila) | 211.52 | 207.03 | 0.03 | 9.55E-01 |
| NULL | GSMG0043564 | NULL | NULL | 920.71 | 864.08 | 0.09 | 7.38E-01 |
| NULL | GSMG0043577 | NULL | NULL | 1129.88 | 1395.41 | -0.30 | 8.25E-02 |
| ENSMUSG00000042757 | GSMG0043578 | Tmem108 | transmembrane protein 108 | 179.73 | 387.92 | -1.11 | 7.31E-08 |
| ENSMUSG00000079469 | GSMG0043579 | Pigb | phosphatidylinositol glycan anchor biosynthesis, class B | 227.29 | 216.72 | 0.07 | 8.68E-01 |
| NULL | GSMG0043580 | NULL | NULL | 498.85 | 341.25 | 0.55 | 7.17E-03 |
| ENSMUSG00000032355 | GSMG0043582 | Mlip | muscular LMNA-interacting protein | 131.82 | 169.43 | -0.35 | 3.23E-01 |
| ENSMUSG00000025241 | GSMG0043589 | Fyco1 | FYVE and coiled-coil domain containing 1 | 3854.44 | 2641.52 | 0.55 | 2.39E-03 |
| NULL | GSMG0043595 | NULL | NULL | 495.73 | 309.15 | 0.68 | 4.37E-04 |
| ENSMUSG00000059146 | GSMG0043600 | Ntrk3 | neurotrophic tyrosine kinase, receptor, type 3 | 446.23 | 590.51 | -0.40 | 1.43E-01 |
| ENSMUSG00000030986 | GSMG0043605 | Dhx32 | DEAH (Asp-Glu-Ala-His) box polypeptide 32 | 1243.03 | 1264.29 | -0.02 | 9.34E-01 |
| ENSMUSG00000031985 | GSMG0043613 | Gnpat | glyceronephosphate O-acyltransferase | 2376.24 | 2489.12 | -0.07 | 7.98E-01 |
| ENSMUSG00000078908 | GSMG0043616 | Mon1b | MON1 homolog B, secretory traffciking associated | 259.08 | 253.75 | 0.03 | 9.51E-01 |
| ENSMUSG00000031652 | GSMG0043620 | N4bp1 | NEDD4 binding protein 1 | 1018.88 | 871.39 | 0.23 | 2.55E-01 |
| NULL | GSMG0043621 | NULL | NULL | 228.05 | 186.27 | 0.29 | 3.76E-01 |
| G00000031916,ENSMUSG000 | GSMG0043622 | Pdf // Cog8 | peptide deformylase (mitochondrial) // component of oligomeric golgi c | 1012.02 | 1060.66 | -0.07 | 8.03E-01 |
| ENSMUSG00000052676 | GSMG0043626 | Zmat1 | zinc finger, matrin type 1 | 457.70 | 436.92 | 0.07 | 8.60E-01 |
| ENSMUSG00000031357 | GSMG0043630 | Syap1 | synapse associated protein 1 | 851.67 | 920.22 | -0.11 | 7.31E-01 |
| ENSMUSG00000037509 | GSMG0043634 | Arhgef4 | Rho guanine nucleotide exchange factor (GEF) 4 | 684.21 | 647.29 | 0.08 | 8.31E-01 |
| ENSMUSG00000040225 | GSMG0043639 | Prrc2c | proline-rich coiled-coil 2C | 2010.22 | 2378.87 | -0.24 | 1.94E-01 |
| ENSMUSG00000060733 | GSMG0043641 | Ipmk | inositol polyphosphate multikinase | 1532.19 | 1698.42 | -0.15 | 5.29E-01 |
| ENSMUSG00000026269 | GSMG0043647 | Rnpepl1 | arginyl aminopeptidase (aminopeptidase B)-like 1 | 3322.35 | 2673.54 | 0.31 | 2.23E-01 |
| ENSMUSG00000026024 | GSMG0043653 | Als2 | amyotrophic lateral sclerosis 2 (juvenile) | 1007.10 | 961.65 | 0.07 | 8.44E-01 |
| ENSMUSG00000048234 | GSMG0043656 | Rnf149 | ring finger protein 149 | 503.01 | 432.47 | 0.22 | 3.80E-01 |
| ENSMUSG00000067653 | GSMG0043658 | Ankrd23 | ankyrin repeat domain 23 | 204.88 | 184.16 | 0.15 | 7.03E-01 |
| ENSMUSG00000079610 | GSMG0043659 | Ankrd39 | ankyrin repeat domain 39 | 1107.03 | 936.36 | 0.24 | 2.27E-01 |
| ENSMUSG00000009418 | GSMG0043660 | Nav1 | neuron navigator 1 | 1214.25 | 2101.62 | -0.79 | 3.34E-04 |

| | | | | | | | |
|---|---|---|---|---|---|---|---|
| NULL | GSMG0043665 | NULL | NULL | 203.64 | 227.64 | -0.16 | 6.34E-01 |
| NULL | GSMG0043666 | NULL | NULL | 109.61 | 121.83 | -0.15 | 7.58E-01 |
| ENSMUSG00000039640 | GSMG0043667 | Mrpl12 | mitochondrial ribosomal protein L12 | 1276.66 | 1679.79 | -0.40 | 2.64E-02 |
| NULL | GSMG0043668 | NULL | NULL | 133.08 | 186.76 | -0.49 | 8.17E-02 |
| ENSMUSG00000039275 | GSMG0043669 | Foxk2 | forkhead box K2 | 382.58 | 401.97 | -0.07 | 8.81E-01 |
| ENSMUSG00000000056 | GSMG0043670 | Narf | nuclear prelamin A recognition factor | 876.47 | 764.32 | 0.20 | 5.00E-01 |
| ENSMUSG00000040699 | GSMG0043677 | Limd2 | LIM domain containing 2 | 1203.75 | 1734.64 | -0.53 | 1.02E-03 |
| NULL | GSMG0043678 | NULL | NULL | 225.34 | 301.76 | -0.42 | 8.31E-02 |
| ENSMUSG00000052949 | GSMG0043679 | Rnf157 | ring finger protein 157 | 424.21 | 483.52 | -0.19 | 5.99E-01 |
| ENSMUSG00000020743 | GSMG0043683 | Mif4gd | MIF4G domain containing | 1544.44 | 1229.98 | 0.33 | 7.20E-02 |
| ENSMUSG00000020744 | GSMG0043684 | Slc25a19 | solute carrier family 25 (mitochondrial thiamine pyrophosphate carrier) | 242.19 | 272.46 | -0.17 | 6.21E-01 |
| ENSMUSG00000003348 | GSMG0043685 | Mob3a | MOB kinase activator 3A | 307.14 | 360.22 | -0.23 | 3.78E-01 |
| ENSMUSG00000020027 | GSMG0043692 | Socs2 | suppressor of cytokine signaling 2 | 404.13 | 416.05 | -0.04 | 9.04E-01 |
| ENSMUSG00000040441 | GSMG0043694 | Slc26a10 | solute carrier family 26, member 10 | 357.59 | 41.54 | 3.11 | 2.25E-34 |
| ENSMUSG00000014245 | GSMG0043698 | Pigl | phosphatidylinositol glycan anchor biosynthesis, class L | 343.33 | 362.43 | -0.08 | 8.12E-01 |
| NULL | GSMG0043702 | NULL | NULL | 367.88 | 344.80 | 0.09 | 8.71E-01 |
| ENSMUSG00000000631 | GSMG0043703 | Myo18a | myosin XVIIIA | 3388.40 | 3829.44 | -0.18 | 6.27E-01 |
| ENSMUSG00000000686 | GSMG0043704 | Abhd15 | abhydrolase domain containing 15 | 254.71 | 312.04 | -0.29 | 3.17E-01 |
| ENSMUSG00000047789 | GSMG0043708 | Slc38a9 | solute carrier family 38, member 9 | 466.84 | 402.24 | 0.22 | 4.52E-01 |
| ENSMUSG00000021699 | GSMG0043709 | Pde4d | phosphodiesterase 4D, cAMP specific | 283.48 | 332.93 | -0.23 | 4.32E-01 |
| NULL | GSMG0043710 | NULL | NULL | 135.67 | 75.86 | 0.85 | 1.50E-02 |
| ENSMUSG00000021066 | GSMG0043722 | Atl1 | atlastin GTPase 1 | 496.58 | 632.21 | -0.35 | 7.99E-02 |
| NULL | GSMG0043726 | NULL | NULL | 136.76 | 159.64 | -0.23 | 5.69E-01 |
| ENSMUSG00000033713 | GSMG0043727 | Foxn3 | forkhead box N3 | 1021.43 | 1139.78 | -0.16 | 5.02E-01 |
| ENSMUSG00000021038 | GSMG0043731 | Vipas39 | VPS33B interacting protein, apical-basolateral polarity regulator, spe-3 | 899.66 | 979.37 | -0.12 | 5.98E-01 |
| ENSMUSG00000021120 | GSMG0043734 | Pigh | phosphatidylinositol glycan anchor biosynthesis, class H | 960.63 | 886.81 | 0.12 | 6.71E-01 |
| ENSMUSG00000006378 | GSMG0043737 | Gcat | glycine C-acetyltransferase (2-amino-3-ketobutyrate-coenzyme A ligase | 72.80 | 120.06 | -0.72 | 3.71E-02 |
| ENSMUSG00000037075 | GSMG0043742 | Rnf139 | ring finger protein 139 | 1650.47 | 1598.78 | 0.05 | 8.77E-01 |
| ENSMUSG00000048546 | GSMG0043744 | Tob2 | transducer of ERBB2, 2 | 1244.78 | 860.59 | 0.53 | 7.65E-03 |
| ENSMUSG00000022557 | GSMG0043746 | Bop1 | block of proliferation 1 | 1844.42 | 2719.19 | -0.56 | 1.90E-04 |
| NULL | GSMG0043748 | C030006K11Rik | RIKEN cDNA C030006K11 gene | 247.64 | 261.60 | -0.08 | 8.32E-01 |
| ENSMUSG00000033707 | GSMG0043749 | Lrrc24 | leucine rich repeat containing 24 | 232.98 | 260.25 | -0.16 | 6.44E-01 |
| ENSMUSG00000079557 | GSMG0043770 | March2 | membrane-associated ring finger (C3HC4) 2 | 2320.73 | 1784.25 | 0.38 | 2.11E-02 |

| | | | | | | | |
|---|---|---|---|---|---|---|---|
| ENSMUSG00000059208 | GSMG0043772 | Hnrnpm | heterogeneous nuclear ribonucleoprotein M | 5011.33 | 5781.53 | -0.21 | 3.61E-01 |
| ENSMUSG00000034673 | GSMG0043778 | Pbx2 | pre B cell leukemia homeobox 2 | 222.65 | 199.90 | 0.15 | 8.07E-01 |
| G00000025733,ENSMUSG000 | GSMG0043781 | Rhbdl1 // Rhot2 | rhomboid, veinlet-like 1 (Drosophila) // ras homolog family member T2 | 907.86 | 566.72 | 0.68 | 9.63E-04 |
| ENSMUSG00000073434 | GSMG0043782 | Wdr90 | WD repeat domain 90 | 1754.53 | 1646.08 | 0.09 | 8.18E-01 |
| NULL | GSMG0043783 | Ntn3 | netrin 3 | 102.34 | 82.09 | 0.32 | 5.89E-01 |
| ENSMUSG00000036473 | GSMG0043784 | Tbc1d24 | TBC1 domain family, member 24 | 296.70 | 354.51 | -0.26 | 4.94E-01 |
| ENSMUSG00000052752 | GSMG0043785 | Traf7 | TNF receptor-associated factor 7 | 2579.17 | 2969.48 | -0.20 | 4.33E-01 |
| ENSMUSG00000079657 | GSMG0043786 | Rab26 | RAB26, member RAS oncogene family | 910.92 | 326.05 | 1.48 | 2.18E-12 |
| ENSMUSG00000046962 | GSMG0043788 | Zbtb21 | zinc finger and BTB domain containing 21 | 240.96 | 262.99 | -0.12 | 7.89E-01 |
| ENSMUSG00000059003 | GSMG0043791 | Grin2a | glutamate receptor, ionotropic, NMDA2A (epsilon 1) | 382.83 | 281.72 | 0.45 | 1.37E-01 |
| ENSMUSG00000035033 | GSMG0043799 | Tbr1 | T-box brain gene 1 | 149.40 | 256.24 | -0.77 | 7.72E-04 |
| ENSMUSG00000036792 | GSMG0043801 | Mbd5 | methyl-CpG binding domain protein 5 | 435.80 | 366.90 | 0.25 | 3.35E-01 |
| NULL | GSMG0043806 | Tmem179b | transmembrane protein 179B | 768.36 | 677.01 | 0.18 | 4.17E-01 |
| ENSMUSG00000003680 | GSMG0043807 | Taf6l | TATA-box binding protein associated factor 6 like | 201.33 | 272.63 | -0.43 | 1.15E-01 |
| G00000024483,ENSMUSG000 | GSMG0043812 | Ankhd1 // Eif4ebp | ankyrin repeat and KH domain containing 1 // eukaryotic translation ini | 3351.26 | 3249.11 | 0.05 | 8.76E-01 |
| ENSMUSG00000051375 | GSMG0043819 | Pcdh1 | protocadherin 1 | 1024.11 | 1202.17 | -0.23 | 4.66E-01 |
| NULL | GSMG0043820 | NULL | NULL | 111.60 | 131.77 | -0.25 | 6.19E-01 |
| ENSMUSG00000036160 | GSMG0043841 | Surf6 | surfeit gene 6 | 776.41 | 641.98 | 0.27 | 2.30E-01 |
| ENSMUSG00000078485 | GSMG0043854 | Plekhn1 | pleckstrin homology domain containing, family N member 1 | 132.25 | 140.35 | -0.08 | 8.72E-01 |
| NULL | GSMG0043855 | Klhl17 | kelch-like 17 | 1579.39 | 1140.49 | 0.47 | 8.60E-02 |
| ENSMUSG00000028729 | GSMG0043870 | Ebna1bp2 | EBNA1 binding protein 2 | 1516.97 | 2061.72 | -0.44 | 3.36E-02 |
| NULL | GSMG0043871 | D4Ertd617e | DNA segment, Chr 4, ERATO Doi 617, expressed | 67.00 | 80.77 | -0.28 | 5.71E-01 |
| ENSMUSG00000074264 | GSMG0043876 | Amy1 | amylase 1, salivary | 876.44 | 809.77 | 0.11 | 6.82E-01 |
| ENSMUSG00000027981 | GSMG0043877 | Rnpc3 | RNA-binding region (RNP1, RRM) containing 3 | 1849.62 | 1774.56 | 0.06 | 8.55E-01 |
| ENSMUSG00000028073 | GSMG0043879 | Pear1 | platelet endothelial aggregation receptor 1 | 3937.29 | 3627.41 | 0.12 | 6.81E-01 |
| NULL | GSMG0044468 | NULL | NULL | 223.83 | 186.23 | 0.26 | 4.52E-01 |
| ENSMUSG00000076316 | GSMG0044750 | Mir673 | microRNA 673 | 36.27 | 57.38 | -0.67 | 2.73E-01 |
| NULL | GSMG0045547 | NULL | NULL | 591.65 | 637.45 | -0.11 | 7.49E-01 |
| NULL | GSMG0045556 | NULL | NULL | 226.22 | 218.31 | 0.06 | 8.94E-01 |
| NULL | GSMG0046001 | NULL | NULL | 284.70 | 292.92 | -0.04 | 9.16E-01 |
| NULL | GSMG0046925 | NULL | NULL | 694.62 | 697.79 | -0.01 | 9.81E-01 |
| NULL | GSMG0047294 | NULL | NULL | 279.98 | 347.14 | -0.31 | 4.27E-01 |
| NULL | GSMG0047996 | NULL | NULL | 37.89 | 53.14 | -0.50 | 4.66E-01 |

| | | | | | | | |
|---|---|---|---|---|---|---|---|
| NULL | GSMG0048009 | Msx1os | msh homeobox 1 opposite strand | 259.84 | 174.14 | 0.58 | 6.31E-02 |
| NULL | GSMG0049013 | Peg3os | Peg3 opposite strand | 424.21 | 310.37 | 0.45 | 6.23E-02 |
| NULL | GSMG0050043 | NULL | NULL | 67.06 | 91.30 | -0.45 | 3.78E-01 |
| ENSMUSG00000042581 | GSMG0050824 | Thsd7b | thrombospondin, type I, domain containing 7B | 118.70 | 126.52 | -0.09 | 8.60E-01 |
| ENSMUSG00000055322 | GSMG0050827 | Tns1 | tensin 1 | 2336.85 | 1858.55 | 0.33 | 2.05E-01 |
| ENSMUSG00000067851 | GSMG0050830 | Arfgef1 | ADP-ribosylation factor guanine nucleotide-exchange factor 1(brefeldi | 218.39 | 241.66 | -0.15 | 6.64E-01 |
| ENSMUSG00000050229 | GSMG0050832 | Pigm | phosphatidylinositol glycan anchor biosynthesis, class M | 674.17 | 1147.73 | -0.77 | 5.69E-05 |
| ENSMUSG00000004356 | GSMG0050834 | Utp20 | UTP20 small subunit processome component | 186.52 | 289.26 | -0.64 | 9.38E-03 |
| ENSMUSG00000038594 | GSMG0050835 | Cep85l | centrosomal protein 85-like | 641.07 | 429.88 | 0.58 | 4.69E-03 |
| ENSMUSG00000019856 | GSMG0050836 | Fam184a | family with sequence similarity 184, member A | 78.19 | 103.61 | -0.40 | 3.05E-01 |
| ENSMUSG00000019866 | GSMG0050837 | Aim1 | absent in melanoma 1 | 176.97 | 61.31 | 1.52 | 4.62E-08 |
| ENSMUSG00000055670 | GSMG0050840 | Zzef1 | zinc finger, ZZ-type with EF hand domain 1 | 2573.59 | 2592.72 | -0.01 | 9.77E-01 |
| ENSMUSG00000034667 | GSMG0050843 | Xpot | exportin, tRNA (nuclear export receptor for tRNAs) | 273.98 | 422.54 | -0.62 | 1.36E-03 |
| ENSMUSG00000020181 | GSMG0050844 | Nav3 | neuron navigator 3 | 3389.78 | 2838.18 | 0.26 | 2.43E-01 |
| ENSMUSG00000056342 | GSMG0050845 | Usp34 | ubiquitin specific peptidase 34 | 1746.82 | 1927.45 | -0.14 | 6.00E-01 |
| ENSMUSG00000057335 | GSMG0050848 | Cep170 | centrosomal protein 170 | 190.64 | 362.33 | -0.93 | 6.85E-06 |
| ENSMUSG00000039835 | GSMG0050850 | Nhsl1 | NHS-like 1 | 1757.52 | 1324.14 | 0.41 | 3.85E-02 |
| ENSMUSG00000026721 | GSMG0050852 | Rabgap1l | RAB GTPase activating protein 1-like | 1217.15 | 1273.17 | -0.07 | 8.10E-01 |
| ENSMUSG00000033671 | GSMG0050853 | Cep350 | centrosomal protein 350 | 250.01 | 194.99 | 0.36 | 3.04E-01 |
| ENSMUSG00000032666 | GSMG0050854 | 1700025G04Rik | RIKEN cDNA 1700025G04 gene | 369.70 | 431.02 | -0.22 | 3.95E-01 |
| ENSMUSG00000079283 | GSMG0050855 | 2310009B15Rik | RIKEN cDNA 2310009B15 gene | 256.69 | 295.69 | -0.21 | 5.33E-01 |
| ENSMUSG00000038774 | GSMG0050858 | Ascc3 | activating signal cointegrator 1 complex subunit 3 | 540.55 | 513.83 | 0.07 | 8.12E-01 |
| ENSMUSG00000035021 | GSMG0050859 | Baz1a | bromodomain adjacent to zinc finger domain 1A | 223.33 | 249.11 | -0.15 | 6.46E-01 |
| ENSMUSG00000035247 | GSMG0050860 | Hectd1 | HECT domain containing 1 | 5264.01 | 5920.04 | -0.17 | 3.92E-01 |
| ENSMUSG00000063632 | GSMG0050861 | Sox11 | SRY (sex determining region Y)-box 11 | 348.18 | 1634.14 | -2.24 | 3.42E-28 |
| ENSMUSG00000039989 | GSMG0050863 | Cbx4 | chromobox 4 | 464.53 | 437.51 | 0.09 | 7.90E-01 |
| ENSMUSG00000052812 | GSMG0050864 | Atad2b | ATPase family, AAA domain containing 2B | 230.45 | 215.48 | 0.09 | 8.35E-01 |
| ENSMUSG00000020576 | GSMG0050866 | Nbas | neuroblastoma amplified sequence | 243.79 | 276.87 | -0.19 | 5.86E-01 |
| ENSMUSG00000063450 | GSMG0050868 | Syne2 | spectrin repeat containing, nuclear envelope 2 | 822.90 | 961.77 | -0.22 | 2.65E-01 |
| ENSMUSG00000029878 | GSMG0050869 | Dbpht2 | DNA binding protein with his-thr domain | 923.15 | 1143.38 | -0.31 | 1.84E-01 |
| ENSMUSG00000035431 | GSMG0050870 | Sstr1 | somatostatin receptor 1 | 552.39 | 799.85 | -0.54 | 5.42E-03 |
| ENSMUSG00000018474 | GSMG0050875 | Chd3 | chromodomain helicase DNA binding protein 3 | 1198.86 | 1380.18 | -0.20 | 3.32E-01 |
| ENSMUSG00000020899 | GSMG0050876 | Pfas | phosphoribosylformylglycinamidine synthase (FGAR amidotransferase | 735.86 | 988.43 | -0.43 | 1.78E-02 |

| | | | | | | | |
|---|---|---|---|---|---|---|---|
| ENSMUSG00000044072 | GSMG0050878 | Eml6 | echinoderm microtubule associated protein like 6 | 379.36 | 295.35 | 0.36 | 1.78E-01 |
| ENSMUSG00000034156 | GSMG0050879 | Bzrap1 | benzodiazepine receptor associated protein 1 | 1304.70 | 1263.51 | 0.05 | 9.20E-01 |
| ENSMUSG00000094483 | GSMG0050880 | Purb | purine rich element binding protein B | 4174.10 | 2515.63 | 0.73 | 8.89E-05 |
| ENSMUSG00000020728 | GSMG0050883 | Cep112 | centrosomal protein 112 | 99.21 | 104.23 | -0.07 | 8.90E-01 |
| ENSMUSG00000060012 | GSMG0050884 | Kif13b | kinesin family member 13B | 1274.70 | 998.67 | 0.35 | 1.84E-01 |
| ENSMUSG00000034235 | GSMG0050887 | Usp54 | ubiquitin specific peptidase 54 | 1898.95 | 1031.72 | 0.88 | 6.05E-08 |
| ENSMUSG00000025278 | GSMG0050890 | Flnb | filamin, beta | 1289.48 | 1049.57 | 0.30 | 1.66E-01 |
| ENSMUSG00000060147 | GSMG0050895 | Serpinb6a | serine (or cysteine) peptidase inhibitor, clade B, member 6a | 862.07 | 630.52 | 0.45 | 2.32E-02 |
| ENSMUSG00000021597 | GSMG0050897 | Slf1 | SMC5-SMC6 complex localization factor 1 | 126.79 | 213.39 | -0.75 | 2.28E-02 |
| ENSMUSG00000034525 | GSMG0050898 | Ice1 | interactor of little elongation complex ELL subunit 1 | 523.18 | 586.63 | -0.16 | 6.04E-01 |
| ENSMUSG00000021470 | GSMG0050900 | Ercc6l2 | excision repair cross-complementing rodent repair deficiency, complem | 392.60 | 429.14 | -0.13 | 6.88E-01 |
| ENSMUSG00000039929 | GSMG0050901 | Urb1 | URB1 ribosome biogenesis 1 homolog (S. cerevisiae) | 293.88 | 361.10 | -0.29 | 2.98E-01 |
| ENSMUSG00000060657 | GSMG0050902 | Marf1 | meiosis arrest female 1 | 394.89 | 364.52 | 0.12 | 7.30E-01 |
| ENSMUSG00000000326 | GSMG0050903 | Comt | catechol-O-methyltransferase | 1144.70 | 1087.77 | 0.07 | 8.00E-01 |
| ENSMUSG00000022533 | GSMG0050904 | Atp13a3 | ATPase type 13A3 | 1905.44 | 1376.64 | 0.47 | 2.10E-02 |
| ENSMUSG00000061751 | GSMG0050905 | Kalrn | kalirin, RhoGEF kinase | 3023.20 | 927.19 | 1.71 | 1.53E-16 |
| ENSMUSG00000022957 | GSMG0050906 | Itsn1 | intersectin 1 (SH3 domain protein 1A) | 423.96 | 539.90 | -0.35 | 9.79E-02 |
| ENSMUSG00000039789 | GSMG0050907 | Zfp597 | zinc finger protein 597 | 190.97 | 180.41 | 0.09 | 8.33E-01 |
| ENSMUSG00000046613 | GSMG0050910 | Vwa5b2 | von Willebrand factor A domain containing 5B2 | 405.13 | 343.94 | 0.23 | 4.70E-01 |
| ENSMUSG00000075254 | GSMG0050911 | Heg1 | heart development protein with EGF-like domains 1 | 19955.25 | 12111.03 | 0.72 | 9.73E-05 |
| ENSMUSG00000048154 | GSMG0050913 | Kmt2d | lysine (K)-specific methyltransferase 2D | 1930.51 | 1803.57 | 0.10 | 7.75E-01 |
| ENSMUSG00000051786 | GSMG0050915 | Tubgcp6 | tubulin, gamma complex associated protein 6 | 2456.13 | 2528.33 | -0.04 | 9.17E-01 |
| ENSMUSG00000036800 | GSMG0050916 | Fam135b | family with sequence similarity 135, member B | 201.88 | 195.41 | 0.05 | 9.13E-01 |
| ENSMUSG00000022311 | GSMG0050917 | Csmd3 | CUB and Sushi multiple domains 3 | 1413.34 | 1787.27 | -0.34 | 5.86E-02 |
| ENSMUSG00000044224 | GSMG0050918 | Dnajc21 | DnaJ heat shock protein family (Hsp40) member C21 | 671.02 | 552.06 | 0.28 | 3.01E-01 |
| ENSMUSG00000033237 | GSMG0050919 | Arid2 | AT rich interactive domain 2 (ARID, RFX-like) | 288.57 | 316.62 | -0.13 | 6.69E-01 |
| ENSMUSG00000037646 | GSMG0050920 | Vps13b | vacuolar protein sorting 13B | 646.26 | 580.82 | 0.16 | 6.91E-01 |
| ENSMUSG00000039801 | GSMG0050921 | 2410089E03Rik | RIKEN cDNA 2410089E03 gene | 879.49 | 1030.96 | -0.23 | 3.09E-01 |
| ENSMUSG00000042104 | GSMG0050923 | Uggt2 | UDP-glucose glycoprotein glucosyltransferase 2 | 136.95 | 169.87 | -0.31 | 3.36E-01 |
| ENSMUSG00000000197 | GSMG0050925 | Nalcn | sodium leak channel, non-selective | 87.37 | 97.40 | -0.15 | 7.90E-01 |
| ENSMUSG00000022016 | GSMG0050926 | Akap11 | A kinase (PRKA) anchor protein 11 | 524.38 | 512.31 | 0.04 | 9.38E-01 |
| G00000025427,ENSMUSG000 | GSMG0050927 | Rnf165 // 8030462 | ring finger protein 165 // RIKEN cDNA 8030462N17 gene | 220.71 | 150.42 | 0.55 | 9.59E-02 |
| ENSMUSG00000048410 | GSMG0050928 | Zfp407 | zinc finger protein 407 | 44.78 | 50.74 | -0.17 | 8.11E-01 |

| | | | | | | | |
|---|---|---|---|---|---|---|---|
| ENSMUSG00000024350 | GSMG0050929 | Dnajc18 | DnaJ heat shock protein family (Hsp40) member C18 | 575.34 | 633.79 | -0.14 | 5.88E-01 |
| ENSMUSG00000033632 | GSMG0050930 | AW554918 | expressed sequence AW554918 | 93.15 | 122.19 | -0.40 | 2.76E-01 |
| ENSMUSG00000073563 | GSMG0050931 | Csnk1g3 | casein kinase 1, gamma 3 | 1937.68 | 2044.63 | -0.08 | 7.72E-01 |
| ENSMUSG00000024542 | GSMG0050932 | Cep192 | centrosomal protein 192 | 254.40 | 378.09 | -0.57 | 5.21E-03 |
| ENSMUSG00000024054 | GSMG0050935 | Smchd1 | SMC hinge domain containing 1 | 356.33 | 408.58 | -0.20 | 4.57E-01 |
| ENSMUSG00000024135 | GSMG0050936 | Srbd1 | S1 RNA binding domain 1 | 42.51 | 60.43 | -0.50 | 3.92E-01 |
| ENSMUSG00000071984 | GSMG0050939 | Fndc1 | fibronectin type III domain containing 1 | 100.30 | 208.13 | -1.05 | 8.61E-05 |
| NULL | GSMG0050945 | NULL | NULL | 80088.83 | 80307.06 | 0.00 | 9.92E-01 |
| ENSMUSG00000041565 | GSMG0050946 | L3mbtl4 | l(3)mbt-like 4 (Drosophila) | 131.58 | 470.73 | -1.84 | 4.43E-19 |
| ENSMUSG00000071054 | GSMG0050948 | Safb | scaffold attachment factor B | 4526.04 | 4594.60 | -0.02 | 9.38E-01 |
| ENSMUSG00000043683 | GSMG0050949 | Fem1a | feminization 1 homolog a (C. elegans) | 1558.22 | 1860.98 | -0.26 | 2.68E-01 |
| ENSMUSG00000023826 | GSMG0050954 | Park2 | Parkinson disease (autosomal recessive, juvenile) 2, parkin | 81.75 | 68.28 | 0.26 | 6.11E-01 |
| G00000048731,ENSMUSG000 | GSMG0050955 | Zbtb9 // Ggnbp1 | zinc finger and BTB domain containing 9 // gametogenetin binding prot | 73.92 | 48.26 | 0.61 | 1.83E-01 |
| ENSMUSG00000027288 | GSMG0050956 | Zfp106 | zinc finger protein 106 | 867.41 | 900.51 | -0.05 | 8.97E-01 |
| ENSMUSG00000037110 | GSMG0050959 | Ralgapa2 | Ral GTPase activating protein, alpha subunit 2 (catalytic) | 288.33 | 397.56 | -0.46 | 9.77E-02 |
| ENSMUSG00000057378 | GSMG0050960 | Ryr3 | ryanodine receptor 3 | 1433.63 | 1713.63 | -0.26 | 2.73E-01 |
| ENSMUSG00000028053 | GSMG0050962 | Ash1l | ash1 (absent, small, or homeotic)-like (Drosophila) | 3044.69 | 3243.67 | -0.09 | 7.14E-01 |
| ENSMUSG00000037270 | GSMG0050964 | 4932438A13Rik | RIKEN cDNA 4932438A13 gene | 4423.65 | 4397.78 | 0.01 | 9.78E-01 |
| ENSMUSG00000028030 | GSMG0050974 | Tbck | TBC1 domain containing kinase | 179.74 | 195.53 | -0.11 | 7.88E-01 |
| ENSMUSG00000041235 | GSMG0050976 | Chd7 | chromodomain helicase DNA binding protein 7 | 1323.43 | 1453.36 | -0.14 | 5.30E-01 |
| ENSMUSG00000039735 | GSMG0050977 | Fnbp1l | formin binding protein 1-like | 1191.14 | 2233.72 | -0.91 | 5.61E-05 |
| ENSMUSG00000024811 | GSMG0050982 | Tnks2 | tankyrase, TRF1-interacting ankyrin-related ADP-ribose polymerase 2 | 2205.37 | 2144.82 | 0.04 | 9.03E-01 |
| ENSMUSG00000075318 | GSMG0050984 | Scn2a1 | sodium channel, voltage-gated, type II, alpha 1 | 1001.05 | 1605.01 | -0.68 | 1.89E-05 |
| ENSMUSG00000035168 | GSMG0050985 | Tanc1 | tetratricopeptide repeat, ankyrin repeat and coiled-coil containing 1 | 1752.44 | 1442.42 | 0.28 | 2.25E-01 |
| ENSMUSG00000027254 | GSMG0050988 | Map1a | microtubule-associated protein 1 A | 980.05 | 603.89 | 0.70 | 1.99E-04 |
| ENSMUSG00000068205 | GSMG0050989 | Macrod2 | MACRO domain containing 2 | 998.76 | 1471.95 | -0.56 | 2.70E-03 |
| ENSMUSG00000026924 | GSMG0050990 | Sec16a | SEC16 homolog A, endoplasmic reticulum export factor | 499.07 | 560.89 | -0.17 | 5.31E-01 |
| ENSMUSG00000064329 | GSMG0050992 | Scn1a | sodium channel, voltage-gated, type I, alpha | 411.63 | 346.17 | 0.25 | 3.45E-01 |
| ENSMUSG00000057182 | GSMG0050993 | Scn3a | sodium channel, voltage-gated, type III, alpha | 738.18 | 1588.50 | -1.11 | 1.96E-12 |
| ENSMUSG00000029202 | GSMG0050996 | Pds5a | PDS5 cohesin associated factor A | 1898.94 | 1870.42 | 0.02 | 9.50E-01 |
| ENSMUSG00000054280 | GSMG0050997 | Prr14l | proline rich 14-like | 1294.84 | 1238.96 | 0.06 | 8.35E-01 |
| ENSMUSG00000061755 | GSMG0050998 | Bod1l | biorientation of chromosomes in cell division 1-like | 3289.15 | 3012.53 | 0.13 | 6.20E-01 |
| ENSMUSG00000033542 | GSMG0051002 | Arhgef5 | Rho guanine nucleotide exchange factor (GEF) 5 | 3041.42 | 3314.52 | -0.12 | 5.93E-01 |

| | | | | | | | |
|---|---|---|---|---|---|---|---|
| ENSMUSG00000029563 | GSMG0051003 | Foxp2 | forkhead box P2 | 128.45 | 190.97 | -0.57 | 7.25E-02 |
| ENSMUSG00000039578 | GSMG0051004 | Ccser1 | coiled-coil serine rich 1 | 473.01 | 528.80 | -0.16 | 6.33E-01 |
| ENSMUSG00000034312 | GSMG0051007 | Iqsec1 | IQ motif and Sec7 domain 1 | 466.11 | 411.65 | 0.18 | 6.42E-01 |
| ENSMUSG00000034832 | GSMG0051008 | Tet3 | tet methylcytosine dioxygenase 3 | 1616.25 | 1503.30 | 0.11 | 7.00E-01 |
| ENSMUSG00000052861 | GSMG0051009 | Dnah6 | dynein, axonemal, heavy chain 6 | 378.33 | 272.98 | 0.47 | 6.61E-02 |
| ENSMUSG00000038070 | GSMG0051011 | Cntln | centlein, centrosomal protein | 893.26 | 979.87 | -0.13 | 5.85E-01 |
| ENSMUSG00000028514 | GSMG0051012 | Usp24 | ubiquitin specific peptidase 24 | 516.04 | 517.83 | 0.00 | 9.90E-01 |
| ENSMUSG00000028634 | GSMG0051013 | Hivep3 | human immunodeficiency virus type I enhancer binding protein 3 | 2000.11 | 2215.19 | -0.15 | 6.09E-01 |
| ENSMUSG00000043333 | GSMG0051014 | Rhbdl2 | rhomboid, veinlet-like 2 (Drosophila) | 295.97 | 481.45 | -0.70 | 5.09E-04 |
| ENSMUSG00000028876 | GSMG0051016 | Epha10 | Eph receptor A10 | 274.87 | 116.12 | 1.24 | 8.24E-06 |
| ENSMUSG00000028820 | GSMG0051017 | Sfpq | splicing factor proline/glutamine rich (polypyrimidine tract binding pro | 4328.08 | 5315.38 | -0.30 | 1.73E-01 |
| G00000066036,ENSMUSG000 | GSMG0051018 | Emc1 // Ubr4 | ER membrane protein complex subunit 1 // ubiquitin protein ligase E3 | 8592.90 | 10506.32 | -0.29 | 1.46E-01 |
| ENSMUSG00000001089 | GSMG0051019 | Luzp1 | leucine zipper protein 1 | 7730.49 | 6247.02 | 0.31 | 5.29E-02 |
| ENSMUSG00000057751 | GSMG0051022 | Megf6 | multiple EGF-like-domains 6 | 6145.79 | 6443.46 | -0.07 | 8.67E-01 |
| ENSMUSG00000073910 | GSMG0051023 | Mob3b | MOB kinase activator 3B | 238.76 | 187.63 | 0.35 | 2.47E-01 |
| ENSMUSG00000045083 | GSMG0051025 | Lingo2 | leucine rich repeat and Ig domain containing 2 | 2678.72 | 2448.71 | 0.13 | 5.76E-01 |
| ENSMUSG00000014592 | GSMG0051028 | Camta1 | calmodulin binding transcription activator 1 | 1255.02 | 2005.80 | -0.68 | 6.06E-06 |
| ENSMUSG00000020220 | GSMG0051029 | Vps13d | vacuolar protein sorting 13D | 70.64 | 62.19 | 0.18 | 7.39E-01 |
| ENSMUSG00000048271 | GSMG0051032 | Rbm33 | RNA binding motif protein 33 | 687.08 | 755.14 | -0.14 | 7.64E-01 |
| ENSMUSG00000029376 | GSMG0051034 | Mthfd2l | methylenetetrahydrofolate dehydrogenase (NADP+ dependent) 2-like | 208.49 | 280.36 | -0.43 | 8.86E-02 |
| ENSMUSG00000042744 | GSMG0051035 | Gm15800 | predicted gene 15800 | 279.88 | 286.52 | -0.03 | 9.43E-01 |
| ENSMUSG00000029263 | GSMG0051036 | Pigg | phosphatidylinositol glycan anchor biosynthesis, class G | 271.59 | 189.90 | 0.52 | 5.91E-02 |
| ENSMUSG00000033209 | GSMG0051037 | Ttc28 | tetratricopeptide repeat domain 28 | 516.40 | 685.95 | -0.41 | 1.77E-01 |
| ENSMUSG00000056602 | GSMG0051040 | Fry | FRY microtubule binding protein | 1537.62 | 1072.79 | 0.52 | 7.66E-04 |
| ENSMUSG00000031841 | GSMG0051041 | Cdh13 | cadherin 13 | 769.83 | 1324.99 | -0.78 | 9.26E-06 |
| ENSMUSG00000030655 | GSMG0051044 | Smg1 | SMG1 homolog, phosphatidylinositol 3-kinase-related kinase (C. elegan | 290.59 | 242.76 | 0.26 | 3.94E-01 |
| ENSMUSG00000036862 | GSMG0051045 | Dchs1 | dachsous 1 (Drosophila) | 256.82 | 541.40 | -1.08 | 1.47E-01 |
| ENSMUSG00000038371 | GSMG0051046 | Sbf2 | SET binding factor 2 | 2531.94 | 5045.80 | -0.99 | 4.72E-12 |
| ENSMUSG00000034973 | GSMG0051047 | Dopey1 | dopey family member 1 | 1926.85 | 1729.01 | 0.16 | 4.84E-01 |
| ENSMUSG00000036972 | GSMG0051048 | Zic4 | zinc finger protein of the cerebellum 4 | 232.64 | 194.63 | 0.26 | 4.04E-01 |
| ENSMUSG00000032409 | GSMG0051049 | Atr | ataxia telangiectasia and Rad3 related | 231.62 | 293.86 | -0.34 | 1.80E-01 |
| ENSMUSG00000053641 | GSMG0051050 | Dennd4a | DENN/MADD domain containing 4A | 177.78 | 142.43 | 0.33 | 3.46E-01 |
| ENSMUSG00000038664 | GSMG0051051 | Herc1 | HECT and RLD domain containing E3 ubiquitin protein ligase family r | 5417.07 | 4926.52 | 0.14 | 5.42E-01 |

| | | | | | | | |
|---|---|---|---|---|---|---|---|
| ENSMUSG00000040204 | GSMG0051052 | 2810417H13Rik | RIKEN cDNA 2810417H13 gene | 248.04 | 698.93 | -1.49 | 3.42E-07 |
| NULL | GSMG0051053 | 9530091C08Rik | RIKEN cDNA 9530091C08 gene | 2033.38 | 1653.55 | 0.30 | 1.62E-01 |
| ENSMUSG00000039585 | GSMG0051054 | Myo9a | myosin IXa | 101.67 | 116.43 | -0.20 | 6.53E-01 |
| ENSMUSG00000074030 | GSMG0051056 | Exoc8 | exocyst complex component 8 | 1761.67 | 1410.27 | 0.32 | 2.19E-01 |
| ENSMUSG00000051451 | GSMG0051060 | Crebzf | CREB/ATF bZIP transcription factor | 4752.42 | 3292.39 | 0.53 | 5.75E-03 |
| ENSMUSG00000052572 | GSMG0051061 | Dlg2 | discs, large homolog 2 (Drosophila) | 219.51 | 269.36 | -0.29 | 3.16E-01 |
| ENSMUSG00000066406 | GSMG0051063 | Akap13 | A kinase (PRKA) anchor protein 13 | 5168.02 | 5076.01 | 0.03 | 9.21E-01 |
| ENSMUSG00000052512 | GSMG0051064 | Nav2 | neuron navigator 2 | 753.83 | 1312.96 | -0.80 | 1.51E-04 |
| ENSMUSG00000038738 | GSMG0051065 | Shank1 | SH3/ankyrin domain gene 1 | 392.16 | 283.94 | 0.46 | 1.55E-01 |
| ENSMUSG00000035713 | GSMG0051070 | Usp35 | ubiquitin specific peptidase 35 | 231.11 | 187.09 | 0.30 | 3.48E-01 |
| G00000078671,ENSMUSG000 | GSMG0051071 | Chd2 // 1810026B | chromodomain helicase DNA binding protein 2 // RIKEN cDNA 18100 | 1797.42 | 1716.44 | 0.07 | 7.93E-01 |
| ENSMUSG00000030880 | GSMG0051077 | Polr3e | polymerase (RNA) III (DNA directed) polypeptide E | 916.98 | 999.61 | -0.12 | 6.14E-01 |
| ENSMUSG00000030779 | GSMG0051078 | Rbbp6 | retinoblastoma binding protein 6 | 2423.14 | 2581.69 | -0.09 | 7.33E-01 |
| ENSMUSG00000041268 | GSMG0051081 | Dmxl2 | Dmx-like 2 | 478.86 | 402.35 | 0.25 | 4.20E-01 |
| ENSMUSG00000043154 | GSMG0051082 | Ppp2r3a | protein phosphatase 2, regulatory subunit B", alpha | 208.00 | 215.99 | -0.06 | 8.90E-01 |
| ENSMUSG00000062296 | GSMG0051090 | Trank1 | tetratricopeptide repeat and ankyrin repeat containing 1 | 6877.37 | 4595.16 | 0.58 | 9.89E-04 |
| ENSMUSG00000044791 | GSMG0051091 | Setd2 | SET domain containing 2 | 199.31 | 210.08 | -0.07 | 8.57E-01 |
| ENSMUSG00000046111 | GSMG0051092 | Cep295 | centrosomal protein 295 | 242.74 | 297.56 | -0.29 | 3.21E-01 |
| ENSMUSG00000039488 | GSMG0051093 | Cntn5 | contactin 5 | 422.69 | 787.14 | -0.90 | 5.96E-07 |
| ENSMUSG00000037475 | GSMG0051094 | Thoc2 | THO complex 2 | 578.89 | 628.19 | -0.12 | 6.99E-01 |
| ENSMUSG00000086503 | GSMG0051096 | Xist | inactive X specific transcripts | 758.28 | 795.00 | -0.07 | 8.26E-01 |
| ENSMUSG00000038323 | GSMG0051104 | 1700066M21Rik | RIKEN cDNA 1700066M21 gene | 439.29 | 554.12 | -0.33 | 1.24E-01 |
| ENSMUSG00000026014 | GSMG0051119 | Raph1 | Ras association (RalGDS/AF-6) and pleckstrin homology domains 1 | 1807.27 | 2391.48 | -0.40 | 4.35E-02 |
| ENSMUSG00000026162 | GSMG0051120 | Nhej1 | nonhomologous end-joining factor 1 | 95.69 | 88.81 | 0.11 | 8.33E-01 |
| ENSMUSG00000111356 | GSMG0051136 | Rpl32l | ribosomal protein L32-like | 174.72 | 255.86 | -0.56 | 4.76E-02 |
| ENSMUSG00000048701 | GSMG0051140 | Ccdc6 | coiled-coil domain containing 6 | 512.83 | 590.39 | -0.21 | 4.09E-01 |
| ENSMUSG00000112639 | GSMG0051164 | A730063M14Rik | RIKEN cDNA A730063M14 gene | 77.98 | 99.14 | -0.35 | 4.34E-01 |
| ENSMUSG00000063760 | GSMG0051181 | Rnf217 | ring finger protein 217 | 264.42 | 250.34 | 0.08 | 8.57E-01 |
| ENSMUSG00000096687 | GSMG0051185 | Mfsd4b4 | major facilitator superfamily domain containing 4B4 | 52.26 | 70.76 | -0.44 | 3.60E-01 |
| ENSMUSG00000020272 | GSMG0051219 | Stk10 | serine/threonine kinase 10 | 331.11 | 290.03 | 0.19 | 5.22E-01 |
| ENSMUSG00000065126 | GSMG0051248 | Snord104 | small nucleolar RNA, C/D box 104 | 108.78 | 91.13 | 0.25 | 6.91E-01 |
| ENSMUSG00000060923 | GSMG0051256 | Acyp2 | acylphosphatase 2, muscle type | 173.74 | 135.30 | 0.36 | 4.63E-01 |
| ENSMUSG00000041449 | GSMG0051317 | Serpina3h | serine (or cysteine) peptidase inhibitor, clade A, member 3H | 188.92 | 531.30 | -1.49 | 3.71E-08 |

| | | | | | | | |
|---|---|---|---|---|---|---|---|
| ENSMUSG00000087370 | GSMG0051357 | Tmem170b | transmembrane protein 170B | 1043.17 | 1258.57 | -0.27 | 2.02E-01 |
| ENSMUSG00000038132 | GSMG0051358 | Rbm24 | RNA binding motif protein 24 | 460.84 | 386.26 | 0.25 | 4.03E-01 |
| ENSMUSG00000100891 | GSMG0051361 | 2810049E08Rik | RIKEN cDNA 2810049E08 gene | 114.54 | 77.86 | 0.56 | 1.31E-01 |
| ENSMUSG00000092741 | GSMG0051385 | Mir3074-1 | microRNA 3074-1 | 53.21 | 30.21 | 0.82 | 1.20E-01 |
| ENSMUSG00000078958 | GSMG0051386 | Atp6ap1l | ATPase, H+ transporting, lysosomal accessory protein 1-like | 119.87 | 27.77 | 2.12 | 6.72E-09 |
| ENSMUSG00000021619 | GSMG0051387 | Atg10 | autophagy related 10 | 238.72 | 214.51 | 0.16 | 6.88E-01 |
| ENSMUSG00000021708 | GSMG0051388 | Rasgrf2 | RAS protein-specific guanine nucleotide-releasing factor 2 | 776.97 | 1429.86 | -0.88 | 8.17E-05 |
| NULL | GSMG0051407 | Gm9199 | predicted gene 9199 | 142.35 | 40.85 | 1.80 | 8.74E-09 |
| ENSMUSG00000025370 | GSMG0051439 | Cdh9 | cadherin 9 | 335.55 | 369.91 | -0.15 | 6.66E-01 |
| ENSMUSG00000058656 | GSMG0051457 | Samd12 | sterile alpha motif domain containing 12 | 204.21 | 122.82 | 0.74 | 6.93E-03 |
| ENSMUSG00000098176 | GSMG0051459 | Ccdc166 | coiled-coil domain containing 166 | 258.25 | 247.89 | 0.06 | 8.90E-01 |
| ENSMUSG00000071561 | GSMG0051488 | BC100530 | cDNA sequence BC100530 | 12.92 | 52.64 | -2.08 | 6.68E-04 |
| ENSMUSG00000069729 | GSMG0051508 | Arid1b | AT rich interactive domain 1B (SWI-like) | 1872.12 | 2098.09 | -0.16 | 5.84E-01 |
| ENSMUSG00000067722 | GSMG0051534 | BC003965 | cDNA sequence BC003965 | 1608.75 | 1674.47 | -0.06 | 8.47E-01 |
| ENSMUSG00000091586 | GSMG0051535 | Cyp4f17 | cytochrome P450, family 4, subfamily f, polypeptide 17 | 193.41 | 201.49 | -0.06 | 8.83E-01 |
| ENSMUSG00000059852 | GSMG0051600 | Kcng2 | potassium voltage-gated channel, subfamily G, member 2 | 188.85 | 329.80 | -0.81 | 1.00E-03 |
| ENSMUSG00000087679 | GSMG0051745 | C330006A16Rik | RIKEN cDNA C330006A16 gene | 1507.09 | 1888.70 | -0.33 | 1.23E-01 |
| NULL | GSMG0051913 | NULL | NULL | 570.70 | 529.70 | 0.11 | 7.38E-01 |
| ENSMUSG00000070720 | GSMG0051922 | Tmem200b | transmembrane protein 200B | 849.27 | 138.40 | 2.61 | 1.18E-29 |
| ENSMUSG00000070583 | GSMG0051930 | Fv1 | Friend virus susceptibility 1 | 209.64 | 128.36 | 0.71 | 8.47E-03 |
| ENSMUSG00000078719 | GSMG0051936 | Msmp | microseminoprotein, prostate associated | 97.39 | 66.80 | 0.54 | 1.54E-01 |
| ENSMUSG00000029359 | GSMG0051991 | Tesc | tescalcin | 205.93 | 36.42 | 2.49 | 1.20E-14 |
| ENSMUSG00000050822 | GSMG0051995 | Slc29a4 | solute carrier family 29 (nucleoside transporters), member 4 | 1775.65 | 2940.16 | -0.73 | 1.07E-05 |
| ENSMUSG00000084235 | GSMG0052007 | Gm15421 | predicted gene 15421 | 344.03 | 476.82 | -0.47 | 1.43E-01 |
| ENSMUSG00000029196 | GSMG0052010 | Tada2b | transcriptional adaptor 2B | 1006.30 | 457.73 | 1.14 | 8.95E-02 |
| ENSMUSG00000045790 | GSMG0052012 | Ccdc149 | coiled-coil domain containing 149 | 1136.79 | 877.57 | 0.37 | 6.77E-02 |
| ENSMUSG00000024437 | GSMG0052034 | Gm8615 | predicted pseudogene 8615 | 956.38 | 745.39 | 0.36 | 5.89E-02 |
| ENSMUSG00000106738 | GSMG0052035 | Gm5 | predicted gene 5 | 207.97 | 108.71 | 0.93 | 7.08E-04 |
| ENSMUSG00000079598 | GSMG0052040 | Clec2l | C-type lectin domain family 2, member L | 1018.13 | 702.38 | 0.54 | 2.68E-02 |
| ENSMUSG00000090698 | GSMG0052074 | Apold1 | apolipoprotein L domain containing 1 | 7362.42 | 2513.49 | 1.55 | 1.56E-14 |
| ENSMUSG00000053367 | GSMG0052114 | Gm6792 | predicted gene 6792 | 63.76 | 96.68 | -0.59 | 1.76E-01 |
| ENSMUSG00000041117 | GSMG0052129 | Ccdc8 | coiled-coil domain containing 8 | 607.14 | 844.36 | -0.48 | 1.08E-02 |
| ENSMUSG00000044997 | GSMG0052158 | E130304I02Rik | RIKEN cDNA E130304I02 gene | 68.18 | 72.94 | -0.10 | 8.65E-01 |

| | | | | | | | |
|---|---|---|---|---|---|---|---|
| ENSMUSG00000057706 | GSMG0052164 | Mex3b | mex3 RNA binding family member B | 298.37 | 437.20 | -0.55 | 8.23E-03 |
| ENSMUSG00000090958 | GSMG0052176 | Lrrc32 | leucine rich repeat containing 32 | 801.21 | 575.50 | 0.48 | 2.65E-02 |
| ENSMUSG00000089987 | GSMG0052178 | Gpx2-ps1 | glutathione peroxidase 2, pseudogene 1 | 105.66 | 60.85 | 0.80 | 1.65E-02 |
| ENSMUSG00000078591 | GSMG0052212 | Hs3st4 | heparan sulfate (glucosamine) 3-O-sulfotransferase 4 | 416.87 | 284.03 | 0.55 | 2.16E-02 |
| ENSMUSG00000064767 | GSMG0052303 | Snord35b | small nucleolar RNA, C/D box 35B | 80.99 | 17.56 | 2.20 | 4.01E-08 |
| ENSMUSG00000079003 | GSMG0052397 | Samd1 | sterile alpha motif domain containing 1 | 216.25 | 181.50 | 0.25 | 6.11E-01 |
| ENSMUSG00000019362 | GSMG0052416 | D8Ertd738e | DNA segment, Chr 8, ERATO Doi 738, expressed | 2015.59 | 2085.18 | -0.05 | 8.60E-01 |
| ENSMUSG00000069895 | GSMG0052420 | Atxn1l | ataxin 1-like | 703.71 | 699.97 | 0.01 | 9.88E-01 |
| ENSMUSG00000014633 | GSMG0052421 | Cmc2 | COX assembly mitochondrial protein 2 | 447.57 | 471.46 | -0.08 | 8.37E-01 |
| ENSMUSG00000090290 | GSMG0052423 | Gm17296 | predicted gene, 17296 | 522.92 | 453.32 | 0.21 | 5.23E-01 |
| ENSMUSG00000051495 | GSMG0052424 | Irf2bp2 | interferon regulatory factor 2 binding protein 2 | 1715.88 | 1149.55 | 0.58 | 2.55E-02 |
| ENSMUSG00000035048 | GSMG0052490 | Anapc13 | anaphase promoting complex subunit 13 | 415.21 | 453.90 | -0.13 | 7.17E-01 |
| ENSMUSG00000066456 | GSMG0052518 | Hmgn3 | high mobility group nucleosomal binding domain 3 | 2388.87 | 2442.32 | -0.03 | 9.36E-01 |
| ENSMUSG00000070287 | GSMG0052521 | Slc35g2 | solute carrier family 35, member G2 | 182.78 | 113.55 | 0.69 | 1.53E-02 |
| ENSMUSG00000039607 | GSMG0052524 | Rbms3 | RNA binding motif, single stranded interacting protein | 1034.97 | 885.36 | 0.22 | 4.23E-01 |
| ENSMUSG00000047485 | GSMG0052586 | Klhl34 | kelch-like 34 | 302.78 | 278.88 | 0.12 | 7.69E-01 |
| ENSMUSG00000090546 | GSMG0052621 | Cdr1 | cerebellar degeneration related antigen 1 | 1869.34 | 1093.32 | 0.77 | 6.21E-03 |
| ENSMUSG00000041670 | GSMG0052680 | Rims1 | regulating synaptic membrane exocytosis 1 | 769.42 | 486.98 | 0.66 | 1.10E-02 |
| ENSMUSG00000041670 | GSMG0052681 | Rims1 | regulating synaptic membrane exocytosis 1 | 1226.69 | 927.08 | 0.40 | 3.00E-02 |
| ENSMUSG00000022536 | GSMG0052896 | Glyr1 | glyoxylate reductase 1 homolog (Arabidopsis) | 1306.27 | 1430.36 | -0.13 | 5.82E-01 |
| ENSMUSG00000066057 | GSMG0052996 | Gm1976 | predicted gene 1976 | 169.77 | 182.62 | -0.11 | 7.82E-01 |
| ENSMUSG00000006740 | GSMG0053004 | Kif5b | kinesin family member 5B | 2681.62 | 2588.82 | 0.05 | 8.76E-01 |
| ENSMUSG00000025195 | GSMG0053034 | Dnmbp | dynamin binding protein | 369.64 | 443.91 | -0.27 | 2.83E-01 |
| ENSMUSG00000072704 | GSMG0053234 | Smim10l1 | small integral membrane protein 10 like 1 | 1036.35 | 917.16 | 0.18 | 4.72E-01 |
| ENSMUSG00000071302 | GSMG0053253 | 2610044O15Rik8 | RIKEN cDNA 2610044O15 gene | 315.20 | 256.92 | 0.30 | 2.51E-01 |
| NULL | GSMG0053277 | Pisd-ps3 | phosphatidylserine decarboxylase, pseudogene 3 | 8891.98 | 4820.46 | 0.88 | 6.65E-08 |
| ENSMUSG00000031791 | GSMG0053359 | Tmem38a | transmembrane protein 38A | 333.86 | 153.51 | 1.12 | 4.05E-06 |
| ENSMUSG00000065512 | GSMG0053367 | Mir138-2 | microRNA 138-2 | 613.07 | 269.93 | 1.18 | 1.99E-11 |
| NULL | GSMG0054384 | NULL | NULL | 152.73 | 145.60 | 0.07 | 8.90E-01 |
| NULL | GSMG0054663 | NULL | NULL | 656.25 | 1091.87 | -0.74 | 3.75E-04 |
| NULL | GSMG0055599 | NULL | NULL | 4777.15 | 2915.08 | 0.71 | 1.30E-07 |
| NULL | GSMG0056159 | NULL | NULL | 180.24 | 162.42 | 0.15 | 7.02E-01 |
| NULL | GSMG0056413 | NULL | NULL | 174.86 | 236.32 | -0.44 | 1.31E-01 |

| | | | | | | | |
|---|---|---|---|---|---|---|---|
| NULL | GSMG0056921 | NULL | NULL | 46.12 | 54.84 | -0.26 | 6.59E-01 |
| ENSMUSG00000095567 | GSMG0057008 | Noc2l | NOC2 like nucleolar associated transcriptional repressor | 900.59 | 1015.39 | -0.17 | 5.36E-01 |
| NULL | GSMG0057010 | NULL | NULL | 28.92 | 59.55 | -1.05 | 2.11E-02 |
| NULL | GSMG0057015 | NULL | NULL | 149.18 | 290.30 | -0.96 | 1.63E-04 |
| NULL | GSMG0057363 | NULL | NULL | 820.98 | 545.23 | 0.59 | 2.49E-02 |
| ENSMUSG00000072692 | GSMG0057602 | Rpl37rt | ribosomal protein L37, retrotransposed | 1323.04 | 1789.36 | -0.44 | 1.65E-01 |
| NULL | GSMG0058042 | NULL | NULL | 53.06 | 69.32 | -0.38 | 4.31E-01 |
| NULL | GSMG0058591 | NULL | NULL | 528.44 | 534.39 | -0.02 | 9.65E-01 |
| NULL | GSMG0058592 | NULL | NULL | 501.52 | 563.17 | -0.17 | 5.02E-01 |
| NULL | GSMG0058593 | NULL | NULL | 475.30 | 455.78 | 0.06 | 8.61E-01 |
| NULL | GSMG0058594 | NULL | NULL | 481.25 | 378.92 | 0.34 | 2.08E-01 |
| NULL | GSMG0058653 | NULL | NULL | 270.34 | 586.45 | -1.12 | 8.62E-02 |
| NULL | GSMG0058663 | NULL | NULL | 112.75 | 102.56 | 0.14 | 7.67E-01 |
| NULL | GSMG0058929 | NULL | NULL | 147.76 | 248.30 | -0.76 | 1.00E-02 |
| NULL | GSMG0059278 | NULL | NULL | 9.31 | 57.60 | -2.62 | 1.85E-07 |
| NULL | GSMG0059302 | NULL | NULL | 28.01 | 152.15 | -2.43 | 7.86E-15 |
| NULL | GSMG0059306 | NULL | NULL | 587.54 | 618.27 | -0.07 | 8.23E-01 |
| NULL | GSMG0059917 | NULL | NULL | 91.96 | 139.22 | -0.60 | 5.13E-02 |
| ENSMUSG00000048424 | GSMG0060093 | Ranbp3l | RAN binding protein 3-like | 100.16 | 112.82 | -0.17 | 6.99E-01 |
| G00000024387,ENSMUSG000 | GSMG0060094 | Ly6g5b // Csnk2b | lymphocyte antigen 6 complex, locus G5B // casein kinase 2, beta polyp | 1289.19 | 1593.77 | -0.31 | 8.33E-02 |
| ENSMUSG00000029447 | GSMG0060095 | Cct6a | chaperonin containing Tcp1, subunit 6a (zeta) | 4630.40 | 6692.10 | -0.53 | 7.36E-04 |
| ENSMUSG00000038023 | GSMG0060096 | Atp6v0a2 | ATPase, H+ transporting, lysosomal V0 subunit A2 | 2758.21 | 2711.14 | 0.03 | 9.38E-01 |
| ENSMUSG00000028399 | GSMG0060097 | Ptprd | protein tyrosine phosphatase, receptor type, D | 4624.66 | 5010.37 | -0.12 | 6.73E-01 |
| ENSMUSG00000056856 | GSMG0060100 | Jakmip3 | janus kinase and microtubule interacting protein 3 | 121.34 | 87.13 | 0.48 | 2.46E-01 |
| NULL | GSMG0060187 | Ggnbp2os | gametogenetin binding protein 2, opposite strand | 343.19 | 327.28 | 0.07 | 8.63E-01 |
| NULL | GSMG0060261 | NULL | NULL | 3271.90 | 2612.00 | 0.33 | 4.20E-02 |
| ENSMUSG00000105010 | GSMG0060272 | Mir219b | microRNA 219b | 401.55 | 40.38 | 3.31 | 1.98E-27 |
| NULL | GSMG0060284 | NULL | NULL | 372.43 | 452.99 | -0.29 | 2.82E-01 |
| ENSMUSG00000098973 | GSMG0060383 | Mir6236 | microRNA 6236 | 4484.44 | 4556.75 | -0.02 | 9.81E-01 |
| 29,ENSMUSG00000089945,EI | GSMG0060428 | Pakap // Palm2 // A | paralemmin A kinase anchor protein // paralemmin 2 // A kinase (PRKA | 9896.65 | 8042.64 | 0.30 | 1.58E-01 |
| ENSMUSG00000046793 | GSMG0060429 | Gpr61 | G protein-coupled receptor 61 | 422.86 | 274.45 | 0.62 | 4.79E-03 |
| ENSMUSG00000039463 | GSMG0060441 | Slc9a8 | solute carrier family 9 (sodium/hydrogen exchanger), member 8 | 346.20 | 455.12 | -0.40 | 1.21E-01 |
| ENSMUSG00000057110 | GSMG0060443 | Cntrl | centriolin | 224.79 | 232.77 | -0.05 | 9.01E-01 |

| | | | | | | | |
|---|---|---|---|---|---|---|---|
| ENSMUSG00000005442 | GSMG0060461 | Cic | capicua transcriptional repressor | 424.24 | 601.22 | -0.51 | 7.53E-02 |
| ENSMUSG00000031028 | GSMG0060463 | Tub | tubby candidate gene | 575.24 | 639.53 | -0.15 | 7.18E-01 |
| ENSMUSG00000036099 | GSMG0060473 | Vezt | vezatin, adherens junctions transmembrane protein | 937.02 | 1044.66 | -0.16 | 4.66E-01 |
| ENSMUSG00000026708 | GSMG0060476 | Cenpl | centromere protein L | 127.53 | 182.59 | -0.51 | 9.46E-02 |
| ENSMUSG00000005886 | GSMG0060478 | Ncoa2 | nuclear receptor coactivator 2 | 175.36 | 207.13 | -0.23 | 5.93E-01 |
| ENSMUSG00000001755 | GSMG0060485 | Coasy | Coenzyme A synthase | 507.74 | 582.95 | -0.20 | 4.06E-01 |
| ENSMUSG00000045983 | GSMG0060492 | Eif4g1 | eukaryotic translation initiation factor 4, gamma 1 | 1725.73 | 2104.99 | -0.29 | 3.44E-01 |
| ENSMUSG00000052917 | GSMG0060494 | Senp7 | SUMO1/sentrin specific peptidase 7 | 210.47 | 228.49 | -0.12 | 7.39E-01 |
| ENSMUSG00000022768 | GSMG0060496 | Ccdc116 | coiled-coil domain containing 116 | 28.91 | 65.01 | -1.17 | 1.03E-02 |
| ENSMUSG00000058351 | GSMG0060499 | Smim4 | small integral membrane protein 4 | 179.98 | 204.49 | -0.19 | 6.11E-01 |
| ENSMUSG00000102224 | GSMG0060599 | 4930447F24Rik | RIKEN cDNA 4930447F24 gene | 211.67 | 106.65 | 0.99 | 2.36E-03 |
| G00000024854,ENSMUSG000 | GSMG0060613 | Clcf1-pold4 // Clcf | Clcf1-pold4 readthrough // cardiotrophin-like cytokine factor 1 // polym | 415.62 | 383.57 | 0.11 | 6.92E-01 |
| G00000015522,ENSMUSG000 | GSMG0060615 | Gm4349 // Arnt | predicted gene 4349 // aryl hydrocarbon receptor nuclear translocator | 1367.50 | 1321.14 | 0.05 | 8.54E-01 |
| ENSMUSG00000027584 | GSMG0060619 | Oprl1 | opioid receptor-like 1 | 73.27 | 127.44 | -0.79 | 1.24E-02 |
| ENSMUSG00000013465 | GSMG0060621 | Nelfb | negative elongation factor complex member B | 3715.70 | 3683.08 | 0.01 | 9.62E-01 |
| 45,ENSMUSG00000071350,EI | GSMG0060628 | Phf11d // Setdb2-p | PHD finger protein 11D // Setdb2-Phf11c readthrough // PHD finger pr | 283.23 | 344.64 | -0.29 | 2.80E-01 |
| ENSMUSG00000057367 | GSMG0060643 | Birc2 | baculoviral IAP repeat-containing 2 | 357.26 | 298.37 | 0.26 | 3.27E-01 |
| ENSMUSG00000032604 | GSMG0060645 | Qars | glutaminyl-tRNA synthetase | 1358.76 | 1464.88 | -0.11 | 6.98E-01 |
| ENSMUSG00000070371 | GSMG0060649 | Prss36 | protease, serine 36 | 124.35 | 102.16 | 0.28 | 4.65E-01 |
| ENSMUSG00000039057 | GSMG0060652 | Myo16 | myosin XVI | 341.33 | 862.50 | -1.34 | 6.91E-11 |
| ENSMUSG00000031626 | GSMG0060653 | Sorbs2 | sorbin and SH3 domain containing 2 | 3107.69 | 1890.97 | 0.72 | 2.17E-07 |
| G00000002345,ENSMUSG000 | GSMG0060654 | 2310045n01rik-me | 2310045N01Rik-mef2b readthrough // BLOC-1 related complex subun | 328.85 | 279.01 | 0.24 | 3.77E-01 |
| 19,ENSMUSG00000043687,EI | GSMG0060655 | Emc8-1190005i06 | Emc8-1190005I06Rik readthrough // RIKEN cDNA 1190005I06 gene / | 2419.79 | 2672.95 | -0.14 | 4.82E-01 |
| ENSMUSG00000074415 | GSMG0060658 | 3110039I08Rik | RIKEN cDNA 3110039I08 gene | 1689.91 | 2565.58 | -0.60 | 9.63E-05 |
| ENSMUSG00000093073 | GSMG0064891 | Mir124a-2 | microRNA 124a-2 | 612.57 | 790.02 | -0.37 | 1.13E-01 |
| ENSMUSG00000034275 | GSMG0064899 | Igsf9b | immunoglobulin superfamily, member 9B | 523.33 | 266.00 | 0.98 | 8.11E-06 |
| ENSMUSG00000101942 | GSMG0064904 | Gm19582 | predicted gene, 19582 | 148.66 | 116.52 | 0.35 | 3.07E-01 |
| ENSMUSG00000072761 | GSMG0064950 | Gm6712 | predicted gene 6712 | 48.61 | 50.44 | -0.05 | 9.52E-01 |
| ENSMUSG00000046994 | GSMG0064999 | Mars2 | methionine-tRNA synthetase 2 (mitochondrial) | 347.47 | 415.98 | -0.26 | 3.27E-01 |
| ENSMUSG00000020023 | GSMG0065007 | Tmcc3 | transmembrane and coiled coil domains 3 | 537.20 | 265.84 | 1.02 | 3.82E-08 |
| ENSMUSG00000096054 | GSMG0065008 | Syne1 | spectrin repeat containing, nuclear envelope 1 | 517.16 | 221.72 | 1.22 | 2.32E-09 |
| G00000069515,ENSMUSG000 | GSMG0065013 | Lyz2 // Lyz1 | lysozyme 2 // lysozyme 1 | 179.60 | 121.39 | 0.57 | 5.57E-02 |
| ENSMUSG00000001240 | GSMG0065019 | Ramp2 | receptor (calcitonin) activity modifying protein 2 | 14598.07 | 13687.23 | 0.09 | 6.95E-01 |

| | | | | | | | |
|---|---|---|---|---|---|---|---|
| ENSMUSG00000078656 | GSMG0065020 | Vps25 | vacuolar protein sorting 25 | 2526.88 | 2629.25 | -0.06 | 8.32E-01 |
| ENSMUSG00000046756 | GSMG0065022 | Mrps7 | mitchondrial ribosomal protein S7 | 1288.19 | 1464.98 | -0.19 | 3.64E-01 |
| NULL | GSMG0065023 | NULL | NULL | 186.03 | 221.97 | -0.26 | 5.09E-01 |
| ENSMUSG00000020950 | GSMG0065025 | Foxg1 | forkhead box G1 | 504.71 | 812.82 | -0.69 | 9.56E-03 |
| ENSMUSG00000021000 | GSMG0065027 | Mia2 // Ctage5 | melanoma inhibitory activity 2 // CTAGE family, member 5 | 1966.41 | 2095.46 | -0.09 | 7.83E-01 |
| ENSMUSG00000021268 | GSMG0065028 | Meg3 | maternally expressed 3 | 55908.90 | 37714.32 | 0.57 | 2.06E-03 |
| NULL | GSMG0065041 | NULL | NULL | 813.69 | 30.59 | 4.72 | 2.29E-63 |
| ENSMUSG00000051359 | GSMG0065043 | Ncald | neurocalcin delta | 450.05 | 548.12 | -0.29 | 2.17E-01 |
| ENSMUSG00000072214 | GSMG0065046 | Sept5 | septin 5 | 1263.57 | 1083.90 | 0.22 | 4.42E-01 |
| ENSMUSG00000050761 | GSMG0065047 | Gp1bb | glycoprotein Ib, beta polypeptide | 355.07 | 268.83 | 0.40 | 1.78E-01 |
| NULL | GSMG0065049 | NULL | NULL | 158.03 | 146.59 | 0.11 | 8.01E-01 |
| ENSMUSG00000023852 | GSMG0065052 | Chd1 | chromodomain helicase DNA binding protein 1 | 510.29 | 617.70 | -0.28 | 2.71E-01 |
| NULL | GSMG0065053 | Riok2 | RIO kinase 2 (yeast) | 475.51 | 524.51 | -0.14 | 5.88E-01 |
| ENSMUSG00000019432 | GSMG0065055 | Ddx39b | DEAD (Asp-Glu-Ala-Asp) box polypeptide 39B | 4076.19 | 5089.47 | -0.32 | 1.08E-01 |
| ENSMUSG00000110185 | GSMG0065063 | Igip | IgA inducing protein | 971.98 | 646.42 | 0.59 | 6.01E-03 |
| NULL | GSMG0065065 | NULL | NULL | 482.99 | 437.28 | 0.14 | 6.21E-01 |
| ENSMUSG00000069833 | GSMG0065066 | Ahnak | AHNAK nucleoprotein (desmoyokin) | 200.62 | 137.04 | 0.55 | 4.48E-02 |
| ENSMUSG00000053279 | GSMG0065067 | E030003E18Rik // | RIKEN cDNA E030003E18 gene // aldehyde dehydrogenase family 1, s | 861.58 | 178.53 | 2.27 | 1.73E-28 |
| ENSMUSG00000075289 | GSMG0065068 | Carns1 | carnosine synthase 1 | 402.32 | 137.55 | 1.55 | 5.57E-11 |
| NULL | GSMG0065070 | NULL | NULL | 7822.92 | 5264.37 | 0.57 | 3.91E-03 |
| NULL | GSMG0065072 | NULL | NULL | 1887.14 | 571.52 | 1.72 | 4.05E-32 |
| NULL | GSMG0065075 | NULL | NULL | 286.68 | 271.72 | 0.08 | 8.47E-01 |
| ENSMUSG00000039943 | GSMG0065076 | Plcb4 | phospholipase C, beta 4 | 1703.04 | 1081.84 | 0.65 | 5.54E-05 |
| ENSMUSG00000027210 | GSMG0065080 | Meis2 | Meis homeobox 2 | 231.40 | 388.44 | -0.75 | 4.16E-03 |
| ENSMUSG00000027408 | GSMG0065081 | Cpxm1 | carboxypeptidase X 1 (M14 family) | 502.19 | 824.77 | -0.71 | 1.66E-03 |
| ENSMUSG00000037773 | GSMG0065082 | Pced1a | PC-esterase domain containing 1A | 229.49 | 265.05 | -0.21 | 5.02E-01 |
| ENSMUSG00000027827 | GSMG0065087 | Kcnab1 | potassium voltage-gated channel, shaker-related subfamily, beta membe | 170.64 | 102.61 | 0.73 | 7.15E-03 |
| ENSMUSG00000074513 | GSMG0065093 | Arfip1 | ADP-ribosylation factor interacting protein 1 | 650.69 | 635.51 | 0.03 | 9.39E-01 |
| NULL | GSMG0065100 | NULL | NULL | 662.89 | 775.29 | -0.23 | 3.48E-01 |
| NULL | GSMG0065105 | NULL | NULL | 987.39 | 907.85 | 0.12 | 6.31E-01 |
| NULL | GSMG0065112 | NULL | NULL | 459.10 | 612.95 | -0.42 | 5.96E-02 |
| ENSMUSG00000029713 | GSMG0065117 | Gnb2 | guanine nucleotide binding protein (G protein), beta 2 | 1967.16 | 2227.21 | -0.18 | 5.33E-01 |
| ENSMUSG00000029798 | GSMG0065122 | Herc6 | hect domain and RLD 6 | 123.56 | 114.79 | 0.10 | 8.30E-01 |

| | | | | | | | |
|---|---|---|---|---|---|---|---|
| NULL | GSMG0065128 | NULL | NULL | 1208.93 | 1109.12 | 0.12 | 5.82E-01 |
| ENSMUSG00000030967 | GSMG0065130 | Zranb1 | zinc finger, RAN-binding domain containing 1 | 588.65 | 529.66 | 0.15 | 6.36E-01 |
| G00000074129,ENSMUSG000 | GSMG0065139 | Rpl13a // Flt3l | ribosomal protein L13A // FMS-like tyrosine kinase 3 ligand | 1618.26 | 2270.62 | -0.49 | 3.68E-02 |
| G00000102252,ENSMUSG000 | GSMG0065140 | Ipw // Snurf // Snrp | imprinted gene in the Prader-Willi syndrome region // SNRPN upstrean | 4466.41 | 4314.15 | 0.05 | 8.60E-01 |
| NULL | GSMG0065144 | NULL | NULL | 7703.92 | 4123.78 | 0.90 | 3.34E-11 |
| ENSMUSG00000070348 | GSMG0065147 | Ccnd1 | cyclin D1 | 2502.91 | 3833.37 | -0.61 | 4.68E-03 |
| ENSMUSG00000049184 | GSMG0065150 | Purg | purine-rich element binding protein G | 250.87 | 283.30 | -0.18 | 5.96E-01 |
| ENSMUSG00000039530 | GSMG0065152 | Tusc3 | tumor suppressor candidate 3 | 1899.18 | 3004.32 | -0.66 | 6.57E-05 |
| NULL | GSMG0065158 | NULL | NULL | 1445.73 | 1423.35 | 0.02 | 9.53E-01 |
| ENSMUSG00000098120 | GSMG0065163 | Gm5914 | predicted gene 5914 | 83.08 | 171.76 | -1.06 | 3.09E-04 |
| ENSMUSG00000032323 | GSMG0065174 | Cyp11a1 | cytochrome P450, family 11, subfamily a, polypeptide 1 | 196.99 | 33.90 | 2.55 | 1.31E-16 |
| NULL | GSMG0065176 | NULL | NULL | 3808.74 | 3634.32 | 0.07 | 8.31E-01 |
| ENSMUSG00000050397 | GSMG0065181 | Foxl2 | forkhead box L2 | 1544.15 | 1665.29 | -0.11 | 7.60E-01 |
| ENSMUSG00000025794 | GSMG0065182 | Rpl14 | ribosomal protein L14 | 1033.71 | 1538.71 | -0.57 | 1.22E-02 |
| ENSMUSG00000046721 | GSMG0065183 | Rpl14-ps1 | ribosomal protein L14, pseudogene 1 | 2278.18 | 3781.26 | -0.73 | 3.15E-06 |
| NULL | GSMG0065187 | NULL | NULL | 2457.76 | 1872.14 | 0.39 | 6.72E-02 |
| ENSMUSG00000072964 | GSMG0065201 | Bhlhb9 | basic helix-loop-helix domain containing, class B9 | 2156.28 | 2223.41 | -0.04 | 8.72E-01 |
| ENSMUSG00000072966 | GSMG0065202 | Gprasp2 | G protein-coupled receptor associated sorting protein 2 | 2871.70 | 2424.68 | 0.24 | 2.64E-01 |
| ENSMUSG00000044550 | GSMG0065203 | Tceal3 | transcription elongation factor A (SII)-like 3 | 271.46 | 324.16 | -0.26 | 4.01E-01 |
| ENSMUSG00000079418 | GSMG0065204 | Atg4a | autophagy related 4A, cysteine peptidase | 713.48 | 684.85 | 0.06 | 8.51E-01 |
| G00000065462,ENSMUSG000 | GSMG0065213 | Mir141 // Mir200c | microRNA 141 // microRNA 200c | 1245.87 | 1394.84 | -0.16 | 4.19E-01 |
| 55,ENSMUSG00000089768,EI | GSMG0065221 | Tmsb15l // Tmsb15 | thymosin beta 15b like // thymosin beta 15b2 // thymosin beta 15b1 | 203.88 | 227.47 | -0.16 | 6.98E-01 |
| NULL | GSMG0065239 | NULL | NULL | 1731.11 | 1576.72 | 0.14 | 5.44E-01 |
| NULL | GSMG0065258 | NULL | NULL | 546.08 | 510.95 | 0.10 | 7.62E-01 |
| NULL | GSMG0065267 | NULL | NULL | 300.96 | 194.39 | 0.63 | 1.72E-02 |
| NULL | GSMG0065295 | NULL | NULL | 1024.20 | 1145.47 | -0.16 | 5.10E-01 |
| NULL | GSMG0065315 | NULL | NULL | 59.91 | 53.03 | 0.19 | 7.77E-01 |
| NULL | GSMG0065335 | NULL | NULL | 506.27 | 417.81 | 0.28 | 2.06E-01 |
| NULL | GSMG0065336 | NULL | NULL | 441.61 | 287.75 | 0.62 | 7.78E-03 |
| NULL | GSMG0065342 | NULL | NULL | 149.47 | 101.11 | 0.57 | 5.76E-02 |
| NULL | GSMG0065351 | NULL | NULL | 2068.07 | 1704.77 | 0.28 | 3.07E-01 |
| NULL | GSMG0065364 | NULL | NULL | 775.14 | 826.22 | -0.09 | 7.31E-01 |
| NULL | GSMG0065365 | NULL | NULL | 1198.65 | 1223.73 | -0.03 | 9.43E-01 |

| | | | | | | | |
|---|---|---|---|---|---|---|---|
| NULL | GSMG0065367 | NULL | NULL | 888.44 | 891.89 | -0.01 | 9.86E-01 |
| NULL | GSMG0065384 | NULL | NULL | 525.92 | 740.65 | -0.50 | 6.07E-02 |
| NULL | GSMG0065399 | NULL | NULL | 571.28 | 654.00 | -0.19 | 4.06E-01 |
| NULL | GSMG0065441 | NULL | NULL | 2260.16 | 2083.15 | 0.12 | 6.54E-01 |
| NULL | GSMG0065443 | NULL | NULL | 646.72 | 673.77 | -0.06 | 8.43E-01 |
| NULL | GSMG0065446 | NULL | NULL | 119.23 | 170.74 | -0.52 | 8.74E-02 |
| NULL | GSMG0065449 | NULL | NULL | 575.81 | 503.11 | 0.19 | 5.02E-01 |
| ENSMUSG00000085327 | GSMG0065450 | Gm16104 | predicted gene 16104 | 29.46 | 50.23 | -0.79 | 1.30E-01 |
| NULL | GSMG0065454 | NULL | NULL | 945.05 | 1188.24 | -0.33 | 5.92E-02 |
| NULL | GSMG0065457 | NULL | NULL | 294.38 | 441.61 | -0.59 | 5.65E-02 |
| NULL | GSMG0065489 | NULL | NULL | 471.82 | 419.58 | 0.17 | 5.74E-01 |
| NULL | GSMG0065501 | NULL | NULL | 1044.30 | 1084.79 | -0.06 | 8.57E-01 |
| NULL | GSMG0065520 | NULL | NULL | 53247.69 | 35040.31 | 0.60 | 6.47E-05 |
| NULL | GSMG0065525 | NULL | NULL | 1137.77 | 928.81 | 0.29 | 1.10E-01 |
| NULL | GSMG0065527 | NULL | NULL | 73.13 | 98.38 | -0.44 | 2.85E-01 |
| NULL | GSMG0065528 | NULL | NULL | 260.33 | 122.75 | 1.08 | 8.11E-05 |
| NULL | GSMG0065543 | NULL | NULL | 2381.19 | 1776.65 | 0.42 | 2.79E-02 |
| NULL | GSMG0065555 | NULL | NULL | 477.65 | 457.79 | 0.06 | 8.70E-01 |
| NULL | GSMG0065559 | NULL | NULL | 337.41 | 101.26 | 1.74 | 1.31E-11 |
| NULL | GSMG0065580 | NULL | NULL | 2765.15 | 3029.80 | -0.13 | 5.03E-01 |
| NULL | GSMG0065593 | NULL | NULL | 152.50 | 139.64 | 0.12 | 7.82E-01 |
| NULL | GSMG0065595 | NULL | NULL | 1756.25 | 2936.08 | -0.74 | 1.30E-05 |
| NULL | GSMG0065605 | NULL | NULL | 173.26 | 408.04 | -1.23 | 3.65E-07 |
| NULL | GSMG0065606 | NULL | NULL | 2953.37 | 2237.51 | 0.40 | 4.50E-02 |
| NULL | GSMG0065611 | NULL | NULL | 266.76 | 221.90 | 0.27 | 4.19E-01 |
| NULL | GSMG0065613 | NULL | NULL | 1489.21 | 1330.54 | 0.16 | 4.48E-01 |
| NULL | GSMG0065614 | NULL | NULL | 88.23 | 57.74 | 0.61 | 1.34E-01 |
| NULL | GSMG0065624 | NULL | NULL | 783.37 | 495.09 | 0.66 | 2.03E-04 |
| NULL | GSMG0065632 | NULL | NULL | 2017.93 | 1652.19 | 0.29 | 1.87E-01 |
| NULL | GSMG0065635 | NULL | NULL | 528.36 | 667.93 | -0.34 | 2.69E-01 |
| NULL | GSMG0065638 | NULL | NULL | 1385.01 | 1428.12 | -0.04 | 8.89E-01 |
| NULL | GSMG0065646 | NULL | NULL | 1756.93 | 1034.17 | 0.77 | 1.08E-05 |
| NULL | GSMG0065648 | NULL | NULL | 83.75 | 88.50 | -0.08 | 8.93E-01 |

| | | | | | | | |
|---|---|---|---|---|---|---|---|
| NULL | GSMG0065659 | NULL | NULL | 1001.69 | 928.24 | 0.11 | 6.42E-01 |
| NULL | GSMG0065660 | NULL | NULL | 776.10 | 656.43 | 0.24 | 2.28E-01 |
| NULL | GSMG0065661 | NULL | NULL | 1378.97 | 1103.29 | 0.32 | 7.66E-02 |
| NULL | GSMG0065667 | NULL | NULL | 254.20 | 231.98 | 0.13 | 7.47E-01 |
| NULL | GSMG0065690 | NULL | NULL | 645.21 | 862.26 | -0.42 | 5.43E-02 |
| NULL | GSMG0065694 | NULL | NULL | 1077.59 | 1120.58 | -0.06 | 8.50E-01 |
| NULL | GSMG0065698 | NULL | NULL | 194.16 | 179.41 | 0.12 | 7.93E-01 |
| NULL | GSMG0065709 | NULL | NULL | 71.01 | 80.15 | -0.18 | 7.34E-01 |
| NULL | GSMG0065714 | NULL | NULL | 171.28 | 218.52 | -0.36 | 2.44E-01 |
| NULL | GSMG0065723 | NULL | NULL | 392.09 | 407.16 | -0.06 | 8.90E-01 |
| NULL | GSMG0065724 | NULL | NULL | 537.75 | 526.93 | 0.03 | 9.32E-01 |
| NULL | GSMG0065726 | NULL | NULL | 1090.14 | 984.56 | 0.15 | 5.13E-01 |
| NULL | GSMG0065727 | NULL | NULL | 374.71 | 400.04 | -0.10 | 7.65E-01 |
| NULL | GSMG0065730 | NULL | NULL | 1480.08 | 1784.23 | -0.27 | 1.25E-01 |
| NULL | GSMG0065752 | NULL | NULL | 710.70 | 230.49 | 1.63 | 2.32E-18 |
| NULL | GSMG0065755 | NULL | NULL | 2312.57 | 2737.19 | -0.24 | 4.20E-01 |
| NULL | GSMG0065769 | NULL | NULL | 167.71 | 338.57 | -1.02 | 6.52E-06 |
| NULL | GSMG0065793 | NULL | NULL | 1022.67 | 848.07 | 0.27 | 2.55E-01 |
| NULL | GSMG0065797 | NULL | NULL | 53.06 | 56.86 | -0.11 | 8.80E-01 |
| NULL | GSMG0065803 | NULL | NULL | 125.70 | 128.70 | -0.03 | 9.50E-01 |
| NULL | GSMG0065808 | NULL | NULL | 356.84 | 600.13 | -0.75 | 5.83E-05 |
| NULL | GSMG0065812 | NULL | NULL | 909.78 | 1120.14 | -0.30 | 1.38E-01 |
| NULL | GSMG0065831 | NULL | NULL | 843.09 | 1089.81 | -0.37 | 5.79E-02 |
| NULL | GSMG0065835 | NULL | NULL | 1927.73 | 1854.19 | 0.06 | 8.61E-01 |
| NULL | GSMG0065836 | NULL | NULL | 142.01 | 115.96 | 0.28 | 5.14E-01 |
| NULL | GSMG0065916 | NULL | NULL | 262.53 | 297.59 | -0.19 | 6.21E-01 |
| NULL | GSMG0065923 | NULL | NULL | 701.15 | 759.88 | -0.12 | 7.85E-01 |
| NULL | GSMG0065935 | NULL | NULL | 1733.31 | 1842.26 | -0.09 | 7.64E-01 |
| NULL | GSMG0065936 | NULL | NULL | 367.67 | 310.10 | 0.25 | 3.60E-01 |
| NULL | GSMG0065962 | NULL | NULL | 75.93 | 73.46 | 0.05 | 9.44E-01 |
| NULL | GSMG0065963 | NULL | NULL | 199.11 | 170.65 | 0.22 | 4.80E-01 |
| NULL | GSMG0065977 | NULL | NULL | 3046.15 | 3527.98 | -0.21 | 3.63E-01 |
| NULL | GSMG0065978 | NULL | NULL | 1327.13 | 1292.76 | 0.04 | 8.97E-01 |

| | | | | | | | |
|---|---|---|---|---|---|---|---|
| NULL | GSMG0066023 | NULL | NULL | 9524.93 | 13515.68 | -0.50 | 5.05E-04 |
| NULL | GSMG0066047 | NULL | NULL | 374.06 | 509.76 | -0.45 | 4.36E-02 |
| NULL | GSMG0066073 | NULL | NULL | 839.85 | 900.71 | -0.10 | 7.33E-01 |
| NULL | GSMG0066078 | NULL | NULL | 256.24 | 234.63 | 0.13 | 7.39E-01 |
| NULL | GSMG0066108 | NULL | NULL | 273.53 | 167.37 | 0.71 | 2.86E-03 |
| NULL | GSMG0066141 | NULL | NULL | 433.64 | 467.03 | -0.11 | 7.34E-01 |
| NULL | GSMG0066142 | NULL | NULL | 68.94 | 22.92 | 1.59 | 1.08E-04 |
| NULL | GSMG0066144 | NULL | NULL | 401.79 | 318.18 | 0.34 | 1.71E-01 |
| NULL | GSMG0066154 | NULL | NULL | 105.71 | 52.78 | 1.00 | 7.44E-03 |
| NULL | GSMG0066162 | NULL | NULL | 459.15 | 337.41 | 0.44 | 3.77E-02 |
| NULL | GSMG0066180 | NULL | NULL | 87.17 | 90.53 | -0.05 | 9.33E-01 |
| NULL | GSMG0066199 | NULL | NULL | 2271.50 | 1609.44 | 0.50 | 5.41E-03 |
| NULL | GSMG0066234 | NULL | NULL | 213.98 | 311.47 | -0.54 | 2.43E-02 |
| NULL | GSMG0066242 | NULL | NULL | 234.87 | 240.76 | -0.04 | 9.52E-01 |
| NULL | GSMG0066243 | NULL | NULL | 77.43 | 72.40 | 0.10 | 8.60E-01 |
| NULL | GSMG0066292 | NULL | NULL | 845.78 | 1236.36 | -0.55 | 7.19E-04 |
| NULL | GSMG0066299 | NULL | NULL | 559.43 | 663.47 | -0.25 | 3.28E-01 |
| NULL | GSMG0066317 | NULL | NULL | 603.14 | 859.80 | -0.51 | 3.12E-03 |
| NULL | GSMG0066348 | NULL | NULL | 286.76 | 183.97 | 0.64 | 7.41E-03 |
| NULL | GSMG0066354 | NULL | NULL | 157.68 | 195.16 | -0.31 | 4.55E-01 |
| NULL | GSMG0066367 | NULL | NULL | 2509.45 | 3540.30 | -0.50 | 4.43E-02 |
| NULL | GSMG0066368 | NULL | NULL | 148.75 | 161.03 | -0.11 | 7.96E-01 |
| NULL | GSMG0066389 | NULL | NULL | 1038.22 | 951.99 | 0.13 | 7.00E-01 |
| NULL | GSMG0066391 | NULL | NULL | 1812.67 | 1763.18 | 0.04 | 8.96E-01 |
| NULL | GSMG0066392 | NULL | NULL | 92.39 | 112.06 | -0.28 | 5.24E-01 |
| NULL | GSMG0066395 | NULL | NULL | 5519.18 | 15964.72 | -1.53 | 7.66E-29 |
| NULL | GSMG0066432 | NULL | NULL | 159.53 | 309.45 | -0.95 | 4.88E-04 |
| NULL | GSMG0066440 | NULL | NULL | 334.67 | 642.71 | -0.94 | 1.79E-07 |
| NULL | GSMG0066452 | NULL | NULL | 121.36 | 106.97 | 0.18 | 6.63E-01 |
| NULL | GSMG0066462 | NULL | NULL | 6990.95 | 2445.87 | 1.52 | 2.56E-23 |
| NULL | GSMG0066463 | NULL | NULL | 151.36 | 180.64 | -0.25 | 4.69E-01 |
| NULL | GSMG0066467 | NULL | NULL | 2879.52 | 2732.63 | 0.08 | 7.42E-01 |
| NULL | GSMG0066468 | NULL | NULL | 2014.53 | 1960.85 | 0.04 | 8.86E-01 |

| | | | | | | | |
|---|---|---|---|---|---|---|---|
| NULL | GSMG0066469 | NULL | NULL | 2531.97 | 2526.82 | 0.00 | 9.92E-01 |
| NULL | GSMG0066470 | NULL | NULL | 239.34 | 255.57 | -0.10 | 7.96E-01 |
| NULL | GSMG0066477 | NULL | NULL | 2564.63 | 4622.43 | -0.85 | 3.27E-07 |
| NULL | GSMG0066502 | NULL | NULL | 2759.24 | 2442.69 | 0.18 | 3.83E-01 |
| NULL | GSMG0066508 | NULL | NULL | 2034.14 | 2321.07 | -0.19 | 3.27E-01 |
| NULL | GSMG0066533 | NULL | NULL | 1529.07 | 1068.95 | 0.52 | 4.87E-02 |
| NULL | GSMG0066537 | NULL | NULL | 52.51 | 71.50 | -0.44 | 3.32E-01 |
| NULL | GSMG0066541 | NULL | NULL | 161.70 | 222.58 | -0.46 | 1.31E-01 |
| NULL | GSMG0066562 | NULL | NULL | 348.49 | 145.73 | 1.26 | 2.64E-08 |
| NULL | GSMG0066624 | NULL | NULL | 347.94 | 192.35 | 0.85 | 2.65E-04 |
| NULL | GSMG0066629 | NULL | NULL | 55.48 | 64.03 | -0.20 | 8.04E-01 |
| ENSMUSG00000073448 | GSMG0066672 | Gm10509 | predicted gene 10509 | 224.05 | 286.61 | -0.35 | 1.89E-01 |
| NULL | GSMG0066680 | NULL | NULL | 3519.01 | 797.33 | 2.14 | 7.61E-51 |
| NULL | GSMG0066688 | NULL | NULL | 1559.44 | 723.05 | 1.11 | 8.73E-09 |
| NULL | GSMG0066694 | NULL | NULL | 422.49 | 267.99 | 0.65 | 1.10E-02 |
| NULL | GSMG0066695 | NULL | NULL | 2971.08 | 7044.66 | -1.25 | 3.71E-16 |
| NULL | GSMG0066719 | NULL | NULL | 4651.46 | 5329.03 | -0.20 | 3.84E-01 |
| NULL | GSMG0066752 | NULL | NULL | 462.43 | 381.80 | 0.28 | 2.86E-01 |
| NULL | GSMG0066753 | NULL | NULL | 452.54 | 559.83 | -0.31 | 1.51E-01 |
| NULL | GSMG0066767 | NULL | NULL | 6616.17 | 5198.41 | 0.35 | 2.44E-02 |
| NULL | GSMG0066773 | NULL | NULL | 1163.41 | 1387.13 | -0.25 | 3.25E-01 |
| NULL | GSMG0066777 | NULL | NULL | 48.90 | 84.24 | -0.78 | 3.80E-02 |
| NULL | GSMG0066789 | NULL | NULL | 441.26 | 778.43 | -0.82 | 3.48E-07 |
| NULL | GSMG0066796 | NULL | NULL | 305.62 | 400.88 | -0.39 | 1.09E-01 |
| NULL | GSMG0066807 | NULL | NULL | 127.59 | 138.90 | -0.12 | 7.89E-01 |
| NULL | GSMG0066847 | NULL | NULL | 483.26 | 351.30 | 0.46 | 1.01E-01 |
| NULL | GSMG0066856 | NULL | NULL | 52.26 | 67.93 | -0.38 | 4.29E-01 |
| NULL | GSMG0066864 | NULL | NULL | 646.62 | 601.91 | 0.10 | 7.27E-01 |
| NULL | GSMG0066878 | NULL | NULL | 368.96 | 402.75 | -0.13 | 7.14E-01 |
| NULL | GSMG0066890 | NULL | NULL | 733.19 | 704.17 | 0.06 | 8.42E-01 |
| NULL | GSMG0066894 | NULL | NULL | 245.45 | 158.86 | 0.63 | 4.99E-02 |
| NULL | GSMG0066897 | NULL | NULL | 1030.99 | 701.85 | 0.55 | 1.95E-02 |
| NULL | GSMG0066907 | NULL | NULL | 258.32 | 256.18 | 0.01 | 9.81E-01 |

| | | | | | | | |
|---|---|---|---|---|---|---|---|
| NULL | GSMG0066912 | NULL | NULL | 1699.62 | 1659.33 | 0.03 | 9.02E-01 |
| NULL | GSMG0066922 | NULL | NULL | 93.10 | 89.81 | 0.05 | 9.41E-01 |
| NULL | GSMG0066933 | NULL | NULL | 487.61 | 321.08 | 0.60 | 2.45E-03 |
| NULL | GSMG0066934 | NULL | NULL | 870.31 | 1006.99 | -0.21 | 3.73E-01 |
| NULL | GSMG0066939 | NULL | NULL | 5055.33 | 7346.85 | -0.54 | 6.45E-04 |
| NULL | GSMG0066974 | NULL | NULL | 129.17 | 140.00 | -0.12 | 8.19E-01 |
| NULL | GSMG0066978 | NULL | NULL | 562.53 | 407.16 | 0.47 | 1.79E-02 |
| NULL | GSMG0066983 | NULL | NULL | 423.37 | 455.03 | -0.11 | 7.54E-01 |
| NULL | GSMG0066991 | NULL | NULL | 452.89 | 548.60 | -0.28 | 2.40E-01 |
| NULL | GSMG0066992 | NULL | NULL | 709.62 | 747.34 | -0.08 | 7.84E-01 |
| NULL | GSMG0067031 | NULL | NULL | 1673.00 | 1368.77 | 0.29 | 9.40E-02 |
| NULL | GSMG0067033 | NULL | NULL | 730.88 | 800.68 | -0.13 | 6.84E-01 |
| NULL | GSMG0067051 | NULL | NULL | 119.89 | 190.79 | -0.67 | 1.81E-02 |
| NULL | GSMG0067059 | NULL | NULL | 1417.96 | 796.56 | 0.83 | 5.97E-04 |
| NULL | GSMG0067087 | NULL | NULL | 951.07 | 1149.34 | -0.27 | 2.00E-01 |
| NULL | GSMG0067125 | NULL | NULL | 1062.10 | 959.53 | 0.15 | 5.95E-01 |
| NULL | GSMG0067169 | NULL | NULL | 8539.50 | 5817.66 | 0.55 | 3.11E-05 |
| NULL | GSMG0067172 | NULL | NULL | 2236.13 | 2151.95 | 0.06 | 8.58E-01 |
| NULL | GSMG0067185 | NULL | NULL | 148.56 | 419.60 | -1.50 | 1.38E-08 |
| NULL | GSMG0067197 | NULL | NULL | 511.35 | 565.09 | -0.15 | 5.99E-01 |
| NULL | GSMG0067203 | NULL | NULL | 63.92 | 65.11 | -0.03 | 9.64E-01 |
| NULL | GSMG0067213 | NULL | NULL | 2472.94 | 2328.67 | 0.09 | 7.34E-01 |
| NULL | GSMG0067224 | NULL | NULL | 2325.03 | 1930.19 | 0.27 | 1.59E-01 |
| NULL | GSMG0067229 | NULL | NULL | 1047.04 | 1185.21 | -0.18 | 4.84E-01 |
| NULL | GSMG0067236 | NULL | NULL | 316.41 | 352.57 | -0.16 | 6.94E-01 |
| NULL | GSMG0067237 | NULL | NULL | 1525.31 | 1820.53 | -0.25 | 2.33E-01 |
| NULL | GSMG0067314 | NULL | NULL | 3048.52 | 3489.19 | -0.20 | 3.76E-01 |
| NULL | GSMG0067315 | NULL | NULL | 3491.63 | 3715.68 | -0.09 | 7.17E-01 |
| NULL | GSMG0067320 | NULL | NULL | 57.03 | 66.09 | -0.21 | 7.18E-01 |
| NULL | GSMG0067324 | NULL | NULL | 647.42 | 866.76 | -0.42 | 2.45E-02 |
| NULL | GSMG0067330 | NULL | NULL | 1243.80 | 572.14 | 1.12 | 1.82E-09 |
| NULL | GSMG0067348 | NULL | NULL | 409.10 | 847.07 | -1.05 | 2.53E-07 |
| NULL | GSMG0067351 | NULL | NULL | 87.30 | 71.09 | 0.30 | 5.05E-01 |

| | | | | | | | |
|---|---|---|---|---|---|---|---|
| NULL | GSMG0067355 | NULL | NULL | 342.62 | 433.22 | -0.34 | 2.36E-01 |
| NULL | GSMG0067360 | NULL | NULL | 111.46 | 151.56 | -0.45 | 1.66E-01 |
| NULL | GSMG0067378 | NULL | NULL | 3608.41 | 1919.21 | 0.91 | 2.10E-06 |
| NULL | GSMG0067387 | NULL | NULL | 1733.58 | 935.69 | 0.89 | 2.04E-08 |
| NULL | GSMG0067507 | NULL | NULL | 819.10 | 1127.31 | -0.46 | 9.59E-03 |
| NULL | GSMG0067534 | NULL | NULL | 1210.62 | 1489.53 | -0.30 | 1.32E-01 |
| NULL | GSMG0067549 | NULL | NULL | 42.13 | 67.04 | -0.68 | 2.37E-01 |
| NULL | GSMG0067554 | NULL | NULL | 184.68 | 320.87 | -0.80 | 3.26E-03 |
| NULL | GSMG0067595 | NULL | NULL | 1720.70 | 1807.82 | -0.07 | 8.33E-01 |
| NULL | GSMG0067599 | NULL | NULL | 579.70 | 491.05 | 0.24 | 4.11E-01 |
| NULL | GSMG0067602 | NULL | NULL | 80.87 | 99.83 | -0.32 | 4.99E-01 |
| NULL | GSMG0067608 | NULL | NULL | 224.87 | 163.71 | 0.45 | 1.62E-01 |
| NULL | GSMG0067621 | NULL | NULL | 5849.51 | 6398.41 | -0.13 | 5.90E-01 |
| NULL | GSMG0067623 | NULL | NULL | 269.87 | 655.63 | -1.28 | 4.84E-11 |
| NULL | GSMG0067649 | NULL | NULL | 1488.29 | 1803.58 | -0.28 | 1.52E-01 |
| NULL | GSMG0067650 | NULL | NULL | 1365.99 | 1020.91 | 0.42 | 1.81E-02 |
| NULL | GSMG0067671 | NULL | NULL | 4957.75 | 3701.58 | 0.42 | 5.58E-03 |
| NULL | GSMG0067688 | NULL | NULL | 35315.29 | 21648.16 | 0.71 | 3.61E-05 |
| NULL | GSMG0067691 | NULL | NULL | 574.56 | 583.21 | -0.02 | 9.45E-01 |
| NULL | GSMG0067696 | NULL | NULL | 309.84 | 456.43 | -0.56 | 9.33E-03 |
| NULL | GSMG0067730 | NULL | NULL | 3849.58 | 3313.96 | 0.22 | 3.95E-01 |
| NULL | GSMG0067755 | NULL | NULL | 306.59 | 330.28 | -0.11 | 7.41E-01 |
| NULL | GSMG0067757 | NULL | NULL | 361.60 | 683.65 | -0.92 | 8.41E-05 |
| NULL | GSMG0067764 | NULL | NULL | 77.42 | 103.94 | -0.42 | 3.25E-01 |
| NULL | GSMG0067771 | NULL | NULL | 87.05 | 90.70 | -0.06 | 9.21E-01 |
| NULL | GSMG0067776 | NULL | NULL | 233.61 | 283.67 | -0.28 | 3.51E-01 |
| NULL | GSMG0067784 | NULL | NULL | 145.62 | 178.48 | -0.30 | 4.20E-01 |
| NULL | GSMG0067796 | NULL | NULL | 737.86 | 847.07 | -0.20 | 4.48E-01 |
| NULL | GSMG0067809 | NULL | NULL | 765.16 | 815.78 | -0.09 | 7.86E-01 |
| NULL | GSMG0067818 | NULL | NULL | 4765.40 | 3186.44 | 0.58 | 1.82E-05 |
| NULL | GSMG0067819 | NULL | NULL | 60.01 | 141.48 | -1.24 | 2.88E-04 |
| NULL | GSMG0067843 | NULL | NULL | 219.13 | 34.57 | 2.65 | 1.53E-16 |
| NULL | GSMG0067844 | NULL | NULL | 1474.23 | 1928.91 | -0.39 | 5.95E-02 |

| | | | | | | | |
|---|---|---|---|---|---|---|---|
| NULL | GSMG0067849 | NULL | NULL | 831.20 | 715.52 | 0.21 | 4.46E-01 |
| NULL | GSMG0067851 | NULL | NULL | 1754.46 | 1801.63 | -0.04 | 8.88E-01 |
| NULL | GSMG0067862 | NULL | NULL | 195.96 | 141.93 | 0.47 | 8.42E-02 |
| NULL | GSMG0067865 | NULL | NULL | 6845.03 | 6932.30 | -0.02 | 9.55E-01 |
| NULL | GSMG0067866 | NULL | NULL | 365.89 | 616.13 | -0.75 | 1.15E-03 |
| NULL | GSMG0067872 | NULL | NULL | 342.81 | 450.21 | -0.40 | 1.10E-01 |
| NULL | GSMG0067874 | NULL | NULL | 1106.72 | 1260.37 | -0.19 | 3.88E-01 |
| NULL | GSMG0067896 | NULL | NULL | 1220.81 | 785.11 | 0.64 | 2.49E-02 |
| NULL | GSMG0067919 | NULL | NULL | 204.52 | 225.33 | -0.14 | 6.85E-01 |
| NULL | GSMG0067920 | NULL | NULL | 911.96 | 1035.02 | -0.18 | 4.64E-01 |
| NULL | GSMG0067947 | NULL | NULL | 305.40 | 273.79 | 0.16 | 6.09E-01 |
| NULL | GSMG0067974 | NULL | NULL | 7190.04 | 4335.00 | 0.73 | 1.30E-05 |
| NULL | GSMG0067989 | NULL | NULL | 6026.48 | 7180.15 | -0.25 | 3.33E-01 |
| NULL | GSMG0068012 | NULL | NULL | 337.30 | 555.55 | -0.72 | 6.59E-04 |
| NULL | GSMG0068024 | NULL | NULL | 288.77 | 354.43 | -0.29 | 3.27E-01 |
| NULL | GSMG0068025 | NULL | NULL | 597.68 | 963.58 | -0.69 | 4.30E-05 |
| ENSMUSG00000086606 | GSMG0068027 | Gm13205 | predicted gene 13205 | 181.61 | 109.96 | 0.72 | 1.23E-02 |
| NULL | GSMG0068072 | NULL | NULL | 547.10 | 339.69 | 0.69 | 1.23E-03 |
| NULL | GSMG0068090 | NULL | NULL | 551.58 | 339.08 | 0.70 | 2.98E-03 |
| NULL | GSMG0068116 | NULL | NULL | 146.44 | 89.45 | 0.72 | 1.63E-02 |
| NULL | GSMG0068133 | NULL | NULL | 112.15 | 62.81 | 0.83 | 8.87E-02 |
| NULL | GSMG0068160 | NULL | NULL | 926.12 | 798.64 | 0.21 | 3.18E-01 |
| NULL | GSMG0068193 | NULL | NULL | 137.01 | 92.21 | 0.56 | 1.27E-01 |
| NULL | GSMG0068199 | NULL | NULL | 641.04 | 942.49 | -0.56 | 1.20E-02 |
| NULL | GSMG0068202 | NULL | NULL | 1064.92 | 1070.10 | -0.01 | 9.85E-01 |
| NULL | GSMG0068208 | NULL | NULL | 301.05 | 133.30 | 1.18 | 3.99E-08 |
| NULL | GSMG0068215 | NULL | NULL | 1000.55 | 954.00 | 0.07 | 8.28E-01 |
| NULL | GSMG0068216 | NULL | NULL | 2799.04 | 1317.94 | 1.09 | 4.55E-14 |
| NULL | GSMG0068221 | NULL | NULL | 421.53 | 429.40 | -0.03 | 9.38E-01 |
| NULL | GSMG0068225 | NULL | NULL | 201.10 | 280.59 | -0.48 | 5.29E-02 |
| NULL | GSMG0068289 | NULL | NULL | 568.90 | 536.55 | 0.08 | 8.08E-01 |
| NULL | GSMG0068290 | NULL | NULL | 897.06 | 1005.04 | -0.16 | 4.56E-01 |
| NULL | GSMG0068301 | NULL | NULL | 1399.45 | 1451.27 | -0.05 | 8.66E-01 |

| | | | | | | | |
|---|---|---|---|---|---|---|---|
| NULL | GSMG0068308 | NULL | NULL | 156.69 | 170.96 | -0.13 | 7.78E-01 |
| NULL | GSMG0068310 | NULL | NULL | 362.12 | 268.48 | 0.43 | 6.99E-02 |
| NULL | GSMG0068312 | NULL | NULL | 1905.14 | 1246.29 | 0.61 | 1.00E-04 |
| NULL | GSMG0068355 | NULL | NULL | 537.56 | 478.86 | 0.17 | 5.16E-01 |
| NULL | GSMG0068359 | NULL | NULL | 207.85 | 24.63 | 3.07 | 2.55E-21 |
| NULL | GSMG0068418 | NULL | NULL | 186.34 | 245.85 | -0.40 | 1.40E-01 |
| NULL | GSMG0068442 | NULL | NULL | 2359.90 | 1705.82 | 0.47 | 2.06E-03 |
| NULL | GSMG0068446 | NULL | NULL | 1724.70 | 2041.99 | -0.24 | 1.64E-01 |
| NULL | GSMG0068449 | NULL | NULL | 4536.91 | 4842.55 | -0.09 | 6.94E-01 |
| NULL | GSMG0068450 | NULL | NULL | 3163.56 | 3076.71 | 0.04 | 8.77E-01 |
| NULL | GSMG0068452 | NULL | NULL | 1151.14 | 875.87 | 0.39 | 9.64E-02 |
| NULL | GSMG0068469 | NULL | NULL | 62.23 | 18.10 | 1.78 | 1.37E-04 |
| NULL | GSMG0068484 | NULL | NULL | 1761.71 | 2047.99 | -0.22 | 2.63E-01 |
| NULL | GSMG0068485 | NULL | NULL | 168.84 | 83.14 | 1.02 | 1.56E-04 |
| NULL | GSMG0068486 | NULL | NULL | 1265.33 | 897.29 | 0.50 | 7.63E-02 |
| NULL | GSMG0068515 | NULL | NULL | 2617.52 | 3825.80 | -0.55 | 4.13E-01 |
| NULL | GSMG0068523 | NULL | NULL | 1258.03 | 819.79 | 0.62 | 3.13E-02 |
| NULL | GSMG0068526 | NULL | NULL | 64.13 | 77.75 | -0.28 | 6.32E-01 |
| NULL | GSMG0068544 | NULL | NULL | 1445.53 | 1350.32 | 0.10 | 7.43E-01 |
| NULL | GSMG0068586 | NULL | NULL | 215.43 | 252.34 | -0.23 | 5.38E-01 |
| NULL | GSMG0068591 | NULL | NULL | 40.97 | 67.00 | -0.73 | 2.11E-01 |
| NULL | GSMG0068610 | NULL | NULL | 733.13 | 653.30 | 0.17 | 6.17E-01 |
| NULL | GSMG0068635 | NULL | NULL | 1192.90 | 527.22 | 1.18 | 3.00E-13 |
| NULL | GSMG0068672 | NULL | NULL | 22773.30 | 6362.24 | 1.84 | 1.80E-47 |
| NULL | GSMG0068695 | NULL | NULL | 481.60 | 542.29 | -0.17 | 5.66E-01 |
| NULL | GSMG0068709 | NULL | NULL | 1098.06 | 1180.58 | -0.10 | 6.67E-01 |
| NULL | GSMG0068710 | NULL | NULL | 367.05 | 137.23 | 1.42 | 2.56E-09 |
| NULL | GSMG0068718 | NULL | NULL | 937.20 | 882.61 | 0.08 | 8.12E-01 |
| NULL | GSMG0068721 | NULL | NULL | 2361.54 | 2748.80 | -0.22 | 3.67E-01 |
| NULL | GSMG0068739 | NULL | NULL | 858.26 | 1395.40 | -0.70 | 5.92E-04 |
| NULL | GSMG0068745 | NULL | NULL | 150.37 | 140.96 | 0.09 | 8.27E-01 |
| NULL | GSMG0068754 | NULL | NULL | 1123.59 | 1049.75 | 0.10 | 8.03E-01 |
| NULL | GSMG0068762 | NULL | NULL | 20043.95 | 10629.28 | 0.92 | 1.87E-08 |

| | | | | | | | |
|---|---|---|---|---|---|---|---|
| NULL | GSMG0068766 | NULL | NULL | 142.92 | 92.36 | 0.63 | 3.61E-02 |
| NULL | GSMG0068770 | NULL | NULL | 2097.92 | 1391.06 | 0.59 | 1.72E-04 |
| NULL | GSMG0068783 | NULL | NULL | 415.38 | 337.57 | 0.30 | 2.43E-01 |
| NULL | GSMG0068792 | NULL | NULL | 2630.71 | 3666.95 | -0.48 | 7.70E-03 |
| NULL | GSMG0068796 | NULL | NULL | 353.90 | 179.49 | 0.98 | 1.76E-06 |
| NULL | GSMG0068804 | NULL | NULL | 6188.08 | 3155.16 | 0.97 | 8.21E-10 |
| NULL | GSMG0068806 | NULL | NULL | 4032.07 | 3044.90 | 0.41 | 7.51E-02 |
| NULL | GSMG0068811 | NULL | NULL | 65.72 | 87.80 | -0.43 | 3.52E-01 |
| NULL | GSMG0068817 | NULL | NULL | 984.77 | 1074.47 | -0.12 | 6.31E-01 |
| NULL | GSMG0068819 | NULL | NULL | 112.75 | 102.56 | 0.14 | 7.67E-01 |
| NULL | GSMG0068848 | NULL | NULL | 413.96 | 381.13 | 0.12 | 6.86E-01 |
| NULL | GSMG0068860 | NULL | NULL | 46.96 | 50.96 | -0.12 | 8.61E-01 |
| NULL | GSMG0068861 | NULL | NULL | 55.38 | 59.77 | -0.12 | 8.60E-01 |
| NULL | GSMG0068871 | NULL | NULL | 765.68 | 657.96 | 0.22 | 3.62E-01 |
| NULL | GSMG0068873 | NULL | NULL | 65.88 | 79.04 | -0.27 | 5.79E-01 |
| NULL | GSMG0068874 | NULL | NULL | 53.76 | 49.69 | 0.11 | 8.73E-01 |
| NULL | GSMG0068876 | NULL | NULL | 933.04 | 1126.48 | -0.27 | 1.67E-01 |
| NULL | GSMG0068898 | NULL | NULL | 2235.58 | 2659.84 | -0.25 | 2.67E-01 |
| NULL | GSMG0068900 | NULL | NULL | 2080.13 | 2192.27 | -0.08 | 7.94E-01 |
| NULL | GSMG0068903 | NULL | NULL | 113.94 | 149.11 | -0.39 | 2.91E-01 |
| NULL | GSMG0068938 | NULL | NULL | 105.46 | 125.26 | -0.25 | 5.51E-01 |
| NULL | GSMG0068939 | NULL | NULL | 133.40 | 171.44 | -0.36 | 2.52E-01 |
| NULL | GSMG0068940 | NULL | NULL | 152.42 | 176.17 | -0.21 | 5.59E-01 |
| NULL | GSMG0068941 | NULL | NULL | 94.05 | 129.32 | -0.46 | 1.98E-01 |
| NULL | GSMG0068943 | NULL | NULL | 2127.45 | 1748.97 | 0.28 | 1.94E-01 |
| NULL | GSMG0068954 | NULL | NULL | 2179.74 | 1493.93 | 0.55 | 2.81E-03 |
| NULL | GSMG0068955 | NULL | NULL | 55.70 | 62.94 | -0.17 | 7.71E-01 |
| NULL | GSMG0068956 | NULL | NULL | 51.42 | 65.26 | -0.35 | 5.00E-01 |
| NULL | GSMG0068964 | NULL | NULL | 900.20 | 1302.63 | -0.53 | 5.33E-03 |
| NULL | GSMG0068988 | NULL | NULL | 670.80 | 728.13 | -0.12 | 7.83E-01 |
| NULL | GSMG0068996 | NULL | NULL | 226.39 | 499.53 | -1.14 | 1.43E-01 |
| NULL | GSMG0069027 | NULL | NULL | 113.86 | 133.49 | -0.23 | 5.83E-01 |
| NULL | GSMG0069055 | NULL | NULL | 1345.21 | 759.92 | 0.82 | 1.94E-06 |

| | | | | | | | |
|---|---|---|---|---|---|---|---|
| NULL | GSMG0069061 | NULL | NULL | 1518.56 | 1174.49 | 0.37 | 4.19E-02 |
| NULL | GSMG0069068 | NULL | NULL | 97.47 | 83.90 | 0.22 | 6.63E-01 |
| NULL | GSMG0069076 | NULL | NULL | 984.77 | 1074.47 | -0.12 | 6.31E-01 |
| NULL | GSMG0069096 | NULL | NULL | 9671.59 | 11910.85 | -0.30 | 1.08E-01 |
| NULL | GSMG0069114 | NULL | NULL | 1123.81 | 801.49 | 0.49 | 3.10E-02 |
| NULL | GSMG0069115 | NULL | NULL | 220.34 | 421.81 | -0.94 | 3.64E-06 |
| NULL | GSMG0069138 | NULL | NULL | 843.20 | 769.53 | 0.13 | 6.71E-01 |
| NULL | GSMG0069147 | NULL | NULL | 491.04 | 318.87 | 0.63 | 1.07E-02 |
| NULL | GSMG0069166 | NULL | NULL | 654.35 | 408.33 | 0.68 | 1.64E-03 |
| NULL | GSMG0069168 | NULL | NULL | 181.66 | 151.51 | 0.26 | 4.39E-01 |
| NULL | GSMG0069170 | NULL | NULL | 1134.87 | 1212.33 | -0.09 | 6.89E-01 |
| NULL | GSMG0069207 | NULL | NULL | 3159.66 | 2482.22 | 0.35 | 9.90E-02 |
| NULL | GSMG0069220 | NULL | NULL | 187.29 | 116.99 | 0.68 | 1.08E-02 |
| NULL | GSMG0069233 | NULL | NULL | 815.38 | 1041.34 | -0.35 | 5.83E-02 |
| NULL | GSMG0069255 | NULL | NULL | 1465.47 | 1567.46 | -0.10 | 7.18E-01 |
| NULL | GSMG0069275 | NULL | NULL | 4076.06 | 3682.69 | 0.15 | 6.04E-01 |
| NULL | GSMG0069277 | NULL | NULL | 706.02 | 441.53 | 0.68 | 2.29E-04 |
| NULL | GSMG0069280 | NULL | NULL | 500.56 | 411.03 | 0.28 | 2.03E-01 |
| NULL | GSMG0069284 | NULL | NULL | 835.91 | 636.59 | 0.39 | 1.23E-01 |
| NULL | GSMG0069291 | NULL | NULL | 1184.94 | 1339.25 | -0.18 | 4.00E-01 |
| NULL | GSMG0069301 | NULL | NULL | 168.63 | 189.81 | -0.17 | 6.84E-01 |
| NULL | GSMG0069316 | NULL | NULL | 89.25 | 81.11 | 0.14 | 8.08E-01 |
| NULL | GSMG0069352 | NULL | NULL | 408.62 | 399.09 | 0.03 | 9.24E-01 |
| NULL | GSMG0069363 | NULL | NULL | 1718.35 | 1497.25 | 0.20 | 3.55E-01 |
| NULL | GSMG0069370 | NULL | NULL | 119.57 | 106.79 | 0.16 | 7.25E-01 |
| NULL | GSMG0069404 | NULL | NULL | 513.14 | 541.30 | -0.08 | 7.95E-01 |
| NULL | GSMG0069410 | NULL | NULL | 47.52 | 50.10 | -0.07 | 9.19E-01 |
| NULL | GSMG0069430 | NULL | NULL | 1465.08 | 962.89 | 0.61 | 1.81E-03 |
| NULL | GSMG0069617 | NULL | NULL | 419.19 | 1290.15 | -1.62 | 6.53E-26 |
| ENSMUSG00000048967 | GSMG0069618 | Yjefn3 | YjeF N-terminal domain containing 3 | 192.59 | 107.73 | 0.83 | 7.90E-03 |
| NULL | GSMG0069622 | NULL | NULL | 3485.90 | 2584.01 | 0.43 | 1.43E-02 |
| NULL | GSMG0069624 | NULL | NULL | 501.02 | 706.63 | -0.50 | 5.72E-03 |
| NULL | GSMG0069627 | NULL | NULL | 4317.77 | 4620.41 | -0.10 | 7.52E-01 |

| | | | | | | | |
|---|---|---|---|---|---|---|---|
| NULL | GSMG0069661 | NULL | NULL | 574.06 | 776.80 | -0.44 | 2.25E-02 |
| NULL | GSMG0069663 | NULL | NULL | 721.25 | 699.72 | 0.04 | 8.93E-01 |
| NULL | GSMG0069667 | NULL | NULL | 759.22 | 360.63 | 1.07 | 1.40E-07 |
| NULL | GSMG0069673 | NULL | NULL | 1370.11 | 1499.47 | -0.13 | 6.52E-01 |
| NULL | GSMG0069694 | NULL | NULL | 2334.03 | 2094.95 | 0.16 | 4.67E-01 |
| NULL | GSMG0069802 | NULL | NULL | 392.45 | 487.26 | -0.31 | 1.54E-01 |
| NULL | GSMG0069805 | NULL | NULL | 62.19 | 75.32 | -0.29 | 6.13E-01 |
| NULL | GSMG0069809 | NULL | NULL | 697.43 | 340.04 | 1.03 | 1.24E-04 |
| NULL | GSMG0069820 | NULL | NULL | 1872.53 | 1470.43 | 0.35 | 4.12E-02 |
| NULL | GSMG0069838 | NULL | NULL | 1455.82 | 1158.94 | 0.33 | 1.69E-01 |
| NULL | GSMG0069845 | NULL | NULL | 2831.33 | 1833.60 | 0.63 | 4.94E-04 |
| NULL | GSMG0069921 | NULL | NULL | 1411.93 | 1631.08 | -0.21 | 3.23E-01 |
| NULL | GSMG0069942 | NULL | NULL | 244.25 | 341.63 | -0.48 | 4.15E-02 |
| NULL | GSMG0069946 | NULL | NULL | 2437.48 | 2466.02 | -0.02 | 9.70E-01 |
| NULL | GSMG0069951 | NULL | NULL | 1598.41 | 1932.99 | -0.27 | 1.48E-01 |
| NULL | GSMG0069964 | NULL | NULL | 1324.00 | 1377.01 | -0.06 | 8.51E-01 |
| NULL | GSMG0069969 | NULL | NULL | 266.13 | 332.52 | -0.32 | 2.20E-01 |
| NULL | GSMG0069974 | NULL | NULL | 315.78 | 318.63 | -0.01 | 9.84E-01 |
| NULL | GSMG0070016 | NULL | NULL | 672.98 | 754.92 | -0.17 | 5.30E-01 |
| NULL | GSMG0070020 | NULL | NULL | 1611.22 | 1413.96 | 0.19 | 4.58E-01 |
| NULL | GSMG0070048 | NULL | NULL | 1120.53 | 1287.78 | -0.20 | 4.63E-01 |
| NULL | GSMG0070074 | NULL | NULL | 893.52 | 1046.67 | -0.23 | 2.39E-01 |
| NULL | GSMG0070118 | NULL | NULL | 298.68 | 296.18 | 0.01 | 9.81E-01 |
| NULL | GSMG0070135 | NULL | NULL | 105.38 | 106.43 | -0.02 | 9.78E-01 |
| NULL | GSMG0070202 | NULL | NULL | 729.84 | 687.40 | 0.09 | 7.40E-01 |
| NULL | GSMG0070206 | NULL | NULL | 91.82 | 23.19 | 1.98 | 4.75E-06 |
| NULL | GSMG0070208 | NULL | NULL | 3675.71 | 3722.02 | -0.02 | 9.53E-01 |
| NULL | GSMG0070232 | NULL | NULL | 5195.31 | 6449.01 | -0.31 | 5.32E-02 |
| NULL | GSMG0070245 | NULL | NULL | 482.48 | 620.40 | -0.36 | 1.27E-01 |
| NULL | GSMG0070271 | NULL | NULL | 1766.30 | 1680.14 | 0.07 | 7.94E-01 |
| NULL | GSMG0070337 | NULL | NULL | 262.82 | 369.46 | -0.49 | 8.19E-02 |
| NULL | GSMG0070344 | NULL | NULL | 307.51 | 236.75 | 0.38 | 1.69E-01 |
| NULL | GSMG0070363 | NULL | NULL | 55.47 | 52.97 | 0.06 | 9.29E-01 |

| | | | | | | | |
|---|---|---|---|---|---|---|---|
| NULL | GSMG0070379 | NULL | NULL | 2486.66 | 2001.02 | 0.31 | 1.16E-01 |
| NULL | GSMG0070380 | NULL | NULL | 9342.23 | 13946.67 | -0.58 | 1.52E-02 |
| NULL | GSMG0070415 | NULL | NULL | 912.36 | 743.36 | 0.30 | 1.23E-01 |
| NULL | GSMG0070416 | NULL | NULL | 602.39 | 628.74 | -0.06 | 8.45E-01 |
| NULL | GSMG0070417 | NULL | NULL | 4715.37 | 5534.64 | -0.23 | 1.99E-01 |
| NULL | GSMG0070424 | NULL | NULL | 2257.01 | 2919.95 | -0.37 | 5.34E-02 |
| NULL | GSMG0070439 | NULL | NULL | 1575.78 | 5020.35 | -1.67 | 2.37E-27 |
| NULL | GSMG0070536 | NULL | NULL | 2258.56 | 2807.50 | -0.31 | 6.12E-02 |
| NULL | GSMG0070562 | NULL | NULL | 551.48 | 524.29 | 0.07 | 8.04E-01 |
| NULL | GSMG0070570 | NULL | NULL | 1225.78 | 1174.17 | 0.06 | 8.15E-01 |
| NULL | GSMG0070595 | NULL | NULL | 2094.17 | 2107.92 | -0.01 | 9.77E-01 |
| NULL | GSMG0070603 | NULL | NULL | 259.62 | 512.17 | -0.98 | 5.84E-07 |
| NULL | GSMG0070618 | NULL | NULL | 2331.20 | 2060.18 | 0.18 | 5.18E-01 |
| NULL | GSMG0070628 | NULL | NULL | 1382.07 | 1665.59 | -0.27 | 1.73E-01 |
| NULL | GSMG0070653 | NULL | NULL | 365.55 | 169.52 | 1.11 | 5.40E-07 |
| NULL | GSMG0070670 | NULL | NULL | 2094.73 | 2443.76 | -0.22 | 2.91E-01 |
| NULL | GSMG0070674 | NULL | NULL | 1491.88 | 1184.33 | 0.33 | 9.78E-02 |
| NULL | GSMG0070710 | NULL | NULL | 756.42 | 1152.81 | -0.61 | 9.24E-04 |
| NULL | GSMG0070714 | NULL | NULL | 1342.12 | 1069.18 | 0.33 | 7.99E-02 |
| NULL | GSMG0070734 | NULL | NULL | 54.48 | 77.99 | -0.52 | 2.50E-01 |
| NULL | GSMG0070741 | NULL | NULL | 1287.23 | 1376.98 | -0.10 | 7.07E-01 |
| NULL | GSMG0070821 | NULL | NULL | 2234.80 | 2006.74 | 0.15 | 5.54E-01 |
| NULL | GSMG0070885 | NULL | NULL | 846.86 | 947.13 | -0.16 | 4.90E-01 |
| NULL | GSMG0070892 | NULL | NULL | 781.57 | 900.77 | -0.20 | 4.73E-01 |
| NULL | GSMG0070893 | NULL | NULL | 603.49 | 786.61 | -0.38 | 6.70E-02 |
| NULL | GSMG0070909 | NULL | NULL | 70.28 | 90.69 | -0.37 | 4.00E-01 |
| NULL | GSMG0070913 | NULL | NULL | 275.56 | 341.44 | -0.31 | 2.54E-01 |
| NULL | GSMG0070986 | NULL | NULL | 999.27 | 1019.66 | -0.03 | 9.23E-01 |

Supplementary Table S2

| Biological processes | | | | |
|---|---|---|---|---|
| GO ID with Link | GO Term | Number of genes | Number of regulated Genes (Up / Down) | P-Value |
| GO:0006084 | acetyl-CoA metabolic process | 7 | 4 (1/3) | 3.12E-02 |
| GO:0030036 | actin cytoskeleton organization | 142 | 30 (10/20) | 5.07E-04 |
| GO:0007015 | actin filament organization | 83 | 17 (11/6) | 1.47E-02 |
| GO:0006919 | endopeptidase activity involved in apoptotic process | 67 | 15 (7/8) | 1.09E-02 |
| GO:0030534 | adult behavior | 38 | 11 (3/8) | 5.72E-03 |
| GO:0007628 | adult walking behavior | 42 | 11 (3/8) | 1.19E-02 |
| GO:0007568 | aging | 173 | 31 (20/11) | 5.78E-03 |
| GO:0006865 | amino acid transport | 42 | 11 (3/8) | 1.19E-02 |
| GO:0001525 | angiogenesis | 239 | 36 (27/9) | 3.69E-02 |
| GO:0006915 | apoptotic process | 570 | 89 (46/43) | 3.14E-04 |
| GO:0014002 | astrocyte development | 17 | 7 (4/3) | 6.70E-03 |
| GO:0007411 | axon guidance | 149 | 45 (10/35) | 2.42E-10 |
| GO:0016199 | axon midline choice point recognition | 5 | 4 (1/3) | 1.05E-02 |
| GO:0007413 | axonal fasciculation | 18 | 8 (2/6) | 1.80E-03 |
| GO:0007409 | axonogenesis | 107 | 24 (7/17) | 8.92E-04 |
| GO:0009058 | biosynthetic process | 37 | 10 (1/9) | 1.45E-02 |
| GO:0032060 | bleb assembly | 11 | 5 (5/0) | 2.37E-02 |
| GO:0007420 | brain development | 216 | 52 (21/31) | 4.56E-08 |
| GO:0050873 | brown fat cell differentiation | 34 | 11 (9/2) | 2.36E-03 |
| GO:0055074 | calcium ion homeostasis | 28 | 8 (5/3) | 2.57E-02 |
| GO:0006816 | calcium ion transport | 141 | 24 (17/7) | 2.79E-02 |
| GO:0048791 | neurotransmitter | 28 | 11 (7/4) | 4.32E-04 |
| GO:0016339 | via plasma membrane cell adhesion molecules | 19 | 6 (2/4) | 4.62E-02 |
| GO:0060317 | transition | 13 | 5 (5/0) | 4.32E-02 |
| GO:0007155 | cell adhesion | 485 | 107 (60/47) | 5.75E-13 |
| GO:0007049 | cell cycle | 614 | 146 (20/126) | 9.73E-21 |
| GO:0030154 | cell differentiation | 780 | 116 (56/60) | 2.71E-04 |
| GO:0051301 | cell division | 374 | 88 (8/80) | 2.09E-12 |
| GO:0016049 | cell growth | 49 | 11 (9/2) | 3.36E-02 |
| GO:0016477 | cell migration | 191 | 36 (15/21) | 1.16E-03 |
| GO:0000902 | cell morphogenesis | 81 | 16 (8/8) | 2.50E-02 |
| GO:0008283 | cell proliferation | 220 | 38 (13/25) | 4.02E-03 |
| GO:0007160 | cell-matrix adhesion | 78 | 16 (13/3) | 1.81E-02 |
| GO:0071313 | cellular response to caffeine | 8 | 4 (3/1) | 4.60E-02 |
| GO:0071277 | cellular response to calcium ion | 53 | 13 (9/4) | 9.41E-03 |
| GO:0071320 | cellular response to cAMP | 50 | 11 (6/5) | 3.81E-02 |
| GO:0071409 | cellular response to cycloheximide | 3 | 3 (2/1) | 3.22E-02 |
| GO:0006974 | stimulus | 420 | 59 (7/52) | 2.83E-02 |
| GO:0035690 | cellular response to drug | 75 | 15 (7/8) | 2.79E-02 |
| GO:0071372 | hormone stimulus | 14 | 6 (0/6) | 1.25E-02 |
| GO:1904322 | cellular response to forskolin | 12 | 5 (1/4) | 3.26E-02 |
| GO:0071480 | cellular response to gamma radiation | 24 | 7 (1/6) | 3.79E-02 |
| GO:0070301 | cellular response to hydrogen peroxide | 57 | 15 (10/5) | 2.38E-03 |
| GO:0071456 | cellular response to hypoxia | 103 | 25 (15/10) | 1.96E-04 |
| GO:0032869 | cellular response to insulin stimulus | 88 | 18 (11/7) | 1.19E-02 |
| GO:0071310 | cellular response to organic substance | 33 | 11 (6/5) | 1.84E-03 |

| | | | | |
|---|---|---|---|---|
| GO:0071560 | growth factor beta stimulus | 61 | 16 (9/7) | 1.66E-03 |
| GO:0071294 | cellular response to zinc ion | 12 | 5 (4/1) | 3.26E-02 |
| GO:0006882 | cellular zinc ion homeostasis | 18 | 6 (4/2) | 3.73E-02 |
| GO:0022010 | central nervous system myelination | 9 | 6 (4/2) | 1.24E-03 |
| GO:0021680 | development | 16 | 7 (2/5) | 4.76E-03 |
| GO:0021987 | cerebral cortex development | 68 | 18 (4/14) | 6.89E-04 |
| GO:0007268 | chemical synaptic transmission | 172 | 27 (11/16) | 4.71E-02 |
| GO:0006935 | chemotaxis | 118 | 27 (15/12) | 2.83E-04 |
| GO:1902476 | chloride transmembrane transport | 42 | 11 (6/5) | 1.19E-02 |
| GO:0006821 | chloride transport | 80 | 17 (9/8) | 1.04E-02 |
| GO:0006695 | cholesterol biosynthetic process | 32 | 12 (1/11) | 3.29E-04 |
| GO:0008203 | cholesterol metabolic process | 89 | 20 (7/13) | 2.61E-03 |
| GO:0007059 | chromosome segregation | 89 | 30 (1/29) | 2.43E-08 |
| GO:0090102 | cochlea development | 29 | 8 (4/4) | 3.09E-02 |
| GO:0021902 | specific neuron type in forebrain | 7 | 5 (1/4) | 3.58E-03 |
| GO:0016358 | dendrite development | 36 | 10 (0/10) | 1.21E-02 |
| GO:0048813 | dendrite morphogenesis | 43 | 10 (4/6) | 3.68E-02 |
| GO:0021542 | dentate gyrus development | 18 | 6 (1/5) | 3.73E-02 |
| GO:0016311 | dephosphorylation | 109 | 21 (11/10) | 1.19E-02 |
| GO:0002024 | diet induced thermogenesis | 13 | 5 (5/0) | 4.32E-02 |
| GO:0071897 | DNA biosynthetic process | 19 | 6 (2/4) | 4.62E-02 |
| GO:0006260 | DNA replication | 123 | 44 (1/43) | 7.38E-13 |
| GO:0006270 | DNA replication initiation | 24 | 15 (0/15) | 1.85E-08 |
| GO:0006268 | replication | 10 | 8 (2/6) | 1.48E-05 |
| GO:0006261 | DNA-dependent DNA replication | 16 | 7 (0/7) | 4.76E-03 |
| GO:0035912 | dorsal aorta morphogenesis | 8 | 4 (4/0) | 4.60E-02 |
| GO:0006231 | dTMP biosynthetic process | 3 | 3 (0/3) | 3.22E-02 |
| GO:0042755 | eating behavior | 38 | 9 (4/5) | 4.62E-02 |
| GO:0048251 | elastic fiber assembly | 7 | 4 (4/0) | 3.12E-02 |
| GO:0048013 | ephrin receptor signaling pathway | 43 | 12 (3/9) | 4.84E-03 |
| GO:0010669 | epithelial structure maintenance | 6 | 4 (2/2) | 1.94E-02 |
| GO:0000132 | orientation | 25 | 8 (3/5) | 1.39E-02 |
| GO:0060079 | excitatory postsynaptic potential | 36 | 13 (5/8) | 2.47E-04 |
| GO:0006887 | exocytosis | 105 | 22 (15/7) | 3.68E-03 |
| GO:0030198 | extracellular matrix organization | 114 | 30 (19/11) | 7.54E-06 |
| GO:0045444 | fat cell differentiation | 79 | 17 (11/6) | 9.23E-03 |
| GO:0006633 | fatty acid biosynthetic process | 74 | 15 (6/9) | 2.51E-02 |
| GO:0030900 | forebrain development | 86 | 16 (6/10) | 4.04E-02 |
| GO:0000082 | G1/S transition of mitotic cell cycle | 62 | 20 (5/15) | 1.67E-05 |
| GO:0000086 | G2/M transition of mitotic cell cycle | 33 | 12 (3/9) | 4.46E-04 |
| GO:0007214 | pathway | 25 | 7 (3/4) | 4.55E-02 |
| GO:0014009 | glial cell proliferation | 8 | 5 (2/3) | 6.56E-03 |
| GO:0072112 | differentiation | 7 | 4 (1/3) | 3.12E-02 |
| GO:0007507 | heart development | 261 | 41 (22/19) | 1.38E-02 |
| GO:0060347 | heart trabecula formation | 15 | 6 (6/0) | 1.71E-02 |
| GO:0007157 | plasma membrane cell adhesion molecules | 52 | 13 (3/10) | 8.05E-03 |
| GO:0021766 | hippocampus development | 73 | 14 (3/11) | 4.66E-02 |
| GO:0034109 | homotypic cell-cell adhesion | 8 | 5 (1/4) | 6.56E-03 |
| GO:0060080 | inhibitory postsynaptic potential | 13 | 8 (3/5) | 1.58E-04 |

| | | | | |
|---|---|---|---|---|
| GO:0035556 | intracellular signal transduction | 400 | 59 (36/23) | 1.14E-02 |
| GO:0006811 | ion transport | 584 | 119 (68/51) | 7.28E-12 |
| GO:0008299 | isoprenoid biosynthetic process | 17 | 6 (0/6) | 2.94E-02 |
| GO:0007612 | learning | 66 | 20 (8/12) | 4.43E-05 |
| GO:0007611 | learning or memory | 51 | 16 (5/11) | 2.12E-04 |
| GO:0002523 | inflammatory response | 13 | 5 (2/3) | 4.32E-02 |
| GO:0051938 | L-glutamate import | 6 | 4 (4/0) | 1.94E-02 |
| GO:0006629 | lipid metabolic process | 459 | 84 (42/42) | 1.40E-06 |
| GO:0097421 | liver regeneration | 29 | 9 (2/7) | 9.63E-03 |
| GO:0007616 | long-term memory | 34 | 10 (8/2) | 8.24E-03 |
| GO:0045576 | mast cell activation | 13 | 5 (4/1) | 4.32E-02 |
| GO:0051899 | membrane depolarization | 23 | 8 (4/4) | 8.59E-03 |
| GO:0097320 | membrane tubulation | 12 | 5 (4/1) | 3.26E-02 |
| GO:0007613 | memory | 83 | 23 (13/10) | 4.67E-05 |
| GO:0051310 | metaphase plate congression | 12 | 8 (1/7) | 8.05E-05 |
| GO:0001578 | microtubule bundle formation | 27 | 9 (1/8) | 6.04E-03 |
| GO:0000226 | microtubule cytoskeleton organization | 96 | 21 (5/16) | 2.76E-03 |
| GO:0007019 | microtubule depolymerization | 14 | 8 (1/7) | 2.87E-04 |
| GO:0046785 | microtubule polymerization | 12 | 5 (2/3) | 3.26E-02 |
| GO:0007018 | microtubule-based movement | 78 | 15 (2/13) | 3.77E-02 |
| GO:0098779 | mitochondrial depolarization | 132 | 25 (14/11) | 7.07E-03 |
| GO:0000278 | mitotic cell cycle | 38 | 11 (1/10) | 5.72E-03 |
| GO:0007076 | mitotic chromosome condensation | 13 | 6 (1/5) | 8.82E-03 |
| GO:0000281 | mitotic cytokinesis | 30 | 12 (2/10) | 1.71E-04 |
| GO:0007067 | mitotic nuclear division | 277 | 73 (6/67) | 6.15E-13 |
| GO:0000070 | mitotic sister chromatid segregation | 23 | 10 (0/10) | 3.81E-04 |
| GO:0051256 | mitotic spindle midzone assembly | 6 | 4 (0/4) | 1.94E-02 |
| GO:0003183 | mitral valve morphogenesis | 8 | 4 (2/2) | 4.60E-02 |
| GO:0016331 | epithelium | 13 | 5 (4/1) | 4.32E-02 |
| GO:0007275 | multicellular organism development | 1029 | 135 (55/80) | 1.07E-02 |
| GO:0042552 | myelination | 60 | 15 (12/3) | 3.94E-03 |
| GO:0050919 | negative chemotaxis | 26 | 8 (5/3) | 1.73E-02 |
| GO:0048843 | involved in axon guidance | 26 | 10 (5/5) | 1.08E-03 |
| GO:1902430 | formation | 7 | 5 (5/0) | 3.58E-03 |
| GO:0090090 | signaling pathway | 102 | 18 (10/8) | 4.47E-02 |
| GO:0007162 | negative regulation of cell adhesion | 42 | 11 (4/7) | 1.19E-02 |
| GO:0045786 | negative regulation of cell cycle | 36 | 9 (4/5) | 3.46E-02 |
| GO:0030308 | negative regulation of cell growth | 125 | 26 (17/9) | 1.62E-03 |
| GO:0030336 | negative regulation of cell migration | 106 | 21 (14/7) | 8.80E-03 |
| GO:0008285 | negative regulation of cell proliferation | 384 | 63 (38/25) | 7.42E-04 |
| GO:0051481 | calcium ion concentration | 17 | 7 (4/3) | 6.70E-03 |
| GO:0045806 | negative regulation of endocytosis | 10 | 5 (4/1) | 1.65E-02 |
| GO:0050680 | proliferation | 72 | 14 (10/4) | 4.23E-02 |
| GO:0070373 | cascade | 66 | 14 (11/3) | 2.22E-02 |
| GO:2000270 | apoptotic process | 9 | 5 (3/2) | 1.08E-02 |
| GO:0007026 | depolymerization | 24 | 9 (3/6) | 2.68E-03 |
| GO:0043524 | process | 160 | 35 (17/18) | 7.77E-05 |
| GO:0010977 | projection development | 64 | 15 (6/9) | 7.24E-03 |
| GO:0090331 | aggregation | 12 | 5 (3/2) | 3.26E-02 |
| GO:0010871 | biosynthetic process | 3 | 3 (3/0) | 3.22E-02 |

| | | | | |
|---|---|---|---|---|
| GO:0043433 | specific DNA binding transcription factor activity | 69 | 18 (12/6) | 8.24E-04 |
| GO:0051154 | cell differentiation | 3 | 3 (1/2) | 3.22E-02 |
| GO:0000122 | negative regulation of transcription from RNA polymerase II promoter | 729 | 106 (45/61) | 1.12E-03 |
| GO:2000678 | regulatory region DNA binding | 13 | 7 (5/2) | 1.35E-03 |
| GO:0030512 | growth factor beta receptor signaling pathway | 54 | 13 (10/3) | 1.09E-02 |
| GO:0007399 | nervous system development | 377 | 95 (26/69) | 2.93E-15 |
| GO:0001755 | neural crest cell migration | 49 | 14 (9/5) | 1.59E-03 |
| GO:0060563 | neuroepithelial cell differentiation | 9 | 5 (0/5) | 1.08E-02 |
| GO:0022008 | neurogenesis | 65 | 15 (4/11) | 8.34E-03 |
| GO:0048666 | neuron development | 58 | 13 (2/11) | 1.90E-02 |
| GO:0030182 | neuron differentiation | 127 | 23 (10/13) | 1.65E-02 |
| GO:0001764 | neuron migration | 124 | 28 (9/19) | 2.68E-04 |
| GO:0031175 | neuron projection development | 139 | 31 (8/23) | 1.49E-04 |
| GO:1990138 | neuron projection extension | 24 | 7 (2/5) | 3.79E-02 |
| GO:0019228 | neuronal action potential | 34 | 10 (3/7) | 8.24E-03 |
| GO:0007269 | neurotransmitter secretion | 28 | 8 (3/5) | 2.57E-02 |
| GO:0006836 | neurotransmitter transport | 45 | 10 (6/4) | 4.77E-02 |
| GO:0009143 | process | 7 | 4 (2/2) | 3.12E-02 |
| GO:0048709 | oligodendrocyte differentiation | 34 | 9 (4/5) | 2.51E-02 |
| GO:0009311 | oligosaccharide metabolic process | 18 | 6 (4/2) | 3.73E-02 |
| GO:0006730 | one-carbon metabolic process | 31 | 11 (2/9) | 1.07E-03 |
| GO:0031100 | organ regeneration | 56 | 16 (7/9) | 6.44E-04 |
| GO:0001649 | osteoblast differentiation | 113 | 21 (10/11) | 1.73E-02 |
| GO:0003151 | outflow tract morphogenesis | 54 | 12 (9/3) | 2.69E-02 |
| GO:0019800 | peptide cross-linking via chondroitin 4-sulfate glycosaminoglycan | 8 | 4 (3/1) | 4.60E-02 |
| GO:0030072 | peptide hormone secretion | 5 | 4 (2/2) | 1.05E-02 |
| GO:0035335 | peptidyl-tyrosine dephosphorylation | 24 | 8 (4/4) | 1.10E-02 |
| GO:0014065 | phosphatidylinositol 3-kinase signaling | 25 | 7 (3/4) | 4.55E-02 |
| GO:0009395 | phospholipid catabolic process | 19 | 6 (3/3) | 4.62E-02 |
| GO:0016310 | phosphorylation | 612 | 99 (44/55) | 3.57E-05 |
| GO:0030168 | platelet activation | 39 | 11 (9/2) | 6.96E-03 |
| GO:0044794 | process | 7 | 4 (3/1) | 3.12E-02 |
| GO:0043065 | positive regulation of apoptotic process | 335 | 66 (39/27) | 1.71E-06 |
| GO:0032781 | positive regulation of ATPase activity | 24 | 7 (4/3) | 3.79E-02 |
| GO:0045773 | positive regulation of axon extension | 40 | 10 (3/7) | 2.39E-02 |
| GO:1902004 | formation | 5 | 4 (3/1) | 1.05E-02 |
| GO:0030513 | pathway | 36 | 9 (6/3) | 3.46E-02 |
| GO:0030501 | mineralization | 37 | 11 (6/5) | 4.65E-03 |
| GO:0050850 | mediated signaling | 24 | 7 (1/6) | 3.79E-02 |
| GO:0010666 | cell apoptotic process | 16 | 9 (7/2) | 1.03E-04 |
| GO:0060045 | cell proliferation | 23 | 7 (3/4) | 3.12E-02 |
| GO:0010942 | positive regulation of cell death | 48 | 11 (10/1) | 2.95E-02 |
| GO:0030335 | positive regulation of cell migration | 203 | 31 (19/12) | 4.54E-02 |
| GO:0008284 | positive regulation of cell proliferation | 542 | 77 (42/35) | 9.59E-03 |
| GO:0001954 | adhesion | 26 | 8 (6/2) | 1.73E-02 |
| GO:0007204 | ion concentration | 148 | 26 (13/13) | 1.51E-02 |
| GO:1903861 | extension | 19 | 6 (5/1) | 4.62E-02 |

| | | | | |
|---|---|---|---|---|
| GO:1900264 | DNA polymerase activity | 8 | 5 (0/5) | 6.56E-03 |
| GO:0001938 | proliferation | 67 | 15 (13/2) | 1.09E-02 |
| GO:0010718 | mesenchymal transition | 34 | 9 (4/5) | 2.51E-02 |
| GO:0045600 | differentiation | 49 | 14 (8/6) | 1.59E-03 |
| GO:0048146 | proliferation | 65 | 13 (6/7) | 4.29E-02 |
| GO:0051491 | assembly | 31 | 9 (4/5) | 1.45E-02 |
| GO:0010628 | positive regulation of gene expression | 399 | 63 (39/24) | 1.93E-03 |
| GO:0014049 | secretion | 11 | 5 (2/3) | 2.37E-02 |
| GO:0010460 | positive regulation of heart rate | 21 | 9 (8/1) | 9.97E-04 |
| GO:0050729 | response | 63 | 14 (10/4) | 1.53E-02 |
| GO:0046330 | positive regulation of JNK cascade | 70 | 15 (11/4) | 1.59E-02 |
| GO:0048170 | neuronal synaptic plasticity | 8 | 4 (1/3) | 4.60E-02 |
| GO:0043406 | activity | 54 | 12 (6/6) | 2.69E-02 |
| GO:0051901 | depolarization | 11 | 6 (5/1) | 3.80E-03 |
| GO:0010976 | development | 138 | 29 (13/16) | 6.99E-04 |
| GO:0045669 | differentiation | 68 | 15 (10/5) | 1.24E-02 |
| GO:0050731 | phosphorylation | 101 | 18 (11/7) | 4.12E-02 |
| GO:0045860 | activity | 64 | 13 (6/7) | 3.87E-02 |
| GO:0001934 | phosphorylation | 184 | 31 (17/14) | 1.36E-02 |
| GO:0061098 | kinase activity | 20 | 7 (5/2) | 1.58E-02 |
| GO:0048661 | cell proliferation | 80 | 16 (10/6) | 2.25E-02 |
| GO:0032230 | transmission, GABAergic | 16 | 7 (3/4) | 4.76E-03 |
| GO:0045909 | positive regulation of vasodilation | 38 | 9 (8/1) | 4.62E-02 |
| GO:0010107 | potassium ion import | 26 | 9 (6/3) | 4.69E-03 |
| GO:0071805 | transport | 90 | 28 (16/12) | 4.85E-07 |
| GO:0006813 | potassium ion transport | 127 | 37 (25/12) | 3.31E-08 |
| GO:0006461 | protein complex assembly | 66 | 15 (10/5) | 9.56E-03 |
| GO:0051260 | protein homooligomerization | 197 | 39 (20/19) | 2.59E-04 |
| GO:0051289 | protein homotetramerization | 74 | 15 (6/9) | 2.51E-02 |
| GO:0017038 | protein import | 4 | 4 (3/1) | 4.58E-03 |
| GO:0043491 | protein kinase B signaling | 38 | 11 (10/1) | 5.72E-03 |
| GO:0034501 | protein localization to kinetochore | 10 | 6 (0/6) | 2.27E-03 |
| GO:0051259 | protein oligomerization | 67 | 16 (7/9) | 4.41E-03 |
| GO:0006468 | protein phosphorylation | 576 | 89 (36/53) | 4.46E-04 |
| GO:0051262 | protein tetramerization | 41 | 10 (4/6) | 2.78E-02 |
| GO:0003184 | pulmonary valve morphogenesis | 11 | 5 (4/1) | 2.37E-02 |
| GO:0009113 | process | 3 | 3 (0/3) | 3.22E-02 |
| GO:0006220 | process | 8 | 4 (0/4) | 4.60E-02 |
| GO:0072593 | process | 38 | 11 (8/3) | 5.72E-03 |
| GO:0031623 | receptor internalization | 40 | 11 (8/3) | 8.41E-03 |
| GO:0060371 | membrane depolarization | 8 | 4 (3/1) | 4.60E-02 |
| GO:0051988 | microtubules to kinetochore | 7 | 4 (0/4) | 3.12E-02 |
| GO:0045607 | differentiation | 7 | 5 (2/3) | 3.58E-03 |
| GO:0050770 | regulation of axonogenesis | 28 | 9 (5/4) | 7.68E-03 |
| GO:0008217 | regulation of blood pressure | 62 | 18 (16/2) | 2.12E-04 |
| GO:0051924 | regulation of calcium ion transport | 37 | 10 (7/3) | 1.45E-02 |
| GO:0017158 | exocytosis | 24 | 7 (4/3) | 3.79E-02 |
| GO:0043949 | regulation of cAMP-mediated signaling | 5 | 4 (3/1) | 1.05E-02 |
| GO:0086004 | contraction | 16 | 6 (5/1) | 2.27E-02 |

| | | | | |
|---|---|---|---|---|
| GO:0010881 | contraction by regulation of the release of sequestered calcium ion | 8 | 5 (3/2) | 6.56E-03 |
| GO:0030155 | regulation of cell adhesion | 42 | 11 (9/2) | 1.19E-02 |
| GO:0051726 | regulation of cell cycle | 112 | 26 (9/17) | 2.98E-04 |
| GO:0001558 | regulation of cell growth | 58 | 15 (11/4) | 2.83E-03 |
| GO:0042127 | regulation of cell proliferation | 227 | 37 (23/14) | 1.14E-02 |
| GO:0051480 | concentration | 35 | 9 (5/4) | 2.96E-02 |
| GO:0048814 | regulation of dendrite morphogenesis | 23 | 7 (3/4) | 3.12E-02 |
| GO:0017157 | regulation of exocytosis | 25 | 9 (7/2) | 3.58E-03 |
| GO:0034765 | transport | 135 | 38 (24/14) | 5.66E-08 |
| GO:0048169 | synaptic plasticity | 28 | 9 (6/3) | 7.68E-03 |
| GO:0042391 | regulation of membrane potential | 97 | 19 (13/6) | 1.47E-02 |
| GO:0031110 | polymerization or depolymerization | 10 | 5 (0/5) | 1.65E-02 |
| GO:0007346 | regulation of mitotic cell cycle | 37 | 11 (4/7) | 4.65E-03 |
| GO:0010975 | development | 30 | 8 (2/6) | 3.67E-02 |
| GO:0048168 | plasticity | 25 | 10 (8/2) | 7.81E-04 |
| GO:2000310 | regulation of N-methyl-D-aspartate selective glutamate receptor activity | 16 | 7 (2/5) | 4.76E-03 |
| GO:0014061 | regulation of norepinephrine secretion | 6 | 4 (2/2) | 1.94E-02 |
| GO:0046578 | transduction | 6 | 4 (1/3) | 1.94E-02 |
| GO:0010880 | calcium ion into cytosol by sarcoplasmic reticulum | 17 | 6 (2/4) | 2.94E-02 |
| GO:0035023 | transduction | 77 | 15 (9/6) | 3.42E-02 |
| GO:0051930 | pain | 41 | 15 (9/6) | 5.66E-05 |
| GO:0048167 | regulation of synaptic plasticity | 42 | 10 (6/4) | 3.21E-02 |
| GO:0003081 | regulation of systemic arterial blood pressure by renin-angiotensin | 5 | 4 (4/0) | 1.05E-02 |
| GO:0061469 | proliferation | 7 | 4 (3/1) | 3.12E-02 |
| GO:2001212 | regulation of vasculogenesis | 3 | 3 (3/0) | 3.22E-02 |
| GO:0051592 | response to calcium ion | 68 | 14 (9/5) | 2.79E-02 |
| GO:0009409 | response to cold | 40 | 12 (9/3) | 2.63E-03 |
| GO:0042493 | response to drug | 339 | 48 (19/29) | 4.24E-02 |
| GO:0045471 | response to ethanol | 117 | 21 (9/12) | 2.46E-02 |
| GO:0001666 | response to hypoxia | 192 | 35 (24/11) | 2.44E-03 |
| GO:0002931 | response to ischemia | 36 | 10 (6/4) | 1.21E-02 |
| GO:0034616 | response to laminar fluid shear stress | 7 | 4 (3/1) | 3.12E-02 |
| GO:0032496 | response to lipopolysaccharide | 197 | 31 (22/9) | 3.21E-02 |
| GO:0035094 | response to nicotine | 45 | 10 (8/2) | 4.77E-02 |
| GO:0010033 | response to organic substance | 65 | 13 (6/7) | 4.29E-02 |
| GO:0010243 | response to organonitrogen compound | 23 | 7 (2/5) | 3.12E-02 |
| GO:0009636 | response to toxic substance | 86 | 17 (11/6) | 2.02E-02 |
| GO:0060041 | retina development in camera-type eye | 84 | 18 (9/9) | 7.48E-03 |
| GO:0048511 | rhythmic process | 128 | 22 (14/8) | 3.26E-02 |
| GO:0097264 | self proteolysis | 8 | 4 (2/2) | 4.60E-02 |
| GO:0071526 | semaphorin-plexin signaling pathway | 34 | 12 (4/8) | 5.96E-04 |
| GO:0019233 | sensory perception of pain | 77 | 19 (11/8) | 1.13E-03 |
| GO:0007605 | sensory perception of sound | 138 | 25 (15/10) | 1.21E-02 |
| GO:0023052 | signaling | 8 | 4 (2/2) | 4.60E-02 |
| GO:0001501 | skeletal system development | 107 | 19 (11/8) | 3.66E-02 |
| GO:0007264 | transduction | 236 | 40 (25/15) | 4.37E-03 |
| GO:0035725 | sodium ion transmembrane transport | 36 | 10 (5/5) | 1.21E-02 |

| | | | | |
|---|---|---|---|---|
| GO:0006814 | sodium ion transport | 124 | 22 (12/10) | 2.38E-02 |
| GO:0006665 | sphingolipid metabolic process | 36 | 9 (5/4) | 3.46E-02 |
| GO:0031577 | spindle checkpoint | 3 | 3 (0/3) | 3.22E-02 |
| GO:0007051 | spindle organization | 16 | 6 (1/5) | 2.27E-02 |
| GO:0001964 | startle response | 22 | 8 (4/4) | 6.57E-03 |
| GO:0006694 | steroid biosynthetic process | 62 | 16 (2/14) | 1.98E-03 |
| GO:0008202 | steroid metabolic process | 84 | 20 (7/13) | 1.28E-03 |
| GO:0016126 | sterol biosynthetic process | 27 | 12 (0/12) | 5.54E-05 |
| GO:0021762 | substantia nigra development | 38 | 10 (6/4) | 1.73E-02 |
| GO:0043129 | surfactant homeostasis | 13 | 5 (4/1) | 4.32E-02 |
| GO:0051932 | synaptic transmission, GABAergic | 12 | 6 (3/3) | 5.95E-03 |
| GO:0035249 | synaptic transmission, glutamatergic | 29 | 11 (5/6) | 5.96E-04 |
| GO:0021537 | telencephalon development | 30 | 8 (2/6) | 3.67E-02 |
| GO:0000723 | telomere maintenance | 38 | 10 (1/9) | 1.73E-02 |
| GO:0046654 | tetrahydrofolate biosynthetic process | 4 | 4 (1/3) | 4.58E-03 |
| GO:0042403 | thyroid hormone metabolic process | 10 | 6 (3/3) | 2.27E-03 |
| GO:0007179 | receptor signaling pathway | 75 | 15 (13/2) | 2.79E-02 |
| GO:0007169 | transmembrane receptor protein tyrosine kinase signaling pathway | 100 | 19 (8/11) | 1.98E-02 |
| GO:0055085 | transmembrane transport | 364 | 60 (34/26) | 8.93E-04 |
| GO:0006810 | transport | 1822 | 242 (131/111) | 2.88E-04 |
| GO:0001570 | vasculogenesis | 68 | 14 (11/3) | 2.79E-02 |
| GO:0042310 | vasoconstriction | 19 | 6 (5/1) | 4.62E-02 |
| GO:0060412 | ventricular septum morphogenesis | 38 | 13 (10/3) | 4.36E-04 |
| GO:0034372 | remodeling | 7 | 4 (2/2) | 3.12E-02 |
| GO:0042060 | wound healing | 94 | 21 (17/4) | 2.13E-03 |

| Cellular component | | | | |
|---|---|---|---|---|
| **GO ID with Link** | **GO Term** | **umber of gen** | **er of regulated genes (Up /** | **P-Value** |
| GO:0015629 | actin cytoskeleton | 201 | 31 (14/17) | 2.96E-02 |
| GO:0005912 | adherens junction | 55 | 14 (7/7) | 3.77E-03 |
| GO:0031225 | anchored component of membrane | 141 | 36 (14/22) | 8.81E-07 |
| GO:0045177 | apical part of cell | 115 | 23 (12/11) | 3.77E-03 |
| GO:0016324 | apical plasma membrane | 328 | 52 (37/15) | 2.61E-03 |
| GO:0030424 | axon | 370 | 87 (35/52) | 6.79E-13 |
| GO:0043203 | axon hillock | 8 | 4 (2/2) | 4.28E-02 |
| GO:0043679 | axon terminus | 86 | 21 (16/5) | 4.67E-04 |
| GO:0005605 | basal lamina | 19 | 9 (6/3) | 3.72E-04 |
| GO:0009925 | basal plasma membrane | 42 | 10 (6/4) | 2.74E-02 |
| GO:0005604 | basement membrane | 97 | 28 (18/10) | 1.44E-06 |
| GO:0016323 | basolateral plasma membrane | 203 | 32 (19/13) | 2.05E-02 |
| GO:0005901 | caveola | 86 | 22 (14/8) | 1.65E-04 |
| GO:0044297 | cell body | 102 | 19 (5/14) | 1.85E-02 |
| GO:0030054 | cell junction | 718 | 138 (75/63) | 1.63E-12 |
| GO:0042995 | cell projection | 712 | 115 (52/63) | 2.40E-06 |
| GO:0009986 | cell surface | 629 | 127 (74/53) | 4.03E-13 |
| GO:0005911 | cell-cell junction | 201 | 37 (16/21) | 9.31E-04 |
| GO:0097149 | centralspindlin complex | 3 | 3 (0/3) | 3.05E-02 |
| GO:0005813 | centrosome | 435 | 69 (19/50) | 4.94E-04 |
| GO:0034707 | chloride channel complex | 46 | 14 (8/6) | 6.45E-04 |

| | | | | |
|---|---|---|---|---|
| GO:0000785 | chromatin | 126 | 24 (6/18) | 5.66E-03 |
| GO:0005694 | chromosome | 344 | 67 (3/64) | 8.27E-07 |
| GO:0032133 | chromosome passenger complex | 5 | 4 (0/4) | 9.74E-03 |
| GO:0000775 | chromosome, centromeric region | 141 | 49 (1/48) | 3.91E-14 |
| GO:0005581 | collagen trimer | 83 | 18 (17/1) | 5.02E-03 |
| GO:0000777 | condensed chromosome kinetochore | 84 | 26 (0/26) | 9.01E-07 |
| GO:0000940 | condensed chromosome outer kinetoch | 9 | 6 (0/6) | 1.09E-03 |
| GO:0000778 | condensed nuclear chromosome kineto | 7 | 5 (0/5) | 3.23E-03 |
| GO:0030864 | cortical actin cytoskeleton | 43 | 13 (9/4) | 1.16E-03 |
| GO:0031390 | Ctf18 RFC-like complex | 8 | 4 (0/4) | 4.28E-02 |
| GO:0005737 | cytoplasm | 6631 | 890 (385/505) | 9.34E-22 |
| GO:0031410 | cytoplasmic vesicle | 646 | 86 (54/32) | 1.60E-02 |
| GO:0030659 | cytoplasmic vesicle membrane | 120 | 23 (14/9) | 6.37E-03 |
| GO:0005856 | cytoskeleton | 1113 | 185 (54/131) | 1.25E-10 |
| GO:0005829 | cytosol | 1784 | 215 (109/106) | 1.60E-02 |
| GO:0030425 | dendrite | 490 | 108 (51/57) | 8.32E-14 |
| GO:0043198 | dendritic shaft | 61 | 18 (8/10) | 1.21E-04 |
| GO:0043197 | dendritic spine | 148 | 36 (19/17) | 2.92E-06 |
| GO:0071953 | elastic fiber | 4 | 4 (3/1) | 4.22E-03 |
| GO:0005783 | endoplasmic reticulum | 1323 | 162 (93/69) | 2.28E-02 |
| GO:0070062 | extracellular exosome | 2674 | 359 (203/156) | 1.54E-07 |
| GO:0031012 | extracellular matrix | 294 | 69 (46/23) | 2.20E-10 |
| GO:0005576 | extracellular region | 1753 | 243 (160/83) | 2.48E-06 |
| GO:0005615 | extracellular space | 1504 | 201 (131/70) | 1.73E-04 |
| GO:0031234 | extrinsic component of cytoplasmic side | 68 | 13 (9/4) | 4.85E-02 |
| GO:0030175 | filopodium | 77 | 18 (8/10) | 2.20E-03 |
| GO:1902711 | GABA-A receptor complex | 18 | 7 (3/4) | 8.00E-03 |
| GO:0005794 | Golgi apparatus | 1190 | 164 (81/83) | 1.76E-04 |
| GO:0005796 | Golgi lumen | 21 | 7 (4/3) | 1.78E-02 |
| GO:0030426 | growth cone | 159 | 42 (17/25) | 3.36E-08 |
| GO:0043657 | host cell | 3 | 3 (2/1) | 3.05E-02 |
| GO:0030173 | integral component of Golgi membrane | 50 | 12 (10/2) | 1.28E-02 |
| GO:0005887 | integral component of plasma membrar | 1126 | 174 (94/80) | 1.04E-07 |
| GO:0014704 | intercalated disc | 51 | 12 (4/8) | 1.49E-02 |
| GO:0043231 | intracellular membrane-bounded organe | 751 | 101 (53/48) | 6.95E-03 |
| GO:0044224 | juxtaparanode region of axon | 10 | 6 (5/1) | 2.00E-03 |
| GO:0005871 | kinesin complex | 52 | 15 (2/13) | 6.93E-04 |
| GO:0000776 | kinetochore | 120 | 39 (0/39) | 1.98E-10 |
| GO:0043256 | laminin complex | 3 | 3 (2/1) | 3.05E-02 |
| GO:0042555 | MCM complex | 9 | 7 (0/7) | 8.31E-05 |
| GO:0072687 | meiotic spindle | 9 | 7 (4/3) | 8.31E-05 |
| GO:0016020 | membrane | 6998 | 989 (512/477) | 8.91E-35 |
| GO:0045121 | membrane raft | 262 | 45 (25/20) | 1.09E-03 |
| GO:0001527 | microfibril | 11 | 5 (4/1) | 2.16E-02 |
| GO:0005874 | microtubule | 327 | 78 (17/61) | 5.51E-12 |
| GO:0015630 | microtubule cytoskeleton | 158 | 26 (3/23) | 2.35E-02 |
| GO:0030496 | midbody | 130 | 28 (6/22) | 3.84E-04 |
| GO:0072686 | mitotic spindle | 47 | 14 (3/11) | 8.08E-04 |
| GO:0043209 | myelin sheath | 192 | 42 (28/14) | 6.85E-06 |
| GO:0031262 | Ndc80 complex | 4 | 4 (0/4) | 4.22E-03 |

| | | | | |
|---|---|---|---|---|
| GO:0031594 | neuromuscular junction | 64 | 16 (7/9) | 2.11E-03 |
| GO:0043005 | neuron projection | 420 | 83 (37/46) | 1.79E-08 |
| GO:0043025 | neuronal cell body | 534 | 124 (61/63) | 8.97E-18 |
| GO:0032809 | neuronal cell body membrane | 29 | 10 (7/3) | 2.12E-03 |
| GO:0017146 | NMDA selective glutamate receptor cor | 12 | 6 (3/3) | 5.27E-03 |
| GO:0000784 | nuclear chromosome, telomeric region | 127 | 21 (1/20) | 4.14E-02 |
| GO:0005635 | nuclear envelope | 158 | 27 (17/10) | 1.33E-02 |
| GO:0005720 | nuclear heterochromatin | 27 | 10 (3/7) | 1.20E-03 |
| GO:0031965 | nuclear membrane | 226 | 41 (15/26) | 6.40E-04 |
| GO:0005654 | nucleoplasm | 1935 | 237 (72/165) | 5.92E-03 |
| GO:0005634 | nucleus | 6019 | 685 (242/443) | 3.77E-03 |
| GO:0033270 | paranode region of axon | 15 | 6 (5/1) | 1.53E-02 |
| GO:0043204 | perikaryon | 150 | 37 (26/11) | 1.46E-06 |
| GO:0048471 | perinuclear region of cytoplasm | 692 | 125 (66/59) | 1.21E-09 |
| GO:0005886 | plasma membrane | 4874 | 611 (332/279) | 7.30E-08 |
| GO:0002102 | podosome | 32 | 8 (4/4) | 4.43E-02 |
| GO:0098794 | postsynapse | 26 | 8 (3/5) | 1.50E-02 |
| GO:0014069 | postsynaptic density | 239 | 60 (36/24) | 2.32E-10 |
| GO:0045211 | postsynaptic membrane | 222 | 59 (32/27) | 2.95E-11 |
| GO:0034705 | potassium channel complex | 11 | 7 (5/2) | 3.79E-04 |
| GO:0098793 | presynapse | 63 | 15 (10/5) | 4.90E-03 |
| GO:0042734 | presynaptic membrane | 75 | 22 (10/12) | 1.87E-05 |
| GO:0043234 | protein complex | 628 | 80 (41/39) | 4.92E-02 |
| GO:0005578 | proteinaceous extracellular matrix | 316 | 84 (55/29) | 1.14E-15 |
| GO:0031298 | replication fork protection complex | 6 | 4 (0/4) | 1.79E-02 |
| GO:0032587 | ruffle membrane | 79 | 17 (12/5) | 7.13E-03 |
| GO:0042383 | sarcolemma | 118 | 28 (19/9) | 7.04E-05 |
| GO:0030667 | secretory granule membrane | 26 | 7 (5/2) | 4.81E-02 |
| GO:0005819 | spindle | 116 | 29 (5/24) | 1.85E-05 |
| GO:0005876 | spindle microtubule | 40 | 10 (0/10) | 2.03E-02 |
| GO:0051233 | spindle midzone | 19 | 8 (1/7) | 2.20E-03 |
| GO:0000922 | spindle pole | 106 | 24 (2/22) | 5.29E-04 |
| GO:0001725 | stress fiber | 67 | 16 (7/9) | 3.39E-03 |
| GO:0045202 | synapse | 505 | 116 (65/51) | 3.62E-16 |
| GO:0043083 | synaptic cleft | 8 | 5 (2/3) | 5.93E-03 |
| GO:0008021 | synaptic vesicle | 126 | 23 (15/8) | 1.12E-02 |
| GO:0043195 | terminal bouton | 113 | 20 (10/10) | 2.54E-02 |
| GO:0030133 | transport vesicle | 64 | 15 (10/5) | 5.69E-03 |
| GO:0030315 | T-tubule | 57 | 14 (7/7) | 5.22E-03 |
| GO:0031982 | vesicle | 164 | 28 (16/12) | 1.17E-02 |
| GO:0008076 | voltage-gated potassium channel comp | 79 | 29 (13/16) | 2.76E-09 |
| GO:0001518 | voltage-gated sodium channel complex | 14 | 6 (2/4) | 1.11E-02 |

Supplementary Table S3

| FIBROBLASTS | | | | | | | | | | | | | |
|---|---|---|---|---|---|---|---|---|---|---|---|---|---|
| | | | Expression in single-cell (Vanlandewijck et al., 2018) | | | | | Cortical MV RNA seq analysis: P15 vs P5 | | | | | |
| | | | | | | | | P15 vs P5 | | RPKM MV / RPKM FB | | | |
| Cluster | Gene | Name | % of cells in cluster | % of cells in other clusters | logFC | Adjusted P-value | RPKM FB | log2Fold Change | Adjusted P-value | P5 | P5/FB | P15 | P15/FB |
| FB | Lum | lumican | 87.2% | 1.5% | 3.8 | 0.0E+00 | 42.8 | 2.3 | 9.3E-21 | 2.6 | 0.1 | 13.4 | 0.3 |
| FB | Col6a1 | collagen, type V | 89.5% | 1.1% | 2.1 | 0.0E+00 | 7.7 | 2.1 | 7.6E-33 | 10.9 | 1.4 | 47.0 | 6.1 |
| FB | Gsn | gelsolin | 89.5% | 16.3% | 2.8 | 1.2E-74 | 34.2 | 1.8 | 1.6E-17 | 13.1 | 0.4 | 44.6 | 1.3 |
| FB | Alpl | alkaline phosph | 64.0% | 22.6% | 1.7 | 4.2E-17 | 13.8 | 1.6 | 4.5E-25 | 4.4 | 0.3 | 13.8 | 1.0 |
| FB | Ctgf | connective tissu | 84.9% | 21.1% | 1.6 | 2.7E-47 | 6.2 | 1.6 | 1.6E-12 | 22.1 | 3.6 | 68.8 | 11.1 |
| FB | F3 | coagulation fac | 79.1% | 8.7% | 1.7 | 1.0E-91 | 12.5 | 1.5 | 7.5E-16 | 7.8 | 0.6 | 22.8 | 1.8 |
| FB | Serping1 | serine (or cystei | 94.2% | 18.3% | 2.5 | 2.7E-82 | 23.7 | 1.3 | 1.9E-12 | 11.7 | 0.5 | 29.6 | 1.2 |
| FB | Vcam1 | vascular cell ad | 80.2% | 15.8% | 1.9 | 9.5E-58 | 16.1 | 1.3 | 5.9E-13 | 6.3 | 0.4 | 15.9 | 1.0 |
| FB | Snhg11 | small nucleolar | 73.3% | 3.7% | 1.8 | 3.6E-179 | 5.4 | 1.2 | 2.2E-19 | 284.7 | 53.0 | 649.8 | 121.1 |
| FB | Col5a1 | collagen, type V | 83.7% | 4.3% | 1.6 | 1.2E-201 | 4.3 | 1.1 | 2.0E-10 | 10.9 | 2.5 | 24.3 | 5.7 |
| FB | Lama1 | laminin, alpha 1 | 64.0% | 2.0% | 1.6 | 5.7E-206 | 4.3 | 1.0 | 1.6E-09 | 8.9 | 2.1 | 18.5 | 4.4 |
| FB | Lbp | lipopolysacchar | 67.4% | 2.9% | 1.8 | 2.8E-180 | 5.6 | 1.0 | 5.4E-05 | 9.2 | 1.6 | 18.7 | 3.3 |
| FB | Tgfbi | transforming gr | 83.7% | 8.3% | 3.2 | 9.2E-124 | 33.2 | 0.9 | 4.6E-05 | 2.6 | 0.1 | 5.0 | 0.2 |
| FB | Ptgds | prostaglandin D | 93.0% | 9.8% | 3.2 | 6.1E-112 | 155.4 | 0.9 | 4.2E-05 | 1105.0 | 7.1 | 2054.2 | 13.2 |
| FB | Mfap4 | microfibrillar-a | 66.3% | 0.6% | 2.1 | 0.0E+00 | 7.5 | 0.8 | 5.4E-05 | 9.4 | 1.2 | 16.9 | 2.2 |
| FB | Coch | cochlin | 68.6% | 1.6% | 2.9 | 1.5E-257 | 18.6 | 0.7 | 2.1E-05 | 8.7 | 0.5 | 14.6 | 0.8 |
| FB | Col3a1 | collagen, type I | 87.2% | 7.4% | 2.3 | 2.8E-147 | 11.9 | 0.7 | 1.4E-05 | 61.7 | 5.2 | 99.1 | 8.3 |
| FB | Col6a2 | collagen, type V | 86.0% | 6.1% | 1.6 | 4.7E-159 | 4.7 | 0.6 | 1.1E-05 | 15.7 | 3.3 | 24.2 | 5.2 |
| FB | Dcn | decorin | 96.5% | 9.1% | 4.5 | 1.1E-160 | 124.3 | 0.6 | 6.7E-04 | 48.5 | 0.4 | 73.9 | 0.6 |
| FB | Cygb | cytoglobin | 72.1% | 15.6% | 1.5 | 5.6E-47 | 5.8 | 0.6 | 1.2E-02 | 2.2 | 0.4 | 3.3 | 0.6 |
| FB | Col1a2 | collagen, type I | 98.8% | 23.2% | 2.2 | 2.8E-76 | 15.9 | 0.5 | 1.1E-04 | 39.6 | 2.5 | 56.4 | 3.5 |
| FB | Socs3 | suppressor of cy | 67.4% | 10.0% | 1.6 | 1.1E-60 | 5.7 | 0.4 | 5.2E-02 | 2.1 | 0.4 | 2.8 | 0.5 |
| FB | Itga8 | integrin alpha 8 | 67.4% | 6.7% | 1.7 | 1.6E-91 | 5.4 | 0.4 | 4.0E-02 | 6.2 | 1.1 | 8.3 | 1.5 |
| FB | Serpinf1 | serine (or cystei | 84.9% | 1.9% | 2.4 | 0.0E+00 | 10.7 | 0.4 | 3.7E-02 | 25.9 | 2.4 | 33.5 | 3.1 |
| FB | Islr | immunoglobuli | 86.0% | 4.9% | 1.7 | 3.0E-191 | 5.4 | 0.3 | 1.4E-01 | 15.4 | 2.9 | 18.9 | 3.5 |
| FB | Plekha6 | pleckstrin homo | 68.6% | 2.5% | 1.5 | 4.5E-204 | 3.7 | 0.2 | 3.5E-01 | 3.4 | 0.9 | 4.1 | 1.1 |
| FB | Efemp1 | epidermal grow | 89.5% | 2.8% | 2.9 | 1.5E-299 | 18.4 | 0.2 | 5.5E-01 | 3.0 | 0.2 | 3.4 | 0.2 |
| FB | Olfml3 | olfactomedin-li | 83.7% | 6.5% | 2.0 | 7.0E-148 | 8.4 | 0.0 | 8.2E-01 | 5.7 | 0.7 | 5.9 | 0.7 |
| FB | Igfbp5 | insulin-like gro | 69.8% | 5.9% | 2.6 | 6.6E-113 | 15.6 | -0.1 | 4.6E-01 | 8.4 | 0.5 | 7.7 | 0.5 |

| | | | | | | | | | | | | | |
|---|---|---|---|---|---|---|---|---|---|---|---|---|---|
| FB | Ogn | osteoglycin | 81.4% | 13.8% | 1.5 | 6.1E-66 | 8.7 | -0.2 | 2.4E-01 | 8.9 | 1.0 | 7.7 | 0.9 |
| FB | Col1a1 | collagen, type I | 93.0% | 2.8% | 2.5 | 0.0E+00 | 11.2 | -0.2 | 2.0E-01 | 41.6 | 3.7 | 35.4 | 3.2 |
| FB | Ccdc80 | coiled-coil dom | 65.1% | 8.4% | 1.8 | 1.7E-70 | 6.6 | -0.3 | 1.5E-01 | 4.3 | 0.7 | 3.6 | 0.6 |
| FB | Igfbp2 | insulin-like grow | 61.6% | 3.0% | 2.0 | 2.4E-145 | 6.9 | -0.3 | 1.0E-01 | 88.9 | 12.9 | 72.9 | 10.6 |
| FB | Rbp1 | retinol binding | 83.7% | 13.5% | 1.7 | 4.8E-79 | 6.1 | -0.3 | 5.7E-02 | 25.9 | 4.3 | 20.9 | 3.4 |
| FB | Aebp1 | AE binding pro | 79.1% | 3.6% | 1.6 | 1.3E-203 | 4.3 | -0.5 | 2.5E-02 | 13.2 | 3.1 | 9.6 | 2.2 |
| FB | Cpxm1 | carboxypeptidas | 61.6% | 1.4% | 1.8 | 9.0E-233 | 5.5 | -0.7 | 1.7E-04 | 12.2 | 2.2 | 7.5 | 1.4 |
| FB | Igfbp4 | insulin-like grow | 88.4% | 9.7% | 3.1 | 4.6E-125 | 27.0 | -0.8 | 2.5E-11 | 100.3 | 3.7 | 56.1 | 2.1 |
| FB | Spp1 | secreted phosph | 72.1% | 15.5% | 1.9 | 1.2E-43 | 23.4 | -1.2 | 3.3E-06 | 10.8 | 0.5 | 4.8 | 0.2 |

| PERICYTES | | | | | | | | | | | | | |
|---|---|---|---|---|---|---|---|---|---|---|---|---|---|
| | | | | Expression in single-cell (Vanlandewijck et al., 2018) | | | | Cortical MV RNA seq analysis: P15 vs P5 | | | | | |
| | | | | | | | | P15 vs P5 | | | RPKM MV / RPKM PC | | |
| Cluster | Gene name | Description | % of cells in cluster | % of cells exp in other clusters | logFC | Adjusted P-value | RPKM PC | log2FoldChange | Adjusted P-value | P5 | P5/PC | P15 | P15/PC |
| PC | Slc6a20a | solute carrier family 6 | 99.10% | 13.2% | 2.3 | 0.0E+00 | 16.9 | 1.0 | 4.6E-09 | 41.7 | 2.5 | 86.5 | 5.1 |
| PC | Pla1a | phospholipase A1 me | 87.50% | 9.3% | 1.9 | 0.0E+00 | 8.0 | 0.9 | 9.5E-06 | 5.4 | 0.7 | 9.8 | 1.2 |
| PC | Enpep | glutamyl aminopeptid | 88.30% | 13.1% | 1.8 | 0.0E+00 | 11.0 | 0.8 | 7.3E-06 | 11.5 | 1.0 | 20.7 | 1.9 |
| PC | Ggt1 | gamma-glutamyltrans | 81.60% | 9.1% | 1.9 | 0.0E+00 | 7.0 | 0.7 | 3.8E-03 | 3.4 | 0.5 | 5.7 | 0.8 |
| PC | Ggt5 | gamma-glutamyltrans | 86.60% | 14.3% | 1.8 | 0.0E+00 | 9.2 | 0.6 | 2.9E-03 | 11.5 | 1.3 | 17.4 | 1.9 |
| PC | Art3 | ADP-ribosyltransfera | 90.50% | 12.4% | 2.2 | 0.0E+00 | 16.0 | 0.5 | 1.6E-02 | 66.4 | 4.1 | 94.1 | 5.9 |
| PC | Plxdc1 | plexin domain contai | 82.20% | 8.8% | 1.8 | 0.0E+00 | 7.5 | 0.3 | 1.1E-01 | 21.6 | 2.9 | 27.0 | 3.6 |
| PC | Trpc3 | transient receptor pot | 81.30% | 9.8% | 1.6 | 0.0E+00 | 5.3 | 0.2 | 2.6E-01 | 17.7 | 3.3 | 20.9 | 3.9 |
| PC | Angptl2 | angiopoietin-like 2 | 67.10% | 11.4% | 1.6 | 1.1E-259 | 6.9 | 0.1 | 6.5E-01 | 11.1 | 1.6 | 12.3 | 1.8 |
| PC | Cd248 | CD248 antigen, endo | 88.60% | 14.6% | 1.6 | 0.0E+00 | 8.1 | -0.3 | 9.0E-02 | 69.8 | 8.6 | 58.1 | 7.2 |
| PC | Kcnj8 | potassium inwardly-r | 98.80% | 15.2% | 2.9 | 0.0E+00 | 33.4 | -0.3 | 6.6E-02 | 33.2 | 1.0 | 26.5 | 0.8 |
| PC | Ifi30 | interferon gamma ind | 77.10% | 11.9% | 1.6 | 0.0E+00 | 5.6 | -0.3 | 3.0E-01 | 11.0 | 2.0 | 8.7 | 1.5 |
| PC | Abcc9 | ATP-binding cassette | 97.20% | 14.4% | 2.6 | 0.0E+00 | 21.1 | -0.6 | 1.1E-03 | 67.8 | 3.2 | 43.8 | 2.1 |
| PC | Uchl1 | ubiquitin carboxy-ter | 80.10% | 12.7% | 1.7 | 0.0E+00 | 5.2 | -0.7 | 1.9E-05 | 214.8 | 41.5 | 136.3 | 26.3 |

| VASCULAR SMOOTH MUSCLE CELLS | | | | | | | | | | | | | |
|---|---|---|---|---|---|---|---|---|---|---|---|---|---|
| | | | Expression in single-cell (Vanlandewijck et al., 2018) | | | | | Cortical MV RNA seq analysis: P15 vs P5 | | | | | |
| | | | | | | | | P15 vs P5 | | RPKM MV / RPKM VSMC | | | |
| Cluster | Gene symbol | Description | % of cells in cluster | % of cells in other clusters | logFC | Adjusted P-value | RPKM VSMC | log2FoldChange | Adjusted P-value | P5 | P5/VSMC | P15 | P15/VSMC |
| VSMC | Pdlim3 | PDZ and LIM domain 3 | 62.0% | 2.1% | 1.8 | 0.0E+00 | 4.9 | 3.8 | 9.6E-12 | 0.2 | 0.0 | 3.3 | 0.8 |
| VSMC | Acta2 | actin, alpha 2, smooth muscle, aorta | 95.3% | 36.3% | 5.1 | 1.2E-170 | 191.0 | 2.6 | 2.9E-08 | 29.4 | 0.2 | 182.3 | 2.1 |
| VSMC | Myh11 | myosin, heavy polypeptide 11, smooth muscle | 93.3% | 18.9% | 3.4 | 8.5E-244 | 38.0 | 2.6 | 1.1E-03 | 2.3 | 0.1 | 14.0 | 4.7 |
| VSMC | Tagln | transgelin | 83.2% | 12.4% | 3.4 | 5.8E-244 | 30.6 | 2.6 | 8.3E-32 | 4.0 | 0.1 | 23.9 | 39.4 |
| VSMC | Nr4a1 | nuclear receptor subfamily 4, group A, member 1 | 75.8% | 33.3% | 1.6 | 2.1E-68 | 11.8 | 2.3 | 2.1E-28 | 8.2 | 0.7 | 41.0 | 1.2 |
| VSMC | Mustn1 | musculoskeletal, embryonic nuclear protein 1 | 87.2% | 11.1% | 2.0 | 1.1E-273 | 8.1 | 2.1 | 1.7E-17 | 1.9 | 0.2 | 8.3 | 1.9 |
| VSMC | Nrip2 | nuclear receptor interacting protein 2 | 85.9% | 18.4% | 2.1 | 4.3E-203 | 7.9 | 1.9 | 3.8E-17 | 4.4 | 0.6 | 17.1 | 0.7 |
| VSMC | Ltbp1 | latent transforming growth factor beta binding protein 1 | 62.0% | 8.8% | 1.6 | 7.1E-161 | 4.6 | 1.9 | 1.1E-07 | 1.2 | 0.3 | 4.3 | 3.5 |
| VSMC | Tpm2 | tropomyosin 2, beta | 97.3% | 25.7% | 2.9 | 4.4E-221 | 28.0 | 1.7 | 6.2E-15 | 16.2 | 0.6 | 53.6 | 1.1 |
| VSMC | Pln | phospholamban | 65.3% | 12.6% | 2.5 | 1.3E-145 | 13.2 | 1.0 | 9.0E-06 | 2.1 | 0.2 | 4.5 | 1.0 |
| VSMC | Filip1l | filamin A interacting protein 1-like | 88.6% | 27.3% | 1.7 | 2.0E-139 | 11.2 | 0.9 | 4.8E-07 | 8.8 | 0.8 | 16.3 | 1.0 |
| VSMC | Rcan2 | regulator of calcineurin 2 | 66.7% | 14.7% | 1.7 | 6.6E-124 | 7.5 | 0.8 | 5.2E-05 | 3.1 | 0.4 | 5.4 | 1.7 |
| VSMC | Des | desmin | 86.5% | 19.9% | 1.6 | 1.9E-166 | 7.7 | 0.7 | 1.1E-04 | 22.1 | 2.9 | 36.4 | 5.4 |
| VSMC | Rasl12 | RAS-like, family 12 | 88.6% | 17.7% | 1.6 | 2.9E-188 | 7.7 | 0.7 | 1.2E-05 | 5.9 | 0.8 | 9.6 | 0.9 |
| VSMC | Npy1r | neuropeptide Y receptor Y1 | 71.4% | 8.0% | 1.7 | 2.2E-218 | 6.1 | 0.7 | 3.3E-02 | 6.7 | 1.1 | 10.7 | 2.2 |
| VSMC | Palld | palladin, cytoskeletal associated protein | 62.6% | 6.5% | 1.7 | 2.9E-202 | 4.9 | 0.6 | 6.1E-02 | 3.7 | 0.7 | 5.4 | 0.3 |
| VSMC | Mylk | myosin, light polypeptide kinase | 99.0% | 39.6% | 1.9 | 4.7E-158 | 16.4 | 0.4 | 2.1E-01 | 25.6 | 1.6 | 33.6 | 0.4 |
| VSMC | Uba2 | ubiquitin-like modifier activating enzyme 2 | 91.2% | 37.6% | 1.6 | 8.6E-118 | 11.4 | 0.1 | 6.4E-01 | 56.1 | 4.9 | 61.0 | 0.7 |
| VSMC | Atp1b1 | ATPase, Na+/K+ transporting, beta 1 polypeptide | 69.4% | 7.9% | 1.6 | 9.2E-217 | 4.5 | 0.0 | 9.3E-01 | 178.3 | 39.8 | 176.3 | 4.6 |
| VSMC | Mcam | melanoma cell adhesion molecule | 96.0% | 38.3% | 1.6 | 1.2E-128 | 19.7 | -0.1 | 8.2E-01 | 94.4 | 4.8 | 91.3 | 1.5 |
| VSMC | Map1b | microtubule-associated protein 1B | 71.7% | 29.1% | 1.6 | 1.4E-80 | 7.0 | -1.7 | 1.2E-28 | 61.6 | 8.7 | 18.5 | 2.6 |

| ENDOTHELIAL CELLS | | | | | | | | | | | | | |
|---|---|---|---|---|---|---|---|---|---|---|---|---|---|
| | | | Expression in single-cell (Vanlandewijck et al., 2018) | | | | | | Cortical MV RNA seq analysis: P15 vs P5 | | | | |
| | | | | | | | | | P15 vs P5 | | RPKM MV / RPKM EC | | |
| Cluster | Gene | Name | % of cells in the cluster | % of cells exp in clusters | logFC | Adjusted P-value | RPKM EC | log2FoldChange | Adjusted P-value | P5 | P5/EC | P15 | P15/EC |
| EC | Abcb1a | ATP-binding ca | 98.6% | 8.0% | 2.4 | 0.0E+00 | 11.2 | 1.2 | 7.5E-17 | 39.0 | 3.5 | 91.3 | 8.2 |
| EC | Abcg2 | ATP-binding ca | 94.1% | 22.6% | 2.1 | 0.0E+00 | 12.1 | 0.4 | 5.5E-03 | 77.6 | 6.4 | 104.0 | 8.6 |
| EC | Acer2 | alkaline cerami | 75.7% | 6.1% | 1.9 | 0.0E+00 | 6.4 | 3.0 | 3.4E-62 | 2.5 | 0.4 | 20.4 | 3.2 |
| EC | Acvrl1 | activin A recept | 92.9% | 11.7% | 2.0 | 0.0E+00 | 9.6 | 0.8 | 9.1E-10 | 65.4 | 6.8 | 119.0 | 12.3 |
| EC | Afap1l1 | actin filament a | 76.5% | 6.6% | 1.8 | 0.0E+00 | 5.7 | 0.4 | 6.7E-04 | 98.0 | 17.2 | 130.8 | 23.0 |
| EC | Alas1 | aminolevulinic | 80.0% | 18.0% | 1.5 | 0.0E+00 | 5.5 | 0.5 | 9.9E-05 | 109.2 | 19.7 | 156.6 | 28.3 |
| EC | Anxa3 | annexin A3 | 85.3% | 16.8% | 1.8 | 0.0E+00 | 9.0 | 0.4 | 8.3E-04 | 150.5 | 16.7 | 206.9 | 22.9 |
| EC | Arl4a | ADP-ribosylati | 86.5% | 19.0% | 1.7 | 0.0E+00 | 8.4 | 0.2 | 1.1E-01 | 24.8 | 3.0 | 29.1 | 3.5 |
| EC | AW112010 | expressed seque | 62.6% | 2.7% | 1.5 | 0.0E+00 | 3.8 | 1.4 | 4.8E-06 | 2.7 | 0.7 | 7.0 | 1.8 |
| EC | Cd34 | CD34 antigen | 95.2% | 13.8% | 2.4 | 0.0E+00 | 16.4 | -0.1 | 5.6E-01 | 349.6 | 21.4 | 334.3 | 20.4 |
| EC | Cd93 | CD93 antigen | 79.1% | 5.7% | 1.7 | 0.0E+00 | 5.1 | -0.5 | 1.5E-02 | 90.3 | 17.7 | 64.5 | 12.6 |
| EC | Cdh5 | cadherin 5 | 94.6% | 5.1% | 1.7 | 0.0E+00 | 4.6 | 0.0 | 8.0E-01 | 42.9 | 9.4 | 44.7 | 9.8 |
| EC | Cldn5 | claudin 5 | 99.9% | 15.7% | 5.4 | 0.0E+00 | 262.1 | 0.1 | 4.2E-01 | 1064.1 | 4.1 | 1174.2 | 4.5 |
| EC | Ctla2a | cytotoxic T lym | 79.8% | 3.5% | 2.1 | 0.0E+00 | 7.4 | 0.3 | 2.5E-02 | 229.6 | 31.2 | 281.0 | 38.2 |
| EC | Egfl7 | EGF-like doma | 99.6% | 7.8% | 2.9 | 0.0E+00 | 18.3 | 0.2 | 2.1E-01 | 172.6 | 9.4 | 195.7 | 10.7 |
| EC | Eltd1 | adhesion G prot | 98.6% | 7.1% | 3.0 | 0.0E+00 | 21.6 | -0.2 | 1.7E-03 | 209.0 | 9.7 | 278.3 | 12.9 |
| EC | Eogt | EGF domain-sp | 85.9% | 10.1% | 2.0 | 0.0E+00 | 8.5 | 0.7 | 4.9E-10 | 134.4 | 15.9 | 226.3 | 26.7 |
| EC | Fn1 | fibronectin 1 | 87.9% | 10.8% | 2.1 | 0.0E+00 | 10.5 | 0.4 | 3.4E-02 | 256.3 | 24.4 | 343.7 | 32.7 |
| EC | Gbp3 | guanylate bindi | 66.8% | 5.8% | 1.7 | 0.0E+00 | 5.0 | 0.0 | 9.7E-01 | 20.0 | 4.0 | 20.1 | 4.0 |
| EC | Gpr116 | adhesion G prot | 98.4% | 23.3% | 2.3 | 0.0E+00 | 16.5 | 0.4 | 3.6E-07 | 102.8 | 6.2 | 162.4 | 9.8 |
| EC | Grap | GRB2-related a | 81.2% | 12.0% | 1.6 | 0.0E+00 | 6.2 | 0.2 | 9.4E-02 | 131.5 | 21.0 | 154.5 | 24.7 |
| EC | Gsta4 | glutathione S-tr | 77.5% | 13.1% | 1.6 | 0.0E+00 | 5.5 | 0.8 | 1.2E-06 | 24.3 | 4.4 | 41.4 | 7.6 |
| EC | Hmgcs2 | 3-hydroxy-3-me | 73.2% | 6.0% | 2.0 | 0.0E+00 | 7.6 | -0.1 | 5.0E-01 | 281.0 | 37.2 | 266.1 | 35.2 |
| EC | Icam2 | intercellular adh | 81.9% | 2.9% | 1.7 | 0.0E+00 | 4.7 | 0.5 | 6.4E-04 | 126.5 | 26.8 | 182.3 | 38.7 |
| EC | Id1 | inhibitor of DN | 86.6% | 17.5% | 1.7 | 0.0E+00 | 6.3 | 0.9 | 1.9E-07 | 86.5 | 13.7 | 163.0 | 25.8 |
| EC | Ivns1abp | influenza virus | 70.0% | 18.6% | 1.9 | 2.3E-214 | 9.2 | -0.4 | 1.1E-02 | 112.0 | 12.2 | 87.4 | 9.5 |
| EC | Jam2 | junction adhesi | 86.6% | 19.2% | 1.5 | 0.0E+00 | 8.5 | 0.2 | 2.1E-01 | 74.2 | 8.8 | 84.1 | 9.9 |

| | | | | | | | | | | | | |
|---|---|---|---|---|---|---|---|---|---|---|---|---|
| EC | Kank3 | KN motif and a | 91.2% | 8.3% | 1.9 | 0.0E+00 | 6.9 | 0.2 | 7.0E-02 | 113.9 | 16.5 | 134.8 | 19.6 |
| EC | Kdr | kinase insert do | 83.6% | 4.1% | 2.0 | 0.0E+00 | 6.4 | -0.1 | 5.0E-01 | 305.5 | 47.9 | 288.9 | 45.3 |
| EC | Kitl | kit ligand | 85.5% | 15.2% | 1.7 | 0.0E+00 | 6.9 | 1.2 | 3.1E-16 | 38.6 | 5.6 | 88.5 | 12.8 |
| EC | Lmo2 | LIM domain on | 82.7% | 7.9% | 1.6 | 0.0E+00 | 5.2 | -0.2 | 1.0E-01 | 182.5 | 34.8 | 160.3 | 30.6 |
| EC | Lsr | lipolysis stimula | 89.9% | 6.1% | 2.2 | 0.0E+00 | 8.2 | 0.3 | 1.5E-02 | 178.3 | 21.8 | 222.1 | 27.1 |
| EC | Ly6c1 | lymphocyte ant | 99.9% | 18.0% | 3.7 | 0.0E+00 | 48.5 | 1.9 | 6.3E-43 | 163.9 | 3.4 | 599.6 | 12.4 |
| EC | Ly6e | lymphocyte ant | 99.7% | 24.4% | 3.4 | 0.0E+00 | 50.4 | 1.7 | 5.2E-26 | 61.8 | 1.2 | 198.3 | 3.9 |
| EC | Mfsd2a | major facilitato | 80.4% | 14.0% | 2.0 | 0.0E+00 | 10.6 | -0.3 | 3.0E-02 | 562.4 | 52.8 | 474.7 | 44.6 |
| EC | Mpzl1 | myelin protein z | 88.7% | 16.5% | 2.0 | 0.0E+00 | 10.8 | -0.1 | 2.8E-01 | 235.7 | 21.8 | 215.0 | 19.9 |
| EC | Ocln | occludin | 82.9% | 5.8% | 1.9 | 0.0E+00 | 6.5 | 0.8 | 1.4E-09 | 85.5 | 13.2 | 151.7 | 23.4 |
| EC | Palmd | palmdelphin | 86.8% | 12.8% | 1.7 | 0.0E+00 | 6.2 | 0.1 | 4.6E-01 | 83.4 | 13.5 | 91.3 | 14.8 |
| EC | Paqr5 | progestin and a | 70.0% | 2.7% | 1.6 | 0.0E+00 | 3.8 | 3.0 | 1.1E-58 | 3.3 | 0.8 | 26.0 | 6.8 |
| EC | Pecam1 | platelet/endothe | 95.8% | 6.3% | 2.6 | 0.0E+00 | 13.5 | 0.1 | 3.3E-01 | 182.2 | 13.5 | 200.0 | 14.8 |
| EC | Plk2 | polo-like kinase | 74.0% | 9.3% | 2.0 | 0.0E+00 | 8.0 | 0.5 | 2.8E-04 | 85.3 | 10.6 | 121.8 | 15.2 |
| EC | Podxl | podocalyxin-lik | 97.8% | 6.9% | 2.4 | 0.0E+00 | 11.2 | 0.6 | 1.8E-04 | 63.2 | 5.6 | 94.5 | 8.4 |
| EC | Prkch | protein kinase C | 71.8% | 2.0% | 1.5 | 0.0E+00 | 3.6 | 0.3 | 4.6E-02 | 29.7 | 8.2 | 36.3 | 10.1 |
| EC | Prom1 | prominin 1 | 95.0% | 7.8% | 2.8 | 0.0E+00 | 18.3 | 1.5 | 1.5E-30 | 62.3 | 3.4 | 175.5 | 9.6 |
| EC | Ptprb | protein tyrosine | 98.2% | 8.3% | 2.7 | 0.0E+00 | 15.3 | 0.7 | 4.9E-07 | 68.8 | 4.5 | 115.2 | 7.5 |
| EC | Ramp2 | receptor (calcito | 97.8% | 12.4% | 2.5 | 0.0E+00 | 14.8 | 0.1 | 4.7E-01 | 498.3 | 33.8 | 536.2 | 36.3 |
| EC | Robo4 | roundabout guid | 87.0% | 3.5% | 2.0 | 0.0E+00 | 6.5 | 0.2 | 1.9E-01 | 69.6 | 10.7 | 79.4 | 12.2 |
| EC | Sdpr | serum deprivati | 76.6% | 5.6% | 1.7 | 0.0E+00 | 4.9 | 0.9 | 9.5E-07 | 23.6 | 4.9 | 45.4 | 9.4 |
| EC | Sema3c | sema domain, in | 72.0% | 4.5% | 1.7 | 0.0E+00 | 4.6 | 3.0 | 6.4E-79 | 9.3 | 2.0 | 73.4 | 16.1 |
| EC | Slc16a1 | solute carrier fa | 81.1% | 17.8% | 2.1 | 0.0E+00 | 11.6 | 0.8 | 2.9E-07 | 570.9 | 49.3 | 973.1 | 84.1 |
| EC | Slc16a4 | solute carrier fa | 60.4% | 4.4% | 1.6 | 1.0E-282 | 4.1 | 0.6 | 1.6E-03 | 31.2 | 7.6 | 46.2 | 11.3 |
| EC | Slc38a3 | solute carrier fa | 87.9% | 17.2% | 1.5 | 0.0E+00 | 11.7 | 0.1 | 6.7E-01 | 248.7 | 21.2 | 261.5 | 22.3 |
| EC | Slc39a10 | solute carrier fa | 91.6% | 18.2% | 2.0 | 0.0E+00 | 9.6 | -0.1 | 6.9E-01 | 116.8 | 12.1 | 112.7 | 11.7 |
| EC | Slc39a8 | solute carrier fa | 65.8% | 3.8% | 1.6 | 0.0E+00 | 4.5 | -0.8 | 4.1E-09 | 204.9 | 45.8 | 116.1 | 26.0 |
| EC | Slc40a1 | solute carrier fa | 85.6% | 6.4% | 2.3 | 0.0E+00 | 11.5 | 0.0 | 7.5E-01 | 391.2 | 33.9 | 384.4 | 33.4 |
| EC | Slc7a1 | solute carrier fa | 86.3% | 16.9% | 1.7 | 0.0E+00 | 7.0 | -0.8 | 1.7E-09 | 257.9 | 36.7 | 154.5 | 22.0 |
| EC | Slc7a5 | solute carrier fa | 91.6% | 13.4% | 2.5 | 0.0E+00 | 14.7 | -0.4 | 5.0E-03 | 900.2 | 61.3 | 685.7 | 46.7 |
| EC | Slco1a4 | solute carrier or | 99.5% | 16.9% | 3.7 | 0.0E+00 | 47.5 | 2.4 | 4.5E-66 | 65.5 | 1.4 | 351.6 | 7.4 |
| EC | Slco1c1 | solute carrier or | 97.6% | 17.1% | 2.9 | 0.0E+00 | 25.9 | 0.1 | 5.5E-01 | 801.3 | 31.0 | 858.9 | 33.2 |
| EC | Slco2b1 | solute carrier or | 80.2% | 5.2% | 2.0 | 0.0E+00 | 7.3 | 0.0 | 8.7E-01 | 93.2 | 12.7 | 92.7 | 12.7 |

| | | | | | | | | | | | | | |
|---|---|---|---|---|---|---|---|---|---|---|---|---|---|
| EC | Sox18 | SRY (sex deter | 89.8% | 3.0% | 2.1 | 0.0E+00 | 7.1 | 0.1 | 4.9E-01 | 84.0 | 11.8 | 90.3 | 12.6 |
| EC | Spock2 | sparc/osteonect | 99.4% | 14.9% | 2.6 | 0.0E+00 | 14.1 | -0.1 | 6.5E-01 | 367.4 | 26.2 | 350.0 | 24.9 |
| EC | Srgn | serglycin | 96.4% | 10.8% | 2.2 | 0.0E+00 | 9.8 | 0.3 | 6.7E-02 | 56.1 | 5.8 | 69.5 | 7.1 |
| EC | St3gal6 | ST3 beta-galact | 86.8% | 18.1% | 1.7 | 0.0E+00 | 7.0 | 0.8 | 6.9E-08 | 83.6 | 12.0 | 145.4 | 20.9 |
| EC | Tdrp | testis developm | 84.4% | 4.1% | 2.0 | 0.0E+00 | 7.2 | -0.1 | 5.9E-01 | 4.3 | 0.6 | 4.0 | 0.6 |
| EC | Tek | endothelial-spec | 96.6% | 8.2% | 2.7 | 0.0E+00 | 15.8 | 0.4 | 8.7E-03 | 114.5 | 7.2 | 149.8 | 9.5 |
| EC | Tfrc | transferrin recep | 74.9% | 18.3% | 1.9 | 1.0E-263 | 9.7 | 0.0 | 7.5E-01 | 308.8 | 31.9 | 320.0 | 33.1 |
| EC | Tie1 | tyrosine kinase | 94.7% | 6.3% | 2.1 | 0.0E+00 | 7.8 | -0.1 | 6.4E-01 | 144.1 | 18.5 | 139.3 | 17.9 |
| EC | Tspan13 | tetraspanin 13 | 97.8% | 11.1% | 2.7 | 0.0E+00 | 16.4 | 0.0 | 9.3E-01 | 224.4 | 13.7 | 224.1 | 13.6 |
| EC | Ucp2 | uncoupling prot | 91.3% | 22.2% | 1.8 | 0.0E+00 | 9.8 | 0.5 | 3.4E-04 | 166.0 | 17.0 | 237.5 | 24.3 |
| EC | Ushbp1 | Usher syndrome | 81.7% | 2.6% | 1.6 | 0.0E+00 | 3.9 | 0.0 | 7.8E-01 | 125.8 | 32.4 | 130.1 | 33.5 |
| EC | Vwa1 | von Willebrand | 98.4% | 18.1% | 2.6 | 0.0E+00 | 22.9 | 0.1 | 3.4E-01 | 497.0 | 21.7 | 543.7 | 23.7 |
| EC | Wfdc1 | WAP four-disul | 92.9% | 16.7% | 1.8 | 0.0E+00 | 7.9 | 1.5 | 1.2E-26 | 68.5 | 8.6 | 195.6 | 24.6 |

Supplementary Table S4

### FIGURE 2

| | | P5 | | | | | | | | |
|---|---|---|---|---|---|---|---|---|---|---|
| | | Number of values | Median | Mean | Std. Deviation | Kruskal-Wallis test | Dunn's multiple comparisons test (Adjusted P Value) | | | |
| | | | | | | | EC | PC | FB | VSMC |
| Fig. 2A | EC | 73 | 12.73 | 16.48 | 13.86 | <0,0001 | | | | |
| | PC | 14 | 2.215 | 5.299 | 10.62 | | 0.0035 | | | |
| | FB | 38 | 1.19 | 3.378 | 8.622 | | <0,0001 | >0,9999 | | |
| | VSMC | 21 | 0.69 | 3.306 | 8.652 | | <0,0001 | 0.8036 | >0,9999 | |

| | | P15 | | | | | | | | |
|---|---|---|---|---|---|---|---|---|---|---|
| | | Number of values | Median | Mean | Std. Deviation | Kruskal-Wallis test | Dunn's multiple comparisons test (Adjusted P Value) | | | |
| | | | | | | | EC | PC | FB | VSMC |
| Fig. 2B | EC | 73 | 14.85 | 19.22 | 13.66 | <0,0001 | | | | |
| | PC | 14 | 1.985 | 4.575 | 6.568 | | 0.0001 | | | |
| | FB | 38 | 1.675 | 6.015 | 19.43 | | <0,0001 | >0,9999 | | |
| | VSMC | 21 | 1.46 | 3.702 | 8.314 | | <0,0001 | >0,9999 | >0,9999 | |

| | | | P5 | | | P15 | | | |
|---|---|---|---|---|---|---|---|---|---|
| | | Number of values | Median | Mean | Std. Deviation | Median | Mean | Std. Deviation | Wilcoxon matched-pairs signed rank test |
| Fig. 2C | EC | 73 | 12.73 | 16.48 | 13.86 | 14.85 | 19.22 | 13.66 | <0,0001 |
| | PC | 14 | 2.215 | 5.299 | 10.62 | 1.985 | 4.575 | 6.568 | 0.5016 |
| | FB | 38 | 1.19 | 3.378 | 8.622 | 1.675 | 6.015 | 19.43 | 0.0026 |
| | VSMC | 21 | 0.69 | 3.306 | 8.652 | 1.46 | 3.702 | 8.314 | 0.0014 |

#### FIGURE 3

##### Fig. 3A (FB)

| | | | | | | | Kruskal-Wallis test | Mann Whitney test (one-tailed) | |
|---|---|---|---|---|---|---|---|---|---|
| Col6a1 | | Number of values | Median | Mean | Std. Deviation | Std. Error of Mean | | P5 | P10 |
| | P5 | 4 | 0.6 | 0.58 | 0.1036 | 0.0518 | 0.0012 | | |
| | P10 | 4 | 1.765 | 1.873 | 0.3989 | 0.1994 | | 0.0143 | |
| | P15 | 4 | 1.245 | 1.36 | 0.3685 | 0.1843 | | 0.0143 | 0.0571 |
| | P5 - normalized | 4 | 1.03 | 1.0 | 0.1796 | 0.08981 | | | |
| | P10 - normalized | 4 | 3.04 | 3.223 | 0.6816 | 0.3408 | | | |
| | P15 - normalized | 4 | 2.145 | 2.343 | 0.6356 | 0.3178 | | | |
| Col1a1 | | Number of values | Median | Mean | Std. Deviation | Std. Error of Mean | | P5 | P10 |
| | P5 | 4 | 0.4776 | 0.491 | 0.1737 | 0.08685 | 0.4356 | | |
| | P10 | 5 | 0.5399 | 0.6529 | 0.2267 | 0.1014 | | 0.2063 | |
| | P15 | 4 | 0.4136 | 0.5144 | 0.2909 | 0.1455 | | 0.4429 | 0.1429 |
| | P5 - normalized | 4 | 0.975 | 1.003 | 0.3539 | 0.1769 | | | |
| | P10 - normalized | 5 | 1.1 | 1.33 | 0.4629 | 0.207 | | | |
| | P15 - normalized | 4 | 0.845 | 1.05 | 0.5923 | 0.2962 | | | |
| Lama1 | | Number of values | Median | Mean | Std. Deviation | Std. Error of Mean | | P5 | P10 |
| | P5 | 3 | 1.04 | 1.13 | 0.2287 | 0.132 | 0.0216 | | |
| | P10 | 4 | 2.39 | 2.54 | 1.423 | 0.7117 | | 0.0571 | |
| | P15 | 4 | 4.99 | 4.898 | 1.807 | 0.9034 | | 0.0286 | 0.1 |
| | P5 - normalized | 3 | 0.92 | 1.0 | 0.2022 | 0.1168 | | | |
| | P10 - normalized | 4 | 2.11 | 2.243 | 1.258 | 0.6289 | | | |
| | P15 - normalized | 4 | 4.405 | 4.325 | 1.595 | 0.7974 | | | |
| Lum | | Number of values | Median | Mean | Std. Deviation | Std. Error of Mean | | P5 | P10 |
| | P5 | 3 | 0.86 | 0.8833 | 0.1464 | 0.08452 | 0.0234 | | |
| | P10 | 4 | 1.915 | 2.513 | 1.647 | 0.8233 | | 0.0286 | |
| | P15 | 4 | 2.93 | 3.448 | 1.886 | 0.9429 | | 0.0286 | 0.2429 |
| | P5 - normalized | 3 | 0.97 | 1.0 | 0.167 | 0.09644 | | | |
| | P10 - normalized | 4 | 2.17 | 2.848 | 1.866 | 0.933 | | | |
| | P15 - normalized | 4 | 3.325 | 3.91 | 2.137 | 1.069 | | | |

| Fig. 3B (PC) | | | | | | | | | |
|---|---|---|---|---|---|---|---|---|---|
| | | | | | | | Kruskal-Wallis test | Mann Whitney test (one-tailed) | |
| Abcc9 | | Number of values | Median | Mean | Std. Deviation | Std. Error of Mean | | P5 | P10 |
| | P5 | 4 | 2.999 | 3.303 | 1.768 | 0.8842 | 0.5901 | | |
| | P10 | 4 | 3.512 | 4.213 | 2.187 | 1.094 | | 0.3 | |
| | P15 | 4 | 4.653 | 4.424 | 1.727 | 0.8636 | | 0.1571 | 0.4286 |
| | P5 - normalized | 4 | 0.95 | 1.025 | 0.5058 | 0.2529 | | | |
| | P10 - normalized | 4 | 1.05 | 1.25 | 0.6557 | 0.3279 | | | |
| | P15 - normalized | 4 | 1.4 | 1.325 | 0.5188 | 0.2594 | | | |
| Kcnj8 | | Number of values | Median | Mean | Std. Deviation | Std. Error of Mean | | P5 | P10 |
| | P5 | 4 | 1.35 | 1.226 | 0.5286 | 0.2643 | 0.8263 | | |
| | P10 | 4 | 1.631 | 1.754 | 0.9764 | 0.4882 | | 0.2571 | |
| | P15 | 4 | 1.506 | 1.834 | 1.093 | 0.5465 | | 0.3714 | 0.4857 |
| | P5 - normalized | 4 | 1.1 | 1.00 | 0.4546 | 0.2273 | | | |
| | P10 - normalized | 4 | 1.35 | 1.45 | 0.7937 | 0.3969 | | | |
| | P15 - normalized | 4 | 1.25 | 1.5 | 0.9055 | 0.4528 | | | |

| Fig. 3C (VSMC) | | | | | | | | | |
|---|---|---|---|---|---|---|---|---|---|
| | | | | | | | Kruskal-Wallis test | Mann Whitney test (one-tailed) | |
| Acta2 | | Number of values | Median | Mean | Std. Deviation | Std. Error of Mean | | P5 | P10 |
| | P5 | 3 | 9.85 | 9.813 | 0.4062 | 0.2345 | 0.0038 | | |
| | P10 | 4 | 17.07 | 17.86 | 4.453 | 2.226 | | 0.0286 | |
| | P15 | 4 | 26.44 | 25.49 | 5.177 | 2.589 | | 0.0286 | 0.0571 |
| | P5 - normalized | 3 | 1.00 | 1.00 | 0.04 | 0.02309 | | | |
| | P10 - normalized | 4 | 1.74 | 1.82 | 0.4517 | 0.2258 | | | |
| | P15 - normalized | 4 | 2.695 | 2.598 | 0.5289 | 0.2644 | | | |
| Myh11 | | Number of values | Median | Mean | Std. Deviation | Std. Error of Mean | | P5 | P10 |
| | P5 | 3 | 1.16 | 1.01 | 0.4732 | 0.2732 | 0.0184 | | |
| | P10 | 4 | 3.755 | 3.738 | 0.6641 | 0.3321 | | 0.0286 | |
| | P15 | 4 | 6.005 | 5.658 | 2.067 | 1.033 | | 0.0286 | 0.1714 |
| | P5 - normalized | 3 | 1.15 | 1.00 | 0.4732 | 0.2732 | | | |
| | P10 - normalized | 4 | 3.715 | 3.698 | 0.6561 | 0.3281 | | | |
| | P15 - normalized | 4 | 5.94 | 5.598 | 2.045 | 1.023 | | | |
| Tagln | | Number of values | Median | Mean | Std. Deviation | Std. Error of Mean | | P5 | P10 |
| | P5 | 3 | 1.14 | 0.9367 | 0.3522 | 0.2033 | 0.0014 | | |
| | P10 | 4 | 3.295 | 3.395 | 0.7774 | 0.3887 | | 0.0286 | |
| | P15 | 4 | 6.415 | 5.858 | 1.298 | 0.6489 | | 0.0286 | 0.0286 |
| | P5 - normalized | 3 | 1.22 | 1.003 | 0.3753 | 0.2167 | | | |
| | P10 - normalized | 4 | 3.515 | 3.623 | 0.8292 | 0.4146 | | | |
| | P15 - normalized | 4 | 6.85 | 6.255 | 1.384 | 0.692 | | | |
| Atp1b1 | | Number of values | Median | Mean | Std. Deviation | Std. Error of Mean | | | |
| | P5 | 4 | 0.3495 | 0.3819 | 0.08162 | 0.04081 | 0.8891 | | |
| | P10 | 5 | 0.3686 | 0.4085 | 0.1265 | 0.05657 | | 0.4524 | |
| | P15 | 4 | 0.3479 | 0.3655 | 0.1616 | 0.08081 | | 0.4429 | 0.3651 |
| | P5 - normalized | 4 | 0.915 | 1.00 | 0.2159 | 0.1079 | | | |
| | P10 - normalized | 5 | 0.97 | 1.07 | 0.3301 | 0.1476 | | | |
| | P15 - normalized | 4 | 0.915 | 0.8825 | 0.5428 | 0.2714 | | | |

**Fig. 3D (EC)**

| Gene | | Number of values | Median | Mean | Std. Deviation | Std. Error of Mean | Kruskal-Wallis test | Mann Whitney test (one-tailed) P5 | Mann Whitney test (one-tailed) P10 |
|---|---|---|---|---|---|---|---|---|---|
| Ly6c1 | P5 | 5 | 2.17 | 2.954 | 2.825 | 1.263 | 0.0002 | | |
| | P10 | 4 | 13.23 | 13.61 | 2.081 | 1.04 | | 0.0079 | |
| | P15 | 3 | 50.1 | 53.00 | 16.27 | 9.394 | | 0.0179 | 0.0286 |
| | P5 - normalized | 5 | 0.73 | 0.998 | 0.9571 | 0.428 | | | |
| | P10 - normalized | 4 | 4.475 | 4.608 | 0.7049 | 0.3525 | | | |
| | P15 - normalized | 3 | 16.96 | 17.94 | 5.511 | 3.182 | | | |
| Acer2 | | Number of values | Median | Mean | Std. Deviation | Std. Error of Mean | | P5 | P10 |
| | P5 | 3 | 0.11 | 0.1133 | 0.005774 | 0.003333 | 0.0016 | | |
| | P10 | 4 | 0.2364 | 0.2376 | 0.0819 | 0.04095 | | 0.0286 | |
| | P15 | 4 | 0.5653 | 0.6582 | 0.3968 | 0.1984 | | 0.0286 | 0.0286 |
| | P5 - normalized | 3 | 0.99 | 1.00 | 0.03606 | 0.02082 | | | |
| | P10 - normalized | 4 | 2.085 | 2.07 | 0.7117 | 0.3558 | | | |
| | P15 - normalized | 4 | 4.985 | 5.733 | 3.456 | 1.728 | | | |
| Slco1a4 | | Number of values | Median | Mean | Std. Deviation | Std. Error of Mean | | P5 | P10 |
| | P5 | 5 | 1.699 | 1.62 | 0.53 | 0.237 | <0.0001 | | |
| | P10 | 4 | 4.936 | 4.895 | 0.7761 | 0.3881 | | 0.0079 | |
| | P15 | 4 | 17.88 | 18.46 | 7.22 | 3.61 | | 0.0079 | 0.0143 |
| | P5 - normalized | 5 | 1.05 | 1 | 0.325 | 0.1453 | | | |
| | P10 - normalized | 4 | 3.045 | 3.02 | 0.4767 | 0.2384 | | | |
| | P15 - normalized | 4 | 11.04 | 11.4 | 4.456 | 2.228 | | | |
| Abcb1a | | Number of values | Median | Mean | Std. Deviation | Std. Error of Mean | | P5 | P10 |
| | P5 | 5 | 0.86 | 0.952 | 0.8036 | 0.3594 | <0.0001 | | |
| | P10 | 4 | 7.39 | 7.19 | 0.7954 | 0.3977 | | 0.0079 | |
| | P15 | 4 | 13.54 | 13.37 | 2.315 | 1.158 | | 0.0079 | 0.0143 |
| | P5 - normalized | 5 | 0.9 | 1 | 0.8466 | 0.3786 | | | |
| | P10 - normalized | 4 | 7.755 | 7.548 | 0.8371 | 0.4185 | | | |
| | P15 - normalized | 4 | 14.22 | 14.04 | 2.43 | 1.215 | | | |

### FIGURE 4

#### Western Blot

| | | | | | | | Kruskal-Wallis test | Mann Whitney test (one-tailed) | | |
|---|---|---|---|---|---|---|---|---|---|---|
| | | Number of values | Median | Mean | Std. Deviation | Std. Error of Mean | | P5 | P15 | P30 |
| Claudin-5 | P5 | 4 | 1.0 | 1.0 | 0.3 | 0.1 | <0.0001 | | | |
| | P15 | 4 | 1.6 | 1.6 | 0.4 | 0.2 | | 0.04 | | |
| | P30 | 4 | 3.0 | 2.9 | 0.6 | 0.3 | | 0.01 | 0.01 | |
| | P60 | 3 | 6.0 | 6.1 | 0.8 | 0.5 | | 0.03 | 0.03 | 0.03 |

| | | | | | | | Kruskal-Wallis test | Mann Whitney test (one-tailed) | | |
|---|---|---|---|---|---|---|---|---|---|---|
| | | Number of values | Median | Mean | Std. Deviation | Std. Error of Mean | | P5 | P15 | P30 |
| P-gP | P5 | 4 | 0.0 | 0.0 | 0.0 | 0.0 | <0.0001 | | | |
| | P15 | 4 | 1.0 | 1.0 | 0.1 | 0.1 | | 0.01 | | |
| | P30 | 4 | 11.4 | 10.4 | 2.3 | 1.1 | | 0.01 | 0.01 | |
| | P60 | 4 | 17.0 | 18.1 | 4.9 | 2.5 | | 0.01 | 0.01 | 0.01 |

#### BBB in vivo

| | | | | | | | Kruskal-Wallis test | Mann Whitney test (one-tailed) | |
|---|---|---|---|---|---|---|---|---|---|
| | | Number of values | Median | Mean | Std. Deviation | Std. Error of Mean | | P5 | P15 |
| 14C-Sucrose | P5 | 8 | 23.5 | 27.7 | 10.6 | 3.7 | 0.7 | | |
| | P15 | 7 | 25.6 | 27.2 | 7.6 | 2.9 | | 0.4 | |
| | P30 | 6 | 29.2 | 27.9 | 8.7 | 3.5 | | 0.3 | 0.2 |
| 3H-Vérapamil | P15 | 7 | 108.7 | 110.8 | 22.8 | 8.6 | | | |
| | P30 | 6 | 79.1 | 75.7 | 9.3 | 3.8 | | | 0.01 |
| 3H-Rosuvastatin | P15 | 7 | 11.7 | 11.4 | 4.8 | 1.8 | | | |
| | P30 | 6 | 1.5 | 1.7 | 1.5 | 0.6 | | | 0.001 |

**FIGURE 5**

<table>
<tr><th colspan="9">P-gP IHC on human brain slices</th></tr>
<tr><td colspan="6"></td><td>Kruskal-Wallis test</td><td>Kruskal-Wallis test</td><td>Mann Whitney test (one-tailed)</td></tr>
<tr><td></td><td>Number of values</td><td>Median</td><td>Mean</td><td>Std. Deviation</td><td>Std. Error of Mean</td><td></td><td></td><td>prenatal</td></tr>
<tr><td>prenatal</td><td>6</td><td>0.30</td><td>0.30</td><td>0.03</td><td>0.01</td><td rowspan="4">0.004</td><td></td><td></td></tr>
<tr><td>0-2 years</td><td>5</td><td>0.42</td><td>0.42</td><td>0.03</td><td>0.01</td><td rowspan="3">0.11</td><td>0.004</td></tr>
<tr><td>2-5 years</td><td>4</td><td>0.36</td><td>0.37</td><td>0.04</td><td>0.02</td><td>0.02</td></tr>
<tr><td>>10 years</td><td>5</td><td>0.43</td><td>0.42</td><td>0.04</td><td>0.02</td><td>0.004</td></tr>
</table>

#### FIGURE 6

##### Myh11 FISH detection in MVs

| Proportion of vessel | | Number of values | Median | Mean | Std. Deviation | Std. Error of Mean | Chi² | Mann Whitney test (one-tailed) |
|---|---|---|---|---|---|---|---|---|
| 0-10 | P5 | 3 | 0.1282 | 0.2247 | 0.2025 | 0.1169 | <0.0001 | 0.05 |
| | P15 | 3 | 0.7909 | 0.7985 | 0.02172 | 0.01254 | | |
| 10-100 | P5 | 3 | 0.8673 | 0.757 | 0.1949 | 0.1125 | | 0.05 |
| | P15 | 3 | 0.1149 | 0.11 | 0.008627 | 0.004981 | | |
| >100 | P5 | 3 | 0.01064 | 0.0183 | 0.0231 | 0.01333 | | 0.05 |
| | P15 | 3 | 0.1034 | 0.1053 | 0.003258 | 0.001881 | | |

| | | | | | | | Kruskal-Wallis test | Mann Whitney test (two-tailed) | |
|---|---|---|---|---|---|---|---|---|---|
| | | Number of values | Median | Mean | Std. Deviation | Std. Error of Mean | | 0-10 | 10-100 |
| Diameters | 0-10 | 247 | 5.31 | 5.961 | 3.084 | 0.1962 | <0.0001 | | |
| | 10-100 | 39 | 8.92 | 9.231 | 3.084 | 0.4938 | | <0.0001 | |
| | >100 | 27 | 12.77 | 13.5 | 5.565 | 1.071 | | <0.0001 | 0.0007 |

##### Western Blot in MVs

| | | | | | | | Kruskal-Wallis test | Mann Whitney test (one-tailed) | | |
|---|---|---|---|---|---|---|---|---|---|---|
| Myh11 | | Number of values | Median | Mean | Std. Deviation | Std. Error of Mean | | P5 | P15 | P30 |
| | P5 | 4 | 0.88 | 1.00 | 0.44 | 0.22 | <0.0001 | | | |
| | P15 | 4 | 22.85 | 23.66 | 6.00 | 3.00 | | 0.01 | | |
| | P30 | 3 | 65.41 | 66.40 | 6.61 | 3.81 | | 0.03 | 0.03 | |
| | P60 | 3 | 47.44 | 47.92 | 3.83 | 2.21 | | 0.03 | 0.03 | 0.05 |
| SMA | | | | | | | Kruskal-Wallis test | Mann Whitney test (one-tailed) | | |
| | | Number of values | Median | Mean | Std. Deviation | Std. Error of Mean | | P5 | P15 | P30 |
| | P5 | 4 | 1.09 | 1.00 | 0.22 | 0.11 | 0.00 | | | |
| | P15 | 4 | 8.67 | 10.25 | 5.15 | 2.58 | | 0.01 | | |
| | P30 | 4 | 17.20 | 16.48 | 4.32 | 2.16 | | 0.01 | 0.06 | |
| | P60 | 3 | 19.61 | 20.06 | 6.00 | 3.47 | | 0.03 | 0.06 | 0.31 |

#### Contractility

| | | | | | | | Kruskal-Wallis test | Mann Whitney test (one-tailed) | |
|---|---|---|---|---|---|---|---|---|---|
| Max amplitude | | Number of values | Median | Mean | Std. Deviation | Std. Error of Mean | | P5 | P15 |
| | P5 | 13 | 4.551 | 6.505 | 5.03 | 1.395 | 0.0095 | | |
| | P15 | 9 | 13.65 | 14.15 | 7.823 | 2.608 | | 0.017 | |
| | P60 | 8 | 14.58 | 18.07 | 13.67 | 4.833 | | 0.006 | 0.8148 |

| | | | | | | | Kruskal-Wallis test | Mann Whitney test (one-tailed) | |
|---|---|---|---|---|---|---|---|---|---|
| Slope | | Number of values | Median | Mean | Std. Deviation | Std. Error of Mean | | P5 | P15 |
| | P5 | 13 | -0.7172 | -0.9008 | 1.061 | 0.2941 | 0.0066 | | |
| | P15 | 9 | -1.054 | -1.469 | 1.029 | 0.3431 | | 0.1858 | |
| | P60 | 8 | -3.395 | -3.219 | 1.414 | 0.4999 | | 0.0045 | 0.0111 |

#### qPCR Tbxa2R

| | | | | | | | Kruskal-Wallis test | Mann Whitney test (one-tailed) | |
|---|---|---|---|---|---|---|---|---|---|
| Tbxa2r | | Number of values | Median | Mean | Std. Deviation | Std. Error of Mean | | P5 | P15 |
| | P5 | 3 | 3.445 | 3.453 | 1.278 | 0.738 | 0.5535 | | |
| | P15 | 4 | 6.462 | 7.602 | 4.944 | 2.472 | | 0.2 | |
| | P60 | 5 | 7.234 | 7.524 | 2.519 | 1.126 | | 0.0357 | 0.4524 |
| | P5 - norm | 3 | 1.00 | 1.00 | 0.37 | 0.2136 | | | |
| | P15 - norr | 4 | 1.87 | 2.2 | 1.434 | 0.7172 | | | |
| | P60 - norr | 5 | 2.1 | 2.178 | 0.7308 | 0.3268 | | | |

#### FIGURE 7

| SMA immunohistofluorescent detection in whole mouse brain | | | | | | | | | |
|---|---|---|---|---|---|---|---|---|---|
| | | | | | | | Kruskal-Wallis test | Mann Whitney test (one-tailed) | |
| Network length | | Number of values | Median | Mean | Std. Deviation | Std. Error of Mean | | P5 | P15 |
| | P5 | 3 | 9.33E-06 | 8.99E-06 | 2.30E-06 | 1.33E-06 | 0.0036 | | |
| | P15 | 3 | 1.20E-05 | 1.26E-05 | 1.17E-06 | 6.78E-07 | | 0.05 | |
| | P60 | 3 | 2.40E-05 | 2.17E-05 | 4.93E-06 | 2.85E-06 | | 0.05 | 0.05 |
| | P5 - normalized | 3 | 1.04 | 1.00 | 0.25 | 0.15 | | | |
| | P15 - normalized | 3 | 1.33 | 1.40 | 0.13 | 0.08 | | | |
| | P60 - normalized | 3 | 2.63 | 2.41 | 0.52 | 0.30 | | | |
| Number of primary branches | | Number of values | Median | Mean | Std. Deviation | Std. Error of Mean | | P5 | P15 |
| | P5 | 3 | 5.26E-08 | 5.43E-08 | 4.973E-09 | 2.871E-09 | 0.6286 | | |
| | P15 | 3 | 4.72E-08 | 4.987E-08 | 1.095E-08 | 6.32E-09 | | 0.35 | |
| | P60 | 3 | 5.10E-08 | 4.487E-08 | 1.097E-08 | 6.334E-09 | | 0.2 | 0.5 |
| | P5 - normalized | 3 | 0.97 | 1 | 0.08888 | 0.05132 | | | |
| | P15 - normalized | 3 | 0.87 | 0.92 | 0.1997 | 0.1153 | | | |
| | P60 - normalized | 3 | 0.94 | 0.8267 | 0.205 | 0.1184 | | | |
| Number of branches | | Number of values | Median | Mean | Std. Deviation | Std. Error of Mean | | P5 | P15 |
| | P5 | 3 | 7.74E-08 | 8.19E-08 | 1.49E-08 | 8.59E-09 | 0.0036 | | |
| | P15 | 3 | 1.23E-07 | 1.39E-07 | 3.86E-08 | 2.23E-08 | | 0.050 | |
| | P60 | 3 | 3.45E-07 | 2.92E-07 | 9.24E-08 | 5.33E-08 | | 0.050 | 0.050 |
| | P5 - normalized | 3 | 0.94 | 1.00 | 0.18 | 0.10 | | | |
| | P15 - normalized | 3 | 1.50 | 1.70 | 0.47 | 0.27 | | | |
| | P60 - normalized | 3 | 4.21 | 3.56 | 1.13 | 0.65 | | | |

#### SMA immunohistofluorescent detection in whole mouse brain

| | | Number of values | Median | Mean | Std. Deviation | Std. Error of Mean | | P5 | P15 |
|---|---|---|---|---|---|---|---|---|---|
| Number of ramifications | P5 | 3 | 1.24E-08 | 1.37E-08 | 4.73E-09 | 2.73E-09 | 0.0036 | | |
| | P15 | 3 | 3.53E-08 | 3.66E-08 | 7.19E-09 | 4.15E-09 | | 0.050 | |
| | P60 | 3 | 1.42E-07 | 1.222E-07 | 3.955E-08 | 2.283E-08 | | 0.050 | 0.050 |
| | P5 - normalized | 3 | 0.91 | 1.00 | 0.34 | 0.20 | | | |
| | P15 - normalized | 3 | 2.58 | 2.68 | 0.53 | 0.30 | | | |
| | P60 - normalized | 3 | 10.37 | 8.94 | 2.893 | 1.671 | | | |

| | | Number of values | Median | Mean | Std. Deviation | Std. Error of Mean | | P5 | P15 |
|---|---|---|---|---|---|---|---|---|---|
| Median straigthness | P5 | 3 | 0.95 | 0.95 | 0.01 | 0.00 | 0.3393 | | |
| | P15 | 3 | 0.92 | 0.93 | 0.03 | 0.02 | | 0.350 | |
| | P60 | 3 | 0.93 | 0.93 | 0.01 | 0.01 | | 0.050 | 0.500 |

#### Myh11 immunohistochemistry on human brain slices

| | | Number of values | Median | Mean | Std. Deviation | Std. Error of Mean | Kruskal-Wallis test | Mann Whitney test (one-tailed) | | |
|---|---|---|---|---|---|---|---|---|---|---|
| | | | | | | | | prenatal | 0-2 years | 2-5 years |
| Number of object | Pre-natal | 6 | 9.39E-08 | 1.42E-07 | 1.10E-07 | 4.48E-08 | 0.0014 | | | |
| | 0-2 years | 4 | 7.71E-07 | 8.15E-07 | 2.38E-07 | 1.19E-07 | | 0.0048 | | |
| | 2-5 years | 4 | 1.29E-06 | 1.49E-06 | 5.15E-07 | 2.57E-07 | | 0.0048 | 0.0286 | |
| | >10 years | 8 | 1.30E-06 | 1.53E-06 | 7.40E-07 | 2.61E-07 | | 0.0003 | 0.0162 | 0.4667 |

| | | Number of values | Median | Mean | Std. Deviation | Std. Error of Mean | | prenatal | 0-2 years | 2-5 years |
|---|---|---|---|---|---|---|---|---|---|---|
| Myh11+ surface | Pre-natal | 6 | 0.00 | 0.01 | 0.02 | 0.01 | 0.0011 | | | |
| | 0-2 years | 4 | 0.14 | 0.14 | 0.03 | 0.01 | | 0.0048 | | |
| | 2-5 years | 4 | 0.32 | 0.32 | 0.13 | 0.07 | | 0.0048 | 0.0286 | |
| | >10 years | 7 | 0.33 | 0.35 | 0.17 | 0.06 | | 0.0006 | 0.0061 | 0.3818 |

Supplementary Table S5

| qPCR primers | | |
|---|---|---|
| **Gene** | **Forward/Reverse** | **Sequence** |
| Tagln | F | CCCAGACACCGAAGCTACTC |
| Tagln | R | TCGATCCCTCAGGATACAGG |
| Acta2 | F | GTCCCAGACATCAGGGAGTAA |
| Acta2 | R | TCGGATACTTCAGCGTCAGGA |
| Myh11 | F | AACGCCCTCAAGAGCAAACTCAGA |
| Myh11 | R | TCCCGAGCGTCCATTTCTTCTTCA |
| Atp1b1 | F | GCTGCTAACCATCAGTGAACT |
| Atp1b1 | R | GGGGTCATTAGGACGGAAGGA |
| Abcb1 | F | GATAGGCTGGTTTGATGTGC |
| Abcb1 | R | TCACAAGGGTTAGCTTCCAG |
| Cldn5 | F | TAAGGCACGGGTAGCACTCA |
| Cldn5 | R | GGACAACGATGTTGGCGAAC |
| Gapdh | F | AGGTCGGTGTGAACGGATTTG |
| Gapdh | R | TGTAGACCATGTAGTTGAGGTCA |
| Col1a1 | F | GCTCCTCTTAGGGGCCACT |
| Col1a1 | R | CCACGTCTCACCATTGGGG |
| Acer 2 | F | GTGTGGCATATTCTCATCTG |
| Acer 2 | R | TAAGGGACACCAATAAAAGC |
| Ly6c1 | F | GCAGTGCTACGAGTGCTATGG |
| Ly6c1 | R | ACTGACGGGTCTTTAGTTTCCTT |
| Slco1a4 | F | GCCAGTGCAAGGCCAGAACC |
| Slco1a4 | R | AGGGTGGGGGTTTGCTACAGAA |

| qPCR probes | | |
|---|---|---|
| **Gene** | **Supplier** | **Reference** |
| Gapdh | Thermo Fisher Scientific | Mm99999915_g1 |
| Lama1 | Thermo Fisher Scientific | Mm01226102_m1 |
| Lum | Thermo Fisher Scientific | Mm01248292_m1 |
| Kcnj8 | Thermo Fisher Scientific | Mm00434620_m1 |
| Abcc9 | Thermo Fisher Scientific | Mm00441638_m1 |
| Col6a1 | Thermo Fisher Scientific | Mm00487160_m1 |

| FISH probe | | | |
|---|---|---|---|
| **Gene** | **Supplier** | **Probe** | **Reference** |
| Myh11 | Advanced cell diagnostic | RNAScope® Mm-Myh11 | 316101 |

| Antibodies | | | | | | |
|---|---|---|---|---|---|---|
| | **Protein** | **Species** | **reference** | **supplier** | **Western blot dilution** | **Immunohistology dilution** |
| Primary antibodies | Claudin 5 | rabbit | 34-1600 | THERMO FISHER | 1:500 | / |
| | H3 | mouse | 14269S | OZYME | 1:2000 | / |
| | Myh11 | rabbit | ab53219 | ABCAM | 1:1000 | / |
| | Myh11 | rabbit | HPA015310 | SIGMA | | 1:50 |
| | P-gP | mouse | ALX-801-002-C100 | ENZO | 1:200 | / |
| | P-gP | mouse | 517312 | SIGMA | | 1:25 |
| | SMA_cy3 | mouse | C6198 | SIGMA | 1:1000 | 1:250 |
| | Pecam-1 | goat | AF3628 | R&D SYSTEMS | / | 1:300 |
| Secondary antibodies | anti-goat_alexa 647 | donkey | A-21447 | LIFE | / | 1:2000 |
| | anti-rabbit_HRP | goat | CSA2115 | COHESION | 1:2500 | / |
| | anti-mouse_HRP | goat | CSA2108 | COHESION | 1:2500 | / |
| Others | IB4 | Alexa-conjugated Isolectin (*griffonia simplicifolia*) | I32450 | INVITROGEN | 1:100 | / |
